# Targeting Mitochondrial Metabolism in Clear Cell Carcinoma of the Ovaries

**DOI:** 10.1101/2021.02.17.431266

**Authors:** Xianonan Zhang, Mihir Shetty, Valentino Clemente, Stig Linder, Martina Bazzaro

## Abstract

Ovarian clear cell carcinoma (OCCC) is a rare but chemorefractory tumor. About 50% of all OCCC patients have inactivating mutations of ARID1A a member of the SWI/SNF chromatin remodeling complex. Members of the SWI/SNF remodeling have emerged as regulators of the energetic metabolism of mammalian cells, however the role of ARID1A as a modulator of the mitochondrial metabolism in OCCCs is yet to be defined. Here we show that ARID1A-loss results in increased mitochondrial metabolism and renders ARID1A-mutated cells increasingly and selectively dependent on it. The increase in mitochondrial activity following ARID1A loss is associated to increase of C-myc and to increased mitochondrial number and reduction of their size consistent with a higher mitochondrial cristae/outer membrane ratio. Significantly, preclinical testing of the complex I mitochondrial inhibitor IACS-010759 extends overall survival in a preclinical model of ARID1A-mutated OCCC. These findings provide the for targeting mitochondrial activity in ARID1A-mutanted OCCCs.

## Introduction

Ovarian clear cell carcinoma (OCCC) has a 5-year overall survival of less than 10% and it is often associated with inactivating mutations of ARID1A, a component of SWI/SNF nucleosome remodeling complex. Over half of the patients with OCCC harbor loss-of-function mutations in ARID1A (Fukumoto et al., 2018). Mutations of members of the SWI/SNF remodeling complex are found in approximately 20% of human cancers including another untreatable, although rare, ovarian cancer histotype the cell carcinoma of the ovary, hypercalcemic type (SCCOHT) where inactivating mutations of *SMARCA4* mutation are found in 100% of the patients (Jelinic et al., 2014).

The SWI/SNF remodeling complex has been shown to control the energetic metabolism of mammalian cells including cancer cells. For instance: *a)* a study conducted in lung cancer shows that mutations in SMARCA4, a SWI/SNF complex member, induced targetable dependence upon mitochondrial inhibitors (Deribe et al., 2018; Fukumoto et al., 2018), *b)* a study conducted in ovarian cancer shows that OCCC-derived cells have higher mitochondrial respiration as compared to ovarian cancer cell lines derived from other histotypes (Dier et al., 2014), c) a study conducted in ovarian cancer shows that loss of ARID1A leads to sensitivity to ROS-inducing agents (Kwan et al., 2016), and *d)* a study shows that targeting the vulnerability to glutathione metabolism can be a strategy to treat ARID1A-deficient human cancers (Ogiwara et al., 2019). Today, the role of ARID1A as a modulator of the specific mitochondrial metabolism in OCCCs remains elusive.

Mitochondrial dynamics is the process through which once formed, mitochondria undergo the cycling process of fusion and fission. Tight regulation of mitochondrial dynamics is fundamental for cell functions and its disruption has been associated to aging, neurogenerative diseases and cancer (Anderson et al., 2018; Archer, 2013; Maycotte et al., 2017; Samant et al., 2014; Trotta and Chipuk, 2017). For instance, a number of studies show that fragmented mitochondrial phenotype is essential in many human tumors (Kashatus et al., 2015; Malhotra et al., 2016; Wieder et al., 2015; Zhang et al., 2013; Zhao et al., 2013). A number of recent reports also show that increased mitochondrial fission is associated with higher mitochondrial activity in cancer cells (Anderson et al., 2018; Peiris-Pages et al., 2018; Rao, 2019). This seems to be due to a higher ratio of cristae to outer membrane surface of the mitochondria (Devine and Kittler, 2018; Perkins et al., 2001; Perkins et al., 2010; Picard et al., 2013).

C-myc has been shown to be involved in regulating mitochondrial biogenesis and trafficking (Li et al., 2005; Morrish and Hockenbery, 2014). In prostate cancer, C-myc has been shown to regulate cancer metastasis via regulating mitochondrial trafficking (Agarwal et al., 2019) and to control mitochondrial fission via upregulating the Mitochondrial Fission Factor (MFF) (Seo et al., 2019). In neuroblastomas, N-myc has been shown to regulate both mitochondrial trafficking and dynamics (Casinelli et al., 2016). C-myc amplification is common in ovarian cancer (Zeng et al., 2018) but is particularly common in the OCCC subtype where over 40% of patients have C-myc gain (Dimova et al., 2006). In mouse fibroblasts C-myc has been shown to be a direct target of ARID1A (Nagl et al., 2006). In OCCCs, ARID1A has been shown to control global transcription (Trizzino et al., 2018). In another highly chemoresistant ovarian cancer histotype, high grade serous carcinoma (HGSC), active C-myc transcription has been shown to be dependent upon continuous transcription (Zeng et al., 2018).

Inhibition of mitochondrial bioenergetics has raised increasing interest in the field of cancer therapeutics (Emmings et al., 2019; Molina et al., 2018; Sica et al., 2020; Zhang et al., 2016) (Frattaruolo et al., 2020; Oliveira et al., 2021). A number of different inhibitors are being evaluated, including inhibitors of the respiratory chain, the tricarboxylic acid cycle and mtDNA (Oliveira et al., 2021). IACS-010759 is a clinical-grade small-molecule inhibitor of the Complex I of the electron transport chain (Molina et al., 2018). IACS-010759 has been shown to selectively kill tumor cells that depend on OXPHOS both, *in vitro* and in preclinical models of human cancers with no cytotoxicity at tolerated doses in normal cells (Bajpai et al., 2020; Fischer et al., 2019; Lissanu Deribe et al., 2018; Molina et al., 2018; Panina et al., 2020; Sun et al., 2019; Teh et al., 2020; Tsuji et al., 2020; Vashisht Gopal et al., 2019; Zhang et al., 2019).

Here we show that ARID1A-deficient cells upregulate OXPHOS genes and pathways and that that corresponds to an upregulation of mitochondrial membrane potential and respiration, and to an increase in mitochondrial number and their of cristae to outer membrane surface ratio. This mechanism could be at least in part due to upregulation of C-myc protein and targets. We also show that sensitivity of ARID1A-deficient cells to the Complex I mitochondrial inhibitor IACS-010759 is exquisitely depend upon ARID1A and IACS-010759 severely compromise mitochondrial respiration of ARID1A-deficient cells in a dose-dependent manner in both monolayer culture and multicellular spheroids. Lastly, we show that in a preclinical model of ARID1A-deficient OCCCs, IACS-010759 treated mice have twice the overall survival as compared to vehicle-treated group with no observable toxicity to the host.

## Results

### ARID1A-deficient cells have increased expression of OXPHOS genes and pathways

To begin investigating the possible effects of ARID1A loss on mitochondrial metabolism, ARID1A was knocked down via siRNA in ARID1A WT ES-2 cells (Figure 1A, and Supplementary Figure 1A) and control and knockdown cells were subjected to transcriptomic profiling using RNA-sequencing. Gene set enrichment analysis (GSEA) revealed that E2F and MYC targets were upregulated in ARID1A knockdown cells, suggesting effects on cell cycle progression. Furthermore, and interesting considering previous findings in the literature (Dier et al., 2014), the oxidative phosphorylation pathway is among the most prominently enriched pathways in ARID1A-mutated cells (Figure 1B-D). This includes upregulation of several genes encoding for components of the mitochondrial Electron Transport Chain (ETC) complexes I, III, IV and V (Figure 1E). The full list of the genes differentially expressed in scramble *vs* ARID1A KD ARID1A WT ES-2 cells is available in Supplementary Table 1. Taken together this shows that loss of ARID1A in ARID1A WT cells results in upregulation of OXPHOS pathway by GSEA analysis and biological processes regulating mitochondrial gene expression by GO analysis.

**Figure 1.**
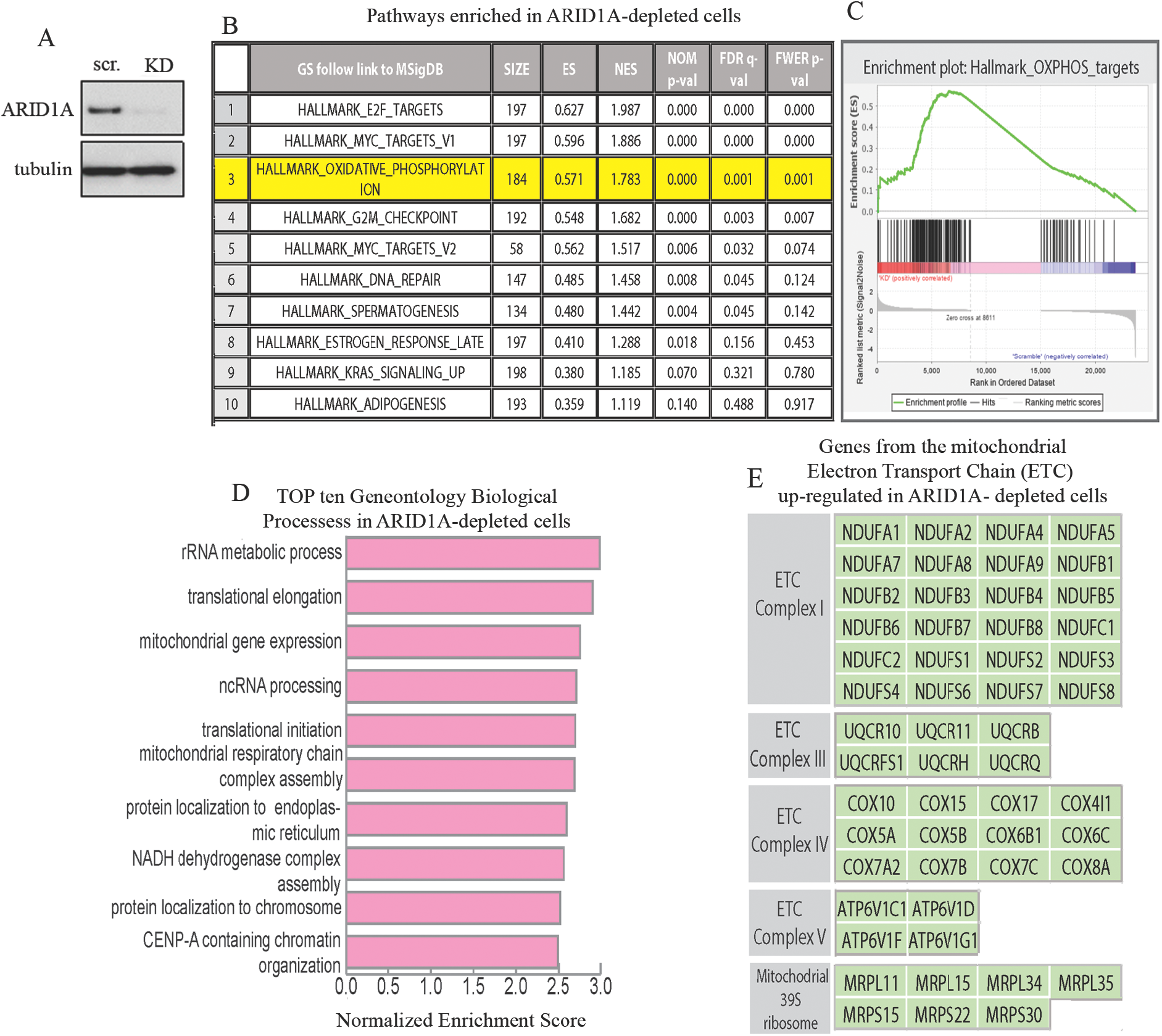
ARID1A-deficient cells upregulate OXPHOX pathways and increase mitochondrial respiration. **A**. Western blot analysis for levels of ARID1A in scramble (scr.) versus ARID1A knockdown (KD) in ARID1A WT ES-2 cells. Tubulin was used as loading control. **B**. GSEA data analysis of microarray transcriptomic profiling in ARID1A KD versus scramble ES-2 cells revealed that the OXPHOS pathway is among the top-most upregulated pathways following ARID1A loss. RNA-sequencing data were obtained using 3 independent samples for each cohort (scramble and KD). **C**. Representative graph for enriched mitochondrial OXPHOS pathway (p=0.000, q=0.001). **D.***Left*, Top ten geneontology (GO) biological process (p<0.001) in ARID1A KD versus scramble ES-2 cells. **E.** Genes coding for mitochondrial Electron Transport Chains (ETC) are upregulated in ARID1A KD versus scramble ES-2 cells. Results are from three independent experiments and are expressed as mean ± SD.

### ARID1A regulates mitochondrial respiration in OCCC cells

To determine whether the enrichment in OXPHOS pathways following ARID1A loss is associated with a functional increase in mitochondrial activity, we measured the mitochondrial oxygen consumption rate (OCR) in presence and in absence of ARID1A. To this end, ARID1A was knocked down via siRNA in ARID1A WT ES-2 cells (Figure 2A, and Supplementary Figure 1B, *left*) and ATP-linked respiration and maximal respiration was measured in scramble versus ARID1A KD equal amounts of cells (Supplementary Figure 1C). We found that loss of ARID1A resulted in increases in OCR (Figure 2B), ATP-linked respiration (Figure 2C) and maximal respiration (Figure 2D). The same results were confirmed when ARID1A was knocked down in an additional ARID1A WT cell line, RMG1 (Figure 2E-H and Supplementary Figure 1B, *right,* and D). To further establish that ARID1A is responsible for increases in mitochondrial respiration, we conducted complementary experiments and tested whether ectopic expression of ARID1A in the ARID1A-mutated TOV-21G ovarian cancer cell line would result in decreases in mitochondrial activity. As shown in Figure 2I-L (and Supplementary Figure 1E and F), we found that cells expressing ARID1A had significantly lower mitochondrial activity as compared to vector alone expressing cells. Noteworthy, the decrease in mitochondrial activity in TOV-21 ARID1A overexpressing cells is less dramatic as compared to the increase in mitochondrial activity following ARID1A KD. This is likely due to the fact that re-expression of ARID1A in TOV-21G induces cell senescence via p21 expression (not shown, personal communication from Dr. Rugang Zhang, Wistar Institute). Taken together this indicates that ARID1A is responsible for controlling mitochondrial activity in OCCC cells and that its’ loss results in increases in mitochondrial respiration.

**Figure 2.**
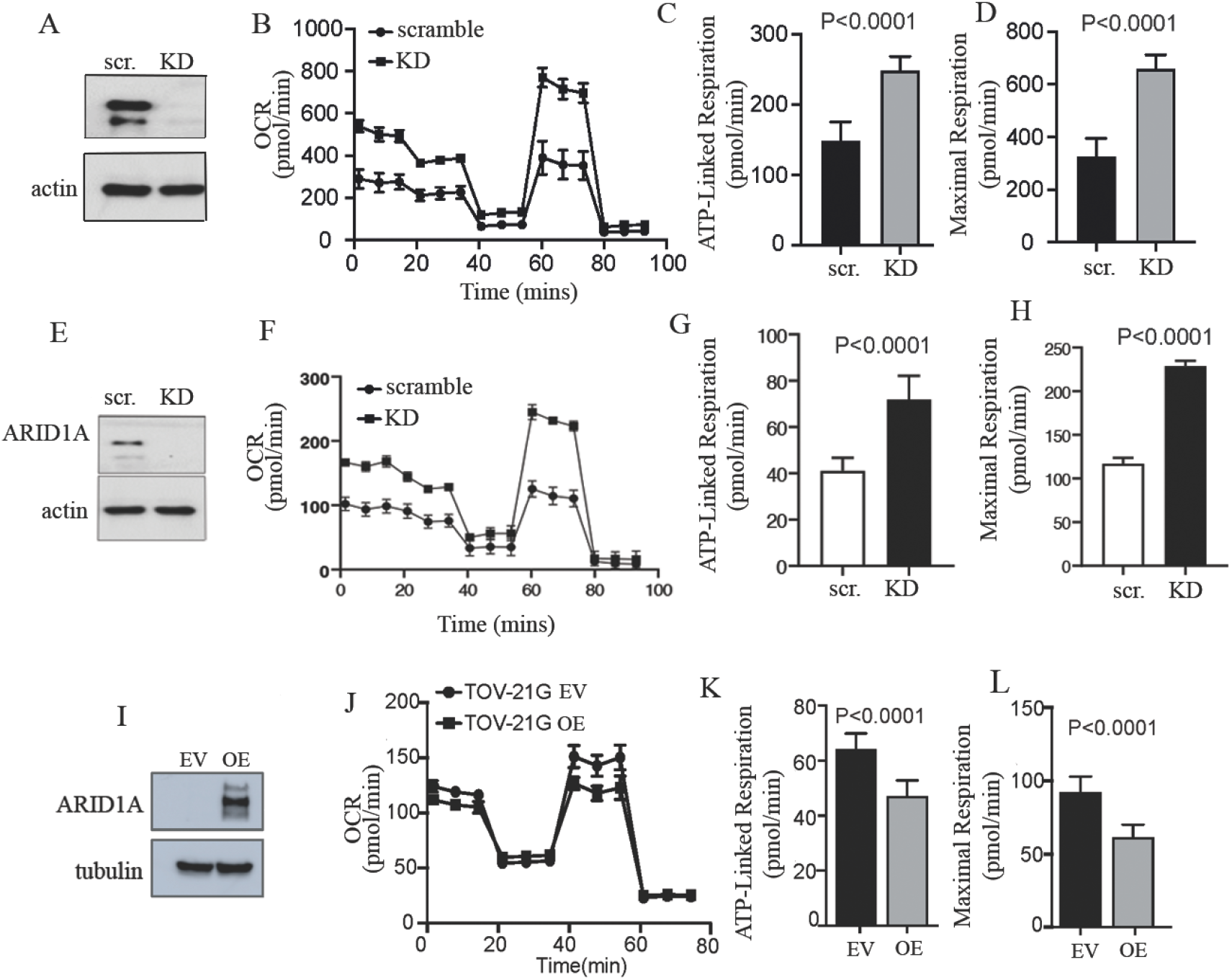
ARID1A controls mitochondria metabolism in OCCC cells. **A.** Western blot analysis for levels of ARID1A in scramble (scr.) versus ARID1A knockdown (KD) ARID1A WT ES-2 cells. Actin was used as a loading control. **B.** Real time Oxygen Consumption Rate (OCR) in scramble versus ARID1A KD ES-2 cells as measured by Seahorse. **C.** ATP-linked respiration in ARID1A in scramble (scr.) versus ARID1A knockdown (KD) ARID1A WT ES-2 cells. **D.** Maximal respiration in scramble versus ARID1A KD ES-2 cells. **E.** Western blot analysis for levels of ARID1A in scramble (scr.) versus ARID1A knockdown (KD) ARID1A WT RMG1 cells. Actin was used as a loading control. **F.** Real time Oxygen Consumption Rate (OCR) in scramble versus ARID1A KD RMG1 cells as measured by Seahorse. **G.** ATP-linked respiration in ARID1A in scramble (scr.) versus ARID1A knockdown (KD) ARID1A WT RMG1 cells. **H.** Maximal respiration in scramble versus ARID1A KD RMG1 cells. **I.** Western blot analysis for levels of ARID1A empty vector (EV) versus ARID1A overexpressing (OE) ARID1A-mutated TOV-21G cells. Tubulin was used as a loading control. **J.** Real time Oxygen Consumption Rate (OCR) in ARID1A empty vector (EV) versus ARID1A overexpressing (OE) ARID1A-mutated TOV-21G cells. **K.** ATP-linked respiration in ARID1A empty vector (EV) versus ARID1A overexpressing (OE) ARID1A-mutated TOV-21G cells. **L.** Maximal respiration in ARID1A empty vector (EV) versus ARID1A overexpressing (OE) ARID1A-mutated TOV-21G cells. Results are from three independent experiments and are expressed as mean ± SD.

### ARID1A controls mitochondrial membrane potential, mitochondrial number and size

The finding that ARID1A regulates mitochondrial respiration in OCCC cells prompted us to examine alterations in mitochondrial membrane potential. To this end, scramble and KD RMG1 and ES-2 ARID1A WT cells were stained with the MitoTracker Deep Red dye which is known to accumulate in mitochondria in a membrane-potential dependent manner (Xiao et al., 2016). As shown in Figure 3A and B, loss of ARID1A resulted in increases in the uptake of MitoTracker Deep Red in both ES2 and RMG1 OCCC cells. We next determined whether loss of ARID1A would result with an increase in mitochondrial content. To this end, scramble versus ARID1A KD ARID1A WT RMG1 cells were subjected to immunostaining using an antibody to the mitochondrial marker TOM20. Mitochondrial content was visualized and evaluated via immunofluorescence analysis of reconstructed mitochondria. As shown in Figure 3C and D, loss of ARID1A resulted in a statistically significant increases of the mitochondrial mass in RMG1 cells.

**Figure 3.**
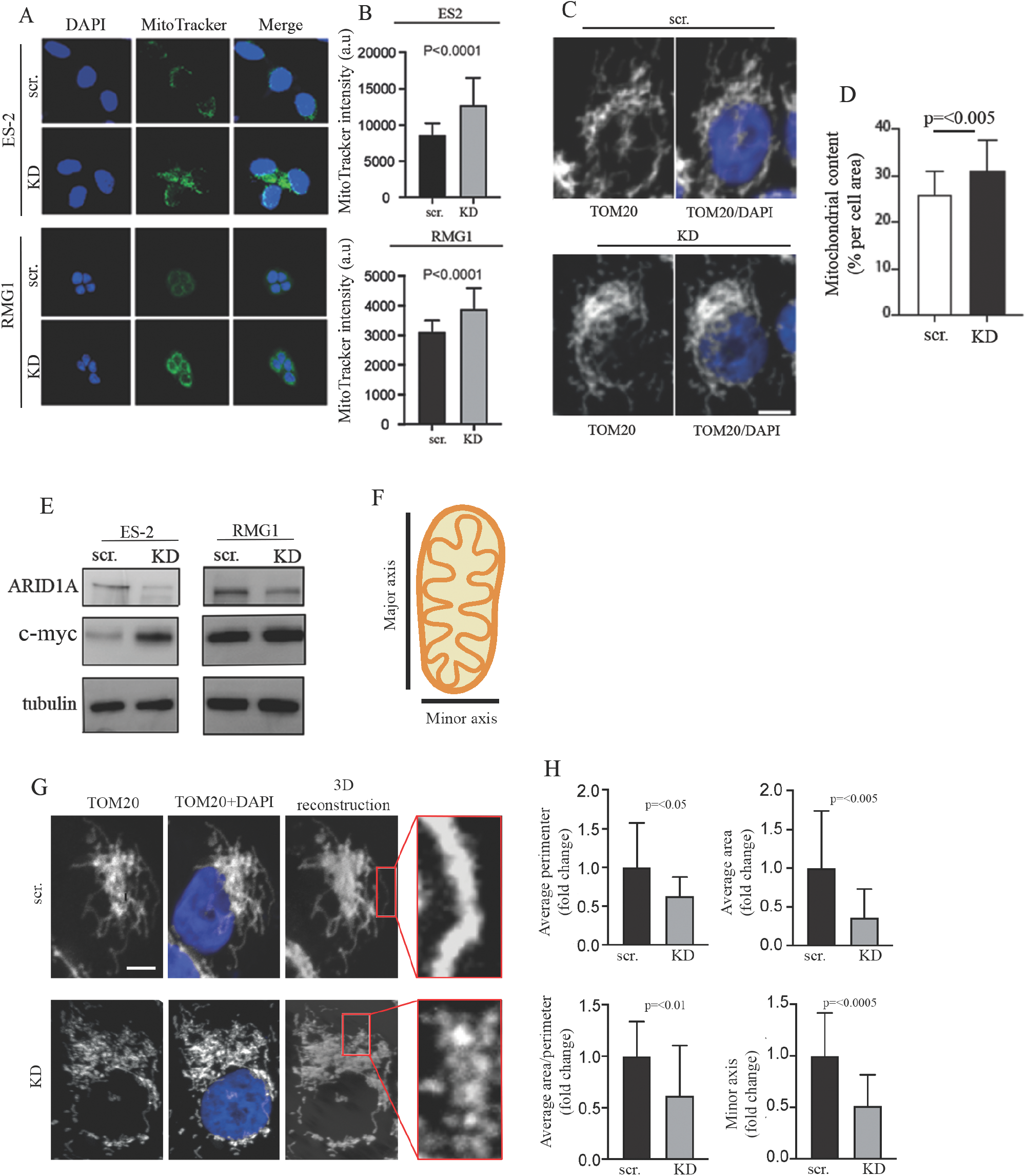
Loss of ARID1A results in increased mitochondrial membrane potential and increased mitochondrial content and fragmentation. **A.** Representative images (original magnification 60X) of scramble (scr.) and ARID1A KD (KD) ES-2 (*top four panels*) and RMG1 (*bottom four panels*) ARID1A WT cells stained with DAPI (blue) and MitoTracker green. Merge image is also shown. **B.** Quantification of MitoTracker fluorescence intensity in ES-2 (*top panel*) or RMG1 (*bottom panel*) cells under the above-mentioned conditions expressed as arbitrary units (a. u). **C.** Representative image of mitochondrial content in scramble (scr.) versus ARID1A knockdown (KD) ARID1A WT RMG1 cells evaluated via TOM20 staining. **D.** Quantification of mitochondrial content expressed per each condition. **E.** *Left,* Western blot analysis for levels of ARID1A and c-myc in scramble (scr.) versus ARID1A knockdown (KD) ARID1A WT ES-2 cells. Tubulin was used as a loading control. *Right*, Western blot analysis for levels of ARID1A and c-myc in scramble (scr.) versus ARID1A knockdown (KD) ARID1A WT RMG1 cells. Tubulin was used as a loading control. **F.** Representation of parameters used to evaluate mitochondrial morphology. **G.** Representative images of TOM20 and DAPI staining in scr. versus ARID1A knockdown (KD) ARID1A WT RMG1 cells followed by 3D reconstruction of mitochondria. **H.** Average mitochondrial perimeter in scr. *vs* KD cells (*top left*), Average mitochondrial area in scr. *vs* KD cells (*top right*), Average mitochondrial area/perimeter in scr. *vs* KD cells (*bottom left*). Minor axis in scr. *vs* KD cells (*bottom right*). Results are from three independent experiments and are expressed as mean ± SD.

A higher ratio of mitochondrial cristae to outer membrane surface has been associated with increased mitochondrial activity (Devine and Kittler, 2018; Perkins et al., 2001; Perkins et al., 2010; Picard et al., 2013). Thus, we investigated whether loss of ARID1A resulted with changes in mitochondrial shape and size in ARID1A scramble versus ARID1A WT RMG1 cells. To this end, we measured both minor and major mitochondrial axes (Figure 3F) as well as in cell perimeter and area in TOM20+DAPI stained cells (Figure 3G) per each condition. As shown in Figure 3H loss of ARID1A led to an overall decrease in mitochondrial size. Taken together this suggests that the increase in mitochondrial activity following ARID1A loss is at least in part due to both, increase in mitochondrial biogenesis and decrease in mitochondrial size. This is consistent with previous reports showing that increase in mitochondrial fragmentation is associated to more aggressive cancer cells’ behavior (Yu et al., 2019).

The increase in mitochondrial content followed by knockdown of ARID1A was accompanied with an increase in the expression levels of C-myc in both ES-2 and RMG1 ovarian cancer cell lines (Figure 3E). This is consistent with a functional antagonism between C-myc and the SWI/SNF complex (Romero et al., 2012) and is also consistent with our RNA-seq data showing that C-myc targets are upregulated following ARID1A loss (Figure 1B).

### ARID1A loss results with selective sensitivity to the mitochondrial inhibitor IACS-010759 in 2D

We next tested whether the increase in mitochondrial activity following ARID1A loss in OCCCs would render these cells selectively dependent upon mitochondrial energy production. To this end we selected a panel of nine mitochondrial inhibitors (Figure 4A) that were in clinical trials at the time when this study was initiated (source: https://www.clinicaltrials.gov) and subjected scramble and ARID1A KD ES-2 ARID1A WT cells to a colony formation assay in absence (mock) and in presence of increasing concentration of the mitochondrial inhibitors indicated in Figure 4A. As shown in Figure 4B, all nine inhibitors induced a dose-dependent decrease in the clonogenicity of ES-2 cells. However, only IACS-010759 (Tsuji et al., 2020) and dapagliflozin (Nasiri et al., 2019) displayed selectivity for ES-2 cells where ARID1A has been knocked down. Of these two, IACS-010759 showed a much higher potency. These results were confirmed using the OCCCs derived ARID1A WT RMG1 cell line. Specifically, using the same colony formation assay, ARID1A KD RMG1 cells (Figure 4C) were more sensitive to the mitochondrial inhibitor IACS-010750 as compared to ARID1A WT (scramble) RMG1 cells (Figure 4D). To confirm that the selective sensitivity to IACS-010759 is dependent upon ARID1A we performed complementary experiment using the ARID1A mutated OCCCs-derived TOV-21G cell line. Specifically, ARID1A was overexpressed in TOV-21 cells (Figure 2I) and sensitivity to IACS-010759 was determined via colony formation assay. As shown in Supplementary Figure 3, expression of ARID1A in the ARID1A-mutated TOV21G cell line results with loss of sensitivity to the mitochondrial inhibitor IACS-010759. Lastly, we confirmed that treatment with IACS-010759 results in dose-dependent decrease in mitochondrial activity in ARID1A KD RMG1 cells as determined via measuring OCR (Figure 4E), ATP-linked respiration (Figure 4F) and maximal respiration (Figure 4G) in scramble and ARID1A KD cells exposed to increasing concentrations of IACS-010759. Taken together this suggests that IACS-010759 selectively affects the cell viability of ARID1A mutated cells via inhibiting their greater demand for mitochondrial activity.

**Figure 4.**
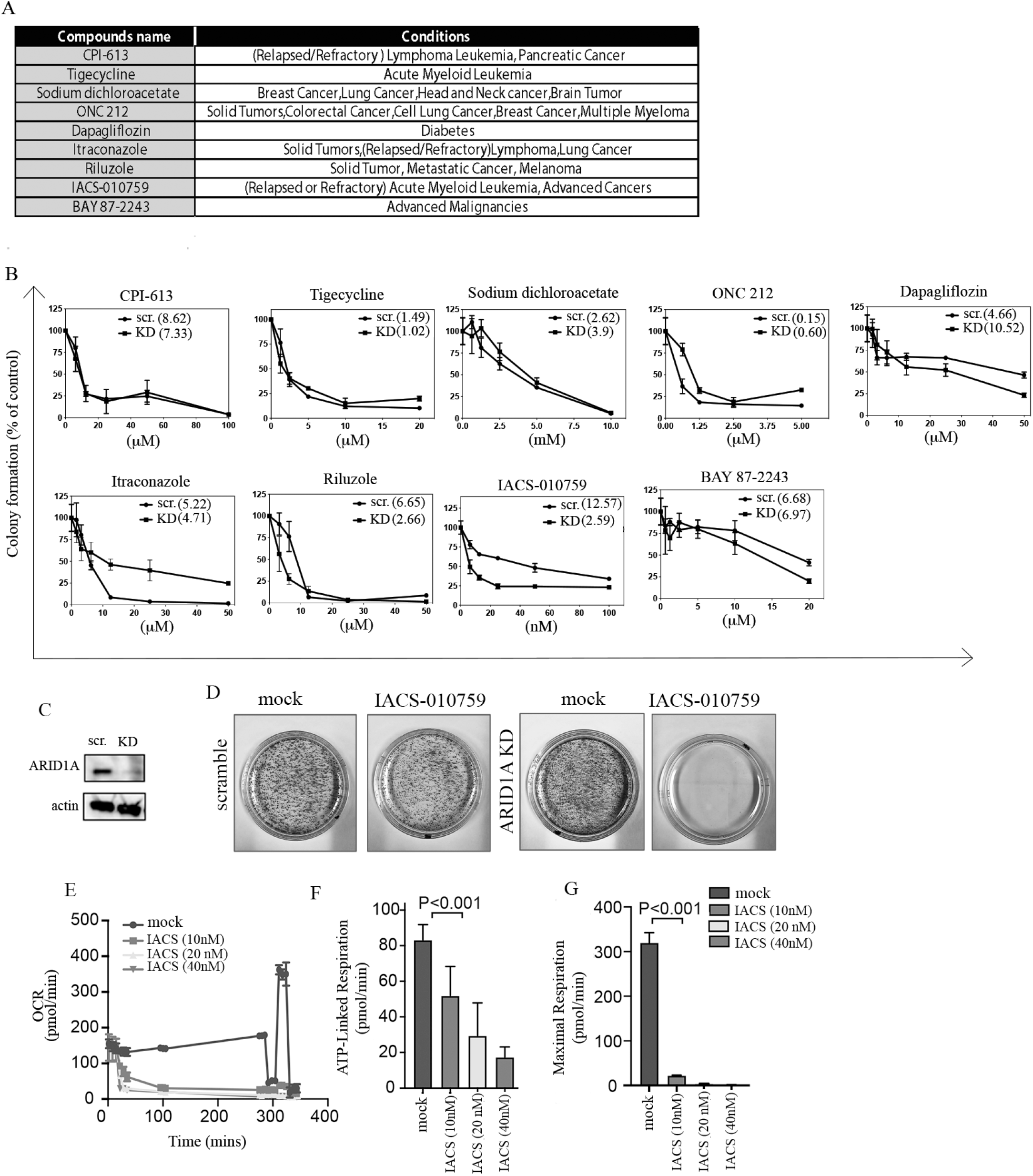
Loss of ARID1A results in selective sensitivity to mitochondrial inhibition in 2D. **A.** Table of compounds with mitochondrial inhibitions properties in clinical trials. Compounds name and conditions for which they are used are indicated. Source: https://clinicaltrials.gov/. **B.** Dose-dependent sensitivity of scramble (scr.) and ARID1A KD (KD) ES-2 WT cells to the panel of mitochondrial inhibitors. Results are expressed as residual colony formation as compared to control. Drug treatment was conducted over the period of 7 days. **C.** Western blot analysis for expression levels of ARID1A in scramble and ARID1A KD RMG1cells obtained by lentiviral-mediated delivery of shRNA-ARID1A. Actin was used as a loading control. **D.** Residual colony formation in scramble and ARID1A KD RMG1 cells exposed to 10 nM of the mitochondrial inhibitor IACS-010759 over the period of 10 days. **E.** Real time Oxygen Consumption Rate (OCR) in scramble versus ARID1A KD RMG1 cells exposed to increasing concentrations of the mitochondrial inhibitor IACS-010759. **F.** ATP-linked respiration in ARID1A in scramble (scr.) versus ARID1A knockdown (KD) ARID1A WT RMG1-2 cells exposed to increasing concentrations on the mitochondrial inhibitor IACS-010759. **G.** Maximal respiration in ARID1A in scramble (scr.) versus ARID1A knockdown (KD) ARID1A WT RMG1-2 cells exposed to increasing concentrations on the mitochondrial inhibitor IACS-010759.

### ARID1A loss results with selective sensitivity to the mitochondrial inhibitor IACS-010759 in three-dimensional culture

Here we sought of determining whether IACS-010759 retains selectivity toward ARID1A-deficient cells in a three-dimensional culture system. To this end ES-2-derived spheroids were either mock treated or treated with increasing concentrations (500 nM and 1 μM) of IACS-010759 over a period of 10 days. Dapagliflozin, which displayed selectivity for ES-2 cells where ARID1A has been knocked down although at micromolar concentrations was also used. As shown in Figure 5A (*left panel*) and Figure B (*top panel*) IACS-010759 treatment led to a decrease in the spheroids’ volume that was greater in the ARID1A-mutated cells as compared to ARID1A WT cells. Dapagliflozin treatment on the other end, had minimal effect on the spheroids’ volume in both ARID1A-mutated or ARID1A WT cells (Figure 5A, *right panel* and Figure 5B, *bottom panel*). Lastly, we determined the effect of IACS-010759 treatment on mitochondrial respiration in spheroids. As shown in Figure 5C, IACS-010759 treatment led to significant reduction of mitochondrial activity as determined via measuring OCR. Taken together this suggests that IACS-010759 selectively affects the cell viability of ARID1A mutated cells in 3D cultures.

**Figure 5.**
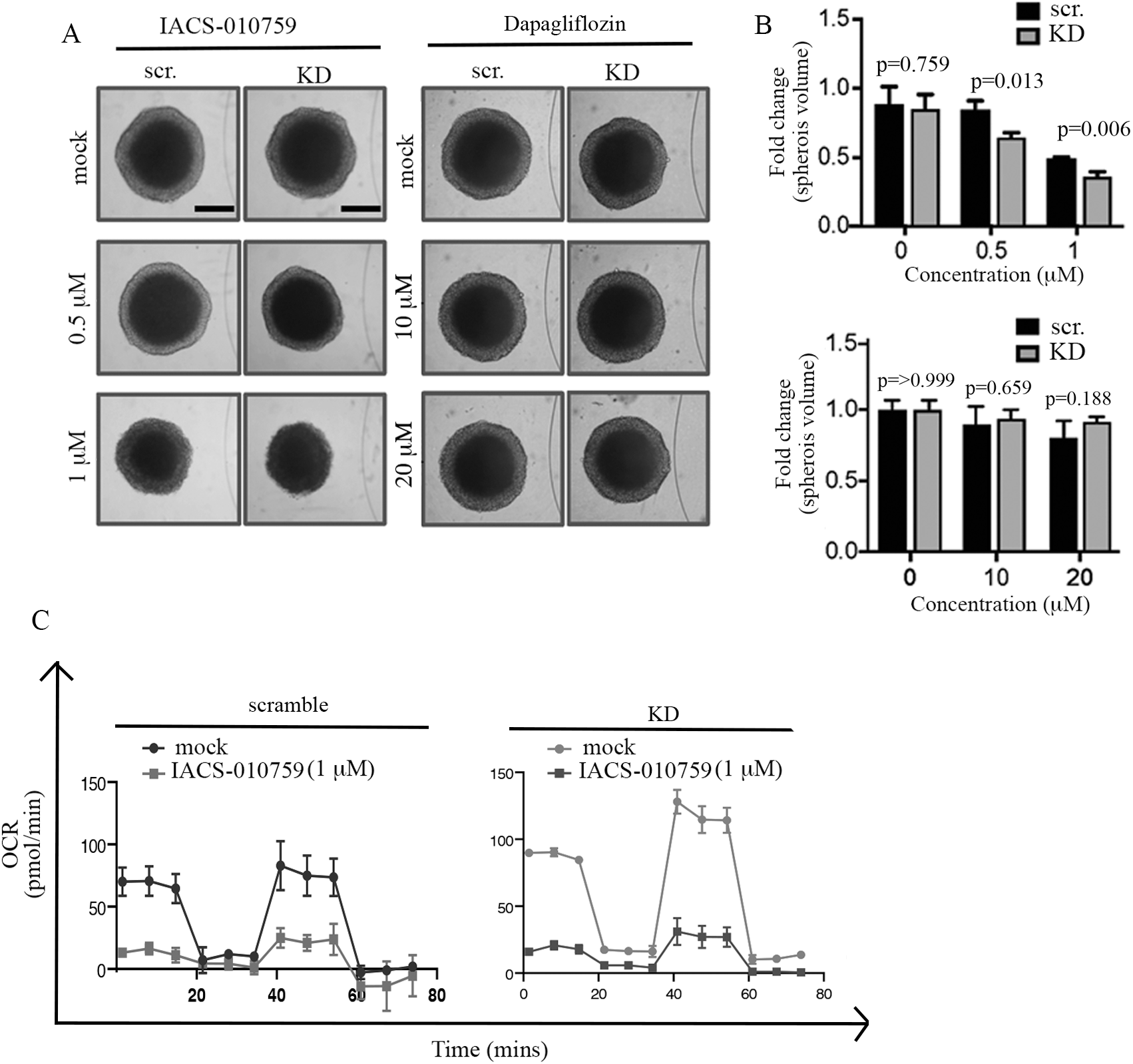
Loss of ARID1A results in selective sensitivity of spheroids to mitochondrial inhibition. **A.** ES-2 derived spheroids exposed to the indicated concentrations of either IACS-010759 (*left*) or Dapagliflozin (right) over a period of 48 hours. **B.** Quantification of residual spheroid volume in cells exposed to IACS-010759 (*top*) or Dapagliflozin (*bottom*). **C.** Real time Oxygen Consumption Rate (OCR) in scramble versus ARID1A KD spheroids derived from ES-2 cells exposed to the indicated concentrations of IACS-010759.

### IACS-010759 treatment prolongs survival in a model of ARID1A-mutated OCCCs

To address the clinical relevance of targeting mitochondrial function as a potential therapeutic strategy for the treatment of ARID1A-deficient OCCC cancers, we tested the ability of IACS-010759 to suppress the growth of ARID1A-deficient OCCC tumors *in vivo*. Specifically, ARID1A-deficient TOV-21G cells were intraperitoneally injected in athymic nude female mice which were randomly assigned to three treatment arms: vehicle alone, IACS-010759 5 mg/kg and IACS-010759 10 mg/kg. These doses were chosen because they were previously described as well tolerated in preclinical studies (Lissanu Deribe et al., 2018; Molina et al., 2018; Zhang et al., 2019). Treatment was performed via oral gavage on a 5 d on/2 d off schedule. As shown in Figure 6A, while mice in the control group (treated with vehicle) had to be sacrificed by week 3 according to the IACUC guideline, mice in both IACS-010750 treated arms survived almost as twice longer and until the experiments were terminated because their reached significance at month two. More specifically and as shown in Figure 6B, the mean days alive in treated groups (mean days alive in 5mg/kg group is 50.1±0.7 days and 10mg/kg group is 44.8±4.9 days) are more than two times of that in the control group (mean days alive is 24.8±1.3 days. More importantly, during the two-month experiment, mice kept a stable growing weight in the treated groups (mice weight increased more than 20%, even dropped but still higher than the starting weight before sacrifice, Figure 6C) which is not due to the increased volume of fluids in abdomen (4.25±0.31 mL in the control group, 1.3±0.88 mL and 0.29±0.25 mL in groups of 5 mg/kg and 10 mg/kg, Figure 6D) which is a characteristic of ovarian cancer (Ahmed and Stenvers, 2013; Penet et al., 2018). Taken together, this suggests that targeting mitochondrial function by IACS-010759 could represent an effective strategy for the treatment of ARID1A-mutated OCCCs.

**Figure 6.**
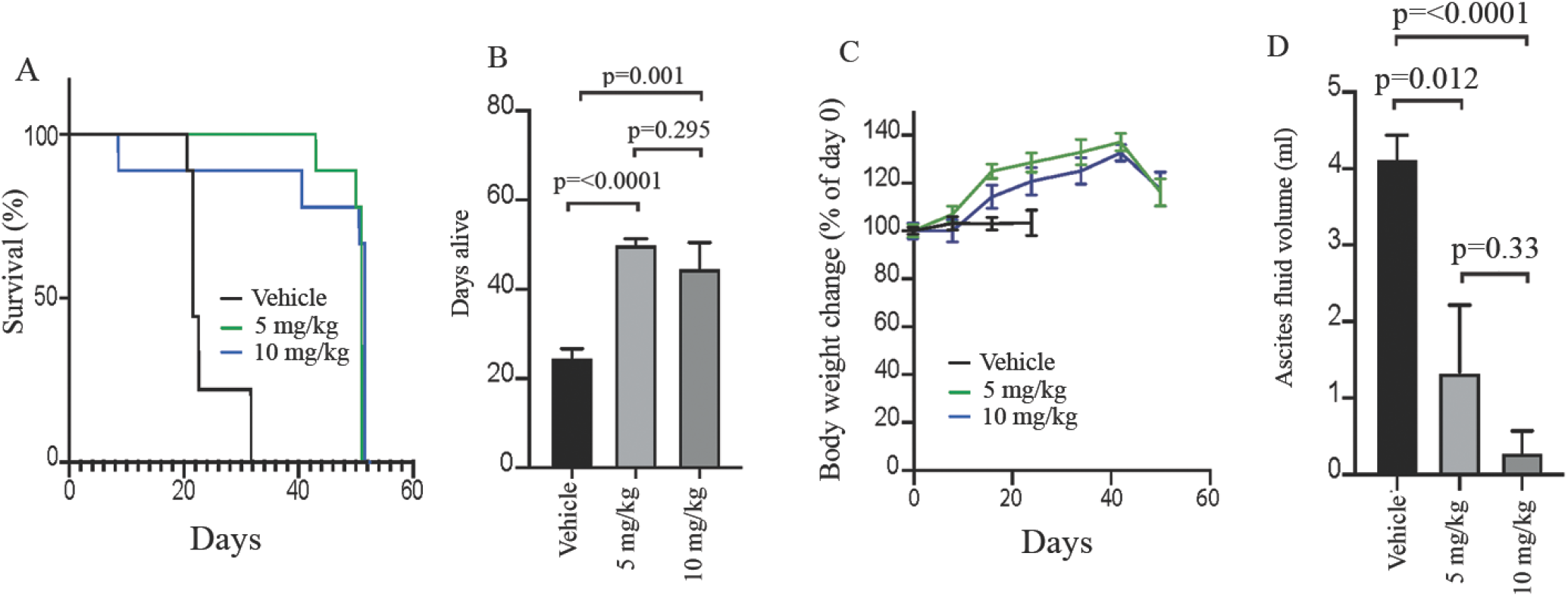
Preclinical testing of IACS-010759 in a xenograft model of OCCC mutated tumor. **A.** The percentage survival of mice. **B.** Days the mice are alive. **C**. Percentage body weight overtime. **D**. Ascitic fluid volume after intraperitoneal injection of TOV21G cells followed by oral dosing of either vehicle control or indicated dose of IACS-010759 (N=9 mice/group).

## Discussion

Here we report that knock-down of ARID1 expression in ovarian cancer cells expressing this protein results in increases in oxidative phosphorylation (OXPHOS). We also show that restoration of ARID1A in previously defective cells results in decreases in OXPHOS. These findings are consistent with those reporting that lung cancer cells with mutations in another component of the SWI/SNF complex (SMARCA4) display enhanced oxygen consumption (Lissanu Deribe et al., 2018). Both studies demonstrate a vulnerability of cells with ARID1A/SMARCA4 mutations/defects to the effects on mitochondrial inhibitors, raising the possibility for therapeutic interventions. The underlying mechanisms are not entirely clear but may relate to C-myc activity. C-myc has been shown to be antagonized by the SWI/SNF complex (Romero et al., 2012) and C-myc is well known to play a major role in metabolic dysregulation of tumors (Dang et al., 2006; Hsieh et al., 2015; Stine et al., 2015). Whereas SMARCA4 mutations were not reported to result in increased of C-myc expression, gene set enrichment analyses (GSEA) showed increase in C-myc target genes (Lissanu Deribe et al., 2018). We observed both a large increase in C-myc expression in ES-2 cells transfected with ARID1A siRNA and also increases in C-myc targets genes in GSEA (Fig. 1). The issue of whether the effect of ARID1A on OXPHOS is mediated by C-myc could theoretically be addressed by knocking down C-myc, but the results would be confounded by the ubiquitious effects of C-myc on cell proliferation and other processes.

The efficacy of glycolysis in terms of ATP production is too low to meet the energy demands of highly active tumor cells, resulting in dependence on mitochondrial energy production (Wallace, 2012). Cells display a high degree of metabolic plasticity, shifting between different energy sources and metabolic pathways. Inhibition of OXPHOS may therefore enable the upregulation of compensatory pathways, such as glycolysis, to support cancer cell survival. It has occasionally been suggested that inhibition of both mitochondrial metabolism and glycolysis may overcome such resistance mechanisms. Such strategies are, however, expected to generate general toxicity. The potential for compensatory glycolysis may in fact be limited in solid tumors due to the low availability of glucose (Hirayama et al., 2009; Yaromina et al., 2009). Cells in solid tumor cells may therefore be more vulnerable to inhibition of OXPHOS compared to normal cells situated in well vascularized regions (Zhang et al., 2016; Zhang et al., 2014). The increasing understanding of the important role of mitochondrial energy production for tumor cell viability has resulted in considerable interest in this process as a target for development of antineoplastic agents (Emmings et al., 2019; Pavlova and Thompson, 2016; Weinberg and Chandel, 2015; Zhang et al., 2014; Zong et al., 2016).

There are several unanswered questions with regard to the optimal characteristics for cancer drugs acting on mitochondrial energetics. Which degree of inhibition of energy production will inhibit tumor cell survival without resulting in general toxicity? Which part of the respiratory chain (or other components) should be inhibited for optimal anti-neoplastic effects? Furthermore, factors related to the pharmacological properties of drugs such as their lipophilicity and ability to penetrate into solid tumors are likely to be important (Fayad et al., 2011; Minchinton and Tannock, 2006).

The first reports of mitochondrial inhibitors showing antitumor effects used the cell penetrating dye Rhodamine 123 (Bernal et al., 1983). Although the mechanism of tumor cell toxicity by Rhodamine 123 was shown to be the impairment of ATP synthesis (Bernal et al., 1983) the larger degree of retention of the dye in carcinoma cells compared to normal cells was believed as an important factor for tumor-specificity (Lampidis et al., 1983). It is possible that the specific mechanism of mitochondrial inhibition may not be critical. Inhibitors of the electron transport chain, inhibitors of the F1F0-ATPase and mitochondrial uncouplers were all reported to be effective in inducing cell death in the core regions of multicellular tumor spheroids (Wenzel et al., 2014). We here tested a number of different mitochondrial inhibitors belonging to different mechanistic classes and found that the complex I inhibitor IACS-010759 showed the best activity in our *in vitro* models. IACS-010759 has been shown to induce apoptosis of OXPHOS-dependent brain cancer and acute myeloid leukemia cells (Molina et al., 2018), lung cancer cell lines and tumors (Lissanu Deribe et al., 2018) and mantle cell lymphoma cells (Zhang et al., 2019). An important factor to be considered in clinical development is the occurrence of acidosis. Lactate levels were increased in patient plasma consistent with inhibition of Complex I, but acidosis was not induced (NCT03291938). IACS-010759 is being evaluated in phase I clinical trials in relapsed/refractory AML and solid tumors (NCT02882321, NCT03291938). It is unclear why IACS-010759 showed selectivity to ARID1A-deficient ovarian cancer cells whereas other inhibitors that were tested showed less or no such selectivity. Further understanding of this question is necessary in order to develop improved compounds.

The field of mitochondrial bioenergetics as a target for antineoplastic agents is expanding rapidly (Frattaruolo et al., 2020; Oliveira et al., 2021; Sica et al., 2020). The identification of metabolic vulnerabilities will be important in expanding this field and is expected to broaden the arsenal of clinically available therapeutics for treatment of advanced disease.

### Limitations of the Study

A potential limitation of the study is that we did not test this approach in a preclinical model of re-constituted ARID1A (i.e TOV-21G cells ectopically expressing ARID1A). However, as per personal communication from Dr. Rugang Zhang (Wistar Institute), re-expression of ARID1A in TOV-21G induces cell senescence via p21 expression rendering this model not suitable for preclinical testing. Further studies are warranted to test this treatment in patient-derived xenograft model of ARID1A-mutated OCCC.

## Materials and methods

### Chemicals and antibodies

IACS-010759 (S8731) was obtained from Selleck Chemicals (Houston, USA). Antibodies used were anti-ARID1A (Santa Cruz catalogue number sc-32761), anti-Actin (Santa Cruz catalogue number sc-8432), anti-Cmyc (Abcam rb anti-cmyc; ab32072), anti-Tubulin (Santa Cruz catalogue number sc-5286), anti-Tom20 (Santa Cruz catalogue number sc-17764). CPI-613 (s2776), Tigecycline (s1403), Sodium Dichloroacetate (DCA, s8615), Onc212 (s8673), Dapagliflozin (s1548), Itraconazole (s2467), Riluzole (s1614), Bay87-2243 (S7309) were all obtained from Selleck Chemicals (Houston, USA).

### Cell culture

ES2, RMG1 and TOV-21G human clear cell carcinoma were maintained in DMEM (Dulbecco’s Modified Eagle Medium, Catalog number 12320032) with 10% FBS and 1% penicillin.

### siRNA

ES2 or RMG1 cells were plated and left to attach for 24h in 6-well plates. Then cells were transfected with ON-TARGET plus human non-targeting control pool (catalog number D-001810-10-05) or ON-TARGET plus human ARID1A siRNAs (accessions Hit: <u>NM_006015, NM_139135</u>, catalog number L-017263-00-0005) followed the general protocol provided by Dharmacon (Cambridge, United Kingdom) for 48hs. Different post-transfection experiments were then performed.

### Western blot

Cells were collected and lysed in ice-cold lysis buffer and run on 4-15% SDS-PAGE gels. Proteins were then transferred to nitrocellulose membranes and incubated at room temperature in 5% milk in 1xPBST buffer for 1 h. Membranes were then incubated overnight with primary antibodies at a dilution of 1:1,000 in 1xPBST buffer. Next day, membranes were washed with 1xPBST buffer and incubated with anti-rabbit or anti-mouse secondary antibodies at a 1:5000 dilution for 1 h. Immunoreactive bands were detected by FluorChem Fluorescent Western Imaging System from ProteinSimple (California, USA).

### RNA-SEQ experiments

ES2 human clear cell carcinoma cells present (wild type) or absent (siRNA) ARID1A were collected and extracted using an RNeasy Plus Micro Kit (Qiagen). Barcoded TruSeq RNA v2 libraries (Illumina) were created, and libraries were sequenced on a HiSeq 2500 (Illumina) as paired-end 100 bp. STAR version 2.4.0d was used to align the RNA-seq reads to the human genome reference build GRCh37 (hg19). The Ensembl gene annotation version 75 was provided as gene transfer format for exon junction support. For each sample, reads were assigned to genes and summarized using featureCounts (subread package version 1.4.5-p1). Raw read counts were read into R (version 3.2.0) and subjected to normalization by the trimmed mean of M-values normalization method implemented in the R/bioconductor edgeR package and variance normalized using voom from the R/bioconductor limma package. All genes with at least one count per million (CPM) mapped reads in at least two samples were analyzed further. Differential gene expression was performed using the R/bioconductor limma package. RNA-seq data can be found under the GEO accession no. GSE117614.

### Measurements of oxygen consumption

The Seahorse XF analyser was used as recommended by the manufacturer (Seahorse Bioscience, North Billerica, MA, USA). Seahorse Cell Mito Stress Test Kit (Agilent Technologies) was used to measure mitochondrial OxPHOS. Briefly, 60,000 cells per condition were plated in 100 μL culture medium in XF24-well cell plates with blank control wells. Before the measurements of OCR the medium was replaced with 500 μl Seahorse assay media (1 mM pyruvate and 2 mM glutamine) at 37 °C without CO2 for 1 h. ATP linked respiration OCR were calculated as average values of basal OCR minus average postoligomycin OCR values. Maximum mitochondrial respiration capacity was calculated as average maximal OCR values minus average postoligomycin OCR values. ATP linked respiration OCR were calculated as average values of basal OCR minus average postoligomycin OCR values. For all Seahorse related experiment, equal cell number per condition was confirmed post-analysis in cell lysates via amido black staining.

### ATP quantification assay

20,000 cells per well were plated one day before the assay in 96-well plates. Next day, glucose metabolic inhibitor 2DG (10 mM or 20 mM) was added 15 min before cell lysis and ATP determination using the ATP Bioluminescence Assay Kit HS II (Sigma catalogue number 11699709001) and analyzed on an Infinite M200 PRO Luminometer (Tecan).

### Confocal microscopy

For mitochondrial membrane potential assay, cells were stained with MitoTracker Deep Red dye (Cell signaling catalogue number 8778) according to the manufacturer’s instructions. Briefly, 500nM MitoTracker Deep Red dye solution was added into cells and incubated for 30 min at 37°C. After staining, medium with MitoTracker solution was removed and cells were then fixed in ice-cold, 100% methanol for 15 min at −20°C and rinse 3 times with PBS for 5 min. Cells were fixed and mounted with DAPI staining dye Hoechst 33342 (ThermoScientific catalogue number H3570). Images were acquired with an Olympus BX upright microscope equipped with a Fluoview 1000 confocal scan head with a 60× oil immersion objective (Olympus). Image processing and fluorescence intensity quantification was performed using Icy open platform software.

### Mitochondrial morphology

For studies of mitochondrial morphology cells were fixed in cold methanol for 5 minutes. After blocking with 5% BSA in PBST, cells were stained with anti-Tom20 followed by Alexa Fluor 594-conjugated secondary antibodies and analyzed via confocal fluorescence microscopy. Images were taken with an Olympus BX2 upright microscope. A UPlanApo N 60X/1.42 NA objective was used. For Alexa Fluor a 594 nm laser was used for excitation and emission collected between 560 and 660 nm. Images were taken with sequential excitation. Further analysis of images was done using the Fiji software mitochondrial morphology plugin and calculated as previously described (Bosch and Calvo, 2019). 3D reconstruction was performed using software Imaris, under the function of volume and 3D view. Procedures in detail could be obtained from the website of Imaris (<u>Imaris for Cell Biologists - Imaris - Oxford Instruments (oxinst.com)</u>.

### Plasmid transfection

TOV-21G cells were seeded in Dulbecco’s Modified Eagle Medium one day before the transfection in 6-well plates and incubated at 37°C, 5% CO2. Transfection was performed with control vector (pcDNA/GW-40/LacZ) or 4 μg pARID1A (Addgene catalogue number 39311) using FuGENE 6 Transfection Reagent (Promega) according to manufacturer protocol. Further experiments were performed after 48 h transfection.

### Generation of spheroids

Spheroids were prepared using a modification of our previously described method^17^. A cell suspension containing 5,000 cells/200 μL was added to each well of 96 well plates. Wells were overfilled by adding 180 ml media to acquire a convex surface curvature and plates inverted in order to allow cells to sediment to the liquid-air interface. Plates were returned to normal position after 24 h incubation, excess media removed by aspiration, and incubated for 4 days before drug exposure. Imagines for each spheroid were taken by microscopy camera and diameters were then measured and volume of spheroids were calculated by equation Volume=4/3*3.14*R^3^(R=radius).

### Crystal violet staining

Cells were fixed with ice cold methanol for 15 minutes then methanol was removed, and crystal violet staining solution (0.05% (w/v)) was added in each well for 30 minutes (the volume should cover the whole surface of each well). Then staining solution was removed and plates were washed with water three times.

### Colony formation assay

For colony formation assay 10,000 or 20,000 cells were plated in 10 cm plates and treated with the indicated drugs at the indicated concentration over a period of 10 days. Medium was changed every 3 three days. At the end of the 10 days period cells were stained with crystal violet and colony were visualized and quantified using imageJ.

### Preclinical testing of IACS-010759

10 million TOV21G cells infected with GFP lentivirus were injected intraperitoneally into athymic nude female mice at 6 weeks of age (Charles River Laboratories, Crl:NU(NCr)-Foxn1nu). Control mice were treated with vehicle (8% DMSO) and the experimental mice were treated with IACS-010759 (5 and 10 mg/kg) orally daily on a 5-on 2-off schedule till the time the mice were either dead or euthanized due to signs of distress or moribundity followed by collection of ascitic fluid/blood from the peritoneal cavity. IACS-010759 was dissolved in DMSO and administered in milk. Mice were weighed and imaged once a week using IVIS. The IACS-010759 administered was started from the same day TOV21G cells were injected. The animal experiments were approved by the Institutional Animal Care and Use Committee at the University of Minnesota (Protocol ID:1807-36121A).

### Statistical analysis

Statistics were calculated with GraphPad Prism (v8.0). Data are expressed as the mean ± SD. Differences between groups were determined by Student’s t-test. P values < 0.05 were considered statistically significant (*P < 0.05, **P < 0.01).

