## Supplementary figures and images for "Targeting Mitochondrial Metabolism in Clear Cell Carcinoma of the Ovaries"

### Supplementary Figure 1

Supplementary Figure 1

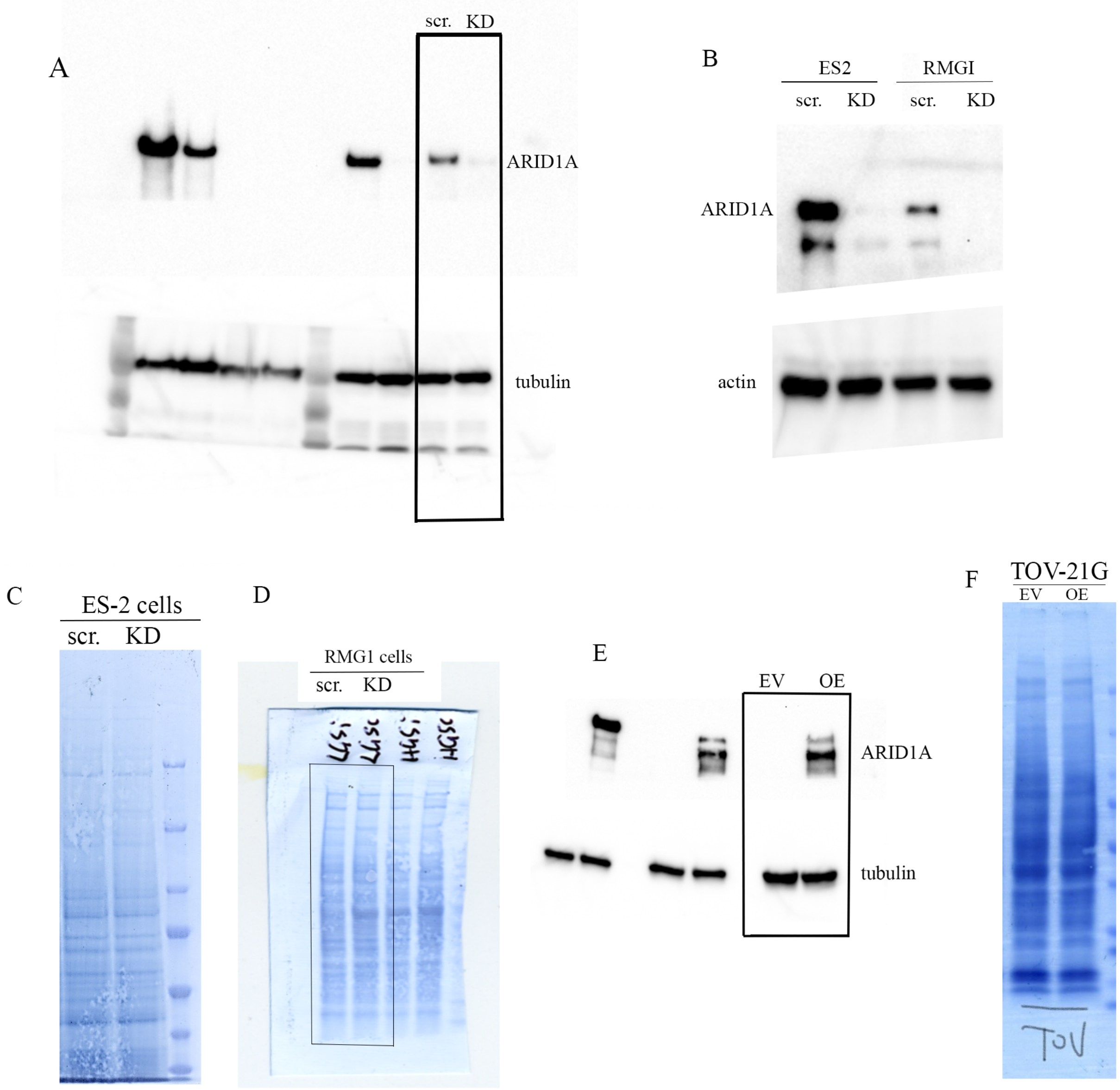

### Supplementary Figure 2

#### Supplementary Figure 2

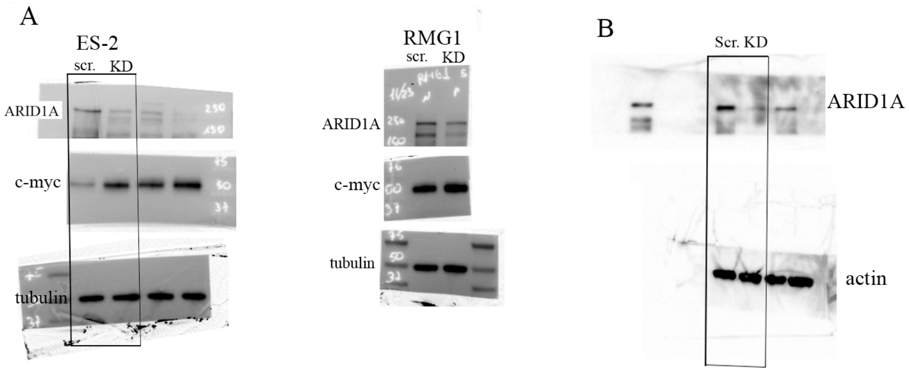

### Supplementary Figure 3

Supplementary Figure 3

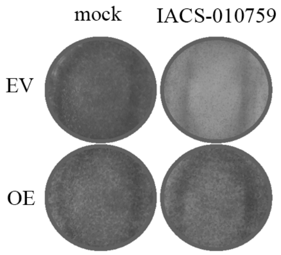
