## Supplementary Table 1 for "Targeting Mitochondrial Metabolism in Clear Cell Carcinoma of the Ovaries"

| Feature ID | ES2 Scramble | | | | |
|---|---|---|---|---|---|
| | Expression value 1 | Expression value 2 | Expression value 3 | Average (Expression value ) | Expression value 1 |
| A1BG | 89.00 | 76.00 | 87.00 | 84.00 | 69.00 |
| A1BG-AS1 | 163.00 | 157.00 | 153.00 | 157.67 | 134.00 |
| A2M | 6.00 | 6.00 | 4.00 | 5.33 | 2.00 |
| A2M-AS1 | 9.00 | 14.00 | 11.00 | 11.33 | 16.00 |
| AAAS | 1566.00 | 1587.00 | 1706.00 | 1619.67 | 1866.00 |
| AACS | 554.00 | 584.00 | 598.00 | 578.67 | 481.00 |
| AACSP1 | 82.00 | 68.00 | 107.00 | 85.67 | 92.00 |
| AADAC | 6.00 | 4.00 | 1.00 | 3.67 | 2.00 |
| AADAT | 383.00 | 310.00 | 397.00 | 363.33 | 465.00 |
| AAED1 | 107.00 | 108.00 | 115.00 | 110.00 | 143.00 |
| AAGAB | 1147.00 | 1109.00 | 1211.00 | 1155.67 | 1622.00 |
| AAK1 | 1773.00 | 1610.00 | 1931.00 | 1771.33 | 1519.00 |
| AAMDC | 48.00 | 53.00 | 59.00 | 53.33 | 79.00 |
| AAMP | 2969.00 | 3063.00 | 3196.00 | 3076.00 | 3491.00 |
| AANAT | 5.00 | 7.00 | 5.00 | 5.67 | 5.00 |
| AAR2 | 830.00 | 827.00 | 900.00 | 852.33 | 878.00 |
| AARD | 15.00 | 10.00 | 21.00 | 15.33 | 20.00 |
| AARS | 10691.00 | 10101.00 | 11170.00 | 10654.00 | 10259.00 |
| AARS2 | 1364.00 | 1380.00 | 1406.00 | 1383.33 | 1358.00 |
| AARSD1 | 7.00 | 17.00 | 11.00 | 11.67 | 20.00 |
| AASDH | 427.00 | 405.00 | 474.00 | 435.33 | 404.00 |
| AASDHPPT | 710.00 | 686.00 | 832.00 | 742.67 | 890.00 |
| AASS | 1932.00 | 1959.00 | 2217.00 | 2036.00 | 1919.00 |
| AATK | 4.00 | 3.00 | 4.00 | 3.67 | 1.00 |
| ABAT | 7.00 | 7.00 | 8.00 | 7.33 | 2.00 |
| ABCA1 | 851.00 | 825.00 | 994.00 | 890.00 | 517.00 |
| ABCA10 | 10.00 | 11.00 | 5.00 | 8.67 | 22.00 |
| ABCA11P | 137.00 | 105.00 | 160.00 | 134.00 | 178.00 |
| ABCA13 | 4.00 | 7.00 | 7.00 | 6.00 | 8.00 |
| ABCA2 | 1793.00 | 1838.00 | 2081.00 | 1904.00 | 1902.00 |
| ABCA3 | 8.00 | 16.00 | 22.00 | 15.33 | 17.00 |
| ABCA4 | 4.00 | 6.00 | 3.00 | 4.33 | 4.00 |
| ABCA5 | 252.00 | 253.00 | 281.00 | 262.00 | 264.00 |
| ABCA7 | 302.00 | 342.00 | 330.00 | 324.67 | 257.00 |
| ABCA8 | 4.00 | 3.00 | 1.00 | 2.67 | 3.00 |
| ABCB1 | 193.00 | 155.00 | 158.00 | 168.67 | 185.00 |
| ABCB10 | 437.00 | 405.00 | 461.00 | 434.33 | 494.00 |
| ABCB4 | 188.00 | 174.00 | 181.00 | 181.00 | 128.00 |
| ABCB6 | 646.00 | 724.00 | 693.00 | 687.67 | 645.00 |
| ABCB7 | 727.00 | 668.00 | 770.00 | 721.67 | 757.00 |
| ABCB8 | 1736.00 | 1753.00 | 1764.00 | 1751.00 | 1852.00 |
| ABCB9 | 384.00 | 335.00 | 306.00 | 341.67 | 244.00 |
| ABCC1 | 364.00 | 355.00 | 367.00 | 362.00 | 339.00 |

| | | | | | |
|---|---|---|---|---|---|
| ABCC10 | 1155.00 | 1104.00 | 1130.00 | 1129.67 | 1005.00 |
| ABCC11 | 8.00 | 4.00 | 9.00 | 7.00 | 4.00 |
| ABCC12 | 4.00 | 2.00 | 0.00 | 2.00 | 2.00 |
| ABCC2 | 817.00 | 829.00 | 833.00 | 826.33 | 861.00 |
| ABCC3 | 699.00 | 774.00 | 931.00 | 801.33 | 166.00 |
| ABCC4 | 1761.00 | 1734.00 | 2020.00 | 1838.33 | 1863.00 |
| ABCC5 | 3057.00 | 2982.00 | 3263.00 | 3100.67 | 2678.00 |
| ABCC9 | 447.00 | 383.00 | 408.00 | 412.67 | 449.00 |
| ABCD1 | 218.00 | 175.00 | 195.00 | 196.00 | 191.00 |
| ABCD3 | 981.00 | 910.00 | 1048.00 | 979.67 | 1097.00 |
| ABCD4 | 1344.00 | 1300.00 | 1314.00 | 1319.33 | 1194.00 |
| ABCE1 | 2943.00 | 2798.00 | 3184.00 | 2975.00 | 3441.00 |
| ABCF1 | 4.00 | 3.00 | 2.00 | 3.00 | 0.00 |
| ABCF2 | 3563.00 | 3577.00 | 3726.00 | 3622.00 | 3920.00 |
| ABCF3 | 1152.00 | 1220.00 | 1138.00 | 1170.00 | 1260.00 |
| ABCG2 | 306.00 | 227.00 | 240.00 | 257.67 | 236.00 |
| ABHD10 | 852.00 | 702.00 | 808.00 | 787.33 | 886.00 |
| ABHD11 | 263.00 | 267.00 | 271.00 | 267.00 | 408.00 |
| ABHD12 | 1178.00 | 1255.00 | 1203.00 | 1212.00 | 1151.00 |
| ABHD13 | 363.00 | 328.00 | 388.00 | 359.67 | 439.00 |
| ABHD14A | 178.00 | 200.00 | 175.00 | 184.33 | 193.00 |
| ABHD14B | 3099.00 | 2884.00 | 3147.00 | 3043.33 | 3547.00 |
| ABHD15 | 228.00 | 243.00 | 253.00 | 241.33 | 251.00 |
| ABHD17A | 392.00 | 470.00 | 528.00 | 463.33 | 541.00 |
| ABHD17B | 194.00 | 159.00 | 213.00 | 188.67 | 201.00 |
| ABHD17C | 254.00 | 245.00 | 265.00 | 254.67 | 297.00 |
| ABHD2 | 4449.00 | 4138.00 | 4289.00 | 4292.00 | 3014.00 |
| ABHD3 | 210.00 | 215.00 | 209.00 | 211.33 | 271.00 |
| ABHD4 | 908.00 | 848.00 | 998.00 | 918.00 | 675.00 |
| ABHD5 | 1089.00 | 1064.00 | 1199.00 | 1117.33 | 1016.00 |
| ABHD6 | 271.00 | 243.00 | 229.00 | 247.67 | 333.00 |
| ABHD8 | 123.00 | 125.00 | 173.00 | 140.33 | 132.00 |
| ABI1 | 1361.00 | 1205.00 | 1515.00 | 1360.33 | 1548.00 |
| ABI2 | 2292.00 | 2327.00 | 2493.00 | 2370.67 | 2540.00 |
| ABI3 | 72.00 | 75.00 | 81.00 | 76.00 | 70.00 |
| ABI3BP | 421.00 | 378.00 | 459.00 | 419.33 | 293.00 |
| ABL1 | 4897.00 | 4562.00 | 4902.00 | 4787.00 | 6225.00 |
| ABL2 | 6297.00 | 5713.00 | 6780.00 | 6263.33 | 6395.00 |
| ABLIM1 | 11.00 | 19.00 | 15.00 | 15.00 | 20.00 |
| ABLIM2 | 4.00 | 4.00 | 5.00 | 4.33 | 11.00 |
| ABLIM3 | 2225.00 | 2281.00 | 2362.00 | 2289.33 | 1396.00 |
| ABRACL | 958.00 | 902.00 | 970.00 | 943.33 | 1161.00 |
| ABT1 | 699.00 | 730.00 | 795.00 | 741.33 | 945.00 |
| ABTB1 | 1013.00 | 1110.00 | 1154.00 | 1092.33 | 1136.00 |
| ABTB2 | 161.00 | 154.00 | 187.00 | 167.33 | 102.00 |
| ACAA1 | 747.00 | 817.00 | 801.00 | 788.33 | 903.00 |
| ACAA2 | 176.00 | 157.00 | 183.00 | 172.00 | 314.00 |
| ACACA | 26.00 | 19.00 | 19.00 | 21.33 | 16.00 |

| | | | | | |
|---|---|---|---|---|---|
| ACACB | 30.00 | 27.00 | 27.00 | 28.00 | 30.00 |
| ACAD10 | 1311.00 | 1297.00 | 1286.00 | 1298.00 | 1246.00 |
| ACAD11 | 182.00 | 238.00 | 254.00 | 224.67 | 194.00 |
| ACAD8 | 465.00 | 520.00 | 571.00 | 518.67 | 452.00 |
| ACAD9 | 804.00 | 868.00 | 877.00 | 849.67 | 905.00 |
| ACADL | 14.00 | 7.00 | 6.00 | 9.00 | 11.00 |
| ACADM | 860.00 | 758.00 | 864.00 | 827.33 | 1119.00 |
| ACADS | 104.00 | 122.00 | 109.00 | 111.67 | 124.00 |
| ACADSB | 612.00 | 557.00 | 673.00 | 614.00 | 587.00 |
| ACADVL | 6792.00 | 6583.00 | 7372.00 | 6915.67 | 7301.00 |
| ACAP1 | 9.00 | 7.00 | 4.00 | 6.67 | 11.00 |
| ACAP2 | 1416.00 | 1189.00 | 1506.00 | 1370.33 | 1621.00 |
| ACAP3 | 813.00 | 897.00 | 916.00 | 875.33 | 755.00 |
| ACAT1 | 1207.00 | 1096.00 | 1186.00 | 1163.00 | 1643.00 |
| ACAT2 | 298.00 | 266.00 | 304.00 | 289.33 | 398.00 |
| ACBD3 | 922.00 | 896.00 | 969.00 | 929.00 | 942.00 |
| ACBD4 | 214.00 | 205.00 | 226.00 | 215.00 | 225.00 |
| ACBD5 | 794.00 | 750.00 | 857.00 | 800.33 | 909.00 |
| ACBD6 | 754.00 | 759.00 | 836.00 | 783.00 | 843.00 |
| ACBD7 | 35.00 | 48.00 | 45.00 | 42.67 | 63.00 |
| ACCS | 377.00 | 374.00 | 365.00 | 372.00 | 433.00 |
| ACD | 601.00 | 552.00 | 556.00 | 569.67 | 700.00 |
| ACER2 | 21.00 | 10.00 | 27.00 | 19.33 | 38.00 |
| ACER3 | 782.00 | 836.00 | 826.00 | 814.67 | 824.00 |
| ACIN1 | 6503.00 | 6554.00 | 6951.00 | 6669.33 | 6151.00 |
| ACLY | 6580.00 | 6019.00 | 6704.00 | 6434.33 | 6542.00 |
| ACO1 | 2611.00 | 2348.00 | 2849.00 | 2602.67 | 2402.00 |
| ACO2 | 2181.00 | 2296.00 | 2165.00 | 2214.00 | 2543.00 |
| ACOT1 | 7.00 | 9.00 | 8.00 | 8.00 | 18.00 |
| ACOT13 | 469.00 | 478.00 | 535.00 | 494.00 | 780.00 |
| ACOT2 | 119.00 | 140.00 | 153.00 | 137.33 | 161.00 |
| ACOT7 | 1062.00 | 1070.00 | 1092.00 | 1074.67 | 1645.00 |
| ACOT8 | 420.00 | 396.00 | 392.00 | 402.67 | 448.00 |
| ACOT9 | 589.00 | 561.00 | 589.00 | 579.67 | 659.00 |
| ACOX1 | 559.00 | 516.00 | 587.00 | 554.00 | 550.00 |
| ACOX2 | 29.00 | 30.00 | 35.00 | 31.33 | 82.00 |
| ACOX3 | 317.00 | 272.00 | 301.00 | 296.67 | 290.00 |
| ACP1 | 1556.00 | 1425.00 | 1837.00 | 1606.00 | 2087.00 |
| ACP2 | 764.00 | 793.00 | 703.00 | 753.33 | 727.00 |
| ACP5 | 261.00 | 313.00 | 248.00 | 274.00 | 257.00 |
| ACP6 | 409.00 | 428.00 | 443.00 | 426.67 | 408.00 |
| ACRBP | 24.00 | 25.00 | 33.00 | 27.33 | 18.00 |
| ACRC | 199.00 | 205.00 | 219.00 | 207.67 | 290.00 |
| ACRV1 | 21.00 | 16.00 | 13.00 | 16.67 | 14.00 |
| ACSF2 | 406.00 | 396.00 | 387.00 | 396.33 | 304.00 |
| ACSF3 | 669.00 | 672.00 | 676.00 | 672.33 | 714.00 |
| ACSL1 | 487.00 | 523.00 | 512.00 | 507.33 | 478.00 |
| ACSL3 | 3659.00 | 3392.00 | 3566.00 | 3539.00 | 3691.00 |

| | | | | | |
|---|---|---|---|---|---|
| ACSL4 | 4029.00 | 3467.00 | 4422.00 | 3972.67 | 4039.00 |
| ACSL5 | 678.00 | 627.00 | 679.00 | 661.33 | 458.00 |
| ACSM3 | 5.00 | 4.00 | 13.00 | 7.33 | 8.00 |
| ACSS2 | 1004.00 | 958.00 | 1051.00 | 1004.33 | 707.00 |
| ACTA2 | 46.00 | 47.00 | 63.00 | 52.00 | 65.00 |
| ACTA2-AS1 | 11.00 | 5.00 | 15.00 | 10.33 | 6.00 |
| ACTB | 44106.00 | 42682.00 | 45460.00 | 44082.67 | 60140.00 |
| ACTG1 | 38194.00 | 36794.00 | 40707.00 | 38565.00 | 46347.00 |
| ACTG1P17 | 33.00 | 13.00 | 24.00 | 23.33 | 36.00 |
| ACTG1P20 | 60.00 | 67.00 | 62.00 | 63.00 | 44.00 |
| ACTG1P4 | 25.00 | 21.00 | 24.00 | 23.33 | 18.00 |
| ACTL10 | 26.00 | 34.00 | 22.00 | 27.33 | 28.00 |
| ACTL6A | 1647.00 | 1550.00 | 1753.00 | 1650.00 | 2190.00 |
| ACTN1 | 7550.00 | 7167.00 | 7514.00 | 7410.33 | 6664.00 |
| ACTN1-AS1 | 4.00 | 2.00 | 0.00 | 2.00 | 1.00 |
| ACTN4 | 16481.00 | 15545.00 | 16400.00 | 16142.00 | 16248.00 |
| ACTR10 | 969.00 | 1001.00 | 1086.00 | 1018.67 | 1267.00 |
| ACTR1A | 2586.00 | 2572.00 | 2640.00 | 2599.33 | 2726.00 |
| ACTR1B | 1255.00 | 1281.00 | 1247.00 | 1261.00 | 1383.00 |
| ACTR2 | 5665.00 | 4986.00 | 5659.00 | 5436.67 | 6875.00 |
| ACTR3 | 5797.00 | 5159.00 | 5781.00 | 5579.00 | 6436.00 |
| ACTR3B | 385.00 | 370.00 | 440.00 | 398.33 | 510.00 |
| ACTR3C | 4.00 | 9.00 | 6.00 | 6.33 | 7.00 |
| ACTR5 | 439.00 | 443.00 | 454.00 | 445.33 | 429.00 |
| ACTR6 | 487.00 | 463.00 | 509.00 | 486.33 | 600.00 |
| ACTR8 | 570.00 | 576.00 | 612.00 | 586.00 | 601.00 |
| ACTRT3 | 100.00 | 75.00 | 79.00 | 84.67 | 88.00 |
| ACVR1 | 1420.00 | 1324.00 | 1501.00 | 1415.00 | 1524.00 |
| ACVR1B | 911.00 | 968.00 | 909.00 | 929.33 | 913.00 |
| ACVR1C | 25.00 | 18.00 | 30.00 | 24.33 | 17.00 |
| ACVR2A | 323.00 | 318.00 | 315.00 | 318.67 | 345.00 |
| ACVR2B | 506.00 | 499.00 | 593.00 | 532.67 | 585.00 |
| ACVR2B-AS1 | 29.00 | 21.00 | 19.00 | 23.00 | 22.00 |
| ACVRL1 | 6.00 | 4.00 | 0.00 | 3.33 | 5.00 |
| ACY1 | 55.00 | 46.00 | 51.00 | 50.67 | 89.00 |
| ACYP1 | 186.00 | 258.00 | 247.00 | 230.33 | 341.00 |
| ACYP2 | 44.00 | 44.00 | 40.00 | 42.67 | 59.00 |
| ADA | 142.00 | 123.00 | 143.00 | 136.00 | 191.00 |
| ADAL | 391.00 | 427.00 | 408.00 | 408.67 | 369.00 |
| ADAM10 | 2859.00 | 2648.00 | 2990.00 | 2832.33 | 2977.00 |
| ADAM11 | 60.00 | 70.00 | 61.00 | 63.67 | 66.00 |
| ADAM12 | 5016.00 | 4677.00 | 4792.00 | 4828.33 | 3050.00 |
| ADAM15 | 5388.00 | 5480.00 | 5665.00 | 5511.00 | 3554.00 |
| ADAM17 | 1535.00 | 1465.00 | 1605.00 | 1535.00 | 1411.00 |
| ADAM19 | 860.00 | 922.00 | 916.00 | 899.33 | 662.00 |
| ADAM1A | 307.00 | 302.00 | 311.00 | 306.67 | 358.00 |
| ADAM20 | 7.00 | 3.00 | 12.00 | 7.33 | 0.00 |
| ADAM20P1 | 37.00 | 29.00 | 55.00 | 40.33 | 39.00 |

| | | | | | |
|---|---|---|---|---|---|
| ADAM21 | 18.00 | 16.00 | 13.00 | 15.67 | 16.00 |
| ADAM22 | 361.00 | 404.00 | 417.00 | 394.00 | 423.00 |
| ADAM23 | 841.00 | 835.00 | 890.00 | 855.33 | 918.00 |
| ADAM28 | 109.00 | 90.00 | 108.00 | 102.33 | 73.00 |
| ADAM32 | 17.00 | 6.00 | 4.00 | 9.00 | 6.00 |
| ADAM8 | 53.00 | 69.00 | 67.00 | 63.00 | 59.00 |
| ADAM9 | 5697.00 | 5365.00 | 5938.00 | 5666.67 | 4709.00 |
| ADAMTS1 | 18464.00 | 18480.00 | 21072.00 | 19338.67 | 14234.00 |
| ADAMTS12 | 1145.00 | 997.00 | 1126.00 | 1089.33 | 729.00 |
| ADAMTS13 | 262.00 | 252.00 | 269.00 | 261.00 | 269.00 |
| ADAMTS15 | 21.00 | 19.00 | 18.00 | 19.33 | 5.00 |
| ADAMTS16 | 411.00 | 341.00 | 394.00 | 382.00 | 338.00 |
| ADAMTS17 | 96.00 | 96.00 | 88.00 | 93.33 | 67.00 |
| ADAMTS19 | 16.00 | 15.00 | 25.00 | 18.67 | 32.00 |
| ADAMTS2 | 3082.00 | 2917.00 | 3185.00 | 3061.33 | 2751.00 |
| ADAMTS20 | 4.00 | 1.00 | 5.00 | 3.33 | 1.00 |
| ADAMTS3 | 209.00 | 213.00 | 257.00 | 226.33 | 185.00 |
| ADAMTS4 | 211.00 | 189.00 | 171.00 | 190.33 | 141.00 |
| ADAMTS5 | 185.00 | 145.00 | 151.00 | 160.33 | 185.00 |
| ADAMTS6 | 2583.00 | 2548.00 | 2887.00 | 2672.67 | 2928.00 |
| ADAMTS9 | 68.00 | 46.00 | 39.00 | 51.00 | 53.00 |
| ADAMTS9-AS2 | 10.00 | 12.00 | 10.00 | 10.67 | 7.00 |
| ADAMTSL1 | 9595.00 | 9557.00 | 9833.00 | 9661.67 | 8692.00 |
| ADAMTSL3 | 38.00 | 31.00 | 31.00 | 33.33 | 24.00 |
| ADAMTSL4 | 134.00 | 137.00 | 139.00 | 136.67 | 88.00 |
| ADAMTSL4-AS1 | 6.00 | 9.00 | 4.00 | 6.33 | 10.00 |
| ADAMTSL5 | 224.00 | 239.00 | 276.00 | 246.33 | 223.00 |
| ADAP1 | 14.00 | 12.00 | 7.00 | 11.00 | 9.00 |
| ADAP2 | 13.00 | 5.00 | 11.00 | 9.67 | 9.00 |
| ADAR | 5809.00 | 5227.00 | 5815.00 | 5617.00 | 5271.00 |
| ADARB1 | 1918.00 | 1879.00 | 1989.00 | 1928.67 | 1647.00 |
| ADAT1 | 596.00 | 579.00 | 564.00 | 579.67 | 416.00 |
| ADAT2 | 993.00 | 1004.00 | 1025.00 | 1007.33 | 1170.00 |
| ADAT3 | 77.00 | 68.00 | 71.00 | 72.00 | 62.00 |
| ADCK1 | 237.00 | 196.00 | 285.00 | 239.33 | 292.00 |
| ADCK2 | 860.00 | 848.00 | 893.00 | 867.00 | 793.00 |
| ADCK3 | 813.00 | 821.00 | 894.00 | 842.67 | 982.00 |
| ADCK4 | 263.00 | 299.00 | 357.00 | 306.33 | 297.00 |
| ADCK5 | 309.00 | 366.00 | 331.00 | 335.33 | 347.00 |
| ADCY1 | 156.00 | 134.00 | 148.00 | 146.00 | 145.00 |
| ADCY10P1 | 608.00 | 597.00 | 718.00 | 641.00 | 496.00 |
| ADCY3 | 1937.00 | 2041.00 | 2107.00 | 2028.33 | 2051.00 |
| ADCY6 | 1831.00 | 1768.00 | 1901.00 | 1833.33 | 1498.00 |
| ADCY7 | 1566.00 | 1478.00 | 1541.00 | 1528.33 | 1666.00 |
| ADCY8 | 4.00 | 3.00 | 4.00 | 3.67 | 6.00 |
| ADCY9 | 554.00 | 602.00 | 583.00 | 579.67 | 532.00 |
| ADD1 | 7705.00 | 7631.00 | 8008.00 | 7781.33 | 7392.00 |
| ADD3 | 615.00 | 532.00 | 619.00 | 588.67 | 549.00 |

| | | | | | |
|---|---|---|---|---|---|
| ADGRA2 | 488.00 | 523.00 | 509.00 | 506.67 | 349.00 |
| ADGRA3 | 935.00 | 906.00 | 1038.00 | 959.67 | 899.00 |
| ADGRB2 | 231.00 | 171.00 | 202.00 | 201.33 | 176.00 |
| ADGRB3 | 134.00 | 141.00 | 117.00 | 130.67 | 128.00 |
| ADGRE1 | 6.00 | 3.00 | 3.00 | 4.00 | 2.00 |
| ADGRE5 | 1282.00 | 1340.00 | 1290.00 | 1304.00 | 1062.00 |
| ADGRF3 | 18.00 | 15.00 | 16.00 | 16.33 | 9.00 |
| ADGRF4 | 142.00 | 152.00 | 131.00 | 141.67 | 44.00 |
| ADGRF5 | 33.00 | 35.00 | 40.00 | 36.00 | 10.00 |
| ADGRG1 | 98.00 | 68.00 | 82.00 | 82.67 | 48.00 |
| ADGRG4 | 24.00 | 28.00 | 25.00 | 25.67 | 8.00 |
| ADGRG6 | 116.00 | 79.00 | 115.00 | 103.33 | 75.00 |
| ADGRL1 | 184.00 | 188.00 | 210.00 | 194.00 | 234.00 |
| ADGRL2 | 1478.00 | 1229.00 | 1473.00 | 1393.33 | 1365.00 |
| ADGRL3 | 4.00 | 0.00 | 0.00 | 1.33 | 2.00 |
| ADGRL4 | 4327.00 | 4042.00 | 4142.00 | 4170.33 | 3464.00 |
| ADGRV1 | 113.00 | 92.00 | 131.00 | 112.00 | 99.00 |
| ADH5 | 1311.00 | 1190.00 | 1346.00 | 1282.33 | 1483.00 |
| ADHFE1 | 84.00 | 104.00 | 103.00 | 97.00 | 74.00 |
| ADI1 | 352.00 | 324.00 | 358.00 | 344.67 | 384.00 |
| ADIPOR1 | 1686.00 | 1575.00 | 1634.00 | 1631.67 | 1985.00 |
| ADIPOR2 | 2796.00 | 2596.00 | 2758.00 | 2716.67 | 2730.00 |
| ADK | 1840.00 | 1595.00 | 1919.00 | 1784.67 | 1785.00 |
| ADM | 14.00 | 9.00 | 8.00 | 10.33 | 10.00 |
| ADM2 | 389.00 | 357.00 | 416.00 | 387.33 | 391.00 |
| ADM5 | 212.00 | 178.00 | 251.00 | 213.67 | 168.00 |
| ADNP | 4332.00 | 4210.00 | 4905.00 | 4482.33 | 4417.00 |
| ADNP2 | 800.00 | 732.00 | 800.00 | 777.33 | 870.00 |
| ADNP-AS1 | 32.00 | 31.00 | 49.00 | 37.33 | 35.00 |
| ADO | 1263.00 | 1164.00 | 1382.00 | 1269.67 | 1511.00 |
| ADORA2A | 9.00 | 17.00 | 15.00 | 13.67 | 7.00 |
| ADORA2B | 3049.00 | 2873.00 | 3070.00 | 2997.33 | 2045.00 |
| ADPGK | 1991.00 | 1824.00 | 2010.00 | 1941.67 | 2019.00 |
| ADPRH | 24.00 | 29.00 | 36.00 | 29.67 | 32.00 |
| ADPRHL1 | 36.00 | 21.00 | 25.00 | 27.33 | 32.00 |
| ADPRHL2 | 560.00 | 572.00 | 547.00 | 559.67 | 620.00 |
| ADPRM | 177.00 | 186.00 | 229.00 | 197.33 | 236.00 |
| ADRB1 | 25.00 | 35.00 | 24.00 | 28.00 | 39.00 |
| ADRB2 | 2312.00 | 2193.00 | 2174.00 | 2226.33 | 2751.00 |
| ADRBK1 | 2329.00 | 2203.00 | 2341.00 | 2291.00 | 2515.00 |
| ADRBK2 | 65.00 | 48.00 | 52.00 | 55.00 | 83.00 |
| ADRM1 | 2800.00 | 2899.00 | 2920.00 | 2873.00 | 3880.00 |
| ADSL | 1703.00 | 1753.00 | 1763.00 | 1739.67 | 2271.00 |
| ADSS | 1430.00 | 1317.00 | 1495.00 | 1414.00 | 1618.00 |
| ADSSL1 | 17.00 | 11.00 | 15.00 | 14.33 | 8.00 |
| ADTRP | 3864.00 | 3744.00 | 3808.00 | 3805.33 | 3779.00 |
| AEBP1 | 6993.00 | 7310.00 | 7334.00 | 7212.33 | 5025.00 |
| AEBP2 | 860.00 | 790.00 | 892.00 | 847.33 | 1045.00 |

| | | | | | |
|---|---|---|---|---|---|
| AEN | 1202.00 | 1210.00 | 1283.00 | 1231.67 | 1286.00 |
| AES | 3509.00 | 3344.00 | 3567.00 | 3473.33 | 4161.00 |
| AFAP1 | 4590.00 | 4200.00 | 4681.00 | 4490.33 | 4009.00 |
| AFAP1L1 | 962.00 | 959.00 | 1026.00 | 982.33 | 958.00 |
| AFAP1L2 | 56.00 | 62.00 | 68.00 | 62.00 | 33.00 |
| AFF1 | 707.00 | 628.00 | 732.00 | 689.00 | 482.00 |
| AFF2 | 15.00 | 15.00 | 17.00 | 15.67 | 7.00 |
| AFF3 | 1462.00 | 1239.00 | 1464.00 | 1388.33 | 1420.00 |
| AFF4 | 3267.00 | 2706.00 | 3493.00 | 3155.33 | 2775.00 |
| AFG3L1P | 1518.00 | 1619.00 | 1638.00 | 1591.67 | 1631.00 |
| AFG3L2 | 2499.00 | 2377.00 | 2641.00 | 2505.67 | 3153.00 |
| AFMID | 623.00 | 521.00 | 560.00 | 568.00 | 678.00 |
| AFP | 6.00 | 0.00 | 1.00 | 2.33 | 1.00 |
| AFTPH | 620.00 | 593.00 | 743.00 | 652.00 | 702.00 |
| AGA | 785.00 | 798.00 | 804.00 | 795.67 | 827.00 |
| AGAP1 | 850.00 | 787.00 | 877.00 | 838.00 | 833.00 |
| AGAP11 | 52.00 | 33.00 | 42.00 | 42.33 | 57.00 |
| AGAP1-IT1 | 7.00 | 0.00 | 3.00 | 3.33 | 2.00 |
| AGAP2 | 21.00 | 13.00 | 16.00 | 16.67 | 20.00 |
| AGAP2-AS1 | 353.00 | 304.00 | 383.00 | 346.67 | 303.00 |
| AGAP3 | 2852.00 | 2958.00 | 3152.00 | 2987.33 | 3134.00 |
| AGAP4 | 250.00 | 214.00 | 295.00 | 253.00 | 231.00 |
| AGAP5 | 163.00 | 185.00 | 214.00 | 187.33 | 176.00 |
| AGAP6 | 848.00 | 896.00 | 949.00 | 897.67 | 918.00 |
| AGAP7P | 97.00 | 87.00 | 107.00 | 97.00 | 106.00 |
| AGAP9 | 488.00 | 521.00 | 473.00 | 494.00 | 462.00 |
| AGBL2 | 56.00 | 55.00 | 77.00 | 62.67 | 57.00 |
| AGBL3 | 86.00 | 74.00 | 77.00 | 79.00 | 99.00 |
| AGBL5 | 839.00 | 739.00 | 767.00 | 781.67 | 793.00 |
| AGFG1 | 1429.00 | 1329.00 | 1574.00 | 1444.00 | 1571.00 |
| AGFG2 | 279.00 | 248.00 | 286.00 | 271.00 | 191.00 |
| AGGF1 | 732.00 | 684.00 | 727.00 | 714.33 | 766.00 |
| AGK | 331.00 | 331.00 | 337.00 | 333.00 | 339.00 |
| AGL | 719.00 | 591.00 | 788.00 | 699.33 | 707.00 |
| AGMAT | 91.00 | 116.00 | 111.00 | 106.00 | 117.00 |
| AGO1 | 659.00 | 619.00 | 655.00 | 644.33 | 568.00 |
| AGO2 | 577.00 | 501.00 | 588.00 | 555.33 | 488.00 |
| AGO3 | 240.00 | 225.00 | 207.00 | 224.00 | 252.00 |
| AGO4 | 800.00 | 830.00 | 957.00 | 862.33 | 727.00 |
| AGPAT1 | 40.00 | 48.00 | 50.00 | 46.00 | 54.00 |
| AGPAT2 | 662.00 | 764.00 | 749.00 | 725.00 | 878.00 |
| AGPAT3 | 2672.00 | 2631.00 | 2717.00 | 2673.33 | 2622.00 |
| AGPAT4 | 3030.00 | 3118.00 | 3488.00 | 3212.00 | 2461.00 |
| AGPAT4-IT1 | 121.00 | 130.00 | 145.00 | 132.00 | 69.00 |
| AGPAT5 | 2363.00 | 2225.00 | 2297.00 | 2295.00 | 2736.00 |
| AGPS | 1420.00 | 1224.00 | 1551.00 | 1398.33 | 1379.00 |
| AGRN | 7531.00 | 7315.00 | 7654.00 | 7500.00 | 4760.00 |
| AGTPBP1 | 522.00 | 453.00 | 513.00 | 496.00 | 691.00 |

| | | | | | |
|---|---|---|---|---|---|
| AGTR1 | 64.00 | 60.00 | 60.00 | 61.33 | 58.00 |
| AGTRAP | 758.00 | 825.00 | 753.00 | 778.67 | 909.00 |
| AHCTF1 | 2414.00 | 2132.00 | 2565.00 | 2370.33 | 2468.00 |
| AHCTF1P1 | 215.00 | 198.00 | 243.00 | 218.67 | 187.00 |
| AHCY | 5401.00 | 5048.00 | 5756.00 | 5401.67 | 7373.00 |
| AHCYL1 | 2545.00 | 2351.00 | 2610.00 | 2502.00 | 2384.00 |
| AHCYL2 | 871.00 | 794.00 | 924.00 | 863.00 | 764.00 |
| AHDC1 | 436.00 | 446.00 | 465.00 | 449.00 | 284.00 |
| AHI1 | 963.00 | 1004.00 | 1095.00 | 1020.67 | 1155.00 |
| AHNAK | 44265.00 | 41138.00 | 45687.00 | 43696.67 | 30121.00 |
| AHNAK2 | 1746.00 | 1739.00 | 1960.00 | 1815.00 | 2125.00 |
| AHR | 339.00 | 273.00 | 429.00 | 347.00 | 283.00 |
| AHRR | 1459.00 | 1347.00 | 1720.00 | 1508.67 | 1411.00 |
| AHSA1 | 2601.00 | 2722.00 | 2800.00 | 2707.67 | 3432.00 |
| AHSA2 | 3294.00 | 3431.00 | 3635.00 | 3453.33 | 3325.00 |
| AIDA | 352.00 | 307.00 | 393.00 | 350.67 | 389.00 |
| AIF1L | 4.00 | 1.00 | 5.00 | 3.33 | 5.00 |
| AIFM1 | 816.00 | 791.00 | 840.00 | 815.67 | 956.00 |
| AIFM2 | 573.00 | 548.00 | 566.00 | 562.33 | 601.00 |
| AIG1 | 1291.00 | 1240.00 | 1203.00 | 1244.67 | 1153.00 |
| AIM1 | 114.00 | 127.00 | 103.00 | 114.67 | 59.00 |
| AIM1L | 13.00 | 18.00 | 4.00 | 11.67 | 9.00 |
| AIM2 | 103.00 | 78.00 | 83.00 | 88.00 | 63.00 |
| AIMP1 | 1493.00 | 1427.00 | 1560.00 | 1493.33 | 1820.00 |
| AIMP2 | 1169.00 | 1104.00 | 1267.00 | 1180.00 | 1424.00 |
| AIP | 868.00 | 857.00 | 814.00 | 846.33 | 1039.00 |
| AJUBA | 4000.00 | 3746.00 | 4086.00 | 3944.00 | 2686.00 |
| AK1 | 540.00 | 516.00 | 514.00 | 523.33 | 565.00 |
| AK2 | 4912.00 | 4837.00 | 5218.00 | 4989.00 | 5591.00 |
| AK3 | 1091.00 | 1107.00 | 1269.00 | 1155.67 | 1352.00 |
| AK4 | 361.00 | 338.00 | 433.00 | 377.33 | 429.00 |
| AK5 | 1043.00 | 901.00 | 1039.00 | 994.33 | 998.00 |
| AK6 | 8.00 | 1.00 | 0.00 | 3.00 | 0.00 |
| AK9 | 129.00 | 131.00 | 125.00 | 128.33 | 125.00 |
| AKAP1 | 2122.00 | 2188.00 | 2225.00 | 2178.33 | 2097.00 |
| AKAP10 | 782.00 | 771.00 | 919.00 | 824.00 | 926.00 |
| AKAP11 | 1890.00 | 1641.00 | 1971.00 | 1834.00 | 1616.00 |
| AKAP12 | 20215.00 | 18166.00 | 20185.00 | 19522.00 | 14538.00 |
| AKAP13 | 2322.00 | 2191.00 | 2501.00 | 2338.00 | 2031.00 |
| AKAP2 | 97.00 | 104.00 | 92.00 | 97.67 | 70.00 |
| AKAP3 | 9.00 | 4.00 | 7.00 | 6.67 | 9.00 |
| AKAP5 | 4.00 | 11.00 | 10.00 | 8.33 | 8.00 |
| AKAP6 | 56.00 | 60.00 | 70.00 | 62.00 | 49.00 |
| AKAP7 | 72.00 | 68.00 | 78.00 | 72.67 | 125.00 |
| AKAP8 | 1039.00 | 1060.00 | 1108.00 | 1069.00 | 1056.00 |
| AKAP8L | 1427.00 | 1496.00 | 1513.00 | 1478.67 | 1431.00 |
| AKAP9 | 1723.00 | 1594.00 | 1795.00 | 1704.00 | 1481.00 |
| AKIP1 | 477.00 | 495.00 | 543.00 | 505.00 | 575.00 |

| | | | | | |
|---|---|---|---|---|---|
| AKIRIN1 | 1465.00 | 1415.00 | 1515.00 | 1465.00 | 1762.00 |
| AKIRIN2 | 1128.00 | 1093.00 | 1168.00 | 1129.67 | 1330.00 |
| AKNA | 422.00 | 360.00 | 397.00 | 393.00 | 381.00 |
| AKNAD1 | 12.00 | 12.00 | 6.00 | 10.00 | 17.00 |
| AKR1A1 | 1467.00 | 1531.00 | 1627.00 | 1541.67 | 2047.00 |
| AKR1B1 | 2030.00 | 1972.00 | 2160.00 | 2054.00 | 2136.00 |
| AKR1C1 | 32.00 | 37.00 | 28.00 | 32.33 | 30.00 |
| AKR1C2 | 7.00 | 8.00 | 5.00 | 6.67 | 8.00 |
| AKR1C3 | 80.00 | 75.00 | 90.00 | 81.67 | 104.00 |
| AKR1C8P | 5.00 | 2.00 | 1.00 | 2.67 | 2.00 |
| AKR1E2 | 54.00 | 46.00 | 53.00 | 51.00 | 60.00 |
| AKR7A2 | 763.00 | 833.00 | 855.00 | 817.00 | 1039.00 |
| AKR7A3 | 18.00 | 12.00 | 11.00 | 13.67 | 9.00 |
| AKR7L | 12.00 | 16.00 | 25.00 | 17.67 | 17.00 |
| AKT1 | 2920.00 | 2830.00 | 2990.00 | 2913.33 | 3137.00 |
| AKT1S1 | 1217.00 | 1177.00 | 1304.00 | 1232.67 | 1437.00 |
| AKT2 | 2596.00 | 2680.00 | 2853.00 | 2709.67 | 2749.00 |
| AKTIP | 371.00 | 327.00 | 401.00 | 366.33 | 438.00 |
| ALAD | 484.00 | 538.00 | 531.00 | 517.67 | 516.00 |
| ALAS1 | 1038.00 | 896.00 | 1034.00 | 989.33 | 1148.00 |
| ALB | 7.00 | 4.00 | 2.00 | 4.33 | 3.00 |
| ALCAM | 6060.00 | 5412.00 | 5944.00 | 5805.33 | 6119.00 |
| ALDH16A1 | 826.00 | 852.00 | 815.00 | 831.00 | 980.00 |
| ALDH18A1 | 4632.00 | 4685.00 | 4527.00 | 4614.67 | 5220.00 |
| ALDH1A3 | 12506.00 | 11741.00 | 13785.00 | 12677.33 | 22207.00 |
| ALDH1B1 | 549.00 | 519.00 | 555.00 | 541.00 | 619.00 |
| ALDH1L1 | 14.00 | 22.00 | 16.00 | 17.33 | 21.00 |
| ALDH1L2 | 1321.00 | 1142.00 | 1197.00 | 1220.00 | 1119.00 |
| ALDH2 | 4.00 | 3.00 | 4.00 | 3.67 | 4.00 |
| ALDH3A2 | 374.00 | 312.00 | 372.00 | 352.67 | 303.00 |
| ALDH3B1 | 211.00 | 221.00 | 198.00 | 210.00 | 88.00 |
| ALDH4A1 | 479.00 | 484.00 | 476.00 | 479.67 | 424.00 |
| ALDH5A1 | 161.00 | 194.00 | 177.00 | 177.33 | 228.00 |
| ALDH6A1 | 354.00 | 351.00 | 382.00 | 362.33 | 324.00 |
| ALDH7A1 | 1145.00 | 1159.00 | 1146.00 | 1150.00 | 1448.00 |
| ALDH8A1 | 13.00 | 9.00 | 21.00 | 14.33 | 13.00 |
| ALDH9A1 | 1297.00 | 1264.00 | 1321.00 | 1294.00 | 1557.00 |
| ALDOA | 11428.00 | 11193.00 | 11490.00 | 11370.33 | 14280.00 |
| ALDOC | 80.00 | 69.00 | 95.00 | 81.33 | 133.00 |
| ALG1 | 1210.00 | 1250.00 | 1299.00 | 1253.00 | 1326.00 |
| ALG10 | 237.00 | 238.00 | 243.00 | 239.33 | 232.00 |
| ALG10B | 1333.00 | 1122.00 | 1227.00 | 1227.33 | 1212.00 |
| ALG11 | 177.00 | 194.00 | 178.00 | 183.00 | 112.00 |
| ALG12 | 502.00 | 508.00 | 500.00 | 503.33 | 538.00 |
| ALG13 | 2429.00 | 2606.00 | 2928.00 | 2654.33 | 2424.00 |
| ALG14 | 139.00 | 161.00 | 163.00 | 154.33 | 276.00 |
| ALG1L9P | 10.00 | 13.00 | 13.00 | 12.00 | 7.00 |
| ALG2 | 1053.00 | 1008.00 | 1090.00 | 1050.33 | 1089.00 |

| | | | | | |
|---|---|---|---|---|---|
| ALG3 | 1771.00 | 1667.00 | 1675.00 | 1704.33 | 1976.00 |
| ALG5 | 626.00 | 588.00 | 530.00 | 581.33 | 791.00 |
| ALG6 | 542.00 | 607.00 | 635.00 | 594.67 | 749.00 |
| ALG8 | 1206.00 | 1191.00 | 1166.00 | 1187.67 | 1235.00 |
| ALG9 | 922.00 | 843.00 | 941.00 | 902.00 | 719.00 |
| ALKBH1 | 243.00 | 199.00 | 245.00 | 229.00 | 267.00 |
| ALKBH2 | 484.00 | 507.00 | 502.00 | 497.67 | 682.00 |
| ALKBH3 | 506.00 | 495.00 | 575.00 | 525.33 | 661.00 |
| ALKBH4 | 200.00 | 199.00 | 237.00 | 212.00 | 259.00 |
| ALKBH5 | 1627.00 | 1455.00 | 1624.00 | 1568.67 | 1778.00 |
| ALKBH6 | 401.00 | 429.00 | 373.00 | 401.00 | 436.00 |
| ALKBH7 | 301.00 | 346.00 | 389.00 | 345.33 | 442.00 |
| ALKBH8 | 317.00 | 311.00 | 326.00 | 318.00 | 354.00 |
| ALMS1 | 556.00 | 494.00 | 571.00 | 540.33 | 512.00 |
| ALMS1-IT1 | 5.00 | 10.00 | 15.00 | 10.00 | 20.00 |
| ALOX12-AS1 | 73.00 | 81.00 | 56.00 | 70.00 | 70.00 |
| ALOX15P1 | 4.00 | 4.00 | 5.00 | 4.33 | 7.00 |
| ALOXE3 | 35.00 | 25.00 | 22.00 | 27.33 | 11.00 |
| ALPK1 | 532.00 | 469.00 | 508.00 | 503.00 | 477.00 |
| ALPK2 | 16.00 | 10.00 | 11.00 | 12.33 | 6.00 |
| ALPK3 | 15.00 | 10.00 | 17.00 | 14.00 | 9.00 |
| ALS2 | 1237.00 | 1170.00 | 1329.00 | 1245.33 | 1320.00 |
| ALS2CR11 | 30.00 | 15.00 | 15.00 | 20.00 | 33.00 |
| ALS2CR12 | 26.00 | 28.00 | 36.00 | 30.00 | 20.00 |
| ALX1 | 160.00 | 185.00 | 174.00 | 173.00 | 215.00 |
| ALYREF | 879.00 | 954.00 | 1050.00 | 961.00 | 1280.00 |
| AMACR | 76.00 | 43.00 | 76.00 | 65.00 | 87.00 |
| AMBRA1 | 939.00 | 939.00 | 936.00 | 938.00 | 713.00 |
| AMD1 | 1520.00 | 1369.00 | 1568.00 | 1485.67 | 1566.00 |
| AMDHD2 | 1219.00 | 1223.00 | 1191.00 | 1211.00 | 1327.00 |
| AMER1 | 421.00 | 462.00 | 552.00 | 478.33 | 396.00 |
| AMFR | 2156.00 | 2125.00 | 2241.00 | 2174.00 | 2383.00 |
| AMH | 458.00 | 486.00 | 577.00 | 507.00 | 453.00 |
| AMIGO1 | 95.00 | 96.00 | 85.00 | 92.00 | 80.00 |
| AMIGO2 | 9225.00 | 8467.00 | 9703.00 | 9131.67 | 7662.00 |
| AMIGO3 | 164.00 | 159.00 | 156.00 | 159.67 | 156.00 |
| AMMECR1 | 396.00 | 377.00 | 457.00 | 410.00 | 443.00 |
| AMMECR1L | 1551.00 | 1509.00 | 1589.00 | 1549.67 | 1448.00 |
| AMN | 5.00 | 5.00 | 2.00 | 4.00 | 2.00 |
| AMN1 | 93.00 | 91.00 | 101.00 | 95.00 | 136.00 |
| AMOTL1 | 2328.00 | 2063.00 | 2360.00 | 2250.33 | 1760.00 |
| AMOTL2 | 8806.00 | 8382.00 | 9225.00 | 8804.33 | 7877.00 |
| AMPD2 | 3201.00 | 3352.00 | 3504.00 | 3352.33 | 3514.00 |
| AMPD3 | 807.00 | 790.00 | 829.00 | 808.67 | 693.00 |
| AMT | 508.00 | 600.00 | 574.00 | 560.67 | 572.00 |
| AMY2B | 194.00 | 195.00 | 213.00 | 200.67 | 213.00 |
| AMZ2 | 1586.00 | 1505.00 | 1698.00 | 1596.33 | 1841.00 |
| AMZ2P1 | 259.00 | 252.00 | 283.00 | 264.67 | 311.00 |

| | | | | | |
|---|---|---|---|---|---|
| ANAPC1 | 1587.00 | 1452.00 | 1674.00 | 1571.00 | 1432.00 |
| ANAPC10 | 119.00 | 121.00 | 141.00 | 127.00 | 168.00 |
| ANAPC11 | 1180.00 | 1187.00 | 1272.00 | 1213.00 | 1614.00 |
| ANAPC13 | 1956.00 | 1775.00 | 2056.00 | 1929.00 | 2403.00 |
| ANAPC15 | 291.00 | 275.00 | 325.00 | 297.00 | 404.00 |
| ANAPC16 | 1387.00 | 1296.00 | 1486.00 | 1389.67 | 1669.00 |
| ANAPC2 | 1114.00 | 1138.00 | 1152.00 | 1134.67 | 935.00 |
| ANAPC4 | 705.00 | 710.00 | 694.00 | 703.00 | 882.00 |
| ANAPC5 | 2505.00 | 2438.00 | 2628.00 | 2523.67 | 2794.00 |
| ANAPC7 | 1879.00 | 1848.00 | 1834.00 | 1853.67 | 2233.00 |
| ANG | 13.00 | 22.00 | 19.00 | 18.00 | 17.00 |
| ANGEL1 | 1240.00 | 1108.00 | 1254.00 | 1200.67 | 1007.00 |
| ANGEL2 | 942.00 | 873.00 | 1026.00 | 947.00 | 1138.00 |
| ANGPT1 | 55.00 | 46.00 | 46.00 | 49.00 | 93.00 |
| ANGPT2 | 462.00 | 431.00 | 437.00 | 443.33 | 845.00 |
| ANGPTL2 | 37.00 | 56.00 | 58.00 | 50.33 | 63.00 |
| ANGPTL4 | 62.00 | 60.00 | 63.00 | 61.67 | 229.00 |
| ANGPTL6 | 7.00 | 3.00 | 0.00 | 3.33 | 4.00 |
| ANK2 | 1005.00 | 900.00 | 1048.00 | 984.33 | 980.00 |
| ANK3 | 1187.00 | 970.00 | 1236.00 | 1131.00 | 1136.00 |
| ANKAR | 88.00 | 82.00 | 108.00 | 92.67 | 87.00 |
| ANKDD1A | 21.00 | 24.00 | 27.00 | 24.00 | 21.00 |
| ANKDD1B | 7.00 | 8.00 | 6.00 | 7.00 | 3.00 |
| ANKEF1 | 109.00 | 110.00 | 112.00 | 110.33 | 127.00 |
| ANKFN1 | 109.00 | 80.00 | 110.00 | 99.67 | 96.00 |
| ANKFY1 | 2685.00 | 2414.00 | 2645.00 | 2581.33 | 2538.00 |
| ANKH | 1067.00 | 1092.00 | 1135.00 | 1098.00 | 1039.00 |
| ANKHD1 | 306.00 | 259.00 | 308.00 | 291.00 | 338.00 |
| ANKHD1-EIF4EBP3 | 14.00 | 26.00 | 41.00 | 27.00 | 27.00 |
| ANKIB1 | 1371.00 | 1296.00 | 1442.00 | 1369.67 | 1446.00 |
| ANKK1 | 22.00 | 13.00 | 11.00 | 15.33 | 12.00 |
| ANKLE1 | 263.00 | 295.00 | 314.00 | 290.67 | 313.00 |
| ANKLE2 | 3045.00 | 2942.00 | 3035.00 | 3007.33 | 2784.00 |
| ANKMY1 | 596.00 | 607.00 | 635.00 | 612.67 | 406.00 |
| ANKMY2 | 493.00 | 468.00 | 473.00 | 478.00 | 543.00 |
| ANKRA2 | 295.00 | 312.00 | 349.00 | 318.67 | 327.00 |
| ANKRD1 | 12519.00 | 12097.00 | 12925.00 | 12513.67 | 12452.00 |
| ANKRD10 | 1498.00 | 1588.00 | 1724.00 | 1603.33 | 1629.00 |
| ANKRD11 | 12039.00 | 11491.00 | 12443.00 | 11991.00 | 8932.00 |
| ANKRD12 | 1286.00 | 1120.00 | 1373.00 | 1259.67 | 1200.00 |
| ANKRD13A | 1545.00 | 1459.00 | 1657.00 | 1553.67 | 1825.00 |
| ANKRD13B | 212.00 | 205.00 | 225.00 | 214.00 | 267.00 |
| ANKRD13C | 623.00 | 617.00 | 663.00 | 634.33 | 612.00 |
| ANKRD13D | 938.00 | 1020.00 | 1107.00 | 1021.67 | 1115.00 |
| ANKRD16 | 293.00 | 322.00 | 316.00 | 310.33 | 346.00 |
| ANKRD17 | 2407.00 | 2235.00 | 2686.00 | 2442.67 | 2157.00 |
| ANKRD18A | 245.00 | 246.00 | 263.00 | 251.33 | 262.00 |
| ANKRD18B | 191.00 | 178.00 | 260.00 | 209.67 | 249.00 |

| | | | | | |
|---|---|---|---|---|---|
| ANKRD19P | 295.00 | 342.00 | 343.00 | 326.67 | 337.00 |
| ANKRD20A5P | 28.00 | 36.00 | 24.00 | 29.33 | 13.00 |
| ANKRD23 | 116.00 | 114.00 | 135.00 | 121.67 | 78.00 |
| ANKRD24 | 76.00 | 93.00 | 113.00 | 94.00 | 66.00 |
| ANKRD26 | 903.00 | 868.00 | 976.00 | 915.67 | 970.00 |
| ANKRD27 | 872.00 | 759.00 | 890.00 | 840.33 | 967.00 |
| ANKRD28 | 3197.00 | 2995.00 | 3486.00 | 3226.00 | 3664.00 |
| ANKRD29 | 265.00 | 330.00 | 346.00 | 313.67 | 241.00 |
| ANKRD31 | 28.00 | 28.00 | 27.00 | 27.67 | 34.00 |
| ANKRD33B | 1472.00 | 1346.00 | 1626.00 | 1481.33 | 676.00 |
| ANKRD34A | 162.00 | 140.00 | 125.00 | 142.33 | 135.00 |
| ANKRD36 | 89.00 | 109.00 | 103.00 | 100.33 | 92.00 |
| ANKRD36B | 69.00 | 53.00 | 59.00 | 60.33 | 70.00 |
| ANKRD36BP2 | 61.00 | 59.00 | 60.00 | 60.00 | 68.00 |
| ANKRD36C | 109.00 | 89.00 | 110.00 | 102.67 | 79.00 |
| ANKRD37 | 35.00 | 40.00 | 45.00 | 40.00 | 45.00 |
| ANKRD39 | 118.00 | 108.00 | 96.00 | 107.33 | 127.00 |
| ANKRD40 | 1688.00 | 1527.00 | 1758.00 | 1657.67 | 1694.00 |
| ANKRD42 | 372.00 | 341.00 | 334.00 | 349.00 | 266.00 |
| ANKRD44 | 332.00 | 325.00 | 405.00 | 354.00 | 382.00 |
| ANKRD45 | 4.00 | 0.00 | 1.00 | 1.67 | 5.00 |
| ANKRD46 | 345.00 | 292.00 | 359.00 | 332.00 | 352.00 |
| ANKRD49 | 360.00 | 292.00 | 389.00 | 347.00 | 313.00 |
| ANKRD50 | 1081.00 | 956.00 | 1197.00 | 1078.00 | 841.00 |
| ANKRD52 | 2945.00 | 2839.00 | 3242.00 | 3008.67 | 2355.00 |
| ANKRD53 | 9.00 | 10.00 | 6.00 | 8.33 | 4.00 |
| ANKRD54 | 314.00 | 367.00 | 375.00 | 352.00 | 440.00 |
| ANKRD6 | 8.00 | 9.00 | 9.00 | 8.67 | 11.00 |
| ANKRD61 | 6.00 | 11.00 | 9.00 | 8.67 | 14.00 |
| ANKRD7 | 42.00 | 26.00 | 41.00 | 36.33 | 43.00 |
| ANKRD9 | 188.00 | 200.00 | 204.00 | 197.33 | 224.00 |
| ANKS1A | 1058.00 | 984.00 | 1125.00 | 1055.67 | 891.00 |
| ANKS3 | 518.00 | 519.00 | 642.00 | 559.67 | 582.00 |
| ANKS6 | 1305.00 | 1305.00 | 1445.00 | 1351.67 | 1323.00 |
| ANKZF1 | 1677.00 | 1655.00 | 1718.00 | 1683.33 | 1650.00 |
| ANLN | 3007.00 | 2759.00 | 2946.00 | 2904.00 | 3719.00 |
| ANO1 | 4.00 | 2.00 | 6.00 | 4.00 | 1.00 |
| ANO10 | 2098.00 | 2050.00 | 2187.00 | 2111.67 | 1612.00 |
| ANO5 | 225.00 | 252.00 | 280.00 | 252.33 | 257.00 |
| ANO6 | 3038.00 | 2786.00 | 3050.00 | 2958.00 | 2364.00 |
| ANO7 | 32.00 | 18.00 | 24.00 | 24.67 | 21.00 |
| ANO8 | 293.00 | 324.00 | 339.00 | 318.67 | 273.00 |
| ANOS1 | 302.00 | 313.00 | 372.00 | 329.00 | 272.00 |
| ANP32A | 2816.00 | 2662.00 | 2978.00 | 2818.67 | 3776.00 |
| ANP32A-IT1 | 31.00 | 24.00 | 27.00 | 27.33 | 9.00 |
| ANP32B | 3507.00 | 3454.00 | 3547.00 | 3502.67 | 4894.00 |
| ANP32E | 1161.00 | 915.00 | 1229.00 | 1101.67 | 1446.00 |
| ANPEP | 15491.00 | 15235.00 | 14621.00 | 15115.67 | 15374.00 |

| | | | | | |
|---|---|---|---|---|---|
| ANTXR1 | 2352.00 | 2152.00 | 2359.00 | 2287.67 | 1508.00 |
| ANTXR2 | 2432.00 | 2156.00 | 2276.00 | 2288.00 | 1712.00 |
| ANXA1 | 4739.00 | 4548.00 | 4758.00 | 4681.67 | 5921.00 |
| ANXA10 | 30.00 | 18.00 | 44.00 | 30.67 | 14.00 |
| ANXA11 | 1529.00 | 1547.00 | 1514.00 | 1530.00 | 1694.00 |
| ANXA2 | 25353.00 | 24446.00 | 26360.00 | 25386.33 | 31926.00 |
| ANXA2P2 | 1274.00 | 1310.00 | 1385.00 | 1323.00 | 1750.00 |
| ANXA2R | 61.00 | 58.00 | 57.00 | 58.67 | 70.00 |
| ANXA3 | 4.00 | 3.00 | 1.00 | 2.67 | 0.00 |
| ANXA4 | 488.00 | 471.00 | 577.00 | 512.00 | 626.00 |
| ANXA5 | 10112.00 | 9664.00 | 10946.00 | 10240.67 | 8855.00 |
| ANXA6 | 2314.00 | 2220.00 | 2449.00 | 2327.67 | 2464.00 |
| ANXA7 | 2755.00 | 2431.00 | 2795.00 | 2660.33 | 2340.00 |
| ANXA9 | 5.00 | 0.00 | 1.00 | 2.00 | 2.00 |
| AOC2 | 93.00 | 84.00 | 120.00 | 99.00 | 84.00 |
| AOC3 | 34.00 | 22.00 | 32.00 | 29.33 | 32.00 |
| AOX1 | 6999.00 | 6353.00 | 7835.00 | 7062.33 | 6765.00 |
| AP1AR | 788.00 | 675.00 | 940.00 | 801.00 | 916.00 |
| AP1B1 | 2162.00 | 2100.00 | 2193.00 | 2151.67 | 2290.00 |
| AP1G1 | 2069.00 | 1898.00 | 2148.00 | 2038.33 | 2125.00 |
| AP1G2 | 391.00 | 359.00 | 384.00 | 378.00 | 292.00 |
| AP1M1 | 1319.00 | 1272.00 | 1263.00 | 1284.67 | 1661.00 |
| AP1S1 | 1483.00 | 1383.00 | 1473.00 | 1446.33 | 1796.00 |
| AP1S2 | 1388.00 | 1203.00 | 1469.00 | 1353.33 | 1243.00 |
| AP1S3 | 58.00 | 50.00 | 62.00 | 56.67 | 91.00 |
| AP2A1 | 1723.00 | 1742.00 | 1833.00 | 1766.00 | 1857.00 |
| AP2A2 | 217.00 | 191.00 | 221.00 | 209.67 | 238.00 |
| AP2B1 | 8329.00 | 7437.00 | 8922.00 | 8229.33 | 7518.00 |
| AP2M1 | 6501.00 | 6235.00 | 6809.00 | 6515.00 | 6318.00 |
| AP2S1 | 1193.00 | 1263.00 | 1311.00 | 1255.67 | 1667.00 |
| AP3B1 | 1244.00 | 1128.00 | 1350.00 | 1240.67 | 1367.00 |
| AP3D1 | 5241.00 | 5123.00 | 5322.00 | 5228.67 | 5215.00 |
| AP3M1 | 1993.00 | 1803.00 | 2126.00 | 1974.00 | 1949.00 |
| AP3M2 | 1600.00 | 1413.00 | 1569.00 | 1527.33 | 1672.00 |
| AP3S1 | 692.00 | 626.00 | 761.00 | 693.00 | 849.00 |
| AP3S2 | 47.00 | 64.00 | 43.00 | 51.33 | 47.00 |
| AP4B1 | 1254.00 | 1158.00 | 1211.00 | 1207.67 | 1397.00 |
| AP4E1 | 527.00 | 479.00 | 581.00 | 529.00 | 573.00 |
| AP4M1 | 505.00 | 443.00 | 465.00 | 471.00 | 795.00 |
| AP4S1 | 236.00 | 210.00 | 223.00 | 223.00 | 254.00 |
| AP5B1 | 1206.00 | 1139.00 | 1292.00 | 1212.33 | 1111.00 |
| AP5M1 | 415.00 | 378.00 | 436.00 | 409.67 | 375.00 |
| AP5S1 | 295.00 | 323.00 | 322.00 | 313.33 | 363.00 |
| AP5Z1 | 1972.00 | 2017.00 | 2184.00 | 2057.67 | 2199.00 |
| APAF1 | 421.00 | 386.00 | 417.00 | 408.00 | 373.00 |
| APBA1 | 10.00 | 3.00 | 9.00 | 7.33 | 1.00 |
| APBA2 | 61.00 | 85.00 | 105.00 | 83.67 | 84.00 |
| APBA3 | 739.00 | 740.00 | 783.00 | 754.00 | 697.00 |

| | | | | | |
|---|---|---|---|---|---|
| APBB1 | 713.00 | 666.00 | 727.00 | 702.00 | 685.00 |
| APBB1IP | 27.00 | 18.00 | 30.00 | 25.00 | 31.00 |
| APBB2 | 4274.00 | 3890.00 | 4674.00 | 4279.33 | 4058.00 |
| APBB3 | 1185.00 | 1314.00 | 1468.00 | 1322.33 | 1102.00 |
| APC | 916.00 | 797.00 | 935.00 | 882.67 | 751.00 |
| APC2 | 8.00 | 9.00 | 11.00 | 9.33 | 12.00 |
| APCDD1L | 4.00 | 1.00 | 0.00 | 1.67 | 0.00 |
| APEH | 1646.00 | 1566.00 | 1700.00 | 1637.33 | 2036.00 |
| APEX1 | 4576.00 | 4426.00 | 4916.00 | 4639.33 | 5569.00 |
| APEX2 | 674.00 | 591.00 | 622.00 | 629.00 | 783.00 |
| APH1A | 2202.00 | 2254.00 | 2173.00 | 2209.67 | 2339.00 |
| APH1B | 1421.00 | 1224.00 | 1217.00 | 1287.33 | 1299.00 |
| API5 | 2300.00 | 2072.00 | 2333.00 | 2235.00 | 2716.00 |
| APIP | 403.00 | 366.00 | 457.00 | 408.67 | 642.00 |
| APITD1 | 50.00 | 39.00 | 30.00 | 39.67 | 57.00 |
| APLF | 160.00 | 166.00 | 163.00 | 163.00 | 168.00 |
| APLN | 222.00 | 194.00 | 189.00 | 201.67 | 494.00 |
| APLP1 | 16.00 | 18.00 | 10.00 | 14.67 | 17.00 |
| APLP2 | 6914.00 | 6797.00 | 7067.00 | 6926.00 | 6829.00 |
| APMAP | 5924.00 | 5960.00 | 5832.00 | 5905.33 | 5365.00 |
| APOA1BP | 876.00 | 905.00 | 901.00 | 894.00 | 1247.00 |
| APOBEC3B | 109.00 | 140.00 | 102.00 | 117.00 | 225.00 |
| APOBEC3C | 151.00 | 150.00 | 159.00 | 153.33 | 182.00 |
| APOBEC3D | 16.00 | 12.00 | 23.00 | 17.00 | 30.00 |
| APOBEC3F | 27.00 | 20.00 | 20.00 | 22.33 | 32.00 |
| APOBR | 5.00 | 3.00 | 3.00 | 3.67 | 1.00 |
| APOC1 | 7.00 | 3.00 | 5.00 | 5.00 | 7.00 |
| APOE | 27.00 | 18.00 | 16.00 | 20.33 | 19.00 |
| APOL1 | 29.00 | 30.00 | 46.00 | 35.00 | 19.00 |
| APOL2 | 210.00 | 242.00 | 231.00 | 227.67 | 277.00 |
| APOL3 | 34.00 | 58.00 | 46.00 | 46.00 | 28.00 |
| APOL6 | 575.00 | 493.00 | 556.00 | 541.33 | 562.00 |
| APOLD1 | 334.00 | 387.00 | 375.00 | 365.33 | 373.00 |
| APOO | 335.00 | 331.00 | 344.00 | 336.67 | 460.00 |
| APOOL | 522.00 | 477.00 | 701.00 | 566.67 | 610.00 |
| APOPT1 | 419.00 | 395.00 | 499.00 | 437.67 | 554.00 |
| APP | 33512.00 | 32910.00 | 35084.00 | 33835.33 | 31640.00 |
| APPBP2 | 1345.00 | 1261.00 | 1525.00 | 1377.00 | 1513.00 |
| APPL1 | 921.00 | 829.00 | 1077.00 | 942.33 | 1083.00 |
| APPL2 | 1001.00 | 963.00 | 1089.00 | 1017.67 | 1090.00 |
| APRT | 2389.00 | 2271.00 | 2374.00 | 2344.67 | 3421.00 |
| APTR | 178.00 | 196.00 | 196.00 | 190.00 | 229.00 |
| APTX | 1029.00 | 1118.00 | 1155.00 | 1100.67 | 1340.00 |
| AQP11 | 54.00 | 71.00 | 68.00 | 64.33 | 69.00 |
| AQP3 | 5.00 | 7.00 | 8.00 | 6.67 | 8.00 |
| AQP7P1 | 95.00 | 97.00 | 120.00 | 104.00 | 116.00 |
| AQP7P3 | 104.00 | 96.00 | 117.00 | 105.67 | 110.00 |
| AQR | 1292.00 | 1244.00 | 1419.00 | 1318.33 | 1330.00 |

| | | | | | |
|---|---|---|---|---|---|
| AR | 593.00 | 516.00 | 696.00 | 601.67 | 278.00 |
| ARAF | 1022.00 | 1049.00 | 1047.00 | 1039.33 | 1148.00 |
| ARAP1 | 2760.00 | 2684.00 | 2882.00 | 2775.33 | 2673.00 |
| ARAP2 | 123.00 | 103.00 | 124.00 | 116.67 | 95.00 |
| ARAP3 | 1687.00 | 1699.00 | 1903.00 | 1763.00 | 1651.00 |
| ARC | 22.00 | 13.00 | 15.00 | 16.67 | 10.00 |
| ARCN1 | 4114.00 | 3780.00 | 4427.00 | 4107.00 | 3838.00 |
| AREG | 73.00 | 73.00 | 66.00 | 70.67 | 100.00 |
| AREL1 | 1499.00 | 1463.00 | 1518.00 | 1493.33 | 1526.00 |
| ARF1 | 5987.00 | 5702.00 | 6483.00 | 6057.33 | 7270.00 |
| ARF3 | 2951.00 | 2768.00 | 3024.00 | 2914.33 | 2937.00 |
| ARF4 | 4031.00 | 3626.00 | 4166.00 | 3941.00 | 4618.00 |
| ARF5 | 935.00 | 1028.00 | 1020.00 | 994.33 | 1295.00 |
| ARF6 | 1894.00 | 1924.00 | 2156.00 | 1991.33 | 2328.00 |
| ARFGAP1 | 3767.00 | 3845.00 | 4165.00 | 3925.67 | 3607.00 |
| ARFGAP2 | 1382.00 | 1323.00 | 1416.00 | 1373.67 | 1636.00 |
| ARFGAP3 | 925.00 | 820.00 | 916.00 | 887.00 | 842.00 |
| ARFGEF1 | 2204.00 | 2012.00 | 2125.00 | 2113.67 | 2018.00 |
| ARFGEF2 | 2739.00 | 2348.00 | 2814.00 | 2633.67 | 2460.00 |
| ARFGEF3 | 418.00 | 392.00 | 451.00 | 420.33 | 364.00 |
| ARFIP1 | 399.00 | 347.00 | 429.00 | 391.67 | 375.00 |
| ARFIP2 | 1417.00 | 1292.00 | 1453.00 | 1387.33 | 1472.00 |
| ARFRP1 | 1116.00 | 1081.00 | 1118.00 | 1105.00 | 1251.00 |
| ARG2 | 326.00 | 278.00 | 332.00 | 312.00 | 460.00 |
| ARGLU1 | 4100.00 | 4232.00 | 4476.00 | 4269.33 | 4284.00 |
| ARHGAP1 | 1147.00 | 1183.00 | 1170.00 | 1166.67 | 1389.00 |
| ARHGAP10 | 235.00 | 247.00 | 262.00 | 248.00 | 261.00 |
| ARHGAP11A | 24.00 | 20.00 | 22.00 | 22.00 | 22.00 |
| ARHGAP11B | 6.00 | 2.00 | 1.00 | 3.00 | 6.00 |
| ARHGAP12 | 1799.00 | 1707.00 | 2012.00 | 1839.33 | 1723.00 |
| ARHGAP17 | 1137.00 | 1076.00 | 1167.00 | 1126.67 | 959.00 |
| ARHGAP18 | 3098.00 | 2899.00 | 3313.00 | 3103.33 | 3152.00 |
| ARHGAP19 | 358.00 | 367.00 | 403.00 | 376.00 | 556.00 |
| ARHGAP20 | 32.00 | 27.00 | 35.00 | 31.33 | 12.00 |
| ARHGAP21 | 2041.00 | 1938.00 | 2238.00 | 2072.33 | 1811.00 |
| ARHGAP22 | 2156.00 | 2200.00 | 2326.00 | 2227.33 | 2179.00 |
| ARHGAP23 | 169.00 | 162.00 | 200.00 | 177.00 | 137.00 |
| ARHGAP24 | 1032.00 | 1097.00 | 1190.00 | 1106.33 | 1478.00 |
| ARHGAP25 | 7.00 | 8.00 | 8.00 | 7.67 | 11.00 |
| ARHGAP26 | 268.00 | 238.00 | 273.00 | 259.67 | 133.00 |
| ARHGAP27 | 24.00 | 20.00 | 12.00 | 18.67 | 18.00 |
| ARHGAP29 | 12036.00 | 10828.00 | 12321.00 | 11728.33 | 11730.00 |
| ARHGAP30 | 249.00 | 283.00 | 256.00 | 262.67 | 216.00 |
| ARHGAP31 | 558.00 | 536.00 | 536.00 | 543.33 | 573.00 |
| ARHGAP31-AS1 | 10.00 | 6.00 | 10.00 | 8.67 | 4.00 |
| ARHGAP32 | 725.00 | 732.00 | 850.00 | 769.00 | 671.00 |
| ARHGAP33 | 208.00 | 259.00 | 190.00 | 219.00 | 254.00 |
| ARHGAP35 | 2641.00 | 2386.00 | 2806.00 | 2611.00 | 1908.00 |

| | | | | | |
|---|---|---|---|---|---|
| ARHGAP39 | 749.00 | 730.00 | 782.00 | 753.67 | 660.00 |
| ARHGAP4 | 18.00 | 21.00 | 17.00 | 18.67 | 19.00 |
| ARHGAP40 | 1826.00 | 1747.00 | 1806.00 | 1793.00 | 2065.00 |
| ARHGAP42 | 898.00 | 711.00 | 965.00 | 858.00 | 770.00 |
| ARHGAP5 | 1322.00 | 1131.00 | 1467.00 | 1306.67 | 1116.00 |
| ARHGAP5-AS1 | 30.00 | 50.00 | 46.00 | 42.00 | 61.00 |
| ARHGAP6 | 177.00 | 182.00 | 183.00 | 180.67 | 165.00 |
| ARHGAP9 | 35.00 | 29.00 | 17.00 | 27.00 | 20.00 |
| ARHGDIA | 8733.00 | 8933.00 | 9363.00 | 9009.67 | 7668.00 |
| ARHGDIB | 2306.00 | 2228.00 | 2382.00 | 2305.33 | 3110.00 |
| ARHGEF1 | 2994.00 | 2979.00 | 2965.00 | 2979.33 | 2966.00 |
| ARHGEF10L | 120.00 | 114.00 | 146.00 | 126.67 | 111.00 |
| ARHGEF11 | 2015.00 | 1973.00 | 2116.00 | 2034.67 | 1906.00 |
| ARHGEF12 | 2087.00 | 1946.00 | 2350.00 | 2127.67 | 1870.00 |
| ARHGEF17 | 1905.00 | 1884.00 | 2046.00 | 1945.00 | 1108.00 |
| ARHGEF18 | 1299.00 | 1234.00 | 1441.00 | 1324.67 | 1015.00 |
| ARHGEF19 | 296.00 | 301.00 | 333.00 | 310.00 | 278.00 |
| ARHGEF2 | 9306.00 | 8858.00 | 9361.00 | 9175.00 | 7586.00 |
| ARHGEF25 | 999.00 | 960.00 | 1059.00 | 1006.00 | 1201.00 |
| ARHGEF26 | 18.00 | 21.00 | 27.00 | 22.00 | 14.00 |
| ARHGEF26-AS1 | 23.00 | 20.00 | 17.00 | 20.00 | 17.00 |
| ARHGEF28 | 3012.00 | 2964.00 | 3315.00 | 3097.00 | 2680.00 |
| ARHGEF3 | 121.00 | 84.00 | 109.00 | 104.67 | 113.00 |
| ARHGEF37 | 5.00 | 5.00 | 6.00 | 5.33 | 7.00 |
| ARHGEF39 | 218.00 | 203.00 | 210.00 | 210.33 | 257.00 |
| ARHGEF40 | 557.00 | 614.00 | 651.00 | 607.33 | 525.00 |
| ARHGEF6 | 331.00 | 301.00 | 344.00 | 325.33 | 244.00 |
| ARHGEF7 | 1032.00 | 924.00 | 1045.00 | 1000.33 | 1037.00 |
| ARHGEF9 | 1144.00 | 1115.00 | 1256.00 | 1171.67 | 1089.00 |
| ARID1A | 2987.00 | 2847.00 | 3013.00 | 2949.00 | 1150.00 |
| ARID1B | 1971.00 | 1815.00 | 2131.00 | 1972.33 | 1810.00 |
| ARID2 | 629.00 | 602.00 | 734.00 | 655.00 | 669.00 |
| ARID3A | 102.00 | 83.00 | 93.00 | 92.67 | 119.00 |
| ARID3B | 474.00 | 466.00 | 576.00 | 505.33 | 380.00 |
| ARID4A | 412.00 | 326.00 | 403.00 | 380.33 | 366.00 |
| ARID4B | 786.00 | 756.00 | 819.00 | 787.00 | 751.00 |
| ARID5A | 130.00 | 129.00 | 119.00 | 126.00 | 73.00 |
| ARID5B | 714.00 | 670.00 | 742.00 | 708.67 | 394.00 |
| ARIH1 | 1382.00 | 1376.00 | 1501.00 | 1419.67 | 1408.00 |
| ARIH2 | 2685.00 | 2711.00 | 2816.00 | 2737.33 | 3283.00 |
| ARIH2OS | 5.00 | 17.00 | 11.00 | 11.00 | 14.00 |
| ARL1 | 1254.00 | 1088.00 | 1333.00 | 1225.00 | 1484.00 |
| ARL10 | 98.00 | 94.00 | 122.00 | 104.67 | 41.00 |
| ARL13B | 378.00 | 317.00 | 338.00 | 344.33 | 395.00 |
| ARL14EP | 633.00 | 518.00 | 566.00 | 572.33 | 727.00 |
| ARL14EPL | 7.00 | 1.00 | 4.00 | 4.00 | 1.00 |
| ARL15 | 210.00 | 140.00 | 189.00 | 179.67 | 203.00 |
| ARL16 | 658.00 | 615.00 | 570.00 | 614.33 | 642.00 |

| | | | | | |
|---|---|---|---|---|---|
| ARL17A | 38.00 | 44.00 | 38.00 | 40.00 | 29.00 |
| ARL2 | 1625.00 | 1603.00 | 1772.00 | 1666.67 | 2190.00 |
| ARL2BP | 1009.00 | 930.00 | 1015.00 | 984.67 | 1095.00 |
| ARL3 | 364.00 | 382.00 | 422.00 | 389.33 | 467.00 |
| ARL4A | 386.00 | 374.00 | 449.00 | 403.00 | 488.00 |
| ARL4C | 708.00 | 672.00 | 784.00 | 721.33 | 658.00 |
| ARL4D | 132.00 | 120.00 | 115.00 | 122.33 | 107.00 |
| ARL5A | 553.00 | 451.00 | 591.00 | 531.67 | 672.00 |
| ARL5B | 294.00 | 240.00 | 326.00 | 286.67 | 276.00 |
| ARL6 | 174.00 | 170.00 | 189.00 | 177.67 | 186.00 |
| ARL6IP1 | 3346.00 | 3092.00 | 3103.00 | 3180.33 | 3655.00 |
| ARL6IP4 | 1874.00 | 1896.00 | 1932.00 | 1900.67 | 2347.00 |
| ARL6IP5 | 1391.00 | 1305.00 | 1335.00 | 1343.67 | 1668.00 |
| ARL6IP6 | 527.00 | 538.00 | 558.00 | 541.00 | 570.00 |
| ARL8A | 395.00 | 408.00 | 423.00 | 408.67 | 432.00 |
| ARL8B | 1492.00 | 1489.00 | 1644.00 | 1541.67 | 1911.00 |
| ARL9 | 7.00 | 4.00 | 8.00 | 6.33 | 7.00 |
| ARMC1 | 827.00 | 804.00 | 908.00 | 846.33 | 994.00 |
| ARMC10 | 234.00 | 167.00 | 206.00 | 202.33 | 219.00 |
| ARMC12 | 22.00 | 21.00 | 15.00 | 19.33 | 22.00 |
| ARMC2 | 59.00 | 46.00 | 39.00 | 48.00 | 51.00 |
| ARMC4 | 626.00 | 560.00 | 685.00 | 623.67 | 1265.00 |
| ARMC5 | 386.00 | 385.00 | 442.00 | 404.33 | 396.00 |
| ARMC6 | 1179.00 | 1233.00 | 1299.00 | 1237.00 | 1494.00 |
| ARMC7 | 169.00 | 185.00 | 185.00 | 179.67 | 237.00 |
| ARMC8 | 850.00 | 816.00 | 885.00 | 850.33 | 881.00 |
| ARMC9 | 722.00 | 723.00 | 803.00 | 749.33 | 665.00 |
| ARMCX1 | 1019.00 | 1089.00 | 1000.00 | 1036.00 | 1094.00 |
| ARMCX2 | 35.00 | 39.00 | 49.00 | 41.00 | 66.00 |
| ARMCX3 | 810.00 | 810.00 | 834.00 | 818.00 | 786.00 |
| ARMCX4 | 943.00 | 927.00 | 1020.00 | 963.33 | 849.00 |
| ARMCX5 | 496.00 | 467.00 | 440.00 | 467.67 | 469.00 |
| ARMCX5-GPRASP2 | 45.00 | 38.00 | 52.00 | 45.00 | 32.00 |
| ARMCX6 | 247.00 | 310.00 | 269.00 | 275.33 | 261.00 |
| ARMT1 | 493.00 | 480.00 | 533.00 | 502.00 | 658.00 |
| ARNT | 660.00 | 621.00 | 723.00 | 668.00 | 615.00 |
| ARNT2 | 175.00 | 165.00 | 172.00 | 170.67 | 185.00 |
| ARNTL | 406.00 | 348.00 | 368.00 | 374.00 | 334.00 |
| ARNTL2 | 2260.00 | 2174.00 | 2485.00 | 2306.33 | 4664.00 |
| ARPC1A | 2815.00 | 2708.00 | 3037.00 | 2853.33 | 4355.00 |
| ARPC1B | 3166.00 | 3190.00 | 3440.00 | 3265.33 | 4387.00 |
| ARPC2 | 4158.00 | 4168.00 | 4446.00 | 4257.33 | 5416.00 |
| ARPC3 | 2497.00 | 2662.00 | 2738.00 | 2632.33 | 3596.00 |
| ARPC4 | 1438.00 | 1364.00 | 1558.00 | 1453.33 | 1881.00 |
| ARPC5 | 2192.00 | 2007.00 | 2160.00 | 2119.67 | 3288.00 |
| ARPC5L | 761.00 | 792.00 | 788.00 | 780.33 | 1056.00 |
| ARPIN | 450.00 | 439.00 | 435.00 | 441.33 | 459.00 |
| ARPP19 | 2097.00 | 1966.00 | 2423.00 | 2162.00 | 2343.00 |

| | | | | | |
|---|---|---|---|---|---|
| ARPP21 | 6.00 | 13.00 | 12.00 | 10.33 | 20.00 |
| ARRB1 | 80.00 | 50.00 | 51.00 | 60.33 | 73.00 |
| ARRB2 | 406.00 | 421.00 | 440.00 | 422.33 | 637.00 |
| ARRDC1 | 560.00 | 496.00 | 548.00 | 534.67 | 615.00 |
| ARRDC1-AS1 | 544.00 | 595.00 | 549.00 | 562.67 | 597.00 |
| ARRDC2 | 387.00 | 393.00 | 381.00 | 387.00 | 441.00 |
| ARRDC3 | 671.00 | 627.00 | 809.00 | 702.33 | 644.00 |
| ARRDC3-AS1 | 88.00 | 70.00 | 63.00 | 73.67 | 84.00 |
| ARRDC4 | 37.00 | 43.00 | 43.00 | 41.00 | 42.00 |
| ARSA | 687.00 | 782.00 | 809.00 | 759.33 | 623.00 |
| ARSB | 464.00 | 482.00 | 488.00 | 478.00 | 449.00 |
| ARSD | 230.00 | 277.00 | 238.00 | 248.33 | 196.00 |
| ARSE | 28.00 | 31.00 | 39.00 | 32.67 | 39.00 |
| ARSG | 178.00 | 178.00 | 209.00 | 188.33 | 134.00 |
| ARSJ | 2033.00 | 1857.00 | 2177.00 | 2022.33 | 1487.00 |
| ARSK | 466.00 | 449.00 | 481.00 | 465.33 | 521.00 |
| ARTN | 58.00 | 82.00 | 92.00 | 77.33 | 87.00 |
| ARV1 | 514.00 | 459.00 | 480.00 | 484.33 | 608.00 |
| ARVCF | 63.00 | 81.00 | 78.00 | 74.00 | 75.00 |
| ASAH1 | 1634.00 | 1638.00 | 1671.00 | 1647.67 | 834.00 |
| ASAH2 | 5.00 | 7.00 | 12.00 | 8.00 | 8.00 |
| ASAH2B | 28.00 | 20.00 | 28.00 | 25.33 | 30.00 |
| ASAP1 | 8593.00 | 8045.00 | 9418.00 | 8685.33 | 6835.00 |
| ASAP1-IT2 | 8.00 | 9.00 | 14.00 | 10.33 | 17.00 |
| ASAP2 | 2632.00 | 2433.00 | 2803.00 | 2622.67 | 3160.00 |
| ASAP3 | 55.00 | 80.00 | 82.00 | 72.33 | 31.00 |
| ASB1 | 17369.00 | 16991.00 | 18783.00 | 17714.33 | 15037.00 |
| ASB13 | 576.00 | 634.00 | 648.00 | 619.33 | 719.00 |
| ASB16-AS1 | 190.00 | 193.00 | 202.00 | 195.00 | 181.00 |
| ASB3 | 43.00 | 29.00 | 35.00 | 35.67 | 39.00 |
| ASB6 | 923.00 | 931.00 | 1015.00 | 956.33 | 1029.00 |
| ASB7 | 469.00 | 459.00 | 514.00 | 480.67 | 480.00 |
| ASB8 | 335.00 | 302.00 | 340.00 | 325.67 | 362.00 |
| ASB9 | 60.00 | 46.00 | 42.00 | 49.33 | 149.00 |
| ASCC1 | 622.00 | 472.00 | 617.00 | 570.33 | 647.00 |
| ASCC2 | 1287.00 | 1245.00 | 1405.00 | 1312.33 | 1395.00 |
| ASCC3 | 1883.00 | 1697.00 | 2126.00 | 1902.00 | 1832.00 |
| ASF1A | 449.00 | 468.00 | 465.00 | 460.67 | 582.00 |
| ASF1B | 437.00 | 410.00 | 346.00 | 397.67 | 461.00 |
| ASGR1 | 78.00 | 83.00 | 77.00 | 79.33 | 66.00 |
| ASH1L | 2028.00 | 1903.00 | 2127.00 | 2019.33 | 1600.00 |
| ASH1L-AS1 | 61.00 | 59.00 | 64.00 | 61.33 | 46.00 |
| ASH2L | 814.00 | 792.00 | 850.00 | 818.67 | 1038.00 |
| ASIC1 | 1008.00 | 1001.00 | 1073.00 | 1027.33 | 1015.00 |
| ASIC2 | 7.00 | 2.00 | 1.00 | 3.33 | 2.00 |
| ASIC3 | 93.00 | 100.00 | 139.00 | 110.67 | 69.00 |
| ASL | 804.00 | 896.00 | 857.00 | 852.33 | 859.00 |
| ASMTL | 5.00 | 2.00 | 0.00 | 2.33 | 1.00 |

| | | | | | |
|---|---|---|---|---|---|
| ASNA1 | 1127.00 | 1158.00 | 1169.00 | 1151.33 | 1421.00 |
| ASNS | 10340.00 | 9684.00 | 10549.00 | 10191.00 | 12124.00 |
| ASNSD1 | 1076.00 | 1004.00 | 1149.00 | 1076.33 | 1359.00 |
| ASPH | 10807.00 | 10239.00 | 11211.00 | 10752.33 | 10800.00 |
| ASPHD1 | 199.00 | 160.00 | 184.00 | 181.00 | 243.00 |
| ASPHD2 | 64.00 | 64.00 | 63.00 | 63.67 | 65.00 |
| ASPM | 2553.00 | 2360.00 | 2459.00 | 2457.33 | 2348.00 |
| ASPRV1 | 15.00 | 12.00 | 5.00 | 10.67 | 7.00 |
| ASPSCR1 | 1069.00 | 1147.00 | 1121.00 | 1112.33 | 1256.00 |
| ASRGL1 | 18.00 | 17.00 | 13.00 | 16.00 | 29.00 |
| ASS1 | 8.00 | 7.00 | 5.00 | 6.67 | 20.00 |
| ASTE1 | 164.00 | 118.00 | 183.00 | 155.00 | 140.00 |
| ASTN2 | 84.00 | 60.00 | 84.00 | 76.00 | 103.00 |
| ASUN | 1708.00 | 1582.00 | 2014.00 | 1768.00 | 2277.00 |
| ASXL1 | 4310.00 | 4269.00 | 4542.00 | 4373.67 | 4062.00 |
| ASXL2 | 809.00 | 631.00 | 798.00 | 746.00 | 520.00 |
| ASXL3 | 12.00 | 8.00 | 4.00 | 8.00 | 3.00 |
| ATAD1 | 2170.00 | 2051.00 | 2295.00 | 2172.00 | 2465.00 |
| ATAD2 | 1285.00 | 1225.00 | 1261.00 | 1257.00 | 1876.00 |
| ATAD2B | 343.00 | 358.00 | 366.00 | 355.67 | 356.00 |
| ATAD3A | 890.00 | 851.00 | 870.00 | 870.33 | 1082.00 |
| ATAD3B | 812.00 | 822.00 | 819.00 | 817.67 | 1011.00 |
| ATAD5 | 397.00 | 374.00 | 425.00 | 398.67 | 432.00 |
| ATE1 | 1050.00 | 767.00 | 973.00 | 930.00 | 998.00 |
| ATE1-AS1 | 43.00 | 33.00 | 36.00 | 37.33 | 32.00 |
| ATF1 | 499.00 | 423.00 | 541.00 | 487.67 | 585.00 |
| ATF2 | 934.00 | 916.00 | 999.00 | 949.67 | 956.00 |
| ATF3 | 153.00 | 157.00 | 167.00 | 159.00 | 177.00 |
| ATF4 | 9670.00 | 9296.00 | 10399.00 | 9788.33 | 12126.00 |
| ATF5 | 213.00 | 196.00 | 253.00 | 220.67 | 258.00 |
| ATF6 | 3179.00 | 2934.00 | 3215.00 | 3109.33 | 2871.00 |
| ATF6B | 94.00 | 111.00 | 102.00 | 102.33 | 134.00 |
| ATF7 | 1134.00 | 1098.00 | 1203.00 | 1145.00 | 881.00 |
| ATF7IP | 1787.00 | 1639.00 | 1784.00 | 1736.67 | 1671.00 |
| ATF7IP2 | 5.00 | 12.00 | 23.00 | 13.33 | 19.00 |
| ATG10 | 90.00 | 94.00 | 120.00 | 101.33 | 134.00 |
| ATG101 | 910.00 | 880.00 | 949.00 | 913.00 | 1103.00 |
| ATG12 | 1097.00 | 1090.00 | 1191.00 | 1126.00 | 1313.00 |
| ATG13 | 1457.00 | 1333.00 | 1477.00 | 1422.33 | 1213.00 |
| ATG14 | 610.00 | 574.00 | 716.00 | 633.33 | 638.00 |
| ATG16L1 | 724.00 | 741.00 | 798.00 | 754.33 | 757.00 |
| ATG16L2 | 1120.00 | 1245.00 | 1255.00 | 1206.67 | 1478.00 |
| ATG2A | 1925.00 | 1883.00 | 1956.00 | 1921.33 | 1596.00 |
| ATG2B | 1376.00 | 1174.00 | 1432.00 | 1327.33 | 1208.00 |
| ATG3 | 1138.00 | 1226.00 | 1293.00 | 1219.00 | 1537.00 |
| ATG4A | 274.00 | 308.00 | 293.00 | 291.67 | 237.00 |
| ATG4B | 1218.00 | 1235.00 | 1239.00 | 1230.67 | 1236.00 |
| ATG4C | 423.00 | 339.00 | 403.00 | 388.33 | 454.00 |

| | | | | | |
|---|---|---|---|---|---|
| ATG4D | 325.00 | 298.00 | 361.00 | 328.00 | 414.00 |
| ATG5 | 687.00 | 640.00 | 721.00 | 682.67 | 561.00 |
| ATG7 | 1031.00 | 980.00 | 1080.00 | 1030.33 | 1240.00 |
| ATG9A | 2276.00 | 2144.00 | 2209.00 | 2209.67 | 2096.00 |
| ATG9B | 11.00 | 13.00 | 9.00 | 11.00 | 9.00 |
| ATHL1 | 1734.00 | 1862.00 | 2034.00 | 1876.67 | 1874.00 |
| ATIC | 4341.00 | 4143.00 | 4696.00 | 4393.33 | 5206.00 |
| ATL1 | 5.00 | 5.00 | 0.00 | 3.33 | 3.00 |
| ATL2 | 1790.00 | 1670.00 | 1840.00 | 1766.67 | 2005.00 |
| ATL3 | 2983.00 | 2911.00 | 3197.00 | 3030.33 | 3315.00 |
| ATM | 2832.00 | 2594.00 | 3159.00 | 2861.67 | 2422.00 |
| ATMIN | 1430.00 | 1317.00 | 1556.00 | 1434.33 | 1383.00 |
| ATN1 | 3626.00 | 3483.00 | 4064.00 | 3724.33 | 3172.00 |
| ATOH8 | 101.00 | 97.00 | 107.00 | 101.67 | 49.00 |
| ATOX1 | 865.00 | 831.00 | 903.00 | 866.33 | 1258.00 |
| ATP10A | 43.00 | 36.00 | 49.00 | 42.67 | 42.00 |
| ATP10D | 809.00 | 817.00 | 864.00 | 830.00 | 617.00 |
| ATP11A | 1250.00 | 1179.00 | 1261.00 | 1230.00 | 943.00 |
| ATP11B | 1755.00 | 1544.00 | 1686.00 | 1661.67 | 1884.00 |
| ATP11C | 995.00 | 847.00 | 1078.00 | 973.33 | 898.00 |
| ATP13A1 | 3042.00 | 3128.00 | 3197.00 | 3122.33 | 2983.00 |
| ATP13A2 | 1065.00 | 1015.00 | 1020.00 | 1033.33 | 848.00 |
| ATP13A3 | 5022.00 | 4452.00 | 5290.00 | 4921.33 | 4622.00 |
| ATP13A4 | 5.00 | 6.00 | 8.00 | 6.33 | 10.00 |
| ATP1A1 | 19183.00 | 19613.00 | 19898.00 | 19564.67 | 17522.00 |
| ATP1A1-AS1 | 36.00 | 40.00 | 46.00 | 40.67 | 46.00 |
| ATP1A2 | 196.00 | 168.00 | 217.00 | 193.67 | 51.00 |
| ATP1B1 | 1353.00 | 1399.00 | 1437.00 | 1396.33 | 1263.00 |
| ATP1B3 | 5319.00 | 5133.00 | 5061.00 | 5171.00 | 5919.00 |
| ATP2A1 | 49.00 | 51.00 | 72.00 | 57.33 | 55.00 |
| ATP2A1-AS1 | 71.00 | 80.00 | 92.00 | 81.00 | 114.00 |
| ATP2A2 | 11257.00 | 10661.00 | 11292.00 | 11070.00 | 10307.00 |
| ATP2B1 | 2895.00 | 2695.00 | 3083.00 | 2891.00 | 2102.00 |
| ATP2B2 | 1771.00 | 1690.00 | 1864.00 | 1775.00 | 1499.00 |
| ATP2B4 | 1393.00 | 1413.00 | 1429.00 | 1411.67 | 985.00 |
| ATP2C1 | 4138.00 | 3992.00 | 4120.00 | 4083.33 | 4418.00 |
| ATP5A1 | 5392.00 | 5112.00 | 5899.00 | 5467.67 | 6405.00 |
| ATP5B | 13786.00 | 13186.00 | 14701.00 | 13891.00 | 17368.00 |
| ATP5C1 | 2371.00 | 2363.00 | 2555.00 | 2429.67 | 2838.00 |
| ATP5D | 1130.00 | 1129.00 | 1178.00 | 1145.67 | 1496.00 |
| ATP5E | 857.00 | 1063.00 | 1023.00 | 981.00 | 1550.00 |
| ATP5EP2 | 12.00 | 6.00 | 9.00 | 9.00 | 19.00 |
| ATP5F1 | 2691.00 | 2614.00 | 2906.00 | 2737.00 | 3783.00 |
| ATP5G1 | 1169.00 | 1201.00 | 1201.00 | 1190.33 | 1545.00 |
| ATP5G2 | 3101.00 | 3019.00 | 3277.00 | 3132.33 | 3913.00 |
| ATP5G3 | 2167.00 | 2212.00 | 2253.00 | 2210.67 | 3224.00 |
| ATP5H | 1370.00 | 1359.00 | 1318.00 | 1349.00 | 1824.00 |
| ATP5I | 556.00 | 552.00 | 523.00 | 543.67 | 765.00 |

| | | | | | |
|---|---|---|---|---|---|
| ATP5J | 1092.00 | 1120.00 | 1136.00 | 1116.00 | 1551.00 |
| ATP5J2 | 691.00 | 757.00 | 788.00 | 745.33 | 1021.00 |
| ATP5L | 1626.00 | 1582.00 | 1608.00 | 1605.33 | 2240.00 |
| ATP5O | 1779.00 | 1794.00 | 1982.00 | 1851.67 | 2484.00 |
| ATP5S | 173.00 | 145.00 | 155.00 | 157.67 | 191.00 |
| ATP5SL | 467.00 | 468.00 | 495.00 | 476.67 | 546.00 |
| ATP6AP1 | 2374.00 | 2556.00 | 2326.00 | 2418.67 | 2506.00 |
| ATP6AP1L | 203.00 | 253.00 | 281.00 | 245.67 | 251.00 |
| ATP6AP2 | 2664.00 | 2702.00 | 2571.00 | 2645.67 | 2528.00 |
| ATP6V0A1 | 2459.00 | 2489.00 | 2651.00 | 2533.00 | 2178.00 |
| ATP6V0A2 | 1334.00 | 1286.00 | 1410.00 | 1343.33 | 1207.00 |
| ATP6V0B | 2408.00 | 2344.00 | 2208.00 | 2320.00 | 2599.00 |
| ATP6V0C | 1558.00 | 1472.00 | 1371.00 | 1467.00 | 1717.00 |
| ATP6V0D1 | 1041.00 | 984.00 | 1019.00 | 1014.67 | 1250.00 |
| ATP6V0E1 | 1747.00 | 1653.00 | 1603.00 | 1667.67 | 1880.00 |
| ATP6V0E2 | 789.00 | 752.00 | 712.00 | 751.00 | 849.00 |
| ATP6V0E2-AS1 | 205.00 | 207.00 | 228.00 | 213.33 | 191.00 |
| ATP6V1A | 976.00 | 877.00 | 1028.00 | 960.33 | 1065.00 |
| ATP6V1B1 | 118.00 | 108.00 | 110.00 | 112.00 | 83.00 |
| ATP6V1B1-AS1 | 90.00 | 88.00 | 84.00 | 87.33 | 98.00 |
| ATP6V1B2 | 1159.00 | 1062.00 | 1213.00 | 1144.67 | 1186.00 |
| ATP6V1C1 | 2297.00 | 2174.00 | 2490.00 | 2320.33 | 2763.00 |
| ATP6V1C2 | 5.00 | 0.00 | 1.00 | 2.00 | 5.00 |
| ATP6V1D | 694.00 | 682.00 | 775.00 | 717.00 | 1001.00 |
| ATP6V1E1 | 1430.00 | 1331.00 | 1474.00 | 1411.67 | 1676.00 |
| ATP6V1E2 | 74.00 | 73.00 | 80.00 | 75.67 | 109.00 |
| ATP6V1F | 1332.00 | 1247.00 | 1328.00 | 1302.33 | 1820.00 |
| ATP6V1G1 | 916.00 | 928.00 | 959.00 | 934.33 | 1321.00 |
| ATP6V1H | 1296.00 | 1261.00 | 1215.00 | 1257.33 | 1445.00 |
| ATP7A | 435.00 | 388.00 | 464.00 | 429.00 | 348.00 |
| ATP7B | 574.00 | 555.00 | 527.00 | 552.00 | 439.00 |
| ATP8A1 | 35.00 | 37.00 | 34.00 | 35.33 | 40.00 |
| ATP8A2 | 37.00 | 33.00 | 38.00 | 36.00 | 31.00 |
| ATP8B1 | 82.00 | 77.00 | 113.00 | 90.67 | 41.00 |
| ATP8B2 | 2119.00 | 2014.00 | 2003.00 | 2045.33 | 1743.00 |
| ATP8B3 | 72.00 | 63.00 | 58.00 | 64.33 | 93.00 |
| ATP8B5P | 10.00 | 14.00 | 15.00 | 13.00 | 8.00 |
| ATP9A | 698.00 | 673.00 | 721.00 | 697.33 | 664.00 |
| ATP9B | 286.00 | 290.00 | 276.00 | 284.00 | 274.00 |
| ATPAF1 | 670.00 | 632.00 | 633.00 | 645.00 | 853.00 |
| ATPAF2 | 200.00 | 207.00 | 195.00 | 200.67 | 258.00 |
| ATPIF1 | 982.00 | 947.00 | 985.00 | 971.33 | 1233.00 |
| ATR | 1940.00 | 1955.00 | 2200.00 | 2031.67 | 1978.00 |
| ATRAID | 2782.00 | 2829.00 | 2769.00 | 2793.33 | 3145.00 |
| ATRIP | 381.00 | 374.00 | 387.00 | 380.67 | 368.00 |
| ATRN | 2893.00 | 2622.00 | 3179.00 | 2898.00 | 2476.00 |
| ATRNL1 | 43.00 | 35.00 | 37.00 | 38.33 | 36.00 |
| ATRX | 2404.00 | 2130.00 | 2744.00 | 2426.00 | 1961.00 |

| | | | | | |
|---|---|---|---|---|---|
| ATXN1 | 790.00 | 702.00 | 868.00 | 786.67 | 516.00 |
| ATXN10 | 1572.00 | 1460.00 | 1650.00 | 1560.67 | 1837.00 |
| ATXN1L | 1004.00 | 961.00 | 1036.00 | 1000.33 | 830.00 |
| ATXN2 | 1472.00 | 1262.00 | 1456.00 | 1396.67 | 1353.00 |
| ATXN2L | 8450.00 | 8335.00 | 8636.00 | 8473.67 | 7851.00 |
| ATXN3 | 334.00 | 341.00 | 392.00 | 355.67 | 381.00 |
| ATXN7 | 717.00 | 722.00 | 795.00 | 744.67 | 734.00 |
| ATXN7L1 | 533.00 | 460.00 | 585.00 | 526.00 | 467.00 |
| ATXN7L2 | 404.00 | 408.00 | 451.00 | 421.00 | 414.00 |
| ATXN7L3 | 1501.00 | 1497.00 | 1499.00 | 1499.00 | 1414.00 |
| ATXN7L3B | 2502.00 | 2395.00 | 2580.00 | 2492.33 | 2830.00 |
| AUH | 115.00 | 124.00 | 130.00 | 123.00 | 119.00 |
| AUNIP | 120.00 | 120.00 | 126.00 | 122.00 | 143.00 |
| AUP1 | 4936.00 | 5142.00 | 5208.00 | 5095.33 | 5806.00 |
| AURKA | 1519.00 | 1411.00 | 1512.00 | 1480.67 | 2231.00 |
| AURKAIP1 | 1159.00 | 1217.00 | 1250.00 | 1208.67 | 1583.00 |
| AURKAPS1 | 35.00 | 44.00 | 30.00 | 36.33 | 28.00 |
| AURKB | 1170.00 | 1186.00 | 1122.00 | 1159.33 | 1737.00 |
| AURKC | 7.00 | 19.00 | 26.00 | 17.33 | 18.00 |
| AUTS2 | 43.00 | 26.00 | 36.00 | 35.00 | 39.00 |
| AVEN | 388.00 | 362.00 | 388.00 | 379.33 | 400.00 |
| AVIL | 77.00 | 105.00 | 120.00 | 100.67 | 86.00 |
| AVL9 | 2825.00 | 2468.00 | 2891.00 | 2728.00 | 2792.00 |
| AVPI1 | 105.00 | 116.00 | 113.00 | 111.33 | 111.00 |
| AWAT2 | 22.00 | 29.00 | 37.00 | 29.33 | 5.00 |
| AXDND1 | 6.00 | 1.00 | 5.00 | 4.00 | 5.00 |
| AXIN1 | 1048.00 | 1040.00 | 1061.00 | 1049.67 | 1159.00 |
| AXIN2 | 19.00 | 11.00 | 12.00 | 14.00 | 22.00 |
| AXL | 27362.00 | 26150.00 | 27807.00 | 27106.33 | 17066.00 |
| AZGP1 | 4.00 | 3.00 | 6.00 | 4.33 | 3.00 |
| AZI2 | 876.00 | 797.00 | 979.00 | 884.00 | 1091.00 |
| AZIN1 | 4309.00 | 3725.00 | 4522.00 | 4185.33 | 4984.00 |
| AZIN1-AS1 | 20.00 | 16.00 | 16.00 | 17.33 | 29.00 |
| AZIN2 | 83.00 | 95.00 | 123.00 | 100.33 | 101.00 |
| B2M | 1269.00 | 1269.00 | 1151.00 | 1229.67 | 1668.00 |
| B3GALNT1 | 214.00 | 251.00 | 224.00 | 229.67 | 267.00 |
| B3GALNT2 | 1163.00 | 1129.00 | 1195.00 | 1162.33 | 1428.00 |
| B3GALT1 | 58.00 | 44.00 | 49.00 | 50.33 | 51.00 |
| B3GALT5 | 68.00 | 60.00 | 68.00 | 65.33 | 43.00 |
| B3GALT5-AS1 | 23.00 | 19.00 | 16.00 | 19.33 | 22.00 |
| B3GALT6 | 1052.00 | 1026.00 | 1097.00 | 1058.33 | 1136.00 |
| B3GAT2 | 6.00 | 4.00 | 1.00 | 3.67 | 5.00 |
| B3GAT3 | 1593.00 | 1641.00 | 1574.00 | 1602.67 | 1733.00 |
| B3GLCT | 333.00 | 354.00 | 316.00 | 334.33 | 360.00 |
| B3GNT2 | 501.00 | 487.00 | 495.00 | 494.33 | 463.00 |
| B3GNT5 | 464.00 | 415.00 | 464.00 | 447.67 | 504.00 |
| B3GNT8 | 4.00 | 1.00 | 0.00 | 1.67 | 1.00 |
| B3GNT9 | 64.00 | 63.00 | 65.00 | 64.00 | 45.00 |

| | | | | | |
|---|---|---|---|---|---|
| B3GNTL1 | 140.00 | 140.00 | 161.00 | 147.00 | 215.00 |
| B4GALNT1 | 51.00 | 48.00 | 40.00 | 46.33 | 51.00 |
| B4GALNT2 | 16.00 | 15.00 | 13.00 | 14.67 | 8.00 |
| B4GALT1 | 1910.00 | 1776.00 | 1759.00 | 1815.00 | 1758.00 |
| B4GALT1-AS1 | 5.00 | 8.00 | 13.00 | 8.67 | 10.00 |
| B4GALT2 | 1308.00 | 1295.00 | 1303.00 | 1302.00 | 1527.00 |
| B4GALT3 | 1541.00 | 1559.00 | 1615.00 | 1571.67 | 1533.00 |
| B4GALT4 | 436.00 | 415.00 | 419.00 | 423.33 | 469.00 |
| B4GALT5 | 2024.00 | 1898.00 | 1932.00 | 1951.33 | 2016.00 |
| B4GALT6 | 577.00 | 492.00 | 649.00 | 572.67 | 707.00 |
| B4GALT7 | 917.00 | 871.00 | 895.00 | 894.33 | 872.00 |
| B4GAT1 | 2203.00 | 2248.00 | 2226.00 | 2225.67 | 1954.00 |
| B9D1 | 319.00 | 318.00 | 310.00 | 315.67 | 393.00 |
| B9D2 | 49.00 | 52.00 | 43.00 | 48.00 | 76.00 |
| BAALC-AS1 | 5.00 | 8.00 | 3.00 | 5.33 | 3.00 |
| BAAT | 4.00 | 3.00 | 1.00 | 2.67 | 2.00 |
| BABAM1 | 1491.00 | 1465.00 | 1562.00 | 1506.00 | 1753.00 |
| BACE1 | 1458.00 | 1489.00 | 1490.00 | 1479.00 | 1127.00 |
| BACE1-AS | 95.00 | 93.00 | 116.00 | 101.33 | 99.00 |
| BACE2 | 3023.00 | 3068.00 | 3004.00 | 3031.67 | 3592.00 |
| BACH1 | 2225.00 | 1932.00 | 2504.00 | 2220.33 | 1990.00 |
| BACH1-IT2 | 5.00 | 5.00 | 10.00 | 6.67 | 3.00 |
| BACH2 | 118.00 | 122.00 | 102.00 | 114.00 | 95.00 |
| BAD | 565.00 | 593.00 | 624.00 | 594.00 | 748.00 |
| BAG1 | 397.00 | 382.00 | 435.00 | 404.67 | 471.00 |
| BAG2 | 662.00 | 586.00 | 681.00 | 643.00 | 865.00 |
| BAG3 | 1316.00 | 1307.00 | 1349.00 | 1324.00 | 1239.00 |
| BAG4 | 356.00 | 322.00 | 352.00 | 343.33 | 385.00 |
| BAG5 | 982.00 | 907.00 | 1061.00 | 983.33 | 1042.00 |
| BAG6 | 7.00 | 2.00 | 6.00 | 5.00 | 4.00 |
| BAHCC1 | 165.00 | 166.00 | 216.00 | 182.33 | 165.00 |
| BAHD1 | 861.00 | 869.00 | 833.00 | 854.33 | 953.00 |
| BAIAP2 | 765.00 | 672.00 | 739.00 | 725.33 | 766.00 |
| BAIAP2-AS1 | 245.00 | 269.00 | 310.00 | 274.67 | 267.00 |
| BAIAP2L1 | 113.00 | 110.00 | 150.00 | 124.33 | 150.00 |
| BAIAP2L2 | 107.00 | 131.00 | 154.00 | 130.67 | 217.00 |
| BAIAP3 | 5.00 | 5.00 | 13.00 | 7.67 | 1.00 |
| BAK1 | 394.00 | 413.00 | 428.00 | 411.67 | 504.00 |
| BAMBI | 84.00 | 87.00 | 82.00 | 84.33 | 77.00 |
| BANF1 | 2600.00 | 2753.00 | 2732.00 | 2695.00 | 3585.00 |
| BANP | 540.00 | 557.00 | 622.00 | 573.00 | 625.00 |
| BAP1 | 1781.00 | 1729.00 | 1804.00 | 1771.33 | 1724.00 |
| BARD1 | 218.00 | 224.00 | 214.00 | 218.67 | 262.00 |
| BARHL2 | 18.00 | 18.00 | 21.00 | 19.00 | 26.00 |
| BASP1 | 4932.00 | 4834.00 | 5454.00 | 5073.33 | 5475.00 |
| BATF | 43.00 | 31.00 | 29.00 | 34.33 | 17.00 |
| BATF2 | 77.00 | 51.00 | 68.00 | 65.33 | 85.00 |
| BATF3 | 16.00 | 21.00 | 27.00 | 21.33 | 22.00 |

| | | | | | |
|---|---|---|---|---|---|
| BAX | 1112.00 | 1147.00 | 1137.00 | 1132.00 | 1356.00 |
| BAZ1A | 1970.00 | 1819.00 | 2022.00 | 1937.00 | 2022.00 |
| BAZ1B | 3399.00 | 3169.00 | 3794.00 | 3454.00 | 3336.00 |
| BAZ2A | 2704.00 | 2783.00 | 3110.00 | 2865.67 | 2224.00 |
| BAZ2B | 688.00 | 618.00 | 719.00 | 675.00 | 514.00 |
| BBC3 | 129.00 | 140.00 | 141.00 | 136.67 | 127.00 |
| BBIP1 | 401.00 | 344.00 | 398.00 | 381.00 | 503.00 |
| BBS1 | 977.00 | 926.00 | 968.00 | 957.00 | 902.00 |
| BBS10 | 917.00 | 750.00 | 898.00 | 855.00 | 815.00 |
| BBS12 | 97.00 | 75.00 | 112.00 | 94.67 | 89.00 |
| BBS2 | 1381.00 | 1387.00 | 1533.00 | 1433.67 | 1285.00 |
| BBS4 | 324.00 | 312.00 | 339.00 | 325.00 | 375.00 |
| BBS5 | 348.00 | 332.00 | 344.00 | 341.33 | 383.00 |
| BBS7 | 487.00 | 478.00 | 540.00 | 501.67 | 464.00 |
| BBS9 | 274.00 | 266.00 | 297.00 | 279.00 | 286.00 |
| BBX | 2476.00 | 2130.00 | 2529.00 | 2378.33 | 2213.00 |
| BCAM | 1378.00 | 1517.00 | 1657.00 | 1517.33 | 1379.00 |
| BCAP29 | 1758.00 | 1546.00 | 1575.00 | 1626.33 | 1800.00 |
| BCAP31 | 7119.00 | 7060.00 | 6762.00 | 6980.33 | 7225.00 |
| BCAR1 | 3621.00 | 3742.00 | 3904.00 | 3755.67 | 5150.00 |
| BCAR3 | 3986.00 | 3803.00 | 4199.00 | 3996.00 | 4253.00 |
| BCAS1 | 6.00 | 7.00 | 9.00 | 7.33 | 7.00 |
| BCAS2 | 656.00 | 635.00 | 653.00 | 648.00 | 857.00 |
| BCAS3 | 427.00 | 364.00 | 461.00 | 417.33 | 393.00 |
| BCAS4 | 280.00 | 352.00 | 294.00 | 308.67 | 326.00 |
| BCAT1 | 6674.00 | 5879.00 | 7008.00 | 6520.33 | 6781.00 |
| BCAT2 | 681.00 | 711.00 | 737.00 | 709.67 | 850.00 |
| BCCIP | 2180.00 | 2201.00 | 2295.00 | 2225.33 | 2599.00 |
| BCDIN3D | 174.00 | 208.00 | 167.00 | 183.00 | 206.00 |
| BCDIN3D-AS1 | 31.00 | 22.00 | 23.00 | 25.33 | 19.00 |
| BCHE | 2000.00 | 1795.00 | 1956.00 | 1917.00 | 2251.00 |
| BCKDHA | 291.00 | 304.00 | 346.00 | 313.67 | 345.00 |
| BCKDHB | 358.00 | 380.00 | 376.00 | 371.33 | 395.00 |
| BCKDK | 766.00 | 773.00 | 781.00 | 773.33 | 803.00 |
| BCL10 | 658.00 | 594.00 | 694.00 | 648.67 | 763.00 |
| BCL11A | 6.00 | 11.00 | 5.00 | 7.33 | 8.00 |
| BCL2 | 278.00 | 281.00 | 368.00 | 309.00 | 453.00 |
| BCL2A1 | 959.00 | 860.00 | 860.00 | 893.00 | 1863.00 |
| BCL2L1 | 1452.00 | 1365.00 | 1472.00 | 1429.67 | 1430.00 |
| BCL2L11 | 121.00 | 132.00 | 143.00 | 132.00 | 175.00 |
| BCL2L12 | 502.00 | 475.00 | 513.00 | 496.67 | 731.00 |
| BCL2L13 | 1376.00 | 1288.00 | 1317.00 | 1327.00 | 1488.00 |
| BCL2L15 | 11.00 | 11.00 | 9.00 | 10.33 | 11.00 |
| BCL2L2 | 1422.00 | 1417.00 | 1416.00 | 1418.33 | 1515.00 |
| BCL2L2-PABPN1 | 7.00 | 1.00 | 1.00 | 3.00 | 7.00 |
| BCL3 | 141.00 | 119.00 | 163.00 | 141.00 | 152.00 |
| BCL6 | 1084.00 | 1037.00 | 1250.00 | 1123.67 | 1165.00 |
| BCL7A | 842.00 | 794.00 | 871.00 | 835.67 | 1369.00 |

| | | | | | |
|---|---|---|---|---|---|
| BCL7B | 801.00 | 713.00 | 765.00 | 759.67 | 858.00 |
| BCL7C | 539.00 | 524.00 | 528.00 | 530.33 | 734.00 |
| BCL9 | 536.00 | 472.00 | 575.00 | 527.67 | 520.00 |
| BCL9L | 2539.00 | 2507.00 | 2549.00 | 2531.67 | 1765.00 |
| BCLAF1 | 3759.00 | 3422.00 | 3917.00 | 3699.33 | 4078.00 |
| BCO2 | 28.00 | 36.00 | 30.00 | 31.33 | 14.00 |
| BCOR | 519.00 | 543.00 | 611.00 | 557.67 | 446.00 |
| BCORL1 | 316.00 | 239.00 | 317.00 | 290.67 | 224.00 |
| BCR | 775.00 | 760.00 | 751.00 | 762.00 | 772.00 |
| BCRP2 | 8.00 | 5.00 | 4.00 | 5.67 | 4.00 |
| BCS1L | 1306.00 | 1414.00 | 1464.00 | 1394.67 | 1611.00 |
| BCYRN1 | 12.00 | 28.00 | 15.00 | 18.33 | 72.00 |
| BDH1 | 29.00 | 16.00 | 22.00 | 22.33 | 14.00 |
| BDH2 | 229.00 | 224.00 | 239.00 | 230.67 | 260.00 |
| BDKRB1 | 12.00 | 13.00 | 18.00 | 14.33 | 12.00 |
| BDKRB2 | 4.00 | 12.00 | 7.00 | 7.67 | 3.00 |
| BDNF | 1180.00 | 1159.00 | 1280.00 | 1206.33 | 1874.00 |
| BDNF-AS | 40.00 | 33.00 | 42.00 | 38.33 | 25.00 |
| BEAN1 | 24.00 | 20.00 | 18.00 | 20.67 | 22.00 |
| BECN1 | 1136.00 | 1242.00 | 1329.00 | 1235.67 | 1485.00 |
| BEND3 | 308.00 | 306.00 | 324.00 | 312.67 | 375.00 |
| BEND4 | 786.00 | 683.00 | 813.00 | 760.67 | 679.00 |
| BEND6 | 75.00 | 74.00 | 111.00 | 86.67 | 95.00 |
| BEND7 | 304.00 | 303.00 | 279.00 | 295.33 | 223.00 |
| BEST1 | 215.00 | 170.00 | 182.00 | 189.00 | 154.00 |
| BEST3 | 71.00 | 57.00 | 97.00 | 75.00 | 119.00 |
| BEST4 | 10.00 | 7.00 | 9.00 | 8.67 | 8.00 |
| BET1 | 668.00 | 623.00 | 705.00 | 665.33 | 790.00 |
| BET1L | 1442.00 | 1366.00 | 1461.00 | 1423.00 | 1388.00 |
| BEX1 | 43.00 | 59.00 | 47.00 | 49.67 | 44.00 |
| BEX2 | 80.00 | 69.00 | 86.00 | 78.33 | 116.00 |
| BEX4 | 619.00 | 655.00 | 678.00 | 650.67 | 707.00 |
| BEX5 | 7.00 | 5.00 | 10.00 | 7.33 | 3.00 |
| BFSP1 | 30.00 | 45.00 | 38.00 | 37.67 | 35.00 |
| BGLAP | 10.00 | 9.00 | 11.00 | 10.00 | 3.00 |
| BGN | 2429.00 | 2460.00 | 2409.00 | 2432.67 | 1703.00 |
| BHLHA15 | 8.00 | 7.00 | 7.00 | 7.33 | 7.00 |
| BHLHB9 | 76.00 | 73.00 | 92.00 | 80.33 | 66.00 |
| BHLHE40 | 384.00 | 397.00 | 401.00 | 394.00 | 438.00 |
| BHLHE40-AS1 | 9.00 | 7.00 | 4.00 | 6.67 | 8.00 |
| BHLHE41 | 595.00 | 573.00 | 651.00 | 606.33 | 543.00 |
| BICC1 | 177.00 | 150.00 | 195.00 | 174.00 | 274.00 |
| BICD1 | 368.00 | 388.00 | 417.00 | 391.00 | 375.00 |
| BICD2 | 1127.00 | 989.00 | 1149.00 | 1088.33 | 1034.00 |
| BID | 403.00 | 408.00 | 465.00 | 425.33 | 578.00 |
| BIN1 | 240.00 | 240.00 | 276.00 | 252.00 | 244.00 |
| BIN3 | 365.00 | 342.00 | 367.00 | 358.00 | 382.00 |
| BIN3-IT1 | 4.00 | 3.00 | 3.00 | 3.33 | 1.00 |

| | | | | | |
|---|---|---|---|---|---|
| BIRC2 | 8106.00 | 7843.00 | 8300.00 | 8083.00 | 10662.00 |
| BIRC3 | 639.00 | 568.00 | 686.00 | 631.00 | 2234.00 |
| BIRC5 | 770.00 | 775.00 | 752.00 | 765.67 | 1153.00 |
| BIRC6 | 2368.00 | 2130.00 | 2464.00 | 2320.67 | 2056.00 |
| BISPR | 13.00 | 5.00 | 18.00 | 12.00 | 17.00 |
| BIVM | 888.00 | 889.00 | 916.00 | 897.67 | 1175.00 |
| BLCAP | 1307.00 | 1270.00 | 1206.00 | 1261.00 | 1453.00 |
| BLID | 4.00 | 10.00 | 10.00 | 8.00 | 7.00 |
| BLK | 1752.00 | 1869.00 | 2002.00 | 1874.33 | 2129.00 |
| BLM | 373.00 | 372.00 | 356.00 | 367.00 | 465.00 |
| BLMH | 656.00 | 611.00 | 653.00 | 640.00 | 668.00 |
| BLOC1S1 | 218.00 | 210.00 | 223.00 | 217.00 | 231.00 |
| BLOC1S2 | 554.00 | 560.00 | 569.00 | 561.00 | 730.00 |
| BLOC1S3 | 295.00 | 240.00 | 299.00 | 278.00 | 355.00 |
| BLOC1S4 | 196.00 | 201.00 | 252.00 | 216.33 | 216.00 |
| BLOC1S6 | 1197.00 | 1100.00 | 1250.00 | 1182.33 | 1443.00 |
| BLVRA | 355.00 | 367.00 | 403.00 | 375.00 | 532.00 |
| BLVRB | 322.00 | 368.00 | 380.00 | 356.67 | 388.00 |
| BLZF1 | 539.00 | 495.00 | 559.00 | 531.00 | 582.00 |
| BMF | 46.00 | 60.00 | 64.00 | 56.67 | 31.00 |
| BMI1 | 77.00 | 59.00 | 88.00 | 74.67 | 80.00 |
| BMP1 | 675.00 | 724.00 | 712.00 | 703.67 | 517.00 |
| BMP2 | 485.00 | 475.00 | 527.00 | 495.67 | 534.00 |
| BMP2K | 238.00 | 224.00 | 252.00 | 238.00 | 200.00 |
| BMP3 | 9.00 | 8.00 | 14.00 | 10.33 | 12.00 |
| BMP4 | 84.00 | 90.00 | 73.00 | 82.33 | 78.00 |
| BMP6 | 264.00 | 218.00 | 255.00 | 245.67 | 631.00 |
| BMPER | 356.00 | 376.00 | 420.00 | 384.00 | 304.00 |
| BMPR1A | 564.00 | 471.00 | 587.00 | 540.67 | 558.00 |
| BMPR1B | 104.00 | 115.00 | 116.00 | 111.67 | 75.00 |
| BMPR2 | 1417.00 | 1346.00 | 1609.00 | 1457.33 | 1285.00 |
| BMS1 | 2836.00 | 2795.00 | 2933.00 | 2854.67 | 2817.00 |
| BMS1P20 | 457.00 | 437.00 | 426.00 | 440.00 | 432.00 |
| BMS1P21 | 19.00 | 18.00 | 20.00 | 19.00 | 9.00 |
| BMS1P4 | 304.00 | 284.00 | 281.00 | 289.67 | 191.00 |
| BMS1P5 | 212.00 | 235.00 | 216.00 | 221.00 | 162.00 |
| BMS1P6 | 99.00 | 82.00 | 112.00 | 97.67 | 68.00 |
| BNC1 | 3219.00 | 2969.00 | 3425.00 | 3204.33 | 3505.00 |
| BNC2 | 537.00 | 526.00 | 532.00 | 531.67 | 308.00 |
| BNIP1 | 344.00 | 323.00 | 326.00 | 331.00 | 444.00 |
| BNIP2 | 1305.00 | 1109.00 | 1294.00 | 1236.00 | 1538.00 |
| BNIP3 | 1710.00 | 1631.00 | 1901.00 | 1747.33 | 2398.00 |
| BNIP3L | 950.00 | 905.00 | 970.00 | 941.67 | 863.00 |
| BNIPL | 40.00 | 39.00 | 44.00 | 41.00 | 34.00 |
| BOC | 369.00 | 399.00 | 398.00 | 388.67 | 295.00 |
| BOD1 | 1497.00 | 1468.00 | 1602.00 | 1522.33 | 1932.00 |
| BOD1L1 | 1394.00 | 1316.00 | 1495.00 | 1401.67 | 1205.00 |
| BOK | 114.00 | 98.00 | 118.00 | 110.00 | 96.00 |

| | | | | | |
|---|---|---|---|---|---|
| BOLA1 | 234.00 | 250.00 | 250.00 | 244.67 | 291.00 |
| BOLA3 | 348.00 | 311.00 | 377.00 | 345.33 | 468.00 |
| BOLA3-AS1 | 77.00 | 94.00 | 68.00 | 79.67 | 76.00 |
| BOP1 | 3303.00 | 3382.00 | 3383.00 | 3356.00 | 4125.00 |
| BORA | 282.00 | 289.00 | 325.00 | 298.67 | 376.00 |
| BPGM | 186.00 | 161.00 | 205.00 | 184.00 | 246.00 |
| BPHL | 317.00 | 320.00 | 343.00 | 326.67 | 368.00 |
| BPNT1 | 472.00 | 467.00 | 488.00 | 475.67 | 595.00 |
| BPTF | 2444.00 | 2091.00 | 2501.00 | 2345.33 | 1944.00 |
| BRAF | 239.00 | 239.00 | 257.00 | 245.00 | 238.00 |
| BRAP | 541.00 | 496.00 | 530.00 | 522.33 | 583.00 |
| BRAT1 | 1871.00 | 1891.00 | 1941.00 | 1901.00 | 2036.00 |
| BRCA1 | 965.00 | 909.00 | 925.00 | 933.00 | 898.00 |
| BRCA2 | 381.00 | 364.00 | 423.00 | 389.33 | 421.00 |
| BRCAT54 | 9.00 | 14.00 | 21.00 | 14.67 | 14.00 |
| BRCC3 | 442.00 | 484.00 | 459.00 | 461.67 | 576.00 |
| BRD1 | 966.00 | 943.00 | 1056.00 | 988.33 | 961.00 |
| BRD2 | 20.00 | 13.00 | 15.00 | 16.00 | 12.00 |
| BRD3 | 969.00 | 938.00 | 995.00 | 967.33 | 832.00 |
| BRD4 | 2057.00 | 1948.00 | 2066.00 | 2023.67 | 1994.00 |
| BRD7 | 1219.00 | 1249.00 | 1296.00 | 1254.67 | 1559.00 |
| BRD8 | 2103.00 | 2200.00 | 2307.00 | 2203.33 | 2319.00 |
| BRD9 | 1166.00 | 1326.00 | 1440.00 | 1310.67 | 1354.00 |
| BRDTP1 | 13.00 | 19.00 | 23.00 | 18.33 | 13.00 |
| BRE | 1967.00 | 1796.00 | 2069.00 | 1944.00 | 1851.00 |
| BRE-AS1 | 24.00 | 33.00 | 29.00 | 28.67 | 18.00 |
| BRF1 | 719.00 | 775.00 | 751.00 | 748.33 | 837.00 |
| BRF2 | 310.00 | 345.00 | 343.00 | 332.67 | 411.00 |
| BRI3 | 1273.00 | 1166.00 | 1207.00 | 1215.33 | 1212.00 |
| BRI3BP | 58.00 | 52.00 | 57.00 | 55.67 | 85.00 |
| BRICD5 | 180.00 | 214.00 | 205.00 | 199.67 | 211.00 |
| BRIP1 | 248.00 | 247.00 | 275.00 | 256.67 | 344.00 |
| BRIX1 | 1162.00 | 1132.00 | 1250.00 | 1181.33 | 1722.00 |
| BRK1 | 1480.00 | 1439.00 | 1593.00 | 1504.00 | 1674.00 |
| BRMS1 | 1169.00 | 1221.00 | 1296.00 | 1228.67 | 1472.00 |
| BRMS1L | 226.00 | 250.00 | 268.00 | 248.00 | 288.00 |
| BROX | 1382.00 | 1237.00 | 1472.00 | 1363.67 | 1365.00 |
| BRPF1 | 845.00 | 873.00 | 811.00 | 843.00 | 835.00 |
| BRPF3 | 1919.00 | 1783.00 | 1971.00 | 1891.00 | 1634.00 |
| BRSK1 | 193.00 | 184.00 | 193.00 | 190.00 | 160.00 |
| BRSK2 | 21.00 | 39.00 | 34.00 | 31.33 | 32.00 |
| BRWD1 | 1350.00 | 1291.00 | 1542.00 | 1394.33 | 1224.00 |
| BRWD1-IT2 | 5.00 | 5.00 | 8.00 | 6.00 | 6.00 |
| BRWD3 | 832.00 | 824.00 | 1009.00 | 888.33 | 815.00 |
| BSCL2 | 163.00 | 192.00 | 167.00 | 174.00 | 147.00 |
| BSDC1 | 1756.00 | 1775.00 | 1897.00 | 1809.33 | 1440.00 |
| BSG | 24092.00 | 25714.00 | 23756.00 | 24520.67 | 24681.00 |
| BSN | 27.00 | 27.00 | 38.00 | 30.67 | 18.00 |

| | | | | | |
|---|---|---|---|---|---|
| BSN-AS2 | 5.00 | 3.00 | 5.00 | 4.33 | 5.00 |
| BST1 | 372.00 | 359.00 | 388.00 | 373.00 | 332.00 |
| BTAF1 | 3487.00 | 3258.00 | 3608.00 | 3451.00 | 2659.00 |
| BTBD1 | 1431.00 | 1300.00 | 1636.00 | 1455.67 | 1689.00 |
| BTBD10 | 828.00 | 677.00 | 828.00 | 777.67 | 901.00 |
| BTBD11 | 351.00 | 357.00 | 416.00 | 374.67 | 255.00 |
| BTBD19 | 509.00 | 566.00 | 563.00 | 546.00 | 387.00 |
| BTBD2 | 1972.00 | 2000.00 | 2122.00 | 2031.33 | 2309.00 |
| BTBD3 | 955.00 | 840.00 | 1114.00 | 969.67 | 1167.00 |
| BTBD6 | 554.00 | 511.00 | 545.00 | 536.67 | 721.00 |
| BTBD8 | 40.00 | 35.00 | 28.00 | 34.33 | 26.00 |
| BTBD9 | 362.00 | 355.00 | 374.00 | 363.67 | 322.00 |
| BTC | 10.00 | 12.00 | 12.00 | 11.33 | 11.00 |
| BTD | 627.00 | 577.00 | 598.00 | 600.67 | 492.00 |
| BTF3 | 8785.00 | 8843.00 | 9573.00 | 9067.00 | 12142.00 |
| BTF3L4 | 1224.00 | 1105.00 | 1325.00 | 1218.00 | 1653.00 |
| BTG1 | 1681.00 | 1504.00 | 1775.00 | 1653.33 | 2171.00 |
| BTG3 | 428.00 | 426.00 | 451.00 | 435.00 | 638.00 |
| BTN2A1 | 1302.00 | 1216.00 | 1288.00 | 1268.67 | 1260.00 |
| BTN2A2 | 462.00 | 436.00 | 457.00 | 451.67 | 452.00 |
| BTN2A3P | 245.00 | 234.00 | 255.00 | 244.67 | 214.00 |
| BTN3A1 | 637.00 | 612.00 | 649.00 | 632.67 | 690.00 |
| BTN3A2 | 564.00 | 606.00 | 604.00 | 591.33 | 581.00 |
| BTN3A3 | 308.00 | 289.00 | 309.00 | 302.00 | 255.00 |
| BTNL8 | 11.00 | 2.00 | 0.00 | 4.33 | 27.00 |
| BTRC | 921.00 | 844.00 | 915.00 | 893.33 | 973.00 |
| BUB1 | 1304.00 | 1210.00 | 1322.00 | 1278.67 | 1627.00 |
| BUB1B | 1129.00 | 1136.00 | 1144.00 | 1136.33 | 1360.00 |
| BUB3 | 5649.00 | 5391.00 | 5958.00 | 5666.00 | 6069.00 |
| BUD13 | 360.00 | 340.00 | 374.00 | 358.00 | 397.00 |
| BUD31 | 1268.00 | 1277.00 | 1341.00 | 1295.33 | 1693.00 |
| BVES | 220.00 | 198.00 | 193.00 | 203.67 | 240.00 |
| BVES-AS1 | 4.00 | 4.00 | 2.00 | 3.33 | 2.00 |
| BYSL | 961.00 | 1003.00 | 1052.00 | 1005.33 | 1159.00 |
| BZRAP1 | 142.00 | 168.00 | 209.00 | 173.00 | 170.00 |
| BZRAP1-AS1 | 69.00 | 56.00 | 48.00 | 57.67 | 79.00 |
| BZW1 | 3669.00 | 3382.00 | 3806.00 | 3619.00 | 4880.00 |
| BZW2 | 2162.00 | 2129.00 | 2401.00 | 2230.67 | 3130.00 |
| C10orf10 | 11.00 | 8.00 | 6.00 | 8.33 | 18.00 |
| C10orf11 | 73.00 | 75.00 | 72.00 | 73.33 | 92.00 |
| C10orf111 | 42.00 | 49.00 | 46.00 | 45.67 | 32.00 |
| C10orf12 | 112.00 | 88.00 | 112.00 | 104.00 | 67.00 |
| C10orf2 | 1173.00 | 1278.00 | 1477.00 | 1309.33 | 1446.00 |
| C10orf25 | 70.00 | 69.00 | 87.00 | 75.33 | 93.00 |
| C10orf32 | 247.00 | 242.00 | 262.00 | 250.33 | 284.00 |
| C10orf54 | 4.00 | 3.00 | 1.00 | 2.67 | 2.00 |
| C10orf55 | 31.00 | 21.00 | 26.00 | 26.00 | 33.00 |
| C10orf67 | 8.00 | 5.00 | 5.00 | 6.00 | 9.00 |

| | | | | | |
|---|---|---|---|---|---|
| C10orf76 | 732.00 | 723.00 | 754.00 | 736.33 | 685.00 |
| C10orf88 | 560.00 | 590.00 | 590.00 | 580.00 | 655.00 |
| C10orf90 | 45.00 | 44.00 | 46.00 | 45.00 | 62.00 |
| C11orf1 | 200.00 | 201.00 | 245.00 | 215.33 | 261.00 |
| C11orf24 | 788.00 | 822.00 | 776.00 | 795.33 | 888.00 |
| C11orf30 | 760.00 | 698.00 | 747.00 | 735.00 | 681.00 |
| C11orf31 | 1062.00 | 981.00 | 1133.00 | 1058.67 | 1281.00 |
| C11orf45 | 74.00 | 66.00 | 70.00 | 70.00 | 77.00 |
| C11orf49 | 396.00 | 421.00 | 456.00 | 424.33 | 494.00 |
| C11orf54 | 514.00 | 398.00 | 573.00 | 495.00 | 490.00 |
| C11orf57 | 727.00 | 636.00 | 749.00 | 704.00 | 789.00 |
| C11orf58 | 2380.00 | 2232.00 | 2578.00 | 2396.67 | 3015.00 |
| C11orf63 | 4.00 | 1.00 | 3.00 | 2.67 | 1.00 |
| C11orf65 | 6.00 | 7.00 | 5.00 | 6.00 | 6.00 |
| C11orf68 | 1809.00 | 1754.00 | 1922.00 | 1828.33 | 2217.00 |
| C11orf70 | 362.00 | 329.00 | 375.00 | 355.33 | 514.00 |
| C11orf71 | 83.00 | 90.00 | 96.00 | 89.67 | 105.00 |
| C11orf73 | 477.00 | 479.00 | 501.00 | 485.67 | 570.00 |
| C11orf74 | 251.00 | 234.00 | 253.00 | 246.00 | 327.00 |
| C11orf80 | 492.00 | 511.00 | 531.00 | 511.33 | 637.00 |
| C11orf84 | 1275.00 | 1340.00 | 1368.00 | 1327.67 | 1442.00 |
| C11orf91 | 118.00 | 104.00 | 149.00 | 123.67 | 149.00 |
| C11orf94 | 6.00 | 8.00 | 9.00 | 7.67 | 5.00 |
| C11orf95 | 1490.00 | 1465.00 | 1820.00 | 1591.67 | 1433.00 |
| C11orf98 | 50.00 | 67.00 | 49.00 | 55.33 | 62.00 |
| C12orf10 | 1183.00 | 1172.00 | 1186.00 | 1180.33 | 1612.00 |
| C12orf29 | 581.00 | 543.00 | 652.00 | 592.00 | 619.00 |
| C12orf4 | 574.00 | 537.00 | 592.00 | 567.67 | 699.00 |
| C12orf43 | 273.00 | 284.00 | 236.00 | 264.33 | 306.00 |
| C12orf45 | 151.00 | 171.00 | 177.00 | 166.33 | 213.00 |
| C12orf49 | 852.00 | 863.00 | 879.00 | 864.67 | 916.00 |
| C12orf56 | 72.00 | 58.00 | 46.00 | 58.67 | 74.00 |
| C12orf57 | 1083.00 | 1054.00 | 1121.00 | 1086.00 | 1161.00 |
| C12orf60 | 62.00 | 46.00 | 51.00 | 53.00 | 59.00 |
| C12orf65 | 305.00 | 318.00 | 343.00 | 322.00 | 366.00 |
| C12orf66 | 275.00 | 216.00 | 278.00 | 256.33 | 236.00 |
| C12orf73 | 243.00 | 227.00 | 245.00 | 238.33 | 252.00 |
| C12orf75 | 710.00 | 584.00 | 729.00 | 674.33 | 896.00 |
| C12orf76 | 185.00 | 209.00 | 227.00 | 207.00 | 272.00 |
| C14orf1 | 440.00 | 400.00 | 427.00 | 422.33 | 445.00 |
| C14orf119 | 918.00 | 987.00 | 1064.00 | 989.67 | 1168.00 |
| C14orf132 | 86.00 | 61.00 | 74.00 | 73.67 | 66.00 |
| C14orf159 | 465.00 | 492.00 | 546.00 | 501.00 | 542.00 |
| C14orf166 | 1954.00 | 1888.00 | 2218.00 | 2020.00 | 2640.00 |
| C14orf169 | 418.00 | 398.00 | 433.00 | 416.33 | 480.00 |
| C14orf2 | 822.00 | 843.00 | 882.00 | 849.00 | 1200.00 |
| C14orf28 | 38.00 | 49.00 | 52.00 | 46.33 | 51.00 |
| C14orf37 | 15.00 | 15.00 | 15.00 | 15.00 | 14.00 |

| | | | | | |
|---|---|---|---|---|---|
| C14orf39 | 4.00 | 3.00 | 3.00 | 3.33 | 1.00 |
| C14orf79 | 426.00 | 407.00 | 421.00 | 418.00 | 423.00 |
| C14orf80 | 303.00 | 270.00 | 234.00 | 269.00 | 354.00 |
| C14orf93 | 292.00 | 311.00 | 357.00 | 320.00 | 352.00 |
| C15orf26 | 24.00 | 12.00 | 14.00 | 16.67 | 14.00 |
| C15orf39 | 367.00 | 291.00 | 301.00 | 319.67 | 320.00 |
| C15orf40 | 316.00 | 296.00 | 280.00 | 297.33 | 359.00 |
| C15orf41 | 432.00 | 416.00 | 448.00 | 432.00 | 510.00 |
| C15orf48 | 6.00 | 9.00 | 9.00 | 8.00 | 6.00 |
| C15orf52 | 5598.00 | 5859.00 | 5863.00 | 5773.33 | 5241.00 |
| C15orf54 | 4.00 | 1.00 | 2.00 | 2.33 | 8.00 |
| C15orf56 | 5.00 | 4.00 | 9.00 | 6.00 | 3.00 |
| C15orf57 | 206.00 | 195.00 | 231.00 | 210.67 | 194.00 |
| C15orf61 | 128.00 | 108.00 | 128.00 | 121.33 | 201.00 |
| C15orf65 | 8.00 | 13.00 | 7.00 | 9.33 | 19.00 |
| C16orf13 | 772.00 | 783.00 | 912.00 | 822.33 | 1090.00 |
| C16orf46 | 91.00 | 82.00 | 68.00 | 80.33 | 88.00 |
| C16orf52 | 1782.00 | 1576.00 | 1924.00 | 1760.67 | 1960.00 |
| C16orf58 | 3383.00 | 3318.00 | 3428.00 | 3376.33 | 3183.00 |
| C16orf59 | 342.00 | 357.00 | 307.00 | 335.33 | 438.00 |
| C16orf62 | 1151.00 | 1041.00 | 1257.00 | 1149.67 | 970.00 |
| C16orf70 | 461.00 | 495.00 | 499.00 | 485.00 | 510.00 |
| C16orf72 | 1020.00 | 814.00 | 1035.00 | 956.33 | 1097.00 |
| C16orf74 | 409.00 | 420.00 | 418.00 | 415.67 | 295.00 |
| C16orf86 | 20.00 | 41.00 | 21.00 | 27.33 | 27.00 |
| C16orf87 | 71.00 | 71.00 | 63.00 | 68.33 | 117.00 |
| C16orf91 | 283.00 | 262.00 | 275.00 | 273.33 | 329.00 |
| C16orf93 | 86.00 | 74.00 | 62.00 | 74.00 | 83.00 |
| C16orf95 | 95.00 | 103.00 | 98.00 | 98.67 | 114.00 |
| C17orf100 | 101.00 | 83.00 | 94.00 | 92.67 | 95.00 |
| C17orf104 | 5.00 | 11.00 | 7.00 | 7.67 | 14.00 |
| C17orf107 | 18.00 | 25.00 | 36.00 | 26.33 | 23.00 |
| C17orf51 | 2929.00 | 2672.00 | 3169.00 | 2923.33 | 2190.00 |
| C17orf53 | 335.00 | 303.00 | 324.00 | 320.67 | 351.00 |
| C17orf58 | 295.00 | 300.00 | 302.00 | 299.00 | 455.00 |
| C17orf59 | 191.00 | 188.00 | 210.00 | 196.33 | 183.00 |
| C17orf62 | 1510.00 | 1432.00 | 1642.00 | 1528.00 | 1736.00 |
| C17orf67 | 51.00 | 56.00 | 67.00 | 58.00 | 59.00 |
| C17orf75 | 518.00 | 505.00 | 518.00 | 513.67 | 676.00 |
| C17orf80 | 839.00 | 795.00 | 900.00 | 844.67 | 892.00 |
| C17orf85 | 2152.00 | 2076.00 | 2328.00 | 2185.33 | 2147.00 |
| C17orf89 | 432.00 | 399.00 | 418.00 | 416.33 | 570.00 |
| C17orf97 | 23.00 | 28.00 | 22.00 | 24.33 | 27.00 |
| C18orf21 | 221.00 | 241.00 | 237.00 | 233.00 | 313.00 |
| C18orf25 | 520.00 | 501.00 | 530.00 | 517.00 | 672.00 |
| C18orf32 | 22.00 | 23.00 | 24.00 | 23.00 | 10.00 |
| C18orf54 | 421.00 | 390.00 | 463.00 | 424.67 | 510.00 |
| C18orf65 | 38.00 | 35.00 | 30.00 | 34.33 | 43.00 |

| | | | | | |
|---|---|---|---|---|---|
| C18orf8 | 612.00 | 524.00 | 601.00 | 579.00 | 599.00 |
| C19orf12 | 455.00 | 444.00 | 480.00 | 459.67 | 599.00 |
| C19orf24 | 469.00 | 459.00 | 483.00 | 470.33 | 637.00 |
| C19orf25 | 382.00 | 332.00 | 426.00 | 380.00 | 446.00 |
| C19orf38 | 7.00 | 6.00 | 6.00 | 6.33 | 10.00 |
| C19orf43 | 1665.00 | 1583.00 | 1711.00 | 1653.00 | 1993.00 |
| C19orf44 | 388.00 | 359.00 | 367.00 | 371.33 | 376.00 |
| C19orf47 | 146.00 | 144.00 | 156.00 | 148.67 | 182.00 |
| C19orf48 | 2048.00 | 2050.00 | 2234.00 | 2110.67 | 2682.00 |
| C19orf52 | 221.00 | 251.00 | 243.00 | 238.33 | 279.00 |
| C19orf53 | 871.00 | 851.00 | 912.00 | 878.00 | 1188.00 |
| C19orf54 | 680.00 | 687.00 | 723.00 | 696.67 | 656.00 |
| C19orf57 | 40.00 | 42.00 | 46.00 | 42.67 | 34.00 |
| C19orf60 | 410.00 | 473.00 | 466.00 | 449.67 | 585.00 |
| C19orf66 | 517.00 | 535.00 | 541.00 | 531.00 | 506.00 |
| C19orf68 | 324.00 | 421.00 | 363.00 | 369.33 | 386.00 |
| C19orf70 | 463.00 | 430.00 | 537.00 | 476.67 | 566.00 |
| C19orf71 | 26.00 | 26.00 | 18.00 | 23.33 | 18.00 |
| C19orf73 | 19.00 | 21.00 | 10.00 | 16.67 | 16.00 |
| C1D | 225.00 | 236.00 | 241.00 | 234.00 | 322.00 |
| C1GALT1 | 1692.00 | 1673.00 | 1921.00 | 1762.00 | 1913.00 |
| C1GALT1C1 | 1089.00 | 1095.00 | 1009.00 | 1064.33 | 1199.00 |
| C1GALT1C1L | 26.00 | 25.00 | 25.00 | 25.33 | 30.00 |
| C1orf101 | 27.00 | 26.00 | 22.00 | 25.00 | 21.00 |
| C1orf109 | 714.00 | 671.00 | 754.00 | 713.00 | 821.00 |
| C1orf110 | 29.00 | 27.00 | 23.00 | 26.33 | 72.00 |
| C1orf112 | 426.00 | 425.00 | 391.00 | 414.00 | 581.00 |
| C1orf122 | 390.00 | 400.00 | 463.00 | 417.67 | 472.00 |
| C1orf123 | 792.00 | 805.00 | 931.00 | 842.67 | 921.00 |
| C1orf131 | 169.00 | 152.00 | 200.00 | 173.67 | 228.00 |
| C1orf159 | 442.00 | 426.00 | 445.00 | 437.67 | 523.00 |
| C1orf162 | 42.00 | 39.00 | 51.00 | 44.00 | 44.00 |
| C1orf174 | 599.00 | 525.00 | 623.00 | 582.33 | 742.00 |
| C1orf189 | 4.00 | 4.00 | 0.00 | 2.67 | 0.00 |
| C1orf198 | 365.00 | 326.00 | 372.00 | 354.33 | 378.00 |
| C1orf204 | 169.00 | 174.00 | 183.00 | 175.33 | 167.00 |
| C1orf21 | 473.00 | 433.00 | 454.00 | 453.33 | 544.00 |
| C1orf216 | 792.00 | 798.00 | 916.00 | 835.33 | 992.00 |
| C1orf220 | 54.00 | 62.00 | 65.00 | 60.33 | 56.00 |
| C1orf226 | 70.00 | 78.00 | 74.00 | 74.00 | 70.00 |
| C1orf228 | 11.00 | 11.00 | 16.00 | 12.67 | 21.00 |
| C1orf27 | 1104.00 | 1075.00 | 1173.00 | 1117.33 | 1200.00 |
| C1orf35 | 392.00 | 422.00 | 428.00 | 414.00 | 455.00 |
| C1orf43 | 6231.00 | 6033.00 | 6295.00 | 6186.33 | 8006.00 |
| C1orf50 | 119.00 | 164.00 | 160.00 | 147.67 | 190.00 |
| C1orf52 | 516.00 | 507.00 | 555.00 | 526.00 | 631.00 |
| C1orf53 | 38.00 | 53.00 | 37.00 | 42.67 | 66.00 |
| C1orf54 | 11.00 | 16.00 | 11.00 | 12.67 | 19.00 |

| | | | | | |
|---|---|---|---|---|---|
| C1orf56 | 158.00 | 174.00 | 176.00 | 169.33 | 202.00 |
| C1orf74 | 205.00 | 198.00 | 209.00 | 204.00 | 247.00 |
| C1QBP | 4781.00 | 4774.00 | 4987.00 | 4847.33 | 6950.00 |
| C1QL4 | 59.00 | 54.00 | 37.00 | 50.00 | 66.00 |
| C1QTNF1 | 75.00 | 70.00 | 74.00 | 73.00 | 29.00 |
| C1QTNF3 | 32.00 | 34.00 | 30.00 | 32.00 | 38.00 |
| C1QTNF6 | 373.00 | 374.00 | 404.00 | 383.67 | 384.00 |
| C1QTNF9B-AS1 | 21.00 | 19.00 | 28.00 | 22.67 | 16.00 |
| C1R | 46.00 | 32.00 | 42.00 | 40.00 | 42.00 |
| C1RL | 274.00 | 288.00 | 277.00 | 279.67 | 244.00 |
| C1RL-AS1 | 640.00 | 621.00 | 677.00 | 646.00 | 377.00 |
| C1S | 49.00 | 32.00 | 32.00 | 37.67 | 29.00 |
| C20orf194 | 1077.00 | 953.00 | 1018.00 | 1016.00 | 920.00 |
| C20orf196 | 36.00 | 42.00 | 37.00 | 38.33 | 46.00 |
| C20orf197 | 4.00 | 5.00 | 5.00 | 4.67 | 14.00 |
| C20orf24 | 238.00 | 225.00 | 206.00 | 223.00 | 318.00 |
| C20orf27 | 1214.00 | 1288.00 | 1252.00 | 1251.33 | 1606.00 |
| C20orf96 | 438.00 | 446.00 | 468.00 | 450.67 | 460.00 |
| C21orf2 | 229.00 | 211.00 | 252.00 | 230.67 | 231.00 |
| C21orf33 | 70.00 | 64.00 | 95.00 | 76.33 | 110.00 |
| C21orf58 | 631.00 | 509.00 | 488.00 | 542.67 | 566.00 |
| C21orf59 | 1068.00 | 1050.00 | 1026.00 | 1048.00 | 1277.00 |
| C21orf91 | 189.00 | 161.00 | 230.00 | 193.33 | 200.00 |
| C22orf23 | 6.00 | 7.00 | 7.00 | 6.67 | 7.00 |
| C22orf29 | 1077.00 | 1024.00 | 1157.00 | 1086.00 | 988.00 |
| C22orf34 | 8.00 | 12.00 | 11.00 | 10.33 | 15.00 |
| C22orf39 | 275.00 | 292.00 | 283.00 | 283.33 | 394.00 |
| C22orf46 | 917.00 | 884.00 | 945.00 | 915.33 | 705.00 |
| C2CD2 | 1609.00 | 1562.00 | 1708.00 | 1626.33 | 1286.00 |
| C2CD2L | 1018.00 | 1053.00 | 1042.00 | 1037.67 | 985.00 |
| C2CD3 | 564.00 | 534.00 | 603.00 | 567.00 | 559.00 |
| C2CD5 | 1402.00 | 1340.00 | 1492.00 | 1411.33 | 1573.00 |
| C2orf16 | 11.00 | 8.00 | 9.00 | 9.33 | 12.00 |
| C2orf27A | 7.00 | 4.00 | 7.00 | 6.00 | 7.00 |
| C2orf42 | 271.00 | 250.00 | 342.00 | 287.67 | 343.00 |
| C2orf44 | 469.00 | 462.00 | 479.00 | 470.00 | 484.00 |
| C2orf47 | 335.00 | 350.00 | 306.00 | 330.33 | 421.00 |
| C2orf48 | 18.00 | 24.00 | 20.00 | 20.67 | 40.00 |
| C2orf49 | 628.00 | 594.00 | 625.00 | 615.67 | 562.00 |
| C2orf61 | 4.00 | 11.00 | 7.00 | 7.33 | 5.00 |
| C2orf66 | 18.00 | 15.00 | 25.00 | 19.33 | 19.00 |
| C2orf68 | 846.00 | 807.00 | 890.00 | 847.67 | 904.00 |
| C2orf69 | 433.00 | 384.00 | 458.00 | 425.00 | 491.00 |
| C2orf72 | 6.00 | 0.00 | 5.00 | 3.67 | 5.00 |
| C2orf74 | 237.00 | 237.00 | 261.00 | 245.00 | 368.00 |
| C2orf76 | 85.00 | 71.00 | 94.00 | 83.33 | 109.00 |
| C2orf81 | 72.00 | 84.00 | 89.00 | 81.67 | 97.00 |
| C2orf82 | 17.00 | 21.00 | 19.00 | 19.00 | 11.00 |

| | | | | | |
|---|---|---|---|---|---|
| C2orf88 | 7.00 | 11.00 | 6.00 | 8.00 | 13.00 |
| C3AR1 | 13.00 | 8.00 | 7.00 | 9.33 | 25.00 |
| C3orf14 | 221.00 | 267.00 | 228.00 | 238.67 | 371.00 |
| C3orf17 | 1529.00 | 1416.00 | 1532.00 | 1492.33 | 1572.00 |
| C3orf18 | 52.00 | 54.00 | 57.00 | 54.33 | 69.00 |
| C3orf20 | 15.00 | 9.00 | 9.00 | 11.00 | 5.00 |
| C3orf33 | 158.00 | 139.00 | 158.00 | 151.67 | 191.00 |
| C3orf35 | 46.00 | 46.00 | 47.00 | 46.33 | 43.00 |
| C3orf38 | 360.00 | 326.00 | 382.00 | 356.00 | 381.00 |
| C3orf52 | 327.00 | 275.00 | 287.00 | 296.33 | 201.00 |
| C3orf58 | 495.00 | 491.00 | 583.00 | 523.00 | 532.00 |
| C3orf62 | 122.00 | 137.00 | 153.00 | 137.33 | 151.00 |
| C3orf67 | 163.00 | 155.00 | 141.00 | 153.00 | 197.00 |
| C3orf70 | 13.00 | 8.00 | 17.00 | 12.67 | 16.00 |
| C3orf80 | 14.00 | 19.00 | 13.00 | 15.33 | 23.00 |
| C4orf19 | 138.00 | 114.00 | 124.00 | 125.33 | 71.00 |
| C4orf22 | 7.00 | 12.00 | 2.00 | 7.00 | 11.00 |
| C4orf27 | 262.00 | 260.00 | 223.00 | 248.33 | 333.00 |
| C4orf29 | 131.00 | 112.00 | 128.00 | 123.67 | 139.00 |
| C4orf3 | 496.00 | 442.00 | 517.00 | 485.00 | 549.00 |
| C4orf32 | 219.00 | 227.00 | 255.00 | 233.67 | 310.00 |
| C4orf33 | 107.00 | 121.00 | 106.00 | 111.33 | 132.00 |
| C4orf36 | 12.00 | 18.00 | 21.00 | 17.00 | 16.00 |
| C4orf46 | 279.00 | 265.00 | 286.00 | 276.67 | 342.00 |
| C4orf47 | 26.00 | 22.00 | 34.00 | 27.33 | 19.00 |
| C4orf48 | 223.00 | 257.00 | 287.00 | 255.67 | 336.00 |
| C5 | 135.00 | 132.00 | 153.00 | 140.00 | 231.00 |
| C5AR1 | 15.00 | 8.00 | 17.00 | 13.33 | 8.00 |
| C5orf15 | 1268.00 | 1227.00 | 1212.00 | 1235.67 | 1481.00 |
| C5orf22 | 645.00 | 538.00 | 662.00 | 615.00 | 662.00 |
| C5orf24 | 1730.00 | 1514.00 | 1850.00 | 1698.00 | 1431.00 |
| C5orf28 | 710.00 | 649.00 | 763.00 | 707.33 | 811.00 |
| C5orf30 | 1025.00 | 852.00 | 1069.00 | 982.00 | 1477.00 |
| C5orf34 | 544.00 | 468.00 | 449.00 | 487.00 | 590.00 |
| C5orf42 | 720.00 | 647.00 | 710.00 | 692.33 | 519.00 |
| C5orf45 | 731.00 | 756.00 | 850.00 | 779.00 | 702.00 |
| C5orf51 | 1484.00 | 1483.00 | 1657.00 | 1541.33 | 1546.00 |
| C5orf56 | 40.00 | 22.00 | 33.00 | 31.67 | 25.00 |
| C5orf63 | 49.00 | 41.00 | 45.00 | 45.00 | 40.00 |
| C5orf66 | 186.00 | 194.00 | 185.00 | 188.33 | 212.00 |
| C6orf1 | 819.00 | 810.00 | 775.00 | 801.33 | 943.00 |
| C6orf106 | 1371.00 | 1285.00 | 1489.00 | 1381.67 | 1484.00 |
| C6orf120 | 2322.00 | 2140.00 | 2263.00 | 2241.67 | 2694.00 |
| C6orf141 | 397.00 | 453.00 | 439.00 | 429.67 | 335.00 |
| C6orf163 | 50.00 | 59.00 | 60.00 | 56.33 | 54.00 |
| C6orf165 | 7.00 | 9.00 | 9.00 | 8.33 | 5.00 |
| C6orf203 | 384.00 | 366.00 | 370.00 | 373.33 | 389.00 |
| C6orf226 | 41.00 | 26.00 | 37.00 | 34.67 | 47.00 |

| | | | | | |
|---|---|---|---|---|---|
| C6orf48 | 8.00 | 9.00 | 6.00 | 7.67 | 13.00 |
| C6orf52 | 21.00 | 23.00 | 21.00 | 21.67 | 24.00 |
| C6orf58 | 10.00 | 1.00 | 9.00 | 6.67 | 35.00 |
| C6orf62 | 3883.00 | 3727.00 | 4293.00 | 3967.67 | 4894.00 |
| C6orf89 | 3096.00 | 2888.00 | 3023.00 | 3002.33 | 2856.00 |
| C7orf13 | 318.00 | 253.00 | 326.00 | 299.00 | 303.00 |
| C7orf25 | 403.00 | 377.00 | 406.00 | 395.33 | 531.00 |
| C7orf26 | 485.00 | 509.00 | 568.00 | 520.67 | 561.00 |
| C7orf31 | 99.00 | 84.00 | 78.00 | 87.00 | 92.00 |
| C7orf43 | 569.00 | 561.00 | 555.00 | 561.67 | 583.00 |
| C7orf49 | 953.00 | 920.00 | 1004.00 | 959.00 | 1075.00 |
| C7orf50 | 1895.00 | 1881.00 | 2080.00 | 1952.00 | 2473.00 |
| C7orf55 | 118.00 | 144.00 | 116.00 | 126.00 | 131.00 |
| C7orf57 | 21.00 | 19.00 | 22.00 | 20.67 | 17.00 |
| C7orf60 | 187.00 | 154.00 | 180.00 | 173.67 | 175.00 |
| C7orf61 | 28.00 | 27.00 | 20.00 | 25.00 | 17.00 |
| C7orf73 | 730.00 | 673.00 | 749.00 | 717.33 | 1012.00 |
| C8G | 8.00 | 24.00 | 17.00 | 16.33 | 22.00 |
| C8orf33 | 1466.00 | 1363.00 | 1432.00 | 1420.33 | 1682.00 |
| C8orf37 | 166.00 | 155.00 | 192.00 | 171.00 | 162.00 |
| C8orf4 | 9.00 | 6.00 | 16.00 | 10.33 | 7.00 |
| C8orf44 | 152.00 | 143.00 | 194.00 | 163.00 | 152.00 |
| C8orf46 | 45.00 | 36.00 | 49.00 | 43.33 | 50.00 |
| C8orf48 | 68.00 | 82.00 | 72.00 | 74.00 | 75.00 |
| C8orf58 | 364.00 | 395.00 | 368.00 | 375.67 | 526.00 |
| C8orf59 | 713.00 | 708.00 | 742.00 | 721.00 | 946.00 |
| C8orf76 | 114.00 | 141.00 | 126.00 | 127.00 | 127.00 |
| C8orf82 | 391.00 | 381.00 | 456.00 | 409.33 | 490.00 |
| C8orf88 | 311.00 | 254.00 | 345.00 | 303.33 | 400.00 |
| C9orf114 | 1636.00 | 1735.00 | 1727.00 | 1699.33 | 1890.00 |
| C9orf116 | 19.00 | 23.00 | 31.00 | 24.33 | 18.00 |
| C9orf142 | 989.00 | 1017.00 | 1013.00 | 1006.33 | 1291.00 |
| C9orf156 | 147.00 | 158.00 | 175.00 | 160.00 | 206.00 |
| C9orf16 | 410.00 | 478.00 | 489.00 | 459.00 | 635.00 |
| C9orf163 | 27.00 | 32.00 | 30.00 | 29.67 | 20.00 |
| C9orf172 | 34.00 | 30.00 | 43.00 | 35.67 | 25.00 |
| C9orf173-AS1 | 23.00 | 31.00 | 20.00 | 24.67 | 29.00 |
| C9orf24 | 6.00 | 11.00 | 8.00 | 8.33 | 10.00 |
| C9orf3 | 144.00 | 134.00 | 163.00 | 147.00 | 124.00 |
| C9orf40 | 382.00 | 374.00 | 418.00 | 391.33 | 501.00 |
| C9orf43 | 16.00 | 22.00 | 23.00 | 20.33 | 21.00 |
| C9orf47 | 13.00 | 11.00 | 17.00 | 13.67 | 17.00 |
| C9orf69 | 1965.00 | 2150.00 | 2067.00 | 2060.67 | 2245.00 |
| C9orf72 | 528.00 | 537.00 | 655.00 | 573.33 | 527.00 |
| C9orf78 | 935.00 | 943.00 | 1013.00 | 963.67 | 1199.00 |
| C9orf85 | 147.00 | 139.00 | 182.00 | 156.00 | 209.00 |
| C9orf89 | 296.00 | 280.00 | 275.00 | 283.67 | 301.00 |
| C9orf9 | 115.00 | 116.00 | 140.00 | 123.67 | 120.00 |

| | | | | | |
|---|---|---|---|---|---|
| C9orf91 | 855.00 | 841.00 | 856.00 | 850.67 | 1053.00 |
| CA11 | 18.00 | 19.00 | 19.00 | 18.67 | 23.00 |
| CA12 | 108.00 | 88.00 | 94.00 | 96.67 | 89.00 |
| CA13 | 379.00 | 373.00 | 384.00 | 378.67 | 310.00 |
| CA2 | 9.00 | 12.00 | 14.00 | 11.67 | 12.00 |
| CA3 | 69.00 | 56.00 | 98.00 | 74.33 | 61.00 |
| CA5B | 197.00 | 163.00 | 188.00 | 182.67 | 194.00 |
| CA5BP1 | 328.00 | 342.00 | 281.00 | 317.00 | 328.00 |
| CA8 | 17.00 | 13.00 | 13.00 | 14.33 | 21.00 |
| CAAP1 | 455.00 | 395.00 | 426.00 | 425.33 | 496.00 |
| CAB39 | 1521.00 | 1260.00 | 1561.00 | 1447.33 | 1494.00 |
| CAB39L | 187.00 | 160.00 | 189.00 | 178.67 | 186.00 |
| CABIN1 | 1415.00 | 1432.00 | 1501.00 | 1449.33 | 1354.00 |
| CABLES1 | 299.00 | 311.00 | 342.00 | 317.33 | 245.00 |
| CABLES2 | 158.00 | 147.00 | 196.00 | 167.00 | 164.00 |
| CABYR | 5.00 | 3.00 | 14.00 | 7.33 | 4.00 |
| CACFD1 | 26.00 | 30.00 | 37.00 | 31.00 | 28.00 |
| CACHD1 | 503.00 | 446.00 | 513.00 | 487.33 | 503.00 |
| CACNA2D1 | 777.00 | 686.00 | 829.00 | 764.00 | 719.00 |
| CACNA2D4 | 4696.00 | 4830.00 | 5203.00 | 4909.67 | 4360.00 |
| CACNB1 | 346.00 | 368.00 | 363.00 | 359.00 | 346.00 |
| CACNB3 | 527.00 | 462.00 | 531.00 | 506.67 | 510.00 |
| CACNB4 | 89.00 | 64.00 | 89.00 | 80.67 | 102.00 |
| CACNG6 | 114.00 | 106.00 | 145.00 | 121.67 | 104.00 |
| CACNG7 | 204.00 | 163.00 | 182.00 | 183.00 | 126.00 |
| CACNG8 | 1389.00 | 1381.00 | 1566.00 | 1445.33 | 955.00 |
| CACTIN | 901.00 | 956.00 | 999.00 | 952.00 | 903.00 |
| CACTIN-AS1 | 22.00 | 13.00 | 25.00 | 20.00 | 23.00 |
| CACUL1 | 1205.00 | 1179.00 | 1303.00 | 1229.00 | 1473.00 |
| CACYBP | 1660.00 | 1591.00 | 1698.00 | 1649.67 | 2542.00 |
| CAD | 6060.00 | 5929.00 | 6429.00 | 6139.33 | 5534.00 |
| CADM4 | 11.00 | 11.00 | 9.00 | 10.33 | 5.00 |
| CADPS | 20.00 | 18.00 | 27.00 | 21.67 | 22.00 |
| CADPS2 | 356.00 | 292.00 | 395.00 | 347.67 | 415.00 |
| CAHM | 43.00 | 43.00 | 33.00 | 39.67 | 35.00 |
| CALCOCO1 | 1570.00 | 1577.00 | 1748.00 | 1631.67 | 1189.00 |
| CALCOCO2 | 1734.00 | 1517.00 | 1736.00 | 1662.33 | 1930.00 |
| CALCRL | 66.00 | 35.00 | 51.00 | 50.67 | 56.00 |
| CALD1 | 3406.00 | 3086.00 | 3593.00 | 3361.67 | 4108.00 |
| CALHM2 | 23.00 | 20.00 | 25.00 | 22.67 | 19.00 |
| CALM1 | 2136.00 | 1987.00 | 2178.00 | 2100.33 | 2743.00 |
| CALM2 | 14641.00 | 13583.00 | 15636.00 | 14620.00 | 14343.00 |
| CALM3 | 2098.00 | 2012.00 | 2205.00 | 2105.00 | 2642.00 |
| CALML4 | 327.00 | 332.00 | 355.00 | 338.00 | 296.00 |
| CALML6 | 15.00 | 20.00 | 14.00 | 16.33 | 13.00 |
| CALR | 18604.00 | 19029.00 | 17844.00 | 18492.33 | 18400.00 |
| CALU | 15901.00 | 15237.00 | 15344.00 | 15494.00 | 15514.00 |
| CAMK1 | 409.00 | 416.00 | 410.00 | 411.67 | 492.00 |

| | | | | | |
|---|---|---|---|---|---|
| CAMK1D | 22.00 | 19.00 | 16.00 | 19.00 | 18.00 |
| CAMK1G | 644.00 | 631.00 | 685.00 | 653.33 | 842.00 |
| CAMK2B | 79.00 | 108.00 | 120.00 | 102.33 | 95.00 |
| CAMK2D | 402.00 | 345.00 | 432.00 | 393.00 | 404.00 |
| CAMK2G | 1256.00 | 1385.00 | 1293.00 | 1311.33 | 1449.00 |
| CAMK2N1 | 503.00 | 485.00 | 596.00 | 528.00 | 615.00 |
| CAMK2N2 | 22.00 | 26.00 | 19.00 | 22.33 | 35.00 |
| CAMK4 | 85.00 | 95.00 | 85.00 | 88.33 | 106.00 |
| CAMKK1 | 318.00 | 321.00 | 369.00 | 336.00 | 280.00 |
| CAMKK2 | 1200.00 | 1082.00 | 1245.00 | 1175.67 | 1252.00 |
| CAMKMT | 122.00 | 108.00 | 148.00 | 126.00 | 164.00 |
| CAMLG | 1249.00 | 1240.00 | 1317.00 | 1268.67 | 1384.00 |
| CAMSAP1 | 1657.00 | 1460.00 | 1753.00 | 1623.33 | 1485.00 |
| CAMSAP2 | 1718.00 | 1498.00 | 1794.00 | 1670.00 | 1626.00 |
| CAMTA1 | 356.00 | 294.00 | 366.00 | 338.67 | 463.00 |
| CAMTA2 | 5684.00 | 5647.00 | 6283.00 | 5871.33 | 5341.00 |
| CAND1 | 2625.00 | 2484.00 | 2756.00 | 2621.67 | 2579.00 |
| CANT1 | 3120.00 | 3196.00 | 3048.00 | 3121.33 | 3098.00 |
| CANX | 30446.00 | 28914.00 | 31528.00 | 30296.00 | 26840.00 |
| CAP1 | 7062.00 | 6624.00 | 7374.00 | 7020.00 | 7961.00 |
| CAP2 | 825.00 | 670.00 | 846.00 | 780.33 | 741.00 |
| CAPG | 7.00 | 8.00 | 15.00 | 10.00 | 18.00 |
| CAPN1 | 2917.00 | 2936.00 | 3014.00 | 2955.67 | 2999.00 |
| CAPN10 | 720.00 | 719.00 | 748.00 | 729.00 | 695.00 |
| CAPN10-AS1 | 216.00 | 214.00 | 195.00 | 208.33 | 154.00 |
| CAPN11 | 19.00 | 15.00 | 14.00 | 16.00 | 12.00 |
| CAPN15 | 2816.00 | 2854.00 | 3079.00 | 2916.33 | 2892.00 |
| CAPN2 | 12275.00 | 11974.00 | 13401.00 | 12550.00 | 11920.00 |
| CAPN3 | 116.00 | 118.00 | 124.00 | 119.33 | 109.00 |
| CAPN5 | 382.00 | 394.00 | 409.00 | 395.00 | 285.00 |
| CAPN7 | 708.00 | 715.00 | 801.00 | 741.33 | 657.00 |
| CAPNS1 | 3485.00 | 3310.00 | 3565.00 | 3453.33 | 4208.00 |
| CAPNS2 | 4.00 | 1.00 | 0.00 | 1.67 | 0.00 |
| CAPRIN1 | 6160.00 | 5614.00 | 6505.00 | 6093.00 | 6869.00 |
| CAPRIN2 | 4503.00 | 4485.00 | 5163.00 | 4717.00 | 5108.00 |
| CAPS | 208.00 | 239.00 | 232.00 | 226.33 | 199.00 |
| CAPS2 | 62.00 | 80.00 | 64.00 | 68.67 | 75.00 |
| CAPZA1 | 4012.00 | 3606.00 | 4231.00 | 3949.67 | 5496.00 |
| CAPZA2 | 1108.00 | 1106.00 | 1276.00 | 1163.33 | 1589.00 |
| CAPZB | 2250.00 | 2203.00 | 2268.00 | 2240.33 | 2398.00 |
| CARD10 | 1566.00 | 1636.00 | 1581.00 | 1594.33 | 1615.00 |
| CARD6 | 172.00 | 150.00 | 185.00 | 169.00 | 152.00 |
| CARD8 | 2281.00 | 2101.00 | 2328.00 | 2236.67 | 3017.00 |
| CARD8-AS1 | 128.00 | 114.00 | 132.00 | 124.67 | 130.00 |
| CARD9 | 46.00 | 56.00 | 38.00 | 46.67 | 25.00 |
| CARF | 317.00 | 299.00 | 308.00 | 308.00 | 276.00 |
| CARHSP1 | 193.00 | 222.00 | 229.00 | 214.67 | 366.00 |
| CARKD | 1105.00 | 1135.00 | 1021.00 | 1087.00 | 1117.00 |

| | | | | | |
|---|---|---|---|---|---|
| CARM1 | 1818.00 | 1644.00 | 1934.00 | 1798.67 | 1962.00 |
| CARNMT1 | 311.00 | 245.00 | 366.00 | 307.33 | 386.00 |
| CARS | 276.00 | 260.00 | 282.00 | 272.67 | 277.00 |
| CARS2 | 1117.00 | 1108.00 | 1284.00 | 1169.67 | 1275.00 |
| CASC10 | 60.00 | 55.00 | 70.00 | 61.67 | 64.00 |
| CASC11 | 6.00 | 3.00 | 5.00 | 4.67 | 2.00 |
| CASC15 | 5.00 | 6.00 | 4.00 | 5.00 | 1.00 |
| CASC2 | 44.00 | 26.00 | 37.00 | 35.67 | 48.00 |
| CASC3 | 2805.00 | 2688.00 | 3027.00 | 2840.00 | 2892.00 |
| CASC4 | 2050.00 | 1856.00 | 2135.00 | 2013.67 | 2020.00 |
| CASC5 | 325.00 | 298.00 | 293.00 | 305.33 | 307.00 |
| CASC9 | 25.00 | 21.00 | 25.00 | 23.67 | 28.00 |
| CASD1 | 531.00 | 497.00 | 646.00 | 558.00 | 548.00 |
| CASK | 386.00 | 349.00 | 402.00 | 379.00 | 398.00 |
| CASKIN2 | 513.00 | 471.00 | 545.00 | 509.67 | 518.00 |
| CASP1 | 59.00 | 63.00 | 56.00 | 59.33 | 69.00 |
| CASP10 | 13.00 | 10.00 | 17.00 | 13.33 | 12.00 |
| CASP2 | 948.00 | 862.00 | 904.00 | 904.67 | 977.00 |
| CASP3 | 469.00 | 426.00 | 512.00 | 469.00 | 711.00 |
| CASP4 | 1997.00 | 2012.00 | 2185.00 | 2064.67 | 2224.00 |
| CASP5 | 12.00 | 5.00 | 0.00 | 5.67 | 0.00 |
| CASP6 | 97.00 | 66.00 | 104.00 | 89.00 | 96.00 |
| CASP7 | 372.00 | 331.00 | 394.00 | 365.67 | 346.00 |
| CASP8 | 837.00 | 785.00 | 825.00 | 815.67 | 862.00 |
| CASP8AP2 | 746.00 | 696.00 | 811.00 | 751.00 | 712.00 |
| CASP9 | 238.00 | 233.00 | 245.00 | 238.67 | 250.00 |
| CASS4 | 68.00 | 58.00 | 78.00 | 68.00 | 125.00 |
| CAST | 3277.00 | 2949.00 | 3425.00 | 3217.00 | 3527.00 |
| CAT | 362.00 | 343.00 | 374.00 | 359.67 | 474.00 |
| CATSPER1 | 165.00 | 179.00 | 191.00 | 178.33 | 232.00 |
| CATSPER2 | 129.00 | 125.00 | 130.00 | 128.00 | 138.00 |
| CATSPER2P1 | 27.00 | 23.00 | 24.00 | 24.67 | 22.00 |
| CATSPER3 | 33.00 | 30.00 | 32.00 | 31.67 | 31.00 |
| CATSPERG | 6.00 | 3.00 | 2.00 | 3.67 | 2.00 |
| CAV1 | 11457.00 | 9991.00 | 11245.00 | 10897.67 | 14862.00 |
| CAV2 | 944.00 | 779.00 | 978.00 | 900.33 | 1135.00 |
| CBARP | 313.00 | 313.00 | 337.00 | 321.00 | 268.00 |
| CBFA2T2 | 1106.00 | 1035.00 | 1129.00 | 1090.00 | 941.00 |
| CBFB | 1525.00 | 1408.00 | 1687.00 | 1540.00 | 2296.00 |
| CBL | 1804.00 | 1631.00 | 1939.00 | 1791.33 | 1442.00 |
| CBLB | 802.00 | 681.00 | 887.00 | 790.00 | 867.00 |
| CBLL1 | 618.00 | 623.00 | 712.00 | 651.00 | 732.00 |
| CBLN3 | 74.00 | 62.00 | 68.00 | 68.00 | 45.00 |
| CBR1 | 935.00 | 954.00 | 1034.00 | 974.33 | 1274.00 |
| CBR3 | 47.00 | 58.00 | 47.00 | 50.67 | 82.00 |
| CBR3-AS1 | 11.00 | 22.00 | 22.00 | 18.33 | 16.00 |
| CBR4 | 485.00 | 409.00 | 489.00 | 461.00 | 551.00 |
| CBS | 171.00 | 206.00 | 191.00 | 189.33 | 189.00 |

| | | | | | |
|---|---|---|---|---|---|
| CBWD1 | 311.00 | 283.00 | 334.00 | 309.33 | 406.00 |
| CBWD2 | 238.00 | 297.00 | 292.00 | 275.67 | 260.00 |
| CBWD3 | 55.00 | 64.00 | 69.00 | 62.67 | 40.00 |
| CBWD5 | 297.00 | 307.00 | 362.00 | 322.00 | 304.00 |
| CBWD6 | 146.00 | 192.00 | 181.00 | 173.00 | 171.00 |
| CBX1 | 1309.00 | 1281.00 | 1415.00 | 1335.00 | 1225.00 |
| CBX2 | 182.00 | 190.00 | 139.00 | 170.33 | 216.00 |
| CBX3 | 4392.00 | 4179.00 | 4552.00 | 4374.33 | 5975.00 |
| CBX3P2 | 31.00 | 30.00 | 32.00 | 31.00 | 18.00 |
| CBX4 | 541.00 | 596.00 | 627.00 | 588.00 | 624.00 |
| CBX5 | 2245.00 | 2007.00 | 2407.00 | 2219.67 | 2502.00 |
| CBX6 | 2186.00 | 2009.00 | 2359.00 | 2184.67 | 2485.00 |
| CBX7 | 301.00 | 351.00 | 365.00 | 339.00 | 324.00 |
| CBX8 | 119.00 | 132.00 | 115.00 | 122.00 | 145.00 |
| CBY1 | 369.00 | 377.00 | 392.00 | 379.33 | 562.00 |
| CC2D1A | 1486.00 | 1458.00 | 1513.00 | 1485.67 | 1445.00 |
| CC2D1B | 1516.00 | 1557.00 | 1582.00 | 1551.67 | 1450.00 |
| CC2D2A | 537.00 | 466.00 | 543.00 | 515.33 | 556.00 |
| CCAR1 | 3962.00 | 3722.00 | 4016.00 | 3900.00 | 3991.00 |
| CCAR2 | 1966.00 | 1993.00 | 2044.00 | 2001.00 | 2176.00 |
| CCBE1 | 5055.00 | 4624.00 | 4785.00 | 4821.33 | 3860.00 |
| CCBL1 | 619.00 | 571.00 | 688.00 | 626.00 | 730.00 |
| CCBL2 | 865.00 | 771.00 | 885.00 | 840.33 | 1090.00 |
| CCDC101 | 209.00 | 210.00 | 205.00 | 208.00 | 257.00 |
| CCDC102A | 296.00 | 284.00 | 283.00 | 287.67 | 338.00 |
| CCDC102B | 25.00 | 32.00 | 32.00 | 29.67 | 35.00 |
| CCDC103 | 16.00 | 15.00 | 13.00 | 14.67 | 28.00 |
| CCDC106 | 261.00 | 269.00 | 274.00 | 268.00 | 292.00 |
| CCDC107 | 194.00 | 225.00 | 194.00 | 204.33 | 244.00 |
| CCDC109B | 170.00 | 132.00 | 157.00 | 153.00 | 258.00 |
| CCDC110 | 7.00 | 7.00 | 15.00 | 9.67 | 12.00 |
| CCDC112 | 256.00 | 221.00 | 241.00 | 239.33 | 320.00 |
| CCDC113 | 219.00 | 199.00 | 230.00 | 216.00 | 215.00 |
| CCDC115 | 387.00 | 399.00 | 420.00 | 402.00 | 440.00 |
| CCDC116 | 6.00 | 9.00 | 13.00 | 9.33 | 11.00 |
| CCDC117 | 309.00 | 310.00 | 316.00 | 311.67 | 398.00 |
| CCDC12 | 351.00 | 376.00 | 354.00 | 360.33 | 473.00 |
| CCDC120 | 152.00 | 174.00 | 183.00 | 169.67 | 183.00 |
| CCDC121 | 14.00 | 13.00 | 9.00 | 12.00 | 17.00 |
| CCDC122 | 96.00 | 115.00 | 112.00 | 107.67 | 128.00 |
| CCDC124 | 975.00 | 966.00 | 995.00 | 978.67 | 1280.00 |
| CCDC125 | 6.00 | 22.00 | 21.00 | 16.33 | 29.00 |
| CCDC126 | 159.00 | 166.00 | 162.00 | 162.33 | 189.00 |
| CCDC127 | 348.00 | 357.00 | 332.00 | 345.67 | 317.00 |
| CCDC130 | 1049.00 | 1115.00 | 1163.00 | 1109.00 | 1026.00 |
| CCDC134 | 73.00 | 76.00 | 81.00 | 76.67 | 82.00 |
| CCDC136 | 759.00 | 750.00 | 750.00 | 753.00 | 556.00 |
| CCDC137 | 1359.00 | 1324.00 | 1358.00 | 1347.00 | 1577.00 |

| | | | | | |
|---|---|---|---|---|---|
| CCDC138 | 140.00 | 139.00 | 159.00 | 146.00 | 208.00 |
| CCDC14 | 4876.00 | 4762.00 | 5088.00 | 4908.67 | 5056.00 |
| CCDC142 | 502.00 | 454.00 | 531.00 | 495.67 | 381.00 |
| CCDC144A | 25.00 | 31.00 | 28.00 | 28.00 | 20.00 |
| CCDC144B | 60.00 | 41.00 | 39.00 | 46.67 | 28.00 |
| CCDC146 | 187.00 | 151.00 | 194.00 | 177.33 | 157.00 |
| CCDC148 | 5.00 | 11.00 | 10.00 | 8.67 | 16.00 |
| CCDC149 | 228.00 | 231.00 | 265.00 | 241.33 | 166.00 |
| CCDC15 | 148.00 | 131.00 | 138.00 | 139.00 | 205.00 |
| CCDC150 | 433.00 | 455.00 | 379.00 | 422.33 | 443.00 |
| CCDC151 | 10.00 | 10.00 | 11.00 | 10.33 | 10.00 |
| CCDC152 | 119.00 | 103.00 | 107.00 | 109.67 | 141.00 |
| CCDC153 | 4.00 | 4.00 | 2.00 | 3.33 | 0.00 |
| CCDC154 | 19.00 | 16.00 | 20.00 | 18.33 | 19.00 |
| CCDC157 | 35.00 | 40.00 | 27.00 | 34.00 | 47.00 |
| CCDC159 | 133.00 | 158.00 | 220.00 | 170.33 | 146.00 |
| CCDC163P | 214.00 | 240.00 | 229.00 | 227.67 | 197.00 |
| CCDC167 | 177.00 | 145.00 | 134.00 | 152.00 | 195.00 |
| CCDC168 | 5.00 | 8.00 | 4.00 | 5.67 | 4.00 |
| CCDC169 | 33.00 | 40.00 | 53.00 | 42.00 | 46.00 |
| CCDC17 | 139.00 | 165.00 | 219.00 | 174.33 | 128.00 |
| CCDC170 | 9.00 | 6.00 | 4.00 | 6.33 | 3.00 |
| CCDC171 | 85.00 | 74.00 | 82.00 | 80.33 | 122.00 |
| CCDC173 | 6.00 | 4.00 | 6.00 | 5.33 | 9.00 |
| CCDC174 | 493.00 | 493.00 | 561.00 | 515.67 | 521.00 |
| CCDC176 | 72.00 | 69.00 | 89.00 | 76.67 | 79.00 |
| CCDC18 | 708.00 | 623.00 | 672.00 | 667.67 | 700.00 |
| CCDC181 | 99.00 | 66.00 | 90.00 | 85.00 | 111.00 |
| CCDC183 | 17.00 | 36.00 | 24.00 | 25.67 | 13.00 |
| CCDC183-AS1 | 66.00 | 51.00 | 56.00 | 57.67 | 50.00 |
| CCDC186 | 300.00 | 279.00 | 327.00 | 302.00 | 278.00 |
| CCDC22 | 569.00 | 589.00 | 571.00 | 576.33 | 661.00 |
| CCDC24 | 90.00 | 89.00 | 82.00 | 87.00 | 122.00 |
| CCDC25 | 520.00 | 508.00 | 561.00 | 529.67 | 608.00 |
| CCDC28A | 328.00 | 322.00 | 338.00 | 329.33 | 364.00 |
| CCDC28B | 58.00 | 67.00 | 62.00 | 62.33 | 104.00 |
| CCDC30 | 22.00 | 12.00 | 20.00 | 18.00 | 10.00 |
| CCDC34 | 243.00 | 257.00 | 306.00 | 268.67 | 361.00 |
| CCDC36 | 153.00 | 110.00 | 101.00 | 121.33 | 109.00 |
| CCDC38 | 4.00 | 0.00 | 0.00 | 1.33 | 2.00 |
| CCDC39 | 4.00 | 0.00 | 0.00 | 1.33 | 4.00 |
| CCDC40 | 42.00 | 54.00 | 64.00 | 53.33 | 36.00 |
| CCDC43 | 456.00 | 458.00 | 488.00 | 467.33 | 570.00 |
| CCDC47 | 5280.00 | 5194.00 | 5469.00 | 5314.33 | 5806.00 |
| CCDC50 | 2275.00 | 2169.00 | 2438.00 | 2294.00 | 2007.00 |
| CCDC51 | 354.00 | 365.00 | 357.00 | 358.67 | 415.00 |
| CCDC53 | 338.00 | 352.00 | 366.00 | 352.00 | 447.00 |
| CCDC57 | 1286.00 | 1274.00 | 1302.00 | 1287.33 | 1102.00 |

| | | | | | |
|---|---|---|---|---|---|
| CCDC58 | 461.00 | 431.00 | 478.00 | 456.67 | 659.00 |
| CCDC59 | 598.00 | 609.00 | 588.00 | 598.33 | 852.00 |
| CCDC6 | 1582.00 | 1376.00 | 1811.00 | 1589.67 | 1884.00 |
| CCDC61 | 239.00 | 246.00 | 268.00 | 251.00 | 337.00 |
| CCDC62 | 23.00 | 13.00 | 29.00 | 21.67 | 22.00 |
| CCDC66 | 641.00 | 702.00 | 748.00 | 697.00 | 680.00 |
| CCDC69 | 119.00 | 101.00 | 83.00 | 101.00 | 90.00 |
| CCDC7 | 142.00 | 153.00 | 164.00 | 153.00 | 175.00 |
| CCDC70 | 6.00 | 16.00 | 15.00 | 12.33 | 9.00 |
| CCDC71 | 413.00 | 465.00 | 465.00 | 447.67 | 566.00 |
| CCDC71L | 550.00 | 465.00 | 629.00 | 548.00 | 752.00 |
| CCDC74A | 17.00 | 19.00 | 30.00 | 22.00 | 50.00 |
| CCDC74B | 8.00 | 1.00 | 5.00 | 4.67 | 13.00 |
| CCDC77 | 528.00 | 573.00 | 594.00 | 565.00 | 722.00 |
| CCDC78 | 561.00 | 588.00 | 628.00 | 592.33 | 694.00 |
| CCDC8 | 56.00 | 77.00 | 70.00 | 67.67 | 63.00 |
| CCDC80 | 5540.00 | 5470.00 | 5702.00 | 5570.67 | 8368.00 |
| CCDC82 | 715.00 | 647.00 | 676.00 | 679.33 | 769.00 |
| CCDC84 | 254.00 | 303.00 | 295.00 | 284.00 | 303.00 |
| CCDC85A | 12.00 | 8.00 | 6.00 | 8.67 | 6.00 |
| CCDC85B | 915.00 | 900.00 | 967.00 | 927.33 | 1840.00 |
| CCDC85C | 433.00 | 423.00 | 463.00 | 439.67 | 450.00 |
| CCDC86 | 1956.00 | 1951.00 | 2036.00 | 1981.00 | 2182.00 |
| CCDC87 | 7.00 | 5.00 | 9.00 | 7.00 | 13.00 |
| CCDC88A | 1583.00 | 1498.00 | 1687.00 | 1589.33 | 1467.00 |
| CCDC88B | 49.00 | 52.00 | 49.00 | 50.00 | 80.00 |
| CCDC88C | 437.00 | 504.00 | 547.00 | 496.00 | 408.00 |
| CCDC9 | 440.00 | 453.00 | 492.00 | 461.67 | 539.00 |
| CCDC90B | 616.00 | 593.00 | 708.00 | 639.00 | 808.00 |
| CCDC91 | 539.00 | 541.00 | 557.00 | 545.67 | 598.00 |
| CCDC92 | 25.00 | 22.00 | 22.00 | 23.00 | 25.00 |
| CCDC93 | 2002.00 | 1962.00 | 2142.00 | 2035.33 | 1788.00 |
| CCDC94 | 258.00 | 230.00 | 229.00 | 239.00 | 317.00 |
| CCDC96 | 13.00 | 16.00 | 19.00 | 16.00 | 9.00 |
| CCDC97 | 514.00 | 498.00 | 490.00 | 500.67 | 520.00 |
| CCHCR1 | 45.00 | 44.00 | 38.00 | 42.33 | 36.00 |
| CCK | 5.00 | 6.00 | 6.00 | 5.67 | 0.00 |
| CCL2 | 5.00 | 7.00 | 5.00 | 5.67 | 7.00 |
| CCL20 | 60.00 | 52.00 | 70.00 | 60.67 | 114.00 |
| CCL28 | 178.00 | 171.00 | 200.00 | 183.00 | 168.00 |
| CCM2 | 687.00 | 819.00 | 782.00 | 762.67 | 875.00 |
| CCNA1 | 26.00 | 35.00 | 35.00 | 32.00 | 51.00 |
| CCNA2 | 555.00 | 498.00 | 572.00 | 541.67 | 713.00 |
| CCNB1 | 2506.00 | 2294.00 | 2453.00 | 2417.67 | 3193.00 |
| CCNB1IP1 | 1669.00 | 1707.00 | 1963.00 | 1779.67 | 1987.00 |
| CCNB2 | 942.00 | 841.00 | 912.00 | 898.33 | 1099.00 |
| CCNB3 | 8.00 | 14.00 | 7.00 | 9.67 | 7.00 |
| CCNC | 1033.00 | 898.00 | 1073.00 | 1001.33 | 1351.00 |

| | | | | | |
|---|---|---|---|---|---|
| CCND1 | 12974.00 | 11982.00 | 13295.00 | 12750.33 | 13447.00 |
| CCND3 | 409.00 | 355.00 | 361.00 | 375.00 | 571.00 |
| CCNDBP1 | 1142.00 | 1102.00 | 1152.00 | 1132.00 | 1200.00 |
| CCNE1 | 115.00 | 122.00 | 104.00 | 113.67 | 157.00 |
| CCNE2 | 137.00 | 131.00 | 144.00 | 137.33 | 240.00 |
| CCNF | 478.00 | 443.00 | 435.00 | 452.00 | 577.00 |
| CCNG1 | 1448.00 | 1269.00 | 1561.00 | 1426.00 | 1643.00 |
| CCNG2 | 257.00 | 262.00 | 313.00 | 277.33 | 278.00 |
| CCNH | 1040.00 | 1052.00 | 1080.00 | 1057.33 | 1417.00 |
| CCNI | 2482.00 | 2206.00 | 2692.00 | 2460.00 | 2894.00 |
| CCNJ | 476.00 | 488.00 | 491.00 | 485.00 | 568.00 |
| CCNJL | 571.00 | 489.00 | 527.00 | 529.00 | 542.00 |
| CCNK | 1014.00 | 891.00 | 1029.00 | 978.00 | 1047.00 |
| CCNL1 | 3733.00 | 3541.00 | 3944.00 | 3739.33 | 4061.00 |
| CCNL2 | 8144.00 | 8096.00 | 8730.00 | 8323.33 | 7788.00 |
| CCNO | 344.00 | 365.00 | 423.00 | 377.33 | 394.00 |
| CCNT1 | 1598.00 | 1401.00 | 1659.00 | 1552.67 | 1530.00 |
| CCNT2 | 1734.00 | 1726.00 | 1932.00 | 1797.33 | 1807.00 |
| CCNT2-AS1 | 17.00 | 14.00 | 21.00 | 17.33 | 8.00 |
| CCNY | 2009.00 | 1972.00 | 2294.00 | 2091.67 | 2501.00 |
| CCNYL1 | 476.00 | 455.00 | 512.00 | 481.00 | 618.00 |
| CCNYL2 | 198.00 | 237.00 | 219.00 | 218.00 | 195.00 |
| CCP110 | 423.00 | 422.00 | 413.00 | 419.33 | 505.00 |
| CCPG1 | 458.00 | 490.00 | 579.00 | 509.00 | 428.00 |
| CCR1 | 29.00 | 13.00 | 20.00 | 20.67 | 31.00 |
| CCR10 | 17.00 | 13.00 | 26.00 | 18.67 | 23.00 |
| CCRL2 | 38.00 | 33.00 | 42.00 | 37.67 | 22.00 |
| CCS | 892.00 | 980.00 | 955.00 | 942.33 | 948.00 |
| CCSAP | 163.00 | 122.00 | 140.00 | 141.67 | 181.00 |
| CCSER1 | 15.00 | 23.00 | 19.00 | 19.00 | 16.00 |
| CCSER2 | 787.00 | 686.00 | 831.00 | 768.00 | 722.00 |
| CCT2 | 6300.00 | 6099.00 | 6476.00 | 6291.67 | 7365.00 |
| CCT3 | 12469.00 | 12089.00 | 12750.00 | 12436.00 | 15530.00 |
| CCT4 | 6958.00 | 6752.00 | 7882.00 | 7197.33 | 9910.00 |
| CCT5 | 10281.00 | 9764.00 | 10709.00 | 10251.33 | 13907.00 |
| CCT6A | 8906.00 | 8551.00 | 9852.00 | 9103.00 | 11110.00 |
| CCT6B | 18.00 | 27.00 | 18.00 | 21.00 | 43.00 |
| CCT6P1 | 188.00 | 189.00 | 192.00 | 189.67 | 242.00 |
| CCT6P3 | 57.00 | 48.00 | 83.00 | 62.67 | 52.00 |
| CCT7 | 8514.00 | 8107.00 | 9108.00 | 8576.33 | 10739.00 |
| CCT8 | 6098.00 | 5823.00 | 6655.00 | 6192.00 | 8842.00 |
| CCZ1 | 478.00 | 423.00 | 470.00 | 457.00 | 595.00 |
| CCZ1B | 176.00 | 134.00 | 197.00 | 169.00 | 228.00 |
| CD101 | 27.00 | 20.00 | 16.00 | 21.00 | 16.00 |
| CD109 | 2390.00 | 2087.00 | 2557.00 | 2344.67 | 2039.00 |
| CD14 | 7.00 | 10.00 | 13.00 | 10.00 | 5.00 |
| CD151 | 8586.00 | 8749.00 | 8440.00 | 8591.67 | 8742.00 |
| CD163L1 | 2780.00 | 2762.00 | 2972.00 | 2838.00 | 3249.00 |

| | | | | | |
|---|---|---|---|---|---|
| CD164 | 7886.00 | 7076.00 | 7851.00 | 7604.33 | 8855.00 |
| CD177 | 110.00 | 125.00 | 138.00 | 124.33 | 108.00 |
| CD200 | 81.00 | 78.00 | 84.00 | 81.00 | 74.00 |
| CD22 | 144.00 | 126.00 | 151.00 | 140.33 | 123.00 |
| CD226 | 14.00 | 18.00 | 12.00 | 14.67 | 20.00 |
| CD24 | 4.00 | 1.00 | 7.00 | 4.00 | 11.00 |
| CD27 | 4.00 | 6.00 | 3.00 | 4.33 | 1.00 |
| CD274 | 2777.00 | 2518.00 | 2666.00 | 2653.67 | 2195.00 |
| CD276 | 1170.00 | 1104.00 | 1169.00 | 1147.67 | 1052.00 |
| CD27-AS1 | 1366.00 | 1474.00 | 1533.00 | 1457.67 | 1654.00 |
| CD2AP | 801.00 | 709.00 | 879.00 | 796.33 | 869.00 |
| CD2BP2 | 1753.00 | 1777.00 | 1881.00 | 1803.67 | 2054.00 |
| CD302 | 27.00 | 25.00 | 18.00 | 23.33 | 28.00 |
| CD320 | 1854.00 | 1939.00 | 1801.00 | 1864.67 | 2934.00 |
| CD33 | 8.00 | 10.00 | 14.00 | 10.67 | 15.00 |
| CD38 | 22.00 | 11.00 | 6.00 | 13.00 | 13.00 |
| CD3EAP | 746.00 | 738.00 | 801.00 | 761.67 | 801.00 |
| CD44 | 24800.00 | 24398.00 | 24999.00 | 24732.33 | 29006.00 |
| CD46 | 3869.00 | 3647.00 | 3792.00 | 3769.33 | 3529.00 |
| CD47 | 2952.00 | 2749.00 | 2865.00 | 2855.33 | 3362.00 |
| CD55 | 2419.00 | 2351.00 | 2435.00 | 2401.67 | 2393.00 |
| CD58 | 394.00 | 350.00 | 298.00 | 347.33 | 305.00 |
| CD59 | 21459.00 | 21158.00 | 20117.00 | 20911.33 | 26405.00 |
| CD63 | 12546.00 | 13395.00 | 12117.00 | 12686.00 | 10008.00 |
| CD68 | 1385.00 | 1435.00 | 1333.00 | 1384.33 | 1564.00 |
| CD70 | 61.00 | 45.00 | 54.00 | 53.33 | 58.00 |
| CD74 | 44.00 | 49.00 | 38.00 | 43.67 | 135.00 |
| CD81 | 5442.00 | 5469.00 | 5026.00 | 5312.33 | 6162.00 |
| CD81-AS1 | 5.00 | 5.00 | 3.00 | 4.33 | 3.00 |
| CD82 | 128.00 | 132.00 | 123.00 | 127.67 | 83.00 |
| CD83 | 178.00 | 205.00 | 198.00 | 193.67 | 423.00 |
| CD9 | 1222.00 | 1250.00 | 1214.00 | 1228.67 | 1240.00 |
| CD93 | 38.00 | 33.00 | 35.00 | 35.33 | 34.00 |
| CD96 | 20.00 | 15.00 | 12.00 | 15.67 | 19.00 |
| CD99 | 23.00 | 25.00 | 37.00 | 28.33 | 34.00 |
| CD99L2 | 1493.00 | 1430.00 | 1347.00 | 1423.33 | 1430.00 |
| CDA | 240.00 | 261.00 | 293.00 | 264.67 | 231.00 |
| CDADC1 | 154.00 | 128.00 | 155.00 | 145.67 | 159.00 |
| CDAN1 | 811.00 | 787.00 | 835.00 | 811.00 | 673.00 |
| CDC123 | 2945.00 | 2552.00 | 2825.00 | 2774.00 | 3628.00 |
| CDC14A | 188.00 | 172.00 | 189.00 | 183.00 | 160.00 |
| CDC14B | 385.00 | 391.00 | 405.00 | 393.67 | 416.00 |
| CDC14C | 4.00 | 2.00 | 2.00 | 2.67 | 1.00 |
| CDC16 | 1607.00 | 1591.00 | 1729.00 | 1642.33 | 1828.00 |
| CDC20 | 2333.00 | 2280.00 | 2214.00 | 2275.67 | 3130.00 |
| CDC20B | 13.00 | 9.00 | 9.00 | 10.33 | 18.00 |
| CDC23 | 944.00 | 892.00 | 957.00 | 931.00 | 1025.00 |
| CDC25A | 292.00 | 291.00 | 279.00 | 287.33 | 454.00 |

| | | | | | |
|---|---|---|---|---|---|
| CDC25B | 1664.00 | 1627.00 | 1694.00 | 1661.67 | 1920.00 |
| CDC25C | 174.00 | 166.00 | 167.00 | 169.00 | 244.00 |
| CDC26 | 164.00 | 149.00 | 196.00 | 169.67 | 190.00 |
| CDC27 | 2914.00 | 2520.00 | 3060.00 | 2831.33 | 3071.00 |
| CDC34 | 1177.00 | 1240.00 | 1280.00 | 1232.33 | 1474.00 |
| CDC37 | 5333.00 | 5715.00 | 5842.00 | 5630.00 | 5770.00 |
| CDC37L1 | 289.00 | 282.00 | 353.00 | 308.00 | 354.00 |
| CDC37L1-AS1 | 15.00 | 13.00 | 8.00 | 12.00 | 17.00 |
| CDC40 | 541.00 | 546.00 | 575.00 | 554.00 | 667.00 |
| CDC42 | 2369.00 | 2172.00 | 2346.00 | 2295.67 | 2664.00 |
| CDC42BPA | 2203.00 | 2063.00 | 2375.00 | 2213.67 | 2136.00 |
| CDC42BPB | 3904.00 | 3686.00 | 4266.00 | 3952.00 | 3328.00 |
| CDC42EP1 | 4500.00 | 4388.00 | 4392.00 | 4426.67 | 4541.00 |
| CDC42EP2 | 1091.00 | 1062.00 | 1046.00 | 1066.33 | 907.00 |
| CDC42EP3 | 6735.00 | 6034.00 | 6877.00 | 6548.67 | 6675.00 |
| CDC42EP4 | 484.00 | 498.00 | 498.00 | 493.33 | 483.00 |
| CDC42SE1 | 2082.00 | 2007.00 | 2194.00 | 2094.33 | 2158.00 |
| CDC42SE2 | 587.00 | 571.00 | 646.00 | 601.33 | 698.00 |
| CDC45 | 301.00 | 381.00 | 324.00 | 335.33 | 530.00 |
| CDC5L | 1700.00 | 1647.00 | 1875.00 | 1740.67 | 1914.00 |
| CDC6 | 444.00 | 442.00 | 439.00 | 441.67 | 618.00 |
| CDC7 | 770.00 | 808.00 | 802.00 | 793.33 | 1110.00 |
| CDC73 | 974.00 | 791.00 | 1032.00 | 932.33 | 1030.00 |
| CDCA2 | 315.00 | 272.00 | 255.00 | 280.67 | 346.00 |
| CDCA3 | 456.00 | 431.00 | 400.00 | 429.00 | 618.00 |
| CDCA4 | 732.00 | 750.00 | 725.00 | 735.67 | 1180.00 |
| CDCA5 | 1193.00 | 1187.00 | 1354.00 | 1244.67 | 1798.00 |
| CDCA7 | 1197.00 | 1223.00 | 1202.00 | 1207.33 | 1683.00 |
| CDCA7L | 1313.00 | 1297.00 | 1487.00 | 1365.67 | 1647.00 |
| CDCA8 | 867.00 | 819.00 | 778.00 | 821.33 | 1126.00 |
| CDCP1 | 55.00 | 60.00 | 59.00 | 58.00 | 42.00 |
| CDH10 | 5.00 | 0.00 | 0.00 | 1.67 | 1.00 |
| CDH12 | 269.00 | 236.00 | 309.00 | 271.33 | 230.00 |
| CDH13 | 4931.00 | 4949.00 | 5242.00 | 5040.67 | 2502.00 |
| CDH15 | 12.00 | 5.00 | 15.00 | 10.67 | 6.00 |
| CDH18 | 78.00 | 58.00 | 108.00 | 81.33 | 68.00 |
| CDH2 | 3934.00 | 3686.00 | 3989.00 | 3869.67 | 3318.00 |
| CDH24 | 2124.00 | 2091.00 | 2267.00 | 2160.67 | 1014.00 |
| CDH26 | 4.00 | 7.00 | 10.00 | 7.00 | 3.00 |
| CDH5 | 3303.00 | 3329.00 | 3186.00 | 3272.67 | 1307.00 |
| CDH6 | 47.00 | 44.00 | 48.00 | 46.33 | 30.00 |
| CDHR3 | 172.00 | 157.00 | 235.00 | 188.00 | 112.00 |
| CDIPT | 1182.00 | 1119.00 | 1219.00 | 1173.33 | 1185.00 |
| CDK1 | 1573.00 | 1405.00 | 1519.00 | 1499.00 | 1978.00 |
| CDK10 | 2315.00 | 2463.00 | 2523.00 | 2433.67 | 2585.00 |
| CDK11A | 142.00 | 138.00 | 154.00 | 144.67 | 137.00 |
| CDK11B | 654.00 | 623.00 | 635.00 | 637.33 | 677.00 |
| CDK12 | 2449.00 | 2322.00 | 2702.00 | 2491.00 | 2173.00 |

| | | | | | |
|---|---|---|---|---|---|
| CDK13 | 1977.00 | 1836.00 | 2141.00 | 1984.67 | 1778.00 |
| CDK14 | 395.00 | 339.00 | 438.00 | 390.67 | 457.00 |
| CDK15 | 19.00 | 20.00 | 25.00 | 21.33 | 7.00 |
| CDK16 | 2816.00 | 2861.00 | 2929.00 | 2868.67 | 3074.00 |
| CDK17 | 2023.00 | 1805.00 | 2108.00 | 1978.67 | 2254.00 |
| CDK18 | 47.00 | 38.00 | 37.00 | 40.67 | 37.00 |
| CDK19 | 646.00 | 579.00 | 730.00 | 651.67 | 682.00 |
| CDK2 | 782.00 | 732.00 | 774.00 | 762.67 | 1081.00 |
| CDK20 | 88.00 | 93.00 | 94.00 | 91.67 | 98.00 |
| CDK2AP1 | 761.00 | 737.00 | 781.00 | 759.67 | 922.00 |
| CDK2AP2 | 520.00 | 415.00 | 465.00 | 466.67 | 504.00 |
| CDK4 | 3752.00 | 3557.00 | 3785.00 | 3698.00 | 4647.00 |
| CDK5 | 423.00 | 358.00 | 400.00 | 393.67 | 508.00 |
| CDK5R1 | 174.00 | 179.00 | 171.00 | 174.67 | 202.00 |
| CDK5R2 | 10.00 | 6.00 | 12.00 | 9.33 | 5.00 |
| CDK5RAP1 | 1229.00 | 1183.00 | 1325.00 | 1245.67 | 1309.00 |
| CDK5RAP2 | 1394.00 | 1239.00 | 1396.00 | 1343.00 | 1427.00 |
| CDK5RAP3 | 5775.00 | 6305.00 | 6651.00 | 6243.67 | 6798.00 |
| CDK6 | 3673.00 | 2995.00 | 3684.00 | 3450.67 | 3737.00 |
| CDK7 | 7.00 | 11.00 | 9.00 | 9.00 | 2.00 |
| CDK8 | 393.00 | 367.00 | 419.00 | 393.00 | 489.00 |
| CDK9 | 889.00 | 819.00 | 853.00 | 853.67 | 1014.00 |
| CDKAL1 | 367.00 | 370.00 | 382.00 | 373.00 | 378.00 |
| CDKL1 | 60.00 | 66.00 | 54.00 | 60.00 | 85.00 |
| CDKL2 | 10.00 | 8.00 | 19.00 | 12.33 | 14.00 |
| CDKL3 | 93.00 | 85.00 | 90.00 | 89.33 | 91.00 |
| CDKL4 | 4.00 | 4.00 | 6.00 | 4.67 | 9.00 |
| CDKL5 | 17.00 | 21.00 | 26.00 | 21.33 | 8.00 |
| CDKN1A | 134.00 | 136.00 | 134.00 | 134.67 | 218.00 |
| CDKN1B | 632.00 | 632.00 | 769.00 | 677.67 | 550.00 |
| CDKN2A | 3818.00 | 3799.00 | 4266.00 | 3961.00 | 4614.00 |
| CDKN2AIP | 273.00 | 227.00 | 273.00 | 257.67 | 318.00 |
| CDKN2AIPNL | 462.00 | 449.00 | 532.00 | 481.00 | 702.00 |
| CDKN2B | 527.00 | 451.00 | 624.00 | 534.00 | 382.00 |
| CDKN2B-AS1 | 132.00 | 112.00 | 123.00 | 122.33 | 76.00 |
| CDKN2C | 68.00 | 68.00 | 66.00 | 67.33 | 110.00 |
| CDKN2D | 70.00 | 80.00 | 68.00 | 72.67 | 92.00 |
| CDKN3 | 308.00 | 260.00 | 334.00 | 300.67 | 543.00 |
| CDNF | 53.00 | 46.00 | 61.00 | 53.33 | 72.00 |
| CDO1 | 155.00 | 118.00 | 149.00 | 140.67 | 187.00 |
| CDON | 113.00 | 113.00 | 118.00 | 114.67 | 140.00 |
| CDPF1 | 350.00 | 366.00 | 397.00 | 371.00 | 473.00 |
| CDR2 | 678.00 | 674.00 | 762.00 | 704.67 | 658.00 |
| CDR2L | 436.00 | 454.00 | 424.00 | 438.00 | 423.00 |
| CDRT1 | 6.00 | 4.00 | 5.00 | 5.00 | 4.00 |
| CDS1 | 13.00 | 14.00 | 7.00 | 11.33 | 6.00 |
| CDS2 | 1601.00 | 1581.00 | 1535.00 | 1572.33 | 1337.00 |
| CDT1 | 481.00 | 514.00 | 533.00 | 509.33 | 711.00 |

| | | | | | |
|---|---|---|---|---|---|
| CDV3 | 5425.00 | 5176.00 | 5903.00 | 5501.33 | 6191.00 |
| CDYL | 861.00 | 853.00 | 950.00 | 888.00 | 1004.00 |
| CDYL2 | 11.00 | 17.00 | 18.00 | 15.33 | 17.00 |
| CEACAM19 | 532.00 | 586.00 | 652.00 | 590.00 | 471.00 |
| CEBPB | 1831.00 | 1656.00 | 1868.00 | 1785.00 | 2570.00 |
| CEBPB-AS1 | 49.00 | 34.00 | 53.00 | 45.33 | 57.00 |
| CEBPD | 49.00 | 70.00 | 67.00 | 62.00 | 55.00 |
| CEBPG | 1694.00 | 1524.00 | 1853.00 | 1690.33 | 1939.00 |
| CEBPZ | 1911.00 | 1865.00 | 2088.00 | 1954.67 | 2256.00 |
| CEBPZOS | 914.00 | 833.00 | 973.00 | 906.67 | 1134.00 |
| CECR5 | 757.00 | 856.00 | 880.00 | 831.00 | 1017.00 |
| CECR5-AS1 | 5.00 | 10.00 | 8.00 | 7.67 | 8.00 |
| CECR6 | 22.00 | 20.00 | 19.00 | 20.33 | 20.00 |
| CECR7 | 249.00 | 227.00 | 286.00 | 254.00 | 240.00 |
| CEL | 22.00 | 34.00 | 24.00 | 26.67 | 19.00 |
| CELF1 | 4222.00 | 4048.00 | 4543.00 | 4271.00 | 4292.00 |
| CELF6 | 75.00 | 56.00 | 66.00 | 65.67 | 55.00 |
| CELSR2 | 142.00 | 125.00 | 181.00 | 149.33 | 79.00 |
| CELSR3 | 841.00 | 862.00 | 961.00 | 888.00 | 527.00 |
| CELSR3-AS1 | 5.00 | 5.00 | 9.00 | 6.33 | 3.00 |
| CEMIP | 70.00 | 78.00 | 102.00 | 83.33 | 55.00 |
| CENPA | 269.00 | 297.00 | 312.00 | 292.67 | 338.00 |
| CENPB | 2714.00 | 2753.00 | 2875.00 | 2780.67 | 3050.00 |
| CENPBD1 | 532.00 | 555.00 | 592.00 | 559.67 | 550.00 |
| CENPBD1P1 | 441.00 | 401.00 | 428.00 | 423.33 | 441.00 |
| CENPC | 326.00 | 294.00 | 327.00 | 315.67 | 359.00 |
| CENPE | 915.00 | 881.00 | 932.00 | 909.33 | 1014.00 |
| CENPF | 3938.00 | 3882.00 | 3945.00 | 3921.67 | 4424.00 |
| CENPH | 349.00 | 365.00 | 376.00 | 363.33 | 471.00 |
| CENPI | 116.00 | 102.00 | 105.00 | 107.67 | 158.00 |
| CENPJ | 771.00 | 826.00 | 822.00 | 806.33 | 739.00 |
| CENPK | 453.00 | 345.00 | 406.00 | 401.33 | 538.00 |
| CENPL | 277.00 | 282.00 | 332.00 | 297.00 | 373.00 |
| CENPM | 112.00 | 92.00 | 96.00 | 100.00 | 161.00 |
| CENPN | 762.00 | 753.00 | 761.00 | 758.67 | 1038.00 |
| CENPO | 683.00 | 573.00 | 565.00 | 607.00 | 744.00 |
| CENPP | 114.00 | 93.00 | 80.00 | 95.67 | 138.00 |
| CENPQ | 248.00 | 262.00 | 282.00 | 264.00 | 356.00 |
| CENPT | 1098.00 | 1230.00 | 1148.00 | 1158.67 | 1297.00 |
| CENPU | 418.00 | 388.00 | 413.00 | 406.33 | 728.00 |
| CENPV | 285.00 | 267.00 | 316.00 | 289.33 | 347.00 |
| CENPW | 318.00 | 390.00 | 320.00 | 342.67 | 520.00 |
| CEP104 | 1002.00 | 923.00 | 1056.00 | 993.67 | 943.00 |
| CEP112 | 70.00 | 88.00 | 80.00 | 79.33 | 111.00 |
| CEP120 | 974.00 | 905.00 | 1061.00 | 980.00 | 1108.00 |
| CEP126 | 870.00 | 869.00 | 845.00 | 861.33 | 937.00 |
| CEP128 | 255.00 | 249.00 | 257.00 | 253.67 | 263.00 |
| CEP131 | 727.00 | 785.00 | 745.00 | 752.33 | 763.00 |

| | | | | | |
|---|---|---|---|---|---|
| CEP135 | 505.00 | 502.00 | 592.00 | 533.00 | 588.00 |
| CEP152 | 1640.00 | 1628.00 | 1682.00 | 1650.00 | 1381.00 |
| CEP162 | 317.00 | 308.00 | 309.00 | 311.33 | 415.00 |
| CEP164 | 2485.00 | 2487.00 | 2628.00 | 2533.33 | 2671.00 |
| CEP170 | 15.00 | 14.00 | 13.00 | 14.00 | 17.00 |
| CEP170B | 2087.00 | 1998.00 | 2237.00 | 2107.33 | 1839.00 |
| CEP170P1 | 26.00 | 16.00 | 24.00 | 22.00 | 23.00 |
| CEP19 | 55.00 | 49.00 | 62.00 | 55.33 | 60.00 |
| CEP192 | 924.00 | 902.00 | 1013.00 | 946.33 | 934.00 |
| CEP250 | 3599.00 | 3810.00 | 4105.00 | 3838.00 | 2861.00 |
| CEP290 | 1411.00 | 1369.00 | 1454.00 | 1411.33 | 1440.00 |
| CEP295 | 655.00 | 622.00 | 696.00 | 657.67 | 652.00 |
| CEP350 | 1239.00 | 1073.00 | 1255.00 | 1189.00 | 1078.00 |
| CEP41 | 1129.00 | 1135.00 | 1076.00 | 1113.33 | 1207.00 |
| CEP44 | 449.00 | 454.00 | 508.00 | 470.33 | 486.00 |
| CEP55 | 922.00 | 854.00 | 904.00 | 893.33 | 1334.00 |
| CEP57 | 3329.00 | 3139.00 | 3725.00 | 3397.67 | 3937.00 |
| CEP57L1 | 288.00 | 244.00 | 255.00 | 262.33 | 302.00 |
| CEP63 | 549.00 | 552.00 | 613.00 | 571.33 | 664.00 |
| CEP68 | 477.00 | 421.00 | 410.00 | 436.00 | 430.00 |
| CEP70 | 401.00 | 396.00 | 398.00 | 398.33 | 443.00 |
| CEP72 | 149.00 | 131.00 | 169.00 | 149.67 | 175.00 |
| CEP76 | 197.00 | 169.00 | 189.00 | 185.00 | 215.00 |
| CEP78 | 415.00 | 442.00 | 374.00 | 410.33 | 459.00 |
| CEP83 | 529.00 | 515.00 | 572.00 | 538.67 | 618.00 |
| CEP83-AS1 | 24.00 | 28.00 | 33.00 | 28.33 | 25.00 |
| CEP85 | 849.00 | 842.00 | 908.00 | 866.33 | 710.00 |
| CEP85L | 390.00 | 350.00 | 434.00 | 391.33 | 321.00 |
| CEP89 | 263.00 | 237.00 | 259.00 | 253.00 | 212.00 |
| CEP95 | 986.00 | 966.00 | 1052.00 | 1001.33 | 1011.00 |
| CEP97 | 463.00 | 413.00 | 520.00 | 465.33 | 469.00 |
| CEPT1 | 1209.00 | 1190.00 | 1271.00 | 1223.33 | 1253.00 |
| CERCAM | 3921.00 | 4025.00 | 3927.00 | 3957.67 | 4352.00 |
| CERK | 2100.00 | 1882.00 | 2150.00 | 2044.00 | 1902.00 |
| CERS1 | 28.00 | 19.00 | 17.00 | 21.33 | 31.00 |
| CERS2 | 5465.00 | 5265.00 | 5321.00 | 5350.33 | 5792.00 |
| CERS5 | 1396.00 | 1534.00 | 1511.00 | 1480.33 | 1629.00 |
| CERS6 | 2244.00 | 2121.00 | 2209.00 | 2191.33 | 2582.00 |
| CES2 | 712.00 | 702.00 | 767.00 | 727.00 | 756.00 |
| CETN2 | 532.00 | 455.00 | 482.00 | 489.67 | 497.00 |
| CETN3 | 610.00 | 588.00 | 682.00 | 626.67 | 836.00 |
| CETN4P | 8.00 | 2.00 | 11.00 | 7.00 | 6.00 |
| CFAP20 | 660.00 | 633.00 | 714.00 | 669.00 | 935.00 |
| CFAP36 | 736.00 | 751.00 | 760.00 | 749.00 | 809.00 |
| CFAP43 | 24.00 | 24.00 | 30.00 | 26.00 | 26.00 |
| CFAP44 | 966.00 | 1005.00 | 1008.00 | 993.00 | 914.00 |
| CFAP45 | 307.00 | 316.00 | 281.00 | 301.33 | 250.00 |
| CFAP53 | 19.00 | 22.00 | 24.00 | 21.67 | 15.00 |

| | | | | | |
|---|---|---|---|---|---|
| CFAP54 | 1108.00 | 1092.00 | 1189.00 | 1129.67 | 891.00 |
| CFAP57 | 58.00 | 68.00 | 59.00 | 61.67 | 84.00 |
| CFAP58 | 65.00 | 58.00 | 68.00 | 63.67 | 43.00 |
| CFAP69 | 253.00 | 253.00 | 225.00 | 243.67 | 213.00 |
| CFAP70 | 98.00 | 100.00 | 82.00 | 93.33 | 75.00 |
| CFAP74 | 4.00 | 5.00 | 2.00 | 3.67 | 6.00 |
| CFAP97 | 1237.00 | 1050.00 | 1413.00 | 1233.33 | 1294.00 |
| CFDP1 | 891.00 | 917.00 | 992.00 | 933.33 | 1198.00 |
| CFH | 557.00 | 501.00 | 598.00 | 552.00 | 582.00 |
| CFL1 | 16357.00 | 16038.00 | 16554.00 | 16316.33 | 22692.00 |
| CFL1P1 | 22.00 | 31.00 | 36.00 | 29.67 | 22.00 |
| CFL2 | 1434.00 | 1209.00 | 1437.00 | 1360.00 | 1293.00 |
| CFLAR | 1375.00 | 1406.00 | 1545.00 | 1442.00 | 1422.00 |
| CFLAR-AS1 | 8.00 | 2.00 | 4.00 | 4.67 | 5.00 |
| CFP | 11.00 | 8.00 | 12.00 | 10.33 | 16.00 |
| CGB | 5.00 | 5.00 | 5.00 | 5.00 | 4.00 |
| CGGBP1 | 2449.00 | 2346.00 | 2643.00 | 2479.33 | 2666.00 |
| CGN | 86.00 | 101.00 | 84.00 | 90.33 | 88.00 |
| CGNL1 | 19.00 | 20.00 | 22.00 | 20.33 | 27.00 |
| CGREF1 | 139.00 | 154.00 | 139.00 | 144.00 | 155.00 |
| CGRRF1 | 193.00 | 176.00 | 211.00 | 193.33 | 226.00 |
| CH17-340M24.3 | 19.00 | 26.00 | 18.00 | 21.00 | 26.00 |
| CH25H | 9.00 | 3.00 | 0.00 | 4.00 | 4.00 |
| CHAC1 | 1180.00 | 1157.00 | 1248.00 | 1195.00 | 1399.00 |
| CHAC2 | 104.00 | 122.00 | 112.00 | 112.67 | 132.00 |
| CHAF1A | 651.00 | 783.00 | 726.00 | 720.00 | 844.00 |
| CHAF1B | 178.00 | 221.00 | 212.00 | 203.67 | 336.00 |
| CHAMP1 | 746.00 | 665.00 | 717.00 | 709.33 | 707.00 |
| CHCHD1 | 585.00 | 669.00 | 737.00 | 663.67 | 894.00 |
| CHCHD2 | 4042.00 | 3914.00 | 4236.00 | 4064.00 | 5380.00 |
| CHCHD3 | 1718.00 | 1576.00 | 1794.00 | 1696.00 | 2253.00 |
| CHCHD4 | 421.00 | 389.00 | 427.00 | 412.33 | 574.00 |
| CHCHD5 | 179.00 | 202.00 | 218.00 | 199.67 | 308.00 |
| CHCHD6 | 241.00 | 246.00 | 299.00 | 262.00 | 334.00 |
| CHCHD7 | 603.00 | 675.00 | 657.00 | 645.00 | 737.00 |
| CHD1 | 1922.00 | 1895.00 | 2209.00 | 2008.67 | 1873.00 |
| CHD1L | 2052.00 | 1940.00 | 2164.00 | 2052.00 | 2361.00 |
| CHD2 | 3394.00 | 3336.00 | 3702.00 | 3477.33 | 2762.00 |
| CHD3 | 5676.00 | 5638.00 | 6117.00 | 5810.33 | 6115.00 |
| CHD4 | 7802.00 | 7204.00 | 8068.00 | 7691.33 | 6491.00 |
| CHD5 | 5.00 | 6.00 | 14.00 | 8.33 | 7.00 |
| CHD6 | 1866.00 | 1760.00 | 1959.00 | 1861.67 | 1398.00 |
| CHD7 | 1825.00 | 1799.00 | 1964.00 | 1862.67 | 1774.00 |
| CHD8 | 2616.00 | 2509.00 | 2585.00 | 2570.00 | 2158.00 |
| CHD9 | 1718.00 | 1553.00 | 1840.00 | 1703.67 | 1528.00 |
| CHEK1 | 329.00 | 306.00 | 341.00 | 325.33 | 486.00 |
| CHEK2 | 495.00 | 484.00 | 512.00 | 497.00 | 747.00 |
| CHERP | 1888.00 | 1810.00 | 1967.00 | 1888.33 | 1863.00 |

| | | | | | |
|---|---|---|---|---|---|
| CHFR | 485.00 | 393.00 | 532.00 | 470.00 | 634.00 |
| CHGB | 18.00 | 15.00 | 25.00 | 19.33 | 44.00 |
| CHIC1 | 522.00 | 436.00 | 525.00 | 494.33 | 499.00 |
| CHIC2 | 289.00 | 257.00 | 351.00 | 299.00 | 346.00 |
| CHID1 | 1732.00 | 1899.00 | 1802.00 | 1811.00 | 1948.00 |
| CHKA | 711.00 | 679.00 | 727.00 | 705.67 | 716.00 |
| CHKB | 23.00 | 14.00 | 15.00 | 17.33 | 20.00 |
| CHKB-AS1 | 32.00 | 27.00 | 25.00 | 28.00 | 21.00 |
| CHKB-CPT1B | 21.00 | 21.00 | 29.00 | 23.67 | 22.00 |
| CHL1 | 7.00 | 4.00 | 5.00 | 5.33 | 5.00 |
| CHM | 1019.00 | 902.00 | 1109.00 | 1010.00 | 1070.00 |
| CHML | 903.00 | 850.00 | 1006.00 | 919.67 | 832.00 |
| CHMP1A | 1727.00 | 1574.00 | 1698.00 | 1666.33 | 1936.00 |
| CHMP1B | 1214.00 | 1137.00 | 1286.00 | 1212.33 | 1391.00 |
| CHMP1B2P | 333.00 | 301.00 | 355.00 | 329.67 | 321.00 |
| CHMP2A | 994.00 | 889.00 | 967.00 | 950.00 | 1170.00 |
| CHMP2B | 672.00 | 648.00 | 749.00 | 689.67 | 808.00 |
| CHMP3 | 75.00 | 77.00 | 97.00 | 83.00 | 112.00 |
| CHMP4A | 645.00 | 652.00 | 641.00 | 646.00 | 783.00 |
| CHMP4B | 1848.00 | 1841.00 | 1986.00 | 1891.67 | 1968.00 |
| CHMP5 | 1309.00 | 1318.00 | 1435.00 | 1354.00 | 1869.00 |
| CHMP6 | 262.00 | 273.00 | 234.00 | 256.33 | 240.00 |
| CHMP7 | 798.00 | 730.00 | 820.00 | 782.67 | 998.00 |
| CHN1 | 337.00 | 316.00 | 356.00 | 336.33 | 260.00 |
| CHN2 | 37.00 | 29.00 | 34.00 | 33.33 | 28.00 |
| CHORDC1 | 2924.00 | 2823.00 | 3393.00 | 3046.67 | 3567.00 |
| CHP1 | 1019.00 | 1008.00 | 1143.00 | 1056.67 | 968.00 |
| CHPF | 4802.00 | 4975.00 | 4860.00 | 4879.00 | 3864.00 |
| CHPF2 | 3910.00 | 4008.00 | 4123.00 | 4013.67 | 2988.00 |
| CHPT1 | 843.00 | 847.00 | 852.00 | 847.33 | 1078.00 |
| CHRAC1 | 1192.00 | 1098.00 | 1251.00 | 1180.33 | 1379.00 |
| CHRDL1 | 2161.00 | 2088.00 | 2229.00 | 2159.33 | 2274.00 |
| CHRFAM7A | 3.00 | 5.00 | 4.00 | 4.00 | 3.00 |
| CHRM3 | 30.00 | 15.00 | 33.00 | 26.00 | 21.00 |
| CHRM3-AS2 | 76.00 | 73.00 | 104.00 | 84.33 | 53.00 |
| CHRNA1 | 1261.00 | 1077.00 | 1150.00 | 1162.67 | 1076.00 |
| CHRNA10 | 53.00 | 64.00 | 64.00 | 60.33 | 57.00 |
| CHRNA5 | 157.00 | 129.00 | 156.00 | 147.33 | 202.00 |
| CHRNA9 | 4.00 | 6.00 | 6.00 | 5.33 | 2.00 |
| CHRNB1 | 40.00 | 47.00 | 45.00 | 44.00 | 48.00 |
| CHRNE | 26.00 | 28.00 | 30.00 | 28.00 | 28.00 |
| CHST10 | 849.00 | 889.00 | 902.00 | 880.00 | 1025.00 |
| CHST11 | 2273.00 | 2090.00 | 2149.00 | 2170.67 | 2080.00 |
| CHST12 | 342.00 | 364.00 | 349.00 | 351.67 | 266.00 |
| CHST14 | 1086.00 | 1111.00 | 1060.00 | 1085.67 | 968.00 |
| CHST2 | 673.00 | 754.00 | 664.00 | 697.00 | 603.00 |
| CHST3 | 34.00 | 35.00 | 20.00 | 29.67 | 26.00 |
| CHST6 | 5.00 | 3.00 | 7.00 | 5.00 | 9.00 |

| | | | | | |
|---|---|---|---|---|---|
| CHST7 | 174.00 | 179.00 | 174.00 | 175.67 | 198.00 |
| CHSY1 | 889.00 | 959.00 | 1012.00 | 953.33 | 996.00 |
| CHSY3 | 511.00 | 509.00 | 508.00 | 509.33 | 514.00 |
| CHTF18 | 1371.00 | 1414.00 | 1402.00 | 1395.67 | 1528.00 |
| CHTF8 | 2050.00 | 1993.00 | 2262.00 | 2101.67 | 2238.00 |
| CHTOP | 2363.00 | 2321.00 | 2547.00 | 2410.33 | 2855.00 |
| CHUK | 900.00 | 814.00 | 936.00 | 883.33 | 813.00 |
| CHURC1 | 716.00 | 616.00 | 697.00 | 676.33 | 764.00 |
| CIAO1 | 2554.00 | 2504.00 | 2754.00 | 2604.00 | 2963.00 |
| CIAPIN1 | 1194.00 | 1140.00 | 1175.00 | 1169.67 | 1429.00 |
| CIART | 220.00 | 244.00 | 234.00 | 232.67 | 415.00 |
| CIB1 | 1192.00 | 1203.00 | 1309.00 | 1234.67 | 1343.00 |
| CIC | 1455.00 | 1435.00 | 1530.00 | 1473.33 | 1194.00 |
| CIDEA | 3.00 | 5.00 | 3.00 | 3.67 | 11.00 |
| CIDECP | 137.00 | 117.00 | 111.00 | 121.67 | 155.00 |
| CIITA | 39.00 | 25.00 | 50.00 | 38.00 | 69.00 |
| CILP | 4.00 | 0.00 | 0.00 | 1.33 | 0.00 |
| CILP2 | 3.00 | 12.00 | 8.00 | 7.67 | 11.00 |
| CINP | 285.00 | 243.00 | 281.00 | 269.67 | 334.00 |
| CIPC | 737.00 | 651.00 | 819.00 | 735.67 | 921.00 |
| CIR1 | 497.00 | 505.00 | 612.00 | 538.00 | 601.00 |
| CIRBP | 4546.00 | 4799.00 | 4865.00 | 4736.67 | 5445.00 |
| CIRBP-AS1 | 18.00 | 19.00 | 24.00 | 20.33 | 38.00 |
| CIRH1A | 469.00 | 473.00 | 517.00 | 486.33 | 465.00 |
| CISD1 | 703.00 | 596.00 | 707.00 | 668.67 | 815.00 |
| CISD2 | 400.00 | 390.00 | 376.00 | 388.67 | 475.00 |
| CISH | 9.00 | 5.00 | 2.00 | 5.33 | 9.00 |
| CIT | 1582.00 | 1513.00 | 1634.00 | 1576.33 | 1647.00 |
| CITED1 | 7.00 | 8.00 | 8.00 | 7.67 | 10.00 |
| CITED2 | 1323.00 | 1173.00 | 1353.00 | 1283.00 | 1707.00 |
| CITED4 | 23.00 | 31.00 | 23.00 | 25.67 | 50.00 |
| CIZ1 | 3448.00 | 3498.00 | 3709.00 | 3551.67 | 3396.00 |
| CKAP2 | 1108.00 | 999.00 | 1125.00 | 1077.33 | 1363.00 |
| CKAP2L | 1123.00 | 1064.00 | 1084.00 | 1090.33 | 1259.00 |
| CKAP4 | 10190.00 | 10066.00 | 10052.00 | 10102.67 | 8798.00 |
| CKAP5 | 5080.00 | 4527.00 | 5139.00 | 4915.33 | 4666.00 |
| CKB | 12.00 | 16.00 | 24.00 | 17.33 | 5.00 |
| CKLF | 248.00 | 257.00 | 208.00 | 237.67 | 311.00 |
| CKMT2-AS1 | 282.00 | 315.00 | 315.00 | 304.00 | 331.00 |
| CKS1B | 650.00 | 590.00 | 649.00 | 629.67 | 987.00 |
| CKS2 | 672.00 | 663.00 | 735.00 | 690.00 | 1247.00 |
| CLASP1 | 1743.00 | 1534.00 | 1773.00 | 1683.33 | 1542.00 |
| CLASP2 | 1163.00 | 1024.00 | 1165.00 | 1117.33 | 1097.00 |
| CLASRP | 1186.00 | 1335.00 | 1279.00 | 1266.67 | 1257.00 |
| CLCA3P | 8.00 | 6.00 | 5.00 | 6.33 | 20.00 |
| CLCC1 | 1570.00 | 1447.00 | 1524.00 | 1513.67 | 1818.00 |
| CLCF1 | 331.00 | 324.00 | 332.00 | 329.00 | 173.00 |
| CLCN2 | 305.00 | 346.00 | 297.00 | 316.00 | 289.00 |

| | | | | | |
|---|---|---|---|---|---|
| CLCN3 | 2026.00 | 1898.00 | 2185.00 | 2036.33 | 1889.00 |
| CLCN4 | 468.00 | 472.00 | 496.00 | 478.67 | 331.00 |
| CLCN5 | 591.00 | 533.00 | 610.00 | 578.00 | 571.00 |
| CLCN6 | 1278.00 | 1356.00 | 1406.00 | 1346.67 | 1041.00 |
| CLCN7 | 4665.00 | 4777.00 | 4963.00 | 4801.67 | 4587.00 |
| CLDN1 | 418.00 | 389.00 | 375.00 | 394.00 | 1020.00 |
| CLDN11 | 688.00 | 630.00 | 633.00 | 650.33 | 787.00 |
| CLDN12 | 3584.00 | 3269.00 | 3749.00 | 3534.00 | 3990.00 |
| CLDN14 | 94.00 | 79.00 | 88.00 | 87.00 | 122.00 |
| CLDN15 | 1132.00 | 1206.00 | 1242.00 | 1193.33 | 945.00 |
| CLDN16 | 6.00 | 1.00 | 1.00 | 2.67 | 10.00 |
| CLDN23 | 7.00 | 7.00 | 16.00 | 10.00 | 3.00 |
| CLDN4 | 15.00 | 12.00 | 8.00 | 11.67 | 16.00 |
| CLDN7 | 10.00 | 13.00 | 5.00 | 9.33 | 10.00 |
| CLDN9 | 4.00 | 5.00 | 6.00 | 5.00 | 3.00 |
| CLDND1 | 2422.00 | 2302.00 | 2441.00 | 2388.33 | 2546.00 |
| CLDND2 | 7.00 | 6.00 | 10.00 | 7.67 | 10.00 |
| CLEC11A | 2704.00 | 3164.00 | 3060.00 | 2976.00 | 3763.00 |
| CLEC12B | 16.00 | 4.00 | 12.00 | 10.67 | 12.00 |
| CLEC16A | 741.00 | 756.00 | 819.00 | 772.00 | 702.00 |
| CLEC18A | 5.00 | 11.00 | 7.00 | 7.67 | 0.00 |
| CLEC18B | 3.00 | 6.00 | 3.00 | 4.00 | 2.00 |
| CLEC2B | 274.00 | 272.00 | 280.00 | 275.33 | 324.00 |
| CLEC2D | 902.00 | 909.00 | 992.00 | 934.33 | 805.00 |
| CLEC4A | 14.00 | 11.00 | 8.00 | 11.00 | 23.00 |
| CLEC4GP1 | 14.00 | 15.00 | 9.00 | 12.67 | 39.00 |
| CLEC4M | 3.00 | 5.00 | 0.00 | 2.67 | 4.00 |
| CLEC5A | 5.00 | 2.00 | 2.00 | 3.00 | 1.00 |
| CLGN | 630.00 | 607.00 | 694.00 | 643.67 | 825.00 |
| CLHC1 | 434.00 | 436.00 | 424.00 | 431.33 | 446.00 |
| CLIC1 | 5.00 | 8.00 | 2.00 | 5.00 | 2.00 |
| CLIC2 | 28.00 | 20.00 | 21.00 | 23.00 | 29.00 |
| CLIC4 | 6085.00 | 5068.00 | 6657.00 | 5936.67 | 7056.00 |
| CLINT1 | 2228.00 | 2031.00 | 2335.00 | 2198.00 | 2228.00 |
| CLIP1 | 2442.00 | 2181.00 | 2622.00 | 2415.00 | 2107.00 |
| CLIP2 | 1367.00 | 1285.00 | 1513.00 | 1388.33 | 971.00 |
| CLIP3 | 84.00 | 65.00 | 74.00 | 74.33 | 38.00 |
| CLIP4 | 2494.00 | 2292.00 | 2842.00 | 2542.67 | 2539.00 |
| CLK1 | 2195.00 | 2144.00 | 2558.00 | 2299.00 | 2462.00 |
| CLK2 | 4.00 | 1.00 | 3.00 | 2.67 | 0.00 |
| CLK2P1 | 14.00 | 10.00 | 14.00 | 12.67 | 17.00 |
| CLK3 | 1183.00 | 1193.00 | 1226.00 | 1200.67 | 1211.00 |
| CLK4 | 478.00 | 436.00 | 499.00 | 471.00 | 506.00 |
| CLMN | 1674.00 | 1600.00 | 1831.00 | 1701.67 | 1227.00 |
| CLMP | 2458.00 | 2228.00 | 2457.00 | 2381.00 | 2187.00 |
| CLN3 | 1112.00 | 1151.00 | 1099.00 | 1120.67 | 1100.00 |
| CLN5 | 922.00 | 921.00 | 882.00 | 908.33 | 1000.00 |
| CLN6 | 906.00 | 881.00 | 911.00 | 899.33 | 1001.00 |

| | | | | | |
|---|---|---|---|---|---|
| CLNS1A | 2143.00 | 2025.00 | 2105.00 | 2091.00 | 2692.00 |
| CLOCK | 637.00 | 598.00 | 698.00 | 644.33 | 603.00 |
| CLP1 | 342.00 | 335.00 | 299.00 | 325.33 | 420.00 |
| CLPB | 475.00 | 485.00 | 490.00 | 483.33 | 472.00 |
| CLPP | 664.00 | 647.00 | 710.00 | 673.67 | 897.00 |
| CLPTM1 | 2522.00 | 2557.00 | 2399.00 | 2492.67 | 2410.00 |
| CLPTM1L | 864.00 | 934.00 | 855.00 | 884.33 | 1056.00 |
| CLPX | 748.00 | 737.00 | 816.00 | 767.00 | 858.00 |
| CLSPN | 294.00 | 298.00 | 272.00 | 288.00 | 423.00 |
| CLSTN1 | 6424.00 | 6291.00 | 6532.00 | 6415.67 | 5405.00 |
| CLSTN2 | 3.00 | 4.00 | 22.00 | 9.67 | 8.00 |
| CLSTN3 | 1176.00 | 1241.00 | 1311.00 | 1242.67 | 1058.00 |
| CLTA | 2670.00 | 2663.00 | 2753.00 | 2695.33 | 3430.00 |
| CLTB | 618.00 | 579.00 | 652.00 | 616.33 | 777.00 |
| CLTC | 13452.00 | 11914.00 | 14068.00 | 13144.67 | 13305.00 |
| CLTCL1 | 387.00 | 379.00 | 399.00 | 388.33 | 227.00 |
| CLU | 118.00 | 100.00 | 103.00 | 107.00 | 85.00 |
| CLUAP1 | 519.00 | 508.00 | 555.00 | 527.33 | 652.00 |
| CLUH | 4891.00 | 4863.00 | 5176.00 | 4976.67 | 4952.00 |
| CLUHP3 | 11.00 | 8.00 | 10.00 | 9.67 | 7.00 |
| CLVS2 | 39.00 | 40.00 | 42.00 | 40.33 | 37.00 |
| CLYBL | 57.00 | 59.00 | 57.00 | 57.67 | 58.00 |
| CMAHP | 22.00 | 22.00 | 20.00 | 21.33 | 23.00 |
| CMAS | 1083.00 | 1019.00 | 1169.00 | 1090.33 | 1317.00 |
| CMBL | 70.00 | 83.00 | 62.00 | 71.67 | 113.00 |
| CMC1 | 488.00 | 541.00 | 513.00 | 514.00 | 702.00 |
| CMC2 | 448.00 | 481.00 | 433.00 | 454.00 | 655.00 |
| CMC4 | 95.00 | 90.00 | 103.00 | 96.00 | 133.00 |
| CMIP | 2091.00 | 1967.00 | 2249.00 | 2102.33 | 2520.00 |
| CMPK1 | 1842.00 | 1609.00 | 1940.00 | 1797.00 | 2209.00 |
| CMSS1 | 1577.00 | 1548.00 | 1626.00 | 1583.67 | 2135.00 |
| CMTM1 | 43.00 | 34.00 | 43.00 | 40.00 | 50.00 |
| CMTM3 | 1050.00 | 1074.00 | 1067.00 | 1063.67 | 1010.00 |
| CMTM4 | 955.00 | 1012.00 | 1116.00 | 1027.67 | 870.00 |
| CMTM6 | 2550.00 | 2304.00 | 2308.00 | 2387.33 | 2566.00 |
| CMTM7 | 1500.00 | 1550.00 | 1446.00 | 1498.67 | 1659.00 |
| CMTM8 | 38.00 | 37.00 | 45.00 | 40.00 | 43.00 |
| CMTR1 | 2241.00 | 2134.00 | 2253.00 | 2209.33 | 2016.00 |
| CMTR2 | 1389.00 | 1136.00 | 1347.00 | 1290.67 | 1254.00 |
| CMYA5 | 17.00 | 21.00 | 21.00 | 19.67 | 11.00 |
| CNBP | 7713.00 | 7585.00 | 8434.00 | 7910.67 | 10057.00 |
| CNDP2 | 1904.00 | 1897.00 | 2021.00 | 1940.67 | 2122.00 |
| CNEP1R1 | 370.00 | 336.00 | 399.00 | 368.33 | 444.00 |
| CNGA1 | 45.00 | 39.00 | 47.00 | 43.67 | 23.00 |
| CNGA4 | 25.00 | 22.00 | 19.00 | 22.00 | 18.00 |
| CNIH1 | 2148.00 | 2029.00 | 2039.00 | 2072.00 | 2686.00 |
| CNIH2 | 13.00 | 7.00 | 17.00 | 12.33 | 10.00 |
| CNIH3 | 272.00 | 287.00 | 333.00 | 297.33 | 200.00 |

| | | | | | |
|---|---|---|---|---|---|
| CNIH4 | 1006.00 | 986.00 | 987.00 | 993.00 | 1332.00 |
| CNKSR1 | 62.00 | 82.00 | 62.00 | 68.67 | 51.00 |
| CNKSR2 | 60.00 | 51.00 | 75.00 | 62.00 | 71.00 |
| CNKSR3 | 215.00 | 208.00 | 209.00 | 210.67 | 219.00 |
| CNN1 | 5.00 | 5.00 | 3.00 | 4.33 | 1.00 |
| CNN2 | 828.00 | 802.00 | 890.00 | 840.00 | 851.00 |
| CNN3 | 3916.00 | 3818.00 | 4078.00 | 3937.33 | 5816.00 |
| CNNM1 | 164.00 | 197.00 | 172.00 | 177.67 | 158.00 |
| CNNM2 | 327.00 | 350.00 | 369.00 | 348.67 | 322.00 |
| CNNM3 | 746.00 | 718.00 | 817.00 | 760.33 | 709.00 |
| CNNM4 | 564.00 | 543.00 | 574.00 | 560.33 | 510.00 |
| CNOT1 | 5330.00 | 4640.00 | 5584.00 | 5184.67 | 4724.00 |
| CNOT10 | 779.00 | 688.00 | 771.00 | 746.00 | 930.00 |
| CNOT11 | 1582.00 | 1541.00 | 1602.00 | 1575.00 | 1754.00 |
| CNOT2 | 1427.00 | 1266.00 | 1432.00 | 1375.00 | 1529.00 |
| CNOT3 | 69.00 | 73.00 | 95.00 | 79.00 | 56.00 |
| CNOT4 | 649.00 | 607.00 | 730.00 | 662.00 | 670.00 |
| CNOT6 | 1136.00 | 979.00 | 1252.00 | 1122.33 | 1260.00 |
| CNOT6L | 405.00 | 390.00 | 421.00 | 405.33 | 421.00 |
| CNOT7 | 1087.00 | 1003.00 | 1216.00 | 1102.00 | 1303.00 |
| CNOT8 | 630.00 | 694.00 | 772.00 | 698.67 | 778.00 |
| CNP | 1010.00 | 949.00 | 979.00 | 979.33 | 1137.00 |
| CNPPD1 | 1229.00 | 1142.00 | 1164.00 | 1178.33 | 1470.00 |
| CNPY2 | 2588.00 | 2702.00 | 2633.00 | 2641.00 | 3059.00 |
| CNPY3 | 1307.00 | 1346.00 | 1309.00 | 1320.67 | 1518.00 |
| CNPY4 | 250.00 | 261.00 | 245.00 | 252.00 | 232.00 |
| CNR1 | 14.00 | 22.00 | 23.00 | 19.67 | 5.00 |
| CNRIP1 | 1063.00 | 1040.00 | 1008.00 | 1037.00 | 1404.00 |
| CNST | 1391.00 | 1287.00 | 1399.00 | 1359.00 | 1733.00 |
| CNTF | 4.00 | 0.00 | 3.00 | 2.33 | 2.00 |
| CNTLN | 445.00 | 393.00 | 450.00 | 429.33 | 439.00 |
| CNTN1 | 28.00 | 24.00 | 29.00 | 27.00 | 34.00 |
| CNTN5 | 122.00 | 101.00 | 108.00 | 110.33 | 67.00 |
| CNTNAP1 | 1596.00 | 1656.00 | 1694.00 | 1648.67 | 1707.00 |
| CNTNAP3 | 124.00 | 145.00 | 157.00 | 142.00 | 99.00 |
| CNTNAP3B | 190.00 | 184.00 | 213.00 | 195.67 | 171.00 |
| CNTNAP3P2 | 81.00 | 66.00 | 82.00 | 76.33 | 49.00 |
| CNTRL | 592.00 | 502.00 | 593.00 | 562.33 | 744.00 |
| CNTROB | 1733.00 | 1803.00 | 1855.00 | 1797.00 | 1891.00 |
| COA1 | 2435.00 | 2354.00 | 2449.00 | 2412.67 | 2620.00 |
| COA3 | 218.00 | 224.00 | 211.00 | 217.67 | 367.00 |
| COA4 | 832.00 | 795.00 | 807.00 | 811.33 | 1069.00 |
| COA5 | 613.00 | 572.00 | 700.00 | 628.33 | 745.00 |
| COA6 | 717.00 | 745.00 | 761.00 | 741.00 | 1024.00 |
| COA7 | 593.00 | 583.00 | 697.00 | 624.33 | 677.00 |
| COASY | 2639.00 | 2497.00 | 2547.00 | 2561.00 | 2645.00 |
| COBLL1 | 150.00 | 155.00 | 149.00 | 151.33 | 158.00 |
| COCH | 26.00 | 22.00 | 25.00 | 24.33 | 22.00 |

| | | | | | |
|---|---|---|---|---|---|
| COG1 | 830.00 | 860.00 | 899.00 | 863.00 | 673.00 |
| COG2 | 647.00 | 605.00 | 734.00 | 662.00 | 723.00 |
| COG3 | 941.00 | 913.00 | 1048.00 | 967.33 | 923.00 |
| COG4 | 1634.00 | 1548.00 | 1659.00 | 1613.67 | 1627.00 |
| COG5 | 1527.00 | 1347.00 | 1608.00 | 1494.00 | 1663.00 |
| COG6 | 577.00 | 452.00 | 594.00 | 541.00 | 587.00 |
| COG7 | 483.00 | 500.00 | 580.00 | 521.00 | 576.00 |
| COG8 | 429.00 | 366.00 | 427.00 | 407.33 | 350.00 |
| COIL | 619.00 | 579.00 | 707.00 | 635.00 | 735.00 |
| COL10A1 | 13.00 | 10.00 | 9.00 | 10.67 | 21.00 |
| COL11A1 | 13.00 | 8.00 | 9.00 | 10.00 | 3.00 |
| COL12A1 | 7061.00 | 6429.00 | 7238.00 | 6909.33 | 2674.00 |
| COL13A1 | 7480.00 | 7946.00 | 7291.00 | 7572.33 | 4528.00 |
| COL16A1 | 1025.00 | 1083.00 | 1083.00 | 1063.67 | 1200.00 |
| COL17A1 | 193.00 | 213.00 | 222.00 | 209.33 | 160.00 |
| COL18A1 | 3226.00 | 3247.00 | 3254.00 | 3242.33 | 3163.00 |
| COL19A1 | 26.00 | 34.00 | 34.00 | 31.33 | 25.00 |
| COL1A1 | 22824.00 | 22238.00 | 21503.00 | 22188.33 | 12853.00 |
| COL1A2 | 7.00 | 0.00 | 1.00 | 2.67 | 1.00 |
| COL21A1 | 13.00 | 17.00 | 22.00 | 17.33 | 3.00 |
| COL24A1 | 4.00 | 10.00 | 4.00 | 6.00 | 3.00 |
| COL25A1 | 55.00 | 58.00 | 67.00 | 60.00 | 72.00 |
| COL27A1 | 4131.00 | 4065.00 | 4231.00 | 4142.33 | 4191.00 |
| COL28A1 | 5.00 | 6.00 | 5.00 | 5.33 | 1.00 |
| COL4A1 | 2921.00 | 2918.00 | 2936.00 | 2925.00 | 1968.00 |
| COL4A2 | 5624.00 | 5954.00 | 5831.00 | 5803.00 | 4412.00 |
| COL4A3 | 3.00 | 0.00 | 2.00 | 1.67 | 0.00 |
| COL4A3BP | 515.00 | 525.00 | 593.00 | 544.33 | 501.00 |
| COL4A4 | 13.00 | 6.00 | 10.00 | 9.67 | 8.00 |
| COL4A5 | 2474.00 | 2417.00 | 2729.00 | 2540.00 | 1967.00 |
| COL4A6 | 1605.00 | 1525.00 | 1702.00 | 1610.67 | 1077.00 |
| COL5A1 | 3715.00 | 3847.00 | 4017.00 | 3859.67 | 1592.00 |
| COL5A2 | 1921.00 | 1930.00 | 1920.00 | 1923.67 | 2107.00 |
| COL6A1 | 16283.00 | 17183.00 | 17365.00 | 16943.67 | 18208.00 |
| COL6A2 | 32979.00 | 35177.00 | 34474.00 | 34210.00 | 34640.00 |
| COL6A3 | 15337.00 | 15031.00 | 16065.00 | 15477.67 | 9061.00 |
| COL7A1 | 291.00 | 361.00 | 391.00 | 347.67 | 219.00 |
| COL8A1 | 24.00 | 28.00 | 23.00 | 25.00 | 8.00 |
| COL9A2 | 39.00 | 46.00 | 45.00 | 43.33 | 49.00 |
| COL9A3 | 122.00 | 125.00 | 147.00 | 131.33 | 157.00 |
| COLEC12 | 13442.00 | 13205.00 | 13785.00 | 13477.33 | 14791.00 |
| COLGALT1 | 2672.00 | 2676.00 | 2599.00 | 2649.00 | 2649.00 |
| COLGALT2 | 350.00 | 319.00 | 394.00 | 354.33 | 383.00 |
| COLQ | 44.00 | 30.00 | 46.00 | 40.00 | 27.00 |
| COMMD1 | 184.00 | 198.00 | 201.00 | 194.33 | 213.00 |
| COMMD10 | 319.00 | 337.00 | 380.00 | 345.33 | 404.00 |
| COMMD2 | 616.00 | 495.00 | 640.00 | 583.67 | 669.00 |
| COMMD3 | 490.00 | 487.00 | 550.00 | 509.00 | 439.00 |

| | | | | | |
|---|---|---|---|---|---|
| COMMD4 | 1177.00 | 1182.00 | 1198.00 | 1185.67 | 1583.00 |
| COMMD5 | 446.00 | 485.00 | 483.00 | 471.33 | 509.00 |
| COMMD6 | 933.00 | 985.00 | 1078.00 | 998.67 | 1325.00 |
| COMMD7 | 597.00 | 597.00 | 678.00 | 624.00 | 611.00 |
| COMMD8 | 306.00 | 239.00 | 317.00 | 287.33 | 389.00 |
| COMMD9 | 392.00 | 368.00 | 370.00 | 376.67 | 495.00 |
| COMT | 3953.00 | 3989.00 | 3971.00 | 3971.00 | 4747.00 |
| COMTD1 | 168.00 | 169.00 | 183.00 | 173.33 | 199.00 |
| COPA | 7984.00 | 7374.00 | 8247.00 | 7868.33 | 7137.00 |
| COPB1 | 5079.00 | 4763.00 | 5353.00 | 5065.00 | 5493.00 |
| COPB2 | 4489.00 | 4151.00 | 4768.00 | 4469.33 | 4550.00 |
| COPE | 1835.00 | 1819.00 | 1915.00 | 1856.33 | 2221.00 |
| COPG1 | 4123.00 | 3841.00 | 4350.00 | 4104.67 | 4148.00 |
| COPG2 | 1613.00 | 1476.00 | 1625.00 | 1571.33 | 1782.00 |
| COPRS | 616.00 | 676.00 | 747.00 | 679.67 | 1046.00 |
| COPS2 | 1353.00 | 1225.00 | 1474.00 | 1350.67 | 1642.00 |
| COPS3 | 1338.00 | 1321.00 | 1346.00 | 1335.00 | 1867.00 |
| COPS4 | 543.00 | 496.00 | 534.00 | 524.33 | 591.00 |
| COPS5 | 1486.00 | 1515.00 | 1599.00 | 1533.33 | 1972.00 |
| COPS6 | 2357.00 | 2137.00 | 2432.00 | 2308.67 | 3092.00 |
| COPS7A | 1949.00 | 1819.00 | 2018.00 | 1928.67 | 2126.00 |
| COPS7B | 1025.00 | 1025.00 | 1096.00 | 1048.67 | 1110.00 |
| COPS8 | 1388.00 | 1178.00 | 1329.00 | 1298.33 | 1514.00 |
| COPZ1 | 1968.00 | 1838.00 | 2130.00 | 1978.67 | 2370.00 |
| COPZ2 | 323.00 | 322.00 | 330.00 | 325.00 | 334.00 |
| COQ10A | 85.00 | 96.00 | 121.00 | 100.67 | 123.00 |
| COQ10B | 573.00 | 503.00 | 642.00 | 572.67 | 767.00 |
| COQ2 | 301.00 | 297.00 | 345.00 | 314.33 | 427.00 |
| COQ3 | 142.00 | 132.00 | 139.00 | 137.67 | 236.00 |
| COQ4 | 959.00 | 960.00 | 1001.00 | 973.33 | 1109.00 |
| COQ5 | 513.00 | 565.00 | 574.00 | 550.67 | 717.00 |
| COQ6 | 449.00 | 449.00 | 383.00 | 427.00 | 477.00 |
| COQ7 | 271.00 | 255.00 | 282.00 | 269.33 | 329.00 |
| COQ9 | 1038.00 | 998.00 | 1100.00 | 1045.33 | 1169.00 |
| CORIN | 15.00 | 8.00 | 6.00 | 9.67 | 11.00 |
| CORO1A | 28.00 | 27.00 | 21.00 | 25.33 | 25.00 |
| CORO1B | 2129.00 | 2282.00 | 2276.00 | 2229.00 | 2516.00 |
| CORO1C | 4941.00 | 4657.00 | 5269.00 | 4955.67 | 5536.00 |
| CORO2A | 34.00 | 27.00 | 38.00 | 33.00 | 46.00 |
| CORO2B | 537.00 | 557.00 | 530.00 | 541.33 | 242.00 |
| CORO6 | 20.00 | 23.00 | 25.00 | 22.67 | 18.00 |
| COTL1 | 7010.00 | 6708.00 | 7017.00 | 6911.67 | 10063.00 |
| COX10 | 266.00 | 271.00 | 296.00 | 277.67 | 314.00 |
| COX10-AS1 | 292.00 | 258.00 | 273.00 | 274.33 | 232.00 |
| COX11 | 701.00 | 567.00 | 809.00 | 692.33 | 738.00 |
| COX14 | 166.00 | 210.00 | 185.00 | 187.00 | 254.00 |
| COX15 | 1097.00 | 1071.00 | 1114.00 | 1094.00 | 1276.00 |
| COX16 | 168.00 | 162.00 | 157.00 | 162.33 | 239.00 |

| | | | | | |
|---|---|---|---|---|---|
| COX17 | 303.00 | 340.00 | 357.00 | 333.33 | 494.00 |
| COX18 | 233.00 | 244.00 | 276.00 | 251.00 | 282.00 |
| COX19 | 594.00 | 664.00 | 656.00 | 638.00 | 705.00 |
| COX20 | 416.00 | 402.00 | 439.00 | 419.00 | 480.00 |
| COX4I1 | 5300.00 | 5278.00 | 5573.00 | 5383.67 | 7050.00 |
| COX5A | 1344.00 | 1358.00 | 1357.00 | 1353.00 | 1866.00 |
| COX5B | 1744.00 | 1828.00 | 1804.00 | 1792.00 | 2415.00 |
| COX6A1 | 2601.00 | 2599.00 | 2569.00 | 2589.67 | 1511.00 |
| COX6B1 | 1024.00 | 1029.00 | 1128.00 | 1060.33 | 1429.00 |
| COX6B2 | 3.00 | 1.00 | 0.00 | 1.33 | 1.00 |
| COX6C | 1823.00 | 1830.00 | 1763.00 | 1805.33 | 2499.00 |
| COX7A2 | 1365.00 | 1402.00 | 1459.00 | 1408.67 | 2031.00 |
| COX7A2L | 1796.00 | 1761.00 | 2004.00 | 1853.67 | 2370.00 |
| COX7B | 1312.00 | 1319.00 | 1444.00 | 1358.33 | 1991.00 |
| COX7C | 3485.00 | 3582.00 | 3664.00 | 3577.00 | 4840.00 |
| COX8A | 3262.00 | 3172.00 | 3198.00 | 3210.67 | 4285.00 |
| CPA4 | 25657.00 | 25375.00 | 27258.00 | 26096.67 | 10917.00 |
| CPA5 | 11.00 | 10.00 | 16.00 | 12.33 | 3.00 |
| CPA6 | 4.00 | 7.00 | 3.00 | 4.67 | 4.00 |
| CPB2-AS1 | 10.00 | 9.00 | 8.00 | 9.00 | 7.00 |
| CPD | 3147.00 | 2759.00 | 3233.00 | 3046.33 | 2518.00 |
| CPE | 37.00 | 37.00 | 30.00 | 34.67 | 47.00 |
| CPEB1 | 11.00 | 15.00 | 24.00 | 16.67 | 11.00 |
| CPEB1-AS1 | 11.00 | 3.00 | 6.00 | 6.67 | 6.00 |
| CPEB2 | 319.00 | 258.00 | 382.00 | 319.67 | 296.00 |
| CPEB2-AS1 | 4.00 | 4.00 | 1.00 | 3.00 | 0.00 |
| CPEB3 | 27.00 | 31.00 | 31.00 | 29.67 | 36.00 |
| CPEB4 | 318.00 | 261.00 | 338.00 | 305.67 | 224.00 |
| CPED1 | 1012.00 | 925.00 | 1079.00 | 1005.33 | 1296.00 |
| CPLX3 | 5.00 | 1.00 | 3.00 | 3.00 | 5.00 |
| CPM | 290.00 | 283.00 | 337.00 | 303.33 | 218.00 |
| CPNE1 | 4916.00 | 4676.00 | 5101.00 | 4897.67 | 5527.00 |
| CPNE2 | 302.00 | 315.00 | 307.00 | 308.00 | 407.00 |
| CPNE3 | 2330.00 | 1948.00 | 2398.00 | 2225.33 | 2076.00 |
| CPNE4 | 50.00 | 35.00 | 41.00 | 42.00 | 67.00 |
| CPNE5 | 27.00 | 23.00 | 33.00 | 27.67 | 19.00 |
| CPNE7 | 65.00 | 64.00 | 90.00 | 73.00 | 35.00 |
| CPNE8 | 745.00 | 692.00 | 801.00 | 746.00 | 977.00 |
| CPOX | 940.00 | 1015.00 | 1080.00 | 1011.67 | 1116.00 |
| CPQ | 393.00 | 382.00 | 415.00 | 396.67 | 332.00 |
| CPS1 | 728.00 | 687.00 | 713.00 | 709.33 | 695.00 |
| CPSF1 | 3224.00 | 3389.00 | 3605.00 | 3406.00 | 3530.00 |
| CPSF2 | 2231.00 | 1992.00 | 2374.00 | 2199.00 | 2501.00 |
| CPSF3 | 1115.00 | 1143.00 | 1291.00 | 1183.00 | 1510.00 |
| CPSF3L | 2267.00 | 2374.00 | 2421.00 | 2354.00 | 2486.00 |
| CPSF4 | 985.00 | 1015.00 | 990.00 | 996.67 | 1248.00 |
| CPSF4L | 3.00 | 1.00 | 3.00 | 2.33 | 1.00 |
| CPSF6 | 3148.00 | 2981.00 | 3490.00 | 3206.33 | 3460.00 |

| | | | | | |
|---|---|---|---|---|---|
| CPSF7 | 2677.00 | 2671.00 | 2746.00 | 2698.00 | 2603.00 |
| CPT1A | 43.00 | 36.00 | 36.00 | 38.33 | 61.00 |
| CPT1B | 75.00 | 69.00 | 69.00 | 71.00 | 53.00 |
| CPT1C | 42.00 | 25.00 | 30.00 | 32.33 | 36.00 |
| CPT2 | 453.00 | 443.00 | 461.00 | 452.33 | 596.00 |
| CPTP | 441.00 | 427.00 | 438.00 | 435.33 | 616.00 |
| CPVL | 694.00 | 652.00 | 661.00 | 669.00 | 584.00 |
| CR2 | 16.00 | 12.00 | 14.00 | 14.00 | 12.00 |
| CRABP2 | 5.00 | 7.00 | 5.00 | 5.67 | 22.00 |
| CRACR2A | 503.00 | 396.00 | 507.00 | 468.67 | 282.00 |
| CRADD | 275.00 | 270.00 | 301.00 | 282.00 | 375.00 |
| CRAMP1L | 1540.00 | 1562.00 | 1696.00 | 1599.33 | 1493.00 |
| CRAT | 636.00 | 588.00 | 662.00 | 628.67 | 562.00 |
| CRBN | 571.00 | 586.00 | 648.00 | 601.67 | 651.00 |
| CRCP | 891.00 | 846.00 | 938.00 | 891.67 | 928.00 |
| CREB1 | 1358.00 | 1251.00 | 1471.00 | 1360.00 | 1500.00 |
| CREB3 | 597.00 | 648.00 | 641.00 | 628.67 | 619.00 |
| CREB3L1 | 841.00 | 823.00 | 801.00 | 821.67 | 375.00 |
| CREB3L2 | 8963.00 | 8046.00 | 8816.00 | 8608.33 | 7975.00 |
| CREB3L4 | 412.00 | 431.00 | 401.00 | 414.67 | 476.00 |
| CREB5 | 1079.00 | 1040.00 | 1157.00 | 1092.00 | 844.00 |
| CREBBP | 2158.00 | 1840.00 | 2210.00 | 2069.33 | 1700.00 |
| CREBL2 | 814.00 | 719.00 | 781.00 | 771.33 | 894.00 |
| CREBRF | 577.00 | 506.00 | 759.00 | 614.00 | 491.00 |
| CREBZF | 2439.00 | 2377.00 | 2676.00 | 2497.33 | 2265.00 |
| CREG1 | 317.00 | 311.00 | 299.00 | 309.00 | 394.00 |
| CREG2 | 7.00 | 5.00 | 6.00 | 6.00 | 21.00 |
| CRELD1 | 729.00 | 779.00 | 750.00 | 752.67 | 677.00 |
| CRELD2 | 666.00 | 734.00 | 678.00 | 692.67 | 734.00 |
| CREM | 699.00 | 687.00 | 762.00 | 716.00 | 867.00 |
| CRHBP | 5.00 | 1.00 | 5.00 | 3.67 | 2.00 |
| CRHR1-IT1 | 64.00 | 37.00 | 91.00 | 64.00 | 41.00 |
| CRIM1 | 8369.00 | 7542.00 | 8554.00 | 8155.00 | 5781.00 |
| CRIP1 | 5.00 | 0.00 | 1.00 | 2.00 | 0.00 |
| CRIP2 | 37.00 | 53.00 | 48.00 | 46.00 | 54.00 |
| CRIPAK | 386.00 | 381.00 | 386.00 | 384.33 | 338.00 |
| CRIPT | 261.00 | 239.00 | 307.00 | 269.00 | 306.00 |
| CRISPLD1 | 313.00 | 300.00 | 299.00 | 304.00 | 284.00 |
| CRISPLD2 | 190.00 | 178.00 | 235.00 | 201.00 | 60.00 |
| CRK | 112.00 | 66.00 | 95.00 | 91.00 | 72.00 |
| CRKL | 1595.00 | 1427.00 | 1613.00 | 1545.00 | 1697.00 |
| CRLF3 | 440.00 | 351.00 | 433.00 | 408.00 | 541.00 |
| CRLS1 | 833.00 | 849.00 | 902.00 | 861.33 | 1103.00 |
| CRMP1 | 3.00 | 1.00 | 1.00 | 1.67 | 2.00 |
| CRNDE | 482.00 | 488.00 | 513.00 | 494.33 | 470.00 |
| CRNKL1 | 826.00 | 844.00 | 954.00 | 874.67 | 1133.00 |
| CROCC | 1156.00 | 1282.00 | 1322.00 | 1253.33 | 1075.00 |
| CROCCP2 | 790.00 | 712.00 | 802.00 | 768.00 | 832.00 |

| | | | | | |
|---|---|---|---|---|---|
| CROCCP3 | 317.00 | 353.00 | 345.00 | 338.33 | 306.00 |
| CROT | 340.00 | 329.00 | 333.00 | 334.00 | 295.00 |
| CRSP8P | 54.00 | 55.00 | 59.00 | 56.00 | 47.00 |
| CRTAP | 8962.00 | 8902.00 | 8929.00 | 8931.00 | 8979.00 |
| CRTC1 | 476.00 | 494.00 | 507.00 | 492.33 | 419.00 |
| CRTC2 | 1462.00 | 1492.00 | 1575.00 | 1509.67 | 1435.00 |
| CRTC3 | 818.00 | 707.00 | 890.00 | 805.00 | 758.00 |
| CRTC3-AS1 | 6.00 | 2.00 | 6.00 | 4.67 | 5.00 |
| CRY1 | 835.00 | 795.00 | 838.00 | 822.67 | 879.00 |
| CRY2 | 312.00 | 283.00 | 328.00 | 307.67 | 305.00 |
| CRYBB2P1 | 267.00 | 272.00 | 258.00 | 265.67 | 282.00 |
| CRYBB3 | 9.00 | 8.00 | 3.00 | 6.67 | 4.00 |
| CRYBG3 | 1058.00 | 965.00 | 1086.00 | 1036.33 | 966.00 |
| CRYGS | 86.00 | 134.00 | 99.00 | 106.33 | 109.00 |
| CRYL1 | 471.00 | 379.00 | 421.00 | 423.67 | 597.00 |
| CRYM | 7.00 | 5.00 | 10.00 | 7.33 | 6.00 |
| CRYM-AS1 | 36.00 | 16.00 | 30.00 | 27.33 | 21.00 |
| CRYZ | 625.00 | 604.00 | 715.00 | 648.00 | 753.00 |
| CRYZL1 | 406.00 | 354.00 | 409.00 | 389.67 | 393.00 |
| CS | 3883.00 | 3748.00 | 3949.00 | 3860.00 | 4259.00 |
| CSAD | 402.00 | 430.00 | 473.00 | 435.00 | 597.00 |
| CSAG1 | 22.00 | 37.00 | 41.00 | 33.33 | 43.00 |
| CSAG3 | 7.00 | 11.00 | 7.00 | 8.33 | 17.00 |
| CSDE1 | 11189.00 | 9979.00 | 12109.00 | 11092.33 | 11342.00 |
| CSE1L | 5335.00 | 4973.00 | 5570.00 | 5292.67 | 6462.00 |
| CSF1 | 314.00 | 299.00 | 282.00 | 298.33 | 165.00 |
| CSF1R | 3.00 | 2.00 | 4.00 | 3.00 | 2.00 |
| CSF2 | 5.00 | 7.00 | 1.00 | 4.33 | 1.00 |
| CSF3 | 210.00 | 246.00 | 230.00 | 228.67 | 312.00 |
| CSGALNACT1 | 3.00 | 3.00 | 0.00 | 2.00 | 1.00 |
| CSGALNACT2 | 1399.00 | 1284.00 | 1450.00 | 1377.67 | 1271.00 |
| CSK | 1785.00 | 1838.00 | 1857.00 | 1826.67 | 2227.00 |
| CSMD2 | 17.00 | 11.00 | 12.00 | 13.33 | 13.00 |
| CSMD3 | 26.00 | 26.00 | 32.00 | 28.00 | 74.00 |
| CSNK1A1 | 3923.00 | 3433.00 | 3986.00 | 3780.67 | 4790.00 |
| CSNK1D | 5208.00 | 5232.00 | 5512.00 | 5317.33 | 4930.00 |
| CSNK1E | 143.00 | 122.00 | 144.00 | 136.33 | 115.00 |
| CSNK1G1 | 837.00 | 752.00 | 799.00 | 796.00 | 549.00 |
| CSNK1G2 | 1826.00 | 1778.00 | 1904.00 | 1836.00 | 1936.00 |
| CSNK1G3 | 1121.00 | 967.00 | 1230.00 | 1106.00 | 1357.00 |
| CSNK2A1 | 3509.00 | 3321.00 | 3734.00 | 3521.33 | 3967.00 |
| CSNK2A2 | 986.00 | 875.00 | 1008.00 | 956.33 | 1004.00 |
| CSNK2A3 | 453.00 | 391.00 | 472.00 | 438.67 | 469.00 |
| CSPG4 | 639.00 | 647.00 | 635.00 | 640.33 | 218.00 |
| CSPG5 | 7.00 | 3.00 | 14.00 | 8.00 | 3.00 |
| CSPP1 | 817.00 | 750.00 | 878.00 | 815.00 | 856.00 |
| CSRNP1 | 401.00 | 387.00 | 464.00 | 417.33 | 284.00 |
| CSRNP2 | 634.00 | 602.00 | 787.00 | 674.33 | 814.00 |

| | | | | | |
|---|---|---|---|---|---|
| CSRP1 | 7043.00 | 6707.00 | 6972.00 | 6907.33 | 9970.00 |
| CSRP2 | 73.00 | 65.00 | 74.00 | 70.67 | 117.00 |
| CSRP2BP | 569.00 | 537.00 | 639.00 | 581.67 | 644.00 |
| CST3 | 2053.00 | 2115.00 | 2088.00 | 2085.33 | 2418.00 |
| CSTA | 15.00 | 16.00 | 8.00 | 13.00 | 15.00 |
| CSTB | 1753.00 | 1648.00 | 1721.00 | 1707.33 | 2124.00 |
| CSTF1 | 942.00 | 885.00 | 1058.00 | 961.67 | 1126.00 |
| CSTF2 | 465.00 | 488.00 | 510.00 | 487.67 | 630.00 |
| CSTF2T | 1257.00 | 1177.00 | 1285.00 | 1239.67 | 1461.00 |
| CSTF3 | 1924.00 | 1768.00 | 1884.00 | 1858.67 | 1739.00 |
| CSTF3-AS1 | 10.00 | 11.00 | 11.00 | 10.67 | 16.00 |
| CTAGE1 | 6.00 | 11.00 | 10.00 | 9.00 | 7.00 |
| CTAGE11P | 5.00 | 2.00 | 6.00 | 4.33 | 10.00 |
| CTAGE5 | 1164.00 | 1145.00 | 1159.00 | 1156.00 | 1251.00 |
| CTAGE6 | 4.00 | 0.00 | 0.00 | 1.33 | 2.00 |
| CTB-12O2.1 | 3.00 | 5.00 | 1.00 | 3.00 | 3.00 |
| CTB-178M22.2 | 7.00 | 4.00 | 5.00 | 5.33 | 9.00 |
| CTBP1 | 1874.00 | 1962.00 | 2084.00 | 1973.33 | 2283.00 |
| CTBP1-AS2 | 165.00 | 192.00 | 178.00 | 178.33 | 188.00 |
| CTBP2 | 1490.00 | 1425.00 | 1629.00 | 1514.67 | 1640.00 |
| CTBS | 823.00 | 752.00 | 875.00 | 816.67 | 750.00 |
| CTC1 | 700.00 | 729.00 | 812.00 | 747.00 | 772.00 |
| CTC-338M12.4 | 435.00 | 445.00 | 428.00 | 436.00 | 517.00 |
| CTC-436P18.1 | 7.00 | 2.00 | 7.00 | 5.33 | 3.00 |
| CTCF | 1161.00 | 1027.00 | 1222.00 | 1136.67 | 1216.00 |
| CTDNEP1 | 2791.00 | 2757.00 | 2930.00 | 2826.00 | 3281.00 |
| CTDP1 | 206.00 | 190.00 | 199.00 | 198.33 | 195.00 |
| CTDSP1 | 857.00 | 847.00 | 910.00 | 871.33 | 1018.00 |
| CTDSP2 | 3420.00 | 3178.00 | 3721.00 | 3439.67 | 3163.00 |
| CTDSPL | 319.00 | 299.00 | 312.00 | 310.00 | 424.00 |
| CTDSPL2 | 423.00 | 389.00 | 485.00 | 432.33 | 507.00 |
| CTF1 | 4.00 | 7.00 | 3.00 | 4.67 | 7.00 |
| CTGF | 47683.00 | 45238.00 | 45448.00 | 46123.00 | 74244.00 |
| CTGLF12P | 151.00 | 140.00 | 188.00 | 159.67 | 135.00 |
| CTH | 889.00 | 882.00 | 902.00 | 891.00 | 976.00 |
| CTHRC1 | 1271.00 | 1228.00 | 1212.00 | 1237.00 | 1471.00 |
| CTIF | 1071.00 | 1027.00 | 1196.00 | 1098.00 | 1260.00 |
| CTNNA1 | 5690.00 | 5285.00 | 5818.00 | 5597.67 | 4802.00 |
| CTNNAL1 | 2046.00 | 1859.00 | 2119.00 | 2008.00 | 2526.00 |
| CTNNB1 | 5898.00 | 5764.00 | 6580.00 | 6080.67 | 6418.00 |
| CTNNBIP1 | 319.00 | 292.00 | 280.00 | 297.00 | 385.00 |
| CTNNBL1 | 1070.00 | 1109.00 | 1161.00 | 1113.33 | 1329.00 |
| CTNND1 | 80.00 | 55.00 | 69.00 | 68.00 | 49.00 |
| CTNND2 | 3.00 | 14.00 | 15.00 | 10.67 | 9.00 |
| CTNS | 1159.00 | 1161.00 | 1129.00 | 1149.67 | 1040.00 |
| CTPS1 | 2600.00 | 2465.00 | 2567.00 | 2544.00 | 2499.00 |
| CTPS2 | 1874.00 | 1717.00 | 2025.00 | 1872.00 | 1999.00 |
| CTR9 | 1708.00 | 1578.00 | 1702.00 | 1662.67 | 1580.00 |

| | | | | | |
|---|---|---|---|---|---|
| CTRL | 39.00 | 62.00 | 49.00 | 50.00 | 39.00 |
| CTSA | 7406.00 | 7462.00 | 7013.00 | 7293.67 | 7439.00 |
| CTSB | 31922.00 | 32320.00 | 32152.00 | 32131.33 | 32900.00 |
| CTSC | 2964.00 | 2816.00 | 2973.00 | 2917.67 | 4609.00 |
| CTSD | 9359.00 | 9788.00 | 8909.00 | 9352.00 | 8850.00 |
| CTSF | 910.00 | 975.00 | 909.00 | 931.33 | 764.00 |
| CTSH | 705.00 | 722.00 | 657.00 | 694.67 | 905.00 |
| CTSK | 666.00 | 701.00 | 694.00 | 687.00 | 930.00 |
| CTSL | 4562.00 | 4708.00 | 4332.00 | 4534.00 | 4759.00 |
| CTSS | 80.00 | 79.00 | 85.00 | 81.33 | 110.00 |
| CTSV | 16.00 | 29.00 | 31.00 | 25.33 | 36.00 |
| CTSW | 7.00 | 7.00 | 6.00 | 6.67 | 1.00 |
| CTSZ | 2208.00 | 2168.00 | 2099.00 | 2158.33 | 2005.00 |
| CTTN | 39428.00 | 39815.00 | 42179.00 | 40474.00 | 40044.00 |
| CTTNBP2 | 196.00 | 190.00 | 204.00 | 196.67 | 189.00 |
| CTTNBP2NL | 882.00 | 809.00 | 977.00 | 889.33 | 946.00 |
| CTU1 | 239.00 | 248.00 | 303.00 | 263.33 | 334.00 |
| CTU2 | 985.00 | 883.00 | 951.00 | 939.67 | 1079.00 |
| CTXN1 | 68.00 | 58.00 | 69.00 | 65.00 | 116.00 |
| CUBN | 239.00 | 214.00 | 285.00 | 246.00 | 288.00 |
| CUEDC1 | 1469.00 | 1528.00 | 1609.00 | 1535.33 | 1563.00 |
| CUEDC2 | 1335.00 | 1437.00 | 1517.00 | 1429.67 | 1740.00 |
| CUL1 | 2772.00 | 2621.00 | 3123.00 | 2838.67 | 3127.00 |
| CUL2 | 2464.00 | 2466.00 | 2773.00 | 2567.67 | 2996.00 |
| CUL3 | 1771.00 | 1730.00 | 1946.00 | 1815.67 | 2152.00 |
| CUL4A | 2033.00 | 1811.00 | 2102.00 | 1982.00 | 2135.00 |
| CUL4B | 2266.00 | 2266.00 | 2551.00 | 2361.00 | 2419.00 |
| CUL5 | 597.00 | 496.00 | 641.00 | 578.00 | 637.00 |
| CUL7 | 2309.00 | 2206.00 | 2229.00 | 2248.00 | 2095.00 |
| CUL9 | 2237.00 | 2243.00 | 2333.00 | 2271.00 | 1897.00 |
| CUTA | 4.00 | 2.00 | 5.00 | 3.67 | 3.00 |
| CUTC | 468.00 | 455.00 | 509.00 | 477.33 | 562.00 |
| CUX1 | 1696.00 | 1620.00 | 1850.00 | 1722.00 | 1493.00 |
| CWC15 | 447.00 | 445.00 | 452.00 | 448.00 | 624.00 |
| CWC22 | 1001.00 | 898.00 | 1133.00 | 1010.67 | 1088.00 |
| CWC27 | 626.00 | 618.00 | 738.00 | 660.67 | 873.00 |
| CWF19L1 | 733.00 | 600.00 | 678.00 | 670.33 | 799.00 |
| CWF19L2 | 209.00 | 201.00 | 199.00 | 203.00 | 235.00 |
| CXCL1 | 991.00 | 874.00 | 820.00 | 895.00 | 1973.00 |
| CXCL2 | 653.00 | 659.00 | 670.00 | 660.67 | 1142.00 |
| CXCL3 | 788.00 | 752.00 | 825.00 | 788.33 | 1251.00 |
| CXCL5 | 26210.00 | 23267.00 | 23874.00 | 24450.33 | 45113.00 |
| CXCL8 | 12741.00 | 11589.00 | 10367.00 | 11565.67 | 25570.00 |
| CXorf23 | 634.00 | 608.00 | 657.00 | 633.00 | 572.00 |
| CXorf38 | 410.00 | 360.00 | 405.00 | 391.67 | 388.00 |
| CXorf40A | 175.00 | 180.00 | 152.00 | 169.00 | 212.00 |
| CXorf40B | 338.00 | 328.00 | 333.00 | 333.00 | 368.00 |
| CXorf56 | 257.00 | 190.00 | 239.00 | 228.67 | 305.00 |

| | | | | | |
|---|---|---|---|---|---|
| CXorf57 | 252.00 | 255.00 | 286.00 | 264.33 | 243.00 |
| CXXC1 | 1138.00 | 1116.00 | 1223.00 | 1159.00 | 1342.00 |
| CXXC4 | 27.00 | 24.00 | 33.00 | 28.00 | 23.00 |
| CXXC5 | 6.00 | 9.00 | 10.00 | 8.33 | 6.00 |
| CYB561 | 627.00 | 616.00 | 615.00 | 619.33 | 660.00 |
| CYB561A3 | 1297.00 | 1295.00 | 1306.00 | 1299.33 | 1088.00 |
| CYB561D1 | 654.00 | 584.00 | 674.00 | 637.33 | 477.00 |
| CYB561D2 | 306.00 | 265.00 | 267.00 | 279.33 | 323.00 |
| CYB5A | 114.00 | 123.00 | 126.00 | 121.00 | 191.00 |
| CYB5B | 1092.00 | 1018.00 | 1008.00 | 1039.33 | 1248.00 |
| CYB5D1 | 1300.00 | 1233.00 | 1343.00 | 1292.00 | 1255.00 |
| CYB5D2 | 667.00 | 648.00 | 724.00 | 679.67 | 717.00 |
| CYB5R1 | 745.00 | 705.00 | 673.00 | 707.67 | 803.00 |
| CYB5R2 | 4.00 | 2.00 | 3.00 | 3.00 | 5.00 |
| CYB5R3 | 3058.00 | 2881.00 | 2797.00 | 2912.00 | 2728.00 |
| CYB5R4 | 317.00 | 283.00 | 344.00 | 314.67 | 384.00 |
| CYB5RL | 297.00 | 288.00 | 301.00 | 295.33 | 217.00 |
| CYBA | 2581.00 | 2697.00 | 2338.00 | 2538.67 | 2635.00 |
| CYBRD1 | 2633.00 | 2571.00 | 2603.00 | 2602.33 | 2858.00 |
| CYC1 | 2824.00 | 2763.00 | 2912.00 | 2833.00 | 3522.00 |
| CYCS | 2997.00 | 2719.00 | 3236.00 | 2984.00 | 4342.00 |
| CYFIP1 | 1180.00 | 1082.00 | 1159.00 | 1140.33 | 1639.00 |
| CYFIP2 | 330.00 | 322.00 | 433.00 | 361.67 | 337.00 |
| CYHR1 | 1497.00 | 1490.00 | 1581.00 | 1522.67 | 1427.00 |
| CYLD | 1076.00 | 1037.00 | 1158.00 | 1090.33 | 876.00 |
| CYP11A1 | 22.00 | 31.00 | 26.00 | 26.33 | 19.00 |
| CYP1A1 | 7.00 | 11.00 | 10.00 | 9.33 | 4.00 |
| CYP1B1 | 15.00 | 9.00 | 9.00 | 11.00 | 5.00 |
| CYP20A1 | 953.00 | 910.00 | 938.00 | 933.67 | 1083.00 |
| CYP24A1 | 9.00 | 14.00 | 14.00 | 12.33 | 23.00 |
| CYP27A1 | 298.00 | 331.00 | 307.00 | 312.00 | 303.00 |
| CYP27B1 | 267.00 | 293.00 | 279.00 | 279.67 | 280.00 |
| CYP2E1 | 3.00 | 3.00 | 7.00 | 4.33 | 7.00 |
| CYP2J2 | 189.00 | 205.00 | 198.00 | 197.33 | 284.00 |
| CYP2R1 | 518.00 | 534.00 | 581.00 | 544.33 | 587.00 |
| CYP2S1 | 12.00 | 11.00 | 12.00 | 11.67 | 3.00 |
| CYP2U1 | 514.00 | 500.00 | 523.00 | 512.33 | 473.00 |
| CYP39A1 | 28.00 | 26.00 | 25.00 | 26.33 | 23.00 |
| CYP4F29P | 62.00 | 51.00 | 59.00 | 57.33 | 112.00 |
| CYP4V2 | 508.00 | 501.00 | 517.00 | 508.67 | 581.00 |
| CYP51A1 | 1333.00 | 1285.00 | 1275.00 | 1297.67 | 1550.00 |
| CYR61 | 53773.00 | 55920.00 | 54736.00 | 54809.67 | 59644.00 |
| CYSRT1 | 16.00 | 17.00 | 16.00 | 16.33 | 22.00 |
| CYSTM1 | 365.00 | 317.00 | 372.00 | 351.33 | 437.00 |
| CYTH1 | 411.00 | 431.00 | 430.00 | 424.00 | 427.00 |
| CYTH2 | 2895.00 | 3117.00 | 3285.00 | 3099.00 | 2518.00 |
| CYTH3 | 939.00 | 857.00 | 938.00 | 911.33 | 802.00 |
| CYTH4 | 7.00 | 10.00 | 6.00 | 7.67 | 12.00 |

| | | | | | |
|---|---|---|---|---|---|
| D2HGDH | 742.00 | 814.00 | 808.00 | 788.00 | 875.00 |
| DAAM1 | 197.00 | 174.00 | 211.00 | 194.00 | 147.00 |
| DAAM2 | 3.00 | 5.00 | 5.00 | 4.33 | 1.00 |
| DAB2 | 1578.00 | 1439.00 | 1580.00 | 1532.33 | 855.00 |
| DAB2IP | 542.00 | 517.00 | 565.00 | 541.33 | 536.00 |
| DACH1 | 36.00 | 24.00 | 44.00 | 34.67 | 41.00 |
| DACH2 | 54.00 | 44.00 | 73.00 | 57.00 | 69.00 |
| DACT1 | 16.00 | 43.00 | 21.00 | 26.67 | 10.00 |
| DACT3 | 3.00 | 2.00 | 3.00 | 2.67 | 3.00 |
| DAD1 | 2601.00 | 2576.00 | 2357.00 | 2511.33 | 3109.00 |
| DAG1 | 1697.00 | 1625.00 | 1791.00 | 1704.33 | 1555.00 |
| DAGLA | 630.00 | 683.00 | 675.00 | 662.67 | 489.00 |
| DAGLB | 1268.00 | 1337.00 | 1403.00 | 1336.00 | 1120.00 |
| DALRD3 | 585.00 | 532.00 | 610.00 | 575.67 | 627.00 |
| DANCR | 973.00 | 995.00 | 1033.00 | 1000.33 | 1129.00 |
| DAP | 3745.00 | 3473.00 | 3738.00 | 3652.00 | 3884.00 |
| DAP3 | 2850.00 | 2727.00 | 2858.00 | 2811.67 | 3653.00 |
| DAPK2 | 13.00 | 8.00 | 7.00 | 9.33 | 13.00 |
| DAPK3 | 1308.00 | 1402.00 | 1413.00 | 1374.33 | 1395.00 |
| DARS | 2319.00 | 2102.00 | 2529.00 | 2316.67 | 2689.00 |
| DARS2 | 2231.00 | 2012.00 | 2279.00 | 2174.00 | 2849.00 |
| DARS-AS1 | 37.00 | 40.00 | 38.00 | 38.33 | 36.00 |
| DAW1 | 765.00 | 670.00 | 704.00 | 713.00 | 1641.00 |
| DAXX | 23.00 | 27.00 | 16.00 | 22.00 | 32.00 |
| DAZAP1 | 4533.00 | 4403.00 | 4713.00 | 4549.67 | 5794.00 |
| DAZAP2 | 1751.00 | 1536.00 | 1917.00 | 1734.67 | 2070.00 |
| DBF4 | 627.00 | 610.00 | 730.00 | 655.67 | 947.00 |
| DBF4B | 801.00 | 870.00 | 860.00 | 843.67 | 889.00 |
| DBH-AS1 | 63.00 | 64.00 | 53.00 | 60.00 | 18.00 |
| DBI | 1012.00 | 925.00 | 956.00 | 964.33 | 1433.00 |
| DBIL5P | 9.00 | 12.00 | 6.00 | 9.00 | 11.00 |
| DBN1 | 6600.00 | 6480.00 | 6555.00 | 6545.00 | 5510.00 |
| DBNDD1 | 110.00 | 119.00 | 128.00 | 119.00 | 155.00 |
| DBNDD2 | 74.00 | 50.00 | 55.00 | 59.67 | 64.00 |
| DBNL | 1021.00 | 1002.00 | 954.00 | 992.33 | 1276.00 |
| DBP | 17.00 | 17.00 | 28.00 | 20.67 | 34.00 |
| DBR1 | 338.00 | 353.00 | 332.00 | 341.00 | 410.00 |
| DBT | 577.00 | 546.00 | 652.00 | 591.67 | 593.00 |
| DCAF10 | 1037.00 | 1076.00 | 1251.00 | 1121.33 | 1089.00 |
| DCAF11 | 1044.00 | 995.00 | 1120.00 | 1053.00 | 1131.00 |
| DCAF12 | 658.00 | 622.00 | 674.00 | 651.33 | 787.00 |
| DCAF13 | 2435.00 | 2179.00 | 2414.00 | 2342.67 | 3062.00 |
| DCAF13P3 | 6.00 | 2.00 | 3.00 | 3.67 | 3.00 |
| DCAF15 | 903.00 | 950.00 | 944.00 | 932.33 | 1109.00 |
| DCAF16 | 906.00 | 936.00 | 1041.00 | 961.00 | 953.00 |
| DCAF17 | 1056.00 | 888.00 | 1177.00 | 1040.33 | 933.00 |
| DCAF4 | 559.00 | 549.00 | 635.00 | 581.00 | 655.00 |
| DCAF4L1 | 120.00 | 123.00 | 124.00 | 122.33 | 119.00 |

| | | | | | |
|---|---|---|---|---|---|
| DCAF5 | 1595.00 | 1543.00 | 1614.00 | 1584.00 | 1475.00 |
| DCAF6 | 925.00 | 923.00 | 1005.00 | 951.00 | 1213.00 |
| DCAF7 | 3358.00 | 3015.00 | 3408.00 | 3260.33 | 3163.00 |
| DCAF8 | 4879.00 | 4882.00 | 5166.00 | 4975.67 | 4636.00 |
| DCAF8L2 | 15.00 | 12.00 | 7.00 | 11.33 | 21.00 |
| DCAKD | 694.00 | 613.00 | 694.00 | 667.00 | 704.00 |
| DCBLD1 | 864.00 | 866.00 | 847.00 | 859.00 | 789.00 |
| DCBLD2 | 24445.00 | 22359.00 | 25005.00 | 23936.33 | 25903.00 |
| DCC | 183.00 | 181.00 | 214.00 | 192.67 | 154.00 |
| DCDC2 | 10.00 | 12.00 | 12.00 | 11.33 | 12.00 |
| DCHS1 | 906.00 | 855.00 | 796.00 | 852.33 | 684.00 |
| DCHS2 | 12.00 | 15.00 | 19.00 | 15.33 | 24.00 |
| DCK | 303.00 | 287.00 | 334.00 | 308.00 | 409.00 |
| DCLK1 | 4.00 | 2.00 | 4.00 | 3.33 | 5.00 |
| DCLK2 | 169.00 | 168.00 | 180.00 | 172.33 | 233.00 |
| DCLRE1A | 606.00 | 584.00 | 626.00 | 605.33 | 796.00 |
| DCLRE1B | 354.00 | 358.00 | 368.00 | 360.00 | 431.00 |
| DCLRE1C | 645.00 | 559.00 | 650.00 | 618.00 | 715.00 |
| DCP1A | 927.00 | 796.00 | 886.00 | 869.67 | 839.00 |
| DCP1B | 337.00 | 330.00 | 383.00 | 350.00 | 336.00 |
| DCP2 | 741.00 | 633.00 | 809.00 | 727.67 | 744.00 |
| DCPS | 109.00 | 112.00 | 99.00 | 106.67 | 143.00 |
| DCST2 | 15.00 | 14.00 | 17.00 | 15.33 | 19.00 |
| DCTD | 1848.00 | 1692.00 | 2054.00 | 1864.67 | 2201.00 |
| DCTN1 | 4958.00 | 4809.00 | 5075.00 | 4947.33 | 4866.00 |
| DCTN2 | 2567.00 | 2411.00 | 2656.00 | 2544.67 | 2775.00 |
| DCTN3 | 484.00 | 517.00 | 483.00 | 494.67 | 675.00 |
| DCTN4 | 1625.00 | 1343.00 | 1729.00 | 1565.67 | 1759.00 |
| DCTN5 | 1276.00 | 1189.00 | 1295.00 | 1253.33 | 1164.00 |
| DCTN6 | 257.00 | 259.00 | 282.00 | 266.00 | 339.00 |
| DCTPP1 | 381.00 | 397.00 | 398.00 | 392.00 | 434.00 |
| DCUN1D1 | 651.00 | 636.00 | 834.00 | 707.00 | 837.00 |
| DCUN1D2 | 355.00 | 335.00 | 386.00 | 358.67 | 352.00 |
| DCUN1D3 | 226.00 | 224.00 | 230.00 | 226.67 | 232.00 |
| DCUN1D4 | 1020.00 | 866.00 | 1050.00 | 978.67 | 1034.00 |
| DCUN1D5 | 3410.00 | 3192.00 | 3493.00 | 3365.00 | 4553.00 |
| DCXR | 768.00 | 819.00 | 978.00 | 855.00 | 1025.00 |
| DDA1 | 887.00 | 823.00 | 861.00 | 857.00 | 810.00 |
| DDAH1 | 3152.00 | 2910.00 | 3579.00 | 3213.67 | 4492.00 |
| DDB1 | 5251.00 | 5016.00 | 5512.00 | 5259.67 | 4970.00 |
| DDB2 | 243.00 | 235.00 | 236.00 | 238.00 | 330.00 |
| DDHD1 | 1519.00 | 1431.00 | 1630.00 | 1526.67 | 1345.00 |
| DDHD2 | 1154.00 | 1152.00 | 1342.00 | 1216.00 | 1297.00 |
| DDI2 | 74.00 | 58.00 | 62.00 | 64.67 | 51.00 |
| DDIAS | 267.00 | 234.00 | 230.00 | 243.67 | 297.00 |
| DDIT3 | 1292.00 | 1333.00 | 1333.00 | 1319.33 | 1428.00 |
| DDIT4 | 1597.00 | 1612.00 | 1590.00 | 1599.67 | 1889.00 |
| DDN | 18.00 | 16.00 | 22.00 | 18.67 | 43.00 |

| | | | | | |
|---|---|---|---|---|---|
| DDO | 10.00 | 19.00 | 15.00 | 14.67 | 16.00 |
| DDOST | 11752.00 | 11703.00 | 11430.00 | 11628.33 | 11922.00 |
| DDR1 | 50.00 | 39.00 | 58.00 | 49.00 | 36.00 |
| DDR2 | 3311.00 | 2819.00 | 3153.00 | 3094.33 | 2523.00 |
| DDRGK1 | 1048.00 | 1105.00 | 993.00 | 1048.67 | 1275.00 |
| DDX1 | 3294.00 | 3090.00 | 3217.00 | 3200.33 | 4020.00 |
| DDX10 | 1426.00 | 1373.00 | 1565.00 | 1454.67 | 1130.00 |
| DDX11 | 1845.00 | 1899.00 | 1873.00 | 1872.33 | 2154.00 |
| DDX11-AS1 | 58.00 | 71.00 | 65.00 | 64.67 | 81.00 |
| DDX11L2 | 74.00 | 84.00 | 90.00 | 82.67 | 96.00 |
| DDX11L9 | 3.00 | 1.00 | 0.00 | 1.33 | 0.00 |
| DDX12P | 969.00 | 1004.00 | 990.00 | 987.67 | 999.00 |
| DDX17 | 13164.00 | 12372.00 | 13337.00 | 12957.67 | 13328.00 |
| DDX18 | 2984.00 | 2681.00 | 3043.00 | 2902.67 | 3515.00 |
| DDX19A | 775.00 | 803.00 | 872.00 | 816.67 | 794.00 |
| DDX19B | 359.00 | 380.00 | 436.00 | 391.67 | 573.00 |
| DDX20 | 853.00 | 805.00 | 912.00 | 856.67 | 969.00 |
| DDX21 | 8965.00 | 7633.00 | 9155.00 | 8584.33 | 9233.00 |
| DDX23 | 2009.00 | 1978.00 | 2084.00 | 2023.67 | 2124.00 |
| DDX24 | 3.00 | 1.00 | 1.00 | 1.67 | 3.00 |
| DDX26B | 277.00 | 261.00 | 306.00 | 281.33 | 271.00 |
| DDX27 | 2298.00 | 2268.00 | 2334.00 | 2300.00 | 2486.00 |
| DDX28 | 299.00 | 292.00 | 339.00 | 310.00 | 378.00 |
| DDX31 | 1491.00 | 1443.00 | 1602.00 | 1512.00 | 1522.00 |
| DDX39A | 1790.00 | 1749.00 | 1847.00 | 1795.33 | 2377.00 |
| DDX39B | 3.00 | 4.00 | 3.00 | 3.33 | 2.00 |
| DDX3X | 9379.00 | 8532.00 | 9659.00 | 9190.00 | 9485.00 |
| DDX4 | 4.00 | 1.00 | 4.00 | 3.00 | 4.00 |
| DDX41 | 1602.00 | 1656.00 | 1705.00 | 1654.33 | 1713.00 |
| DDX42 | 4938.00 | 4841.00 | 5122.00 | 4967.00 | 5316.00 |
| DDX46 | 2390.00 | 2140.00 | 2553.00 | 2361.00 | 2619.00 |
| DDX47 | 1361.00 | 1303.00 | 1420.00 | 1361.33 | 1554.00 |
| DDX49 | 725.00 | 683.00 | 742.00 | 716.67 | 998.00 |
| DDX5 | 27591.00 | 26984.00 | 28461.00 | 27678.67 | 30919.00 |
| DDX50 | 1401.00 | 1320.00 | 1570.00 | 1430.33 | 1885.00 |
| DDX51 | 902.00 | 952.00 | 1006.00 | 953.33 | 938.00 |
| DDX52 | 112.00 | 102.00 | 126.00 | 113.33 | 119.00 |
| DDX53 | 20.00 | 20.00 | 9.00 | 16.33 | 19.00 |
| DDX54 | 2256.00 | 2257.00 | 2389.00 | 2300.67 | 2306.00 |
| DDX55 | 1247.00 | 1295.00 | 1235.00 | 1259.00 | 1503.00 |
| DDX56 | 3005.00 | 3143.00 | 3195.00 | 3114.33 | 3413.00 |
| DDX58 | 552.00 | 577.00 | 592.00 | 573.67 | 696.00 |
| DDX59 | 449.00 | 394.00 | 497.00 | 446.67 | 514.00 |
| DDX6 | 2383.00 | 2217.00 | 2637.00 | 2412.33 | 2398.00 |
| DDX60 | 114.00 | 91.00 | 117.00 | 107.33 | 128.00 |
| DDX60L | 169.00 | 160.00 | 190.00 | 173.00 | 151.00 |
| DEAF1 | 477.00 | 475.00 | 460.00 | 470.67 | 499.00 |
| DECR1 | 1215.00 | 1166.00 | 1231.00 | 1204.00 | 1566.00 |

| | | | | | |
|---|---|---|---|---|---|
| DEDD | 848.00 | 769.00 | 868.00 | 828.33 | 928.00 |
| DEDD2 | 307.00 | 258.00 | 301.00 | 288.67 | 310.00 |
| DEF8 | 2529.00 | 2605.00 | 2839.00 | 2657.67 | 2672.00 |
| DEGS1 | 2675.00 | 2551.00 | 2619.00 | 2615.00 | 2635.00 |
| DEK | 2760.00 | 2394.00 | 2893.00 | 2682.33 | 3540.00 |
| DENND1A | 445.00 | 369.00 | 490.00 | 434.67 | 514.00 |
| DENND1B | 147.00 | 138.00 | 166.00 | 150.33 | 155.00 |
| DENND1C | 3.00 | 0.00 | 1.00 | 1.33 | 1.00 |
| DENND2A | 716.00 | 670.00 | 654.00 | 680.00 | 523.00 |
| DENND2C | 49.00 | 31.00 | 47.00 | 42.33 | 39.00 |
| DENND3 | 1222.00 | 1148.00 | 1250.00 | 1206.67 | 1374.00 |
| DENND4A | 643.00 | 614.00 | 799.00 | 685.33 | 665.00 |
| DENND4B | 2963.00 | 2710.00 | 2956.00 | 2876.33 | 2461.00 |
| DENND4C | 810.00 | 726.00 | 825.00 | 787.00 | 704.00 |
| DENND5A | 1822.00 | 1835.00 | 2014.00 | 1890.33 | 1619.00 |
| DENND5B | 721.00 | 667.00 | 808.00 | 732.00 | 607.00 |
| DENND6A | 498.00 | 465.00 | 585.00 | 516.00 | 533.00 |
| DENND6B | 19.00 | 22.00 | 24.00 | 21.67 | 22.00 |
| DENR | 2697.00 | 2456.00 | 2792.00 | 2648.33 | 3362.00 |
| DEPDC1 | 1465.00 | 1256.00 | 1367.00 | 1362.67 | 1706.00 |
| DEPDC1-AS1 | 5.00 | 2.00 | 3.00 | 3.33 | 6.00 |
| DEPDC1B | 272.00 | 214.00 | 282.00 | 256.00 | 335.00 |
| DEPDC4 | 16.00 | 22.00 | 17.00 | 18.33 | 29.00 |
| DEPDC5 | 317.00 | 343.00 | 355.00 | 338.33 | 368.00 |
| DEPDC7 | 177.00 | 147.00 | 139.00 | 154.33 | 206.00 |
| DEPTOR | 19.00 | 38.00 | 38.00 | 31.67 | 43.00 |
| DERA | 530.00 | 482.00 | 535.00 | 515.67 | 737.00 |
| DERL1 | 2745.00 | 2574.00 | 2658.00 | 2659.00 | 2574.00 |
| DERL2 | 1628.00 | 1533.00 | 1609.00 | 1590.00 | 1736.00 |
| DESI1 | 1845.00 | 1776.00 | 1957.00 | 1859.33 | 1819.00 |
| DESI2 | 1444.00 | 1277.00 | 1562.00 | 1427.67 | 1848.00 |
| DET1 | 237.00 | 211.00 | 257.00 | 235.00 | 244.00 |
| DEXI | 543.00 | 486.00 | 519.00 | 516.00 | 630.00 |
| DFFA | 972.00 | 874.00 | 924.00 | 923.33 | 1035.00 |
| DFFB | 364.00 | 361.00 | 386.00 | 370.33 | 397.00 |
| DFNA5 | 2010.00 | 1752.00 | 2145.00 | 1969.00 | 1900.00 |
| DFNB31 | 77.00 | 98.00 | 94.00 | 89.67 | 82.00 |
| DFNB59 | 147.00 | 155.00 | 164.00 | 155.33 | 118.00 |
| DGAT1 | 856.00 | 904.00 | 954.00 | 904.67 | 910.00 |
| DGAT2 | 2445.00 | 2564.00 | 2441.00 | 2483.33 | 2794.00 |
| DGCR11 | 17.00 | 31.00 | 23.00 | 23.67 | 25.00 |
| DGCR14 | 390.00 | 389.00 | 446.00 | 408.33 | 391.00 |
| DGCR2 | 1960.00 | 1946.00 | 1956.00 | 1954.00 | 1896.00 |
| DGCR6L | 361.00 | 365.00 | 373.00 | 366.33 | 470.00 |
| DGCR8 | 1354.00 | 1348.00 | 1527.00 | 1409.67 | 1463.00 |
| DGKA | 419.00 | 414.00 | 509.00 | 447.33 | 409.00 |
| DGKD | 1490.00 | 1480.00 | 1460.00 | 1476.67 | 1115.00 |
| DGKE | 151.00 | 134.00 | 153.00 | 146.00 | 131.00 |

| | | | | | |
|---|---|---|---|---|---|
| DGKG | 105.00 | 128.00 | 104.00 | 112.33 | 71.00 |
| DGKH | 916.00 | 835.00 | 997.00 | 916.00 | 684.00 |
| DGKI | 59.00 | 71.00 | 46.00 | 58.67 | 58.00 |
| DGKQ | 602.00 | 595.00 | 622.00 | 606.33 | 563.00 |
| DGKZ | 1046.00 | 1026.00 | 1085.00 | 1052.33 | 1160.00 |
| DGUOK | 1091.00 | 1042.00 | 1092.00 | 1075.00 | 1494.00 |
| DHCR24 | 3201.00 | 2924.00 | 2889.00 | 3004.67 | 2460.00 |
| DHCR7 | 16037.00 | 15728.00 | 15874.00 | 15879.67 | 15222.00 |
| DHDDS | 689.00 | 682.00 | 747.00 | 706.00 | 678.00 |
| DHDH | 3.00 | 1.00 | 8.00 | 4.00 | 3.00 |
| DHFR | 248.00 | 251.00 | 243.00 | 247.33 | 312.00 |
| DHFRL1 | 287.00 | 252.00 | 260.00 | 266.33 | 342.00 |
| DHODH | 408.00 | 408.00 | 422.00 | 412.67 | 537.00 |
| DHPS | 1370.00 | 1290.00 | 1417.00 | 1359.00 | 1493.00 |
| DHRS1 | 360.00 | 329.00 | 349.00 | 346.00 | 375.00 |
| DHRS12 | 54.00 | 56.00 | 62.00 | 57.33 | 95.00 |
| DHRS13 | 77.00 | 94.00 | 90.00 | 87.00 | 145.00 |
| DHRS2 | 3.00 | 2.00 | 2.00 | 2.33 | 1.00 |
| DHRS3 | 185.00 | 207.00 | 211.00 | 201.00 | 199.00 |
| DHRS4 | 155.00 | 142.00 | 132.00 | 143.00 | 186.00 |
| DHRS4-AS1 | 248.00 | 224.00 | 265.00 | 245.67 | 277.00 |
| DHRS4L2 | 68.00 | 72.00 | 85.00 | 75.00 | 112.00 |
| DHRS7 | 1490.00 | 1587.00 | 1584.00 | 1553.67 | 2049.00 |
| DHRS7B | 326.00 | 358.00 | 312.00 | 332.00 | 427.00 |
| DHTKD1 | 634.00 | 545.00 | 676.00 | 618.33 | 553.00 |
| DHX15 | 3272.00 | 3209.00 | 3423.00 | 3301.33 | 3776.00 |
| DHX29 | 1095.00 | 1082.00 | 1277.00 | 1151.33 | 1118.00 |
| DHX30 | 2618.00 | 2694.00 | 2733.00 | 2681.67 | 2886.00 |
| DHX32 | 1167.00 | 1004.00 | 1195.00 | 1122.00 | 1296.00 |
| DHX33 | 2729.00 | 2572.00 | 2909.00 | 2736.67 | 2795.00 |
| DHX34 | 1543.00 | 1516.00 | 1548.00 | 1535.67 | 1365.00 |
| DHX35 | 643.00 | 679.00 | 732.00 | 684.67 | 695.00 |
| DHX36 | 1174.00 | 1078.00 | 1328.00 | 1193.33 | 1291.00 |
| DHX37 | 1207.00 | 1301.00 | 1295.00 | 1267.67 | 1276.00 |
| DHX38 | 2209.00 | 2391.00 | 2247.00 | 2282.33 | 2207.00 |
| DHX40 | 1289.00 | 1198.00 | 1375.00 | 1287.33 | 1426.00 |
| DHX57 | 1023.00 | 964.00 | 1034.00 | 1007.00 | 1110.00 |
| DHX58 | 42.00 | 47.00 | 43.00 | 44.00 | 45.00 |
| DHX8 | 991.00 | 997.00 | 993.00 | 993.67 | 879.00 |
| DHX9 | 5110.00 | 4918.00 | 5388.00 | 5138.67 | 5998.00 |
| DIABLO | 707.00 | 696.00 | 701.00 | 701.33 | 840.00 |
| DIAPH1 | 3367.00 | 3109.00 | 3344.00 | 3273.33 | 3302.00 |
| DIAPH2 | 413.00 | 333.00 | 360.00 | 368.67 | 434.00 |
| DIAPH2-AS1 | 6.00 | 4.00 | 4.00 | 4.67 | 0.00 |
| DIAPH3 | 371.00 | 312.00 | 339.00 | 340.67 | 374.00 |
| DICER1 | 1991.00 | 1734.00 | 2039.00 | 1921.33 | 1825.00 |
| DICER1-AS1 | 175.00 | 183.00 | 157.00 | 171.67 | 158.00 |
| DIDO1 | 4247.00 | 4114.00 | 4439.00 | 4266.67 | 3696.00 |

| | | | | | |
|---|---|---|---|---|---|
| DIEXF | 1288.00 | 1200.00 | 1339.00 | 1275.67 | 1334.00 |
| DIMT1 | 506.00 | 464.00 | 521.00 | 497.00 | 641.00 |
| DIO2 | 17.00 | 17.00 | 15.00 | 16.33 | 12.00 |
| DIP2A | 1779.00 | 1826.00 | 2003.00 | 1869.33 | 1496.00 |
| DIP2A-IT1 | 10.00 | 4.00 | 1.00 | 5.00 | 1.00 |
| DIP2B | 1143.00 | 1005.00 | 1168.00 | 1105.33 | 961.00 |
| DIP2C | 499.00 | 584.00 | 578.00 | 553.67 | 518.00 |
| DIRAS3 | 7.00 | 10.00 | 17.00 | 11.33 | 7.00 |
| DIRC2 | 742.00 | 726.00 | 664.00 | 710.67 | 792.00 |
| DIRC3 | 3.00 | 0.00 | 0.00 | 1.00 | 1.00 |
| DIS3 | 1598.00 | 1445.00 | 1737.00 | 1593.33 | 1770.00 |
| DIS3L | 1191.00 | 1144.00 | 1243.00 | 1192.67 | 1241.00 |
| DIS3L2 | 953.00 | 888.00 | 977.00 | 939.33 | 865.00 |
| DISC1 | 4.00 | 7.00 | 5.00 | 5.33 | 0.00 |
| DISC1FP1 | 8.00 | 7.00 | 13.00 | 9.33 | 7.00 |
| DISP1 | 294.00 | 264.00 | 250.00 | 269.33 | 191.00 |
| DISP2 | 256.00 | 309.00 | 304.00 | 289.67 | 248.00 |
| DIXDC1 | 271.00 | 223.00 | 303.00 | 265.67 | 321.00 |
| DKC1 | 3217.00 | 3121.00 | 3439.00 | 3259.00 | 4342.00 |
| DKFZP434A062 | 41.00 | 30.00 | 37.00 | 36.00 | 29.00 |
| DKFZP434I0714 | 151.00 | 172.00 | 203.00 | 175.33 | 162.00 |
| DKFZP586I1420 | 66.00 | 69.00 | 71.00 | 68.67 | 76.00 |
| DKK1 | 8141.00 | 7899.00 | 8556.00 | 8198.67 | 9122.00 |
| DKK2 | 182.00 | 137.00 | 180.00 | 166.33 | 173.00 |
| DKK3 | 20483.00 | 20219.00 | 20554.00 | 20418.67 | 13594.00 |
| DLAT | 1035.00 | 860.00 | 1059.00 | 984.67 | 1159.00 |
| DLC1 | 4957.00 | 4472.00 | 5307.00 | 4912.00 | 4950.00 |
| DLD | 1448.00 | 1422.00 | 1602.00 | 1490.67 | 1751.00 |
| DLEU1 | 258.00 | 232.00 | 306.00 | 265.33 | 330.00 |
| DLEU2 | 44.00 | 56.00 | 56.00 | 52.00 | 55.00 |
| DLEU2L | 11.00 | 11.00 | 15.00 | 12.33 | 14.00 |
| DLG1 | 3010.00 | 2733.00 | 3147.00 | 2963.33 | 3168.00 |
| DLG2 | 15.00 | 9.00 | 19.00 | 14.33 | 23.00 |
| DLG3 | 654.00 | 613.00 | 685.00 | 650.67 | 627.00 |
| DLG4 | 1388.00 | 1344.00 | 1568.00 | 1433.33 | 1389.00 |
| DLG5 | 5.00 | 2.00 | 6.00 | 4.33 | 1.00 |
| DLGAP1-AS1 | 370.00 | 374.00 | 387.00 | 377.00 | 404.00 |
| DLGAP1-AS2 | 375.00 | 345.00 | 335.00 | 351.67 | 350.00 |
| DLGAP3 | 3.00 | 0.00 | 1.00 | 1.33 | 0.00 |
| DLGAP4 | 2706.00 | 2582.00 | 2904.00 | 2730.67 | 2251.00 |
| DLGAP4-AS1 | 3.00 | 2.00 | 0.00 | 1.67 | 7.00 |
| DLGAP5 | 579.00 | 502.00 | 544.00 | 541.67 | 775.00 |
| DLK2 | 3.00 | 5.00 | 3.00 | 3.67 | 2.00 |
| DLL3 | 87.00 | 88.00 | 92.00 | 89.00 | 139.00 |
| DLL4 | 7.00 | 10.00 | 7.00 | 8.00 | 9.00 |
| DLST | 2431.00 | 2358.00 | 2415.00 | 2401.33 | 2619.00 |
| DLSTP1 | 4.00 | 4.00 | 3.00 | 3.67 | 1.00 |
| DLX1 | 786.00 | 707.00 | 787.00 | 760.00 | 961.00 |

| | | | | | |
|---|---|---|---|---|---|
| DLX2 | 61.00 | 66.00 | 75.00 | 67.33 | 100.00 |
| DLX2-AS1 | 4.00 | 3.00 | 7.00 | 4.67 | 12.00 |
| DLX5 | 3.00 | 1.00 | 7.00 | 3.67 | 11.00 |
| DLX6 | 21.00 | 16.00 | 19.00 | 18.67 | 35.00 |
| DMAP1 | 642.00 | 643.00 | 672.00 | 652.33 | 709.00 |
| DMD | 46.00 | 63.00 | 78.00 | 62.33 | 49.00 |
| DMGDH | 34.00 | 32.00 | 24.00 | 30.00 | 18.00 |
| DMPK | 1459.00 | 1544.00 | 1719.00 | 1574.00 | 1464.00 |
| DMRTA1 | 72.00 | 59.00 | 77.00 | 69.33 | 69.00 |
| DMTF1 | 3294.00 | 3115.00 | 3589.00 | 3332.67 | 3301.00 |
| DMTN | 4.00 | 2.00 | 2.00 | 2.67 | 6.00 |
| DMWD | 1020.00 | 1024.00 | 1154.00 | 1066.00 | 873.00 |
| DMXL1 | 743.00 | 726.00 | 860.00 | 776.33 | 668.00 |
| DMXL2 | 1214.00 | 1033.00 | 1210.00 | 1152.33 | 1049.00 |
| DNA2 | 441.00 | 450.00 | 452.00 | 447.67 | 577.00 |
| DNAAF2 | 375.00 | 417.00 | 429.00 | 407.00 | 441.00 |
| DNAAF3 | 4.00 | 14.00 | 14.00 | 10.67 | 12.00 |
| DNAAF5 | 1791.00 | 1591.00 | 1708.00 | 1696.67 | 1794.00 |
| DNAH1 | 217.00 | 221.00 | 249.00 | 229.00 | 150.00 |
| DNAH11 | 684.00 | 671.00 | 757.00 | 704.00 | 508.00 |
| DNAH12 | 26.00 | 19.00 | 34.00 | 26.33 | 15.00 |
| DNAH14 | 661.00 | 666.00 | 664.00 | 663.67 | 727.00 |
| DNAH2 | 219.00 | 223.00 | 246.00 | 229.33 | 191.00 |
| DNAH5 | 388.00 | 358.00 | 445.00 | 397.00 | 301.00 |
| DNAH6 | 41.00 | 23.00 | 30.00 | 31.33 | 25.00 |
| DNAH7 | 17.00 | 12.00 | 9.00 | 12.67 | 12.00 |
| DNAJA1 | 3537.00 | 3371.00 | 3720.00 | 3542.67 | 4141.00 |
| DNAJA2 | 1874.00 | 1778.00 | 2058.00 | 1903.33 | 2333.00 |
| DNAJA3 | 5.00 | 3.00 | 5.00 | 4.33 | 5.00 |
| DNAJB1 | 1547.00 | 1472.00 | 1587.00 | 1535.33 | 1623.00 |
| DNAJB11 | 2594.00 | 2652.00 | 2546.00 | 2597.33 | 2795.00 |
| DNAJB12 | 1806.00 | 1793.00 | 1774.00 | 1791.00 | 2302.00 |
| DNAJB13 | 10.00 | 12.00 | 11.00 | 11.00 | 8.00 |
| DNAJB14 | 739.00 | 615.00 | 801.00 | 718.33 | 635.00 |
| DNAJB2 | 594.00 | 673.00 | 759.00 | 675.33 | 698.00 |
| DNAJB4 | 376.00 | 324.00 | 357.00 | 352.33 | 305.00 |
| DNAJB5 | 141.00 | 148.00 | 131.00 | 140.00 | 159.00 |
| DNAJB6 | 5689.00 | 5396.00 | 5868.00 | 5651.00 | 7369.00 |
| DNAJB9 | 655.00 | 561.00 | 627.00 | 614.33 | 553.00 |
| DNAJC1 | 1411.00 | 1433.00 | 1469.00 | 1437.67 | 1385.00 |
| DNAJC10 | 6140.00 | 5565.00 | 6307.00 | 6004.00 | 5819.00 |
| DNAJC11 | 2228.00 | 2056.00 | 2246.00 | 2176.67 | 2442.00 |
| DNAJC12 | 3.00 | 6.00 | 4.00 | 4.33 | 9.00 |
| DNAJC13 | 2789.00 | 2582.00 | 2875.00 | 2748.67 | 2729.00 |
| DNAJC14 | 1322.00 | 1341.00 | 1415.00 | 1359.33 | 1342.00 |
| DNAJC15 | 305.00 | 307.00 | 326.00 | 312.67 | 411.00 |
| DNAJC16 | 1114.00 | 1064.00 | 1195.00 | 1124.33 | 995.00 |
| DNAJC17 | 252.00 | 241.00 | 263.00 | 252.00 | 334.00 |

| | | | | | |
|---|---|---|---|---|---|
| DNAJC18 | 905.00 | 749.00 | 858.00 | 837.33 | 832.00 |
| DNAJC19 | 719.00 | 699.00 | 705.00 | 707.67 | 939.00 |
| DNAJC2 | 1224.00 | 1219.00 | 1356.00 | 1266.33 | 1462.00 |
| DNAJC21 | 877.00 | 851.00 | 956.00 | 894.67 | 1123.00 |
| DNAJC22 | 308.00 | 300.00 | 296.00 | 301.33 | 333.00 |
| DNAJC24 | 461.00 | 437.00 | 443.00 | 447.00 | 572.00 |
| DNAJC25 | 236.00 | 249.00 | 242.00 | 242.33 | 278.00 |
| DNAJC27 | 194.00 | 158.00 | 185.00 | 179.00 | 166.00 |
| DNAJC3 | 1224.00 | 1155.00 | 1163.00 | 1180.67 | 1127.00 |
| DNAJC30 | 175.00 | 164.00 | 155.00 | 164.67 | 207.00 |
| DNAJC3-AS1 | 64.00 | 68.00 | 86.00 | 72.67 | 55.00 |
| DNAJC4 | 925.00 | 903.00 | 935.00 | 921.00 | 958.00 |
| DNAJC5 | 2474.00 | 2318.00 | 2500.00 | 2430.67 | 2564.00 |
| DNAJC5B | 3.00 | 5.00 | 0.00 | 2.67 | 7.00 |
| DNAJC6 | 216.00 | 171.00 | 192.00 | 193.00 | 205.00 |
| DNAJC7 | 1716.00 | 1619.00 | 1880.00 | 1738.33 | 2117.00 |
| DNAJC8 | 1762.00 | 1739.00 | 1850.00 | 1783.67 | 2411.00 |
| DNAJC9 | 1271.00 | 1194.00 | 1344.00 | 1269.67 | 1513.00 |
| DNAJC9-AS1 | 4.00 | 2.00 | 5.00 | 3.67 | 3.00 |
| DNAL1 | 284.00 | 305.00 | 331.00 | 306.67 | 294.00 |
| DNAL4 | 262.00 | 248.00 | 261.00 | 257.00 | 318.00 |
| DNASE1 | 164.00 | 222.00 | 224.00 | 203.33 | 158.00 |
| DNASE1L1 | 355.00 | 396.00 | 383.00 | 378.00 | 464.00 |
| DNASE1L2 | 92.00 | 91.00 | 91.00 | 91.33 | 81.00 |
| DNASE2 | 1203.00 | 1110.00 | 1118.00 | 1143.67 | 1081.00 |
| DND1 | 96.00 | 84.00 | 83.00 | 87.67 | 111.00 |
| DNER | 1341.00 | 1399.00 | 1415.00 | 1385.00 | 1430.00 |
| DNHD1 | 2292.00 | 2175.00 | 2407.00 | 2291.33 | 1757.00 |
| DNLZ | 184.00 | 206.00 | 237.00 | 209.00 | 221.00 |
| DNM1 | 1425.00 | 1429.00 | 1512.00 | 1455.33 | 1274.00 |
| DNM1L | 2329.00 | 2173.00 | 2528.00 | 2343.33 | 2759.00 |
| DNM1P46 | 19.00 | 21.00 | 18.00 | 19.33 | 19.00 |
| DNM2 | 3072.00 | 2967.00 | 3170.00 | 3069.67 | 3113.00 |
| DNM3 | 19.00 | 22.00 | 19.00 | 20.00 | 24.00 |
| DNMBP | 8211.00 | 7643.00 | 7921.00 | 7925.00 | 7331.00 |
| DNMBP-AS1 | 12.00 | 8.00 | 10.00 | 10.00 | 1.00 |
| DNMT1 | 4093.00 | 3969.00 | 4141.00 | 4067.67 | 4809.00 |
| DNMT3A | 484.00 | 498.00 | 549.00 | 510.33 | 422.00 |
| DNMT3B | 250.00 | 231.00 | 271.00 | 250.67 | 202.00 |
| DNPEP | 913.00 | 972.00 | 1099.00 | 994.67 | 1047.00 |
| DNPH1 | 721.00 | 743.00 | 804.00 | 756.00 | 1139.00 |
| DNTTIP1 | 860.00 | 837.00 | 916.00 | 871.00 | 1131.00 |
| DNTTIP2 | 1888.00 | 1768.00 | 1867.00 | 1841.00 | 2039.00 |
| DOC2A | 6.00 | 5.00 | 6.00 | 5.67 | 5.00 |
| DOCK1 | 2357.00 | 2162.00 | 2422.00 | 2313.67 | 2057.00 |
| DOCK10 | 2026.00 | 1895.00 | 2030.00 | 1983.67 | 1928.00 |
| DOCK11 | 439.00 | 394.00 | 472.00 | 435.00 | 360.00 |
| DOCK2 | 270.00 | 216.00 | 260.00 | 248.67 | 139.00 |

| | | | | | |
|---|---|---|---|---|---|
| DOCK3 | 2029.00 | 2102.00 | 2288.00 | 2139.67 | 1484.00 |
| DOCK4 | 3008.00 | 2486.00 | 2868.00 | 2787.33 | 2664.00 |
| DOCK5 | 1520.00 | 1442.00 | 1510.00 | 1490.67 | 1320.00 |
| DOCK6 | 954.00 | 944.00 | 1134.00 | 1010.67 | 849.00 |
| DOCK7 | 1661.00 | 1491.00 | 1742.00 | 1631.33 | 1608.00 |
| DOCK9 | 1215.00 | 1149.00 | 1321.00 | 1228.33 | 792.00 |
| DOCK9-AS2 | 7.00 | 6.00 | 6.00 | 6.33 | 7.00 |
| DOHH | 381.00 | 398.00 | 393.00 | 390.67 | 439.00 |
| DOK1 | 100.00 | 142.00 | 117.00 | 119.67 | 149.00 |
| DOK2 | 7.00 | 1.00 | 3.00 | 3.67 | 5.00 |
| DOK3 | 232.00 | 216.00 | 236.00 | 228.00 | 203.00 |
| DOK4 | 45.00 | 34.00 | 22.00 | 33.67 | 37.00 |
| DOK6 | 17.00 | 8.00 | 22.00 | 15.67 | 8.00 |
| DOK7 | 8.00 | 7.00 | 14.00 | 9.67 | 3.00 |
| DOLK | 688.00 | 695.00 | 754.00 | 712.33 | 691.00 |
| DOLPP1 | 666.00 | 661.00 | 585.00 | 637.33 | 672.00 |
| DONSON | 417.00 | 446.00 | 402.00 | 421.67 | 470.00 |
| DOPEY1 | 539.00 | 490.00 | 527.00 | 518.67 | 456.00 |
| DOPEY2 | 552.00 | 491.00 | 587.00 | 543.33 | 500.00 |
| DOT1L | 1648.00 | 1707.00 | 1870.00 | 1741.67 | 1709.00 |
| DPAGT1 | 2687.00 | 2662.00 | 2813.00 | 2720.67 | 2591.00 |
| DPCD | 268.00 | 267.00 | 289.00 | 274.67 | 366.00 |
| DPF1 | 60.00 | 85.00 | 76.00 | 73.67 | 62.00 |
| DPF2 | 2206.00 | 2165.00 | 2393.00 | 2254.67 | 2381.00 |
| DPF3 | 434.00 | 383.00 | 477.00 | 431.33 | 399.00 |
| DPH1 | 613.00 | 713.00 | 723.00 | 683.00 | 740.00 |
| DPH2 | 1378.00 | 1315.00 | 1329.00 | 1340.67 | 1459.00 |
| DPH3 | 635.00 | 598.00 | 718.00 | 650.33 | 824.00 |
| DPH5 | 1222.00 | 1295.00 | 1426.00 | 1314.33 | 1633.00 |
| DPH6 | 258.00 | 270.00 | 237.00 | 255.00 | 291.00 |
| DPH6-AS1 | 10.00 | 15.00 | 13.00 | 12.67 | 18.00 |
| DPH7 | 1311.00 | 1339.00 | 1486.00 | 1378.67 | 1401.00 |
| DPM1 | 1197.00 | 1060.00 | 1187.00 | 1148.00 | 1515.00 |
| DPM2 | 965.00 | 949.00 | 865.00 | 926.33 | 1153.00 |
| DPM3 | 276.00 | 280.00 | 241.00 | 265.67 | 315.00 |
| DPP3 | 1451.00 | 1509.00 | 1594.00 | 1518.00 | 1652.00 |
| DPP7 | 2836.00 | 2970.00 | 3113.00 | 2973.00 | 3143.00 |
| DPP8 | 635.00 | 513.00 | 642.00 | 596.67 | 489.00 |
| DPP9 | 6745.00 | 6719.00 | 7205.00 | 6889.67 | 6697.00 |
| DPPA4 | 17.00 | 8.00 | 12.00 | 12.33 | 8.00 |
| DPRXP4 | 5.00 | 9.00 | 10.00 | 8.00 | 8.00 |
| DPY19L1 | 4222.00 | 3824.00 | 4219.00 | 4088.33 | 3347.00 |
| DPY19L1P1 | 111.00 | 108.00 | 128.00 | 115.67 | 119.00 |
| DPY19L2 | 734.00 | 688.00 | 697.00 | 706.33 | 689.00 |
| DPY19L2P1 | 123.00 | 133.00 | 134.00 | 130.00 | 189.00 |
| DPY19L2P2 | 185.00 | 188.00 | 199.00 | 190.67 | 205.00 |
| DPY19L2P3 | 5.00 | 6.00 | 5.00 | 5.33 | 8.00 |
| DPY19L2P4 | 48.00 | 43.00 | 36.00 | 42.33 | 41.00 |

| | | | | | |
|---|---|---|---|---|---|
| DPY19L3 | 1448.00 | 1463.00 | 1577.00 | 1496.00 | 1311.00 |
| DPY19L4 | 1349.00 | 1241.00 | 1424.00 | 1338.00 | 1295.00 |
| DPY30 | 554.00 | 525.00 | 608.00 | 562.33 | 747.00 |
| DPYD | 385.00 | 312.00 | 352.00 | 349.67 | 429.00 |
| DPYS | 8.00 | 2.00 | 0.00 | 3.33 | 3.00 |
| DPYSL2 | 822.00 | 825.00 | 774.00 | 807.00 | 853.00 |
| DPYSL3 | 1741.00 | 1566.00 | 1742.00 | 1683.00 | 1325.00 |
| DQX1 | 15.00 | 12.00 | 18.00 | 15.00 | 15.00 |
| DR1 | 1243.00 | 1154.00 | 1267.00 | 1221.33 | 1425.00 |
| DRAIC | 3.00 | 3.00 | 3.00 | 3.00 | 0.00 |
| DRAM1 | 1011.00 | 1079.00 | 1106.00 | 1065.33 | 1163.00 |
| DRAM2 | 1219.00 | 1151.00 | 1164.00 | 1178.00 | 1391.00 |
| DRAP1 | 11689.00 | 12784.00 | 12804.00 | 12425.67 | 10649.00 |
| DRD1 | 20.00 | 23.00 | 32.00 | 25.00 | 15.00 |
| DRD2 | 3.00 | 11.00 | 7.00 | 7.00 | 4.00 |
| DRD4 | 4.00 | 2.00 | 6.00 | 4.00 | 3.00 |
| DRG1 | 968.00 | 958.00 | 1100.00 | 1008.67 | 1259.00 |
| DRG2 | 934.00 | 946.00 | 969.00 | 949.67 | 1091.00 |
| DROSHA | 2220.00 | 2107.00 | 2261.00 | 2196.00 | 2036.00 |
| DRP2 | 92.00 | 75.00 | 78.00 | 81.67 | 60.00 |
| DSC2 | 48.00 | 48.00 | 50.00 | 48.67 | 40.00 |
| DSCAS | 6.00 | 1.00 | 0.00 | 2.33 | 4.00 |
| DSCC1 | 174.00 | 177.00 | 168.00 | 173.00 | 300.00 |
| DSCR3 | 1134.00 | 1022.00 | 1199.00 | 1118.33 | 1164.00 |
| DSCR4 | 5.00 | 2.00 | 7.00 | 4.67 | 7.00 |
| DSCR8 | 31.00 | 53.00 | 42.00 | 42.00 | 38.00 |
| DSE | 987.00 | 897.00 | 1099.00 | 994.33 | 946.00 |
| DSG2 | 322.00 | 307.00 | 350.00 | 326.33 | 304.00 |
| DSN1 | 738.00 | 720.00 | 803.00 | 753.67 | 811.00 |
| DST | 13194.00 | 12474.00 | 13486.00 | 13051.33 | 10791.00 |
| DSTN | 2144.00 | 2182.00 | 2434.00 | 2253.33 | 2879.00 |
| DSTNP2 | 88.00 | 62.00 | 67.00 | 72.33 | 87.00 |
| DSTYK | 650.00 | 615.00 | 679.00 | 648.00 | 519.00 |
| DTD1 | 1010.00 | 1049.00 | 1200.00 | 1086.33 | 1399.00 |
| DTD2 | 335.00 | 269.00 | 350.00 | 318.00 | 381.00 |
| DTL | 610.00 | 585.00 | 550.00 | 581.67 | 799.00 |
| DTNA | 710.00 | 643.00 | 672.00 | 675.00 | 690.00 |
| DTNB | 194.00 | 182.00 | 210.00 | 195.33 | 269.00 |
| DTNBP1 | 378.00 | 342.00 | 351.00 | 357.00 | 498.00 |
| DTWD1 | 581.00 | 496.00 | 600.00 | 559.00 | 646.00 |
| DTWD2 | 57.00 | 48.00 | 60.00 | 55.00 | 56.00 |
| DTX2 | 173.00 | 148.00 | 160.00 | 160.33 | 154.00 |
| 2P1-UPK3BP1-PMS2 | 30.00 | 44.00 | 38.00 | 37.33 | 32.00 |
| DTX3 | 1830.00 | 1919.00 | 2060.00 | 1936.33 | 1819.00 |
| DTX3L | 732.00 | 595.00 | 799.00 | 708.67 | 582.00 |
| DTYMK | 1077.00 | 1066.00 | 1078.00 | 1073.67 | 1415.00 |
| DUS1L | 3432.00 | 3628.00 | 3571.00 | 3543.67 | 3766.00 |
| DUS2 | 477.00 | 503.00 | 444.00 | 474.67 | 564.00 |

| | | | | | |
|---|---|---|---|---|---|
| DUS3L | 1064.00 | 1003.00 | 1061.00 | 1042.67 | 1281.00 |
| DUS4L | 525.00 | 478.00 | 488.00 | 497.00 | 618.00 |
| DUSP1 | 3564.00 | 3421.00 | 3862.00 | 3615.67 | 4872.00 |
| DUSP10 | 274.00 | 248.00 | 263.00 | 261.67 | 182.00 |
| DUSP11 | 359.00 | 320.00 | 344.00 | 341.00 | 397.00 |
| DUSP12 | 786.00 | 777.00 | 840.00 | 801.00 | 950.00 |
| DUSP14 | 9.00 | 11.00 | 10.00 | 10.00 | 11.00 |
| DUSP16 | 475.00 | 494.00 | 577.00 | 515.33 | 420.00 |
| DUSP18 | 72.00 | 63.00 | 58.00 | 64.33 | 86.00 |
| DUSP19 | 41.00 | 37.00 | 58.00 | 45.33 | 65.00 |
| DUSP2 | 18.00 | 25.00 | 22.00 | 21.67 | 28.00 |
| DUSP22 | 503.00 | 500.00 | 529.00 | 510.67 | 558.00 |
| DUSP23 | 116.00 | 109.00 | 110.00 | 111.67 | 122.00 |
| DUSP28 | 53.00 | 67.00 | 79.00 | 66.33 | 74.00 |
| DUSP3 | 2580.00 | 2465.00 | 2712.00 | 2585.67 | 2006.00 |
| DUSP4 | 1537.00 | 1516.00 | 1660.00 | 1571.00 | 617.00 |
| DUSP5 | 4551.00 | 4425.00 | 4726.00 | 4567.33 | 5664.00 |
| DUSP5P1 | 5.00 | 1.00 | 4.00 | 3.33 | 1.00 |
| DUSP6 | 4915.00 | 4560.00 | 5012.00 | 4829.00 | 4153.00 |
| DUSP7 | 941.00 | 887.00 | 1069.00 | 965.67 | 1133.00 |
| DUT | 1572.00 | 1428.00 | 1530.00 | 1510.00 | 1909.00 |
| DUXAP10 | 3.00 | 4.00 | 7.00 | 4.67 | 2.00 |
| DUXAP8 | 183.00 | 156.00 | 167.00 | 168.67 | 143.00 |
| DVL1 | 2808.00 | 2814.00 | 3037.00 | 2886.33 | 3100.00 |
| DVL2 | 1703.00 | 1710.00 | 1725.00 | 1712.67 | 1939.00 |
| DVL3 | 2595.00 | 2664.00 | 2949.00 | 2736.00 | 2387.00 |
| DXO | 41.00 | 37.00 | 55.00 | 44.33 | 55.00 |
| DYM | 551.00 | 508.00 | 655.00 | 571.33 | 645.00 |
| DYNAP | 9.00 | 8.00 | 3.00 | 6.67 | 11.00 |
| DYNC1H1 | 29813.00 | 26977.00 | 30524.00 | 29104.67 | 26552.00 |
| DYNC1I1 | 38.00 | 37.00 | 45.00 | 40.00 | 54.00 |
| DYNC1I2 | 1886.00 | 1720.00 | 1885.00 | 1830.33 | 2052.00 |
| DYNC1LI1 | 984.00 | 921.00 | 1000.00 | 968.33 | 1316.00 |
| DYNC1LI2 | 2922.00 | 2724.00 | 2981.00 | 2875.67 | 2753.00 |
| DYNC2H1 | 3988.00 | 3622.00 | 4191.00 | 3933.67 | 3308.00 |
| DYNC2LI1 | 446.00 | 400.00 | 516.00 | 454.00 | 508.00 |
| DYNLL1 | 2442.00 | 2416.00 | 2614.00 | 2490.67 | 3138.00 |
| DYNLL2 | 922.00 | 916.00 | 929.00 | 922.33 | 862.00 |
| DYNLRB1 | 1218.00 | 1129.00 | 1356.00 | 1234.33 | 1609.00 |
| DYNLT1 | 630.00 | 716.00 | 712.00 | 686.00 | 808.00 |
| DYNLT3 | 293.00 | 252.00 | 330.00 | 291.67 | 337.00 |
| DYRK1A | 1071.00 | 985.00 | 1187.00 | 1081.00 | 1097.00 |
| DYRK1B | 134.00 | 133.00 | 130.00 | 132.33 | 119.00 |
| DYRK2 | 807.00 | 690.00 | 929.00 | 808.67 | 652.00 |
| DYRK3 | 558.00 | 602.00 | 621.00 | 593.67 | 523.00 |
| DYRK4 | 361.00 | 387.00 | 401.00 | 383.00 | 417.00 |
| DYSF | 4.00 | 0.00 | 0.00 | 1.33 | 1.00 |
| DYX1C1 | 39.00 | 40.00 | 31.00 | 36.67 | 52.00 |

| | | | | | |
|---|---|---|---|---|---|
| DZANK1 | 180.00 | 197.00 | 196.00 | 191.00 | 174.00 |
| DZIP1 | 965.00 | 884.00 | 1026.00 | 958.33 | 1049.00 |
| DZIP1L | 326.00 | 286.00 | 307.00 | 306.33 | 314.00 |
| DZIP3 | 862.00 | 851.00 | 879.00 | 864.00 | 1138.00 |
| E2F1 | 510.00 | 522.00 | 483.00 | 505.00 | 860.00 |
| E2F2 | 15.00 | 10.00 | 11.00 | 12.00 | 43.00 |
| E2F3 | 1883.00 | 1711.00 | 1895.00 | 1829.67 | 2168.00 |
| E2F4 | 2364.00 | 2361.00 | 2511.00 | 2412.00 | 2767.00 |
| E2F5 | 755.00 | 813.00 | 859.00 | 809.00 | 1003.00 |
| E2F6 | 457.00 | 446.00 | 447.00 | 450.00 | 567.00 |
| E2F7 | 5157.00 | 4948.00 | 5345.00 | 5150.00 | 3100.00 |
| E2F8 | 50.00 | 40.00 | 45.00 | 45.00 | 121.00 |
| E4F1 | 931.00 | 1042.00 | 1066.00 | 1013.00 | 1037.00 |
| EAF1 | 745.00 | 593.00 | 757.00 | 698.33 | 725.00 |
| EAF2 | 31.00 | 22.00 | 20.00 | 24.33 | 47.00 |
| EAPP | 413.00 | 415.00 | 438.00 | 422.00 | 484.00 |
| EARS2 | 830.00 | 803.00 | 916.00 | 849.67 | 810.00 |
| EBAG9 | 500.00 | 478.00 | 480.00 | 486.00 | 557.00 |
| EBF1 | 33.00 | 17.00 | 25.00 | 25.00 | 14.00 |
| EBF2 | 20.00 | 13.00 | 9.00 | 14.00 | 15.00 |
| EBF4 | 17.00 | 16.00 | 19.00 | 17.33 | 10.00 |
| EBI3 | 3.00 | 0.00 | 2.00 | 1.67 | 1.00 |
| EBLN1 | 3.00 | 0.00 | 0.00 | 1.00 | 1.00 |
| EBLN2 | 94.00 | 95.00 | 100.00 | 96.33 | 83.00 |
| EBLN3 | 2097.00 | 1968.00 | 2373.00 | 2146.00 | 2155.00 |
| EBNA1BP2 | 3037.00 | 3023.00 | 3073.00 | 3044.33 | 4725.00 |
| EBP | 559.00 | 576.00 | 507.00 | 547.33 | 614.00 |
| EBPL | 878.00 | 842.00 | 829.00 | 849.67 | 1106.00 |
| ECD | 1189.00 | 1124.00 | 1282.00 | 1198.33 | 1292.00 |
| ECE1 | 3117.00 | 3030.00 | 2883.00 | 3010.00 | 2565.00 |
| ECE2 | 104.00 | 98.00 | 130.00 | 110.67 | 141.00 |
| ECH1 | 302.00 | 350.00 | 389.00 | 347.00 | 510.00 |
| ECHDC1 | 1125.00 | 1095.00 | 1170.00 | 1130.00 | 1517.00 |
| ECHDC2 | 647.00 | 728.00 | 693.00 | 689.33 | 723.00 |
| ECHDC3 | 107.00 | 96.00 | 103.00 | 102.00 | 127.00 |
| ECHS1 | 1714.00 | 1683.00 | 1756.00 | 1717.67 | 1645.00 |
| ECI1 | 513.00 | 518.00 | 509.00 | 513.33 | 659.00 |
| ECI2 | 7689.00 | 7525.00 | 8281.00 | 7831.67 | 9476.00 |
| ECM1 | 264.00 | 253.00 | 243.00 | 253.33 | 456.00 |
| ECSCR | 511.00 | 500.00 | 519.00 | 510.00 | 855.00 |
| ECSIT | 531.00 | 479.00 | 539.00 | 516.33 | 642.00 |
| ECT2 | 1542.00 | 1316.00 | 1592.00 | 1483.33 | 2058.00 |
| ECT2L | 3.00 | 3.00 | 1.00 | 2.33 | 0.00 |
| EDA | 13.00 | 10.00 | 10.00 | 11.00 | 13.00 |
| EDA2R | 402.00 | 375.00 | 493.00 | 423.33 | 349.00 |
| EDC3 | 1159.00 | 1168.00 | 1259.00 | 1195.33 | 1190.00 |
| EDC4 | 2071.00 | 2122.00 | 2206.00 | 2133.00 | 2072.00 |
| EDEM1 | 3027.00 | 2794.00 | 3101.00 | 2974.00 | 3157.00 |

| | | | | | |
|---|---|---|---|---|---|
| EDEM2 | 1594.00 | 1541.00 | 1681.00 | 1605.33 | 1616.00 |
| EDEM3 | 2354.00 | 2072.00 | 2464.00 | 2296.67 | 2281.00 |
| EDF1 | 2685.00 | 2639.00 | 2618.00 | 2647.33 | 3351.00 |
| EDIL3 | 101.00 | 97.00 | 116.00 | 104.67 | 13.00 |
| EDN1 | 1816.00 | 1942.00 | 1981.00 | 1913.00 | 3268.00 |
| EDNRA | 47.00 | 32.00 | 35.00 | 38.00 | 63.00 |
| EDNRB | 122.00 | 135.00 | 123.00 | 126.67 | 141.00 |
| EDNRB-AS1 | 4.00 | 1.00 | 3.00 | 2.67 | 2.00 |
| EDRF1 | 1296.00 | 1264.00 | 1289.00 | 1283.00 | 1228.00 |
| EEA1 | 1190.00 | 986.00 | 1236.00 | 1137.33 | 1002.00 |
| EED | 334.00 | 333.00 | 318.00 | 328.33 | 401.00 |
| EEF1A1 | 101709.00 | 93779.00 | 113343.00 | 102943.67 | 134787.00 |
| EEF1A2 | 11.00 | 11.00 | 13.00 | 11.67 | 11.00 |
| EEF1B2 | 6348.00 | 6228.00 | 6855.00 | 6477.00 | 9076.00 |
| EEF1D | 28.00 | 10.00 | 22.00 | 20.00 | 29.00 |
| EEF1DP3 | 3.00 | 2.00 | 6.00 | 3.67 | 1.00 |
| EEF1E1 | 396.00 | 382.00 | 453.00 | 410.33 | 593.00 |
| EEF1G | 22992.00 | 22243.00 | 24838.00 | 23357.67 | 30013.00 |
| EEF2 | 65845.00 | 62362.00 | 71160.00 | 66455.67 | 68269.00 |
| EEF2K | 831.00 | 843.00 | 1043.00 | 905.67 | 1094.00 |
| EEF2KMT | 362.00 | 370.00 | 317.00 | 349.67 | 438.00 |
| EEFSEC | 343.00 | 397.00 | 406.00 | 382.00 | 407.00 |
| EEPD1 | 44.00 | 44.00 | 67.00 | 51.67 | 102.00 |
| EFCAB11 | 143.00 | 167.00 | 162.00 | 157.33 | 159.00 |
| EFCAB12 | 3.00 | 2.00 | 3.00 | 2.67 | 1.00 |
| EFCAB13 | 246.00 | 222.00 | 257.00 | 241.67 | 224.00 |
| EFCAB14 | 1712.00 | 1629.00 | 1768.00 | 1703.00 | 1655.00 |
| EFCAB2 | 120.00 | 112.00 | 116.00 | 116.00 | 143.00 |
| EFCAB5 | 8.00 | 7.00 | 6.00 | 7.00 | 8.00 |
| EFCAB7 | 191.00 | 192.00 | 211.00 | 198.00 | 231.00 |
| EFEMP1 | 1209.00 | 1134.00 | 1237.00 | 1193.33 | 1810.00 |
| EFEMP2 | 524.00 | 572.00 | 573.00 | 556.33 | 577.00 |
| EFHC1 | 1405.00 | 1431.00 | 1455.00 | 1430.33 | 1427.00 |
| EFHC2 | 11.00 | 17.00 | 5.00 | 11.00 | 13.00 |
| EFHD2 | 2337.00 | 2296.00 | 2326.00 | 2319.67 | 3155.00 |
| EFNA2 | 5.00 | 0.00 | 4.00 | 3.00 | 4.00 |
| EFNA3 | 27.00 | 28.00 | 30.00 | 28.33 | 35.00 |
| EFNA4 | 110.00 | 112.00 | 92.00 | 104.67 | 98.00 |
| EFNA5 | 357.00 | 358.00 | 404.00 | 373.00 | 417.00 |
| EFNB1 | 1033.00 | 1106.00 | 1123.00 | 1087.33 | 755.00 |
| EFNB2 | 320.00 | 328.00 | 300.00 | 316.00 | 264.00 |
| EFNB3 | 9.00 | 6.00 | 15.00 | 10.00 | 8.00 |
| EFR3A | 1731.00 | 1570.00 | 1933.00 | 1744.67 | 1788.00 |
| EFTUD1 | 981.00 | 956.00 | 1110.00 | 1015.67 | 1089.00 |
| EFTUD1P1 | 7.00 | 3.00 | 4.00 | 4.67 | 2.00 |
| EFTUD2 | 3499.00 | 3294.00 | 3670.00 | 3487.67 | 3900.00 |
| EGF | 17.00 | 19.00 | 23.00 | 19.67 | 4.00 |
| EGFEM1P | 7.00 | 3.00 | 3.00 | 4.33 | 5.00 |

| | | | | | |
|---|---|---|---|---|---|
| EGFL7 | 30.00 | 22.00 | 33.00 | 28.33 | 34.00 |
| EGFR | 10026.00 | 8684.00 | 10292.00 | 9667.33 | 5998.00 |
| EGLN1 | 1200.00 | 1016.00 | 1220.00 | 1145.33 | 1413.00 |
| EGLN2 | 19.00 | 13.00 | 16.00 | 16.00 | 31.00 |
| EGR1 | 1389.00 | 1233.00 | 1691.00 | 1437.67 | 1888.00 |
| EGR2 | 17.00 | 19.00 | 27.00 | 21.00 | 17.00 |
| EHBP1 | 1742.00 | 1641.00 | 1772.00 | 1718.33 | 1867.00 |
| EHBP1L1 | 2519.00 | 2721.00 | 2606.00 | 2615.33 | 2674.00 |
| EHD1 | 2309.00 | 2151.00 | 2292.00 | 2250.67 | 2715.00 |
| EHD2 | 1151.00 | 1079.00 | 1239.00 | 1156.33 | 858.00 |
| EHD3 | 8.00 | 21.00 | 16.00 | 15.00 | 22.00 |
| EHD4 | 2277.00 | 1974.00 | 2080.00 | 2110.33 | 3075.00 |
| EHF | 3.00 | 2.00 | 1.00 | 2.00 | 1.00 |
| EHHADH | 73.00 | 72.00 | 70.00 | 71.67 | 86.00 |
| EHMT1 | 1462.00 | 1401.00 | 1563.00 | 1475.33 | 1401.00 |
| EI24 | 1048.00 | 1028.00 | 1066.00 | 1047.33 | 1300.00 |
| EID1 | 2270.00 | 2143.00 | 2435.00 | 2282.67 | 2601.00 |
| EID2 | 204.00 | 197.00 | 210.00 | 203.67 | 225.00 |
| EID2B | 69.00 | 69.00 | 75.00 | 71.00 | 87.00 |
| EID3 | 23.00 | 30.00 | 32.00 | 28.33 | 29.00 |
| EIF1 | 13289.00 | 13226.00 | 14966.00 | 13827.00 | 17460.00 |
| EIF1AD | 902.00 | 900.00 | 921.00 | 907.67 | 771.00 |
| EIF1AX | 1949.00 | 1690.00 | 2078.00 | 1905.67 | 2403.00 |
| EIF1B | 1106.00 | 1088.00 | 1216.00 | 1136.67 | 1456.00 |
| EIF1B-AS1 | 9.00 | 11.00 | 10.00 | 10.00 | 14.00 |
| EIF2A | 2913.00 | 2882.00 | 3331.00 | 3042.00 | 3828.00 |
| EIF2AK1 | 2367.00 | 2259.00 | 2504.00 | 2376.67 | 2452.00 |
| EIF2AK2 | 518.00 | 463.00 | 555.00 | 512.00 | 460.00 |
| EIF2AK3 | 533.00 | 500.00 | 520.00 | 517.67 | 476.00 |
| EIF2AK4 | 2681.00 | 2418.00 | 2802.00 | 2633.67 | 2554.00 |
| EIF2B1 | 1176.00 | 1065.00 | 1155.00 | 1132.00 | 1365.00 |
| EIF2B2 | 1018.00 | 956.00 | 1055.00 | 1009.67 | 1059.00 |
| EIF2B3 | 698.00 | 678.00 | 683.00 | 686.33 | 909.00 |
| EIF2B4 | 891.00 | 892.00 | 928.00 | 903.67 | 905.00 |
| EIF2B5 | 1728.00 | 1628.00 | 1765.00 | 1707.00 | 1680.00 |
| EIF2D | 1914.00 | 1906.00 | 2040.00 | 1953.33 | 2057.00 |
| EIF2S1 | 2108.00 | 1949.00 | 2219.00 | 2092.00 | 3006.00 |
| EIF2S2 | 5109.00 | 4717.00 | 5404.00 | 5076.67 | 4374.00 |
| EIF2S3 | 10383.00 | 9676.00 | 10740.00 | 10266.33 | 11354.00 |
| EIF3A | 13971.00 | 12996.00 | 14911.00 | 13959.33 | 13257.00 |
| EIF3B | 6563.00 | 6580.00 | 7015.00 | 6719.33 | 7941.00 |
| EIF3D | 6324.00 | 6110.00 | 6978.00 | 6470.67 | 8135.00 |
| EIF3E | 10866.00 | 10419.00 | 11640.00 | 10975.00 | 14071.00 |
| EIF3F | 1908.00 | 1905.00 | 2124.00 | 1979.00 | 2568.00 |
| EIF3G | 3124.00 | 3036.00 | 3196.00 | 3118.67 | 3833.00 |
| EIF3H | 4872.00 | 4561.00 | 5277.00 | 4903.33 | 6657.00 |
| EIF3I | 3863.00 | 3621.00 | 4216.00 | 3900.00 | 4929.00 |
| EIF3J | 1964.00 | 1913.00 | 2015.00 | 1964.00 | 2455.00 |

| | | | | | |
|---|---|---|---|---|---|
| EIF3J-AS1 | 277.00 | 302.00 | 316.00 | 298.33 | 345.00 |
| EIF3K | 1873.00 | 1883.00 | 1962.00 | 1906.00 | 2615.00 |
| EIF3L | 5768.00 | 5268.00 | 6118.00 | 5718.00 | 6762.00 |
| EIF3M | 4181.00 | 4060.00 | 4592.00 | 4277.67 | 5851.00 |
| EIF4A2 | 14151.00 | 14038.00 | 15565.00 | 14584.67 | 16342.00 |
| EIF4A3 | 2219.00 | 2121.00 | 2287.00 | 2209.00 | 2678.00 |
| EIF4B | 8228.00 | 7489.00 | 9249.00 | 8322.00 | 9014.00 |
| EIF4E | 436.00 | 376.00 | 495.00 | 435.67 | 563.00 |
| EIF4E2 | 818.00 | 834.00 | 916.00 | 856.00 | 1015.00 |
| EIF4E3 | 29.00 | 28.00 | 15.00 | 24.00 | 27.00 |
| EIF4EBP1 | 3338.00 | 3194.00 | 3434.00 | 3322.00 | 4454.00 |
| EIF4EBP2 | 1434.00 | 1316.00 | 1536.00 | 1428.67 | 1570.00 |
| EIF4EBP3 | 40.00 | 43.00 | 47.00 | 43.33 | 85.00 |
| EIF4ENIF1 | 569.00 | 548.00 | 566.00 | 561.00 | 552.00 |
| EIF4G1 | 18903.00 | 18701.00 | 19210.00 | 18938.00 | 18856.00 |
| EIF4G2 | 18153.00 | 16485.00 | 19704.00 | 18114.00 | 19658.00 |
| EIF4G3 | 3933.00 | 3759.00 | 4092.00 | 3928.00 | 3699.00 |
| EIF4H | 4720.00 | 4717.00 | 5005.00 | 4814.00 | 5747.00 |
| EIF5 | 5314.00 | 5141.00 | 5469.00 | 5308.00 | 5681.00 |
| EIF5A | 10337.00 | 10063.00 | 10826.00 | 10408.67 | 14207.00 |
| EIF5A2 | 256.00 | 263.00 | 330.00 | 283.00 | 298.00 |
| EIF5AL1 | 332.00 | 278.00 | 331.00 | 313.67 | 466.00 |
| EIF5B | 5373.00 | 5194.00 | 5702.00 | 5423.00 | 7015.00 |
| EIF6 | 4339.00 | 4247.00 | 4626.00 | 4404.00 | 5512.00 |
| ELAC1 | 186.00 | 184.00 | 141.00 | 170.33 | 226.00 |
| ELAC2 | 1603.00 | 1720.00 | 1638.00 | 1653.67 | 1838.00 |
| ELAVL1 | 1853.00 | 1737.00 | 1875.00 | 1821.67 | 2300.00 |
| ELAVL2 | 54.00 | 64.00 | 55.00 | 57.67 | 75.00 |
| ELF1 | 669.00 | 566.00 | 642.00 | 625.67 | 560.00 |
| ELF2 | 312.00 | 291.00 | 309.00 | 304.00 | 343.00 |
| ELF4 | 363.00 | 334.00 | 337.00 | 344.67 | 296.00 |
| ELFN1 | 9.00 | 11.00 | 3.00 | 7.67 | 8.00 |
| ELFN1-AS1 | 38.00 | 58.00 | 45.00 | 47.00 | 67.00 |
| ELFN2 | 3022.00 | 3013.00 | 3154.00 | 3063.00 | 2015.00 |
| ELK1 | 753.00 | 730.00 | 786.00 | 756.33 | 897.00 |
| ELK3 | 5447.00 | 4784.00 | 5510.00 | 5247.00 | 6694.00 |
| ELK4 | 2022.00 | 1920.00 | 2204.00 | 2048.67 | 1851.00 |
| ELL | 724.00 | 663.00 | 713.00 | 700.00 | 758.00 |
| ELL2 | 3273.00 | 2746.00 | 3606.00 | 3208.33 | 3827.00 |
| ELL3 | 14.00 | 14.00 | 12.00 | 13.33 | 16.00 |
| ELMO2 | 882.00 | 803.00 | 916.00 | 867.00 | 849.00 |
| ELMO3 | 128.00 | 144.00 | 139.00 | 137.00 | 138.00 |
| ELMOD1 | 130.00 | 144.00 | 136.00 | 136.67 | 168.00 |
| ELMOD2 | 785.00 | 664.00 | 765.00 | 738.00 | 727.00 |
| ELMOD3 | 645.00 | 672.00 | 660.00 | 659.00 | 687.00 |
| ELMSAN1 | 949.00 | 775.00 | 923.00 | 882.33 | 770.00 |
| ELOF1 | 625.00 | 650.00 | 728.00 | 667.67 | 751.00 |
| ELOVL1 | 3626.00 | 3744.00 | 3700.00 | 3690.00 | 4441.00 |

| | | | | | |
|---|---|---|---|---|---|
| ELOVL2 | 191.00 | 198.00 | 195.00 | 194.67 | 327.00 |
| ELOVL3 | 8.00 | 11.00 | 15.00 | 11.33 | 21.00 |
| ELOVL4 | 508.00 | 501.00 | 543.00 | 517.33 | 581.00 |
| ELOVL5 | 4136.00 | 3942.00 | 4431.00 | 4169.67 | 4968.00 |
| ELOVL6 | 1492.00 | 1405.00 | 1350.00 | 1415.67 | 1541.00 |
| ELOVL7 | 61.00 | 62.00 | 58.00 | 60.33 | 62.00 |
| ELP2 | 1272.00 | 1256.00 | 1387.00 | 1305.00 | 1421.00 |
| ELP3 | 552.00 | 507.00 | 597.00 | 552.00 | 662.00 |
| ELP4 | 230.00 | 236.00 | 280.00 | 248.67 | 294.00 |
| ELP5 | 1284.00 | 1217.00 | 1324.00 | 1275.00 | 1660.00 |
| ELP6 | 593.00 | 641.00 | 679.00 | 637.67 | 811.00 |
| EMBP1 | 4.00 | 6.00 | 7.00 | 5.67 | 4.00 |
| EMC1 | 4530.00 | 4501.00 | 4752.00 | 4594.33 | 4005.00 |
| EMC10 | 1227.00 | 1300.00 | 1199.00 | 1242.00 | 1149.00 |
| EMC2 | 756.00 | 690.00 | 806.00 | 750.67 | 996.00 |
| EMC3 | 1135.00 | 1141.00 | 1036.00 | 1104.00 | 1185.00 |
| EMC3-AS1 | 384.00 | 483.00 | 441.00 | 436.00 | 484.00 |
| EMC4 | 1685.00 | 1731.00 | 1619.00 | 1678.33 | 2353.00 |
| EMC6 | 592.00 | 582.00 | 591.00 | 588.33 | 735.00 |
| EMC7 | 1698.00 | 1769.00 | 1585.00 | 1684.00 | 2086.00 |
| EMC8 | 727.00 | 684.00 | 800.00 | 737.00 | 747.00 |
| EMC9 | 219.00 | 232.00 | 207.00 | 219.33 | 246.00 |
| EMD | 1162.00 | 1177.00 | 1192.00 | 1177.00 | 1543.00 |
| EME1 | 762.00 | 735.00 | 761.00 | 752.67 | 820.00 |
| EME2 | 236.00 | 244.00 | 255.00 | 245.00 | 198.00 |
| EMG1 | 896.00 | 858.00 | 962.00 | 905.33 | 1140.00 |
| EMILIN1 | 271.00 | 268.00 | 237.00 | 258.67 | 184.00 |
| EMILIN2 | 689.00 | 677.00 | 715.00 | 693.67 | 791.00 |
| EML1 | 99.00 | 95.00 | 89.00 | 94.33 | 78.00 |
| EML2 | 972.00 | 1047.00 | 1005.00 | 1008.00 | 980.00 |
| EML2-AS1 | 202.00 | 192.00 | 208.00 | 200.67 | 127.00 |
| EML3 | 1881.00 | 1914.00 | 2131.00 | 1975.33 | 1890.00 |
| EML4 | 1394.00 | 1421.00 | 1574.00 | 1463.00 | 1508.00 |
| EML5 | 76.00 | 51.00 | 72.00 | 66.33 | 69.00 |
| EML6 | 219.00 | 200.00 | 202.00 | 207.00 | 154.00 |
| EMP1 | 24554.00 | 23959.00 | 22949.00 | 23820.67 | 36952.00 |
| EMP2 | 191.00 | 200.00 | 169.00 | 186.67 | 276.00 |
| EMP3 | 1675.00 | 1591.00 | 1539.00 | 1601.67 | 2128.00 |
| ENAH | 1687.00 | 1482.00 | 1935.00 | 1701.33 | 1676.00 |
| ENC1 | 5278.00 | 4880.00 | 5178.00 | 5112.00 | 5775.00 |
| ENDOD1 | 818.00 | 830.00 | 826.00 | 824.67 | 634.00 |
| ENDOG | 185.00 | 125.00 | 171.00 | 160.33 | 206.00 |
| ENDOU | 3.00 | 9.00 | 2.00 | 4.67 | 11.00 |
| ENDOV | 371.00 | 401.00 | 459.00 | 410.33 | 347.00 |
| ENG | 16228.00 | 16187.00 | 15077.00 | 15830.67 | 12864.00 |
| ENGASE | 1002.00 | 1101.00 | 1116.00 | 1073.00 | 996.00 |
| ENHO | 4.00 | 4.00 | 2.00 | 3.33 | 10.00 |
| ENKD1 | 471.00 | 542.00 | 505.00 | 506.00 | 613.00 |

| | | | | | |
|---|---|---|---|---|---|
| ENKUR | 9.00 | 11.00 | 9.00 | 9.67 | 13.00 |
| ENO1 | 33552.00 | 31589.00 | 35324.00 | 33488.33 | 38214.00 |
| ENO1-AS1 | 7.00 | 2.00 | 5.00 | 4.67 | 4.00 |
| ENO2 | 1263.00 | 1433.00 | 1560.00 | 1418.67 | 1560.00 |
| ENO3 | 636.00 | 629.00 | 700.00 | 655.00 | 812.00 |
| ENOPH1 | 687.00 | 744.00 | 833.00 | 754.67 | 967.00 |
| ENOSF1 | 3546.00 | 3491.00 | 3760.00 | 3599.00 | 4091.00 |
| ENOX1 | 40.00 | 54.00 | 58.00 | 50.67 | 96.00 |
| ENOX2 | 236.00 | 214.00 | 252.00 | 234.00 | 264.00 |
| ENPEP | 118.00 | 118.00 | 122.00 | 119.33 | 94.00 |
| ENPP1 | 1880.00 | 1722.00 | 1739.00 | 1780.33 | 1787.00 |
| ENPP2 | 51.00 | 42.00 | 50.00 | 47.67 | 98.00 |
| ENPP3 | 5.00 | 6.00 | 1.00 | 4.00 | 3.00 |
| ENPP4 | 349.00 | 348.00 | 363.00 | 353.33 | 304.00 |
| ENSA | 1640.00 | 1556.00 | 1683.00 | 1626.33 | 2033.00 |
| ENTHD2 | 1151.00 | 1159.00 | 1199.00 | 1169.67 | 1167.00 |
| ENTPD1-AS1 | 76.00 | 77.00 | 90.00 | 81.00 | 85.00 |
| ENTPD3 | 498.00 | 463.00 | 518.00 | 493.00 | 125.00 |
| ENTPD3-AS1 | 23.00 | 24.00 | 24.00 | 23.67 | 41.00 |
| ENTPD4 | 3551.00 | 3436.00 | 3635.00 | 3540.67 | 3566.00 |
| ENTPD5 | 263.00 | 265.00 | 235.00 | 254.33 | 216.00 |
| ENTPD6 | 3182.00 | 3296.00 | 3220.00 | 3232.67 | 3144.00 |
| ENTPD7 | 878.00 | 826.00 | 1009.00 | 904.33 | 996.00 |
| ENY2 | 1004.00 | 957.00 | 1045.00 | 1002.00 | 1268.00 |
| EOGT | 1234.00 | 1208.00 | 1290.00 | 1244.00 | 1429.00 |
| EP300 | 2240.00 | 1956.00 | 2388.00 | 2194.67 | 1594.00 |
| EP300-AS1 | 3.00 | 2.00 | 5.00 | 3.33 | 3.00 |
| EP400 | 2210.00 | 2147.00 | 2366.00 | 2241.00 | 1780.00 |
| EP400NL | 271.00 | 267.00 | 264.00 | 267.33 | 203.00 |
| EPAS1 | 380.00 | 346.00 | 375.00 | 367.00 | 286.00 |
| EPB41 | 890.00 | 901.00 | 1073.00 | 954.67 | 812.00 |
| EPB41L1 | 567.00 | 519.00 | 555.00 | 547.00 | 562.00 |
| EPB41L2 | 3597.00 | 3408.00 | 3495.00 | 3500.00 | 4042.00 |
| EPB41L3 | 1605.00 | 1361.00 | 1688.00 | 1551.33 | 1945.00 |
| EPB41L4A | 10.00 | 14.00 | 27.00 | 17.00 | 20.00 |
| EPB41L4A-AS1 | 683.00 | 676.00 | 734.00 | 697.67 | 845.00 |
| EPB41L4B | 109.00 | 90.00 | 118.00 | 105.67 | 145.00 |
| EPB41L5 | 488.00 | 453.00 | 492.00 | 477.67 | 455.00 |
| EPC1 | 790.00 | 714.00 | 887.00 | 797.00 | 752.00 |
| EPC2 | 499.00 | 441.00 | 594.00 | 511.33 | 486.00 |
| EPCAM | 4.00 | 0.00 | 0.00 | 1.33 | 6.00 |
| EPDR1 | 11.00 | 13.00 | 9.00 | 11.00 | 8.00 |
| EPG5 | 6143.00 | 5720.00 | 6573.00 | 6145.33 | 6716.00 |
| EPGN | 4.00 | 8.00 | 8.00 | 6.67 | 6.00 |
| EPHA1 | 3.00 | 1.00 | 3.00 | 2.33 | 0.00 |
| EPHA2 | 16661.00 | 16783.00 | 16280.00 | 16574.67 | 13645.00 |
| EPHA4 | 2244.00 | 2049.00 | 2231.00 | 2174.67 | 3299.00 |
| EPHA5 | 117.00 | 92.00 | 135.00 | 114.67 | 132.00 |

| | | | | | |
|---|---|---|---|---|---|
| EPHA5-AS1 | 6.00 | 5.00 | 6.00 | 5.67 | 25.00 |
| EPHA7 | 17.00 | 30.00 | 34.00 | 27.00 | 31.00 |
| EPHB1 | 121.00 | 121.00 | 121.00 | 121.00 | 162.00 |
| EPHB2 | 3199.00 | 3180.00 | 3105.00 | 3161.33 | 1927.00 |
| EPHB4 | 1955.00 | 1922.00 | 2013.00 | 1963.33 | 2023.00 |
| EPHB6 | 24.00 | 13.00 | 14.00 | 17.00 | 7.00 |
| EPHX1 | 565.00 | 575.00 | 514.00 | 551.33 | 481.00 |
| EPHX3 | 4.00 | 3.00 | 3.00 | 3.33 | 6.00 |
| EPHX4 | 206.00 | 247.00 | 213.00 | 222.00 | 286.00 |
| EPM2A | 125.00 | 144.00 | 140.00 | 136.33 | 112.00 |
| EPM2AIP1 | 1374.00 | 1297.00 | 1443.00 | 1371.33 | 1139.00 |
| EPN1 | 1863.00 | 1878.00 | 1976.00 | 1905.67 | 2113.00 |
| EPN2 | 1333.00 | 1205.00 | 1320.00 | 1286.00 | 1087.00 |
| EPOR | 304.00 | 334.00 | 321.00 | 319.67 | 298.00 |
| EPPK1 | 7.00 | 4.00 | 0.00 | 3.67 | 0.00 |
| EPRS | 9899.00 | 9035.00 | 10334.00 | 9756.00 | 9640.00 |
| EPS15 | 1258.00 | 1123.00 | 1355.00 | 1245.33 | 1441.00 |
| EPS15L1 | 881.00 | 800.00 | 834.00 | 838.33 | 795.00 |
| EPS8 | 445.00 | 502.00 | 550.00 | 499.00 | 500.00 |
| EPS8L2 | 15.00 | 15.00 | 14.00 | 14.67 | 20.00 |
| EPT1 | 2336.00 | 2231.00 | 2420.00 | 2329.00 | 2191.00 |
| ERAL1 | 1185.00 | 1084.00 | 1063.00 | 1110.67 | 1345.00 |
| ERAP1 | 2425.00 | 2369.00 | 2597.00 | 2463.67 | 2403.00 |
| ERAP2 | 272.00 | 296.00 | 318.00 | 295.33 | 281.00 |
| ERBB2 | 48.00 | 59.00 | 35.00 | 47.33 | 76.00 |
| ERBB2IP | 2132.00 | 1851.00 | 2181.00 | 2054.67 | 2056.00 |
| ERBB3 | 6.00 | 4.00 | 6.00 | 5.33 | 3.00 |
| ERBB4 | 21.00 | 29.00 | 18.00 | 22.67 | 20.00 |
| ERC1 | 1622.00 | 1448.00 | 1728.00 | 1599.33 | 1273.00 |
| ERCC1 | 2412.00 | 2309.00 | 2521.00 | 2414.00 | 2442.00 |
| ERCC2 | 2120.00 | 2003.00 | 2226.00 | 2116.33 | 1600.00 |
| ERCC3 | 1445.00 | 1412.00 | 1613.00 | 1490.00 | 1644.00 |
| ERCC4 | 494.00 | 439.00 | 557.00 | 496.67 | 425.00 |
| ERCC5 | 58.00 | 59.00 | 66.00 | 61.00 | 49.00 |
| ERCC6 | 280.00 | 291.00 | 326.00 | 299.00 | 285.00 |
| ERCC6L | 306.00 | 288.00 | 300.00 | 298.00 | 435.00 |
| ERCC6L2 | 547.00 | 555.00 | 631.00 | 577.67 | 539.00 |
| ERCC8 | 244.00 | 238.00 | 261.00 | 247.67 | 280.00 |
| EREG | 2051.00 | 1728.00 | 1983.00 | 1920.67 | 2508.00 |
| ERF | 1344.00 | 1378.00 | 1432.00 | 1384.67 | 1566.00 |
| ERGIC1 | 7317.00 | 7140.00 | 7710.00 | 7389.00 | 8106.00 |
| ERGIC2 | 1812.00 | 1673.00 | 1771.00 | 1752.00 | 2045.00 |
| ERGIC3 | 5739.00 | 5589.00 | 5473.00 | 5600.33 | 6212.00 |
| ERH | 1639.00 | 1530.00 | 1592.00 | 1587.00 | 2593.00 |
| ERI1 | 570.00 | 494.00 | 606.00 | 556.67 | 652.00 |
| ERI2 | 487.00 | 439.00 | 502.00 | 476.00 | 591.00 |
| ERI3 | 1235.00 | 1109.00 | 1246.00 | 1196.67 | 1374.00 |
| ERI3-IT1 | 58.00 | 50.00 | 52.00 | 53.33 | 42.00 |

| | | | | | |
|---|---|---|---|---|---|
| ERICH1 | 225.00 | 245.00 | 228.00 | 232.67 | 259.00 |
| ERICH2 | 168.00 | 163.00 | 193.00 | 174.67 | 236.00 |
| ERICH3 | 5.00 | 1.00 | 1.00 | 2.33 | 1.00 |
| ERICH6 | 3.00 | 5.00 | 6.00 | 4.67 | 0.00 |
| ERICH6-AS1 | 55.00 | 49.00 | 45.00 | 49.67 | 47.00 |
| ERLEC1 | 1522.00 | 1419.00 | 1484.00 | 1475.00 | 1581.00 |
| ERLIN1 | 2735.00 | 2676.00 | 2663.00 | 2691.33 | 3384.00 |
| ERLIN2 | 4527.00 | 4040.00 | 4159.00 | 4242.00 | 3730.00 |
| ERMAP | 411.00 | 447.00 | 461.00 | 439.67 | 506.00 |
| ERMP1 | 1073.00 | 1082.00 | 1177.00 | 1110.67 | 1074.00 |
| ERN1 | 1096.00 | 870.00 | 1052.00 | 1006.00 | 570.00 |
| ERO1A | 3331.00 | 3397.00 | 3503.00 | 3410.33 | 4048.00 |
| ERO1B | 325.00 | 273.00 | 316.00 | 304.67 | 266.00 |
| ERP29 | 5643.00 | 6090.00 | 5486.00 | 5739.67 | 7218.00 |
| ERP44 | 1747.00 | 1728.00 | 1757.00 | 1744.00 | 1964.00 |
| ERRFI1 | 17530.00 | 16098.00 | 18762.00 | 17463.33 | 17507.00 |
| ERV3-1 | 28.00 | 21.00 | 28.00 | 25.67 | 22.00 |
| ERVK13-1 | 244.00 | 220.00 | 239.00 | 234.33 | 216.00 |
| ERVMER34-1 | 10.00 | 13.00 | 14.00 | 12.33 | 15.00 |
| ESAM | 194.00 | 196.00 | 213.00 | 201.00 | 118.00 |
| ESCO1 | 364.00 | 346.00 | 366.00 | 358.67 | 374.00 |
| ESCO2 | 79.00 | 68.00 | 81.00 | 76.00 | 105.00 |
| ESD | 1533.00 | 1506.00 | 1583.00 | 1540.67 | 1985.00 |
| ESF1 | 1381.00 | 1375.00 | 1563.00 | 1439.67 | 1573.00 |
| ESM1 | 18740.00 | 17505.00 | 17922.00 | 18055.67 | 27557.00 |
| ESPL1 | 1588.00 | 1503.00 | 1535.00 | 1542.00 | 1854.00 |
| ESPNL | 7.00 | 5.00 | 3.00 | 5.00 | 10.00 |
| ESR1 | 16.00 | 16.00 | 16.00 | 16.00 | 23.00 |
| ESR2 | 21.00 | 28.00 | 18.00 | 22.33 | 22.00 |
| ESRP1 | 14.00 | 11.00 | 14.00 | 13.00 | 7.00 |
| ESRP2 | 11.00 | 13.00 | 14.00 | 12.67 | 25.00 |
| ESRRA | 2157.00 | 2125.00 | 2405.00 | 2229.00 | 2333.00 |
| ESRRG | 10.00 | 9.00 | 10.00 | 9.67 | 17.00 |
| ESYT1 | 4239.00 | 4108.00 | 4045.00 | 4130.67 | 3628.00 |
| ESYT2 | 4426.00 | 4005.00 | 4393.00 | 4274.67 | 4368.00 |
| ESYT3 | 3.00 | 6.00 | 5.00 | 4.67 | 1.00 |
| ETAA1 | 504.00 | 513.00 | 594.00 | 537.00 | 561.00 |
| ETF1 | 3774.00 | 3461.00 | 3919.00 | 3718.00 | 4741.00 |
| ETFA | 1665.00 | 1507.00 | 1733.00 | 1635.00 | 1853.00 |
| ETFB | 607.00 | 688.00 | 697.00 | 664.00 | 740.00 |
| ETFDH | 383.00 | 372.00 | 395.00 | 383.33 | 427.00 |
| ETHE1 | 476.00 | 490.00 | 548.00 | 504.67 | 588.00 |
| ETNK1 | 2411.00 | 2173.00 | 2761.00 | 2448.33 | 2413.00 |
| ETS1 | 5210.00 | 4538.00 | 5300.00 | 5016.00 | 5139.00 |
| ETS2 | 1204.00 | 1164.00 | 1355.00 | 1241.00 | 1323.00 |
| ETV1 | 1008.00 | 883.00 | 1008.00 | 966.33 | 1053.00 |
| ETV2 | 9.00 | 4.00 | 5.00 | 6.00 | 9.00 |
| ETV3 | 195.00 | 144.00 | 185.00 | 174.67 | 123.00 |

| | | | | | |
|---|---|---|---|---|---|
| ETV4 | 770.00 | 710.00 | 837.00 | 772.33 | 947.00 |
| ETV5 | 2784.00 | 2548.00 | 2484.00 | 2605.33 | 3113.00 |
| ETV6 | 698.00 | 655.00 | 733.00 | 695.33 | 663.00 |
| EVA1A | 1465.00 | 1385.00 | 1405.00 | 1418.33 | 2025.00 |
| EVA1B | 319.00 | 288.00 | 389.00 | 332.00 | 355.00 |
| EVA1C | 65.00 | 65.00 | 74.00 | 68.00 | 72.00 |
| EVC | 5866.00 | 5443.00 | 6306.00 | 5871.67 | 3473.00 |
| EVI2A | 763.00 | 716.00 | 681.00 | 720.00 | 815.00 |
| EVI2B | 236.00 | 228.00 | 216.00 | 226.67 | 261.00 |
| EVI5 | 753.00 | 692.00 | 748.00 | 731.00 | 760.00 |
| EVI5L | 573.00 | 542.00 | 542.00 | 552.33 | 633.00 |
| EVL | 458.00 | 445.00 | 462.00 | 455.00 | 545.00 |
| EWSR1 | 7941.00 | 7877.00 | 8059.00 | 7959.00 | 9371.00 |
| EXD1 | 8.00 | 4.00 | 3.00 | 5.00 | 8.00 |
| EXD2 | 801.00 | 846.00 | 866.00 | 837.67 | 741.00 |
| EXD3 | 434.00 | 493.00 | 511.00 | 479.33 | 401.00 |
| EXO1 | 261.00 | 207.00 | 273.00 | 247.00 | 349.00 |
| EXO5 | 81.00 | 96.00 | 85.00 | 87.33 | 89.00 |
| EXOC1 | 976.00 | 824.00 | 1005.00 | 935.00 | 1014.00 |
| EXOC2 | 1377.00 | 1414.00 | 1445.00 | 1412.00 | 1357.00 |
| EXOC3 | 1158.00 | 1164.00 | 1246.00 | 1189.33 | 1096.00 |
| EXOC3-AS1 | 86.00 | 76.00 | 57.00 | 73.00 | 95.00 |
| EXOC4 | 1870.00 | 1826.00 | 1975.00 | 1890.33 | 1822.00 |
| EXOC5 | 947.00 | 854.00 | 1007.00 | 936.00 | 988.00 |
| EXOC6 | 277.00 | 257.00 | 320.00 | 284.67 | 299.00 |
| EXOC6B | 201.00 | 203.00 | 239.00 | 214.33 | 143.00 |
| EXOC7 | 2446.00 | 2433.00 | 2489.00 | 2456.00 | 2113.00 |
| EXOC8 | 593.00 | 574.00 | 622.00 | 596.33 | 539.00 |
| EXOG | 612.00 | 609.00 | 665.00 | 628.67 | 627.00 |
| EXOSC1 | 701.00 | 691.00 | 768.00 | 720.00 | 963.00 |
| EXOSC10 | 2294.00 | 2264.00 | 2462.00 | 2340.00 | 2365.00 |
| EXOSC2 | 1133.00 | 1087.00 | 1147.00 | 1122.33 | 1265.00 |
| EXOSC3 | 799.00 | 684.00 | 837.00 | 773.33 | 1042.00 |
| EXOSC4 | 446.00 | 479.00 | 490.00 | 471.67 | 579.00 |
| EXOSC5 | 572.00 | 562.00 | 563.00 | 565.67 | 670.00 |
| EXOSC6 | 347.00 | 326.00 | 340.00 | 337.67 | 383.00 |
| EXOSC7 | 834.00 | 918.00 | 902.00 | 884.67 | 1195.00 |
| EXOSC8 | 1116.00 | 1079.00 | 1123.00 | 1106.00 | 1535.00 |
| EXOSC9 | 616.00 | 593.00 | 587.00 | 598.67 | 692.00 |
| EXPH5 | 142.00 | 117.00 | 145.00 | 134.67 | 116.00 |
| EXT1 | 5693.00 | 5678.00 | 5852.00 | 5741.00 | 5119.00 |
| EXT2 | 3520.00 | 3607.00 | 3706.00 | 3611.00 | 3266.00 |
| EXTL2 | 2726.00 | 2570.00 | 2544.00 | 2613.33 | 3001.00 |
| EXTL3 | 2033.00 | 2024.00 | 2044.00 | 2033.67 | 1680.00 |
| EXTL3-AS1 | 13.00 | 7.00 | 3.00 | 7.67 | 4.00 |
| EYA1 | 12.00 | 7.00 | 2.00 | 7.00 | 7.00 |
| EYA3 | 769.00 | 689.00 | 741.00 | 733.00 | 713.00 |
| EYA4 | 901.00 | 852.00 | 988.00 | 913.67 | 876.00 |

| | | | | | |
|---|---|---|---|---|---|
| EYS | 8.00 | 3.00 | 8.00 | 6.33 | 16.00 |
| EZH1 | 2354.00 | 2535.00 | 2728.00 | 2539.00 | 2465.00 |
| EZH2 | 856.00 | 830.00 | 815.00 | 833.67 | 1280.00 |
| EZR | 2696.00 | 2750.00 | 2739.00 | 2728.33 | 3294.00 |
| F11-AS1 | 5.00 | 23.00 | 20.00 | 16.00 | 17.00 |
| F11R | 107.00 | 122.00 | 108.00 | 112.33 | 149.00 |
| F12 | 69.00 | 60.00 | 86.00 | 71.67 | 80.00 |
| F2R | 4246.00 | 3891.00 | 4365.00 | 4167.33 | 2485.00 |
| F2RL1 | 1089.00 | 997.00 | 1117.00 | 1067.67 | 475.00 |
| F2RL2 | 151.00 | 155.00 | 185.00 | 163.67 | 176.00 |
| F3 | 759.00 | 758.00 | 797.00 | 771.33 | 561.00 |
| F8 | 100.00 | 71.00 | 89.00 | 86.67 | 107.00 |
| F8A1 | 30.00 | 40.00 | 58.00 | 42.67 | 58.00 |
| FAAH | 15.00 | 9.00 | 19.00 | 14.33 | 10.00 |
| FAAHP1 | 3.00 | 0.00 | 1.00 | 1.33 | 0.00 |
| FAAP100 | 957.00 | 887.00 | 900.00 | 914.67 | 1079.00 |
| FAAP20 | 1202.00 | 1229.00 | 1320.00 | 1250.33 | 1306.00 |
| FAAP24 | 57.00 | 62.00 | 69.00 | 62.67 | 67.00 |
| FABP5 | 437.00 | 466.00 | 431.00 | 444.67 | 1323.00 |
| FABP6 | 10.00 | 12.00 | 20.00 | 14.00 | 58.00 |
| FADD | 3327.00 | 3311.00 | 3464.00 | 3367.33 | 4281.00 |
| FADS1 | 3668.00 | 3460.00 | 3573.00 | 3567.00 | 3566.00 |
| FADS2 | 1083.00 | 1067.00 | 1029.00 | 1059.67 | 1022.00 |
| FADS3 | 713.00 | 845.00 | 819.00 | 792.33 | 550.00 |
| FAF1 | 1520.00 | 1399.00 | 1530.00 | 1483.00 | 2058.00 |
| FAF2 | 2771.00 | 2437.00 | 2547.00 | 2585.00 | 2592.00 |
| FAH | 1153.00 | 1046.00 | 1134.00 | 1111.00 | 1273.00 |
| FAHD1 | 393.00 | 423.00 | 470.00 | 428.67 | 466.00 |
| FAHD2A | 738.00 | 768.00 | 710.00 | 738.67 | 756.00 |
| FAHD2B | 289.00 | 301.00 | 310.00 | 300.00 | 365.00 |
| FAHD2CP | 57.00 | 94.00 | 94.00 | 81.67 | 98.00 |
| FAIM | 226.00 | 219.00 | 239.00 | 228.00 | 255.00 |
| FALEC | 8.00 | 20.00 | 8.00 | 12.00 | 6.00 |
| FAM101B | 580.00 | 555.00 | 625.00 | 586.67 | 784.00 |
| FAM102A | 916.00 | 855.00 | 936.00 | 902.33 | 795.00 |
| FAM102B | 223.00 | 204.00 | 226.00 | 217.67 | 271.00 |
| FAM103A1 | 157.00 | 134.00 | 200.00 | 163.67 | 185.00 |
| FAM104A | 661.00 | 701.00 | 751.00 | 704.33 | 745.00 |
| FAM104B | 213.00 | 171.00 | 175.00 | 186.33 | 330.00 |
| FAM105A | 60.00 | 59.00 | 55.00 | 58.00 | 33.00 |
| FAM106A | 3.00 | 5.00 | 4.00 | 4.00 | 3.00 |
| FAM106B | 5.00 | 5.00 | 5.00 | 5.00 | 5.00 |
| FAM106CP | 45.00 | 47.00 | 58.00 | 50.00 | 26.00 |
| FAM107A | 9.00 | 6.00 | 4.00 | 6.33 | 6.00 |
| FAM107B | 1099.00 | 901.00 | 1162.00 | 1054.00 | 824.00 |
| FAM109A | 293.00 | 292.00 | 286.00 | 290.33 | 215.00 |
| FAM109B | 98.00 | 136.00 | 88.00 | 107.33 | 93.00 |
| FAM110A | 147.00 | 149.00 | 115.00 | 137.00 | 204.00 |

| | | | | | |
|---|---|---|---|---|---|
| FAM111A | 577.00 | 510.00 | 566.00 | 551.00 | 778.00 |
| FAM111B | 82.00 | 75.00 | 69.00 | 75.33 | 152.00 |
| FAM114A1 | 1404.00 | 1348.00 | 1453.00 | 1401.67 | 1331.00 |
| FAM114A2 | 550.00 | 519.00 | 558.00 | 542.33 | 685.00 |
| FAM117A | 7.00 | 8.00 | 7.00 | 7.33 | 8.00 |
| FAM117B | 179.00 | 175.00 | 214.00 | 189.33 | 221.00 |
| FAM118A | 408.00 | 347.00 | 404.00 | 386.33 | 359.00 |
| FAM118B | 259.00 | 243.00 | 247.00 | 249.67 | 283.00 |
| FAM120A | 4526.00 | 4261.00 | 4935.00 | 4574.00 | 4603.00 |
| FAM120AOS | 312.00 | 336.00 | 344.00 | 330.67 | 434.00 |
| FAM120C | 409.00 | 362.00 | 425.00 | 398.67 | 411.00 |
| FAM122A | 278.00 | 294.00 | 364.00 | 312.00 | 276.00 |
| FAM122B | 987.00 | 1029.00 | 1031.00 | 1015.67 | 1135.00 |
| FAM122C | 24.00 | 30.00 | 24.00 | 26.00 | 37.00 |
| FAM124A | 23.00 | 25.00 | 33.00 | 27.00 | 40.00 |
| FAM126A | 1120.00 | 1000.00 | 1153.00 | 1091.00 | 1351.00 |
| FAM126B | 366.00 | 364.00 | 392.00 | 374.00 | 350.00 |
| FAM127A | 584.00 | 625.00 | 584.00 | 597.67 | 622.00 |
| FAM127B | 504.00 | 502.00 | 523.00 | 509.67 | 532.00 |
| FAM127C | 169.00 | 163.00 | 173.00 | 168.33 | 162.00 |
| FAM129A | 1420.00 | 1256.00 | 1396.00 | 1357.33 | 1616.00 |
| FAM129B | 12296.00 | 12109.00 | 12357.00 | 12254.00 | 11590.00 |
| FAM129C | 4.00 | 3.00 | 6.00 | 4.33 | 7.00 |
| FAM131A | 771.00 | 837.00 | 788.00 | 798.67 | 863.00 |
| FAM131B | 28.00 | 28.00 | 34.00 | 30.00 | 31.00 |
| FAM131C | 14.00 | 15.00 | 11.00 | 13.33 | 9.00 |
| FAM132A | 75.00 | 81.00 | 89.00 | 81.67 | 77.00 |
| FAM132B | 93.00 | 103.00 | 60.00 | 85.33 | 73.00 |
| FAM133A | 48.00 | 61.00 | 73.00 | 60.67 | 114.00 |
| FAM133B | 324.00 | 253.00 | 300.00 | 292.33 | 355.00 |
| FAM133CP | 12.00 | 12.00 | 13.00 | 12.33 | 24.00 |
| FAM133DP | 9.00 | 18.00 | 16.00 | 14.33 | 9.00 |
| FAM134A | 2468.00 | 2431.00 | 2453.00 | 2450.67 | 2563.00 |
| FAM134B | 37.00 | 39.00 | 41.00 | 39.00 | 43.00 |
| FAM134C | 1198.00 | 1131.00 | 1213.00 | 1180.67 | 1156.00 |
| FAM135A | 429.00 | 374.00 | 493.00 | 432.00 | 418.00 |
| FAM136A | 1213.00 | 1192.00 | 1293.00 | 1232.67 | 1868.00 |
| FAM138B | 14.00 | 12.00 | 5.00 | 10.33 | 5.00 |
| FAM138E | 3.00 | 6.00 | 0.00 | 3.00 | 6.00 |
| FAM13A | 409.00 | 369.00 | 462.00 | 413.33 | 410.00 |
| FAM13A-AS1 | 27.00 | 40.00 | 49.00 | 38.67 | 30.00 |
| FAM13B | 859.00 | 793.00 | 839.00 | 830.33 | 797.00 |
| FAM149A | 19.00 | 14.00 | 20.00 | 17.67 | 29.00 |
| FAM149B1 | 507.00 | 449.00 | 486.00 | 480.67 | 558.00 |
| FAM151B | 25.00 | 25.00 | 21.00 | 23.67 | 23.00 |
| FAM155A | 269.00 | 258.00 | 305.00 | 277.33 | 276.00 |
| FAM156A | 21.00 | 12.00 | 26.00 | 19.67 | 28.00 |
| FAM160A1 | 36.00 | 32.00 | 26.00 | 31.33 | 20.00 |

| | | | | | |
|---|---|---|---|---|---|
| FAM160A2 | 926.00 | 913.00 | 1067.00 | 968.67 | 864.00 |
| FAM160B1 | 605.00 | 564.00 | 747.00 | 638.67 | 607.00 |
| FAM160B2 | 1164.00 | 1144.00 | 1185.00 | 1164.33 | 1132.00 |
| FAM161A | 323.00 | 345.00 | 352.00 | 340.00 | 316.00 |
| FAM161B | 126.00 | 92.00 | 117.00 | 111.67 | 109.00 |
| FAM162A | 619.00 | 676.00 | 714.00 | 669.67 | 1070.00 |
| FAM166B | 19.00 | 13.00 | 21.00 | 17.67 | 11.00 |
| FAM167A | 536.00 | 508.00 | 630.00 | 558.00 | 585.00 |
| FAM167B | 19.00 | 17.00 | 27.00 | 21.00 | 10.00 |
| FAM168A | 1291.00 | 1270.00 | 1282.00 | 1281.00 | 1276.00 |
| FAM168B | 3988.00 | 3727.00 | 4303.00 | 4006.00 | 4236.00 |
| FAM169A | 109.00 | 99.00 | 125.00 | 111.00 | 126.00 |
| FAM171A1 | 6236.00 | 6098.00 | 6370.00 | 6234.67 | 5681.00 |
| FAM171A2 | 154.00 | 127.00 | 146.00 | 142.33 | 124.00 |
| FAM171B | 179.00 | 161.00 | 197.00 | 179.00 | 156.00 |
| FAM172A | 619.00 | 604.00 | 691.00 | 638.00 | 705.00 |
| FAM172BP | 11.00 | 9.00 | 11.00 | 10.33 | 6.00 |
| FAM173A | 315.00 | 304.00 | 312.00 | 310.33 | 307.00 |
| FAM173B | 281.00 | 296.00 | 288.00 | 288.33 | 413.00 |
| FAM174A | 240.00 | 212.00 | 250.00 | 234.00 | 273.00 |
| FAM175A | 525.00 | 529.00 | 578.00 | 544.00 | 552.00 |
| FAM175B | 608.00 | 580.00 | 628.00 | 605.33 | 694.00 |
| FAM177A1 | 370.00 | 363.00 | 423.00 | 385.33 | 459.00 |
| FAM179A | 13.00 | 3.00 | 13.00 | 9.67 | 9.00 |
| FAM179B | 463.00 | 408.00 | 479.00 | 450.00 | 345.00 |
| FAM180A | 25.00 | 20.00 | 21.00 | 22.00 | 27.00 |
| FAM182B | 26.00 | 14.00 | 34.00 | 24.67 | 17.00 |
| FAM184A | 169.00 | 126.00 | 159.00 | 151.33 | 179.00 |
| FAM185A | 123.00 | 122.00 | 144.00 | 129.67 | 114.00 |
| FAM186B | 9.00 | 16.00 | 21.00 | 15.33 | 15.00 |
| FAM188A | 591.00 | 592.00 | 620.00 | 601.00 | 776.00 |
| FAM188B | 8.00 | 6.00 | 8.00 | 7.33 | 7.00 |
| FAM189B | 3.00 | 8.00 | 7.00 | 6.00 | 1.00 |
| FAM192A | 1914.00 | 1963.00 | 2310.00 | 2062.33 | 2155.00 |
| FAM193A | 601.00 | 544.00 | 614.00 | 586.33 | 588.00 |
| FAM193B | 1378.00 | 1456.00 | 1430.00 | 1421.33 | 1406.00 |
| FAM195A | 412.00 | 468.00 | 526.00 | 468.67 | 612.00 |
| FAM195B | 1630.00 | 1630.00 | 1632.00 | 1630.67 | 2004.00 |
| FAM196B | 1009.00 | 871.00 | 1071.00 | 983.67 | 696.00 |
| FAM199X | 1817.00 | 1700.00 | 2088.00 | 1868.33 | 1757.00 |
| FAM19A1 | 3.00 | 1.00 | 8.00 | 4.00 | 3.00 |
| FAM19A2 | 12.00 | 17.00 | 21.00 | 16.67 | 28.00 |
| FAM200A | 215.00 | 206.00 | 217.00 | 212.67 | 233.00 |
| FAM200B | 597.00 | 550.00 | 596.00 | 581.00 | 507.00 |
| FAM201A | 248.00 | 245.00 | 245.00 | 246.00 | 265.00 |
| FAM204A | 631.00 | 599.00 | 707.00 | 645.67 | 855.00 |
| FAM206A | 483.00 | 455.00 | 500.00 | 479.33 | 598.00 |
| FAM207A | 521.00 | 485.00 | 546.00 | 517.33 | 676.00 |

| | | | | | |
|---|---|---|---|---|---|
| FAM208A | 2769.00 | 2487.00 | 2962.00 | 2739.33 | 2811.00 |
| FAM208B | 4625.00 | 4172.00 | 4861.00 | 4552.67 | 4110.00 |
| FAM20B | 1520.00 | 1399.00 | 1531.00 | 1483.33 | 1572.00 |
| FAM20C | 2348.00 | 2591.00 | 2651.00 | 2530.00 | 1641.00 |
| FAM210A | 463.00 | 462.00 | 530.00 | 485.00 | 591.00 |
| FAM210B | 3041.00 | 2965.00 | 3235.00 | 3080.33 | 3803.00 |
| FAM212A | 3.00 | 0.00 | 1.00 | 1.33 | 3.00 |
| FAM212B | 30.00 | 36.00 | 39.00 | 35.00 | 17.00 |
| FAM212B-AS1 | 4.00 | 3.00 | 4.00 | 3.67 | 0.00 |
| FAM213A | 1715.00 | 1770.00 | 1705.00 | 1730.00 | 1782.00 |
| FAM214A | 348.00 | 363.00 | 368.00 | 359.67 | 273.00 |
| FAM214B | 670.00 | 653.00 | 707.00 | 676.67 | 615.00 |
| FAM216A | 735.00 | 780.00 | 852.00 | 789.00 | 1054.00 |
| FAM217B | 408.00 | 380.00 | 460.00 | 416.00 | 478.00 |
| FAM218A | 21.00 | 34.00 | 31.00 | 28.67 | 29.00 |
| FAM219A | 796.00 | 722.00 | 817.00 | 778.33 | 812.00 |
| FAM219B | 2173.00 | 2132.00 | 2345.00 | 2216.67 | 2011.00 |
| FAM21A | 2026.00 | 2051.00 | 2202.00 | 2093.00 | 1908.00 |
| FAM21C | 1686.00 | 1677.00 | 1795.00 | 1719.33 | 1543.00 |
| FAM21EP | 9.00 | 13.00 | 6.00 | 9.33 | 5.00 |
| FAM220A | 449.00 | 372.00 | 492.00 | 437.67 | 528.00 |
| FAM221A | 107.00 | 78.00 | 94.00 | 93.00 | 110.00 |
| FAM222A | 10.00 | 4.00 | 5.00 | 6.33 | 6.00 |
| FAM222B | 344.00 | 307.00 | 355.00 | 335.33 | 393.00 |
| FAM225A | 13.00 | 12.00 | 22.00 | 15.67 | 24.00 |
| FAM225B | 3.00 | 12.00 | 2.00 | 5.67 | 4.00 |
| FAM227A | 311.00 | 335.00 | 323.00 | 323.00 | 286.00 |
| FAM227B | 46.00 | 47.00 | 47.00 | 46.67 | 43.00 |
| FAM228B | 133.00 | 173.00 | 171.00 | 159.00 | 166.00 |
| FAM229A | 82.00 | 92.00 | 96.00 | 90.00 | 79.00 |
| FAM229B | 182.00 | 154.00 | 182.00 | 172.67 | 216.00 |
| FAM24B | 311.00 | 246.00 | 295.00 | 284.00 | 345.00 |
| FAM26E | 83.00 | 101.00 | 75.00 | 86.33 | 126.00 |
| FAM26F | 46.00 | 56.00 | 61.00 | 54.33 | 98.00 |
| FAM27B | 14.00 | 19.00 | 10.00 | 14.33 | 17.00 |
| FAM27E3 | 34.00 | 29.00 | 41.00 | 34.67 | 50.00 |
| FAM32A | 883.00 | 943.00 | 923.00 | 916.33 | 1105.00 |
| FAM35A | 667.00 | 598.00 | 731.00 | 665.33 | 735.00 |
| FAM35BP | 16.00 | 15.00 | 22.00 | 17.67 | 22.00 |
| FAM35DP | 5.00 | 17.00 | 11.00 | 11.00 | 18.00 |
| FAM3A | 844.00 | 837.00 | 797.00 | 826.00 | 836.00 |
| FAM3C | 1209.00 | 1229.00 | 1310.00 | 1249.33 | 1614.00 |
| FAM43A | 483.00 | 467.00 | 555.00 | 501.67 | 458.00 |
| FAM45A | 319.00 | 304.00 | 385.00 | 336.00 | 416.00 |
| FAM45B | 67.00 | 63.00 | 49.00 | 59.67 | 66.00 |
| FAM46A | 223.00 | 205.00 | 276.00 | 234.67 | 195.00 |
| FAM46B | 37.00 | 30.00 | 29.00 | 32.00 | 28.00 |
| FAM46D | 5.00 | 3.00 | 8.00 | 5.33 | 5.00 |

| | | | | | |
|---|---|---|---|---|---|
| FAM47E | 8.00 | 2.00 | 1.00 | 3.67 | 1.00 |
| FAM49A | 12.00 | 21.00 | 13.00 | 15.33 | 6.00 |
| FAM49B | 1306.00 | 1255.00 | 1317.00 | 1292.67 | 1665.00 |
| FAM50A | 2128.00 | 2276.00 | 2331.00 | 2245.00 | 2397.00 |
| FAM50B | 175.00 | 169.00 | 168.00 | 170.67 | 267.00 |
| FAM53A | 121.00 | 129.00 | 167.00 | 139.00 | 193.00 |
| FAM53B | 529.00 | 580.00 | 635.00 | 581.33 | 632.00 |
| FAM53C | 1293.00 | 1145.00 | 1312.00 | 1250.00 | 1150.00 |
| FAM57A | 918.00 | 952.00 | 893.00 | 921.00 | 988.00 |
| FAM58A | 206.00 | 214.00 | 272.00 | 230.67 | 349.00 |
| FAM60A | 8.00 | 2.00 | 3.00 | 4.33 | 10.00 |
| FAM63A | 236.00 | 238.00 | 230.00 | 234.67 | 226.00 |
| FAM63B | 140.00 | 110.00 | 170.00 | 140.00 | 108.00 |
| FAM64A | 243.00 | 212.00 | 184.00 | 213.00 | 292.00 |
| FAM65A | 2630.00 | 2586.00 | 2744.00 | 2653.33 | 2242.00 |
| FAM65C | 5.00 | 6.00 | 4.00 | 5.00 | 7.00 |
| FAM66C | 84.00 | 107.00 | 92.00 | 94.33 | 99.00 |
| FAM66D | 42.00 | 39.00 | 32.00 | 37.67 | 33.00 |
| FAM66E | 3.00 | 0.00 | 1.00 | 1.33 | 1.00 |
| FAM69A | 264.00 | 240.00 | 249.00 | 251.00 | 214.00 |
| FAM71E2 | 5.00 | 1.00 | 2.00 | 2.67 | 0.00 |
| FAM71F2 | 286.00 | 298.00 | 342.00 | 308.67 | 304.00 |
| FAM72A | 135.00 | 115.00 | 116.00 | 122.00 | 171.00 |
| FAM72B | 171.00 | 191.00 | 217.00 | 193.00 | 299.00 |
| FAM72C | 23.00 | 35.00 | 35.00 | 31.00 | 47.00 |
| FAM72D | 63.00 | 38.00 | 44.00 | 48.33 | 62.00 |
| FAM73A | 738.00 | 610.00 | 682.00 | 676.67 | 676.00 |
| FAM73B | 1480.00 | 1472.00 | 1442.00 | 1464.67 | 1402.00 |
| FAM76A | 157.00 | 138.00 | 172.00 | 155.67 | 160.00 |
| FAM76B | 695.00 | 675.00 | 739.00 | 703.00 | 762.00 |
| FAM78A | 54.00 | 73.00 | 99.00 | 75.33 | 83.00 |
| FAM78B | 4.00 | 4.00 | 3.00 | 3.67 | 4.00 |
| FAM81A | 139.00 | 134.00 | 175.00 | 149.33 | 210.00 |
| FAM83A | 4.00 | 3.00 | 3.00 | 3.33 | 1.00 |
| FAM83B | 9.00 | 11.00 | 11.00 | 10.33 | 13.00 |
| FAM83D | 604.00 | 521.00 | 598.00 | 574.33 | 771.00 |
| FAM83E | 4.00 | 0.00 | 4.00 | 2.67 | 5.00 |
| FAM83G | 662.00 | 653.00 | 759.00 | 691.33 | 573.00 |
| FAM84A | 50.00 | 38.00 | 51.00 | 46.33 | 107.00 |
| FAM84B | 345.00 | 286.00 | 359.00 | 330.00 | 291.00 |
| FAM86B1 | 5.00 | 3.00 | 6.00 | 4.67 | 1.00 |
| FAM86B2 | 5.00 | 5.00 | 2.00 | 4.00 | 3.00 |
| FAM86B3P | 448.00 | 416.00 | 449.00 | 437.67 | 352.00 |
| FAM86C1 | 512.00 | 498.00 | 508.00 | 506.00 | 554.00 |
| FAM86C2P | 88.00 | 78.00 | 79.00 | 81.67 | 111.00 |
| FAM86DP | 404.00 | 410.00 | 419.00 | 411.00 | 331.00 |
| FAM86EP | 245.00 | 205.00 | 218.00 | 222.67 | 238.00 |
| FAM86FP | 47.00 | 58.00 | 51.00 | 52.00 | 46.00 |

| | | | | | |
|---|---|---|---|---|---|
| FAM86HP | 165.00 | 197.00 | 160.00 | 174.00 | 149.00 |
| FAM86JP | 90.00 | 82.00 | 90.00 | 87.33 | 96.00 |
| FAM89A | 230.00 | 231.00 | 261.00 | 240.67 | 281.00 |
| FAM89B | 636.00 | 729.00 | 673.00 | 679.33 | 752.00 |
| FAM8A1 | 577.00 | 558.00 | 602.00 | 579.00 | 677.00 |
| FAM91A1 | 1733.00 | 1485.00 | 1836.00 | 1684.67 | 1764.00 |
| FAM92A1 | 683.00 | 693.00 | 729.00 | 701.67 | 823.00 |
| FAM95B1 | 12.00 | 8.00 | 10.00 | 10.00 | 8.00 |
| FAM95C | 496.00 | 493.00 | 589.00 | 526.00 | 532.00 |
| FAM96A | 785.00 | 793.00 | 819.00 | 799.00 | 1051.00 |
| FAM96B | 656.00 | 701.00 | 724.00 | 693.67 | 949.00 |
| FAM98A | 1597.00 | 1442.00 | 1576.00 | 1538.33 | 1737.00 |
| FAM98B | 870.00 | 825.00 | 878.00 | 857.67 | 1122.00 |
| FAM98C | 234.00 | 216.00 | 241.00 | 230.33 | 280.00 |
| FAN1 | 426.00 | 424.00 | 424.00 | 424.67 | 455.00 |
| FANCA | 1565.00 | 1579.00 | 1461.00 | 1535.00 | 1704.00 |
| FANCB | 91.00 | 64.00 | 70.00 | 75.00 | 72.00 |
| FANCC | 359.00 | 309.00 | 356.00 | 341.33 | 404.00 |
| FANCD2 | 898.00 | 882.00 | 813.00 | 864.33 | 1020.00 |
| FANCE | 283.00 | 307.00 | 319.00 | 303.00 | 367.00 |
| FANCF | 432.00 | 405.00 | 460.00 | 432.33 | 567.00 |
| FANCG | 976.00 | 1040.00 | 913.00 | 976.33 | 1286.00 |
| FANCI | 1287.00 | 1201.00 | 1159.00 | 1215.67 | 1721.00 |
| FANCL | 595.00 | 642.00 | 642.00 | 626.33 | 827.00 |
| FANCM | 300.00 | 280.00 | 309.00 | 296.33 | 306.00 |
| FANK1 | 18.00 | 9.00 | 12.00 | 13.00 | 5.00 |
| FAP | 209.00 | 176.00 | 200.00 | 195.00 | 191.00 |
| FAR1 | 1752.00 | 1579.00 | 1928.00 | 1753.00 | 1982.00 |
| FAR2 | 1268.00 | 1254.00 | 1363.00 | 1295.00 | 1161.00 |
| FAR2P2 | 3.00 | 0.00 | 1.00 | 1.33 | 1.00 |
| FARP1 | 680.00 | 722.00 | 693.00 | 698.33 | 789.00 |
| FARP2 | 203.00 | 191.00 | 203.00 | 199.00 | 159.00 |
| FARS2 | 196.00 | 185.00 | 212.00 | 197.67 | 288.00 |
| FARSA | 1728.00 | 1743.00 | 1795.00 | 1755.33 | 2016.00 |
| FARSB | 2623.00 | 2409.00 | 2834.00 | 2622.00 | 2902.00 |
| FAS | 134.00 | 127.00 | 101.00 | 120.67 | 141.00 |
| FASLG | 25.00 | 30.00 | 19.00 | 24.67 | 20.00 |
| FASN | 10535.00 | 10177.00 | 10780.00 | 10497.33 | 9370.00 |
| FASTK | 2398.00 | 2488.00 | 2432.00 | 2439.33 | 2793.00 |
| FASTKD1 | 1188.00 | 1180.00 | 1301.00 | 1223.00 | 1250.00 |
| FASTKD2 | 2120.00 | 1926.00 | 2222.00 | 2089.33 | 2016.00 |
| FASTKD3 | 433.00 | 446.00 | 459.00 | 446.00 | 479.00 |
| FASTKD5 | 557.00 | 581.00 | 540.00 | 559.33 | 616.00 |
| FAT1 | 2217.00 | 2052.00 | 2237.00 | 2168.67 | 1744.00 |
| FAT3 | 3.00 | 1.00 | 3.00 | 2.33 | 1.00 |
| FAT4 | 665.00 | 626.00 | 712.00 | 667.67 | 1217.00 |
| FAU | 8415.00 | 8625.00 | 8929.00 | 8656.33 | 10774.00 |
| FAXC | 54.00 | 47.00 | 59.00 | 53.33 | 64.00 |

| | | | | | |
|---|---|---|---|---|---|
| FAXDC2 | 14.00 | 22.00 | 19.00 | 18.33 | 13.00 |
| FBF1 | 667.00 | 664.00 | 738.00 | 689.67 | 743.00 |
| FBL | 2342.00 | 2432.00 | 2491.00 | 2421.67 | 3646.00 |
| FBLIM1 | 1556.00 | 1498.00 | 1642.00 | 1565.33 | 1130.00 |
| FBLN2 | 25.00 | 29.00 | 38.00 | 30.67 | 33.00 |
| FBLN7 | 35.00 | 30.00 | 38.00 | 34.33 | 20.00 |
| FBN1 | 2982.00 | 2666.00 | 3321.00 | 2989.67 | 1609.00 |
| FBN2 | 10489.00 | 9487.00 | 11144.00 | 10373.33 | 5081.00 |
| FBRS | 2306.00 | 2223.00 | 2438.00 | 2322.33 | 1933.00 |
| FBRSL1 | 1129.00 | 1057.00 | 1163.00 | 1116.33 | 1259.00 |
| FBXL12 | 461.00 | 439.00 | 457.00 | 452.33 | 508.00 |
| FBXL13 | 15.00 | 12.00 | 29.00 | 18.67 | 19.00 |
| FBXL14 | 369.00 | 342.00 | 406.00 | 372.33 | 393.00 |
| FBXL15 | 404.00 | 398.00 | 431.00 | 411.00 | 426.00 |
| FBXL17 | 596.00 | 475.00 | 599.00 | 556.67 | 538.00 |
| FBXL18 | 827.00 | 724.00 | 887.00 | 812.67 | 735.00 |
| FBXL19 | 1196.00 | 1314.00 | 1401.00 | 1303.67 | 1204.00 |
| FBXL19-AS1 | 784.00 | 679.00 | 809.00 | 757.33 | 509.00 |
| FBXL2 | 375.00 | 400.00 | 369.00 | 381.33 | 431.00 |
| FBXL20 | 487.00 | 468.00 | 537.00 | 497.33 | 465.00 |
| FBXL21 | 12.00 | 11.00 | 20.00 | 14.33 | 9.00 |
| FBXL22 | 8.00 | 6.00 | 3.00 | 5.67 | 9.00 |
| FBXL3 | 674.00 | 609.00 | 783.00 | 688.67 | 725.00 |
| FBXL4 | 682.00 | 607.00 | 666.00 | 651.67 | 800.00 |
| FBXL5 | 596.00 | 559.00 | 663.00 | 606.00 | 542.00 |
| FBXL6 | 1043.00 | 1095.00 | 1155.00 | 1097.67 | 1187.00 |
| FBXL7 | 203.00 | 192.00 | 237.00 | 210.67 | 245.00 |
| FBXL8 | 316.00 | 324.00 | 363.00 | 334.33 | 275.00 |
| FBXO10 | 562.00 | 481.00 | 502.00 | 515.00 | 490.00 |
| FBXO11 | 1464.00 | 1412.00 | 1654.00 | 1510.00 | 1787.00 |
| FBXO15 | 22.00 | 16.00 | 18.00 | 18.67 | 21.00 |
| FBXO16 | 37.00 | 42.00 | 43.00 | 40.67 | 44.00 |
| FBXO17 | 813.00 | 844.00 | 927.00 | 861.33 | 964.00 |
| FBXO18 | 2841.00 | 2712.00 | 2930.00 | 2827.67 | 2834.00 |
| FBXO21 | 580.00 | 472.00 | 534.00 | 528.67 | 472.00 |
| FBXO22 | 717.00 | 657.00 | 785.00 | 719.67 | 867.00 |
| FBXO24 | 14.00 | 12.00 | 11.00 | 12.33 | 21.00 |
| FBXO25 | 93.00 | 68.00 | 122.00 | 94.33 | 129.00 |
| FBXO28 | 976.00 | 877.00 | 1088.00 | 980.33 | 1060.00 |
| FBXO3 | 350.00 | 304.00 | 355.00 | 336.33 | 349.00 |
| FBXO30 | 1030.00 | 857.00 | 1080.00 | 989.00 | 962.00 |
| FBXO31 | 1557.00 | 1588.00 | 1639.00 | 1594.67 | 1663.00 |
| FBXO32 | 561.00 | 531.00 | 579.00 | 557.00 | 724.00 |
| FBXO33 | 222.00 | 254.00 | 281.00 | 252.33 | 291.00 |
| FBXO34 | 597.00 | 530.00 | 641.00 | 589.33 | 686.00 |
| FBXO36 | 72.00 | 89.00 | 105.00 | 88.67 | 79.00 |
| FBXO38 | 1063.00 | 1020.00 | 1158.00 | 1080.33 | 1065.00 |
| FBXO4 | 333.00 | 307.00 | 358.00 | 332.67 | 482.00 |

| | | | | | |
|---|---|---|---|---|---|
| FBXO41 | 905.00 | 976.00 | 974.00 | 951.67 | 917.00 |
| FBXO42 | 574.00 | 546.00 | 599.00 | 573.00 | 625.00 |
| FBXO43 | 203.00 | 203.00 | 184.00 | 196.67 | 254.00 |
| FBXO44 | 341.00 | 411.00 | 411.00 | 387.67 | 420.00 |
| FBXO45 | 617.00 | 536.00 | 646.00 | 599.67 | 682.00 |
| FBXO46 | 474.00 | 490.00 | 580.00 | 514.67 | 602.00 |
| FBXO48 | 66.00 | 58.00 | 76.00 | 66.67 | 45.00 |
| FBXO5 | 381.00 | 349.00 | 391.00 | 373.67 | 510.00 |
| FBXO6 | 8.00 | 11.00 | 17.00 | 12.00 | 32.00 |
| FBXO7 | 1175.00 | 1168.00 | 1270.00 | 1204.33 | 1295.00 |
| FBXO8 | 149.00 | 177.00 | 221.00 | 182.33 | 203.00 |
| FBXO9 | 1410.00 | 1240.00 | 1469.00 | 1373.00 | 1391.00 |
| FBXW11 | 1714.00 | 1496.00 | 1920.00 | 1710.00 | 1476.00 |
| FBXW2 | 1832.00 | 1755.00 | 1991.00 | 1859.33 | 1746.00 |
| FBXW4 | 759.00 | 755.00 | 790.00 | 768.00 | 859.00 |
| FBXW4P1 | 5.00 | 5.00 | 8.00 | 6.00 | 2.00 |
| FBXW5 | 1840.00 | 1827.00 | 2008.00 | 1891.67 | 2188.00 |
| FBXW7 | 287.00 | 306.00 | 324.00 | 305.67 | 334.00 |
| FBXW8 | 773.00 | 723.00 | 796.00 | 764.00 | 630.00 |
| FBXW9 | 283.00 | 244.00 | 284.00 | 270.33 | 272.00 |
| FCER1G | 29.00 | 23.00 | 13.00 | 21.67 | 18.00 |
| FCF1 | 877.00 | 806.00 | 923.00 | 868.67 | 1046.00 |
| FCGBP | 15.00 | 7.00 | 13.00 | 11.67 | 12.00 |
| FCGRT | 496.00 | 544.00 | 520.00 | 520.00 | 548.00 |
| FCHO1 | 9.00 | 18.00 | 20.00 | 15.67 | 5.00 |
| FCHO2 | 667.00 | 615.00 | 746.00 | 676.00 | 740.00 |
| FCHSD1 | 1065.00 | 1144.00 | 1173.00 | 1127.33 | 934.00 |
| FCHSD2 | 694.00 | 617.00 | 714.00 | 675.00 | 656.00 |
| FCMR | 20.00 | 19.00 | 18.00 | 19.00 | 8.00 |
| FCRLA | 9.00 | 13.00 | 14.00 | 12.00 | 3.00 |
| FCRLB | 105.00 | 82.00 | 96.00 | 94.33 | 103.00 |
| FDFT1 | 1179.00 | 1096.00 | 1175.00 | 1150.00 | 1171.00 |
| FDPS | 2313.00 | 2344.00 | 2365.00 | 2340.67 | 2944.00 |
| FDX1 | 581.00 | 531.00 | 585.00 | 565.67 | 654.00 |
| FDX1L | 367.00 | 388.00 | 371.00 | 375.33 | 402.00 |
| FDXACB1 | 182.00 | 182.00 | 205.00 | 189.67 | 264.00 |
| FDXR | 182.00 | 191.00 | 204.00 | 192.33 | 165.00 |
| FECH | 288.00 | 295.00 | 292.00 | 291.67 | 317.00 |
| FEM1A | 728.00 | 720.00 | 755.00 | 734.33 | 848.00 |
| FEM1B | 1048.00 | 1010.00 | 1286.00 | 1114.67 | 1145.00 |
| FEM1C | 670.00 | 594.00 | 700.00 | 654.67 | 571.00 |
| FEN1 | 826.00 | 783.00 | 818.00 | 809.00 | 1083.00 |
| FENDRR | 3.00 | 3.00 | 5.00 | 3.67 | 7.00 |
| FER | 690.00 | 604.00 | 757.00 | 683.67 | 672.00 |
| FER1L4 | 85.00 | 91.00 | 90.00 | 88.67 | 42.00 |
| FER1L6 | 15.00 | 21.00 | 17.00 | 17.67 | 1.00 |
| FERMT1 | 1139.00 | 1085.00 | 1276.00 | 1166.67 | 1343.00 |
| FERMT2 | 1738.00 | 1599.00 | 1916.00 | 1751.00 | 1605.00 |

| | | | | | |
|---|---|---|---|---|---|
| FERMT3 | 577.00 | 554.00 | 557.00 | 562.67 | 373.00 |
| FES | 8.00 | 6.00 | 5.00 | 6.33 | 4.00 |
| FEZ1 | 6.00 | 11.00 | 5.00 | 7.33 | 11.00 |
| FEZ2 | 2461.00 | 2147.00 | 2632.00 | 2413.33 | 3937.00 |
| FGD1 | 705.00 | 618.00 | 707.00 | 676.67 | 713.00 |
| FGD4 | 11.00 | 12.00 | 12.00 | 11.67 | 13.00 |
| FGD5-AS1 | 2055.00 | 1934.00 | 2190.00 | 2059.67 | 2406.00 |
| FGD6 | 348.00 | 328.00 | 344.00 | 340.00 | 474.00 |
| FGF1 | 108.00 | 78.00 | 84.00 | 90.00 | 213.00 |
| FGF11 | 25.00 | 20.00 | 14.00 | 19.67 | 10.00 |
| FGF12 | 65.00 | 75.00 | 68.00 | 69.33 | 99.00 |
| FGF13 | 98.00 | 76.00 | 118.00 | 97.33 | 129.00 |
| FGF18 | 16.00 | 6.00 | 10.00 | 10.67 | 4.00 |
| FGF2 | 881.00 | 696.00 | 926.00 | 834.33 | 659.00 |
| FGF20 | 3.00 | 1.00 | 4.00 | 2.67 | 4.00 |
| FGF4 | 37.00 | 39.00 | 51.00 | 42.33 | 36.00 |
| FGF5 | 2425.00 | 2167.00 | 2581.00 | 2391.00 | 2854.00 |
| FGFBP3 | 126.00 | 117.00 | 112.00 | 118.33 | 140.00 |
| FGFR1 | 4821.00 | 4946.00 | 5066.00 | 4944.33 | 3323.00 |
| FGFR1OP | 929.00 | 909.00 | 953.00 | 930.33 | 1060.00 |
| FGFR1OP2 | 974.00 | 859.00 | 1022.00 | 951.67 | 1165.00 |
| FGFR4 | 13.00 | 6.00 | 8.00 | 9.00 | 11.00 |
| FGFRL1 | 478.00 | 534.00 | 573.00 | 528.33 | 465.00 |
| FGGY | 280.00 | 289.00 | 306.00 | 291.67 | 380.00 |
| FH | 1443.00 | 1455.00 | 1482.00 | 1460.00 | 1899.00 |
| FHAD1 | 38.00 | 30.00 | 29.00 | 32.33 | 32.00 |
| FHDC1 | 61.00 | 46.00 | 55.00 | 54.00 | 43.00 |
| FHIT | 98.00 | 77.00 | 98.00 | 91.00 | 81.00 |
| FHL1 | 1220.00 | 1085.00 | 1236.00 | 1180.33 | 1053.00 |
| FHL2 | 3630.00 | 3492.00 | 3995.00 | 3705.67 | 3467.00 |
| FHL3 | 245.00 | 246.00 | 277.00 | 256.00 | 280.00 |
| FHOD1 | 1488.00 | 1555.00 | 1589.00 | 1544.00 | 1692.00 |
| FHOD3 | 608.00 | 592.00 | 703.00 | 634.33 | 832.00 |
| FIBCD1 | 8.00 | 5.00 | 11.00 | 8.00 | 3.00 |
| FIBP | 1921.00 | 1788.00 | 1994.00 | 1901.00 | 2292.00 |
| FICD | 135.00 | 121.00 | 138.00 | 131.33 | 113.00 |
| FIG4 | 704.00 | 655.00 | 733.00 | 697.33 | 645.00 |
| FIGN | 481.00 | 460.00 | 520.00 | 487.00 | 401.00 |
| FIGNL1 | 1056.00 | 938.00 | 984.00 | 992.67 | 1026.00 |
| FIGNL2 | 12.00 | 8.00 | 13.00 | 11.00 | 4.00 |
| FIP1L1 | 1041.00 | 1039.00 | 1193.00 | 1091.00 | 1296.00 |
| FIRRE | 232.00 | 224.00 | 271.00 | 242.33 | 170.00 |
| FIS1 | 664.00 | 694.00 | 668.00 | 675.33 | 736.00 |
| FITM1 | 4.00 | 3.00 | 5.00 | 4.00 | 3.00 |
| FITM2 | 82.00 | 88.00 | 97.00 | 89.00 | 74.00 |
| FIZ1 | 223.00 | 216.00 | 256.00 | 231.67 | 299.00 |
| FJX1 | 3734.00 | 3695.00 | 3842.00 | 3757.00 | 4373.00 |
| FKBP10 | 14626.00 | 14924.00 | 14739.00 | 14763.00 | 12942.00 |

| | | | | | |
|---|---|---|---|---|---|
| FKBP11 | 2091.00 | 2088.00 | 1963.00 | 2047.33 | 2334.00 |
| FKBP14 | 1326.00 | 1211.00 | 1341.00 | 1292.67 | 1133.00 |
| FKBP15 | 1699.00 | 1655.00 | 1805.00 | 1719.67 | 1659.00 |
| FKBP1A | 7545.00 | 7037.00 | 7201.00 | 7261.00 | 7881.00 |
| FKBP1AP1 | 37.00 | 41.00 | 39.00 | 39.00 | 63.00 |
| FKBP1B | 10.00 | 13.00 | 5.00 | 9.33 | 16.00 |
| FKBP2 | 1483.00 | 1680.00 | 1514.00 | 1559.00 | 1840.00 |
| FKBP3 | 1421.00 | 1410.00 | 1529.00 | 1453.33 | 1832.00 |
| FKBP4 | 3526.00 | 3337.00 | 3732.00 | 3531.67 | 4309.00 |
| FKBP5 | 436.00 | 380.00 | 456.00 | 424.00 | 729.00 |
| FKBP7 | 545.00 | 492.00 | 505.00 | 514.00 | 566.00 |
| FKBP8 | 3591.00 | 3528.00 | 4180.00 | 3766.33 | 4122.00 |
| FKBP9 | 1909.00 | 1983.00 | 1936.00 | 1942.67 | 1751.00 |
| FKBP9P1 | 42.00 | 40.00 | 32.00 | 38.00 | 33.00 |
| FKRP | 959.00 | 835.00 | 910.00 | 901.33 | 762.00 |
| FKTN | 1822.00 | 1593.00 | 1806.00 | 1740.33 | 1629.00 |
| FLAD1 | 1181.00 | 1154.00 | 1232.00 | 1189.00 | 1279.00 |
| FLCN | 1215.00 | 1220.00 | 1289.00 | 1241.33 | 1161.00 |
| FLG-AS1 | 378.00 | 366.00 | 332.00 | 358.67 | 370.00 |
| FLI1 | 832.00 | 852.00 | 948.00 | 877.33 | 932.00 |
| FLII | 4070.00 | 4053.00 | 4351.00 | 4158.00 | 3654.00 |
| FLJ10038 | 298.00 | 285.00 | 312.00 | 298.33 | 322.00 |
| FLJ20021 | 28.00 | 37.00 | 41.00 | 35.33 | 33.00 |
| FLJ21408 | 32.00 | 25.00 | 27.00 | 28.00 | 17.00 |
| FLJ22447 | 123.00 | 80.00 | 96.00 | 99.67 | 46.00 |
| FLJ23867 | 1071.00 | 1133.00 | 1063.00 | 1089.00 | 709.00 |
| FLJ27354 | 31.00 | 20.00 | 28.00 | 26.33 | 17.00 |
| FLJ30679 | 3.00 | 3.00 | 2.00 | 2.67 | 1.00 |
| FLJ31104 | 28.00 | 23.00 | 35.00 | 28.67 | 33.00 |
| FLJ31356 | 11.00 | 13.00 | 16.00 | 13.33 | 13.00 |
| FLJ32255 | 36.00 | 24.00 | 38.00 | 32.67 | 31.00 |
| FLJ37035 | 35.00 | 27.00 | 35.00 | 32.33 | 34.00 |
| FLJ37201 | 70.00 | 67.00 | 68.00 | 68.33 | 78.00 |
| FLJ37453 | 74.00 | 87.00 | 97.00 | 86.00 | 68.00 |
| FLJ41200 | 87.00 | 108.00 | 97.00 | 97.33 | 80.00 |
| FLJ42351 | 105.00 | 80.00 | 95.00 | 93.33 | 103.00 |
| FLJ42627 | 123.00 | 120.00 | 126.00 | 123.00 | 93.00 |
| FLJ45513 | 14.00 | 30.00 | 34.00 | 26.00 | 35.00 |
| FLJ46906 | 282.00 | 308.00 | 325.00 | 305.00 | 376.00 |
| FLNA | 40919.00 | 39191.00 | 43079.00 | 41063.00 | 29995.00 |
| FLNB | 17827.00 | 16972.00 | 17902.00 | 17567.00 | 14733.00 |
| FLNC | 15968.00 | 15739.00 | 15940.00 | 15882.33 | 5783.00 |
| FLOT1 | 10.00 | 6.00 | 17.00 | 11.00 | 14.00 |
| FLOT2 | 849.00 | 844.00 | 856.00 | 849.67 | 907.00 |
| FLRT2 | 690.00 | 685.00 | 749.00 | 708.00 | 728.00 |
| FLRT3 | 13.00 | 16.00 | 24.00 | 17.67 | 6.00 |
| FLT3LG | 66.00 | 67.00 | 60.00 | 64.33 | 71.00 |
| FLVCR1 | 963.00 | 914.00 | 969.00 | 948.67 | 951.00 |

| | | | | | |
|---|---|---|---|---|---|
| FLVCR1-AS1 | 14.00 | 16.00 | 21.00 | 17.00 | 21.00 |
| FLVCR2 | 7.00 | 19.00 | 14.00 | 13.33 | 12.00 |
| FLYWCH1 | 2623.00 | 2645.00 | 2608.00 | 2625.33 | 2682.00 |
| FLYWCH2 | 387.00 | 367.00 | 404.00 | 386.00 | 446.00 |
| FMN1 | 491.00 | 422.00 | 498.00 | 470.33 | 324.00 |
| FMN2 | 211.00 | 159.00 | 218.00 | 196.00 | 278.00 |
| FMNL1 | 1708.00 | 1727.00 | 1722.00 | 1719.00 | 1008.00 |
| FMNL2 | 2855.00 | 2697.00 | 2978.00 | 2843.33 | 3222.00 |
| FMNL3 | 2404.00 | 2259.00 | 2540.00 | 2401.00 | 1879.00 |
| FMO4 | 60.00 | 70.00 | 58.00 | 62.67 | 84.00 |
| FMR1 | 722.00 | 661.00 | 709.00 | 697.33 | 734.00 |
| FN1 | 65121.00 | 59217.00 | 69496.00 | 64611.33 | 76876.00 |
| FN3K | 68.00 | 59.00 | 66.00 | 64.33 | 69.00 |
| FN3KRP | 593.00 | 539.00 | 606.00 | 579.33 | 640.00 |
| FNBP1 | 1122.00 | 946.00 | 1116.00 | 1061.33 | 1061.00 |
| FNBP1L | 736.00 | 695.00 | 786.00 | 739.00 | 831.00 |
| FNBP4 | 3482.00 | 3448.00 | 3796.00 | 3575.33 | 3229.00 |
| FNDC3A | 1564.00 | 1401.00 | 1729.00 | 1564.67 | 1683.00 |
| FNDC3B | 1606.00 | 1340.00 | 1600.00 | 1515.33 | 1310.00 |
| FNDC4 | 75.00 | 95.00 | 111.00 | 93.67 | 72.00 |
| FNDC7 | 5.00 | 2.00 | 0.00 | 2.33 | 3.00 |
| FNIP1 | 1174.00 | 1001.00 | 1216.00 | 1130.33 | 1043.00 |
| FNIP2 | 156.00 | 172.00 | 180.00 | 169.33 | 149.00 |
| FNTA | 1272.00 | 1249.00 | 1431.00 | 1317.33 | 1519.00 |
| FNTB | 23.00 | 24.00 | 21.00 | 22.67 | 32.00 |
| FOCAD | 1858.00 | 1725.00 | 1774.00 | 1785.67 | 1723.00 |
| FOLH1 | 4.00 | 4.00 | 3.00 | 3.67 | 3.00 |
| FOPNL | 48.00 | 51.00 | 48.00 | 49.00 | 68.00 |
| FOS | 29.00 | 45.00 | 38.00 | 37.33 | 32.00 |
| FOSB | 118.00 | 113.00 | 118.00 | 116.33 | 107.00 |
| FOSL1 | 14136.00 | 13800.00 | 14620.00 | 14185.33 | 14209.00 |
| FOSL2 | 2910.00 | 2939.00 | 3237.00 | 3028.67 | 2526.00 |
| FOXA1 | 24.00 | 29.00 | 24.00 | 25.67 | 28.00 |
| FOXA2 | 874.00 | 927.00 | 1019.00 | 940.00 | 735.00 |
| FOXC1 | 320.00 | 307.00 | 351.00 | 326.00 | 330.00 |
| FOXC2 | 118.00 | 105.00 | 123.00 | 115.33 | 164.00 |
| FOXC2-AS1 | 22.00 | 21.00 | 21.00 | 21.33 | 39.00 |
| FOXD1 | 2874.00 | 2938.00 | 3002.00 | 2938.00 | 4973.00 |
| FOXD2 | 9.00 | 13.00 | 11.00 | 11.00 | 19.00 |
| FOXD2-AS1 | 82.00 | 93.00 | 84.00 | 86.33 | 82.00 |
| FOXD3 | 36.00 | 41.00 | 31.00 | 36.00 | 30.00 |
| FOXD3-AS1 | 63.00 | 68.00 | 53.00 | 61.33 | 64.00 |
| FOXD4 | 21.00 | 20.00 | 29.00 | 23.33 | 14.00 |
| FOXD4L1 | 9.00 | 14.00 | 7.00 | 10.00 | 11.00 |
| FOXF1 | 232.00 | 233.00 | 267.00 | 244.00 | 540.00 |
| FOXF2 | 523.00 | 562.00 | 586.00 | 557.00 | 718.00 |
| FOXG1 | 110.00 | 114.00 | 111.00 | 111.67 | 129.00 |
| FOXH1 | 16.00 | 24.00 | 23.00 | 21.00 | 18.00 |

| | | | | | |
|---|---|---|---|---|---|
| FOXJ2 | 852.00 | 772.00 | 900.00 | 841.33 | 663.00 |
| FOXJ3 | 2658.00 | 2500.00 | 2811.00 | 2656.33 | 2497.00 |
| FOXK1 | 3164.00 | 2926.00 | 2476.00 | 2855.33 | 2285.00 |
| FOXK2 | 2599.00 | 2511.00 | 2705.00 | 2605.00 | 2865.00 |
| FOXL1 | 1337.00 | 1277.00 | 1423.00 | 1345.67 | 1458.00 |
| FOXM1 | 2326.00 | 2205.00 | 2113.00 | 2214.67 | 2661.00 |
| FOXN2 | 630.00 | 530.00 | 635.00 | 598.33 | 756.00 |
| FOXN3 | 697.00 | 622.00 | 771.00 | 696.67 | 592.00 |
| FOXN3-AS1 | 35.00 | 25.00 | 21.00 | 27.00 | 43.00 |
| FOXO1 | 169.00 | 180.00 | 200.00 | 183.00 | 184.00 |
| FOXO3 | 767.00 | 720.00 | 803.00 | 763.33 | 645.00 |
| FOXO3B | 309.00 | 281.00 | 317.00 | 302.33 | 313.00 |
| FOXO4 | 229.00 | 260.00 | 285.00 | 258.00 | 205.00 |
| FOXO6 | 3.00 | 4.00 | 11.00 | 6.00 | 0.00 |
| FOXP1 | 1620.00 | 1526.00 | 1818.00 | 1654.67 | 1731.00 |
| FOXP2 | 137.00 | 138.00 | 177.00 | 150.67 | 117.00 |
| FOXP3 | 3.00 | 0.00 | 1.00 | 1.33 | 0.00 |
| FOXP4 | 852.00 | 909.00 | 1007.00 | 922.67 | 921.00 |
| FOXP4-AS1 | 26.00 | 21.00 | 19.00 | 22.00 | 36.00 |
| FOXQ1 | 4.00 | 2.00 | 1.00 | 2.33 | 4.00 |
| FOXRED1 | 726.00 | 726.00 | 798.00 | 750.00 | 748.00 |
| FOXRED2 | 1019.00 | 1010.00 | 1021.00 | 1016.67 | 980.00 |
| FPGS | 1146.00 | 1207.00 | 1194.00 | 1182.33 | 1174.00 |
| FPGT | 248.00 | 223.00 | 254.00 | 241.67 | 281.00 |
| FRA10AC1 | 978.00 | 975.00 | 1066.00 | 1006.33 | 1164.00 |
| FRAS1 | 817.00 | 852.00 | 942.00 | 870.33 | 445.00 |
| FRAT1 | 83.00 | 85.00 | 97.00 | 88.33 | 115.00 |
| FRAT2 | 242.00 | 218.00 | 257.00 | 239.00 | 303.00 |
| FREM1 | 3.00 | 0.00 | 2.00 | 1.67 | 3.00 |
| FREM2 | 48.00 | 49.00 | 50.00 | 49.00 | 43.00 |
| FRG1 | 3.00 | 2.00 | 6.00 | 3.67 | 1.00 |
| FRG1BP | 248.00 | 230.00 | 245.00 | 241.00 | 250.00 |
| FRG1CP | 368.00 | 390.00 | 403.00 | 387.00 | 317.00 |
| FRG1DP | 70.00 | 58.00 | 66.00 | 64.67 | 66.00 |
| FRG1HP | 340.00 | 320.00 | 387.00 | 349.00 | 364.00 |
| FRG1JP | 67.00 | 74.00 | 90.00 | 77.00 | 82.00 |
| FRK | 9.00 | 4.00 | 6.00 | 6.33 | 11.00 |
| FRMD3 | 30.00 | 20.00 | 30.00 | 26.67 | 28.00 |
| FRMD4A | 3693.00 | 3529.00 | 3845.00 | 3689.00 | 2786.00 |
| FRMD4B | 233.00 | 166.00 | 188.00 | 195.67 | 256.00 |
| FRMD5 | 1047.00 | 945.00 | 1070.00 | 1020.67 | 866.00 |
| FRMD6 | 3757.00 | 3513.00 | 4289.00 | 3853.00 | 4695.00 |
| FRMD6-AS1 | 17.00 | 14.00 | 16.00 | 15.67 | 16.00 |
| FRMD8 | 1204.00 | 1236.00 | 1134.00 | 1191.33 | 1278.00 |
| FRMPD2 | 3.00 | 1.00 | 2.00 | 2.00 | 0.00 |
| FRMPD4 | 8.00 | 7.00 | 10.00 | 8.33 | 9.00 |
| FRRS1 | 120.00 | 122.00 | 173.00 | 138.33 | 151.00 |
| FRS2 | 408.00 | 423.00 | 464.00 | 431.67 | 433.00 |

| | | | | | |
|---|---|---|---|---|---|
| FRS3 | 188.00 | 152.00 | 235.00 | 191.67 | 154.00 |
| FRY | 118.00 | 104.00 | 134.00 | 118.67 | 131.00 |
| FRYL | 1674.00 | 1504.00 | 1630.00 | 1602.67 | 1570.00 |
| FRZB | 16.00 | 16.00 | 15.00 | 15.67 | 12.00 |
| FSCN1 | 1503.00 | 1547.00 | 1680.00 | 1576.67 | 2192.00 |
| FSD1 | 768.00 | 770.00 | 773.00 | 770.33 | 905.00 |
| FSD1L | 211.00 | 184.00 | 219.00 | 204.67 | 198.00 |
| FSIP1 | 12.00 | 20.00 | 19.00 | 17.00 | 21.00 |
| FSIP2 | 81.00 | 75.00 | 80.00 | 78.67 | 32.00 |
| FST | 2564.00 | 2325.00 | 2583.00 | 2490.67 | 3640.00 |
| FSTL1 | 5931.00 | 5787.00 | 5681.00 | 5799.67 | 5694.00 |
| FSTL3 | 453.00 | 473.00 | 512.00 | 479.33 | 332.00 |
| FTCD | 21.00 | 12.00 | 17.00 | 16.67 | 10.00 |
| FTCDNL1 | 30.00 | 38.00 | 24.00 | 30.67 | 43.00 |
| FTH1 | 6401.00 | 6379.00 | 6836.00 | 6538.67 | 7264.00 |
| FTH1P3 | 79.00 | 63.00 | 66.00 | 69.33 | 72.00 |
| FTL | 14798.00 | 15157.00 | 16649.00 | 15534.67 | 13052.00 |
| FTO | 749.00 | 717.00 | 765.00 | 743.67 | 743.00 |
| FTO-IT1 | 6.00 | 7.00 | 7.00 | 6.67 | 3.00 |
| FTSJ1 | 1162.00 | 1191.00 | 1152.00 | 1168.33 | 1236.00 |
| FTSJ2 | 639.00 | 638.00 | 708.00 | 661.67 | 894.00 |
| FTSJ3 | 2464.00 | 2453.00 | 2480.00 | 2465.67 | 2519.00 |
| FTX | 42.00 | 38.00 | 42.00 | 40.67 | 39.00 |
| FUBP1 | 3838.00 | 3551.00 | 4034.00 | 3807.67 | 3627.00 |
| FUBP3 | 1732.00 | 1642.00 | 1928.00 | 1767.33 | 1739.00 |
| FUCA1 | 145.00 | 118.00 | 124.00 | 129.00 | 165.00 |
| FUCA2 | 5411.00 | 5416.00 | 5089.00 | 5305.33 | 5487.00 |
| FUK | 594.00 | 589.00 | 658.00 | 613.67 | 574.00 |
| FUNDC1 | 250.00 | 207.00 | 254.00 | 237.00 | 316.00 |
| FUNDC2 | 327.00 | 377.00 | 393.00 | 365.67 | 448.00 |
| FUOM | 42.00 | 35.00 | 38.00 | 38.33 | 52.00 |
| FURIN | 2217.00 | 2212.00 | 2289.00 | 2239.33 | 1733.00 |
| FUS | 9417.00 | 9965.00 | 9999.00 | 9793.67 | 12138.00 |
| FUT1 | 8.00 | 3.00 | 4.00 | 5.00 | 8.00 |
| FUT10 | 145.00 | 121.00 | 153.00 | 139.67 | 160.00 |
| FUT11 | 221.00 | 181.00 | 232.00 | 211.33 | 160.00 |
| FUT4 | 407.00 | 442.00 | 386.00 | 411.67 | 414.00 |
| FUT8 | 3891.00 | 3934.00 | 4082.00 | 3969.00 | 4756.00 |
| FUT8-AS1 | 181.00 | 176.00 | 162.00 | 173.00 | 214.00 |
| FUZ | 401.00 | 379.00 | 419.00 | 399.67 | 540.00 |
| FXN | 241.00 | 237.00 | 246.00 | 241.33 | 315.00 |
| FXR1 | 3519.00 | 3388.00 | 3793.00 | 3566.67 | 4524.00 |
| FXR2 | 1377.00 | 1444.00 | 1646.00 | 1489.00 | 1403.00 |
| FXYD5 | 2795.00 | 2765.00 | 2685.00 | 2748.33 | 3463.00 |
| FYB | 4.00 | 0.00 | 0.00 | 1.33 | 1.00 |
| FYCO1 | 1422.00 | 1197.00 | 1396.00 | 1338.33 | 1080.00 |
| FYN | 2337.00 | 2183.00 | 2182.00 | 2234.00 | 2537.00 |
| FYTTD1 | 1057.00 | 1027.00 | 1123.00 | 1069.00 | 1137.00 |

| | | | | | |
|---|---|---|---|---|---|
| FZD1 | 444.00 | 417.00 | 396.00 | 419.00 | 394.00 |
| FZD2 | 1145.00 | 1173.00 | 1194.00 | 1170.67 | 1217.00 |
| FZD3 | 379.00 | 345.00 | 420.00 | 381.33 | 327.00 |
| FZD4 | 191.00 | 211.00 | 218.00 | 206.67 | 151.00 |
| FZD5 | 184.00 | 172.00 | 202.00 | 186.00 | 180.00 |
| FZD6 | 2857.00 | 2626.00 | 2965.00 | 2816.00 | 2831.00 |
| FZD7 | 349.00 | 342.00 | 379.00 | 356.67 | 304.00 |
| FZD8 | 881.00 | 887.00 | 932.00 | 900.00 | 854.00 |
| FZD9 | 9.00 | 4.00 | 7.00 | 6.67 | 4.00 |
| FZR1 | 900.00 | 842.00 | 892.00 | 878.00 | 854.00 |
| G0S2 | 6820.00 | 7039.00 | 6563.00 | 6807.33 | 8080.00 |
| G2E3 | 804.00 | 743.00 | 853.00 | 800.00 | 806.00 |
| G3BP1 | 4115.00 | 3584.00 | 4049.00 | 3916.00 | 4611.00 |
| G3BP2 | 1918.00 | 1740.00 | 2076.00 | 1911.33 | 1574.00 |
| G6PC3 | 2145.00 | 2219.00 | 1990.00 | 2118.00 | 2413.00 |
| G6PD | 1095.00 | 1075.00 | 1083.00 | 1084.33 | 1076.00 |
| GAA | 4659.00 | 4682.00 | 4678.00 | 4673.00 | 4472.00 |
| GAB1 | 236.00 | 217.00 | 251.00 | 234.67 | 179.00 |
| GAB2 | 409.00 | 373.00 | 436.00 | 406.00 | 298.00 |
| GAB3 | 7.00 | 6.00 | 7.00 | 6.67 | 13.00 |
| GABARAP | 2287.00 | 2270.00 | 2374.00 | 2310.33 | 3014.00 |
| GABARAPL1 | 568.00 | 557.00 | 644.00 | 589.67 | 663.00 |
| GABARAPL2 | 809.00 | 785.00 | 887.00 | 827.00 | 1028.00 |
| GABBR2 | 26.00 | 10.00 | 25.00 | 20.33 | 25.00 |
| GABPA | 770.00 | 727.00 | 883.00 | 793.33 | 842.00 |
| GABPB1 | 478.00 | 425.00 | 524.00 | 475.67 | 699.00 |
| GABPB1-AS1 | 1295.00 | 1159.00 | 1169.00 | 1207.67 | 671.00 |
| GABPB2 | 163.00 | 145.00 | 168.00 | 158.67 | 132.00 |
| GABRA2 | 27.00 | 37.00 | 38.00 | 34.00 | 33.00 |
| GABRA3 | 75.00 | 66.00 | 72.00 | 71.00 | 78.00 |
| GABRB2 | 8.00 | 6.00 | 2.00 | 5.33 | 16.00 |
| GABRE | 761.00 | 749.00 | 824.00 | 778.00 | 326.00 |
| GABRG1 | 48.00 | 31.00 | 49.00 | 42.67 | 76.00 |
| GABRG2 | 27.00 | 18.00 | 37.00 | 27.33 | 33.00 |
| GABRQ | 11.00 | 14.00 | 8.00 | 11.00 | 7.00 |
| GABRR2 | 4.00 | 3.00 | 10.00 | 5.67 | 6.00 |
| GACAT2 | 46.00 | 48.00 | 73.00 | 55.67 | 105.00 |
| GAD1 | 72.00 | 80.00 | 69.00 | 73.67 | 97.00 |
| GADD45A | 1846.00 | 1741.00 | 1915.00 | 1834.00 | 2346.00 |
| GADD45B | 975.00 | 1024.00 | 1072.00 | 1023.67 | 1169.00 |
| GADD45GIP1 | 821.00 | 815.00 | 894.00 | 843.33 | 1064.00 |
| GAK | 1686.00 | 1686.00 | 1791.00 | 1721.00 | 1458.00 |
| GAL3ST4 | 39.00 | 57.00 | 35.00 | 43.67 | 42.00 |
| GALC | 321.00 | 317.00 | 334.00 | 324.00 | 229.00 |
| GALE | 1832.00 | 1765.00 | 1897.00 | 1831.33 | 1945.00 |
| GALK1 | 263.00 | 321.00 | 319.00 | 301.00 | 459.00 |
| GALK2 | 535.00 | 503.00 | 533.00 | 523.67 | 530.00 |
| GALM | 10.00 | 10.00 | 13.00 | 11.00 | 9.00 |

| | | | | | |
|---|---|---|---|---|---|
| GALNS | 1317.00 | 1298.00 | 1184.00 | 1266.33 | 1374.00 |
| GALNT1 | 1173.00 | 989.00 | 1100.00 | 1087.33 | 1285.00 |
| GALNT10 | 2569.00 | 2296.00 | 2726.00 | 2530.33 | 2011.00 |
| GALNT11 | 1115.00 | 1050.00 | 1043.00 | 1069.33 | 1271.00 |
| GALNT12 | 150.00 | 125.00 | 135.00 | 136.67 | 181.00 |
| GALNT13 | 4.00 | 1.00 | 9.00 | 4.67 | 2.00 |
| GALNT15 | 20.00 | 23.00 | 30.00 | 24.33 | 12.00 |
| GALNT16 | 150.00 | 169.00 | 191.00 | 170.00 | 296.00 |
| GALNT18 | 190.00 | 202.00 | 228.00 | 206.67 | 346.00 |
| GALNT2 | 5099.00 | 4953.00 | 4797.00 | 4949.67 | 3886.00 |
| GALNT3 | 3.00 | 8.00 | 6.00 | 5.67 | 7.00 |
| GALNT4 | 17.00 | 14.00 | 15.00 | 15.33 | 3.00 |
| GALNT5 | 8.00 | 6.00 | 7.00 | 7.00 | 1.00 |
| GALNT6 | 3432.00 | 3334.00 | 3412.00 | 3392.67 | 3153.00 |
| GALNT7 | 1223.00 | 1098.00 | 1208.00 | 1176.33 | 813.00 |
| GALNTL6 | 6.00 | 7.00 | 17.00 | 10.00 | 11.00 |
| GALR3 | 4.00 | 9.00 | 8.00 | 7.00 | 8.00 |
| GALT | 359.00 | 386.00 | 342.00 | 362.33 | 390.00 |
| GAMT | 444.00 | 392.00 | 390.00 | 408.67 | 559.00 |
| GAN | 47.00 | 40.00 | 64.00 | 50.33 | 70.00 |
| GANAB | 25839.00 | 25179.00 | 26236.00 | 25751.33 | 21023.00 |
| GANC | 345.00 | 249.00 | 359.00 | 317.67 | 296.00 |
| GAP43 | 5.00 | 2.00 | 4.00 | 3.67 | 9.00 |
| GAPDH | 62649.00 | 60444.00 | 69670.00 | 64254.33 | 82176.00 |
| GAPLINC | 86.00 | 93.00 | 71.00 | 83.33 | 189.00 |
| GAPVD1 | 1328.00 | 1262.00 | 1345.00 | 1311.67 | 1254.00 |
| GAR1 | 717.00 | 663.00 | 715.00 | 698.33 | 946.00 |
| GAREM | 91.00 | 82.00 | 101.00 | 91.33 | 82.00 |
| GAREML | 16.00 | 17.00 | 15.00 | 16.00 | 24.00 |
| GARNL3 | 169.00 | 128.00 | 172.00 | 156.33 | 122.00 |
| GARS | 14840.00 | 13949.00 | 15101.00 | 14630.00 | 14810.00 |
| GART | 3178.00 | 2901.00 | 3309.00 | 3129.33 | 3507.00 |
| GAS2 | 12.00 | 9.00 | 14.00 | 11.67 | 23.00 |
| GAS2L1 | 304.00 | 337.00 | 324.00 | 321.67 | 347.00 |
| GAS2L3 | 535.00 | 475.00 | 595.00 | 535.00 | 633.00 |
| GAS5 | 10950.00 | 11507.00 | 12257.00 | 11571.33 | 15524.00 |
| GAS6 | 2259.00 | 2399.00 | 2500.00 | 2386.00 | 2336.00 |
| GAS6-AS1 | 20.00 | 18.00 | 17.00 | 18.33 | 6.00 |
| GAS6-AS2 | 83.00 | 64.00 | 77.00 | 74.67 | 61.00 |
| GAS8 | 726.00 | 762.00 | 782.00 | 756.67 | 861.00 |
| GATA2 | 216.00 | 207.00 | 200.00 | 207.67 | 243.00 |
| GATA2-AS1 | 54.00 | 60.00 | 58.00 | 57.33 | 30.00 |
| GATA6 | 425.00 | 442.00 | 504.00 | 457.00 | 312.00 |
| GATA6-AS1 | 53.00 | 53.00 | 52.00 | 52.67 | 36.00 |
| GATAD1 | 1929.00 | 1907.00 | 2242.00 | 2026.00 | 2143.00 |
| GATAD2A | 2684.00 | 2643.00 | 2784.00 | 2703.67 | 3028.00 |
| GATAD2B | 17.00 | 14.00 | 8.00 | 13.00 | 6.00 |
| GATB | 545.00 | 496.00 | 569.00 | 536.67 | 642.00 |

| | | | | | |
|---|---|---|---|---|---|
| GATC | 853.00 | 842.00 | 903.00 | 866.00 | 995.00 |
| GATM | 13.00 | 9.00 | 3.00 | 8.33 | 13.00 |
| GATS | 113.00 | 95.00 | 94.00 | 100.67 | 76.00 |
| GATSL2 | 97.00 | 90.00 | 86.00 | 91.00 | 114.00 |
| GATSL3 | 223.00 | 229.00 | 237.00 | 229.67 | 232.00 |
| GBA | 16.00 | 13.00 | 19.00 | 16.00 | 18.00 |
| GBA2 | 2328.00 | 2248.00 | 2510.00 | 2362.00 | 2039.00 |
| GBAP1 | 45.00 | 70.00 | 75.00 | 63.33 | 62.00 |
| GBAS | 919.00 | 847.00 | 999.00 | 921.67 | 1151.00 |
| GBAT2 | 7.00 | 6.00 | 3.00 | 5.33 | 8.00 |
| GBE1 | 1638.00 | 1550.00 | 1785.00 | 1657.67 | 1917.00 |
| GBF1 | 4241.00 | 3957.00 | 4387.00 | 4195.00 | 3668.00 |
| GBP1 | 121.00 | 114.00 | 154.00 | 129.67 | 187.00 |
| GBP2 | 10.00 | 8.00 | 20.00 | 12.67 | 25.00 |
| GBP3 | 95.00 | 100.00 | 102.00 | 99.00 | 137.00 |
| GBP5 | 6.00 | 8.00 | 9.00 | 7.67 | 8.00 |
| GBX2 | 21.00 | 15.00 | 20.00 | 18.67 | 38.00 |
| GCA | 224.00 | 197.00 | 199.00 | 206.67 | 264.00 |
| GCAT | 445.00 | 435.00 | 499.00 | 459.67 | 647.00 |
| GCC1 | 668.00 | 685.00 | 788.00 | 713.67 | 690.00 |
| GCC2 | 1183.00 | 1024.00 | 1198.00 | 1135.00 | 1027.00 |
| GCDH | 419.00 | 441.00 | 405.00 | 421.67 | 470.00 |
| GCFC2 | 609.00 | 622.00 | 739.00 | 656.67 | 858.00 |
| GCH1 | 39.00 | 51.00 | 63.00 | 51.00 | 70.00 |
| GCLC | 794.00 | 805.00 | 908.00 | 835.67 | 859.00 |
| GCLM | 713.00 | 638.00 | 746.00 | 699.00 | 1009.00 |
| GCN1 | 7009.00 | 6711.00 | 7168.00 | 6962.67 | 5830.00 |
| GCNT1 | 15.00 | 18.00 | 25.00 | 19.33 | 20.00 |
| GCNT2 | 806.00 | 738.00 | 701.00 | 748.33 | 760.00 |
| GCNT3 | 4.00 | 6.00 | 4.00 | 4.67 | 11.00 |
| GCSH | 244.00 | 243.00 | 257.00 | 248.00 | 342.00 |
| GCSHP3 | 107.00 | 94.00 | 92.00 | 97.67 | 128.00 |
| GDAP1 | 375.00 | 369.00 | 366.00 | 370.00 | 431.00 |
| GDAP1L1 | 3.00 | 1.00 | 7.00 | 3.67 | 7.00 |
| GDAP2 | 439.00 | 392.00 | 468.00 | 433.00 | 460.00 |
| GDE1 | 1118.00 | 1114.00 | 1121.00 | 1117.67 | 1368.00 |
| GDF11 | 348.00 | 359.00 | 337.00 | 348.00 | 288.00 |
| GDF15 | 1063.00 | 1068.00 | 1015.00 | 1048.67 | 510.00 |
| GDF9 | 39.00 | 33.00 | 27.00 | 33.00 | 31.00 |
| GDI1 | 2447.00 | 2534.00 | 2675.00 | 2552.00 | 2706.00 |
| GDI2 | 6354.00 | 5879.00 | 6981.00 | 6404.67 | 7983.00 |
| GDNF | 481.00 | 518.00 | 589.00 | 529.33 | 945.00 |
| GDNF-AS1 | 724.00 | 752.00 | 784.00 | 753.33 | 750.00 |
| GDPD1 | 314.00 | 337.00 | 343.00 | 331.33 | 309.00 |
| GDPD3 | 63.00 | 71.00 | 62.00 | 65.33 | 51.00 |
| GDPD5 | 206.00 | 205.00 | 213.00 | 208.00 | 161.00 |
| GDPGP1 | 27.00 | 25.00 | 37.00 | 29.67 | 30.00 |
| GEM | 870.00 | 850.00 | 976.00 | 898.67 | 857.00 |

| | | | | | |
|---|---|---|---|---|---|
| GEMIN2 | 200.00 | 220.00 | 233.00 | 217.67 | 229.00 |
| GEMIN4 | 1234.00 | 1219.00 | 1204.00 | 1219.00 | 1265.00 |
| GEMIN5 | 1291.00 | 1217.00 | 1279.00 | 1262.33 | 1339.00 |
| GEMIN6 | 281.00 | 251.00 | 278.00 | 270.00 | 333.00 |
| GEMIN7 | 228.00 | 233.00 | 264.00 | 241.67 | 337.00 |
| GEMIN8 | 538.00 | 578.00 | 658.00 | 591.33 | 669.00 |
| GEMIN8P4 | 14.00 | 8.00 | 16.00 | 12.67 | 25.00 |
| GEN1 | 887.00 | 840.00 | 802.00 | 843.00 | 1010.00 |
| GET4 | 1325.00 | 1358.00 | 1465.00 | 1382.67 | 1550.00 |
| GFER | 482.00 | 503.00 | 515.00 | 500.00 | 574.00 |
| GFI1 | 40.00 | 34.00 | 30.00 | 34.67 | 38.00 |
| GFM1 | 1526.00 | 1345.00 | 1592.00 | 1487.67 | 1771.00 |
| GFM2 | 1185.00 | 1161.00 | 1163.00 | 1169.67 | 1290.00 |
| GFOD1 | 96.00 | 106.00 | 115.00 | 105.67 | 99.00 |
| GFOD2 | 555.00 | 537.00 | 560.00 | 550.67 | 568.00 |
| GFPT1 | 2577.00 | 2296.00 | 2792.00 | 2555.00 | 2534.00 |
| GFPT2 | 1326.00 | 1330.00 | 1378.00 | 1344.67 | 1164.00 |
| GGA1 | 3215.00 | 3230.00 | 3469.00 | 3304.67 | 3003.00 |
| GGA2 | 1215.00 | 1255.00 | 1398.00 | 1289.33 | 1304.00 |
| GGA3 | 1527.00 | 1531.00 | 1643.00 | 1567.00 | 1394.00 |
| GGACT | 54.00 | 35.00 | 49.00 | 46.00 | 58.00 |
| GGCT | 759.00 | 708.00 | 787.00 | 751.33 | 1097.00 |
| GGCX | 1427.00 | 1449.00 | 1365.00 | 1413.67 | 1257.00 |
| GGH | 2354.00 | 2318.00 | 2113.00 | 2261.67 | 2836.00 |
| GGN | 14.00 | 13.00 | 15.00 | 14.00 | 14.00 |
| GGNBP1 | 4.00 | 2.00 | 1.00 | 2.33 | 5.00 |
| GGNBP2 | 5.00 | 5.00 | 10.00 | 6.67 | 7.00 |
| GGPS1 | 603.00 | 496.00 | 637.00 | 578.67 | 636.00 |
| GGT1 | 673.00 | 650.00 | 628.00 | 650.33 | 632.00 |
| GGT5 | 2198.00 | 2259.00 | 2189.00 | 2215.33 | 1245.00 |
| GGT7 | 2823.00 | 2965.00 | 3028.00 | 2938.67 | 2493.00 |
| GHDC | 1168.00 | 1139.00 | 1267.00 | 1191.33 | 1090.00 |
| GHET1 | 7.00 | 4.00 | 10.00 | 7.00 | 9.00 |
| GHITM | 4177.00 | 3646.00 | 4307.00 | 4043.33 | 4897.00 |
| GHR | 97.00 | 69.00 | 82.00 | 82.67 | 94.00 |
| GHRL | 7.00 | 3.00 | 2.00 | 4.00 | 3.00 |
| GHRLOS | 85.00 | 98.00 | 82.00 | 88.33 | 105.00 |
| GID4 | 512.00 | 461.00 | 606.00 | 526.33 | 487.00 |
| GID8 | 1157.00 | 1066.00 | 1161.00 | 1128.00 | 1087.00 |
| GIGYF1 | 4740.00 | 4922.00 | 5257.00 | 4973.00 | 5266.00 |
| GIGYF2 | 1297.00 | 1239.00 | 1353.00 | 1296.33 | 1207.00 |
| GIMAP2 | 13.00 | 11.00 | 17.00 | 13.67 | 24.00 |
| GIN1 | 170.00 | 150.00 | 185.00 | 168.33 | 149.00 |
| GINM1 | 684.00 | 649.00 | 660.00 | 664.33 | 796.00 |
| GINS1 | 415.00 | 392.00 | 407.00 | 404.67 | 521.00 |
| GINS2 | 184.00 | 203.00 | 179.00 | 188.67 | 299.00 |
| GINS3 | 201.00 | 235.00 | 209.00 | 215.00 | 353.00 |
| GINS4 | 499.00 | 517.00 | 465.00 | 493.67 | 624.00 |

| | | | | | |
|---|---|---|---|---|---|
| GIPC1 | 1460.00 | 1381.00 | 1532.00 | 1457.67 | 1428.00 |
| GIPR | 5.00 | 5.00 | 8.00 | 6.00 | 1.00 |
| GIT1 | 2095.00 | 2065.00 | 2309.00 | 2156.33 | 2214.00 |
| GIT2 | 1250.00 | 1250.00 | 1325.00 | 1275.00 | 1286.00 |
| GJA3 | 129.00 | 121.00 | 147.00 | 132.33 | 165.00 |
| GJB2 | 27.00 | 19.00 | 11.00 | 19.00 | 37.00 |
| GJC1 | 1095.00 | 1080.00 | 1188.00 | 1121.00 | 1018.00 |
| GJC2 | 14.00 | 12.00 | 13.00 | 13.00 | 16.00 |
| GJC3 | 3.00 | 1.00 | 4.00 | 2.67 | 0.00 |
| GJD4 | 4.00 | 3.00 | 1.00 | 2.67 | 7.00 |
| GK | 223.00 | 235.00 | 249.00 | 235.67 | 231.00 |
| GK5 | 906.00 | 821.00 | 931.00 | 886.00 | 872.00 |
| GKAP1 | 99.00 | 95.00 | 100.00 | 98.00 | 121.00 |
| GLA | 388.00 | 401.00 | 371.00 | 386.67 | 422.00 |
| GLB1 | 3321.00 | 3380.00 | 3449.00 | 3383.33 | 3050.00 |
| GLB1L | 159.00 | 207.00 | 184.00 | 183.33 | 185.00 |
| GLCCI1 | 316.00 | 260.00 | 333.00 | 303.00 | 276.00 |
| GLCE | 1382.00 | 1257.00 | 1431.00 | 1356.67 | 884.00 |
| GLDC | 14.00 | 19.00 | 21.00 | 18.00 | 47.00 |
| GLE1 | 1505.00 | 1330.00 | 1566.00 | 1467.00 | 1668.00 |
| GLG1 | 7417.00 | 6894.00 | 7740.00 | 7350.33 | 5657.00 |
| GLI1 | 7.00 | 5.00 | 8.00 | 6.67 | 6.00 |
| GLI2 | 1275.00 | 1272.00 | 1487.00 | 1344.67 | 1062.00 |
| GLI3 | 1163.00 | 1093.00 | 1177.00 | 1144.33 | 1053.00 |
| GLI4 | 630.00 | 620.00 | 770.00 | 673.33 | 650.00 |
| GLIDR | 138.00 | 109.00 | 119.00 | 122.00 | 119.00 |
| GLIPR1 | 1445.00 | 1339.00 | 1399.00 | 1394.33 | 1209.00 |
| GLIPR1L2 | 6.00 | 13.00 | 6.00 | 8.33 | 7.00 |
| GLIPR2 | 186.00 | 221.00 | 219.00 | 208.67 | 144.00 |
| GLIS1 | 69.00 | 82.00 | 96.00 | 82.33 | 48.00 |
| GLIS2 | 18.00 | 7.00 | 33.00 | 19.33 | 23.00 |
| GLIS3 | 1189.00 | 1071.00 | 1266.00 | 1175.33 | 948.00 |
| GLMN | 506.00 | 468.00 | 509.00 | 494.33 | 610.00 |
| GLMP | 3039.00 | 2987.00 | 2896.00 | 2974.00 | 2937.00 |
| GLO1 | 2801.00 | 2504.00 | 3033.00 | 2779.33 | 4520.00 |
| GLOD4 | 901.00 | 950.00 | 1011.00 | 954.00 | 1189.00 |
| GLRB | 464.00 | 413.00 | 467.00 | 448.00 | 503.00 |
| GLRX | 962.00 | 845.00 | 913.00 | 906.67 | 1450.00 |
| GLRX2 | 373.00 | 357.00 | 381.00 | 370.33 | 510.00 |
| GLRX3 | 1740.00 | 1644.00 | 1908.00 | 1764.00 | 2325.00 |
| GLRX5 | 511.00 | 540.00 | 531.00 | 527.33 | 669.00 |
| GLS | 3497.00 | 3146.00 | 3838.00 | 3493.67 | 4321.00 |
| GLT8D1 | 1549.00 | 1506.00 | 1475.00 | 1510.00 | 1666.00 |
| GLT8D2 | 338.00 | 322.00 | 364.00 | 341.33 | 291.00 |
| GLTP | 830.00 | 810.00 | 854.00 | 831.33 | 1200.00 |
| GLTSCR1 | 421.00 | 387.00 | 402.00 | 403.33 | 379.00 |
| GLTSCR1L | 593.00 | 620.00 | 607.00 | 606.67 | 522.00 |
| GLTSCR2 | 3180.00 | 3272.00 | 3439.00 | 3297.00 | 4013.00 |

| | | | | | |
|---|---|---|---|---|---|
| GLUD1 | 2437.00 | 2350.00 | 2635.00 | 2474.00 | 3008.00 |
| GLUD1P3 | 343.00 | 361.00 | 346.00 | 350.00 | 246.00 |
| GLUD1P7 | 13.00 | 5.00 | 11.00 | 9.67 | 11.00 |
| GLUD2 | 30.00 | 39.00 | 47.00 | 38.67 | 47.00 |
| GLUL | 726.00 | 772.00 | 793.00 | 763.67 | 982.00 |
| GLYATL1 | 6.00 | 3.00 | 5.00 | 4.67 | 7.00 |
| GLYATL2 | 5.00 | 4.00 | 4.00 | 4.33 | 4.00 |
| GLYCTK | 534.00 | 498.00 | 502.00 | 511.33 | 705.00 |
| GLYR1 | 2433.00 | 2357.00 | 2623.00 | 2471.00 | 2567.00 |
| GM2A | 822.00 | 725.00 | 749.00 | 765.33 | 919.00 |
| GMCL1 | 358.00 | 314.00 | 368.00 | 346.67 | 382.00 |
| GMDS | 339.00 | 344.00 | 348.00 | 343.67 | 398.00 |
| GMDS-AS1 | 30.00 | 40.00 | 32.00 | 34.00 | 32.00 |
| GMEB1 | 273.00 | 292.00 | 251.00 | 272.00 | 287.00 |
| GMEB2 | 878.00 | 928.00 | 903.00 | 903.00 | 1164.00 |
| GMFB | 1229.00 | 1136.00 | 1422.00 | 1262.33 | 1582.00 |
| GMFG | 10.00 | 9.00 | 7.00 | 8.67 | 0.00 |
| GMIP | 527.00 | 509.00 | 455.00 | 497.00 | 412.00 |
| GMNN | 333.00 | 307.00 | 386.00 | 342.00 | 528.00 |
| GMPPA | 1005.00 | 978.00 | 1133.00 | 1038.67 | 996.00 |
| GMPPB | 323.00 | 307.00 | 340.00 | 323.33 | 332.00 |
| GMPR | 54.00 | 39.00 | 54.00 | 49.00 | 58.00 |
| GMPR2 | 1586.00 | 1560.00 | 1815.00 | 1653.67 | 1621.00 |
| GMPS | 1970.00 | 1819.00 | 2089.00 | 1959.33 | 2416.00 |
| GNA11 | 1543.00 | 1440.00 | 1598.00 | 1527.00 | 1570.00 |
| GNA12 | 1967.00 | 1950.00 | 2158.00 | 2025.00 | 1984.00 |
| GNA13 | 1169.00 | 1049.00 | 1372.00 | 1196.67 | 1294.00 |
| GNA14 | 3.00 | 2.00 | 0.00 | 1.67 | 2.00 |
| GNAI1 | 273.00 | 211.00 | 245.00 | 243.00 | 287.00 |
| GNAI2 | 3358.00 | 3248.00 | 3395.00 | 3333.67 | 3575.00 |
| GNAI3 | 1403.00 | 1149.00 | 1309.00 | 1287.00 | 1731.00 |
| GNAQ | 816.00 | 667.00 | 907.00 | 796.67 | 908.00 |
| GNAS | 8791.00 | 8587.00 | 8864.00 | 8747.33 | 10450.00 |
| GNAS-AS1 | 24.00 | 18.00 | 25.00 | 22.33 | 16.00 |
| GNAT2 | 4.00 | 7.00 | 6.00 | 5.67 | 2.00 |
| GNAZ | 7.00 | 8.00 | 11.00 | 8.67 | 10.00 |
| GNB1 | 5557.00 | 5235.00 | 5670.00 | 5487.33 | 6672.00 |
| GNB1L | 197.00 | 208.00 | 180.00 | 195.00 | 204.00 |
| GNB2 | 2351.00 | 2306.00 | 2481.00 | 2379.33 | 2618.00 |
| GNB2L1 | 33460.00 | 32101.00 | 37496.00 | 34352.33 | 39910.00 |
| GNB3 | 970.00 | 1001.00 | 1112.00 | 1027.67 | 906.00 |
| GNB4 | 833.00 | 626.00 | 827.00 | 762.00 | 846.00 |
| GNB5 | 356.00 | 346.00 | 365.00 | 355.67 | 352.00 |
| GNE | 1009.00 | 924.00 | 1003.00 | 978.67 | 804.00 |
| GNG10 | 37.00 | 46.00 | 56.00 | 46.33 | 75.00 |
| GNG11 | 738.00 | 744.00 | 818.00 | 766.67 | 989.00 |
| GNG12 | 2767.00 | 2308.00 | 2982.00 | 2685.67 | 2441.00 |
| GNG12-AS1 | 6.00 | 17.00 | 10.00 | 11.00 | 8.00 |

| | | | | | |
|---|---|---|---|---|---|
| GNG2 | 41.00 | 36.00 | 52.00 | 43.00 | 56.00 |
| GNG4 | 124.00 | 129.00 | 170.00 | 141.00 | 222.00 |
| GNG5 | 1545.00 | 1528.00 | 1696.00 | 1589.67 | 2098.00 |
| GNGT2 | 21.00 | 13.00 | 5.00 | 13.00 | 18.00 |
| GNL2 | 2041.00 | 1898.00 | 2216.00 | 2051.67 | 2772.00 |
| GNL3 | 3366.00 | 3354.00 | 3610.00 | 3443.33 | 3857.00 |
| GNL3L | 1148.00 | 1014.00 | 1087.00 | 1083.00 | 1237.00 |
| GNMT | 10.00 | 5.00 | 3.00 | 6.00 | 5.00 |
| GNPAT | 1205.00 | 1116.00 | 1229.00 | 1183.33 | 1448.00 |
| GNPDA1 | 2736.00 | 2628.00 | 2938.00 | 2767.33 | 3087.00 |
| GNPDA2 | 740.00 | 687.00 | 772.00 | 733.00 | 819.00 |
| GNPNAT1 | 2658.00 | 2330.00 | 2942.00 | 2643.33 | 2859.00 |
| GNPTAB | 1009.00 | 983.00 | 1144.00 | 1045.33 | 952.00 |
| GNPTG | 1050.00 | 1180.00 | 1164.00 | 1131.33 | 1156.00 |
| GNRH1 | 44.00 | 59.00 | 56.00 | 53.00 | 59.00 |
| GNRH2 | 8.00 | 7.00 | 6.00 | 7.00 | 2.00 |
| GNRHR2 | 9.00 | 12.00 | 22.00 | 14.33 | 19.00 |
| GNS | 6513.00 | 5977.00 | 6165.00 | 6218.33 | 3400.00 |
| GOLGA1 | 806.00 | 789.00 | 796.00 | 797.00 | 770.00 |
| GOLGA2 | 2936.00 | 2780.00 | 2972.00 | 2896.00 | 2869.00 |
| GOLGA2P5 | 47.00 | 42.00 | 39.00 | 42.67 | 32.00 |
| GOLGA2P7 | 102.00 | 105.00 | 93.00 | 100.00 | 98.00 |
| GOLGA3 | 3332.00 | 3268.00 | 3329.00 | 3309.67 | 2925.00 |
| GOLGA4 | 2586.00 | 2379.00 | 2804.00 | 2589.67 | 2381.00 |
| GOLGA5 | 975.00 | 862.00 | 1067.00 | 968.00 | 1131.00 |
| GOLGA6L10 | 13.00 | 18.00 | 13.00 | 14.67 | 13.00 |
| GOLGA6L4 | 3.00 | 4.00 | 3.00 | 3.33 | 1.00 |
| GOLGA6L5P | 4.00 | 5.00 | 7.00 | 5.33 | 9.00 |
| GOLGA6L9 | 43.00 | 49.00 | 57.00 | 49.67 | 40.00 |
| GOLGA7 | 701.00 | 631.00 | 726.00 | 686.00 | 873.00 |
| GOLGA7B | 4.00 | 5.00 | 2.00 | 3.67 | 5.00 |
| GOLGA8A | 4380.00 | 4550.00 | 4858.00 | 4596.00 | 4617.00 |
| GOLGA8B | 2993.00 | 3117.00 | 3406.00 | 3172.00 | 3278.00 |
| GOLGA8J | 8.00 | 2.00 | 7.00 | 5.67 | 1.00 |
| GOLGA8O | 5.00 | 6.00 | 4.00 | 5.00 | 5.00 |
| GOLGA8R | 6.00 | 6.00 | 5.00 | 5.67 | 4.00 |
| GOLGB1 | 3498.00 | 3169.00 | 3484.00 | 3383.67 | 2517.00 |
| GOLIM4 | 1751.00 | 1619.00 | 1800.00 | 1723.33 | 1696.00 |
| GOLM1 | 3581.00 | 3501.00 | 3512.00 | 3531.33 | 3517.00 |
| GOLPH3 | 989.00 | 877.00 | 1104.00 | 990.00 | 1191.00 |
| GOLPH3L | 566.00 | 499.00 | 596.00 | 553.67 | 675.00 |
| GOLT1B | 2669.00 | 2415.00 | 2939.00 | 2674.33 | 3073.00 |
| GON4L | 1561.00 | 1419.00 | 1559.00 | 1513.00 | 1158.00 |
| GOPC | 1624.00 | 1552.00 | 1687.00 | 1621.00 | 1647.00 |
| GORAB | 580.00 | 593.00 | 611.00 | 594.67 | 723.00 |
| GORASP1 | 1300.00 | 1301.00 | 1486.00 | 1362.33 | 1319.00 |
| GORASP2 | 1826.00 | 1642.00 | 1884.00 | 1784.00 | 2022.00 |
| GOSR1 | 826.00 | 799.00 | 927.00 | 850.67 | 933.00 |

| | | | | | |
|---|---|---|---|---|---|
| GOSR2 | 1147.00 | 1145.00 | 1202.00 | 1164.67 | 1119.00 |
| GOT1 | 1842.00 | 1724.00 | 1787.00 | 1784.33 | 2179.00 |
| GOT2 | 2589.00 | 2433.00 | 2590.00 | 2537.33 | 3069.00 |
| GPAA1 | 3483.00 | 3430.00 | 3214.00 | 3375.67 | 3457.00 |
| GPALPP1 | 320.00 | 302.00 | 318.00 | 313.33 | 335.00 |
| GPAM | 1689.00 | 1508.00 | 1700.00 | 1632.33 | 1159.00 |
| GPAT3 | 248.00 | 263.00 | 254.00 | 255.00 | 228.00 |
| GPAT4 | 3610.00 | 3630.00 | 3679.00 | 3639.67 | 3475.00 |
| GPATCH1 | 213.00 | 227.00 | 227.00 | 222.33 | 240.00 |
| GPATCH11 | 435.00 | 352.00 | 428.00 | 405.00 | 397.00 |
| GPATCH2 | 493.00 | 432.00 | 445.00 | 456.67 | 479.00 |
| GPATCH2L | 1594.00 | 1484.00 | 1694.00 | 1590.67 | 1319.00 |
| GPATCH3 | 264.00 | 303.00 | 315.00 | 294.00 | 323.00 |
| GPATCH4 | 1113.00 | 1171.00 | 1209.00 | 1164.33 | 1493.00 |
| GPATCH8 | 2151.00 | 2044.00 | 2266.00 | 2153.67 | 1752.00 |
| GPBAR1 | 4.00 | 3.00 | 1.00 | 2.67 | 6.00 |
| GPBP1 | 1612.00 | 1640.00 | 1803.00 | 1685.00 | 1774.00 |
| GPBP1L1 | 2974.00 | 3013.00 | 3230.00 | 3072.33 | 3196.00 |
| GPC1 | 98.00 | 103.00 | 90.00 | 97.00 | 107.00 |
| GPC2 | 237.00 | 245.00 | 246.00 | 242.67 | 294.00 |
| GPC5 | 8.00 | 7.00 | 8.00 | 7.67 | 9.00 |
| GPC6 | 158.00 | 132.00 | 154.00 | 148.00 | 83.00 |
| GPCPD1 | 1394.00 | 1448.00 | 1591.00 | 1477.67 | 1458.00 |
| GPD1 | 12.00 | 15.00 | 9.00 | 12.00 | 20.00 |
| GPD1L | 531.00 | 464.00 | 598.00 | 531.00 | 578.00 |
| GPD2 | 799.00 | 708.00 | 857.00 | 788.00 | 1012.00 |
| GPER1 | 6.00 | 5.00 | 5.00 | 5.33 | 10.00 |
| GPHN | 560.00 | 560.00 | 582.00 | 567.33 | 545.00 |
| GPI | 2342.00 | 2208.00 | 2377.00 | 2309.00 | 2949.00 |
| GPKOW | 549.00 | 567.00 | 576.00 | 564.00 | 664.00 |
| GPLD1 | 33.00 | 27.00 | 24.00 | 28.00 | 41.00 |
| GPM6A | 13.00 | 13.00 | 11.00 | 12.33 | 15.00 |
| GPM6B | 88.00 | 94.00 | 98.00 | 93.33 | 142.00 |
| GPN1 | 771.00 | 706.00 | 849.00 | 775.33 | 834.00 |
| GPN2 | 512.00 | 438.00 | 541.00 | 497.00 | 467.00 |
| GPN3 | 546.00 | 478.00 | 530.00 | 518.00 | 678.00 |
| GPNMB | 1703.00 | 1692.00 | 1627.00 | 1674.00 | 892.00 |
| GPR1 | 5.00 | 5.00 | 10.00 | 6.67 | 4.00 |
| GPR107 | 3105.00 | 2992.00 | 3187.00 | 3094.67 | 2944.00 |
| GPR108 | 1742.00 | 1764.00 | 1750.00 | 1752.00 | 1754.00 |
| GPR135 | 306.00 | 311.00 | 374.00 | 330.33 | 215.00 |
| GPR137 | 1622.00 | 1605.00 | 1653.00 | 1626.67 | 1655.00 |
| GPR137B | 544.00 | 519.00 | 499.00 | 520.67 | 497.00 |
| GPR137C | 62.00 | 66.00 | 53.00 | 60.33 | 52.00 |
| GPR143 | 1271.00 | 1257.00 | 1295.00 | 1274.33 | 1562.00 |
| GPR146 | 24.00 | 14.00 | 24.00 | 20.67 | 13.00 |
| GPR153 | 151.00 | 181.00 | 177.00 | 169.67 | 116.00 |
| GPR155 | 169.00 | 149.00 | 165.00 | 161.00 | 118.00 |

| | | | | | |
|---|---|---|---|---|---|
| GPR156 | 30.00 | 35.00 | 32.00 | 32.33 | 27.00 |
| GPR157 | 68.00 | 52.00 | 70.00 | 63.33 | 45.00 |
| GPR158 | 4.00 | 0.00 | 6.00 | 3.33 | 9.00 |
| GPR160 | 23.00 | 16.00 | 23.00 | 20.67 | 27.00 |
| GPR161 | 866.00 | 878.00 | 963.00 | 902.33 | 773.00 |
| GPR162 | 126.00 | 128.00 | 110.00 | 121.33 | 117.00 |
| GPR173 | 605.00 | 672.00 | 661.00 | 646.00 | 588.00 |
| GPR176 | 882.00 | 822.00 | 955.00 | 886.33 | 905.00 |
| GPR180 | 870.00 | 845.00 | 941.00 | 885.33 | 967.00 |
| GPR19 | 46.00 | 67.00 | 46.00 | 53.00 | 48.00 |
| GPR3 | 306.00 | 289.00 | 341.00 | 312.00 | 293.00 |
| GPR35 | 15.00 | 10.00 | 14.00 | 13.00 | 16.00 |
| GPR37 | 181.00 | 193.00 | 177.00 | 183.67 | 211.00 |
| GPR39 | 1624.00 | 1628.00 | 1749.00 | 1667.00 | 1635.00 |
| GPR4 | 243.00 | 194.00 | 212.00 | 216.33 | 103.00 |
| GPR50 | 3.00 | 0.00 | 1.00 | 1.33 | 0.00 |
| GPR63 | 453.00 | 496.00 | 519.00 | 489.33 | 451.00 |
| GPR65 | 3.00 | 3.00 | 1.00 | 2.33 | 7.00 |
| GPR68 | 58.00 | 62.00 | 61.00 | 60.33 | 39.00 |
| GPR75 | 155.00 | 175.00 | 168.00 | 166.00 | 125.00 |
| GPR83 | 5.00 | 4.00 | 5.00 | 4.67 | 5.00 |
| GPR85 | 74.00 | 57.00 | 84.00 | 71.67 | 72.00 |
| GPR88 | 4.00 | 3.00 | 6.00 | 4.33 | 3.00 |
| GPR89A | 157.00 | 128.00 | 136.00 | 140.33 | 144.00 |
| GPR89B | 102.00 | 102.00 | 121.00 | 108.33 | 96.00 |
| GPRASP1 | 469.00 | 436.00 | 486.00 | 463.67 | 414.00 |
| GPRASP2 | 27.00 | 18.00 | 23.00 | 22.67 | 15.00 |
| GPRC5A | 786.00 | 821.00 | 914.00 | 840.33 | 598.00 |
| GPRIN1 | 162.00 | 162.00 | 188.00 | 170.67 | 163.00 |
| GPRIN3 | 3.00 | 3.00 | 2.00 | 2.67 | 1.00 |
| GPS1 | 2308.00 | 2184.00 | 2317.00 | 2269.67 | 2506.00 |
| GPS2 | 778.00 | 849.00 | 790.00 | 805.67 | 882.00 |
| GPSM1 | 2164.00 | 2222.00 | 2403.00 | 2263.00 | 1930.00 |
| GPSM2 | 342.00 | 352.00 | 429.00 | 374.33 | 472.00 |
| GPT | 12.00 | 27.00 | 30.00 | 23.00 | 18.00 |
| GPT2 | 2304.00 | 2373.00 | 2399.00 | 2358.67 | 2422.00 |
| GPX1 | 2222.00 | 2196.00 | 2204.00 | 2207.33 | 2788.00 |
| GPX4 | 1782.00 | 1741.00 | 1830.00 | 1784.33 | 2412.00 |
| GPX8 | 2381.00 | 2452.00 | 2296.00 | 2376.33 | 3135.00 |
| GRAMD1A | 1237.00 | 1224.00 | 1226.00 | 1229.00 | 1179.00 |
| GRAMD1C | 132.00 | 114.00 | 113.00 | 119.67 | 117.00 |
| GRAMD3 | 112.00 | 119.00 | 108.00 | 113.00 | 117.00 |
| GRAMD4 | 388.00 | 338.00 | 359.00 | 361.67 | 412.00 |
| GRB10 | 3107.00 | 3012.00 | 3144.00 | 3087.67 | 2949.00 |
| GRB14 | 85.00 | 71.00 | 90.00 | 82.00 | 101.00 |
| GRB2 | 1890.00 | 1739.00 | 2001.00 | 1876.67 | 2068.00 |
| GREB1 | 51.00 | 54.00 | 71.00 | 58.67 | 56.00 |
| GREB1L | 67.00 | 83.00 | 87.00 | 79.00 | 69.00 |

| | | | | | |
|---|---|---|---|---|---|
| GRHL1 | 38.00 | 23.00 | 29.00 | 30.00 | 31.00 |
| GRHPR | 1525.00 | 1566.00 | 1723.00 | 1604.67 | 2030.00 |
| GRIA1 | 4.00 | 4.00 | 4.00 | 4.00 | 7.00 |
| GRID2 | 8.00 | 8.00 | 13.00 | 9.67 | 5.00 |
| GRID2IP | 3.00 | 0.00 | 1.00 | 1.33 | 0.00 |
| GRIK1-AS1 | 3.00 | 4.00 | 2.00 | 3.00 | 1.00 |
| GRIK1-AS2 | 28.00 | 30.00 | 46.00 | 34.67 | 18.00 |
| GRIK2 | 25.00 | 28.00 | 31.00 | 28.00 | 16.00 |
| GRIK5 | 14.00 | 20.00 | 26.00 | 20.00 | 13.00 |
| GRIN2D | 109.00 | 100.00 | 85.00 | 98.00 | 73.00 |
| GRINA | 4891.00 | 4885.00 | 4873.00 | 4883.00 | 4680.00 |
| GRIP1 | 23.00 | 31.00 | 28.00 | 27.33 | 24.00 |
| GRIPAP1 | 1641.00 | 1675.00 | 1667.00 | 1661.00 | 1557.00 |
| GRK4 | 213.00 | 165.00 | 189.00 | 189.00 | 183.00 |
| GRK5 | 277.00 | 262.00 | 259.00 | 266.00 | 245.00 |
| GRK6 | 797.00 | 761.00 | 779.00 | 779.00 | 1166.00 |
| GRK7 | 5.00 | 2.00 | 3.00 | 3.33 | 3.00 |
| GRM1 | 12.00 | 10.00 | 7.00 | 9.67 | 5.00 |
| GRM5 | 3.00 | 2.00 | 0.00 | 1.67 | 9.00 |
| GRM8 | 3.00 | 3.00 | 14.00 | 6.67 | 10.00 |
| GRN | 12724.00 | 13356.00 | 12765.00 | 12948.33 | 10451.00 |
| GRPEL1 | 1045.00 | 998.00 | 995.00 | 1012.67 | 1379.00 |
| GRPEL2 | 986.00 | 927.00 | 1023.00 | 978.67 | 1353.00 |
| GRPR | 459.00 | 436.00 | 441.00 | 445.33 | 438.00 |
| GRSF1 | 2191.00 | 1978.00 | 2275.00 | 2148.00 | 2627.00 |
| GRWD1 | 844.00 | 757.00 | 831.00 | 810.67 | 867.00 |
| GS1-124K5.11 | 392.00 | 391.00 | 428.00 | 403.67 | 388.00 |
| GS1-259H13.2 | 11.00 | 14.00 | 10.00 | 11.67 | 9.00 |
| GSAP | 586.00 | 622.00 | 684.00 | 630.67 | 882.00 |
| GSDMB | 500.00 | 580.00 | 663.00 | 581.00 | 532.00 |
| GSE1 | 1826.00 | 1705.00 | 2006.00 | 1845.67 | 1292.00 |
| GSG1 | 12.00 | 22.00 | 14.00 | 16.00 | 21.00 |
| GSG2 | 271.00 | 232.00 | 254.00 | 252.33 | 273.00 |
| GSK3A | 1023.00 | 950.00 | 1054.00 | 1009.00 | 1130.00 |
| GSK3B | 1734.00 | 1596.00 | 1940.00 | 1756.67 | 1664.00 |
| GSKIP | 241.00 | 214.00 | 281.00 | 245.33 | 340.00 |
| GSN | 282.00 | 298.00 | 277.00 | 285.67 | 378.00 |
| GSPT1 | 5498.00 | 4993.00 | 5611.00 | 5367.33 | 5695.00 |
| GSPT2 | 374.00 | 430.00 | 450.00 | 418.00 | 418.00 |
| GSR | 672.00 | 598.00 | 677.00 | 649.00 | 846.00 |
| GSS | 2922.00 | 2954.00 | 3084.00 | 2986.67 | 3130.00 |
| GSTA4 | 150.00 | 152.00 | 151.00 | 151.00 | 191.00 |
| GSTCD | 383.00 | 308.00 | 375.00 | 355.33 | 401.00 |
| GSTK1 | 2315.00 | 2352.00 | 2547.00 | 2404.67 | 2689.00 |
| GSTM2 | 61.00 | 56.00 | 60.00 | 59.00 | 78.00 |
| GSTM3 | 678.00 | 700.00 | 727.00 | 701.67 | 777.00 |
| GSTM4 | 186.00 | 140.00 | 214.00 | 180.00 | 209.00 |
| GSTO1 | 3849.00 | 3599.00 | 4118.00 | 3855.33 | 4852.00 |

| | | | | | |
|---|---|---|---|---|---|
| GSTO2 | 62.00 | 56.00 | 84.00 | 67.33 | 94.00 |
| GSTP1 | 3490.00 | 3379.00 | 3416.00 | 3428.33 | 4543.00 |
| GSTZ1 | 186.00 | 257.00 | 310.00 | 251.00 | 276.00 |
| GTDC1 | 414.00 | 378.00 | 453.00 | 415.00 | 606.00 |
| GTF2A1 | 985.00 | 937.00 | 1156.00 | 1026.00 | 1228.00 |
| GTF2A2 | 529.00 | 521.00 | 584.00 | 544.67 | 657.00 |
| GTF2B | 518.00 | 487.00 | 563.00 | 522.67 | 652.00 |
| GTF2E1 | 575.00 | 513.00 | 566.00 | 551.33 | 659.00 |
| GTF2E2 | 519.00 | 479.00 | 548.00 | 515.33 | 639.00 |
| GTF2F1 | 1751.00 | 1730.00 | 1807.00 | 1762.67 | 1669.00 |
| GTF2F2 | 1053.00 | 1082.00 | 1184.00 | 1106.33 | 1510.00 |
| GTF2H1 | 1105.00 | 1036.00 | 1206.00 | 1115.67 | 1436.00 |
| GTF2H2B | 11.00 | 10.00 | 13.00 | 11.33 | 15.00 |
| GTF2H2C | 10.00 | 5.00 | 5.00 | 6.67 | 3.00 |
| GTF2H3 | 1418.00 | 1302.00 | 1531.00 | 1417.00 | 1692.00 |
| GTF2H4 | 141.00 | 106.00 | 143.00 | 130.00 | 114.00 |
| GTF2H5 | 455.00 | 398.00 | 554.00 | 469.00 | 581.00 |
| GTF2I | 1037.00 | 982.00 | 1077.00 | 1032.00 | 852.00 |
| GTF2IP1 | 96.00 | 99.00 | 98.00 | 97.67 | 117.00 |
| GTF2IP20 | 524.00 | 490.00 | 531.00 | 515.00 | 485.00 |
| GTF2IRD1 | 541.00 | 535.00 | 596.00 | 557.33 | 589.00 |
| GTF2IRD2 | 34.00 | 34.00 | 21.00 | 29.67 | 16.00 |
| GTF2IRD2B | 54.00 | 50.00 | 59.00 | 54.33 | 43.00 |
| GTF3A | 1608.00 | 1556.00 | 1826.00 | 1663.33 | 2135.00 |
| GTF3C1 | 3826.00 | 3717.00 | 3862.00 | 3801.67 | 3316.00 |
| GTF3C2 | 1800.00 | 1725.00 | 1915.00 | 1813.33 | 2027.00 |
| GTF3C3 | 2039.00 | 1823.00 | 2051.00 | 1971.00 | 1975.00 |
| GTF3C4 | 689.00 | 561.00 | 637.00 | 629.00 | 699.00 |
| GTF3C5 | 1292.00 | 1294.00 | 1302.00 | 1296.00 | 1478.00 |
| GTF3C6 | 747.00 | 649.00 | 725.00 | 707.00 | 990.00 |
| GTPBP1 | 1193.00 | 1058.00 | 1204.00 | 1151.67 | 872.00 |
| GTPBP10 | 746.00 | 686.00 | 774.00 | 735.33 | 766.00 |
| GTPBP2 | 6801.00 | 6809.00 | 6866.00 | 6825.33 | 6464.00 |
| GTPBP3 | 922.00 | 961.00 | 964.00 | 949.00 | 961.00 |
| GTPBP4 | 3846.00 | 3470.00 | 3874.00 | 3730.00 | 4288.00 |
| GTPBP6 | 6.00 | 3.00 | 2.00 | 3.67 | 5.00 |
| GTPBP8 | 351.00 | 322.00 | 319.00 | 330.67 | 447.00 |
| GTSE1 | 568.00 | 512.00 | 501.00 | 527.00 | 545.00 |
| GTSE1-AS1 | 5.00 | 7.00 | 6.00 | 6.00 | 13.00 |
| GUCA1B | 55.00 | 48.00 | 69.00 | 57.33 | 52.00 |
| GUCD1 | 909.00 | 872.00 | 860.00 | 880.33 | 927.00 |
| GUCY1A2 | 8.00 | 6.00 | 11.00 | 8.33 | 7.00 |
| GUCY1A3 | 23.00 | 13.00 | 24.00 | 20.00 | 28.00 |
| GUCY1B3 | 139.00 | 112.00 | 133.00 | 128.00 | 165.00 |
| GUF1 | 1940.00 | 1857.00 | 2090.00 | 1962.33 | 2068.00 |
| GUK1 | 1587.00 | 1680.00 | 1730.00 | 1665.67 | 1919.00 |
| GULP1 | 198.00 | 183.00 | 218.00 | 199.67 | 230.00 |
| GUSB | 2601.00 | 2724.00 | 2568.00 | 2631.00 | 2928.00 |

| | | | | | |
|---|---|---|---|---|---|
| GUSBP1 | 212.00 | 202.00 | 192.00 | 202.00 | 230.00 |
| GUSBP11 | 527.00 | 550.00 | 592.00 | 556.33 | 430.00 |
| GUSBP2 | 20.00 | 15.00 | 17.00 | 17.33 | 23.00 |
| GUSBP4 | 299.00 | 343.00 | 398.00 | 346.67 | 399.00 |
| GVINP1 | 7.00 | 16.00 | 14.00 | 12.33 | 8.00 |
| GXYLT1 | 1562.00 | 1379.00 | 1566.00 | 1502.33 | 1415.00 |
| GXYLT2 | 79.00 | 74.00 | 87.00 | 80.00 | 95.00 |
| GYG1 | 985.00 | 974.00 | 1135.00 | 1031.33 | 1382.00 |
| GYPC | 392.00 | 434.00 | 484.00 | 436.67 | 460.00 |
| GYPE | 35.00 | 29.00 | 32.00 | 32.00 | 38.00 |
| GYS1 | 1126.00 | 1057.00 | 1053.00 | 1078.67 | 1119.00 |
| GZF1 | 861.00 | 794.00 | 961.00 | 872.00 | 873.00 |
| GZMA | 83.00 | 93.00 | 102.00 | 92.67 | 77.00 |
| H1F0 | 1415.00 | 1326.00 | 1420.00 | 1387.00 | 1620.00 |
| H1FX | 1143.00 | 1216.00 | 1244.00 | 1201.00 | 1252.00 |
| H1FX-AS1 | 129.00 | 106.00 | 130.00 | 121.67 | 131.00 |
| H2AFJ | 388.00 | 444.00 | 416.00 | 416.00 | 526.00 |
| H2AFV | 2124.00 | 2102.00 | 2131.00 | 2119.00 | 2659.00 |
| H2AFX | 977.00 | 1010.00 | 900.00 | 962.33 | 1188.00 |
| H2AFY | 2604.00 | 2708.00 | 2642.00 | 2651.33 | 2806.00 |
| H2AFZ | 4042.00 | 3765.00 | 3938.00 | 3915.00 | 6020.00 |
| H3F3AP4 | 78.00 | 57.00 | 59.00 | 64.67 | 132.00 |
| H3F3B | 8154.00 | 7702.00 | 8966.00 | 8274.00 | 11633.00 |
| H3F3C | 53.00 | 45.00 | 47.00 | 48.33 | 54.00 |
| H6PD | 3609.00 | 3559.00 | 3747.00 | 3638.33 | 2599.00 |
| HABP4 | 425.00 | 425.00 | 428.00 | 426.00 | 453.00 |
| HACD1 | 433.00 | 469.00 | 517.00 | 473.00 | 567.00 |
| HACD2 | 514.00 | 362.00 | 409.00 | 428.33 | 343.00 |
| HACD3 | 3651.00 | 3560.00 | 3683.00 | 3631.33 | 4185.00 |
| HACD4 | 168.00 | 167.00 | 138.00 | 157.67 | 212.00 |
| HACE1 | 479.00 | 379.00 | 443.00 | 433.67 | 475.00 |
| HACL1 | 583.00 | 533.00 | 590.00 | 568.67 | 721.00 |
| HADH | 456.00 | 427.00 | 464.00 | 449.00 | 594.00 |
| HADHA | 3956.00 | 3822.00 | 4004.00 | 3927.33 | 4707.00 |
| HADHB | 1898.00 | 1778.00 | 2004.00 | 1893.33 | 2369.00 |
| HAGH | 524.00 | 523.00 | 539.00 | 528.67 | 628.00 |
| HAGHL | 528.00 | 548.00 | 658.00 | 578.00 | 574.00 |
| HAGLR | 208.00 | 214.00 | 219.00 | 213.67 | 156.00 |
| HAL | 4.00 | 4.00 | 3.00 | 3.67 | 7.00 |
| HAND2 | 64.00 | 46.00 | 50.00 | 53.33 | 79.00 |
| HAND2-AS1 | 4.00 | 3.00 | 6.00 | 4.33 | 6.00 |
| HAPLN3 | 46.00 | 39.00 | 38.00 | 41.00 | 56.00 |
| HARBI1 | 82.00 | 74.00 | 74.00 | 76.67 | 100.00 |
| HARS | 2152.00 | 2116.00 | 2144.00 | 2137.33 | 2340.00 |
| HARS2 | 1176.00 | 1245.00 | 1239.00 | 1220.00 | 1338.00 |
| HAS2 | 55.00 | 54.00 | 74.00 | 61.00 | 89.00 |
| HAS2-AS1 | 4.00 | 7.00 | 6.00 | 5.67 | 14.00 |
| HAS3 | 103.00 | 116.00 | 77.00 | 98.67 | 100.00 |

| | | | | | |
|---|---|---|---|---|---|
| HAT1 | 872.00 | 842.00 | 886.00 | 866.67 | 1305.00 |
| HAUS1 | 251.00 | 227.00 | 279.00 | 252.33 | 383.00 |
| HAUS2 | 404.00 | 359.00 | 428.00 | 397.00 | 504.00 |
| HAUS3 | 427.00 | 431.00 | 419.00 | 425.67 | 462.00 |
| HAUS4 | 698.00 | 653.00 | 737.00 | 696.00 | 845.00 |
| HAUS5 | 731.00 | 781.00 | 800.00 | 770.67 | 787.00 |
| HAUS6 | 1641.00 | 1471.00 | 1745.00 | 1619.00 | 1831.00 |
| HAUS7 | 516.00 | 546.00 | 598.00 | 553.33 | 662.00 |
| HAUS8 | 138.00 | 166.00 | 174.00 | 159.33 | 247.00 |
| HAVCR2 | 4.00 | 3.00 | 6.00 | 4.33 | 6.00 |
| HAX1 | 2654.00 | 2571.00 | 2712.00 | 2645.67 | 3642.00 |
| HBBP1 | 111.00 | 96.00 | 130.00 | 112.33 | 241.00 |
| HBD | 3.00 | 1.00 | 3.00 | 2.33 | 3.00 |
| HBE1 | 16.00 | 17.00 | 18.00 | 17.00 | 23.00 |
| HBEGF | 7187.00 | 6902.00 | 6884.00 | 6991.00 | 7283.00 |
| HBP1 | 647.00 | 643.00 | 730.00 | 673.33 | 698.00 |
| HBQ1 | 3.00 | 5.00 | 7.00 | 5.00 | 3.00 |
| HBS1L | 1202.00 | 1115.00 | 1271.00 | 1196.00 | 1611.00 |
| HCAR1 | 96.00 | 105.00 | 107.00 | 102.67 | 99.00 |
| HCCS | 452.00 | 414.00 | 457.00 | 441.00 | 518.00 |
| HCFC1 | 2815.00 | 2549.00 | 2905.00 | 2756.33 | 2502.00 |
| HCFC1R1 | 447.00 | 418.00 | 443.00 | 436.00 | 707.00 |
| HCFC2 | 227.00 | 214.00 | 236.00 | 225.67 | 250.00 |
| HCG11 | 18.00 | 14.00 | 10.00 | 14.00 | 15.00 |
| HCG2040054 | 3.00 | 3.00 | 5.00 | 3.67 | 2.00 |
| HCG8 | 7.00 | 16.00 | 20.00 | 14.33 | 18.00 |
| HCLS1 | 3684.00 | 3715.00 | 3989.00 | 3796.00 | 4385.00 |
| HCN2 | 127.00 | 138.00 | 136.00 | 133.67 | 64.00 |
| HCN4 | 3.00 | 0.00 | 4.00 | 2.33 | 0.00 |
| HDAC1 | 2504.00 | 2297.00 | 2589.00 | 2463.33 | 2652.00 |
| HDAC10 | 1245.00 | 1276.00 | 1413.00 | 1311.33 | 1174.00 |
| HDAC11 | 61.00 | 50.00 | 52.00 | 54.33 | 71.00 |
| HDAC2 | 3323.00 | 3112.00 | 3514.00 | 3316.33 | 4260.00 |
| HDAC3 | 1199.00 | 1212.00 | 1311.00 | 1240.67 | 1367.00 |
| HDAC4 | 459.00 | 425.00 | 497.00 | 460.33 | 454.00 |
| HDAC5 | 463.00 | 507.00 | 538.00 | 502.67 | 392.00 |
| HDAC6 | 1292.00 | 1346.00 | 1389.00 | 1342.33 | 1164.00 |
| HDAC7 | 3756.00 | 3696.00 | 3765.00 | 3739.00 | 3804.00 |
| HDAC8 | 371.00 | 396.00 | 424.00 | 397.00 | 440.00 |
| HDAC9 | 664.00 | 613.00 | 741.00 | 672.67 | 961.00 |
| HDDC2 | 964.00 | 863.00 | 1052.00 | 959.67 | 1345.00 |
| HDDC3 | 397.00 | 371.00 | 414.00 | 394.00 | 473.00 |
| HDGF | 7301.00 | 6978.00 | 7293.00 | 7190.67 | 8383.00 |
| HDGFRP2 | 1264.00 | 1402.00 | 1387.00 | 1351.00 | 1546.00 |
| HDGFRP3 | 801.00 | 676.00 | 885.00 | 787.33 | 1106.00 |
| HDHD1 | 511.00 | 430.00 | 461.00 | 467.33 | 556.00 |
| HDHD2 | 456.00 | 534.00 | 612.00 | 534.00 | 565.00 |
| HDHD3 | 264.00 | 280.00 | 282.00 | 275.33 | 314.00 |

| | | | | | |
|---|---|---|---|---|---|
| HDLBP | 12703.00 | 11932.00 | 13877.00 | 12837.33 | 12900.00 |
| HDX | 118.00 | 127.00 | 126.00 | 123.67 | 118.00 |
| HEATR1 | 4326.00 | 3917.00 | 4641.00 | 4294.67 | 4172.00 |
| HEATR3 | 627.00 | 586.00 | 630.00 | 614.33 | 605.00 |
| HEATR5A | 618.00 | 580.00 | 709.00 | 635.67 | 481.00 |
| HEATR5B | 1275.00 | 1284.00 | 1437.00 | 1332.00 | 1222.00 |
| HEATR6 | 2284.00 | 2210.00 | 2497.00 | 2330.33 | 2169.00 |
| HEBP1 | 1053.00 | 979.00 | 1078.00 | 1036.67 | 1333.00 |
| HEBP2 | 470.00 | 446.00 | 496.00 | 470.67 | 514.00 |
| HECA | 487.00 | 482.00 | 514.00 | 494.33 | 528.00 |
| HECTD1 | 4192.00 | 3868.00 | 4516.00 | 4192.00 | 3653.00 |
| HECTD2 | 422.00 | 358.00 | 435.00 | 405.00 | 398.00 |
| HECTD3 | 1138.00 | 1130.00 | 1052.00 | 1106.67 | 971.00 |
| HECTD4 | 1830.00 | 1795.00 | 1997.00 | 1874.00 | 1259.00 |
| HECW2 | 842.00 | 744.00 | 811.00 | 799.00 | 997.00 |
| HEG1 | 2425.00 | 2184.00 | 2646.00 | 2418.33 | 1466.00 |
| HEIH | 400.00 | 395.00 | 396.00 | 397.00 | 483.00 |
| HELB | 67.00 | 72.00 | 69.00 | 69.33 | 73.00 |
| HELLS | 247.00 | 272.00 | 261.00 | 260.00 | 449.00 |
| HELQ | 239.00 | 225.00 | 238.00 | 234.00 | 278.00 |
| HELZ | 968.00 | 862.00 | 981.00 | 937.00 | 846.00 |
| HELZ2 | 1047.00 | 965.00 | 1159.00 | 1057.00 | 1126.00 |
| HEMK1 | 1826.00 | 1844.00 | 1840.00 | 1836.67 | 1461.00 |
| HEPACAM2 | 4.00 | 2.00 | 7.00 | 4.33 | 2.00 |
| HERC1 | 1259.00 | 1232.00 | 1405.00 | 1298.67 | 1099.00 |
| HERC2 | 1869.00 | 1900.00 | 1960.00 | 1909.67 | 1407.00 |
| HERC2P2 | 577.00 | 599.00 | 574.00 | 583.33 | 526.00 |
| HERC2P3 | 266.00 | 242.00 | 298.00 | 268.67 | 165.00 |
| HERC2P4 | 51.00 | 48.00 | 60.00 | 53.00 | 51.00 |
| HERC2P7 | 19.00 | 19.00 | 25.00 | 21.00 | 23.00 |
| HERC3 | 177.00 | 151.00 | 177.00 | 168.33 | 132.00 |
| HERC4 | 3173.00 | 3070.00 | 3437.00 | 3226.67 | 3180.00 |
| HERC5 | 52.00 | 46.00 | 73.00 | 57.00 | 87.00 |
| HERC6 | 137.00 | 99.00 | 116.00 | 117.33 | 130.00 |
| HERPUD1 | 2225.00 | 1990.00 | 2163.00 | 2126.00 | 2348.00 |
| HERPUD2 | 931.00 | 835.00 | 962.00 | 909.33 | 857.00 |
| HES1 | 158.00 | 194.00 | 211.00 | 187.67 | 123.00 |
| HES2 | 3.00 | 1.00 | 0.00 | 1.33 | 0.00 |
| HES4 | 67.00 | 68.00 | 92.00 | 75.67 | 96.00 |
| HES6 | 72.00 | 86.00 | 80.00 | 79.33 | 149.00 |
| HES7 | 29.00 | 16.00 | 32.00 | 25.67 | 26.00 |
| HESX1 | 11.00 | 22.00 | 26.00 | 19.67 | 39.00 |
| HEXA | 5530.00 | 5563.00 | 5466.00 | 5519.67 | 5359.00 |
| HEXA-AS1 | 14.00 | 12.00 | 11.00 | 12.33 | 8.00 |
| HEXB | 4749.00 | 4920.00 | 4583.00 | 4750.67 | 5327.00 |
| HEXDC | 498.00 | 484.00 | 515.00 | 499.00 | 513.00 |
| HEXIM1 | 587.00 | 621.00 | 665.00 | 624.33 | 731.00 |
| HEXIM2 | 75.00 | 109.00 | 89.00 | 91.00 | 108.00 |

| | | | | | |
|---|---|---|---|---|---|
| HEY1 | 496.00 | 505.00 | 527.00 | 509.33 | 567.00 |
| HEY2 | 68.00 | 62.00 | 67.00 | 65.67 | 73.00 |
| HFE | 322.00 | 376.00 | 338.00 | 345.33 | 393.00 |
| HFM1 | 38.00 | 40.00 | 48.00 | 42.00 | 60.00 |
| HGD | 4.00 | 2.00 | 0.00 | 2.00 | 1.00 |
| HGFAC | 8.00 | 12.00 | 19.00 | 13.00 | 6.00 |
| HGH1 | 710.00 | 658.00 | 706.00 | 691.33 | 757.00 |
| HGS | 9123.00 | 8879.00 | 9482.00 | 9161.33 | 8775.00 |
| HGSNAT | 1586.00 | 1562.00 | 1615.00 | 1587.67 | 1548.00 |
| HHAT | 262.00 | 300.00 | 283.00 | 281.67 | 212.00 |
| HHEX | 222.00 | 208.00 | 228.00 | 219.33 | 358.00 |
| HHIP | 456.00 | 429.00 | 467.00 | 450.67 | 496.00 |
| HHIP-AS1 | 103.00 | 102.00 | 137.00 | 114.00 | 115.00 |
| HHIPL1 | 19.00 | 23.00 | 25.00 | 22.33 | 16.00 |
| HHIPL2 | 10.00 | 15.00 | 21.00 | 15.33 | 19.00 |
| HHLA3 | 207.00 | 210.00 | 211.00 | 209.33 | 290.00 |
| HIAT1 | 1249.00 | 1199.00 | 1263.00 | 1237.00 | 1572.00 |
| HIATL1 | 1384.00 | 1422.00 | 1404.00 | 1403.33 | 1595.00 |
| HIATL2 | 97.00 | 92.00 | 74.00 | 87.67 | 71.00 |
| HIBADH | 1331.00 | 1218.00 | 1484.00 | 1344.33 | 1846.00 |
| HIBCH | 425.00 | 398.00 | 408.00 | 410.33 | 487.00 |
| HIC1 | 147.00 | 174.00 | 193.00 | 171.33 | 133.00 |
| HIC2 | 165.00 | 172.00 | 174.00 | 170.33 | 175.00 |
| HID1 | 9.00 | 15.00 | 8.00 | 10.67 | 22.00 |
| HIF1A | 8733.00 | 7513.00 | 9423.00 | 8556.33 | 6408.00 |
| HIF1A-AS1 | 8.00 | 3.00 | 13.00 | 8.00 | 13.00 |
| HIF1AN | 1982.00 | 1995.00 | 2178.00 | 2051.67 | 1915.00 |
| HIGD1A | 1086.00 | 1067.00 | 1122.00 | 1091.67 | 1514.00 |
| HIGD2A | 574.00 | 597.00 | 631.00 | 600.67 | 822.00 |
| HILPDA | 349.00 | 333.00 | 327.00 | 336.33 | 433.00 |
| HINFP | 478.00 | 504.00 | 511.00 | 497.67 | 555.00 |
| HINT1 | 4453.00 | 4387.00 | 4720.00 | 4520.00 | 5937.00 |
| HINT2 | 473.00 | 527.00 | 487.00 | 495.67 | 612.00 |
| HINT3 | 538.00 | 522.00 | 552.00 | 537.33 | 637.00 |
| HIP1 | 1636.00 | 1568.00 | 1698.00 | 1634.00 | 1179.00 |
| HIP1R | 781.00 | 834.00 | 826.00 | 813.67 | 673.00 |
| HIPK1 | 1123.00 | 989.00 | 1298.00 | 1136.67 | 932.00 |
| HIPK1-AS1 | 18.00 | 13.00 | 13.00 | 14.67 | 9.00 |
| HIPK2 | 5935.00 | 5161.00 | 6070.00 | 5722.00 | 5217.00 |
| HIPK3 | 862.00 | 773.00 | 883.00 | 839.33 | 779.00 |
| HIRA | 1121.00 | 1127.00 | 1130.00 | 1126.00 | 1262.00 |
| HIRIP3 | 427.00 | 446.00 | 408.00 | 427.00 | 566.00 |
| HIST1H1C | 24.00 | 20.00 | 20.00 | 21.33 | 23.00 |
| HIST1H2AC | 13.00 | 9.00 | 15.00 | 12.33 | 12.00 |
| HIST1H2AE | 3.00 | 1.00 | 6.00 | 3.33 | 2.00 |
| HIST1H2AK | 6.00 | 5.00 | 4.00 | 5.00 | 8.00 |
| HIST1H2AM | 3.00 | 0.00 | 0.00 | 1.00 | 3.00 |
| HIST1H2BC | 3.00 | 1.00 | 4.00 | 2.67 | 1.00 |

| | | | | | |
|---|---|---|---|---|---|
| HIST1H2BD | 23.00 | 16.00 | 15.00 | 18.00 | 23.00 |
| HIST1H2BH | 7.00 | 2.00 | 5.00 | 4.67 | 2.00 |
| HIST1H2BK | 217.00 | 214.00 | 247.00 | 226.00 | 244.00 |
| HIST1H2BN | 7.00 | 17.00 | 12.00 | 12.00 | 16.00 |
| HIST1H3B | 3.00 | 1.00 | 1.00 | 1.67 | 0.00 |
| HIST1H3D | 4.00 | 3.00 | 1.00 | 2.67 | 1.00 |
| HIST1H3E | 17.00 | 4.00 | 12.00 | 11.00 | 12.00 |
| HIST2H2BA | 10.00 | 21.00 | 21.00 | 17.33 | 9.00 |
| HIST2H2BC | 8.00 | 8.00 | 2.00 | 6.00 | 3.00 |
| HIST2H2BE | 9.00 | 8.00 | 18.00 | 11.67 | 22.00 |
| HIST2H2BF | 4.00 | 1.00 | 1.00 | 2.00 | 2.00 |
| HIST3H2A | 41.00 | 45.00 | 50.00 | 45.33 | 44.00 |
| HIST4H4 | 6.00 | 9.00 | 10.00 | 8.33 | 4.00 |
| HIVEP1 | 1076.00 | 875.00 | 1155.00 | 1035.33 | 645.00 |
| HIVEP2 | 1408.00 | 1199.00 | 1449.00 | 1352.00 | 1074.00 |
| HIVEP3 | 473.00 | 466.00 | 453.00 | 464.00 | 489.00 |
| HJURP | 1378.00 | 1465.00 | 1468.00 | 1437.00 | 1879.00 |
| HK1 | 4946.00 | 4884.00 | 5177.00 | 5002.33 | 4923.00 |
| HK2 | 4882.00 | 4636.00 | 5048.00 | 4855.33 | 5632.00 |
| HKDC1 | 203.00 | 185.00 | 208.00 | 198.67 | 132.00 |
| HKR1 | 513.00 | 575.00 | 508.00 | 532.00 | 587.00 |
| HLA-A | 2475.00 | 2996.00 | 2587.00 | 2686.00 | 2678.00 |
| HLA-B | 381.00 | 376.00 | 329.00 | 362.00 | 365.00 |
| HLA-C | 883.00 | 804.00 | 831.00 | 839.33 | 782.00 |
| HLA-DMA | 10.00 | 8.00 | 5.00 | 7.67 | 11.00 |
| HLA-DRB1 | 74.00 | 94.00 | 85.00 | 84.33 | 133.00 |
| HLA-DRB5 | 5.00 | 6.00 | 6.00 | 5.67 | 7.00 |
| HLA-H | 4.00 | 0.00 | 1.00 | 1.67 | 2.00 |
| HLCS | 1280.00 | 1146.00 | 1242.00 | 1222.67 | 1237.00 |
| HLF | 30.00 | 16.00 | 18.00 | 21.33 | 32.00 |
| HLTF | 2526.00 | 2376.00 | 2642.00 | 2514.67 | 2854.00 |
| HLX | 246.00 | 284.00 | 290.00 | 273.33 | 260.00 |
| HLX-AS1 | 11.00 | 15.00 | 14.00 | 13.33 | 9.00 |
| HM13 | 5835.00 | 6027.00 | 6018.00 | 5960.00 | 5487.00 |
| HMBOX1 | 111.00 | 87.00 | 102.00 | 100.00 | 53.00 |
| HMBS | 692.00 | 658.00 | 731.00 | 693.67 | 861.00 |
| HMCES | 563.00 | 561.00 | 567.00 | 563.67 | 815.00 |
| HMCN1 | 78.00 | 48.00 | 77.00 | 67.67 | 61.00 |
| HMCN2 | 7.00 | 5.00 | 0.00 | 4.00 | 8.00 |
| HMG20A | 961.00 | 823.00 | 931.00 | 905.00 | 917.00 |
| HMG20B | 1467.00 | 1608.00 | 1699.00 | 1591.33 | 1974.00 |
| HMGA1 | 41750.00 | 40059.00 | 43831.00 | 41880.00 | 35150.00 |
| HMGA2 | 5156.00 | 5100.00 | 5891.00 | 5382.33 | 5253.00 |
| HMGB1 | 3412.00 | 3475.00 | 3497.00 | 3461.33 | 4932.00 |
| HMGB2 | 2454.00 | 2342.00 | 2513.00 | 2436.33 | 3811.00 |
| HMGB3 | 500.00 | 470.00 | 514.00 | 494.67 | 697.00 |
| HMGCL | 674.00 | 651.00 | 719.00 | 681.33 | 720.00 |
| HMGCR | 1289.00 | 1216.00 | 1276.00 | 1260.33 | 1336.00 |

| | | | | | |
|---|---|---|---|---|---|
| HMGCS1 | 629.00 | 573.00 | 673.00 | 625.00 | 688.00 |
| HMGN1 | 2659.00 | 2689.00 | 2807.00 | 2718.33 | 3645.00 |
| HMGN2 | 2329.00 | 2272.00 | 2424.00 | 2341.67 | 3068.00 |
| HMGN3 | 912.00 | 852.00 | 935.00 | 899.67 | 939.00 |
| HMGN3-AS1 | 32.00 | 19.00 | 17.00 | 22.67 | 17.00 |
| HMGN4 | 1974.00 | 1798.00 | 2174.00 | 1982.00 | 2162.00 |
| HMGN5 | 295.00 | 287.00 | 357.00 | 313.00 | 483.00 |
| HMGXB3 | 1874.00 | 1780.00 | 1971.00 | 1875.00 | 1845.00 |
| HMGXB4 | 900.00 | 824.00 | 946.00 | 890.00 | 913.00 |
| HMHA1 | 81.00 | 94.00 | 109.00 | 94.67 | 63.00 |
| HMMR | 930.00 | 798.00 | 890.00 | 872.67 | 1022.00 |
| HMOX1 | 305.00 | 369.00 | 354.00 | 342.67 | 622.00 |
| HN1 | 787.00 | 706.00 | 843.00 | 778.67 | 1056.00 |
| HN1L | 1346.00 | 1307.00 | 1480.00 | 1377.67 | 1416.00 |
| HNF4G | 69.00 | 67.00 | 80.00 | 72.00 | 121.00 |
| HNMT | 3.00 | 3.00 | 0.00 | 2.00 | 2.00 |
| HNRNPA0 | 3181.00 | 3094.00 | 3243.00 | 3172.67 | 4017.00 |
| HNRNPA1 | 9531.00 | 9447.00 | 10328.00 | 9768.67 | 12823.00 |
| HNRNPA1L2 | 252.00 | 212.00 | 223.00 | 229.00 | 222.00 |
| HNRNPA1P10 | 1833.00 | 1858.00 | 1954.00 | 1881.67 | 2808.00 |
| HNRNPA2B1 | 23910.00 | 23772.00 | 24633.00 | 24105.00 | 30997.00 |
| HNRNPA3 | 9827.00 | 9893.00 | 10344.00 | 10021.33 | 11659.00 |
| HNRNPAB | 3987.00 | 3910.00 | 4257.00 | 4051.33 | 5608.00 |
| HNRNPC | 12066.00 | 11414.00 | 12741.00 | 12073.67 | 15797.00 |
| HNRNPD | 4735.00 | 4526.00 | 4669.00 | 4643.33 | 6272.00 |
| HNRNPDL | 6041.00 | 6065.00 | 6258.00 | 6121.33 | 6983.00 |
| HNRNPF | 6650.00 | 6482.00 | 7164.00 | 6765.33 | 8291.00 |
| HNRNPH1 | 14022.00 | 13790.00 | 14890.00 | 14234.00 | 15269.00 |
| HNRNPH2 | 31.00 | 23.00 | 34.00 | 29.33 | 40.00 |
| HNRNPH3 | 5234.00 | 5229.00 | 5558.00 | 5340.33 | 6680.00 |
| HNRNPK | 13327.00 | 12734.00 | 14074.00 | 13378.33 | 15680.00 |
| HNRNPL | 4309.00 | 4224.00 | 4464.00 | 4332.33 | 5409.00 |
| HNRNPLL | 628.00 | 601.00 | 685.00 | 638.00 | 696.00 |
| HNRNPM | 6154.00 | 6037.00 | 6322.00 | 6171.00 | 7446.00 |
| HNRNPR | 3903.00 | 3852.00 | 3992.00 | 3915.67 | 4672.00 |
| HNRNPU | 13045.00 | 12455.00 | 13402.00 | 12967.33 | 16826.00 |
| HNRNPU-AS1 | 2390.00 | 2494.00 | 2872.00 | 2585.33 | 2306.00 |
| HNRNPUL1 | 4264.00 | 4023.00 | 4423.00 | 4236.67 | 4638.00 |
| HNRNPUL2 | 1260.00 | 1222.00 | 1349.00 | 1277.00 | 1440.00 |
| HOGA1 | 40.00 | 33.00 | 36.00 | 36.33 | 42.00 |
| HOMER1 | 281.00 | 237.00 | 298.00 | 272.00 | 287.00 |
| HOMER2 | 57.00 | 32.00 | 51.00 | 46.67 | 32.00 |
| HOMER3 | 1087.00 | 1143.00 | 1137.00 | 1122.33 | 1348.00 |
| HOMEZ | 312.00 | 273.00 | 275.00 | 286.67 | 255.00 |
| HOOK2 | 749.00 | 782.00 | 806.00 | 779.00 | 676.00 |
| HOOK3 | 809.00 | 697.00 | 839.00 | 781.67 | 798.00 |
| HOPX | 1257.00 | 1193.00 | 1330.00 | 1260.00 | 1459.00 |
| HOTAIR | 21.00 | 26.00 | 32.00 | 26.33 | 18.00 |

| | | | | | |
|---|---|---|---|---|---|
| HOTAIRM1 | 812.00 | 796.00 | 874.00 | 827.33 | 815.00 |
| HOXA1 | 128.00 | 111.00 | 136.00 | 125.00 | 182.00 |
| HOXA10 | 201.00 | 206.00 | 229.00 | 212.00 | 306.00 |
| HOXA11 | 59.00 | 65.00 | 61.00 | 61.67 | 99.00 |
| HOXA11-AS | 168.00 | 123.00 | 134.00 | 141.67 | 165.00 |
| HOXA13 | 13.00 | 2.00 | 12.00 | 9.00 | 11.00 |
| HOXA2 | 26.00 | 36.00 | 22.00 | 28.00 | 20.00 |
| HOXA3 | 157.00 | 142.00 | 137.00 | 145.33 | 171.00 |
| HOXA4 | 116.00 | 150.00 | 142.00 | 136.00 | 172.00 |
| HOXA5 | 83.00 | 93.00 | 79.00 | 85.00 | 88.00 |
| HOXA6 | 15.00 | 22.00 | 19.00 | 18.67 | 19.00 |
| HOXA7 | 120.00 | 152.00 | 104.00 | 125.33 | 154.00 |
| HOXA9 | 5.00 | 4.00 | 0.00 | 3.00 | 12.00 |
| HOXA-AS2 | 49.00 | 62.00 | 47.00 | 52.67 | 56.00 |
| HOXB2 | 1053.00 | 1041.00 | 1080.00 | 1058.00 | 1651.00 |
| HOXB3 | 1117.00 | 1088.00 | 1166.00 | 1123.67 | 1170.00 |
| HOXB4 | 454.00 | 514.00 | 479.00 | 482.33 | 577.00 |
| HOXB5 | 385.00 | 334.00 | 403.00 | 374.00 | 398.00 |
| HOXB6 | 828.00 | 788.00 | 867.00 | 827.67 | 785.00 |
| HOXB7 | 467.00 | 448.00 | 484.00 | 466.33 | 613.00 |
| HOXB8 | 235.00 | 219.00 | 205.00 | 219.67 | 246.00 |
| HOXB9 | 2348.00 | 2242.00 | 2452.00 | 2347.33 | 2516.00 |
| HOXB-AS1 | 523.00 | 641.00 | 656.00 | 606.67 | 626.00 |
| HOXC10 | 22.00 | 28.00 | 21.00 | 23.67 | 37.00 |
| HOXC11 | 36.00 | 41.00 | 37.00 | 38.00 | 56.00 |
| HOXC13 | 48.00 | 39.00 | 40.00 | 42.33 | 55.00 |
| HOXC4 | 236.00 | 242.00 | 246.00 | 241.33 | 266.00 |
| HOXC5 | 75.00 | 58.00 | 64.00 | 65.67 | 66.00 |
| HOXC6 | 426.00 | 417.00 | 418.00 | 420.33 | 398.00 |
| HOXC8 | 433.00 | 458.00 | 518.00 | 469.67 | 418.00 |
| HOXC9 | 152.00 | 143.00 | 167.00 | 154.00 | 205.00 |
| HOXC-AS1 | 22.00 | 29.00 | 47.00 | 32.67 | 35.00 |
| HOXD1 | 10.00 | 9.00 | 13.00 | 10.67 | 11.00 |
| HOXD13 | 79.00 | 61.00 | 78.00 | 72.67 | 93.00 |
| HOXD3 | 232.00 | 265.00 | 281.00 | 259.33 | 250.00 |
| HOXD4 | 187.00 | 181.00 | 196.00 | 188.00 | 205.00 |
| HOXD8 | 274.00 | 276.00 | 315.00 | 288.33 | 330.00 |
| HOXD9 | 221.00 | 281.00 | 253.00 | 251.67 | 303.00 |
| HOXD-AS2 | 90.00 | 99.00 | 95.00 | 94.67 | 137.00 |
| HP09053 | 4.00 | 12.00 | 5.00 | 7.00 | 6.00 |
| HP1BP3 | 6932.00 | 6635.00 | 7408.00 | 6991.67 | 6334.00 |
| HPCAL1 | 1498.00 | 1449.00 | 1529.00 | 1492.00 | 1478.00 |
| HPDL | 320.00 | 310.00 | 325.00 | 318.33 | 510.00 |
| HPRT1 | 514.00 | 451.00 | 555.00 | 506.67 | 729.00 |
| HPS1 | 1100.00 | 1110.00 | 1213.00 | 1141.00 | 1233.00 |
| HPS3 | 1474.00 | 1374.00 | 1494.00 | 1447.33 | 1422.00 |
| HPS4 | 1451.00 | 1400.00 | 1551.00 | 1467.33 | 1316.00 |
| HPS5 | 690.00 | 603.00 | 670.00 | 654.33 | 784.00 |

| | | | | | |
|---|---|---|---|---|---|
| HPS6 | 592.00 | 579.00 | 632.00 | 601.00 | 635.00 |
| HPSE | 264.00 | 245.00 | 295.00 | 268.00 | 308.00 |
| HPX | 8.00 | 9.00 | 7.00 | 8.00 | 14.00 |
| HRASLS | 96.00 | 93.00 | 99.00 | 96.00 | 123.00 |
| HRASLS2 | 3.00 | 0.00 | 0.00 | 1.00 | 0.00 |
| HRAT5 | 4.00 | 3.00 | 2.00 | 3.00 | 8.00 |
| HRAT92 | 225.00 | 266.00 | 259.00 | 250.00 | 131.00 |
| HRH1 | 770.00 | 741.00 | 752.00 | 754.33 | 565.00 |
| HRSP12 | 348.00 | 326.00 | 419.00 | 364.33 | 527.00 |
| HS1BP3 | 481.00 | 513.00 | 541.00 | 511.67 | 530.00 |
| HS1BP3-IT1 | 3.00 | 1.00 | 5.00 | 3.00 | 3.00 |
| HS2ST1 | 1486.00 | 1375.00 | 1610.00 | 1490.33 | 1687.00 |
| HS3ST1 | 29.00 | 47.00 | 18.00 | 31.33 | 26.00 |
| HS3ST3A1 | 9.00 | 3.00 | 4.00 | 5.33 | 6.00 |
| HS3ST5 | 20.00 | 20.00 | 29.00 | 23.00 | 24.00 |
| HS6ST1 | 266.00 | 298.00 | 295.00 | 286.33 | 337.00 |
| HS6ST2 | 5.00 | 1.00 | 9.00 | 5.00 | 3.00 |
| HS6ST3 | 57.00 | 57.00 | 56.00 | 56.67 | 27.00 |
| HSBP1 | 1940.00 | 1836.00 | 1947.00 | 1907.67 | 2703.00 |
| HSBP1L1 | 53.00 | 38.00 | 41.00 | 44.00 | 34.00 |
| HSCB | 207.00 | 203.00 | 185.00 | 198.33 | 271.00 |
| HSD11B1 | 17.00 | 16.00 | 13.00 | 15.33 | 7.00 |
| HSD11B1L | 200.00 | 190.00 | 186.00 | 192.00 | 238.00 |
| HSD17B1 | 30.00 | 32.00 | 44.00 | 35.33 | 36.00 |
| HSD17B10 | 676.00 | 662.00 | 683.00 | 673.67 | 906.00 |
| HSD17B11 | 968.00 | 930.00 | 1035.00 | 977.67 | 1268.00 |
| HSD17B12 | 1964.00 | 1966.00 | 1991.00 | 1973.67 | 2382.00 |
| HSD17B13 | 3.00 | 5.00 | 5.00 | 4.33 | 3.00 |
| HSD17B14 | 55.00 | 71.00 | 63.00 | 63.00 | 46.00 |
| HSD17B3 | 4.00 | 2.00 | 4.00 | 3.33 | 7.00 |
| HSD17B4 | 1546.00 | 1571.00 | 1801.00 | 1639.33 | 1792.00 |
| HSD17B6 | 19.00 | 20.00 | 29.00 | 22.67 | 20.00 |
| HSD17B7 | 84.00 | 72.00 | 76.00 | 77.33 | 92.00 |
| HSD17B7P2 | 56.00 | 68.00 | 61.00 | 61.67 | 40.00 |
| HSD3B7 | 112.00 | 106.00 | 153.00 | 123.67 | 149.00 |
| HSD52 | 5.00 | 6.00 | 6.00 | 5.67 | 7.00 |
| HSDL1 | 1043.00 | 941.00 | 1231.00 | 1071.67 | 1074.00 |
| HSDL2 | 484.00 | 475.00 | 565.00 | 508.00 | 583.00 |
| HSF1 | 2685.00 | 2597.00 | 2805.00 | 2695.67 | 2679.00 |
| HSF2 | 654.00 | 607.00 | 777.00 | 679.33 | 744.00 |
| HSF2BP | 49.00 | 31.00 | 48.00 | 42.67 | 36.00 |
| HSF4 | 1315.00 | 1278.00 | 1427.00 | 1340.00 | 1148.00 |
| HSH2D | 6.00 | 10.00 | 11.00 | 9.00 | 6.00 |
| HSP90AA1 | 21616.00 | 19963.00 | 21903.00 | 21160.67 | 27257.00 |
| HSP90AB1 | 48643.00 | 45977.00 | 50481.00 | 48367.00 | 55257.00 |
| HSP90B1 | 38685.00 | 39220.00 | 39708.00 | 39204.33 | 38475.00 |
| HSP90B2P | 186.00 | 161.00 | 175.00 | 174.00 | 157.00 |
| HSPA12A | 49.00 | 51.00 | 45.00 | 48.33 | 38.00 |

| | | | | | |
|---|---|---|---|---|---|
| HSPA13 | 3762.00 | 3278.00 | 3567.00 | 3535.67 | 3454.00 |
| HSPA14 | 1384.00 | 1270.00 | 1381.00 | 1345.00 | 1617.00 |
| HSPA1A | 22.00 | 29.00 | 33.00 | 28.00 | 17.00 |
| HSPA1B | 7.00 | 11.00 | 11.00 | 9.67 | 11.00 |
| HSPA2 | 105.00 | 104.00 | 84.00 | 97.67 | 108.00 |
| HSPA4 | 4258.00 | 3919.00 | 4540.00 | 4239.00 | 5537.00 |
| HSPA4L | 243.00 | 245.00 | 286.00 | 258.00 | 297.00 |
| HSPA5 | 30237.00 | 30313.00 | 30545.00 | 30365.00 | 25526.00 |
| HSPA8 | 26188.00 | 26586.00 | 30814.00 | 27862.67 | 29937.00 |
| HSPA9 | 15888.00 | 14728.00 | 16612.00 | 15742.67 | 18843.00 |
| HSPB1 | 2109.00 | 1992.00 | 2195.00 | 2098.67 | 3300.00 |
| HSPB11 | 224.00 | 233.00 | 257.00 | 238.00 | 330.00 |
| HSPBAP1 | 417.00 | 429.00 | 531.00 | 459.00 | 612.00 |
| HSPBP1 | 833.00 | 848.00 | 875.00 | 852.00 | 972.00 |
| HSPD1 | 14166.00 | 13240.00 | 15067.00 | 14157.67 | 16246.00 |
| HSPE1 | 1100.00 | 1095.00 | 1090.00 | 1095.00 | 1546.00 |
| HSPG2 | 2354.00 | 2208.00 | 2244.00 | 2268.67 | 1548.00 |
| HSPH1 | 3167.00 | 3089.00 | 3418.00 | 3224.67 | 3939.00 |
| HTATIP2 | 899.00 | 914.00 | 990.00 | 934.33 | 1037.00 |
| HTATSF1 | 1257.00 | 1292.00 | 1335.00 | 1294.67 | 1519.00 |
| HTATSF1P2 | 281.00 | 299.00 | 269.00 | 283.00 | 279.00 |
| HTR1B | 7.00 | 3.00 | 0.00 | 3.33 | 2.00 |
| HTR4 | 3.00 | 4.00 | 2.00 | 3.00 | 0.00 |
| HTR7 | 624.00 | 582.00 | 643.00 | 616.33 | 618.00 |
| HTR7P1 | 253.00 | 249.00 | 256.00 | 252.67 | 264.00 |
| HTRA1 | 261.00 | 280.00 | 300.00 | 280.33 | 339.00 |
| HTRA2 | 621.00 | 682.00 | 722.00 | 675.00 | 767.00 |
| HTRA4 | 3.00 | 0.00 | 4.00 | 2.33 | 4.00 |
| HTT | 4048.00 | 3893.00 | 4190.00 | 4043.67 | 3150.00 |
| HULC | 4.00 | 7.00 | 8.00 | 6.33 | 4.00 |
| HUS1 | 355.00 | 385.00 | 432.00 | 390.67 | 395.00 |
| HUWE1 | 9571.00 | 8615.00 | 9698.00 | 9294.67 | 7128.00 |
| HVCN1 | 4.00 | 5.00 | 4.00 | 4.33 | 4.00 |
| HYAL1 | 10.00 | 13.00 | 14.00 | 12.33 | 6.00 |
| HYAL2 | 1171.00 | 1260.00 | 1131.00 | 1187.33 | 1213.00 |
| HYAL3 | 216.00 | 218.00 | 259.00 | 231.00 | 176.00 |
| HYI | 2118.00 | 2077.00 | 2278.00 | 2157.67 | 1868.00 |
| HYKK | 54.00 | 56.00 | 85.00 | 65.00 | 72.00 |
| HYLS1 | 187.00 | 189.00 | 188.00 | 188.00 | 262.00 |
| HYOU1 | 7543.00 | 7577.00 | 8007.00 | 7709.00 | 6495.00 |
| IAH1 | 492.00 | 459.00 | 527.00 | 492.67 | 602.00 |
| IARS | 9773.00 | 9102.00 | 9944.00 | 9606.33 | 9604.00 |
| IARS2 | 3465.00 | 3202.00 | 3549.00 | 3405.33 | 3941.00 |
| IBA57 | 249.00 | 212.00 | 202.00 | 221.00 | 198.00 |
| IBA57-AS1 | 4.00 | 4.00 | 1.00 | 3.00 | 2.00 |
| IBTK | 2113.00 | 1822.00 | 2180.00 | 2038.33 | 1798.00 |
| ICA1 | 507.00 | 445.00 | 496.00 | 482.67 | 586.00 |
| ICA1L | 111.00 | 120.00 | 126.00 | 119.00 | 142.00 |

| | | | | | |
|---|---|---|---|---|---|
| ICAM1 | 5162.00 | 5203.00 | 4895.00 | 5086.67 | 5768.00 |
| ICAM3 | 137.00 | 150.00 | 131.00 | 139.33 | 198.00 |
| ICAM4 | 7.00 | 5.00 | 5.00 | 5.67 | 10.00 |
| ICAM5 | 1482.00 | 1601.00 | 1774.00 | 1619.00 | 1487.00 |
| ICE1 | 2504.00 | 2320.00 | 2690.00 | 2504.67 | 2296.00 |
| ICE2 | 1389.00 | 1364.00 | 1521.00 | 1424.67 | 1408.00 |
| ICK | 299.00 | 275.00 | 354.00 | 309.33 | 284.00 |
| ICMT | 1937.00 | 1758.00 | 1889.00 | 1861.33 | 1943.00 |
| ICT1 | 419.00 | 379.00 | 430.00 | 409.33 | 531.00 |
| ID1 | 234.00 | 290.00 | 240.00 | 254.67 | 344.00 |
| ID2 | 5.00 | 10.00 | 4.00 | 6.33 | 6.00 |
| ID3 | 442.00 | 468.00 | 479.00 | 463.00 | 518.00 |
| IDE | 1316.00 | 1163.00 | 1422.00 | 1300.33 | 1281.00 |
| IDH1 | 1553.00 | 1533.00 | 1626.00 | 1570.67 | 1816.00 |
| IDH1-AS1 | 4.00 | 6.00 | 4.00 | 4.67 | 8.00 |
| IDH2 | 345.00 | 344.00 | 312.00 | 333.67 | 484.00 |
| IDH3A | 942.00 | 912.00 | 1017.00 | 957.00 | 1047.00 |
| IDH3B | 1572.00 | 1614.00 | 1657.00 | 1614.33 | 1763.00 |
| IDH3G | 661.00 | 671.00 | 721.00 | 684.33 | 763.00 |
| IDI1 | 427.00 | 467.00 | 506.00 | 466.67 | 545.00 |
| IDI2-AS1 | 50.00 | 46.00 | 43.00 | 46.33 | 55.00 |
| IDNK | 71.00 | 66.00 | 53.00 | 63.33 | 66.00 |
| IDS | 2531.00 | 2267.00 | 2726.00 | 2508.00 | 1684.00 |
| IDUA | 370.00 | 430.00 | 447.00 | 415.67 | 357.00 |
| IER2 | 2360.00 | 2295.00 | 2367.00 | 2340.67 | 3024.00 |
| IER3IP1 | 732.00 | 695.00 | 772.00 | 733.00 | 1072.00 |
| IER5 | 1166.00 | 1179.00 | 1223.00 | 1189.33 | 1161.00 |
| IER5L | 62.00 | 56.00 | 84.00 | 67.33 | 65.00 |
| IFFO1 | 1198.00 | 1213.00 | 1327.00 | 1246.00 | 1193.00 |
| IFFO2 | 549.00 | 482.00 | 632.00 | 554.33 | 581.00 |
| IFI16 | 6518.00 | 6128.00 | 6709.00 | 6451.67 | 5927.00 |
| IFI27L1 | 14.00 | 7.00 | 14.00 | 11.67 | 17.00 |
| IFI30 | 40.00 | 42.00 | 43.00 | 41.67 | 23.00 |
| IFI35 | 136.00 | 122.00 | 111.00 | 123.00 | 177.00 |
| IFI44 | 579.00 | 599.00 | 560.00 | 579.33 | 555.00 |
| IFI44L | 90.00 | 101.00 | 109.00 | 100.00 | 50.00 |
| IFI6 | 105.00 | 98.00 | 82.00 | 95.00 | 150.00 |
| IFIH1 | 80.00 | 85.00 | 96.00 | 87.00 | 118.00 |
| IFIT1 | 17.00 | 7.00 | 13.00 | 12.33 | 9.00 |
| IFIT2 | 59.00 | 54.00 | 67.00 | 60.00 | 58.00 |
| IFIT3 | 157.00 | 173.00 | 192.00 | 174.00 | 196.00 |
| IFIT5 | 681.00 | 636.00 | 772.00 | 696.33 | 808.00 |
| IFITM1 | 53.00 | 64.00 | 66.00 | 61.00 | 67.00 |
| IFITM2 | 724.00 | 718.00 | 802.00 | 748.00 | 990.00 |
| IFITM3 | 870.00 | 796.00 | 869.00 | 845.00 | 1071.00 |
| IFNAR1 | 1061.00 | 902.00 | 886.00 | 949.67 | 1064.00 |
| IFNAR2 | 310.00 | 309.00 | 343.00 | 320.67 | 436.00 |
| IFNE | 382.00 | 416.00 | 473.00 | 423.67 | 314.00 |

| | | | | | |
|---|---|---|---|---|---|
| IFNGR1 | 1166.00 | 1138.00 | 1212.00 | 1172.00 | 1223.00 |
| IFNGR2 | 33.00 | 44.00 | 57.00 | 44.67 | 40.00 |
| IFRD1 | 2182.00 | 1949.00 | 2202.00 | 2111.00 | 2547.00 |
| IFRD2 | 2234.00 | 2288.00 | 2281.00 | 2267.67 | 2797.00 |
| IFT122 | 338.00 | 400.00 | 393.00 | 377.00 | 424.00 |
| IFT140 | 900.00 | 931.00 | 971.00 | 934.00 | 912.00 |
| IFT172 | 1397.00 | 1381.00 | 1421.00 | 1399.67 | 1266.00 |
| IFT20 | 420.00 | 437.00 | 439.00 | 432.00 | 404.00 |
| IFT22 | 162.00 | 152.00 | 169.00 | 161.00 | 214.00 |
| IFT27 | 366.00 | 370.00 | 377.00 | 371.00 | 402.00 |
| IFT43 | 294.00 | 298.00 | 271.00 | 287.67 | 290.00 |
| IFT46 | 464.00 | 490.00 | 542.00 | 498.67 | 559.00 |
| IFT52 | 337.00 | 360.00 | 373.00 | 356.67 | 421.00 |
| IFT57 | 773.00 | 673.00 | 814.00 | 753.33 | 985.00 |
| IFT74 | 440.00 | 418.00 | 450.00 | 436.00 | 545.00 |
| IFT80 | 1444.00 | 1417.00 | 1511.00 | 1457.33 | 1324.00 |
| IFT81 | 571.00 | 616.00 | 621.00 | 602.67 | 663.00 |
| IFT88 | 358.00 | 351.00 | 335.00 | 348.00 | 429.00 |
| IGBP1 | 1367.00 | 1386.00 | 1487.00 | 1413.33 | 1594.00 |
| IGDCC4 | 3.00 | 8.00 | 3.00 | 4.67 | 3.00 |
| IGF1R | 2273.00 | 2342.00 | 2432.00 | 2349.00 | 2093.00 |
| IGF2BP1 | 2516.00 | 2359.00 | 2662.00 | 2512.33 | 2357.00 |
| IGF2BP2 | 6153.00 | 5968.00 | 6301.00 | 6140.67 | 6383.00 |
| IGF2BP2-AS1 | 55.00 | 74.00 | 52.00 | 60.33 | 77.00 |
| IGF2BP3 | 3026.00 | 2671.00 | 3072.00 | 2923.00 | 3185.00 |
| IGF2R | 12203.00 | 11518.00 | 12384.00 | 12035.00 | 8516.00 |
| IGFBP1 | 6.00 | 14.00 | 3.00 | 7.67 | 16.00 |
| IGFBP3 | 126.00 | 104.00 | 134.00 | 121.33 | 785.00 |
| IGFBP4 | 41.00 | 36.00 | 45.00 | 40.67 | 38.00 |
| IGFBP6 | 392.00 | 503.00 | 444.00 | 446.33 | 576.00 |
| IGFBP7 | 471.00 | 459.00 | 464.00 | 464.67 | 276.00 |
| IGFLR1 | 22.00 | 32.00 | 22.00 | 25.33 | 28.00 |
| IGFN1 | 235.00 | 239.00 | 246.00 | 240.00 | 124.00 |
| IGHMBP2 | 543.00 | 552.00 | 610.00 | 568.33 | 619.00 |
| IGIP | 186.00 | 172.00 | 248.00 | 202.00 | 174.00 |
| IGSF1 | 5.00 | 5.00 | 4.00 | 4.67 | 5.00 |
| IGSF10 | 1061.00 | 1006.00 | 1106.00 | 1057.67 | 738.00 |
| IGSF23 | 6.00 | 4.00 | 8.00 | 6.00 | 1.00 |
| IGSF3 | 94.00 | 91.00 | 109.00 | 98.00 | 86.00 |
| IGSF5 | 8.00 | 3.00 | 7.00 | 6.00 | 3.00 |
| IGSF8 | 1392.00 | 1444.00 | 1265.00 | 1367.00 | 1188.00 |
| IK | 2230.00 | 2222.00 | 2409.00 | 2287.00 | 2567.00 |
| IKBIP | 1264.00 | 1234.00 | 1130.00 | 1209.33 | 1516.00 |
| IKBKAP | 1931.00 | 1759.00 | 2055.00 | 1915.00 | 1890.00 |
| IKBKB | 3242.00 | 3180.00 | 3283.00 | 3235.00 | 2921.00 |
| IKBKE | 199.00 | 206.00 | 236.00 | 213.67 | 170.00 |
| IKBKG | 346.00 | 308.00 | 337.00 | 330.33 | 362.00 |
| IKZF2 | 82.00 | 73.00 | 72.00 | 75.67 | 44.00 |

| | | | | | |
|---|---|---|---|---|---|
| IKZF4 | 277.00 | 289.00 | 307.00 | 291.00 | 248.00 |
| IKZF5 | 628.00 | 526.00 | 666.00 | 606.67 | 616.00 |
| IL10RB | 427.00 | 428.00 | 431.00 | 428.67 | 542.00 |
| IL11 | 678.00 | 625.00 | 659.00 | 654.00 | 98.00 |
| IL11RA | 102.00 | 102.00 | 127.00 | 110.33 | 77.00 |
| IL12A | 388.00 | 379.00 | 409.00 | 392.00 | 803.00 |
| IL12A-AS1 | 10.00 | 11.00 | 12.00 | 11.00 | 10.00 |
| IL12RB1 | 3.00 | 6.00 | 4.00 | 4.33 | 0.00 |
| IL12RB2 | 239.00 | 232.00 | 212.00 | 227.67 | 191.00 |
| IL13RA1 | 1605.00 | 1557.00 | 1619.00 | 1593.67 | 1537.00 |
| IL13RA2 | 176.00 | 203.00 | 159.00 | 179.33 | 148.00 |
| IL15 | 8.00 | 8.00 | 14.00 | 10.00 | 13.00 |
| IL15RA | 24.00 | 18.00 | 26.00 | 22.67 | 9.00 |
| IL16 | 14.00 | 13.00 | 21.00 | 16.00 | 19.00 |
| IL17D | 68.00 | 74.00 | 81.00 | 74.33 | 83.00 |
| IL17RA | 662.00 | 692.00 | 668.00 | 674.00 | 676.00 |
| IL17RC | 462.00 | 455.00 | 488.00 | 468.33 | 539.00 |
| IL17RD | 484.00 | 442.00 | 486.00 | 470.67 | 462.00 |
| IL17RE | 15.00 | 10.00 | 18.00 | 14.33 | 11.00 |
| IL18BP | 907.00 | 768.00 | 965.00 | 880.00 | 494.00 |
| IL18R1 | 35.00 | 35.00 | 49.00 | 39.67 | 42.00 |
| IL1A | 101.00 | 94.00 | 108.00 | 101.00 | 148.00 |
| IL1B | 6601.00 | 6221.00 | 6349.00 | 6390.33 | 12035.00 |
| IL1R1 | 42.00 | 52.00 | 48.00 | 47.33 | 48.00 |
| IL1RAP | 1050.00 | 1011.00 | 1159.00 | 1073.33 | 1013.00 |
| IL1RAPL1 | 114.00 | 101.00 | 123.00 | 112.67 | 77.00 |
| IL1RL1 | 36.00 | 36.00 | 47.00 | 39.67 | 26.00 |
| IL20RB | 104.00 | 136.00 | 113.00 | 117.67 | 103.00 |
| IL21R | 52.00 | 35.00 | 44.00 | 43.67 | 89.00 |
| IL23A | 9.00 | 13.00 | 7.00 | 9.67 | 15.00 |
| IL24 | 75.00 | 74.00 | 129.00 | 92.67 | 59.00 |
| IL27RA | 616.00 | 585.00 | 613.00 | 604.67 | 727.00 |
| IL2RB | 3.00 | 4.00 | 3.00 | 3.33 | 4.00 |
| IL2RG | 6.00 | 0.00 | 0.00 | 2.00 | 5.00 |
| IL31RA | 1167.00 | 952.00 | 1130.00 | 1083.00 | 754.00 |
| IL32 | 19.00 | 17.00 | 31.00 | 22.33 | 66.00 |
| IL37 | 17.00 | 11.00 | 14.00 | 14.00 | 17.00 |
| IL4I1 | 25.00 | 24.00 | 43.00 | 30.67 | 33.00 |
| IL4R | 985.00 | 960.00 | 905.00 | 950.00 | 852.00 |
| IL6 | 35.00 | 33.00 | 36.00 | 34.67 | 69.00 |
| IL6R | 33.00 | 32.00 | 33.00 | 32.67 | 9.00 |
| IL6ST | 5644.00 | 5052.00 | 5436.00 | 5377.33 | 5132.00 |
| IL7 | 28.00 | 38.00 | 44.00 | 36.67 | 52.00 |
| IL7R | 562.00 | 452.00 | 443.00 | 485.67 | 126.00 |
| ILF2 | 4002.00 | 3819.00 | 4026.00 | 3949.00 | 5112.00 |
| ILF3 | 11707.00 | 11906.00 | 12463.00 | 12025.33 | 12564.00 |
| ILF3-AS1 | 318.00 | 359.00 | 371.00 | 349.33 | 347.00 |
| ILK | 2796.00 | 2759.00 | 2958.00 | 2837.67 | 3730.00 |

| | | | | | |
|---|---|---|---|---|---|
| ILKAP | 677.00 | 735.00 | 772.00 | 728.00 | 847.00 |
| ILVBL | 1334.00 | 1317.00 | 1453.00 | 1368.00 | 1499.00 |
| IMMP1L | 64.00 | 57.00 | 76.00 | 65.67 | 88.00 |
| IMMP2L | 330.00 | 299.00 | 350.00 | 326.33 | 371.00 |
| IMMT | 2212.00 | 2122.00 | 2282.00 | 2205.33 | 2733.00 |
| IMP3 | 1121.00 | 1068.00 | 1141.00 | 1110.00 | 1447.00 |
| IMP4 | 1260.00 | 1254.00 | 1251.00 | 1255.00 | 1430.00 |
| IMPA1 | 750.00 | 695.00 | 821.00 | 755.33 | 885.00 |
| IMPA2 | 63.00 | 59.00 | 77.00 | 66.33 | 98.00 |
| IMPACT | 894.00 | 794.00 | 976.00 | 888.00 | 1165.00 |
| IMPAD1 | 3689.00 | 3249.00 | 3654.00 | 3530.67 | 3520.00 |
| IMPDH1 | 1830.00 | 1740.00 | 1886.00 | 1818.67 | 2177.00 |
| IMPDH2 | 8930.00 | 8677.00 | 9989.00 | 9198.67 | 9965.00 |
| IMPG2 | 50.00 | 32.00 | 42.00 | 41.33 | 28.00 |
| INA | 66.00 | 72.00 | 66.00 | 68.00 | 35.00 |
| INAFM1 | 66.00 | 66.00 | 72.00 | 68.00 | 69.00 |
| INAFM2 | 620.00 | 631.00 | 630.00 | 627.00 | 686.00 |
| INCA1 | 15.00 | 16.00 | 17.00 | 16.00 | 13.00 |
| INCENP | 935.00 | 839.00 | 918.00 | 897.33 | 980.00 |
| INE1 | 134.00 | 120.00 | 133.00 | 129.00 | 91.00 |
| INF2 | 2554.00 | 2660.00 | 2865.00 | 2693.00 | 2445.00 |
| ING1 | 207.00 | 207.00 | 217.00 | 210.33 | 245.00 |
| ING2 | 85.00 | 81.00 | 90.00 | 85.33 | 116.00 |
| ING3 | 316.00 | 312.00 | 340.00 | 322.67 | 424.00 |
| ING4 | 598.00 | 601.00 | 633.00 | 610.67 | 587.00 |
| ING5 | 985.00 | 1022.00 | 1156.00 | 1054.33 | 1135.00 |
| INHA | 30.00 | 27.00 | 36.00 | 31.00 | 30.00 |
| INHBA | 2539.00 | 2471.00 | 2762.00 | 2590.67 | 2529.00 |
| INHBA-AS1 | 13.00 | 7.00 | 14.00 | 11.33 | 11.00 |
| INHBC | 9.00 | 4.00 | 4.00 | 5.67 | 4.00 |
| INHBE | 491.00 | 461.00 | 481.00 | 477.67 | 264.00 |
| INIP | 178.00 | 127.00 | 209.00 | 171.33 | 160.00 |
| INO80 | 1486.00 | 1363.00 | 1559.00 | 1469.33 | 1374.00 |
| INO80B | 68.00 | 57.00 | 73.00 | 66.00 | 36.00 |
| INO80C | 130.00 | 117.00 | 135.00 | 127.33 | 211.00 |
| INO80D | 659.00 | 570.00 | 738.00 | 655.67 | 676.00 |
| INO80E | 1224.00 | 1218.00 | 1250.00 | 1230.67 | 1482.00 |
| INPP1 | 798.00 | 733.00 | 810.00 | 780.33 | 804.00 |
| INPP4A | 1109.00 | 1054.00 | 1186.00 | 1116.33 | 1134.00 |
| INPP4B | 457.00 | 372.00 | 444.00 | 424.33 | 226.00 |
| INPP5A | 478.00 | 448.00 | 491.00 | 472.33 | 506.00 |
| INPP5B | 721.00 | 649.00 | 668.00 | 679.33 | 751.00 |
| INPP5E | 1135.00 | 1167.00 | 1282.00 | 1194.67 | 1105.00 |
| INPP5F | 744.00 | 747.00 | 828.00 | 773.00 | 675.00 |
| INPP5J | 20.00 | 33.00 | 34.00 | 29.00 | 24.00 |
| INPP5K | 462.00 | 475.00 | 521.00 | 486.00 | 468.00 |
| INPPL1 | 5877.00 | 5778.00 | 6415.00 | 6023.33 | 5490.00 |
| INSC | 24.00 | 22.00 | 29.00 | 25.00 | 66.00 |

| | | | | | |
|---|---|---|---|---|---|
| INSIG1 | 879.00 | 836.00 | 894.00 | 869.67 | 1235.00 |
| INSIG2 | 432.00 | 407.00 | 392.00 | 410.33 | 362.00 |
| INSR | 3.00 | 2.00 | 6.00 | 3.67 | 8.00 |
| INTS1 | 4176.00 | 3917.00 | 4027.00 | 4040.00 | 3668.00 |
| INTS10 | 839.00 | 789.00 | 848.00 | 825.33 | 958.00 |
| INTS12 | 437.00 | 405.00 | 440.00 | 427.33 | 474.00 |
| INTS2 | 1145.00 | 1057.00 | 1250.00 | 1150.67 | 1156.00 |
| INTS3 | 7.00 | 6.00 | 7.00 | 6.67 | 13.00 |
| INTS4 | 495.00 | 434.00 | 531.00 | 486.67 | 479.00 |
| INTS4P2 | 9.00 | 9.00 | 13.00 | 10.33 | 7.00 |
| INTS5 | 646.00 | 607.00 | 600.00 | 617.67 | 556.00 |
| INTS6 | 1026.00 | 959.00 | 1097.00 | 1027.33 | 1144.00 |
| INTS6-AS1 | 41.00 | 59.00 | 41.00 | 47.00 | 52.00 |
| INTS7 | 821.00 | 699.00 | 764.00 | 761.33 | 890.00 |
| INTS8 | 1619.00 | 1603.00 | 1794.00 | 1672.00 | 1740.00 |
| INTS9 | 416.00 | 430.00 | 394.00 | 413.33 | 482.00 |
| INTU | 154.00 | 153.00 | 163.00 | 156.67 | 221.00 |
| INVS | 488.00 | 436.00 | 440.00 | 454.67 | 472.00 |
| IP6K1 | 974.00 | 890.00 | 1006.00 | 956.67 | 994.00 |
| IP6K2 | 1960.00 | 1938.00 | 1998.00 | 1965.33 | 2197.00 |
| IPCEF1 | 5.00 | 4.00 | 12.00 | 7.00 | 4.00 |
| IPMK | 295.00 | 244.00 | 295.00 | 278.00 | 248.00 |
| IPO11 | 268.00 | 236.00 | 269.00 | 257.67 | 252.00 |
| IPO13 | 1244.00 | 1292.00 | 1271.00 | 1269.00 | 1160.00 |
| IPO4 | 2165.00 | 2189.00 | 2055.00 | 2136.33 | 2458.00 |
| IPO5 | 5851.00 | 5233.00 | 6116.00 | 5733.33 | 6678.00 |
| IPO5P1 | 8.00 | 15.00 | 11.00 | 11.33 | 12.00 |
| IPO7 | 8688.00 | 7729.00 | 9405.00 | 8607.33 | 9079.00 |
| IPO8 | 1979.00 | 1772.00 | 1955.00 | 1902.00 | 1909.00 |
| IPO9 | 2656.00 | 2629.00 | 2785.00 | 2690.00 | 2363.00 |
| IPO9-AS1 | 6.00 | 5.00 | 0.00 | 3.67 | 9.00 |
| IPP | 316.00 | 277.00 | 282.00 | 291.67 | 310.00 |
| IPPK | 473.00 | 452.00 | 485.00 | 470.00 | 598.00 |
| IPW | 814.00 | 827.00 | 983.00 | 874.67 | 852.00 |
| IQCB1 | 1261.00 | 1216.00 | 1330.00 | 1269.00 | 1491.00 |
| IQCC | 142.00 | 155.00 | 168.00 | 155.00 | 203.00 |
| IQCD | 24.00 | 25.00 | 22.00 | 23.67 | 17.00 |
| IQCE | 382.00 | 402.00 | 494.00 | 426.00 | 388.00 |
| IQCG | 465.00 | 413.00 | 455.00 | 444.33 | 629.00 |
| IQCH | 114.00 | 128.00 | 121.00 | 121.00 | 118.00 |
| IQCH-AS1 | 245.00 | 253.00 | 254.00 | 250.67 | 191.00 |
| IQCJ | 16.00 | 22.00 | 18.00 | 18.67 | 24.00 |
| IQCK | 69.00 | 65.00 | 67.00 | 67.00 | 81.00 |
| IQGAP1 | 7749.00 | 7020.00 | 8309.00 | 7692.67 | 6382.00 |
| IQGAP2 | 17.00 | 18.00 | 13.00 | 16.00 | 6.00 |
| IQGAP3 | 3222.00 | 3057.00 | 3077.00 | 3118.67 | 2840.00 |
| IQSEC1 | 687.00 | 669.00 | 810.00 | 722.00 | 574.00 |
| IQSEC2 | 73.00 | 77.00 | 63.00 | 71.00 | 69.00 |

| | | | | | |
|---|---|---|---|---|---|
| IQUB | 9.00 | 8.00 | 10.00 | 9.00 | 12.00 |
| IRAIN | 18.00 | 9.00 | 10.00 | 12.33 | 17.00 |
| IRAK1 | 5417.00 | 5680.00 | 5783.00 | 5626.67 | 6788.00 |
| IRAK1BP1 | 174.00 | 159.00 | 198.00 | 177.00 | 181.00 |
| IRAK2 | 259.00 | 261.00 | 249.00 | 256.33 | 230.00 |
| IRAK3 | 12.00 | 9.00 | 8.00 | 9.67 | 8.00 |
| IRAK4 | 605.00 | 631.00 | 651.00 | 629.00 | 857.00 |
| IREB2 | 1257.00 | 1184.00 | 1427.00 | 1289.33 | 1010.00 |
| IRF1 | 270.00 | 259.00 | 320.00 | 283.00 | 294.00 |
| IRF2 | 309.00 | 282.00 | 345.00 | 312.00 | 348.00 |
| IRF2BP1 | 595.00 | 616.00 | 720.00 | 643.67 | 821.00 |
| IRF2BP2 | 2625.00 | 2486.00 | 2990.00 | 2700.33 | 2927.00 |
| IRF2BPL | 1253.00 | 1176.00 | 1476.00 | 1301.67 | 1051.00 |
| IRF3 | 1598.00 | 1603.00 | 1655.00 | 1618.67 | 1732.00 |
| IRF7 | 6.00 | 6.00 | 2.00 | 4.67 | 4.00 |
| IRF9 | 331.00 | 303.00 | 330.00 | 321.33 | 340.00 |
| IRG1 | 5.00 | 7.00 | 6.00 | 6.00 | 8.00 |
| IRGQ | 919.00 | 868.00 | 996.00 | 927.67 | 716.00 |
| IRS1 | 3445.00 | 3351.00 | 3869.00 | 3555.00 | 4152.00 |
| IRS2 | 572.00 | 523.00 | 649.00 | 581.33 | 672.00 |
| IRX3 | 148.00 | 128.00 | 175.00 | 150.33 | 129.00 |
| IRX5 | 34.00 | 36.00 | 44.00 | 38.00 | 32.00 |
| ISCA1 | 299.00 | 332.00 | 366.00 | 332.33 | 421.00 |
| ISCA2 | 208.00 | 167.00 | 170.00 | 181.67 | 207.00 |
| ISCU | 830.00 | 924.00 | 932.00 | 895.33 | 941.00 |
| ISG15 | 55.00 | 59.00 | 69.00 | 61.00 | 48.00 |
| ISG20 | 415.00 | 462.00 | 530.00 | 469.00 | 615.00 |
| ISG20L2 | 886.00 | 873.00 | 971.00 | 910.00 | 798.00 |
| ISL1 | 43.00 | 49.00 | 37.00 | 43.00 | 44.00 |
| ISL2 | 102.00 | 79.00 | 93.00 | 91.33 | 108.00 |
| ISM1 | 111.00 | 110.00 | 128.00 | 116.33 | 136.00 |
| ISOC1 | 375.00 | 325.00 | 373.00 | 357.67 | 542.00 |
| ISOC2 | 395.00 | 421.00 | 486.00 | 434.00 | 547.00 |
| ISPD | 49.00 | 50.00 | 61.00 | 53.33 | 57.00 |
| IST1 | 2287.00 | 2277.00 | 2425.00 | 2329.67 | 2465.00 |
| ISY1 | 346.00 | 334.00 | 397.00 | 359.00 | 406.00 |
| ISYNA1 | 212.00 | 186.00 | 210.00 | 202.67 | 175.00 |
| ITCH | 1799.00 | 1585.00 | 1886.00 | 1756.67 | 1904.00 |
| ITFG1 | 1649.00 | 1582.00 | 1668.00 | 1633.00 | 1682.00 |
| ITFG2 | 464.00 | 471.00 | 489.00 | 474.67 | 488.00 |
| ITFG3 | 2146.00 | 2153.00 | 1995.00 | 2098.00 | 2085.00 |
| ITGA1 | 517.00 | 447.00 | 497.00 | 487.00 | 272.00 |
| ITGA10 | 900.00 | 918.00 | 948.00 | 922.00 | 1574.00 |
| ITGA11 | 21.00 | 21.00 | 24.00 | 22.00 | 5.00 |
| ITGA2 | 4388.00 | 4102.00 | 4628.00 | 4372.67 | 2355.00 |
| ITGA2B | 3.00 | 6.00 | 4.00 | 4.33 | 2.00 |
| ITGA3 | 42824.00 | 42922.00 | 44349.00 | 43365.00 | 22400.00 |
| ITGA4 | 2256.00 | 2093.00 | 2323.00 | 2224.00 | 2284.00 |

| | | | | | |
|---|---|---|---|---|---|
| ITGA5 | 7517.00 | 7724.00 | 7633.00 | 7624.67 | 5894.00 |
| ITGA6 | 17752.00 | 16096.00 | 17542.00 | 17130.00 | 20076.00 |
| ITGA7 | 62.00 | 47.00 | 71.00 | 60.00 | 61.00 |
| ITGA9-AS1 | 7.00 | 4.00 | 6.00 | 5.67 | 13.00 |
| ITGAE | 389.00 | 351.00 | 391.00 | 377.00 | 456.00 |
| ITGAL | 3.00 | 1.00 | 0.00 | 1.33 | 1.00 |
| ITGAV | 2369.00 | 2164.00 | 2511.00 | 2348.00 | 1979.00 |
| ITGAX | 115.00 | 122.00 | 105.00 | 114.00 | 74.00 |
| ITGB1 | 34834.00 | 33200.00 | 34581.00 | 34205.00 | 32367.00 |
| ITGB1BP1 | 595.00 | 549.00 | 661.00 | 601.67 | 794.00 |
| ITGB1BP2 | 30.00 | 27.00 | 40.00 | 32.33 | 29.00 |
| ITGB2 | 3707.00 | 3743.00 | 4030.00 | 3826.67 | 3070.00 |
| ITGB2-AS1 | 686.00 | 635.00 | 731.00 | 684.00 | 727.00 |
| ITGB3 | 802.00 | 847.00 | 837.00 | 828.67 | 184.00 |
| ITGB3BP | 346.00 | 324.00 | 370.00 | 346.67 | 465.00 |
| ITGB4 | 15.00 | 15.00 | 16.00 | 15.33 | 7.00 |
| ITGB5 | 4942.00 | 5177.00 | 5090.00 | 5069.67 | 2762.00 |
| ITGB7 | 6.00 | 2.00 | 6.00 | 4.67 | 2.00 |
| ITGB8 | 155.00 | 150.00 | 200.00 | 168.33 | 369.00 |
| ITGBL1 | 5427.00 | 5054.00 | 5389.00 | 5290.00 | 4955.00 |
| ITIH4 | 27.00 | 30.00 | 31.00 | 29.33 | 22.00 |
| ITM2B | 2451.00 | 2461.00 | 2402.00 | 2438.00 | 2642.00 |
| ITM2C | 3773.00 | 3791.00 | 3651.00 | 3738.33 | 3444.00 |
| ITPA | 1282.00 | 1356.00 | 1431.00 | 1356.33 | 1862.00 |
| ITPKA | 168.00 | 163.00 | 193.00 | 174.67 | 177.00 |
| ITPKB | 334.00 | 349.00 | 368.00 | 350.33 | 448.00 |
| ITPKB-IT1 | 3.00 | 0.00 | 1.00 | 1.33 | 1.00 |
| ITPKC | 281.00 | 295.00 | 338.00 | 304.67 | 310.00 |
| ITPR1 | 410.00 | 389.00 | 477.00 | 425.33 | 465.00 |
| ITPR1-AS1 | 17.00 | 13.00 | 13.00 | 14.33 | 16.00 |
| ITPR2 | 1526.00 | 1374.00 | 1548.00 | 1482.67 | 1294.00 |
| ITPR3 | 3270.00 | 3157.00 | 3605.00 | 3344.00 | 2209.00 |
| ITPRIP | 5632.00 | 5648.00 | 5660.00 | 5646.67 | 4752.00 |
| ITPRIPL1 | 10.00 | 17.00 | 17.00 | 14.67 | 13.00 |
| ITPRIPL2 | 4132.00 | 3790.00 | 4101.00 | 4007.67 | 3343.00 |
| ITSN1 | 924.00 | 883.00 | 912.00 | 906.33 | 1160.00 |
| ITSN2 | 1132.00 | 1012.00 | 1204.00 | 1116.00 | 970.00 |
| IVD | 1800.00 | 1590.00 | 1782.00 | 1724.00 | 1728.00 |
| IVNS1ABP | 2589.00 | 2659.00 | 2977.00 | 2741.67 | 3001.00 |
| IWS1 | 1350.00 | 1254.00 | 1364.00 | 1322.67 | 1553.00 |
| IZUMO4 | 49.00 | 40.00 | 72.00 | 53.67 | 48.00 |
| JADE1 | 661.00 | 596.00 | 746.00 | 667.67 | 786.00 |
| JADE2 | 851.00 | 735.00 | 938.00 | 841.33 | 774.00 |
| JADE3 | 495.00 | 420.00 | 481.00 | 465.33 | 461.00 |
| JAG1 | 2544.00 | 2460.00 | 2744.00 | 2582.67 | 1059.00 |
| JAGN1 | 614.00 | 600.00 | 620.00 | 611.33 | 599.00 |
| JAK1 | 4067.00 | 3788.00 | 4181.00 | 4012.00 | 3860.00 |
| JAK2 | 540.00 | 470.00 | 542.00 | 517.33 | 445.00 |

| | | | | | |
|---|---|---|---|---|---|
| JAK3 | 8.00 | 4.00 | 5.00 | 5.67 | 3.00 |
| JAM2 | 185.00 | 191.00 | 200.00 | 192.00 | 190.00 |
| JAM3 | 1112.00 | 1115.00 | 1161.00 | 1129.33 | 1144.00 |
| JARID2 | 691.00 | 699.00 | 799.00 | 729.67 | 618.00 |
| JAZF1 | 166.00 | 150.00 | 179.00 | 165.00 | 177.00 |
| JAZF1-AS1 | 6.00 | 0.00 | 3.00 | 3.00 | 0.00 |
| JDP2 | 3.00 | 0.00 | 1.00 | 1.33 | 4.00 |
| JHDM1D-AS1 | 65.00 | 90.00 | 82.00 | 79.00 | 87.00 |
| JKAMP | 1358.00 | 1346.00 | 1344.00 | 1349.33 | 1529.00 |
| JMJD1C | 4074.00 | 3439.00 | 3921.00 | 3811.33 | 3663.00 |
| JMJD4 | 394.00 | 354.00 | 309.00 | 352.33 | 450.00 |
| JMJD6 | 678.00 | 652.00 | 683.00 | 671.00 | 705.00 |
| JMJD7 | 104.00 | 90.00 | 105.00 | 99.67 | 100.00 |
| JMJD8 | 822.00 | 872.00 | 856.00 | 850.00 | 1005.00 |
| JMY | 847.00 | 623.00 | 809.00 | 759.67 | 576.00 |
| JOSD1 | 1590.00 | 1568.00 | 1751.00 | 1636.33 | 1863.00 |
| JOSD2 | 291.00 | 290.00 | 312.00 | 297.67 | 336.00 |
| JPH1 | 219.00 | 194.00 | 243.00 | 218.67 | 270.00 |
| JPX | 165.00 | 138.00 | 174.00 | 159.00 | 217.00 |
| JRK | 2422.00 | 2285.00 | 2547.00 | 2418.00 | 2044.00 |
| JRKL | 875.00 | 885.00 | 957.00 | 905.67 | 965.00 |
| JTB | 1650.00 | 1617.00 | 1759.00 | 1675.33 | 2118.00 |
| JUN | 7846.00 | 7757.00 | 7953.00 | 7852.00 | 9217.00 |
| JUNB | 493.00 | 542.00 | 515.00 | 516.67 | 741.00 |
| JUND | 326.00 | 274.00 | 375.00 | 325.00 | 268.00 |
| KALRN | 17.00 | 17.00 | 17.00 | 17.00 | 7.00 |
| KANK1 | 1187.00 | 1163.00 | 1288.00 | 1212.67 | 909.00 |
| KANK2 | 3559.00 | 3603.00 | 3715.00 | 3625.67 | 1478.00 |
| KANSL1 | 289.00 | 312.00 | 309.00 | 303.33 | 298.00 |
| KANSL1L | 248.00 | 212.00 | 301.00 | 253.67 | 242.00 |
| KANSL2 | 543.00 | 522.00 | 600.00 | 555.00 | 641.00 |
| KANSL3 | 2122.00 | 2103.00 | 2416.00 | 2213.67 | 2337.00 |
| KANTR | 7.00 | 9.00 | 6.00 | 7.33 | 19.00 |
| KARS | 3131.00 | 2906.00 | 3271.00 | 3102.67 | 3708.00 |
| KAT2A | 4664.00 | 4928.00 | 4993.00 | 4861.67 | 4615.00 |
| KAT2B | 236.00 | 202.00 | 236.00 | 224.67 | 312.00 |
| KAT5 | 1314.00 | 1212.00 | 1461.00 | 1329.00 | 1421.00 |
| KAT6A | 1476.00 | 1195.00 | 1522.00 | 1397.67 | 833.00 |
| KAT6B | 1052.00 | 983.00 | 1182.00 | 1072.33 | 881.00 |
| KAT7 | 959.00 | 912.00 | 960.00 | 943.67 | 884.00 |
| KAT8 | 689.00 | 712.00 | 815.00 | 738.67 | 701.00 |
| KATNA1 | 570.00 | 496.00 | 495.00 | 520.33 | 685.00 |
| KATNAL1 | 237.00 | 207.00 | 258.00 | 234.00 | 219.00 |
| KATNAL2 | 111.00 | 137.00 | 145.00 | 131.00 | 130.00 |
| KATNB1 | 1056.00 | 1008.00 | 1063.00 | 1042.33 | 1195.00 |
| KATNBL1 | 521.00 | 492.00 | 543.00 | 518.67 | 631.00 |
| KATNBL1P6 | 17.00 | 24.00 | 43.00 | 28.00 | 27.00 |
| KAZALD1 | 103.00 | 92.00 | 75.00 | 90.00 | 99.00 |

| | | | | | |
|---|---|---|---|---|---|
| KBTBD2 | 1855.00 | 1931.00 | 2044.00 | 1943.33 | 2307.00 |
| KBTBD3 | 42.00 | 31.00 | 50.00 | 41.00 | 47.00 |
| KBTBD4 | 244.00 | 247.00 | 246.00 | 245.67 | 302.00 |
| KBTBD6 | 364.00 | 340.00 | 388.00 | 364.00 | 375.00 |
| KBTBD7 | 81.00 | 48.00 | 81.00 | 70.00 | 98.00 |
| KBTBD8 | 238.00 | 254.00 | 290.00 | 260.67 | 279.00 |
| KC6 | 287.00 | 322.00 | 366.00 | 325.00 | 269.00 |
| KCMF1 | 1106.00 | 1091.00 | 1284.00 | 1160.33 | 1540.00 |
| KCNA6 | 3.00 | 0.00 | 1.00 | 1.33 | 2.00 |
| KCNAB1 | 10.00 | 11.00 | 14.00 | 11.67 | 23.00 |
| KCNAB2 | 3508.00 | 3577.00 | 3708.00 | 3597.67 | 3118.00 |
| KCNAB3 | 163.00 | 149.00 | 154.00 | 155.33 | 136.00 |
| KCNC4 | 111.00 | 87.00 | 117.00 | 105.00 | 95.00 |
| KCND1 | 164.00 | 183.00 | 184.00 | 177.00 | 104.00 |
| KCNE2 | 4.00 | 3.00 | 2.00 | 3.00 | 1.00 |
| KCNE4 | 136.00 | 123.00 | 125.00 | 128.00 | 139.00 |
| KCNE5 | 37.00 | 32.00 | 30.00 | 33.00 | 35.00 |
| KCNG1 | 3443.00 | 3522.00 | 3639.00 | 3534.67 | 4022.00 |
| KCNG3 | 12.00 | 14.00 | 9.00 | 11.67 | 19.00 |
| KCNH1 | 157.00 | 158.00 | 201.00 | 172.00 | 79.00 |
| KCNH3 | 16.00 | 17.00 | 23.00 | 18.67 | 26.00 |
| KCNH5 | 3.00 | 1.00 | 6.00 | 3.33 | 2.00 |
| KCNH8 | 12.00 | 9.00 | 4.00 | 8.33 | 8.00 |
| KCNIP2 | 235.00 | 266.00 | 274.00 | 258.33 | 183.00 |
| KCNIP2-AS1 | 20.00 | 21.00 | 13.00 | 18.00 | 20.00 |
| KCNIP3 | 630.00 | 620.00 | 664.00 | 638.00 | 584.00 |
| KCNJ14 | 248.00 | 251.00 | 311.00 | 270.00 | 248.00 |
| KCNJ15 | 9.00 | 14.00 | 11.00 | 11.33 | 8.00 |
| KCNJ2 | 277.00 | 260.00 | 283.00 | 273.33 | 351.00 |
| KCNJ2-AS1 | 11.00 | 21.00 | 19.00 | 17.00 | 29.00 |
| KCNJ3 | 4.00 | 3.00 | 8.00 | 5.00 | 7.00 |
| KCNJ4 | 5.00 | 3.00 | 4.00 | 4.00 | 4.00 |
| KCNJ8 | 76.00 | 76.00 | 73.00 | 75.00 | 154.00 |
| KCNK1 | 114.00 | 109.00 | 131.00 | 118.00 | 121.00 |
| KCNK2 | 157.00 | 153.00 | 181.00 | 163.67 | 460.00 |
| KCNK4 | 6.00 | 4.00 | 12.00 | 7.33 | 5.00 |
| KCNK6 | 155.00 | 157.00 | 190.00 | 167.33 | 103.00 |
| KCNK7 | 4.00 | 2.00 | 3.00 | 3.00 | 0.00 |
| KCNMA1 | 4500.00 | 4121.00 | 4763.00 | 4461.33 | 4037.00 |
| KCNMB2-AS1 | 22.00 | 16.00 | 18.00 | 18.67 | 19.00 |
| KCNMB3 | 148.00 | 147.00 | 161.00 | 152.00 | 175.00 |
| KCNMB4 | 24.00 | 21.00 | 23.00 | 22.67 | 26.00 |
| KCNN2 | 83.00 | 78.00 | 94.00 | 85.00 | 85.00 |
| KCNN4 | 445.00 | 495.00 | 498.00 | 479.33 | 525.00 |
| KCNQ1OT1 | 306.00 | 305.00 | 313.00 | 308.00 | 193.00 |
| KCNQ3 | 11.00 | 15.00 | 11.00 | 12.33 | 7.00 |
| KCNQ5 | 1736.00 | 1497.00 | 1678.00 | 1637.00 | 1398.00 |
| KCNQ5-IT1 | 4.00 | 2.00 | 6.00 | 4.00 | 6.00 |

| | | | | | |
|---|---|---|---|---|---|
| KCNRG | 8.00 | 17.00 | 12.00 | 12.33 | 6.00 |
| KCNS3 | 59.00 | 60.00 | 62.00 | 60.33 | 48.00 |
| KCNT2 | 261.00 | 187.00 | 231.00 | 226.33 | 206.00 |
| KCNU1 | 133.00 | 123.00 | 144.00 | 133.33 | 111.00 |
| KCP | 22.00 | 17.00 | 22.00 | 20.33 | 9.00 |
| KCTD1 | 407.00 | 375.00 | 440.00 | 407.33 | 403.00 |
| KCTD10 | 1394.00 | 1379.00 | 1543.00 | 1438.67 | 1330.00 |
| KCTD11 | 461.00 | 442.00 | 489.00 | 464.00 | 454.00 |
| KCTD12 | 900.00 | 827.00 | 1010.00 | 912.33 | 1134.00 |
| KCTD13 | 577.00 | 569.00 | 605.00 | 583.67 | 657.00 |
| KCTD15 | 207.00 | 223.00 | 266.00 | 232.00 | 214.00 |
| KCTD16 | 19.00 | 13.00 | 11.00 | 14.33 | 13.00 |
| KCTD17 | 284.00 | 291.00 | 303.00 | 292.67 | 284.00 |
| KCTD18 | 659.00 | 675.00 | 751.00 | 695.00 | 755.00 |
| KCTD19 | 13.00 | 9.00 | 13.00 | 11.67 | 16.00 |
| KCTD2 | 582.00 | 568.00 | 622.00 | 590.67 | 649.00 |
| KCTD20 | 1250.00 | 1184.00 | 1394.00 | 1276.00 | 1355.00 |
| KCTD21 | 149.00 | 185.00 | 168.00 | 167.33 | 164.00 |
| KCTD21-AS1 | 13.00 | 6.00 | 14.00 | 11.00 | 9.00 |
| KCTD3 | 908.00 | 799.00 | 958.00 | 888.33 | 938.00 |
| KCTD4 | 218.00 | 253.00 | 251.00 | 240.67 | 227.00 |
| KCTD5 | 670.00 | 578.00 | 701.00 | 649.67 | 686.00 |
| KCTD6 | 205.00 | 194.00 | 213.00 | 204.00 | 250.00 |
| KCTD7 | 508.00 | 573.00 | 645.00 | 575.33 | 489.00 |
| KCTD8 | 3.00 | 9.00 | 7.00 | 6.33 | 9.00 |
| KCTD9 | 414.00 | 403.00 | 503.00 | 440.00 | 478.00 |
| KDELC1 | 329.00 | 328.00 | 323.00 | 326.67 | 426.00 |
| KDELC2 | 1513.00 | 1372.00 | 1489.00 | 1458.00 | 1588.00 |
| KDELR1 | 4050.00 | 4054.00 | 3777.00 | 3960.33 | 4071.00 |
| KDELR2 | 5614.00 | 5220.00 | 5248.00 | 5360.67 | 5626.00 |
| KDELR3 | 527.00 | 483.00 | 465.00 | 491.67 | 501.00 |
| KDM1A | 2275.00 | 2197.00 | 2326.00 | 2266.00 | 2642.00 |
| KDM1B | 749.00 | 612.00 | 764.00 | 708.33 | 951.00 |
| KDM2A | 2640.00 | 2484.00 | 2722.00 | 2615.33 | 2589.00 |
| KDM2B | 698.00 | 656.00 | 774.00 | 709.33 | 743.00 |
| KDM3A | 1421.00 | 1314.00 | 1513.00 | 1416.00 | 1466.00 |
| KDM3B | 2310.00 | 2139.00 | 2450.00 | 2299.67 | 2192.00 |
| KDM4A | 2267.00 | 2228.00 | 2295.00 | 2263.33 | 2549.00 |
| KDM4A-AS1 | 48.00 | 38.00 | 53.00 | 46.33 | 70.00 |
| KDM4B | 870.00 | 795.00 | 801.00 | 822.00 | 750.00 |
| KDM4C | 950.00 | 945.00 | 1034.00 | 976.33 | 993.00 |
| KDM4D | 48.00 | 50.00 | 64.00 | 54.00 | 62.00 |
| KDM5A | 1356.00 | 1331.00 | 1373.00 | 1353.33 | 1288.00 |
| KDM5B | 1887.00 | 1694.00 | 1921.00 | 1834.00 | 1863.00 |
| KDM5C | 6262.00 | 6344.00 | 6509.00 | 6371.67 | 5644.00 |
| KDM6A | 671.00 | 695.00 | 785.00 | 717.00 | 706.00 |
| KDM6B | 1381.00 | 1271.00 | 1612.00 | 1421.33 | 1058.00 |
| KDM7A | 1176.00 | 1038.00 | 1323.00 | 1179.00 | 1067.00 |

| | | | | | |
|---|---|---|---|---|---|
| KDM8 | 78.00 | 78.00 | 96.00 | 84.00 | 77.00 |
| KDR | 64.00 | 58.00 | 69.00 | 63.67 | 61.00 |
| KDSR | 2027.00 | 1924.00 | 2006.00 | 1985.67 | 2049.00 |
| KEAP1 | 809.00 | 862.00 | 872.00 | 847.67 | 976.00 |
| KGFLP1 | 6.00 | 6.00 | 2.00 | 4.67 | 5.00 |
| KGFLP2 | 8.00 | 14.00 | 12.00 | 11.33 | 22.00 |
| KHDC1 | 131.00 | 151.00 | 147.00 | 143.00 | 135.00 |
| KHDRBS1 | 3523.00 | 3396.00 | 3544.00 | 3487.67 | 4558.00 |
| KHDRBS3 | 125.00 | 123.00 | 125.00 | 124.33 | 170.00 |
| KHK | 14.00 | 21.00 | 22.00 | 19.00 | 30.00 |
| KHNYN | 2509.00 | 2562.00 | 2853.00 | 2641.33 | 2199.00 |
| KHSRP | 4495.00 | 4213.00 | 4459.00 | 4389.00 | 4644.00 |
| KIAA0020 | 2423.00 | 2245.00 | 2523.00 | 2397.00 | 2874.00 |
| KIAA0100 | 5845.00 | 5576.00 | 6109.00 | 5843.33 | 4642.00 |
| KIAA0101 | 243.00 | 204.00 | 265.00 | 237.33 | 429.00 |
| KIAA0141 | 2186.00 | 2131.00 | 2374.00 | 2230.33 | 2191.00 |
| KIAA0195 | 2386.00 | 2394.00 | 2451.00 | 2410.33 | 1961.00 |
| KIAA0196 | 1624.00 | 1576.00 | 1815.00 | 1671.67 | 1769.00 |
| KIAA0226 | 907.00 | 794.00 | 975.00 | 892.00 | 998.00 |
| KIAA0226L | 24.00 | 20.00 | 19.00 | 21.00 | 5.00 |
| KIAA0232 | 847.00 | 800.00 | 1014.00 | 887.00 | 834.00 |
| KIAA0319L | 2607.00 | 2406.00 | 2315.00 | 2442.67 | 2363.00 |
| KIAA0355 | 200.00 | 180.00 | 216.00 | 198.67 | 136.00 |
| KIAA0368 | 3967.00 | 3728.00 | 4171.00 | 3955.33 | 3742.00 |
| KIAA0391 | 659.00 | 567.00 | 635.00 | 620.33 | 699.00 |
| KIAA0513 | 232.00 | 237.00 | 247.00 | 238.67 | 248.00 |
| KIAA0556 | 969.00 | 848.00 | 1000.00 | 939.00 | 689.00 |
| KIAA0586 | 737.00 | 609.00 | 691.00 | 679.00 | 670.00 |
| KIAA0753 | 1246.00 | 1164.00 | 1194.00 | 1201.33 | 1165.00 |
| KIAA0754 | 2059.00 | 1511.00 | 1940.00 | 1836.67 | 886.00 |
| KIAA0825 | 20.00 | 24.00 | 16.00 | 20.00 | 18.00 |
| KIAA0895 | 168.00 | 183.00 | 207.00 | 186.00 | 202.00 |
| KIAA0895L | 741.00 | 788.00 | 742.00 | 757.00 | 717.00 |
| KIAA0907 | 4459.00 | 4606.00 | 5037.00 | 4700.67 | 4480.00 |
| KIAA0922 | 737.00 | 720.00 | 750.00 | 735.67 | 737.00 |
| KIAA0930 | 478.00 | 505.00 | 560.00 | 514.33 | 557.00 |
| KIAA1024 | 121.00 | 158.00 | 146.00 | 141.67 | 113.00 |
| KIAA1033 | 1746.00 | 1679.00 | 1859.00 | 1761.33 | 1567.00 |
| KIAA1107 | 274.00 | 242.00 | 288.00 | 268.00 | 254.00 |
| KIAA1109 | 2405.00 | 2179.00 | 2470.00 | 2351.33 | 1890.00 |
| KIAA1143 | 420.00 | 346.00 | 484.00 | 416.67 | 429.00 |
| KIAA1161 | 21.00 | 26.00 | 13.00 | 20.00 | 21.00 |
| KIAA1191 | 1701.00 | 1622.00 | 1828.00 | 1717.00 | 1945.00 |
| KIAA1211 | 242.00 | 242.00 | 235.00 | 239.67 | 196.00 |
| KIAA1217 | 471.00 | 473.00 | 522.00 | 488.67 | 402.00 |
| KIAA1279 | 867.00 | 865.00 | 962.00 | 898.00 | 1061.00 |
| KIAA1324 | 4.00 | 10.00 | 6.00 | 6.67 | 1.00 |
| KIAA1324L | 75.00 | 86.00 | 77.00 | 79.33 | 54.00 |

| | | | | | |
|---|---|---|---|---|---|
| KIAA1328 | 127.00 | 102.00 | 122.00 | 117.00 | 90.00 |
| KIAA1407 | 410.00 | 347.00 | 387.00 | 381.33 | 461.00 |
| KIAA1429 | 2663.00 | 2505.00 | 2871.00 | 2679.67 | 2597.00 |
| KIAA1462 | 1006.00 | 949.00 | 1167.00 | 1040.67 | 505.00 |
| KIAA1467 | 955.00 | 866.00 | 905.00 | 908.67 | 930.00 |
| KIAA1468 | 759.00 | 744.00 | 849.00 | 784.00 | 644.00 |
| KIAA1522 | 48.00 | 39.00 | 50.00 | 45.67 | 42.00 |
| KIAA1524 | 844.00 | 746.00 | 799.00 | 796.33 | 1051.00 |
| KIAA1549 | 1374.00 | 1329.00 | 1445.00 | 1382.67 | 1203.00 |
| KIAA1549L | 4757.00 | 4485.00 | 4776.00 | 4672.67 | 4113.00 |
| KIAA1551 | 853.00 | 825.00 | 948.00 | 875.33 | 855.00 |
| KIAA1586 | 216.00 | 190.00 | 216.00 | 207.33 | 248.00 |
| KIAA1656 | 144.00 | 171.00 | 145.00 | 153.33 | 264.00 |
| KIAA1671 | 1240.00 | 1101.00 | 1249.00 | 1196.67 | 972.00 |
| KIAA1683 | 5.00 | 4.00 | 2.00 | 3.67 | 0.00 |
| KIAA1715 | 919.00 | 838.00 | 963.00 | 906.67 | 976.00 |
| KIAA1755 | 128.00 | 107.00 | 155.00 | 130.00 | 166.00 |
| KIAA1804 | 407.00 | 407.00 | 455.00 | 423.00 | 469.00 |
| KIAA1841 | 138.00 | 124.00 | 137.00 | 133.00 | 158.00 |
| KIAA1875 | 89.00 | 100.00 | 110.00 | 99.67 | 71.00 |
| KIAA1919 | 206.00 | 198.00 | 185.00 | 196.33 | 126.00 |
| KIAA1958 | 155.00 | 175.00 | 210.00 | 180.00 | 146.00 |
| KIAA2013 | 1732.00 | 1694.00 | 1735.00 | 1720.33 | 1748.00 |
| KIAA2018 | 376.00 | 276.00 | 359.00 | 337.00 | 315.00 |
| KIAA2026 | 595.00 | 603.00 | 695.00 | 631.00 | 535.00 |
| KIDINS220 | 2451.00 | 2140.00 | 2409.00 | 2333.33 | 2034.00 |
| KIF11 | 1254.00 | 1040.00 | 1167.00 | 1153.67 | 1518.00 |
| KIF12 | 3.00 | 0.00 | 0.00 | 1.00 | 0.00 |
| KIF13A | 1223.00 | 1141.00 | 1365.00 | 1243.00 | 1216.00 |
| KIF13B | 447.00 | 416.00 | 487.00 | 450.00 | 378.00 |
| KIF14 | 1057.00 | 979.00 | 1072.00 | 1036.00 | 1219.00 |
| KIF15 | 453.00 | 399.00 | 399.00 | 417.00 | 596.00 |
| KIF16B | 515.00 | 501.00 | 535.00 | 517.00 | 450.00 |
| KIF17 | 46.00 | 43.00 | 36.00 | 41.67 | 28.00 |
| KIF18A | 855.00 | 873.00 | 859.00 | 862.33 | 1170.00 |
| KIF18B | 1009.00 | 985.00 | 949.00 | 981.00 | 1337.00 |
| KIF1B | 1438.00 | 1294.00 | 1596.00 | 1442.67 | 1207.00 |
| KIF1C | 4882.00 | 4955.00 | 4924.00 | 4920.33 | 5158.00 |
| KIF20A | 559.00 | 558.00 | 555.00 | 557.33 | 695.00 |
| KIF20B | 1447.00 | 1323.00 | 1462.00 | 1410.67 | 1403.00 |
| KIF21A | 1228.00 | 1175.00 | 1338.00 | 1247.00 | 1221.00 |
| KIF21B | 1084.00 | 905.00 | 1119.00 | 1036.00 | 612.00 |
| KIF22 | 1307.00 | 1273.00 | 1296.00 | 1292.00 | 1676.00 |
| KIF23 | 1292.00 | 1224.00 | 1233.00 | 1249.67 | 1508.00 |
| KIF24 | 286.00 | 241.00 | 273.00 | 266.67 | 344.00 |
| KIF27 | 196.00 | 173.00 | 143.00 | 170.67 | 145.00 |
| KIF2A | 912.00 | 874.00 | 928.00 | 904.67 | 1062.00 |
| KIF2C | 1365.00 | 1351.00 | 1407.00 | 1374.33 | 1791.00 |

| | | | | | |
|---|---|---|---|---|---|
| KIF3A | 716.00 | 691.00 | 675.00 | 694.00 | 635.00 |
| KIF3B | 1343.00 | 1142.00 | 1345.00 | 1276.67 | 1300.00 |
| KIF3C | 582.00 | 478.00 | 564.00 | 541.33 | 598.00 |
| KIF4A | 1228.00 | 1101.00 | 1121.00 | 1150.00 | 1425.00 |
| KIF4B | 17.00 | 10.00 | 18.00 | 15.00 | 18.00 |
| KIF5B | 4622.00 | 3961.00 | 4949.00 | 4510.67 | 4078.00 |
| KIF5C | 246.00 | 241.00 | 268.00 | 251.67 | 175.00 |
| KIF6 | 60.00 | 41.00 | 63.00 | 54.67 | 52.00 |
| KIF7 | 940.00 | 915.00 | 914.00 | 923.00 | 754.00 |
| KIF9 | 120.00 | 157.00 | 166.00 | 147.67 | 191.00 |
| KIF9-AS1 | 49.00 | 61.00 | 46.00 | 52.00 | 29.00 |
| KIFAP3 | 615.00 | 563.00 | 665.00 | 614.33 | 658.00 |
| KIFC1 | 8.00 | 5.00 | 9.00 | 7.33 | 4.00 |
| KIFC2 | 2638.00 | 2940.00 | 3087.00 | 2888.33 | 2685.00 |
| KIFC3 | 2000.00 | 2091.00 | 2101.00 | 2064.00 | 2223.00 |
| KIN | 598.00 | 608.00 | 557.00 | 587.67 | 639.00 |
| KIRREL | 5736.00 | 5305.00 | 5783.00 | 5608.00 | 4839.00 |
| KISS1 | 40.00 | 49.00 | 40.00 | 43.00 | 21.00 |
| KIT | 64.00 | 67.00 | 53.00 | 61.33 | 10.00 |
| KITLG | 207.00 | 187.00 | 190.00 | 194.67 | 192.00 |
| KIZ | 453.00 | 434.00 | 520.00 | 469.00 | 480.00 |
| KLC1 | 1841.00 | 1834.00 | 1896.00 | 1857.00 | 2067.00 |
| KLC2 | 2586.00 | 2561.00 | 2809.00 | 2652.00 | 2702.00 |
| KLC4 | 742.00 | 767.00 | 855.00 | 788.00 | 672.00 |
| KLF10 | 719.00 | 692.00 | 797.00 | 736.00 | 799.00 |
| KLF11 | 60.00 | 67.00 | 96.00 | 74.33 | 95.00 |
| KLF12 | 349.00 | 326.00 | 373.00 | 349.33 | 348.00 |
| KLF13 | 31.00 | 31.00 | 41.00 | 34.33 | 43.00 |
| KLF14 | 14.00 | 14.00 | 8.00 | 12.00 | 23.00 |
| KLF15 | 147.00 | 124.00 | 154.00 | 141.67 | 151.00 |
| KLF16 | 1138.00 | 1259.00 | 1268.00 | 1221.67 | 1344.00 |
| KLF2 | 345.00 | 328.00 | 298.00 | 323.67 | 625.00 |
| KLF3 | 1574.00 | 1365.00 | 1734.00 | 1557.67 | 1627.00 |
| KLF3-AS1 | 55.00 | 56.00 | 37.00 | 49.33 | 68.00 |
| KLF4 | 414.00 | 390.00 | 407.00 | 403.67 | 372.00 |
| KLF5 | 46.00 | 29.00 | 38.00 | 37.67 | 45.00 |
| KLF6 | 2295.00 | 2242.00 | 2511.00 | 2349.33 | 2404.00 |
| KLF7 | 1115.00 | 987.00 | 1243.00 | 1115.00 | 1055.00 |
| KLF9 | 230.00 | 227.00 | 276.00 | 244.33 | 269.00 |
| KLHDC1 | 38.00 | 36.00 | 43.00 | 39.00 | 41.00 |
| KLHDC10 | 1366.00 | 1245.00 | 1493.00 | 1368.00 | 1352.00 |
| KLHDC2 | 618.00 | 592.00 | 589.00 | 599.67 | 605.00 |
| KLHDC3 | 1091.00 | 1117.00 | 1181.00 | 1129.67 | 1404.00 |
| KLHDC4 | 874.00 | 917.00 | 903.00 | 898.00 | 891.00 |
| KLHDC7B | 28.00 | 15.00 | 33.00 | 25.33 | 10.00 |
| KLHDC8B | 92.00 | 135.00 | 98.00 | 108.33 | 142.00 |
| KLHL11 | 292.00 | 286.00 | 373.00 | 317.00 | 346.00 |
| KLHL12 | 1385.00 | 1326.00 | 1463.00 | 1391.33 | 1620.00 |

| | | | | | |
|---|---|---|---|---|---|
| KLHL13 | 17.00 | 24.00 | 18.00 | 19.67 | 22.00 |
| KLHL14 | 15.00 | 16.00 | 10.00 | 13.67 | 34.00 |
| KLHL15 | 209.00 | 181.00 | 261.00 | 217.00 | 247.00 |
| KLHL17 | 1090.00 | 1212.00 | 1198.00 | 1166.67 | 1172.00 |
| KLHL18 | 703.00 | 697.00 | 709.00 | 703.00 | 794.00 |
| KLHL2 | 328.00 | 327.00 | 374.00 | 343.00 | 340.00 |
| KLHL20 | 635.00 | 541.00 | 591.00 | 589.00 | 589.00 |
| KLHL21 | 2070.00 | 1993.00 | 2127.00 | 2063.33 | 1751.00 |
| KLHL22 | 442.00 | 441.00 | 513.00 | 465.33 | 502.00 |
| KLHL23 | 16.00 | 9.00 | 10.00 | 11.67 | 18.00 |
| KLHL24 | 477.00 | 412.00 | 501.00 | 463.33 | 365.00 |
| KLHL25 | 142.00 | 128.00 | 131.00 | 133.67 | 144.00 |
| KLHL26 | 169.00 | 143.00 | 197.00 | 169.67 | 189.00 |
| KLHL28 | 205.00 | 171.00 | 215.00 | 197.00 | 180.00 |
| KLHL29 | 1316.00 | 1281.00 | 1369.00 | 1322.00 | 1067.00 |
| KLHL3 | 868.00 | 796.00 | 919.00 | 861.00 | 754.00 |
| KLHL31 | 61.00 | 59.00 | 61.00 | 60.33 | 65.00 |
| KLHL35 | 49.00 | 28.00 | 45.00 | 40.67 | 50.00 |
| KLHL36 | 1721.00 | 1694.00 | 1930.00 | 1781.67 | 1542.00 |
| KLHL4 | 320.00 | 315.00 | 424.00 | 353.00 | 304.00 |
| KLHL41 | 22.00 | 14.00 | 12.00 | 16.00 | 17.00 |
| KLHL42 | 2396.00 | 2331.00 | 2699.00 | 2475.33 | 2519.00 |
| KLHL5 | 2935.00 | 2613.00 | 3184.00 | 2910.67 | 3013.00 |
| KLHL7 | 816.00 | 749.00 | 774.00 | 779.67 | 847.00 |
| KLHL7-AS1 | 37.00 | 46.00 | 48.00 | 43.67 | 51.00 |
| KLHL8 | 372.00 | 314.00 | 401.00 | 362.33 | 417.00 |
| KLHL9 | 2182.00 | 1968.00 | 2370.00 | 2173.33 | 2399.00 |
| KLK14 | 52.00 | 41.00 | 65.00 | 52.67 | 37.00 |
| KLKB1 | 7.00 | 12.00 | 10.00 | 9.67 | 15.00 |
| KLLN | 4.00 | 10.00 | 8.00 | 7.33 | 12.00 |
| KLRAP1 | 71.00 | 71.00 | 89.00 | 77.00 | 91.00 |
| KLRC2 | 42.00 | 46.00 | 45.00 | 44.33 | 38.00 |
| KLRC3 | 10.00 | 8.00 | 22.00 | 13.33 | 4.00 |
| KLRC4 | 4.00 | 11.00 | 5.00 | 6.67 | 1.00 |
| KLRG1 | 43.00 | 45.00 | 48.00 | 45.33 | 83.00 |
| KMT2A | 3497.00 | 3251.00 | 3795.00 | 3514.33 | 2651.00 |
| KMT2B | 2202.00 | 2215.00 | 2407.00 | 2274.67 | 1858.00 |
| KMT2C | 1477.00 | 1282.00 | 1626.00 | 1461.67 | 1307.00 |
| KMT2D | 5618.00 | 4994.00 | 6102.00 | 5571.33 | 3455.00 |
| KMT2E | 1319.00 | 1176.00 | 1401.00 | 1298.67 | 968.00 |
| KMT2E-AS1 | 90.00 | 84.00 | 84.00 | 86.00 | 90.00 |
| KNOP1 | 373.00 | 365.00 | 370.00 | 369.33 | 450.00 |
| KNSTRN | 605.00 | 535.00 | 590.00 | 576.67 | 810.00 |
| KNTC1 | 1318.00 | 1282.00 | 1274.00 | 1291.33 | 1493.00 |
| KPNA1 | 2288.00 | 2168.00 | 2507.00 | 2321.00 | 2495.00 |
| KPNA2 | 3254.00 | 2992.00 | 3234.00 | 3160.00 | 4373.00 |
| KPNA3 | 1113.00 | 1001.00 | 1119.00 | 1077.67 | 1406.00 |
| KPNA4 | 3031.00 | 2695.00 | 3228.00 | 2984.67 | 3721.00 |

| | | | | | |
|---|---|---|---|---|---|
| KPNA5 | 160.00 | 140.00 | 178.00 | 159.33 | 141.00 |
| KPNA6 | 2303.00 | 2138.00 | 2430.00 | 2290.33 | 2196.00 |
| KPNB1 | 9061.00 | 8356.00 | 9279.00 | 8898.67 | 9731.00 |
| KPRP | 4.00 | 4.00 | 6.00 | 4.67 | 6.00 |
| KPTN | 275.00 | 270.00 | 294.00 | 279.67 | 288.00 |
| KRAS | 918.00 | 779.00 | 998.00 | 898.33 | 913.00 |
| KRBA1 | 335.00 | 343.00 | 369.00 | 349.00 | 348.00 |
| KRBA2 | 81.00 | 64.00 | 78.00 | 74.33 | 42.00 |
| KRBOX4 | 696.00 | 739.00 | 729.00 | 721.33 | 890.00 |
| KRCC1 | 634.00 | 638.00 | 715.00 | 662.33 | 832.00 |
| KREMEN1 | 342.00 | 333.00 | 337.00 | 337.33 | 253.00 |
| KREMEN2 | 6.00 | 5.00 | 3.00 | 4.67 | 3.00 |
| KRI1 | 1955.00 | 1996.00 | 1936.00 | 1962.33 | 2257.00 |
| KRIT1 | 1347.00 | 1307.00 | 1533.00 | 1395.67 | 1328.00 |
| KRR1 | 1019.00 | 869.00 | 1053.00 | 980.33 | 1101.00 |
| KRT10 | 452.00 | 424.00 | 448.00 | 441.33 | 511.00 |
| KRT15 | 117.00 | 115.00 | 104.00 | 112.00 | 294.00 |
| KRT18 | 3996.00 | 3906.00 | 4071.00 | 3991.00 | 3701.00 |
| KRT7 | 5781.00 | 5804.00 | 6249.00 | 5944.67 | 3828.00 |
| KRT79 | 331.00 | 321.00 | 372.00 | 341.33 | 298.00 |
| KRT8 | 1022.00 | 953.00 | 1120.00 | 1031.67 | 971.00 |
| KRT80 | 739.00 | 747.00 | 810.00 | 765.33 | 648.00 |
| KRT81 | 17011.00 | 16825.00 | 17276.00 | 17037.33 | 10223.00 |
| KRT83 | 20.00 | 32.00 | 21.00 | 24.33 | 15.00 |
| KRT85 | 7.00 | 5.00 | 6.00 | 6.00 | 0.00 |
| KRT86 | 173.00 | 201.00 | 164.00 | 179.33 | 108.00 |
| KRTAP2-3 | 28.00 | 41.00 | 21.00 | 30.00 | 312.00 |
| KRTAP2-4 | 38.00 | 58.00 | 25.00 | 40.33 | 98.00 |
| KRTAP4-12 | 11.00 | 21.00 | 11.00 | 14.33 | 14.00 |
| KRTAP4-7 | 34.00 | 34.00 | 29.00 | 32.33 | 106.00 |
| KRTAP4-9 | 5.00 | 4.00 | 14.00 | 7.67 | 22.00 |
| KRTAP5-11 | 7.00 | 5.00 | 6.00 | 6.00 | 2.00 |
| KRTAP5-9 | 10.00 | 8.00 | 4.00 | 7.33 | 0.00 |
| KRTCAP2 | 1305.00 | 1311.00 | 1324.00 | 1313.33 | 1706.00 |
| KRTCAP3 | 38.00 | 33.00 | 47.00 | 39.33 | 47.00 |
| KSR1 | 400.00 | 361.00 | 441.00 | 400.67 | 405.00 |
| KTN1 | 5729.00 | 5125.00 | 5747.00 | 5533.67 | 5628.00 |
| KTN1-AS1 | 59.00 | 67.00 | 54.00 | 60.00 | 64.00 |
| KXD1 | 1674.00 | 1583.00 | 1805.00 | 1687.33 | 1990.00 |
| KYNU | 4.00 | 2.00 | 5.00 | 3.67 | 7.00 |
| L1CAM | 5.00 | 3.00 | 1.00 | 3.00 | 1.00 |
| L2HGDH | 271.00 | 252.00 | 285.00 | 269.33 | 290.00 |
| L3HYPDH | 870.00 | 862.00 | 952.00 | 894.67 | 741.00 |
| L3MBTL1 | 422.00 | 389.00 | 382.00 | 397.67 | 256.00 |
| L3MBTL2 | 619.00 | 602.00 | 608.00 | 609.67 | 721.00 |
| L3MBTL3 | 684.00 | 699.00 | 734.00 | 705.67 | 802.00 |
| LACC1 | 413.00 | 414.00 | 463.00 | 430.00 | 438.00 |
| LACE1 | 113.00 | 80.00 | 93.00 | 95.33 | 103.00 |

| | | | | | |
|---|---|---|---|---|---|
| LACTB | 1018.00 | 960.00 | 1052.00 | 1010.00 | 1032.00 |
| LACTB2 | 501.00 | 449.00 | 537.00 | 495.67 | 636.00 |
| LAG3 | 7.00 | 7.00 | 8.00 | 7.33 | 3.00 |
| LAGE3 | 166.00 | 186.00 | 211.00 | 187.67 | 287.00 |
| LAMA1 | 136.00 | 130.00 | 158.00 | 141.33 | 147.00 |
| LAMA2 | 15.00 | 16.00 | 14.00 | 15.00 | 36.00 |
| LAMA3 | 887.00 | 823.00 | 985.00 | 898.33 | 385.00 |
| LAMA4 | 5358.00 | 5011.00 | 5671.00 | 5346.67 | 5162.00 |
| LAMA5 | 3667.00 | 3525.00 | 3760.00 | 3650.67 | 2757.00 |
| LAMB1 | 27462.00 | 26647.00 | 29298.00 | 27802.33 | 20644.00 |
| LAMB2 | 6895.00 | 6936.00 | 7434.00 | 7088.33 | 5391.00 |
| LAMB2P1 | 5.00 | 3.00 | 5.00 | 4.33 | 2.00 |
| LAMB3 | 28972.00 | 29501.00 | 28361.00 | 28944.67 | 27079.00 |
| LAMB4 | 7.00 | 3.00 | 4.00 | 4.67 | 7.00 |
| LAMC1 | 21091.00 | 19798.00 | 21223.00 | 20704.00 | 20158.00 |
| LAMC2 | 1058.00 | 1115.00 | 1116.00 | 1096.33 | 1303.00 |
| LAMP1 | 8565.00 | 8691.00 | 8527.00 | 8594.33 | 5928.00 |
| LAMP2 | 3408.00 | 3120.00 | 3403.00 | 3310.33 | 2904.00 |
| LAMTOR1 | 1714.00 | 1727.00 | 1703.00 | 1714.67 | 2039.00 |
| LAMTOR2 | 507.00 | 484.00 | 522.00 | 504.33 | 653.00 |
| LAMTOR3 | 318.00 | 377.00 | 382.00 | 359.00 | 490.00 |
| LAMTOR4 | 669.00 | 754.00 | 781.00 | 734.67 | 901.00 |
| LAMTOR5 | 930.00 | 1022.00 | 1093.00 | 1015.00 | 1251.00 |
| LAMTOR5-AS1 | 4.00 | 11.00 | 4.00 | 6.33 | 12.00 |
| LANCL1 | 1617.00 | 1459.00 | 1683.00 | 1586.33 | 1714.00 |
| LANCL2 | 404.00 | 380.00 | 438.00 | 407.33 | 495.00 |
| LAP3 | 696.00 | 692.00 | 759.00 | 715.67 | 856.00 |
| LAPTM4A | 3381.00 | 3296.00 | 3329.00 | 3335.33 | 3346.00 |
| LAPTM4B | 5424.00 | 5297.00 | 5084.00 | 5268.33 | 6173.00 |
| LARP1 | 10728.00 | 10401.00 | 11104.00 | 10744.33 | 10981.00 |
| LARP1B | 351.00 | 286.00 | 357.00 | 331.33 | 327.00 |
| LARP4 | 1938.00 | 1759.00 | 2180.00 | 1959.00 | 2087.00 |
| LARP4B | 2543.00 | 2376.00 | 2540.00 | 2486.33 | 2831.00 |
| LARP6 | 1542.00 | 1587.00 | 1729.00 | 1619.33 | 1700.00 |
| LARP7 | 672.00 | 709.00 | 786.00 | 722.33 | 879.00 |
| LARS | 10977.00 | 10308.00 | 11909.00 | 11064.67 | 11099.00 |
| LARS2 | 1436.00 | 1383.00 | 1402.00 | 1407.00 | 1425.00 |
| LAS1L | 2776.00 | 2728.00 | 2972.00 | 2825.33 | 3031.00 |
| LASP1 | 4497.00 | 4092.00 | 4217.00 | 4268.67 | 3541.00 |
| LAT | 626.00 | 616.00 | 747.00 | 663.00 | 455.00 |
| LAT2 | 30.00 | 37.00 | 45.00 | 37.33 | 211.00 |
| LATS1 | 791.00 | 690.00 | 882.00 | 787.67 | 770.00 |
| LATS2 | 511.00 | 452.00 | 484.00 | 482.33 | 521.00 |
| LAYN | 628.00 | 589.00 | 667.00 | 628.00 | 842.00 |
| LBHD1 | 567.00 | 553.00 | 599.00 | 573.00 | 809.00 |
| LBR | 2435.00 | 2095.00 | 2221.00 | 2250.33 | 2311.00 |
| LBX2 | 78.00 | 74.00 | 87.00 | 79.67 | 87.00 |
| LBX2-AS1 | 153.00 | 139.00 | 135.00 | 142.33 | 143.00 |

| | | | | | |
|---|---|---|---|---|---|
| LCA5 | 142.00 | 121.00 | 148.00 | 137.00 | 146.00 |
| LCA5L | 30.00 | 33.00 | 32.00 | 31.67 | 31.00 |
| LCAT | 204.00 | 163.00 | 181.00 | 182.67 | 204.00 |
| LCLAT1 | 585.00 | 503.00 | 615.00 | 567.67 | 695.00 |
| LCMT1 | 362.00 | 358.00 | 358.00 | 359.33 | 424.00 |
| LCMT1-AS1 | 5.00 | 11.00 | 11.00 | 9.00 | 9.00 |
| LCMT2 | 264.00 | 279.00 | 340.00 | 294.33 | 308.00 |
| LCOR | 537.00 | 521.00 | 650.00 | 569.33 | 505.00 |
| LCORL | 144.00 | 123.00 | 139.00 | 135.33 | 125.00 |
| LCP1 | 65.00 | 78.00 | 80.00 | 74.33 | 43.00 |
| LCT | 3.00 | 1.00 | 1.00 | 1.67 | 0.00 |
| LCTL | 69.00 | 65.00 | 60.00 | 64.67 | 107.00 |
| LDAH | 581.00 | 566.00 | 626.00 | 591.00 | 642.00 |
| LDB1 | 1794.00 | 1702.00 | 1996.00 | 1830.67 | 1573.00 |
| LDHA | 8179.00 | 7646.00 | 8420.00 | 8081.67 | 13030.00 |
| LDHB | 16543.00 | 15348.00 | 17751.00 | 16547.33 | 23576.00 |
| LDHC | 72.00 | 59.00 | 73.00 | 68.00 | 113.00 |
| LDHD | 15.00 | 11.00 | 13.00 | 13.00 | 20.00 |
| LDLR | 2008.00 | 1988.00 | 1991.00 | 1995.67 | 1257.00 |
| LDLRAD3 | 1079.00 | 940.00 | 981.00 | 1000.00 | 1273.00 |
| LDLRAP1 | 277.00 | 291.00 | 308.00 | 292.00 | 335.00 |
| LDOC1L | 647.00 | 607.00 | 687.00 | 647.00 | 496.00 |
| LEAP2 | 56.00 | 60.00 | 47.00 | 54.33 | 66.00 |
| LEF1 | 238.00 | 214.00 | 274.00 | 242.00 | 352.00 |
| LEF1-AS1 | 13.00 | 4.00 | 10.00 | 9.00 | 12.00 |
| LEMD2 | 2337.00 | 2283.00 | 2353.00 | 2324.33 | 2259.00 |
| LEMD3 | 856.00 | 776.00 | 890.00 | 840.67 | 723.00 |
| LENG8 | 187.00 | 176.00 | 121.00 | 161.33 | 80.00 |
| LENG8-AS1 | 6.00 | 9.00 | 11.00 | 8.67 | 13.00 |
| LEO1 | 671.00 | 634.00 | 645.00 | 650.00 | 795.00 |
| LEPR | 362.00 | 323.00 | 384.00 | 356.33 | 397.00 |
| LEPROT | 2236.00 | 2137.00 | 2189.00 | 2187.33 | 2374.00 |
| LEPROTL1 | 809.00 | 721.00 | 738.00 | 756.00 | 893.00 |
| LETM1 | 2751.00 | 2757.00 | 2955.00 | 2821.00 | 2996.00 |
| LETM2 | 565.00 | 512.00 | 575.00 | 550.67 | 692.00 |
| LETMD1 | 2593.00 | 2404.00 | 2652.00 | 2549.67 | 2592.00 |
| LFNG | 42.00 | 41.00 | 39.00 | 40.67 | 35.00 |
| LGALS1 | 6169.00 | 6047.00 | 6559.00 | 6258.33 | 7989.00 |
| LGALS3 | 835.00 | 851.00 | 941.00 | 875.67 | 953.00 |
| LGALS3BP | 17787.00 | 18539.00 | 18567.00 | 18297.67 | 17045.00 |
| LGALS8 | 1148.00 | 1117.00 | 1186.00 | 1150.33 | 1052.00 |
| LGALS9 | 5.00 | 3.00 | 0.00 | 2.67 | 2.00 |
| LGALSL | 68.00 | 83.00 | 103.00 | 84.67 | 121.00 |
| LGI2 | 3.00 | 5.00 | 0.00 | 2.67 | 1.00 |
| LGMN | 1381.00 | 1422.00 | 1298.00 | 1367.00 | 1581.00 |
| LGR4 | 152.00 | 158.00 | 185.00 | 165.00 | 218.00 |
| LGR5 | 702.00 | 657.00 | 741.00 | 700.00 | 458.00 |
| LGR6 | 4.00 | 1.00 | 2.00 | 2.33 | 0.00 |

| | | | | | |
|---|---|---|---|---|---|
| LHB | 26.00 | 23.00 | 16.00 | 21.67 | 24.00 |
| LHFP | 495.00 | 472.00 | 460.00 | 475.67 | 404.00 |
| LHFPL2 | 1374.00 | 1278.00 | 1492.00 | 1381.33 | 1433.00 |
| LHPP | 114.00 | 103.00 | 118.00 | 111.67 | 128.00 |
| LHX4 | 3.00 | 1.00 | 4.00 | 2.67 | 1.00 |
| LHX4-AS1 | 736.00 | 791.00 | 832.00 | 786.33 | 799.00 |
| LHX9 | 4.00 | 0.00 | 7.00 | 3.67 | 2.00 |
| LIAS | 953.00 | 1014.00 | 1108.00 | 1025.00 | 1071.00 |
| LIF | 8994.00 | 9013.00 | 9582.00 | 9196.33 | 8807.00 |
| LIFR | 836.00 | 711.00 | 937.00 | 828.00 | 570.00 |
| LIFR-AS1 | 17.00 | 16.00 | 22.00 | 18.33 | 24.00 |
| LIG1 | 685.00 | 822.00 | 788.00 | 765.00 | 929.00 |
| LIG3 | 868.00 | 826.00 | 823.00 | 839.00 | 768.00 |
| LIG4 | 584.00 | 504.00 | 596.00 | 561.33 | 545.00 |
| LIMA1 | 1228.00 | 1183.00 | 1359.00 | 1256.67 | 1211.00 |
| LIMCH1 | 4.00 | 1.00 | 1.00 | 2.00 | 1.00 |
| LIMD1 | 2068.00 | 1999.00 | 2228.00 | 2098.33 | 2077.00 |
| LIMD1-AS1 | 3.00 | 7.00 | 7.00 | 5.67 | 6.00 |
| LIMD2 | 658.00 | 698.00 | 657.00 | 671.00 | 716.00 |
| LIME1 | 994.00 | 1127.00 | 1191.00 | 1104.00 | 1148.00 |
| LIMK1 | 1923.00 | 1961.00 | 1988.00 | 1957.33 | 1513.00 |
| LIMK2 | 332.00 | 276.00 | 303.00 | 303.67 | 352.00 |
| LIMS1 | 832.00 | 755.00 | 904.00 | 830.33 | 1050.00 |
| LIN28B | 74.00 | 57.00 | 65.00 | 65.33 | 71.00 |
| LIN37 | 211.00 | 187.00 | 197.00 | 198.33 | 266.00 |
| LIN52 | 197.00 | 179.00 | 193.00 | 189.67 | 210.00 |
| LIN54 | 235.00 | 197.00 | 248.00 | 226.67 | 266.00 |
| LIN7B | 63.00 | 74.00 | 83.00 | 73.33 | 65.00 |
| LIN7C | 356.00 | 365.00 | 430.00 | 383.67 | 534.00 |
| LIN9 | 233.00 | 219.00 | 233.00 | 228.33 | 298.00 |
| LINC00094 | 661.00 | 639.00 | 696.00 | 665.33 | 713.00 |
| LINC00115 | 24.00 | 28.00 | 36.00 | 29.33 | 20.00 |
| LINC00116 | 223.00 | 241.00 | 220.00 | 228.00 | 332.00 |
| LINC00152 | 298.00 | 275.00 | 306.00 | 293.00 | 303.00 |
| LINC00173 | 41.00 | 50.00 | 39.00 | 43.33 | 67.00 |
| LINC00174 | 789.00 | 755.00 | 861.00 | 801.67 | 762.00 |
| LINC00176 | 99.00 | 96.00 | 96.00 | 97.00 | 80.00 |
| LINC00184 | 11.00 | 7.00 | 11.00 | 9.67 | 15.00 |
| LINC00189 | 40.00 | 45.00 | 71.00 | 52.00 | 56.00 |
| LINC00202-1 | 3.00 | 23.00 | 22.00 | 16.00 | 14.00 |
| LINC00211 | 20.00 | 26.00 | 16.00 | 20.67 | 30.00 |
| LINC00240 | 9.00 | 13.00 | 14.00 | 12.00 | 20.00 |
| LINC00260 | 6.00 | 4.00 | 5.00 | 5.00 | 4.00 |
| LINC00261 | 561.00 | 567.00 | 567.00 | 565.00 | 370.00 |
| LINC00265 | 239.00 | 218.00 | 235.00 | 230.67 | 221.00 |
| LINC00271 | 3.00 | 2.00 | 4.00 | 3.00 | 1.00 |
| LINC00282 | 8.00 | 5.00 | 2.00 | 5.00 | 4.00 |
| LINC00294 | 375.00 | 369.00 | 423.00 | 389.00 | 418.00 |

| | | | | | |
|---|---|---|---|---|---|
| LINC00304 | 3.00 | 2.00 | 2.00 | 2.33 | 2.00 |
| LINC00310 | 11.00 | 8.00 | 14.00 | 11.00 | 28.00 |
| LINC00311 | 5.00 | 5.00 | 2.00 | 4.00 | 5.00 |
| LINC00312 | 3.00 | 2.00 | 4.00 | 3.00 | 4.00 |
| LINC00317 | 3.00 | 3.00 | 7.00 | 4.33 | 6.00 |
| LINC00324 | 10.00 | 10.00 | 9.00 | 9.67 | 6.00 |
| LINC00326 | 33.00 | 33.00 | 38.00 | 34.67 | 64.00 |
| LINC00327 | 9.00 | 11.00 | 7.00 | 9.00 | 8.00 |
| LINC00339 | 287.00 | 324.00 | 312.00 | 307.67 | 402.00 |
| LINC00341 | 171.00 | 170.00 | 209.00 | 183.33 | 185.00 |
| LINC00342 | 707.00 | 697.00 | 710.00 | 704.67 | 401.00 |
| LINC00346 | 290.00 | 263.00 | 265.00 | 272.67 | 221.00 |
| LINC00412 | 17.00 | 8.00 | 6.00 | 10.33 | 17.00 |
| LINC00431 | 6.00 | 7.00 | 10.00 | 7.67 | 6.00 |
| LINC00441 | 4.00 | 3.00 | 0.00 | 2.33 | 2.00 |
| LINC00452 | 6.00 | 4.00 | 3.00 | 4.33 | 4.00 |
| LINC00460 | 3.00 | 3.00 | 1.00 | 2.33 | 1.00 |
| LINC00461 | 29.00 | 27.00 | 26.00 | 27.33 | 27.00 |
| LINC00467 | 157.00 | 138.00 | 151.00 | 148.67 | 212.00 |
| LINC00470 | 127.00 | 134.00 | 130.00 | 130.33 | 132.00 |
| LINC00471 | 45.00 | 58.00 | 55.00 | 52.67 | 51.00 |
| LINC00472 | 447.00 | 442.00 | 513.00 | 467.33 | 512.00 |
| LINC00475 | 3.00 | 1.00 | 0.00 | 1.33 | 1.00 |
| LINC00476 | 96.00 | 78.00 | 94.00 | 89.33 | 95.00 |
| LINC00486 | 3.00 | 1.00 | 0.00 | 1.33 | 0.00 |
| LINC00493 | 579.00 | 659.00 | 608.00 | 615.33 | 906.00 |
| LINC00515 | 3.00 | 0.00 | 1.00 | 1.33 | 3.00 |
| LINC00518 | 11.00 | 9.00 | 13.00 | 11.00 | 18.00 |
| LINC00520 | 5.00 | 9.00 | 1.00 | 5.00 | 1.00 |
| LINC00526 | 82.00 | 93.00 | 82.00 | 85.67 | 99.00 |
| LINC00536 | 3.00 | 2.00 | 0.00 | 1.67 | 0.00 |
| LINC00562 | 5.00 | 6.00 | 8.00 | 6.33 | 8.00 |
| LINC00565 | 773.00 | 766.00 | 785.00 | 774.67 | 734.00 |
| LINC00571 | 5.00 | 1.00 | 1.00 | 2.33 | 2.00 |
| LINC00592 | 15.00 | 16.00 | 14.00 | 15.00 | 24.00 |
| LINC00598 | 12.00 | 3.00 | 5.00 | 6.67 | 0.00 |
| LINC00601 | 36.00 | 53.00 | 44.00 | 44.33 | 43.00 |
| LINC00615 | 3.00 | 1.00 | 3.00 | 2.33 | 0.00 |
| LINC00622 | 58.00 | 62.00 | 51.00 | 57.00 | 44.00 |
| LINC00623 | 151.00 | 157.00 | 141.00 | 149.67 | 158.00 |
| LINC00629 | 77.00 | 75.00 | 61.00 | 71.00 | 66.00 |
| LINC00630 | 86.00 | 80.00 | 89.00 | 85.00 | 71.00 |
| LINC00632 | 80.00 | 88.00 | 102.00 | 90.00 | 57.00 |
| LINC00634 | 3.00 | 2.00 | 4.00 | 3.00 | 4.00 |
| LINC00638 | 145.00 | 127.00 | 168.00 | 146.67 | 113.00 |
| LINC00640 | 11.00 | 14.00 | 9.00 | 11.33 | 8.00 |
| LINC00641 | 582.00 | 554.00 | 659.00 | 598.33 | 369.00 |
| LINC00643 | 77.00 | 66.00 | 64.00 | 69.00 | 101.00 |

| | | | | | |
|---|---|---|---|---|---|
| LINC00645 | 5.00 | 2.00 | 4.00 | 3.67 | 4.00 |
| LINC00648 | 129.00 | 126.00 | 150.00 | 135.00 | 152.00 |
| LINC00654 | 19.00 | 11.00 | 10.00 | 13.33 | 12.00 |
| LINC00657 | 9769.00 | 8800.00 | 10775.00 | 9781.33 | 9603.00 |
| LINC00659 | 3.00 | 0.00 | 0.00 | 1.00 | 7.00 |
| LINC00662 | 100.00 | 88.00 | 94.00 | 94.00 | 113.00 |
| LINC00663 | 36.00 | 20.00 | 38.00 | 31.33 | 36.00 |
| LINC00665 | 35.00 | 34.00 | 42.00 | 37.00 | 41.00 |
| LINC00667 | 1447.00 | 1411.00 | 1594.00 | 1484.00 | 1547.00 |
| LINC00669 | 14.00 | 15.00 | 8.00 | 12.33 | 18.00 |
| LINC00672 | 16.00 | 11.00 | 19.00 | 15.33 | 14.00 |
| LINC00673 | 60.00 | 56.00 | 61.00 | 59.00 | 44.00 |
| LINC00674 | 813.00 | 802.00 | 941.00 | 852.00 | 729.00 |
| LINC00682 | 3.00 | 0.00 | 0.00 | 1.00 | 0.00 |
| LINC00685 | 8.00 | 18.00 | 9.00 | 11.67 | 11.00 |
| LINC00696 | 3.00 | 0.00 | 1.00 | 1.33 | 0.00 |
| LINC00702 | 43.00 | 46.00 | 48.00 | 45.67 | 33.00 |
| LINC00704 | 3258.00 | 3161.00 | 3383.00 | 3267.33 | 2197.00 |
| LINC00706 | 8.00 | 8.00 | 14.00 | 10.00 | 13.00 |
| LINC00707 | 948.00 | 823.00 | 899.00 | 890.00 | 2106.00 |
| LINC00839 | 848.00 | 912.00 | 954.00 | 904.67 | 1060.00 |
| LINC00842 | 25.00 | 21.00 | 25.00 | 23.67 | 15.00 |
| LINC00847 | 239.00 | 225.00 | 287.00 | 250.33 | 243.00 |
| LINC00852 | 35.00 | 51.00 | 51.00 | 45.67 | 38.00 |
| LINC00857 | 15.00 | 19.00 | 12.00 | 15.33 | 14.00 |
| LINC00858 | 39.00 | 35.00 | 40.00 | 38.00 | 36.00 |
| LINC00862 | 6.00 | 5.00 | 14.00 | 8.33 | 10.00 |
| LINC00869 | 48.00 | 54.00 | 49.00 | 50.33 | 43.00 |
| LINC00880 | 6.00 | 4.00 | 11.00 | 7.00 | 9.00 |
| LINC00882 | 18.00 | 17.00 | 11.00 | 15.33 | 19.00 |
| LINC00883 | 226.00 | 200.00 | 218.00 | 214.67 | 240.00 |
| LINC00886 | 43.00 | 20.00 | 23.00 | 28.67 | 25.00 |
| LINC00888 | 145.00 | 145.00 | 166.00 | 152.00 | 147.00 |
| LINC00889 | 4.00 | 9.00 | 7.00 | 6.67 | 4.00 |
| LINC00893 | 20.00 | 15.00 | 22.00 | 19.00 | 13.00 |
| LINC00894 | 96.00 | 92.00 | 108.00 | 98.67 | 93.00 |
| LINC00896 | 8.00 | 11.00 | 12.00 | 10.33 | 8.00 |
| LINC00899 | 181.00 | 167.00 | 177.00 | 175.00 | 193.00 |
| LINC00909 | 234.00 | 215.00 | 251.00 | 233.33 | 318.00 |
| LINC00910 | 68.00 | 83.00 | 56.00 | 69.00 | 72.00 |
| LINC00917 | 39.00 | 36.00 | 42.00 | 39.00 | 69.00 |
| LINC00920 | 42.00 | 31.00 | 33.00 | 35.33 | 55.00 |
| LINC00921 | 45.00 | 51.00 | 46.00 | 47.33 | 56.00 |
| LINC00922 | 54.00 | 44.00 | 44.00 | 47.33 | 52.00 |
| LINC00926 | 35.00 | 40.00 | 28.00 | 34.33 | 44.00 |
| LINC00936 | 26.00 | 26.00 | 25.00 | 25.67 | 11.00 |
| LINC00938 | 231.00 | 218.00 | 229.00 | 226.00 | 244.00 |
| LINC00941 | 343.00 | 386.00 | 359.00 | 362.67 | 450.00 |

| | | | | | |
|---|---|---|---|---|---|
| LINC00942 | 78.00 | 93.00 | 89.00 | 86.67 | 113.00 |
| LINC00950 | 12.00 | 6.00 | 13.00 | 10.33 | 3.00 |
| LINC00957 | 10.00 | 2.00 | 13.00 | 8.33 | 10.00 |
| LINC00959 | 48.00 | 28.00 | 44.00 | 40.00 | 57.00 |
| LINC00960 | 24.00 | 32.00 | 33.00 | 29.67 | 29.00 |
| LINC00963 | 324.00 | 304.00 | 322.00 | 316.67 | 292.00 |
| LINC00987 | 3.00 | 1.00 | 2.00 | 2.00 | 1.00 |
| LINC00997 | 131.00 | 147.00 | 166.00 | 148.00 | 183.00 |
| LINC00998 | 857.00 | 835.00 | 870.00 | 854.00 | 1078.00 |
| LINC00999 | 210.00 | 195.00 | 226.00 | 210.33 | 156.00 |
| LINC01000 | 3545.00 | 3234.00 | 3644.00 | 3474.33 | 2737.00 |
| LINC01003 | 67.00 | 57.00 | 55.00 | 59.67 | 73.00 |
| LINC01004 | 577.00 | 649.00 | 725.00 | 650.33 | 494.00 |
| LINC01011 | 56.00 | 46.00 | 48.00 | 50.00 | 75.00 |
| LINC01012 | 4.00 | 5.00 | 6.00 | 5.00 | 5.00 |
| LINC01021 | 113.00 | 98.00 | 125.00 | 112.00 | 87.00 |
| LINC01023 | 14.00 | 21.00 | 14.00 | 16.33 | 31.00 |
| LINC01024 | 56.00 | 70.00 | 75.00 | 67.00 | 81.00 |
| LINC01029 | 24.00 | 31.00 | 19.00 | 24.67 | 27.00 |
| LINC01032 | 3.00 | 6.00 | 9.00 | 6.00 | 7.00 |
| LINC01057 | 3.00 | 5.00 | 3.00 | 3.67 | 3.00 |
| LINC01061 | 292.00 | 302.00 | 330.00 | 308.00 | 243.00 |
| LINC01085 | 7.00 | 3.00 | 6.00 | 5.33 | 3.00 |
| LINC01088 | 7.00 | 6.00 | 7.00 | 6.67 | 5.00 |
| LINC01089 | 626.00 | 607.00 | 662.00 | 631.67 | 605.00 |
| LINC01094 | 35.00 | 28.00 | 53.00 | 38.67 | 29.00 |
| LINC01096 | 3.00 | 7.00 | 5.00 | 5.00 | 4.00 |
| LINC01098 | 7.00 | 3.00 | 5.00 | 5.00 | 5.00 |
| LINC01111 | 47.00 | 51.00 | 70.00 | 56.00 | 79.00 |
| LINC01116 | 385.00 | 328.00 | 388.00 | 367.00 | 478.00 |
| LINC01117 | 48.00 | 23.00 | 45.00 | 38.67 | 48.00 |
| LINC01118 | 6.00 | 10.00 | 8.00 | 8.00 | 9.00 |
| LINC01119 | 37.00 | 43.00 | 46.00 | 42.00 | 45.00 |
| LINC01124 | 3.00 | 3.00 | 3.00 | 3.00 | 0.00 |
| LINC01125 | 34.00 | 28.00 | 40.00 | 34.00 | 37.00 |
| LINC01126 | 134.00 | 154.00 | 122.00 | 136.67 | 74.00 |
| LINC01128 | 300.00 | 286.00 | 283.00 | 289.67 | 248.00 |
| LINC01134 | 6.00 | 9.00 | 12.00 | 9.00 | 2.00 |
| LINC01135 | 9.00 | 7.00 | 7.00 | 7.67 | 9.00 |
| LINC01137 | 41.00 | 28.00 | 48.00 | 39.00 | 60.00 |
| LINC01138 | 196.00 | 167.00 | 185.00 | 182.67 | 141.00 |
| LINC01139 | 225.00 | 214.00 | 285.00 | 241.33 | 298.00 |
| LINC01140 | 18.00 | 22.00 | 27.00 | 22.33 | 31.00 |
| LINC01144 | 45.00 | 53.00 | 45.00 | 47.67 | 45.00 |
| LINC01151 | 5.00 | 0.00 | 1.00 | 2.00 | 1.00 |
| LINC01152 | 14.00 | 7.00 | 17.00 | 12.67 | 28.00 |
| LINC01160 | 6.00 | 4.00 | 7.00 | 5.67 | 6.00 |
| LINC01176 | 13.00 | 16.00 | 22.00 | 17.00 | 9.00 |

| | | | | | |
|---|---|---|---|---|---|
| LINC01184 | 374.00 | 317.00 | 368.00 | 353.00 | 437.00 |
| LINC01202 | 3.00 | 2.00 | 7.00 | 4.00 | 0.00 |
| LINC01204 | 356.00 | 361.00 | 378.00 | 365.00 | 370.00 |
| LINC01212 | 4.00 | 1.00 | 0.00 | 1.67 | 2.00 |
| LINC01232 | 15.00 | 9.00 | 19.00 | 14.33 | 14.00 |
| LINC01234 | 111.00 | 139.00 | 141.00 | 130.33 | 204.00 |
| LINC01239 | 93.00 | 98.00 | 93.00 | 94.67 | 83.00 |
| LINC01252 | 62.00 | 54.00 | 69.00 | 61.67 | 79.00 |
| LINC01270 | 105.00 | 83.00 | 102.00 | 96.67 | 186.00 |
| LINC01277 | 6.00 | 10.00 | 5.00 | 7.00 | 3.00 |
| LINC01278 | 1084.00 | 960.00 | 1051.00 | 1031.67 | 1017.00 |
| LINC01279 | 379.00 | 348.00 | 348.00 | 358.33 | 556.00 |
| LINC01291 | 22.00 | 23.00 | 33.00 | 26.00 | 37.00 |
| LINC01296 | 146.00 | 144.00 | 194.00 | 161.33 | 136.00 |
| LINC01301 | 11.00 | 15.00 | 15.00 | 13.67 | 15.00 |
| LINC01311 | 97.00 | 107.00 | 109.00 | 104.33 | 103.00 |
| LINC01317 | 18.00 | 15.00 | 14.00 | 15.67 | 15.00 |
| LINC01322 | 152.00 | 141.00 | 130.00 | 141.00 | 161.00 |
| LINC01336 | 3.00 | 0.00 | 0.00 | 1.00 | 0.00 |
| LINC01340 | 6.00 | 2.00 | 3.00 | 3.67 | 2.00 |
| LINC01341 | 14.00 | 14.00 | 14.00 | 14.00 | 18.00 |
| LINC01347 | 3.00 | 5.00 | 6.00 | 4.67 | 3.00 |
| LINC01348 | 3.00 | 5.00 | 1.00 | 3.00 | 2.00 |
| LINC01355 | 655.00 | 620.00 | 736.00 | 670.33 | 643.00 |
| LINC01356 | 10.00 | 16.00 | 20.00 | 15.33 | 13.00 |
| LINC01358 | 58.00 | 61.00 | 93.00 | 70.67 | 145.00 |
| LINC01359 | 7.00 | 7.00 | 11.00 | 8.33 | 4.00 |
| LINC01372 | 26.00 | 22.00 | 16.00 | 21.33 | 16.00 |
| LINC01389 | 4.00 | 4.00 | 1.00 | 3.00 | 2.00 |
| LINC01397 | 4.00 | 3.00 | 2.00 | 3.00 | 2.00 |
| LINC01399 | 76.00 | 86.00 | 92.00 | 84.67 | 58.00 |
| LINC01410 | 512.00 | 470.00 | 501.00 | 494.33 | 408.00 |
| LINC01420 | 630.00 | 563.00 | 637.00 | 610.00 | 747.00 |
| LINC01424 | 19.00 | 34.00 | 19.00 | 24.00 | 27.00 |
| LINC01425 | 22.00 | 16.00 | 29.00 | 22.33 | 25.00 |
| LINC01426 | 300.00 | 328.00 | 311.00 | 313.00 | 346.00 |
| LINC01431 | 6.00 | 9.00 | 7.00 | 7.33 | 8.00 |
| LINC01433 | 11.00 | 5.00 | 8.00 | 8.00 | 5.00 |
| LINC01435 | 6.00 | 4.00 | 8.00 | 6.00 | 15.00 |
| LINC01436 | 419.00 | 412.00 | 385.00 | 405.33 | 472.00 |
| LINC01461 | 11.00 | 12.00 | 15.00 | 12.67 | 9.00 |
| LINC01465 | 5.00 | 14.00 | 10.00 | 9.67 | 11.00 |
| LINC01468 | 42.00 | 37.00 | 40.00 | 39.67 | 38.00 |
| LINC01473 | 14.00 | 7.00 | 14.00 | 11.67 | 12.00 |
| LINC01481 | 71.00 | 88.00 | 77.00 | 78.67 | 66.00 |
| LINC01503 | 3.00 | 0.00 | 0.00 | 1.00 | 0.00 |
| LINC01508 | 4.00 | 1.00 | 0.00 | 1.67 | 0.00 |
| LINC01512 | 14.00 | 20.00 | 18.00 | 17.33 | 8.00 |

| | | | | | |
|---|---|---|---|---|---|
| LINC01518 | 62.00 | 45.00 | 69.00 | 58.67 | 38.00 |
| LINC01521 | 79.00 | 88.00 | 93.00 | 86.67 | 109.00 |
| LINC01530 | 4.00 | 5.00 | 8.00 | 5.67 | 3.00 |
| LINC01534 | 14.00 | 7.00 | 16.00 | 12.33 | 5.00 |
| LINC01535 | 38.00 | 30.00 | 32.00 | 33.33 | 30.00 |
| LINC01543 | 3.00 | 3.00 | 1.00 | 2.33 | 2.00 |
| LINC01545 | 6.00 | 1.00 | 5.00 | 4.00 | 6.00 |
| LINC01547 | 14.00 | 8.00 | 13.00 | 11.67 | 6.00 |
| LINC01551 | 48.00 | 77.00 | 66.00 | 63.67 | 76.00 |
| LINC01554 | 4.00 | 5.00 | 5.00 | 4.67 | 5.00 |
| LINC01560 | 57.00 | 54.00 | 49.00 | 53.33 | 68.00 |
| LINC01561 | 5.00 | 3.00 | 4.00 | 4.00 | 3.00 |
| LINC01569 | 97.00 | 136.00 | 132.00 | 121.67 | 136.00 |
| LINC01572 | 57.00 | 64.00 | 58.00 | 59.67 | 58.00 |
| LINC01573 | 60.00 | 56.00 | 59.00 | 58.33 | 86.00 |
| LINC01578 | 468.00 | 432.00 | 489.00 | 463.00 | 666.00 |
| LINC01583 | 1519.00 | 1509.00 | 1676.00 | 1568.00 | 1223.00 |
| LINC01588 | 36.00 | 33.00 | 57.00 | 42.00 | 42.00 |
| LINC01590 | 57.00 | 63.00 | 65.00 | 61.67 | 57.00 |
| LINC01599 | 7.00 | 9.00 | 14.00 | 10.00 | 3.00 |
| LINC01600 | 7.00 | 3.00 | 13.00 | 7.67 | 7.00 |
| LINC01602 | 3.00 | 2.00 | 2.00 | 2.33 | 4.00 |
| LINC01604 | 245.00 | 246.00 | 223.00 | 238.00 | 227.00 |
| LINC01605 | 905.00 | 776.00 | 842.00 | 841.00 | 833.00 |
| LINC01611 | 5.00 | 12.00 | 17.00 | 11.33 | 11.00 |
| LINC-PINT | 1176.00 | 1164.00 | 1270.00 | 1203.33 | 1047.00 |
| LINCR-0001 | 6.00 | 6.00 | 14.00 | 8.67 | 4.00 |
| LINGO2 | 45.00 | 40.00 | 79.00 | 54.67 | 39.00 |
| LINS | 280.00 | 243.00 | 234.00 | 252.33 | 253.00 |
| LIPA | 3840.00 | 3586.00 | 3744.00 | 3723.33 | 3639.00 |
| LIPE | 16.00 | 16.00 | 23.00 | 18.33 | 24.00 |
| LIPE-AS1 | 17.00 | 16.00 | 12.00 | 15.00 | 22.00 |
| LIPT1 | 292.00 | 246.00 | 304.00 | 280.67 | 290.00 |
| LIPT2 | 17.00 | 6.00 | 9.00 | 10.67 | 33.00 |
| LITAF | 45.00 | 38.00 | 57.00 | 46.67 | 79.00 |
| LIX1L | 1073.00 | 966.00 | 1063.00 | 1034.00 | 1193.00 |
| LLGL1 | 1449.00 | 1434.00 | 1487.00 | 1456.67 | 1347.00 |
| LLGL2 | 19.00 | 40.00 | 29.00 | 29.33 | 30.00 |
| LLPH | 519.00 | 483.00 | 515.00 | 505.67 | 622.00 |
| LLPH-AS1 | 12.00 | 10.00 | 14.00 | 12.00 | 14.00 |
| LMAN1 | 6717.00 | 6004.00 | 6781.00 | 6500.67 | 5975.00 |
| LMAN2 | 6008.00 | 6196.00 | 5767.00 | 5990.33 | 4661.00 |
| LMAN2L | 800.00 | 889.00 | 900.00 | 863.00 | 809.00 |
| LMBR1 | 1180.00 | 1110.00 | 1227.00 | 1172.33 | 1189.00 |
| LMBR1L | 1232.00 | 1236.00 | 1317.00 | 1261.67 | 1113.00 |
| LMBRD1 | 661.00 | 653.00 | 721.00 | 678.33 | 705.00 |
| LMBRD2 | 341.00 | 280.00 | 316.00 | 312.33 | 200.00 |
| LMCD1 | 54.00 | 70.00 | 72.00 | 65.33 | 59.00 |

| | | | | | |
|---|---|---|---|---|---|
| LMF2 | 1959.00 | 2047.00 | 1997.00 | 2001.00 | 1973.00 |
| LMLN | 869.00 | 845.00 | 848.00 | 854.00 | 816.00 |
| LMLN-AS1 | 15.00 | 20.00 | 15.00 | 16.67 | 20.00 |
| LMNA | 3470.00 | 3437.00 | 3433.00 | 3446.67 | 3962.00 |
| LMNB1 | 874.00 | 768.00 | 878.00 | 840.00 | 1287.00 |
| LMNB2 | 7016.00 | 7012.00 | 7178.00 | 7068.67 | 6347.00 |
| LMNTD2 | 3.00 | 2.00 | 3.00 | 2.67 | 1.00 |
| LMO4 | 1623.00 | 1542.00 | 1836.00 | 1667.00 | 2278.00 |
| LMO7 | 1033.00 | 1000.00 | 1053.00 | 1028.67 | 947.00 |
| LMOD3 | 4.00 | 2.00 | 1.00 | 2.33 | 1.00 |
| LMTK2 | 1177.00 | 1203.00 | 1168.00 | 1182.67 | 860.00 |
| LNP1 | 110.00 | 94.00 | 131.00 | 111.67 | 153.00 |
| LNPEP | 391.00 | 309.00 | 344.00 | 348.00 | 206.00 |
| LNX1 | 42.00 | 37.00 | 26.00 | 35.00 | 43.00 |
| LNX2 | 197.00 | 155.00 | 190.00 | 180.67 | 159.00 |
| LOC100049716 | 3.00 | 0.00 | 0.00 | 1.00 | 2.00 |
| LOC100126784 | 169.00 | 196.00 | 210.00 | 191.67 | 202.00 |
| LOC100128076 | 3.00 | 3.00 | 7.00 | 4.33 | 13.00 |
| LOC100128164 | 5.00 | 11.00 | 3.00 | 6.33 | 8.00 |
| LOC100128233 | 17.00 | 11.00 | 9.00 | 12.33 | 3.00 |
| LOC100128288 | 74.00 | 52.00 | 80.00 | 68.67 | 68.00 |
| LOC100128361 | 44.00 | 38.00 | 34.00 | 38.67 | 44.00 |
| LOC100128398 | 9.00 | 11.00 | 3.00 | 7.67 | 5.00 |
| LOC100128494 | 82.00 | 73.00 | 69.00 | 74.67 | 57.00 |
| LOC100128568 | 19.00 | 13.00 | 12.00 | 14.67 | 10.00 |
| LOC100128573 | 3.00 | 1.00 | 1.00 | 1.67 | 2.00 |
| LOC100128770 | 10.00 | 10.00 | 11.00 | 10.33 | 2.00 |
| LOC100129027 | 16.00 | 25.00 | 27.00 | 22.67 | 12.00 |
| LOC100129034 | 1813.00 | 1818.00 | 1840.00 | 1823.67 | 1329.00 |
| LOC100129203 | 23.00 | 26.00 | 30.00 | 26.33 | 15.00 |
| LOC100129434 | 5.00 | 1.00 | 4.00 | 3.33 | 1.00 |
| LOC100129461 | 90.00 | 69.00 | 92.00 | 83.67 | 59.00 |
| LOC100129518 | 5.00 | 1.00 | 1.00 | 2.33 | 7.00 |
| LOC100129534 | 108.00 | 89.00 | 86.00 | 94.33 | 77.00 |
| LOC100129550 | 386.00 | 348.00 | 383.00 | 372.33 | 307.00 |
| LOC100129617 | 27.00 | 28.00 | 26.00 | 27.00 | 31.00 |
| LOC100129917 | 92.00 | 85.00 | 94.00 | 90.33 | 33.00 |
| LOC100129924 | 6.00 | 6.00 | 3.00 | 5.00 | 2.00 |
| LOC100130357 | 119.00 | 121.00 | 134.00 | 124.67 | 92.00 |
| LOC100130451 | 5.00 | 11.00 | 0.00 | 5.33 | 2.00 |
| LOC100130691 | 204.00 | 188.00 | 207.00 | 199.67 | 201.00 |
| LOC100130705 | 31.00 | 31.00 | 44.00 | 35.33 | 24.00 |
| LOC100130744 | 225.00 | 228.00 | 265.00 | 239.33 | 213.00 |
| LOC100130950 | 62.00 | 81.00 | 58.00 | 67.00 | 61.00 |
| LOC100130987 | 28.00 | 35.00 | 25.00 | 29.33 | 38.00 |
| LOC100130992 | 25.00 | 22.00 | 31.00 | 26.00 | 22.00 |
| LOC100131564 | 1945.00 | 1940.00 | 2128.00 | 2004.33 | 1788.00 |
| LOC100131655 | 17.00 | 10.00 | 7.00 | 11.33 | 13.00 |

| | | | | | |
|---|---|---|---|---|---|
| LOC100132057 | 11.00 | 13.00 | 16.00 | 13.33 | 9.00 |
| LOC100132356 | 22.00 | 23.00 | 18.00 | 21.00 | 26.00 |
| LOC100133050 | 8.00 | 3.00 | 2.00 | 4.33 | 3.00 |
| LOC100133077 | 3.00 | 0.00 | 0.00 | 1.00 | 0.00 |
| LOC100133315 | 67.00 | 96.00 | 90.00 | 84.33 | 82.00 |
| LOC100133331 | 85.00 | 72.00 | 96.00 | 84.33 | 79.00 |
| LOC100133985 | 9.00 | 4.00 | 17.00 | 10.00 | 4.00 |
| LOC100134368 | 18.00 | 11.00 | 13.00 | 14.00 | 9.00 |
| LOC100134868 | 35.00 | 42.00 | 45.00 | 40.67 | 40.00 |
| LOC100147773 | 18.00 | 15.00 | 10.00 | 14.33 | 14.00 |
| LOC100190986 | 37.00 | 41.00 | 37.00 | 38.33 | 13.00 |
| LOC100233156 | 232.00 | 225.00 | 280.00 | 245.67 | 225.00 |
| LOC100268168 | 215.00 | 229.00 | 203.00 | 215.67 | 268.00 |
| LOC100270746 | 25.00 | 17.00 | 27.00 | 23.00 | 28.00 |
| LOC100270804 | 130.00 | 125.00 | 114.00 | 123.00 | 117.00 |
| LOC100272217 | 72.00 | 61.00 | 75.00 | 69.33 | 48.00 |
| LOC100287015 | 71.00 | 61.00 | 53.00 | 61.67 | 83.00 |
| LOC100287036 | 94.00 | 91.00 | 92.00 | 92.33 | 36.00 |
| LOC100287042 | 103.00 | 99.00 | 77.00 | 93.00 | 100.00 |
| LOC100287072 | 6.00 | 6.00 | 10.00 | 7.33 | 4.00 |
| LOC100287225 | 5.00 | 7.00 | 5.00 | 5.67 | 5.00 |
| LOC100288069 | 54.00 | 50.00 | 63.00 | 55.67 | 30.00 |
| LOC100288152 | 387.00 | 427.00 | 474.00 | 429.33 | 378.00 |
| LOC100288181 | 18.00 | 13.00 | 21.00 | 17.33 | 30.00 |
| LOC100288637 | 67.00 | 76.00 | 71.00 | 71.33 | 92.00 |
| LOC100288778 | 9.00 | 22.00 | 23.00 | 18.00 | 17.00 |
| LOC100288798 | 31.00 | 31.00 | 38.00 | 33.33 | 30.00 |
| LOC100288842 | 30.00 | 23.00 | 25.00 | 26.00 | 34.00 |
| LOC100288846 | 8.00 | 10.00 | 5.00 | 7.67 | 6.00 |
| LOC100289230 | 73.00 | 111.00 | 99.00 | 94.33 | 116.00 |
| LOC100289361 | 17.00 | 9.00 | 13.00 | 13.00 | 17.00 |
| LOC100289473 | 5.00 | 2.00 | 3.00 | 3.33 | 1.00 |
| LOC100289495 | 6.00 | 17.00 | 25.00 | 16.00 | 10.00 |
| LOC100289561 | 27.00 | 28.00 | 23.00 | 26.00 | 46.00 |
| LOC100289650 | 40.00 | 37.00 | 36.00 | 37.67 | 29.00 |
| LOC100289673 | 6.00 | 1.00 | 5.00 | 4.00 | 1.00 |
| LOC100294362 | 50.00 | 37.00 | 46.00 | 44.33 | 39.00 |
| LOC100335030 | 11.00 | 8.00 | 13.00 | 10.67 | 26.00 |
| LOC100379224 | 5.00 | 11.00 | 15.00 | 10.33 | 17.00 |
| LOC100499484 | 58.00 | 58.00 | 48.00 | 54.67 | 44.00 |
| LOC100499489 | 68.00 | 92.00 | 71.00 | 77.00 | 109.00 |
| LOC100505549 | 70.00 | 51.00 | 60.00 | 60.33 | 78.00 |
| LOC100505622 | 54.00 | 53.00 | 50.00 | 52.33 | 54.00 |
| LOC100505635 | 7.00 | 4.00 | 3.00 | 4.67 | 3.00 |
| LOC100505666 | 44.00 | 50.00 | 60.00 | 51.33 | 67.00 |
| LOC100505715 | 38.00 | 36.00 | 44.00 | 39.33 | 35.00 |
| LOC100505912 | 19.00 | 15.00 | 18.00 | 17.33 | 6.00 |
| LOC100505938 | 33.00 | 37.00 | 27.00 | 32.33 | 22.00 |

| | | | | | |
|---|---|---|---|---|---|
| LOC100505942 | 6.00 | 8.00 | 2.00 | 5.33 | 2.00 |
| LOC100505984 | 4.00 | 9.00 | 12.00 | 8.33 | 9.00 |
| LOC100506022 | 10.00 | 6.00 | 19.00 | 11.67 | 16.00 |
| LOC100506023 | 20.00 | 24.00 | 23.00 | 22.33 | 18.00 |
| LOC100506076 | 4.00 | 3.00 | 4.00 | 3.67 | 1.00 |
| LOC100506082 | 5.00 | 4.00 | 3.00 | 4.00 | 5.00 |
| LOC100506083 | 19.00 | 39.00 | 23.00 | 27.00 | 29.00 |
| LOC100506100 | 24.00 | 25.00 | 19.00 | 22.67 | 38.00 |
| LOC100506124 | 43.00 | 46.00 | 48.00 | 45.67 | 48.00 |
| LOC100506125 | 3.00 | 2.00 | 0.00 | 1.67 | 0.00 |
| LOC100506127 | 369.00 | 313.00 | 390.00 | 357.33 | 277.00 |
| LOC100506136 | 24.00 | 22.00 | 26.00 | 24.00 | 31.00 |
| LOC100506178 | 44.00 | 60.00 | 56.00 | 53.33 | 103.00 |
| LOC100506188 | 5.00 | 3.00 | 10.00 | 6.00 | 0.00 |
| LOC100506207 | 19.00 | 28.00 | 21.00 | 22.67 | 31.00 |
| LOC100506271 | 5.00 | 8.00 | 10.00 | 7.67 | 2.00 |
| LOC100506302 | 207.00 | 197.00 | 243.00 | 215.67 | 222.00 |
| LOC100506368 | 6.00 | 4.00 | 2.00 | 4.00 | 10.00 |
| LOC100506472 | 38.00 | 40.00 | 42.00 | 40.00 | 52.00 |
| LOC100506476 | 48.00 | 52.00 | 57.00 | 52.33 | 50.00 |
| LOC100506548 | 2782.00 | 2772.00 | 3175.00 | 2909.67 | 2928.00 |
| LOC100506603 | 256.00 | 281.00 | 234.00 | 257.00 | 255.00 |
| LOC100506606 | 49.00 | 41.00 | 44.00 | 44.67 | 34.00 |
| LOC100506639 | 196.00 | 171.00 | 210.00 | 192.33 | 156.00 |
| LOC100506730 | 61.00 | 68.00 | 79.00 | 69.33 | 64.00 |
| LOC100506746 | 52.00 | 49.00 | 40.00 | 47.00 | 42.00 |
| LOC100506801 | 5.00 | 8.00 | 13.00 | 8.67 | 12.00 |
| LOC100506804 | 21.00 | 19.00 | 20.00 | 20.00 | 16.00 |
| LOC100506844 | 133.00 | 156.00 | 137.00 | 142.00 | 170.00 |
| LOC100506860 | 72.00 | 58.00 | 54.00 | 61.33 | 33.00 |
| LOC100506990 | 34.00 | 37.00 | 36.00 | 35.67 | 33.00 |
| LOC100507002 | 8.00 | 12.00 | 10.00 | 10.00 | 3.00 |
| LOC100507053 | 86.00 | 108.00 | 82.00 | 92.00 | 117.00 |
| LOC100507065 | 1818.00 | 1658.00 | 1979.00 | 1818.33 | 1350.00 |
| LOC100507144 | 3.00 | 6.00 | 3.00 | 4.00 | 7.00 |
| LOC100507156 | 30.00 | 22.00 | 27.00 | 26.33 | 20.00 |
| LOC100507250 | 5.00 | 16.00 | 6.00 | 9.00 | 16.00 |
| LOC100507283 | 33.00 | 39.00 | 34.00 | 35.33 | 32.00 |
| LOC100507291 | 33.00 | 39.00 | 47.00 | 39.67 | 23.00 |
| LOC100507373 | 193.00 | 223.00 | 235.00 | 217.00 | 247.00 |
| LOC100507412 | 20.00 | 58.00 | 13.00 | 30.33 | 10.00 |
| LOC100507424 | 84.00 | 85.00 | 101.00 | 90.00 | 111.00 |
| LOC100507437 | 39.00 | 28.00 | 32.00 | 33.00 | 24.00 |
| LOC100507462 | 71.00 | 55.00 | 71.00 | 65.67 | 91.00 |
| LOC100507487 | 93.00 | 85.00 | 124.00 | 100.67 | 111.00 |
| LOC100507506 | 45.00 | 37.00 | 42.00 | 41.33 | 45.00 |
| LOC100507557 | 14.00 | 10.00 | 20.00 | 14.67 | 7.00 |
| LOC100507564 | 28.00 | 23.00 | 23.00 | 24.67 | 24.00 |

| | | | | | |
|---|---|---|---|---|---|
| LOC100507577 | 286.00 | 286.00 | 298.00 | 290.00 | 308.00 |
| LOC100507600 | 6.00 | 4.00 | 10.00 | 6.67 | 19.00 |
| LOC100507634 | 21.00 | 12.00 | 33.00 | 22.00 | 18.00 |
| LOC100630923 | 10.00 | 8.00 | 15.00 | 11.00 | 16.00 |
| LOC100652758 | 32.00 | 30.00 | 41.00 | 34.33 | 47.00 |
| LOC100652768 | 33.00 | 17.00 | 27.00 | 25.67 | 15.00 |
| LOC100996351 | 4.00 | 1.00 | 2.00 | 2.33 | 2.00 |
| LOC100996437 | 79.00 | 78.00 | 86.00 | 81.00 | 95.00 |
| LOC100996634 | 4.00 | 0.00 | 1.00 | 1.67 | 0.00 |
| LOC101059948 | 11.00 | 23.00 | 14.00 | 16.00 | 21.00 |
| LOC101060091 | 9.00 | 7.00 | 8.00 | 8.00 | 12.00 |
| LOC101241902 | 6.00 | 9.00 | 17.00 | 10.67 | 5.00 |
| LOC101409256 | 33.00 | 18.00 | 35.00 | 28.67 | 22.00 |
| LOC101448202 | 2689.00 | 2502.00 | 2942.00 | 2711.00 | 2474.00 |
| LOC101559451 | 11.00 | 11.00 | 8.00 | 10.00 | 19.00 |
| LOC101926911 | 4.00 | 2.00 | 4.00 | 3.33 | 8.00 |
| LOC101926913 | 5.00 | 4.00 | 6.00 | 5.00 | 10.00 |
| LOC101926935 | 32.00 | 24.00 | 24.00 | 26.67 | 24.00 |
| LOC101926940 | 10.00 | 12.00 | 10.00 | 10.67 | 10.00 |
| LOC101926941 | 47.00 | 47.00 | 43.00 | 45.67 | 37.00 |
| LOC101926963 | 102.00 | 91.00 | 98.00 | 97.00 | 101.00 |
| LOC101926975 | 6.00 | 4.00 | 13.00 | 7.67 | 5.00 |
| LOC101927021 | 170.00 | 185.00 | 170.00 | 175.00 | 257.00 |
| LOC101927027 | 649.00 | 618.00 | 611.00 | 626.00 | 488.00 |
| LOC101927040 | 310.00 | 304.00 | 341.00 | 318.33 | 219.00 |
| LOC101927043 | 5.00 | 1.00 | 1.00 | 2.33 | 0.00 |
| LOC101927045 | 132.00 | 134.00 | 124.00 | 130.00 | 117.00 |
| LOC101927051 | 4.00 | 6.00 | 6.00 | 5.33 | 6.00 |
| LOC101927056 | 25.00 | 24.00 | 33.00 | 27.33 | 18.00 |
| LOC101927087 | 9.00 | 10.00 | 15.00 | 11.33 | 0.00 |
| LOC101927124 | 3.00 | 1.00 | 0.00 | 1.33 | 5.00 |
| LOC101927132 | 3.00 | 8.00 | 5.00 | 5.33 | 3.00 |
| LOC101927151 | 128.00 | 99.00 | 130.00 | 119.00 | 108.00 |
| LOC101927178 | 11.00 | 7.00 | 7.00 | 8.33 | 6.00 |
| LOC101927181 | 69.00 | 51.00 | 69.00 | 63.00 | 49.00 |
| LOC101927196 | 5.00 | 6.00 | 2.00 | 4.33 | 2.00 |
| LOC101927204 | 24.00 | 15.00 | 33.00 | 24.00 | 14.00 |
| LOC101927230 | 67.00 | 64.00 | 75.00 | 68.67 | 41.00 |
| LOC101927237 | 6.00 | 7.00 | 2.00 | 5.00 | 5.00 |
| LOC101927282 | 3.00 | 1.00 | 3.00 | 2.33 | 0.00 |
| LOC101927322 | 20.00 | 11.00 | 14.00 | 15.00 | 15.00 |
| LOC101927354 | 7.00 | 8.00 | 12.00 | 9.00 | 5.00 |
| LOC101927359 | 7.00 | 2.00 | 7.00 | 5.33 | 4.00 |
| LOC101927365 | 53.00 | 50.00 | 54.00 | 52.33 | 67.00 |
| LOC101927391 | 22.00 | 24.00 | 22.00 | 22.67 | 18.00 |
| LOC101927415 | 37.00 | 34.00 | 36.00 | 35.67 | 19.00 |
| LOC101927481 | 8.00 | 3.00 | 12.00 | 7.67 | 10.00 |
| LOC101927482 | 23.00 | 8.00 | 18.00 | 16.33 | 17.00 |

| | | | | | |
|---|---|---|---|---|---|
| LOC101927497 | 155.00 | 141.00 | 124.00 | 140.00 | 347.00 |
| LOC101927549 | 7.00 | 6.00 | 5.00 | 6.00 | 8.00 |
| LOC101927550 | 58.00 | 50.00 | 55.00 | 54.33 | 50.00 |
| LOC101927571 | 3.00 | 1.00 | 5.00 | 3.00 | 3.00 |
| LOC101927572 | 25.00 | 30.00 | 25.00 | 26.67 | 42.00 |
| LOC101927588 | 3.00 | 2.00 | 3.00 | 2.67 | 7.00 |
| LOC101927604 | 5.00 | 3.00 | 3.00 | 3.67 | 2.00 |
| LOC101927620 | 10.00 | 2.00 | 4.00 | 5.33 | 1.00 |
| LOC101927701 | 10.00 | 6.00 | 11.00 | 9.00 | 23.00 |
| LOC101927746 | 91.00 | 93.00 | 88.00 | 90.67 | 135.00 |
| LOC101927755 | 9.00 | 5.00 | 9.00 | 7.67 | 9.00 |
| LOC101927762 | 3.00 | 2.00 | 5.00 | 3.33 | 2.00 |
| LOC101927765 | 37.00 | 40.00 | 55.00 | 44.00 | 44.00 |
| LOC101927768 | 8.00 | 6.00 | 11.00 | 8.33 | 5.00 |
| LOC101927780 | 17.00 | 23.00 | 32.00 | 24.00 | 13.00 |
| LOC101927787 | 15.00 | 8.00 | 13.00 | 12.00 | 11.00 |
| LOC101927811 | 15.00 | 13.00 | 15.00 | 14.33 | 29.00 |
| LOC101927843 | 47.00 | 28.00 | 37.00 | 37.33 | 36.00 |
| LOC101927865 | 61.00 | 54.00 | 59.00 | 58.00 | 79.00 |
| LOC101927911 | 4.00 | 5.00 | 2.00 | 3.67 | 8.00 |
| LOC101927919 | 24.00 | 17.00 | 19.00 | 20.00 | 23.00 |
| LOC101927932 | 5.00 | 3.00 | 15.00 | 7.67 | 12.00 |
| LOC101928020 | 43.00 | 54.00 | 47.00 | 48.00 | 38.00 |
| LOC101928053 | 14.00 | 14.00 | 20.00 | 16.00 | 18.00 |
| LOC101928068 | 116.00 | 126.00 | 120.00 | 120.67 | 115.00 |
| LOC101928069 | 163.00 | 150.00 | 128.00 | 147.00 | 152.00 |
| LOC101928103 | 12.00 | 6.00 | 8.00 | 8.67 | 12.00 |
| LOC101928161 | 3.00 | 3.00 | 9.00 | 5.00 | 8.00 |
| LOC101928201 | 22.00 | 33.00 | 20.00 | 25.00 | 15.00 |
| LOC101928222 | 16.00 | 15.00 | 16.00 | 15.67 | 16.00 |
| LOC101928269 | 24.00 | 34.00 | 28.00 | 28.67 | 38.00 |
| LOC101928307 | 5.00 | 11.00 | 6.00 | 7.33 | 14.00 |
| LOC101928323 | 14.00 | 7.00 | 6.00 | 9.00 | 2.00 |
| LOC101928324 | 4.00 | 12.00 | 6.00 | 7.33 | 7.00 |
| LOC101928402 | 4.00 | 8.00 | 8.00 | 6.67 | 3.00 |
| LOC101928414 | 7.00 | 4.00 | 7.00 | 6.00 | 8.00 |
| LOC101928416 | 3.00 | 1.00 | 0.00 | 1.33 | 0.00 |
| LOC101928438 | 6.00 | 8.00 | 1.00 | 5.00 | 6.00 |
| LOC101928445 | 8.00 | 10.00 | 9.00 | 9.00 | 13.00 |
| LOC101928446 | 3.00 | 3.00 | 3.00 | 3.00 | 2.00 |
| LOC101928453 | 141.00 | 156.00 | 165.00 | 154.00 | 174.00 |
| LOC101928461 | 3.00 | 1.00 | 2.00 | 2.00 | 5.00 |
| LOC101928489 | 16.00 | 12.00 | 20.00 | 16.00 | 11.00 |
| LOC101928509 | 8.00 | 9.00 | 25.00 | 14.00 | 9.00 |
| LOC101928530 | 5.00 | 4.00 | 2.00 | 3.67 | 2.00 |
| LOC101928673 | 34.00 | 47.00 | 42.00 | 41.00 | 60.00 |
| LOC101928674 | 5.00 | 4.00 | 10.00 | 6.33 | 7.00 |
| LOC101928710 | 21.00 | 17.00 | 23.00 | 20.33 | 16.00 |

| | | | | | |
|---|---|---|---|---|---|
| LOC101928766 | 3.00 | 0.00 | 1.00 | 1.33 | 0.00 |
| LOC101928767 | 6.00 | 2.00 | 2.00 | 3.33 | 7.00 |
| LOC101928794 | 5.00 | 9.00 | 6.00 | 6.67 | 6.00 |
| LOC101928796 | 30.00 | 32.00 | 38.00 | 33.33 | 20.00 |
| LOC101928837 | 6.00 | 5.00 | 8.00 | 6.33 | 4.00 |
| LOC101928841 | 4.00 | 1.00 | 4.00 | 3.00 | 1.00 |
| LOC101928865 | 35.00 | 30.00 | 26.00 | 30.33 | 17.00 |
| LOC101928979 | 9.00 | 2.00 | 8.00 | 6.33 | 1.00 |
| LOC101928994 | 18.00 | 17.00 | 19.00 | 18.00 | 10.00 |
| LOC101929057 | 3.00 | 5.00 | 12.00 | 6.67 | 2.00 |
| LOC101929066 | 17.00 | 27.00 | 25.00 | 23.00 | 38.00 |
| LOC101929125 | 22.00 | 19.00 | 27.00 | 22.67 | 19.00 |
| LOC101929140 | 14.00 | 19.00 | 13.00 | 15.33 | 27.00 |
| LOC101929147 | 614.00 | 700.00 | 716.00 | 676.67 | 783.00 |
| LOC101929217 | 3.00 | 0.00 | 0.00 | 1.00 | 0.00 |
| LOC101929224 | 5.00 | 10.00 | 8.00 | 7.67 | 8.00 |
| LOC101929234 | 30.00 | 34.00 | 47.00 | 37.00 | 32.00 |
| LOC101929268 | 37.00 | 37.00 | 37.00 | 37.00 | 31.00 |
| LOC101929295 | 8.00 | 12.00 | 8.00 | 9.33 | 5.00 |
| LOC101929378 | 45.00 | 28.00 | 33.00 | 35.33 | 32.00 |
| LOC101929380 | 41.00 | 30.00 | 30.00 | 33.67 | 84.00 |
| LOC101929406 | 4.00 | 0.00 | 1.00 | 1.67 | 1.00 |
| LOC101929464 | 5.00 | 2.00 | 2.00 | 3.00 | 3.00 |
| LOC101929555 | 3.00 | 1.00 | 3.00 | 2.33 | 8.00 |
| LOC101929574 | 4.00 | 8.00 | 3.00 | 5.00 | 10.00 |
| LOC101929577 | 3.00 | 5.00 | 7.00 | 5.00 | 5.00 |
| LOC101929679 | 12.00 | 11.00 | 7.00 | 10.00 | 6.00 |
| LOC101929705 | 66.00 | 48.00 | 77.00 | 63.67 | 20.00 |
| LOC101929709 | 220.00 | 239.00 | 250.00 | 236.33 | 226.00 |
| LOC101929710 | 7.00 | 4.00 | 13.00 | 8.00 | 5.00 |
| LOC101929715 | 51.00 | 61.00 | 45.00 | 52.33 | 61.00 |
| LOC101929719 | 23.00 | 26.00 | 16.00 | 21.67 | 25.00 |
| LOC101929741 | 7.00 | 10.00 | 8.00 | 8.33 | 1.00 |
| LOC101929762 | 6.00 | 12.00 | 5.00 | 7.67 | 9.00 |
| LOC101929767 | 90.00 | 74.00 | 107.00 | 90.33 | 72.00 |
| LOC101930071 | 10.00 | 10.00 | 11.00 | 10.33 | 9.00 |
| LOC101930452 | 17.00 | 26.00 | 19.00 | 20.67 | 21.00 |
| LOC102477328 | 26.00 | 20.00 | 18.00 | 21.33 | 27.00 |
| LOC102546294 | 14.00 | 12.00 | 15.00 | 13.67 | 24.00 |
| LOC102606465 | 360.00 | 383.00 | 416.00 | 386.33 | 464.00 |
| LOC102723354 | 41.00 | 36.00 | 25.00 | 34.00 | 23.00 |
| LOC102723505 | 3.00 | 1.00 | 5.00 | 3.00 | 5.00 |
| LOC102723582 | 8.00 | 4.00 | 11.00 | 7.67 | 7.00 |
| LOC102723701 | 5.00 | 5.00 | 4.00 | 4.67 | 4.00 |
| LOC102723703 | 20.00 | 21.00 | 31.00 | 24.00 | 17.00 |
| LOC102723780 | 8.00 | 6.00 | 7.00 | 7.00 | 5.00 |
| LOC102723968 | 4.00 | 5.00 | 6.00 | 5.00 | 0.00 |
| LOC102724297 | 3.00 | 1.00 | 0.00 | 1.33 | 3.00 |

| | | | | | |
|---|---|---|---|---|---|
| LOC102724301 | 7.00 | 3.00 | 3.00 | 4.33 | 8.00 |
| LOC102724434 | 145.00 | 134.00 | 150.00 | 143.00 | 210.00 |
| LOC102724450 | 53.00 | 43.00 | 56.00 | 50.67 | 71.00 |
| LOC102724467 | 29.00 | 27.00 | 24.00 | 26.67 | 27.00 |
| LOC102724552 | 7.00 | 5.00 | 3.00 | 5.00 | 4.00 |
| LOC102724596 | 18.00 | 20.00 | 20.00 | 19.33 | 16.00 |
| LOC102724623 | 4.00 | 0.00 | 5.00 | 3.00 | 11.00 |
| LOC102724699 | 226.00 | 227.00 | 224.00 | 225.67 | 237.00 |
| LOC102724814 | 675.00 | 758.00 | 759.00 | 730.67 | 483.00 |
| LOC102724919 | 4.00 | 4.00 | 2.00 | 3.33 | 2.00 |
| LOC103091866 | 32.00 | 24.00 | 49.00 | 35.00 | 23.00 |
| LOC103344931 | 414.00 | 456.00 | 434.00 | 434.67 | 372.00 |
| LOC103611081 | 282.00 | 287.00 | 277.00 | 282.00 | 232.00 |
| LOC103908605 | 60.00 | 82.00 | 90.00 | 77.33 | 99.00 |
| LOC104968399 | 5.00 | 3.00 | 0.00 | 2.67 | 3.00 |
| LOC105274304 | 19.00 | 7.00 | 9.00 | 11.67 | 18.00 |
| LOC105373300 | 14.00 | 9.00 | 12.00 | 11.67 | 9.00 |
| LOC105373383 | 3.00 | 4.00 | 5.00 | 4.00 | 5.00 |
| LOC105377348 | 13.00 | 17.00 | 26.00 | 18.67 | 20.00 |
| LOC105378732 | 13.00 | 9.00 | 11.00 | 11.00 | 12.00 |
| LOC105447645 | 9.00 | 7.00 | 14.00 | 10.00 | 8.00 |
| LOC105747689 | 395.00 | 414.00 | 422.00 | 410.33 | 343.00 |
| LOC105748977 | 7.00 | 8.00 | 3.00 | 6.00 | 8.00 |
| LOC106099130 | 9.00 | 7.00 | 2.00 | 6.00 | 3.00 |
| LOC113230 | 8.00 | 9.00 | 12.00 | 9.67 | 8.00 |
| LOC145783 | 43.00 | 53.00 | 46.00 | 47.33 | 57.00 |
| LOC146880 | 2000.00 | 2046.00 | 2255.00 | 2100.33 | 2130.00 |
| LOC148413 | 617.00 | 658.00 | 687.00 | 654.00 | 698.00 |
| LOC150776 | 945.00 | 982.00 | 1032.00 | 986.33 | 936.00 |
| LOC152225 | 170.00 | 176.00 | 169.00 | 171.67 | 174.00 |
| LOC153684 | 95.00 | 92.00 | 113.00 | 100.00 | 75.00 |
| LOC154761 | 4.00 | 2.00 | 5.00 | 3.67 | 2.00 |
| LOC155060 | 1028.00 | 1122.00 | 1197.00 | 1115.67 | 783.00 |
| LOC171391 | 211.00 | 228.00 | 203.00 | 214.00 | 223.00 |
| LOC202181 | 73.00 | 89.00 | 85.00 | 82.33 | 61.00 |
| LOC220729 | 530.00 | 527.00 | 548.00 | 535.00 | 467.00 |
| LOC254896 | 57.00 | 66.00 | 83.00 | 68.67 | 84.00 |
| LOC255187 | 14.00 | 14.00 | 13.00 | 13.67 | 40.00 |
| LOC257396 | 231.00 | 214.00 | 195.00 | 213.33 | 253.00 |
| LOC283140 | 3.00 | 3.00 | 5.00 | 3.67 | 4.00 |
| LOC283335 | 12.00 | 17.00 | 16.00 | 15.00 | 6.00 |
| LOC283352 | 13.00 | 15.00 | 15.00 | 14.33 | 13.00 |
| LOC283440 | 4.00 | 2.00 | 2.00 | 2.67 | 0.00 |
| LOC283575 | 9.00 | 6.00 | 8.00 | 7.67 | 3.00 |
| LOC283683 | 3.00 | 11.00 | 5.00 | 6.33 | 7.00 |
| LOC283788 | 484.00 | 394.00 | 474.00 | 450.67 | 442.00 |
| LOC283922 | 153.00 | 142.00 | 162.00 | 152.33 | 128.00 |
| LOC284009 | 25.00 | 32.00 | 26.00 | 27.67 | 33.00 |

| | | | | | |
|---|---|---|---|---|---|
| LOC284023 | 119.00 | 133.00 | 150.00 | 134.00 | 208.00 |
| LOC284080 | 192.00 | 176.00 | 189.00 | 185.67 | 126.00 |
| LOC284454 | 701.00 | 760.00 | 816.00 | 759.00 | 780.00 |
| LOC284581 | 487.00 | 504.00 | 559.00 | 516.67 | 440.00 |
| LOC284930 | 22.00 | 9.00 | 12.00 | 14.33 | 15.00 |
| LOC284950 | 3.00 | 0.00 | 3.00 | 2.00 | 0.00 |
| LOC285074 | 178.00 | 179.00 | 231.00 | 196.00 | 157.00 |
| LOC285696 | 15.00 | 12.00 | 19.00 | 15.33 | 16.00 |
| LOC285768 | 29.00 | 12.00 | 32.00 | 24.33 | 23.00 |
| LOC286437 | 224.00 | 185.00 | 257.00 | 222.00 | 203.00 |
| LOC339803 | 280.00 | 221.00 | 254.00 | 251.67 | 320.00 |
| LOC339874 | 9.00 | 10.00 | 9.00 | 9.33 | 13.00 |
| LOC344887 | 78.00 | 123.00 | 110.00 | 103.67 | 98.00 |
| LOC374443 | 296.00 | 286.00 | 364.00 | 315.33 | 341.00 |
| LOC388849 | 244.00 | 308.00 | 286.00 | 279.33 | 258.00 |
| LOC389199 | 3.00 | 1.00 | 3.00 | 2.33 | 2.00 |
| LOC389247 | 10.00 | 11.00 | 8.00 | 9.67 | 5.00 |
| LOC389602 | 67.00 | 52.00 | 74.00 | 64.33 | 71.00 |
| LOC389641 | 32.00 | 60.00 | 47.00 | 46.33 | 40.00 |
| LOC389765 | 50.00 | 50.00 | 61.00 | 53.67 | 52.00 |
| LOC389831 | 134.00 | 124.00 | 124.00 | 127.33 | 168.00 |
| LOC389834 | 421.00 | 396.00 | 462.00 | 426.33 | 368.00 |
| LOC389906 | 136.00 | 119.00 | 132.00 | 129.00 | 158.00 |
| LOC391322 | 5.00 | 11.00 | 12.00 | 9.33 | 8.00 |
| LOC399715 | 26.00 | 23.00 | 40.00 | 29.67 | 32.00 |
| LOC399815 | 318.00 | 315.00 | 300.00 | 311.00 | 418.00 |
| LOC400655 | 595.00 | 568.00 | 707.00 | 623.33 | 518.00 |
| LOC400684 | 48.00 | 60.00 | 40.00 | 49.33 | 71.00 |
| LOC400867 | 3.00 | 0.00 | 0.00 | 1.00 | 0.00 |
| LOC400927 | 12.00 | 13.00 | 13.00 | 12.67 | 5.00 |
| LOC401010 | 7.00 | 5.00 | 14.00 | 8.67 | 9.00 |
| LOC401127 | 15.00 | 9.00 | 11.00 | 11.67 | 7.00 |
| LOC401320 | 353.00 | 329.00 | 344.00 | 342.00 | 316.00 |
| LOC403323 | 8.00 | 3.00 | 6.00 | 5.67 | 4.00 |
| LOC407835 | 9.00 | 9.00 | 6.00 | 8.00 | 13.00 |
| LOC439994 | 39.00 | 43.00 | 74.00 | 52.00 | 42.00 |
| LOC440028 | 11.00 | 6.00 | 22.00 | 13.00 | 10.00 |
| LOC440311 | 22.00 | 23.00 | 23.00 | 22.67 | 21.00 |
| LOC440434 | 22.00 | 27.00 | 22.00 | 23.67 | 17.00 |
| LOC440600 | 7.00 | 9.00 | 8.00 | 8.00 | 9.00 |
| LOC440896 | 11.00 | 6.00 | 16.00 | 11.00 | 17.00 |
| LOC440982 | 3.00 | 2.00 | 3.00 | 2.67 | 8.00 |
| LOC441155 | 25.00 | 33.00 | 32.00 | 30.00 | 47.00 |
| LOC441242 | 127.00 | 147.00 | 146.00 | 140.00 | 144.00 |
| LOC441666 | 53.00 | 46.00 | 46.00 | 48.33 | 36.00 |
| LOC541472 | 8.00 | 6.00 | 3.00 | 5.67 | 12.00 |
| LOC550113 | 4.00 | 0.00 | 1.00 | 1.67 | 2.00 |
| LOC553103 | 5.00 | 5.00 | 15.00 | 8.33 | 6.00 |

| | | | | | |
|---|---|---|---|---|---|
| LOC554206 | 32.00 | 30.00 | 30.00 | 30.67 | 33.00 |
| LOC613037 | 16.00 | 10.00 | 24.00 | 16.67 | 83.00 |
| LOC642361 | 30.00 | 43.00 | 39.00 | 37.33 | 51.00 |
| LOC642366 | 31.00 | 38.00 | 34.00 | 34.33 | 22.00 |
| LOC642423 | 18.00 | 21.00 | 16.00 | 18.33 | 19.00 |
| LOC642846 | 719.00 | 752.00 | 724.00 | 731.67 | 750.00 |
| LOC642852 | 2390.00 | 2318.00 | 2617.00 | 2441.67 | 1967.00 |
| LOC643072 | 16.00 | 15.00 | 17.00 | 16.00 | 25.00 |
| LOC643201 | 12.00 | 2.00 | 7.00 | 7.00 | 3.00 |
| LOC643923 | 3.00 | 0.00 | 4.00 | 2.33 | 4.00 |
| LOC644285 | 31.00 | 40.00 | 38.00 | 36.33 | 18.00 |
| LOC644656 | 91.00 | 68.00 | 65.00 | 74.67 | 106.00 |
| LOC645166 | 72.00 | 88.00 | 84.00 | 81.33 | 79.00 |
| LOC645513 | 105.00 | 105.00 | 105.00 | 105.00 | 97.00 |
| LOC646471 | 6.00 | 2.00 | 1.00 | 3.00 | 0.00 |
| LOC646626 | 24.00 | 30.00 | 31.00 | 28.33 | 52.00 |
| LOC646762 | 414.00 | 366.00 | 416.00 | 398.67 | 417.00 |
| LOC646903 | 45.00 | 64.00 | 50.00 | 53.00 | 80.00 |
| LOC648987 | 255.00 | 237.00 | 259.00 | 250.33 | 250.00 |
| LOC652276 | 119.00 | 88.00 | 105.00 | 104.00 | 77.00 |
| LOC653160 | 97.00 | 105.00 | 95.00 | 99.00 | 121.00 |
| LOC653513 | 3.00 | 12.00 | 17.00 | 10.67 | 25.00 |
| LOC653602 | 72.00 | 65.00 | 54.00 | 63.67 | 90.00 |
| LOC653653 | 63.00 | 34.00 | 64.00 | 53.67 | 66.00 |
| LOC653712 | 20.00 | 23.00 | 24.00 | 22.33 | 19.00 |
| LOC654342 | 140.00 | 144.00 | 144.00 | 142.67 | 151.00 |
| LOC654841 | 35.00 | 31.00 | 33.00 | 33.00 | 27.00 |
| LOC727751 | 3.00 | 1.00 | 4.00 | 2.67 | 2.00 |
| LOC727896 | 111.00 | 92.00 | 104.00 | 102.33 | 145.00 |
| LOC728024 | 11.00 | 13.00 | 15.00 | 13.00 | 18.00 |
| LOC728175 | 4.00 | 3.00 | 8.00 | 5.00 | 1.00 |
| LOC728323 | 32.00 | 29.00 | 33.00 | 31.33 | 28.00 |
| LOC728392 | 466.00 | 450.00 | 464.00 | 460.00 | 445.00 |
| LOC728673 | 30.00 | 29.00 | 33.00 | 30.67 | 21.00 |
| LOC728730 | 124.00 | 125.00 | 121.00 | 123.33 | 118.00 |
| LOC728743 | 91.00 | 93.00 | 121.00 | 101.67 | 119.00 |
| LOC728752 | 19.00 | 21.00 | 31.00 | 23.67 | 28.00 |
| LOC729218 | 937.00 | 810.00 | 936.00 | 894.33 | 537.00 |
| LOC729506 | 7.00 | 6.00 | 8.00 | 7.00 | 9.00 |
| LOC729603 | 157.00 | 134.00 | 173.00 | 154.67 | 132.00 |
| LOC729683 | 33.00 | 24.00 | 23.00 | 26.67 | 18.00 |
| LOC729732 | 3.00 | 1.00 | 1.00 | 1.67 | 2.00 |
| LOC729970 | 116.00 | 98.00 | 100.00 | 104.67 | 103.00 |
| LOC729987 | 126.00 | 122.00 | 143.00 | 130.33 | 173.00 |
| LOC730101 | 250.00 | 284.00 | 288.00 | 274.00 | 292.00 |
| LOC730102 | 476.00 | 440.00 | 436.00 | 450.67 | 573.00 |
| LOC730183 | 18.00 | 21.00 | 8.00 | 15.67 | 22.00 |
| LOC730202 | 23.00 | 37.00 | 27.00 | 29.00 | 40.00 |

| | | | | | |
|---|---|---|---|---|---|
| LOC730668 | 7.00 | 8.00 | 8.00 | 7.67 | 5.00 |
| LOC731157 | 4.00 | 4.00 | 0.00 | 2.67 | 2.00 |
| LOC79160 | 18.00 | 26.00 | 18.00 | 20.67 | 20.00 |
| LOC81691 | 212.00 | 218.00 | 204.00 | 211.33 | 302.00 |
| LOC90246 | 24.00 | 22.00 | 36.00 | 27.33 | 34.00 |
| LOC90768 | 215.00 | 225.00 | 204.00 | 214.67 | 269.00 |
| LOC90784 | 289.00 | 296.00 | 363.00 | 316.00 | 328.00 |
| LOC93622 | 270.00 | 318.00 | 336.00 | 308.00 | 388.00 |
| LOH12CR1 | 121.00 | 95.00 | 132.00 | 116.00 | 122.00 |
| LOH12CR2 | 5.00 | 7.00 | 4.00 | 5.33 | 13.00 |
| LONP1 | 5030.00 | 5227.00 | 5198.00 | 5151.67 | 5453.00 |
| LONP2 | 1645.00 | 1554.00 | 1779.00 | 1659.33 | 1548.00 |
| LONRF1 | 490.00 | 538.00 | 616.00 | 548.00 | 625.00 |
| LONRF2 | 815.00 | 775.00 | 862.00 | 817.33 | 639.00 |
| LOX | 11571.00 | 10686.00 | 10701.00 | 10986.00 | 28674.00 |
| LOXL1 | 3185.00 | 3187.00 | 3111.00 | 3161.00 | 2923.00 |
| LOXL1-AS1 | 608.00 | 562.00 | 619.00 | 596.33 | 715.00 |
| LOXL2 | 14020.00 | 13988.00 | 13937.00 | 13981.67 | 11800.00 |
| LOXL3 | 534.00 | 531.00 | 544.00 | 536.33 | 341.00 |
| LPAL2 | 224.00 | 225.00 | 276.00 | 241.67 | 196.00 |
| LPAR1 | 1201.00 | 1145.00 | 1193.00 | 1179.67 | 1038.00 |
| LPAR2 | 153.00 | 130.00 | 168.00 | 150.33 | 164.00 |
| LPAR3 | 209.00 | 159.00 | 183.00 | 183.67 | 194.00 |
| LPAR6 | 68.00 | 74.00 | 92.00 | 78.00 | 58.00 |
| LPCAT1 | 64.00 | 35.00 | 46.00 | 48.33 | 40.00 |
| LPCAT2 | 422.00 | 361.00 | 426.00 | 403.00 | 361.00 |
| LPCAT3 | 460.00 | 463.00 | 457.00 | 460.00 | 367.00 |
| LPCAT4 | 1408.00 | 1490.00 | 1425.00 | 1441.00 | 1686.00 |
| LPGAT1 | 2110.00 | 1889.00 | 2107.00 | 2035.33 | 2451.00 |
| LPIN1 | 1393.00 | 1309.00 | 1515.00 | 1405.67 | 1566.00 |
| LPIN2 | 4369.00 | 4110.00 | 4418.00 | 4299.00 | 3475.00 |
| LPIN3 | 8.00 | 10.00 | 14.00 | 10.67 | 13.00 |
| LPP | 2744.00 | 2353.00 | 2928.00 | 2675.00 | 2552.00 |
| LPP-AS2 | 37.00 | 35.00 | 41.00 | 37.67 | 55.00 |
| LPPR2 | 635.00 | 617.00 | 652.00 | 634.67 | 625.00 |
| LPPR4 | 6.00 | 4.00 | 6.00 | 5.33 | 1.00 |
| LPPR5 | 4.00 | 2.00 | 5.00 | 3.67 | 12.00 |
| LPXN | 2701.00 | 2670.00 | 2954.00 | 2775.00 | 1399.00 |
| LRBA | 1139.00 | 1030.00 | 1183.00 | 1117.33 | 973.00 |
| LRCH1 | 708.00 | 635.00 | 735.00 | 692.67 | 782.00 |
| LRCH2 | 73.00 | 67.00 | 69.00 | 69.67 | 76.00 |
| LRCH3 | 442.00 | 426.00 | 487.00 | 451.67 | 360.00 |
| LRCH4 | 1277.00 | 1385.00 | 1496.00 | 1386.00 | 1056.00 |
| LRFN1 | 97.00 | 113.00 | 84.00 | 98.00 | 80.00 |
| LRFN3 | 234.00 | 251.00 | 259.00 | 248.00 | 273.00 |
| LRFN4 | 2089.00 | 2098.00 | 2153.00 | 2113.33 | 2545.00 |
| LRFN5 | 214.00 | 234.00 | 237.00 | 228.33 | 364.00 |
| LRGUK | 13.00 | 19.00 | 23.00 | 18.33 | 14.00 |

| | | | | | |
|---|---|---|---|---|---|
| LRIF1 | 662.00 | 699.00 | 769.00 | 710.00 | 879.00 |
| LRIG1 | 823.00 | 746.00 | 825.00 | 798.00 | 646.00 |
| LRIG2 | 667.00 | 695.00 | 730.00 | 697.33 | 616.00 |
| LRIG3 | 246.00 | 201.00 | 270.00 | 239.00 | 282.00 |
| LRIT3 | 12.00 | 15.00 | 11.00 | 12.67 | 17.00 |
| LRP1 | 3723.00 | 3533.00 | 3796.00 | 3684.00 | 2310.00 |
| LRP10 | 5377.00 | 5236.00 | 5320.00 | 5311.00 | 4265.00 |
| LRP11 | 959.00 | 917.00 | 1025.00 | 967.00 | 1175.00 |
| LRP12 | 1502.00 | 1362.00 | 1434.00 | 1432.67 | 1420.00 |
| LRP1-AS | 4.00 | 3.00 | 1.00 | 2.67 | 0.00 |
| LRP1B | 20.00 | 19.00 | 17.00 | 18.67 | 12.00 |
| LRP2BP | 13.00 | 24.00 | 20.00 | 19.00 | 25.00 |
| LRP4 | 480.00 | 484.00 | 523.00 | 495.67 | 390.00 |
| LRP4-AS1 | 7.00 | 5.00 | 6.00 | 6.00 | 8.00 |
| LRP5 | 370.00 | 352.00 | 372.00 | 364.67 | 368.00 |
| LRP5L | 129.00 | 148.00 | 164.00 | 147.00 | 134.00 |
| LRP6 | 1466.00 | 1373.00 | 1416.00 | 1418.33 | 1277.00 |
| LRP8 | 2217.00 | 2206.00 | 2240.00 | 2221.00 | 2138.00 |
| LRPAP1 | 6619.00 | 6892.00 | 6511.00 | 6674.00 | 6681.00 |
| LRPPRC | 4891.00 | 4399.00 | 5327.00 | 4872.33 | 5150.00 |
| LRR1 | 359.00 | 333.00 | 350.00 | 347.33 | 430.00 |
| LRRC1 | 146.00 | 152.00 | 175.00 | 157.67 | 163.00 |
| LRRC14 | 1800.00 | 1857.00 | 1903.00 | 1853.33 | 1862.00 |
| LRRC16A | 158.00 | 173.00 | 174.00 | 168.33 | 204.00 |
| LRRC16B | 10.00 | 7.00 | 10.00 | 9.00 | 15.00 |
| LRRC17 | 1576.00 | 1519.00 | 1693.00 | 1596.00 | 1485.00 |
| LRRC2 | 176.00 | 118.00 | 166.00 | 153.33 | 228.00 |
| LRRC20 | 1045.00 | 1034.00 | 1114.00 | 1064.33 | 1230.00 |
| LRRC23 | 82.00 | 103.00 | 96.00 | 93.67 | 87.00 |
| LRRC24 | 109.00 | 128.00 | 136.00 | 124.33 | 142.00 |
| LRRC27 | 306.00 | 280.00 | 303.00 | 296.33 | 307.00 |
| LRRC28 | 118.00 | 105.00 | 130.00 | 117.67 | 123.00 |
| LRRC29 | 43.00 | 49.00 | 44.00 | 45.33 | 43.00 |
| LRRC2-AS1 | 22.00 | 19.00 | 15.00 | 18.67 | 17.00 |
| LRRC34 | 148.00 | 175.00 | 202.00 | 175.00 | 204.00 |
| LRRC36 | 4.00 | 6.00 | 2.00 | 4.00 | 1.00 |
| LRRC37A3 | 114.00 | 119.00 | 166.00 | 133.00 | 155.00 |
| LRRC37A4P | 30.00 | 24.00 | 18.00 | 24.00 | 25.00 |
| LRRC37B | 99.00 | 86.00 | 107.00 | 97.33 | 125.00 |
| LRRC37BP1 | 294.00 | 272.00 | 324.00 | 296.67 | 279.00 |
| LRRC39 | 25.00 | 30.00 | 23.00 | 26.00 | 25.00 |
| LRRC40 | 530.00 | 498.00 | 611.00 | 546.33 | 667.00 |
| LRRC41 | 2878.00 | 2820.00 | 3092.00 | 2930.00 | 2724.00 |
| LRRC42 | 671.00 | 628.00 | 698.00 | 665.67 | 787.00 |
| LRRC45 | 470.00 | 445.00 | 449.00 | 454.67 | 474.00 |
| LRRC46 | 24.00 | 15.00 | 10.00 | 16.33 | 27.00 |
| LRRC47 | 1238.00 | 1215.00 | 1223.00 | 1225.33 | 1385.00 |
| LRRC48 | 30.00 | 13.00 | 18.00 | 20.33 | 18.00 |

| | | | | | |
|---|---|---|---|---|---|
| LRRC49 | 197.00 | 168.00 | 224.00 | 196.33 | 207.00 |
| LRRC56 | 56.00 | 58.00 | 66.00 | 60.00 | 46.00 |
| LRRC57 | 423.00 | 391.00 | 344.00 | 386.00 | 412.00 |
| LRRC58 | 1604.00 | 1330.00 | 1792.00 | 1575.33 | 1768.00 |
| LRRC59 | 4777.00 | 4454.00 | 4815.00 | 4682.00 | 5408.00 |
| LRRC6 | 71.00 | 65.00 | 67.00 | 67.67 | 57.00 |
| LRRC63 | 24.00 | 22.00 | 26.00 | 24.00 | 22.00 |
| LRRC66 | 12.00 | 10.00 | 14.00 | 12.00 | 15.00 |
| LRRC69 | 81.00 | 79.00 | 64.00 | 74.67 | 44.00 |
| LRRC71 | 256.00 | 260.00 | 297.00 | 271.00 | 238.00 |
| LRRC73 | 3.00 | 3.00 | 4.00 | 3.33 | 10.00 |
| LRRC75A-AS1 | 15065.00 | 15008.00 | 16966.00 | 15679.67 | 19048.00 |
| LRRC75B | 68.00 | 92.00 | 64.00 | 74.67 | 92.00 |
| LRRC8A | 3329.00 | 3434.00 | 3434.00 | 3399.00 | 2522.00 |
| LRRC8B | 369.00 | 309.00 | 411.00 | 363.00 | 402.00 |
| LRRC8C | 1072.00 | 918.00 | 984.00 | 991.33 | 896.00 |
| LRRC8D | 2335.00 | 2139.00 | 2357.00 | 2277.00 | 2340.00 |
| LRRC8E | 624.00 | 633.00 | 629.00 | 628.67 | 579.00 |
| LRRCC1 | 339.00 | 310.00 | 348.00 | 332.33 | 493.00 |
| LRRD1 | 3.00 | 2.00 | 1.00 | 2.00 | 4.00 |
| LRRFIP1 | 2943.00 | 2613.00 | 3122.00 | 2892.67 | 2934.00 |
| LRRFIP2 | 1438.00 | 1351.00 | 1482.00 | 1423.67 | 2080.00 |
| LRRIQ1 | 3.00 | 7.00 | 8.00 | 6.00 | 5.00 |
| LRRIQ3 | 24.00 | 18.00 | 12.00 | 18.00 | 33.00 |
| LRRIQ4 | 7.00 | 4.00 | 3.00 | 4.67 | 11.00 |
| LRRK1 | 1281.00 | 1281.00 | 1450.00 | 1337.33 | 1064.00 |
| LRRK2 | 117.00 | 89.00 | 115.00 | 107.00 | 125.00 |
| LRRTM2 | 7.00 | 11.00 | 7.00 | 8.33 | 10.00 |
| LRRTM3 | 5.00 | 0.00 | 0.00 | 1.67 | 0.00 |
| LRSAM1 | 949.00 | 960.00 | 1018.00 | 975.67 | 941.00 |
| LRTM2 | 5.00 | 6.00 | 4.00 | 5.00 | 3.00 |
| LRTOMT | 113.00 | 94.00 | 88.00 | 98.33 | 143.00 |
| LRWD1 | 736.00 | 676.00 | 731.00 | 714.33 | 826.00 |
| LSAMP | 4.00 | 4.00 | 7.00 | 5.00 | 4.00 |
| LSG1 | 2233.00 | 2207.00 | 2380.00 | 2273.33 | 2233.00 |
| LSM1 | 515.00 | 485.00 | 502.00 | 500.67 | 715.00 |
| LSM10 | 436.00 | 477.00 | 490.00 | 467.67 | 620.00 |
| LSM11 | 387.00 | 385.00 | 411.00 | 394.33 | 292.00 |
| LSM12 | 252.00 | 201.00 | 210.00 | 221.00 | 297.00 |
| LSM14A | 27.00 | 33.00 | 25.00 | 28.33 | 27.00 |
| LSM14B | 1661.00 | 1580.00 | 1657.00 | 1632.67 | 1753.00 |
| LSM3 | 374.00 | 365.00 | 379.00 | 372.67 | 514.00 |
| LSM4 | 1884.00 | 1975.00 | 2089.00 | 1982.67 | 2530.00 |
| LSM5 | 969.00 | 983.00 | 1134.00 | 1028.67 | 1535.00 |
| LSM6 | 251.00 | 209.00 | 243.00 | 234.33 | 309.00 |
| LSM7 | 925.00 | 962.00 | 1024.00 | 970.33 | 1228.00 |
| LSM8 | 1289.00 | 1402.00 | 1537.00 | 1409.33 | 1555.00 |
| LSMEM1 | 133.00 | 148.00 | 158.00 | 146.33 | 128.00 |

| | | | | | |
|---|---|---|---|---|---|
| LSR | 18.00 | 25.00 | 21.00 | 21.33 | 31.00 |
| LSS | 143.00 | 127.00 | 138.00 | 136.00 | 101.00 |
| LTA4H | 2125.00 | 1871.00 | 2241.00 | 2079.00 | 2286.00 |
| LTB4R | 2761.00 | 2910.00 | 3122.00 | 2931.00 | 2702.00 |
| LTB4R2 | 677.00 | 649.00 | 837.00 | 721.00 | 441.00 |
| LTBP1 | 305.00 | 284.00 | 339.00 | 309.33 | 210.00 |
| LTBP2 | 7456.00 | 7175.00 | 7967.00 | 7532.67 | 6756.00 |
| LTBP3 | 6706.00 | 6742.00 | 7139.00 | 6862.33 | 4015.00 |
| LTBP4 | 4465.00 | 4493.00 | 4732.00 | 4563.33 | 3682.00 |
| LTBR | 3201.00 | 3513.00 | 3449.00 | 3387.67 | 3279.00 |
| LTK | 7.00 | 16.00 | 15.00 | 12.67 | 8.00 |
| LTN1 | 1575.00 | 1530.00 | 1737.00 | 1614.00 | 1526.00 |
| LTV1 | 1473.00 | 1470.00 | 1602.00 | 1515.00 | 1826.00 |
| LUC7L | 832.00 | 972.00 | 909.00 | 904.33 | 991.00 |
| LUC7L2 | 197.00 | 182.00 | 229.00 | 202.67 | 178.00 |
| LUC7L3 | 10974.00 | 11235.00 | 11942.00 | 11383.67 | 11489.00 |
| LUCAT1 | 34.00 | 35.00 | 40.00 | 36.33 | 17.00 |
| LURAP1 | 28.00 | 32.00 | 37.00 | 32.33 | 33.00 |
| LURAP1L | 315.00 | 339.00 | 366.00 | 340.00 | 465.00 |
| LURAP1L-AS1 | 442.00 | 466.00 | 477.00 | 461.67 | 352.00 |
| LUZP1 | 2109.00 | 1870.00 | 2221.00 | 2066.67 | 1391.00 |
| LUZP2 | 6.00 | 3.00 | 3.00 | 4.00 | 3.00 |
| LVCAT1 | 34.00 | 46.00 | 37.00 | 39.00 | 53.00 |
| LVCAT5 | 6.00 | 7.00 | 9.00 | 7.33 | 6.00 |
| LVRN | 8.00 | 3.00 | 22.00 | 11.00 | 7.00 |
| LXN | 310.00 | 320.00 | 318.00 | 316.00 | 502.00 |
| LY6K | 4189.00 | 4333.00 | 4018.00 | 4180.00 | 4031.00 |
| LY96 | 664.00 | 674.00 | 577.00 | 638.33 | 857.00 |
| LYAR | 692.00 | 680.00 | 710.00 | 694.00 | 919.00 |
| LYG1 | 26.00 | 43.00 | 47.00 | 38.67 | 41.00 |
| LYL1 | 26.00 | 22.00 | 17.00 | 21.67 | 36.00 |
| LYN | 104.00 | 106.00 | 105.00 | 105.00 | 117.00 |
| LYPD1 | 86.00 | 67.00 | 70.00 | 74.33 | 37.00 |
| LYPD6 | 67.00 | 68.00 | 72.00 | 69.00 | 105.00 |
| LYPLA1 | 916.00 | 714.00 | 1000.00 | 876.67 | 1150.00 |
| LYPLA2 | 968.00 | 878.00 | 964.00 | 936.67 | 1114.00 |
| LYPLAL1 | 535.00 | 561.00 | 619.00 | 571.67 | 633.00 |
| LYPLAL1-AS1 | 355.00 | 341.00 | 400.00 | 365.33 | 212.00 |
| LYRM1 | 428.00 | 382.00 | 456.00 | 422.00 | 562.00 |
| LYRM2 | 812.00 | 749.00 | 857.00 | 806.00 | 998.00 |
| LYRM4 | 515.00 | 492.00 | 559.00 | 522.00 | 647.00 |
| LYRM5 | 233.00 | 212.00 | 251.00 | 232.00 | 263.00 |
| LYRM7 | 937.00 | 822.00 | 999.00 | 919.33 | 962.00 |
| LYRM9 | 14.00 | 25.00 | 14.00 | 17.67 | 26.00 |
| LYSMD1 | 283.00 | 286.00 | 379.00 | 316.00 | 314.00 |
| LYSMD2 | 245.00 | 180.00 | 237.00 | 220.67 | 285.00 |
| LYSMD3 | 767.00 | 645.00 | 747.00 | 719.67 | 662.00 |
| LYSMD4 | 264.00 | 282.00 | 300.00 | 282.00 | 261.00 |

| | | | | | |
|---|---|---|---|---|---|
| LYST | 610.00 | 616.00 | 782.00 | 669.33 | 490.00 |
| LYZ | 5.00 | 6.00 | 4.00 | 5.00 | 7.00 |
| LZIC | 321.00 | 321.00 | 370.00 | 337.33 | 455.00 |
| LZTFL1 | 458.00 | 412.00 | 479.00 | 449.67 | 545.00 |
| LZTR1 | 2358.00 | 2381.00 | 2525.00 | 2421.33 | 2322.00 |
| LZTS2 | 1824.00 | 1903.00 | 2035.00 | 1920.67 | 1592.00 |
| LZTS3 | 255.00 | 259.00 | 285.00 | 266.33 | 269.00 |
| M6PR | 4275.00 | 4325.00 | 4337.00 | 4312.33 | 4145.00 |
| MAATS1 | 3.00 | 3.00 | 1.00 | 2.33 | 6.00 |
| MAB21L1 | 7.00 | 4.00 | 16.00 | 9.00 | 14.00 |
| MAB21L3 | 5.00 | 10.00 | 2.00 | 5.67 | 3.00 |
| MACF1 | 20133.00 | 18545.00 | 20739.00 | 19805.67 | 19095.00 |
| MACROD1 | 1177.00 | 1201.00 | 1374.00 | 1250.67 | 1376.00 |
| MACROD2 | 22.00 | 16.00 | 20.00 | 19.33 | 16.00 |
| MAD1L1 | 1025.00 | 1033.00 | 1083.00 | 1047.00 | 1289.00 |
| MAD2L1 | 478.00 | 478.00 | 495.00 | 483.67 | 809.00 |
| MAD2L1BP | 584.00 | 500.00 | 538.00 | 540.67 | 648.00 |
| MAD2L2 | 521.00 | 531.00 | 554.00 | 535.33 | 649.00 |
| MADD | 1565.00 | 1502.00 | 1621.00 | 1562.67 | 1231.00 |
| MAEA | 700.00 | 717.00 | 740.00 | 719.00 | 849.00 |
| MAF1 | 2602.00 | 2845.00 | 2893.00 | 2780.00 | 3236.00 |
| MAFB | 39.00 | 35.00 | 39.00 | 37.67 | 19.00 |
| MAFF | 1635.00 | 1561.00 | 1585.00 | 1593.67 | 2072.00 |
| MAFG | 3752.00 | 3692.00 | 3981.00 | 3808.33 | 3468.00 |
| MAFG-AS1 | 122.00 | 97.00 | 87.00 | 102.00 | 136.00 |
| MAFIP | 351.00 | 382.00 | 458.00 | 397.00 | 386.00 |
| MAFK | 1340.00 | 1377.00 | 1562.00 | 1426.33 | 1468.00 |
| MAGEA1 | 7.00 | 7.00 | 8.00 | 7.33 | 13.00 |
| MAGEA10 | 42.00 | 36.00 | 44.00 | 40.67 | 51.00 |
| MAGEA11 | 46.00 | 36.00 | 65.00 | 49.00 | 56.00 |
| MAGEA12 | 109.00 | 104.00 | 104.00 | 105.67 | 128.00 |
| MAGEA3 | 359.00 | 388.00 | 395.00 | 380.67 | 435.00 |
| MAGEA4 | 4.00 | 7.00 | 6.00 | 5.67 | 1.00 |
| MAGEA6 | 69.00 | 78.00 | 85.00 | 77.33 | 112.00 |
| MAGEB17 | 18.00 | 24.00 | 19.00 | 20.33 | 26.00 |
| MAGEB2 | 27.00 | 24.00 | 38.00 | 29.67 | 21.00 |
| MAGEB3 | 3.00 | 7.00 | 10.00 | 6.67 | 5.00 |
| MAGEC1 | 167.00 | 167.00 | 171.00 | 168.33 | 176.00 |
| MAGEC2 | 25.00 | 17.00 | 12.00 | 18.00 | 26.00 |
| MAGEC3 | 9.00 | 13.00 | 14.00 | 12.00 | 6.00 |
| MAGED1 | 2555.00 | 2352.00 | 2652.00 | 2519.67 | 2593.00 |
| MAGED2 | 3867.00 | 3828.00 | 4063.00 | 3919.33 | 3839.00 |
| MAGEE1 | 38.00 | 47.00 | 53.00 | 46.00 | 60.00 |
| MAGEF1 | 867.00 | 859.00 | 844.00 | 856.67 | 1095.00 |
| MAGEH1 | 59.00 | 89.00 | 67.00 | 71.67 | 83.00 |
| MAGI1 | 1761.00 | 1703.00 | 1988.00 | 1817.33 | 1651.00 |
| MAGI2 | 162.00 | 183.00 | 188.00 | 177.67 | 182.00 |
| MAGI2-AS3 | 1539.00 | 1432.00 | 1634.00 | 1535.00 | 1292.00 |

| | | | | | |
|---|---|---|---|---|---|
| MAGI3 | 384.00 | 346.00 | 404.00 | 378.00 | 424.00 |
| MAGIX | 17.00 | 8.00 | 12.00 | 12.33 | 4.00 |
| MAGOH | 747.00 | 790.00 | 773.00 | 770.00 | 1141.00 |
| MAGOHB | 516.00 | 504.00 | 575.00 | 531.67 | 920.00 |
| MAGT1 | 2846.00 | 2664.00 | 2817.00 | 2775.67 | 2474.00 |
| MAK | 3.00 | 0.00 | 1.00 | 1.33 | 10.00 |
| MAK16 | 763.00 | 665.00 | 785.00 | 737.67 | 885.00 |
| MALAT1 | 5984.00 | 6138.00 | 6286.00 | 6136.00 | 4351.00 |
| MALRD1 | 6.00 | 6.00 | 5.00 | 5.67 | 3.00 |
| MALSU1 | 422.00 | 444.00 | 504.00 | 456.67 | 501.00 |
| MALT1 | 734.00 | 672.00 | 804.00 | 736.67 | 564.00 |
| MAMDC4 | 435.00 | 457.00 | 490.00 | 460.67 | 523.00 |
| MAML1 | 1237.00 | 1236.00 | 1238.00 | 1237.00 | 1179.00 |
| MAML2 | 1480.00 | 1365.00 | 1572.00 | 1472.33 | 1289.00 |
| MAML3 | 504.00 | 398.00 | 490.00 | 464.00 | 339.00 |
| MAMLD1 | 217.00 | 203.00 | 197.00 | 205.67 | 148.00 |
| MAMSTR | 30.00 | 16.00 | 22.00 | 22.67 | 27.00 |
| MAN1A1 | 768.00 | 718.00 | 797.00 | 761.00 | 719.00 |
| MAN1A2 | 940.00 | 836.00 | 969.00 | 915.00 | 717.00 |
| MAN1B1 | 3947.00 | 3986.00 | 3905.00 | 3946.00 | 3623.00 |
| MAN1B1-AS1 | 39.00 | 26.00 | 36.00 | 33.67 | 45.00 |
| MAN1C1 | 4.00 | 0.00 | 1.00 | 1.67 | 0.00 |
| MAN2A1 | 4940.00 | 4412.00 | 5225.00 | 4859.00 | 3751.00 |
| MAN2A2 | 1465.00 | 1518.00 | 1499.00 | 1494.00 | 1182.00 |
| MAN2B1 | 1793.00 | 1848.00 | 1928.00 | 1856.33 | 1705.00 |
| MAN2B2 | 700.00 | 703.00 | 646.00 | 683.00 | 603.00 |
| MAN2C1 | 3105.00 | 3064.00 | 3372.00 | 3180.33 | 2761.00 |
| MANBA | 640.00 | 724.00 | 705.00 | 689.67 | 684.00 |
| MANBAL | 1114.00 | 1042.00 | 1214.00 | 1123.33 | 1287.00 |
| MANEA | 882.00 | 793.00 | 840.00 | 838.33 | 924.00 |
| MANEA-AS1 | 31.00 | 49.00 | 62.00 | 47.33 | 45.00 |
| MANEAL | 69.00 | 77.00 | 84.00 | 76.67 | 72.00 |
| MANF | 1669.00 | 1722.00 | 1580.00 | 1657.00 | 1848.00 |
| MANSC1 | 43.00 | 45.00 | 48.00 | 45.33 | 45.00 |
| MANSC4 | 5.00 | 1.00 | 2.00 | 2.67 | 9.00 |
| MAP1A | 52.00 | 57.00 | 66.00 | 58.33 | 66.00 |
| MAP1B | 3469.00 | 2869.00 | 3539.00 | 3292.33 | 2176.00 |
| MAP1LC3A | 23.00 | 29.00 | 28.00 | 26.67 | 55.00 |
| MAP1LC3B | 2099.00 | 2046.00 | 2096.00 | 2080.33 | 2735.00 |
| MAP1LC3B2 | 61.00 | 47.00 | 70.00 | 59.33 | 91.00 |
| MAP1S | 939.00 | 993.00 | 945.00 | 959.00 | 883.00 |
| MAP2 | 180.00 | 150.00 | 203.00 | 177.67 | 131.00 |
| MAP2K1 | 1453.00 | 1514.00 | 1609.00 | 1525.33 | 1704.00 |
| MAP2K2 | 2407.00 | 2472.00 | 2681.00 | 2520.00 | 2949.00 |
| MAP2K3 | 1308.00 | 1237.00 | 1315.00 | 1286.67 | 1427.00 |
| MAP2K4 | 775.00 | 779.00 | 870.00 | 808.00 | 734.00 |
| MAP2K5 | 453.00 | 453.00 | 531.00 | 479.00 | 490.00 |
| MAP2K7 | 1145.00 | 1218.00 | 1239.00 | 1200.67 | 1312.00 |

| | | | | | |
|---|---|---|---|---|---|
| MAP3K1 | 259.00 | 254.00 | 284.00 | 265.67 | 159.00 |
| MAP3K10 | 545.00 | 554.00 | 595.00 | 564.67 | 595.00 |
| MAP3K11 | 1842.00 | 1806.00 | 1854.00 | 1834.00 | 1736.00 |
| MAP3K12 | 947.00 | 1003.00 | 1003.00 | 984.33 | 951.00 |
| MAP3K13 | 469.00 | 443.00 | 484.00 | 465.33 | 563.00 |
| MAP3K14 | 1477.00 | 1475.00 | 1517.00 | 1489.67 | 1594.00 |
| MAP3K2 | 1137.00 | 1105.00 | 1306.00 | 1182.67 | 931.00 |
| MAP3K3 | 1253.00 | 1271.00 | 1306.00 | 1276.67 | 1154.00 |
| MAP3K4 | 862.00 | 802.00 | 969.00 | 877.67 | 759.00 |
| MAP3K5 | 177.00 | 151.00 | 177.00 | 168.33 | 217.00 |
| MAP3K6 | 389.00 | 458.00 | 434.00 | 427.00 | 590.00 |
| MAP3K7 | 1648.00 | 1659.00 | 1758.00 | 1688.33 | 1774.00 |
| MAP3K7CL | 294.00 | 267.00 | 309.00 | 290.00 | 524.00 |
| MAP3K8 | 15.00 | 19.00 | 31.00 | 21.67 | 21.00 |
| MAP3K9 | 165.00 | 165.00 | 172.00 | 167.33 | 114.00 |
| MAP4 | 8760.00 | 8432.00 | 8996.00 | 8729.33 | 6919.00 |
| MAP4K1 | 18.00 | 7.00 | 19.00 | 14.67 | 22.00 |
| MAP4K2 | 1078.00 | 1028.00 | 1091.00 | 1065.67 | 1211.00 |
| MAP4K3 | 1376.00 | 1195.00 | 1576.00 | 1382.33 | 1309.00 |
| MAP4K4 | 9164.00 | 8650.00 | 9459.00 | 9091.00 | 9760.00 |
| MAP4K5 | 1817.00 | 1764.00 | 2060.00 | 1880.33 | 1991.00 |
| MAP6 | 6.00 | 2.00 | 3.00 | 3.67 | 0.00 |
| MAP6D1 | 80.00 | 73.00 | 98.00 | 83.67 | 106.00 |
| MAP7 | 4.00 | 8.00 | 13.00 | 8.33 | 12.00 |
| MAP7D1 | 2003.00 | 2026.00 | 1993.00 | 2007.33 | 1809.00 |
| MAP7D2 | 3.00 | 4.00 | 4.00 | 3.67 | 9.00 |
| MAP7D3 | 2145.00 | 1923.00 | 2132.00 | 2066.67 | 1730.00 |
| MAP9 | 529.00 | 503.00 | 589.00 | 540.33 | 541.00 |
| MAPK1 | 2358.00 | 2216.00 | 2517.00 | 2363.67 | 2597.00 |
| MAPK10 | 7.00 | 6.00 | 9.00 | 7.33 | 4.00 |
| MAPK11 | 309.00 | 337.00 | 357.00 | 334.33 | 362.00 |
| MAPK12 | 1028.00 | 1051.00 | 1076.00 | 1051.67 | 987.00 |
| MAPK13 | 1214.00 | 1279.00 | 1477.00 | 1323.33 | 1267.00 |
| MAPK14 | 1671.00 | 1437.00 | 1624.00 | 1577.33 | 1692.00 |
| MAPK1IP1L | 1940.00 | 1824.00 | 2094.00 | 1952.67 | 2286.00 |
| MAPK3 | 678.00 | 664.00 | 753.00 | 698.33 | 812.00 |
| MAPK6 | 1232.00 | 1107.00 | 1455.00 | 1264.67 | 1467.00 |
| MAPK7 | 313.00 | 336.00 | 302.00 | 317.00 | 330.00 |
| MAPK8 | 1151.00 | 1102.00 | 1226.00 | 1159.67 | 1266.00 |
| MAPK8IP1 | 227.00 | 197.00 | 224.00 | 216.00 | 224.00 |
| MAPK8IP2 | 17.00 | 15.00 | 11.00 | 14.33 | 20.00 |
| MAPK8IP3 | 3863.00 | 4152.00 | 4404.00 | 4139.67 | 3651.00 |
| MAPK9 | 775.00 | 796.00 | 819.00 | 796.67 | 922.00 |
| MAPKAP1 | 2489.00 | 2262.00 | 2566.00 | 2439.00 | 3298.00 |
| MAPKAPK2 | 2105.00 | 2066.00 | 2331.00 | 2167.33 | 2376.00 |
| MAPKAPK3 | 1228.00 | 1216.00 | 1285.00 | 1243.00 | 906.00 |
| MAPKAPK5 | 692.00 | 677.00 | 740.00 | 703.00 | 803.00 |
| MAPKAPK5-AS1 | 449.00 | 444.00 | 503.00 | 465.33 | 501.00 |

| | | | | | |
|---|---|---|---|---|---|
| MAPKBP1 | 1485.00 | 1309.00 | 1492.00 | 1428.67 | 1312.00 |
| MAPRE1 | 2655.00 | 2306.00 | 2636.00 | 2532.33 | 3401.00 |
| MAPRE2 | 415.00 | 381.00 | 434.00 | 410.00 | 413.00 |
| MAPRE3 | 179.00 | 147.00 | 172.00 | 166.00 | 192.00 |
| MAPT | 16.00 | 15.00 | 21.00 | 17.33 | 14.00 |
| MAR11. | 11.00 | 4.00 | 2.00 | 5.67 | 11.00 |
| MAR3. | 55.00 | 39.00 | 50.00 | 48.00 | 74.00 |
| MAR4. | 2731.00 | 2793.00 | 2907.00 | 2810.33 | 4283.00 |
| MAR5. | 949.00 | 913.00 | 1096.00 | 986.00 | 1129.00 |
| MAR6. | 5533.00 | 5314.00 | 6055.00 | 5634.00 | 5471.00 |
| MAR7. | 2184.00 | 2036.00 | 2465.00 | 2228.33 | 2179.00 |
| MAR8. | 59.00 | 64.00 | 69.00 | 64.00 | 45.00 |
| MAR9. | 608.00 | 547.00 | 637.00 | 597.33 | 595.00 |
| MARCKS | 2504.00 | 2466.00 | 2577.00 | 2515.67 | 2620.00 |
| MARCKSL1 | 1227.00 | 1126.00 | 1219.00 | 1190.67 | 1897.00 |
| MARK1 | 293.00 | 306.00 | 354.00 | 317.67 | 362.00 |
| MARK2 | 1377.00 | 1366.00 | 1487.00 | 1410.00 | 1525.00 |
| MARK3 | 1993.00 | 1989.00 | 2105.00 | 2029.00 | 1954.00 |
| MARK4 | 579.00 | 598.00 | 675.00 | 617.33 | 548.00 |
| MARS | 11869.00 | 11211.00 | 11931.00 | 11670.33 | 12901.00 |
| MARS2 | 845.00 | 815.00 | 916.00 | 858.67 | 860.00 |
| MARVELD1 | 2377.00 | 2358.00 | 2461.00 | 2398.67 | 2154.00 |
| MASP1 | 23.00 | 23.00 | 30.00 | 25.33 | 9.00 |
| MASP2 | 5.00 | 3.00 | 5.00 | 4.33 | 10.00 |
| MAST2 | 3797.00 | 3716.00 | 4058.00 | 3857.00 | 3072.00 |
| MAST3 | 260.00 | 239.00 | 268.00 | 255.67 | 219.00 |
| MAST4 | 601.00 | 508.00 | 614.00 | 574.33 | 628.00 |
| MASTL | 533.00 | 526.00 | 549.00 | 536.00 | 648.00 |
| MAT2A | 13774.00 | 13131.00 | 14376.00 | 13760.33 | 12771.00 |
| MAT2B | 1240.00 | 1198.00 | 1297.00 | 1245.00 | 1545.00 |
| MATN1 | 5.00 | 3.00 | 8.00 | 5.33 | 4.00 |
| MATN1-AS1 | 15.00 | 11.00 | 15.00 | 13.67 | 11.00 |
| MATN2 | 153.00 | 190.00 | 177.00 | 173.33 | 80.00 |
| MATN3 | 50.00 | 48.00 | 57.00 | 51.67 | 71.00 |
| MATN4 | 4.00 | 7.00 | 4.00 | 5.00 | 3.00 |
| MATR3 | 7154.00 | 6377.00 | 7530.00 | 7020.33 | 8106.00 |
| MAU2 | 1523.00 | 1457.00 | 1540.00 | 1506.67 | 1458.00 |
| MAVS | 4094.00 | 3783.00 | 4147.00 | 4008.00 | 2874.00 |
| MAX | 1273.00 | 1178.00 | 1254.00 | 1235.00 | 1304.00 |
| MAZ | 1955.00 | 2090.00 | 2340.00 | 2128.33 | 2664.00 |
| MB21D2 | 388.00 | 315.00 | 321.00 | 341.33 | 314.00 |
| MBD1 | 2260.00 | 2261.00 | 2328.00 | 2283.00 | 2131.00 |
| MBD2 | 675.00 | 771.00 | 810.00 | 752.00 | 885.00 |
| MBD3 | 2866.00 | 2945.00 | 3198.00 | 3003.00 | 3459.00 |
| MBD4 | 819.00 | 771.00 | 816.00 | 802.00 | 852.00 |
| MBD5 | 251.00 | 214.00 | 294.00 | 253.00 | 241.00 |
| MBD6 | 1219.00 | 1210.00 | 1395.00 | 1274.67 | 1241.00 |
| MBIP | 247.00 | 241.00 | 275.00 | 254.33 | 309.00 |

| | | | | | |
|---|---|---|---|---|---|
| MBL1P | 182.00 | 167.00 | 198.00 | 182.33 | 174.00 |
| MBLAC1 | 11.00 | 26.00 | 24.00 | 20.33 | 28.00 |
| MBLAC2 | 435.00 | 406.00 | 467.00 | 436.00 | 472.00 |
| MBNL1 | 3846.00 | 3478.00 | 4305.00 | 3876.33 | 3707.00 |
| MBNL1-AS1 | 49.00 | 36.00 | 46.00 | 43.67 | 48.00 |
| MBNL2 | 515.00 | 393.00 | 482.00 | 463.33 | 364.00 |
| MBNL3 | 210.00 | 187.00 | 204.00 | 200.33 | 254.00 |
| MBOAT1 | 238.00 | 257.00 | 225.00 | 240.00 | 256.00 |
| MBOAT2 | 426.00 | 365.00 | 399.00 | 396.67 | 412.00 |
| MBOAT7 | 39.00 | 43.00 | 33.00 | 38.33 | 24.00 |
| MBP | 908.00 | 865.00 | 905.00 | 892.67 | 650.00 |
| MBTD1 | 1053.00 | 975.00 | 1120.00 | 1049.33 | 1067.00 |
| MBTPS1 | 7117.00 | 6833.00 | 6950.00 | 6966.67 | 6408.00 |
| MBTPS2 | 895.00 | 810.00 | 836.00 | 847.00 | 917.00 |
| MC1R | 1530.00 | 1647.00 | 1725.00 | 1634.00 | 1802.00 |
| MC4R | 3.00 | 0.00 | 0.00 | 1.00 | 4.00 |
| MCAM | 97.00 | 99.00 | 105.00 | 100.33 | 33.00 |
| MCAT | 217.00 | 212.00 | 237.00 | 222.00 | 296.00 |
| MCC | 672.00 | 554.00 | 644.00 | 623.33 | 698.00 |
| MCCC1 | 771.00 | 772.00 | 781.00 | 774.67 | 797.00 |
| MCCC2 | 1344.00 | 1209.00 | 1354.00 | 1302.33 | 1467.00 |
| MCEE | 121.00 | 119.00 | 112.00 | 117.33 | 125.00 |
| MCF2L | 13.00 | 10.00 | 11.00 | 11.33 | 4.00 |
| MCF2L2 | 28.00 | 36.00 | 40.00 | 34.67 | 45.00 |
| MCFD2 | 16094.00 | 14551.00 | 15316.00 | 15320.33 | 15302.00 |
| MCIDAS | 175.00 | 161.00 | 229.00 | 188.33 | 221.00 |
| MCL1 | 14729.00 | 14120.00 | 15201.00 | 14683.33 | 17909.00 |
| MCM10 | 324.00 | 277.00 | 324.00 | 308.33 | 433.00 |
| MCM2 | 1736.00 | 1800.00 | 1770.00 | 1768.67 | 2059.00 |
| MCM3 | 2730.00 | 2643.00 | 2790.00 | 2721.00 | 3355.00 |
| MCM3AP | 3567.00 | 3535.00 | 3614.00 | 3572.00 | 3331.00 |
| MCM3AP-AS1 | 207.00 | 192.00 | 213.00 | 204.00 | 173.00 |
| MCM4 | 2342.00 | 2166.00 | 2326.00 | 2278.00 | 2227.00 |
| MCM5 | 1286.00 | 1289.00 | 1189.00 | 1254.67 | 1192.00 |
| MCM6 | 1408.00 | 1296.00 | 1414.00 | 1372.67 | 1671.00 |
| MCM7 | 4153.00 | 4153.00 | 4112.00 | 4139.33 | 6020.00 |
| MCM8 | 1437.00 | 1471.00 | 1511.00 | 1473.00 | 1783.00 |
| MCM9 | 278.00 | 324.00 | 333.00 | 311.67 | 351.00 |
| MCMBP | 1582.00 | 1395.00 | 1638.00 | 1538.33 | 1922.00 |
| MCMDC2 | 22.00 | 27.00 | 25.00 | 24.67 | 25.00 |
| MCOLN1 | 783.00 | 680.00 | 675.00 | 712.67 | 649.00 |
| MCOLN2 | 776.00 | 770.00 | 827.00 | 791.00 | 926.00 |
| MCOLN3 | 245.00 | 254.00 | 262.00 | 253.67 | 214.00 |
| MCPH1 | 345.00 | 334.00 | 360.00 | 346.33 | 357.00 |
| MCPH1-AS1 | 26.00 | 12.00 | 24.00 | 20.67 | 32.00 |
| MCRS1 | 1248.00 | 1280.00 | 1344.00 | 1290.67 | 1571.00 |
| MCTP1 | 664.00 | 643.00 | 730.00 | 679.00 | 828.00 |
| MCTS1 | 1148.00 | 1069.00 | 1191.00 | 1136.00 | 1281.00 |

| | | | | | |
|---|---|---|---|---|---|
| MCTS2P | 214.00 | 240.00 | 229.00 | 227.67 | 189.00 |
| MCU | 686.00 | 611.00 | 734.00 | 677.00 | 702.00 |
| MCUR1 | 745.00 | 676.00 | 801.00 | 740.67 | 847.00 |
| MDC1 | 3.00 | 4.00 | 1.00 | 2.67 | 1.00 |
| MDFIC | 700.00 | 621.00 | 832.00 | 717.67 | 809.00 |
| MDGA1 | 72.00 | 60.00 | 83.00 | 71.67 | 30.00 |
| MDGA2 | 116.00 | 99.00 | 133.00 | 116.00 | 102.00 |
| MDH1 | 2624.00 | 2432.00 | 2683.00 | 2579.67 | 3565.00 |
| MDH1B | 20.00 | 26.00 | 27.00 | 24.33 | 39.00 |
| MDH2 | 3421.00 | 3300.00 | 3570.00 | 3430.33 | 4650.00 |
| MDK | 1491.00 | 1703.00 | 1543.00 | 1579.00 | 1905.00 |
| MDM1 | 332.00 | 323.00 | 358.00 | 337.67 | 390.00 |
| MDM2 | 1127.00 | 993.00 | 1133.00 | 1084.33 | 1239.00 |
| MDM4 | 2031.00 | 2058.00 | 2196.00 | 2095.00 | 2319.00 |
| MDN1 | 4019.00 | 3650.00 | 4433.00 | 4034.00 | 3503.00 |
| MDP1 | 52.00 | 42.00 | 42.00 | 45.33 | 39.00 |
| ME1 | 731.00 | 682.00 | 760.00 | 724.33 | 935.00 |
| ME2 | 1199.00 | 1109.00 | 1240.00 | 1182.67 | 1482.00 |
| ME3 | 67.00 | 56.00 | 98.00 | 73.67 | 47.00 |
| MEA1 | 939.00 | 997.00 | 1003.00 | 979.67 | 1314.00 |
| MEAF6 | 1212.00 | 1130.00 | 1349.00 | 1230.33 | 1706.00 |
| MECOM | 1079.00 | 838.00 | 1098.00 | 1005.00 | 1020.00 |
| MECP2 | 1408.00 | 1350.00 | 1530.00 | 1429.33 | 1153.00 |
| MECR | 451.00 | 441.00 | 492.00 | 461.33 | 498.00 |
| MED1 | 2623.00 | 2333.00 | 2864.00 | 2606.67 | 2186.00 |
| MED10 | 570.00 | 560.00 | 620.00 | 583.33 | 769.00 |
| MED11 | 229.00 | 209.00 | 204.00 | 214.00 | 282.00 |
| MED12 | 2019.00 | 1868.00 | 2011.00 | 1966.00 | 1599.00 |
| MED12L | 94.00 | 109.00 | 104.00 | 102.33 | 74.00 |
| MED13 | 2217.00 | 1938.00 | 2492.00 | 2215.67 | 1953.00 |
| MED13L | 3008.00 | 2636.00 | 3293.00 | 2979.00 | 2553.00 |
| MED14 | 1477.00 | 1412.00 | 1588.00 | 1492.33 | 1505.00 |
| MED14OS | 6.00 | 0.00 | 2.00 | 2.67 | 0.00 |
| MED15 | 1916.00 | 1866.00 | 2018.00 | 1933.33 | 2015.00 |
| MED16 | 651.00 | 661.00 | 649.00 | 653.67 | 631.00 |
| MED17 | 875.00 | 866.00 | 905.00 | 882.00 | 968.00 |
| MED18 | 310.00 | 276.00 | 323.00 | 303.00 | 381.00 |
| MED19 | 336.00 | 326.00 | 354.00 | 338.67 | 409.00 |
| MED20 | 616.00 | 591.00 | 658.00 | 621.67 | 725.00 |
| MED21 | 603.00 | 524.00 | 577.00 | 568.00 | 645.00 |
| MED22 | 1430.00 | 1367.00 | 1442.00 | 1413.00 | 1592.00 |
| MED23 | 1473.00 | 1411.00 | 1516.00 | 1466.67 | 1353.00 |
| MED24 | 2293.00 | 2235.00 | 2310.00 | 2279.33 | 2245.00 |
| MED25 | 609.00 | 568.00 | 600.00 | 592.33 | 588.00 |
| MED26 | 254.00 | 220.00 | 221.00 | 231.67 | 239.00 |
| MED27 | 477.00 | 459.00 | 497.00 | 477.67 | 607.00 |
| MED28 | 480.00 | 458.00 | 530.00 | 489.33 | 540.00 |
| MED29 | 530.00 | 554.00 | 633.00 | 572.33 | 596.00 |

| | | | | | |
|---|---|---|---|---|---|
| MED30 | 269.00 | 250.00 | 277.00 | 265.33 | 400.00 |
| MED31 | 148.00 | 159.00 | 150.00 | 152.33 | 181.00 |
| MED4 | 435.00 | 401.00 | 400.00 | 412.00 | 541.00 |
| MED6 | 468.00 | 400.00 | 453.00 | 440.33 | 528.00 |
| MED7 | 264.00 | 237.00 | 309.00 | 270.00 | 286.00 |
| MED8 | 1433.00 | 1387.00 | 1521.00 | 1447.00 | 1919.00 |
| MED9 | 222.00 | 251.00 | 210.00 | 227.67 | 223.00 |
| MEF2A | 1251.00 | 1037.00 | 1286.00 | 1191.33 | 1207.00 |
| MEF2B | 6.00 | 3.00 | 5.00 | 4.67 | 1.00 |
| MEF2BNB | 95.00 | 68.00 | 78.00 | 80.33 | 113.00 |
| MEF2BNB-MEF2B | 10.00 | 7.00 | 5.00 | 7.33 | 9.00 |
| MEF2C | 248.00 | 209.00 | 261.00 | 239.33 | 246.00 |
| MEF2C-AS1 | 12.00 | 5.00 | 11.00 | 9.33 | 13.00 |
| MEF2D | 1671.00 | 1644.00 | 1683.00 | 1666.00 | 1442.00 |
| MEGF6 | 11.00 | 12.00 | 7.00 | 10.00 | 2.00 |
| MEGF8 | 2164.00 | 2029.00 | 2173.00 | 2122.00 | 1392.00 |
| MEGF9 | 271.00 | 280.00 | 319.00 | 290.00 | 210.00 |
| MEI1 | 15.00 | 12.00 | 11.00 | 12.67 | 15.00 |
| MEIG1 | 8.00 | 5.00 | 8.00 | 7.00 | 6.00 |
| MEIS1 | 124.00 | 129.00 | 147.00 | 133.33 | 159.00 |
| MEIS1-AS3 | 38.00 | 38.00 | 37.00 | 37.67 | 35.00 |
| MEIS2 | 402.00 | 397.00 | 456.00 | 418.33 | 445.00 |
| MEIS3 | 5.00 | 4.00 | 8.00 | 5.67 | 6.00 |
| MEIS3P1 | 145.00 | 139.00 | 157.00 | 147.00 | 151.00 |
| MELK | 1100.00 | 1056.00 | 997.00 | 1051.00 | 1571.00 |
| MEMO1 | 416.00 | 420.00 | 494.00 | 443.33 | 652.00 |
| MEN1 | 3763.00 | 3527.00 | 3836.00 | 3708.67 | 3627.00 |
| MEPCE | 1982.00 | 1926.00 | 2009.00 | 1972.33 | 1955.00 |
| MERTK | 438.00 | 391.00 | 471.00 | 433.33 | 346.00 |
| MESDC1 | 750.00 | 757.00 | 790.00 | 765.67 | 1057.00 |
| MESDC2 | 1950.00 | 1963.00 | 1856.00 | 1923.00 | 2024.00 |
| MESP1 | 12.00 | 16.00 | 17.00 | 15.00 | 32.00 |
| MESP2 | 9.00 | 7.00 | 8.00 | 8.00 | 11.00 |
| MEST | 16.00 | 17.00 | 26.00 | 19.67 | 18.00 |
| MET | 7276.00 | 6975.00 | 7652.00 | 7301.00 | 5277.00 |
| METAP1 | 1183.00 | 1151.00 | 1302.00 | 1212.00 | 1546.00 |
| METAP1D | 616.00 | 613.00 | 630.00 | 619.67 | 543.00 |
| METAP2 | 2173.00 | 2072.00 | 2375.00 | 2206.67 | 2881.00 |
| METRN | 576.00 | 666.00 | 691.00 | 644.33 | 700.00 |
| METRNL | 166.00 | 176.00 | 182.00 | 174.67 | 183.00 |
| METTL1 | 389.00 | 402.00 | 425.00 | 405.33 | 505.00 |
| METTL10 | 491.00 | 444.00 | 515.00 | 483.33 | 642.00 |
| METTL12 | 43.00 | 46.00 | 34.00 | 41.00 | 49.00 |
| METTL13 | 1626.00 | 1512.00 | 1627.00 | 1588.33 | 1688.00 |
| METTL14 | 374.00 | 417.00 | 409.00 | 400.00 | 453.00 |
| METTL15 | 477.00 | 464.00 | 495.00 | 478.67 | 556.00 |
| METTL16 | 1108.00 | 1082.00 | 1145.00 | 1111.67 | 1169.00 |
| METTL17 | 1418.00 | 1438.00 | 1421.00 | 1425.67 | 1565.00 |

| | | | | | |
|---|---|---|---|---|---|
| METTL18 | 206.00 | 204.00 | 189.00 | 199.67 | 282.00 |
| METTL20 | 55.00 | 42.00 | 34.00 | 43.67 | 42.00 |
| METTL21A | 205.00 | 182.00 | 208.00 | 198.33 | 312.00 |
| METTL21B | 308.00 | 285.00 | 333.00 | 308.67 | 320.00 |
| METTL21EP | 3.00 | 7.00 | 3.00 | 4.33 | 4.00 |
| METTL22 | 433.00 | 419.00 | 434.00 | 428.67 | 416.00 |
| METTL23 | 598.00 | 588.00 | 661.00 | 615.67 | 731.00 |
| METTL25 | 65.00 | 61.00 | 62.00 | 62.67 | 67.00 |
| METTL2A | 479.00 | 457.00 | 513.00 | 483.00 | 595.00 |
| METTL2B | 662.00 | 623.00 | 665.00 | 650.00 | 714.00 |
| METTL3 | 2699.00 | 2844.00 | 2854.00 | 2799.00 | 3041.00 |
| METTL4 | 965.00 | 1020.00 | 1095.00 | 1026.67 | 1111.00 |
| METTL5 | 818.00 | 758.00 | 844.00 | 806.67 | 1098.00 |
| METTL6 | 404.00 | 367.00 | 406.00 | 392.33 | 421.00 |
| METTL7A | 4.00 | 2.00 | 1.00 | 2.33 | 0.00 |
| METTL7B | 18.00 | 12.00 | 9.00 | 13.00 | 9.00 |
| METTL8 | 674.00 | 634.00 | 728.00 | 678.67 | 808.00 |
| METTL9 | 3675.00 | 3560.00 | 3851.00 | 3695.33 | 5476.00 |
| MEX3A | 732.00 | 765.00 | 961.00 | 819.33 | 889.00 |
| MEX3B | 144.00 | 124.00 | 166.00 | 144.67 | 136.00 |
| MEX3C | 981.00 | 901.00 | 1075.00 | 985.67 | 1259.00 |
| MEX3D | 271.00 | 286.00 | 281.00 | 279.33 | 322.00 |
| MFAP1 | 709.00 | 668.00 | 687.00 | 688.00 | 817.00 |
| MFAP2 | 80.00 | 111.00 | 88.00 | 93.00 | 90.00 |
| MFAP3 | 1146.00 | 981.00 | 1182.00 | 1103.00 | 924.00 |
| MFAP3L | 196.00 | 180.00 | 207.00 | 194.33 | 195.00 |
| MFF | 1451.00 | 1369.00 | 1579.00 | 1466.33 | 1909.00 |
| MFGE8 | 1987.00 | 2085.00 | 2095.00 | 2055.67 | 1472.00 |
| MFHAS1 | 609.00 | 542.00 | 650.00 | 600.33 | 468.00 |
| MFI2 | 2272.00 | 2487.00 | 2595.00 | 2451.33 | 2039.00 |
| MFI2-AS1 | 93.00 | 79.00 | 97.00 | 89.67 | 138.00 |
| MFN1 | 993.00 | 953.00 | 1047.00 | 997.67 | 1055.00 |
| MFN2 | 2686.00 | 2699.00 | 2954.00 | 2779.67 | 3026.00 |
| MFNG | 50.00 | 58.00 | 66.00 | 58.00 | 54.00 |
| MFSD1 | 2606.00 | 2519.00 | 2385.00 | 2503.33 | 2574.00 |
| MFSD10 | 6739.00 | 6725.00 | 6858.00 | 6774.00 | 7374.00 |
| MFSD11 | 530.00 | 510.00 | 535.00 | 525.00 | 659.00 |
| MFSD12 | 1794.00 | 1929.00 | 2049.00 | 1924.00 | 1453.00 |
| MFSD2A | 363.00 | 327.00 | 323.00 | 337.67 | 323.00 |
| MFSD3 | 1002.00 | 965.00 | 1050.00 | 1005.67 | 915.00 |
| MFSD5 | 751.00 | 761.00 | 787.00 | 766.33 | 747.00 |
| MFSD6 | 122.00 | 138.00 | 131.00 | 130.33 | 83.00 |
| MFSD8 | 531.00 | 562.00 | 608.00 | 567.00 | 589.00 |
| MFSD9 | 175.00 | 191.00 | 187.00 | 184.33 | 151.00 |
| MGA | 1345.00 | 1180.00 | 1427.00 | 1317.33 | 1259.00 |
| MGAT1 | 2685.00 | 2674.00 | 2653.00 | 2670.67 | 2460.00 |
| MGAT2 | 1358.00 | 1169.00 | 1394.00 | 1307.00 | 1434.00 |
| MGAT4A | 12.00 | 4.00 | 6.00 | 7.33 | 14.00 |

| | | | | | |
|---|---|---|---|---|---|
| MGAT4B | 3949.00 | 4041.00 | 4070.00 | 4020.00 | 4314.00 |
| MGAT5 | 2186.00 | 1930.00 | 2065.00 | 2060.33 | 1924.00 |
| MGAT5B | 789.00 | 774.00 | 818.00 | 793.67 | 445.00 |
| MGC16142 | 28.00 | 17.00 | 17.00 | 20.67 | 15.00 |
| MGC16275 | 3.00 | 4.00 | 1.00 | 2.67 | 7.00 |
| MGC27345 | 784.00 | 750.00 | 801.00 | 778.33 | 769.00 |
| MGC2889 | 21.00 | 19.00 | 21.00 | 20.33 | 17.00 |
| MGC57346 | 103.00 | 96.00 | 127.00 | 108.67 | 106.00 |
| MGC72080 | 118.00 | 141.00 | 142.00 | 133.67 | 187.00 |
| MGEA5 | 7614.00 | 7290.00 | 8237.00 | 7713.67 | 7569.00 |
| MGLL | 5747.00 | 5426.00 | 5711.00 | 5628.00 | 6431.00 |
| MGME1 | 769.00 | 786.00 | 860.00 | 805.00 | 1022.00 |
| MGRN1 | 1035.00 | 983.00 | 1055.00 | 1024.33 | 904.00 |
| MGST1 | 1765.00 | 1732.00 | 1680.00 | 1725.67 | 2020.00 |
| MGST2 | 140.00 | 168.00 | 128.00 | 145.33 | 174.00 |
| MGST3 | 1139.00 | 1139.00 | 1080.00 | 1119.33 | 1154.00 |
| MIA3 | 3552.00 | 3299.00 | 3488.00 | 3446.33 | 2946.00 |
| MIAT | 15.00 | 11.00 | 10.00 | 12.00 | 16.00 |
| MIATNB | 41.00 | 39.00 | 37.00 | 39.00 | 44.00 |
| MIB1 | 1376.00 | 1260.00 | 1667.00 | 1434.33 | 1300.00 |
| MIB2 | 1122.00 | 1218.00 | 1300.00 | 1213.33 | 1010.00 |
| MICA | 216.00 | 233.00 | 273.00 | 240.67 | 291.00 |
| MICAL1 | 1881.00 | 1914.00 | 1993.00 | 1929.33 | 1767.00 |
| MICAL2 | 10066.00 | 9207.00 | 10964.00 | 10079.00 | 8300.00 |
| MICAL3 | 3770.00 | 3682.00 | 3880.00 | 3777.33 | 2904.00 |
| MICALCL | 93.00 | 81.00 | 81.00 | 85.00 | 111.00 |
| MICALL1 | 1357.00 | 1248.00 | 1273.00 | 1292.67 | 1019.00 |
| MICALL2 | 456.00 | 442.00 | 504.00 | 467.33 | 349.00 |
| MICU1 | 875.00 | 925.00 | 876.00 | 892.00 | 1031.00 |
| MICU2 | 694.00 | 694.00 | 748.00 | 712.00 | 796.00 |
| MICU3 | 182.00 | 151.00 | 187.00 | 173.33 | 154.00 |
| MID1 | 618.00 | 572.00 | 680.00 | 623.33 | 515.00 |
| MID1IP1 | 558.00 | 583.00 | 639.00 | 593.33 | 684.00 |
| MID1IP1-AS1 | 23.00 | 22.00 | 23.00 | 22.67 | 14.00 |
| MIDN | 1421.00 | 1390.00 | 1505.00 | 1438.67 | 916.00 |
| MIEF1 | 1547.00 | 1263.00 | 1548.00 | 1452.67 | 1432.00 |
| MIEF2 | 225.00 | 211.00 | 204.00 | 213.33 | 158.00 |
| MIEN1 | 425.00 | 393.00 | 457.00 | 425.00 | 537.00 |
| MIER1 | 602.00 | 545.00 | 669.00 | 605.33 | 699.00 |
| MIER2 | 749.00 | 775.00 | 812.00 | 778.67 | 842.00 |
| MIER3 | 542.00 | 553.00 | 630.00 | 575.00 | 597.00 |
| MIF | 7.00 | 4.00 | 3.00 | 4.67 | 8.00 |
| MIF4GD | 364.00 | 328.00 | 375.00 | 355.67 | 422.00 |
| MIIP | 746.00 | 820.00 | 809.00 | 791.67 | 978.00 |
| MILR1 | 359.00 | 348.00 | 325.00 | 344.00 | 304.00 |
| MINA | 1975.00 | 1861.00 | 2104.00 | 1980.00 | 2266.00 |
| MINK1 | 4053.00 | 3971.00 | 4458.00 | 4160.67 | 4064.00 |
| MINOS1 | 708.00 | 709.00 | 701.00 | 706.00 | 939.00 |

| | | | | | |
|---|---|---|---|---|---|
| MINPP1 | 1236.00 | 1208.00 | 1214.00 | 1219.33 | 1272.00 |
| MIOS | 1130.00 | 1065.00 | 1236.00 | 1143.67 | 1381.00 |
| MIOX | 10.00 | 9.00 | 24.00 | 14.33 | 2.00 |
| MIPEP | 341.00 | 355.00 | 378.00 | 358.00 | 461.00 |
| MIPEPP3 | 51.00 | 54.00 | 71.00 | 58.67 | 60.00 |
| MIPOL1 | 446.00 | 391.00 | 494.00 | 443.67 | 534.00 |
| MIR100HG | 5058.00 | 4985.00 | 6285.00 | 5442.67 | 4338.00 |
| MIR103A2 | 6.00 | 5.00 | 3.00 | 4.67 | 8.00 |
| MIR106B | 8.00 | 12.00 | 7.00 | 9.00 | 4.00 |
| MIR10A | 7.00 | 6.00 | 3.00 | 5.33 | 5.00 |
| MIR10B | 4.00 | 3.00 | 4.00 | 3.67 | 5.00 |
| MIR1204 | 4.00 | 16.00 | 9.00 | 9.67 | 5.00 |
| MIR1254-1 | 63.00 | 74.00 | 72.00 | 69.67 | 53.00 |
| MIR125B1 | 14.00 | 10.00 | 15.00 | 13.00 | 0.00 |
| MIR1268A | 3.00 | 14.00 | 4.00 | 7.00 | 8.00 |
| MIR1285-1 | 8.00 | 6.00 | 6.00 | 6.67 | 5.00 |
| MIR1299 | 3.00 | 0.00 | 0.00 | 1.00 | 0.00 |
| MIR1304 | 101.00 | 107.00 | 122.00 | 110.00 | 110.00 |
| MIR137HG | 8.00 | 11.00 | 11.00 | 10.00 | 8.00 |
| MIR138-1 | 57.00 | 52.00 | 48.00 | 52.33 | 19.00 |
| MIR142 | 73.00 | 60.00 | 86.00 | 73.00 | 96.00 |
| MIR146A | 7.00 | 8.00 | 5.00 | 6.67 | 7.00 |
| MIR155HG | 214.00 | 218.00 | 243.00 | 225.00 | 218.00 |
| MIR17HG | 609.00 | 566.00 | 752.00 | 642.33 | 566.00 |
| MIR186 | 57.00 | 38.00 | 52.00 | 49.00 | 19.00 |
| MIR1908 | 3.00 | 3.00 | 5.00 | 3.67 | 3.00 |
| MIR1910 | 4.00 | 1.00 | 3.00 | 2.67 | 1.00 |
| MIR1914 | 67.00 | 61.00 | 80.00 | 69.33 | 87.00 |
| MIR199A1 | 11.00 | 18.00 | 18.00 | 15.67 | 16.00 |
| MIR21 | 28.00 | 28.00 | 45.00 | 33.67 | 25.00 |
| MIR221 | 786.00 | 850.00 | 880.00 | 838.67 | 739.00 |
| MIR222 | 78.00 | 66.00 | 82.00 | 75.33 | 58.00 |
| MIR224 | 11.00 | 32.00 | 27.00 | 23.33 | 9.00 |
| MIR23A | 4.00 | 2.00 | 8.00 | 4.67 | 2.00 |
| MIR24-1 | 3.00 | 2.00 | 2.00 | 2.33 | 2.00 |
| MIR25 | 49.00 | 35.00 | 46.00 | 43.33 | 70.00 |
| MIR26A1 | 3.00 | 0.00 | 0.00 | 1.00 | 0.00 |
| MIR26A2 | 3.00 | 2.00 | 1.00 | 2.00 | 3.00 |
| MIR27A | 5.00 | 3.00 | 2.00 | 3.33 | 2.00 |
| MIR27B | 4.00 | 2.00 | 6.00 | 4.00 | 2.00 |
| MIR298 | 8.00 | 1.00 | 12.00 | 7.00 | 4.00 |
| MIR29A | 60.00 | 43.00 | 42.00 | 48.33 | 45.00 |
| MIR29B1 | 3.00 | 3.00 | 13.00 | 6.33 | 6.00 |
| MIR29B2 | 7.00 | 10.00 | 24.00 | 13.67 | 12.00 |
| MIR29C | 23.00 | 28.00 | 33.00 | 28.00 | 29.00 |
| MIR3064 | 259.00 | 225.00 | 262.00 | 248.67 | 303.00 |
| MIR30C2 | 25.00 | 18.00 | 29.00 | 24.00 | 26.00 |
| MIR31 | 3.00 | 6.00 | 6.00 | 5.00 | 1.00 |

| | | | | | |
|---|---|---|---|---|---|
| MIR3136 | 3.00 | 3.00 | 3.00 | 3.00 | 1.00 |
| MIR3149 | 8.00 | 1.00 | 7.00 | 5.33 | 1.00 |
| MIR3153 | 6.00 | 11.00 | 9.00 | 8.67 | 3.00 |
| MIR3164 | 43.00 | 26.00 | 43.00 | 37.33 | 36.00 |
| MIR3174 | 3.00 | 2.00 | 0.00 | 1.67 | 0.00 |
| MIR3176 | 41.00 | 42.00 | 48.00 | 43.67 | 22.00 |
| MIR3191 | 12.00 | 10.00 | 4.00 | 8.67 | 2.00 |
| MIR3192 | 5.00 | 4.00 | 5.00 | 4.67 | 5.00 |
| MIR31HG | 443.00 | 465.00 | 523.00 | 477.00 | 395.00 |
| MIR339 | 4.00 | 1.00 | 1.00 | 2.00 | 1.00 |
| MIR34A | 3.00 | 5.00 | 6.00 | 4.67 | 1.00 |
| MIR3605 | 7.00 | 12.00 | 6.00 | 8.33 | 11.00 |
| MIR3607 | 6.00 | 6.00 | 7.00 | 6.33 | 4.00 |
| MIR3611 | 3.00 | 0.00 | 0.00 | 1.00 | 0.00 |
| MIR3654 | 1784.00 | 1689.00 | 1806.00 | 1759.67 | 2210.00 |
| MIR3657 | 5.00 | 3.00 | 1.00 | 3.00 | 3.00 |
| MIR3662 | 4.00 | 2.00 | 7.00 | 4.33 | 0.00 |
| MIR3671 | 8.00 | 5.00 | 9.00 | 7.33 | 2.00 |
| MIR3677 | 6.00 | 1.00 | 6.00 | 4.33 | 7.00 |
| MIR3682 | 37.00 | 23.00 | 44.00 | 34.67 | 51.00 |
| MIR3685 | 5.00 | 10.00 | 4.00 | 6.33 | 11.00 |
| MIR3689B | 6.00 | 5.00 | 2.00 | 4.33 | 0.00 |
| MIR3689C | 6.00 | 4.00 | 6.00 | 5.33 | 0.00 |
| MIR3689F | 4.00 | 4.00 | 3.00 | 3.67 | 1.00 |
| MIR374B | 7.00 | 4.00 | 6.00 | 5.67 | 3.00 |
| MIR378H | 3.00 | 2.00 | 5.00 | 3.33 | 6.00 |
| MIR3911 | 7.00 | 5.00 | 4.00 | 5.33 | 9.00 |
| MIR3916 | 121.00 | 145.00 | 134.00 | 133.33 | 115.00 |
| MIR3942 | 5.00 | 3.00 | 5.00 | 4.33 | 3.00 |
| MIR421 | 10.00 | 13.00 | 4.00 | 9.00 | 2.00 |
| MIR423 | 3.00 | 1.00 | 4.00 | 2.67 | 4.00 |
| MIR4258 | 24.00 | 26.00 | 22.00 | 24.00 | 20.00 |
| MIR4263 | 4.00 | 7.00 | 2.00 | 4.33 | 12.00 |
| MIR4271 | 16.00 | 18.00 | 13.00 | 15.67 | 10.00 |
| MIR4292 | 30.00 | 20.00 | 27.00 | 25.67 | 18.00 |
| MIR4326 | 4.00 | 8.00 | 3.00 | 5.00 | 4.00 |
| MIR4426 | 2697.00 | 2694.00 | 3015.00 | 2802.00 | 3547.00 |
| MIR4435-2HG | 185.00 | 160.00 | 177.00 | 174.00 | 151.00 |
| MIR4442 | 33.00 | 17.00 | 42.00 | 30.67 | 29.00 |
| MIR4458HG | 244.00 | 274.00 | 297.00 | 271.67 | 308.00 |
| MIR4477B | 7.00 | 14.00 | 11.00 | 10.67 | 16.00 |
| MIR4479 | 3.00 | 1.00 | 2.00 | 2.00 | 2.00 |
| MIR4482 | 5.00 | 1.00 | 0.00 | 2.00 | 1.00 |
| MIR4489 | 4.00 | 0.00 | 1.00 | 1.67 | 2.00 |
| MIR4500HG | 5.00 | 9.00 | 10.00 | 8.00 | 22.00 |
| MIR4512 | 5.00 | 3.00 | 1.00 | 3.00 | 1.00 |
| MIR4517 | 51.00 | 23.00 | 32.00 | 35.33 | 26.00 |
| MIR4653 | 12.00 | 9.00 | 10.00 | 10.33 | 7.00 |

| | | | | | |
|---|---|---|---|---|---|
| MIR4668 | 3.00 | 4.00 | 1.00 | 2.67 | 1.00 |
| MIR4691 | 27.00 | 31.00 | 19.00 | 25.67 | 16.00 |
| MIR4697HG | 6.00 | 5.00 | 7.00 | 6.00 | 5.00 |
| MIR4701 | 9.00 | 4.00 | 6.00 | 6.33 | 1.00 |
| MIR4712 | 215.00 | 183.00 | 199.00 | 199.00 | 99.00 |
| MIR4720 | 7.00 | 10.00 | 8.00 | 8.33 | 11.00 |
| MIR4730 | 8.00 | 8.00 | 3.00 | 6.33 | 3.00 |
| MIR4746 | 5.00 | 10.00 | 6.00 | 7.00 | 3.00 |
| MIR4766 | 3.00 | 1.00 | 2.00 | 2.00 | 1.00 |
| MIR4782 | 3.00 | 5.00 | 0.00 | 2.67 | 2.00 |
| MIR4784 | 8.00 | 11.00 | 8.00 | 9.00 | 2.00 |
| MIR4786 | 8.00 | 10.00 | 4.00 | 7.33 | 10.00 |
| MIR4787 | 3.00 | 1.00 | 0.00 | 1.33 | 2.00 |
| MIR491 | 6.00 | 9.00 | 6.00 | 7.00 | 4.00 |
| MIR499B | 8.00 | 5.00 | 1.00 | 4.67 | 4.00 |
| MIR5006 | 4.00 | 3.00 | 2.00 | 3.00 | 4.00 |
| MIR5010 | 6.00 | 20.00 | 19.00 | 15.00 | 9.00 |
| MIR503 | 92.00 | 118.00 | 117.00 | 109.00 | 73.00 |
| MIR503HG | 1054.00 | 1068.00 | 1203.00 | 1108.33 | 1082.00 |
| MIR5047 | 1840.00 | 1895.00 | 1870.00 | 1868.33 | 2251.00 |
| MIR542 | 3.00 | 2.00 | 0.00 | 1.67 | 1.00 |
| MIR548A3 | 6.00 | 2.00 | 1.00 | 3.00 | 0.00 |
| MIR548AM | 4.00 | 3.00 | 1.00 | 2.67 | 0.00 |
| MIR548AN | 4.00 | 2.00 | 3.00 | 3.00 | 1.00 |
| MIR548AR | 16.00 | 5.00 | 3.00 | 8.00 | 5.00 |
| MIR548D1 | 3.00 | 2.00 | 4.00 | 3.00 | 5.00 |
| MIR553 | 5.00 | 2.00 | 4.00 | 3.67 | 3.00 |
| MIR5581 | 14.00 | 12.00 | 4.00 | 10.00 | 13.00 |
| MIR5587 | 14.00 | 15.00 | 15.00 | 14.67 | 4.00 |
| MIR563 | 3.00 | 3.00 | 2.00 | 2.67 | 4.00 |
| MIR573 | 12.00 | 10.00 | 18.00 | 13.33 | 9.00 |
| MIR574 | 3.00 | 1.00 | 2.00 | 2.00 | 0.00 |
| MIR589 | 6.00 | 9.00 | 1.00 | 5.33 | 3.00 |
| MIR590 | 48.00 | 33.00 | 47.00 | 42.67 | 30.00 |
| MIR600HG | 191.00 | 231.00 | 224.00 | 215.33 | 191.00 |
| MIR614 | 42.00 | 45.00 | 52.00 | 46.33 | 37.00 |
| MIR616 | 19.00 | 18.00 | 18.00 | 18.33 | 12.00 |
| MIR622 | 27.00 | 26.00 | 20.00 | 24.33 | 9.00 |
| MIR631 | 17.00 | 12.00 | 25.00 | 18.00 | 22.00 |
| MIR635 | 19.00 | 15.00 | 10.00 | 14.67 | 4.00 |
| MIR637 | 21.00 | 23.00 | 25.00 | 23.00 | 24.00 |
| MIR641 | 30.00 | 24.00 | 23.00 | 25.67 | 7.00 |
| MIR646HG | 25.00 | 17.00 | 21.00 | 21.00 | 10.00 |
| MIR647 | 53.00 | 70.00 | 73.00 | 65.33 | 69.00 |
| MIR6505 | 5.00 | 3.00 | 2.00 | 3.33 | 4.00 |
| MIR6516 | 3.00 | 1.00 | 0.00 | 1.33 | 0.00 |
| MIR661 | 4.00 | 3.00 | 3.00 | 3.33 | 3.00 |
| MIR6728 | 5.00 | 11.00 | 3.00 | 6.33 | 1.00 |

| | | | | | |
|---|---|---|---|---|---|
| MIR6733 | 4.00 | 0.00 | 4.00 | 2.67 | 2.00 |
| MIR6734 | 3.00 | 2.00 | 0.00 | 1.67 | 1.00 |
| MIR6739 | 3.00 | 6.00 | 5.00 | 4.67 | 3.00 |
| MIR6753 | 3.00 | 8.00 | 9.00 | 6.67 | 4.00 |
| MIR6754 | 3.00 | 1.00 | 2.00 | 2.00 | 0.00 |
| MIR6758 | 26.00 | 22.00 | 28.00 | 25.33 | 19.00 |
| MIR6763 | 5.00 | 3.00 | 4.00 | 4.00 | 0.00 |
| MIR6772 | 5.00 | 7.00 | 4.00 | 5.33 | 4.00 |
| MIR6775 | 6.00 | 4.00 | 7.00 | 5.67 | 6.00 |
| MIR6797 | 5.00 | 4.00 | 3.00 | 4.00 | 11.00 |
| MIR6835 | 98.00 | 101.00 | 127.00 | 108.67 | 52.00 |
| MIR6840 | 13.00 | 13.00 | 9.00 | 11.67 | 7.00 |
| MIR6845 | 12.00 | 8.00 | 6.00 | 8.67 | 2.00 |
| MIR6847 | 11.00 | 3.00 | 7.00 | 7.00 | 1.00 |
| MIR6855 | 4.00 | 0.00 | 0.00 | 1.33 | 0.00 |
| MIR6884 | 6.00 | 5.00 | 6.00 | 5.67 | 3.00 |
| MIR6895 | 5.00 | 3.00 | 3.00 | 3.67 | 4.00 |
| MIR7-1 | 38.00 | 48.00 | 44.00 | 43.33 | 58.00 |
| MIR7106 | 4.00 | 1.00 | 2.00 | 2.33 | 5.00 |
| MIR7111 | 12.00 | 6.00 | 14.00 | 10.67 | 18.00 |
| MIR7161 | 9.00 | 5.00 | 7.00 | 7.00 | 7.00 |
| MIR762HG | 26.00 | 58.00 | 52.00 | 45.33 | 39.00 |
| MIR7641-2 | 10.00 | 3.00 | 4.00 | 5.67 | 2.00 |
| MIR765 | 3.00 | 0.00 | 2.00 | 1.67 | 0.00 |
| MIR7851 | 5.00 | 1.00 | 2.00 | 2.67 | 2.00 |
| MIR7856 | 4.00 | 8.00 | 4.00 | 5.33 | 7.00 |
| MIR8058 | 6.00 | 6.00 | 6.00 | 6.00 | 4.00 |
| MIR93 | 11.00 | 10.00 | 14.00 | 11.67 | 11.00 |
| MIR940 | 5.00 | 8.00 | 17.00 | 10.00 | 14.00 |
| MIR99AHG | 133.00 | 123.00 | 163.00 | 139.67 | 103.00 |
| MIRLET7BHG | 346.00 | 334.00 | 459.00 | 379.67 | 272.00 |
| MIRLET7D | 52.00 | 76.00 | 58.00 | 62.00 | 56.00 |
| MIRLET7F1 | 14.00 | 19.00 | 11.00 | 14.67 | 3.00 |
| MIRLET7G | 3.00 | 3.00 | 4.00 | 3.33 | 1.00 |
| MIRLET7I | 8.00 | 18.00 | 11.00 | 12.33 | 11.00 |
| MIS12 | 1493.00 | 1474.00 | 1599.00 | 1522.00 | 1608.00 |
| MIS18A | 364.00 | 368.00 | 364.00 | 365.33 | 505.00 |
| MIS18BP1 | 991.00 | 893.00 | 979.00 | 954.33 | 954.00 |
| MITD1 | 565.00 | 531.00 | 575.00 | 557.00 | 688.00 |
| MITF | 179.00 | 207.00 | 190.00 | 192.00 | 153.00 |
| MIXL1 | 4.00 | 0.00 | 1.00 | 1.67 | 3.00 |
| MKI67 | 5085.00 | 4895.00 | 5014.00 | 4998.00 | 5001.00 |
| MKKS | 1156.00 | 1119.00 | 1189.00 | 1154.67 | 1430.00 |
| MKL1 | 517.00 | 517.00 | 555.00 | 529.67 | 501.00 |
| MKL2 | 655.00 | 569.00 | 618.00 | 614.00 | 548.00 |
| MKLN1 | 2192.00 | 2003.00 | 2351.00 | 2182.00 | 2227.00 |
| MKLN1-AS | 88.00 | 85.00 | 101.00 | 91.33 | 86.00 |
| MKNK1 | 687.00 | 613.00 | 686.00 | 662.00 | 661.00 |

| | | | | | |
|---|---|---|---|---|---|
| MKNK2 | 1792.00 | 1788.00 | 2100.00 | 1893.33 | 1487.00 |
| MKRN1 | 1896.00 | 1875.00 | 2064.00 | 1945.00 | 2087.00 |
| MKRN2 | 993.00 | 942.00 | 1121.00 | 1018.67 | 1101.00 |
| MKS1 | 897.00 | 852.00 | 888.00 | 879.00 | 875.00 |
| MKX | 126.00 | 101.00 | 126.00 | 117.67 | 126.00 |
| MLEC | 5927.00 | 5576.00 | 5878.00 | 5793.67 | 4562.00 |
| MLF1 | 332.00 | 328.00 | 397.00 | 352.33 | 463.00 |
| MLF2 | 2714.00 | 2640.00 | 2836.00 | 2730.00 | 3306.00 |
| MLH1 | 879.00 | 778.00 | 835.00 | 830.67 | 1000.00 |
| MLH3 | 792.00 | 692.00 | 846.00 | 776.67 | 792.00 |
| MLKL | 582.00 | 619.00 | 599.00 | 600.00 | 572.00 |
| MLLT1 | 2408.00 | 2285.00 | 2477.00 | 2390.00 | 2604.00 |
| MLLT10 | 1939.00 | 1765.00 | 2016.00 | 1906.67 | 2097.00 |
| MLLT11 | 148.00 | 131.00 | 129.00 | 136.00 | 174.00 |
| MLLT3 | 134.00 | 124.00 | 144.00 | 134.00 | 149.00 |
| MLLT4 | 3700.00 | 3422.00 | 3907.00 | 3676.33 | 3361.00 |
| MLLT4-AS1 | 21.00 | 22.00 | 23.00 | 22.00 | 31.00 |
| MLLT6 | 6.00 | 4.00 | 9.00 | 6.33 | 2.00 |
| MLPH | 12379.00 | 12737.00 | 13886.00 | 13000.67 | 9807.00 |
| MLST8 | 834.00 | 840.00 | 858.00 | 844.00 | 1191.00 |
| MLX | 1109.00 | 1120.00 | 1058.00 | 1095.67 | 1460.00 |
| MLXIP | 2391.00 | 2426.00 | 2717.00 | 2511.33 | 2049.00 |
| MLYCD | 255.00 | 205.00 | 239.00 | 233.00 | 244.00 |
| MMAA | 43.00 | 42.00 | 52.00 | 45.67 | 70.00 |
| MMAB | 265.00 | 249.00 | 258.00 | 257.33 | 317.00 |
| MMACHC | 263.00 | 228.00 | 279.00 | 256.67 | 254.00 |
| MMADHC | 1960.00 | 1798.00 | 2181.00 | 1979.67 | 2646.00 |
| MMD | 282.00 | 280.00 | 307.00 | 289.67 | 290.00 |
| MME | 7559.00 | 7011.00 | 7765.00 | 7445.00 | 10683.00 |
| MMGT1 | 833.00 | 699.00 | 785.00 | 772.33 | 939.00 |
| MMP1 | 930124.00 | 919814.00 | 938341.00 | 929426.33 | 940851.00 |
| MMP10 | 12.00 | 17.00 | 5.00 | 11.33 | 15.00 |
| MMP12 | 4.00 | 5.00 | 8.00 | 5.67 | 5.00 |
| MMP13 | 4.00 | 5.00 | 5.00 | 4.67 | 5.00 |
| MMP14 | 17728.00 | 18442.00 | 18289.00 | 18153.00 | 15792.00 |
| MMP15 | 15.00 | 9.00 | 11.00 | 11.67 | 13.00 |
| MMP16 | 416.00 | 393.00 | 459.00 | 422.67 | 329.00 |
| MMP17 | 349.00 | 338.00 | 369.00 | 352.00 | 284.00 |
| MMP19 | 207.00 | 184.00 | 228.00 | 206.33 | 172.00 |
| MMP2 | 3870.00 | 3859.00 | 3737.00 | 3822.00 | 2108.00 |
| MMP24-AS1 | 802.00 | 835.00 | 830.00 | 822.33 | 810.00 |
| MMP25 | 31.00 | 27.00 | 39.00 | 32.33 | 22.00 |
| MMP25-AS1 | 105.00 | 101.00 | 116.00 | 107.33 | 70.00 |
| MMP3 | 4626.00 | 4479.00 | 4484.00 | 4529.67 | 2578.00 |
| MMS19 | 3286.00 | 3089.00 | 3410.00 | 3261.67 | 3442.00 |
| MMS22L | 590.00 | 535.00 | 603.00 | 576.00 | 725.00 |
| MN1 | 3.00 | 2.00 | 1.00 | 2.00 | 1.00 |
| MNAT1 | 739.00 | 668.00 | 690.00 | 699.00 | 985.00 |

| | | | | | |
|---|---|---|---|---|---|
| MND1 | 115.00 | 127.00 | 105.00 | 115.67 | 180.00 |
| MNS1 | 96.00 | 95.00 | 68.00 | 86.33 | 122.00 |
| MNT | 1066.00 | 1040.00 | 1167.00 | 1091.00 | 1117.00 |
| MOB1A | 678.00 | 606.00 | 757.00 | 680.33 | 747.00 |
| MOB1B | 580.00 | 581.00 | 578.00 | 579.67 | 686.00 |
| MOB2 | 351.00 | 385.00 | 382.00 | 372.67 | 441.00 |
| MOB3A | 1006.00 | 1018.00 | 1069.00 | 1031.00 | 858.00 |
| MOB3B | 9.00 | 10.00 | 10.00 | 9.67 | 17.00 |
| MOB3C | 216.00 | 274.00 | 274.00 | 254.67 | 250.00 |
| MOB4 | 30.00 | 30.00 | 48.00 | 36.00 | 45.00 |
| MOCOS | 1733.00 | 1600.00 | 1705.00 | 1679.33 | 1178.00 |
| MOCS1 | 568.00 | 565.00 | 554.00 | 562.33 | 577.00 |
| MOCS2 | 1040.00 | 895.00 | 1014.00 | 983.00 | 1311.00 |
| MOCS3 | 400.00 | 380.00 | 433.00 | 404.33 | 465.00 |
| MOGAT1 | 3.00 | 0.00 | 1.00 | 1.33 | 0.00 |
| MOGS | 4099.00 | 4259.00 | 4265.00 | 4207.67 | 3607.00 |
| MOK | 20547.00 | 20739.00 | 22141.00 | 21142.33 | 30034.00 |
| MON1A | 488.00 | 418.00 | 499.00 | 468.33 | 464.00 |
| MON1B | 1085.00 | 1058.00 | 1019.00 | 1054.00 | 905.00 |
| MON2 | 1147.00 | 1031.00 | 1117.00 | 1098.33 | 1083.00 |
| MORC2 | 2226.00 | 2096.00 | 2328.00 | 2216.67 | 2183.00 |
| MORC3 | 632.00 | 551.00 | 632.00 | 605.00 | 627.00 |
| MORC4 | 562.00 | 525.00 | 581.00 | 556.00 | 725.00 |
| MORF4L1 | 2288.00 | 2064.00 | 2274.00 | 2208.67 | 2706.00 |
| MORF4L2 | 8697.00 | 8287.00 | 8805.00 | 8596.33 | 9501.00 |
| MORF4L2-AS1 | 144.00 | 134.00 | 156.00 | 144.67 | 156.00 |
| MORN1 | 166.00 | 166.00 | 136.00 | 156.00 | 140.00 |
| MORN2 | 149.00 | 167.00 | 166.00 | 160.67 | 222.00 |
| MORN3 | 5.00 | 5.00 | 3.00 | 4.33 | 3.00 |
| MORN4 | 539.00 | 506.00 | 532.00 | 525.67 | 469.00 |
| MOSPD1 | 333.00 | 301.00 | 341.00 | 325.00 | 381.00 |
| MOSPD2 | 376.00 | 345.00 | 411.00 | 377.33 | 337.00 |
| MOSPD3 | 263.00 | 316.00 | 270.00 | 283.00 | 327.00 |
| MOV10 | 1430.00 | 1408.00 | 1480.00 | 1439.33 | 1300.00 |
| MOXD1 | 1297.00 | 1287.00 | 1245.00 | 1276.33 | 1720.00 |
| MOXD2P | 12.00 | 2.00 | 11.00 | 8.33 | 0.00 |
| MPC1 | 590.00 | 522.00 | 575.00 | 562.33 | 629.00 |
| MPC2 | 409.00 | 383.00 | 424.00 | 405.33 | 435.00 |
| MPDU1 | 1370.00 | 1415.00 | 1227.00 | 1337.33 | 1599.00 |
| MPDZ | 4080.00 | 3809.00 | 4483.00 | 4124.00 | 3343.00 |
| MPG | 1212.00 | 1203.00 | 1277.00 | 1230.67 | 1120.00 |
| MPHOSPH10 | 933.00 | 927.00 | 1014.00 | 958.00 | 1053.00 |
| MPHOSPH6 | 845.00 | 886.00 | 1004.00 | 911.67 | 1118.00 |
| MPHOSPH8 | 1117.00 | 1036.00 | 1188.00 | 1113.67 | 1003.00 |
| MPHOSPH9 | 518.00 | 498.00 | 630.00 | 548.67 | 499.00 |
| MPI | 571.00 | 560.00 | 568.00 | 566.33 | 605.00 |
| MPL | 5.00 | 2.00 | 3.00 | 3.33 | 9.00 |
| MPLKIP | 775.00 | 793.00 | 848.00 | 805.33 | 1031.00 |

| | | | | | |
|---|---|---|---|---|---|
| MPND | 115.00 | 119.00 | 106.00 | 113.33 | 129.00 |
| MPP1 | 98.00 | 119.00 | 79.00 | 98.67 | 182.00 |
| MPP2 | 51.00 | 27.00 | 45.00 | 41.00 | 46.00 |
| MPP3 | 917.00 | 1036.00 | 1071.00 | 1008.00 | 871.00 |
| MPP4 | 1328.00 | 1253.00 | 1312.00 | 1297.67 | 1327.00 |
| MPP5 | 834.00 | 796.00 | 907.00 | 845.67 | 978.00 |
| MPP6 | 957.00 | 886.00 | 1100.00 | 981.00 | 1210.00 |
| MPP7 | 199.00 | 166.00 | 211.00 | 192.00 | 156.00 |
| MPPE1 | 412.00 | 432.00 | 464.00 | 436.00 | 499.00 |
| MPRIP | 8648.00 | 8026.00 | 8720.00 | 8464.67 | 6645.00 |
| MPST | 568.00 | 594.00 | 622.00 | 594.67 | 645.00 |
| MPV17 | 1489.00 | 1476.00 | 1510.00 | 1491.67 | 1308.00 |
| MPV17L2 | 336.00 | 345.00 | 355.00 | 345.33 | 395.00 |
| MPZ | 51.00 | 35.00 | 32.00 | 39.33 | 14.00 |
| MPZL1 | 2961.00 | 2877.00 | 2775.00 | 2871.00 | 2699.00 |
| MR1 | 351.00 | 320.00 | 321.00 | 330.67 | 284.00 |
| MRAP2 | 6.00 | 7.00 | 4.00 | 5.67 | 13.00 |
| MRAS | 60.00 | 85.00 | 80.00 | 75.00 | 79.00 |
| MRC1 | 5.00 | 6.00 | 2.00 | 4.33 | 4.00 |
| MRC2 | 684.00 | 663.00 | 609.00 | 652.00 | 444.00 |
| MRE11A | 1272.00 | 1133.00 | 1279.00 | 1228.00 | 1372.00 |
| MREG | 120.00 | 126.00 | 123.00 | 123.00 | 127.00 |
| MRFAP1 | 2432.00 | 2401.00 | 2588.00 | 2473.67 | 3312.00 |
| MRFAP1L1 | 753.00 | 703.00 | 755.00 | 737.00 | 900.00 |
| MRGBP | 940.00 | 1032.00 | 1041.00 | 1004.33 | 1268.00 |
| MRI1 | 1289.00 | 1379.00 | 1524.00 | 1397.33 | 1392.00 |
| MRM1 | 18.00 | 13.00 | 18.00 | 16.33 | 13.00 |
| MROH1 | 1758.00 | 1762.00 | 1984.00 | 1834.67 | 1255.00 |
| MROH8 | 5.00 | 5.00 | 9.00 | 6.33 | 11.00 |
| MRPL1 | 398.00 | 384.00 | 430.00 | 404.00 | 516.00 |
| MRPL10 | 1070.00 | 954.00 | 1075.00 | 1033.00 | 1264.00 |
| MRPL11 | 1482.00 | 1527.00 | 1589.00 | 1532.67 | 1983.00 |
| MRPL12 | 1636.00 | 1459.00 | 1707.00 | 1600.67 | 2324.00 |
| MRPL13 | 996.00 | 907.00 | 990.00 | 964.33 | 1299.00 |
| MRPL14 | 570.00 | 629.00 | 667.00 | 622.00 | 760.00 |
| MRPL15 | 1384.00 | 1325.00 | 1417.00 | 1375.33 | 1793.00 |
| MRPL16 | 683.00 | 639.00 | 697.00 | 673.00 | 882.00 |
| MRPL17 | 1033.00 | 1009.00 | 1109.00 | 1050.33 | 1461.00 |
| MRPL18 | 794.00 | 779.00 | 814.00 | 795.67 | 1024.00 |
| MRPL19 | 1102.00 | 933.00 | 1130.00 | 1055.00 | 1358.00 |
| MRPL2 | 713.00 | 687.00 | 767.00 | 722.33 | 955.00 |
| MRPL20 | 755.00 | 757.00 | 823.00 | 778.33 | 1072.00 |
| MRPL21 | 660.00 | 688.00 | 726.00 | 691.33 | 979.00 |
| MRPL22 | 431.00 | 432.00 | 417.00 | 426.67 | 552.00 |
| MRPL23 | 79.00 | 73.00 | 77.00 | 76.33 | 108.00 |
| MRPL24 | 1949.00 | 2037.00 | 2080.00 | 2022.00 | 2461.00 |
| MRPL27 | 629.00 | 636.00 | 644.00 | 636.33 | 768.00 |
| MRPL28 | 806.00 | 875.00 | 886.00 | 855.67 | 1002.00 |

| | | | | | |
|---|---|---|---|---|---|
| MRPL3 | 2907.00 | 2841.00 | 3108.00 | 2952.00 | 3907.00 |
| MRPL30 | 1319.00 | 1183.00 | 1438.00 | 1313.33 | 1587.00 |
| MRPL32 | 795.00 | 818.00 | 939.00 | 850.67 | 999.00 |
| MRPL33 | 1179.00 | 1252.00 | 1171.00 | 1200.67 | 1598.00 |
| MRPL34 | 566.00 | 585.00 | 711.00 | 620.67 | 900.00 |
| MRPL35 | 845.00 | 748.00 | 855.00 | 816.00 | 972.00 |
| MRPL36 | 514.00 | 514.00 | 581.00 | 536.33 | 689.00 |
| MRPL37 | 1679.00 | 1587.00 | 1836.00 | 1700.67 | 2048.00 |
| MRPL38 | 1006.00 | 1050.00 | 1108.00 | 1054.67 | 1273.00 |
| MRPL39 | 547.00 | 439.00 | 519.00 | 501.67 | 658.00 |
| MRPL4 | 1613.00 | 1508.00 | 1637.00 | 1586.00 | 1941.00 |
| MRPL40 | 561.00 | 579.00 | 566.00 | 568.67 | 707.00 |
| MRPL41 | 474.00 | 474.00 | 536.00 | 494.67 | 636.00 |
| MRPL42 | 1074.00 | 955.00 | 1170.00 | 1066.33 | 1149.00 |
| MRPL43 | 927.00 | 937.00 | 971.00 | 945.00 | 1071.00 |
| MRPL44 | 591.00 | 531.00 | 613.00 | 578.33 | 717.00 |
| MRPL45 | 120.00 | 89.00 | 121.00 | 110.00 | 154.00 |
| MRPL45P2 | 63.00 | 56.00 | 59.00 | 59.33 | 64.00 |
| MRPL46 | 436.00 | 429.00 | 422.00 | 429.00 | 516.00 |
| MRPL47 | 927.00 | 827.00 | 904.00 | 886.00 | 1251.00 |
| MRPL48 | 282.00 | 287.00 | 318.00 | 295.67 | 493.00 |
| MRPL49 | 1244.00 | 1188.00 | 1336.00 | 1256.00 | 1320.00 |
| MRPL50 | 419.00 | 350.00 | 401.00 | 390.00 | 557.00 |
| MRPL51 | 1722.00 | 1795.00 | 1883.00 | 1800.00 | 2373.00 |
| MRPL52 | 1115.00 | 1101.00 | 1175.00 | 1130.33 | 1534.00 |
| MRPL53 | 358.00 | 388.00 | 360.00 | 368.67 | 550.00 |
| MRPL54 | 312.00 | 312.00 | 315.00 | 313.00 | 482.00 |
| MRPL55 | 345.00 | 321.00 | 351.00 | 339.00 | 393.00 |
| MRPL57 | 540.00 | 593.00 | 602.00 | 578.33 | 710.00 |
| MRPL9 | 1349.00 | 1271.00 | 1422.00 | 1347.33 | 1758.00 |
| MRPS10 | 1443.00 | 1284.00 | 1517.00 | 1414.67 | 1661.00 |
| MRPS11 | 506.00 | 466.00 | 551.00 | 507.67 | 490.00 |
| MRPS12 | 555.00 | 632.00 | 706.00 | 631.00 | 736.00 |
| MRPS14 | 502.00 | 453.00 | 500.00 | 485.00 | 500.00 |
| MRPS15 | 779.00 | 750.00 | 864.00 | 797.67 | 1018.00 |
| MRPS16 | 2776.00 | 2539.00 | 2888.00 | 2734.33 | 3211.00 |
| MRPS17 | 345.00 | 335.00 | 366.00 | 348.67 | 459.00 |
| MRPS18A | 480.00 | 395.00 | 468.00 | 447.67 | 609.00 |
| MRPS18C | 228.00 | 225.00 | 196.00 | 216.33 | 296.00 |
| MRPS2 | 1381.00 | 1408.00 | 1595.00 | 1461.33 | 1713.00 |
| MRPS21 | 1408.00 | 1476.00 | 1588.00 | 1490.67 | 1914.00 |
| MRPS22 | 1013.00 | 1023.00 | 1106.00 | 1047.33 | 1234.00 |
| MRPS23 | 974.00 | 925.00 | 1086.00 | 995.00 | 1285.00 |
| MRPS24 | 63.00 | 81.00 | 86.00 | 76.67 | 123.00 |
| MRPS25 | 2786.00 | 2782.00 | 2886.00 | 2818.00 | 2738.00 |
| MRPS26 | 797.00 | 835.00 | 843.00 | 825.00 | 1063.00 |
| MRPS27 | 1809.00 | 1821.00 | 2045.00 | 1891.67 | 2018.00 |
| MRPS28 | 475.00 | 473.00 | 512.00 | 486.67 | 653.00 |

| | | | | | |
|---|---|---|---|---|---|
| MRPS30 | 797.00 | 725.00 | 808.00 | 776.67 | 1007.00 |
| MRPS31 | 454.00 | 453.00 | 479.00 | 462.00 | 447.00 |
| MRPS31P5 | 239.00 | 234.00 | 270.00 | 247.67 | 231.00 |
| MRPS33 | 911.00 | 855.00 | 846.00 | 870.67 | 1103.00 |
| MRPS34 | 1416.00 | 1396.00 | 1537.00 | 1449.67 | 1980.00 |
| MRPS35 | 1253.00 | 1121.00 | 1437.00 | 1270.33 | 1665.00 |
| MRPS36 | 117.00 | 105.00 | 89.00 | 103.67 | 138.00 |
| MRPS5 | 1855.00 | 1765.00 | 1958.00 | 1859.33 | 2245.00 |
| MRPS6 | 734.00 | 656.00 | 733.00 | 707.67 | 1047.00 |
| MRPS7 | 722.00 | 721.00 | 816.00 | 753.00 | 1016.00 |
| MRPS9 | 883.00 | 830.00 | 858.00 | 857.00 | 1223.00 |
| MRRF | 785.00 | 768.00 | 866.00 | 806.33 | 927.00 |
| MRS2 | 886.00 | 852.00 | 972.00 | 903.33 | 1063.00 |
| MRTO4 | 1303.00 | 1311.00 | 1390.00 | 1334.67 | 1772.00 |
| MSANTD1 | 10.00 | 9.00 | 9.00 | 9.33 | 13.00 |
| MSANTD2 | 840.00 | 801.00 | 930.00 | 857.00 | 801.00 |
| MSANTD3 | 1105.00 | 1095.00 | 1234.00 | 1144.67 | 1372.00 |
| MSANTD4 | 430.00 | 454.00 | 573.00 | 485.67 | 478.00 |
| MSC | 11.00 | 15.00 | 10.00 | 12.00 | 10.00 |
| MSC-AS1 | 88.00 | 79.00 | 109.00 | 92.00 | 79.00 |
| MSH2 | 1317.00 | 1230.00 | 1324.00 | 1290.33 | 1491.00 |
| MSH3 | 451.00 | 403.00 | 471.00 | 441.67 | 416.00 |
| MSH4 | 20.00 | 15.00 | 20.00 | 18.33 | 22.00 |
| MSH6 | 1785.00 | 1700.00 | 1853.00 | 1779.33 | 1833.00 |
| MSI2 | 372.00 | 344.00 | 387.00 | 367.67 | 323.00 |
| MSL1 | 2025.00 | 1979.00 | 2151.00 | 2051.67 | 2116.00 |
| MSL2 | 758.00 | 719.00 | 1012.00 | 829.67 | 972.00 |
| MSL3 | 1360.00 | 1357.00 | 1457.00 | 1391.33 | 1296.00 |
| MSL3P1 | 357.00 | 324.00 | 409.00 | 363.33 | 364.00 |
| MSMO1 | 696.00 | 632.00 | 676.00 | 668.00 | 706.00 |
| MSMP | 4.00 | 2.00 | 2.00 | 2.67 | 2.00 |
| MSN | 24667.00 | 23073.00 | 24789.00 | 24176.33 | 22150.00 |
| MSR1 | 31.00 | 27.00 | 27.00 | 28.33 | 36.00 |
| MSRA | 127.00 | 146.00 | 157.00 | 143.33 | 217.00 |
| MSRB1 | 160.00 | 185.00 | 179.00 | 174.67 | 201.00 |
| MSRB2 | 364.00 | 424.00 | 377.00 | 388.33 | 442.00 |
| MSRB3 | 816.00 | 778.00 | 947.00 | 847.00 | 906.00 |
| MSS51 | 326.00 | 349.00 | 364.00 | 346.33 | 383.00 |
| MST1 | 173.00 | 162.00 | 168.00 | 167.67 | 168.00 |
| MST1L | 17.00 | 14.00 | 18.00 | 16.33 | 10.00 |
| MST1P2 | 94.00 | 100.00 | 101.00 | 98.33 | 101.00 |
| MST1R | 4.00 | 1.00 | 1.00 | 2.00 | 3.00 |
| MSTN | 5.00 | 1.00 | 1.00 | 2.33 | 1.00 |
| MSTO1 | 84.00 | 96.00 | 98.00 | 92.67 | 93.00 |
| MSTO2P | 101.00 | 121.00 | 118.00 | 113.33 | 132.00 |
| MSX1 | 124.00 | 105.00 | 108.00 | 112.33 | 116.00 |
| MT1E | 2273.00 | 2165.00 | 2219.00 | 2219.00 | 3104.00 |
| MT1F | 3.00 | 0.00 | 1.00 | 1.33 | 0.00 |

| | | | | | |
|---|---|---|---|---|---|
| MT1X | 180.00 | 195.00 | 214.00 | 196.33 | 294.00 |
| MT2A | 6498.00 | 6391.00 | 6833.00 | 6574.00 | 5365.00 |
| MTA1 | 2591.00 | 2613.00 | 2862.00 | 2688.67 | 2937.00 |
| MTA2 | 2687.00 | 2765.00 | 2733.00 | 2728.33 | 3084.00 |
| MTA3 | 682.00 | 607.00 | 715.00 | 668.00 | 799.00 |
| MTAP | 2997.00 | 2694.00 | 3092.00 | 2927.67 | 3504.00 |
| MTBP | 544.00 | 509.00 | 526.00 | 526.33 | 667.00 |
| MTCH1 | 4997.00 | 5124.00 | 5316.00 | 5145.67 | 5443.00 |
| MTCH2 | 1639.00 | 1544.00 | 1619.00 | 1600.67 | 2193.00 |
| MTCL1 | 4150.00 | 4189.00 | 4608.00 | 4315.67 | 4185.00 |
| MTCP1 | 138.00 | 146.00 | 140.00 | 141.33 | 167.00 |
| MTDH | 7320.00 | 6797.00 | 7434.00 | 7183.67 | 7636.00 |
| MTERF1 | 349.00 | 365.00 | 363.00 | 359.00 | 406.00 |
| MTERF2 | 226.00 | 262.00 | 247.00 | 245.00 | 293.00 |
| MTERF3 | 884.00 | 910.00 | 975.00 | 923.00 | 1117.00 |
| MTERF4 | 1093.00 | 1092.00 | 1204.00 | 1129.67 | 1117.00 |
| MTF1 | 587.00 | 547.00 | 620.00 | 584.67 | 547.00 |
| MTF2 | 654.00 | 557.00 | 618.00 | 609.67 | 821.00 |
| MTFMT | 238.00 | 287.00 | 332.00 | 285.67 | 337.00 |
| MTFP1 | 396.00 | 377.00 | 357.00 | 376.67 | 525.00 |
| MTFR1 | 676.00 | 639.00 | 699.00 | 671.33 | 907.00 |
| MTFR1L | 803.00 | 701.00 | 836.00 | 780.00 | 858.00 |
| MTFR2 | 137.00 | 150.00 | 151.00 | 146.00 | 206.00 |
| MTG1 | 1539.00 | 1554.00 | 1560.00 | 1551.00 | 1673.00 |
| MTG2 | 1224.00 | 1364.00 | 1296.00 | 1294.67 | 1223.00 |
| MTHFD1 | 2961.00 | 2760.00 | 2911.00 | 2877.33 | 3564.00 |
| MTHFD1L | 3109.00 | 2958.00 | 3319.00 | 3128.67 | 3608.00 |
| MTHFD2 | 5442.00 | 4835.00 | 5582.00 | 5286.33 | 6915.00 |
| MTHFD2L | 129.00 | 107.00 | 118.00 | 118.00 | 169.00 |
| MTHFR | 219.00 | 210.00 | 249.00 | 226.00 | 218.00 |
| MTHFS | 27.00 | 32.00 | 38.00 | 32.33 | 56.00 |
| MTHFSD | 782.00 | 750.00 | 767.00 | 766.33 | 785.00 |
| MTIF2 | 1813.00 | 1743.00 | 1859.00 | 1805.00 | 2298.00 |
| MTIF3 | 375.00 | 405.00 | 440.00 | 406.67 | 513.00 |
| MTL5 | 104.00 | 106.00 | 81.00 | 97.00 | 108.00 |
| MTM1 | 179.00 | 182.00 | 204.00 | 188.33 | 171.00 |
| MTMR1 | 1599.00 | 1459.00 | 1688.00 | 1582.00 | 1886.00 |
| MTMR10 | 30.00 | 21.00 | 37.00 | 29.33 | 23.00 |
| MTMR11 | 325.00 | 311.00 | 303.00 | 313.00 | 356.00 |
| MTMR12 | 843.00 | 707.00 | 930.00 | 826.67 | 978.00 |
| MTMR14 | 725.00 | 707.00 | 709.00 | 713.67 | 823.00 |
| MTMR2 | 1839.00 | 1722.00 | 2067.00 | 1876.00 | 2171.00 |
| MTMR3 | 1077.00 | 1055.00 | 1174.00 | 1102.00 | 987.00 |
| MTMR4 | 1733.00 | 1603.00 | 1898.00 | 1744.67 | 1881.00 |
| MTMR6 | 722.00 | 516.00 | 688.00 | 642.00 | 736.00 |
| MTMR7 | 8.00 | 5.00 | 8.00 | 7.00 | 3.00 |
| MTMR9 | 518.00 | 489.00 | 488.00 | 498.33 | 484.00 |
| MTMR9LP | 163.00 | 169.00 | 187.00 | 173.00 | 76.00 |

| | | | | | |
|---|---|---|---|---|---|
| MTO1 | 933.00 | 960.00 | 1048.00 | 980.33 | 1125.00 |
| MTOR | 2533.00 | 2316.00 | 2489.00 | 2446.00 | 2263.00 |
| MTPAP | 841.00 | 855.00 | 929.00 | 875.00 | 1035.00 |
| MTR | 1232.00 | 1112.00 | 1220.00 | 1188.00 | 1228.00 |
| MTRF1 | 554.00 | 574.00 | 609.00 | 579.00 | 622.00 |
| MTRF1L | 397.00 | 292.00 | 376.00 | 355.00 | 349.00 |
| MTRNR2L2 | 12.00 | 10.00 | 17.00 | 13.00 | 26.00 |
| MTRR | 1387.00 | 1271.00 | 1490.00 | 1382.67 | 1254.00 |
| MTSS1 | 128.00 | 131.00 | 131.00 | 130.00 | 126.00 |
| MTSS1L | 1212.00 | 1288.00 | 1418.00 | 1306.00 | 942.00 |
| MTURN | 665.00 | 595.00 | 702.00 | 654.00 | 609.00 |
| MTUS1 | 30.00 | 20.00 | 29.00 | 26.33 | 11.00 |
| MTX1 | 272.00 | 313.00 | 294.00 | 293.00 | 351.00 |
| MTX2 | 946.00 | 942.00 | 999.00 | 962.33 | 1267.00 |
| MTX3 | 1147.00 | 1086.00 | 1303.00 | 1178.67 | 1193.00 |
| MUC1 | 91.00 | 57.00 | 81.00 | 76.33 | 65.00 |
| MUL1 | 814.00 | 823.00 | 744.00 | 793.67 | 712.00 |
| MUM1 | 1486.00 | 1407.00 | 1418.00 | 1437.00 | 1506.00 |
| MUM1L1 | 84.00 | 70.00 | 104.00 | 86.00 | 104.00 |
| MURC | 50.00 | 59.00 | 76.00 | 61.67 | 81.00 |
| MUS81 | 2049.00 | 2120.00 | 2069.00 | 2079.33 | 2454.00 |
| MUSK | 43.00 | 35.00 | 50.00 | 42.67 | 39.00 |
| MUSTN1 | 29.00 | 35.00 | 46.00 | 36.67 | 31.00 |
| MUT | 1111.00 | 1037.00 | 1196.00 | 1114.67 | 1219.00 |
| MUTYH | 830.00 | 994.00 | 965.00 | 929.67 | 1142.00 |
| MVB12A | 644.00 | 625.00 | 640.00 | 636.33 | 745.00 |
| MVB12B | 330.00 | 287.00 | 321.00 | 312.67 | 314.00 |
| MVD | 756.00 | 743.00 | 797.00 | 765.33 | 831.00 |
| MVK | 156.00 | 158.00 | 151.00 | 155.00 | 160.00 |
| MVP | 1958.00 | 1939.00 | 1988.00 | 1961.67 | 1950.00 |
| MX1 | 15.00 | 13.00 | 25.00 | 17.67 | 31.00 |
| MXD1 | 563.00 | 564.00 | 630.00 | 585.67 | 648.00 |
| MXD3 | 218.00 | 184.00 | 230.00 | 210.67 | 220.00 |
| MXD4 | 436.00 | 441.00 | 482.00 | 453.00 | 398.00 |
| MXI1 | 753.00 | 723.00 | 890.00 | 788.67 | 958.00 |
| MXRA7 | 1905.00 | 1782.00 | 1905.00 | 1864.00 | 2040.00 |
| MXRA8 | 6.00 | 1.00 | 0.00 | 2.33 | 2.00 |
| MYADM | 690.00 | 611.00 | 569.00 | 623.33 | 597.00 |
| MYB | 18.00 | 8.00 | 6.00 | 10.67 | 32.00 |
| MYBBP1A | 4758.00 | 4855.00 | 4901.00 | 4838.00 | 4560.00 |
| MYBL1 | 917.00 | 826.00 | 919.00 | 887.33 | 1130.00 |
| MYBL2 | 375.00 | 393.00 | 300.00 | 356.00 | 625.00 |
| MYC | 6156.00 | 6092.00 | 6415.00 | 6221.00 | 7550.00 |
| MYCBP | 7.00 | 5.00 | 7.00 | 6.33 | 10.00 |
| MYCBP2 | 2921.00 | 2753.00 | 3187.00 | 2953.67 | 2433.00 |
| MYCBPAP | 7.00 | 5.00 | 10.00 | 7.33 | 7.00 |
| MYCT1 | 238.00 | 231.00 | 193.00 | 220.67 | 241.00 |
| MYD88 | 368.00 | 354.00 | 377.00 | 366.33 | 383.00 |

| | | | | | |
|---|---|---|---|---|---|
| MYDGF | 2620.00 | 2701.00 | 2325.00 | 2548.67 | 2893.00 |
| MYEF2 | 1564.00 | 1452.00 | 1665.00 | 1560.33 | 1804.00 |
| MYEOV | 3460.00 | 3335.00 | 3610.00 | 3468.33 | 2255.00 |
| MYEOV2 | 380.00 | 356.00 | 374.00 | 370.00 | 496.00 |
| MYH10 | 1296.00 | 1153.00 | 1235.00 | 1228.00 | 1163.00 |
| MYH15 | 56.00 | 53.00 | 55.00 | 54.67 | 118.00 |
| MYH16 | 4.00 | 3.00 | 6.00 | 4.33 | 6.00 |
| MYH3 | 133.00 | 163.00 | 152.00 | 149.33 | 132.00 |
| MYH7B | 10.00 | 6.00 | 5.00 | 7.00 | 9.00 |
| MYH9 | 22354.00 | 20953.00 | 22210.00 | 21839.00 | 13558.00 |
| MYL12A | 5513.00 | 5386.00 | 6077.00 | 5658.67 | 7669.00 |
| MYL12B | 5146.00 | 4899.00 | 5268.00 | 5104.33 | 6939.00 |
| MYL5 | 59.00 | 69.00 | 73.00 | 67.00 | 71.00 |
| MYL6 | 3569.00 | 3604.00 | 3598.00 | 3590.33 | 4463.00 |
| MYL6B | 1361.00 | 1359.00 | 1463.00 | 1394.33 | 1513.00 |
| MYLIP | 137.00 | 121.00 | 157.00 | 138.33 | 115.00 |
| MYLK | 4120.00 | 3894.00 | 4144.00 | 4052.67 | 3190.00 |
| MYLK2 | 7.00 | 8.00 | 5.00 | 6.67 | 9.00 |
| MYLK4 | 60.00 | 48.00 | 62.00 | 56.67 | 41.00 |
| MYLK-AS1 | 33.00 | 43.00 | 36.00 | 37.33 | 42.00 |
| MYNN | 916.00 | 866.00 | 928.00 | 903.33 | 1041.00 |
| MYO10 | 8150.00 | 7747.00 | 9185.00 | 8360.67 | 7683.00 |
| MYO15A | 6.00 | 2.00 | 6.00 | 4.67 | 6.00 |
| MYO15B | 22.00 | 34.00 | 38.00 | 31.33 | 25.00 |
| MYO18A | 712.00 | 671.00 | 768.00 | 717.00 | 580.00 |
| MYO19 | 4.00 | 14.00 | 23.00 | 13.67 | 10.00 |
| MYO1A | 7.00 | 1.00 | 2.00 | 3.33 | 0.00 |
| MYO1B | 3019.00 | 2861.00 | 3305.00 | 3061.67 | 3091.00 |
| MYO1C | 2508.00 | 2445.00 | 2726.00 | 2559.67 | 2430.00 |
| MYO1D | 221.00 | 222.00 | 210.00 | 217.67 | 180.00 |
| MYO1E | 1048.00 | 964.00 | 1115.00 | 1042.33 | 878.00 |
| MYO1F | 6.00 | 3.00 | 3.00 | 4.00 | 4.00 |
| MYO5A | 2133.00 | 1992.00 | 2362.00 | 2162.33 | 1521.00 |
| MYO5B | 35.00 | 20.00 | 27.00 | 27.33 | 19.00 |
| MYO5C | 4.00 | 5.00 | 1.00 | 3.33 | 5.00 |
| MYO6 | 1546.00 | 1378.00 | 1654.00 | 1526.00 | 1398.00 |
| MYO7A | 31.00 | 28.00 | 25.00 | 28.00 | 21.00 |
| MYO9A | 466.00 | 396.00 | 490.00 | 450.67 | 299.00 |
| MYO9B | 3773.00 | 3870.00 | 3978.00 | 3873.67 | 3649.00 |
| MYOF | 39067.00 | 34732.00 | 41595.00 | 38464.67 | 39619.00 |
| MYOM1 | 28.00 | 45.00 | 45.00 | 39.33 | 23.00 |
| MYOZ3 | 8.00 | 1.00 | 9.00 | 6.00 | 3.00 |
| MYPN | 643.00 | 653.00 | 782.00 | 692.67 | 414.00 |
| MYPOP | 151.00 | 162.00 | 175.00 | 162.67 | 194.00 |
| MYRF | 207.00 | 221.00 | 205.00 | 211.00 | 50.00 |
| MYRFL | 7.00 | 11.00 | 15.00 | 11.00 | 21.00 |
| MYRIP | 81.00 | 82.00 | 101.00 | 88.00 | 92.00 |
| MYSM1 | 3009.00 | 2853.00 | 3299.00 | 3053.67 | 3066.00 |

| | | | | | |
|---|---|---|---|---|---|
| MZB1 | 4.00 | 1.00 | 4.00 | 3.00 | 0.00 |
| MZF1 | 853.00 | 835.00 | 991.00 | 893.00 | 808.00 |
| MZF1-AS1 | 123.00 | 117.00 | 105.00 | 115.00 | 119.00 |
| MZT1 | 477.00 | 444.00 | 520.00 | 480.33 | 672.00 |
| MZT2A | 461.00 | 463.00 | 452.00 | 458.67 | 596.00 |
| MZT2B | 1164.00 | 1130.00 | 1127.00 | 1140.33 | 1446.00 |
| N4BP1 | 872.00 | 806.00 | 906.00 | 861.33 | 982.00 |
| N4BP2 | 725.00 | 653.00 | 758.00 | 712.00 | 627.00 |
| N4BP2L1 | 30.00 | 18.00 | 31.00 | 26.33 | 24.00 |
| N4BP2L2 | 1848.00 | 1655.00 | 1999.00 | 1834.00 | 1959.00 |
| N4BP2L2-IT2 | 267.00 | 291.00 | 224.00 | 260.67 | 220.00 |
| N6AMT1 | 306.00 | 302.00 | 295.00 | 301.00 | 278.00 |
| N6AMT2 | 106.00 | 112.00 | 149.00 | 122.33 | 142.00 |
| NAA10 | 1494.00 | 1569.00 | 1680.00 | 1581.00 | 2160.00 |
| NAA15 | 1539.00 | 1332.00 | 1507.00 | 1459.33 | 1616.00 |
| NAA16 | 484.00 | 456.00 | 560.00 | 500.00 | 463.00 |
| NAA20 | 1681.00 | 1719.00 | 1722.00 | 1707.33 | 2171.00 |
| NAA25 | 1881.00 | 1713.00 | 1927.00 | 1840.33 | 1833.00 |
| NAA30 | 502.00 | 421.00 | 567.00 | 496.67 | 547.00 |
| NAA35 | 637.00 | 524.00 | 597.00 | 586.00 | 620.00 |
| NAA38 | 768.00 | 828.00 | 800.00 | 798.67 | 1119.00 |
| NAA40 | 1337.00 | 1273.00 | 1297.00 | 1302.33 | 1361.00 |
| NAA50 | 3588.00 | 3191.00 | 3890.00 | 3556.33 | 3857.00 |
| NAA60 | 571.00 | 584.00 | 579.00 | 578.00 | 621.00 |
| NAAA | 179.00 | 191.00 | 159.00 | 176.33 | 160.00 |
| NAALAD2 | 4.00 | 2.00 | 7.00 | 4.33 | 0.00 |
| NAALADL2 | 34.00 | 27.00 | 32.00 | 31.00 | 37.00 |
| NAB1 | 1230.00 | 1179.00 | 1316.00 | 1241.67 | 1517.00 |
| NAB2 | 680.00 | 655.00 | 738.00 | 691.00 | 1240.00 |
| NABP1 | 2638.00 | 2524.00 | 2853.00 | 2671.67 | 2139.00 |
| NABP2 | 678.00 | 633.00 | 736.00 | 682.33 | 798.00 |
| NACA | 10932.00 | 10813.00 | 11957.00 | 11234.00 | 14993.00 |
| NACC1 | 3043.00 | 2870.00 | 3130.00 | 3014.33 | 2889.00 |
| NACC2 | 1065.00 | 1021.00 | 1069.00 | 1051.67 | 934.00 |
| NADK | 1119.00 | 1105.00 | 1257.00 | 1160.33 | 1162.00 |
| NADK2 | 766.00 | 754.00 | 938.00 | 819.33 | 1007.00 |
| NADSYN1 | 8120.00 | 8594.00 | 8946.00 | 8553.33 | 7591.00 |
| NAE1 | 1777.00 | 1601.00 | 1867.00 | 1748.33 | 2195.00 |
| NAF1 | 304.00 | 229.00 | 278.00 | 270.33 | 296.00 |
| NAGA | 882.00 | 848.00 | 926.00 | 885.33 | 786.00 |
| NAGK | 525.00 | 532.00 | 543.00 | 533.33 | 513.00 |
| NAGLU | 1345.00 | 1427.00 | 1383.00 | 1385.00 | 1481.00 |
| NAGPA | 1009.00 | 1008.00 | 981.00 | 999.33 | 1001.00 |
| NAGPA-AS1 | 8.00 | 2.00 | 10.00 | 6.67 | 1.00 |
| NAGS | 89.00 | 78.00 | 72.00 | 79.67 | 69.00 |
| NAIF1 | 289.00 | 288.00 | 337.00 | 304.67 | 286.00 |
| NAIP | 9.00 | 6.00 | 8.00 | 7.67 | 13.00 |
| NALCN | 586.00 | 562.00 | 695.00 | 614.33 | 466.00 |

| | | | | | |
|---|---|---|---|---|---|
| NAMPT | 4547.00 | 4128.00 | 4804.00 | 4493.00 | 4679.00 |
| NANOS1 | 4.00 | 3.00 | 0.00 | 2.33 | 0.00 |
| NANP | 374.00 | 360.00 | 436.00 | 390.00 | 387.00 |
| NANS | 649.00 | 600.00 | 698.00 | 649.00 | 763.00 |
| NAP1L1 | 15044.00 | 14114.00 | 16531.00 | 15229.67 | 18121.00 |
| NAP1L2 | 33.00 | 27.00 | 22.00 | 27.33 | 41.00 |
| NAP1L4 | 68.00 | 54.00 | 70.00 | 64.00 | 75.00 |
| NAP1L5 | 192.00 | 196.00 | 214.00 | 200.67 | 257.00 |
| NAP1L6 | 9.00 | 3.00 | 11.00 | 7.67 | 8.00 |
| NAPA | 898.00 | 844.00 | 995.00 | 912.33 | 984.00 |
| NAPA-AS1 | 10.00 | 8.00 | 13.00 | 10.33 | 21.00 |
| NAPB | 547.00 | 511.00 | 582.00 | 546.67 | 516.00 |
| NAPEPLD | 14.00 | 4.00 | 10.00 | 9.33 | 6.00 |
| NAPG | 1311.00 | 1219.00 | 1377.00 | 1302.33 | 1410.00 |
| NAPSA | 12.00 | 15.00 | 9.00 | 12.00 | 11.00 |
| NARF | 739.00 | 714.00 | 803.00 | 752.00 | 808.00 |
| NARFL | 575.00 | 531.00 | 633.00 | 579.67 | 670.00 |
| NARS | 7349.00 | 6802.00 | 7423.00 | 7191.33 | 8000.00 |
| NARS2 | 540.00 | 442.00 | 528.00 | 503.33 | 522.00 |
| NASP | 2867.00 | 2761.00 | 2935.00 | 2854.33 | 3694.00 |
| NAT1 | 42.00 | 38.00 | 49.00 | 43.00 | 56.00 |
| NAT10 | 2467.00 | 2437.00 | 2609.00 | 2504.33 | 2437.00 |
| NAT14 | 556.00 | 671.00 | 578.00 | 601.67 | 634.00 |
| NAT16 | 4.00 | 1.00 | 0.00 | 1.67 | 0.00 |
| NAT6 | 204.00 | 180.00 | 230.00 | 204.67 | 221.00 |
| NAT8L | 1592.00 | 1667.00 | 1732.00 | 1663.67 | 1665.00 |
| NAT9 | 1974.00 | 1983.00 | 2006.00 | 1987.67 | 2126.00 |
| NATD1 | 18.00 | 19.00 | 19.00 | 18.67 | 20.00 |
| NAV1 | 4550.00 | 4184.00 | 4600.00 | 4444.67 | 2676.00 |
| NAV2 | 4538.00 | 4361.00 | 4901.00 | 4600.00 | 2973.00 |
| NAV3 | 3579.00 | 3170.00 | 3717.00 | 3488.67 | 3818.00 |
| NBAS | 1506.00 | 1494.00 | 1627.00 | 1542.33 | 1523.00 |
| NBEA | 465.00 | 394.00 | 465.00 | 441.33 | 354.00 |
| NBEAL1 | 89.00 | 76.00 | 96.00 | 87.00 | 84.00 |
| NBEAL2 | 2958.00 | 2993.00 | 3461.00 | 3137.33 | 2860.00 |
| NBL1 | 17.00 | 11.00 | 13.00 | 13.67 | 11.00 |
| NBN | 1671.00 | 1589.00 | 1785.00 | 1681.67 | 2056.00 |
| NBPF1 | 441.00 | 337.00 | 462.00 | 413.33 | 304.00 |
| NBPF10 | 98.00 | 56.00 | 75.00 | 76.33 | 60.00 |
| NBPF11 | 375.00 | 353.00 | 380.00 | 369.33 | 338.00 |
| NBPF12 | 495.00 | 479.00 | 531.00 | 501.67 | 433.00 |
| NBPF14 | 435.00 | 385.00 | 416.00 | 412.00 | 350.00 |
| NBPF15 | 2029.00 | 2057.00 | 2206.00 | 2097.33 | 2204.00 |
| NBPF20 | 206.00 | 184.00 | 218.00 | 202.67 | 236.00 |
| NBPF25P | 32.00 | 29.00 | 37.00 | 32.67 | 18.00 |
| NBPF3 | 306.00 | 314.00 | 318.00 | 312.67 | 287.00 |
| NBPF6 | 3.00 | 2.00 | 1.00 | 2.00 | 0.00 |
| NBPF7 | 3.00 | 1.00 | 7.00 | 3.67 | 0.00 |

| | | | | | |
|---|---|---|---|---|---|
| NBPF8 | 1358.00 | 1372.00 | 1459.00 | 1396.33 | 1561.00 |
| NBPF9 | 1065.00 | 964.00 | 1090.00 | 1039.67 | 1197.00 |
| NBR1 | 3554.00 | 3140.00 | 3639.00 | 3444.33 | 3111.00 |
| NBR2 | 280.00 | 273.00 | 325.00 | 292.67 | 238.00 |
| NCAM1 | 69.00 | 46.00 | 54.00 | 56.33 | 71.00 |
| NCAM2 | 275.00 | 267.00 | 353.00 | 298.33 | 445.00 |
| NCAPD2 | 5116.00 | 4704.00 | 4931.00 | 4917.00 | 5115.00 |
| NCAPD3 | 1049.00 | 975.00 | 1067.00 | 1030.33 | 942.00 |
| NCAPG | 782.00 | 687.00 | 763.00 | 744.00 | 1097.00 |
| NCAPG2 | 1831.00 | 1708.00 | 1721.00 | 1753.33 | 2160.00 |
| NCAPH | 616.00 | 551.00 | 587.00 | 584.67 | 781.00 |
| NCAPH2 | 671.00 | 719.00 | 796.00 | 728.67 | 914.00 |
| NCBP1 | 1728.00 | 1596.00 | 1873.00 | 1732.33 | 1800.00 |
| NCBP2 | 2959.00 | 2720.00 | 3126.00 | 2935.00 | 3770.00 |
| NCBP2-AS2 | 332.00 | 319.00 | 364.00 | 338.33 | 424.00 |
| NCDN | 1807.00 | 1778.00 | 1857.00 | 1814.00 | 1720.00 |
| NCEH1 | 2164.00 | 2038.00 | 2217.00 | 2139.67 | 3477.00 |
| NCF2 | 10.00 | 26.00 | 9.00 | 15.00 | 11.00 |
| NCK1 | 642.00 | 628.00 | 756.00 | 675.33 | 711.00 |
| NCK1-AS1 | 13.00 | 20.00 | 21.00 | 18.00 | 32.00 |
| NCK2 | 1022.00 | 992.00 | 1108.00 | 1040.67 | 931.00 |
| NCKAP1 | 4904.00 | 4204.00 | 5387.00 | 4831.67 | 5081.00 |
| NCKAP1L | 5.00 | 2.00 | 6.00 | 4.33 | 6.00 |
| NCKAP5 | 77.00 | 69.00 | 86.00 | 77.33 | 76.00 |
| NCKAP5L | 1604.00 | 1603.00 | 1577.00 | 1594.67 | 1305.00 |
| NCKIPSD | 760.00 | 720.00 | 782.00 | 754.00 | 666.00 |
| NCL | 21252.00 | 20768.00 | 22910.00 | 21643.33 | 25812.00 |
| NCLN | 3986.00 | 4076.00 | 4001.00 | 4021.00 | 4424.00 |
| NCOA1 | 450.00 | 389.00 | 471.00 | 436.67 | 369.00 |
| NCOA2 | 538.00 | 428.00 | 647.00 | 537.67 | 400.00 |
| NCOA3 | 1422.00 | 1285.00 | 1457.00 | 1388.00 | 1117.00 |
| NCOA4 | 3457.00 | 3061.00 | 3716.00 | 3411.33 | 3487.00 |
| NCOA5 | 1485.00 | 1485.00 | 1638.00 | 1536.00 | 1581.00 |
| NCOA6 | 1681.00 | 1569.00 | 1867.00 | 1705.67 | 1372.00 |
| NCOA7 | 828.00 | 694.00 | 885.00 | 802.33 | 667.00 |
| NCOR1 | 2463.00 | 2173.00 | 2457.00 | 2364.33 | 2066.00 |
| NCOR1P1 | 4.00 | 6.00 | 4.00 | 4.67 | 4.00 |
| NCOR2 | 8887.00 | 8612.00 | 9297.00 | 8932.00 | 7215.00 |
| NCR3LG1 | 710.00 | 671.00 | 757.00 | 712.67 | 485.00 |
| NCRNA00250 | 3.00 | 5.00 | 0.00 | 2.67 | 4.00 |
| NCS1 | 2266.00 | 2230.00 | 2284.00 | 2260.00 | 2178.00 |
| NCSTN | 6451.00 | 6216.00 | 6367.00 | 6344.67 | 5504.00 |
| NDC1 | 2790.00 | 2560.00 | 2637.00 | 2662.33 | 3286.00 |
| NDC80 | 812.00 | 847.00 | 860.00 | 839.67 | 1226.00 |
| NDEL1 | 1302.00 | 1327.00 | 1382.00 | 1337.00 | 1320.00 |
| NDFIP1 | 1891.00 | 1664.00 | 1794.00 | 1783.00 | 2204.00 |
| NDFIP2 | 1113.00 | 901.00 | 1115.00 | 1043.00 | 1031.00 |
| NDNF | 87.00 | 92.00 | 82.00 | 87.00 | 63.00 |

| | | | | | |
|---|---|---|---|---|---|
| NDOR1 | 2932.00 | 2924.00 | 3065.00 | 2973.67 | 2710.00 |
| NDP | 8.00 | 6.00 | 6.00 | 6.67 | 2.00 |
| NDRG1 | 464.00 | 469.00 | 443.00 | 458.67 | 403.00 |
| NDRG3 | 932.00 | 1016.00 | 1069.00 | 1005.67 | 1057.00 |
| NDRG4 | 18.00 | 15.00 | 8.00 | 13.67 | 15.00 |
| NDST1 | 4886.00 | 4539.00 | 4719.00 | 4714.67 | 3136.00 |
| NDST2 | 1036.00 | 1043.00 | 1145.00 | 1074.67 | 864.00 |
| NDST3 | 5.00 | 11.00 | 4.00 | 6.67 | 0.00 |
| NDUFA1 | 537.00 | 600.00 | 563.00 | 566.67 | 734.00 |
| NDUFA10 | 1379.00 | 1418.00 | 1502.00 | 1433.00 | 1635.00 |
| NDUFA11 | 1009.00 | 1040.00 | 1083.00 | 1044.00 | 1234.00 |
| NDUFA12 | 557.00 | 509.00 | 500.00 | 522.00 | 735.00 |
| NDUFA13 | 1356.00 | 1445.00 | 1533.00 | 1444.67 | 1918.00 |
| NDUFA2 | 380.00 | 365.00 | 461.00 | 402.00 | 506.00 |
| NDUFA4 | 1846.00 | 1738.00 | 1763.00 | 1782.33 | 2473.00 |
| NDUFA4L2 | 61.00 | 57.00 | 61.00 | 59.67 | 58.00 |
| NDUFA5 | 739.00 | 699.00 | 764.00 | 734.00 | 918.00 |
| NDUFA7 | 615.00 | 556.00 | 645.00 | 605.33 | 978.00 |
| NDUFA8 | 569.00 | 581.00 | 610.00 | 586.67 | 812.00 |
| NDUFA9 | 1521.00 | 1418.00 | 1546.00 | 1495.00 | 1894.00 |
| NDUFAB1 | 1053.00 | 1000.00 | 1079.00 | 1044.00 | 1512.00 |
| NDUFAF1 | 298.00 | 311.00 | 280.00 | 296.33 | 344.00 |
| NDUFAF2 | 542.00 | 521.00 | 532.00 | 531.67 | 745.00 |
| NDUFAF3 | 339.00 | 358.00 | 368.00 | 355.00 | 466.00 |
| NDUFAF4 | 656.00 | 629.00 | 696.00 | 660.33 | 821.00 |
| NDUFAF5 | 243.00 | 213.00 | 232.00 | 229.33 | 320.00 |
| NDUFAF6 | 359.00 | 356.00 | 399.00 | 371.33 | 406.00 |
| NDUFAF7 | 627.00 | 670.00 | 642.00 | 646.33 | 767.00 |
| NDUFB1 | 323.00 | 343.00 | 304.00 | 323.33 | 488.00 |
| NDUFB10 | 1129.00 | 1063.00 | 1113.00 | 1101.67 | 1340.00 |
| NDUFB11 | 849.00 | 804.00 | 795.00 | 816.00 | 1086.00 |
| NDUFB2 | 1107.00 | 1071.00 | 1119.00 | 1099.00 | 1605.00 |
| NDUFB2-AS1 | 33.00 | 46.00 | 33.00 | 37.33 | 40.00 |
| NDUFB3 | 593.00 | 608.00 | 611.00 | 604.00 | 789.00 |
| NDUFB4 | 1234.00 | 1209.00 | 1289.00 | 1244.00 | 1647.00 |
| NDUFB5 | 932.00 | 853.00 | 943.00 | 909.33 | 1308.00 |
| NDUFB6 | 747.00 | 743.00 | 720.00 | 736.67 | 1077.00 |
| NDUFB7 | 902.00 | 965.00 | 955.00 | 940.67 | 1240.00 |
| NDUFB8 | 1744.00 | 1699.00 | 1777.00 | 1740.00 | 2367.00 |
| NDUFB9 | 3596.00 | 3826.00 | 3723.00 | 3715.00 | 4417.00 |
| NDUFC1 | 299.00 | 304.00 | 300.00 | 301.00 | 475.00 |
| NDUFC2 | 277.00 | 271.00 | 280.00 | 276.00 | 331.00 |
| NDUFS1 | 1581.00 | 1433.00 | 1635.00 | 1549.67 | 1891.00 |
| NDUFS2 | 3235.00 | 3122.00 | 3303.00 | 3220.00 | 3992.00 |
| NDUFS3 | 820.00 | 844.00 | 829.00 | 831.00 | 1118.00 |
| NDUFS4 | 694.00 | 672.00 | 747.00 | 704.33 | 947.00 |
| NDUFS5 | 1429.00 | 1321.00 | 1479.00 | 1409.67 | 2107.00 |
| NDUFS6 | 824.00 | 777.00 | 873.00 | 824.67 | 1199.00 |

| | | | | | |
|---|---|---|---|---|---|
| NDUFS7 | 721.00 | 847.00 | 875.00 | 814.33 | 1107.00 |
| NDUFS8 | 824.00 | 874.00 | 910.00 | 869.33 | 1041.00 |
| NDUFV1 | 2785.00 | 2825.00 | 2859.00 | 2823.00 | 3296.00 |
| NDUFV2 | 1798.00 | 1751.00 | 1908.00 | 1819.00 | 2724.00 |
| NDUFV2-AS1 | 12.00 | 18.00 | 18.00 | 16.00 | 9.00 |
| NDUFV3 | 591.00 | 622.00 | 649.00 | 620.67 | 659.00 |
| NEAT1 | 19490.00 | 19969.00 | 19585.00 | 19681.33 | 18060.00 |
| NEB | 10.00 | 12.00 | 12.00 | 11.33 | 14.00 |
| NEBL | 4.00 | 4.00 | 5.00 | 4.33 | 4.00 |
| NECAB2 | 38.00 | 19.00 | 22.00 | 26.33 | 44.00 |
| NECAB3 | 556.00 | 581.00 | 604.00 | 580.33 | 642.00 |
| NECAP1 | 811.00 | 860.00 | 876.00 | 849.00 | 1013.00 |
| NECAP2 | 1033.00 | 945.00 | 1037.00 | 1005.00 | 1147.00 |
| NEDD1 | 661.00 | 613.00 | 725.00 | 666.33 | 804.00 |
| NEDD4 | 2646.00 | 2342.00 | 2664.00 | 2550.67 | 2785.00 |
| NEDD4L | 2070.00 | 1889.00 | 2137.00 | 2032.00 | 1793.00 |
| NEDD8 | 942.00 | 949.00 | 968.00 | 953.00 | 1284.00 |
| NEDD9 | 3.00 | 1.00 | 3.00 | 2.33 | 3.00 |
| NEFL | 257.00 | 230.00 | 297.00 | 261.33 | 327.00 |
| NEGR1 | 227.00 | 196.00 | 257.00 | 226.67 | 190.00 |
| NEIL1 | 46.00 | 61.00 | 59.00 | 55.33 | 69.00 |
| NEIL2 | 424.00 | 419.00 | 394.00 | 412.33 | 552.00 |
| NEIL3 | 107.00 | 82.00 | 106.00 | 98.33 | 149.00 |
| NEK1 | 407.00 | 371.00 | 434.00 | 404.00 | 355.00 |
| NEK10 | 171.00 | 171.00 | 196.00 | 179.33 | 146.00 |
| NEK11 | 101.00 | 107.00 | 90.00 | 99.33 | 103.00 |
| NEK2 | 505.00 | 436.00 | 478.00 | 473.00 | 593.00 |
| NEK3 | 388.00 | 363.00 | 393.00 | 381.33 | 554.00 |
| NEK4 | 479.00 | 525.00 | 583.00 | 529.00 | 538.00 |
| NEK5 | 17.00 | 14.00 | 16.00 | 15.67 | 20.00 |
| NEK6 | 1748.00 | 1634.00 | 1758.00 | 1713.33 | 1375.00 |
| NEK7 | 1014.00 | 875.00 | 1122.00 | 1003.67 | 1182.00 |
| NEK8 | 118.00 | 126.00 | 133.00 | 125.67 | 85.00 |
| NEK9 | 4558.00 | 4188.00 | 4710.00 | 4485.33 | 3880.00 |
| NELFA | 1433.00 | 1509.00 | 1546.00 | 1496.00 | 1684.00 |
| NELFB | 1520.00 | 1590.00 | 1713.00 | 1607.67 | 1816.00 |
| NELFCD | 2521.00 | 2489.00 | 2615.00 | 2541.67 | 2796.00 |
| NELL2 | 101.00 | 134.00 | 131.00 | 122.00 | 113.00 |
| NEMF | 1682.00 | 1670.00 | 1854.00 | 1735.33 | 1835.00 |
| NEMP1 | 2174.00 | 2093.00 | 2188.00 | 2151.67 | 2425.00 |
| NEMP2 | 372.00 | 364.00 | 316.00 | 350.67 | 572.00 |
| NENF | 564.00 | 647.00 | 625.00 | 612.00 | 869.00 |
| NEO1 | 624.00 | 648.00 | 709.00 | 660.33 | 495.00 |
| NES | 93.00 | 97.00 | 103.00 | 97.67 | 110.00 |
| NET1 | 1790.00 | 1827.00 | 1864.00 | 1827.00 | 2509.00 |
| NETO2 | 2195.00 | 2010.00 | 2399.00 | 2201.33 | 1974.00 |
| NEU3 | 380.00 | 347.00 | 412.00 | 379.67 | 296.00 |
| NEURL2 | 244.00 | 223.00 | 231.00 | 232.67 | 199.00 |

| | | | | | |
|---|---|---|---|---|---|
| NEURL4 | 2409.00 | 2309.00 | 2446.00 | 2388.00 | 2331.00 |
| NEUROG2 | 15.00 | 8.00 | 7.00 | 10.00 | 3.00 |
| NEXN | 983.00 | 995.00 | 1062.00 | 1013.33 | 1028.00 |
| NEXN-AS1 | 73.00 | 72.00 | 71.00 | 72.00 | 97.00 |
| NF1 | 2324.00 | 2022.00 | 2586.00 | 2310.67 | 2074.00 |
| NF2 | 3023.00 | 2883.00 | 3125.00 | 3010.33 | 3918.00 |
| NFAT5 | 2879.00 | 2692.00 | 3153.00 | 2908.00 | 2376.00 |
| NFATC2 | 28.00 | 18.00 | 29.00 | 25.00 | 20.00 |
| NFATC2IP | 2276.00 | 2399.00 | 2411.00 | 2362.00 | 2404.00 |
| NFATC3 | 635.00 | 560.00 | 682.00 | 625.67 | 492.00 |
| NFATC4 | 844.00 | 921.00 | 927.00 | 897.33 | 918.00 |
| NFE2L1 | 11525.00 | 11532.00 | 11692.00 | 11583.00 | 10206.00 |
| NFE2L2 | 2521.00 | 2362.00 | 2749.00 | 2544.00 | 2765.00 |
| NFE2L3 | 1804.00 | 1868.00 | 1863.00 | 1845.00 | 3350.00 |
| NFIA | 215.00 | 224.00 | 269.00 | 236.00 | 328.00 |
| NFIB | 1807.00 | 1664.00 | 2021.00 | 1830.67 | 1893.00 |
| NFIC | 1455.00 | 1476.00 | 1650.00 | 1527.00 | 1311.00 |
| NFIL3 | 475.00 | 464.00 | 502.00 | 480.33 | 477.00 |
| NFIX | 503.00 | 455.00 | 515.00 | 491.00 | 486.00 |
| NFKB1 | 1057.00 | 1055.00 | 1141.00 | 1084.33 | 1242.00 |
| NFKB2 | 2004.00 | 1934.00 | 1997.00 | 1978.33 | 2857.00 |
| NFKBIA | 570.00 | 483.00 | 594.00 | 549.00 | 776.00 |
| NFKBIB | 458.00 | 494.00 | 507.00 | 486.33 | 688.00 |
| NFKBID | 111.00 | 131.00 | 121.00 | 121.00 | 111.00 |
| NFKBIE | 214.00 | 245.00 | 294.00 | 251.00 | 281.00 |
| NFKBIL1 | 28.00 | 35.00 | 30.00 | 31.00 | 25.00 |
| NFKBIZ | 1442.00 | 1347.00 | 1527.00 | 1438.67 | 1305.00 |
| NFRKB | 785.00 | 741.00 | 757.00 | 761.00 | 660.00 |
| NFS1 | 887.00 | 807.00 | 897.00 | 863.67 | 943.00 |
| NFU1 | 363.00 | 333.00 | 438.00 | 378.00 | 530.00 |
| NFX1 | 1782.00 | 1743.00 | 1878.00 | 1801.00 | 1813.00 |
| NFXL1 | 704.00 | 644.00 | 700.00 | 682.67 | 673.00 |
| NFYA | 2164.00 | 2156.00 | 2468.00 | 2262.67 | 2034.00 |
| NFYB | 398.00 | 364.00 | 444.00 | 402.00 | 515.00 |
| NFYC | 1011.00 | 931.00 | 1029.00 | 990.33 | 1179.00 |
| NFYC-AS1 | 327.00 | 306.00 | 315.00 | 316.00 | 300.00 |
| NGDN | 641.00 | 653.00 | 666.00 | 653.33 | 831.00 |
| NGF | 182.00 | 204.00 | 189.00 | 191.67 | 88.00 |
| NGFR | 590.00 | 581.00 | 584.00 | 585.00 | 816.00 |
| NGFRAP1 | 3424.00 | 3396.00 | 3830.00 | 3550.00 | 4831.00 |
| NGLY1 | 861.00 | 875.00 | 1044.00 | 926.67 | 971.00 |
| NGRN | 1653.00 | 1548.00 | 1748.00 | 1649.67 | 2105.00 |
| NHEJ1 | 290.00 | 295.00 | 331.00 | 305.33 | 330.00 |
| NHLH1 | 5.00 | 5.00 | 6.00 | 5.33 | 5.00 |
| NHLH2 | 3.00 | 2.00 | 12.00 | 5.67 | 5.00 |
| NHLRC1 | 11.00 | 13.00 | 10.00 | 11.33 | 22.00 |
| NHLRC2 | 413.00 | 365.00 | 411.00 | 396.33 | 367.00 |
| NHLRC3 | 745.00 | 733.00 | 784.00 | 754.00 | 766.00 |

| | | | | | |
|---|---|---|---|---|---|
| NHLRC4 | 8.00 | 11.00 | 16.00 | 11.67 | 14.00 |
| NHP2 | 1554.00 | 1574.00 | 1788.00 | 1638.67 | 2086.00 |
| NHS | 176.00 | 182.00 | 234.00 | 197.33 | 78.00 |
| NHSL1 | 439.00 | 403.00 | 467.00 | 436.33 | 493.00 |
| NHSL2 | 11.00 | 9.00 | 9.00 | 9.67 | 4.00 |
| NICN1 | 773.00 | 821.00 | 849.00 | 814.33 | 662.00 |
| NID1 | 197.00 | 195.00 | 205.00 | 199.00 | 233.00 |
| NID2 | 7.00 | 4.00 | 12.00 | 7.67 | 7.00 |
| NIF3L1 | 905.00 | 919.00 | 1080.00 | 968.00 | 1097.00 |
| NIFK | 1584.00 | 1410.00 | 1689.00 | 1561.00 | 1994.00 |
| NIFK-AS1 | 82.00 | 65.00 | 65.00 | 70.67 | 100.00 |
| NIM1K | 23.00 | 20.00 | 34.00 | 25.67 | 18.00 |
| NIN | 1536.00 | 1460.00 | 1711.00 | 1569.00 | 1595.00 |
| NINJ1 | 377.00 | 368.00 | 378.00 | 374.33 | 415.00 |
| NINL | 904.00 | 977.00 | 1127.00 | 1002.67 | 926.00 |
| NIP7 | 1227.00 | 1219.00 | 1253.00 | 1233.00 | 1637.00 |
| NIPA1 | 1096.00 | 1010.00 | 1158.00 | 1088.00 | 1255.00 |
| NIPA2 | 1912.00 | 1759.00 | 2021.00 | 1897.33 | 2293.00 |
| NIPAL1 | 248.00 | 218.00 | 275.00 | 247.00 | 164.00 |
| NIPAL2 | 148.00 | 131.00 | 118.00 | 132.33 | 120.00 |
| NIPAL3 | 697.00 | 613.00 | 670.00 | 660.00 | 550.00 |
| NIPAL4 | 128.00 | 103.00 | 107.00 | 112.67 | 79.00 |
| NIPBL | 2208.00 | 2137.00 | 2338.00 | 2227.67 | 2024.00 |
| NIPBL-AS1 | 460.00 | 428.00 | 469.00 | 452.33 | 398.00 |
| NIPSNAP1 | 1215.00 | 1179.00 | 1246.00 | 1213.33 | 1749.00 |
| NIPSNAP3A | 238.00 | 192.00 | 244.00 | 224.67 | 226.00 |
| NIPSNAP3B | 44.00 | 64.00 | 64.00 | 57.33 | 53.00 |
| NISCH | 3388.00 | 3433.00 | 3616.00 | 3479.00 | 2896.00 |
| NIT1 | 782.00 | 709.00 | 768.00 | 753.00 | 846.00 |
| NIT2 | 1093.00 | 1054.00 | 1158.00 | 1101.67 | 1436.00 |
| NKAP | 376.00 | 340.00 | 363.00 | 359.67 | 420.00 |
| NKILA | 14.00 | 13.00 | 12.00 | 13.00 | 7.00 |
| NKIRAS1 | 410.00 | 394.00 | 423.00 | 409.00 | 500.00 |
| NKIRAS2 | 703.00 | 680.00 | 744.00 | 709.00 | 668.00 |
| NKRF | 466.00 | 432.00 | 457.00 | 451.67 | 507.00 |
| NKTR | 4611.00 | 4457.00 | 4884.00 | 4650.67 | 4158.00 |
| NKX2-1 | 16.00 | 28.00 | 30.00 | 24.67 | 18.00 |
| NKX2-4 | 93.00 | 76.00 | 67.00 | 78.67 | 108.00 |
| NKX2-6 | 132.00 | 146.00 | 167.00 | 148.33 | 191.00 |
| NKX2-8 | 12.00 | 21.00 | 21.00 | 18.00 | 20.00 |
| NKX3-1 | 109.00 | 129.00 | 136.00 | 124.67 | 105.00 |
| NKX3-2 | 20.00 | 19.00 | 28.00 | 22.33 | 29.00 |
| NKX6-1 | 106.00 | 129.00 | 135.00 | 123.33 | 103.00 |
| NLE1 | 888.00 | 778.00 | 851.00 | 839.00 | 994.00 |
| NLGN1 | 80.00 | 57.00 | 67.00 | 68.00 | 80.00 |
| NLGN2 | 3028.00 | 3236.00 | 3435.00 | 3233.00 | 2991.00 |
| NLGN3 | 10.00 | 11.00 | 9.00 | 10.00 | 10.00 |
| NLK | 775.00 | 730.00 | 844.00 | 783.00 | 585.00 |

| | | | | | |
|---|---|---|---|---|---|
| NLN | 1222.00 | 1145.00 | 1315.00 | 1227.33 | 1428.00 |
| NLRC5 | 420.00 | 405.00 | 478.00 | 434.33 | 344.00 |
| NLRP1 | 2772.00 | 2816.00 | 3026.00 | 2871.33 | 2984.00 |
| NLRP12 | 3.00 | 4.00 | 3.00 | 3.33 | 2.00 |
| NLRP14 | 7.00 | 2.00 | 4.00 | 4.33 | 2.00 |
| NLRP3 | 730.00 | 662.00 | 648.00 | 680.00 | 1037.00 |
| NLRX1 | 414.00 | 407.00 | 400.00 | 407.00 | 452.00 |
| NMB | 197.00 | 183.00 | 196.00 | 192.00 | 257.00 |
| NMBR | 4.00 | 2.00 | 1.00 | 2.33 | 2.00 |
| NMD3 | 2157.00 | 1860.00 | 2156.00 | 2057.67 | 2503.00 |
| NME1 | 1600.00 | 1421.00 | 1610.00 | 1543.67 | 2270.00 |
| NME2 | 511.00 | 522.00 | 569.00 | 534.00 | 675.00 |
| NME3 | 571.00 | 630.00 | 580.00 | 593.67 | 709.00 |
| NME4 | 1446.00 | 1629.00 | 1624.00 | 1566.33 | 2062.00 |
| NME6 | 431.00 | 386.00 | 459.00 | 425.33 | 461.00 |
| NME7 | 556.00 | 542.00 | 569.00 | 555.67 | 625.00 |
| NMI | 228.00 | 210.00 | 258.00 | 232.00 | 273.00 |
| NMNAT1 | 363.00 | 328.00 | 351.00 | 347.33 | 409.00 |
| NMNAT2 | 1703.00 | 1672.00 | 1720.00 | 1698.33 | 2584.00 |
| NMRK1 | 477.00 | 461.00 | 484.00 | 474.00 | 549.00 |
| NMT1 | 2748.00 | 2495.00 | 2844.00 | 2695.67 | 2836.00 |
| NMT2 | 3871.00 | 3523.00 | 4115.00 | 3836.33 | 2825.00 |
| NMU | 7.00 | 11.00 | 3.00 | 7.00 | 24.00 |
| NNMT | 111.00 | 130.00 | 120.00 | 120.33 | 150.00 |
| NNT | 1660.00 | 1438.00 | 1645.00 | 1581.00 | 1404.00 |
| NNT-AS1 | 443.00 | 399.00 | 570.00 | 470.67 | 394.00 |
| NOA1 | 541.00 | 486.00 | 588.00 | 538.33 | 590.00 |
| NOB1 | 1803.00 | 1755.00 | 1975.00 | 1844.33 | 2166.00 |
| NOC2L | 3882.00 | 3790.00 | 3936.00 | 3869.33 | 4281.00 |
| NOC3L | 2490.00 | 2274.00 | 2493.00 | 2419.00 | 2785.00 |
| NOC4L | 602.00 | 558.00 | 644.00 | 601.33 | 707.00 |
| NOCT | 118.00 | 80.00 | 137.00 | 111.67 | 110.00 |
| NOD1 | 257.00 | 233.00 | 280.00 | 256.67 | 313.00 |
| NOG | 15.00 | 23.00 | 23.00 | 20.33 | 18.00 |
| NOL10 | 998.00 | 941.00 | 1095.00 | 1011.33 | 1304.00 |
| NOL11 | 1547.00 | 1506.00 | 1671.00 | 1574.67 | 1841.00 |
| NOL12 | 731.00 | 763.00 | 758.00 | 750.67 | 814.00 |
| NOL3 | 870.00 | 842.00 | 885.00 | 865.67 | 804.00 |
| NOL4 | 7.00 | 12.00 | 20.00 | 13.00 | 4.00 |
| NOL4L | 1920.00 | 1903.00 | 2113.00 | 1978.67 | 1927.00 |
| NOL6 | 3483.00 | 3488.00 | 3590.00 | 3520.33 | 3791.00 |
| NOL7 | 1403.00 | 1272.00 | 1311.00 | 1328.67 | 1827.00 |
| NOL8 | 1584.00 | 1535.00 | 1677.00 | 1598.67 | 1661.00 |
| NOL9 | 845.00 | 762.00 | 668.00 | 758.33 | 955.00 |
| NOLC1 | 5978.00 | 5488.00 | 6210.00 | 5892.00 | 6743.00 |
| NOM1 | 1386.00 | 1265.00 | 1441.00 | 1364.00 | 1263.00 |
| NOMO1 | 178.00 | 202.00 | 222.00 | 200.67 | 186.00 |
| NOMO2 | 733.00 | 622.00 | 684.00 | 679.67 | 529.00 |

| | | | | | |
|---|---|---|---|---|---|
| NONO | 11133.00 | 10453.00 | 11239.00 | 10941.67 | 11917.00 |
| NOP10 | 1254.00 | 1239.00 | 1320.00 | 1271.00 | 2058.00 |
| NOP14 | 5605.00 | 5427.00 | 6029.00 | 5687.00 | 5957.00 |
| NOP14-AS1 | 269.00 | 291.00 | 297.00 | 285.67 | 338.00 |
| NOP16 | 1267.00 | 1221.00 | 1343.00 | 1277.00 | 1628.00 |
| NOP2 | 3307.00 | 3171.00 | 3654.00 | 3377.33 | 3551.00 |
| NOP56 | 7464.00 | 7561.00 | 7751.00 | 7592.00 | 9016.00 |
| NOP58 | 3873.00 | 3722.00 | 3936.00 | 3843.67 | 4868.00 |
| NOP9 | 2659.00 | 2326.00 | 2776.00 | 2587.00 | 1465.00 |
| NOS1AP | 20.00 | 15.00 | 14.00 | 16.33 | 11.00 |
| NOS3 | 185.00 | 185.00 | 170.00 | 180.00 | 166.00 |
| NOSIP | 1229.00 | 1133.00 | 1184.00 | 1182.00 | 1565.00 |
| NOTCH1 | 1836.00 | 1911.00 | 2065.00 | 1937.33 | 1445.00 |
| NOTCH2 | 14010.00 | 12484.00 | 14420.00 | 13638.00 | 10099.00 |
| NOTCH2NL | 160.00 | 139.00 | 114.00 | 137.67 | 129.00 |
| NOTCH3 | 6.00 | 1.00 | 4.00 | 3.67 | 7.00 |
| NOV | 324.00 | 295.00 | 296.00 | 305.00 | 195.00 |
| NOVA1 | 3.00 | 6.00 | 2.00 | 3.67 | 3.00 |
| NOX3 | 3.00 | 0.00 | 0.00 | 1.00 | 0.00 |
| NOX4 | 132.00 | 136.00 | 152.00 | 140.00 | 172.00 |
| NOX5 | 26.00 | 14.00 | 25.00 | 21.67 | 18.00 |
| NOXA1 | 82.00 | 75.00 | 92.00 | 83.00 | 95.00 |
| NOXO1 | 4.00 | 4.00 | 2.00 | 3.33 | 4.00 |
| NOXRED1 | 20.00 | 22.00 | 18.00 | 20.00 | 25.00 |
| NPAS1 | 10.00 | 16.00 | 21.00 | 15.67 | 10.00 |
| NPAS2 | 6783.00 | 6582.00 | 6988.00 | 6784.33 | 8315.00 |
| NPAS3 | 102.00 | 73.00 | 95.00 | 90.00 | 95.00 |
| NPAT | 373.00 | 342.00 | 428.00 | 381.00 | 348.00 |
| NPC1 | 4834.00 | 4729.00 | 5185.00 | 4916.00 | 4188.00 |
| NPC1L1 | 3.00 | 4.00 | 4.00 | 3.67 | 4.00 |
| NPC2 | 734.00 | 745.00 | 650.00 | 709.67 | 602.00 |
| NPDC1 | 3834.00 | 3865.00 | 4076.00 | 3925.00 | 4058.00 |
| NPEPL1 | 28.00 | 36.00 | 42.00 | 35.33 | 42.00 |
| NPEPPS | 2060.00 | 1962.00 | 2149.00 | 2057.00 | 2223.00 |
| NPFF | 63.00 | 83.00 | 59.00 | 68.33 | 86.00 |
| NPFFR2 | 12.00 | 18.00 | 19.00 | 16.33 | 48.00 |
| NPHP1 | 174.00 | 172.00 | 162.00 | 169.33 | 157.00 |
| NPHP3 | 394.00 | 356.00 | 439.00 | 396.33 | 371.00 |
| NPHP3-ACAD11 | 210.00 | 172.00 | 261.00 | 214.33 | 248.00 |
| NPHP4 | 704.00 | 682.00 | 631.00 | 672.33 | 549.00 |
| NPIPA1 | 26.00 | 31.00 | 18.00 | 25.00 | 41.00 |
| NPIPB11 | 88.00 | 89.00 | 107.00 | 94.67 | 75.00 |
| NPIPB15 | 56.00 | 71.00 | 75.00 | 67.33 | 66.00 |
| NPIPB3 | 96.00 | 99.00 | 116.00 | 103.67 | 88.00 |
| NPIPB4 | 188.00 | 169.00 | 160.00 | 172.33 | 150.00 |
| NPIPB5 | 371.00 | 445.00 | 460.00 | 425.33 | 475.00 |
| NPIPB6 | 7.00 | 6.00 | 4.00 | 5.67 | 2.00 |
| NPIPB8 | 14.00 | 21.00 | 17.00 | 17.33 | 19.00 |

| | | | | | |
|---|---|---|---|---|---|
| NPIPB9 | 15.00 | 18.00 | 18.00 | 17.00 | 6.00 |
| NPL | 164.00 | 162.00 | 167.00 | 164.33 | 197.00 |
| NPLOC4 | 3951.00 | 3860.00 | 4128.00 | 3979.67 | 3663.00 |
| NPM1 | 22549.00 | 21879.00 | 24628.00 | 23018.67 | 31981.00 |
| NPM3 | 1099.00 | 1080.00 | 1226.00 | 1135.00 | 1579.00 |
| NPPA-AS1 | 83.00 | 67.00 | 89.00 | 79.67 | 109.00 |
| NPR2 | 622.00 | 602.00 | 675.00 | 633.00 | 572.00 |
| NPR3 | 302.00 | 260.00 | 332.00 | 298.00 | 285.00 |
| NPRL2 | 625.00 | 577.00 | 674.00 | 625.33 | 688.00 |
| NPRL3 | 1346.00 | 1255.00 | 1343.00 | 1314.67 | 1673.00 |
| NPTN | 1939.00 | 1718.00 | 1701.00 | 1786.00 | 1861.00 |
| NPTN-IT1 | 164.00 | 146.00 | 145.00 | 151.67 | 91.00 |
| NPTXR | 16.00 | 17.00 | 23.00 | 18.67 | 5.00 |
| NPW | 78.00 | 87.00 | 89.00 | 84.67 | 99.00 |
| NPY1R | 94.00 | 101.00 | 107.00 | 100.67 | 113.00 |
| NPY5R | 4.00 | 4.00 | 7.00 | 5.00 | 10.00 |
| NQO1 | 2622.00 | 2559.00 | 2492.00 | 2557.67 | 2363.00 |
| NQO2 | 410.00 | 393.00 | 358.00 | 387.00 | 467.00 |
| NR1D1 | 242.00 | 273.00 | 243.00 | 252.67 | 315.00 |
| NR1D2 | 1244.00 | 1172.00 | 1527.00 | 1314.33 | 1550.00 |
| NR1H2 | 971.00 | 921.00 | 1000.00 | 964.00 | 1124.00 |
| NR1H3 | 197.00 | 144.00 | 188.00 | 176.33 | 218.00 |
| NR1I3 | 3.00 | 0.00 | 0.00 | 1.00 | 0.00 |
| NR2C1 | 818.00 | 828.00 | 977.00 | 874.33 | 724.00 |
| NR2C2 | 1785.00 | 1610.00 | 1876.00 | 1757.00 | 1606.00 |
| NR2C2AP | 512.00 | 547.00 | 525.00 | 528.00 | 672.00 |
| NR2E3 | 18.00 | 20.00 | 12.00 | 16.67 | 6.00 |
| NR2F1 | 738.00 | 680.00 | 813.00 | 743.67 | 993.00 |
| NR2F1-AS1 | 480.00 | 409.00 | 551.00 | 480.00 | 561.00 |
| NR2F2 | 3226.00 | 3011.00 | 3605.00 | 3280.67 | 3424.00 |
| NR2F2-AS1 | 215.00 | 199.00 | 227.00 | 213.67 | 124.00 |
| NR2F6 | 828.00 | 845.00 | 972.00 | 881.67 | 901.00 |
| NR3C1 | 1313.00 | 1175.00 | 1463.00 | 1317.00 | 1298.00 |
| NR3C2 | 287.00 | 238.00 | 287.00 | 270.67 | 244.00 |
| NR4A1 | 229.00 | 223.00 | 221.00 | 224.33 | 192.00 |
| NR4A2 | 18.00 | 21.00 | 21.00 | 20.00 | 18.00 |
| NR4A3 | 11.00 | 5.00 | 13.00 | 9.67 | 9.00 |
| NR5A2 | 3.00 | 9.00 | 9.00 | 7.00 | 13.00 |
| NR6A1 | 200.00 | 188.00 | 210.00 | 199.33 | 171.00 |
| NRARP | 33.00 | 33.00 | 55.00 | 40.33 | 53.00 |
| NRAS | 1827.00 | 1582.00 | 1876.00 | 1761.67 | 2336.00 |
| NRAV | 197.00 | 183.00 | 180.00 | 186.67 | 169.00 |
| NRBF2 | 536.00 | 494.00 | 524.00 | 518.00 | 537.00 |
| NRBP1 | 2177.00 | 2029.00 | 2316.00 | 2174.00 | 2316.00 |
| NRBP2 | 1148.00 | 1219.00 | 1350.00 | 1239.00 | 1167.00 |
| NRD1 | 5749.00 | 5627.00 | 6175.00 | 5850.33 | 6412.00 |
| NRDE2 | 359.00 | 378.00 | 353.00 | 363.33 | 354.00 |
| NREP | 377.00 | 370.00 | 418.00 | 388.33 | 510.00 |

| | | | | | |
|---|---|---|---|---|---|
| NRF1 | 474.00 | 481.00 | 495.00 | 483.33 | 476.00 |
| NRG1 | 1974.00 | 1677.00 | 1867.00 | 1839.33 | 1662.00 |
| NRG2 | 89.00 | 82.00 | 85.00 | 85.33 | 51.00 |
| NRGN | 177.00 | 136.00 | 149.00 | 154.00 | 185.00 |
| NRIP1 | 2110.00 | 1895.00 | 2228.00 | 2077.67 | 1607.00 |
| NRIP3 | 958.00 | 932.00 | 982.00 | 957.33 | 1587.00 |
| NRK | 1066.00 | 930.00 | 992.00 | 996.00 | 797.00 |
| NRL | 8.00 | 9.00 | 5.00 | 7.33 | 12.00 |
| NRN1 | 4.00 | 2.00 | 2.00 | 2.67 | 1.00 |
| NRN1L | 9.00 | 12.00 | 4.00 | 8.33 | 5.00 |
| NRP1 | 13376.00 | 12787.00 | 14264.00 | 13475.67 | 8341.00 |
| NRP2 | 2533.00 | 2474.00 | 2588.00 | 2531.67 | 2785.00 |
| NRROS | 475.00 | 535.00 | 555.00 | 521.67 | 515.00 |
| NRSN2 | 1236.00 | 1222.00 | 1180.00 | 1212.67 | 1216.00 |
| NRSN2-AS1 | 149.00 | 124.00 | 135.00 | 136.00 | 105.00 |
| NRXN1 | 6.00 | 2.00 | 1.00 | 3.00 | 4.00 |
| NRXN3 | 142.00 | 117.00 | 153.00 | 137.33 | 86.00 |
| NSA2 | 1284.00 | 1246.00 | 1286.00 | 1272.00 | 1775.00 |
| NSD1 | 2181.00 | 2016.00 | 2292.00 | 2163.00 | 1726.00 |
| NSDHL | 510.00 | 504.00 | 535.00 | 516.33 | 657.00 |
| NSF | 29.00 | 33.00 | 28.00 | 30.00 | 40.00 |
| NSFL1C | 1381.00 | 1398.00 | 1465.00 | 1414.67 | 1580.00 |
| NSFP1 | 5.00 | 5.00 | 8.00 | 6.00 | 4.00 |
| NSL1 | 469.00 | 464.00 | 461.00 | 464.67 | 587.00 |
| NSMAF | 2436.00 | 2335.00 | 2610.00 | 2460.33 | 2725.00 |
| NSMCE1 | 781.00 | 779.00 | 819.00 | 793.00 | 964.00 |
| NSMCE2 | 468.00 | 469.00 | 511.00 | 482.67 | 542.00 |
| NSMCE4A | 838.00 | 870.00 | 910.00 | 872.67 | 1100.00 |
| NSMF | 2379.00 | 2399.00 | 2639.00 | 2472.33 | 2203.00 |
| NSRP1 | 1046.00 | 1077.00 | 1152.00 | 1091.67 | 1395.00 |
| NSUN2 | 3349.00 | 3234.00 | 3679.00 | 3420.67 | 3650.00 |
| NSUN3 | 128.00 | 90.00 | 129.00 | 115.67 | 161.00 |
| NSUN4 | 914.00 | 861.00 | 945.00 | 906.67 | 957.00 |
| NSUN5 | 413.00 | 472.00 | 457.00 | 447.33 | 579.00 |
| NSUN5P1 | 804.00 | 884.00 | 927.00 | 871.67 | 908.00 |
| NSUN5P2 | 651.00 | 737.00 | 788.00 | 725.33 | 767.00 |
| NSUN6 | 643.00 | 622.00 | 666.00 | 643.67 | 641.00 |
| NSUN7 | 8.00 | 4.00 | 2.00 | 4.67 | 3.00 |
| NT5C | 726.00 | 758.00 | 777.00 | 753.67 | 991.00 |
| NT5C2 | 2400.00 | 2385.00 | 2608.00 | 2464.33 | 2310.00 |
| NT5C3A | 813.00 | 841.00 | 825.00 | 826.33 | 1067.00 |
| NT5C3B | 838.00 | 863.00 | 877.00 | 859.33 | 1099.00 |
| NT5DC1 | 560.00 | 516.00 | 617.00 | 564.33 | 639.00 |
| NT5DC2 | 2685.00 | 2667.00 | 2680.00 | 2677.33 | 2586.00 |
| NT5DC3 | 358.00 | 409.00 | 455.00 | 407.33 | 522.00 |
| NT5E | 27695.00 | 26190.00 | 28421.00 | 27435.33 | 25067.00 |
| NT5M | 97.00 | 106.00 | 105.00 | 102.67 | 135.00 |
| NTAN1 | 12.00 | 12.00 | 30.00 | 18.00 | 11.00 |

| | | | | | |
|---|---|---|---|---|---|
| NTHL1 | 455.00 | 502.00 | 469.00 | 475.33 | 582.00 |
| NTM | 3713.00 | 3679.00 | 3973.00 | 3788.33 | 3777.00 |
| NTMT1 | 1086.00 | 1064.00 | 1043.00 | 1064.33 | 1229.00 |
| NTN4 | 2148.00 | 1929.00 | 2075.00 | 2050.67 | 942.00 |
| NTNG1 | 3064.00 | 2929.00 | 3103.00 | 3032.00 | 2389.00 |
| NTNG2 | 29.00 | 37.00 | 36.00 | 34.00 | 8.00 |
| NTPCR | 253.00 | 236.00 | 260.00 | 249.67 | 329.00 |
| NTSR1 | 6.00 | 6.00 | 9.00 | 7.00 | 9.00 |
| NUAK1 | 207.00 | 179.00 | 202.00 | 196.00 | 200.00 |
| NUAK2 | 268.00 | 236.00 | 267.00 | 257.00 | 223.00 |
| NUB1 | 1439.00 | 1231.00 | 1454.00 | 1374.67 | 1216.00 |
| NUBP1 | 299.00 | 326.00 | 341.00 | 322.00 | 351.00 |
| NUBP2 | 724.00 | 759.00 | 854.00 | 779.00 | 915.00 |
| NUBPL | 417.00 | 400.00 | 458.00 | 425.00 | 502.00 |
| NUCB1 | 3677.00 | 3894.00 | 3499.00 | 3690.00 | 3371.00 |
| NUCB2 | 1382.00 | 1367.00 | 1311.00 | 1353.33 | 1867.00 |
| NUCKS1 | 3979.00 | 3692.00 | 4107.00 | 3926.00 | 5001.00 |
| NUDC | 4681.00 | 4892.00 | 5114.00 | 4895.67 | 6394.00 |
| NUDCD1 | 1293.00 | 1184.00 | 1355.00 | 1277.33 | 1670.00 |
| NUDCD2 | 571.00 | 555.00 | 589.00 | 571.67 | 707.00 |
| NUDCD3 | 2648.00 | 2595.00 | 2834.00 | 2692.33 | 2998.00 |
| NUDT1 | 313.00 | 259.00 | 312.00 | 294.67 | 440.00 |
| NUDT10 | 8.00 | 5.00 | 3.00 | 5.33 | 4.00 |
| NUDT11 | 293.00 | 268.00 | 313.00 | 291.33 | 375.00 |
| NUDT12 | 394.00 | 400.00 | 391.00 | 395.00 | 464.00 |
| NUDT13 | 266.00 | 284.00 | 270.00 | 273.33 | 366.00 |
| NUDT14 | 22.00 | 27.00 | 25.00 | 24.67 | 26.00 |
| NUDT15 | 412.00 | 383.00 | 417.00 | 404.00 | 570.00 |
| NUDT16 | 2001.00 | 1949.00 | 2044.00 | 1998.00 | 1823.00 |
| NUDT16L1 | 323.00 | 299.00 | 358.00 | 326.67 | 358.00 |
| NUDT17 | 135.00 | 150.00 | 117.00 | 134.00 | 142.00 |
| NUDT18 | 35.00 | 44.00 | 44.00 | 41.00 | 68.00 |
| NUDT19 | 494.00 | 403.00 | 492.00 | 463.00 | 613.00 |
| NUDT2 | 573.00 | 608.00 | 585.00 | 588.67 | 807.00 |
| NUDT21 | 2884.00 | 2357.00 | 2858.00 | 2699.67 | 3289.00 |
| NUDT22 | 1405.00 | 1445.00 | 1553.00 | 1467.67 | 1636.00 |
| NUDT3 | 62.00 | 102.00 | 94.00 | 86.00 | 121.00 |
| NUDT4 | 536.00 | 529.00 | 606.00 | 557.00 | 612.00 |
| NUDT5 | 3045.00 | 2895.00 | 3046.00 | 2995.33 | 3626.00 |
| NUDT6 | 95.00 | 102.00 | 115.00 | 104.00 | 103.00 |
| NUDT8 | 15.00 | 21.00 | 26.00 | 20.67 | 21.00 |
| NUDT9 | 497.00 | 484.00 | 511.00 | 497.33 | 530.00 |
| NUDT9P1 | 27.00 | 20.00 | 21.00 | 22.67 | 21.00 |
| NUF2 | 706.00 | 768.00 | 725.00 | 733.00 | 1187.00 |
| NUFIP1 | 273.00 | 293.00 | 276.00 | 280.67 | 376.00 |
| NUFIP2 | 4108.00 | 3743.00 | 4754.00 | 4201.67 | 3671.00 |
| NUMA1 | 12363.00 | 12206.00 | 13076.00 | 12548.33 | 10065.00 |
| NUMB | 2146.00 | 1997.00 | 2161.00 | 2101.33 | 2012.00 |

| | | | | | |
|---|---|---|---|---|---|
| NUMBL | 517.00 | 521.00 | 542.00 | 526.67 | 607.00 |
| NUP107 | 2458.00 | 2436.00 | 2533.00 | 2475.67 | 2778.00 |
| NUP133 | 1440.00 | 1260.00 | 1448.00 | 1382.67 | 1628.00 |
| NUP153 | 2912.00 | 2648.00 | 3118.00 | 2892.67 | 2992.00 |
| NUP155 | 1356.00 | 1157.00 | 1302.00 | 1271.67 | 1469.00 |
| NUP160 | 1398.00 | 1181.00 | 1348.00 | 1309.00 | 1419.00 |
| NUP188 | 3250.00 | 3001.00 | 3151.00 | 3134.00 | 3094.00 |
| NUP205 | 4095.00 | 3784.00 | 4276.00 | 4051.67 | 4535.00 |
| NUP214 | 2111.00 | 2033.00 | 2304.00 | 2149.33 | 1721.00 |
| NUP35 | 497.00 | 555.00 | 555.00 | 535.67 | 689.00 |
| NUP37 | 981.00 | 871.00 | 976.00 | 942.67 | 1140.00 |
| NUP43 | 1001.00 | 890.00 | 984.00 | 958.33 | 1235.00 |
| NUP50 | 1352.00 | 1329.00 | 1562.00 | 1414.33 | 1657.00 |
| NUP50-AS1 | 221.00 | 219.00 | 264.00 | 234.67 | 323.00 |
| NUP54 | 700.00 | 670.00 | 718.00 | 696.00 | 833.00 |
| NUP62 | 1589.00 | 1623.00 | 1686.00 | 1632.67 | 1845.00 |
| NUP62CL | 13.00 | 8.00 | 13.00 | 11.33 | 21.00 |
| NUP85 | 1086.00 | 1133.00 | 1069.00 | 1096.00 | 1249.00 |
| NUP88 | 1733.00 | 1546.00 | 1794.00 | 1691.00 | 2052.00 |
| NUP93 | 1953.00 | 1902.00 | 1980.00 | 1945.00 | 2396.00 |
| NUP98 | 3752.00 | 3627.00 | 4029.00 | 3802.67 | 3625.00 |
| NUPL1 | 1317.00 | 1160.00 | 1335.00 | 1270.67 | 1463.00 |
| NUPL2 | 967.00 | 830.00 | 1007.00 | 934.67 | 980.00 |
| NUPR1 | 450.00 | 491.00 | 444.00 | 461.67 | 317.00 |
| NUS1 | 1081.00 | 1023.00 | 1099.00 | 1067.67 | 1288.00 |
| NUSAP1 | 1062.00 | 976.00 | 1034.00 | 1024.00 | 1518.00 |
| NUTF2 | 1429.00 | 1373.00 | 1529.00 | 1443.67 | 1963.00 |
| NUTM2A | 12.00 | 11.00 | 21.00 | 14.67 | 19.00 |
| NUTM2A-AS1 | 349.00 | 300.00 | 337.00 | 328.67 | 384.00 |
| NUTM2B | 9.00 | 8.00 | 4.00 | 7.00 | 13.00 |
| NUTM2B-AS1 | 358.00 | 399.00 | 467.00 | 408.00 | 454.00 |
| NUTM2D | 26.00 | 24.00 | 19.00 | 23.00 | 25.00 |
| NVL | 773.00 | 752.00 | 764.00 | 763.00 | 919.00 |
| NXF1 | 2003.00 | 2078.00 | 2176.00 | 2085.67 | 2032.00 |
| NXF3 | 27.00 | 20.00 | 13.00 | 20.00 | 16.00 |
| NXN | 792.00 | 770.00 | 878.00 | 813.33 | 932.00 |
| NXNL2 | 12.00 | 15.00 | 15.00 | 14.00 | 13.00 |
| NXPE3 | 911.00 | 786.00 | 955.00 | 884.00 | 678.00 |
| NXPH2 | 37.00 | 27.00 | 19.00 | 27.67 | 42.00 |
| NXPH4 | 58.00 | 68.00 | 58.00 | 61.33 | 75.00 |
| NXT1 | 502.00 | 490.00 | 552.00 | 514.67 | 628.00 |
| NXT2 | 123.00 | 94.00 | 131.00 | 116.00 | 130.00 |
| NYAP1 | 34.00 | 59.00 | 29.00 | 40.67 | 32.00 |
| OAF | 2183.00 | 2197.00 | 2035.00 | 2138.33 | 1842.00 |
| OARD1 | 579.00 | 599.00 | 592.00 | 590.00 | 734.00 |
| OAS1 | 3.00 | 3.00 | 2.00 | 2.67 | 7.00 |
| OAS2 | 6.00 | 1.00 | 2.00 | 3.00 | 8.00 |
| OAS3 | 61.00 | 78.00 | 62.00 | 67.00 | 94.00 |

| | | | | | |
|---|---|---|---|---|---|
| OAT | 2113.00 | 1875.00 | 2109.00 | 2032.33 | 2073.00 |
| OAZ1 | 8270.00 | 8255.00 | 8833.00 | 8452.67 | 10465.00 |
| OAZ2 | 852.00 | 855.00 | 914.00 | 873.67 | 1219.00 |
| OAZ3 | 22.00 | 13.00 | 14.00 | 16.33 | 32.00 |
| OBFC1 | 484.00 | 578.00 | 575.00 | 545.67 | 588.00 |
| OBSCN | 219.00 | 224.00 | 223.00 | 222.00 | 132.00 |
| OBSL1 | 1763.00 | 1945.00 | 1948.00 | 1885.33 | 2024.00 |
| OCA2 | 3.00 | 2.00 | 2.00 | 2.33 | 2.00 |
| OCEL1 | 93.00 | 67.00 | 63.00 | 74.33 | 91.00 |
| OCIAD1 | 3966.00 | 3785.00 | 3855.00 | 3868.67 | 4581.00 |
| OCIAD2 | 1816.00 | 1872.00 | 1891.00 | 1859.67 | 2637.00 |
| OCLM | 22.00 | 16.00 | 19.00 | 19.00 | 22.00 |
| OCLN | 18.00 | 5.00 | 20.00 | 14.33 | 25.00 |
| OCRL | 738.00 | 648.00 | 770.00 | 718.67 | 724.00 |
| ODC1 | 8620.00 | 8005.00 | 9092.00 | 8572.33 | 9640.00 |
| ODF2 | 3229.00 | 3368.00 | 3518.00 | 3371.67 | 3529.00 |
| ODF2L | 1674.00 | 1651.00 | 1784.00 | 1703.00 | 1833.00 |
| ODF3B | 8.00 | 5.00 | 5.00 | 6.00 | 3.00 |
| OFD1 | 1841.00 | 1790.00 | 1998.00 | 1876.33 | 1755.00 |
| OGDH | 2421.00 | 2363.00 | 2458.00 | 2414.00 | 2528.00 |
| OGDHL | 55.00 | 52.00 | 44.00 | 50.33 | 82.00 |
| OGFOD1 | 1521.00 | 1578.00 | 1745.00 | 1614.67 | 1782.00 |
| OGFOD2 | 746.00 | 819.00 | 865.00 | 810.00 | 779.00 |
| OGFOD3 | 1138.00 | 1198.00 | 1135.00 | 1157.00 | 1142.00 |
| OGFR | 1053.00 | 1201.00 | 1129.00 | 1127.67 | 1192.00 |
| OGFRL1 | 531.00 | 451.00 | 524.00 | 502.00 | 466.00 |
| OGFRP1 | 4.00 | 7.00 | 7.00 | 6.00 | 10.00 |
| OGG1 | 352.00 | 340.00 | 351.00 | 347.67 | 355.00 |
| OGT | 29893.00 | 29625.00 | 32479.00 | 30665.67 | 30951.00 |
| OIP5 | 143.00 | 122.00 | 132.00 | 132.33 | 166.00 |
| OIP5-AS1 | 259.00 | 250.00 | 319.00 | 276.00 | 274.00 |
| OLA1 | 2556.00 | 2445.00 | 2779.00 | 2593.33 | 3312.00 |
| OLAH | 23.00 | 29.00 | 31.00 | 27.67 | 25.00 |
| OLFML2B | 16.00 | 17.00 | 3.00 | 12.00 | 35.00 |
| OLFML3 | 937.00 | 908.00 | 888.00 | 911.00 | 918.00 |
| OLIG2 | 10.00 | 7.00 | 12.00 | 9.67 | 14.00 |
| OLMALINC | 208.00 | 234.00 | 278.00 | 240.00 | 268.00 |
| OMA1 | 776.00 | 815.00 | 933.00 | 841.33 | 923.00 |
| ONECUT2 | 125.00 | 112.00 | 125.00 | 120.67 | 143.00 |
| OPA1 | 1981.00 | 1943.00 | 2205.00 | 2043.00 | 2095.00 |
| OPA3 | 447.00 | 430.00 | 508.00 | 461.67 | 470.00 |
| OPCML | 5.00 | 5.00 | 5.00 | 5.00 | 26.00 |
| OPHN1 | 323.00 | 285.00 | 317.00 | 308.33 | 264.00 |
| OPLAH | 3.00 | 3.00 | 5.00 | 3.67 | 9.00 |
| OPN3 | 795.00 | 678.00 | 791.00 | 754.67 | 597.00 |
| OPRL1 | 3.00 | 9.00 | 9.00 | 7.00 | 3.00 |
| OPTN | 751.00 | 725.00 | 798.00 | 758.00 | 916.00 |
| OR1F1 | 9.00 | 2.00 | 2.00 | 4.33 | 2.00 |

| | | | | | |
|---|---|---|---|---|---|
| OR2T33 | 3.00 | 0.00 | 1.00 | 1.33 | 2.00 |
| OR2T8 | 17.00 | 10.00 | 4.00 | 10.33 | 6.00 |
| OR2W3 | 40.00 | 33.00 | 36.00 | 36.33 | 35.00 |
| OR51E2 | 29.00 | 31.00 | 29.00 | 29.67 | 37.00 |
| OR51M1 | 37.00 | 35.00 | 23.00 | 31.67 | 14.00 |
| OR6F1 | 19.00 | 17.00 | 24.00 | 20.00 | 20.00 |
| ORAI1 | 714.00 | 634.00 | 651.00 | 666.33 | 745.00 |
| ORAI2 | 1485.00 | 1442.00 | 1500.00 | 1475.67 | 1105.00 |
| ORAI3 | 403.00 | 412.00 | 400.00 | 405.00 | 336.00 |
| ORAOV1 | 4063.00 | 4102.00 | 4344.00 | 4169.67 | 4248.00 |
| ORC1 | 158.00 | 131.00 | 114.00 | 134.33 | 217.00 |
| ORC2 | 1055.00 | 1065.00 | 1188.00 | 1102.67 | 1230.00 |
| ORC3 | 898.00 | 898.00 | 978.00 | 924.67 | 1183.00 |
| ORC4 | 638.00 | 699.00 | 738.00 | 691.67 | 792.00 |
| ORC5 | 844.00 | 840.00 | 829.00 | 837.67 | 1127.00 |
| ORC6 | 944.00 | 1013.00 | 876.00 | 944.33 | 1285.00 |
| ORMDL1 | 2230.00 | 2126.00 | 2321.00 | 2225.67 | 2277.00 |
| ORMDL2 | 243.00 | 231.00 | 221.00 | 231.67 | 270.00 |
| ORMDL3 | 863.00 | 909.00 | 875.00 | 882.33 | 887.00 |
| OS9 | 7123.00 | 7201.00 | 7271.00 | 7198.33 | 6500.00 |
| OSBP | 1711.00 | 1628.00 | 1779.00 | 1706.00 | 1663.00 |
| OSBP2 | 125.00 | 126.00 | 151.00 | 134.00 | 146.00 |
| OSBPL10 | 383.00 | 382.00 | 377.00 | 380.67 | 418.00 |
| OSBPL11 | 443.00 | 407.00 | 465.00 | 438.33 | 516.00 |
| OSBPL1A | 865.00 | 778.00 | 896.00 | 846.33 | 978.00 |
| OSBPL2 | 902.00 | 926.00 | 926.00 | 918.00 | 902.00 |
| OSBPL3 | 1334.00 | 1291.00 | 1452.00 | 1359.00 | 1370.00 |
| OSBPL5 | 371.00 | 428.00 | 427.00 | 408.67 | 424.00 |
| OSBPL6 | 818.00 | 774.00 | 902.00 | 831.33 | 1126.00 |
| OSBPL7 | 666.00 | 634.00 | 758.00 | 686.00 | 503.00 |
| OSBPL8 | 1170.00 | 1007.00 | 1360.00 | 1179.00 | 1077.00 |
| OSBPL9 | 1708.00 | 1716.00 | 1841.00 | 1755.00 | 1780.00 |
| OSCAR | 76.00 | 80.00 | 94.00 | 83.33 | 47.00 |
| OSCP1 | 56.00 | 69.00 | 82.00 | 69.00 | 80.00 |
| OSER1 | 1072.00 | 1033.00 | 1170.00 | 1091.67 | 1175.00 |
| OSER1-AS1 | 54.00 | 41.00 | 44.00 | 46.33 | 57.00 |
| OSGEP | 1094.00 | 1023.00 | 1083.00 | 1066.67 | 1054.00 |
| OSGEPL1 | 470.00 | 434.00 | 468.00 | 457.33 | 504.00 |
| OSGEPL1-AS1 | 15.00 | 8.00 | 14.00 | 12.33 | 18.00 |
| OSGIN1 | 198.00 | 206.00 | 239.00 | 214.33 | 213.00 |
| OSGIN2 | 668.00 | 594.00 | 721.00 | 661.00 | 754.00 |
| OSMR | 7943.00 | 7474.00 | 8317.00 | 7911.33 | 6130.00 |
| OSMR-AS1 | 9.00 | 19.00 | 11.00 | 13.00 | 13.00 |
| OSR2 | 31.00 | 33.00 | 55.00 | 39.67 | 48.00 |
| OST4 | 1112.00 | 1088.00 | 1146.00 | 1115.33 | 1557.00 |
| OSTC | 1252.00 | 1170.00 | 1274.00 | 1232.00 | 1476.00 |
| OSTF1 | 347.00 | 380.00 | 350.00 | 359.00 | 584.00 |
| OSTM1 | 1150.00 | 1101.00 | 1238.00 | 1163.00 | 1323.00 |

| | | | | | |
|---|---|---|---|---|---|
| OTOGL | 11.00 | 17.00 | 22.00 | 16.67 | 7.00 |
| OTOS | 3.00 | 1.00 | 0.00 | 1.33 | 0.00 |
| OTUB1 | 2618.00 | 2551.00 | 2546.00 | 2571.67 | 2936.00 |
| OTUD1 | 161.00 | 132.00 | 198.00 | 163.67 | 177.00 |
| OTUD3 | 858.00 | 829.00 | 937.00 | 874.67 | 788.00 |
| OTUD4 | 1563.00 | 1523.00 | 1858.00 | 1648.00 | 1559.00 |
| OTUD5 | 1216.00 | 1226.00 | 1326.00 | 1256.00 | 1324.00 |
| OTUD6B | 800.00 | 681.00 | 857.00 | 779.33 | 878.00 |
| OTUD6B-AS1 | 663.00 | 542.00 | 669.00 | 624.67 | 735.00 |
| OTUD7B | 717.00 | 661.00 | 706.00 | 694.67 | 653.00 |
| OTULIN | 682.00 | 583.00 | 713.00 | 659.33 | 586.00 |
| OTX1 | 52.00 | 62.00 | 54.00 | 56.00 | 64.00 |
| OVAAL | 342.00 | 362.00 | 367.00 | 357.00 | 378.00 |
| OVCA2 | 60.00 | 63.00 | 56.00 | 59.67 | 69.00 |
| OVGP1 | 44.00 | 81.00 | 66.00 | 63.67 | 40.00 |
| OXA1L | 2311.00 | 2282.00 | 2501.00 | 2364.67 | 2763.00 |
| OXCT1 | 1659.00 | 1636.00 | 1814.00 | 1703.00 | 2190.00 |
| OXCT1-AS1 | 11.00 | 3.00 | 5.00 | 6.33 | 10.00 |
| OXCT2 | 7.00 | 4.00 | 0.00 | 3.67 | 3.00 |
| OXGR1 | 7.00 | 5.00 | 2.00 | 4.67 | 3.00 |
| OXLD1 | 1341.00 | 1327.00 | 1453.00 | 1373.67 | 1528.00 |
| OXNAD1 | 361.00 | 370.00 | 435.00 | 388.67 | 422.00 |
| OXR1 | 663.00 | 597.00 | 764.00 | 674.67 | 801.00 |
| OXSM | 560.00 | 529.00 | 596.00 | 561.67 | 580.00 |
| OXSR1 | 1680.00 | 1605.00 | 1769.00 | 1684.67 | 1595.00 |
| OXTR | 459.00 | 479.00 | 468.00 | 468.67 | 424.00 |
| P2RX4 | 270.00 | 298.00 | 299.00 | 289.00 | 230.00 |
| P2RX5 | 500.00 | 501.00 | 463.00 | 488.00 | 543.00 |
| P2RX5-TAX1BP3 | 28.00 | 41.00 | 37.00 | 35.33 | 35.00 |
| P2RX6 | 13.00 | 5.00 | 15.00 | 11.00 | 14.00 |
| P2RX7 | 4.00 | 7.00 | 7.00 | 6.00 | 7.00 |
| P2RY1 | 161.00 | 128.00 | 133.00 | 140.67 | 97.00 |
| P2RY11 | 119.00 | 141.00 | 145.00 | 135.00 | 119.00 |
| P2RY2 | 8.00 | 4.00 | 9.00 | 7.00 | 5.00 |
| P2RY4 | 6.00 | 7.00 | 15.00 | 9.33 | 2.00 |
| P3H1 | 2837.00 | 2844.00 | 2739.00 | 2806.67 | 2563.00 |
| P3H2 | 1558.00 | 1502.00 | 1496.00 | 1518.67 | 1501.00 |
| P3H3 | 3942.00 | 4202.00 | 4041.00 | 4061.67 | 3884.00 |
| P3H4 | 1138.00 | 1283.00 | 1213.00 | 1211.33 | 1276.00 |
| P4HA1 | 2675.00 | 2559.00 | 2625.00 | 2619.67 | 2826.00 |
| P4HA2 | 899.00 | 909.00 | 891.00 | 899.67 | 890.00 |
| P4HA3 | 35.00 | 38.00 | 35.00 | 36.00 | 55.00 |
| P4HB | 23486.00 | 23451.00 | 22398.00 | 23111.67 | 21999.00 |
| P4HTM | 338.00 | 348.00 | 353.00 | 346.33 | 330.00 |
| PA2G4 | 5483.00 | 5116.00 | 5581.00 | 5393.33 | 6988.00 |
| PA2G4P4 | 155.00 | 125.00 | 129.00 | 136.33 | 162.00 |
| PAAF1 | 474.00 | 493.00 | 489.00 | 485.33 | 543.00 |
| PABPC1 | 37071.00 | 35224.00 | 40570.00 | 37621.67 | 43031.00 |

| | | | | | |
|---|---|---|---|---|---|
| PABPC1L | 4559.00 | 4578.00 | 4836.00 | 4657.67 | 4771.00 |
| PABPC3 | 169.00 | 123.00 | 143.00 | 145.00 | 170.00 |
| PABPC4 | 8543.00 | 8265.00 | 9046.00 | 8618.00 | 9453.00 |
| PABPN1 | 829.00 | 928.00 | 1073.00 | 943.33 | 1035.00 |
| PACERR | 38.00 | 36.00 | 36.00 | 36.67 | 24.00 |
| PACRG | 4.00 | 3.00 | 2.00 | 3.00 | 5.00 |
| PACRGL | 291.00 | 268.00 | 328.00 | 295.67 | 316.00 |
| PACS1 | 2281.00 | 2212.00 | 2376.00 | 2289.67 | 1804.00 |
| PACS2 | 1463.00 | 1366.00 | 1610.00 | 1479.67 | 1321.00 |
| PACSIN1 | 28.00 | 20.00 | 40.00 | 29.33 | 29.00 |
| PACSIN2 | 1166.00 | 1053.00 | 1197.00 | 1138.67 | 1140.00 |
| PACSIN3 | 530.00 | 627.00 | 571.00 | 576.00 | 610.00 |
| PADI1 | 4.00 | 2.00 | 5.00 | 3.67 | 2.00 |
| PAF1 | 827.00 | 828.00 | 894.00 | 849.67 | 894.00 |
| PAFAH1B1 | 4246.00 | 3910.00 | 4504.00 | 4220.00 | 4460.00 |
| PAFAH1B2 | 1211.00 | 1002.00 | 1135.00 | 1116.00 | 1395.00 |
| PAFAH1B3 | 298.00 | 285.00 | 308.00 | 297.00 | 437.00 |
| PAFAH2 | 398.00 | 373.00 | 425.00 | 398.67 | 400.00 |
| PAG1 | 817.00 | 724.00 | 862.00 | 801.00 | 496.00 |
| PAGE1 | 3.00 | 0.00 | 3.00 | 2.00 | 12.00 |
| PAGR1 | 505.00 | 521.00 | 539.00 | 521.67 | 570.00 |
| PAICS | 3745.00 | 3493.00 | 3977.00 | 3738.33 | 4767.00 |
| PAIP1 | 1563.00 | 1443.00 | 1743.00 | 1583.00 | 1959.00 |
| PAIP2 | 930.00 | 910.00 | 1104.00 | 981.33 | 1269.00 |
| PAIP2B | 73.00 | 85.00 | 107.00 | 88.33 | 82.00 |
| PAK1 | 1013.00 | 950.00 | 1057.00 | 1006.67 | 1178.00 |
| PAK1IP1 | 1187.00 | 1156.00 | 1273.00 | 1205.33 | 1505.00 |
| PAK2 | 2496.00 | 2053.00 | 2436.00 | 2328.33 | 2420.00 |
| PAK3 | 145.00 | 158.00 | 156.00 | 153.00 | 127.00 |
| PAK4 | 594.00 | 602.00 | 636.00 | 610.67 | 763.00 |
| PAK6 | 3.00 | 2.00 | 2.00 | 2.33 | 2.00 |
| PALB2 | 569.00 | 555.00 | 584.00 | 569.33 | 618.00 |
| PALLD | 269.00 | 300.00 | 328.00 | 299.00 | 420.00 |
| PALM2 | 75.00 | 77.00 | 87.00 | 79.67 | 71.00 |
| PALM2-AKAP2 | 14.00 | 5.00 | 5.00 | 8.00 | 4.00 |
| PAM | 4612.00 | 4606.00 | 4652.00 | 4623.33 | 3668.00 |
| PAM16 | 3.00 | 2.00 | 1.00 | 2.00 | 0.00 |
| PAMR1 | 2810.00 | 2798.00 | 2950.00 | 2852.67 | 1800.00 |
| PAN2 | 3000.00 | 3022.00 | 3404.00 | 3142.00 | 2561.00 |
| PAN3 | 1068.00 | 1033.00 | 1179.00 | 1093.33 | 1111.00 |
| PAN3-AS1 | 68.00 | 66.00 | 79.00 | 71.00 | 64.00 |
| PANK1 | 251.00 | 238.00 | 262.00 | 250.33 | 329.00 |
| PANK2 | 595.00 | 572.00 | 652.00 | 606.33 | 717.00 |
| PANK3 | 369.00 | 292.00 | 332.00 | 331.00 | 223.00 |
| PANO1 | 17.00 | 27.00 | 21.00 | 21.67 | 17.00 |
| PANX1 | 501.00 | 469.00 | 490.00 | 486.67 | 500.00 |
| PANX2 | 191.00 | 209.00 | 195.00 | 198.33 | 151.00 |
| PAOX | 28.00 | 30.00 | 39.00 | 32.33 | 41.00 |

| | | | | | |
|---|---|---|---|---|---|
| PAPD4 | 831.00 | 760.00 | 860.00 | 817.00 | 854.00 |
| PAPD5 | 505.00 | 415.00 | 587.00 | 502.33 | 486.00 |
| PAPD7 | 1372.00 | 1390.00 | 1474.00 | 1412.00 | 1330.00 |
| PAPLN | 277.00 | 255.00 | 293.00 | 275.00 | 366.00 |
| PAPOLA | 3587.00 | 3138.00 | 3797.00 | 3507.33 | 4030.00 |
| PAPOLG | 456.00 | 441.00 | 495.00 | 464.00 | 490.00 |
| PAPPA | 262.00 | 251.00 | 298.00 | 270.33 | 124.00 |
| PAPPA2 | 2120.00 | 1754.00 | 2108.00 | 1994.00 | 1047.00 |
| PAPSS1 | 721.00 | 675.00 | 820.00 | 738.67 | 802.00 |
| PAPSS2 | 695.00 | 649.00 | 728.00 | 690.67 | 759.00 |
| PAQR3 | 774.00 | 624.00 | 767.00 | 721.67 | 685.00 |
| PAQR4 | 371.00 | 396.00 | 392.00 | 386.33 | 572.00 |
| PAQR5 | 48.00 | 45.00 | 47.00 | 46.67 | 41.00 |
| PAQR6 | 285.00 | 252.00 | 323.00 | 286.67 | 237.00 |
| PAQR7 | 922.00 | 1008.00 | 928.00 | 952.67 | 1149.00 |
| PAQR8 | 939.00 | 919.00 | 878.00 | 912.00 | 651.00 |
| PARD3 | 2075.00 | 2067.00 | 2283.00 | 2141.67 | 2030.00 |
| PARD3-AS1 | 20.00 | 11.00 | 22.00 | 17.67 | 16.00 |
| PARD3B | 104.00 | 116.00 | 120.00 | 113.33 | 102.00 |
| PARD6A | 25.00 | 31.00 | 20.00 | 25.33 | 25.00 |
| PARD6B | 76.00 | 56.00 | 64.00 | 65.33 | 91.00 |
| PARD6G | 6.00 | 2.00 | 3.00 | 3.67 | 5.00 |
| PARD6G-AS1 | 5.00 | 9.00 | 5.00 | 6.33 | 6.00 |
| PARG | 658.00 | 672.00 | 683.00 | 671.00 | 730.00 |
| PARGP1 | 176.00 | 146.00 | 183.00 | 168.33 | 95.00 |
| PARK2 | 34.00 | 36.00 | 40.00 | 36.67 | 44.00 |
| PARK7 | 2855.00 | 2821.00 | 2978.00 | 2884.67 | 3813.00 |
| PARL | 728.00 | 732.00 | 680.00 | 713.33 | 869.00 |
| PARM1 | 4.00 | 4.00 | 4.00 | 4.00 | 1.00 |
| PARN | 6.00 | 6.00 | 7.00 | 6.33 | 1.00 |
| PARP1 | 2814.00 | 2690.00 | 2929.00 | 2811.00 | 3476.00 |
| PARP10 | 731.00 | 871.00 | 901.00 | 834.33 | 827.00 |
| PARP11 | 227.00 | 210.00 | 214.00 | 217.00 | 224.00 |
| PARP12 | 362.00 | 308.00 | 337.00 | 335.67 | 381.00 |
| PARP14 | 834.00 | 847.00 | 1015.00 | 898.67 | 845.00 |
| PARP16 | 219.00 | 181.00 | 201.00 | 200.33 | 258.00 |
| PARP2 | 778.00 | 680.00 | 845.00 | 767.67 | 861.00 |
| PARP3 | 1083.00 | 1150.00 | 1268.00 | 1167.00 | 1120.00 |
| PARP4 | 2771.00 | 2492.00 | 2794.00 | 2685.67 | 2146.00 |
| PARP6 | 1679.00 | 1737.00 | 1753.00 | 1723.00 | 1631.00 |
| PARP8 | 563.00 | 553.00 | 614.00 | 576.67 | 531.00 |
| PARP9 | 291.00 | 270.00 | 287.00 | 282.67 | 240.00 |
| PARPBP | 395.00 | 363.00 | 402.00 | 386.67 | 484.00 |
| PARS2 | 187.00 | 200.00 | 207.00 | 198.00 | 198.00 |
| PART1 | 4.00 | 3.00 | 4.00 | 3.67 | 5.00 |
| PARTICL | 61.00 | 55.00 | 47.00 | 54.33 | 67.00 |
| PARVA | 2222.00 | 2095.00 | 2504.00 | 2273.67 | 2011.00 |
| PARVB | 857.00 | 842.00 | 941.00 | 880.00 | 1138.00 |

| | | | | | |
|---|---|---|---|---|---|
| PASK | 415.00 | 379.00 | 459.00 | 417.67 | 467.00 |
| PATE2 | 9.00 | 8.00 | 7.00 | 8.00 | 7.00 |
| PATL1 | 1147.00 | 1170.00 | 1266.00 | 1194.33 | 1195.00 |
| PATL2 | 21.00 | 31.00 | 33.00 | 28.33 | 38.00 |
| PATZ1 | 1080.00 | 1106.00 | 1215.00 | 1133.67 | 1177.00 |
| PAWR | 393.00 | 393.00 | 476.00 | 420.67 | 349.00 |
| PAX3 | 614.00 | 544.00 | 604.00 | 587.33 | 573.00 |
| PAX8 | 24.00 | 28.00 | 24.00 | 25.33 | 21.00 |
| PAX8-AS1 | 11.00 | 4.00 | 17.00 | 10.67 | 10.00 |
| PAX9 | 134.00 | 136.00 | 131.00 | 133.67 | 150.00 |
| PAXBP1 | 1734.00 | 1759.00 | 1926.00 | 1806.33 | 1819.00 |
| PAXBP1-AS1 | 32.00 | 33.00 | 33.00 | 32.67 | 26.00 |
| PAXIP1 | 734.00 | 773.00 | 774.00 | 760.33 | 818.00 |
| PAXIP1-AS1 | 319.00 | 268.00 | 329.00 | 305.33 | 322.00 |
| PAXIP1-AS2 | 124.00 | 132.00 | 134.00 | 130.00 | 117.00 |
| PBDC1 | 485.00 | 497.00 | 569.00 | 517.00 | 615.00 |
| PBK | 335.00 | 322.00 | 329.00 | 328.67 | 545.00 |
| PBLD | 111.00 | 126.00 | 135.00 | 124.00 | 152.00 |
| PBRM1 | 1557.00 | 1458.00 | 1734.00 | 1583.00 | 1385.00 |
| PBX2 | 19.00 | 22.00 | 25.00 | 22.00 | 17.00 |
| PBX3 | 273.00 | 289.00 | 360.00 | 307.33 | 587.00 |
| PBXIP1 | 1193.00 | 1236.00 | 1242.00 | 1223.67 | 1011.00 |
| PC | 835.00 | 873.00 | 865.00 | 857.67 | 926.00 |
| PCAT6 | 11.00 | 18.00 | 8.00 | 12.33 | 9.00 |
| PCBD1 | 976.00 | 894.00 | 943.00 | 937.67 | 1427.00 |
| PCBD2 | 295.00 | 277.00 | 280.00 | 284.00 | 321.00 |
| PCBP1 | 4871.00 | 4687.00 | 4669.00 | 4742.33 | 6029.00 |
| PCBP1-AS1 | 237.00 | 268.00 | 273.00 | 259.33 | 261.00 |
| PCBP2 | 10352.00 | 9988.00 | 11012.00 | 10450.67 | 12642.00 |
| PCBP2-OT1 | 19.00 | 6.00 | 8.00 | 11.00 | 8.00 |
| PCBP3 | 12.00 | 11.00 | 16.00 | 13.00 | 6.00 |
| PCBP4 | 637.00 | 666.00 | 705.00 | 669.33 | 697.00 |
| PCCA | 435.00 | 372.00 | 426.00 | 411.00 | 350.00 |
| PCCB | 1005.00 | 972.00 | 1050.00 | 1009.00 | 1135.00 |
| PCDH1 | 3.00 | 2.00 | 0.00 | 1.67 | 2.00 |
| PCDH10 | 50.00 | 48.00 | 47.00 | 48.33 | 37.00 |
| PCDH15 | 257.00 | 208.00 | 259.00 | 241.33 | 268.00 |
| PCDH18 | 12.00 | 17.00 | 12.00 | 13.67 | 18.00 |
| PCDH20 | 5.00 | 6.00 | 6.00 | 5.67 | 8.00 |
| PCDH7 | 36.00 | 28.00 | 34.00 | 32.67 | 14.00 |
| PCDH9 | 60.00 | 70.00 | 54.00 | 61.33 | 43.00 |
| PCDHA4 | 3.00 | 5.00 | 3.00 | 3.67 | 3.00 |
| PCDHA6 | 11.00 | 5.00 | 6.00 | 7.33 | 1.00 |
| PCDHB10 | 12.00 | 14.00 | 17.00 | 14.33 | 15.00 |
| PCDHB11 | 3.00 | 5.00 | 9.00 | 5.67 | 7.00 |
| PCDHB12 | 10.00 | 19.00 | 13.00 | 14.00 | 18.00 |
| PCDHB13 | 30.00 | 41.00 | 18.00 | 29.67 | 35.00 |
| PCDHB14 | 69.00 | 56.00 | 50.00 | 58.33 | 63.00 |

| | | | | | |
|---|---|---|---|---|---|
| PCDHB15 | 60.00 | 80.00 | 69.00 | 69.67 | 58.00 |
| PCDHB16 | 41.00 | 45.00 | 39.00 | 41.67 | 38.00 |
| PCDHB18P | 3.00 | 1.00 | 4.00 | 2.67 | 0.00 |
| PCDHB19P | 4.00 | 6.00 | 2.00 | 4.00 | 3.00 |
| PCDHB2 | 55.00 | 55.00 | 48.00 | 52.67 | 49.00 |
| PCDHB3 | 3.00 | 3.00 | 1.00 | 2.33 | 5.00 |
| PCDHB4 | 3.00 | 8.00 | 8.00 | 6.33 | 8.00 |
| PCDHB5 | 46.00 | 42.00 | 38.00 | 42.00 | 59.00 |
| PCDHB6 | 34.00 | 34.00 | 31.00 | 33.00 | 49.00 |
| PCDHB7 | 11.00 | 11.00 | 11.00 | 11.00 | 16.00 |
| PCDHB8 | 12.00 | 13.00 | 11.00 | 12.00 | 13.00 |
| PCDHB9 | 15.00 | 11.00 | 7.00 | 11.00 | 10.00 |
| PCDHGA1 | 30.00 | 33.00 | 36.00 | 33.00 | 17.00 |
| PCDHGA10 | 96.00 | 121.00 | 110.00 | 109.00 | 42.00 |
| PCDHGA11 | 314.00 | 347.00 | 333.00 | 331.33 | 173.00 |
| PCDHGA12 | 11.00 | 3.00 | 8.00 | 7.33 | 4.00 |
| PCDHGA2 | 26.00 | 25.00 | 20.00 | 23.67 | 25.00 |
| PCDHGA3 | 13.00 | 7.00 | 16.00 | 12.00 | 7.00 |
| PCDHGA4 | 20.00 | 17.00 | 21.00 | 19.33 | 9.00 |
| PCDHGA5 | 11.00 | 13.00 | 17.00 | 13.67 | 11.00 |
| PCDHGA6 | 27.00 | 30.00 | 31.00 | 29.33 | 32.00 |
| PCDHGA7 | 11.00 | 13.00 | 12.00 | 12.00 | 9.00 |
| PCDHGA9 | 18.00 | 6.00 | 17.00 | 13.67 | 4.00 |
| PCDHGB1 | 54.00 | 57.00 | 64.00 | 58.33 | 36.00 |
| PCDHGB2 | 24.00 | 28.00 | 24.00 | 25.33 | 9.00 |
| PCDHGB3 | 15.00 | 18.00 | 24.00 | 19.00 | 4.00 |
| PCDHGB4 | 41.00 | 39.00 | 34.00 | 38.00 | 26.00 |
| PCDHGB5 | 227.00 | 201.00 | 212.00 | 213.33 | 119.00 |
| PCDHGB6 | 12.00 | 13.00 | 26.00 | 17.00 | 21.00 |
| PCDHGB7 | 19.00 | 27.00 | 27.00 | 24.33 | 10.00 |
| PCDHGC3 | 970.00 | 834.00 | 832.00 | 878.67 | 404.00 |
| PCDHGC4 | 16.00 | 15.00 | 10.00 | 13.67 | 5.00 |
| PCDHGC5 | 23.00 | 15.00 | 10.00 | 16.00 | 4.00 |
| PCED1A | 3414.00 | 3699.00 | 3815.00 | 3642.67 | 3894.00 |
| PCED1B | 433.00 | 406.00 | 433.00 | 424.00 | 348.00 |
| PCF11 | 1995.00 | 1883.00 | 2172.00 | 2016.67 | 1815.00 |
| PCGF1 | 415.00 | 455.00 | 452.00 | 440.67 | 561.00 |
| PCGF3 | 694.00 | 681.00 | 751.00 | 708.67 | 755.00 |
| PCGF5 | 827.00 | 790.00 | 958.00 | 858.33 | 812.00 |
| PCGF6 | 328.00 | 314.00 | 315.00 | 319.00 | 435.00 |
| PCID2 | 1402.00 | 1429.00 | 1604.00 | 1478.33 | 1670.00 |
| PCIF1 | 612.00 | 521.00 | 518.00 | 550.33 | 595.00 |
| PCK2 | 4655.00 | 4387.00 | 4287.00 | 4443.00 | 4869.00 |
| PCLO | 484.00 | 420.00 | 505.00 | 469.67 | 343.00 |
| PCM1 | 1716.00 | 1597.00 | 1922.00 | 1745.00 | 1857.00 |
| PCMT1 | 1505.00 | 1356.00 | 1342.00 | 1401.00 | 1835.00 |
| PCMTD1 | 790.00 | 777.00 | 922.00 | 829.67 | 809.00 |
| PCMTD2 | 658.00 | 583.00 | 771.00 | 670.67 | 761.00 |

| | | | | | |
|---|---|---|---|---|---|
| PCNA | 997.00 | 970.00 | 960.00 | 975.67 | 1535.00 |
| PCNP | 1901.00 | 1660.00 | 2071.00 | 1877.33 | 2286.00 |
| PCNT | 2470.00 | 2306.00 | 2457.00 | 2411.00 | 2164.00 |
| PCNX | 2475.00 | 2187.00 | 2612.00 | 2424.67 | 2021.00 |
| PCNXL2 | 1248.00 | 1209.00 | 1232.00 | 1229.67 | 966.00 |
| PCNXL3 | 4854.00 | 4668.00 | 4951.00 | 4824.33 | 3489.00 |
| PCNXL4 | 2536.00 | 2270.00 | 2581.00 | 2462.33 | 2454.00 |
| PCOLCE | 6304.00 | 6439.00 | 5908.00 | 6217.00 | 5669.00 |
| PCOLCE2 | 503.00 | 475.00 | 511.00 | 496.33 | 640.00 |
| PCOLCE-AS1 | 55.00 | 53.00 | 62.00 | 56.67 | 51.00 |
| PCSK1 | 9.00 | 8.00 | 6.00 | 7.67 | 4.00 |
| PCSK4 | 131.00 | 146.00 | 186.00 | 154.33 | 124.00 |
| PCSK7 | 1263.00 | 1214.00 | 1238.00 | 1238.33 | 1219.00 |
| PCTP | 295.00 | 270.00 | 312.00 | 292.33 | 375.00 |
| PCYOX1 | 1763.00 | 1654.00 | 1948.00 | 1788.33 | 1403.00 |
| PCYOX1L | 507.00 | 471.00 | 464.00 | 480.67 | 484.00 |
| PCYT1A | 544.00 | 531.00 | 562.00 | 545.67 | 495.00 |
| PCYT2 | 654.00 | 686.00 | 664.00 | 668.00 | 726.00 |
| PDAP1 | 2559.00 | 2699.00 | 2537.00 | 2598.33 | 3385.00 |
| PDC | 4.00 | 1.00 | 4.00 | 3.00 | 0.00 |
| PDCD10 | 732.00 | 630.00 | 718.00 | 693.33 | 910.00 |
| PDCD11 | 6300.00 | 6047.00 | 6457.00 | 6268.00 | 5444.00 |
| PDCD1LG2 | 1066.00 | 944.00 | 1097.00 | 1035.67 | 1210.00 |
| PDCD2 | 1467.00 | 1357.00 | 1616.00 | 1480.00 | 1982.00 |
| PDCD2L | 214.00 | 230.00 | 227.00 | 223.67 | 356.00 |
| PDCD4 | 2667.00 | 2551.00 | 2899.00 | 2705.67 | 2459.00 |
| PDCD4-AS1 | 145.00 | 186.00 | 148.00 | 159.67 | 179.00 |
| PDCD5 | 961.00 | 979.00 | 1032.00 | 990.67 | 1449.00 |
| PDCD6 | 2941.00 | 2886.00 | 3048.00 | 2958.33 | 4000.00 |
| PDCD6IP | 4568.00 | 4231.00 | 4700.00 | 4499.67 | 5035.00 |
| PDCD6IPP2 | 37.00 | 35.00 | 23.00 | 31.67 | 29.00 |
| PDCD7 | 874.00 | 849.00 | 910.00 | 877.67 | 1041.00 |
| PDCL | 252.00 | 252.00 | 279.00 | 261.00 | 244.00 |
| PDCL3 | 356.00 | 330.00 | 355.00 | 347.00 | 430.00 |
| PDCL3P4 | 45.00 | 44.00 | 30.00 | 39.67 | 54.00 |
| PDDC1 | 2800.00 | 2989.00 | 2993.00 | 2927.33 | 2653.00 |
| PDE10A | 12.00 | 17.00 | 26.00 | 18.33 | 15.00 |
| PDE11A | 19.00 | 10.00 | 14.00 | 14.33 | 9.00 |
| PDE12 | 708.00 | 661.00 | 738.00 | 702.33 | 759.00 |
| PDE1C | 1150.00 | 1097.00 | 1202.00 | 1149.67 | 1263.00 |
| PDE2A | 152.00 | 154.00 | 166.00 | 157.33 | 142.00 |
| PDE3A | 20.00 | 20.00 | 21.00 | 20.33 | 9.00 |
| PDE3B | 462.00 | 436.00 | 533.00 | 477.00 | 608.00 |
| PDE4A | 563.00 | 482.00 | 553.00 | 532.67 | 496.00 |
| PDE4B | 224.00 | 183.00 | 203.00 | 203.33 | 232.00 |
| PDE4C | 10.00 | 12.00 | 19.00 | 13.67 | 10.00 |
| PDE4D | 792.00 | 784.00 | 934.00 | 836.67 | 824.00 |
| PDE4DIP | 2917.00 | 2948.00 | 2951.00 | 2938.67 | 2678.00 |

| | | | | | |
|---|---|---|---|---|---|
| PDE5A | 82.00 | 93.00 | 103.00 | 92.67 | 98.00 |
| PDE6D | 461.00 | 432.00 | 444.00 | 445.67 | 530.00 |
| PDE6G | 3.00 | 5.00 | 9.00 | 5.67 | 4.00 |
| PDE7A | 697.00 | 755.00 | 810.00 | 754.00 | 829.00 |
| PDE7B | 102.00 | 77.00 | 100.00 | 93.00 | 206.00 |
| PDE8A | 909.00 | 880.00 | 959.00 | 916.00 | 877.00 |
| PDF | 71.00 | 63.00 | 54.00 | 62.67 | 77.00 |
| PDGFA | 1349.00 | 1377.00 | 1454.00 | 1393.33 | 854.00 |
| PDGFB | 25.00 | 36.00 | 25.00 | 28.67 | 16.00 |
| PDGFC | 2641.00 | 2374.00 | 2591.00 | 2535.33 | 2435.00 |
| PDGFD | 125.00 | 96.00 | 113.00 | 111.33 | 123.00 |
| PDGFRA | 1397.00 | 1233.00 | 1371.00 | 1333.67 | 988.00 |
| PDGFRB | 58.00 | 63.00 | 68.00 | 63.00 | 29.00 |
| PDGFRL | 105.00 | 113.00 | 104.00 | 107.33 | 77.00 |
| PDHA1 | 1176.00 | 1123.00 | 1286.00 | 1195.00 | 1361.00 |
| PDHB | 840.00 | 817.00 | 853.00 | 836.67 | 1123.00 |
| PDHX | 718.00 | 669.00 | 703.00 | 696.67 | 903.00 |
| PDIA3 | 15725.00 | 15204.00 | 15473.00 | 15467.33 | 14544.00 |
| PDIA3P1 | 1449.00 | 1358.00 | 1476.00 | 1427.67 | 1394.00 |
| PDIA4 | 14463.00 | 14599.00 | 13884.00 | 14315.33 | 13957.00 |
| PDIA5 | 1296.00 | 1275.00 | 1222.00 | 1264.33 | 1399.00 |
| PDIA6 | 13719.00 | 13599.00 | 13451.00 | 13589.67 | 14895.00 |
| PDIK1L | 169.00 | 136.00 | 159.00 | 154.67 | 166.00 |
| PDK1 | 502.00 | 519.00 | 614.00 | 545.00 | 755.00 |
| PDK2 | 291.00 | 258.00 | 301.00 | 283.33 | 293.00 |
| PDK3 | 205.00 | 170.00 | 210.00 | 195.00 | 200.00 |
| PDK4 | 6.00 | 7.00 | 10.00 | 7.67 | 14.00 |
| PDLIM1 | 887.00 | 920.00 | 1006.00 | 937.67 | 896.00 |
| PDLIM2 | 449.00 | 410.00 | 397.00 | 418.67 | 532.00 |
| PDLIM3 | 67.00 | 50.00 | 61.00 | 59.33 | 28.00 |
| PDLIM4 | 635.00 | 628.00 | 699.00 | 654.00 | 723.00 |
| PDLIM5 | 498.00 | 480.00 | 520.00 | 499.33 | 546.00 |
| PDLIM7 | 1645.00 | 1700.00 | 1865.00 | 1736.67 | 2202.00 |
| PDP1 | 4212.00 | 3924.00 | 4676.00 | 4270.67 | 3958.00 |
| PDP2 | 737.00 | 692.00 | 888.00 | 772.33 | 626.00 |
| PDPK1 | 785.00 | 725.00 | 815.00 | 775.00 | 575.00 |
| PDPN | 3.00 | 4.00 | 8.00 | 5.00 | 2.00 |
| PDPR | 2419.00 | 2202.00 | 2482.00 | 2367.67 | 1890.00 |
| PDRG1 | 497.00 | 480.00 | 511.00 | 496.00 | 649.00 |
| PDS5A | 5309.00 | 4691.00 | 5784.00 | 5261.33 | 5756.00 |
| PDS5B | 624.00 | 603.00 | 682.00 | 636.33 | 647.00 |
| PDSS1 | 342.00 | 313.00 | 343.00 | 332.67 | 478.00 |
| PDSS2 | 430.00 | 371.00 | 421.00 | 407.33 | 472.00 |
| PDXDC1 | 96.00 | 98.00 | 121.00 | 105.00 | 111.00 |
| PDXDC2P | 560.00 | 591.00 | 580.00 | 577.00 | 406.00 |
| PDXK | 2030.00 | 2049.00 | 2214.00 | 2097.67 | 1902.00 |
| PDXP | 486.00 | 441.00 | 437.00 | 454.67 | 588.00 |
| PDZD11 | 998.00 | 875.00 | 1014.00 | 962.33 | 1190.00 |

| | | | | | |
|---|---|---|---|---|---|
| PDZD2 | 26.00 | 23.00 | 22.00 | 23.67 | 7.00 |
| PDZD7 | 5.00 | 3.00 | 6.00 | 4.67 | 4.00 |
| PDZD8 | 1821.00 | 1682.00 | 1981.00 | 1828.00 | 1815.00 |
| PDZK1 | 8.00 | 4.00 | 13.00 | 8.33 | 4.00 |
| PDZRN3 | 20.00 | 27.00 | 24.00 | 23.67 | 23.00 |
| PDZRN3-AS1 | 3.00 | 0.00 | 2.00 | 1.67 | 1.00 |
| PEA15 | 6188.00 | 5932.00 | 6443.00 | 6187.67 | 5751.00 |
| PEAK1 | 1334.00 | 1171.00 | 1281.00 | 1262.00 | 1050.00 |
| PEAR1 | 7441.00 | 7554.00 | 7433.00 | 7476.00 | 6172.00 |
| PEBP1 | 3314.00 | 3157.00 | 3529.00 | 3333.33 | 4116.00 |
| PECAM1 | 3.00 | 4.00 | 6.00 | 4.33 | 5.00 |
| PECR | 136.00 | 109.00 | 150.00 | 131.67 | 179.00 |
| PEF1 | 962.00 | 921.00 | 939.00 | 940.67 | 1057.00 |
| PEG10 | 1201.00 | 1022.00 | 1242.00 | 1155.00 | 1389.00 |
| PELI1 | 229.00 | 210.00 | 261.00 | 233.33 | 275.00 |
| PELI2 | 29.00 | 27.00 | 37.00 | 31.00 | 40.00 |
| PELI3 | 178.00 | 160.00 | 169.00 | 169.00 | 111.00 |
| PELO | 314.00 | 335.00 | 351.00 | 333.33 | 362.00 |
| PELP1 | 2730.00 | 2806.00 | 2733.00 | 2756.33 | 2970.00 |
| PEMT | 717.00 | 716.00 | 745.00 | 726.00 | 839.00 |
| PEPD | 846.00 | 785.00 | 830.00 | 820.33 | 932.00 |
| PER1 | 726.00 | 636.00 | 753.00 | 705.00 | 674.00 |
| PER2 | 412.00 | 368.00 | 395.00 | 391.67 | 306.00 |
| PER3 | 359.00 | 293.00 | 381.00 | 344.33 | 414.00 |
| PERP | 2276.00 | 2251.00 | 2189.00 | 2238.67 | 2563.00 |
| PES1 | 2612.00 | 2555.00 | 2838.00 | 2668.33 | 3241.00 |
| PET100 | 211.00 | 311.00 | 291.00 | 271.00 | 370.00 |
| PET117 | 113.00 | 117.00 | 125.00 | 118.33 | 135.00 |
| PEX1 | 897.00 | 947.00 | 983.00 | 942.33 | 847.00 |
| PEX10 | 664.00 | 716.00 | 718.00 | 699.33 | 823.00 |
| PEX11A | 171.00 | 185.00 | 205.00 | 187.00 | 187.00 |
| PEX11B | 408.00 | 441.00 | 433.00 | 427.33 | 536.00 |
| PEX11G | 26.00 | 37.00 | 30.00 | 31.00 | 45.00 |
| PEX12 | 289.00 | 273.00 | 276.00 | 279.33 | 312.00 |
| PEX13 | 393.00 | 316.00 | 378.00 | 362.33 | 428.00 |
| PEX14 | 505.00 | 497.00 | 499.00 | 500.33 | 572.00 |
| PEX16 | 407.00 | 423.00 | 431.00 | 420.33 | 438.00 |
| PEX19 | 1080.00 | 906.00 | 1046.00 | 1010.67 | 1083.00 |
| PEX2 | 566.00 | 608.00 | 739.00 | 637.67 | 772.00 |
| PEX26 | 591.00 | 521.00 | 587.00 | 566.33 | 550.00 |
| PEX3 | 464.00 | 468.00 | 489.00 | 473.67 | 546.00 |
| PEX5 | 1482.00 | 1390.00 | 1571.00 | 1481.00 | 1407.00 |
| PEX5L | 142.00 | 136.00 | 119.00 | 132.33 | 370.00 |
| PEX6 | 3136.00 | 3096.00 | 3284.00 | 3172.00 | 2883.00 |
| PEX7 | 186.00 | 194.00 | 205.00 | 195.00 | 277.00 |
| PF4 | 8.00 | 8.00 | 13.00 | 9.67 | 8.00 |
| PFAS | 4929.00 | 4553.00 | 4899.00 | 4793.67 | 4693.00 |
| PFDN1 | 2272.00 | 2141.00 | 2273.00 | 2228.67 | 3021.00 |

| | | | | | |
|---|---|---|---|---|---|
| PFDN2 | 1855.00 | 1925.00 | 1902.00 | 1894.00 | 2690.00 |
| PFDN4 | 551.00 | 538.00 | 608.00 | 565.67 | 749.00 |
| PFDN5 | 3495.00 | 3643.00 | 3879.00 | 3672.33 | 4862.00 |
| PFDN6 | 3.00 | 0.00 | 3.00 | 2.00 | 0.00 |
| PFKFB2 | 438.00 | 389.00 | 467.00 | 431.33 | 417.00 |
| PFKFB3 | 1411.00 | 1484.00 | 1645.00 | 1513.33 | 1809.00 |
| PFKFB4 | 144.00 | 170.00 | 158.00 | 157.33 | 112.00 |
| PFKL | 2112.00 | 2103.00 | 2221.00 | 2145.33 | 2200.00 |
| PFKM | 3462.00 | 3326.00 | 3886.00 | 3558.00 | 3829.00 |
| PFKP | 11303.00 | 10832.00 | 11510.00 | 11215.00 | 10140.00 |
| PFN1 | 18361.00 | 17762.00 | 18757.00 | 18293.33 | 25412.00 |
| PFN2 | 4499.00 | 4404.00 | 4846.00 | 4583.00 | 6404.00 |
| PFN4 | 29.00 | 19.00 | 22.00 | 23.33 | 21.00 |
| PGAM1 | 1441.00 | 1304.00 | 1496.00 | 1413.67 | 1964.00 |
| PGAM2 | 4.00 | 7.00 | 10.00 | 7.00 | 12.00 |
| PGAM4 | 3.00 | 9.00 | 10.00 | 7.33 | 8.00 |
| PGAM5 | 812.00 | 714.00 | 836.00 | 787.33 | 1032.00 |
| PGAP1 | 560.00 | 485.00 | 611.00 | 552.00 | 450.00 |
| PGAP2 | 621.00 | 649.00 | 651.00 | 640.33 | 843.00 |
| PGAP3 | 258.00 | 189.00 | 237.00 | 228.00 | 153.00 |
| PGBD1 | 284.00 | 278.00 | 333.00 | 298.33 | 336.00 |
| PGBD2 | 243.00 | 284.00 | 245.00 | 257.33 | 292.00 |
| PGBD4 | 261.00 | 250.00 | 263.00 | 258.00 | 238.00 |
| PGD | 3550.00 | 3313.00 | 3618.00 | 3493.67 | 3615.00 |
| PGF | 594.00 | 593.00 | 553.00 | 580.00 | 533.00 |
| PGGT1B | 180.00 | 175.00 | 199.00 | 184.67 | 185.00 |
| PGK1 | 9451.00 | 8839.00 | 9724.00 | 9338.00 | 12212.00 |
| PGLS | 1651.00 | 1674.00 | 1840.00 | 1721.67 | 2026.00 |
| PGM1 | 678.00 | 690.00 | 692.00 | 686.67 | 737.00 |
| PGM2 | 1568.00 | 1406.00 | 1638.00 | 1537.33 | 1969.00 |
| PGM2L1 | 170.00 | 159.00 | 160.00 | 163.00 | 141.00 |
| PGM3 | 1833.00 | 1779.00 | 2020.00 | 1877.33 | 1882.00 |
| PGM5 | 62.00 | 19.00 | 34.00 | 38.33 | 21.00 |
| PGM5P2 | 1099.00 | 1081.00 | 1264.00 | 1148.00 | 553.00 |
| PGM5P3-AS1 | 32.00 | 52.00 | 41.00 | 41.67 | 45.00 |
| PGP | 806.00 | 847.00 | 850.00 | 834.33 | 840.00 |
| PGPEP1 | 623.00 | 676.00 | 717.00 | 672.00 | 503.00 |
| PGR | 112.00 | 87.00 | 129.00 | 109.33 | 88.00 |
| PGRMC1 | 1085.00 | 1174.00 | 1118.00 | 1125.67 | 1523.00 |
| PGRMC2 | 1052.00 | 1107.00 | 1106.00 | 1088.33 | 1147.00 |
| PGS1 | 1584.00 | 1603.00 | 1710.00 | 1632.33 | 1494.00 |
| PHACTR1 | 274.00 | 238.00 | 251.00 | 254.33 | 278.00 |
| PHACTR2 | 555.00 | 492.00 | 536.00 | 527.67 | 594.00 |
| PHACTR4 | 881.00 | 846.00 | 866.00 | 864.33 | 876.00 |
| PHAX | 653.00 | 621.00 | 720.00 | 664.67 | 780.00 |
| PHB | 2912.00 | 2887.00 | 3090.00 | 2963.00 | 3940.00 |
| PHB2 | 4535.00 | 4829.00 | 4988.00 | 4784.00 | 6146.00 |
| PHC1 | 689.00 | 655.00 | 812.00 | 718.67 | 551.00 |

| | | | | | |
|---|---|---|---|---|---|
| PHC2 | 3457.00 | 3304.00 | 3502.00 | 3421.00 | 3569.00 |
| PHC3 | 1315.00 | 1204.00 | 1573.00 | 1364.00 | 1152.00 |
| PHEX | 53.00 | 50.00 | 69.00 | 57.33 | 75.00 |
| PHF1 | 79.00 | 76.00 | 101.00 | 85.33 | 84.00 |
| PHF10 | 807.00 | 745.00 | 901.00 | 817.67 | 791.00 |
| PHF11 | 142.00 | 126.00 | 138.00 | 135.33 | 147.00 |
| PHF12 | 1207.00 | 1144.00 | 1138.00 | 1163.00 | 944.00 |
| PHF13 | 559.00 | 563.00 | 570.00 | 564.00 | 619.00 |
| PHF14 | 1366.00 | 1355.00 | 1400.00 | 1373.67 | 1726.00 |
| PHF19 | 917.00 | 838.00 | 898.00 | 884.33 | 1125.00 |
| PHF2 | 848.00 | 849.00 | 877.00 | 858.00 | 757.00 |
| PHF20 | 1388.00 | 1329.00 | 1570.00 | 1429.00 | 1374.00 |
| PHF20L1 | 2099.00 | 2043.00 | 2292.00 | 2144.67 | 2169.00 |
| PHF21A | 932.00 | 867.00 | 895.00 | 898.00 | 883.00 |
| PHF23 | 1388.00 | 1417.00 | 1625.00 | 1476.67 | 1759.00 |
| PHF3 | 1779.00 | 1675.00 | 2101.00 | 1851.67 | 1651.00 |
| PHF5A | 531.00 | 527.00 | 546.00 | 534.67 | 793.00 |
| PHF6 | 926.00 | 783.00 | 947.00 | 885.33 | 1010.00 |
| PHF7 | 124.00 | 148.00 | 133.00 | 135.00 | 127.00 |
| PHF8 | 1423.00 | 1351.00 | 1448.00 | 1407.33 | 1074.00 |
| PHGDH | 12180.00 | 11948.00 | 12431.00 | 12186.33 | 14692.00 |
| PHIP | 1772.00 | 1723.00 | 1988.00 | 1827.67 | 1764.00 |
| PHKA1 | 725.00 | 697.00 | 777.00 | 733.00 | 659.00 |
| PHKA2 | 725.00 | 737.00 | 843.00 | 768.33 | 671.00 |
| PHKB | 1035.00 | 1108.00 | 1149.00 | 1097.33 | 1088.00 |
| PHKG1 | 21.00 | 24.00 | 33.00 | 26.00 | 13.00 |
| PHKG2 | 734.00 | 777.00 | 840.00 | 783.67 | 789.00 |
| PHLDA1 | 8017.00 | 7583.00 | 8807.00 | 8135.67 | 7430.00 |
| PHLDA3 | 217.00 | 247.00 | 239.00 | 234.33 | 218.00 |
| PHLDB1 | 1899.00 | 1993.00 | 1941.00 | 1944.33 | 1857.00 |
| PHLDB2 | 1019.00 | 865.00 | 1040.00 | 974.67 | 261.00 |
| PHLDB3 | 127.00 | 107.00 | 114.00 | 116.00 | 129.00 |
| PHLPP1 | 374.00 | 388.00 | 398.00 | 386.67 | 249.00 |
| PHLPP2 | 694.00 | 607.00 | 650.00 | 650.33 | 605.00 |
| PHOSPHO2 | 66.00 | 62.00 | 63.00 | 63.67 | 46.00 |
| PHPT1 | 1072.00 | 1078.00 | 1091.00 | 1080.33 | 1272.00 |
| PHRF1 | 31.00 | 30.00 | 31.00 | 30.67 | 50.00 |
| PHTF1 | 512.00 | 503.00 | 482.00 | 499.00 | 537.00 |
| PHTF2 | 757.00 | 688.00 | 730.00 | 725.00 | 901.00 |
| PHYH | 502.00 | 477.00 | 561.00 | 513.33 | 577.00 |
| PHYHD1 | 137.00 | 125.00 | 112.00 | 124.67 | 141.00 |
| PHYHIP | 31.00 | 14.00 | 18.00 | 21.00 | 15.00 |
| PHYKPL | 787.00 | 681.00 | 745.00 | 737.67 | 696.00 |
| PI15 | 5.00 | 1.00 | 6.00 | 4.00 | 2.00 |
| PI4K2A | 1036.00 | 1013.00 | 1111.00 | 1053.33 | 1021.00 |
| PI4K2B | 239.00 | 183.00 | 239.00 | 220.33 | 313.00 |
| PI4KA | 1700.00 | 1608.00 | 1783.00 | 1697.00 | 1419.00 |
| PI4KAP1 | 60.00 | 82.00 | 109.00 | 83.67 | 74.00 |

| | | | | | |
|---|---|---|---|---|---|
| PI4KAP2 | 157.00 | 140.00 | 166.00 | 154.33 | 160.00 |
| PI4KB | 1967.00 | 1769.00 | 1976.00 | 1904.00 | 2013.00 |
| PIANP | 8.00 | 12.00 | 8.00 | 9.33 | 3.00 |
| PIAS1 | 332.00 | 316.00 | 368.00 | 338.67 | 326.00 |
| PIAS2 | 589.00 | 578.00 | 599.00 | 588.67 | 619.00 |
| PIAS3 | 879.00 | 854.00 | 905.00 | 879.33 | 1135.00 |
| PIAS4 | 449.00 | 443.00 | 512.00 | 468.00 | 478.00 |
| PIBF1 | 476.00 | 443.00 | 468.00 | 462.33 | 589.00 |
| PICALM | 2710.00 | 2439.00 | 2891.00 | 2680.00 | 2735.00 |
| PICK1 | 224.00 | 276.00 | 204.00 | 234.67 | 243.00 |
| PID1 | 30.00 | 19.00 | 25.00 | 24.67 | 36.00 |
| PIDD1 | 608.00 | 578.00 | 635.00 | 607.00 | 567.00 |
| PIEZO1 | 8079.00 | 7865.00 | 8297.00 | 8080.33 | 6811.00 |
| PIF1 | 809.00 | 776.00 | 778.00 | 787.67 | 963.00 |
| PIGA | 198.00 | 192.00 | 225.00 | 205.00 | 233.00 |
| PIGB | 559.00 | 551.00 | 507.00 | 539.00 | 561.00 |
| PIGBOS1 | 180.00 | 183.00 | 191.00 | 184.67 | 250.00 |
| PIGC | 694.00 | 699.00 | 625.00 | 672.67 | 836.00 |
| PIGF | 574.00 | 628.00 | 545.00 | 582.33 | 708.00 |
| PIGG | 1176.00 | 1146.00 | 1193.00 | 1171.67 | 1009.00 |
| PIGH | 375.00 | 392.00 | 366.00 | 377.67 | 491.00 |
| PIGK | 2716.00 | 2520.00 | 2585.00 | 2607.00 | 2650.00 |
| PIGL | 232.00 | 234.00 | 266.00 | 244.00 | 262.00 |
| PIGM | 684.00 | 638.00 | 683.00 | 668.33 | 646.00 |
| PIGN | 838.00 | 767.00 | 868.00 | 824.33 | 699.00 |
| PIGO | 2097.00 | 1912.00 | 1964.00 | 1991.00 | 1757.00 |
| PIGP | 220.00 | 205.00 | 227.00 | 217.33 | 300.00 |
| PIGQ | 991.00 | 975.00 | 1038.00 | 1001.33 | 1090.00 |
| PIGS | 1062.00 | 1057.00 | 1012.00 | 1043.67 | 1077.00 |
| PIGT | 4609.00 | 4483.00 | 4664.00 | 4585.33 | 4388.00 |
| PIGU | 1365.00 | 1331.00 | 1348.00 | 1348.00 | 1349.00 |
| PIGV | 784.00 | 740.00 | 769.00 | 764.33 | 841.00 |
| PIGX | 269.00 | 266.00 | 268.00 | 267.67 | 289.00 |
| PIGZ | 187.00 | 171.00 | 183.00 | 180.33 | 205.00 |
| PIH1D1 | 812.00 | 730.00 | 795.00 | 779.00 | 918.00 |
| PIH1D2 | 44.00 | 36.00 | 52.00 | 44.00 | 41.00 |
| PIK3C2A | 1358.00 | 1242.00 | 1476.00 | 1358.67 | 1106.00 |
| PIK3C2B | 21.00 | 16.00 | 8.00 | 15.00 | 16.00 |
| PIK3C3 | 570.00 | 535.00 | 584.00 | 563.00 | 545.00 |
| PIK3CA | 625.00 | 486.00 | 575.00 | 562.00 | 538.00 |
| PIK3CB | 698.00 | 594.00 | 706.00 | 666.00 | 679.00 |
| PIK3CD | 1578.00 | 1435.00 | 1543.00 | 1518.67 | 1271.00 |
| PIK3CD-AS2 | 19.00 | 18.00 | 16.00 | 17.67 | 37.00 |
| PIK3CG | 139.00 | 115.00 | 140.00 | 131.33 | 134.00 |
| PIK3IP1 | 19.00 | 11.00 | 14.00 | 14.67 | 15.00 |
| PIK3R1 | 523.00 | 499.00 | 581.00 | 534.33 | 500.00 |
| PIK3R2 | 1574.00 | 1523.00 | 1606.00 | 1567.67 | 1416.00 |
| PIK3R3 | 61.00 | 59.00 | 53.00 | 57.67 | 54.00 |

| | | | | | |
|---|---|---|---|---|---|
| PIK3R4 | 1501.00 | 1575.00 | 1679.00 | 1585.00 | 1510.00 |
| PIKFYVE | 1912.00 | 1766.00 | 2010.00 | 1896.00 | 1696.00 |
| PILRA | 10.00 | 12.00 | 6.00 | 9.33 | 8.00 |
| PILRB | 21.00 | 28.00 | 20.00 | 23.00 | 35.00 |
| PIM1 | 874.00 | 816.00 | 914.00 | 868.00 | 1090.00 |
| PIM2 | 140.00 | 117.00 | 145.00 | 134.00 | 167.00 |
| PIM3 | 1125.00 | 1059.00 | 1205.00 | 1129.67 | 1492.00 |
| PIN1 | 829.00 | 736.00 | 860.00 | 808.33 | 933.00 |
| PIN4 | 359.00 | 378.00 | 370.00 | 369.00 | 503.00 |
| PIN4P1 | 13.00 | 19.00 | 17.00 | 16.33 | 8.00 |
| PINK1 | 1248.00 | 1273.00 | 1287.00 | 1269.33 | 1039.00 |
| PINK1-AS | 339.00 | 296.00 | 359.00 | 331.33 | 301.00 |
| PINLYP | 65.00 | 65.00 | 87.00 | 72.33 | 35.00 |
| PINX1 | 222.00 | 256.00 | 251.00 | 243.00 | 279.00 |
| PIP4K2A | 1689.00 | 1593.00 | 1866.00 | 1716.00 | 1484.00 |
| PIP4K2C | 783.00 | 786.00 | 865.00 | 811.33 | 834.00 |
| PIP5K1A | 2713.00 | 2378.00 | 2777.00 | 2622.67 | 2693.00 |
| PIP5K1C | 1209.00 | 1103.00 | 1268.00 | 1193.33 | 1059.00 |
| PIP5KL1 | 31.00 | 40.00 | 40.00 | 37.00 | 30.00 |
| PIPSL | 40.00 | 21.00 | 37.00 | 32.67 | 23.00 |
| PIR | 35.00 | 31.00 | 47.00 | 37.67 | 50.00 |
| PISD | 1346.00 | 1282.00 | 1321.00 | 1316.33 | 1246.00 |
| PITHD1 | 611.00 | 553.00 | 664.00 | 609.33 | 934.00 |
| PITPNA | 1678.00 | 1554.00 | 1599.00 | 1610.33 | 1624.00 |
| PITPNA-AS1 | 164.00 | 185.00 | 180.00 | 176.33 | 210.00 |
| PITPNB | 1857.00 | 1750.00 | 1953.00 | 1853.33 | 2235.00 |
| PITPNC1 | 295.00 | 294.00 | 350.00 | 313.00 | 349.00 |
| PITPNM1 | 1118.00 | 1128.00 | 1083.00 | 1109.67 | 907.00 |
| PITPNM2 | 1043.00 | 1077.00 | 1304.00 | 1141.33 | 920.00 |
| PITPNM3 | 29.00 | 29.00 | 42.00 | 33.33 | 33.00 |
| PITRM1 | 4258.00 | 4168.00 | 4751.00 | 4392.33 | 3857.00 |
| PITRM1-AS1 | 13.00 | 10.00 | 14.00 | 12.33 | 13.00 |
| PITX1 | 186.00 | 178.00 | 175.00 | 179.67 | 252.00 |
| PITX2 | 47.00 | 46.00 | 40.00 | 44.33 | 50.00 |
| PITX3 | 3.00 | 1.00 | 0.00 | 1.33 | 1.00 |
| PIWIL4 | 58.00 | 38.00 | 52.00 | 49.33 | 45.00 |
| PJA1 | 901.00 | 788.00 | 891.00 | 860.00 | 933.00 |
| PJA2 | 1876.00 | 1546.00 | 1914.00 | 1778.67 | 1590.00 |
| PKD1 | 4981.00 | 4889.00 | 5496.00 | 5122.00 | 3685.00 |
| PKD1L1 | 3.00 | 7.00 | 6.00 | 5.33 | 8.00 |
| PKD1L2 | 47.00 | 45.00 | 58.00 | 50.00 | 48.00 |
| PKD1P1 | 153.00 | 155.00 | 149.00 | 152.33 | 116.00 |
| PKD1P6 | 262.00 | 332.00 | 317.00 | 303.67 | 266.00 |
| PKD2 | 1409.00 | 1368.00 | 1551.00 | 1442.67 | 1497.00 |
| PKD2L2 | 5.00 | 2.00 | 5.00 | 4.00 | 5.00 |
| PKDCC | 34.00 | 45.00 | 58.00 | 45.67 | 66.00 |
| PKI55 | 302.00 | 297.00 | 316.00 | 305.00 | 317.00 |
| PKIA | 805.00 | 800.00 | 886.00 | 830.33 | 1243.00 |

| | | | | | |
|---|---|---|---|---|---|
| PKIB | 20.00 | 21.00 | 35.00 | 25.33 | 28.00 |
| PKIG | 152.00 | 176.00 | 198.00 | 175.33 | 259.00 |
| PKM | 26262.00 | 24961.00 | 26148.00 | 25790.33 | 26940.00 |
| PKMYT1 | 498.00 | 431.00 | 412.00 | 447.00 | 600.00 |
| PKN1 | 3957.00 | 4027.00 | 4252.00 | 4078.67 | 4204.00 |
| PKN2 | 1933.00 | 1631.00 | 1994.00 | 1852.67 | 1848.00 |
| PKN2-AS1 | 8.00 | 5.00 | 5.00 | 6.00 | 14.00 |
| PKN3 | 673.00 | 755.00 | 768.00 | 732.00 | 886.00 |
| PKNOX1 | 619.00 | 629.00 | 686.00 | 644.67 | 589.00 |
| PKP3 | 39.00 | 37.00 | 61.00 | 45.67 | 64.00 |
| PKP4 | 1972.00 | 1845.00 | 2057.00 | 1958.00 | 2092.00 |
| PLA2G12A | 313.00 | 277.00 | 304.00 | 298.00 | 307.00 |
| PLA2G12B | 3.00 | 1.00 | 0.00 | 1.33 | 0.00 |
| PLA2G15 | 675.00 | 640.00 | 681.00 | 665.33 | 601.00 |
| PLA2G16 | 156.00 | 147.00 | 157.00 | 153.33 | 133.00 |
| PLA2G4A | 389.00 | 369.00 | 356.00 | 371.33 | 305.00 |
| PLA2G4B | 24.00 | 12.00 | 23.00 | 19.67 | 23.00 |
| PLA2G4C | 50.00 | 38.00 | 45.00 | 44.33 | 54.00 |
| PLA2G6 | 316.00 | 339.00 | 347.00 | 334.00 | 285.00 |
| PLA2G7 | 7.00 | 7.00 | 11.00 | 8.33 | 10.00 |
| PLA2R1 | 4.00 | 6.00 | 5.00 | 5.00 | 3.00 |
| PLAA | 1038.00 | 1012.00 | 1078.00 | 1042.67 | 1360.00 |
| PLAC1 | 63.00 | 64.00 | 64.00 | 63.67 | 82.00 |
| PLAC8 | 5.00 | 5.00 | 3.00 | 4.33 | 10.00 |
| PLAC8L1 | 53.00 | 51.00 | 51.00 | 51.67 | 72.00 |
| PLAG1 | 711.00 | 659.00 | 799.00 | 723.00 | 631.00 |
| PLAGL1 | 1188.00 | 1160.00 | 1263.00 | 1203.67 | 1010.00 |
| PLAGL2 | 911.00 | 786.00 | 867.00 | 854.67 | 847.00 |
| PLAT | 42487.00 | 41434.00 | 39963.00 | 41294.67 | 48035.00 |
| PLAU | 23078.00 | 22743.00 | 21550.00 | 22457.00 | 16792.00 |
| PLAUR | 12406.00 | 12817.00 | 11642.00 | 12288.33 | 12789.00 |
| PLB1 | 6.00 | 6.00 | 8.00 | 6.67 | 6.00 |
| PLBD2 | 2127.00 | 2052.00 | 2034.00 | 2071.00 | 1286.00 |
| PLCB1 | 766.00 | 603.00 | 791.00 | 720.00 | 601.00 |
| PLCB2 | 67.00 | 113.00 | 96.00 | 92.00 | 123.00 |
| PLCB3 | 2963.00 | 2918.00 | 3055.00 | 2978.67 | 2418.00 |
| PLCB4 | 184.00 | 215.00 | 194.00 | 197.67 | 180.00 |
| PLCD1 | 30.00 | 63.00 | 39.00 | 44.00 | 38.00 |
| PLCD3 | 294.00 | 249.00 | 271.00 | 271.33 | 255.00 |
| PLCD4 | 12.00 | 16.00 | 28.00 | 18.67 | 46.00 |
| PLCE1 | 780.00 | 682.00 | 780.00 | 747.33 | 545.00 |
| PLCG1 | 4279.00 | 4299.00 | 4836.00 | 4471.33 | 4054.00 |
| PLCG1-AS1 | 20.00 | 7.00 | 14.00 | 13.67 | 9.00 |
| PLCG2 | 7.00 | 3.00 | 2.00 | 4.00 | 4.00 |
| PLCH1 | 79.00 | 67.00 | 64.00 | 70.00 | 71.00 |
| PLCL2 | 191.00 | 193.00 | 208.00 | 197.33 | 185.00 |
| PLCXD2 | 78.00 | 64.00 | 71.00 | 71.00 | 31.00 |
| PLCZ1 | 47.00 | 50.00 | 55.00 | 50.67 | 48.00 |

| | | | | | |
|---|---|---|---|---|---|
| PLD1 | 116.00 | 78.00 | 118.00 | 104.00 | 132.00 |
| PLD2 | 1196.00 | 1275.00 | 1283.00 | 1251.33 | 1151.00 |
| PLD3 | 4135.00 | 4400.00 | 4355.00 | 4296.67 | 3780.00 |
| PLD6 | 465.00 | 405.00 | 403.00 | 424.33 | 536.00 |
| PLEC | 30576.00 | 30685.00 | 32591.00 | 31284.00 | 17659.00 |
| PLEK2 | 33.00 | 41.00 | 47.00 | 40.33 | 34.00 |
| PLEKHA1 | 450.00 | 473.00 | 525.00 | 482.67 | 472.00 |
| PLEKHA2 | 1383.00 | 1161.00 | 1259.00 | 1267.67 | 1518.00 |
| PLEKHA3 | 309.00 | 279.00 | 355.00 | 314.33 | 352.00 |
| PLEKHA4 | 217.00 | 175.00 | 195.00 | 195.67 | 235.00 |
| PLEKHA5 | 1266.00 | 1186.00 | 1310.00 | 1254.00 | 996.00 |
| PLEKHA6 | 55.00 | 63.00 | 58.00 | 58.67 | 23.00 |
| PLEKHA7 | 6.00 | 5.00 | 7.00 | 6.00 | 8.00 |
| PLEKHA8 | 658.00 | 679.00 | 790.00 | 709.00 | 613.00 |
| PLEKHA8P1 | 101.00 | 104.00 | 115.00 | 106.67 | 109.00 |
| PLEKHB2 | 2716.00 | 2631.00 | 2975.00 | 2774.00 | 3162.00 |
| PLEKHF1 | 146.00 | 141.00 | 130.00 | 139.00 | 157.00 |
| PLEKHF2 | 334.00 | 310.00 | 354.00 | 332.67 | 316.00 |
| PLEKHG2 | 1028.00 | 979.00 | 1060.00 | 1022.33 | 952.00 |
| PLEKHG3 | 536.00 | 601.00 | 594.00 | 577.00 | 599.00 |
| PLEKHG4 | 2842.00 | 2931.00 | 3064.00 | 2945.67 | 1954.00 |
| PLEKHG4B | 51.00 | 36.00 | 47.00 | 44.67 | 34.00 |
| PLEKHG5 | 1527.00 | 1565.00 | 1653.00 | 1581.67 | 1568.00 |
| PLEKHH1 | 502.00 | 429.00 | 473.00 | 468.00 | 473.00 |
| PLEKHH2 | 129.00 | 155.00 | 164.00 | 149.33 | 134.00 |
| PLEKHH3 | 438.00 | 482.00 | 498.00 | 472.67 | 535.00 |
| PLEKHJ1 | 897.00 | 989.00 | 972.00 | 952.67 | 1093.00 |
| PLEKHM1 | 149.00 | 142.00 | 152.00 | 147.67 | 147.00 |
| PLEKHM1P | 758.00 | 685.00 | 780.00 | 741.00 | 442.00 |
| PLEKHM2 | 2412.00 | 2297.00 | 2522.00 | 2410.33 | 2151.00 |
| PLEKHM3 | 343.00 | 382.00 | 391.00 | 372.00 | 329.00 |
| PLEKHN1 | 178.00 | 179.00 | 204.00 | 187.00 | 199.00 |
| PLEKHO1 | 517.00 | 522.00 | 565.00 | 534.67 | 570.00 |
| PLEKHO2 | 743.00 | 782.00 | 777.00 | 767.33 | 898.00 |
| PLGRKT | 634.00 | 610.00 | 632.00 | 625.33 | 752.00 |
| PLIN1 | 9.00 | 9.00 | 5.00 | 7.67 | 6.00 |
| PLIN2 | 539.00 | 549.00 | 602.00 | 563.33 | 596.00 |
| PLIN3 | 1517.00 | 1448.00 | 1507.00 | 1490.67 | 1856.00 |
| PLK1 | 1238.00 | 1118.00 | 1208.00 | 1188.00 | 1547.00 |
| PLK2 | 1868.00 | 1737.00 | 1839.00 | 1814.67 | 2418.00 |
| PLK3 | 464.00 | 439.00 | 448.00 | 450.33 | 503.00 |
| PLK4 | 422.00 | 398.00 | 440.00 | 420.00 | 462.00 |
| PLK5 | 3.00 | 4.00 | 1.00 | 2.67 | 1.00 |
| PLOD1 | 7099.00 | 7354.00 | 7159.00 | 7204.00 | 5803.00 |
| PLOD2 | 18904.00 | 17494.00 | 19513.00 | 18637.00 | 18583.00 |
| PLOD3 | 6203.00 | 6405.00 | 6334.00 | 6314.00 | 6204.00 |
| PLP2 | 1778.00 | 1809.00 | 1673.00 | 1753.33 | 1990.00 |
| PLRG1 | 1765.00 | 1627.00 | 1820.00 | 1737.33 | 2034.00 |

| | | | | | |
|---|---|---|---|---|---|
| PLS1 | 177.00 | 133.00 | 192.00 | 167.33 | 206.00 |
| PLS3 | 8023.00 | 7137.00 | 8391.00 | 7850.33 | 8869.00 |
| PLS3-AS1 | 64.00 | 91.00 | 77.00 | 77.33 | 94.00 |
| PLSCR1 | 666.00 | 608.00 | 719.00 | 664.33 | 786.00 |
| PLSCR3 | 8.00 | 16.00 | 17.00 | 13.67 | 20.00 |
| PLSCR4 | 156.00 | 124.00 | 140.00 | 140.00 | 113.00 |
| PLTP | 3450.00 | 3537.00 | 3495.00 | 3494.00 | 3833.00 |
| PLXDC2 | 5.00 | 2.00 | 6.00 | 4.33 | 3.00 |
| PLXNA1 | 5790.00 | 5476.00 | 5818.00 | 5694.67 | 3975.00 |
| PLXNA2 | 4274.00 | 3997.00 | 3826.00 | 4032.33 | 5292.00 |
| PLXNA3 | 732.00 | 748.00 | 904.00 | 794.67 | 694.00 |
| PLXNA4 | 3475.00 | 3216.00 | 3493.00 | 3394.67 | 2005.00 |
| PLXNB1 | 137.00 | 155.00 | 156.00 | 149.33 | 109.00 |
| PLXNB2 | 2072.00 | 2100.00 | 2057.00 | 2076.33 | 1294.00 |
| PLXNB3 | 985.00 | 1043.00 | 1071.00 | 1033.00 | 772.00 |
| PLXNC1 | 140.00 | 121.00 | 154.00 | 138.33 | 131.00 |
| PLXND1 | 3365.00 | 3357.00 | 3583.00 | 3435.00 | 2195.00 |
| PM20D2 | 374.00 | 310.00 | 436.00 | 373.33 | 443.00 |
| PMAIP1 | 1359.00 | 1154.00 | 1476.00 | 1329.67 | 1928.00 |
| PMEL | 4.00 | 6.00 | 10.00 | 6.67 | 7.00 |
| PMEPA1 | 222.00 | 229.00 | 296.00 | 249.00 | 189.00 |
| PMF1 | 222.00 | 180.00 | 202.00 | 201.33 | 265.00 |
| PMF1-BGLAP | 9.00 | 9.00 | 9.00 | 9.00 | 7.00 |
| PMFBP1 | 20.00 | 26.00 | 25.00 | 23.67 | 18.00 |
| PML | 801.00 | 763.00 | 784.00 | 782.67 | 772.00 |
| PMM1 | 420.00 | 437.00 | 400.00 | 419.00 | 541.00 |
| PMM2 | 903.00 | 894.00 | 970.00 | 922.33 | 929.00 |
| PMP22 | 487.00 | 439.00 | 486.00 | 470.67 | 464.00 |
| PMPCA | 2055.00 | 2008.00 | 2027.00 | 2030.00 | 2178.00 |
| PMPCB | 1318.00 | 1292.00 | 1380.00 | 1330.00 | 1457.00 |
| PMS1 | 1259.00 | 1196.00 | 1315.00 | 1256.67 | 1466.00 |
| PMS2 | 538.00 | 507.00 | 561.00 | 535.33 | 533.00 |
| PMS2CL | 130.00 | 150.00 | 164.00 | 148.00 | 165.00 |
| PMS2P1 | 246.00 | 234.00 | 277.00 | 252.33 | 373.00 |
| PMS2P2 | 3.00 | 2.00 | 2.00 | 2.33 | 5.00 |
| PMS2P3 | 167.00 | 200.00 | 181.00 | 182.67 | 153.00 |
| PMS2P4 | 19.00 | 24.00 | 20.00 | 21.00 | 39.00 |
| PMS2P5 | 51.00 | 56.00 | 61.00 | 56.00 | 64.00 |
| PMS2P7 | 11.00 | 15.00 | 9.00 | 11.67 | 11.00 |
| PMS2P9 | 9.00 | 14.00 | 8.00 | 10.33 | 10.00 |
| PMVK | 496.00 | 503.00 | 540.00 | 513.00 | 729.00 |
| PNCK | 4.00 | 1.00 | 0.00 | 1.67 | 3.00 |
| PNISR | 6218.00 | 6223.00 | 6869.00 | 6436.67 | 5898.00 |
| PNKD | 462.00 | 504.00 | 513.00 | 493.00 | 507.00 |
| PNKP | 517.00 | 567.00 | 539.00 | 541.00 | 538.00 |
| PNLDC1 | 3.00 | 6.00 | 3.00 | 4.00 | 2.00 |
| PNMA1 | 1226.00 | 1178.00 | 1257.00 | 1220.33 | 1312.00 |
| PNMA2 | 257.00 | 220.00 | 273.00 | 250.00 | 158.00 |

| | | | | | |
|---|---|---|---|---|---|
| PNMA6A | 14.00 | 9.00 | 6.00 | 9.67 | 9.00 |
| PNMAL1 | 523.00 | 547.00 | 559.00 | 543.00 | 711.00 |
| PNN | 9432.00 | 9454.00 | 9810.00 | 9565.33 | 10340.00 |
| PNO1 | 908.00 | 816.00 | 981.00 | 901.67 | 1048.00 |
| PNP | 2046.00 | 2022.00 | 2183.00 | 2083.67 | 3276.00 |
| PNPLA2 | 1454.00 | 1390.00 | 1515.00 | 1453.00 | 1229.00 |
| PNPLA3 | 286.00 | 256.00 | 354.00 | 298.67 | 183.00 |
| PNPLA4 | 334.00 | 358.00 | 417.00 | 369.67 | 441.00 |
| PNPLA6 | 3120.00 | 3216.00 | 3121.00 | 3152.33 | 2775.00 |
| PNPLA7 | 27.00 | 37.00 | 31.00 | 31.67 | 38.00 |
| PNPLA8 | 526.00 | 461.00 | 545.00 | 510.67 | 466.00 |
| PNPO | 750.00 | 633.00 | 703.00 | 695.33 | 776.00 |
| PNPT1 | 1503.00 | 1444.00 | 1539.00 | 1495.33 | 1754.00 |
| PNRC1 | 278.00 | 249.00 | 299.00 | 275.33 | 226.00 |
| PNRC2 | 1456.00 | 1430.00 | 1503.00 | 1463.00 | 1833.00 |
| POC1A | 413.00 | 424.00 | 439.00 | 425.33 | 558.00 |
| POC1B | 661.00 | 541.00 | 678.00 | 626.67 | 622.00 |
| POC5 | 370.00 | 352.00 | 383.00 | 368.33 | 454.00 |
| PODNL1 | 3.00 | 9.00 | 7.00 | 6.33 | 5.00 |
| PODXL | 4382.00 | 4329.00 | 4514.00 | 4408.33 | 4206.00 |
| PODXL2 | 1890.00 | 2111.00 | 2201.00 | 2067.33 | 2466.00 |
| POFUT1 | 3909.00 | 3561.00 | 3858.00 | 3776.00 | 2811.00 |
| POFUT2 | 1993.00 | 2142.00 | 1982.00 | 2039.00 | 1731.00 |
| POGK | 2700.00 | 2555.00 | 2890.00 | 2715.00 | 3086.00 |
| POGLUT1 | 1295.00 | 1227.00 | 1290.00 | 1270.67 | 1013.00 |
| POGZ | 3539.00 | 3422.00 | 3787.00 | 3582.67 | 3584.00 |
| POLA1 | 526.00 | 475.00 | 511.00 | 504.00 | 586.00 |
| POLA2 | 985.00 | 898.00 | 905.00 | 929.33 | 1221.00 |
| POLB | 420.00 | 413.00 | 424.00 | 419.00 | 491.00 |
| POLD1 | 1271.00 | 1217.00 | 1341.00 | 1276.33 | 1476.00 |
| POLD2 | 2731.00 | 2645.00 | 2778.00 | 2718.00 | 3585.00 |
| POLD3 | 442.00 | 412.00 | 449.00 | 434.33 | 585.00 |
| POLD4 | 633.00 | 727.00 | 727.00 | 695.67 | 827.00 |
| POLDIP2 | 2050.00 | 2079.00 | 2287.00 | 2138.67 | 2630.00 |
| POLDIP3 | 1337.00 | 1268.00 | 1440.00 | 1348.33 | 1225.00 |
| POLE | 1764.00 | 1804.00 | 1801.00 | 1789.67 | 1869.00 |
| POLE2 | 135.00 | 124.00 | 118.00 | 125.67 | 199.00 |
| POLE3 | 1055.00 | 976.00 | 1052.00 | 1027.67 | 1450.00 |
| POLE4 | 482.00 | 500.00 | 537.00 | 506.33 | 751.00 |
| POLG | 2020.00 | 1918.00 | 2201.00 | 2046.33 | 1944.00 |
| POLG2 | 1040.00 | 1073.00 | 1117.00 | 1076.67 | 1028.00 |
| POLH | 314.00 | 264.00 | 320.00 | 299.33 | 297.00 |
| POLI | 917.00 | 878.00 | 1008.00 | 934.33 | 959.00 |
| POLK | 995.00 | 928.00 | 1016.00 | 979.67 | 1009.00 |
| POLL | 1022.00 | 1066.00 | 1024.00 | 1037.33 | 1107.00 |
| POLM | 1656.00 | 1705.00 | 1827.00 | 1729.33 | 1982.00 |
| POLN | 136.00 | 123.00 | 142.00 | 133.67 | 103.00 |
| POLQ | 488.00 | 456.00 | 454.00 | 466.00 | 590.00 |

| | | | | | |
|---|---|---|---|---|---|
| POLR1A | 2166.00 | 1910.00 | 2187.00 | 2087.67 | 1825.00 |
| POLR1B | 1983.00 | 1826.00 | 2124.00 | 1977.67 | 2194.00 |
| POLR1C | 1146.00 | 1165.00 | 1289.00 | 1200.00 | 1398.00 |
| POLR1D | 1709.00 | 1677.00 | 1847.00 | 1744.33 | 2244.00 |
| POLR1E | 652.00 | 690.00 | 749.00 | 697.00 | 951.00 |
| POLR2A | 6623.00 | 6244.00 | 6403.00 | 6423.33 | 5016.00 |
| POLR2B | 3317.00 | 3101.00 | 3630.00 | 3349.33 | 3764.00 |
| POLR2C | 1399.00 | 1247.00 | 1407.00 | 1351.00 | 1905.00 |
| POLR2D | 605.00 | 626.00 | 654.00 | 628.33 | 813.00 |
| POLR2E | 1829.00 | 1841.00 | 1969.00 | 1879.67 | 2282.00 |
| POLR2F | 440.00 | 467.00 | 504.00 | 470.33 | 571.00 |
| POLR2G | 906.00 | 965.00 | 1004.00 | 958.33 | 1280.00 |
| POLR2H | 1468.00 | 1379.00 | 1521.00 | 1456.00 | 1775.00 |
| POLR2I | 265.00 | 260.00 | 296.00 | 273.67 | 392.00 |
| POLR2J | 194.00 | 180.00 | 206.00 | 193.33 | 263.00 |
| POLR2J3 | 44.00 | 30.00 | 27.00 | 33.67 | 23.00 |
| POLR2J4 | 256.00 | 271.00 | 291.00 | 272.67 | 267.00 |
| POLR2K | 747.00 | 749.00 | 838.00 | 778.00 | 1094.00 |
| POLR2L | 909.00 | 899.00 | 889.00 | 899.00 | 1205.00 |
| POLR2M | 51.00 | 36.00 | 45.00 | 44.00 | 48.00 |
| POLR3A | 2615.00 | 2487.00 | 2851.00 | 2651.00 | 2543.00 |
| POLR3B | 642.00 | 580.00 | 633.00 | 618.33 | 644.00 |
| POLR3C | 985.00 | 954.00 | 909.00 | 949.33 | 1154.00 |
| POLR3D | 871.00 | 791.00 | 930.00 | 864.00 | 864.00 |
| POLR3E | 1466.00 | 1439.00 | 1655.00 | 1520.00 | 1638.00 |
| POLR3F | 492.00 | 432.00 | 536.00 | 486.67 | 566.00 |
| POLR3G | 861.00 | 730.00 | 1012.00 | 867.67 | 978.00 |
| POLR3GL | 315.00 | 264.00 | 278.00 | 285.67 | 364.00 |
| POLR3H | 1061.00 | 1132.00 | 1079.00 | 1090.67 | 1196.00 |
| POLR3K | 258.00 | 288.00 | 283.00 | 276.33 | 424.00 |
| POLRMT | 2452.00 | 2508.00 | 2688.00 | 2549.33 | 2357.00 |
| POM121 | 1126.00 | 1098.00 | 1076.00 | 1100.00 | 834.00 |
| POM121C | 1546.00 | 1370.00 | 1435.00 | 1450.33 | 1262.00 |
| POM121L10P | 3.00 | 6.00 | 7.00 | 5.33 | 2.00 |
| POM121L9P | 105.00 | 109.00 | 107.00 | 107.00 | 68.00 |
| POMGNT1 | 2798.00 | 2758.00 | 2808.00 | 2788.00 | 2618.00 |
| POMGNT2 | 477.00 | 489.00 | 554.00 | 506.67 | 510.00 |
| POMK | 25.00 | 34.00 | 29.00 | 29.33 | 19.00 |
| POMP | 1098.00 | 1090.00 | 1057.00 | 1081.67 | 1492.00 |
| POMT1 | 1382.00 | 1429.00 | 1427.00 | 1412.67 | 1307.00 |
| POMT2 | 1650.00 | 1626.00 | 1658.00 | 1644.67 | 1219.00 |
| PON2 | 2183.00 | 2206.00 | 2148.00 | 2179.00 | 2474.00 |
| POP1 | 719.00 | 631.00 | 732.00 | 694.00 | 775.00 |
| POP4 | 430.00 | 423.00 | 475.00 | 442.67 | 564.00 |
| POP5 | 370.00 | 447.00 | 475.00 | 430.67 | 565.00 |
| POP7 | 510.00 | 487.00 | 559.00 | 518.67 | 667.00 |
| POPDC2 | 36.00 | 19.00 | 14.00 | 23.00 | 22.00 |
| POPDC3 | 554.00 | 561.00 | 602.00 | 572.33 | 847.00 |

| | | | | | |
|---|---|---|---|---|---|
| POR | 2246.00 | 2506.00 | 2380.00 | 2377.33 | 2364.00 |
| PORCN | 437.00 | 520.00 | 509.00 | 488.67 | 406.00 |
| POT1 | 848.00 | 808.00 | 957.00 | 871.00 | 1026.00 |
| POT1-AS1 | 5.00 | 13.00 | 6.00 | 8.00 | 20.00 |
| POU2F1 | 940.00 | 844.00 | 944.00 | 909.33 | 788.00 |
| POU2F2 | 1922.00 | 1988.00 | 2086.00 | 1998.67 | 2243.00 |
| POU3F2 | 29.00 | 29.00 | 28.00 | 28.67 | 40.00 |
| POU4F1 | 1077.00 | 1052.00 | 1140.00 | 1089.67 | 1301.00 |
| POU6F1 | 129.00 | 147.00 | 184.00 | 153.33 | 125.00 |
| PP7080 | 152.00 | 166.00 | 174.00 | 164.00 | 180.00 |
| PPA1 | 2768.00 | 2613.00 | 3025.00 | 2802.00 | 3636.00 |
| PPA2 | 669.00 | 639.00 | 710.00 | 672.67 | 830.00 |
| PPAN | 228.00 | 227.00 | 259.00 | 238.00 | 201.00 |
| PPAP2A | 356.00 | 313.00 | 305.00 | 324.67 | 361.00 |
| PPAP2B | 2271.00 | 2123.00 | 2264.00 | 2219.33 | 1831.00 |
| PPAP2C | 1602.00 | 1627.00 | 1513.00 | 1580.67 | 1879.00 |
| PPAPDC1A | 45.00 | 51.00 | 33.00 | 43.00 | 36.00 |
| PPAPDC1B | 1075.00 | 1042.00 | 1103.00 | 1073.33 | 1130.00 |
| PPAPDC2 | 617.00 | 566.00 | 588.00 | 590.33 | 549.00 |
| PPAPDC3 | 62.00 | 55.00 | 72.00 | 63.00 | 61.00 |
| PPARA | 472.00 | 424.00 | 449.00 | 448.33 | 294.00 |
| PPARD | 3129.00 | 3064.00 | 3170.00 | 3121.00 | 3433.00 |
| PPARG | 522.00 | 505.00 | 549.00 | 525.33 | 434.00 |
| PPARGC1A | 6.00 | 13.00 | 21.00 | 13.33 | 17.00 |
| PPARGC1B | 347.00 | 369.00 | 396.00 | 370.67 | 347.00 |
| PPAT | 617.00 | 589.00 | 674.00 | 626.67 | 752.00 |
| PPCDC | 202.00 | 232.00 | 209.00 | 214.33 | 323.00 |
| PPCS | 692.00 | 617.00 | 733.00 | 680.67 | 922.00 |
| PPDPF | 1206.00 | 1106.00 | 1244.00 | 1185.33 | 1566.00 |
| PPFIA1 | 8155.00 | 7894.00 | 8815.00 | 8288.00 | 7574.00 |
| PPFIA2 | 187.00 | 189.00 | 150.00 | 175.33 | 167.00 |
| PPFIA3 | 235.00 | 246.00 | 300.00 | 260.33 | 231.00 |
| PPFIA4 | 10.00 | 7.00 | 10.00 | 9.00 | 8.00 |
| PPFIBP1 | 4256.00 | 3915.00 | 4419.00 | 4196.67 | 3024.00 |
| PPFIBP2 | 24.00 | 26.00 | 25.00 | 25.00 | 34.00 |
| PPHLN1 | 1205.00 | 1167.00 | 1370.00 | 1247.33 | 1249.00 |
| PPIA | 8984.00 | 8812.00 | 9372.00 | 9056.00 | 12491.00 |
| PPIAL4A | 19.00 | 34.00 | 24.00 | 25.67 | 35.00 |
| PPIAP30 | 3.00 | 0.00 | 1.00 | 1.33 | 0.00 |
| PPIB | 12832.00 | 13090.00 | 12226.00 | 12716.00 | 12334.00 |
| PPIC | 677.00 | 701.00 | 646.00 | 674.67 | 784.00 |
| PPID | 815.00 | 780.00 | 894.00 | 829.67 | 1151.00 |
| PPIE | 1011.00 | 1000.00 | 1041.00 | 1017.33 | 1316.00 |
| PPIEL | 162.00 | 177.00 | 176.00 | 171.67 | 140.00 |
| PPIF | 2556.00 | 2476.00 | 2671.00 | 2567.67 | 2894.00 |
| PPIG | 1039.00 | 991.00 | 1122.00 | 1050.67 | 1207.00 |
| PPIH | 703.00 | 671.00 | 670.00 | 681.33 | 964.00 |
| PPIL1 | 984.00 | 854.00 | 1013.00 | 950.33 | 1340.00 |

| | | | | | |
|---|---|---|---|---|---|
| PPIL2 | 881.00 | 791.00 | 844.00 | 838.67 | 924.00 |
| PPIL3 | 497.00 | 451.00 | 524.00 | 490.67 | 644.00 |
| PPIL4 | 784.00 | 741.00 | 786.00 | 770.33 | 956.00 |
| PPIL6 | 50.00 | 33.00 | 61.00 | 48.00 | 58.00 |
| PPIP5K1 | 640.00 | 601.00 | 638.00 | 626.33 | 616.00 |
| PPIP5K2 | 771.00 | 675.00 | 922.00 | 789.33 | 892.00 |
| PPM1A | 1098.00 | 991.00 | 1177.00 | 1088.67 | 1250.00 |
| PPM1B | 650.00 | 595.00 | 689.00 | 644.67 | 714.00 |
| PPM1D | 245.00 | 245.00 | 248.00 | 246.00 | 325.00 |
| PPM1E | 431.00 | 361.00 | 435.00 | 409.00 | 450.00 |
| PPM1F | 2219.00 | 2195.00 | 2239.00 | 2217.67 | 2085.00 |
| PPM1G | 3238.00 | 3082.00 | 3343.00 | 3221.00 | 4054.00 |
| PPM1H | 149.00 | 130.00 | 169.00 | 149.33 | 146.00 |
| PPM1J | 31.00 | 22.00 | 21.00 | 24.67 | 31.00 |
| PPM1K | 464.00 | 356.00 | 484.00 | 434.67 | 493.00 |
| PPM1L | 78.00 | 70.00 | 101.00 | 83.00 | 36.00 |
| PPM1M | 653.00 | 576.00 | 645.00 | 624.67 | 636.00 |
| PPM1N | 17.00 | 35.00 | 22.00 | 24.67 | 23.00 |
| PPME1 | 1053.00 | 1007.00 | 1075.00 | 1045.00 | 1029.00 |
| PPOX | 941.00 | 924.00 | 936.00 | 933.67 | 986.00 |
| PPP1CA | 1871.00 | 1770.00 | 1913.00 | 1851.33 | 2531.00 |
| PPP1CB | 1829.00 | 1562.00 | 2008.00 | 1799.67 | 2202.00 |
| PPP1CC | 4025.00 | 3721.00 | 4328.00 | 4024.67 | 5153.00 |
| PPP1R11 | 85.00 | 86.00 | 85.00 | 85.33 | 83.00 |
| PPP1R12A | 1452.00 | 1316.00 | 1542.00 | 1436.67 | 1423.00 |
| PPP1R12B | 514.00 | 459.00 | 518.00 | 497.00 | 363.00 |
| PPP1R12C | 2112.00 | 2214.00 | 2299.00 | 2208.33 | 2236.00 |
| PPP1R13B | 344.00 | 340.00 | 352.00 | 345.33 | 319.00 |
| PPP1R13L | 323.00 | 329.00 | 305.00 | 319.00 | 295.00 |
| PPP1R14B | 4357.00 | 4366.00 | 4590.00 | 4437.67 | 5589.00 |
| PPP1R15A | 11212.00 | 11454.00 | 11626.00 | 11430.67 | 10516.00 |
| PPP1R15B | 2562.00 | 2338.00 | 2806.00 | 2568.67 | 2696.00 |
| PPP1R16A | 1476.00 | 1482.00 | 1520.00 | 1492.67 | 1674.00 |
| PPP1R18 | 3.00 | 3.00 | 3.00 | 3.00 | 4.00 |
| PPP1R1C | 3.00 | 4.00 | 5.00 | 4.00 | 4.00 |
| PPP1R2 | 702.00 | 632.00 | 721.00 | 685.00 | 778.00 |
| PPP1R21 | 413.00 | 438.00 | 487.00 | 446.00 | 498.00 |
| PPP1R26 | 1392.00 | 1420.00 | 1425.00 | 1412.33 | 1236.00 |
| PPP1R26-AS1 | 59.00 | 59.00 | 64.00 | 60.67 | 54.00 |
| PPP1R2P3 | 24.00 | 15.00 | 18.00 | 19.00 | 24.00 |
| PPP1R32 | 17.00 | 27.00 | 20.00 | 21.33 | 25.00 |
| PPP1R35 | 882.00 | 807.00 | 852.00 | 847.00 | 1154.00 |
| PPP1R36 | 3.00 | 5.00 | 4.00 | 4.00 | 7.00 |
| PPP1R37 | 722.00 | 762.00 | 677.00 | 720.33 | 738.00 |
| PPP1R3B | 551.00 | 484.00 | 640.00 | 558.33 | 865.00 |
| PPP1R3D | 135.00 | 135.00 | 158.00 | 142.67 | 147.00 |
| PPP1R3E | 689.00 | 734.00 | 792.00 | 738.33 | 660.00 |
| PPP1R3F | 228.00 | 255.00 | 284.00 | 255.67 | 306.00 |

| | | | | | |
|---|---|---|---|---|---|
| PPP1R7 | 717.00 | 638.00 | 705.00 | 686.67 | 877.00 |
| PPP1R8 | 980.00 | 855.00 | 1031.00 | 955.33 | 1140.00 |
| PPP1R9A | 357.00 | 318.00 | 367.00 | 347.33 | 376.00 |
| PPP1R9B | 2811.00 | 3048.00 | 3251.00 | 3036.67 | 3324.00 |
| PPP2CA | 3237.00 | 3166.00 | 3422.00 | 3275.00 | 3832.00 |
| PPP2CB | 864.00 | 886.00 | 954.00 | 901.33 | 1089.00 |
| PPP2R1A | 3169.00 | 3033.00 | 3366.00 | 3189.33 | 3801.00 |
| PPP2R1B | 523.00 | 548.00 | 585.00 | 552.00 | 523.00 |
| PPP2R2A | 1430.00 | 1452.00 | 1635.00 | 1505.67 | 1933.00 |
| PPP2R2B | 10.00 | 16.00 | 15.00 | 13.67 | 10.00 |
| PPP2R2D | 2273.00 | 2169.00 | 2412.00 | 2284.67 | 2191.00 |
| PPP2R3A | 332.00 | 291.00 | 400.00 | 341.00 | 294.00 |
| PPP2R3B | 121.00 | 146.00 | 152.00 | 139.67 | 140.00 |
| PPP2R3C | 555.00 | 522.00 | 605.00 | 560.67 | 733.00 |
| PPP2R4 | 2138.00 | 2100.00 | 2262.00 | 2166.67 | 2333.00 |
| PPP2R5A | 599.00 | 547.00 | 641.00 | 595.67 | 673.00 |
| PPP2R5B | 710.00 | 715.00 | 842.00 | 755.67 | 809.00 |
| PPP2R5C | 1474.00 | 1380.00 | 1630.00 | 1494.67 | 2024.00 |
| PPP2R5D | 1088.00 | 1050.00 | 1094.00 | 1077.33 | 1234.00 |
| PPP2R5E | 864.00 | 797.00 | 899.00 | 853.33 | 1033.00 |
| PPP3CA | 712.00 | 687.00 | 754.00 | 717.67 | 610.00 |
| PPP3CB | 1380.00 | 1421.00 | 1485.00 | 1428.67 | 1792.00 |
| PPP3CB-AS1 | 333.00 | 324.00 | 329.00 | 328.67 | 309.00 |
| PPP3CC | 109.00 | 130.00 | 107.00 | 115.33 | 126.00 |
| PPP3R1 | 1140.00 | 1067.00 | 1166.00 | 1124.33 | 1478.00 |
| PPP4C | 1583.00 | 1580.00 | 1576.00 | 1579.67 | 1967.00 |
| PPP4R1 | 2183.00 | 2148.00 | 2442.00 | 2257.67 | 2439.00 |
| PPP4R1L | 340.00 | 318.00 | 362.00 | 340.00 | 255.00 |
| PPP4R2 | 675.00 | 529.00 | 651.00 | 618.33 | 725.00 |
| PPP4R3A | 1711.00 | 1665.00 | 1778.00 | 1718.00 | 1875.00 |
| PPP4R3B | 1673.00 | 1546.00 | 1743.00 | 1654.00 | 1793.00 |
| PPP5C | 1018.00 | 1106.00 | 1144.00 | 1089.33 | 1368.00 |
| PPP6C | 788.00 | 762.00 | 845.00 | 798.33 | 1039.00 |
| PPP6R1 | 2857.00 | 2824.00 | 3015.00 | 2898.67 | 3025.00 |
| PPP6R2 | 1481.00 | 1398.00 | 1468.00 | 1449.00 | 1568.00 |
| PPP6R3 | 2164.00 | 2034.00 | 2302.00 | 2166.67 | 2295.00 |
| PPRC1 | 3981.00 | 3879.00 | 4146.00 | 4002.00 | 3481.00 |
| PPT1 | 4030.00 | 3980.00 | 3936.00 | 3982.00 | 5070.00 |
| PPTC7 | 537.00 | 473.00 | 580.00 | 530.00 | 571.00 |
| PPWD1 | 1073.00 | 1041.00 | 1160.00 | 1091.33 | 1313.00 |
| PQBP1 | 968.00 | 999.00 | 1006.00 | 991.00 | 1173.00 |
| PQLC1 | 552.00 | 540.00 | 572.00 | 554.67 | 641.00 |
| PQLC2 | 675.00 | 694.00 | 693.00 | 687.33 | 736.00 |
| PQLC3 | 154.00 | 149.00 | 132.00 | 145.00 | 171.00 |
| PRADC1 | 124.00 | 130.00 | 134.00 | 129.33 | 193.00 |
| PRAF2 | 715.00 | 788.00 | 694.00 | 732.33 | 891.00 |
| PRAME | 7.00 | 4.00 | 3.00 | 4.67 | 1.00 |
| PRC1 | 1834.00 | 1721.00 | 1761.00 | 1772.00 | 2281.00 |

| | | | | | |
|---|---|---|---|---|---|
| PRC1-AS1 | 20.00 | 19.00 | 27.00 | 22.00 | 17.00 |
| PRCAT47 | 395.00 | 423.00 | 415.00 | 411.00 | 492.00 |
| PRCC | 1329.00 | 1254.00 | 1277.00 | 1286.67 | 1628.00 |
| PRCD | 236.00 | 232.00 | 242.00 | 236.67 | 140.00 |
| PRCP | 1906.00 | 1770.00 | 1894.00 | 1856.67 | 1574.00 |
| PRDM1 | 39.00 | 36.00 | 40.00 | 38.33 | 107.00 |
| PRDM10 | 276.00 | 273.00 | 299.00 | 282.67 | 314.00 |
| PRDM11 | 27.00 | 21.00 | 18.00 | 22.00 | 16.00 |
| PRDM12 | 13.00 | 12.00 | 10.00 | 11.67 | 8.00 |
| PRDM13 | 45.00 | 45.00 | 61.00 | 50.33 | 74.00 |
| PRDM15 | 573.00 | 563.00 | 544.00 | 560.00 | 614.00 |
| PRDM2 | 811.00 | 779.00 | 835.00 | 808.33 | 656.00 |
| PRDM4 | 1735.00 | 1685.00 | 1877.00 | 1765.67 | 1675.00 |
| PRDM5 | 179.00 | 186.00 | 196.00 | 187.00 | 167.00 |
| PRDM6 | 10.00 | 15.00 | 26.00 | 17.00 | 17.00 |
| PRDM8 | 615.00 | 616.00 | 607.00 | 612.67 | 602.00 |
| PRDX1 | 5395.00 | 5357.00 | 5426.00 | 5392.67 | 7643.00 |
| PRDX2 | 1466.00 | 1448.00 | 1513.00 | 1475.67 | 2067.00 |
| PRDX3 | 2729.00 | 2613.00 | 2796.00 | 2712.67 | 3476.00 |
| PRDX4 | 4211.00 | 4307.00 | 4141.00 | 4219.67 | 5015.00 |
| PRDX5 | 4591.00 | 4584.00 | 4691.00 | 4622.00 | 6188.00 |
| PRDX6 | 4073.00 | 4195.00 | 4441.00 | 4236.33 | 5659.00 |
| PREB | 910.00 | 891.00 | 874.00 | 891.67 | 973.00 |
| PRELID1 | 2005.00 | 2021.00 | 2201.00 | 2075.67 | 2984.00 |
| PRELID2 | 130.00 | 125.00 | 127.00 | 127.33 | 140.00 |
| PREP | 785.00 | 809.00 | 902.00 | 832.00 | 908.00 |
| PREPL | 2708.00 | 2445.00 | 2856.00 | 2669.67 | 2645.00 |
| PREX1 | 3181.00 | 3056.00 | 3112.00 | 3116.33 | 3131.00 |
| PRG2 | 6.00 | 2.00 | 9.00 | 5.67 | 1.00 |
| PRICKLE1 | 77.00 | 108.00 | 70.00 | 85.00 | 87.00 |
| PRICKLE2 | 176.00 | 156.00 | 179.00 | 170.33 | 144.00 |
| PRICKLE3 | 340.00 | 358.00 | 343.00 | 347.00 | 255.00 |
| PRICKLE4 | 195.00 | 206.00 | 216.00 | 205.67 | 137.00 |
| PRIM1 | 220.00 | 239.00 | 267.00 | 242.00 | 342.00 |
| PRIM2 | 456.00 | 487.00 | 492.00 | 478.33 | 695.00 |
| PRIMPOL | 335.00 | 363.00 | 338.00 | 345.33 | 496.00 |
| PRKAA1 | 2626.00 | 2153.00 | 2762.00 | 2513.67 | 2388.00 |
| PRKAA2 | 583.00 | 532.00 | 646.00 | 587.00 | 530.00 |
| PRKAB1 | 511.00 | 497.00 | 575.00 | 527.67 | 528.00 |
| PRKAB2 | 1457.00 | 1279.00 | 1379.00 | 1371.67 | 1268.00 |
| PRKACA | 724.00 | 765.00 | 729.00 | 739.33 | 748.00 |
| PRKACB | 1154.00 | 1025.00 | 1202.00 | 1127.00 | 1166.00 |
| PRKAG1 | 1574.00 | 1531.00 | 1672.00 | 1592.33 | 1738.00 |
| PRKAG2 | 2505.00 | 2349.00 | 2645.00 | 2499.67 | 2805.00 |
| PRKAR1A | 3682.00 | 3211.00 | 3756.00 | 3549.67 | 2840.00 |
| PRKAR1B | 402.00 | 449.00 | 407.00 | 419.33 | 411.00 |
| PRKAR2A | 369.00 | 334.00 | 417.00 | 373.33 | 277.00 |
| PRKAR2A-AS1 | 40.00 | 27.00 | 31.00 | 32.67 | 21.00 |

| | | | | | |
|---|---|---|---|---|---|
| PRKAR2B | 120.00 | 119.00 | 128.00 | 122.33 | 197.00 |
| PRKCA | 5762.00 | 5441.00 | 5852.00 | 5685.00 | 6099.00 |
| PRKCD | 786.00 | 739.00 | 765.00 | 763.33 | 882.00 |
| PRKCDBP | 178.00 | 157.00 | 165.00 | 166.67 | 344.00 |
| PRKCE | 685.00 | 609.00 | 660.00 | 651.33 | 760.00 |
| PRKCG | 5.00 | 6.00 | 4.00 | 5.00 | 0.00 |
| PRKCH | 254.00 | 269.00 | 295.00 | 272.67 | 210.00 |
| PRKCI | 1436.00 | 1348.00 | 1599.00 | 1461.00 | 1680.00 |
| PRKCQ-AS1 | 79.00 | 74.00 | 82.00 | 78.33 | 97.00 |
| PRKCSH | 12016.00 | 12505.00 | 11779.00 | 12100.00 | 12437.00 |
| PRKD1 | 236.00 | 211.00 | 233.00 | 226.67 | 215.00 |
| PRKD2 | 472.00 | 460.00 | 484.00 | 472.00 | 377.00 |
| PRKD3 | 1759.00 | 1581.00 | 1862.00 | 1734.00 | 1994.00 |
| PRKDC | 15109.00 | 13788.00 | 15573.00 | 14823.33 | 14128.00 |
| PRKG1 | 151.00 | 133.00 | 193.00 | 159.00 | 112.00 |
| PRKG1-AS1 | 42.00 | 30.00 | 41.00 | 37.67 | 33.00 |
| PRKG2 | 34.00 | 25.00 | 34.00 | 31.00 | 35.00 |
| PRKRA | 389.00 | 422.00 | 454.00 | 421.67 | 520.00 |
| PRKRIP1 | 45.00 | 53.00 | 48.00 | 48.67 | 45.00 |
| PRKRIR | 1390.00 | 1264.00 | 1459.00 | 1371.00 | 1458.00 |
| PRKX | 209.00 | 186.00 | 209.00 | 201.33 | 176.00 |
| PRKXP1 | 438.00 | 443.00 | 484.00 | 455.00 | 380.00 |
| PRLR | 70.00 | 85.00 | 68.00 | 74.33 | 87.00 |
| PRMT1 | 3099.00 | 3047.00 | 3292.00 | 3146.00 | 4214.00 |
| PRMT2 | 1697.00 | 1689.00 | 1680.00 | 1688.67 | 1575.00 |
| PRMT3 | 888.00 | 796.00 | 935.00 | 873.00 | 1066.00 |
| PRMT5 | 2577.00 | 2325.00 | 2776.00 | 2559.33 | 2773.00 |
| PRMT6 | 640.00 | 636.00 | 762.00 | 679.33 | 777.00 |
| PRMT7 | 781.00 | 805.00 | 889.00 | 825.00 | 921.00 |
| PRMT9 | 260.00 | 261.00 | 292.00 | 271.00 | 276.00 |
| PRNP | 12037.00 | 11811.00 | 11433.00 | 11760.33 | 16616.00 |
| PROB1 | 80.00 | 100.00 | 114.00 | 98.00 | 45.00 |
| PROCA1 | 84.00 | 88.00 | 73.00 | 81.67 | 99.00 |
| PROCR | 1960.00 | 1914.00 | 1943.00 | 1939.00 | 2443.00 |
| PROM1 | 4.00 | 1.00 | 3.00 | 2.67 | 3.00 |
| PROM2 | 43.00 | 39.00 | 44.00 | 42.00 | 41.00 |
| PRORSD1P | 47.00 | 64.00 | 40.00 | 50.33 | 64.00 |
| PROS1 | 621.00 | 599.00 | 657.00 | 625.67 | 634.00 |
| PROSC | 826.00 | 810.00 | 929.00 | 855.00 | 1104.00 |
| PROSER1 | 1459.00 | 1237.00 | 1577.00 | 1424.33 | 1469.00 |
| PROSER2 | 68.00 | 65.00 | 75.00 | 69.33 | 61.00 |
| PROSER3 | 362.00 | 364.00 | 382.00 | 369.33 | 344.00 |
| PROX1 | 9.00 | 8.00 | 10.00 | 9.00 | 5.00 |
| PRPF18 | 311.00 | 281.00 | 327.00 | 306.33 | 362.00 |
| PRPF19 | 3481.00 | 3312.00 | 3520.00 | 3437.67 | 4053.00 |
| PRPF3 | 1486.00 | 1522.00 | 1576.00 | 1528.00 | 1672.00 |
| PRPF31 | 21.00 | 36.00 | 25.00 | 27.33 | 15.00 |
| PRPF38A | 1150.00 | 1149.00 | 1296.00 | 1198.33 | 1332.00 |

| | | | | | |
|---|---|---|---|---|---|
| PRPF38B | 3068.00 | 3242.00 | 3325.00 | 3211.67 | 3628.00 |
| PRPF39 | 1740.00 | 1722.00 | 1909.00 | 1790.33 | 1904.00 |
| PRPF4 | 805.00 | 761.00 | 816.00 | 794.00 | 868.00 |
| PRPF40A | 3399.00 | 3093.00 | 3623.00 | 3371.67 | 3749.00 |
| PRPF40B | 466.00 | 502.00 | 526.00 | 498.00 | 504.00 |
| PRPF4B | 3780.00 | 3650.00 | 4160.00 | 3863.33 | 3533.00 |
| PRPF6 | 3076.00 | 2968.00 | 3243.00 | 3095.67 | 3339.00 |
| PRPF8 | 14.00 | 21.00 | 16.00 | 17.00 | 12.00 |
| PRPS1 | 809.00 | 780.00 | 924.00 | 837.67 | 1059.00 |
| PRPS2 | 369.00 | 361.00 | 411.00 | 380.33 | 463.00 |
| PRPSAP1 | 627.00 | 581.00 | 639.00 | 615.67 | 752.00 |
| PRPSAP2 | 674.00 | 555.00 | 693.00 | 640.67 | 731.00 |
| PRR11 | 2330.00 | 2070.00 | 2351.00 | 2250.33 | 2557.00 |
| PRR12 | 922.00 | 928.00 | 966.00 | 938.67 | 875.00 |
| PRR13 | 942.00 | 873.00 | 922.00 | 912.33 | 1268.00 |
| PRR14 | 753.00 | 770.00 | 869.00 | 797.33 | 899.00 |
| PRR14L | 1521.00 | 1515.00 | 1686.00 | 1574.00 | 1477.00 |
| PRR16 | 207.00 | 154.00 | 195.00 | 185.33 | 163.00 |
| PRR19 | 11.00 | 14.00 | 24.00 | 16.33 | 18.00 |
| PRR22 | 172.00 | 186.00 | 212.00 | 190.00 | 242.00 |
| PRR29 | 4.00 | 9.00 | 6.00 | 6.33 | 11.00 |
| PRR34 | 5.00 | 3.00 | 4.00 | 4.00 | 9.00 |
| PRR34-AS1 | 63.00 | 63.00 | 57.00 | 61.00 | 83.00 |
| PRR5L | 41.00 | 32.00 | 34.00 | 35.67 | 57.00 |
| PRR7 | 240.00 | 229.00 | 280.00 | 249.67 | 317.00 |
| PRR7-AS1 | 132.00 | 121.00 | 103.00 | 118.67 | 122.00 |
| PRRC1 | 1345.00 | 1245.00 | 1428.00 | 1339.33 | 1447.00 |
| PRRC2A | 24.00 | 16.00 | 17.00 | 19.00 | 19.00 |
| PRRC2B | 10053.00 | 9416.00 | 10540.00 | 10003.00 | 9027.00 |
| PRRC2C | 10074.00 | 8801.00 | 10441.00 | 9772.00 | 7480.00 |
| PRRG1 | 245.00 | 194.00 | 247.00 | 228.67 | 232.00 |
| PRRG2 | 9.00 | 9.00 | 7.00 | 8.33 | 9.00 |
| PRRG4 | 9.00 | 1.00 | 6.00 | 5.33 | 11.00 |
| PRRT2 | 11.00 | 15.00 | 17.00 | 14.33 | 13.00 |
| PRRT3 | 75.00 | 83.00 | 54.00 | 70.67 | 51.00 |
| PRRT3-AS1 | 61.00 | 66.00 | 54.00 | 60.33 | 64.00 |
| PRRX1 | 1375.00 | 1239.00 | 1539.00 | 1384.33 | 1289.00 |
| PRSS12 | 1220.00 | 1194.00 | 1281.00 | 1231.67 | 1261.00 |
| PRSS23 | 6088.00 | 5836.00 | 6018.00 | 5980.67 | 4407.00 |
| PRSS27 | 24.00 | 16.00 | 21.00 | 20.33 | 20.00 |
| PRSS3 | 736.00 | 685.00 | 807.00 | 742.67 | 859.00 |
| PRSS35 | 5.00 | 3.00 | 5.00 | 4.33 | 6.00 |
| PRSS36 | 6.00 | 8.00 | 3.00 | 5.67 | 3.00 |
| PRSS53 | 470.00 | 504.00 | 517.00 | 497.00 | 472.00 |
| PRTFDC1 | 3.00 | 4.00 | 3.00 | 3.33 | 2.00 |
| PRTG | 704.00 | 703.00 | 801.00 | 736.00 | 684.00 |
| PRUNE | 235.00 | 210.00 | 249.00 | 231.33 | 218.00 |
| PRUNE2 | 11.00 | 12.00 | 7.00 | 10.00 | 10.00 |

| | | | | | |
|---|---|---|---|---|---|
| PRX | 121.00 | 114.00 | 141.00 | 125.33 | 142.00 |
| PSAP | 34245.00 | 35506.00 | 34985.00 | 34912.00 | 29818.00 |
| PSAT1 | 7383.00 | 6540.00 | 7421.00 | 7114.67 | 9158.00 |
| PSCA | 22.00 | 15.00 | 36.00 | 24.33 | 27.00 |
| PSD | 22.00 | 12.00 | 23.00 | 19.00 | 18.00 |
| PSD3 | 824.00 | 755.00 | 793.00 | 790.67 | 455.00 |
| PSD4 | 848.00 | 808.00 | 916.00 | 857.33 | 754.00 |
| PSEN1 | 1536.00 | 1486.00 | 1624.00 | 1548.67 | 1593.00 |
| PSEN2 | 1076.00 | 1087.00 | 1136.00 | 1099.67 | 1273.00 |
| PSENEN | 807.00 | 846.00 | 745.00 | 799.33 | 1033.00 |
| PSG1 | 30.00 | 35.00 | 54.00 | 39.67 | 36.00 |
| PSG11 | 3.00 | 9.00 | 5.00 | 5.67 | 29.00 |
| PSG2 | 6.00 | 5.00 | 4.00 | 5.00 | 7.00 |
| PSG4 | 2456.00 | 2363.00 | 2520.00 | 2446.33 | 2340.00 |
| PSG5 | 102.00 | 99.00 | 105.00 | 102.00 | 116.00 |
| PSG6 | 29.00 | 17.00 | 29.00 | 25.00 | 27.00 |
| PSG7 | 7.00 | 13.00 | 8.00 | 9.33 | 32.00 |
| PSG9 | 33.00 | 26.00 | 27.00 | 28.67 | 45.00 |
| PSIP1 | 2041.00 | 1919.00 | 2264.00 | 2074.67 | 2820.00 |
| PSKH1 | 549.00 | 500.00 | 534.00 | 527.67 | 660.00 |
| PSMA1 | 3122.00 | 3067.00 | 3330.00 | 3173.00 | 4378.00 |
| PSMA2 | 1949.00 | 1862.00 | 2053.00 | 1954.67 | 2783.00 |
| PSMA3 | 1250.00 | 1285.00 | 1268.00 | 1267.67 | 1659.00 |
| PSMA3-AS1 | 1436.00 | 1437.00 | 1482.00 | 1451.67 | 1648.00 |
| PSMA4 | 2067.00 | 1889.00 | 2086.00 | 2014.00 | 2682.00 |
| PSMA5 | 1997.00 | 1937.00 | 1993.00 | 1975.67 | 2575.00 |
| PSMA6 | 1825.00 | 1730.00 | 1803.00 | 1786.00 | 2174.00 |
| PSMA7 | 4480.00 | 4487.00 | 4670.00 | 4545.67 | 5805.00 |
| PSMB1 | 2018.00 | 1945.00 | 2097.00 | 2020.00 | 2803.00 |
| PSMB10 | 274.00 | 254.00 | 281.00 | 269.67 | 320.00 |
| PSMB2 | 1751.00 | 1617.00 | 1802.00 | 1723.33 | 2258.00 |
| PSMB4 | 3634.00 | 3548.00 | 3688.00 | 3623.33 | 5077.00 |
| PSMB5 | 2257.00 | 2289.00 | 2331.00 | 2292.33 | 3019.00 |
| PSMB6 | 2096.00 | 2115.00 | 2064.00 | 2091.67 | 2938.00 |
| PSMB7 | 1871.00 | 1876.00 | 1981.00 | 1909.33 | 2520.00 |
| PSMC1 | 1116.00 | 1085.00 | 1207.00 | 1136.00 | 1482.00 |
| PSMC2 | 1550.00 | 1513.00 | 1591.00 | 1551.33 | 2115.00 |
| PSMC3 | 1975.00 | 1884.00 | 1971.00 | 1943.33 | 2480.00 |
| PSMC3IP | 306.00 | 339.00 | 301.00 | 315.33 | 538.00 |
| PSMC4 | 1608.00 | 1575.00 | 1768.00 | 1650.33 | 2147.00 |
| PSMC5 | 3214.00 | 3127.00 | 3395.00 | 3245.33 | 4458.00 |
| PSMC6 | 1461.00 | 1373.00 | 1449.00 | 1427.67 | 1681.00 |
| PSMD1 | 2433.00 | 2319.00 | 2564.00 | 2438.67 | 2647.00 |
| PSMD10 | 1246.00 | 1067.00 | 1121.00 | 1144.67 | 1412.00 |
| PSMD11 | 1904.00 | 1828.00 | 1895.00 | 1875.67 | 2267.00 |
| PSMD12 | 1525.00 | 1343.00 | 1485.00 | 1451.00 | 1704.00 |
| PSMD13 | 1912.00 | 1947.00 | 1929.00 | 1929.33 | 2623.00 |
| PSMD14 | 2459.00 | 2441.00 | 2633.00 | 2511.00 | 3457.00 |

| | | | | | |
|---|---|---|---|---|---|
| PSMD2 | 13390.00 | 12925.00 | 13843.00 | 13386.00 | 12614.00 |
| PSMD3 | 2449.00 | 2486.00 | 2573.00 | 2502.67 | 3231.00 |
| PSMD4 | 2507.00 | 2342.00 | 2616.00 | 2488.33 | 3414.00 |
| PSMD5 | 748.00 | 723.00 | 808.00 | 759.67 | 938.00 |
| PSMD5-AS1 | 703.00 | 668.00 | 635.00 | 668.67 | 736.00 |
| PSMD6 | 1415.00 | 1242.00 | 1455.00 | 1370.67 | 1802.00 |
| PSMD6-AS2 | 129.00 | 102.00 | 138.00 | 123.00 | 90.00 |
| PSMD7 | 1767.00 | 1797.00 | 1894.00 | 1819.33 | 2438.00 |
| PSMD8 | 1773.00 | 1730.00 | 1938.00 | 1813.67 | 2369.00 |
| PSMD9 | 871.00 | 807.00 | 923.00 | 867.00 | 1209.00 |
| PSME1 | 1016.00 | 1020.00 | 1129.00 | 1055.00 | 1235.00 |
| PSME2 | 1035.00 | 988.00 | 1045.00 | 1022.67 | 1325.00 |
| PSME3 | 2443.00 | 2179.00 | 2488.00 | 2370.00 | 2572.00 |
| PSME4 | 2415.00 | 2335.00 | 2826.00 | 2525.33 | 2417.00 |
| PSMF1 | 1964.00 | 1901.00 | 1956.00 | 1940.33 | 2187.00 |
| PSMG1 | 762.00 | 716.00 | 782.00 | 753.33 | 980.00 |
| PSMG2 | 1075.00 | 1023.00 | 1197.00 | 1098.33 | 1367.00 |
| PSMG3 | 1505.00 | 1455.00 | 1619.00 | 1526.33 | 1931.00 |
| PSMG3-AS1 | 148.00 | 151.00 | 174.00 | 157.67 | 130.00 |
| PSMG4 | 293.00 | 328.00 | 342.00 | 321.00 | 418.00 |
| PSPC1 | 659.00 | 656.00 | 629.00 | 648.00 | 737.00 |
| PSPH | 1077.00 | 1044.00 | 1053.00 | 1058.00 | 1459.00 |
| PSPN | 17.00 | 19.00 | 15.00 | 17.00 | 13.00 |
| PSRC1 | 628.00 | 669.00 | 664.00 | 653.67 | 826.00 |
| PSTK | 128.00 | 139.00 | 127.00 | 131.33 | 138.00 |
| PSTPIP2 | 143.00 | 114.00 | 149.00 | 135.33 | 184.00 |
| PTAFR | 4.00 | 3.00 | 12.00 | 6.33 | 2.00 |
| PTAR1 | 1532.00 | 1344.00 | 1641.00 | 1505.67 | 1234.00 |
| PTBP1 | 7521.00 | 6933.00 | 7602.00 | 7352.00 | 8475.00 |
| PTBP2 | 739.00 | 692.00 | 840.00 | 757.00 | 887.00 |
| PTBP3 | 1251.00 | 1033.00 | 1367.00 | 1217.00 | 1240.00 |
| PTCD1 | 19.00 | 14.00 | 19.00 | 17.33 | 9.00 |
| PTCD2 | 361.00 | 309.00 | 361.00 | 343.67 | 311.00 |
| PTCD3 | 2421.00 | 2325.00 | 2669.00 | 2471.67 | 2645.00 |
| PTCH1 | 144.00 | 141.00 | 171.00 | 152.00 | 128.00 |
| PTCH2 | 9.00 | 27.00 | 13.00 | 16.33 | 8.00 |
| PTDSS1 | 3763.00 | 3834.00 | 3718.00 | 3771.67 | 3851.00 |
| PTDSS2 | 188.00 | 162.00 | 214.00 | 188.00 | 175.00 |
| PTEN | 1199.00 | 1051.00 | 1336.00 | 1195.33 | 1450.00 |
| PTENP1 | 3.00 | 8.00 | 10.00 | 7.00 | 11.00 |
| PTENP1-AS | 7.00 | 3.00 | 3.00 | 4.33 | 6.00 |
| PTER | 578.00 | 557.00 | 601.00 | 578.67 | 696.00 |
| PTGER1 | 23.00 | 22.00 | 20.00 | 21.67 | 23.00 |
| PTGER2 | 111.00 | 104.00 | 121.00 | 112.00 | 66.00 |
| PTGER3 | 8.00 | 5.00 | 11.00 | 8.00 | 5.00 |
| PTGER4 | 188.00 | 156.00 | 140.00 | 161.33 | 236.00 |
| PTGES2 | 1375.00 | 1351.00 | 1514.00 | 1413.33 | 1544.00 |
| PTGES2-AS1 | 8.00 | 20.00 | 9.00 | 12.33 | 9.00 |

| | | | | | |
|---|---|---|---|---|---|
| PTGES3 | 6096.00 | 5810.00 | 6060.00 | 5988.67 | 8294.00 |
| PTGFR | 1266.00 | 1124.00 | 1266.00 | 1218.67 | 1032.00 |
| PTGFRN | 2468.00 | 2355.00 | 2407.00 | 2410.00 | 2196.00 |
| PTGIR | 19.00 | 15.00 | 14.00 | 16.00 | 15.00 |
| PTGR1 | 848.00 | 919.00 | 950.00 | 905.67 | 1050.00 |
| PTGR2 | 498.00 | 474.00 | 490.00 | 487.33 | 501.00 |
| PTGS1 | 765.00 | 781.00 | 714.00 | 753.33 | 209.00 |
| PTGS2 | 465.00 | 454.00 | 441.00 | 453.33 | 477.00 |
| PTH2R | 3.00 | 0.00 | 1.00 | 1.33 | 0.00 |
| PTHLH | 65.00 | 54.00 | 63.00 | 60.67 | 45.00 |
| PTK2 | 3550.00 | 3407.00 | 3873.00 | 3610.00 | 3401.00 |
| PTK2B | 70.00 | 71.00 | 72.00 | 71.00 | 69.00 |
| PTK6 | 8.00 | 8.00 | 9.00 | 8.33 | 7.00 |
| PTK7 | 484.00 | 551.00 | 509.00 | 514.67 | 444.00 |
| PTMA | 9008.00 | 9123.00 | 9528.00 | 9219.67 | 14112.00 |
| PTMS | 1455.00 | 1543.00 | 1473.00 | 1490.33 | 1891.00 |
| PTOV1 | 1209.00 | 1275.00 | 1363.00 | 1282.33 | 1391.00 |
| PTOV1-AS1 | 7.00 | 9.00 | 11.00 | 9.00 | 7.00 |
| PTOV1-AS2 | 265.00 | 319.00 | 298.00 | 294.00 | 303.00 |
| PTP4A1 | 1938.00 | 1772.00 | 2087.00 | 1932.33 | 2565.00 |
| PTP4A2 | 4515.00 | 4124.00 | 4854.00 | 4497.67 | 5941.00 |
| PTP4A3 | 7.00 | 16.00 | 16.00 | 13.00 | 9.00 |
| PTPDC1 | 1118.00 | 999.00 | 1095.00 | 1070.67 | 1062.00 |
| PTPMT1 | 645.00 | 671.00 | 622.00 | 646.00 | 777.00 |
| PTPN1 | 2339.00 | 2160.00 | 2346.00 | 2281.67 | 2310.00 |
| PTPN11 | 4085.00 | 3683.00 | 4290.00 | 4019.33 | 4478.00 |
| PTPN12 | 2627.00 | 2317.00 | 2781.00 | 2575.00 | 3216.00 |
| PTPN13 | 898.00 | 777.00 | 938.00 | 871.00 | 833.00 |
| PTPN14 | 4892.00 | 4296.00 | 5194.00 | 4794.00 | 3942.00 |
| PTPN18 | 1411.00 | 1511.00 | 1536.00 | 1486.00 | 1519.00 |
| PTPN2 | 1172.00 | 1041.00 | 1246.00 | 1153.00 | 1534.00 |
| PTPN20 | 7.00 | 4.00 | 5.00 | 5.33 | 8.00 |
| PTPN21 | 521.00 | 512.00 | 579.00 | 537.33 | 548.00 |
| PTPN22 | 441.00 | 453.00 | 486.00 | 460.00 | 542.00 |
| PTPN23 | 2158.00 | 2098.00 | 2223.00 | 2159.67 | 1694.00 |
| PTPN3 | 308.00 | 331.00 | 359.00 | 332.67 | 311.00 |
| PTPN4 | 295.00 | 291.00 | 341.00 | 309.00 | 383.00 |
| PTPN7 | 17.00 | 8.00 | 15.00 | 13.33 | 32.00 |
| PTPN9 | 1103.00 | 1024.00 | 1228.00 | 1118.33 | 1129.00 |
| PTPRA | 1944.00 | 1881.00 | 1981.00 | 1935.33 | 2247.00 |
| PTPRB | 523.00 | 523.00 | 610.00 | 552.00 | 525.00 |
| PTPRCAP | 68.00 | 59.00 | 76.00 | 67.67 | 61.00 |
| PTPRD | 164.00 | 140.00 | 172.00 | 158.67 | 122.00 |
| PTPRE | 670.00 | 673.00 | 652.00 | 665.00 | 575.00 |
| PTPRF | 17001.00 | 16229.00 | 17306.00 | 16845.33 | 15142.00 |
| PTPRG | 1607.00 | 1374.00 | 1616.00 | 1532.33 | 1361.00 |
| PTPRG-AS1 | 50.00 | 40.00 | 60.00 | 50.00 | 67.00 |
| PTPRH | 103.00 | 99.00 | 132.00 | 111.33 | 145.00 |

| | | | | | |
|---|---|---|---|---|---|
| PTPRJ | 789.00 | 896.00 | 825.00 | 836.67 | 717.00 |
| PTPRK | 6.00 | 6.00 | 2.00 | 4.67 | 3.00 |
| PTPRM | 964.00 | 915.00 | 1012.00 | 963.67 | 736.00 |
| PTPRN | 3.00 | 1.00 | 0.00 | 1.33 | 0.00 |
| PTPRN2 | 305.00 | 281.00 | 294.00 | 293.33 | 318.00 |
| PTPRO | 414.00 | 362.00 | 409.00 | 395.00 | 302.00 |
| PTPRQ | 4.00 | 2.00 | 3.00 | 3.00 | 3.00 |
| PTPRR | 18.00 | 18.00 | 20.00 | 18.67 | 16.00 |
| PTPRS | 2111.00 | 2051.00 | 2282.00 | 2148.00 | 1864.00 |
| PTPRU | 4.00 | 1.00 | 1.00 | 2.00 | 1.00 |
| PTPRZ1 | 102.00 | 96.00 | 81.00 | 93.00 | 246.00 |
| PTRF | 3687.00 | 3516.00 | 3710.00 | 3637.67 | 4580.00 |
| PTRH1 | 152.00 | 180.00 | 147.00 | 159.67 | 172.00 |
| PTRH2 | 1135.00 | 1140.00 | 1079.00 | 1118.00 | 1383.00 |
| PTRHD1 | 170.00 | 134.00 | 162.00 | 155.33 | 229.00 |
| PTS | 372.00 | 362.00 | 370.00 | 368.00 | 497.00 |
| PTTG1 | 1129.00 | 1046.00 | 1036.00 | 1070.33 | 1575.00 |
| PTTG1IP | 13974.00 | 13442.00 | 13149.00 | 13521.67 | 16638.00 |
| PTTG3P | 7.00 | 4.00 | 4.00 | 5.00 | 9.00 |
| PTX3 | 35851.00 | 36057.00 | 34630.00 | 35512.67 | 45384.00 |
| PUF60 | 76.00 | 64.00 | 56.00 | 65.33 | 108.00 |
| PUM1 | 2205.00 | 2093.00 | 2304.00 | 2200.67 | 2154.00 |
| PUM2 | 2676.00 | 2454.00 | 2944.00 | 2691.33 | 3014.00 |
| PURA | 493.00 | 434.00 | 545.00 | 490.67 | 474.00 |
| PURB | 2734.00 | 2523.00 | 2960.00 | 2739.00 | 2626.00 |
| PURG | 20.00 | 18.00 | 31.00 | 23.00 | 35.00 |
| PUS1 | 849.00 | 932.00 | 997.00 | 926.00 | 1117.00 |
| PUS10 | 169.00 | 140.00 | 159.00 | 156.00 | 137.00 |
| PUS3 | 144.00 | 126.00 | 167.00 | 145.67 | 212.00 |
| PUS7 | 1050.00 | 1017.00 | 1117.00 | 1061.33 | 1176.00 |
| PUS7L | 424.00 | 402.00 | 448.00 | 424.67 | 542.00 |
| PUSL1 | 142.00 | 117.00 | 172.00 | 143.67 | 174.00 |
| PVR | 9265.00 | 9129.00 | 9374.00 | 9256.00 | 5842.00 |
| PVRIG | 5.00 | 4.00 | 2.00 | 3.67 | 2.00 |
| PVRL1 | 733.00 | 747.00 | 727.00 | 735.67 | 889.00 |
| PVRL2 | 2477.00 | 2549.00 | 2343.00 | 2456.33 | 2305.00 |
| PVRL3 | 977.00 | 855.00 | 974.00 | 935.33 | 1025.00 |
| PVRL3-AS1 | 5.00 | 2.00 | 2.00 | 3.00 | 15.00 |
| PVRL4 | 3.00 | 6.00 | 3.00 | 4.00 | 1.00 |
| PVT1 | 1347.00 | 1226.00 | 1387.00 | 1320.00 | 1402.00 |
| PWAR1 | 8.00 | 7.00 | 9.00 | 8.00 | 4.00 |
| PWAR5 | 71.00 | 58.00 | 60.00 | 63.00 | 52.00 |
| PWARSN | 21.00 | 17.00 | 21.00 | 19.67 | 17.00 |
| PWP1 | 1553.00 | 1409.00 | 1635.00 | 1532.33 | 1745.00 |
| PWP2 | 318.00 | 274.00 | 325.00 | 305.67 | 377.00 |
| PWWP2A | 897.00 | 864.00 | 1022.00 | 927.67 | 1485.00 |
| PWWP2B | 309.00 | 336.00 | 396.00 | 347.00 | 340.00 |
| PXDC1 | 1085.00 | 1095.00 | 1213.00 | 1131.00 | 1221.00 |

| | | | | | |
|---|---|---|---|---|---|
| PXDN | 7453.00 | 7252.00 | 7474.00 | 7393.00 | 6092.00 |
| PXDNL | 4.00 | 6.00 | 9.00 | 6.33 | 5.00 |
| PXK | 943.00 | 831.00 | 887.00 | 887.00 | 681.00 |
| PXMP2 | 171.00 | 177.00 | 202.00 | 183.33 | 255.00 |
| PXMP4 | 204.00 | 204.00 | 201.00 | 203.00 | 193.00 |
| PXN | 9075.00 | 8847.00 | 9442.00 | 9121.33 | 8735.00 |
| PXN-AS1 | 28.00 | 23.00 | 26.00 | 25.67 | 41.00 |
| PXYLP1 | 200.00 | 180.00 | 225.00 | 201.67 | 231.00 |
| PYCARD-AS1 | 3.00 | 2.00 | 0.00 | 1.67 | 0.00 |
| PYCR1 | 4941.00 | 4762.00 | 5231.00 | 4978.00 | 5166.00 |
| PYCR2 | 2502.00 | 2577.00 | 2628.00 | 2569.00 | 3380.00 |
| PYGB | 4864.00 | 4783.00 | 5286.00 | 4977.67 | 4545.00 |
| PYGL | 95.00 | 98.00 | 109.00 | 100.67 | 200.00 |
| PYGM | 11.00 | 23.00 | 17.00 | 17.00 | 12.00 |
| PYGO1 | 338.00 | 316.00 | 410.00 | 354.67 | 340.00 |
| PYGO2 | 1184.00 | 1090.00 | 1219.00 | 1164.33 | 1317.00 |
| PYROXD1 | 469.00 | 435.00 | 560.00 | 488.00 | 646.00 |
| PYROXD2 | 454.00 | 487.00 | 470.00 | 470.33 | 441.00 |
| PYY | 18.00 | 16.00 | 21.00 | 18.33 | 30.00 |
| PYY2 | 6.00 | 11.00 | 4.00 | 7.00 | 1.00 |
| QARS | 4002.00 | 4089.00 | 4274.00 | 4121.67 | 4358.00 |
| QDPR | 364.00 | 375.00 | 348.00 | 362.33 | 474.00 |
| QKI | 3013.00 | 2678.00 | 3281.00 | 2990.67 | 2783.00 |
| QPCT | 893.00 | 902.00 | 946.00 | 913.67 | 863.00 |
| QPCTL | 639.00 | 629.00 | 671.00 | 646.33 | 625.00 |
| QPRT | 174.00 | 142.00 | 171.00 | 162.33 | 224.00 |
| QRICH1 | 1976.00 | 1945.00 | 2010.00 | 1977.00 | 2090.00 |
| QRICH2 | 217.00 | 223.00 | 262.00 | 234.00 | 185.00 |
| QRSL1 | 554.00 | 536.00 | 575.00 | 555.00 | 625.00 |
| QSER1 | 1737.00 | 1570.00 | 1797.00 | 1701.33 | 1917.00 |
| QSOX1 | 39887.00 | 39340.00 | 38776.00 | 39334.33 | 29417.00 |
| QSOX2 | 3662.00 | 3820.00 | 3846.00 | 3776.00 | 3468.00 |
| QTRT1 | 1497.00 | 1643.00 | 1651.00 | 1597.00 | 1528.00 |
| QTRTD1 | 1384.00 | 1201.00 | 1339.00 | 1308.00 | 1540.00 |
| R3HCC1 | 233.00 | 222.00 | 271.00 | 242.00 | 351.00 |
| R3HCC1L | 536.00 | 567.00 | 556.00 | 553.00 | 607.00 |
| R3HDM1 | 1144.00 | 1121.00 | 1281.00 | 1182.00 | 1186.00 |
| R3HDM2 | 2420.00 | 2420.00 | 2798.00 | 2546.00 | 2493.00 |
| R3HDM4 | 1140.00 | 1233.00 | 1270.00 | 1214.33 | 1323.00 |
| RAB10 | 2613.00 | 2325.00 | 2882.00 | 2606.67 | 2804.00 |
| RAB11A | 1143.00 | 1032.00 | 1167.00 | 1114.00 | 1445.00 |
| RAB11B | 928.00 | 962.00 | 978.00 | 956.00 | 1269.00 |
| RAB11B-AS1 | 26.00 | 27.00 | 39.00 | 30.67 | 41.00 |
| RAB11FIP1 | 277.00 | 256.00 | 271.00 | 268.00 | 293.00 |
| RAB11FIP2 | 831.00 | 812.00 | 1020.00 | 887.67 | 895.00 |
| RAB11FIP3 | 16.00 | 22.00 | 12.00 | 16.67 | 30.00 |
| RAB11FIP4 | 4.00 | 7.00 | 10.00 | 7.00 | 7.00 |
| RAB11FIP5 | 1637.00 | 1555.00 | 1792.00 | 1661.33 | 1441.00 |

| | | | | | |
|---|---|---|---|---|---|
| RAB12 | 1149.00 | 1044.00 | 1257.00 | 1150.00 | 1361.00 |
| RAB13 | 2948.00 | 2994.00 | 3302.00 | 3081.33 | 2991.00 |
| RAB14 | 1605.00 | 1307.00 | 1665.00 | 1525.67 | 1802.00 |
| RAB15 | 199.00 | 189.00 | 214.00 | 200.67 | 245.00 |
| RAB18 | 1292.00 | 1122.00 | 1379.00 | 1264.33 | 1536.00 |
| RAB1A | 1866.00 | 1748.00 | 2080.00 | 1898.00 | 2393.00 |
| RAB1B | 3526.00 | 3428.00 | 3718.00 | 3557.33 | 4028.00 |
| RAB20 | 18.00 | 19.00 | 27.00 | 21.33 | 25.00 |
| RAB21 | 635.00 | 629.00 | 686.00 | 650.00 | 721.00 |
| RAB22A | 1363.00 | 1265.00 | 1379.00 | 1335.67 | 1314.00 |
| RAB23 | 684.00 | 579.00 | 737.00 | 666.67 | 647.00 |
| RAB24 | 1328.00 | 1300.00 | 1392.00 | 1340.00 | 1509.00 |
| RAB26 | 32.00 | 37.00 | 46.00 | 38.33 | 34.00 |
| RAB27A | 176.00 | 127.00 | 161.00 | 154.67 | 172.00 |
| RAB27B | 149.00 | 136.00 | 156.00 | 147.00 | 207.00 |
| RAB28 | 295.00 | 294.00 | 328.00 | 305.67 | 328.00 |
| RAB29 | 1200.00 | 1204.00 | 1239.00 | 1214.33 | 1240.00 |
| RAB2A | 1827.00 | 1659.00 | 2021.00 | 1835.67 | 2233.00 |
| RAB2B | 440.00 | 420.00 | 495.00 | 451.67 | 509.00 |
| RAB30 | 29.00 | 27.00 | 31.00 | 29.00 | 60.00 |
| RAB30-AS1 | 58.00 | 63.00 | 76.00 | 65.67 | 77.00 |
| RAB31 | 1296.00 | 1114.00 | 1411.00 | 1273.67 | 1349.00 |
| RAB32 | 868.00 | 780.00 | 908.00 | 852.00 | 1071.00 |
| RAB33B | 127.00 | 118.00 | 168.00 | 137.67 | 161.00 |
| RAB34 | 1642.00 | 1640.00 | 1821.00 | 1701.00 | 1935.00 |
| RAB35 | 1349.00 | 1248.00 | 1303.00 | 1300.00 | 1526.00 |
| RAB36 | 128.00 | 95.00 | 82.00 | 101.67 | 80.00 |
| RAB38 | 228.00 | 202.00 | 248.00 | 226.00 | 237.00 |
| RAB39A | 18.00 | 20.00 | 20.00 | 19.33 | 22.00 |
| RAB3A | 16.00 | 23.00 | 20.00 | 19.67 | 35.00 |
| RAB3B | 6810.00 | 6231.00 | 6979.00 | 6673.33 | 7781.00 |
| RAB3C | 8.00 | 7.00 | 9.00 | 8.00 | 7.00 |
| RAB3D | 43.00 | 38.00 | 34.00 | 38.33 | 60.00 |
| RAB3GAP1 | 1835.00 | 1771.00 | 1945.00 | 1850.33 | 1960.00 |
| RAB3GAP2 | 1755.00 | 1576.00 | 1821.00 | 1717.33 | 1672.00 |
| RAB3IL1 | 10.00 | 9.00 | 5.00 | 8.00 | 12.00 |
| RAB3IP | 594.00 | 541.00 | 683.00 | 606.00 | 775.00 |
| RAB40A | 8.00 | 5.00 | 14.00 | 9.00 | 7.00 |
| RAB40AL | 6.00 | 2.00 | 2.00 | 3.33 | 5.00 |
| RAB40B | 178.00 | 153.00 | 163.00 | 164.67 | 197.00 |
| RAB40C | 519.00 | 486.00 | 508.00 | 504.33 | 465.00 |
| RAB43 | 35.00 | 43.00 | 61.00 | 46.33 | 20.00 |
| RAB4A | 340.00 | 280.00 | 392.00 | 337.33 | 514.00 |
| RAB5A | 1153.00 | 963.00 | 1158.00 | 1091.33 | 1268.00 |
| RAB5B | 1405.00 | 1257.00 | 1452.00 | 1371.33 | 1326.00 |
| RAB5C | 1845.00 | 1784.00 | 2044.00 | 1891.00 | 2091.00 |
| RAB6A | 1828.00 | 1644.00 | 2000.00 | 1824.00 | 1904.00 |
| RAB6B | 49.00 | 44.00 | 43.00 | 45.33 | 78.00 |

| | | | | | |
|---|---|---|---|---|---|
| RAB7A | 5507.00 | 5211.00 | 5701.00 | 5473.00 | 6588.00 |
| RAB8A | 815.00 | 780.00 | 809.00 | 801.33 | 978.00 |
| RAB8B | 858.00 | 718.00 | 957.00 | 844.33 | 641.00 |
| RAB9A | 565.00 | 600.00 | 607.00 | 590.67 | 625.00 |
| RAB9B | 14.00 | 20.00 | 7.00 | 13.67 | 36.00 |
| RABAC1 | 764.00 | 779.00 | 731.00 | 758.00 | 745.00 |
| RABEP1 | 2404.00 | 2213.00 | 2605.00 | 2407.33 | 2844.00 |
| RABEP2 | 606.00 | 597.00 | 634.00 | 612.33 | 751.00 |
| RABEPK | 681.00 | 698.00 | 819.00 | 732.67 | 943.00 |
| RABGAP1 | 1291.00 | 1183.00 | 1322.00 | 1265.33 | 1192.00 |
| RABGAP1L | 440.00 | 369.00 | 469.00 | 426.00 | 527.00 |
| RABGEF1 | 1218.00 | 1110.00 | 1208.00 | 1178.67 | 1353.00 |
| RABGGTA | 482.00 | 493.00 | 522.00 | 499.00 | 531.00 |
| RABGGTB | 6672.00 | 6421.00 | 6845.00 | 6646.00 | 8030.00 |
| RABIF | 202.00 | 177.00 | 189.00 | 189.33 | 224.00 |
| RABL2A | 98.00 | 87.00 | 94.00 | 93.00 | 82.00 |
| RABL2B | 417.00 | 440.00 | 486.00 | 447.67 | 486.00 |
| RABL3 | 522.00 | 471.00 | 608.00 | 533.67 | 565.00 |
| RABL6 | 2331.00 | 2346.00 | 2512.00 | 2396.33 | 2818.00 |
| RAC1 | 5122.00 | 4718.00 | 5557.00 | 5132.33 | 6602.00 |
| RAC2 | 1681.00 | 1649.00 | 1721.00 | 1683.67 | 1798.00 |
| RAC3 | 748.00 | 804.00 | 841.00 | 797.67 | 1048.00 |
| RACGAP1 | 1213.00 | 1062.00 | 1205.00 | 1160.00 | 1502.00 |
| RAD1 | 730.00 | 644.00 | 684.00 | 686.00 | 805.00 |
| RAD17 | 60.00 | 65.00 | 70.00 | 65.00 | 69.00 |
| RAD18 | 847.00 | 840.00 | 885.00 | 857.33 | 863.00 |
| RAD21 | 5613.00 | 5242.00 | 6211.00 | 5688.67 | 6731.00 |
| RAD21-AS1 | 9.00 | 5.00 | 10.00 | 8.00 | 8.00 |
| RAD21L1 | 17.00 | 12.00 | 15.00 | 14.67 | 29.00 |
| RAD23A | 2481.00 | 2417.00 | 2477.00 | 2458.33 | 3107.00 |
| RAD23B | 2562.00 | 2341.00 | 2756.00 | 2553.00 | 3146.00 |
| RAD50 | 2242.00 | 2332.00 | 2377.00 | 2317.00 | 2210.00 |
| RAD51 | 115.00 | 102.00 | 92.00 | 103.00 | 189.00 |
| RAD51AP1 | 401.00 | 384.00 | 357.00 | 380.67 | 392.00 |
| RAD51-AS1 | 138.00 | 142.00 | 184.00 | 154.67 | 56.00 |
| RAD51B | 219.00 | 221.00 | 198.00 | 212.67 | 241.00 |
| RAD51C | 546.00 | 605.00 | 660.00 | 603.67 | 813.00 |
| RAD51D | 266.00 | 255.00 | 240.00 | 253.67 | 285.00 |
| RAD52 | 1037.00 | 1054.00 | 1071.00 | 1054.00 | 1028.00 |
| RAD54B | 271.00 | 277.00 | 301.00 | 283.00 | 379.00 |
| RAD54L | 363.00 | 362.00 | 363.00 | 362.67 | 426.00 |
| RAD54L2 | 547.00 | 542.00 | 650.00 | 579.67 | 328.00 |
| RAD9A | 280.00 | 336.00 | 282.00 | 299.33 | 305.00 |
| RAD9B | 12.00 | 14.00 | 16.00 | 14.00 | 21.00 |
| RAE1 | 1332.00 | 1324.00 | 1471.00 | 1375.67 | 1731.00 |
| RAET1E | 31.00 | 22.00 | 17.00 | 23.33 | 18.00 |
| RAET1G | 11.00 | 3.00 | 9.00 | 7.67 | 9.00 |
| RAET1K | 27.00 | 28.00 | 26.00 | 27.00 | 37.00 |

| | | | | | |
|---|---|---|---|---|---|
| RAF1 | 1911.00 | 1793.00 | 1927.00 | 1877.00 | 1953.00 |
| RAG1 | 24.00 | 12.00 | 22.00 | 19.33 | 13.00 |
| RAI1 | 1798.00 | 1690.00 | 1868.00 | 1785.33 | 1391.00 |
| RAI14 | 8129.00 | 7168.00 | 8571.00 | 7956.00 | 8355.00 |
| RALA | 1236.00 | 1177.00 | 1368.00 | 1260.33 | 1472.00 |
| RALB | 990.00 | 862.00 | 972.00 | 941.33 | 1152.00 |
| RALBP1 | 2220.00 | 2029.00 | 2429.00 | 2226.00 | 2373.00 |
| RALGAPA1 | 359.00 | 365.00 | 413.00 | 379.00 | 343.00 |
| RALGAPA1P | 25.00 | 21.00 | 33.00 | 26.33 | 27.00 |
| RALGAPA2 | 546.00 | 508.00 | 619.00 | 557.67 | 551.00 |
| RALGAPB | 2763.00 | 2650.00 | 2949.00 | 2787.33 | 2629.00 |
| RALGDS | 708.00 | 802.00 | 876.00 | 795.33 | 978.00 |
| RALGPS1 | 53.00 | 64.00 | 74.00 | 63.67 | 62.00 |
| RALGPS2 | 925.00 | 830.00 | 975.00 | 910.00 | 654.00 |
| RALY | 2778.00 | 2599.00 | 2883.00 | 2753.33 | 3413.00 |
| RALY-AS1 | 12.00 | 10.00 | 9.00 | 10.33 | 7.00 |
| RAMP2 | 5.00 | 5.00 | 7.00 | 5.67 | 2.00 |
| RAMP2-AS1 | 11.00 | 4.00 | 10.00 | 8.33 | 7.00 |
| RAN | 5751.00 | 5533.00 | 5911.00 | 5731.67 | 7961.00 |
| RANBP1 | 1372.00 | 1305.00 | 1354.00 | 1343.67 | 1802.00 |
| RANBP10 | 746.00 | 734.00 | 698.00 | 726.00 | 725.00 |
| RANBP17 | 369.00 | 334.00 | 379.00 | 360.67 | 346.00 |
| RANBP2 | 4231.00 | 3838.00 | 4357.00 | 4142.00 | 4066.00 |
| RANBP3 | 1055.00 | 1125.00 | 1097.00 | 1092.33 | 1236.00 |
| RANBP6 | 1638.00 | 1529.00 | 1803.00 | 1656.67 | 1658.00 |
| RANBP9 | 945.00 | 977.00 | 1003.00 | 975.00 | 1117.00 |
| RANGAP1 | 5172.00 | 5199.00 | 5273.00 | 5214.67 | 5680.00 |
| RANGRF | 1025.00 | 1025.00 | 1133.00 | 1061.00 | 1185.00 |
| RAP1A | 610.00 | 533.00 | 636.00 | 593.00 | 783.00 |
| RAP1B | 1358.00 | 1200.00 | 1300.00 | 1286.00 | 1570.00 |
| RAP1GAP2 | 4859.00 | 4644.00 | 5335.00 | 4946.00 | 5907.00 |
| RAP1GDS1 | 960.00 | 892.00 | 1031.00 | 961.00 | 1078.00 |
| RAP2A | 445.00 | 390.00 | 482.00 | 439.00 | 516.00 |
| RAP2B | 1972.00 | 1818.00 | 2090.00 | 1960.00 | 3600.00 |
| RAP2C | 444.00 | 437.00 | 506.00 | 462.33 | 566.00 |
| RAP2C-AS1 | 31.00 | 29.00 | 31.00 | 30.33 | 11.00 |
| RAPGEF1 | 1305.00 | 1317.00 | 1436.00 | 1352.67 | 1490.00 |
| RAPGEF2 | 844.00 | 783.00 | 892.00 | 839.67 | 545.00 |
| RAPGEF3 | 389.00 | 389.00 | 412.00 | 396.67 | 387.00 |
| RAPGEF4 | 12.00 | 3.00 | 10.00 | 8.33 | 7.00 |
| RAPGEF5 | 147.00 | 165.00 | 170.00 | 160.67 | 223.00 |
| RAPGEF6 | 705.00 | 611.00 | 716.00 | 677.33 | 576.00 |
| RAPGEFL1 | 178.00 | 191.00 | 206.00 | 191.67 | 218.00 |
| RAPH1 | 2517.00 | 2190.00 | 2485.00 | 2397.33 | 2890.00 |
| RAPSN | 8.00 | 15.00 | 17.00 | 13.33 | 3.00 |
| RARA | 843.00 | 845.00 | 924.00 | 870.67 | 676.00 |
| RARA-AS1 | 19.00 | 29.00 | 26.00 | 24.67 | 46.00 |
| RARB | 124.00 | 90.00 | 116.00 | 110.00 | 86.00 |

| | | | | | |
|---|---|---|---|---|---|
| RARG | 291.00 | 288.00 | 311.00 | 296.67 | 249.00 |
| RARRES1 | 52.00 | 47.00 | 39.00 | 46.00 | 92.00 |
| RARRES3 | 6.00 | 6.00 | 6.00 | 6.00 | 10.00 |
| RARS | 2469.00 | 2299.00 | 2582.00 | 2450.00 | 2884.00 |
| RARS2 | 680.00 | 665.00 | 693.00 | 679.33 | 795.00 |
| RASA1 | 2605.00 | 2389.00 | 2871.00 | 2621.67 | 3058.00 |
| RASA2 | 1291.00 | 1243.00 | 1426.00 | 1320.00 | 1676.00 |
| RASA3 | 1610.00 | 1503.00 | 1757.00 | 1623.33 | 1387.00 |
| RASA4B | 4.00 | 5.00 | 4.00 | 4.33 | 1.00 |
| RASA4CP | 8.00 | 11.00 | 4.00 | 7.67 | 9.00 |
| RASAL2 | 1723.00 | 1666.00 | 1785.00 | 1724.67 | 1732.00 |
| RASAL2-AS1 | 47.00 | 53.00 | 42.00 | 47.33 | 59.00 |
| RASAL3 | 11.00 | 9.00 | 5.00 | 8.33 | 5.00 |
| RASD2 | 6.00 | 3.00 | 4.00 | 4.33 | 5.00 |
| RASEF | 7.00 | 3.00 | 3.00 | 4.33 | 5.00 |
| RASGRF1 | 693.00 | 766.00 | 762.00 | 740.33 | 663.00 |
| RASGRF2 | 321.00 | 321.00 | 389.00 | 343.67 | 211.00 |
| RASGRP3 | 8.00 | 7.00 | 6.00 | 7.00 | 8.00 |
| RASL10B | 55.00 | 46.00 | 59.00 | 53.33 | 60.00 |
| RASL11A | 8.00 | 4.00 | 8.00 | 6.67 | 11.00 |
| RASL11B | 8.00 | 2.00 | 4.00 | 4.67 | 6.00 |
| RASSF1 | 647.00 | 628.00 | 658.00 | 644.33 | 786.00 |
| RASSF1-AS1 | 5.00 | 3.00 | 3.00 | 3.67 | 2.00 |
| RASSF3 | 836.00 | 786.00 | 864.00 | 828.67 | 802.00 |
| RASSF4 | 379.00 | 371.00 | 391.00 | 380.33 | 501.00 |
| RASSF5 | 7.00 | 2.00 | 3.00 | 4.00 | 3.00 |
| RASSF7 | 4.00 | 6.00 | 7.00 | 5.67 | 13.00 |
| RASSF8 | 2122.00 | 1693.00 | 2186.00 | 2000.33 | 1678.00 |
| RASSF8-AS1 | 135.00 | 101.00 | 130.00 | 122.00 | 131.00 |
| RAVER1 | 1757.00 | 1624.00 | 1796.00 | 1725.67 | 1774.00 |
| RAVER2 | 596.00 | 552.00 | 765.00 | 637.67 | 640.00 |
| RB1 | 770.00 | 640.00 | 730.00 | 713.33 | 799.00 |
| RB1CC1 | 2317.00 | 2142.00 | 2574.00 | 2344.33 | 2308.00 |
| RBAK | 874.00 | 794.00 | 923.00 | 863.67 | 804.00 |
| RBBP4 | 2461.00 | 2449.00 | 2711.00 | 2540.33 | 3113.00 |
| RBBP5 | 545.00 | 570.00 | 622.00 | 579.00 | 679.00 |
| RBBP6 | 2624.00 | 2610.00 | 3023.00 | 2752.33 | 2659.00 |
| RBBP7 | 3567.00 | 3413.00 | 3669.00 | 3549.67 | 4181.00 |
| RBBP8 | 991.00 | 874.00 | 1066.00 | 977.00 | 1306.00 |
| RBBP9 | 380.00 | 368.00 | 386.00 | 378.00 | 475.00 |
| RBCK1 | 3242.00 | 3347.00 | 3503.00 | 3364.00 | 2594.00 |
| RBFA | 537.00 | 566.00 | 637.00 | 580.00 | 564.00 |
| RBFADN | 14.00 | 26.00 | 26.00 | 22.00 | 15.00 |
| RBFOX1 | 4.00 | 3.00 | 4.00 | 3.67 | 2.00 |
| RBFOX2 | 102.00 | 80.00 | 106.00 | 96.00 | 86.00 |
| RBKS | 68.00 | 58.00 | 67.00 | 64.33 | 97.00 |
| RBL1 | 477.00 | 439.00 | 488.00 | 468.00 | 598.00 |
| RBL2 | 3023.00 | 2760.00 | 3106.00 | 2963.00 | 2774.00 |

| | | | | | |
|---|---|---|---|---|---|
| RBM10 | 1934.00 | 1753.00 | 1835.00 | 1840.67 | 2037.00 |
| RBM12 | 2460.00 | 2286.00 | 2615.00 | 2453.67 | 2604.00 |
| RBM12B | 983.00 | 922.00 | 947.00 | 950.67 | 869.00 |
| RBM14 | 1115.00 | 1022.00 | 1036.00 | 1057.67 | 1321.00 |
| RBM15 | 492.00 | 535.00 | 575.00 | 534.00 | 592.00 |
| RBM15B | 1706.00 | 1665.00 | 1932.00 | 1767.67 | 1825.00 |
| RBM17 | 2241.00 | 2317.00 | 2343.00 | 2300.33 | 2645.00 |
| RBM18 | 492.00 | 457.00 | 509.00 | 486.00 | 569.00 |
| RBM19 | 1750.00 | 1823.00 | 1924.00 | 1832.33 | 1801.00 |
| RBM20 | 9.00 | 12.00 | 18.00 | 13.00 | 17.00 |
| RBM22 | 1311.00 | 1302.00 | 1476.00 | 1363.00 | 1416.00 |
| RBM23 | 927.00 | 892.00 | 947.00 | 922.00 | 884.00 |
| RBM24 | 29.00 | 28.00 | 16.00 | 24.33 | 18.00 |
| RBM25 | 4600.00 | 4455.00 | 4790.00 | 4615.00 | 4884.00 |
| RBM26 | 1759.00 | 1649.00 | 1842.00 | 1750.00 | 1839.00 |
| RBM26-AS1 | 97.00 | 110.00 | 106.00 | 104.33 | 114.00 |
| RBM27 | 859.00 | 736.00 | 915.00 | 836.67 | 969.00 |
| RBM28 | 1574.00 | 1504.00 | 1663.00 | 1580.33 | 1792.00 |
| RBM3 | 2015.00 | 1993.00 | 2103.00 | 2037.00 | 2850.00 |
| RBM33 | 3018.00 | 3115.00 | 3367.00 | 3166.67 | 2473.00 |
| RBM34 | 796.00 | 728.00 | 742.00 | 755.33 | 913.00 |
| RBM38 | 393.00 | 400.00 | 384.00 | 392.33 | 470.00 |
| RBM39 | 11567.00 | 11997.00 | 12963.00 | 12175.67 | 12440.00 |
| RBM4 | 1518.00 | 1578.00 | 1608.00 | 1568.00 | 1654.00 |
| RBM41 | 844.00 | 712.00 | 742.00 | 766.00 | 797.00 |
| RBM42 | 1108.00 | 1116.00 | 1125.00 | 1116.33 | 1354.00 |
| RBM43 | 128.00 | 103.00 | 172.00 | 134.33 | 182.00 |
| RBM44 | 35.00 | 39.00 | 33.00 | 35.67 | 25.00 |
| RBM45 | 229.00 | 232.00 | 245.00 | 235.33 | 301.00 |
| RBM47 | 10.00 | 2.00 | 13.00 | 8.33 | 4.00 |
| RBM48 | 545.00 | 479.00 | 551.00 | 525.00 | 540.00 |
| RBM4B | 601.00 | 592.00 | 706.00 | 633.00 | 677.00 |
| RBM5 | 2833.00 | 2977.00 | 3038.00 | 2949.33 | 2929.00 |
| RBM6 | 2153.00 | 2298.00 | 2478.00 | 2309.67 | 2359.00 |
| RBM7 | 506.00 | 373.00 | 471.00 | 450.00 | 484.00 |
| RBM8A | 1672.00 | 1669.00 | 1660.00 | 1667.00 | 2153.00 |
| RBMS1 | 1586.00 | 1377.00 | 1757.00 | 1573.33 | 1803.00 |
| RBMS2 | 752.00 | 750.00 | 805.00 | 769.00 | 637.00 |
| RBMS3 | 412.00 | 389.00 | 502.00 | 434.33 | 431.00 |
| RBMS3-AS3 | 29.00 | 14.00 | 27.00 | 23.33 | 25.00 |
| RBMX | 5567.00 | 5323.00 | 5703.00 | 5531.00 | 6667.00 |
| RBMX2 | 345.00 | 321.00 | 333.00 | 333.00 | 389.00 |
| RBMXL1 | 519.00 | 483.00 | 546.00 | 516.00 | 631.00 |
| RBP4 | 5.00 | 2.00 | 2.00 | 3.00 | 5.00 |
| RBPJ | 1076.00 | 1041.00 | 1136.00 | 1084.33 | 935.00 |
| RBPMS | 114.00 | 92.00 | 129.00 | 111.67 | 122.00 |
| RBSN | 996.00 | 942.00 | 1084.00 | 1007.33 | 946.00 |
| RBX1 | 438.00 | 384.00 | 393.00 | 405.00 | 566.00 |

| | | | | | |
|---|---|---|---|---|---|
| RC3H1 | 727.00 | 624.00 | 695.00 | 682.00 | 690.00 |
| RC3H2 | 317.00 | 301.00 | 353.00 | 323.67 | 286.00 |
| RCAN1 | 1438.00 | 1280.00 | 1490.00 | 1402.67 | 2252.00 |
| RCAN2 | 10.00 | 13.00 | 6.00 | 9.67 | 13.00 |
| RCAN3 | 35.00 | 22.00 | 26.00 | 27.67 | 14.00 |
| RCBTB1 | 862.00 | 765.00 | 844.00 | 823.67 | 881.00 |
| RCBTB2 | 140.00 | 154.00 | 151.00 | 148.33 | 126.00 |
| RCC1 | 2149.00 | 2029.00 | 2200.00 | 2126.00 | 2523.00 |
| RCC2 | 2197.00 | 2222.00 | 2205.00 | 2208.00 | 2842.00 |
| RCCD1 | 646.00 | 664.00 | 714.00 | 674.67 | 748.00 |
| RCE1 | 898.00 | 1010.00 | 875.00 | 927.67 | 1037.00 |
| RCHY1 | 275.00 | 216.00 | 306.00 | 265.67 | 298.00 |
| RCL1 | 587.00 | 571.00 | 659.00 | 605.67 | 796.00 |
| RCN1 | 9555.00 | 9400.00 | 9312.00 | 9422.33 | 10236.00 |
| RCN2 | 2594.00 | 2462.00 | 2554.00 | 2536.67 | 2910.00 |
| RCN3 | 6023.00 | 6379.00 | 5805.00 | 6069.00 | 5546.00 |
| RCOR1 | 622.00 | 597.00 | 653.00 | 624.00 | 628.00 |
| RCOR2 | 32.00 | 23.00 | 23.00 | 26.00 | 29.00 |
| RCOR3 | 890.00 | 894.00 | 1017.00 | 933.67 | 898.00 |
| RDH10 | 366.00 | 358.00 | 438.00 | 387.33 | 443.00 |
| RDH11 | 4229.00 | 4027.00 | 3926.00 | 4060.67 | 4898.00 |
| RDH12 | 5.00 | 5.00 | 6.00 | 5.33 | 8.00 |
| RDH14 | 62.00 | 59.00 | 57.00 | 59.33 | 76.00 |
| RDH16 | 5.00 | 10.00 | 20.00 | 11.67 | 18.00 |
| RDH5 | 35.00 | 42.00 | 43.00 | 40.00 | 16.00 |
| RDX | 2843.00 | 2545.00 | 3077.00 | 2821.67 | 3207.00 |
| REC8 | 54.00 | 80.00 | 72.00 | 68.67 | 87.00 |
| RECK | 267.00 | 251.00 | 268.00 | 262.00 | 242.00 |
| RECQL | 2676.00 | 2283.00 | 2827.00 | 2595.33 | 3173.00 |
| RECQL4 | 1166.00 | 1109.00 | 1022.00 | 1099.00 | 1412.00 |
| RECQL5 | 1354.00 | 1493.00 | 1444.00 | 1430.33 | 1428.00 |
| REEP1 | 4.00 | 1.00 | 2.00 | 2.33 | 3.00 |
| REEP2 | 70.00 | 65.00 | 80.00 | 71.67 | 87.00 |
| REEP3 | 921.00 | 790.00 | 963.00 | 891.33 | 1056.00 |
| REEP4 | 991.00 | 948.00 | 898.00 | 945.67 | 1134.00 |
| REEP5 | 3034.00 | 3126.00 | 2896.00 | 3018.67 | 3281.00 |
| REEP6 | 68.00 | 93.00 | 87.00 | 82.67 | 90.00 |
| REL | 99.00 | 82.00 | 105.00 | 95.33 | 67.00 |
| RELA | 2628.00 | 2410.00 | 2588.00 | 2542.00 | 2906.00 |
| RELB | 444.00 | 441.00 | 471.00 | 452.00 | 838.00 |
| RELL1 | 235.00 | 292.00 | 225.00 | 250.67 | 320.00 |
| RELL2 | 198.00 | 227.00 | 215.00 | 213.33 | 166.00 |
| RELN | 116.00 | 95.00 | 117.00 | 109.33 | 40.00 |
| RELT | 623.00 | 585.00 | 587.00 | 598.33 | 490.00 |
| REM2 | 13.00 | 23.00 | 15.00 | 17.00 | 13.00 |
| RENBP | 52.00 | 62.00 | 61.00 | 58.33 | 43.00 |
| REP15 | 40.00 | 47.00 | 47.00 | 44.67 | 31.00 |
| REPIN1 | 3369.00 | 3414.00 | 3569.00 | 3450.67 | 3933.00 |

| | | | | | |
|---|---|---|---|---|---|
| REPS1 | 1483.00 | 1445.00 | 1627.00 | 1518.33 | 1493.00 |
| REPS2 | 29.00 | 34.00 | 34.00 | 32.33 | 18.00 |
| RER1 | 3365.00 | 3451.00 | 3430.00 | 3415.33 | 3417.00 |
| RERE | 2423.00 | 2392.00 | 2704.00 | 2506.33 | 2251.00 |
| RERGL | 61.00 | 63.00 | 84.00 | 69.33 | 54.00 |
| REST | 984.00 | 898.00 | 1048.00 | 976.67 | 1087.00 |
| RETSAT | 1748.00 | 1815.00 | 1846.00 | 1803.00 | 1630.00 |
| REV1 | 995.00 | 1029.00 | 1089.00 | 1037.67 | 1104.00 |
| REV3L | 649.00 | 591.00 | 698.00 | 646.00 | 507.00 |
| REXO1 | 1371.00 | 1430.00 | 1485.00 | 1428.67 | 1525.00 |
| REXO2 | 1717.00 | 1790.00 | 1959.00 | 1822.00 | 2030.00 |
| REXO4 | 794.00 | 773.00 | 821.00 | 796.00 | 982.00 |
| RFC1 | 2385.00 | 2129.00 | 2438.00 | 2317.33 | 2650.00 |
| RFC2 | 449.00 | 396.00 | 414.00 | 419.67 | 684.00 |
| RFC3 | 520.00 | 480.00 | 518.00 | 506.00 | 668.00 |
| RFC4 | 952.00 | 912.00 | 978.00 | 947.33 | 1295.00 |
| RFC5 | 644.00 | 655.00 | 735.00 | 678.00 | 925.00 |
| RFESD | 58.00 | 74.00 | 72.00 | 68.00 | 81.00 |
| RFFL | 113.00 | 111.00 | 124.00 | 116.00 | 101.00 |
| RFK | 453.00 | 356.00 | 398.00 | 402.33 | 507.00 |
| RFNG | 1736.00 | 1842.00 | 1801.00 | 1793.00 | 1794.00 |
| RFPL3S | 13.00 | 10.00 | 10.00 | 11.00 | 11.00 |
| RFT1 | 1706.00 | 1656.00 | 1733.00 | 1698.33 | 1691.00 |
| RFTN1 | 1223.00 | 1096.00 | 1329.00 | 1216.00 | 1369.00 |
| RFTN2 | 3.00 | 2.00 | 2.00 | 2.33 | 8.00 |
| RFWD2 | 770.00 | 799.00 | 872.00 | 813.67 | 1089.00 |
| RFWD3 | 1513.00 | 1483.00 | 1771.00 | 1589.00 | 1939.00 |
| RFX1 | 467.00 | 473.00 | 442.00 | 460.67 | 423.00 |
| RFX2 | 7.00 | 8.00 | 11.00 | 8.67 | 16.00 |
| RFX3 | 514.00 | 426.00 | 634.00 | 524.67 | 600.00 |
| RFX3-AS1 | 41.00 | 40.00 | 38.00 | 39.67 | 44.00 |
| RFX5 | 1208.00 | 1127.00 | 1207.00 | 1180.67 | 1554.00 |
| RFX7 | 617.00 | 555.00 | 586.00 | 586.00 | 537.00 |
| RFX8 | 2809.00 | 2696.00 | 3011.00 | 2838.67 | 3276.00 |
| RFXANK | 783.00 | 765.00 | 823.00 | 790.33 | 948.00 |
| RFXAP | 206.00 | 245.00 | 273.00 | 241.33 | 239.00 |
| RGAG1 | 4.00 | 3.00 | 6.00 | 4.33 | 2.00 |
| RGAG4 | 27.00 | 32.00 | 36.00 | 31.67 | 27.00 |
| RGCC | 22.00 | 25.00 | 34.00 | 27.00 | 51.00 |
| RGL1 | 418.00 | 387.00 | 426.00 | 410.33 | 432.00 |
| RGL2 | 9.00 | 14.00 | 22.00 | 15.00 | 16.00 |
| RGL3 | 520.00 | 464.00 | 559.00 | 514.33 | 422.00 |
| RGMB | 6427.00 | 6003.00 | 6494.00 | 6308.00 | 7462.00 |
| RGMB-AS1 | 6.00 | 5.00 | 3.00 | 4.67 | 3.00 |
| RGP1 | 430.00 | 361.00 | 390.00 | 393.67 | 206.00 |
| RGPD3 | 18.00 | 11.00 | 16.00 | 15.00 | 6.00 |
| RGS10 | 1575.00 | 1627.00 | 1728.00 | 1643.33 | 1531.00 |
| RGS12 | 1452.00 | 1529.00 | 1504.00 | 1495.00 | 1217.00 |

| | | | | | |
|---|---|---|---|---|---|
| RGS14 | 238.00 | 199.00 | 254.00 | 230.33 | 218.00 |
| RGS16 | 15.00 | 26.00 | 21.00 | 20.67 | 25.00 |
| RGS17 | 91.00 | 79.00 | 96.00 | 88.67 | 85.00 |
| RGS19 | 273.00 | 251.00 | 232.00 | 252.00 | 340.00 |
| RGS2 | 94.00 | 110.00 | 133.00 | 112.33 | 129.00 |
| RGS20 | 549.00 | 521.00 | 584.00 | 551.33 | 699.00 |
| RGS22 | 5.00 | 9.00 | 2.00 | 5.33 | 4.00 |
| RGS3 | 1853.00 | 1833.00 | 2048.00 | 1911.33 | 3129.00 |
| RGS4 | 1124.00 | 1000.00 | 1162.00 | 1095.33 | 3246.00 |
| RGS5 | 177.00 | 149.00 | 162.00 | 162.67 | 183.00 |
| RGS9 | 9.00 | 4.00 | 3.00 | 5.33 | 2.00 |
| RGS9BP | 11.00 | 3.00 | 8.00 | 7.33 | 7.00 |
| RHBDD1 | 1799.00 | 1762.00 | 1970.00 | 1843.67 | 1824.00 |
| RHBDD2 | 518.00 | 453.00 | 468.00 | 479.67 | 560.00 |
| RHBDD3 | 620.00 | 677.00 | 624.00 | 640.33 | 737.00 |
| RHBDF1 | 1440.00 | 1446.00 | 1609.00 | 1498.33 | 975.00 |
| RHBDF2 | 1231.00 | 1215.00 | 1351.00 | 1265.67 | 1091.00 |
| RHBDL1 | 123.00 | 128.00 | 139.00 | 130.00 | 94.00 |
| RHCE | 8.00 | 8.00 | 6.00 | 7.33 | 4.00 |
| RHEB | 1750.00 | 1630.00 | 1850.00 | 1743.33 | 2327.00 |
| RHEBL1 | 135.00 | 117.00 | 138.00 | 130.00 | 132.00 |
| RHNO1 | 582.00 | 579.00 | 606.00 | 589.00 | 838.00 |
| RHOA | 5353.00 | 5034.00 | 5538.00 | 5308.33 | 6709.00 |
| RHOB | 193.00 | 156.00 | 180.00 | 176.33 | 93.00 |
| RHOBTB1 | 493.00 | 504.00 | 527.00 | 508.00 | 490.00 |
| RHOBTB2 | 262.00 | 276.00 | 267.00 | 268.33 | 317.00 |
| RHOBTB3 | 3360.00 | 3148.00 | 3676.00 | 3394.67 | 3238.00 |
| RHOC | 3014.00 | 3066.00 | 3189.00 | 3089.67 | 4016.00 |
| RHOF | 81.00 | 95.00 | 103.00 | 93.00 | 135.00 |
| RHOG | 605.00 | 643.00 | 617.00 | 621.67 | 739.00 |
| RHOH | 8.00 | 5.00 | 5.00 | 6.00 | 2.00 |
| RHOJ | 224.00 | 209.00 | 272.00 | 235.00 | 319.00 |
| RHOQ | 960.00 | 999.00 | 1071.00 | 1010.00 | 1133.00 |
| RHOT1 | 811.00 | 683.00 | 922.00 | 805.33 | 929.00 |
| RHOT2 | 2641.00 | 2827.00 | 2886.00 | 2784.67 | 2942.00 |
| RHPN1 | 111.00 | 85.00 | 116.00 | 104.00 | 134.00 |
| RHPN1-AS1 | 9.00 | 12.00 | 5.00 | 8.67 | 12.00 |
| RHPN2 | 101.00 | 73.00 | 93.00 | 89.00 | 95.00 |
| RIBC1 | 9.00 | 12.00 | 10.00 | 10.33 | 3.00 |
| RIBC2 | 9.00 | 2.00 | 8.00 | 6.33 | 16.00 |
| RIC1 | 1121.00 | 1030.00 | 1169.00 | 1106.67 | 955.00 |
| RIC8A | 1883.00 | 1819.00 | 1954.00 | 1885.33 | 2085.00 |
| RIC8B | 412.00 | 428.00 | 469.00 | 436.33 | 466.00 |
| RICTOR | 1437.00 | 1347.00 | 1617.00 | 1467.00 | 1155.00 |
| RIF1 | 1571.00 | 1475.00 | 1708.00 | 1584.67 | 1568.00 |
| RILPL1 | 601.00 | 631.00 | 660.00 | 630.67 | 543.00 |
| RILPL2 | 196.00 | 225.00 | 243.00 | 221.33 | 232.00 |
| RIMKLA | 5.00 | 1.00 | 1.00 | 2.33 | 3.00 |

| | | | | | |
|---|---|---|---|---|---|
| RIMKLB | 1115.00 | 1047.00 | 1151.00 | 1104.33 | 1109.00 |
| RIMS1 | 26.00 | 25.00 | 33.00 | 28.00 | 14.00 |
| RIMS2 | 79.00 | 51.00 | 98.00 | 76.00 | 83.00 |
| RIMS3 | 3.00 | 0.00 | 3.00 | 2.00 | 4.00 |
| RIN1 | 1390.00 | 1354.00 | 1396.00 | 1380.00 | 1203.00 |
| RIN2 | 1764.00 | 1675.00 | 1937.00 | 1792.00 | 2519.00 |
| RIN3 | 993.00 | 974.00 | 1003.00 | 990.00 | 751.00 |
| RINL | 67.00 | 77.00 | 62.00 | 68.67 | 58.00 |
| RINT1 | 650.00 | 658.00 | 696.00 | 668.00 | 788.00 |
| RIOK1 | 1006.00 | 939.00 | 1001.00 | 982.00 | 1280.00 |
| RIOK2 | 1162.00 | 1060.00 | 1283.00 | 1168.33 | 1255.00 |
| RIOK3 | 1580.00 | 1470.00 | 1780.00 | 1610.00 | 1629.00 |
| RIPK1 | 1026.00 | 979.00 | 1024.00 | 1009.67 | 1112.00 |
| RIPK2 | 1123.00 | 1051.00 | 1332.00 | 1168.67 | 1250.00 |
| RIPPLY2 | 54.00 | 61.00 | 78.00 | 64.33 | 85.00 |
| RIT1 | 580.00 | 486.00 | 567.00 | 544.33 | 642.00 |
| RITA1 | 533.00 | 504.00 | 618.00 | 551.67 | 676.00 |
| RLF | 472.00 | 508.00 | 562.00 | 514.00 | 500.00 |
| RLIM | 1874.00 | 1663.00 | 2053.00 | 1863.33 | 1885.00 |
| RLTPR | 10.00 | 19.00 | 8.00 | 12.33 | 15.00 |
| RMDN1 | 1117.00 | 1096.00 | 1194.00 | 1135.67 | 1405.00 |
| RMDN2 | 103.00 | 79.00 | 97.00 | 93.00 | 107.00 |
| RMDN3 | 1045.00 | 1043.00 | 1062.00 | 1050.00 | 1268.00 |
| RMI1 | 213.00 | 168.00 | 167.00 | 182.67 | 293.00 |
| RMI2 | 166.00 | 165.00 | 167.00 | 166.00 | 216.00 |
| RMND1 | 634.00 | 682.00 | 790.00 | 702.00 | 733.00 |
| RMND5A | 646.00 | 568.00 | 675.00 | 629.67 | 807.00 |
| RMND5B | 308.00 | 296.00 | 327.00 | 310.33 | 416.00 |
| RMRP | 18.00 | 15.00 | 6.00 | 13.00 | 27.00 |
| RN7SK | 28.00 | 46.00 | 31.00 | 35.00 | 64.00 |
| RN7SL1 | 440.00 | 427.00 | 114.00 | 327.00 | 301.00 |
| RN7SL2 | 643.00 | 529.00 | 100.00 | 424.00 | 301.00 |
| RNA28S5 | 3.00 | 2.00 | 4.00 | 3.00 | 6.00 |
| RNA45S5 | 10.00 | 9.00 | 12.00 | 10.33 | 12.00 |
| RNASE4 | 13.00 | 21.00 | 21.00 | 18.33 | 32.00 |
| RNASEH1 | 646.00 | 692.00 | 645.00 | 661.00 | 678.00 |
| RNASEH1-AS1 | 222.00 | 256.00 | 274.00 | 250.67 | 344.00 |
| RNASEH2A | 726.00 | 664.00 | 697.00 | 695.67 | 944.00 |
| RNASEH2B | 423.00 | 392.00 | 444.00 | 419.67 | 568.00 |
| RNASEH2B-AS1 | 5.00 | 8.00 | 11.00 | 8.00 | 3.00 |
| RNASEH2C | 1756.00 | 1786.00 | 1917.00 | 1819.67 | 1815.00 |
| RNASEK | 86.00 | 124.00 | 112.00 | 107.33 | 130.00 |
| RNASEL | 43.00 | 50.00 | 58.00 | 50.33 | 65.00 |
| RNASET2 | 502.00 | 516.00 | 541.00 | 519.67 | 587.00 |
| RND2 | 4.00 | 0.00 | 0.00 | 1.33 | 0.00 |
| RND3 | 5162.00 | 4932.00 | 5572.00 | 5222.00 | 7042.00 |
| RNF10 | 2037.00 | 1878.00 | 2105.00 | 2006.67 | 1876.00 |
| RNF103 | 775.00 | 695.00 | 769.00 | 746.33 | 710.00 |

| | | | | | |
|---|---|---|---|---|---|
| RNF11 | 1117.00 | 947.00 | 1161.00 | 1075.00 | 911.00 |
| RNF111 | 998.00 | 816.00 | 985.00 | 933.00 | 788.00 |
| RNF112 | 6.00 | 2.00 | 4.00 | 4.00 | 2.00 |
| RNF113A | 235.00 | 244.00 | 239.00 | 239.33 | 276.00 |
| RNF114 | 1776.00 | 1707.00 | 1923.00 | 1802.00 | 2339.00 |
| RNF115 | 472.00 | 394.00 | 457.00 | 441.00 | 389.00 |
| RNF121 | 1583.00 | 1723.00 | 1637.00 | 1647.67 | 1464.00 |
| RNF122 | 19.00 | 9.00 | 9.00 | 12.33 | 14.00 |
| RNF123 | 1466.00 | 1376.00 | 1540.00 | 1460.67 | 1424.00 |
| RNF125 | 167.00 | 122.00 | 162.00 | 150.33 | 235.00 |
| RNF126 | 931.00 | 892.00 | 993.00 | 938.67 | 1169.00 |
| RNF128 | 98.00 | 103.00 | 100.00 | 100.33 | 85.00 |
| RNF13 | 1089.00 | 947.00 | 1119.00 | 1051.67 | 1181.00 |
| RNF130 | 652.00 | 668.00 | 647.00 | 655.67 | 692.00 |
| RNF133 | 3.00 | 1.00 | 2.00 | 2.00 | 3.00 |
| RNF135 | 592.00 | 594.00 | 680.00 | 622.00 | 788.00 |
| RNF138 | 348.00 | 313.00 | 423.00 | 361.33 | 470.00 |
| RNF139 | 1191.00 | 1191.00 | 1262.00 | 1214.67 | 1226.00 |
| RNF139-AS1 | 95.00 | 82.00 | 81.00 | 86.00 | 113.00 |
| RNF14 | 825.00 | 687.00 | 845.00 | 785.67 | 944.00 |
| RNF141 | 443.00 | 424.00 | 473.00 | 446.67 | 546.00 |
| RNF144B | 105.00 | 92.00 | 135.00 | 110.67 | 123.00 |
| RNF145 | 1778.00 | 1678.00 | 1711.00 | 1722.33 | 1754.00 |
| RNF146 | 1326.00 | 1303.00 | 1495.00 | 1374.67 | 1421.00 |
| RNF149 | 3097.00 | 3103.00 | 2989.00 | 3063.00 | 3137.00 |
| RNF150 | 45.00 | 43.00 | 60.00 | 49.33 | 34.00 |
| RNF152 | 13.00 | 19.00 | 27.00 | 19.67 | 10.00 |
| RNF157 | 125.00 | 118.00 | 129.00 | 124.00 | 89.00 |
| RNF166 | 913.00 | 905.00 | 1005.00 | 941.00 | 1050.00 |
| RNF167 | 2103.00 | 2104.00 | 2183.00 | 2130.00 | 2172.00 |
| RNF168 | 421.00 | 371.00 | 432.00 | 408.00 | 414.00 |
| RNF169 | 732.00 | 729.00 | 806.00 | 755.67 | 732.00 |
| RNF170 | 697.00 | 645.00 | 743.00 | 695.00 | 790.00 |
| RNF180 | 14.00 | 7.00 | 8.00 | 9.67 | 2.00 |
| RNF181 | 945.00 | 921.00 | 944.00 | 936.67 | 1098.00 |
| RNF182 | 185.00 | 170.00 | 194.00 | 183.00 | 178.00 |
| RNF185 | 697.00 | 699.00 | 756.00 | 717.33 | 753.00 |
| RNF187 | 1600.00 | 1585.00 | 1667.00 | 1617.33 | 1667.00 |
| RNF19A | 856.00 | 770.00 | 950.00 | 858.67 | 898.00 |
| RNF19B | 529.00 | 512.00 | 588.00 | 543.00 | 605.00 |
| RNF2 | 483.00 | 404.00 | 502.00 | 463.00 | 661.00 |
| RNF20 | 1381.00 | 1308.00 | 1508.00 | 1399.00 | 1282.00 |
| RNF207 | 710.00 | 717.00 | 775.00 | 734.00 | 766.00 |
| RNF208 | 33.00 | 45.00 | 46.00 | 41.33 | 52.00 |
| RNF213 | 6097.00 | 5336.00 | 6485.00 | 5972.67 | 4482.00 |
| RNF214 | 511.00 | 443.00 | 530.00 | 494.67 | 459.00 |
| RNF215 | 811.00 | 893.00 | 835.00 | 846.33 | 1070.00 |
| RNF216 | 2910.00 | 2894.00 | 3199.00 | 3001.00 | 2858.00 |

| | | | | | |
|---|---|---|---|---|---|
| RNF216P1 | 715.00 | 683.00 | 724.00 | 707.33 | 799.00 |
| RNF217 | 304.00 | 268.00 | 300.00 | 290.67 | 289.00 |
| RNF217-AS1 | 6.00 | 13.00 | 15.00 | 11.33 | 16.00 |
| RNF219 | 806.00 | 786.00 | 829.00 | 807.00 | 929.00 |
| RNF219-AS1 | 3.00 | 1.00 | 0.00 | 1.33 | 0.00 |
| RNF220 | 1734.00 | 1722.00 | 1946.00 | 1800.67 | 1526.00 |
| RNF24 | 550.00 | 462.00 | 515.00 | 509.00 | 367.00 |
| RNF25 | 506.00 | 481.00 | 548.00 | 511.67 | 629.00 |
| RNF26 | 1804.00 | 1813.00 | 1717.00 | 1778.00 | 1864.00 |
| RNF31 | 1121.00 | 1091.00 | 1218.00 | 1143.33 | 1139.00 |
| RNF32 | 110.00 | 108.00 | 119.00 | 112.33 | 144.00 |
| RNF34 | 585.00 | 611.00 | 573.00 | 589.67 | 723.00 |
| RNF38 | 467.00 | 424.00 | 467.00 | 452.67 | 459.00 |
| RNF4 | 641.00 | 676.00 | 723.00 | 680.00 | 781.00 |
| RNF40 | 2563.00 | 2512.00 | 2554.00 | 2543.00 | 2371.00 |
| RNF41 | 992.00 | 851.00 | 970.00 | 937.67 | 967.00 |
| RNF44 | 1617.00 | 1558.00 | 1852.00 | 1675.67 | 1695.00 |
| RNF5P1 | 19.00 | 20.00 | 17.00 | 18.67 | 24.00 |
| RNF6 | 951.00 | 928.00 | 1084.00 | 987.67 | 1094.00 |
| RNF7 | 1266.00 | 1330.00 | 1366.00 | 1320.67 | 1655.00 |
| RNF8 | 485.00 | 470.00 | 489.00 | 481.33 | 529.00 |
| RNFT1 | 623.00 | 613.00 | 587.00 | 607.67 | 700.00 |
| RNFT2 | 119.00 | 135.00 | 129.00 | 127.67 | 167.00 |
| RNGTT | 700.00 | 545.00 | 645.00 | 630.00 | 668.00 |
| RNH1 | 5.00 | 6.00 | 10.00 | 7.00 | 13.00 |
| RNLS | 56.00 | 42.00 | 43.00 | 47.00 | 42.00 |
| RNMT | 2180.00 | 1963.00 | 2367.00 | 2170.00 | 1957.00 |
| RNMTL1 | 372.00 | 364.00 | 334.00 | 356.67 | 431.00 |
| RNPC3 | 670.00 | 641.00 | 740.00 | 683.67 | 636.00 |
| RNPEP | 1385.00 | 1454.00 | 1438.00 | 1425.67 | 1693.00 |
| RNPEPL1 | 1028.00 | 1111.00 | 1155.00 | 1098.00 | 1116.00 |
| RNPS1 | 2801.00 | 2600.00 | 2810.00 | 2737.00 | 3558.00 |
| RNU11 | 4.00 | 1.00 | 3.00 | 2.67 | 2.00 |
| RNU2-1 | 3.00 | 3.00 | 2.00 | 2.67 | 4.00 |
| RNU6-26P | 48.00 | 44.00 | 61.00 | 51.00 | 18.00 |
| RNU86 | 73.00 | 77.00 | 67.00 | 72.33 | 77.00 |
| ROBO1 | 1511.00 | 1365.00 | 1561.00 | 1479.00 | 1314.00 |
| ROBO2 | 71.00 | 77.00 | 94.00 | 80.67 | 71.00 |
| ROBO3 | 392.00 | 448.00 | 405.00 | 415.00 | 589.00 |
| ROBO4 | 1806.00 | 1820.00 | 1796.00 | 1807.33 | 1840.00 |
| ROCK1 | 1556.00 | 1428.00 | 1689.00 | 1557.67 | 1492.00 |
| ROCK1P1 | 34.00 | 29.00 | 53.00 | 38.67 | 36.00 |
| ROCK2 | 1218.00 | 1065.00 | 1290.00 | 1191.00 | 966.00 |
| ROGDI | 85.00 | 103.00 | 113.00 | 100.33 | 112.00 |
| ROM1 | 75.00 | 88.00 | 97.00 | 86.67 | 68.00 |
| ROMO1 | 697.00 | 656.00 | 607.00 | 653.33 | 839.00 |
| ROR1 | 29.00 | 21.00 | 35.00 | 28.33 | 21.00 |
| RORA | 141.00 | 135.00 | 169.00 | 148.33 | 188.00 |

| | | | | | |
|---|---|---|---|---|---|
| RORA-AS1 | 23.00 | 18.00 | 33.00 | 24.67 | 25.00 |
| RORB | 5.00 | 17.00 | 12.00 | 11.33 | 6.00 |
| ROS1 | 184.00 | 216.00 | 183.00 | 194.33 | 88.00 |
| RP1 | 14.00 | 15.00 | 9.00 | 12.67 | 10.00 |
| RP1L1 | 55.00 | 50.00 | 49.00 | 51.33 | 40.00 |
| RP2 | 254.00 | 218.00 | 256.00 | 242.67 | 270.00 |
| RP9 | 284.00 | 277.00 | 324.00 | 295.00 | 405.00 |
| RP9P | 70.00 | 84.00 | 67.00 | 73.67 | 94.00 |
| RPA1 | 1293.00 | 1331.00 | 1427.00 | 1350.33 | 1633.00 |
| RPA2 | 643.00 | 666.00 | 701.00 | 670.00 | 870.00 |
| RPA3 | 562.00 | 507.00 | 545.00 | 538.00 | 736.00 |
| RPA4 | 7.00 | 8.00 | 4.00 | 6.33 | 11.00 |
| RPAIN | 1400.00 | 1256.00 | 1437.00 | 1364.33 | 1496.00 |
| RPAP1 | 1646.00 | 1604.00 | 1738.00 | 1662.67 | 1484.00 |
| RPAP2 | 490.00 | 411.00 | 481.00 | 460.67 | 511.00 |
| RPAP3 | 883.00 | 748.00 | 875.00 | 835.33 | 1053.00 |
| RPARP-AS1 | 233.00 | 231.00 | 255.00 | 239.67 | 311.00 |
| RPE | 1320.00 | 1219.00 | 1520.00 | 1353.00 | 1507.00 |
| RPEL1 | 16.00 | 26.00 | 19.00 | 20.33 | 27.00 |
| RPF1 | 1041.00 | 955.00 | 1065.00 | 1020.33 | 1325.00 |
| RPF2 | 1364.00 | 1231.00 | 1468.00 | 1354.33 | 1748.00 |
| RPGR | 227.00 | 198.00 | 182.00 | 202.33 | 305.00 |
| RPGRIP1 | 5.00 | 3.00 | 5.00 | 4.33 | 1.00 |
| RPGRIP1L | 514.00 | 498.00 | 522.00 | 511.33 | 488.00 |
| RPIA | 778.00 | 716.00 | 859.00 | 784.33 | 1061.00 |
| RPL10 | 16704.00 | 16769.00 | 17960.00 | 17144.33 | 22462.00 |
| RPL10A | 11534.00 | 11336.00 | 12685.00 | 11851.67 | 16055.00 |
| RPL11 | 16077.00 | 16285.00 | 17698.00 | 16686.67 | 21672.00 |
| RPL12 | 17826.00 | 18042.00 | 20353.00 | 18740.33 | 25954.00 |
| RPL13 | 22691.00 | 22775.00 | 25683.00 | 23716.33 | 29859.00 |
| RPL13A | 28221.00 | 27354.00 | 31178.00 | 28917.67 | 36255.00 |
| RPL13AP20 | 75.00 | 79.00 | 65.00 | 73.00 | 81.00 |
| RPL13AP5 | 2211.00 | 1987.00 | 2337.00 | 2178.33 | 3127.00 |
| RPL13AP6 | 3.00 | 0.00 | 1.00 | 1.33 | 3.00 |
| RPL13P5 | 35.00 | 40.00 | 44.00 | 39.67 | 40.00 |
| RPL14 | 9584.00 | 9006.00 | 10142.00 | 9577.33 | 12468.00 |
| RPL15 | 16311.00 | 15915.00 | 18021.00 | 16749.00 | 21376.00 |
| RPL17 | 1078.00 | 1080.00 | 1076.00 | 1078.00 | 1347.00 |
| RPL18 | 10901.00 | 11099.00 | 11844.00 | 11281.33 | 14495.00 |
| RPL18A | 6662.00 | 6679.00 | 7250.00 | 6863.67 | 8960.00 |
| RPL19 | 21590.00 | 21513.00 | 23939.00 | 22347.33 | 28649.00 |
| RPL19P12 | 27.00 | 31.00 | 30.00 | 29.33 | 34.00 |
| RPL21 | 672.00 | 673.00 | 793.00 | 712.67 | 996.00 |
| RPL21P44 | 11.00 | 7.00 | 9.00 | 9.00 | 6.00 |
| RPL22 | 5198.00 | 5048.00 | 5664.00 | 5303.33 | 7107.00 |
| RPL22L1 | 3808.00 | 3912.00 | 3940.00 | 3886.67 | 5181.00 |
| RPL23 | 14274.00 | 14404.00 | 15522.00 | 14733.33 | 19851.00 |
| RPL23A | 7338.00 | 7488.00 | 8276.00 | 7700.67 | 9967.00 |

| | | | | | |
|---|---|---|---|---|---|
| RPL23AP53 | 125.00 | 99.00 | 113.00 | 112.33 | 106.00 |
| RPL23AP7 | 175.00 | 179.00 | 154.00 | 169.33 | 179.00 |
| RPL23AP82 | 114.00 | 115.00 | 94.00 | 107.67 | 128.00 |
| RPL23P8 | 63.00 | 68.00 | 69.00 | 66.67 | 94.00 |
| RPL24 | 6349.00 | 6508.00 | 7181.00 | 6679.33 | 9087.00 |
| RPL26 | 19431.00 | 19283.00 | 21782.00 | 20165.33 | 27991.00 |
| RPL26L1 | 764.00 | 703.00 | 771.00 | 746.00 | 1136.00 |
| RPL27 | 12275.00 | 12251.00 | 13370.00 | 12632.00 | 17463.00 |
| RPL27A | 13338.00 | 13336.00 | 14583.00 | 13752.33 | 17444.00 |
| RPL28 | 7978.00 | 7963.00 | 9019.00 | 8320.00 | 9991.00 |
| RPL29 | 10705.00 | 10697.00 | 12172.00 | 11191.33 | 14537.00 |
| RPL3 | 38683.00 | 39014.00 | 43704.00 | 40467.00 | 50408.00 |
| RPL30 | 14397.00 | 14235.00 | 15797.00 | 14809.67 | 19381.00 |
| RPL31 | 9661.00 | 9537.00 | 10751.00 | 9983.00 | 14280.00 |
| RPL32 | 10609.00 | 10376.00 | 11934.00 | 10973.00 | 15400.00 |
| RPL32P3 | 405.00 | 329.00 | 361.00 | 365.00 | 351.00 |
| RPL34 | 4670.00 | 4667.00 | 5056.00 | 4797.67 | 6235.00 |
| RPL34-AS1 | 5.00 | 5.00 | 12.00 | 7.33 | 16.00 |
| RPL35 | 6393.00 | 6487.00 | 6824.00 | 6568.00 | 9064.00 |
| RPL35A | 9039.00 | 8966.00 | 9936.00 | 9313.67 | 13096.00 |
| RPL36 | 6935.00 | 7276.00 | 7706.00 | 7305.67 | 9986.00 |
| RPL36A | 435.00 | 428.00 | 416.00 | 426.33 | 431.00 |
| RPL36AL | 1491.00 | 1617.00 | 1664.00 | 1590.67 | 1917.00 |
| RPL37 | 18044.00 | 19039.00 | 18742.00 | 18608.33 | 26566.00 |
| RPL37A | 17856.00 | 18718.00 | 19453.00 | 18675.67 | 25570.00 |
| RPL38 | 4821.00 | 4866.00 | 5432.00 | 5039.67 | 7654.00 |
| RPL39 | 3940.00 | 4070.00 | 4345.00 | 4118.33 | 5454.00 |
| RPL39L | 559.00 | 522.00 | 533.00 | 538.00 | 834.00 |
| RPL4 | 30515.00 | 28968.00 | 33783.00 | 31088.67 | 39424.00 |
| RPL41 | 1043.00 | 1055.00 | 1172.00 | 1090.00 | 1592.00 |
| RPL5 | 26750.00 | 26422.00 | 29311.00 | 27494.33 | 37013.00 |
| RPL6 | 15494.00 | 15034.00 | 17409.00 | 15979.00 | 21336.00 |
| RPL7 | 10527.00 | 10273.00 | 11576.00 | 10792.00 | 15195.00 |
| RPL7A | 23684.00 | 23666.00 | 26252.00 | 24534.00 | 31753.00 |
| RPL7L1 | 1292.00 | 1140.00 | 1382.00 | 1271.33 | 1410.00 |
| RPL8 | 31110.00 | 31494.00 | 34722.00 | 32442.00 | 45259.00 |
| RPL9 | 10687.00 | 10896.00 | 11622.00 | 11068.33 | 15019.00 |
| RPLP0 | 32502.00 | 31320.00 | 35830.00 | 33217.33 | 41796.00 |
| RPLP0P2 | 75.00 | 54.00 | 72.00 | 67.00 | 63.00 |
| RPLP1 | 21062.00 | 22391.00 | 23372.00 | 22275.00 | 29408.00 |
| RPLP2 | 10483.00 | 10381.00 | 10727.00 | 10530.33 | 14251.00 |
| RPN1 | 9007.00 | 8772.00 | 9038.00 | 8939.00 | 9139.00 |
| RPN2 | 14400.00 | 13707.00 | 14109.00 | 14072.00 | 13456.00 |
| RPP14 | 633.00 | 574.00 | 708.00 | 638.33 | 669.00 |
| RPP25 | 24.00 | 24.00 | 22.00 | 23.33 | 45.00 |
| RPP25L | 485.00 | 464.00 | 513.00 | 487.33 | 634.00 |
| RPP30 | 1018.00 | 981.00 | 1103.00 | 1034.00 | 1264.00 |
| RPP38 | 282.00 | 268.00 | 325.00 | 291.67 | 429.00 |

| | | | | | |
|---|---|---|---|---|---|
| RPP40 | 294.00 | 289.00 | 335.00 | 306.00 | 373.00 |
| RPPH1 | 16.00 | 21.00 | 4.00 | 13.67 | 20.00 |
| RPRD1A | 1249.00 | 1272.00 | 1353.00 | 1291.33 | 1521.00 |
| RPRD1B | 1351.00 | 1281.00 | 1307.00 | 1313.00 | 1366.00 |
| RPRD2 | 1491.00 | 1354.00 | 1519.00 | 1454.67 | 1345.00 |
| RPRM | 4.00 | 6.00 | 6.00 | 5.33 | 3.00 |
| RPS10 | 698.00 | 704.00 | 731.00 | 711.00 | 1007.00 |
| RPS10P7 | 68.00 | 80.00 | 76.00 | 74.67 | 94.00 |
| RPS11 | 11790.00 | 11638.00 | 13193.00 | 12207.00 | 16453.00 |
| RPS12 | 17272.00 | 17050.00 | 18930.00 | 17750.67 | 23754.00 |
| RPS13 | 8452.00 | 8501.00 | 9280.00 | 8744.33 | 11642.00 |
| RPS14 | 14014.00 | 14027.00 | 15177.00 | 14406.00 | 19691.00 |
| RPS14P3 | 46.00 | 56.00 | 68.00 | 56.67 | 56.00 |
| RPS15 | 4136.00 | 4097.00 | 4616.00 | 4283.00 | 5531.00 |
| RPS15A | 10377.00 | 10414.00 | 11203.00 | 10664.67 | 14363.00 |
| RPS15AP10 | 12.00 | 13.00 | 14.00 | 13.00 | 6.00 |
| RPS16 | 10765.00 | 10501.00 | 11724.00 | 10996.67 | 13938.00 |
| RPS17 | 14.00 | 19.00 | 16.00 | 16.33 | 15.00 |
| RPS18 | 160.00 | 172.00 | 154.00 | 162.00 | 229.00 |
| RPS18P9 | 83.00 | 104.00 | 88.00 | 91.67 | 128.00 |
| RPS19 | 12577.00 | 13242.00 | 14280.00 | 13366.33 | 16404.00 |
| RPS19BP1 | 1044.00 | 1115.00 | 1157.00 | 1105.33 | 1314.00 |
| RPS2 | 24143.00 | 26175.00 | 28009.00 | 26109.00 | 39103.00 |
| RPS20 | 13852.00 | 13961.00 | 15194.00 | 14335.67 | 19380.00 |
| RPS21 | 8449.00 | 9010.00 | 9192.00 | 8883.67 | 11758.00 |
| RPS23 | 12441.00 | 12567.00 | 13621.00 | 12876.33 | 18374.00 |
| RPS24 | 12434.00 | 12369.00 | 13446.00 | 12749.67 | 18131.00 |
| RPS25 | 8830.00 | 9135.00 | 9776.00 | 9247.00 | 12905.00 |
| RPS26 | 204.00 | 207.00 | 225.00 | 212.00 | 303.00 |
| RPS27 | 5458.00 | 5774.00 | 6235.00 | 5822.33 | 7529.00 |
| RPS27A | 10617.00 | 10699.00 | 11553.00 | 10956.33 | 14683.00 |
| RPS27L | 902.00 | 824.00 | 984.00 | 903.33 | 1191.00 |
| RPS28 | 1966.00 | 1959.00 | 2204.00 | 2043.00 | 2585.00 |
| RPS29 | 5633.00 | 5851.00 | 6100.00 | 5861.33 | 8411.00 |
| RPS2P32 | 30.00 | 29.00 | 18.00 | 25.67 | 16.00 |
| RPS3 | 35876.00 | 35899.00 | 38969.00 | 36914.67 | 49092.00 |
| RPS3A | 7996.00 | 7980.00 | 9115.00 | 8363.67 | 11071.00 |
| RPS4X | 34099.00 | 33747.00 | 38450.00 | 35432.00 | 47539.00 |
| RPS5 | 10641.00 | 10448.00 | 11703.00 | 10930.67 | 13287.00 |
| RPS6 | 28439.00 | 27842.00 | 31938.00 | 29406.33 | 37153.00 |
| RPS6KA1 | 347.00 | 330.00 | 417.00 | 364.67 | 385.00 |
| RPS6KA2 | 4220.00 | 3883.00 | 4370.00 | 4157.67 | 4630.00 |
| RPS6KA2-IT1 | 15.00 | 17.00 | 15.00 | 15.67 | 18.00 |
| RPS6KA3 | 1277.00 | 1169.00 | 1321.00 | 1255.67 | 1152.00 |
| RPS6KA4 | 1930.00 | 1850.00 | 1932.00 | 1904.00 | 2202.00 |
| RPS6KA5 | 76.00 | 66.00 | 80.00 | 74.00 | 59.00 |
| RPS6KA6 | 134.00 | 134.00 | 163.00 | 143.67 | 127.00 |
| RPS6KB1 | 906.00 | 862.00 | 978.00 | 915.33 | 888.00 |

| | | | | | |
|---|---|---|---|---|---|
| RPS6KB2 | 591.00 | 583.00 | 605.00 | 593.00 | 715.00 |
| RPS6KC1 | 664.00 | 633.00 | 669.00 | 655.33 | 598.00 |
| RPS6KL1 | 142.00 | 112.00 | 141.00 | 131.67 | 116.00 |
| RPS7 | 13795.00 | 14448.00 | 15504.00 | 14582.33 | 20205.00 |
| RPS8 | 18653.00 | 18021.00 | 20375.00 | 19016.33 | 24859.00 |
| RPS9 | 350.00 | 365.00 | 358.00 | 357.67 | 433.00 |
| RPSA | 12595.00 | 12413.00 | 14296.00 | 13101.33 | 16892.00 |
| RPSAP52 | 67.00 | 55.00 | 53.00 | 58.33 | 42.00 |
| RPSAP58 | 1847.00 | 1879.00 | 2104.00 | 1943.33 | 2623.00 |
| RPSAP9 | 42.00 | 49.00 | 64.00 | 51.67 | 59.00 |
| RPTOR | 1761.00 | 1702.00 | 1800.00 | 1754.33 | 1629.00 |
| RPUSD1 | 1015.00 | 1042.00 | 1057.00 | 1038.00 | 1142.00 |
| RPUSD2 | 317.00 | 366.00 | 354.00 | 345.67 | 370.00 |
| RPUSD3 | 715.00 | 679.00 | 689.00 | 694.33 | 818.00 |
| RPUSD4 | 699.00 | 625.00 | 719.00 | 681.00 | 815.00 |
| RQCD1 | 1608.00 | 1551.00 | 1655.00 | 1604.67 | 2206.00 |
| RRAD | 8.00 | 18.00 | 12.00 | 12.67 | 13.00 |
| RRAGA | 799.00 | 754.00 | 816.00 | 789.67 | 954.00 |
| RRAGB | 330.00 | 330.00 | 380.00 | 346.67 | 305.00 |
| RRAGC | 686.00 | 689.00 | 732.00 | 702.33 | 656.00 |
| RRAGD | 7.00 | 11.00 | 15.00 | 11.00 | 14.00 |
| RRAS | 313.00 | 333.00 | 365.00 | 337.00 | 409.00 |
| RRAS2 | 1406.00 | 1140.00 | 1517.00 | 1354.33 | 1784.00 |
| RRBP1 | 8205.00 | 8752.00 | 8480.00 | 8479.00 | 6543.00 |
| RREB1 | 1112.00 | 1010.00 | 1179.00 | 1100.33 | 931.00 |
| RRH | 4.00 | 3.00 | 3.00 | 3.33 | 4.00 |
| RRM1 | 1473.00 | 1414.00 | 1419.00 | 1435.33 | 2250.00 |
| RRM2 | 1326.00 | 1269.00 | 1367.00 | 1320.67 | 2406.00 |
| RRM2B | 516.00 | 457.00 | 566.00 | 513.00 | 625.00 |
| RRN3 | 11.00 | 6.00 | 10.00 | 9.00 | 13.00 |
| RRN3P1 | 341.00 | 293.00 | 317.00 | 317.00 | 320.00 |
| RRN3P2 | 23.00 | 18.00 | 28.00 | 23.00 | 12.00 |
| RRN3P3 | 109.00 | 108.00 | 102.00 | 106.33 | 117.00 |
| RRNAD1 | 1106.00 | 1083.00 | 1163.00 | 1117.33 | 1107.00 |
| RRP1 | 1130.00 | 1231.00 | 1114.00 | 1158.33 | 1341.00 |
| RRP12 | 3168.00 | 3048.00 | 3284.00 | 3166.67 | 3011.00 |
| RRP15 | 979.00 | 911.00 | 1082.00 | 990.67 | 1139.00 |
| RRP1B | 2245.00 | 2115.00 | 2419.00 | 2259.67 | 2235.00 |
| RRP36 | 767.00 | 709.00 | 868.00 | 781.33 | 936.00 |
| RRP7A | 700.00 | 694.00 | 695.00 | 696.33 | 814.00 |
| RRP7BP | 224.00 | 263.00 | 208.00 | 231.67 | 197.00 |
| RRP8 | 383.00 | 422.00 | 398.00 | 401.00 | 442.00 |
| RRP9 | 1163.00 | 1234.00 | 1318.00 | 1238.33 | 1377.00 |
| RRS1 | 1775.00 | 1847.00 | 1767.00 | 1796.33 | 2206.00 |
| RRS1-AS1 | 8.00 | 3.00 | 2.00 | 4.33 | 4.00 |
| RSAD1 | 2839.00 | 2922.00 | 3224.00 | 2995.00 | 3253.00 |
| RSBN1 | 430.00 | 364.00 | 448.00 | 414.00 | 380.00 |
| RSBN1L | 400.00 | 359.00 | 414.00 | 391.00 | 410.00 |

| | | | | | |
|---|---|---|---|---|---|
| RSC1A1 | 76.00 | 65.00 | 80.00 | 73.67 | 40.00 |
| RSF1 | 711.00 | 606.00 | 786.00 | 701.00 | 639.00 |
| RSG1 | 107.00 | 81.00 | 108.00 | 98.67 | 107.00 |
| RSL1D1 | 4055.00 | 4064.00 | 4645.00 | 4254.67 | 5121.00 |
| RSL24D1 | 3061.00 | 2988.00 | 3402.00 | 3150.33 | 4390.00 |
| RSPH3 | 106.00 | 98.00 | 113.00 | 105.67 | 76.00 |
| RSPH4A | 8.00 | 8.00 | 3.00 | 6.33 | 6.00 |
| RSPO2 | 16.00 | 9.00 | 6.00 | 10.33 | 12.00 |
| RSPRY1 | 1289.00 | 1141.00 | 1353.00 | 1261.00 | 1726.00 |
| RSRC1 | 371.00 | 389.00 | 404.00 | 388.00 | 473.00 |
| RSRC2 | 1557.00 | 1615.00 | 1631.00 | 1601.00 | 1961.00 |
| RSRP1 | 1596.00 | 1813.00 | 1855.00 | 1754.67 | 1690.00 |
| RSU1 | 1856.00 | 1682.00 | 1933.00 | 1823.67 | 2257.00 |
| RTCA | 1359.00 | 1262.00 | 1432.00 | 1351.00 | 1738.00 |
| RTCA-AS1 | 12.00 | 14.00 | 20.00 | 15.33 | 21.00 |
| RTCB | 1685.00 | 1686.00 | 1788.00 | 1719.67 | 2073.00 |
| RTEL1 | 151.00 | 157.00 | 143.00 | 150.33 | 144.00 |
| RTEL1-TNFRSF6B | 8.00 | 5.00 | 7.00 | 6.67 | 17.00 |
| RTF1 | 976.00 | 888.00 | 1094.00 | 986.00 | 1059.00 |
| RTFDC1 | 2161.00 | 1980.00 | 2250.00 | 2130.33 | 2728.00 |
| RTKN | 398.00 | 319.00 | 369.00 | 362.00 | 436.00 |
| RTKN2 | 140.00 | 132.00 | 169.00 | 147.00 | 196.00 |
| RTN1 | 388.00 | 392.00 | 415.00 | 398.33 | 652.00 |
| RTN2 | 142.00 | 144.00 | 116.00 | 134.00 | 103.00 |
| RTN3 | 6626.00 | 6273.00 | 6136.00 | 6345.00 | 7021.00 |
| RTN4 | 24993.00 | 23999.00 | 23762.00 | 24251.33 | 28744.00 |
| RTN4IP1 | 239.00 | 210.00 | 263.00 | 237.33 | 303.00 |
| RTN4RL2 | 32.00 | 34.00 | 27.00 | 31.00 | 37.00 |
| RTP4 | 6.00 | 3.00 | 4.00 | 4.33 | 3.00 |
| RTTN | 473.00 | 375.00 | 426.00 | 424.67 | 419.00 |
| RUFY1 | 787.00 | 768.00 | 841.00 | 798.67 | 824.00 |
| RUFY2 | 653.00 | 555.00 | 672.00 | 626.67 | 548.00 |
| RUFY3 | 1221.00 | 1167.00 | 1314.00 | 1234.00 | 1035.00 |
| RUNDC1 | 448.00 | 411.00 | 436.00 | 431.67 | 384.00 |
| RUNDC3A-AS1 | 9.00 | 4.00 | 8.00 | 7.00 | 9.00 |
| RUNDC3B | 27.00 | 27.00 | 23.00 | 25.67 | 34.00 |
| RUNX1 | 2161.00 | 1927.00 | 2270.00 | 2119.33 | 2123.00 |
| RUNX1-IT1 | 3.00 | 4.00 | 1.00 | 2.67 | 5.00 |
| RUNX1T1 | 234.00 | 193.00 | 228.00 | 218.33 | 323.00 |
| RUNX2 | 1098.00 | 999.00 | 1196.00 | 1097.67 | 959.00 |
| RUSC1 | 647.00 | 710.00 | 715.00 | 690.67 | 681.00 |
| RUSC1-AS1 | 31.00 | 25.00 | 22.00 | 26.00 | 19.00 |
| RUSC2 | 3799.00 | 3789.00 | 3870.00 | 3819.33 | 2813.00 |
| RUVBL1 | 2706.00 | 2583.00 | 2781.00 | 2690.00 | 3263.00 |
| RUVBL2 | 2265.00 | 2246.00 | 2401.00 | 2304.00 | 2944.00 |
| RWDD1 | 786.00 | 788.00 | 794.00 | 789.33 | 987.00 |
| RWDD2A | 260.00 | 242.00 | 309.00 | 270.33 | 280.00 |
| RWDD2B | 522.00 | 464.00 | 513.00 | 499.67 | 711.00 |

| | | | | | |
|---|---|---|---|---|---|
| RWDD3 | 273.00 | 326.00 | 266.00 | 288.33 | 339.00 |
| RWDD4 | 227.00 | 203.00 | 217.00 | 215.67 | 264.00 |
| RXFP2 | 136.00 | 149.00 | 167.00 | 150.67 | 172.00 |
| RXRA | 320.00 | 344.00 | 370.00 | 344.67 | 351.00 |
| RXRB | 11.00 | 1.00 | 11.00 | 7.67 | 6.00 |
| RYBP | 710.00 | 600.00 | 721.00 | 677.00 | 750.00 |
| RYK | 2399.00 | 2281.00 | 2370.00 | 2350.00 | 2822.00 |
| RYR2 | 101.00 | 84.00 | 81.00 | 88.67 | 77.00 |
| S100A10 | 373.00 | 344.00 | 409.00 | 375.33 | 394.00 |
| S100A11 | 1792.00 | 1734.00 | 1912.00 | 1812.67 | 2269.00 |
| S100A13 | 1074.00 | 1090.00 | 1148.00 | 1104.00 | 1454.00 |
| S100A14 | 7.00 | 5.00 | 14.00 | 8.67 | 11.00 |
| S100A16 | 3530.00 | 3334.00 | 3764.00 | 3542.67 | 4800.00 |
| S100A2 | 63.00 | 71.00 | 75.00 | 69.67 | 65.00 |
| S100A3 | 10.00 | 7.00 | 5.00 | 7.33 | 6.00 |
| S100A4 | 6.00 | 3.00 | 2.00 | 3.67 | 6.00 |
| S100A5 | 4.00 | 1.00 | 3.00 | 2.67 | 5.00 |
| S100A6 | 5180.00 | 5068.00 | 5458.00 | 5235.33 | 4884.00 |
| S100G | 14.00 | 8.00 | 7.00 | 9.67 | 19.00 |
| S100PBP | 863.00 | 818.00 | 952.00 | 877.67 | 862.00 |
| S1PR1 | 237.00 | 246.00 | 179.00 | 220.67 | 233.00 |
| S1PR2 | 105.00 | 89.00 | 103.00 | 99.00 | 78.00 |
| S1PR3 | 714.00 | 625.00 | 675.00 | 671.33 | 585.00 |
| SAAL1 | 525.00 | 512.00 | 514.00 | 517.00 | 660.00 |
| SAC3D1 | 238.00 | 275.00 | 286.00 | 266.33 | 377.00 |
| SACM1L | 1065.00 | 1099.00 | 1183.00 | 1115.67 | 1272.00 |
| SACS | 4761.00 | 4269.00 | 5236.00 | 4755.33 | 3888.00 |
| SAE1 | 2175.00 | 2082.00 | 2291.00 | 2182.67 | 2693.00 |
| SAFB | 2967.00 | 2770.00 | 3010.00 | 2915.67 | 3190.00 |
| SAFB2 | 1471.00 | 1481.00 | 1593.00 | 1515.00 | 1547.00 |
| SALL2 | 92.00 | 66.00 | 111.00 | 89.67 | 76.00 |
| SAMD1 | 397.00 | 407.00 | 446.00 | 416.67 | 474.00 |
| SAMD10 | 238.00 | 247.00 | 191.00 | 225.33 | 283.00 |
| SAMD12 | 296.00 | 254.00 | 358.00 | 302.67 | 238.00 |
| SAMD13 | 41.00 | 41.00 | 36.00 | 39.33 | 42.00 |
| SAMD15 | 22.00 | 18.00 | 14.00 | 18.00 | 25.00 |
| SAMD3 | 514.00 | 516.00 | 575.00 | 535.00 | 556.00 |
| SAMD4A | 2884.00 | 2787.00 | 2992.00 | 2887.67 | 3271.00 |
| SAMD4B | 2255.00 | 2046.00 | 2398.00 | 2233.00 | 1880.00 |
| SAMD5 | 138.00 | 128.00 | 162.00 | 142.67 | 153.00 |
| SAMD8 | 1471.00 | 1425.00 | 1550.00 | 1482.00 | 1420.00 |
| SAMD9 | 603.00 | 525.00 | 576.00 | 568.00 | 502.00 |
| SAMD9L | 367.00 | 372.00 | 362.00 | 367.00 | 305.00 |
| SAMHD1 | 570.00 | 513.00 | 529.00 | 537.33 | 522.00 |
| SAMM50 | 718.00 | 726.00 | 751.00 | 731.67 | 875.00 |
| SAMSN1 | 9.00 | 6.00 | 10.00 | 8.33 | 11.00 |
| SAP130 | 639.00 | 617.00 | 729.00 | 661.67 | 666.00 |
| SAP18 | 1247.00 | 1154.00 | 1438.00 | 1279.67 | 1724.00 |

| | | | | | |
|---|---|---|---|---|---|
| SAP25 | 113.00 | 120.00 | 143.00 | 125.33 | 107.00 |
| SAP30 | 136.00 | 160.00 | 160.00 | 152.00 | 217.00 |
| SAP30BP | 933.00 | 1053.00 | 1051.00 | 1012.33 | 1186.00 |
| SAP30L | 550.00 | 501.00 | 630.00 | 560.33 | 504.00 |
| SAP30L-AS1 | 20.00 | 15.00 | 15.00 | 16.67 | 11.00 |
| SAPCD2 | 543.00 | 599.00 | 724.00 | 622.00 | 682.00 |
| SAR1A | 1990.00 | 1879.00 | 2038.00 | 1969.00 | 2414.00 |
| SAR1B | 975.00 | 853.00 | 963.00 | 930.33 | 1115.00 |
| SARAF | 1906.00 | 1861.00 | 1888.00 | 1885.00 | 1769.00 |
| SARDH | 80.00 | 62.00 | 90.00 | 77.33 | 60.00 |
| SARM1 | 820.00 | 770.00 | 832.00 | 807.33 | 840.00 |
| SARNP | 613.00 | 618.00 | 606.00 | 612.33 | 773.00 |
| SARS | 9673.00 | 9144.00 | 10046.00 | 9621.00 | 10974.00 |
| SARS2 | 401.00 | 405.00 | 427.00 | 411.00 | 495.00 |
| SART1 | 3119.00 | 3283.00 | 3159.00 | 3187.00 | 3765.00 |
| SART3 | 1385.00 | 1392.00 | 1524.00 | 1433.67 | 1495.00 |
| SASH1 | 2207.00 | 2020.00 | 2298.00 | 2175.00 | 2715.00 |
| SASS6 | 472.00 | 419.00 | 487.00 | 459.33 | 517.00 |
| SAT1 | 401.00 | 399.00 | 515.00 | 438.33 | 371.00 |
| SAT2 | 841.00 | 888.00 | 972.00 | 900.33 | 961.00 |
| SATB1 | 629.00 | 547.00 | 595.00 | 590.33 | 560.00 |
| SATB1-AS1 | 7.00 | 2.00 | 4.00 | 4.33 | 10.00 |
| SATB2 | 985.00 | 1040.00 | 1108.00 | 1044.33 | 1033.00 |
| SAV1 | 288.00 | 242.00 | 303.00 | 277.67 | 307.00 |
| SAXO2 | 12.00 | 11.00 | 7.00 | 10.00 | 8.00 |
| SAYSD1 | 341.00 | 282.00 | 355.00 | 326.00 | 373.00 |
| SBDS | 1819.00 | 1788.00 | 1839.00 | 1815.33 | 1942.00 |
| SBDSP1 | 626.00 | 634.00 | 686.00 | 648.67 | 837.00 |
| SBF1 | 2382.00 | 2432.00 | 2514.00 | 2442.67 | 2187.00 |
| SBF2 | 1298.00 | 1169.00 | 1385.00 | 1284.00 | 1123.00 |
| SBF2-AS1 | 77.00 | 84.00 | 75.00 | 78.67 | 77.00 |
| SBK1 | 5.00 | 1.00 | 11.00 | 5.67 | 7.00 |
| SBNO1 | 1461.00 | 1416.00 | 1702.00 | 1526.33 | 1647.00 |
| SBNO2 | 51.00 | 56.00 | 68.00 | 58.33 | 37.00 |
| SBSN | 267.00 | 287.00 | 253.00 | 269.00 | 225.00 |
| SC5D | 658.00 | 637.00 | 609.00 | 634.67 | 749.00 |
| SCAF1 | 1608.00 | 1526.00 | 1604.00 | 1579.33 | 1468.00 |
| SCAF11 | 2532.00 | 2420.00 | 2687.00 | 2546.33 | 2344.00 |
| SCAF4 | 1267.00 | 1247.00 | 1358.00 | 1290.67 | 1296.00 |
| SCAF8 | 1863.00 | 1759.00 | 1861.00 | 1827.67 | 1997.00 |
| SCAI | 294.00 | 276.00 | 354.00 | 308.00 | 269.00 |
| SCAMP1 | 871.00 | 717.00 | 844.00 | 810.67 | 861.00 |
| SCAMP1-AS1 | 151.00 | 182.00 | 166.00 | 166.33 | 185.00 |
| SCAMP2 | 1355.00 | 1373.00 | 1290.00 | 1339.33 | 1246.00 |
| SCAMP4 | 1856.00 | 1928.00 | 1743.00 | 1842.33 | 1791.00 |
| SCAMP5 | 38.00 | 48.00 | 60.00 | 48.67 | 84.00 |
| SCAND1 | 543.00 | 546.00 | 538.00 | 542.33 | 718.00 |
| SCAND2P | 223.00 | 197.00 | 179.00 | 199.67 | 200.00 |

| | | | | | |
|---|---|---|---|---|---|
| SCAP | 3015.00 | 3094.00 | 2964.00 | 3024.33 | 2689.00 |
| SCAPER | 248.00 | 203.00 | 235.00 | 228.67 | 276.00 |
| SCARA3 | 25.00 | 17.00 | 24.00 | 22.00 | 15.00 |
| SCARB1 | 2517.00 | 2412.00 | 2313.00 | 2414.00 | 2475.00 |
| SCARB2 | 4356.00 | 4169.00 | 4389.00 | 4304.67 | 4740.00 |
| SCARF2 | 900.00 | 969.00 | 922.00 | 930.33 | 797.00 |
| SCARNA1 | 3.00 | 2.00 | 0.00 | 1.67 | 0.00 |
| SCARNA12 | 147.00 | 139.00 | 128.00 | 138.00 | 151.00 |
| SCARNA15 | 41.00 | 30.00 | 26.00 | 32.33 | 34.00 |
| SCARNA16 | 17.00 | 20.00 | 18.00 | 18.33 | 15.00 |
| SCARNA17 | 9.00 | 2.00 | 2.00 | 4.33 | 6.00 |
| SCARNA2 | 3.00 | 1.00 | 2.00 | 2.00 | 5.00 |
| SCARNA22 | 15.00 | 19.00 | 21.00 | 18.33 | 18.00 |
| SCARNA27 | 39.00 | 40.00 | 33.00 | 37.33 | 22.00 |
| SCARNA4 | 4.00 | 5.00 | 1.00 | 3.33 | 5.00 |
| SCARNA5 | 3.00 | 4.00 | 4.00 | 3.67 | 4.00 |
| SCARNA6 | 4.00 | 3.00 | 1.00 | 2.67 | 5.00 |
| SCARNA7 | 4.00 | 1.00 | 1.00 | 2.00 | 6.00 |
| SCARNA9 | 7.00 | 6.00 | 4.00 | 5.67 | 7.00 |
| SCART1 | 189.00 | 221.00 | 225.00 | 211.67 | 236.00 |
| SCCPDH | 1258.00 | 1159.00 | 1219.00 | 1212.00 | 1543.00 |
| SCD | 9500.00 | 8804.00 | 9105.00 | 9136.33 | 10727.00 |
| SCD5 | 211.00 | 164.00 | 185.00 | 186.67 | 189.00 |
| SCEL | 25.00 | 22.00 | 13.00 | 20.00 | 21.00 |
| SCFD1 | 978.00 | 946.00 | 902.00 | 942.00 | 1124.00 |
| SCFD2 | 470.00 | 397.00 | 497.00 | 454.67 | 496.00 |
| SCG2 | 46.00 | 39.00 | 55.00 | 46.67 | 53.00 |
| SCHIP1 | 4.00 | 3.00 | 3.00 | 3.33 | 2.00 |
| SCLT1 | 391.00 | 382.00 | 366.00 | 379.67 | 398.00 |
| SCLY | 22.00 | 12.00 | 23.00 | 19.00 | 21.00 |
| SCMH1 | 1460.00 | 1361.00 | 1350.00 | 1390.33 | 1457.00 |
| SCML1 | 284.00 | 271.00 | 290.00 | 281.67 | 389.00 |
| SCML2 | 148.00 | 145.00 | 149.00 | 147.33 | 162.00 |
| SCN11A | 3.00 | 0.00 | 0.00 | 1.00 | 0.00 |
| SCN1B | 10.00 | 4.00 | 7.00 | 7.00 | 4.00 |
| SCN2A | 33.00 | 29.00 | 29.00 | 30.33 | 46.00 |
| SCN3A | 6.00 | 5.00 | 6.00 | 5.67 | 4.00 |
| SCN5A | 317.00 | 361.00 | 359.00 | 345.67 | 146.00 |
| SCN7A | 8.00 | 12.00 | 4.00 | 8.00 | 2.00 |
| SCN8A | 982.00 | 1003.00 | 1010.00 | 998.33 | 690.00 |
| SCN9A | 27.00 | 30.00 | 22.00 | 26.33 | 16.00 |
| SCNM1 | 86.00 | 77.00 | 103.00 | 88.67 | 122.00 |
| SCNN1A | 16.00 | 12.00 | 19.00 | 15.67 | 16.00 |
| SCNN1D | 518.00 | 586.00 | 520.00 | 541.33 | 484.00 |
| SCO1 | 852.00 | 765.00 | 853.00 | 823.33 | 940.00 |
| SCO2 | 296.00 | 282.00 | 298.00 | 292.00 | 283.00 |
| SCOC | 859.00 | 737.00 | 903.00 | 833.00 | 1073.00 |
| SCOC-AS1 | 6.00 | 14.00 | 7.00 | 9.00 | 23.00 |

| | | | | | |
|---|---|---|---|---|---|
| SCP2 | 954.00 | 819.00 | 923.00 | 898.67 | 1081.00 |
| SCPEP1 | 1330.00 | 1357.00 | 1361.00 | 1349.33 | 1472.00 |
| SCRIB | 6.00 | 15.00 | 3.00 | 8.00 | 16.00 |
| SCRN1 | 8947.00 | 7939.00 | 9152.00 | 8679.33 | 8465.00 |
| SCRN2 | 734.00 | 687.00 | 663.00 | 694.67 | 789.00 |
| SCRN3 | 537.00 | 515.00 | 527.00 | 526.33 | 588.00 |
| SCUBE2 | 7.00 | 7.00 | 5.00 | 6.33 | 10.00 |
| SCUBE3 | 44.00 | 47.00 | 38.00 | 43.00 | 54.00 |
| SCX | 76.00 | 109.00 | 105.00 | 96.67 | 116.00 |
| SCYL1 | 3083.00 | 3145.00 | 3216.00 | 3148.00 | 3371.00 |
| SCYL2 | 761.00 | 742.00 | 821.00 | 774.67 | 843.00 |
| SCYL3 | 380.00 | 319.00 | 401.00 | 366.67 | 374.00 |
| SDAD1 | 1120.00 | 1033.00 | 1191.00 | 1114.67 | 1335.00 |
| SDC1 | 5692.00 | 5998.00 | 5727.00 | 5805.67 | 8906.00 |
| SDC2 | 84.00 | 105.00 | 87.00 | 92.00 | 63.00 |
| SDC3 | 1234.00 | 1197.00 | 1232.00 | 1221.00 | 1058.00 |
| SDC4 | 1007.00 | 966.00 | 900.00 | 957.67 | 1380.00 |
| SDCBP | 3092.00 | 2791.00 | 3290.00 | 3057.67 | 3264.00 |
| SDCBP2-AS1 | 84.00 | 105.00 | 61.00 | 83.33 | 122.00 |
| SDCCAG3 | 2670.00 | 2697.00 | 2822.00 | 2729.67 | 3271.00 |
| SDCCAG8 | 7.00 | 28.00 | 12.00 | 15.67 | 22.00 |
| SDE2 | 606.00 | 550.00 | 591.00 | 582.33 | 585.00 |
| SDF2 | 751.00 | 686.00 | 733.00 | 723.33 | 758.00 |
| SDF2L1 | 366.00 | 369.00 | 333.00 | 356.00 | 471.00 |
| SDF4 | 4672.00 | 4885.00 | 4347.00 | 4634.67 | 4887.00 |
| SDHA | 2398.00 | 2443.00 | 2569.00 | 2470.00 | 2683.00 |
| SDHAF1 | 149.00 | 157.00 | 174.00 | 160.00 | 173.00 |
| SDHAF2 | 538.00 | 458.00 | 572.00 | 522.67 | 658.00 |
| SDHAF3 | 143.00 | 137.00 | 148.00 | 142.67 | 198.00 |
| SDHAF4 | 141.00 | 140.00 | 176.00 | 152.33 | 232.00 |
| SDHAP1 | 416.00 | 402.00 | 414.00 | 410.67 | 324.00 |
| SDHAP2 | 8.00 | 1.00 | 1.00 | 3.33 | 0.00 |
| SDHAP3 | 25.00 | 16.00 | 19.00 | 20.00 | 13.00 |
| SDHB | 1139.00 | 1142.00 | 1198.00 | 1159.67 | 1526.00 |
| SDHC | 1283.00 | 1187.00 | 1321.00 | 1263.67 | 1305.00 |
| SDHD | 1249.00 | 1171.00 | 1361.00 | 1260.33 | 1073.00 |
| SDK1 | 24.00 | 18.00 | 28.00 | 23.33 | 20.00 |
| SDR39U1 | 605.00 | 699.00 | 699.00 | 667.67 | 806.00 |
| SDSL | 135.00 | 188.00 | 166.00 | 163.00 | 173.00 |
| SEC11A | 3087.00 | 3071.00 | 3049.00 | 3069.00 | 4007.00 |
| SEC11C | 431.00 | 465.00 | 442.00 | 446.00 | 588.00 |
| SEC13 | 2129.00 | 2008.00 | 2290.00 | 2142.33 | 2385.00 |
| SEC14L1 | 2680.00 | 2418.00 | 2757.00 | 2618.33 | 2325.00 |
| SEC14L1P1 | 58.00 | 55.00 | 49.00 | 54.00 | 61.00 |
| SEC14L2 | 245.00 | 294.00 | 284.00 | 274.33 | 298.00 |
| SEC16A | 5157.00 | 4647.00 | 5003.00 | 4935.67 | 3996.00 |
| SEC16B | 138.00 | 140.00 | 162.00 | 146.67 | 164.00 |
| SEC22A | 508.00 | 471.00 | 513.00 | 497.33 | 565.00 |

| | | | | | |
|---|---|---|---|---|---|
| SEC22B | 1666.00 | 1527.00 | 1693.00 | 1628.67 | 1390.00 |
| SEC22C | 1272.00 | 1299.00 | 1332.00 | 1301.00 | 1247.00 |
| SEC23A | 1636.00 | 1429.00 | 1792.00 | 1619.00 | 1619.00 |
| SEC23B | 1767.00 | 1578.00 | 1862.00 | 1735.67 | 1848.00 |
| SEC23IP | 1769.00 | 1578.00 | 1814.00 | 1720.33 | 1760.00 |
| SEC24A | 960.00 | 756.00 | 1026.00 | 914.00 | 650.00 |
| SEC24B | 886.00 | 765.00 | 954.00 | 868.33 | 894.00 |
| SEC24B-AS1 | 26.00 | 27.00 | 13.00 | 22.00 | 13.00 |
| SEC24C | 4496.00 | 4296.00 | 4639.00 | 4477.00 | 4010.00 |
| SEC24D | 1263.00 | 1198.00 | 1283.00 | 1248.00 | 996.00 |
| SEC31A | 2574.00 | 2379.00 | 2765.00 | 2572.67 | 2295.00 |
| SEC31B | 923.00 | 1003.00 | 1036.00 | 987.33 | 762.00 |
| SEC61A1 | 18381.00 | 17967.00 | 18341.00 | 18229.67 | 15624.00 |
| SEC61A2 | 439.00 | 409.00 | 482.00 | 443.33 | 429.00 |
| SEC61B | 1033.00 | 954.00 | 907.00 | 964.67 | 1194.00 |
| SEC61G | 838.00 | 829.00 | 871.00 | 846.00 | 1129.00 |
| SEC62 | 3241.00 | 3112.00 | 3287.00 | 3213.33 | 3434.00 |
| SEC63 | 5058.00 | 4766.00 | 4814.00 | 4879.33 | 5457.00 |
| SECISBP2 | 1435.00 | 1404.00 | 1477.00 | 1438.67 | 1665.00 |
| SECISBP2L | 871.00 | 726.00 | 937.00 | 844.67 | 749.00 |
| SECTM1 | 4.00 | 9.00 | 11.00 | 8.00 | 9.00 |
| SEH1L | 1632.00 | 1647.00 | 1942.00 | 1740.33 | 2355.00 |
| SEL1L | 3938.00 | 3574.00 | 4067.00 | 3859.67 | 2821.00 |
| SEL1L3 | 717.00 | 655.00 | 719.00 | 697.00 | 405.00 |
| SELENBP1 | 24.00 | 22.00 | 20.00 | 22.00 | 42.00 |
| SELK | 449.00 | 500.00 | 486.00 | 478.33 | 600.00 |
| SELM | 16.00 | 23.00 | 19.00 | 19.33 | 19.00 |
| SELO | 562.00 | 592.00 | 643.00 | 599.00 | 660.00 |
| SELPLG | 33.00 | 18.00 | 22.00 | 24.33 | 19.00 |
| SELT | 1764.00 | 1590.00 | 1755.00 | 1703.00 | 2226.00 |
| SEMA3A | 2075.00 | 1811.00 | 2152.00 | 2012.67 | 1638.00 |
| SEMA3B | 140.00 | 159.00 | 141.00 | 146.67 | 78.00 |
| SEMA3C | 843.00 | 755.00 | 965.00 | 854.33 | 1493.00 |
| SEMA3D | 29.00 | 29.00 | 34.00 | 30.67 | 20.00 |
| SEMA3F | 48.00 | 59.00 | 49.00 | 52.00 | 24.00 |
| SEMA4A | 90.00 | 103.00 | 79.00 | 90.67 | 128.00 |
| SEMA4B | 1457.00 | 1470.00 | 1335.00 | 1420.67 | 880.00 |
| SEMA4C | 3004.00 | 3220.00 | 3248.00 | 3157.33 | 2181.00 |
| SEMA4D | 169.00 | 145.00 | 161.00 | 158.33 | 143.00 |
| SEMA4F | 812.00 | 775.00 | 800.00 | 795.67 | 801.00 |
| SEMA4G | 64.00 | 47.00 | 53.00 | 54.67 | 54.00 |
| SEMA5A | 836.00 | 783.00 | 877.00 | 832.00 | 1074.00 |
| SEMA6A | 35.00 | 13.00 | 21.00 | 23.00 | 20.00 |
| SEMA6A-AS1 | 17.00 | 28.00 | 38.00 | 27.67 | 24.00 |
| SEMA6B | 97.00 | 110.00 | 107.00 | 104.67 | 47.00 |
| SEMA6C | 231.00 | 226.00 | 256.00 | 237.67 | 260.00 |
| SEMA6D | 23.00 | 19.00 | 17.00 | 19.67 | 9.00 |
| SEMA7A | 8845.00 | 8953.00 | 8553.00 | 8783.67 | 4378.00 |

| | | | | | |
|---|---|---|---|---|---|
| SENCR | 72.00 | 57.00 | 73.00 | 67.33 | 89.00 |
| SENP1 | 579.00 | 516.00 | 627.00 | 574.00 | 593.00 |
| SENP2 | 823.00 | 756.00 | 819.00 | 799.33 | 963.00 |
| SENP3 | 382.00 | 337.00 | 363.00 | 360.67 | 450.00 |
| SENP3-EIF4A1 | 28.00 | 26.00 | 29.00 | 27.67 | 18.00 |
| SENP5 | 1371.00 | 1271.00 | 1440.00 | 1360.67 | 1349.00 |
| SENP6 | 2341.00 | 2354.00 | 2611.00 | 2435.33 | 2785.00 |
| SENP7 | 325.00 | 327.00 | 353.00 | 335.00 | 314.00 |
| SENP8 | 62.00 | 56.00 | 52.00 | 56.67 | 58.00 |
| SEP1. | 16.00 | 15.00 | 18.00 | 16.33 | 18.00 |
| SEP10. | 1286.00 | 1165.00 | 1370.00 | 1273.67 | 1570.00 |
| SEP11. | 2363.00 | 2067.00 | 2416.00 | 2282.00 | 2759.00 |
| SEP15. | 3099.00 | 3111.00 | 2992.00 | 3067.33 | 4094.00 |
| SEP2. | 7394.00 | 6868.00 | 7878.00 | 7380.00 | 8219.00 |
| SEP6. | 55.00 | 59.00 | 55.00 | 56.33 | 51.00 |
| SEP7. | 3122.00 | 2861.00 | 3144.00 | 3042.33 | 3668.00 |
| SEP8. | 1845.00 | 1854.00 | 1801.00 | 1833.33 | 1838.00 |
| SEP9. | 2675.00 | 2554.00 | 2738.00 | 2655.67 | 2446.00 |
| SEPHS1 | 1847.00 | 1743.00 | 1782.00 | 1790.67 | 2241.00 |
| SEPHS2 | 1399.00 | 1316.00 | 1432.00 | 1382.33 | 1739.00 |
| SEPN1 | 3063.00 | 2970.00 | 2992.00 | 3008.33 | 2894.00 |
| SEPSECS | 260.00 | 216.00 | 285.00 | 253.67 | 267.00 |
| SEPSECS-AS1 | 3.00 | 6.00 | 10.00 | 6.33 | 8.00 |
| SEPT7-AS1 | 25.00 | 26.00 | 21.00 | 24.00 | 23.00 |
| SEPT7P2 | 425.00 | 449.00 | 482.00 | 452.00 | 372.00 |
| SEPT7P9 | 6.00 | 10.00 | 8.00 | 8.00 | 5.00 |
| SEPW1 | 351.00 | 306.00 | 360.00 | 339.00 | 452.00 |
| SERAC1 | 139.00 | 139.00 | 144.00 | 140.67 | 126.00 |
| SERBP1 | 7410.00 | 6989.00 | 7973.00 | 7457.33 | 9721.00 |
| SERF2 | 3509.00 | 3331.00 | 3571.00 | 3470.33 | 4286.00 |
| SERGEF | 221.00 | 204.00 | 261.00 | 228.67 | 271.00 |
| SERINC1 | 6220.00 | 5981.00 | 6391.00 | 6197.33 | 6231.00 |
| SERINC2 | 5170.00 | 4787.00 | 4725.00 | 4894.00 | 967.00 |
| SERINC3 | 5113.00 | 4642.00 | 4965.00 | 4906.67 | 4260.00 |
| SERINC4 | 3.00 | 5.00 | 8.00 | 5.33 | 3.00 |
| SERINC5 | 400.00 | 453.00 | 385.00 | 412.67 | 328.00 |
| SERP1 | 2850.00 | 2664.00 | 3142.00 | 2885.33 | 3342.00 |
| SERP2 | 3.00 | 6.00 | 9.00 | 6.00 | 10.00 |
| SERPINA1 | 15.00 | 10.00 | 12.00 | 12.33 | 35.00 |
| SERPINA3 | 38.00 | 34.00 | 38.00 | 36.67 | 110.00 |
| SERPINA5 | 71.00 | 76.00 | 86.00 | 77.67 | 146.00 |
| SERPINB1 | 724.00 | 659.00 | 859.00 | 747.33 | 863.00 |
| SERPINB2 | 13807.00 | 11529.00 | 13951.00 | 13095.67 | 20160.00 |
| SERPINB6 | 1885.00 | 1798.00 | 1991.00 | 1891.33 | 2276.00 |
| SERPINB7 | 1193.00 | 1087.00 | 1091.00 | 1123.67 | 1367.00 |
| SERPINB8 | 1629.00 | 1516.00 | 1610.00 | 1585.00 | 2294.00 |
| SERPINB9P1 | 5.00 | 5.00 | 10.00 | 6.67 | 6.00 |
| SERPIND1 | 166.00 | 145.00 | 145.00 | 152.00 | 171.00 |

| | | | | | |
|---|---|---|---|---|---|
| SERPINE1 | 82134.00 | 78740.00 | 84922.00 | 81932.00 | 63676.00 |
| SERPINE2 | 729.00 | 739.00 | 644.00 | 704.00 | 569.00 |
| SERPINF1 | 582.00 | 588.00 | 667.00 | 612.33 | 838.00 |
| SERPINH1 | 10152.00 | 10495.00 | 9441.00 | 10029.33 | 8992.00 |
| SERPINI1 | 136.00 | 121.00 | 118.00 | 125.00 | 147.00 |
| SERTAD1 | 225.00 | 245.00 | 282.00 | 250.67 | 312.00 |
| SERTAD2 | 1236.00 | 1173.00 | 1448.00 | 1285.67 | 1490.00 |
| SERTAD3 | 177.00 | 189.00 | 193.00 | 186.33 | 220.00 |
| SERTAD4 | 10.00 | 10.00 | 8.00 | 9.33 | 5.00 |
| SERTAD4-AS1 | 13.00 | 13.00 | 18.00 | 14.67 | 6.00 |
| SESN1 | 145.00 | 121.00 | 169.00 | 145.00 | 158.00 |
| SESN2 | 4167.00 | 3870.00 | 4141.00 | 4059.33 | 4361.00 |
| SESN3 | 301.00 | 255.00 | 317.00 | 291.00 | 244.00 |
| SESTD1 | 239.00 | 233.00 | 267.00 | 246.33 | 219.00 |
| SET | 8724.00 | 8318.00 | 9143.00 | 8728.33 | 11367.00 |
| SETBP1 | 125.00 | 103.00 | 126.00 | 118.00 | 69.00 |
| SETD1A | 1234.00 | 1173.00 | 1338.00 | 1248.33 | 1063.00 |
| SETD1B | 1003.00 | 976.00 | 1015.00 | 998.00 | 808.00 |
| SETD2 | 1972.00 | 1944.00 | 2175.00 | 2030.33 | 1653.00 |
| SETD3 | 914.00 | 808.00 | 888.00 | 870.00 | 1029.00 |
| SETD4 | 824.00 | 855.00 | 857.00 | 845.33 | 845.00 |
| SETD5 | 4510.00 | 4518.00 | 4952.00 | 4660.00 | 4225.00 |
| SETD6 | 746.00 | 780.00 | 774.00 | 766.67 | 803.00 |
| SETD7 | 1387.00 | 1255.00 | 1475.00 | 1372.33 | 1351.00 |
| SETD8 | 1482.00 | 1424.00 | 1489.00 | 1465.00 | 1511.00 |
| SETD9 | 134.00 | 148.00 | 168.00 | 150.00 | 223.00 |
| SETDB1 | 1643.00 | 1433.00 | 1615.00 | 1563.67 | 1508.00 |
| SETDB2 | 342.00 | 330.00 | 378.00 | 350.00 | 355.00 |
| SETMAR | 188.00 | 132.00 | 147.00 | 155.67 | 178.00 |
| SETSIP | 33.00 | 26.00 | 30.00 | 29.67 | 44.00 |
| SETX | 1722.00 | 1614.00 | 1882.00 | 1739.33 | 1694.00 |
| SEZ6L2 | 777.00 | 795.00 | 838.00 | 803.33 | 660.00 |
| SF1 | 8138.00 | 8231.00 | 8738.00 | 8369.00 | 8693.00 |
| SF3A1 | 2940.00 | 2687.00 | 2931.00 | 2852.67 | 2741.00 |
| SF3A2 | 1542.00 | 1571.00 | 1737.00 | 1616.67 | 1659.00 |
| SF3A3 | 2608.00 | 2541.00 | 2736.00 | 2628.33 | 3101.00 |
| SF3B1 | 10804.00 | 10071.00 | 11521.00 | 10798.67 | 10977.00 |
| SF3B2 | 7886.00 | 7461.00 | 8019.00 | 7788.67 | 8683.00 |
| SF3B3 | 7363.00 | 6954.00 | 7763.00 | 7360.00 | 8079.00 |
| SF3B4 | 1643.00 | 1513.00 | 1599.00 | 1585.00 | 1843.00 |
| SF3B5 | 1105.00 | 1080.00 | 1086.00 | 1090.33 | 1396.00 |
| SF3B6 | 1919.00 | 1837.00 | 1887.00 | 1881.00 | 3510.00 |
| SFI1 | 808.00 | 862.00 | 801.00 | 823.67 | 919.00 |
| SFMBT1 | 379.00 | 367.00 | 394.00 | 380.00 | 380.00 |
| SFMBT2 | 601.00 | 553.00 | 645.00 | 599.67 | 580.00 |
| SFPQ | 6921.00 | 6421.00 | 6966.00 | 6769.33 | 8404.00 |
| SFR1 | 164.00 | 135.00 | 170.00 | 156.33 | 205.00 |
| SFRP1 | 12762.00 | 12019.00 | 12886.00 | 12555.67 | 13839.00 |

| | | | | | |
|---|---|---|---|---|---|
| SFSWAP | 1017.00 | 1031.00 | 1100.00 | 1049.33 | 994.00 |
| SFT2D1 | 1816.00 | 1874.00 | 1891.00 | 1860.33 | 1723.00 |
| SFT2D2 | 86.00 | 79.00 | 100.00 | 88.33 | 101.00 |
| SFT2D3 | 176.00 | 192.00 | 186.00 | 184.67 | 276.00 |
| SFTA1P | 8.00 | 8.00 | 9.00 | 8.33 | 2.00 |
| SFXN1 | 988.00 | 890.00 | 954.00 | 944.00 | 1122.00 |
| SFXN2 | 255.00 | 224.00 | 227.00 | 235.33 | 235.00 |
| SFXN3 | 602.00 | 589.00 | 675.00 | 622.00 | 977.00 |
| SFXN4 | 1155.00 | 1089.00 | 1279.00 | 1174.33 | 1565.00 |
| SFXN5 | 360.00 | 353.00 | 347.00 | 353.33 | 276.00 |
| SGCB | 687.00 | 675.00 | 621.00 | 661.00 | 718.00 |
| SGCE | 1004.00 | 992.00 | 996.00 | 997.33 | 976.00 |
| SGCG | 3.00 | 3.00 | 1.00 | 2.33 | 1.00 |
| SGCZ | 12.00 | 11.00 | 18.00 | 13.67 | 14.00 |
| SGK1 | 3693.00 | 3748.00 | 4095.00 | 3845.33 | 6173.00 |
| SGK2 | 8.00 | 7.00 | 7.00 | 7.33 | 9.00 |
| SGK223 | 689.00 | 661.00 | 747.00 | 699.00 | 390.00 |
| SGK3 | 3.00 | 0.00 | 3.00 | 2.00 | 6.00 |
| SGK494 | 807.00 | 787.00 | 859.00 | 817.67 | 643.00 |
| SGMS1 | 520.00 | 517.00 | 641.00 | 559.33 | 524.00 |
| SGMS1-AS1 | 52.00 | 50.00 | 55.00 | 52.33 | 53.00 |
| SGMS2 | 342.00 | 288.00 | 320.00 | 316.67 | 232.00 |
| SGOL1 | 256.00 | 256.00 | 253.00 | 255.00 | 319.00 |
| SGOL2 | 380.00 | 342.00 | 387.00 | 369.67 | 479.00 |
| SGPL1 | 1909.00 | 1822.00 | 1850.00 | 1860.33 | 2380.00 |
| SGPP1 | 1061.00 | 1033.00 | 1062.00 | 1052.00 | 824.00 |
| SGSH | 1108.00 | 1176.00 | 1084.00 | 1122.67 | 1026.00 |
| SGSM2 | 3675.00 | 3904.00 | 4183.00 | 3920.67 | 3543.00 |
| SGSM3 | 1503.00 | 1509.00 | 1622.00 | 1544.67 | 1569.00 |
| SGTA | 1644.00 | 1665.00 | 1732.00 | 1680.33 | 1930.00 |
| SGTB | 455.00 | 381.00 | 485.00 | 440.33 | 425.00 |
| SH2B1 | 1930.00 | 1899.00 | 1982.00 | 1937.00 | 1782.00 |
| SH2B2 | 68.00 | 74.00 | 72.00 | 71.33 | 57.00 |
| SH2B3 | 4394.00 | 4020.00 | 4466.00 | 4293.33 | 3138.00 |
| SH2D1B | 3.00 | 5.00 | 4.00 | 4.00 | 1.00 |
| SH2D2A | 9.00 | 11.00 | 11.00 | 10.33 | 18.00 |
| SH2D5 | 1265.00 | 1276.00 | 1369.00 | 1303.33 | 1310.00 |
| SH2D6 | 5.00 | 5.00 | 3.00 | 4.33 | 1.00 |
| SH3BGR | 36.00 | 28.00 | 30.00 | 31.33 | 31.00 |
| SH3BGRL | 1131.00 | 931.00 | 1247.00 | 1103.00 | 1277.00 |
| SH3BGRL2 | 267.00 | 246.00 | 254.00 | 255.67 | 241.00 |
| SH3BGRL3 | 2798.00 | 2639.00 | 2868.00 | 2768.33 | 3195.00 |
| SH3BP1 | 234.00 | 236.00 | 277.00 | 249.00 | 261.00 |
| SH3BP2 | 9948.00 | 9773.00 | 10895.00 | 10205.33 | 7814.00 |
| SH3BP4 | 4188.00 | 3862.00 | 4539.00 | 4196.33 | 2261.00 |
| SH3BP5 | 1226.00 | 1223.00 | 1351.00 | 1266.67 | 1504.00 |
| SH3BP5-AS1 | 327.00 | 393.00 | 420.00 | 380.00 | 326.00 |
| SH3BP5L | 3643.00 | 3522.00 | 3447.00 | 3537.33 | 3152.00 |

| | | | | | |
|---|---|---|---|---|---|
| SH3D19 | 524.00 | 510.00 | 572.00 | 535.33 | 558.00 |
| SH3D21 | 485.00 | 520.00 | 503.00 | 502.67 | 312.00 |
| SH3GL1 | 1157.00 | 1108.00 | 1154.00 | 1139.67 | 1440.00 |
| SH3GL1P1 | 27.00 | 29.00 | 21.00 | 25.67 | 13.00 |
| SH3GL2 | 7.00 | 13.00 | 16.00 | 12.00 | 22.00 |
| SH3GL3 | 26.00 | 28.00 | 36.00 | 30.00 | 53.00 |
| SH3GLB1 | 1440.00 | 1355.00 | 1574.00 | 1456.33 | 1568.00 |
| SH3GLB2 | 1283.00 | 1349.00 | 1441.00 | 1357.67 | 1575.00 |
| SH3KBP1 | 5039.00 | 4853.00 | 4917.00 | 4936.33 | 4449.00 |
| SH3PXD2A | 5203.00 | 4899.00 | 5550.00 | 5217.33 | 4523.00 |
| SH3PXD2B | 1199.00 | 1096.00 | 1263.00 | 1186.00 | 782.00 |
| SH3RF1 | 753.00 | 638.00 | 740.00 | 710.33 | 629.00 |
| SH3RF2 | 134.00 | 126.00 | 111.00 | 123.67 | 122.00 |
| SH3RF3 | 908.00 | 830.00 | 887.00 | 875.00 | 704.00 |
| SH3RF3-AS1 | 7.00 | 11.00 | 9.00 | 9.00 | 11.00 |
| SH3TC1 | 193.00 | 185.00 | 249.00 | 209.00 | 214.00 |
| SH3TC2 | 1024.00 | 952.00 | 1048.00 | 1008.00 | 1688.00 |
| SH3YL1 | 1095.00 | 1047.00 | 1173.00 | 1105.00 | 1033.00 |
| SHANK1 | 981.00 | 914.00 | 1095.00 | 996.67 | 799.00 |
| SHANK2 | 1366.00 | 1266.00 | 1439.00 | 1357.00 | 1112.00 |
| SHARPIN | 1031.00 | 1064.00 | 1133.00 | 1076.00 | 1070.00 |
| SHB | 773.00 | 694.00 | 786.00 | 751.00 | 818.00 |
| SHBG | 7.00 | 5.00 | 6.00 | 6.00 | 3.00 |
| SHC1 | 6559.00 | 6365.00 | 6925.00 | 6616.33 | 6563.00 |
| SHC2 | 67.00 | 60.00 | 73.00 | 66.67 | 40.00 |
| SHC3 | 49.00 | 58.00 | 50.00 | 52.33 | 42.00 |
| SHC4 | 35.00 | 30.00 | 41.00 | 35.33 | 16.00 |
| SHCBP1 | 638.00 | 600.00 | 638.00 | 625.33 | 733.00 |
| SHF | 28.00 | 24.00 | 26.00 | 26.00 | 21.00 |
| SHFM1 | 1042.00 | 1145.00 | 1149.00 | 1112.00 | 1471.00 |
| SHISA2 | 124.00 | 134.00 | 178.00 | 145.33 | 279.00 |
| SHISA3 | 169.00 | 173.00 | 174.00 | 172.00 | 201.00 |
| SHISA4 | 102.00 | 103.00 | 112.00 | 105.67 | 142.00 |
| SHISA5 | 3001.00 | 2997.00 | 2839.00 | 2945.67 | 3580.00 |
| SHKBP1 | 1241.00 | 1206.00 | 1326.00 | 1257.67 | 1436.00 |
| SHMT1 | 332.00 | 345.00 | 340.00 | 339.00 | 484.00 |
| SHMT2 | 10573.00 | 10092.00 | 10698.00 | 10454.33 | 11866.00 |
| SHOC2 | 655.00 | 603.00 | 774.00 | 677.33 | 742.00 |
| SHOX2 | 549.00 | 599.00 | 582.00 | 576.67 | 636.00 |
| SHPK | 534.00 | 471.00 | 551.00 | 518.67 | 389.00 |
| SHPRH | 624.00 | 631.00 | 702.00 | 652.33 | 590.00 |
| SHQ1 | 462.00 | 432.00 | 504.00 | 466.00 | 591.00 |
| SHROOM1 | 407.00 | 373.00 | 408.00 | 396.00 | 397.00 |
| SHROOM2 | 713.00 | 631.00 | 619.00 | 654.33 | 714.00 |
| SHROOM3 | 16.00 | 32.00 | 23.00 | 23.67 | 45.00 |
| SHROOM4 | 23.00 | 31.00 | 27.00 | 27.00 | 22.00 |
| SHTN1 | 1478.00 | 1434.00 | 1605.00 | 1505.67 | 1614.00 |
| SIAE | 911.00 | 830.00 | 976.00 | 905.67 | 695.00 |

| | | | | | |
|---|---|---|---|---|---|
| SIAH1 | 456.00 | 411.00 | 464.00 | 443.67 | 563.00 |
| SIAH2 | 637.00 | 590.00 | 625.00 | 617.33 | 791.00 |
| SIDT2 | 449.00 | 484.00 | 470.00 | 467.67 | 327.00 |
| SIGIRR | 320.00 | 303.00 | 284.00 | 302.33 | 505.00 |
| SIGLEC15 | 414.00 | 428.00 | 446.00 | 429.33 | 216.00 |
| SIGLEC9 | 9.00 | 10.00 | 7.00 | 8.67 | 3.00 |
| SIGMAR1 | 4642.00 | 4584.00 | 4305.00 | 4510.33 | 5095.00 |
| SIK1 | 81.00 | 64.00 | 64.00 | 69.67 | 70.00 |
| SIK2 | 504.00 | 459.00 | 526.00 | 496.33 | 412.00 |
| SIK3 | 669.00 | 629.00 | 745.00 | 681.00 | 626.00 |
| SIKE1 | 1498.00 | 1393.00 | 1669.00 | 1520.00 | 1471.00 |
| SIL1 | 1783.00 | 1681.00 | 1582.00 | 1682.00 | 1831.00 |
| SIM1 | 12.00 | 7.00 | 16.00 | 11.67 | 11.00 |
| SIM2 | 82.00 | 74.00 | 71.00 | 75.67 | 80.00 |
| SIMC1 | 317.00 | 301.00 | 303.00 | 307.00 | 343.00 |
| SIN3A | 1462.00 | 1435.00 | 1439.00 | 1445.33 | 1361.00 |
| SIN3B | 2291.00 | 2309.00 | 2519.00 | 2373.00 | 2002.00 |
| SIPA1 | 908.00 | 882.00 | 865.00 | 885.00 | 835.00 |
| SIPA1L1 | 2235.00 | 1956.00 | 2249.00 | 2146.67 | 1516.00 |
| SIPA1L2 | 53.00 | 65.00 | 67.00 | 61.67 | 47.00 |
| SIPA1L3 | 2891.00 | 2894.00 | 3246.00 | 3010.33 | 2612.00 |
| SIRPA | 695.00 | 723.00 | 733.00 | 717.00 | 615.00 |
| SIRPB1 | 228.00 | 253.00 | 253.00 | 244.67 | 123.00 |
| SIRPD | 3.00 | 2.00 | 0.00 | 1.67 | 2.00 |
| SIRT1 | 581.00 | 613.00 | 751.00 | 648.33 | 765.00 |
| SIRT2 | 664.00 | 625.00 | 642.00 | 643.67 | 695.00 |
| SIRT3 | 400.00 | 426.00 | 448.00 | 424.67 | 416.00 |
| SIRT4 | 8.00 | 5.00 | 9.00 | 7.33 | 9.00 |
| SIRT5 | 685.00 | 586.00 | 668.00 | 646.33 | 670.00 |
| SIRT6 | 819.00 | 638.00 | 713.00 | 723.33 | 759.00 |
| SIRT7 | 805.00 | 879.00 | 958.00 | 880.67 | 957.00 |
| SIVA1 | 608.00 | 595.00 | 560.00 | 587.67 | 687.00 |
| SIX1 | 449.00 | 397.00 | 406.00 | 417.33 | 436.00 |
| SIX4 | 716.00 | 560.00 | 832.00 | 702.67 | 631.00 |
| SIX5 | 257.00 | 303.00 | 310.00 | 290.00 | 333.00 |
| SKA2 | 513.00 | 523.00 | 541.00 | 525.67 | 825.00 |
| SKA3 | 400.00 | 380.00 | 484.00 | 421.33 | 539.00 |
| SKAP1 | 10.00 | 20.00 | 9.00 | 13.00 | 12.00 |
| SKAP2 | 1277.00 | 1157.00 | 1354.00 | 1262.67 | 1008.00 |
| SKI | 1843.00 | 1799.00 | 1859.00 | 1833.67 | 1983.00 |
| SKIDA1 | 64.00 | 71.00 | 58.00 | 64.33 | 59.00 |
| SKIL | 827.00 | 744.00 | 949.00 | 840.00 | 832.00 |
| SKIV2L | 5.00 | 2.00 | 4.00 | 3.67 | 6.00 |
| SKIV2L2 | 2611.00 | 2303.00 | 2728.00 | 2547.33 | 2873.00 |
| SKOR1 | 11.00 | 20.00 | 16.00 | 15.67 | 13.00 |
| SKP1 | 2362.00 | 2235.00 | 2478.00 | 2358.33 | 2979.00 |
| SKP1P2 | 22.00 | 25.00 | 21.00 | 22.67 | 20.00 |
| SKP2 | 652.00 | 604.00 | 743.00 | 666.33 | 854.00 |

| | | | | | |
|---|---|---|---|---|---|
| SLA | 5.00 | 2.00 | 5.00 | 4.00 | 2.00 |
| SLAIN1 | 463.00 | 391.00 | 489.00 | 447.67 | 522.00 |
| SLAIN2 | 924.00 | 850.00 | 990.00 | 921.33 | 946.00 |
| SLBP | 4056.00 | 3774.00 | 4140.00 | 3990.00 | 5071.00 |
| SLC10A3 | 523.00 | 573.00 | 566.00 | 554.00 | 539.00 |
| SLC10A5 | 7.00 | 5.00 | 2.00 | 4.67 | 9.00 |
| SLC10A7 | 260.00 | 219.00 | 287.00 | 255.33 | 252.00 |
| SLC11A2 | 1761.00 | 1668.00 | 1844.00 | 1757.67 | 1621.00 |
| SLC12A2 | 690.00 | 597.00 | 675.00 | 654.00 | 661.00 |
| SLC12A4 | 1169.00 | 1186.00 | 1220.00 | 1191.67 | 1233.00 |
| SLC12A6 | 2078.00 | 1912.00 | 2160.00 | 2050.00 | 1968.00 |
| SLC12A8 | 1368.00 | 1306.00 | 1396.00 | 1356.67 | 1526.00 |
| SLC12A9 | 1854.00 | 1887.00 | 1928.00 | 1889.67 | 2010.00 |
| SLC13A3 | 921.00 | 792.00 | 898.00 | 870.33 | 708.00 |
| SLC14A1 | 4.00 | 8.00 | 4.00 | 5.33 | 1.00 |
| SLC15A2 | 27.00 | 30.00 | 34.00 | 30.33 | 35.00 |
| SLC15A4 | 558.00 | 471.00 | 542.00 | 523.67 | 591.00 |
| SLC16A1 | 4860.00 | 4554.00 | 5120.00 | 4844.67 | 4670.00 |
| SLC16A10 | 42.00 | 56.00 | 71.00 | 56.33 | 86.00 |
| SLC16A12 | 89.00 | 59.00 | 64.00 | 70.67 | 36.00 |
| SLC16A13 | 65.00 | 67.00 | 76.00 | 69.33 | 53.00 |
| SLC16A14 | 66.00 | 61.00 | 61.00 | 62.67 | 50.00 |
| SLC16A1-AS1 | 206.00 | 193.00 | 201.00 | 200.00 | 184.00 |
| SLC16A2 | 526.00 | 496.00 | 483.00 | 501.67 | 597.00 |
| SLC16A3 | 535.00 | 535.00 | 581.00 | 550.33 | 527.00 |
| SLC16A4 | 21.00 | 34.00 | 27.00 | 27.33 | 28.00 |
| SLC16A5 | 108.00 | 95.00 | 124.00 | 109.00 | 88.00 |
| SLC16A6 | 192.00 | 184.00 | 230.00 | 202.00 | 231.00 |
| SLC16A7 | 594.00 | 458.00 | 601.00 | 551.00 | 784.00 |
| SLC16A9 | 1955.00 | 1815.00 | 1945.00 | 1905.00 | 1604.00 |
| SLC17A5 | 569.00 | 530.00 | 579.00 | 559.33 | 289.00 |
| SLC17A7 | 3.00 | 6.00 | 3.00 | 4.00 | 8.00 |
| SLC18B1 | 749.00 | 769.00 | 714.00 | 744.00 | 717.00 |
| SLC19A1 | 1164.00 | 1231.00 | 1245.00 | 1213.33 | 1244.00 |
| SLC19A2 | 919.00 | 842.00 | 1030.00 | 930.33 | 902.00 |
| SLC19A3 | 260.00 | 222.00 | 280.00 | 254.00 | 352.00 |
| SLC1A1 | 232.00 | 245.00 | 267.00 | 248.00 | 314.00 |
| SLC1A2 | 60.00 | 61.00 | 56.00 | 59.00 | 28.00 |
| SLC1A3 | 1247.00 | 1266.00 | 1448.00 | 1320.33 | 1569.00 |
| SLC1A4 | 2649.00 | 2559.00 | 2682.00 | 2630.00 | 2627.00 |
| SLC1A5 | 16906.00 | 17051.00 | 17914.00 | 17290.33 | 16333.00 |
| SLC20A1 | 9019.00 | 8499.00 | 8821.00 | 8779.67 | 11988.00 |
| SLC20A2 | 1074.00 | 1005.00 | 1106.00 | 1061.67 | 931.00 |
| SLC22A1 | 32.00 | 27.00 | 37.00 | 32.00 | 23.00 |
| SLC22A13 | 16.00 | 9.00 | 9.00 | 11.33 | 2.00 |
| SLC22A14 | 5.00 | 9.00 | 3.00 | 5.67 | 1.00 |
| SLC22A15 | 548.00 | 495.00 | 537.00 | 526.67 | 392.00 |
| SLC22A17 | 3.00 | 2.00 | 2.00 | 2.33 | 3.00 |

| | | | | | |
|---|---|---|---|---|---|
| SLC22A18 | 57.00 | 39.00 | 37.00 | 44.33 | 61.00 |
| SLC22A18AS | 5.00 | 4.00 | 6.00 | 5.00 | 0.00 |
| SLC22A20 | 14.00 | 15.00 | 10.00 | 13.00 | 2.00 |
| SLC22A3 | 698.00 | 630.00 | 749.00 | 692.33 | 512.00 |
| SLC22A4 | 95.00 | 98.00 | 99.00 | 97.33 | 101.00 |
| SLC22A5 | 220.00 | 251.00 | 237.00 | 236.00 | 160.00 |
| SLC23A2 | 2635.00 | 2469.00 | 2713.00 | 2605.67 | 2265.00 |
| SLC23A3 | 160.00 | 185.00 | 213.00 | 186.00 | 115.00 |
| SLC24A1 | 727.00 | 666.00 | 675.00 | 689.33 | 568.00 |
| SLC24A2 | 3.00 | 6.00 | 2.00 | 3.67 | 0.00 |
| SLC25A1 | 402.00 | 401.00 | 458.00 | 420.33 | 606.00 |
| SLC25A10 | 410.00 | 358.00 | 340.00 | 369.33 | 475.00 |
| SLC25A11 | 1089.00 | 1053.00 | 1062.00 | 1068.00 | 1373.00 |
| SLC25A12 | 950.00 | 881.00 | 991.00 | 940.67 | 1034.00 |
| SLC25A13 | 857.00 | 751.00 | 886.00 | 831.33 | 1170.00 |
| SLC25A14 | 164.00 | 218.00 | 209.00 | 197.00 | 199.00 |
| SLC25A15 | 468.00 | 434.00 | 461.00 | 454.33 | 609.00 |
| SLC25A16 | 451.00 | 455.00 | 490.00 | 465.33 | 554.00 |
| SLC25A17 | 453.00 | 382.00 | 460.00 | 431.67 | 598.00 |
| SLC25A19 | 327.00 | 330.00 | 338.00 | 331.67 | 395.00 |
| SLC25A20 | 132.00 | 122.00 | 105.00 | 119.67 | 160.00 |
| SLC25A21 | 61.00 | 55.00 | 62.00 | 59.33 | 112.00 |
| SLC25A21-AS1 | 92.00 | 83.00 | 107.00 | 94.00 | 94.00 |
| SLC25A22 | 815.00 | 868.00 | 1005.00 | 896.00 | 1032.00 |
| SLC25A23 | 190.00 | 172.00 | 178.00 | 180.00 | 181.00 |
| SLC25A24 | 1438.00 | 1253.00 | 1451.00 | 1380.67 | 1668.00 |
| SLC25A25 | 458.00 | 503.00 | 444.00 | 468.33 | 510.00 |
| SLC25A25-AS1 | 473.00 | 535.00 | 659.00 | 555.67 | 423.00 |
| SLC25A26 | 278.00 | 252.00 | 295.00 | 275.00 | 344.00 |
| SLC25A27 | 939.00 | 922.00 | 1076.00 | 979.00 | 771.00 |
| SLC25A28 | 643.00 | 801.00 | 757.00 | 733.67 | 780.00 |
| SLC25A29 | 1114.00 | 1126.00 | 1276.00 | 1172.00 | 1151.00 |
| SLC25A3 | 8427.00 | 8420.00 | 9468.00 | 8771.67 | 11197.00 |
| SLC25A30 | 322.00 | 290.00 | 366.00 | 326.00 | 308.00 |
| SLC25A32 | 1326.00 | 1201.00 | 1402.00 | 1309.67 | 1617.00 |
| SLC25A33 | 328.00 | 336.00 | 335.00 | 333.00 | 442.00 |
| SLC25A34 | 56.00 | 66.00 | 55.00 | 59.00 | 55.00 |
| SLC25A35 | 255.00 | 265.00 | 256.00 | 258.67 | 311.00 |
| SLC25A36 | 2220.00 | 2096.00 | 2542.00 | 2286.00 | 2526.00 |
| SLC25A37 | 1343.00 | 1340.00 | 1490.00 | 1391.00 | 1088.00 |
| SLC25A38 | 1016.00 | 969.00 | 1121.00 | 1035.33 | 1141.00 |
| SLC25A39 | 2481.00 | 2522.00 | 2651.00 | 2551.33 | 2721.00 |
| SLC25A4 | 295.00 | 237.00 | 301.00 | 277.67 | 375.00 |
| SLC25A40 | 274.00 | 236.00 | 269.00 | 259.67 | 374.00 |
| SLC25A42 | 105.00 | 144.00 | 106.00 | 118.33 | 114.00 |
| SLC25A43 | 328.00 | 303.00 | 318.00 | 316.33 | 361.00 |
| SLC25A44 | 763.00 | 765.00 | 750.00 | 759.33 | 866.00 |
| SLC25A45 | 73.00 | 79.00 | 87.00 | 79.67 | 229.00 |

| | | | | | |
|---|---|---|---|---|---|
| SLC25A46 | 1227.00 | 1018.00 | 1260.00 | 1168.33 | 1265.00 |
| SLC25A5 | 3739.00 | 3523.00 | 4018.00 | 3760.00 | 4856.00 |
| SLC25A51 | 254.00 | 221.00 | 229.00 | 234.67 | 273.00 |
| SLC25A53 | 110.00 | 116.00 | 110.00 | 112.00 | 108.00 |
| SLC25A5-AS1 | 28.00 | 28.00 | 21.00 | 25.67 | 32.00 |
| SLC25A6 | 18.00 | 19.00 | 15.00 | 17.33 | 30.00 |
| SLC26A1 | 6.00 | 4.00 | 4.00 | 4.67 | 4.00 |
| SLC26A10 | 178.00 | 165.00 | 170.00 | 171.00 | 162.00 |
| SLC26A11 | 461.00 | 442.00 | 457.00 | 453.33 | 487.00 |
| SLC26A2 | 1106.00 | 1026.00 | 1174.00 | 1102.00 | 1063.00 |
| SLC26A4 | 14.00 | 11.00 | 7.00 | 10.67 | 9.00 |
| SLC26A6 | 1062.00 | 1112.00 | 1197.00 | 1123.67 | 810.00 |
| SLC26A7 | 16.00 | 16.00 | 25.00 | 19.00 | 16.00 |
| SLC27A1 | 328.00 | 339.00 | 379.00 | 348.67 | 291.00 |
| SLC27A4 | 1655.00 | 1699.00 | 1567.00 | 1640.33 | 1708.00 |
| SLC27A5 | 55.00 | 63.00 | 60.00 | 59.33 | 69.00 |
| SLC27A6 | 56.00 | 65.00 | 55.00 | 58.67 | 60.00 |
| SLC29A1 | 2522.00 | 2628.00 | 2471.00 | 2540.33 | 3184.00 |
| SLC29A2 | 68.00 | 59.00 | 99.00 | 75.33 | 93.00 |
| SLC29A3 | 9.00 | 5.00 | 16.00 | 10.00 | 12.00 |
| SLC29A4 | 23.00 | 21.00 | 26.00 | 23.33 | 26.00 |
| SLC2A1 | 2621.00 | 2552.00 | 2619.00 | 2597.33 | 2369.00 |
| SLC2A10 | 89.00 | 111.00 | 104.00 | 101.33 | 69.00 |
| SLC2A11 | 5.00 | 4.00 | 0.00 | 3.00 | 2.00 |
| SLC2A12 | 37.00 | 38.00 | 47.00 | 40.67 | 50.00 |
| SLC2A13 | 807.00 | 743.00 | 898.00 | 816.00 | 623.00 |
| SLC2A1-AS1 | 12.00 | 13.00 | 18.00 | 14.33 | 10.00 |
| SLC2A3 | 803.00 | 881.00 | 881.00 | 855.00 | 873.00 |
| SLC2A4 | 5.00 | 3.00 | 3.00 | 3.67 | 1.00 |
| SLC2A4RG | 1305.00 | 1306.00 | 1374.00 | 1328.33 | 946.00 |
| SLC2A6 | 253.00 | 222.00 | 203.00 | 226.00 | 206.00 |
| SLC2A8 | 409.00 | 457.00 | 414.00 | 426.67 | 366.00 |
| SLC30A1 | 487.00 | 414.00 | 516.00 | 472.33 | 422.00 |
| SLC30A4 | 71.00 | 44.00 | 70.00 | 61.67 | 40.00 |
| SLC30A5 | 1865.00 | 1783.00 | 1851.00 | 1833.00 | 2023.00 |
| SLC30A6 | 1171.00 | 1031.00 | 1195.00 | 1132.33 | 1160.00 |
| SLC30A7 | 2355.00 | 2167.00 | 2428.00 | 2316.67 | 2200.00 |
| SLC30A9 | 2399.00 | 2053.00 | 2457.00 | 2303.00 | 2697.00 |
| SLC31A1 | 1552.00 | 1490.00 | 1424.00 | 1488.67 | 1319.00 |
| SLC31A2 | 168.00 | 168.00 | 150.00 | 162.00 | 178.00 |
| SLC33A1 | 1801.00 | 1641.00 | 1908.00 | 1783.33 | 1614.00 |
| SLC35A1 | 834.00 | 835.00 | 764.00 | 811.00 | 895.00 |
| SLC35A2 | 813.00 | 848.00 | 706.00 | 789.00 | 891.00 |
| SLC35A3 | 822.00 | 760.00 | 781.00 | 787.67 | 817.00 |
| SLC35A4 | 1198.00 | 1074.00 | 1128.00 | 1133.33 | 1171.00 |
| SLC35A5 | 1524.00 | 1349.00 | 1383.00 | 1418.67 | 1586.00 |
| SLC35B1 | 901.00 | 886.00 | 914.00 | 900.33 | 1029.00 |
| SLC35B2 | 2576.00 | 2573.00 | 2443.00 | 2530.67 | 2669.00 |

| | | | | | |
|---|---|---|---|---|---|
| SLC35B3 | 775.00 | 764.00 | 817.00 | 785.33 | 877.00 |
| SLC35B4 | 1324.00 | 1244.00 | 1291.00 | 1286.33 | 1200.00 |
| SLC35C1 | 727.00 | 829.00 | 703.00 | 753.00 | 726.00 |
| SLC35C2 | 2609.00 | 2659.00 | 2658.00 | 2642.00 | 2576.00 |
| SLC35D1 | 696.00 | 660.00 | 717.00 | 691.00 | 728.00 |
| SLC35D2 | 714.00 | 676.00 | 746.00 | 712.00 | 871.00 |
| SLC35D3 | 3.00 | 2.00 | 4.00 | 3.00 | 1.00 |
| SLC35E1 | 1277.00 | 1258.00 | 1271.00 | 1268.67 | 1383.00 |
| SLC35E2 | 559.00 | 540.00 | 580.00 | 559.67 | 593.00 |
| SLC35E2B | 1719.00 | 1656.00 | 1826.00 | 1733.67 | 1617.00 |
| SLC35E3 | 272.00 | 263.00 | 254.00 | 263.00 | 265.00 |
| SLC35E4 | 536.00 | 595.00 | 570.00 | 567.00 | 446.00 |
| SLC35F1 | 7.00 | 6.00 | 2.00 | 5.00 | 11.00 |
| SLC35F2 | 1656.00 | 1698.00 | 1621.00 | 1658.33 | 1759.00 |
| SLC35F3 | 8.00 | 9.00 | 11.00 | 9.33 | 8.00 |
| SLC35F5 | 1251.00 | 1085.00 | 1130.00 | 1155.33 | 1316.00 |
| SLC35F6 | 1164.00 | 1112.00 | 1059.00 | 1111.67 | 1091.00 |
| SLC35G1 | 229.00 | 251.00 | 253.00 | 244.33 | 288.00 |
| SLC35G2 | 1111.00 | 1146.00 | 1141.00 | 1132.67 | 1419.00 |
| SLC36A1 | 866.00 | 808.00 | 782.00 | 818.67 | 752.00 |
| SLC36A4 | 358.00 | 348.00 | 397.00 | 367.67 | 389.00 |
| SLC37A2 | 877.00 | 824.00 | 883.00 | 861.33 | 470.00 |
| SLC37A3 | 1729.00 | 1494.00 | 1695.00 | 1639.33 | 1844.00 |
| SLC37A4 | 679.00 | 724.00 | 686.00 | 696.33 | 803.00 |
| SLC38A1 | 20450.00 | 18781.00 | 21173.00 | 20134.67 | 16952.00 |
| SLC38A10 | 5066.00 | 5240.00 | 5455.00 | 5253.67 | 4797.00 |
| SLC38A2 | 25457.00 | 23590.00 | 28223.00 | 25756.67 | 24361.00 |
| SLC38A4 | 52.00 | 59.00 | 61.00 | 57.33 | 80.00 |
| SLC38A5 | 12.00 | 26.00 | 20.00 | 19.33 | 10.00 |
| SLC38A6 | 1237.00 | 1232.00 | 1258.00 | 1242.33 | 1373.00 |
| SLC38A7 | 1394.00 | 1307.00 | 1376.00 | 1359.00 | 1178.00 |
| SLC38A9 | 751.00 | 685.00 | 722.00 | 719.33 | 660.00 |
| SLC39A1 | 3329.00 | 3423.00 | 3348.00 | 3366.67 | 3441.00 |
| SLC39A10 | 1505.00 | 1301.00 | 1646.00 | 1484.00 | 1631.00 |
| SLC39A11 | 544.00 | 539.00 | 531.00 | 538.00 | 589.00 |
| SLC39A13 | 3302.00 | 3588.00 | 3630.00 | 3506.67 | 3159.00 |
| SLC39A14 | 6077.00 | 5540.00 | 5890.00 | 5835.67 | 5026.00 |
| SLC39A3 | 723.00 | 770.00 | 732.00 | 741.67 | 830.00 |
| SLC39A4 | 212.00 | 205.00 | 166.00 | 194.33 | 303.00 |
| SLC39A5 | 5.00 | 6.00 | 5.00 | 5.33 | 6.00 |
| SLC39A6 | 2233.00 | 2111.00 | 2290.00 | 2211.33 | 2133.00 |
| SLC39A8 | 410.00 | 389.00 | 413.00 | 404.00 | 427.00 |
| SLC39A9 | 1622.00 | 1521.00 | 1643.00 | 1595.33 | 1644.00 |
| SLC3A2 | 15204.00 | 16012.00 | 15747.00 | 15654.33 | 15408.00 |
| SLC41A1 | 2162.00 | 2145.00 | 2183.00 | 2163.33 | 1627.00 |
| SLC41A2 | 187.00 | 145.00 | 200.00 | 177.33 | 138.00 |
| SLC41A3 | 1100.00 | 1089.00 | 1076.00 | 1088.33 | 1293.00 |
| SLC43A1 | 1198.00 | 1241.00 | 1257.00 | 1232.00 | 1433.00 |

| | | | | | |
|---|---|---|---|---|---|
| SLC43A2 | 3.00 | 1.00 | 1.00 | 1.67 | 0.00 |
| SLC43A3 | 7706.00 | 7548.00 | 7716.00 | 7656.67 | 7408.00 |
| SLC44A1 | 1027.00 | 926.00 | 1018.00 | 990.33 | 1048.00 |
| SLC44A2 | 1108.00 | 1122.00 | 1145.00 | 1125.00 | 795.00 |
| SLC44A3 | 15.00 | 7.00 | 6.00 | 9.33 | 11.00 |
| SLC44A5 | 769.00 | 615.00 | 670.00 | 684.67 | 631.00 |
| SLC45A1 | 78.00 | 99.00 | 100.00 | 92.33 | 109.00 |
| SLC45A3 | 701.00 | 658.00 | 778.00 | 712.33 | 579.00 |
| SLC45A4 | 785.00 | 802.00 | 827.00 | 804.67 | 931.00 |
| SLC46A1 | 197.00 | 190.00 | 211.00 | 199.33 | 178.00 |
| SLC46A3 | 206.00 | 178.00 | 163.00 | 182.33 | 119.00 |
| SLC48A1 | 340.00 | 390.00 | 398.00 | 376.00 | 398.00 |
| SLC4A11 | 26.00 | 24.00 | 25.00 | 25.00 | 60.00 |
| SLC4A1AP | 840.00 | 778.00 | 921.00 | 846.33 | 899.00 |
| SLC4A2 | 7129.00 | 7234.00 | 7225.00 | 7196.00 | 7374.00 |
| SLC4A3 | 158.00 | 157.00 | 161.00 | 158.67 | 133.00 |
| SLC4A4 | 51.00 | 37.00 | 54.00 | 47.33 | 56.00 |
| SLC4A5 | 7.00 | 11.00 | 11.00 | 9.67 | 3.00 |
| SLC4A7 | 3432.00 | 2887.00 | 3959.00 | 3426.00 | 2777.00 |
| SLC4A8 | 556.00 | 535.00 | 611.00 | 567.33 | 564.00 |
| SLC4A9 | 14.00 | 26.00 | 24.00 | 21.33 | 20.00 |
| SLC50A1 | 1123.00 | 1247.00 | 1120.00 | 1163.33 | 1216.00 |
| SLC52A2 | 3506.00 | 3467.00 | 3520.00 | 3497.67 | 3669.00 |
| SLC5A3 | 1303.00 | 1194.00 | 1273.00 | 1256.67 | 1229.00 |
| SLC5A6 | 2686.00 | 2704.00 | 2898.00 | 2762.67 | 2624.00 |
| SLC6A13 | 4.00 | 7.00 | 5.00 | 5.33 | 9.00 |
| SLC6A15 | 4647.00 | 4411.00 | 5052.00 | 4703.33 | 4550.00 |
| SLC6A17 | 265.00 | 287.00 | 292.00 | 281.33 | 212.00 |
| SLC6A6 | 1231.00 | 1260.00 | 1180.00 | 1223.67 | 935.00 |
| SLC6A8 | 1468.00 | 1538.00 | 1666.00 | 1557.33 | 1698.00 |
| SLC6A9 | 544.00 | 587.00 | 568.00 | 566.33 | 511.00 |
| SLC7A1 | 9818.00 | 9371.00 | 10103.00 | 9764.00 | 8702.00 |
| SLC7A11 | 5136.00 | 4513.00 | 5022.00 | 4890.33 | 2991.00 |
| SLC7A11-AS1 | 241.00 | 237.00 | 266.00 | 248.00 | 293.00 |
| SLC7A2 | 1287.00 | 1183.00 | 1441.00 | 1303.67 | 883.00 |
| SLC7A3 | 4.00 | 2.00 | 4.00 | 3.33 | 1.00 |
| SLC7A5 | 25633.00 | 25253.00 | 24306.00 | 25064.00 | 24375.00 |
| SLC7A5P1 | 45.00 | 39.00 | 34.00 | 39.33 | 33.00 |
| SLC7A5P2 | 72.00 | 56.00 | 62.00 | 63.33 | 47.00 |
| SLC7A6 | 2410.00 | 2458.00 | 2624.00 | 2497.33 | 2130.00 |
| SLC7A6OS | 318.00 | 318.00 | 396.00 | 344.00 | 338.00 |
| SLC7A7 | 30.00 | 29.00 | 40.00 | 33.00 | 25.00 |
| SLC7A8 | 60.00 | 67.00 | 75.00 | 67.33 | 32.00 |
| SLC7A9 | 6.00 | 2.00 | 4.00 | 4.00 | 4.00 |
| SLC8A1 | 1504.00 | 1396.00 | 1536.00 | 1478.67 | 1409.00 |
| SLC8A1-AS1 | 6.00 | 8.00 | 14.00 | 9.33 | 3.00 |
| SLC8A2 | 3.00 | 3.00 | 3.00 | 3.00 | 0.00 |
| SLC8B1 | 344.00 | 361.00 | 330.00 | 345.00 | 264.00 |

| | | | | | |
|---|---|---|---|---|---|
| SLC9A1 | 2044.00 | 2003.00 | 2049.00 | 2032.00 | 1607.00 |
| SLC9A2 | 18.00 | 17.00 | 21.00 | 18.67 | 23.00 |
| SLC9A3R1 | 375.00 | 371.00 | 422.00 | 389.33 | 413.00 |
| SLC9A3R2 | 2098.00 | 2054.00 | 2107.00 | 2086.33 | 1115.00 |
| SLC9A5 | 710.00 | 704.00 | 729.00 | 714.33 | 659.00 |
| SLC9A6 | 714.00 | 671.00 | 680.00 | 688.33 | 588.00 |
| SLC9A7 | 3828.00 | 3795.00 | 3956.00 | 3859.67 | 3854.00 |
| SLC9A8 | 1324.00 | 1313.00 | 1329.00 | 1322.00 | 1113.00 |
| SLC9A9 | 38.00 | 49.00 | 50.00 | 45.67 | 53.00 |
| SLC9B2 | 467.00 | 517.00 | 536.00 | 506.67 | 523.00 |
| SLCO2A1 | 3.00 | 4.00 | 2.00 | 3.00 | 2.00 |
| SLCO3A1 | 59.00 | 82.00 | 66.00 | 69.00 | 55.00 |
| SLCO4A1 | 1604.00 | 1773.00 | 1642.00 | 1673.00 | 1679.00 |
| SLF1 | 259.00 | 249.00 | 255.00 | 254.33 | 375.00 |
| SLF2 | 1631.00 | 1460.00 | 1654.00 | 1581.67 | 1622.00 |
| SLFN11 | 828.00 | 789.00 | 892.00 | 836.33 | 1111.00 |
| SLFN12 | 402.00 | 389.00 | 378.00 | 389.67 | 364.00 |
| SLFN12L | 8.00 | 3.00 | 10.00 | 7.00 | 5.00 |
| SLFN13 | 6.00 | 13.00 | 6.00 | 8.33 | 7.00 |
| SLFN5 | 3407.00 | 2991.00 | 3529.00 | 3309.00 | 2500.00 |
| SLFNL1 | 8.00 | 8.00 | 9.00 | 8.33 | 12.00 |
| SLFNL1-AS1 | 177.00 | 183.00 | 156.00 | 172.00 | 151.00 |
| SLIRP | 745.00 | 744.00 | 809.00 | 766.00 | 1041.00 |
| SLIT2 | 889.00 | 844.00 | 914.00 | 882.33 | 1204.00 |
| SLIT3 | 2102.00 | 2107.00 | 2156.00 | 2121.67 | 1506.00 |
| SLITRK3 | 6.00 | 7.00 | 2.00 | 5.00 | 5.00 |
| SLITRK5 | 676.00 | 690.00 | 746.00 | 704.00 | 824.00 |
| SLK | 1774.00 | 1519.00 | 1929.00 | 1740.67 | 1673.00 |
| SLMAP | 1077.00 | 975.00 | 1163.00 | 1071.67 | 1101.00 |
| SLMO1 | 350.00 | 322.00 | 362.00 | 344.67 | 376.00 |
| SLMO2 | 965.00 | 837.00 | 1084.00 | 962.00 | 1102.00 |
| SLTM | 3630.00 | 3515.00 | 3817.00 | 3654.00 | 3838.00 |
| SLU7 | 1171.00 | 1094.00 | 1310.00 | 1191.67 | 1237.00 |
| SLX4 | 641.00 | 636.00 | 636.00 | 637.67 | 627.00 |
| SLX4IP | 121.00 | 77.00 | 78.00 | 92.00 | 71.00 |
| SMA4 | 27.00 | 15.00 | 17.00 | 19.67 | 14.00 |
| SMAD1 | 217.00 | 198.00 | 233.00 | 216.00 | 255.00 |
| SMAD2 | 980.00 | 870.00 | 1109.00 | 986.33 | 1228.00 |
| SMAD3 | 7307.00 | 7016.00 | 7721.00 | 7348.00 | 4253.00 |
| SMAD4 | 1111.00 | 1138.00 | 1336.00 | 1195.00 | 1169.00 |
| SMAD5 | 1951.00 | 1757.00 | 2204.00 | 1970.67 | 1854.00 |
| SMAD6 | 14.00 | 12.00 | 26.00 | 17.33 | 24.00 |
| SMAD7 | 142.00 | 158.00 | 176.00 | 158.67 | 101.00 |
| SMAD9 | 106.00 | 116.00 | 135.00 | 119.00 | 137.00 |
| SMAGP | 1105.00 | 1108.00 | 1130.00 | 1114.33 | 1361.00 |
| SMAP1 | 690.00 | 639.00 | 758.00 | 695.67 | 851.00 |
| SMAP2 | 1417.00 | 1293.00 | 1359.00 | 1356.33 | 1904.00 |
| SMARCA1 | 845.00 | 809.00 | 905.00 | 853.00 | 1079.00 |

| | | | | | |
|---|---|---|---|---|---|
| SMARCA2 | 965.00 | 916.00 | 885.00 | 922.00 | 957.00 |
| SMARCA4 | 4221.00 | 4263.00 | 4320.00 | 4268.00 | 4407.00 |
| SMARCA5 | 1989.00 | 1832.00 | 2129.00 | 1983.33 | 2219.00 |
| SMARCAD1 | 1459.00 | 1478.00 | 1681.00 | 1539.33 | 1605.00 |
| SMARCAL1 | 823.00 | 727.00 | 835.00 | 795.00 | 882.00 |
| SMARCC1 | 3136.00 | 2785.00 | 3239.00 | 3053.33 | 2734.00 |
| SMARCC2 | 2868.00 | 2789.00 | 2954.00 | 2870.33 | 2592.00 |
| SMARCD1 | 3114.00 | 3041.00 | 3252.00 | 3135.67 | 3802.00 |
| SMARCD2 | 2477.00 | 2627.00 | 2742.00 | 2615.33 | 2983.00 |
| SMARCD3 | 169.00 | 207.00 | 198.00 | 191.33 | 151.00 |
| SMARCE1 | 1934.00 | 1724.00 | 2010.00 | 1889.33 | 2396.00 |
| SMC1A | 4478.00 | 4193.00 | 4403.00 | 4358.00 | 4385.00 |
| SMC2 | 1106.00 | 1040.00 | 1075.00 | 1073.67 | 1268.00 |
| SMC3 | 2713.00 | 2475.00 | 2753.00 | 2647.00 | 3032.00 |
| SMC4 | 4807.00 | 4413.00 | 4775.00 | 4665.00 | 6040.00 |
| SMC5 | 1096.00 | 939.00 | 1018.00 | 1017.67 | 989.00 |
| SMC5-AS1 | 13.00 | 7.00 | 6.00 | 8.67 | 11.00 |
| SMC6 | 1735.00 | 1586.00 | 1849.00 | 1723.33 | 1832.00 |
| SMCHD1 | 3010.00 | 2714.00 | 3122.00 | 2948.67 | 2878.00 |
| SMCO2 | 58.00 | 50.00 | 36.00 | 48.00 | 71.00 |
| SMCO4 | 81.00 | 72.00 | 67.00 | 73.33 | 135.00 |
| SMCR8 | 1921.00 | 1671.00 | 2022.00 | 1871.33 | 1452.00 |
| SMG1 | 4039.00 | 3575.00 | 4424.00 | 4012.67 | 3679.00 |
| SMG1P1 | 87.00 | 57.00 | 91.00 | 78.33 | 44.00 |
| SMG1P2 | 33.00 | 28.00 | 31.00 | 30.67 | 25.00 |
| SMG1P3 | 74.00 | 66.00 | 56.00 | 65.33 | 28.00 |
| SMG1P5 | 13.00 | 11.00 | 16.00 | 13.33 | 8.00 |
| SMG1P7 | 176.00 | 146.00 | 186.00 | 169.33 | 222.00 |
| SMG5 | 4068.00 | 4091.00 | 4472.00 | 4210.33 | 4025.00 |
| SMG6 | 1829.00 | 1760.00 | 1873.00 | 1820.67 | 2322.00 |
| SMG7 | 2590.00 | 2385.00 | 2680.00 | 2551.67 | 2428.00 |
| SMG7-AS1 | 5.00 | 4.00 | 5.00 | 4.67 | 3.00 |
| SMG8 | 1051.00 | 1041.00 | 1070.00 | 1054.00 | 1113.00 |
| SMG9 | 1561.00 | 1526.00 | 1561.00 | 1549.33 | 1958.00 |
| SMIM10 | 108.00 | 83.00 | 99.00 | 96.67 | 124.00 |
| SMIM10L1 | 108.00 | 133.00 | 127.00 | 122.67 | 130.00 |
| SMIM12 | 723.00 | 721.00 | 742.00 | 728.67 | 674.00 |
| SMIM13 | 729.00 | 715.00 | 877.00 | 773.67 | 1127.00 |
| SMIM14 | 144.00 | 141.00 | 149.00 | 144.67 | 159.00 |
| SMIM15 | 1230.00 | 1116.00 | 1368.00 | 1238.00 | 1595.00 |
| SMIM17 | 6.00 | 9.00 | 6.00 | 7.00 | 6.00 |
| SMIM19 | 359.00 | 349.00 | 397.00 | 368.33 | 460.00 |
| SMIM20 | 289.00 | 280.00 | 319.00 | 296.00 | 356.00 |
| SMIM2-AS1 | 24.00 | 17.00 | 18.00 | 19.67 | 24.00 |
| SMIM3 | 85.00 | 96.00 | 96.00 | 92.33 | 89.00 |
| SMIM4 | 213.00 | 231.00 | 185.00 | 209.67 | 244.00 |
| SMIM7 | 854.00 | 773.00 | 859.00 | 828.67 | 1119.00 |
| SMIM8 | 130.00 | 116.00 | 144.00 | 130.00 | 161.00 |

| | | | | | |
|---|---|---|---|---|---|
| SMKR1 | 45.00 | 36.00 | 49.00 | 43.33 | 52.00 |
| SMNDC1 | 1165.00 | 1098.00 | 1230.00 | 1164.33 | 1312.00 |
| SMO | 173.00 | 204.00 | 211.00 | 196.00 | 197.00 |
| SMOC1 | 509.00 | 496.00 | 501.00 | 502.00 | 596.00 |
| SMOC2 | 47.00 | 52.00 | 37.00 | 45.33 | 39.00 |
| SMOX | 719.00 | 780.00 | 828.00 | 775.67 | 633.00 |
| SMPD1 | 1288.00 | 1434.00 | 1297.00 | 1339.67 | 1327.00 |
| SMPD2 | 189.00 | 175.00 | 174.00 | 179.33 | 235.00 |
| SMPD3 | 5.00 | 6.00 | 8.00 | 6.33 | 2.00 |
| SMPD4 | 1679.00 | 1591.00 | 1677.00 | 1649.00 | 1522.00 |
| SMPDL3A | 194.00 | 147.00 | 185.00 | 175.33 | 198.00 |
| SMS | 2725.00 | 2764.00 | 2956.00 | 2815.00 | 4297.00 |
| SMTN | 7797.00 | 7865.00 | 7912.00 | 7858.00 | 7814.00 |
| SMU1 | 1500.00 | 1493.00 | 1593.00 | 1528.67 | 1784.00 |
| SMUG1 | 557.00 | 558.00 | 589.00 | 568.00 | 620.00 |
| SMURF1 | 1100.00 | 1025.00 | 1228.00 | 1117.67 | 1031.00 |
| SMURF2 | 8200.00 | 7657.00 | 8872.00 | 8243.00 | 11001.00 |
| SMYD2 | 611.00 | 565.00 | 595.00 | 590.33 | 689.00 |
| SMYD3 | 450.00 | 405.00 | 537.00 | 464.00 | 590.00 |
| SMYD4 | 499.00 | 492.00 | 444.00 | 478.33 | 448.00 |
| SMYD5 | 1082.00 | 1055.00 | 1140.00 | 1092.33 | 1127.00 |
| SNAI1 | 34.00 | 29.00 | 36.00 | 33.00 | 50.00 |
| SNAI2 | 1173.00 | 1104.00 | 1277.00 | 1184.67 | 1101.00 |
| SNAI3 | 3.00 | 0.00 | 3.00 | 2.00 | 0.00 |
| SNAI3-AS1 | 87.00 | 83.00 | 69.00 | 79.67 | 77.00 |
| SNAP23 | 1501.00 | 1426.00 | 1568.00 | 1498.33 | 1597.00 |
| SNAP25 | 137.00 | 119.00 | 122.00 | 126.00 | 185.00 |
| SNAP29 | 504.00 | 504.00 | 492.00 | 500.00 | 552.00 |
| SNAP47 | 513.00 | 563.00 | 645.00 | 573.67 | 678.00 |
| SNAP91 | 67.00 | 64.00 | 56.00 | 62.33 | 67.00 |
| SNAPC1 | 1981.00 | 1818.00 | 2058.00 | 1952.33 | 1956.00 |
| SNAPC2 | 675.00 | 687.00 | 683.00 | 681.67 | 782.00 |
| SNAPC3 | 661.00 | 712.00 | 735.00 | 702.67 | 744.00 |
| SNAPC4 | 1439.00 | 1496.00 | 1485.00 | 1473.33 | 1472.00 |
| SNAPC5 | 173.00 | 175.00 | 156.00 | 168.00 | 212.00 |
| SNAPIN | 395.00 | 364.00 | 426.00 | 395.00 | 466.00 |
| SNCA | 306.00 | 293.00 | 319.00 | 306.00 | 234.00 |
| SNCAIP | 44.00 | 43.00 | 67.00 | 51.33 | 93.00 |
| SNCG | 39.00 | 41.00 | 38.00 | 39.33 | 30.00 |
| SND1 | 7450.00 | 6983.00 | 7837.00 | 7423.33 | 7045.00 |
| SNF8 | 813.00 | 742.00 | 813.00 | 789.33 | 975.00 |
| SNHG1 | 5536.00 | 5507.00 | 5716.00 | 5586.33 | 6789.00 |
| SNHG10 | 290.00 | 375.00 | 374.00 | 346.33 | 395.00 |
| SNHG11 | 494.00 | 474.00 | 512.00 | 493.33 | 550.00 |
| SNHG12 | 850.00 | 865.00 | 935.00 | 883.33 | 976.00 |
| SNHG15 | 1115.00 | 1141.00 | 1190.00 | 1148.67 | 1571.00 |
| SNHG16 | 2266.00 | 2129.00 | 2436.00 | 2277.00 | 2693.00 |
| SNHG17 | 1592.00 | 1590.00 | 1708.00 | 1630.00 | 2247.00 |

| | | | | | |
|---|---|---|---|---|---|
| SNHG18 | 270.00 | 229.00 | 234.00 | 244.33 | 267.00 |
| SNHG19 | 493.00 | 554.00 | 519.00 | 522.00 | 552.00 |
| SNHG20 | 510.00 | 493.00 | 543.00 | 515.33 | 517.00 |
| SNHG21 | 220.00 | 229.00 | 231.00 | 226.67 | 237.00 |
| SNHG22 | 8.00 | 10.00 | 9.00 | 9.00 | 13.00 |
| SNHG3 | 5222.00 | 5257.00 | 5735.00 | 5404.67 | 5662.00 |
| SNHG4 | 1615.00 | 1500.00 | 1659.00 | 1591.33 | 1411.00 |
| SNHG5 | 1134.00 | 1177.00 | 1238.00 | 1183.00 | 1490.00 |
| SNHG6 | 1699.00 | 1687.00 | 1670.00 | 1685.33 | 2431.00 |
| SNHG7 | 2964.00 | 2929.00 | 3103.00 | 2998.67 | 3356.00 |
| SNHG8 | 2044.00 | 1986.00 | 2199.00 | 2076.33 | 2520.00 |
| SNHG9 | 49.00 | 54.00 | 48.00 | 50.33 | 62.00 |
| SNIP1 | 459.00 | 365.00 | 428.00 | 417.33 | 455.00 |
| SNN | 165.00 | 117.00 | 141.00 | 141.00 | 162.00 |
| SNORA1 | 45.00 | 36.00 | 47.00 | 42.67 | 48.00 |
| SNORA10 | 24.00 | 30.00 | 25.00 | 26.33 | 46.00 |
| SNORA11 | 4.00 | 7.00 | 5.00 | 5.33 | 2.00 |
| SNORA12 | 7.00 | 3.00 | 2.00 | 4.00 | 6.00 |
| SNORA13 | 9.00 | 13.00 | 9.00 | 10.33 | 18.00 |
| SNORA14B | 12.00 | 13.00 | 15.00 | 13.33 | 20.00 |
| SNORA15 | 3.00 | 5.00 | 1.00 | 3.00 | 4.00 |
| SNORA16A | 54.00 | 57.00 | 52.00 | 54.33 | 49.00 |
| SNORA16B | 3.00 | 0.00 | 1.00 | 1.33 | 0.00 |
| SNORA18 | 207.00 | 226.00 | 207.00 | 213.33 | 209.00 |
| SNORA20 | 6.00 | 6.00 | 3.00 | 5.00 | 7.00 |
| SNORA21 | 31.00 | 21.00 | 35.00 | 29.00 | 28.00 |
| SNORA22 | 5.00 | 5.00 | 6.00 | 5.33 | 3.00 |
| SNORA23 | 6.00 | 2.00 | 6.00 | 4.67 | 3.00 |
| SNORA24 | 38.00 | 45.00 | 40.00 | 41.00 | 41.00 |
| SNORA25 | 125.00 | 127.00 | 135.00 | 129.00 | 125.00 |
| SNORA26 | 25.00 | 22.00 | 25.00 | 24.00 | 24.00 |
| SNORA27 | 40.00 | 45.00 | 32.00 | 39.00 | 43.00 |
| SNORA29 | 7.00 | 1.00 | 6.00 | 4.67 | 5.00 |
| SNORA2A | 6.00 | 3.00 | 10.00 | 6.33 | 6.00 |
| SNORA31 | 11.00 | 15.00 | 22.00 | 16.00 | 17.00 |
| SNORA32 | 128.00 | 166.00 | 137.00 | 143.67 | 159.00 |
| SNORA33 | 114.00 | 112.00 | 108.00 | 111.33 | 111.00 |
| SNORA4 | 44.00 | 28.00 | 49.00 | 40.33 | 65.00 |
| SNORA40 | 224.00 | 265.00 | 251.00 | 246.67 | 283.00 |
| SNORA41 | 21.00 | 24.00 | 27.00 | 24.00 | 22.00 |
| SNORA44 | 60.00 | 45.00 | 54.00 | 53.00 | 52.00 |
| SNORA45A | 46.00 | 52.00 | 50.00 | 49.33 | 38.00 |
| SNORA45B | 58.00 | 67.00 | 52.00 | 59.00 | 67.00 |
| SNORA47 | 11.00 | 14.00 | 12.00 | 12.33 | 12.00 |
| SNORA48 | 66.00 | 59.00 | 68.00 | 64.33 | 34.00 |
| SNORA51 | 30.00 | 34.00 | 35.00 | 33.00 | 29.00 |
| SNORA52 | 66.00 | 63.00 | 41.00 | 56.67 | 58.00 |
| SNORA53 | 7.00 | 8.00 | 12.00 | 9.00 | 11.00 |

| | | | | | |
|---|---|---|---|---|---|
| SNORA55 | 41.00 | 25.00 | 33.00 | 33.00 | 33.00 |
| SNORA57 | 18.00 | 24.00 | 25.00 | 22.33 | 25.00 |
| SNORA58 | 3.00 | 2.00 | 2.00 | 2.33 | 0.00 |
| SNORA5A | 9.00 | 18.00 | 13.00 | 13.33 | 15.00 |
| SNORA5B | 19.00 | 31.00 | 26.00 | 25.33 | 18.00 |
| SNORA5C | 55.00 | 43.00 | 51.00 | 49.67 | 44.00 |
| SNORA6 | 54.00 | 66.00 | 61.00 | 60.33 | 39.00 |
| SNORA60 | 18.00 | 27.00 | 29.00 | 24.67 | 24.00 |
| SNORA61 | 165.00 | 219.00 | 182.00 | 188.67 | 181.00 |
| SNORA62 | 84.00 | 83.00 | 76.00 | 81.00 | 83.00 |
| SNORA63 | 74.00 | 51.00 | 64.00 | 63.00 | 65.00 |
| SNORA64 | 55.00 | 66.00 | 61.00 | 60.67 | 66.00 |
| SNORA65 | 33.00 | 35.00 | 33.00 | 33.67 | 36.00 |
| SNORA66 | 37.00 | 27.00 | 33.00 | 32.33 | 48.00 |
| SNORA67 | 100.00 | 124.00 | 119.00 | 114.33 | 119.00 |
| SNORA68 | 39.00 | 47.00 | 44.00 | 43.33 | 36.00 |
| SNORA69 | 28.00 | 21.00 | 12.00 | 20.33 | 18.00 |
| SNORA70 | 281.00 | 300.00 | 320.00 | 300.33 | 332.00 |
| SNORA71A | 46.00 | 39.00 | 35.00 | 40.00 | 37.00 |
| SNORA71B | 44.00 | 54.00 | 51.00 | 49.67 | 49.00 |
| SNORA71C | 51.00 | 54.00 | 70.00 | 58.33 | 61.00 |
| SNORA71E | 9.00 | 16.00 | 12.00 | 12.33 | 5.00 |
| SNORA72 | 41.00 | 31.00 | 31.00 | 34.33 | 43.00 |
| SNORA73A | 243.00 | 259.00 | 257.00 | 253.00 | 262.00 |
| SNORA73B | 145.00 | 125.00 | 135.00 | 135.00 | 155.00 |
| SNORA74A | 6.00 | 6.00 | 10.00 | 7.33 | 7.00 |
| SNORA75 | 43.00 | 19.00 | 31.00 | 31.00 | 36.00 |
| SNORA76C | 20.00 | 9.00 | 16.00 | 15.00 | 17.00 |
| SNORA78 | 5.00 | 1.00 | 1.00 | 2.33 | 1.00 |
| SNORA7B | 10.00 | 9.00 | 10.00 | 9.67 | 5.00 |
| SNORA8 | 196.00 | 191.00 | 212.00 | 199.67 | 242.00 |
| SNORA80B | 13.00 | 6.00 | 10.00 | 9.67 | 8.00 |
| SNORA80E | 10.00 | 4.00 | 4.00 | 6.00 | 2.00 |
| SNORA81 | 87.00 | 84.00 | 74.00 | 81.67 | 58.00 |
| SNORA9 | 52.00 | 67.00 | 86.00 | 68.33 | 57.00 |
| SNORD10 | 81.00 | 89.00 | 85.00 | 85.00 | 92.00 |
| SNORD100 | 45.00 | 44.00 | 27.00 | 38.67 | 56.00 |
| SNORD101 | 63.00 | 49.00 | 57.00 | 56.33 | 71.00 |
| SNORD102 | 42.00 | 41.00 | 42.00 | 41.67 | 46.00 |
| SNORD104 | 83.00 | 116.00 | 90.00 | 96.33 | 111.00 |
| SNORD11 | 9.00 | 10.00 | 10.00 | 9.67 | 9.00 |
| SNORD110 | 113.00 | 109.00 | 126.00 | 116.00 | 101.00 |
| SNORD116-13 | 5.00 | 4.00 | 1.00 | 3.33 | 0.00 |
| SNORD116-2 | 4.00 | 1.00 | 1.00 | 2.00 | 2.00 |
| SNORD116-20 | 4.00 | 4.00 | 11.00 | 6.33 | 3.00 |
| SNORD116-21 | 14.00 | 7.00 | 4.00 | 8.33 | 7.00 |
| SNORD116-22 | 4.00 | 14.00 | 11.00 | 9.67 | 6.00 |
| SNORD116-23 | 5.00 | 4.00 | 2.00 | 3.67 | 2.00 |

| | | | | | |
|---|---|---|---|---|---|
| SNORD116-24 | 9.00 | 5.00 | 4.00 | 6.00 | 5.00 |
| SNORD116-28 | 12.00 | 8.00 | 13.00 | 11.00 | 8.00 |
| SNORD116-4 | 41.00 | 27.00 | 49.00 | 39.00 | 16.00 |
| SNORD119 | 5.00 | 7.00 | 6.00 | 6.00 | 13.00 |
| SNORD11B | 3.00 | 1.00 | 3.00 | 2.33 | 2.00 |
| SNORD12 | 73.00 | 73.00 | 60.00 | 68.67 | 44.00 |
| SNORD121A | 3.00 | 3.00 | 2.00 | 2.67 | 2.00 |
| SNORD123 | 7.00 | 2.00 | 4.00 | 4.33 | 8.00 |
| SNORD124 | 7.00 | 8.00 | 12.00 | 9.00 | 10.00 |
| SNORD12B | 60.00 | 46.00 | 55.00 | 53.67 | 43.00 |
| SNORD12C | 9.00 | 12.00 | 11.00 | 10.67 | 5.00 |
| SNORD14A | 69.00 | 75.00 | 63.00 | 69.00 | 74.00 |
| SNORD14B | 91.00 | 102.00 | 111.00 | 101.33 | 112.00 |
| SNORD14C | 11.00 | 5.00 | 5.00 | 7.00 | 7.00 |
| SNORD14D | 16.00 | 18.00 | 24.00 | 19.33 | 30.00 |
| SNORD14E | 16.00 | 17.00 | 15.00 | 16.00 | 11.00 |
| SNORD15A | 21.00 | 21.00 | 22.00 | 21.33 | 17.00 |
| SNORD15B | 30.00 | 29.00 | 27.00 | 28.67 | 26.00 |
| SNORD16 | 16.00 | 15.00 | 30.00 | 20.33 | 23.00 |
| SNORD17 | 21.00 | 16.00 | 20.00 | 19.00 | 29.00 |
| SNORD18B | 11.00 | 12.00 | 15.00 | 12.67 | 9.00 |
| SNORD18C | 50.00 | 50.00 | 38.00 | 46.00 | 49.00 |
| SNORD19 | 7.00 | 19.00 | 17.00 | 14.33 | 15.00 |
| SNORD19B | 25.00 | 38.00 | 36.00 | 33.00 | 38.00 |
| SNORD1A | 18.00 | 12.00 | 18.00 | 16.00 | 9.00 |
| SNORD1B | 8.00 | 5.00 | 10.00 | 7.67 | 5.00 |
| SNORD1C | 19.00 | 33.00 | 24.00 | 25.33 | 18.00 |
| SNORD2 | 67.00 | 54.00 | 67.00 | 62.67 | 64.00 |
| SNORD20 | 17.00 | 20.00 | 19.00 | 18.67 | 21.00 |
| SNORD21 | 27.00 | 31.00 | 25.00 | 27.67 | 25.00 |
| SNORD22 | 140.00 | 182.00 | 171.00 | 164.33 | 186.00 |
| SNORD23 | 16.00 | 17.00 | 12.00 | 15.00 | 23.00 |
| SNORD24 | 15.00 | 15.00 | 18.00 | 16.00 | 15.00 |
| SNORD25 | 57.00 | 70.00 | 54.00 | 60.33 | 69.00 |
| SNORD26 | 37.00 | 39.00 | 20.00 | 32.00 | 32.00 |
| SNORD27 | 5.00 | 5.00 | 11.00 | 7.00 | 17.00 |
| SNORD28 | 60.00 | 57.00 | 80.00 | 65.67 | 85.00 |
| SNORD29 | 24.00 | 14.00 | 29.00 | 22.33 | 21.00 |
| SNORD30 | 30.00 | 28.00 | 33.00 | 30.33 | 43.00 |
| SNORD31 | 213.00 | 228.00 | 223.00 | 221.33 | 216.00 |
| SNORD32A | 26.00 | 21.00 | 22.00 | 23.00 | 20.00 |
| SNORD35A | 36.00 | 44.00 | 31.00 | 37.00 | 39.00 |
| SNORD35B | 29.00 | 37.00 | 44.00 | 36.67 | 34.00 |
| SNORD36A | 196.00 | 188.00 | 188.00 | 190.67 | 280.00 |
| SNORD36C | 37.00 | 27.00 | 30.00 | 31.33 | 29.00 |
| SNORD37 | 5.00 | 4.00 | 3.00 | 4.00 | 6.00 |
| SNORD38A | 10.00 | 13.00 | 8.00 | 10.33 | 17.00 |
| SNORD38B | 14.00 | 29.00 | 31.00 | 24.67 | 39.00 |

| | | | | | |
|---|---|---|---|---|---|
| SNORD3A | 5.00 | 3.00 | 1.00 | 3.00 | 3.00 |
| SNORD42A | 9.00 | 20.00 | 11.00 | 13.33 | 18.00 |
| SNORD42B | 16.00 | 10.00 | 9.00 | 11.67 | 2.00 |
| SNORD43 | 26.00 | 34.00 | 30.00 | 30.00 | 32.00 |
| SNORD44 | 137.00 | 129.00 | 121.00 | 129.00 | 156.00 |
| SNORD45A | 137.00 | 124.00 | 166.00 | 142.33 | 113.00 |
| SNORD45B | 207.00 | 213.00 | 222.00 | 214.00 | 191.00 |
| SNORD45C | 109.00 | 120.00 | 126.00 | 118.33 | 86.00 |
| SNORD46 | 18.00 | 21.00 | 22.00 | 20.33 | 28.00 |
| SNORD47 | 171.00 | 196.00 | 184.00 | 183.67 | 156.00 |
| SNORD49A | 99.00 | 91.00 | 107.00 | 99.00 | 95.00 |
| SNORD4A | 34.00 | 30.00 | 35.00 | 33.00 | 35.00 |
| SNORD4B | 5.00 | 5.00 | 3.00 | 4.33 | 5.00 |
| SNORD5 | 87.00 | 76.00 | 112.00 | 91.67 | 88.00 |
| SNORD50A | 3.00 | 3.00 | 4.00 | 3.33 | 1.00 |
| SNORD50B | 29.00 | 45.00 | 41.00 | 38.33 | 37.00 |
| SNORD51 | 9.00 | 14.00 | 7.00 | 10.00 | 16.00 |
| SNORD53 | 3.00 | 3.00 | 2.00 | 2.67 | 2.00 |
| SNORD54 | 26.00 | 36.00 | 43.00 | 35.00 | 40.00 |
| SNORD55 | 24.00 | 23.00 | 30.00 | 25.67 | 26.00 |
| SNORD57 | 5.00 | 6.00 | 2.00 | 4.33 | 8.00 |
| SNORD58A | 50.00 | 46.00 | 57.00 | 51.00 | 52.00 |
| SNORD58C | 43.00 | 91.00 | 57.00 | 63.67 | 72.00 |
| SNORD59A | 3.00 | 3.00 | 2.00 | 2.67 | 3.00 |
| SNORD59B | 6.00 | 6.00 | 5.00 | 5.67 | 2.00 |
| SNORD6 | 43.00 | 27.00 | 41.00 | 37.00 | 37.00 |
| SNORD60 | 8.00 | 7.00 | 18.00 | 11.00 | 17.00 |
| SNORD63 | 22.00 | 27.00 | 26.00 | 25.00 | 22.00 |
| SNORD65 | 10.00 | 11.00 | 12.00 | 11.00 | 11.00 |
| SNORD67 | 5.00 | 3.00 | 1.00 | 3.00 | 2.00 |
| SNORD68 | 31.00 | 23.00 | 24.00 | 26.00 | 34.00 |
| SNORD69 | 5.00 | 9.00 | 6.00 | 6.67 | 9.00 |
| SNORD7 | 10.00 | 10.00 | 5.00 | 8.33 | 11.00 |
| SNORD72 | 9.00 | 4.00 | 6.00 | 6.33 | 3.00 |
| SNORD74 | 45.00 | 58.00 | 34.00 | 45.67 | 45.00 |
| SNORD75 | 4.00 | 3.00 | 4.00 | 3.67 | 9.00 |
| SNORD76 | 50.00 | 43.00 | 43.00 | 45.33 | 36.00 |
| SNORD77 | 80.00 | 76.00 | 74.00 | 76.67 | 75.00 |
| SNORD78 | 70.00 | 99.00 | 88.00 | 85.67 | 101.00 |
| SNORD79 | 26.00 | 22.00 | 26.00 | 24.67 | 26.00 |
| SNORD80 | 443.00 | 458.00 | 415.00 | 438.67 | 456.00 |
| SNORD81 | 23.00 | 31.00 | 25.00 | 26.33 | 28.00 |
| SNORD83A | 97.00 | 97.00 | 82.00 | 92.00 | 79.00 |
| SNORD83B | 118.00 | 118.00 | 127.00 | 121.00 | 99.00 |
| SNORD85 | 3.00 | 1.00 | 2.00 | 2.00 | 0.00 |
| SNORD87 | 31.00 | 28.00 | 31.00 | 30.00 | 25.00 |
| SNORD88B | 14.00 | 26.00 | 13.00 | 17.67 | 28.00 |
| SNORD88C | 3.00 | 2.00 | 10.00 | 5.00 | 16.00 |

| | | | | | |
|---|---|---|---|---|---|
| SNORD89 | 44.00 | 36.00 | 28.00 | 36.00 | 46.00 |
| SNORD93 | 12.00 | 10.00 | 12.00 | 11.33 | 13.00 |
| SNORD94 | 18.00 | 16.00 | 19.00 | 17.67 | 13.00 |
| SNORD95 | 28.00 | 20.00 | 18.00 | 22.00 | 10.00 |
| SNORD96A | 289.00 | 307.00 | 280.00 | 292.00 | 301.00 |
| SNORD97 | 18.00 | 16.00 | 25.00 | 19.67 | 21.00 |
| SNORD98 | 4.00 | 8.00 | 1.00 | 4.33 | 1.00 |
| SNORD99 | 16.00 | 21.00 | 22.00 | 19.67 | 20.00 |
| SNPH | 493.00 | 547.00 | 555.00 | 531.67 | 615.00 |
| SNRK | 626.00 | 560.00 | 623.00 | 603.00 | 588.00 |
| SNRK-AS1 | 383.00 | 361.00 | 438.00 | 394.00 | 306.00 |
| SNRNP200 | 8198.00 | 7855.00 | 8655.00 | 8236.00 | 7408.00 |
| SNRNP25 | 249.00 | 210.00 | 174.00 | 211.00 | 310.00 |
| SNRNP27 | 487.00 | 427.00 | 548.00 | 487.33 | 704.00 |
| SNRNP35 | 259.00 | 231.00 | 275.00 | 255.00 | 336.00 |
| SNRNP40 | 1169.00 | 1055.00 | 1153.00 | 1125.67 | 1385.00 |
| SNRNP48 | 750.00 | 648.00 | 718.00 | 705.33 | 726.00 |
| SNRNP70 | 3760.00 | 3920.00 | 4024.00 | 3901.33 | 4247.00 |
| SNRPA | 1921.00 | 1875.00 | 1920.00 | 1905.33 | 2497.00 |
| SNRPA1 | 942.00 | 990.00 | 989.00 | 973.67 | 1235.00 |
| SNRPB | 5331.00 | 5178.00 | 5576.00 | 5361.67 | 7324.00 |
| SNRPB2 | 1658.00 | 1588.00 | 1621.00 | 1622.33 | 2194.00 |
| SNRPC | 1870.00 | 1935.00 | 1904.00 | 1903.00 | 2823.00 |
| SNRPD1 | 1227.00 | 1144.00 | 1195.00 | 1188.67 | 1882.00 |
| SNRPD2 | 1912.00 | 1933.00 | 2018.00 | 1954.33 | 2744.00 |
| SNRPD3 | 1647.00 | 1612.00 | 1754.00 | 1671.00 | 2198.00 |
| SNRPE | 1242.00 | 1183.00 | 1237.00 | 1220.67 | 1949.00 |
| SNRPF | 1043.00 | 1071.00 | 1077.00 | 1063.67 | 1577.00 |
| SNRPG | 967.00 | 934.00 | 1018.00 | 973.00 | 1459.00 |
| SNTA1 | 107.00 | 122.00 | 130.00 | 119.67 | 148.00 |
| SNTB1 | 100.00 | 122.00 | 103.00 | 108.33 | 108.00 |
| SNTB2 | 930.00 | 837.00 | 944.00 | 903.67 | 1024.00 |
| SNU13 | 1739.00 | 1639.00 | 1794.00 | 1724.00 | 2514.00 |
| SNUPN | 547.00 | 518.00 | 552.00 | 539.00 | 666.00 |
| SNW1 | 1832.00 | 1778.00 | 1857.00 | 1822.33 | 2122.00 |
| SNX1 | 2671.00 | 2525.00 | 2776.00 | 2657.33 | 2493.00 |
| SNX10 | 56.00 | 47.00 | 62.00 | 55.00 | 56.00 |
| SNX11 | 367.00 | 374.00 | 412.00 | 384.33 | 383.00 |
| SNX12 | 1044.00 | 893.00 | 1094.00 | 1010.33 | 1303.00 |
| SNX13 | 1130.00 | 1041.00 | 1134.00 | 1101.67 | 1166.00 |
| SNX14 | 2726.00 | 2548.00 | 2551.00 | 2608.33 | 2791.00 |
| SNX16 | 244.00 | 227.00 | 291.00 | 254.00 | 320.00 |
| SNX17 | 1834.00 | 1816.00 | 1772.00 | 1807.33 | 2276.00 |
| SNX18 | 356.00 | 320.00 | 316.00 | 330.67 | 407.00 |
| SNX19 | 1889.00 | 1732.00 | 1889.00 | 1836.67 | 1534.00 |
| SNX2 | 1444.00 | 1324.00 | 1606.00 | 1458.00 | 1918.00 |
| SNX21 | 451.00 | 433.00 | 461.00 | 448.33 | 408.00 |
| SNX22 | 3.00 | 1.00 | 0.00 | 1.33 | 1.00 |

| | | | | | |
|---|---|---|---|---|---|
| SNX24 | 239.00 | 265.00 | 295.00 | 266.33 | 358.00 |
| SNX25 | 417.00 | 433.00 | 469.00 | 439.67 | 576.00 |
| SNX27 | 1308.00 | 1257.00 | 1379.00 | 1314.67 | 1265.00 |
| SNX29 | 539.00 | 494.00 | 500.00 | 511.00 | 365.00 |
| SNX29P1 | 12.00 | 9.00 | 17.00 | 12.67 | 6.00 |
| SNX29P2 | 5.00 | 2.00 | 5.00 | 4.00 | 2.00 |
| SNX3 | 1832.00 | 1735.00 | 1871.00 | 1812.67 | 2166.00 |
| SNX30 | 511.00 | 395.00 | 544.00 | 483.33 | 433.00 |
| SNX32 | 64.00 | 52.00 | 58.00 | 58.00 | 61.00 |
| SNX33 | 1558.00 | 1557.00 | 1651.00 | 1588.67 | 1171.00 |
| SNX4 | 296.00 | 325.00 | 298.00 | 306.33 | 427.00 |
| SNX5 | 2676.00 | 2756.00 | 2751.00 | 2727.67 | 3371.00 |
| SNX6 | 1708.00 | 1504.00 | 1735.00 | 1649.00 | 1839.00 |
| SNX7 | 481.00 | 463.00 | 495.00 | 479.67 | 632.00 |
| SNX8 | 1080.00 | 1049.00 | 1130.00 | 1086.33 | 1140.00 |
| SNX9 | 1692.00 | 1691.00 | 1861.00 | 1748.00 | 1581.00 |
| SOAT1 | 2707.00 | 2561.00 | 2752.00 | 2673.33 | 2842.00 |
| SOBP | 163.00 | 180.00 | 173.00 | 172.00 | 159.00 |
| SOCS1 | 5.00 | 13.00 | 6.00 | 8.00 | 11.00 |
| SOCS2 | 515.00 | 469.00 | 568.00 | 517.33 | 658.00 |
| SOCS2-AS1 | 81.00 | 50.00 | 64.00 | 65.00 | 59.00 |
| SOCS3 | 1080.00 | 1031.00 | 1196.00 | 1102.33 | 1099.00 |
| SOCS4 | 1267.00 | 1245.00 | 1403.00 | 1305.00 | 1284.00 |
| SOCS5 | 604.00 | 526.00 | 672.00 | 600.67 | 556.00 |
| SOCS6 | 385.00 | 370.00 | 432.00 | 395.67 | 337.00 |
| SOCS7 | 65.00 | 80.00 | 67.00 | 70.67 | 61.00 |
| SOD1 | 1884.00 | 1872.00 | 1936.00 | 1897.33 | 2540.00 |
| SOD2 | 493.00 | 508.00 | 617.00 | 539.33 | 689.00 |
| SOGA1 | 955.00 | 919.00 | 1026.00 | 966.67 | 626.00 |
| SOGA3 | 515.00 | 467.00 | 532.00 | 504.67 | 325.00 |
| SOHLH2 | 5.00 | 2.00 | 9.00 | 5.33 | 3.00 |
| SON | 7891.00 | 7151.00 | 7894.00 | 7645.33 | 6748.00 |
| SORBS1 | 8.00 | 5.00 | 6.00 | 6.33 | 4.00 |
| SORBS2 | 107.00 | 111.00 | 130.00 | 116.00 | 102.00 |
| SORBS3 | 794.00 | 776.00 | 811.00 | 793.67 | 895.00 |
| SORCS1 | 12.00 | 8.00 | 13.00 | 11.00 | 6.00 |
| SORCS2 | 424.00 | 426.00 | 446.00 | 432.00 | 276.00 |
| SORCS3 | 759.00 | 754.00 | 780.00 | 764.33 | 572.00 |
| SORD | 946.00 | 921.00 | 921.00 | 929.33 | 1211.00 |
| SORT1 | 852.00 | 803.00 | 777.00 | 810.67 | 779.00 |
| SOS1 | 1391.00 | 1170.00 | 1382.00 | 1314.33 | 1225.00 |
| SOS2 | 857.00 | 699.00 | 835.00 | 797.00 | 758.00 |
| SOST | 3.00 | 0.00 | 0.00 | 1.00 | 1.00 |
| SOWAHC | 264.00 | 295.00 | 313.00 | 290.67 | 370.00 |
| SOX12 | 1481.00 | 1511.00 | 1549.00 | 1513.67 | 1847.00 |
| SOX13 | 161.00 | 136.00 | 127.00 | 141.33 | 113.00 |
| SOX15 | 16.00 | 9.00 | 17.00 | 14.00 | 14.00 |
| SOX21-AS1 | 15.00 | 16.00 | 14.00 | 15.00 | 14.00 |

| | | | | | |
|---|---|---|---|---|---|
| SOX2-OT | 3.00 | 11.00 | 7.00 | 7.00 | 12.00 |
| SOX30 | 3.00 | 1.00 | 7.00 | 3.67 | 3.00 |
| SOX4 | 576.00 | 515.00 | 676.00 | 589.00 | 629.00 |
| SOX5 | 13.00 | 8.00 | 29.00 | 16.67 | 12.00 |
| SOX6 | 10.00 | 12.00 | 23.00 | 15.00 | 7.00 |
| SOX7 | 44.00 | 53.00 | 65.00 | 54.00 | 42.00 |
| SOX9 | 1559.00 | 1583.00 | 1583.00 | 1575.00 | 2439.00 |
| SOX9-AS1 | 74.00 | 71.00 | 63.00 | 69.33 | 115.00 |
| SP1 | 2264.00 | 2119.00 | 2306.00 | 2229.67 | 1965.00 |
| SP100 | 2068.00 | 1986.00 | 2221.00 | 2091.67 | 2069.00 |
| SP110 | 333.00 | 306.00 | 345.00 | 328.00 | 347.00 |
| SP140 | 152.00 | 171.00 | 167.00 | 163.33 | 132.00 |
| SP140L | 780.00 | 783.00 | 835.00 | 799.33 | 818.00 |
| SP2 | 481.00 | 437.00 | 468.00 | 462.00 | 495.00 |
| SP2-AS1 | 27.00 | 32.00 | 38.00 | 32.33 | 39.00 |
| SP3 | 1525.00 | 1315.00 | 1639.00 | 1493.00 | 1647.00 |
| SP4 | 408.00 | 298.00 | 402.00 | 369.33 | 392.00 |
| SP5 | 8.00 | 7.00 | 9.00 | 8.00 | 11.00 |
| SP6 | 4.00 | 4.00 | 10.00 | 6.00 | 7.00 |
| SP8 | 13.00 | 13.00 | 15.00 | 13.67 | 17.00 |
| SPA17 | 126.00 | 116.00 | 134.00 | 125.33 | 190.00 |
| SPACA6P | 92.00 | 91.00 | 85.00 | 89.33 | 88.00 |
| SPAG1 | 180.00 | 153.00 | 157.00 | 163.33 | 222.00 |
| SPAG16 | 286.00 | 291.00 | 318.00 | 298.33 | 378.00 |
| SPAG4 | 219.00 | 215.00 | 222.00 | 218.67 | 140.00 |
| SPAG5 | 1332.00 | 1274.00 | 1302.00 | 1302.67 | 1450.00 |
| SPAG5-AS1 | 3.00 | 4.00 | 5.00 | 4.00 | 7.00 |
| SPAG7 | 2530.00 | 2705.00 | 2825.00 | 2686.67 | 2767.00 |
| SPAG8 | 38.00 | 48.00 | 41.00 | 42.33 | 43.00 |
| SPAG9 | 3976.00 | 3656.00 | 4231.00 | 3954.33 | 3545.00 |
| SPANXA2-OT1 | 3.00 | 3.00 | 3.00 | 3.00 | 3.00 |
| SPANXB1 | 277.00 | 239.00 | 245.00 | 253.67 | 335.00 |
| SPANXC | 30.00 | 23.00 | 24.00 | 25.67 | 34.00 |
| SPARC | 6149.00 | 6098.00 | 5637.00 | 5961.33 | 5932.00 |
| SPAST | 412.00 | 316.00 | 430.00 | 386.00 | 528.00 |
| SPATA1 | 5.00 | 2.00 | 6.00 | 4.33 | 8.00 |
| SPATA13 | 1162.00 | 1127.00 | 1144.00 | 1144.33 | 886.00 |
| SPATA17 | 70.00 | 49.00 | 59.00 | 59.33 | 62.00 |
| SPATA2 | 516.00 | 530.00 | 517.00 | 521.00 | 641.00 |
| SPATA20 | 2113.00 | 2109.00 | 2310.00 | 2177.33 | 1985.00 |
| SPATA24 | 77.00 | 87.00 | 60.00 | 74.67 | 85.00 |
| SPATA25 | 214.00 | 179.00 | 174.00 | 189.00 | 191.00 |
| SPATA2L | 201.00 | 196.00 | 172.00 | 189.67 | 308.00 |
| SPATA33 | 205.00 | 240.00 | 207.00 | 217.33 | 206.00 |
| SPATA3-AS1 | 5.00 | 8.00 | 4.00 | 5.67 | 14.00 |
| SPATA4 | 21.00 | 9.00 | 13.00 | 14.33 | 21.00 |
| SPATA5 | 208.00 | 176.00 | 243.00 | 209.00 | 205.00 |
| SPATA5L1 | 549.00 | 524.00 | 579.00 | 550.67 | 570.00 |

| | | | | | |
|---|---|---|---|---|---|
| SPATA6 | 55.00 | 64.00 | 80.00 | 66.33 | 92.00 |
| SPATA6L | 55.00 | 64.00 | 52.00 | 57.00 | 34.00 |
| SPATA7 | 178.00 | 182.00 | 185.00 | 181.67 | 189.00 |
| SPATA9 | 9.00 | 6.00 | 11.00 | 8.67 | 11.00 |
| SPATS2 | 4381.00 | 4124.00 | 4574.00 | 4359.67 | 6027.00 |
| SPATS2L | 2094.00 | 2057.00 | 2266.00 | 2139.00 | 2244.00 |
| SPC24 | 95.00 | 97.00 | 109.00 | 100.33 | 127.00 |
| SPC25 | 5.00 | 6.00 | 2.00 | 4.33 | 9.00 |
| SPCS1 | 1882.00 | 1877.00 | 1779.00 | 1846.00 | 2465.00 |
| SPCS2 | 581.00 | 550.00 | 483.00 | 538.00 | 698.00 |
| SPCS3 | 3117.00 | 2856.00 | 3001.00 | 2991.33 | 3870.00 |
| SPDL1 | 1227.00 | 1185.00 | 1318.00 | 1243.33 | 1367.00 |
| SPDYA | 8.00 | 4.00 | 10.00 | 7.33 | 6.00 |
| SPDYC | 3.00 | 4.00 | 5.00 | 4.00 | 3.00 |
| SPECC1 | 82.00 | 69.00 | 91.00 | 80.67 | 109.00 |
| SPECC1L | 1077.00 | 1002.00 | 1076.00 | 1051.67 | 1087.00 |
| SPEF1 | 5.00 | 19.00 | 14.00 | 12.67 | 7.00 |
| SPEF2 | 169.00 | 130.00 | 191.00 | 163.33 | 165.00 |
| SPEG | 3021.00 | 3202.00 | 3443.00 | 3222.00 | 2410.00 |
| SPEN | 4031.00 | 3654.00 | 3996.00 | 3893.67 | 2374.00 |
| SPESP1 | 144.00 | 158.00 | 121.00 | 141.00 | 157.00 |
| SPG11 | 2134.00 | 1851.00 | 2068.00 | 2017.67 | 1789.00 |
| SPG20 | 1730.00 | 1562.00 | 1844.00 | 1712.00 | 1910.00 |
| SPG20-AS1 | 23.00 | 31.00 | 27.00 | 27.00 | 30.00 |
| SPG21 | 1704.00 | 1668.00 | 1851.00 | 1741.00 | 2069.00 |
| SPG7 | 3228.00 | 3400.00 | 3468.00 | 3365.33 | 3395.00 |
| SPHK1 | 3568.00 | 3607.00 | 3765.00 | 3646.67 | 3216.00 |
| SPHK2 | 607.00 | 604.00 | 659.00 | 623.33 | 694.00 |
| SPICE1 | 453.00 | 420.00 | 448.00 | 440.33 | 409.00 |
| SPIDR | 937.00 | 964.00 | 1089.00 | 996.67 | 961.00 |
| SPIN1 | 1962.00 | 1721.00 | 2019.00 | 1900.67 | 1889.00 |
| SPIN2B | 27.00 | 29.00 | 37.00 | 31.00 | 42.00 |
| SPIN3 | 857.00 | 883.00 | 886.00 | 875.33 | 867.00 |
| SPIN4 | 408.00 | 367.00 | 464.00 | 413.00 | 566.00 |
| SPINK1 | 4.00 | 7.00 | 6.00 | 5.67 | 2.00 |
| SPINK2 | 3.00 | 2.00 | 1.00 | 2.00 | 1.00 |
| SPINT2 | 33.00 | 35.00 | 26.00 | 31.33 | 22.00 |
| SPIRE1 | 2586.00 | 2432.00 | 2874.00 | 2630.67 | 2851.00 |
| SPIRE2 | 365.00 | 392.00 | 390.00 | 382.33 | 365.00 |
| SPNS1 | 1640.00 | 1697.00 | 1631.00 | 1656.00 | 1803.00 |
| SPNS2 | 7.00 | 2.00 | 3.00 | 4.00 | 7.00 |
| SPNS3 | 30.00 | 39.00 | 40.00 | 36.33 | 21.00 |
| SPOCD1 | 2476.00 | 2571.00 | 2766.00 | 2604.33 | 2167.00 |
| SPOCK1 | 361.00 | 328.00 | 438.00 | 375.67 | 139.00 |
| SPOCK2 | 3.00 | 7.00 | 12.00 | 7.33 | 4.00 |
| SPOCK3 | 7.00 | 7.00 | 10.00 | 8.00 | 6.00 |
| SPOP | 1116.00 | 1069.00 | 1100.00 | 1095.00 | 1339.00 |
| SPOPL | 580.00 | 513.00 | 636.00 | 576.33 | 702.00 |

| | | | | | |
|---|---|---|---|---|---|
| SPP1 | 76.00 | 81.00 | 84.00 | 80.33 | 141.00 |
| SPPL2A | 873.00 | 787.00 | 801.00 | 820.33 | 1026.00 |
| SPPL2B | 1674.00 | 1753.00 | 1717.00 | 1714.67 | 1655.00 |
| SPPL3 | 889.00 | 899.00 | 920.00 | 902.67 | 1100.00 |
| SPR | 414.00 | 373.00 | 440.00 | 409.00 | 657.00 |
| SPRED1 | 2545.00 | 2293.00 | 2840.00 | 2559.33 | 1938.00 |
| SPRED2 | 1782.00 | 1667.00 | 1768.00 | 1739.00 | 2150.00 |
| SPRED3 | 299.00 | 280.00 | 303.00 | 294.00 | 307.00 |
| SPRTN | 225.00 | 228.00 | 252.00 | 235.00 | 305.00 |
| SPRY1 | 75.00 | 64.00 | 86.00 | 75.00 | 114.00 |
| SPRY2 | 4612.00 | 4263.00 | 4930.00 | 4601.67 | 5536.00 |
| SPRY4 | 3603.00 | 3363.00 | 3934.00 | 3633.33 | 3466.00 |
| SPRY4-IT1 | 123.00 | 85.00 | 105.00 | 104.33 | 50.00 |
| SPRYD3 | 525.00 | 545.00 | 604.00 | 558.00 | 620.00 |
| SPRYD4 | 255.00 | 244.00 | 244.00 | 247.67 | 206.00 |
| SPRYD7 | 249.00 | 239.00 | 252.00 | 246.67 | 338.00 |
| SPSB1 | 864.00 | 839.00 | 984.00 | 895.67 | 868.00 |
| SPSB2 | 276.00 | 283.00 | 303.00 | 287.33 | 304.00 |
| SPSB3 | 653.00 | 662.00 | 726.00 | 680.33 | 678.00 |
| SPTAN1 | 5716.00 | 5339.00 | 5890.00 | 5648.33 | 4827.00 |
| SPTB | 4.00 | 3.00 | 3.00 | 3.33 | 2.00 |
| SPTBN1 | 4415.00 | 4109.00 | 4438.00 | 4320.67 | 2938.00 |
| SPTBN2 | 3631.00 | 3759.00 | 3929.00 | 3773.00 | 3689.00 |
| SPTBN4 | 8.00 | 3.00 | 6.00 | 5.67 | 0.00 |
| SPTBN5 | 193.00 | 212.00 | 204.00 | 203.00 | 190.00 |
| SPTLC1 | 2168.00 | 2121.00 | 1984.00 | 2091.00 | 2186.00 |
| SPTLC2 | 1268.00 | 1223.00 | 1252.00 | 1247.67 | 1082.00 |
| SPTLC3 | 464.00 | 459.00 | 513.00 | 478.67 | 418.00 |
| SPTSSA | 359.00 | 315.00 | 417.00 | 363.67 | 467.00 |
| SPTY2D1 | 567.00 | 546.00 | 598.00 | 570.33 | 550.00 |
| SPTY2D1-AS1 | 41.00 | 31.00 | 38.00 | 36.67 | 44.00 |
| SPX | 703.00 | 575.00 | 764.00 | 680.67 | 644.00 |
| SQLE | 1670.00 | 1597.00 | 1686.00 | 1651.00 | 1844.00 |
| SQRDL | 318.00 | 281.00 | 300.00 | 299.67 | 300.00 |
| SQSTM1 | 8120.00 | 8042.00 | 8386.00 | 8182.67 | 6556.00 |
| SRA1 | 885.00 | 903.00 | 947.00 | 911.67 | 1112.00 |
| SRBD1 | 397.00 | 356.00 | 428.00 | 393.67 | 433.00 |
| SRC | 2204.00 | 2194.00 | 2291.00 | 2229.67 | 2176.00 |
| SRCAP | 4390.00 | 4100.00 | 4737.00 | 4409.00 | 3116.00 |
| SRD5A1 | 430.00 | 436.00 | 419.00 | 428.33 | 443.00 |
| SRD5A3 | 187.00 | 148.00 | 163.00 | 166.00 | 281.00 |
| SRD5A3-AS1 | 14.00 | 6.00 | 15.00 | 11.67 | 7.00 |
| SREBF1 | 3682.00 | 3603.00 | 3799.00 | 3694.67 | 3800.00 |
| SREBF2 | 2232.00 | 2310.00 | 2248.00 | 2263.33 | 2281.00 |
| SREK1 | 2911.00 | 2908.00 | 3322.00 | 3047.00 | 3272.00 |
| SREK1IP1 | 602.00 | 495.00 | 672.00 | 589.67 | 636.00 |
| SRF | 1579.00 | 1619.00 | 1700.00 | 1632.67 | 1897.00 |
| SRFBP1 | 467.00 | 448.00 | 456.00 | 457.00 | 527.00 |

| | | | | | |
|---|---|---|---|---|---|
| SRGAP1 | 2226.00 | 1983.00 | 2319.00 | 2176.00 | 2211.00 |
| SRGAP2 | 1784.00 | 1744.00 | 1939.00 | 1822.33 | 1612.00 |
| SRGAP2B | 108.00 | 99.00 | 109.00 | 105.33 | 79.00 |
| SRGAP2C | 72.00 | 41.00 | 74.00 | 62.33 | 58.00 |
| SRGAP2D | 25.00 | 29.00 | 51.00 | 35.00 | 21.00 |
| SRGAP3 | 11.00 | 8.00 | 17.00 | 12.00 | 6.00 |
| SRGN | 7099.00 | 6573.00 | 6338.00 | 6670.00 | 9712.00 |
| SRI | 1342.00 | 1308.00 | 1320.00 | 1323.33 | 1899.00 |
| SRM | 1645.00 | 1516.00 | 1635.00 | 1598.67 | 2104.00 |
| SRP14 | 1926.00 | 1822.00 | 1984.00 | 1910.67 | 2330.00 |
| SRP14-AS1 | 228.00 | 264.00 | 219.00 | 237.00 | 152.00 |
| SRP19 | 701.00 | 689.00 | 651.00 | 680.33 | 740.00 |
| SRP54 | 1801.00 | 1531.00 | 1596.00 | 1642.67 | 1755.00 |
| SRP68 | 1764.00 | 1674.00 | 1794.00 | 1744.00 | 1990.00 |
| SRP72 | 2145.00 | 1852.00 | 2195.00 | 2064.00 | 2328.00 |
| SRP9 | 2795.00 | 2533.00 | 2814.00 | 2714.00 | 3697.00 |
| SRPK1 | 3626.00 | 3372.00 | 3769.00 | 3589.00 | 4275.00 |
| SRPK2 | 684.00 | 695.00 | 807.00 | 728.67 | 852.00 |
| SRPR | 3423.00 | 3208.00 | 3290.00 | 3307.00 | 3436.00 |
| SRPRB | 2419.00 | 2414.00 | 2320.00 | 2384.33 | 2710.00 |
| SRPX | 5974.00 | 6053.00 | 5754.00 | 5927.00 | 7331.00 |
| SRPX2 | 1290.00 | 1328.00 | 1311.00 | 1309.67 | 1116.00 |
| SRR | 151.00 | 194.00 | 165.00 | 170.00 | 176.00 |
| SRRD | 322.00 | 328.00 | 353.00 | 334.33 | 409.00 |
| SRRM1 | 3058.00 | 3074.00 | 3525.00 | 3219.00 | 2957.00 |
| SRRM2 | 31849.00 | 30292.00 | 34391.00 | 32177.33 | 22692.00 |
| SRRM2-AS1 | 9.00 | 9.00 | 7.00 | 8.33 | 6.00 |
| SRRM5 | 18.00 | 21.00 | 22.00 | 20.33 | 25.00 |
| SRRT | 2958.00 | 3166.00 | 3104.00 | 3076.00 | 3427.00 |
| SRSF1 | 8553.00 | 7998.00 | 9010.00 | 8520.33 | 10303.00 |
| SRSF10 | 3095.00 | 2960.00 | 3266.00 | 3107.00 | 3662.00 |
| SRSF11 | 6920.00 | 6713.00 | 7529.00 | 7054.00 | 8062.00 |
| SRSF2 | 7266.00 | 7214.00 | 7820.00 | 7433.33 | 8586.00 |
| SRSF3 | 5458.00 | 5320.00 | 5911.00 | 5563.00 | 7168.00 |
| SRSF4 | 2570.00 | 2544.00 | 2791.00 | 2635.00 | 2777.00 |
| SRSF5 | 5397.00 | 5487.00 | 5756.00 | 5546.67 | 5610.00 |
| SRSF6 | 7999.00 | 8287.00 | 8946.00 | 8410.67 | 8852.00 |
| SRSF7 | 3468.00 | 3464.00 | 3527.00 | 3486.33 | 4288.00 |
| SRSF8 | 747.00 | 628.00 | 811.00 | 728.67 | 801.00 |
| SRSF9 | 2573.00 | 2573.00 | 2668.00 | 2604.67 | 3209.00 |
| SRXN1 | 1620.00 | 1458.00 | 1619.00 | 1565.67 | 2010.00 |
| SS18 | 2376.00 | 2342.00 | 2521.00 | 2413.00 | 2230.00 |
| SS18L1 | 1219.00 | 1237.00 | 1328.00 | 1261.33 | 1315.00 |
| SS18L2 | 302.00 | 281.00 | 274.00 | 285.67 | 367.00 |
| SSB | 3062.00 | 2930.00 | 3260.00 | 3084.00 | 3986.00 |
| SSBP1 | 601.00 | 528.00 | 585.00 | 571.33 | 781.00 |
| SSBP2 | 114.00 | 95.00 | 109.00 | 106.00 | 148.00 |
| SSBP3 | 1100.00 | 1122.00 | 1148.00 | 1123.33 | 1268.00 |

| | | | | | |
|---|---|---|---|---|---|
| SSBP4 | 993.00 | 904.00 | 1049.00 | 982.00 | 1095.00 |
| SSC4D | 10.00 | 13.00 | 18.00 | 13.67 | 14.00 |
| SSC5D | 40.00 | 38.00 | 45.00 | 41.00 | 28.00 |
| SSFA2 | 3347.00 | 2792.00 | 3620.00 | 3253.00 | 3320.00 |
| SSH1 | 1916.00 | 1692.00 | 2017.00 | 1875.00 | 1513.00 |
| SSH2 | 702.00 | 594.00 | 712.00 | 669.33 | 610.00 |
| SSH3 | 211.00 | 202.00 | 213.00 | 208.67 | 216.00 |
| SSNA1 | 663.00 | 687.00 | 710.00 | 686.67 | 937.00 |
| SSPN | 10.00 | 8.00 | 15.00 | 11.00 | 13.00 |
| SSPO | 7.00 | 2.00 | 6.00 | 5.00 | 1.00 |
| SSR1 | 7603.00 | 7179.00 | 7338.00 | 7373.33 | 7616.00 |
| SSR2 | 5530.00 | 5488.00 | 5207.00 | 5408.33 | 6936.00 |
| SSR3 | 21072.00 | 19343.00 | 19538.00 | 19984.33 | 28925.00 |
| SSR4 | 2263.00 | 2179.00 | 2030.00 | 2157.33 | 2542.00 |
| SSR4P1 | 74.00 | 71.00 | 76.00 | 73.67 | 86.00 |
| SSRP1 | 5247.00 | 4987.00 | 5656.00 | 5296.67 | 6486.00 |
| SSSCA1 | 639.00 | 622.00 | 628.00 | 629.67 | 858.00 |
| SSSCA1-AS1 | 59.00 | 43.00 | 44.00 | 48.67 | 43.00 |
| SSU72 | 1501.00 | 1497.00 | 1633.00 | 1543.67 | 2072.00 |
| SSUH2 | 14.00 | 14.00 | 14.00 | 14.00 | 9.00 |
| SSX2IP | 486.00 | 434.00 | 535.00 | 485.00 | 615.00 |
| ST13 | 2815.00 | 2713.00 | 3009.00 | 2845.67 | 3566.00 |
| ST13P4 | 41.00 | 31.00 | 43.00 | 38.33 | 77.00 |
| ST20 | 365.00 | 377.00 | 383.00 | 375.00 | 413.00 |
| ST20-AS1 | 40.00 | 58.00 | 58.00 | 52.00 | 72.00 |
| ST20-MTHFS | 7.00 | 6.00 | 6.00 | 6.33 | 10.00 |
| ST3GAL1 | 2931.00 | 2791.00 | 3025.00 | 2915.67 | 1944.00 |
| ST3GAL2 | 568.00 | 579.00 | 575.00 | 574.00 | 572.00 |
| ST3GAL3 | 595.00 | 595.00 | 648.00 | 612.67 | 666.00 |
| ST3GAL4 | 289.00 | 260.00 | 241.00 | 263.33 | 269.00 |
| ST3GAL4-AS1 | 39.00 | 41.00 | 27.00 | 35.67 | 30.00 |
| ST3GAL5 | 128.00 | 121.00 | 129.00 | 126.00 | 112.00 |
| ST3GAL6 | 853.00 | 887.00 | 993.00 | 911.00 | 564.00 |
| ST3GAL6-AS1 | 78.00 | 86.00 | 95.00 | 86.33 | 59.00 |
| ST5 | 668.00 | 638.00 | 666.00 | 657.33 | 552.00 |
| ST6GAL1 | 349.00 | 372.00 | 402.00 | 374.33 | 434.00 |
| ST6GAL2 | 14.00 | 10.00 | 14.00 | 12.67 | 17.00 |
| ST6GALNAC2 | 8.00 | 3.00 | 3.00 | 4.67 | 1.00 |
| ST6GALNAC3 | 298.00 | 232.00 | 290.00 | 273.33 | 229.00 |
| ST6GALNAC4 | 554.00 | 551.00 | 501.00 | 535.33 | 597.00 |
| ST6GALNAC6 | 832.00 | 902.00 | 961.00 | 898.33 | 1147.00 |
| ST7 | 968.00 | 929.00 | 1030.00 | 975.67 | 880.00 |
| ST7-AS1 | 50.00 | 47.00 | 48.00 | 48.33 | 35.00 |
| ST7L | 1136.00 | 1115.00 | 1173.00 | 1141.33 | 1260.00 |
| ST7-OT4 | 11.00 | 9.00 | 19.00 | 13.00 | 7.00 |
| ST8SIA1 | 246.00 | 208.00 | 244.00 | 232.67 | 268.00 |
| ST8SIA4 | 83.00 | 92.00 | 81.00 | 85.33 | 90.00 |
| ST8SIA5 | 55.00 | 39.00 | 39.00 | 44.33 | 33.00 |

| | | | | | |
|---|---|---|---|---|---|
| STAB1 | 5.00 | 4.00 | 5.00 | 4.67 | 2.00 |
| STAC | 154.00 | 134.00 | 167.00 | 151.67 | 118.00 |
| STAC3 | 71.00 | 81.00 | 72.00 | 74.67 | 70.00 |
| STAG1 | 1076.00 | 974.00 | 1141.00 | 1063.67 | 1011.00 |
| STAG2 | 2315.00 | 1952.00 | 2466.00 | 2244.33 | 2254.00 |
| STAG3 | 239.00 | 265.00 | 246.00 | 250.00 | 303.00 |
| STAG3L1 | 39.00 | 22.00 | 24.00 | 28.33 | 20.00 |
| STAG3L2 | 258.00 | 236.00 | 197.00 | 230.33 | 167.00 |
| STAG3L3 | 77.00 | 98.00 | 107.00 | 94.00 | 75.00 |
| STAG3L4 | 184.00 | 166.00 | 206.00 | 185.33 | 204.00 |
| STAG3L5P | 5.00 | 4.00 | 8.00 | 5.67 | 8.00 |
| AG3L5P-PVRIG2P-PIL | 820.00 | 850.00 | 882.00 | 850.67 | 782.00 |
| STAM | 906.00 | 883.00 | 1009.00 | 932.67 | 1055.00 |
| STAM2 | 632.00 | 600.00 | 715.00 | 649.00 | 704.00 |
| STAM-AS1 | 77.00 | 85.00 | 110.00 | 90.67 | 78.00 |
| STAMBP | 1027.00 | 877.00 | 1062.00 | 988.67 | 1147.00 |
| STAMBPL1 | 974.00 | 846.00 | 1004.00 | 941.33 | 1054.00 |
| STAP2 | 76.00 | 63.00 | 97.00 | 78.67 | 119.00 |
| STARD10 | 179.00 | 194.00 | 238.00 | 203.67 | 261.00 |
| STARD13 | 2744.00 | 2516.00 | 2905.00 | 2721.67 | 2582.00 |
| STARD3 | 1601.00 | 1604.00 | 1585.00 | 1596.67 | 1619.00 |
| STARD3NL | 1332.00 | 1208.00 | 1348.00 | 1296.00 | 1286.00 |
| STARD4 | 331.00 | 297.00 | 357.00 | 328.33 | 451.00 |
| STARD5 | 58.00 | 50.00 | 55.00 | 54.33 | 45.00 |
| STARD7 | 3226.00 | 3009.00 | 3365.00 | 3200.00 | 3767.00 |
| STARD7-AS1 | 228.00 | 222.00 | 249.00 | 233.00 | 205.00 |
| STARD8 | 483.00 | 441.00 | 511.00 | 478.33 | 460.00 |
| STARD9 | 756.00 | 685.00 | 778.00 | 739.67 | 448.00 |
| STAT1 | 1482.00 | 1387.00 | 1651.00 | 1506.67 | 1669.00 |
| STAT2 | 1298.00 | 1299.00 | 1336.00 | 1311.00 | 1101.00 |
| STAT3 | 1151.00 | 1075.00 | 1111.00 | 1112.33 | 1156.00 |
| STAT4 | 30.00 | 35.00 | 40.00 | 35.00 | 22.00 |
| STAT5A | 146.00 | 148.00 | 170.00 | 154.67 | 158.00 |
| STAT5B | 1669.00 | 1589.00 | 1788.00 | 1682.00 | 1579.00 |
| STAT6 | 4107.00 | 4100.00 | 4461.00 | 4222.67 | 3822.00 |
| STAU1 | 2526.00 | 2361.00 | 2631.00 | 2506.00 | 2896.00 |
| STAU2 | 657.00 | 653.00 | 647.00 | 652.33 | 702.00 |
| STBD1 | 7.00 | 17.00 | 12.00 | 12.00 | 11.00 |
| STC1 | 4321.00 | 4087.00 | 4611.00 | 4339.67 | 3521.00 |
| STC2 | 15862.00 | 16470.00 | 16653.00 | 16328.33 | 13437.00 |
| STEAP1 | 1293.00 | 1271.00 | 1181.00 | 1248.33 | 2130.00 |
| STEAP1B | 934.00 | 853.00 | 863.00 | 883.33 | 1551.00 |
| STEAP2 | 813.00 | 798.00 | 955.00 | 855.33 | 1092.00 |
| STEAP3 | 5071.00 | 4978.00 | 5338.00 | 5129.00 | 4132.00 |
| STIL | 575.00 | 593.00 | 607.00 | 591.67 | 782.00 |
| STIM1 | 1303.00 | 1161.00 | 1222.00 | 1228.67 | 1030.00 |
| STIM2 | 372.00 | 379.00 | 360.00 | 370.33 | 412.00 |
| STIP1 | 9115.00 | 9098.00 | 10202.00 | 9471.67 | 10739.00 |

| | | | | | |
|---|---|---|---|---|---|
| STK10 | 1153.00 | 1133.00 | 1125.00 | 1137.00 | 832.00 |
| STK11 | 1261.00 | 1268.00 | 1392.00 | 1307.00 | 1483.00 |
| STK11IP | 672.00 | 699.00 | 741.00 | 704.00 | 869.00 |
| STK16 | 661.00 | 635.00 | 687.00 | 661.00 | 763.00 |
| STK17A | 1533.00 | 1446.00 | 1650.00 | 1543.00 | 1850.00 |
| STK17B | 550.00 | 505.00 | 593.00 | 549.33 | 467.00 |
| STK24 | 1877.00 | 1784.00 | 2023.00 | 1894.67 | 2119.00 |
| STK25 | 2575.00 | 2471.00 | 2628.00 | 2558.00 | 3029.00 |
| STK26 | 373.00 | 305.00 | 424.00 | 367.33 | 420.00 |
| STK3 | 527.00 | 472.00 | 513.00 | 504.00 | 608.00 |
| STK32A | 37.00 | 54.00 | 70.00 | 53.67 | 48.00 |
| STK32C | 36.00 | 32.00 | 40.00 | 36.00 | 40.00 |
| STK33 | 201.00 | 183.00 | 252.00 | 212.00 | 199.00 |
| STK35 | 1530.00 | 1450.00 | 1584.00 | 1521.33 | 1448.00 |
| STK36 | 1460.00 | 1435.00 | 1482.00 | 1459.00 | 1304.00 |
| STK38 | 1169.00 | 1252.00 | 1255.00 | 1225.33 | 1241.00 |
| STK38L | 496.00 | 470.00 | 559.00 | 508.33 | 637.00 |
| STK39 | 1225.00 | 1122.00 | 1225.00 | 1190.67 | 1277.00 |
| STK4 | 1933.00 | 1668.00 | 1944.00 | 1848.33 | 1528.00 |
| STK40 | 1429.00 | 1337.00 | 1351.00 | 1372.33 | 1139.00 |
| STK4-AS1 | 7.00 | 8.00 | 12.00 | 9.00 | 20.00 |
| STKLD1 | 11.00 | 14.00 | 8.00 | 11.00 | 15.00 |
| STMN1 | 2286.00 | 2057.00 | 2218.00 | 2187.00 | 3811.00 |
| STMN3 | 964.00 | 993.00 | 1041.00 | 999.33 | 1490.00 |
| STOM | 1295.00 | 1201.00 | 1299.00 | 1265.00 | 1611.00 |
| STOML1 | 322.00 | 324.00 | 285.00 | 310.33 | 331.00 |
| STOML2 | 3867.00 | 3597.00 | 4090.00 | 3851.33 | 5625.00 |
| STON1 | 109.00 | 99.00 | 100.00 | 102.67 | 87.00 |
| STOX1 | 3.00 | 4.00 | 4.00 | 3.67 | 5.00 |
| STPG1 | 400.00 | 323.00 | 288.00 | 337.00 | 299.00 |
| STPG2 | 3.00 | 4.00 | 1.00 | 2.67 | 4.00 |
| STRA13 | 608.00 | 581.00 | 675.00 | 621.33 | 922.00 |
| STRADA | 927.00 | 956.00 | 1053.00 | 978.67 | 1123.00 |
| STRADB | 673.00 | 620.00 | 722.00 | 671.67 | 677.00 |
| STRAP | 4272.00 | 4058.00 | 4562.00 | 4297.33 | 4040.00 |
| STRBP | 529.00 | 553.00 | 592.00 | 558.00 | 813.00 |
| STRIP1 | 1149.00 | 1095.00 | 1162.00 | 1135.33 | 988.00 |
| STRIP2 | 2363.00 | 2335.00 | 2411.00 | 2369.67 | 1399.00 |
| STRN | 833.00 | 782.00 | 954.00 | 856.33 | 767.00 |
| STRN3 | 567.00 | 564.00 | 614.00 | 581.67 | 642.00 |
| STRN4 | 1859.00 | 1811.00 | 1999.00 | 1889.67 | 2106.00 |
| STS | 224.00 | 237.00 | 249.00 | 236.67 | 200.00 |
| STT3A | 5703.00 | 5653.00 | 6018.00 | 5791.33 | 5464.00 |
| STT3B | 7968.00 | 7584.00 | 8124.00 | 7892.00 | 7989.00 |
| STUB1 | 1980.00 | 1977.00 | 2094.00 | 2017.00 | 2139.00 |
| STX10 | 825.00 | 879.00 | 934.00 | 879.33 | 1173.00 |
| STX11 | 38.00 | 35.00 | 24.00 | 32.33 | 26.00 |
| STX12 | 682.00 | 644.00 | 762.00 | 696.00 | 825.00 |

| | | | | | |
|---|---|---|---|---|---|
| STX16 | 4597.00 | 4451.00 | 4965.00 | 4671.00 | 4462.00 |
| STX16-NPEPL1 | 48.00 | 32.00 | 42.00 | 40.67 | 39.00 |
| STX17 | 539.00 | 447.00 | 514.00 | 500.00 | 396.00 |
| STX18 | 441.00 | 435.00 | 417.00 | 431.00 | 498.00 |
| STX18-AS1 | 25.00 | 34.00 | 35.00 | 31.33 | 20.00 |
| STX19 | 4.00 | 1.00 | 1.00 | 2.00 | 2.00 |
| STX1A | 1512.00 | 1484.00 | 1470.00 | 1488.67 | 1017.00 |
| STX1B | 788.00 | 831.00 | 845.00 | 821.33 | 551.00 |
| STX2 | 784.00 | 790.00 | 880.00 | 818.00 | 909.00 |
| STX3 | 1241.00 | 1156.00 | 1459.00 | 1285.33 | 1206.00 |
| STX4 | 921.00 | 928.00 | 897.00 | 915.33 | 960.00 |
| STX5 | 529.00 | 525.00 | 577.00 | 543.67 | 537.00 |
| STX6 | 990.00 | 862.00 | 902.00 | 918.00 | 880.00 |
| STX7 | 848.00 | 748.00 | 939.00 | 845.00 | 1060.00 |
| STX8 | 302.00 | 316.00 | 344.00 | 320.67 | 435.00 |
| STXBP1 | 1541.00 | 1452.00 | 1567.00 | 1520.00 | 1511.00 |
| STXBP2 | 5.00 | 7.00 | 4.00 | 5.33 | 2.00 |
| STXBP3 | 763.00 | 695.00 | 863.00 | 773.67 | 917.00 |
| STXBP4 | 196.00 | 194.00 | 186.00 | 192.00 | 233.00 |
| STXBP5 | 755.00 | 700.00 | 889.00 | 781.33 | 671.00 |
| STXBP5-AS1 | 273.00 | 287.00 | 333.00 | 297.67 | 310.00 |
| STYK1 | 87.00 | 72.00 | 53.00 | 70.67 | 285.00 |
| STYX | 405.00 | 347.00 | 392.00 | 381.33 | 440.00 |
| STYXL1 | 870.00 | 845.00 | 921.00 | 878.67 | 1071.00 |
| SUB1 | 2012.00 | 1897.00 | 2219.00 | 2042.67 | 2597.00 |
| SUCLA2 | 408.00 | 417.00 | 505.00 | 443.33 | 482.00 |
| SUCLG1 | 999.00 | 955.00 | 952.00 | 968.67 | 1300.00 |
| SUCLG2 | 2064.00 | 1757.00 | 2094.00 | 1971.67 | 1919.00 |
| SUCLG2-AS1 | 156.00 | 141.00 | 189.00 | 162.00 | 117.00 |
| SUCO | 1963.00 | 1757.00 | 1916.00 | 1878.67 | 1663.00 |
| SUDS3 | 1127.00 | 1054.00 | 1113.00 | 1098.00 | 1338.00 |
| SUFU | 654.00 | 591.00 | 669.00 | 638.00 | 516.00 |
| SUGCT | 92.00 | 90.00 | 115.00 | 99.00 | 136.00 |
| SUGP1 | 725.00 | 778.00 | 736.00 | 746.33 | 816.00 |
| SUGP2 | 4166.00 | 3936.00 | 4323.00 | 4141.67 | 3508.00 |
| SUGT1 | 691.00 | 670.00 | 720.00 | 693.67 | 995.00 |
| SUGT1P1 | 142.00 | 153.00 | 185.00 | 160.00 | 186.00 |
| SUGT1P3 | 9.00 | 0.00 | 6.00 | 5.00 | 10.00 |
| SULT1C2 | 159.00 | 185.00 | 208.00 | 184.00 | 62.00 |
| SUMF1 | 771.00 | 728.00 | 699.00 | 732.67 | 715.00 |
| SUMF2 | 6814.00 | 6890.00 | 6631.00 | 6778.33 | 7988.00 |
| SUMO1 | 1357.00 | 1224.00 | 1462.00 | 1347.67 | 1971.00 |
| SUMO1P3 | 22.00 | 23.00 | 24.00 | 23.00 | 33.00 |
| SUMO2 | 1468.00 | 1363.00 | 1548.00 | 1459.67 | 2045.00 |
| SUMO3 | 1682.00 | 1643.00 | 1792.00 | 1705.67 | 2340.00 |
| SUN1 | 4347.00 | 4374.00 | 4641.00 | 4454.00 | 4054.00 |
| SUN2 | 1864.00 | 2003.00 | 1851.00 | 1906.00 | 1762.00 |
| SUOX | 299.00 | 350.00 | 325.00 | 324.67 | 378.00 |

| | | | | | |
|---|---|---|---|---|---|
| SUPT16H | 4697.00 | 4517.00 | 4881.00 | 4698.33 | 5388.00 |
| SUPT20H | 1191.00 | 1147.00 | 1282.00 | 1206.67 | 1279.00 |
| SUPT3H | 610.00 | 570.00 | 685.00 | 621.67 | 729.00 |
| SUPT4H1 | 1026.00 | 921.00 | 1012.00 | 986.33 | 1144.00 |
| SUPT5H | 3113.00 | 3197.00 | 3449.00 | 3253.00 | 2940.00 |
| SUPT6H | 4196.00 | 4152.00 | 4396.00 | 4248.00 | 3878.00 |
| SUPT7L | 2840.00 | 2825.00 | 2983.00 | 2882.67 | 2812.00 |
| SUPV3L1 | 1228.00 | 1147.00 | 1278.00 | 1217.67 | 1387.00 |
| SURF1 | 507.00 | 622.00 | 513.00 | 547.33 | 644.00 |
| SURF2 | 285.00 | 314.00 | 340.00 | 313.00 | 412.00 |
| SURF4 | 9243.00 | 9438.00 | 8871.00 | 9184.00 | 8481.00 |
| SURF6 | 1205.00 | 1218.00 | 1288.00 | 1237.00 | 1550.00 |
| SUSD1 | 144.00 | 114.00 | 112.00 | 123.33 | 130.00 |
| SUSD2 | 7.00 | 5.00 | 10.00 | 7.33 | 7.00 |
| SUSD5 | 106.00 | 87.00 | 105.00 | 99.33 | 82.00 |
| SUSD6 | 532.00 | 539.00 | 528.00 | 533.00 | 506.00 |
| SUV39H1 | 439.00 | 391.00 | 424.00 | 418.00 | 535.00 |
| SUV39H2 | 438.00 | 387.00 | 404.00 | 409.67 | 516.00 |
| SUV420H1 | 857.00 | 746.00 | 921.00 | 841.33 | 840.00 |
| SUV420H2 | 239.00 | 266.00 | 273.00 | 259.33 | 260.00 |
| SUZ12 | 1056.00 | 971.00 | 1130.00 | 1052.33 | 1046.00 |
| SUZ12P1 | 203.00 | 216.00 | 220.00 | 213.00 | 240.00 |
| SV2A | 1184.00 | 1119.00 | 1165.00 | 1156.00 | 861.00 |
| SVBP | 74.00 | 79.00 | 79.00 | 77.33 | 124.00 |
| SVEP1 | 94.00 | 89.00 | 90.00 | 91.00 | 103.00 |
| SVIL | 972.00 | 835.00 | 949.00 | 918.67 | 954.00 |
| SVIL-AS1 | 624.00 | 551.00 | 611.00 | 595.33 | 608.00 |
| SVILP1 | 79.00 | 91.00 | 103.00 | 91.00 | 53.00 |
| SVIP | 290.00 | 288.00 | 336.00 | 304.67 | 515.00 |
| SWAP70 | 966.00 | 868.00 | 1018.00 | 950.67 | 1183.00 |
| SWI5 | 725.00 | 664.00 | 732.00 | 707.00 | 915.00 |
| SWSAP1 | 33.00 | 38.00 | 22.00 | 31.00 | 49.00 |
| SWT1 | 174.00 | 160.00 | 179.00 | 171.00 | 169.00 |
| SYAP1 | 1147.00 | 1081.00 | 1156.00 | 1128.00 | 1398.00 |
| SYBU | 24.00 | 39.00 | 40.00 | 34.33 | 51.00 |
| SYCE1L | 3.00 | 2.00 | 1.00 | 2.00 | 4.00 |
| SYCP2 | 25.00 | 27.00 | 42.00 | 31.33 | 46.00 |
| SYCP2L | 57.00 | 34.00 | 38.00 | 43.00 | 51.00 |
| SYCP3 | 4.00 | 5.00 | 4.00 | 4.33 | 2.00 |
| SYDE1 | 912.00 | 918.00 | 837.00 | 889.00 | 932.00 |
| SYDE2 | 101.00 | 92.00 | 120.00 | 104.33 | 129.00 |
| SYF2 | 855.00 | 811.00 | 992.00 | 886.00 | 1141.00 |
| SYMPK | 2763.00 | 2845.00 | 2940.00 | 2849.33 | 3031.00 |
| SYNC | 329.00 | 318.00 | 319.00 | 322.00 | 433.00 |
| SYNCRIP | 7457.00 | 6707.00 | 7622.00 | 7262.00 | 8218.00 |
| SYNE1 | 1408.00 | 1395.00 | 1587.00 | 1463.33 | 1347.00 |
| SYNE2 | 373.00 | 355.00 | 354.00 | 360.67 | 415.00 |
| SYNE3 | 208.00 | 200.00 | 212.00 | 206.67 | 111.00 |

| | | | | | |
|---|---|---|---|---|---|
| SYNE4 | 39.00 | 41.00 | 46.00 | 42.00 | 57.00 |
| SYNGAP1 | 176.00 | 106.00 | 136.00 | 139.33 | 66.00 |
| SYNGR1 | 310.00 | 298.00 | 299.00 | 302.33 | 220.00 |
| SYNGR2 | 1195.00 | 1207.00 | 1058.00 | 1153.33 | 1305.00 |
| SYNGR3 | 11.00 | 11.00 | 4.00 | 8.67 | 10.00 |
| SYNJ1 | 393.00 | 321.00 | 360.00 | 358.00 | 306.00 |
| SYNJ2 | 1976.00 | 2029.00 | 2191.00 | 2065.33 | 1476.00 |
| SYNJ2BP | 714.00 | 672.00 | 742.00 | 709.33 | 718.00 |
| SYNM | 433.00 | 385.00 | 452.00 | 423.33 | 397.00 |
| SYNPO | 1610.00 | 1536.00 | 1634.00 | 1593.33 | 930.00 |
| SYNPO2 | 12.00 | 22.00 | 16.00 | 16.67 | 7.00 |
| SYNPR-AS1 | 6.00 | 3.00 | 3.00 | 4.00 | 1.00 |
| SYNRG | 5.00 | 4.00 | 4.00 | 4.33 | 7.00 |
| SYP | 38.00 | 45.00 | 47.00 | 43.33 | 54.00 |
| SYPL1 | 2746.00 | 2801.00 | 2776.00 | 2774.33 | 3289.00 |
| SYPL2 | 10.00 | 7.00 | 8.00 | 8.33 | 12.00 |
| SYS1 | 711.00 | 703.00 | 642.00 | 685.33 | 759.00 |
| SYT1 | 501.00 | 443.00 | 542.00 | 495.33 | 531.00 |
| SYT11 | 866.00 | 832.00 | 910.00 | 869.33 | 883.00 |
| SYT14 | 90.00 | 105.00 | 118.00 | 104.33 | 104.00 |
| SYT16 | 52.00 | 41.00 | 51.00 | 48.00 | 61.00 |
| SYT17 | 3.00 | 2.00 | 2.00 | 2.33 | 0.00 |
| SYT6 | 4.00 | 7.00 | 6.00 | 5.67 | 10.00 |
| SYT9 | 116.00 | 76.00 | 111.00 | 101.00 | 141.00 |
| SYTL1 | 10.00 | 10.00 | 10.00 | 10.00 | 41.00 |
| SYTL2 | 654.00 | 628.00 | 741.00 | 674.33 | 841.00 |
| SYTL3 | 244.00 | 254.00 | 227.00 | 241.67 | 195.00 |
| SYTL4 | 123.00 | 112.00 | 112.00 | 115.67 | 122.00 |
| SYVN1 | 3217.00 | 3097.00 | 3100.00 | 3138.00 | 2595.00 |
| SZRD1 | 3259.00 | 3266.00 | 3401.00 | 3308.67 | 3752.00 |
| SZT2 | 1841.00 | 1824.00 | 2015.00 | 1893.33 | 1380.00 |
| TAB1 | 584.00 | 619.00 | 613.00 | 605.33 | 592.00 |
| TAB2 | 1632.00 | 1454.00 | 1743.00 | 1609.67 | 1686.00 |
| TAB3 | 445.00 | 420.00 | 516.00 | 460.33 | 449.00 |
| TACC1 | 3288.00 | 3142.00 | 3368.00 | 3266.00 | 1935.00 |
| TACC2 | 1237.00 | 1189.00 | 1292.00 | 1239.33 | 1219.00 |
| TACC3 | 8151.00 | 8261.00 | 7850.00 | 8087.33 | 9437.00 |
| TACO1 | 580.00 | 501.00 | 619.00 | 566.67 | 697.00 |
| TACSTD2 | 19.00 | 26.00 | 33.00 | 26.00 | 4.00 |
| TADA1 | 254.00 | 266.00 | 300.00 | 273.33 | 317.00 |
| TADA2A | 10.00 | 7.00 | 22.00 | 13.00 | 14.00 |
| TADA2B | 530.00 | 485.00 | 578.00 | 531.00 | 531.00 |
| TADA3 | 2324.00 | 2170.00 | 2352.00 | 2282.00 | 2827.00 |
| TAF1 | 2265.00 | 1995.00 | 2299.00 | 2186.33 | 1724.00 |
| TAF10 | 707.00 | 618.00 | 692.00 | 672.33 | 888.00 |
| TAF11 | 633.00 | 608.00 | 640.00 | 627.00 | 720.00 |
| TAF12 | 476.00 | 485.00 | 507.00 | 489.33 | 532.00 |
| TAF13 | 186.00 | 158.00 | 207.00 | 183.67 | 236.00 |

| | | | | | |
|---|---|---|---|---|---|
| TAF15 | 22.00 | 10.00 | 7.00 | 13.00 | 13.00 |
| TAF1A | 399.00 | 350.00 | 350.00 | 366.33 | 409.00 |
| TAF1A-AS1 | 47.00 | 30.00 | 30.00 | 35.67 | 38.00 |
| TAF1B | 456.00 | 380.00 | 398.00 | 411.33 | 594.00 |
| TAF1C | 2321.00 | 2405.00 | 2414.00 | 2380.00 | 2377.00 |
| TAF1D | 1904.00 | 1951.00 | 2181.00 | 2012.00 | 2115.00 |
| TAF1L | 3.00 | 4.00 | 5.00 | 4.00 | 2.00 |
| TAF2 | 1674.00 | 1493.00 | 1837.00 | 1668.00 | 1594.00 |
| TAF3 | 296.00 | 245.00 | 270.00 | 270.33 | 272.00 |
| TAF4 | 119.00 | 119.00 | 129.00 | 122.33 | 110.00 |
| TAF4B | 608.00 | 540.00 | 612.00 | 586.67 | 690.00 |
| TAF5 | 219.00 | 177.00 | 219.00 | 205.00 | 253.00 |
| TAF5L | 699.00 | 673.00 | 710.00 | 694.00 | 760.00 |
| TAF6 | 1650.00 | 1528.00 | 1707.00 | 1628.33 | 1675.00 |
| TAF6L | 408.00 | 414.00 | 431.00 | 417.67 | 476.00 |
| TAF7 | 2024.00 | 1864.00 | 2077.00 | 1988.33 | 2171.00 |
| TAF8 | 613.00 | 507.00 | 534.00 | 551.33 | 563.00 |
| TAF9 | 11.00 | 5.00 | 0.00 | 5.33 | 0.00 |
| TAF9B | 676.00 | 562.00 | 712.00 | 650.00 | 846.00 |
| TAGLN | 61.00 | 68.00 | 94.00 | 74.33 | 61.00 |
| TAGLN2 | 4455.00 | 4362.00 | 4348.00 | 4388.33 | 4844.00 |
| TAGLN3 | 640.00 | 591.00 | 649.00 | 626.67 | 919.00 |
| TAL2 | 5.00 | 7.00 | 3.00 | 5.00 | 7.00 |
| TALDO1 | 2862.00 | 2634.00 | 2851.00 | 2782.33 | 3506.00 |
| TAMM41 | 226.00 | 207.00 | 237.00 | 223.33 | 262.00 |
| TANC1 | 851.00 | 821.00 | 955.00 | 875.67 | 674.00 |
| TANC2 | 1571.00 | 1433.00 | 1752.00 | 1585.33 | 974.00 |
| TANGO2 | 385.00 | 374.00 | 370.00 | 376.33 | 400.00 |
| TANGO6 | 413.00 | 429.00 | 459.00 | 433.67 | 463.00 |
| TANK | 653.00 | 677.00 | 745.00 | 691.67 | 838.00 |
| TAOK1 | 1730.00 | 1569.00 | 2017.00 | 1772.00 | 1611.00 |
| TAOK2 | 1952.00 | 1946.00 | 2048.00 | 1982.00 | 1536.00 |
| TAOK3 | 1580.00 | 1449.00 | 1551.00 | 1526.67 | 2146.00 |
| TAP1 | 3.00 | 1.00 | 1.00 | 1.67 | 1.00 |
| TAP2 | 14.00 | 12.00 | 5.00 | 10.33 | 9.00 |
| TAPBP | 30.00 | 22.00 | 29.00 | 27.00 | 23.00 |
| TAPBPL | 241.00 | 244.00 | 225.00 | 236.67 | 211.00 |
| TAPT1 | 388.00 | 362.00 | 366.00 | 372.00 | 348.00 |
| TAPT1-AS1 | 73.00 | 62.00 | 82.00 | 72.33 | 77.00 |
| TARBP1 | 1787.00 | 1576.00 | 1956.00 | 1773.00 | 1626.00 |
| TARBP2 | 723.00 | 719.00 | 872.00 | 771.33 | 829.00 |
| TARDBP | 4284.00 | 4292.00 | 4442.00 | 4339.33 | 4382.00 |
| TARID | 8.00 | 5.00 | 11.00 | 8.00 | 11.00 |
| TARS | 10354.00 | 9634.00 | 11297.00 | 10428.33 | 11752.00 |
| TARS2 | 1252.00 | 1170.00 | 1306.00 | 1242.67 | 1236.00 |
| TARSL2 | 230.00 | 236.00 | 258.00 | 241.33 | 217.00 |
| TAS2R19 | 4.00 | 3.00 | 1.00 | 2.67 | 10.00 |
| TAS2R4 | 10.00 | 22.00 | 18.00 | 16.67 | 12.00 |

| | | | | | |
|---|---|---|---|---|---|
| TAS2R5 | 112.00 | 109.00 | 132.00 | 117.67 | 111.00 |
| TASP1 | 216.00 | 180.00 | 205.00 | 200.33 | 253.00 |
| TATDN1 | 973.00 | 880.00 | 998.00 | 950.33 | 1332.00 |
| TATDN2 | 1058.00 | 1000.00 | 1080.00 | 1046.00 | 944.00 |
| TATDN3 | 267.00 | 241.00 | 274.00 | 260.67 | 374.00 |
| TAX1BP1 | 2735.00 | 2466.00 | 2759.00 | 2653.33 | 2937.00 |
| TAX1BP3 | 151.00 | 170.00 | 153.00 | 158.00 | 199.00 |
| TAZ | 1255.00 | 1212.00 | 1312.00 | 1259.67 | 1281.00 |
| TBC1D1 | 1452.00 | 1513.00 | 1517.00 | 1494.00 | 1451.00 |
| TBC1D10A | 250.00 | 282.00 | 214.00 | 248.67 | 302.00 |
| TBC1D10B | 1220.00 | 1291.00 | 1302.00 | 1271.00 | 1222.00 |
| TBC1D12 | 305.00 | 275.00 | 290.00 | 290.00 | 293.00 |
| TBC1D13 | 990.00 | 949.00 | 977.00 | 972.00 | 1012.00 |
| TBC1D14 | 1649.00 | 1596.00 | 1796.00 | 1680.33 | 1762.00 |
| TBC1D15 | 745.00 | 712.00 | 851.00 | 769.33 | 776.00 |
| TBC1D16 | 2227.00 | 2395.00 | 2426.00 | 2349.33 | 1795.00 |
| TBC1D17 | 513.00 | 477.00 | 557.00 | 515.67 | 593.00 |
| TBC1D19 | 51.00 | 43.00 | 53.00 | 49.00 | 64.00 |
| TBC1D2 | 206.00 | 229.00 | 211.00 | 215.33 | 234.00 |
| TBC1D20 | 1591.00 | 1620.00 | 1751.00 | 1654.00 | 1319.00 |
| TBC1D22A | 423.00 | 425.00 | 476.00 | 441.33 | 465.00 |
| TBC1D22B | 316.00 | 334.00 | 380.00 | 343.33 | 459.00 |
| TBC1D23 | 1906.00 | 1651.00 | 1873.00 | 1810.00 | 2902.00 |
| TBC1D24 | 839.00 | 797.00 | 828.00 | 821.33 | 775.00 |
| TBC1D25 | 374.00 | 399.00 | 401.00 | 391.33 | 452.00 |
| TBC1D2B | 1407.00 | 1417.00 | 1547.00 | 1457.00 | 966.00 |
| TBC1D30 | 177.00 | 178.00 | 181.00 | 178.67 | 207.00 |
| TBC1D31 | 466.00 | 398.00 | 469.00 | 444.33 | 518.00 |
| TBC1D32 | 287.00 | 289.00 | 349.00 | 308.33 | 303.00 |
| ΓBC1D3P1-DHX40P1 | 3.00 | 3.00 | 2.00 | 2.67 | 1.00 |
| TBC1D4 | 878.00 | 816.00 | 949.00 | 881.00 | 1013.00 |
| TBC1D5 | 2464.00 | 2231.00 | 2644.00 | 2446.33 | 2219.00 |
| TBC1D7 | 236.00 | 209.00 | 232.00 | 225.67 | 214.00 |
| TBC1D8 | 663.00 | 618.00 | 722.00 | 667.67 | 515.00 |
| TBC1D8B | 370.00 | 322.00 | 419.00 | 370.33 | 344.00 |
| TBC1D9 | 818.00 | 716.00 | 856.00 | 796.67 | 763.00 |
| TBC1D9B | 4399.00 | 4371.00 | 4793.00 | 4521.00 | 4148.00 |
| TBCA | 1416.00 | 1362.00 | 1428.00 | 1402.00 | 2109.00 |
| TBCB | 992.00 | 949.00 | 992.00 | 977.67 | 1268.00 |
| TBCC | 532.00 | 503.00 | 547.00 | 527.33 | 665.00 |
| TBCCD1 | 250.00 | 243.00 | 233.00 | 242.00 | 273.00 |
| TBCD | 985.00 | 930.00 | 997.00 | 970.67 | 850.00 |
| TBCE | 688.00 | 671.00 | 695.00 | 684.67 | 888.00 |
| TBCEL | 600.00 | 636.00 | 670.00 | 635.33 | 621.00 |
| TBCK | 551.00 | 569.00 | 650.00 | 590.00 | 498.00 |
| TBK1 | 883.00 | 882.00 | 1002.00 | 922.33 | 1123.00 |
| TBKBP1 | 123.00 | 146.00 | 160.00 | 143.00 | 131.00 |
| TBL1X | 815.00 | 749.00 | 875.00 | 813.00 | 780.00 |

| | | | | | |
|---|---|---|---|---|---|
| TBL1XR1 | 2199.00 | 2071.00 | 2557.00 | 2275.67 | 2686.00 |
| TBL2 | 1371.00 | 1373.00 | 1256.00 | 1333.33 | 1439.00 |
| TBL3 | 1100.00 | 1100.00 | 1033.00 | 1077.67 | 1182.00 |
| TBP | 390.00 | 363.00 | 403.00 | 385.33 | 496.00 |
| TBPL1 | 564.00 | 527.00 | 573.00 | 554.67 | 752.00 |
| TBRG1 | 1139.00 | 1093.00 | 1203.00 | 1145.00 | 1116.00 |
| TBRG4 | 3438.00 | 3331.00 | 3642.00 | 3470.33 | 3762.00 |
| TBX18 | 848.00 | 840.00 | 935.00 | 874.33 | 883.00 |
| TBX19 | 82.00 | 85.00 | 95.00 | 87.33 | 80.00 |
| TBX2 | 348.00 | 385.00 | 362.00 | 365.00 | 371.00 |
| TBX2-AS1 | 74.00 | 114.00 | 104.00 | 97.33 | 97.00 |
| TBX3 | 2829.00 | 2856.00 | 3208.00 | 2964.33 | 2264.00 |
| TBX6 | 34.00 | 51.00 | 52.00 | 45.67 | 54.00 |
| TBXA2R | 24.00 | 24.00 | 27.00 | 25.00 | 22.00 |
| TBXAS1 | 223.00 | 214.00 | 232.00 | 223.00 | 218.00 |
| TCAF1 | 1299.00 | 1322.00 | 1508.00 | 1376.33 | 1084.00 |
| TCAF2 | 47.00 | 57.00 | 43.00 | 49.00 | 77.00 |
| TCAIM | 585.00 | 513.00 | 605.00 | 567.67 | 562.00 |
| TCAM1P | 36.00 | 41.00 | 39.00 | 38.67 | 51.00 |
| TCAP | 14.00 | 18.00 | 13.00 | 15.00 | 3.00 |
| TCEA1 | 3286.00 | 2997.00 | 3717.00 | 3333.33 | 4183.00 |
| TCEA2 | 449.00 | 440.00 | 517.00 | 468.67 | 412.00 |
| TCEA3 | 4.00 | 6.00 | 10.00 | 6.67 | 12.00 |
| TCEAL1 | 448.00 | 459.00 | 461.00 | 456.00 | 391.00 |
| TCEAL3 | 385.00 | 404.00 | 414.00 | 401.00 | 398.00 |
| TCEAL4 | 2084.00 | 2140.00 | 2240.00 | 2154.67 | 2609.00 |
| TCEAL8 | 1239.00 | 1206.00 | 1374.00 | 1273.00 | 1644.00 |
| TCEANC | 59.00 | 80.00 | 98.00 | 79.00 | 87.00 |
| TCEANC2 | 504.00 | 438.00 | 444.00 | 462.00 | 400.00 |
| TCEB1 | 1255.00 | 1174.00 | 1307.00 | 1245.33 | 1683.00 |
| TCEB2 | 1295.00 | 1260.00 | 1344.00 | 1299.67 | 1641.00 |
| TCEB3 | 1958.00 | 1873.00 | 2086.00 | 1972.33 | 1997.00 |
| TCEB3-AS1 | 78.00 | 101.00 | 90.00 | 89.67 | 96.00 |
| TCERG1 | 3118.00 | 3083.00 | 3222.00 | 3141.00 | 3455.00 |
| TCF12 | 1910.00 | 1577.00 | 2015.00 | 1834.00 | 1886.00 |
| TCF24 | 10.00 | 12.00 | 13.00 | 11.67 | 11.00 |
| TCF25 | 2591.00 | 2545.00 | 2728.00 | 2621.33 | 2558.00 |
| TCF3 | 2310.00 | 2229.00 | 2437.00 | 2325.33 | 2734.00 |
| TCF4 | 378.00 | 313.00 | 359.00 | 350.00 | 319.00 |
| TCF7 | 29.00 | 31.00 | 31.00 | 30.33 | 39.00 |
| TCF7L1 | 5.00 | 7.00 | 14.00 | 8.67 | 12.00 |
| TCF7L2 | 513.00 | 491.00 | 559.00 | 521.00 | 519.00 |
| TCFL5 | 889.00 | 942.00 | 1030.00 | 953.67 | 1157.00 |
| TCHP | 885.00 | 928.00 | 934.00 | 915.67 | 1035.00 |
| TCIRG1 | 1082.00 | 1136.00 | 1106.00 | 1108.00 | 1014.00 |
| TCN1 | 3.00 | 2.00 | 2.00 | 2.33 | 15.00 |
| TCN2 | 43.00 | 72.00 | 55.00 | 56.67 | 138.00 |
| TCOF1 | 4253.00 | 4180.00 | 4323.00 | 4252.00 | 4229.00 |

| | | | | | |
|---|---|---|---|---|---|
| TCONS_00029157 | 4.00 | 6.00 | 6.00 | 5.33 | 3.00 |
| TCP1 | 5651.00 | 5363.00 | 5868.00 | 5627.33 | 7730.00 |
| TCP11 | 8.00 | 5.00 | 6.00 | 6.33 | 11.00 |
| TCP11L1 | 328.00 | 276.00 | 268.00 | 290.67 | 273.00 |
| TCP11L2 | 127.00 | 95.00 | 91.00 | 104.33 | 74.00 |
| TCTA | 436.00 | 392.00 | 434.00 | 420.67 | 382.00 |
| TCTEX1D2 | 192.00 | 179.00 | 207.00 | 192.67 | 237.00 |
| TCTN1 | 1074.00 | 930.00 | 998.00 | 1000.67 | 983.00 |
| TCTN2 | 211.00 | 214.00 | 206.00 | 210.33 | 193.00 |
| TCTN3 | 1823.00 | 1653.00 | 1731.00 | 1735.67 | 1746.00 |
| TDG | 496.00 | 534.00 | 581.00 | 537.00 | 544.00 |
| TDP1 | 921.00 | 880.00 | 967.00 | 922.67 | 842.00 |
| TDP2 | 1017.00 | 974.00 | 1113.00 | 1034.67 | 1278.00 |
| TDRD3 | 272.00 | 316.00 | 361.00 | 316.33 | 354.00 |
| TDRD6 | 31.00 | 50.00 | 39.00 | 40.00 | 29.00 |
| TDRD7 | 115.00 | 122.00 | 120.00 | 119.00 | 127.00 |
| TDRKH | 217.00 | 224.00 | 240.00 | 227.00 | 318.00 |
| TDRP | 135.00 | 124.00 | 140.00 | 133.00 | 127.00 |
| TEAD1 | 2812.00 | 2553.00 | 3058.00 | 2807.67 | 2071.00 |
| TEAD2 | 78.00 | 85.00 | 86.00 | 83.00 | 96.00 |
| TEAD3 | 359.00 | 341.00 | 345.00 | 348.33 | 321.00 |
| TEAD4 | 733.00 | 812.00 | 919.00 | 821.33 | 916.00 |
| TEC | 125.00 | 108.00 | 120.00 | 117.67 | 93.00 |
| TECPR1 | 1008.00 | 998.00 | 1124.00 | 1043.33 | 1047.00 |
| TECPR2 | 884.00 | 839.00 | 914.00 | 879.00 | 680.00 |
| TECR | 2197.00 | 2355.00 | 1978.00 | 2176.67 | 2631.00 |
| TECTA | 10.00 | 21.00 | 27.00 | 19.33 | 23.00 |
| TECTB | 4.00 | 5.00 | 4.00 | 4.33 | 2.00 |
| TEF | 110.00 | 97.00 | 121.00 | 109.33 | 125.00 |
| TEFM | 208.00 | 223.00 | 214.00 | 215.00 | 222.00 |
| TEKT4P2 | 60.00 | 45.00 | 51.00 | 52.00 | 46.00 |
| TELO2 | 1047.00 | 1078.00 | 1066.00 | 1063.67 | 1163.00 |
| TEN1-CDK3 | 51.00 | 71.00 | 61.00 | 61.00 | 68.00 |
| TENM1 | 375.00 | 354.00 | 438.00 | 389.00 | 260.00 |
| TENM2 | 1081.00 | 893.00 | 1087.00 | 1020.33 | 665.00 |
| TENM3 | 1223.00 | 1183.00 | 1272.00 | 1226.00 | 1153.00 |
| TENM4 | 261.00 | 199.00 | 284.00 | 248.00 | 170.00 |
| TEP1 | 1493.00 | 1369.00 | 1477.00 | 1446.33 | 1200.00 |
| TERF1 | 574.00 | 550.00 | 616.00 | 580.00 | 676.00 |
| TERF2 | 671.00 | 642.00 | 736.00 | 683.00 | 784.00 |
| TERF2IP | 1064.00 | 1006.00 | 1113.00 | 1061.00 | 1350.00 |
| TERT | 220.00 | 238.00 | 235.00 | 231.00 | 281.00 |
| TES | 345.00 | 358.00 | 341.00 | 348.00 | 317.00 |
| TESK1 | 461.00 | 515.00 | 531.00 | 502.33 | 481.00 |
| TESK2 | 86.00 | 83.00 | 108.00 | 92.33 | 60.00 |
| TET1 | 65.00 | 63.00 | 55.00 | 61.00 | 58.00 |
| TET2 | 316.00 | 268.00 | 321.00 | 301.67 | 293.00 |
| TET3 | 1277.00 | 1143.00 | 1396.00 | 1272.00 | 1062.00 |

| | | | | | |
|---|---|---|---|---|---|
| TEX10 | 1423.00 | 1251.00 | 1417.00 | 1363.67 | 1641.00 |
| TEX12 | 5.00 | 4.00 | 6.00 | 5.00 | 5.00 |
| TEX14 | 7.00 | 4.00 | 7.00 | 6.00 | 8.00 |
| TEX15 | 63.00 | 66.00 | 98.00 | 75.67 | 128.00 |
| TEX19 | 33.00 | 31.00 | 40.00 | 34.67 | 18.00 |
| TEX2 | 1365.00 | 1283.00 | 1315.00 | 1321.00 | 1251.00 |
| TEX21P | 24.00 | 14.00 | 16.00 | 18.00 | 14.00 |
| TEX22 | 33.00 | 32.00 | 31.00 | 32.00 | 44.00 |
| TEX261 | 2837.00 | 2799.00 | 2856.00 | 2830.67 | 2680.00 |
| TEX264 | 1294.00 | 1296.00 | 1232.00 | 1274.00 | 1353.00 |
| TEX30 | 344.00 | 344.00 | 374.00 | 354.00 | 541.00 |
| TEX41 | 16.00 | 13.00 | 16.00 | 15.00 | 7.00 |
| TEX9 | 70.00 | 50.00 | 79.00 | 66.33 | 101.00 |
| TF | 3.00 | 3.00 | 4.00 | 3.33 | 1.00 |
| TFAM | 1773.00 | 1627.00 | 1712.00 | 1704.00 | 2203.00 |
| TFAP2A | 338.00 | 320.00 | 385.00 | 347.67 | 386.00 |
| TFAP2A-AS1 | 19.00 | 20.00 | 23.00 | 20.67 | 15.00 |
| TFAP2C | 287.00 | 313.00 | 308.00 | 302.67 | 315.00 |
| TFAP2E | 7.00 | 10.00 | 9.00 | 8.67 | 3.00 |
| TFAP4 | 653.00 | 686.00 | 771.00 | 703.33 | 815.00 |
| TFB1M | 526.00 | 518.00 | 550.00 | 531.33 | 638.00 |
| TFB2M | 660.00 | 619.00 | 738.00 | 672.33 | 825.00 |
| TFCP2 | 900.00 | 913.00 | 958.00 | 923.67 | 1086.00 |
| TFDP1 | 1745.00 | 1660.00 | 1859.00 | 1754.67 | 2295.00 |
| TFDP2 | 787.00 | 724.00 | 809.00 | 773.33 | 878.00 |
| TFE3 | 2476.00 | 2442.00 | 2426.00 | 2448.00 | 2817.00 |
| TFEB | 12.00 | 13.00 | 11.00 | 12.00 | 7.00 |
| TFG | 2832.00 | 2696.00 | 2940.00 | 2822.67 | 3064.00 |
| TFIP11 | 910.00 | 931.00 | 966.00 | 935.67 | 908.00 |
| TFPI | 17535.00 | 17024.00 | 18755.00 | 17771.33 | 14801.00 |
| TFPI2 | 5412.00 | 5301.00 | 5137.00 | 5283.33 | 7130.00 |
| TFPT | 10.00 | 7.00 | 11.00 | 9.33 | 5.00 |
| TFR2 | 44.00 | 65.00 | 54.00 | 54.33 | 51.00 |
| TFRC | 6775.00 | 6402.00 | 6537.00 | 6571.33 | 5802.00 |
| TG | 4.00 | 4.00 | 12.00 | 6.67 | 6.00 |
| TGDS | 308.00 | 289.00 | 328.00 | 308.33 | 401.00 |
| TGFB1 | 1687.00 | 1764.00 | 1675.00 | 1708.67 | 1501.00 |
| TGFB1I1 | 1096.00 | 1065.00 | 1083.00 | 1081.33 | 1116.00 |
| TGFB2 | 544.00 | 457.00 | 496.00 | 499.00 | 361.00 |
| TGFB2-AS1 | 18.00 | 7.00 | 15.00 | 13.33 | 18.00 |
| TGFB3 | 13.00 | 23.00 | 8.00 | 14.67 | 16.00 |
| TGFBI | 2113.00 | 1991.00 | 2307.00 | 2137.00 | 1176.00 |
| TGFBR1 | 652.00 | 570.00 | 632.00 | 618.00 | 620.00 |
| TGFBR2 | 7410.00 | 6669.00 | 7245.00 | 7108.00 | 6374.00 |
| TGFBR3 | 298.00 | 322.00 | 346.00 | 322.00 | 268.00 |
| TGFBR3L | 17.00 | 16.00 | 36.00 | 23.00 | 26.00 |
| TGFBRAP1 | 1050.00 | 981.00 | 1090.00 | 1040.33 | 1129.00 |
| TGIF1 | 1100.00 | 1069.00 | 1256.00 | 1141.67 | 1058.00 |

| | | | | | |
|---|---|---|---|---|---|
| TGIF2 | 462.00 | 497.00 | 445.00 | 468.00 | 767.00 |
| TGM1 | 31.00 | 39.00 | 39.00 | 36.33 | 23.00 |
| TGM2 | 18265.00 | 17660.00 | 19049.00 | 18324.67 | 18381.00 |
| TGM4 | 5.00 | 5.00 | 8.00 | 6.00 | 1.00 |
| TGOLN2 | 9540.00 | 8718.00 | 9347.00 | 9201.67 | 6681.00 |
| TGS1 | 806.00 | 777.00 | 834.00 | 805.67 | 834.00 |
| THADA | 1656.00 | 1665.00 | 1781.00 | 1700.67 | 1567.00 |
| THAP1 | 202.00 | 228.00 | 251.00 | 227.00 | 264.00 |
| THAP10 | 147.00 | 129.00 | 132.00 | 136.00 | 142.00 |
| THAP11 | 630.00 | 574.00 | 630.00 | 611.33 | 707.00 |
| THAP2 | 148.00 | 126.00 | 144.00 | 139.33 | 150.00 |
| THAP3 | 363.00 | 338.00 | 392.00 | 364.33 | 404.00 |
| THAP4 | 1227.00 | 1246.00 | 1284.00 | 1252.33 | 1475.00 |
| THAP5 | 369.00 | 427.00 | 439.00 | 411.67 | 516.00 |
| THAP6 | 221.00 | 223.00 | 272.00 | 238.67 | 315.00 |
| THAP7 | 251.00 | 272.00 | 277.00 | 266.67 | 316.00 |
| THAP7-AS1 | 40.00 | 34.00 | 37.00 | 37.00 | 36.00 |
| THAP8 | 71.00 | 48.00 | 51.00 | 56.67 | 69.00 |
| THAP9 | 95.00 | 83.00 | 100.00 | 92.67 | 75.00 |
| THAP9-AS1 | 1211.00 | 1210.00 | 1394.00 | 1271.67 | 1626.00 |
| THBS1 | 1680.00 | 1556.00 | 1805.00 | 1680.33 | 1641.00 |
| THBS2 | 5.00 | 7.00 | 16.00 | 9.33 | 8.00 |
| THBS3 | 861.00 | 841.00 | 935.00 | 879.00 | 991.00 |
| THBS4 | 24.00 | 22.00 | 25.00 | 23.67 | 18.00 |
| THCAT158 | 7.00 | 3.00 | 8.00 | 6.00 | 8.00 |
| THEGL | 22.00 | 16.00 | 19.00 | 19.00 | 4.00 |
| THEM4 | 509.00 | 526.00 | 580.00 | 538.33 | 576.00 |
| THEM6 | 976.00 | 1123.00 | 1176.00 | 1091.67 | 1170.00 |
| THEMIS2 | 16.00 | 28.00 | 30.00 | 24.67 | 8.00 |
| THG1L | 306.00 | 377.00 | 395.00 | 359.33 | 457.00 |
| THNSL1 | 712.00 | 667.00 | 744.00 | 707.67 | 730.00 |
| THNSL2 | 83.00 | 90.00 | 76.00 | 83.00 | 74.00 |
| THOC1 | 2521.00 | 2626.00 | 2761.00 | 2636.00 | 3040.00 |
| THOC2 | 1671.00 | 1489.00 | 1694.00 | 1618.00 | 1548.00 |
| THOC3 | 255.00 | 282.00 | 252.00 | 263.00 | 316.00 |
| THOC5 | 1239.00 | 1157.00 | 1325.00 | 1240.33 | 1413.00 |
| THOC6 | 774.00 | 814.00 | 794.00 | 794.00 | 1033.00 |
| THOC7 | 900.00 | 873.00 | 908.00 | 893.67 | 1154.00 |
| THOP1 | 2029.00 | 2029.00 | 2115.00 | 2057.67 | 2313.00 |
| THRA | 270.00 | 275.00 | 327.00 | 290.67 | 294.00 |
| THRAP3 | 4020.00 | 3883.00 | 4189.00 | 4030.67 | 4161.00 |
| THRSP | 3.00 | 1.00 | 2.00 | 2.00 | 1.00 |
| THSD1 | 96.00 | 73.00 | 110.00 | 93.00 | 124.00 |
| THSD4 | 1933.00 | 1823.00 | 2144.00 | 1966.67 | 1110.00 |
| THSD7A | 3.00 | 4.00 | 5.00 | 4.00 | 3.00 |
| THTPA | 354.00 | 441.00 | 444.00 | 413.00 | 304.00 |
| THUMPD1 | 978.00 | 918.00 | 1064.00 | 986.67 | 1147.00 |
| THUMPD2 | 449.00 | 452.00 | 571.00 | 490.67 | 541.00 |

| | | | | | |
|---|---|---|---|---|---|
| THUMPD3 | 968.00 | 901.00 | 1047.00 | 972.00 | 1059.00 |
| THUMPD3-AS1 | 575.00 | 541.00 | 610.00 | 575.33 | 525.00 |
| THY1 | 825.00 | 767.00 | 684.00 | 758.67 | 736.00 |
| THYN1 | 388.00 | 408.00 | 405.00 | 400.33 | 472.00 |
| TIA1 | 2450.00 | 2562.00 | 2794.00 | 2602.00 | 2467.00 |
| TIAF1 | 18.00 | 13.00 | 19.00 | 16.67 | 5.00 |
| TIAL1 | 3884.00 | 3938.00 | 4266.00 | 4029.33 | 4727.00 |
| TIAM1 | 404.00 | 321.00 | 432.00 | 385.67 | 331.00 |
| TIAM2 | 24.00 | 20.00 | 34.00 | 26.00 | 15.00 |
| TICAM1 | 389.00 | 400.00 | 449.00 | 412.67 | 458.00 |
| TICAM2 | 3.00 | 2.00 | 2.00 | 2.33 | 1.00 |
| TICRR | 578.00 | 524.00 | 513.00 | 538.33 | 612.00 |
| TIE1 | 14927.00 | 15295.00 | 14959.00 | 15060.33 | 13062.00 |
| TIFA | 192.00 | 171.00 | 174.00 | 179.00 | 243.00 |
| TIGAR | 368.00 | 332.00 | 372.00 | 357.33 | 448.00 |
| TIGD1 | 272.00 | 263.00 | 322.00 | 285.67 | 288.00 |
| TIGD2 | 182.00 | 151.00 | 203.00 | 178.67 | 190.00 |
| TIGD3 | 6.00 | 6.00 | 10.00 | 7.33 | 10.00 |
| TIGD4 | 18.00 | 25.00 | 27.00 | 23.33 | 24.00 |
| TIGD6 | 156.00 | 160.00 | 161.00 | 159.00 | 166.00 |
| TIGD7 | 21.00 | 26.00 | 20.00 | 22.33 | 28.00 |
| TIMELESS | 1264.00 | 1233.00 | 1250.00 | 1249.00 | 1518.00 |
| TIMM10 | 407.00 | 383.00 | 437.00 | 409.00 | 548.00 |
| TIMM10B | 639.00 | 658.00 | 680.00 | 659.00 | 673.00 |
| TIMM13 | 686.00 | 693.00 | 788.00 | 722.33 | 1019.00 |
| TIMM17A | 919.00 | 913.00 | 886.00 | 906.00 | 1323.00 |
| TIMM17B | 717.00 | 804.00 | 722.00 | 747.67 | 833.00 |
| TIMM21 | 516.00 | 532.00 | 539.00 | 529.00 | 701.00 |
| TIMM23 | 1310.00 | 1273.00 | 1349.00 | 1310.67 | 1809.00 |
| TIMM23B | 391.00 | 333.00 | 422.00 | 382.00 | 358.00 |
| TIMM44 | 1569.00 | 1512.00 | 1637.00 | 1572.67 | 1768.00 |
| TIMM50 | 967.00 | 875.00 | 945.00 | 929.00 | 1091.00 |
| TIMM8A | 215.00 | 182.00 | 231.00 | 209.33 | 245.00 |
| TIMM8B | 304.00 | 319.00 | 311.00 | 311.33 | 428.00 |
| TIMM9 | 575.00 | 630.00 | 642.00 | 615.67 | 843.00 |
| TIMMDC1 | 2100.00 | 1972.00 | 1981.00 | 2017.67 | 2272.00 |
| TIMP1 | 8176.00 | 8339.00 | 7602.00 | 8039.00 | 9081.00 |
| TIMP2 | 6350.00 | 5973.00 | 6010.00 | 6111.00 | 5336.00 |
| TIMP3 | 4655.00 | 4360.00 | 4095.00 | 4370.00 | 3938.00 |
| TINAGL1 | 20.00 | 23.00 | 31.00 | 24.67 | 9.00 |
| TINCR | 13.00 | 33.00 | 20.00 | 22.00 | 22.00 |
| TINF2 | 593.00 | 599.00 | 636.00 | 609.33 | 666.00 |
| TIPARP | 756.00 | 685.00 | 782.00 | 741.00 | 801.00 |
| TIPARP-AS1 | 45.00 | 32.00 | 38.00 | 38.33 | 46.00 |
| TIPIN | 237.00 | 245.00 | 242.00 | 241.33 | 320.00 |
| TIPRL | 1217.00 | 1162.00 | 1244.00 | 1207.67 | 1626.00 |
| TIRAP | 70.00 | 109.00 | 86.00 | 88.33 | 68.00 |
| TJAP1 | 1716.00 | 1629.00 | 1732.00 | 1692.33 | 1736.00 |

| | | | | | |
|---|---|---|---|---|---|
| TJP1 | 85.00 | 83.00 | 79.00 | 82.33 | 70.00 |
| TJP2 | 1593.00 | 1532.00 | 1711.00 | 1612.00 | 2031.00 |
| TK1 | 285.00 | 287.00 | 237.00 | 269.67 | 516.00 |
| TK2 | 1011.00 | 997.00 | 1110.00 | 1039.33 | 1047.00 |
| TKFC | 1154.00 | 1214.00 | 1257.00 | 1208.33 | 1337.00 |
| TKT | 7574.00 | 7314.00 | 7840.00 | 7576.00 | 6870.00 |
| TLCD1 | 115.00 | 125.00 | 98.00 | 112.67 | 145.00 |
| TLDC1 | 2539.00 | 2430.00 | 2623.00 | 2530.67 | 2408.00 |
| TLE1 | 2156.00 | 2114.00 | 2195.00 | 2155.00 | 2315.00 |
| TLE3 | 793.00 | 739.00 | 814.00 | 782.00 | 693.00 |
| TLE4 | 188.00 | 221.00 | 189.00 | 199.33 | 248.00 |
| TLK1 | 898.00 | 761.00 | 1026.00 | 895.00 | 1025.00 |
| TLK2 | 1060.00 | 1051.00 | 1168.00 | 1093.00 | 1130.00 |
| TLL1 | 7.00 | 2.00 | 5.00 | 4.67 | 4.00 |
| TLL2 | 3.00 | 1.00 | 5.00 | 3.00 | 3.00 |
| TLN1 | 19645.00 | 18730.00 | 20019.00 | 19464.67 | 12875.00 |
| TLN2 | 285.00 | 262.00 | 303.00 | 283.33 | 388.00 |
| TLR1 | 16.00 | 17.00 | 24.00 | 19.00 | 22.00 |
| TLR3 | 15.00 | 26.00 | 18.00 | 19.67 | 24.00 |
| TLR4 | 48.00 | 57.00 | 55.00 | 53.33 | 9.00 |
| TLR6 | 445.00 | 428.00 | 485.00 | 452.67 | 552.00 |
| TLR9 | 5.00 | 1.00 | 2.00 | 2.67 | 1.00 |
| TM2D1 | 390.00 | 366.00 | 391.00 | 382.33 | 457.00 |
| TM2D2 | 945.00 | 929.00 | 908.00 | 927.33 | 1018.00 |
| TM2D3 | 541.00 | 494.00 | 532.00 | 522.33 | 656.00 |
| TM4SF1 | 11375.00 | 10737.00 | 9965.00 | 10692.33 | 25975.00 |
| TM4SF18 | 364.00 | 314.00 | 319.00 | 332.33 | 480.00 |
| TM4SF19 | 58.00 | 64.00 | 72.00 | 64.67 | 32.00 |
| TM4SF19-AS1 | 24.00 | 24.00 | 23.00 | 23.67 | 40.00 |
| TM4SF1-AS1 | 5.00 | 3.00 | 3.00 | 3.67 | 10.00 |
| TM6SF1 | 9.00 | 10.00 | 14.00 | 11.00 | 3.00 |
| TM7SF2 | 215.00 | 233.00 | 231.00 | 226.33 | 224.00 |
| TM7SF3 | 2903.00 | 2792.00 | 2952.00 | 2882.33 | 3571.00 |
| TM9SF1 | 2645.00 | 2601.00 | 2644.00 | 2630.00 | 2630.00 |
| TM9SF2 | 4427.00 | 4217.00 | 4520.00 | 4388.00 | 4263.00 |
| TM9SF3 | 9624.00 | 8955.00 | 9711.00 | 9430.00 | 10531.00 |
| TM9SF4 | 4727.00 | 4657.00 | 4800.00 | 4728.00 | 4180.00 |
| TMA16 | 401.00 | 399.00 | 425.00 | 408.33 | 597.00 |
| TMA7 | 983.00 | 1020.00 | 1103.00 | 1035.33 | 1548.00 |
| TMBIM1 | 2820.00 | 2610.00 | 2700.00 | 2710.00 | 2921.00 |
| TMBIM4 | 1208.00 | 1155.00 | 1172.00 | 1178.33 | 1453.00 |
| TMBIM6 | 28741.00 | 27001.00 | 27334.00 | 27692.00 | 25434.00 |
| TMC3 | 3.00 | 6.00 | 5.00 | 4.67 | 1.00 |
| TMC3-AS1 | 25.00 | 31.00 | 22.00 | 26.00 | 23.00 |
| TMC6 | 42.00 | 35.00 | 36.00 | 37.67 | 32.00 |
| TMC7 | 245.00 | 262.00 | 270.00 | 259.00 | 364.00 |
| TMC8 | 6.00 | 5.00 | 1.00 | 4.00 | 0.00 |
| TMCC1 | 407.00 | 379.00 | 468.00 | 418.00 | 466.00 |

| | | | | | |
|---|---|---|---|---|---|
| TMCC1-AS1 | 668.00 | 675.00 | 706.00 | 683.00 | 769.00 |
| TMCC2 | 73.00 | 62.00 | 62.00 | 65.67 | 86.00 |
| TMCC3 | 334.00 | 277.00 | 346.00 | 319.00 | 290.00 |
| TMCO1 | 2405.00 | 2300.00 | 2270.00 | 2325.00 | 2795.00 |
| TMCO3 | 1002.00 | 976.00 | 1010.00 | 996.00 | 814.00 |
| TMCO4 | 263.00 | 268.00 | 323.00 | 284.67 | 329.00 |
| TMCO6 | 669.00 | 636.00 | 676.00 | 660.33 | 618.00 |
| TMED1 | 814.00 | 852.00 | 764.00 | 810.00 | 832.00 |
| TMED10 | 6545.00 | 6094.00 | 6225.00 | 6288.00 | 6632.00 |
| TMED2 | 4304.00 | 4072.00 | 4161.00 | 4179.00 | 4677.00 |
| TMED3 | 1727.00 | 1707.00 | 1655.00 | 1696.33 | 2243.00 |
| TMED4 | 3330.00 | 3399.00 | 3285.00 | 3338.00 | 3569.00 |
| TMED5 | 2505.00 | 2344.00 | 2725.00 | 2524.67 | 2603.00 |
| TMED6 | 17.00 | 17.00 | 6.00 | 13.33 | 8.00 |
| TMED7 | 1278.00 | 1188.00 | 1300.00 | 1255.33 | 1384.00 |
| TMED7-TICAM2 | 11.00 | 16.00 | 17.00 | 14.67 | 9.00 |
| TMED8 | 66.00 | 45.00 | 57.00 | 56.00 | 49.00 |
| TMED9 | 5304.00 | 5613.00 | 5099.00 | 5338.67 | 5554.00 |
| TMEFF2 | 40.00 | 20.00 | 23.00 | 27.67 | 25.00 |
| TMEM101 | 1283.00 | 1264.00 | 1315.00 | 1287.33 | 1544.00 |
| TMEM102 | 65.00 | 50.00 | 48.00 | 54.33 | 43.00 |
| TMEM104 | 846.00 | 910.00 | 852.00 | 869.33 | 712.00 |
| TMEM106A | 167.00 | 181.00 | 162.00 | 170.00 | 178.00 |
| TMEM106B | 2168.00 | 1844.00 | 2135.00 | 2049.00 | 1844.00 |
| TMEM106C | 1181.00 | 1106.00 | 1196.00 | 1161.00 | 1430.00 |
| TMEM107 | 498.00 | 494.00 | 428.00 | 473.33 | 558.00 |
| TMEM108 | 197.00 | 175.00 | 200.00 | 190.67 | 183.00 |
| TMEM108-AS1 | 380.00 | 404.00 | 342.00 | 375.33 | 316.00 |
| TMEM109 | 3000.00 | 3002.00 | 2779.00 | 2927.00 | 3539.00 |
| TMEM11 | 338.00 | 329.00 | 374.00 | 347.00 | 406.00 |
| TMEM110 | 617.00 | 621.00 | 662.00 | 633.33 | 530.00 |
| TMEM115 | 1891.00 | 1993.00 | 1919.00 | 1934.33 | 1860.00 |
| TMEM116 | 218.00 | 203.00 | 209.00 | 210.00 | 294.00 |
| TMEM117 | 61.00 | 88.00 | 91.00 | 80.00 | 63.00 |
| TMEM120A | 421.00 | 416.00 | 436.00 | 424.33 | 522.00 |
| TMEM120B | 2049.00 | 2178.00 | 2163.00 | 2130.00 | 1795.00 |
| TMEM121 | 11.00 | 12.00 | 26.00 | 16.33 | 30.00 |
| TMEM123 | 14927.00 | 13162.00 | 14887.00 | 14325.33 | 13421.00 |
| TMEM126A | 250.00 | 238.00 | 259.00 | 249.00 | 302.00 |
| TMEM126B | 576.00 | 514.00 | 629.00 | 573.00 | 737.00 |
| TMEM127 | 1702.00 | 1676.00 | 1644.00 | 1674.00 | 1492.00 |
| TMEM128 | 280.00 | 304.00 | 303.00 | 295.67 | 366.00 |
| TMEM129 | 6501.00 | 6620.00 | 6398.00 | 6506.33 | 7167.00 |
| TMEM131 | 2968.00 | 2841.00 | 3076.00 | 2961.67 | 2354.00 |
| TMEM132A | 2418.00 | 2431.00 | 2412.00 | 2420.33 | 2532.00 |
| TMEM132B | 4.00 | 8.00 | 1.00 | 4.33 | 4.00 |
| TMEM133 | 488.00 | 473.00 | 525.00 | 495.33 | 523.00 |
| TMEM134 | 203.00 | 199.00 | 239.00 | 213.67 | 260.00 |

| | | | | | |
|---|---|---|---|---|---|
| TMEM135 | 284.00 | 219.00 | 310.00 | 271.00 | 289.00 |
| TMEM136 | 470.00 | 431.00 | 503.00 | 468.00 | 528.00 |
| TMEM138 | 861.00 | 864.00 | 900.00 | 875.00 | 1000.00 |
| TMEM140 | 30.00 | 29.00 | 41.00 | 33.33 | 32.00 |
| TMEM141 | 323.00 | 320.00 | 351.00 | 331.33 | 443.00 |
| TMEM143 | 118.00 | 122.00 | 139.00 | 126.33 | 134.00 |
| TMEM144 | 149.00 | 118.00 | 143.00 | 136.67 | 230.00 |
| TMEM145 | 24.00 | 26.00 | 27.00 | 25.67 | 29.00 |
| TMEM147 | 1430.00 | 1419.00 | 1487.00 | 1445.33 | 1694.00 |
| TMEM147-AS1 | 950.00 | 1095.00 | 1067.00 | 1037.33 | 1209.00 |
| TMEM14A | 250.00 | 234.00 | 213.00 | 232.33 | 407.00 |
| TMEM14B | 1663.00 | 1558.00 | 1491.00 | 1570.67 | 2162.00 |
| TMEM14C | 2168.00 | 2164.00 | 2160.00 | 2164.00 | 2734.00 |
| TMEM150A | 419.00 | 433.00 | 444.00 | 432.00 | 434.00 |
| TMEM150C | 16.00 | 22.00 | 16.00 | 18.00 | 33.00 |
| TMEM154 | 595.00 | 478.00 | 504.00 | 525.67 | 466.00 |
| TMEM156 | 3708.00 | 3438.00 | 3604.00 | 3583.33 | 3463.00 |
| TMEM158 | 1934.00 | 2034.00 | 2193.00 | 2053.67 | 3125.00 |
| TMEM159 | 385.00 | 330.00 | 351.00 | 355.33 | 330.00 |
| TMEM160 | 169.00 | 129.00 | 181.00 | 159.67 | 240.00 |
| TMEM161A | 1271.00 | 1375.00 | 1337.00 | 1327.67 | 1234.00 |
| TMEM161B | 1249.00 | 1227.00 | 1368.00 | 1281.33 | 1200.00 |
| TMEM161B-AS1 | 257.00 | 289.00 | 282.00 | 276.00 | 335.00 |
| TMEM164 | 1643.00 | 1529.00 | 1689.00 | 1620.33 | 1352.00 |
| TMEM165 | 1696.00 | 1545.00 | 1674.00 | 1638.33 | 1789.00 |
| TMEM167A | 1823.00 | 1685.00 | 2015.00 | 1841.00 | 2172.00 |
| TMEM167B | 842.00 | 834.00 | 984.00 | 886.67 | 1120.00 |
| TMEM168 | 645.00 | 580.00 | 605.00 | 610.00 | 654.00 |
| TMEM169 | 7.00 | 2.00 | 4.00 | 4.33 | 2.00 |
| TMEM17 | 237.00 | 255.00 | 231.00 | 241.00 | 312.00 |
| TMEM170A | 813.00 | 710.00 | 833.00 | 785.33 | 883.00 |
| TMEM170B | 187.00 | 159.00 | 224.00 | 190.00 | 274.00 |
| TMEM171 | 72.00 | 75.00 | 82.00 | 76.33 | 128.00 |
| TMEM173 | 456.00 | 381.00 | 377.00 | 404.67 | 526.00 |
| TMEM175 | 440.00 | 460.00 | 438.00 | 446.00 | 535.00 |
| TMEM177 | 362.00 | 364.00 | 345.00 | 357.00 | 438.00 |
| TMEM178A | 44.00 | 32.00 | 47.00 | 41.00 | 35.00 |
| TMEM179B | 623.00 | 615.00 | 562.00 | 600.00 | 703.00 |
| TMEM18 | 682.00 | 728.00 | 749.00 | 719.67 | 919.00 |
| TMEM180 | 320.00 | 357.00 | 366.00 | 347.67 | 278.00 |
| TMEM181 | 3478.00 | 3577.00 | 3739.00 | 3598.00 | 3718.00 |
| TMEM182 | 115.00 | 98.00 | 121.00 | 111.33 | 169.00 |
| TMEM183A | 487.00 | 496.00 | 584.00 | 522.33 | 558.00 |
| TMEM183B | 43.00 | 20.00 | 54.00 | 39.00 | 43.00 |
| TMEM184A | 20.00 | 20.00 | 28.00 | 22.67 | 24.00 |
| TMEM184B | 1864.00 | 1844.00 | 1743.00 | 1817.00 | 1743.00 |
| TMEM184C | 800.00 | 745.00 | 807.00 | 784.00 | 739.00 |
| TMEM185A | 212.00 | 195.00 | 189.00 | 198.67 | 199.00 |

| | | | | | |
|---|---|---|---|---|---|
| TMEM185B | 3737.00 | 3444.00 | 3583.00 | 3588.00 | 3777.00 |
| TMEM186 | 368.00 | 373.00 | 388.00 | 376.33 | 409.00 |
| TMEM187 | 287.00 | 294.00 | 268.00 | 283.00 | 322.00 |
| TMEM189 | 646.00 | 686.00 | 584.00 | 638.67 | 817.00 |
| TMEM19 | 601.00 | 541.00 | 558.00 | 566.67 | 647.00 |
| TMEM191A | 3.00 | 6.00 | 2.00 | 3.67 | 5.00 |
| TMEM191C | 4.00 | 3.00 | 2.00 | 3.00 | 0.00 |
| TMEM192 | 805.00 | 742.00 | 824.00 | 790.33 | 777.00 |
| TMEM198 | 162.00 | 157.00 | 166.00 | 161.67 | 111.00 |
| TMEM198B | 1426.00 | 1407.00 | 1580.00 | 1471.00 | 1128.00 |
| TMEM199 | 537.00 | 522.00 | 540.00 | 533.00 | 612.00 |
| TMEM2 | 404.00 | 448.00 | 409.00 | 420.33 | 424.00 |
| TMEM200A | 4366.00 | 4136.00 | 4377.00 | 4293.00 | 3837.00 |
| TMEM200B | 44.00 | 61.00 | 52.00 | 52.33 | 76.00 |
| TMEM201 | 965.00 | 941.00 | 928.00 | 944.67 | 1422.00 |
| TMEM203 | 798.00 | 789.00 | 849.00 | 812.00 | 949.00 |
| TMEM204 | 462.00 | 511.00 | 497.00 | 490.00 | 483.00 |
| TMEM205 | 743.00 | 770.00 | 745.00 | 752.67 | 835.00 |
| TMEM206 | 993.00 | 976.00 | 945.00 | 971.33 | 1054.00 |
| TMEM208 | 632.00 | 663.00 | 540.00 | 611.67 | 718.00 |
| TMEM209 | 2544.00 | 2371.00 | 2660.00 | 2525.00 | 2764.00 |
| TMEM214 | 2669.00 | 2502.00 | 2408.00 | 2526.33 | 1671.00 |
| TMEM216 | 213.00 | 210.00 | 198.00 | 207.00 | 158.00 |
| TMEM217 | 33.00 | 23.00 | 15.00 | 23.67 | 26.00 |
| TMEM218 | 265.00 | 292.00 | 280.00 | 279.00 | 287.00 |
| TMEM219 | 1085.00 | 1120.00 | 1191.00 | 1132.00 | 1359.00 |
| TMEM220 | 15.00 | 13.00 | 21.00 | 16.33 | 15.00 |
| TMEM222 | 713.00 | 767.00 | 791.00 | 757.00 | 815.00 |
| TMEM223 | 232.00 | 265.00 | 230.00 | 242.33 | 303.00 |
| TMEM230 | 2351.00 | 2141.00 | 2456.00 | 2316.00 | 2921.00 |
| TMEM231 | 446.00 | 434.00 | 461.00 | 447.00 | 450.00 |
| TMEM232 | 44.00 | 47.00 | 37.00 | 42.67 | 31.00 |
| TMEM234 | 139.00 | 132.00 | 142.00 | 137.67 | 154.00 |
| TMEM236 | 86.00 | 74.00 | 96.00 | 85.33 | 62.00 |
| TMEM237 | 1098.00 | 1053.00 | 1106.00 | 1085.67 | 1204.00 |
| TMEM238 | 4.00 | 1.00 | 2.00 | 2.33 | 4.00 |
| TMEM240 | 18.00 | 19.00 | 15.00 | 17.33 | 11.00 |
| TMEM241 | 208.00 | 201.00 | 253.00 | 220.67 | 248.00 |
| TMEM242 | 443.00 | 371.00 | 422.00 | 412.00 | 594.00 |
| TMEM243 | 581.00 | 554.00 | 571.00 | 568.67 | 613.00 |
| TMEM244 | 5.00 | 5.00 | 14.00 | 8.00 | 34.00 |
| TMEM245 | 2259.00 | 2095.00 | 2389.00 | 2247.67 | 2269.00 |
| TMEM246 | 1124.00 | 1075.00 | 1090.00 | 1096.33 | 1088.00 |
| TMEM248 | 2570.00 | 2515.00 | 2582.00 | 2555.67 | 2355.00 |
| TMEM249 | 102.00 | 131.00 | 128.00 | 120.33 | 89.00 |
| TMEM25 | 6.00 | 0.00 | 5.00 | 3.67 | 1.00 |
| TMEM254 | 677.00 | 575.00 | 629.00 | 627.00 | 667.00 |
| TMEM254-AS1 | 53.00 | 60.00 | 81.00 | 64.67 | 50.00 |

| | | | | | |
|---|---|---|---|---|---|
| TMEM255B | 710.00 | 687.00 | 638.00 | 678.33 | 343.00 |
| TMEM256 | 374.00 | 356.00 | 437.00 | 389.00 | 474.00 |
| TMEM258 | 816.00 | 745.00 | 851.00 | 804.00 | 1085.00 |
| TMEM259 | 6710.00 | 6776.00 | 6887.00 | 6791.00 | 6897.00 |
| TMEM26 | 21.00 | 19.00 | 10.00 | 16.67 | 17.00 |
| TMEM260 | 531.00 | 557.00 | 568.00 | 552.00 | 626.00 |
| TMEM261 | 1191.00 | 1135.00 | 1205.00 | 1177.00 | 1509.00 |
| TMEM263 | 1177.00 | 1129.00 | 1361.00 | 1222.33 | 1424.00 |
| TMEM265 | 314.00 | 335.00 | 340.00 | 329.67 | 320.00 |
| TMEM27 | 17.00 | 17.00 | 15.00 | 16.33 | 11.00 |
| TMEM30A | 6835.00 | 6201.00 | 6660.00 | 6565.33 | 5138.00 |
| TMEM33 | 6521.00 | 6202.00 | 6809.00 | 6510.67 | 7243.00 |
| TMEM38A | 21.00 | 18.00 | 18.00 | 19.00 | 20.00 |
| TMEM38B | 838.00 | 756.00 | 791.00 | 795.00 | 928.00 |
| TMEM39A | 2029.00 | 1981.00 | 2185.00 | 2065.00 | 1882.00 |
| TMEM39B | 1212.00 | 1127.00 | 1120.00 | 1153.00 | 1252.00 |
| TMEM41A | 1047.00 | 950.00 | 935.00 | 977.33 | 1015.00 |
| TMEM41B | 1128.00 | 1067.00 | 1075.00 | 1090.00 | 1192.00 |
| TMEM42 | 532.00 | 548.00 | 579.00 | 553.00 | 582.00 |
| TMEM43 | 5693.00 | 5705.00 | 5517.00 | 5638.33 | 4571.00 |
| TMEM44 | 1220.00 | 1265.00 | 1289.00 | 1258.00 | 1019.00 |
| TMEM44-AS1 | 120.00 | 87.00 | 114.00 | 107.00 | 124.00 |
| TMEM45A | 175.00 | 197.00 | 200.00 | 190.67 | 181.00 |
| TMEM47 | 5.00 | 4.00 | 4.00 | 4.33 | 6.00 |
| TMEM5 | 863.00 | 882.00 | 917.00 | 887.33 | 1126.00 |
| TMEM50A | 1888.00 | 1858.00 | 1788.00 | 1844.67 | 2367.00 |
| TMEM50B | 6.00 | 1.00 | 13.00 | 6.67 | 2.00 |
| TMEM51 | 213.00 | 210.00 | 205.00 | 209.33 | 191.00 |
| TMEM52 | 3.00 | 0.00 | 3.00 | 2.00 | 3.00 |
| TMEM53 | 78.00 | 64.00 | 93.00 | 78.33 | 83.00 |
| TMEM54 | 458.00 | 462.00 | 459.00 | 459.67 | 541.00 |
| TMEM55A | 398.00 | 327.00 | 390.00 | 371.67 | 510.00 |
| TMEM55B | 581.00 | 553.00 | 633.00 | 589.00 | 596.00 |
| TMEM56 | 31.00 | 29.00 | 29.00 | 29.67 | 48.00 |
| TMEM56-RWDD3 | 50.00 | 29.00 | 44.00 | 41.00 | 22.00 |
| TMEM57 | 806.00 | 824.00 | 753.00 | 794.33 | 814.00 |
| TMEM59 | 2637.00 | 2532.00 | 2568.00 | 2579.00 | 2714.00 |
| TMEM59L | 73.00 | 78.00 | 70.00 | 73.67 | 36.00 |
| TMEM60 | 296.00 | 292.00 | 301.00 | 296.33 | 350.00 |
| TMEM62 | 359.00 | 271.00 | 297.00 | 309.00 | 440.00 |
| TMEM63A | 1136.00 | 1125.00 | 1185.00 | 1148.67 | 952.00 |
| TMEM63B | 1492.00 | 1451.00 | 1337.00 | 1426.67 | 1358.00 |
| TMEM64 | 547.00 | 500.00 | 574.00 | 540.33 | 686.00 |
| TMEM65 | 341.00 | 299.00 | 378.00 | 339.33 | 309.00 |
| TMEM67 | 721.00 | 692.00 | 727.00 | 713.33 | 735.00 |
| TMEM68 | 1058.00 | 980.00 | 1097.00 | 1045.00 | 1385.00 |
| TMEM69 | 878.00 | 831.00 | 840.00 | 849.67 | 1145.00 |
| TMEM70 | 663.00 | 663.00 | 763.00 | 696.33 | 851.00 |

| | | | | | |
|---|---|---|---|---|---|
| TMEM71 | 167.00 | 184.00 | 202.00 | 184.33 | 209.00 |
| TMEM74 | 24.00 | 30.00 | 25.00 | 26.33 | 27.00 |
| TMEM79 | 298.00 | 264.00 | 306.00 | 289.33 | 322.00 |
| TMEM80 | 684.00 | 622.00 | 637.00 | 647.67 | 710.00 |
| TMEM81 | 65.00 | 87.00 | 91.00 | 81.00 | 90.00 |
| TMEM86A | 13.00 | 5.00 | 16.00 | 11.33 | 7.00 |
| TMEM86B | 141.00 | 141.00 | 179.00 | 153.67 | 140.00 |
| TMEM87A | 1596.00 | 1606.00 | 1758.00 | 1653.33 | 1656.00 |
| TMEM87B | 1616.00 | 1459.00 | 1655.00 | 1576.67 | 1331.00 |
| TMEM88 | 18.00 | 23.00 | 25.00 | 22.00 | 15.00 |
| TMEM8A | 1550.00 | 1717.00 | 1779.00 | 1682.00 | 1727.00 |
| TMEM8B | 398.00 | 373.00 | 382.00 | 384.33 | 310.00 |
| TMEM9 | 1779.00 | 1852.00 | 1682.00 | 1771.00 | 1866.00 |
| TMEM91 | 188.00 | 194.00 | 236.00 | 206.00 | 182.00 |
| TMEM92-AS1 | 29.00 | 32.00 | 23.00 | 28.00 | 34.00 |
| TMEM97 | 620.00 | 560.00 | 554.00 | 578.00 | 844.00 |
| TMEM98 | 485.00 | 471.00 | 479.00 | 478.33 | 618.00 |
| TMEM99 | 338.00 | 304.00 | 345.00 | 329.00 | 399.00 |
| TMEM9B | 765.00 | 736.00 | 686.00 | 729.00 | 833.00 |
| TMEM9B-AS1 | 260.00 | 229.00 | 292.00 | 260.33 | 140.00 |
| TMF1 | 1140.00 | 1055.00 | 1181.00 | 1125.33 | 1117.00 |
| TMLHE | 146.00 | 128.00 | 152.00 | 142.00 | 135.00 |
| TMOD2 | 338.00 | 317.00 | 416.00 | 357.00 | 319.00 |
| TMOD3 | 825.00 | 686.00 | 828.00 | 779.67 | 859.00 |
| TMOD4 | 20.00 | 21.00 | 26.00 | 22.33 | 19.00 |
| TMPO | 1791.00 | 1590.00 | 1933.00 | 1771.33 | 2470.00 |
| TMPO-AS1 | 32.00 | 42.00 | 40.00 | 38.00 | 67.00 |
| TMPPE | 52.00 | 50.00 | 44.00 | 48.67 | 22.00 |
| TMPRSS15 | 896.00 | 790.00 | 1072.00 | 919.33 | 796.00 |
| TMPRSS5 | 11.00 | 14.00 | 12.00 | 12.33 | 10.00 |
| TMPRSS7 | 5.00 | 3.00 | 5.00 | 4.33 | 3.00 |
| TMSB10 | 11371.00 | 11523.00 | 11851.00 | 11581.67 | 18708.00 |
| TMSB15A | 7.00 | 7.00 | 5.00 | 6.33 | 12.00 |
| TMSB4X | 2875.00 | 3052.00 | 3173.00 | 3033.33 | 5257.00 |
| TMTC1 | 5.00 | 6.00 | 6.00 | 5.67 | 3.00 |
| TMTC2 | 302.00 | 300.00 | 337.00 | 313.00 | 283.00 |
| TMTC3 | 657.00 | 598.00 | 720.00 | 658.33 | 620.00 |
| TMTC4 | 536.00 | 535.00 | 538.00 | 536.33 | 674.00 |
| TMUB1 | 1065.00 | 1060.00 | 1045.00 | 1056.67 | 1306.00 |
| TMUB2 | 1200.00 | 1286.00 | 1281.00 | 1255.67 | 1196.00 |
| TMX1 | 1822.00 | 1600.00 | 1617.00 | 1679.67 | 2064.00 |
| TMX2 | 2513.00 | 2447.00 | 2387.00 | 2449.00 | 2905.00 |
| TMX3 | 3491.00 | 3405.00 | 3519.00 | 3471.67 | 3593.00 |
| TMX4 | 1062.00 | 919.00 | 930.00 | 970.33 | 1020.00 |
| TNC | 10415.00 | 9684.00 | 10955.00 | 10351.33 | 6783.00 |
| TNFAIP1 | 670.00 | 582.00 | 698.00 | 650.00 | 657.00 |
| TNFAIP2 | 1428.00 | 1307.00 | 1352.00 | 1362.33 | 1060.00 |
| TNFAIP3 | 760.00 | 816.00 | 835.00 | 803.67 | 703.00 |

| | | | | | |
|---|---|---|---|---|---|
| TNFAIP6 | 3.00 | 1.00 | 7.00 | 3.67 | 0.00 |
| TNFAIP8 | 105.00 | 64.00 | 101.00 | 90.00 | 116.00 |
| TNFAIP8L1 | 244.00 | 220.00 | 269.00 | 244.33 | 376.00 |
| TNFRSF10A | 465.00 | 473.00 | 494.00 | 477.33 | 401.00 |
| TNFRSF10B | 5984.00 | 5867.00 | 5841.00 | 5897.33 | 4943.00 |
| TNFRSF10C | 8.00 | 3.00 | 5.00 | 5.33 | 1.00 |
| TNFRSF10D | 5816.00 | 5574.00 | 5277.00 | 5555.67 | 4324.00 |
| TNFRSF11A | 170.00 | 142.00 | 147.00 | 153.00 | 145.00 |
| TNFRSF11B | 14.00 | 9.00 | 13.00 | 12.00 | 11.00 |
| TNFRSF12A | 7975.00 | 8905.00 | 8461.00 | 8447.00 | 10347.00 |
| TNFRSF19 | 717.00 | 657.00 | 693.00 | 689.00 | 858.00 |
| TNFRSF1A | 2843.00 | 2864.00 | 2643.00 | 2783.33 | 2517.00 |
| TNFRSF21 | 913.00 | 914.00 | 927.00 | 918.00 | 760.00 |
| TNFRSF25 | 594.00 | 706.00 | 743.00 | 681.00 | 801.00 |
| TNFRSF6B | 283.00 | 320.00 | 257.00 | 286.67 | 653.00 |
| TNFRSF9 | 40.00 | 32.00 | 28.00 | 33.33 | 25.00 |
| TNFSF12 | 168.00 | 164.00 | 175.00 | 169.00 | 215.00 |
| TNFSF13 | 7.00 | 20.00 | 11.00 | 12.67 | 8.00 |
| TNFSF13B | 43.00 | 47.00 | 54.00 | 48.00 | 43.00 |
| TNFSF15 | 25.00 | 18.00 | 30.00 | 24.33 | 21.00 |
| TNFSF18 | 7.00 | 0.00 | 2.00 | 3.00 | 11.00 |
| TNFSF4 | 59.00 | 82.00 | 78.00 | 73.00 | 96.00 |
| TNFSF9 | 5.00 | 7.00 | 8.00 | 6.67 | 5.00 |
| TNIK | 435.00 | 419.00 | 424.00 | 426.00 | 342.00 |
| TNIP1 | 2324.00 | 2291.00 | 2395.00 | 2336.67 | 1929.00 |
| TNIP2 | 543.00 | 564.00 | 608.00 | 571.67 | 683.00 |
| TNIP3 | 4.00 | 2.00 | 3.00 | 3.00 | 4.00 |
| TNK2 | 1864.00 | 1920.00 | 2046.00 | 1943.33 | 1878.00 |
| TNKS | 1267.00 | 1116.00 | 1365.00 | 1249.33 | 1239.00 |
| TNKS1BP1 | 3898.00 | 3981.00 | 4080.00 | 3986.33 | 3687.00 |
| TNKS2 | 2407.00 | 2171.00 | 2559.00 | 2379.00 | 2402.00 |
| TNKS2-AS1 | 14.00 | 10.00 | 15.00 | 13.00 | 16.00 |
| TNNC1 | 5.00 | 3.00 | 6.00 | 4.67 | 2.00 |
| TNNT1 | 51.00 | 65.00 | 76.00 | 64.00 | 54.00 |
| TNPO1 | 5390.00 | 4614.00 | 6209.00 | 5404.33 | 5182.00 |
| TNPO2 | 4144.00 | 4194.00 | 4360.00 | 4232.67 | 4411.00 |
| TNPO3 | 2686.00 | 2575.00 | 2804.00 | 2688.33 | 2482.00 |
| TNRC18 | 5872.00 | 5685.00 | 5995.00 | 5850.67 | 5335.00 |
| TNRC6A | 3006.00 | 2869.00 | 3118.00 | 2997.67 | 2735.00 |
| TNRC6B | 721.00 | 718.00 | 787.00 | 742.00 | 618.00 |
| TNRC6C | 402.00 | 400.00 | 439.00 | 413.67 | 351.00 |
| TNRC6C-AS1 | 7.00 | 3.00 | 3.00 | 4.33 | 4.00 |
| TNS2 | 753.00 | 733.00 | 767.00 | 751.00 | 402.00 |
| TNS3 | 2445.00 | 2229.00 | 2374.00 | 2349.33 | 1990.00 |
| TOB1 | 176.00 | 194.00 | 207.00 | 192.33 | 133.00 |
| TOB1-AS1 | 6.00 | 13.00 | 14.00 | 11.00 | 10.00 |
| TOB2 | 1212.00 | 1211.00 | 1249.00 | 1224.00 | 1191.00 |
| TOB2P1 | 38.00 | 32.00 | 26.00 | 32.00 | 45.00 |

| | | | | | |
|---|---|---|---|---|---|
| TOE1 | 655.00 | 606.00 | 682.00 | 647.67 | 814.00 |
| TOLLIP | 885.00 | 831.00 | 823.00 | 846.33 | 839.00 |
| TOLLIP-AS1 | 11.00 | 12.00 | 11.00 | 11.33 | 12.00 |
| TOM1 | 536.00 | 527.00 | 577.00 | 546.67 | 609.00 |
| TOM1L1 | 831.00 | 781.00 | 805.00 | 805.67 | 1097.00 |
| TOM1L2 | 1027.00 | 1011.00 | 1066.00 | 1034.67 | 885.00 |
| TOMM20 | 3874.00 | 3812.00 | 4182.00 | 3956.00 | 5047.00 |
| TOMM22 | 960.00 | 968.00 | 962.00 | 963.33 | 1394.00 |
| TOMM34 | 6789.00 | 6238.00 | 7114.00 | 6713.67 | 7563.00 |
| TOMM40 | 1464.00 | 1476.00 | 1503.00 | 1481.00 | 1973.00 |
| TOMM40L | 484.00 | 432.00 | 457.00 | 457.67 | 526.00 |
| TOMM5 | 1120.00 | 1187.00 | 1242.00 | 1183.00 | 1794.00 |
| TOMM6 | 1263.00 | 1205.00 | 1413.00 | 1293.67 | 1159.00 |
| TOMM7 | 1801.00 | 1851.00 | 2042.00 | 1898.00 | 2593.00 |
| TOMM70A | 3129.00 | 2720.00 | 3144.00 | 2997.67 | 3423.00 |
| TONSL | 1357.00 | 1555.00 | 1529.00 | 1480.33 | 1680.00 |
| TONSL-AS1 | 21.00 | 26.00 | 32.00 | 26.33 | 21.00 |
| TOP1 | 5010.00 | 4719.00 | 5138.00 | 4955.67 | 4653.00 |
| TOP1MT | 1358.00 | 1412.00 | 1658.00 | 1476.00 | 1670.00 |
| TOP1P1 | 49.00 | 51.00 | 60.00 | 53.33 | 45.00 |
| TOP1P2 | 3.00 | 0.00 | 2.00 | 1.67 | 2.00 |
| TOP2A | 3991.00 | 3497.00 | 3679.00 | 3722.33 | 4298.00 |
| TOP2B | 3312.00 | 3029.00 | 3669.00 | 3336.67 | 3632.00 |
| TOP3A | 815.00 | 805.00 | 838.00 | 819.33 | 807.00 |
| TOP3B | 796.00 | 805.00 | 876.00 | 825.67 | 762.00 |
| TOPBP1 | 1567.00 | 1443.00 | 1687.00 | 1565.67 | 1637.00 |
| TOPORS | 791.00 | 728.00 | 838.00 | 785.67 | 874.00 |
| TOPORS-AS1 | 29.00 | 52.00 | 38.00 | 39.67 | 33.00 |
| TOR1A | 1747.00 | 1682.00 | 1542.00 | 1657.00 | 1877.00 |
| TOR1AIP1 | 1275.00 | 1230.00 | 1213.00 | 1239.33 | 1412.00 |
| TOR1AIP2 | 1730.00 | 1670.00 | 1832.00 | 1744.00 | 1983.00 |
| TOR1B | 1152.00 | 1211.00 | 1116.00 | 1159.67 | 1132.00 |
| TOR2A | 506.00 | 510.00 | 486.00 | 500.67 | 527.00 |
| TOR3A | 784.00 | 779.00 | 741.00 | 768.00 | 915.00 |
| TOR4A | 1519.00 | 1636.00 | 1789.00 | 1648.00 | 1866.00 |
| TOX | 56.00 | 63.00 | 66.00 | 61.67 | 84.00 |
| TOX2 | 1830.00 | 1734.00 | 1886.00 | 1816.67 | 1547.00 |
| TOX4 | 1190.00 | 1168.00 | 1215.00 | 1191.00 | 1142.00 |
| TP53 | 1264.00 | 1234.00 | 1360.00 | 1286.00 | 1937.00 |
| TP53AIP1 | 3.00 | 2.00 | 2.00 | 2.33 | 0.00 |
| TP53BP1 | 2384.00 | 2185.00 | 2385.00 | 2318.00 | 2093.00 |
| TP53BP2 | 1499.00 | 1406.00 | 1534.00 | 1479.67 | 1424.00 |
| TP53I13 | 763.00 | 759.00 | 700.00 | 740.67 | 786.00 |
| TP53I3 | 247.00 | 232.00 | 280.00 | 253.00 | 287.00 |
| TP53INP1 | 168.00 | 169.00 | 192.00 | 176.33 | 124.00 |
| TP53INP2 | 267.00 | 272.00 | 331.00 | 290.00 | 166.00 |
| TP53RK | 525.00 | 496.00 | 552.00 | 524.33 | 654.00 |
| TP53TG1 | 40.00 | 30.00 | 57.00 | 42.33 | 63.00 |

| | | | | | |
|---|---|---|---|---|---|
| TP53TG3D | 30.00 | 24.00 | 23.00 | 25.67 | 20.00 |
| TP63 | 707.00 | 614.00 | 649.00 | 656.67 | 871.00 |
| TPBG | 4716.00 | 4549.00 | 4891.00 | 4718.67 | 2179.00 |
| TPCN1 | 5013.00 | 4932.00 | 5099.00 | 5014.67 | 3364.00 |
| TPCN2 | 3667.00 | 3403.00 | 3506.00 | 3525.33 | 3302.00 |
| TPD52 | 693.00 | 632.00 | 780.00 | 701.67 | 942.00 |
| TPD52L1 | 21.00 | 23.00 | 23.00 | 22.33 | 33.00 |
| TPD52L2 | 4384.00 | 4299.00 | 4448.00 | 4377.00 | 5153.00 |
| TPGS1 | 149.00 | 145.00 | 136.00 | 143.33 | 168.00 |
| TPGS2 | 881.00 | 906.00 | 907.00 | 898.00 | 949.00 |
| TPH1 | 26.00 | 16.00 | 17.00 | 19.67 | 31.00 |
| TPI1 | 9473.00 | 9209.00 | 9896.00 | 9526.00 | 12547.00 |
| TPI1P2 | 46.00 | 51.00 | 39.00 | 45.33 | 58.00 |
| TPK1 | 82.00 | 85.00 | 123.00 | 96.67 | 132.00 |
| TPM1 | 5774.00 | 5763.00 | 6318.00 | 5951.67 | 7601.00 |
| TPM2 | 1367.00 | 1483.00 | 1661.00 | 1503.67 | 1678.00 |
| TPM3 | 6335.00 | 5829.00 | 6321.00 | 6161.67 | 8346.00 |
| TPM3P9 | 191.00 | 201.00 | 221.00 | 204.33 | 202.00 |
| TPM4 | 10368.00 | 9984.00 | 11165.00 | 10505.67 | 13278.00 |
| TPMT | 381.00 | 395.00 | 462.00 | 412.67 | 500.00 |
| TPP1 | 19207.00 | 19267.00 | 19813.00 | 19429.00 | 14348.00 |
| TPP2 | 2040.00 | 1920.00 | 2218.00 | 2059.33 | 2162.00 |
| TPPP | 5.00 | 5.00 | 3.00 | 4.33 | 1.00 |
| TPPP3 | 5.00 | 11.00 | 5.00 | 7.00 | 9.00 |
| TPR | 5775.00 | 5501.00 | 6063.00 | 5779.67 | 4949.00 |
| TPRA1 | 1599.00 | 1660.00 | 1755.00 | 1671.33 | 1536.00 |
| TPRG1 | 8.00 | 1.00 | 1.00 | 3.33 | 3.00 |
| TPRG1-AS1 | 4.00 | 6.00 | 3.00 | 4.33 | 2.00 |
| TPRG1L | 719.00 | 744.00 | 807.00 | 756.67 | 724.00 |
| TPRKB | 421.00 | 445.00 | 468.00 | 444.67 | 570.00 |
| TPRN | 687.00 | 732.00 | 809.00 | 742.67 | 731.00 |
| TPST1 | 846.00 | 879.00 | 844.00 | 856.33 | 936.00 |
| TPST2 | 503.00 | 417.00 | 496.00 | 472.00 | 420.00 |
| TPT1 | 25252.00 | 24733.00 | 27115.00 | 25700.00 | 31328.00 |
| TPT1-AS1 | 158.00 | 138.00 | 170.00 | 155.33 | 178.00 |
| TPTE2 | 4.00 | 2.00 | 6.00 | 4.00 | 3.00 |
| TPTE2P1 | 7.00 | 8.00 | 15.00 | 10.00 | 9.00 |
| TPTE2P5 | 19.00 | 14.00 | 22.00 | 18.33 | 23.00 |
| TPTE2P6 | 19.00 | 17.00 | 21.00 | 19.00 | 7.00 |
| TPX2 | 4366.00 | 3995.00 | 4109.00 | 4156.67 | 4358.00 |
| TRA2A | 1876.00 | 1857.00 | 2108.00 | 1947.00 | 2510.00 |
| TRA2B | 3549.00 | 3412.00 | 3803.00 | 3588.00 | 4612.00 |
| TRABD | 1456.00 | 1475.00 | 1620.00 | 1517.00 | 1765.00 |
| TRABD2A | 705.00 | 669.00 | 704.00 | 692.67 | 684.00 |
| TRADD | 249.00 | 289.00 | 282.00 | 273.33 | 278.00 |
| TRAF1 | 48.00 | 91.00 | 75.00 | 71.33 | 121.00 |
| TRAF2 | 740.00 | 844.00 | 874.00 | 819.33 | 974.00 |
| TRAF3 | 1455.00 | 1393.00 | 1498.00 | 1448.67 | 1407.00 |

| | | | | | |
|---|---|---|---|---|---|
| TRAF3IP1 | 392.00 | 412.00 | 487.00 | 430.33 | 400.00 |
| TRAF3IP2 | 344.00 | 281.00 | 313.00 | 312.67 | 293.00 |
| TRAF3IP2-AS1 | 205.00 | 177.00 | 234.00 | 205.33 | 199.00 |
| TRAF4 | 705.00 | 716.00 | 737.00 | 719.33 | 751.00 |
| TRAF5 | 778.00 | 672.00 | 662.00 | 704.00 | 867.00 |
| TRAF6 | 405.00 | 389.00 | 417.00 | 403.67 | 342.00 |
| TRAF7 | 2687.00 | 2715.00 | 2956.00 | 2786.00 | 2108.00 |
| TRAFD1 | 513.00 | 445.00 | 475.00 | 477.67 | 487.00 |
| TRAIP | 313.00 | 384.00 | 287.00 | 328.00 | 424.00 |
| TRAK1 | 1077.00 | 1105.00 | 1137.00 | 1106.33 | 1172.00 |
| TRAK2 | 1151.00 | 1033.00 | 1198.00 | 1127.33 | 909.00 |
| TRAM1 | 3805.00 | 3425.00 | 4015.00 | 3748.33 | 3320.00 |
| TRAM1L1 | 77.00 | 62.00 | 91.00 | 76.67 | 92.00 |
| TRAM2 | 6544.00 | 6013.00 | 6496.00 | 6351.00 | 4253.00 |
| TRAM2-AS1 | 207.00 | 156.00 | 206.00 | 189.67 | 248.00 |
| TRANK1 | 47.00 | 56.00 | 36.00 | 46.33 | 40.00 |
| TRAP1 | 3415.00 | 3381.00 | 3618.00 | 3471.33 | 4065.00 |
| TRAPPC1 | 1225.00 | 1260.00 | 1212.00 | 1232.33 | 1725.00 |
| TRAPPC10 | 857.00 | 877.00 | 882.00 | 872.00 | 756.00 |
| TRAPPC11 | 763.00 | 716.00 | 758.00 | 745.67 | 673.00 |
| TRAPPC12 | 331.00 | 346.00 | 392.00 | 356.33 | 280.00 |
| TRAPPC13 | 654.00 | 550.00 | 626.00 | 610.00 | 789.00 |
| TRAPPC2 | 404.00 | 406.00 | 428.00 | 412.67 | 547.00 |
| TRAPPC2B | 61.00 | 57.00 | 80.00 | 66.00 | 69.00 |
| TRAPPC2L | 539.00 | 536.00 | 593.00 | 556.00 | 747.00 |
| TRAPPC3 | 1090.00 | 1170.00 | 1223.00 | 1161.00 | 1260.00 |
| TRAPPC3L | 3.00 | 0.00 | 0.00 | 1.00 | 0.00 |
| TRAPPC4 | 669.00 | 605.00 | 660.00 | 644.67 | 761.00 |
| TRAPPC5 | 396.00 | 380.00 | 390.00 | 388.67 | 495.00 |
| TRAPPC6A | 75.00 | 65.00 | 98.00 | 79.33 | 76.00 |
| TRAPPC6B | 694.00 | 548.00 | 646.00 | 629.33 | 682.00 |
| TRAPPC8 | 1007.00 | 917.00 | 1042.00 | 988.67 | 999.00 |
| TRAPPC9 | 385.00 | 373.00 | 429.00 | 395.67 | 379.00 |
| TRDMT1 | 266.00 | 288.00 | 292.00 | 282.00 | 290.00 |
| TRERF1 | 1393.00 | 1223.00 | 1414.00 | 1343.33 | 1154.00 |
| TREX1 | 243.00 | 265.00 | 252.00 | 253.33 | 268.00 |
| TREX2 | 33.00 | 31.00 | 32.00 | 32.00 | 33.00 |
| TRG-AS1 | 3.00 | 6.00 | 13.00 | 7.33 | 7.00 |
| TRHDE | 1532.00 | 1337.00 | 1606.00 | 1491.67 | 1157.00 |
| TRHDE-AS1 | 798.00 | 796.00 | 841.00 | 811.67 | 655.00 |
| TRIAP1 | 308.00 | 325.00 | 355.00 | 329.33 | 450.00 |
| TRIB1 | 524.00 | 518.00 | 548.00 | 530.00 | 658.00 |
| TRIB2 | 728.00 | 684.00 | 755.00 | 722.33 | 582.00 |
| TRIB3 | 6311.00 | 5836.00 | 6224.00 | 6123.67 | 5130.00 |
| TRIL | 3.00 | 7.00 | 20.00 | 10.00 | 19.00 |
| TRIM11 | 923.00 | 915.00 | 992.00 | 943.33 | 1116.00 |
| TRIM13 | 592.00 | 581.00 | 674.00 | 615.67 | 660.00 |
| TRIM14 | 791.00 | 684.00 | 777.00 | 750.67 | 808.00 |

| | | | | | |
|---|---|---|---|---|---|
| TRIM16 | 575.00 | 605.00 | 651.00 | 610.33 | 692.00 |
| TRIM16L | 229.00 | 199.00 | 231.00 | 219.67 | 257.00 |
| TRIM2 | 16.00 | 22.00 | 18.00 | 18.67 | 26.00 |
| TRIM21 | 260.00 | 238.00 | 238.00 | 245.33 | 302.00 |
| TRIM23 | 296.00 | 241.00 | 289.00 | 275.33 | 302.00 |
| TRIM24 | 400.00 | 361.00 | 405.00 | 388.67 | 461.00 |
| TRIM25 | 4058.00 | 3858.00 | 4339.00 | 4085.00 | 3459.00 |
| TRIM28 | 6988.00 | 6860.00 | 6837.00 | 6895.00 | 7718.00 |
| TRIM3 | 521.00 | 512.00 | 515.00 | 516.00 | 505.00 |
| TRIM32 | 650.00 | 657.00 | 698.00 | 668.33 | 610.00 |
| TRIM33 | 1336.00 | 1199.00 | 1385.00 | 1306.67 | 1174.00 |
| TRIM34 | 13.00 | 10.00 | 13.00 | 12.00 | 8.00 |
| TRIM35 | 299.00 | 271.00 | 341.00 | 303.67 | 349.00 |
| TRIM36 | 144.00 | 148.00 | 165.00 | 152.33 | 190.00 |
| TRIM37 | 1110.00 | 984.00 | 1107.00 | 1067.00 | 1171.00 |
| TRIM38 | 1296.00 | 1215.00 | 1305.00 | 1272.00 | 1288.00 |
| TRIM4 | 309.00 | 264.00 | 294.00 | 289.00 | 283.00 |
| TRIM41 | 2112.00 | 2089.00 | 2170.00 | 2123.67 | 2321.00 |
| TRIM44 | 5124.00 | 4625.00 | 5418.00 | 5055.67 | 4574.00 |
| TRIM45 | 561.00 | 548.00 | 491.00 | 533.33 | 567.00 |
| TRIM46 | 303.00 | 315.00 | 345.00 | 321.00 | 316.00 |
| TRIM47 | 30.00 | 39.00 | 37.00 | 35.33 | 54.00 |
| TRIM5 | 908.00 | 821.00 | 940.00 | 889.67 | 1030.00 |
| TRIM52 | 443.00 | 449.00 | 477.00 | 456.33 | 352.00 |
| TRIM52-AS1 | 222.00 | 199.00 | 253.00 | 224.67 | 237.00 |
| TRIM55 | 2184.00 | 2004.00 | 2235.00 | 2141.00 | 2917.00 |
| TRIM56 | 772.00 | 736.00 | 874.00 | 794.00 | 437.00 |
| TRIM58 | 884.00 | 872.00 | 921.00 | 892.33 | 799.00 |
| TRIM59 | 393.00 | 332.00 | 390.00 | 371.67 | 438.00 |
| TRIM6 | 139.00 | 147.00 | 151.00 | 145.67 | 167.00 |
| TRIM62 | 192.00 | 191.00 | 198.00 | 193.67 | 186.00 |
| TRIM65 | 1470.00 | 1508.00 | 1752.00 | 1576.67 | 2098.00 |
| TRIM66 | 1234.00 | 1129.00 | 1231.00 | 1198.00 | 909.00 |
| TRIM68 | 374.00 | 389.00 | 413.00 | 392.00 | 385.00 |
| TRIM7 | 87.00 | 76.00 | 83.00 | 82.00 | 95.00 |
| TRIM8 | 3747.00 | 3675.00 | 4096.00 | 3839.33 | 3989.00 |
| TRIM9 | 138.00 | 104.00 | 123.00 | 121.67 | 197.00 |
| TRIO | 13442.00 | 12723.00 | 14462.00 | 13542.33 | 9261.00 |
| TRIOBP | 2575.00 | 2488.00 | 2529.00 | 2530.67 | 2289.00 |
| TRIP10 | 1552.00 | 1537.00 | 1639.00 | 1576.00 | 1752.00 |
| TRIP11 | 1739.00 | 1672.00 | 1777.00 | 1729.33 | 1593.00 |
| TRIP12 | 5677.00 | 5176.00 | 5923.00 | 5592.00 | 5362.00 |
| TRIP13 | 934.00 | 826.00 | 910.00 | 890.00 | 1265.00 |
| TRIP4 | 462.00 | 462.00 | 553.00 | 492.33 | 567.00 |
| TRIP6 | 2737.00 | 2818.00 | 3037.00 | 2864.00 | 3362.00 |
| TRIQK | 696.00 | 609.00 | 845.00 | 716.67 | 854.00 |
| TRIT1 | 865.00 | 863.00 | 1040.00 | 922.67 | 1009.00 |
| TRMT1 | 3526.00 | 3736.00 | 3799.00 | 3687.00 | 4060.00 |

| | | | | | |
|---|---|---|---|---|---|
| TRMT10A | 165.00 | 157.00 | 167.00 | 163.00 | 182.00 |
| TRMT10B | 303.00 | 279.00 | 317.00 | 299.67 | 335.00 |
| TRMT10C | 736.00 | 699.00 | 758.00 | 731.00 | 879.00 |
| TRMT11 | 654.00 | 595.00 | 674.00 | 641.00 | 689.00 |
| TRMT112 | 3345.00 | 3305.00 | 3463.00 | 3371.00 | 4349.00 |
| TRMT12 | 378.00 | 378.00 | 390.00 | 382.00 | 377.00 |
| TRMT13 | 1022.00 | 1042.00 | 1204.00 | 1089.33 | 1054.00 |
| TRMT1L | 757.00 | 778.00 | 832.00 | 789.00 | 813.00 |
| TRMT2A | 1021.00 | 1089.00 | 1020.00 | 1043.33 | 1161.00 |
| TRMT2B | 351.00 | 348.00 | 404.00 | 367.67 | 483.00 |
| TRMT44 | 356.00 | 334.00 | 327.00 | 339.00 | 322.00 |
| TRMT5 | 1006.00 | 875.00 | 969.00 | 950.00 | 1062.00 |
| TRMT6 | 1095.00 | 896.00 | 1027.00 | 1006.00 | 1296.00 |
| TRMT61A | 999.00 | 990.00 | 1027.00 | 1005.33 | 1237.00 |
| TRMT61B | 507.00 | 436.00 | 535.00 | 492.67 | 503.00 |
| TRMU | 704.00 | 716.00 | 803.00 | 741.00 | 763.00 |
| TRNAU1AP | 215.00 | 244.00 | 258.00 | 239.00 | 278.00 |
| TRNP1 | 1660.00 | 1463.00 | 1627.00 | 1583.33 | 1097.00 |
| TRNT1 | 827.00 | 751.00 | 817.00 | 798.33 | 823.00 |
| TRO | 3.00 | 2.00 | 2.00 | 2.33 | 0.00 |
| TROAP | 1102.00 | 1124.00 | 968.00 | 1064.67 | 1314.00 |
| TROVE2 | 402.00 | 365.00 | 433.00 | 400.00 | 407.00 |
| TRPA1 | 148.00 | 123.00 | 161.00 | 144.00 | 147.00 |
| TRPC1 | 837.00 | 863.00 | 919.00 | 873.00 | 726.00 |
| TRPC3 | 17.00 | 18.00 | 39.00 | 24.67 | 24.00 |
| TRPC4 | 45.00 | 40.00 | 40.00 | 41.67 | 58.00 |
| TRPC4AP | 3211.00 | 3214.00 | 3530.00 | 3318.33 | 3291.00 |
| TRPC6 | 98.00 | 120.00 | 142.00 | 120.00 | 119.00 |
| TRPM2 | 679.00 | 729.00 | 725.00 | 711.00 | 851.00 |
| TRPM2-AS | 3.00 | 1.00 | 3.00 | 2.33 | 1.00 |
| TRPM4 | 935.00 | 916.00 | 981.00 | 944.00 | 905.00 |
| TRPM6 | 3.00 | 6.00 | 2.00 | 3.67 | 1.00 |
| TRPM7 | 2946.00 | 2683.00 | 3232.00 | 2953.67 | 3038.00 |
| TRPS1 | 496.00 | 463.00 | 578.00 | 512.33 | 428.00 |
| TRPT1 | 865.00 | 916.00 | 899.00 | 893.33 | 1070.00 |
| TRPV1 | 545.00 | 601.00 | 662.00 | 602.67 | 528.00 |
| TRPV2 | 1966.00 | 1988.00 | 1766.00 | 1906.67 | 1627.00 |
| TRRAP | 2878.00 | 2543.00 | 2887.00 | 2769.33 | 2272.00 |
| TRUB1 | 848.00 | 699.00 | 875.00 | 807.33 | 881.00 |
| TRUB2 | 711.00 | 690.00 | 784.00 | 728.33 | 731.00 |
| TSACC | 15.00 | 8.00 | 14.00 | 12.33 | 18.00 |
| TSC1 | 1815.00 | 1851.00 | 1886.00 | 1850.67 | 1715.00 |
| TSC2 | 2539.00 | 2356.00 | 2554.00 | 2483.00 | 2041.00 |
| TSC22D1 | 1651.00 | 1611.00 | 1973.00 | 1745.00 | 1222.00 |
| TSC22D1-AS1 | 36.00 | 25.00 | 40.00 | 33.67 | 35.00 |
| TSC22D2 | 976.00 | 986.00 | 1164.00 | 1042.00 | 1064.00 |
| TSC22D3 | 530.00 | 480.00 | 546.00 | 518.67 | 611.00 |
| TSC22D4 | 1180.00 | 1218.00 | 1293.00 | 1230.33 | 1048.00 |

| | | | | | |
|---|---|---|---|---|---|
| TSEN15 | 1372.00 | 1364.00 | 1454.00 | 1396.67 | 1817.00 |
| TSEN2 | 673.00 | 673.00 | 692.00 | 679.33 | 583.00 |
| TSEN54 | 515.00 | 613.00 | 628.00 | 585.33 | 630.00 |
| TSFM | 529.00 | 491.00 | 537.00 | 519.00 | 664.00 |
| TSG101 | 1344.00 | 1214.00 | 1331.00 | 1296.33 | 1465.00 |
| TSGA10 | 63.00 | 78.00 | 79.00 | 73.33 | 73.00 |
| TSGA10IP | 14.00 | 13.00 | 10.00 | 12.33 | 13.00 |
| TSHZ1 | 249.00 | 233.00 | 271.00 | 251.00 | 228.00 |
| TSHZ3 | 507.00 | 459.00 | 470.00 | 478.67 | 531.00 |
| TSIX | 9.00 | 9.00 | 8.00 | 8.67 | 0.00 |
| TSKU | 1008.00 | 936.00 | 1045.00 | 996.33 | 958.00 |
| TSLP | 597.00 | 458.00 | 630.00 | 561.67 | 349.00 |
| TSN | 1812.00 | 1649.00 | 1848.00 | 1769.67 | 2255.00 |
| TSNARE1 | 229.00 | 231.00 | 253.00 | 237.67 | 253.00 |
| TSNAX | 573.00 | 487.00 | 594.00 | 551.33 | 701.00 |
| TSNAXIP1 | 30.00 | 32.00 | 44.00 | 35.33 | 28.00 |
| TSPAN1 | 6.00 | 2.00 | 3.00 | 3.67 | 5.00 |
| TSPAN10 | 70.00 | 65.00 | 91.00 | 75.33 | 119.00 |
| TSPAN12 | 134.00 | 98.00 | 110.00 | 114.00 | 143.00 |
| TSPAN13 | 105.00 | 99.00 | 94.00 | 99.33 | 132.00 |
| TSPAN14 | 4059.00 | 4064.00 | 3987.00 | 4036.67 | 2459.00 |
| TSPAN15 | 141.00 | 133.00 | 129.00 | 134.33 | 110.00 |
| TSPAN17 | 1118.00 | 1102.00 | 1222.00 | 1147.33 | 1178.00 |
| TSPAN2 | 7.00 | 5.00 | 1.00 | 4.33 | 12.00 |
| TSPAN3 | 3647.00 | 3870.00 | 3571.00 | 3696.00 | 4155.00 |
| TSPAN31 | 489.00 | 515.00 | 561.00 | 521.67 | 538.00 |
| TSPAN33 | 20.00 | 28.00 | 16.00 | 21.33 | 50.00 |
| TSPAN4 | 4052.00 | 4106.00 | 4159.00 | 4105.67 | 4719.00 |
| TSPAN5 | 2157.00 | 2081.00 | 2150.00 | 2129.33 | 1787.00 |
| TSPAN6 | 880.00 | 891.00 | 908.00 | 893.00 | 1030.00 |
| TSPAN7 | 437.00 | 390.00 | 379.00 | 402.00 | 348.00 |
| TSPAN8 | 3.00 | 6.00 | 2.00 | 3.67 | 4.00 |
| TSPAN9 | 785.00 | 781.00 | 778.00 | 781.33 | 614.00 |
| TSPO | 2128.00 | 2106.00 | 2080.00 | 2104.67 | 2792.00 |
| TSPYL1 | 1123.00 | 1038.00 | 1171.00 | 1110.67 | 1269.00 |
| TSPYL2 | 1255.00 | 1377.00 | 1418.00 | 1350.00 | 1097.00 |
| TSPYL4 | 1096.00 | 1031.00 | 1191.00 | 1106.00 | 1104.00 |
| TSR1 | 4497.00 | 4310.00 | 4632.00 | 4479.67 | 5295.00 |
| TSR2 | 546.00 | 521.00 | 530.00 | 532.33 | 675.00 |
| TSR3 | 755.00 | 775.00 | 827.00 | 785.67 | 979.00 |
| TSSC1 | 499.00 | 549.00 | 550.00 | 532.67 | 682.00 |
| TSSC2 | 153.00 | 148.00 | 213.00 | 171.33 | 115.00 |
| TSSC4 | 769.00 | 811.00 | 838.00 | 806.00 | 906.00 |
| TSSK3 | 4.00 | 3.00 | 1.00 | 2.67 | 5.00 |
| TSSK6 | 60.00 | 83.00 | 73.00 | 72.00 | 75.00 |
| TST | 194.00 | 183.00 | 168.00 | 181.67 | 240.00 |
| TSTA3 | 20.00 | 35.00 | 39.00 | 31.33 | 30.00 |
| TSTD2 | 719.00 | 722.00 | 726.00 | 722.33 | 672.00 |

| | | | | | |
|---|---|---|---|---|---|
| TSTD3 | 104.00 | 80.00 | 88.00 | 90.67 | 85.00 |
| TTBK2 | 277.00 | 238.00 | 326.00 | 280.33 | 197.00 |
| TTC1 | 1284.00 | 1251.00 | 1249.00 | 1261.33 | 1605.00 |
| TTC12 | 416.00 | 471.00 | 445.00 | 444.00 | 426.00 |
| TTC13 | 785.00 | 805.00 | 871.00 | 820.33 | 918.00 |
| TTC14 | 2168.00 | 2094.00 | 2369.00 | 2210.33 | 2214.00 |
| TTC17 | 3554.00 | 3513.00 | 3650.00 | 3572.33 | 3110.00 |
| TTC19 | 1654.00 | 1640.00 | 1631.00 | 1641.67 | 1608.00 |
| TTC21A | 131.00 | 132.00 | 134.00 | 132.33 | 84.00 |
| TTC21B | 726.00 | 683.00 | 797.00 | 735.33 | 772.00 |
| TTC23 | 361.00 | 376.00 | 391.00 | 376.00 | 405.00 |
| TTC23L | 4.00 | 1.00 | 0.00 | 1.67 | 0.00 |
| TTC25 | 21.00 | 30.00 | 21.00 | 24.00 | 26.00 |
| TTC26 | 526.00 | 514.00 | 576.00 | 538.67 | 577.00 |
| TTC27 | 515.00 | 511.00 | 525.00 | 517.00 | 564.00 |
| TTC28 | 734.00 | 640.00 | 782.00 | 718.67 | 701.00 |
| TTC28-AS1 | 199.00 | 184.00 | 261.00 | 214.67 | 183.00 |
| TTC29 | 3.00 | 4.00 | 2.00 | 3.00 | 3.00 |
| TTC3 | 3704.00 | 3569.00 | 3946.00 | 3739.67 | 3616.00 |
| TTC30A | 136.00 | 123.00 | 132.00 | 130.33 | 129.00 |
| TTC30B | 84.00 | 64.00 | 74.00 | 74.00 | 82.00 |
| TTC31 | 1099.00 | 1100.00 | 1092.00 | 1097.00 | 1069.00 |
| TTC32 | 143.00 | 121.00 | 143.00 | 135.67 | 143.00 |
| TTC33 | 391.00 | 338.00 | 432.00 | 387.00 | 489.00 |
| TTC37 | 4663.00 | 4363.00 | 5163.00 | 4729.67 | 4447.00 |
| TTC38 | 456.00 | 486.00 | 556.00 | 499.33 | 448.00 |
| TTC39A | 3.00 | 0.00 | 0.00 | 1.00 | 0.00 |
| TTC39B | 104.00 | 97.00 | 117.00 | 106.00 | 147.00 |
| TTC39C | 215.00 | 199.00 | 222.00 | 212.00 | 272.00 |
| TTC3P1 | 40.00 | 36.00 | 33.00 | 36.33 | 33.00 |
| TTC4 | 88.00 | 69.00 | 79.00 | 78.67 | 79.00 |
| TTC41P | 32.00 | 15.00 | 16.00 | 21.00 | 21.00 |
| TTC5 | 313.00 | 291.00 | 338.00 | 314.00 | 320.00 |
| TTC6 | 8.00 | 1.00 | 7.00 | 5.33 | 3.00 |
| TTC7A | 440.00 | 469.00 | 472.00 | 460.33 | 455.00 |
| TTC7B | 924.00 | 811.00 | 920.00 | 885.00 | 1162.00 |
| TTC8 | 565.00 | 528.00 | 645.00 | 579.33 | 623.00 |
| TTC9B | 3.00 | 3.00 | 2.00 | 2.67 | 2.00 |
| TTC9C | 440.00 | 466.00 | 490.00 | 465.33 | 492.00 |
| TTF1 | 418.00 | 378.00 | 457.00 | 417.67 | 478.00 |
| TTF2 | 1390.00 | 1326.00 | 1335.00 | 1350.33 | 1267.00 |
| TTI1 | 1077.00 | 1043.00 | 1136.00 | 1085.33 | 1032.00 |
| TTI2 | 246.00 | 221.00 | 207.00 | 224.67 | 255.00 |
| TTK | 779.00 | 793.00 | 853.00 | 808.33 | 1003.00 |
| TTL | 2519.00 | 2389.00 | 2605.00 | 2504.33 | 3004.00 |
| TTLL1 | 78.00 | 74.00 | 93.00 | 81.67 | 91.00 |
| TTLL11 | 415.00 | 385.00 | 413.00 | 404.33 | 268.00 |
| TTLL12 | 1361.00 | 1278.00 | 1510.00 | 1383.00 | 1577.00 |

| | | | | | |
|---|---|---|---|---|---|
| TTLL13P | 12.00 | 1.00 | 14.00 | 9.00 | 8.00 |
| TTLL3 | 163.00 | 155.00 | 178.00 | 165.33 | 83.00 |
| TTLL4 | 1451.00 | 1392.00 | 1547.00 | 1463.33 | 1242.00 |
| TTLL5 | 910.00 | 876.00 | 999.00 | 928.33 | 845.00 |
| TTLL7 | 236.00 | 198.00 | 235.00 | 223.00 | 148.00 |
| TTN | 5.00 | 4.00 | 4.00 | 4.33 | 2.00 |
| TTN-AS1 | 249.00 | 218.00 | 229.00 | 232.00 | 178.00 |
| TTPA | 19.00 | 32.00 | 29.00 | 26.67 | 28.00 |
| TTPAL | 1260.00 | 1067.00 | 1230.00 | 1185.67 | 1147.00 |
| TTYH2 | 321.00 | 308.00 | 329.00 | 319.33 | 213.00 |
| TTYH3 | 3278.00 | 3375.00 | 3539.00 | 3397.33 | 2415.00 |
| TUBA1A | 483.00 | 482.00 | 660.00 | 541.67 | 461.00 |
| TUBA1B | 5097.00 | 4766.00 | 5705.00 | 5189.33 | 6869.00 |
| TUBA1C | 6459.00 | 6198.00 | 6757.00 | 6471.33 | 7712.00 |
| TUBA3FP | 3.00 | 3.00 | 6.00 | 4.00 | 2.00 |
| TUBA4A | 3.00 | 1.00 | 1.00 | 1.67 | 1.00 |
| TUBA8 | 23.00 | 17.00 | 15.00 | 18.33 | 20.00 |
| TUBB | 13.00 | 10.00 | 12.00 | 11.67 | 7.00 |
| TUBB1 | 3.00 | 4.00 | 3.00 | 3.33 | 0.00 |
| TUBB2A | 60.00 | 64.00 | 66.00 | 63.33 | 71.00 |
| TUBB3 | 701.00 | 758.00 | 747.00 | 735.33 | 833.00 |
| TUBB4B | 4458.00 | 4400.00 | 4135.00 | 4331.00 | 5211.00 |
| TUBB6 | 6759.00 | 6536.00 | 6963.00 | 6752.67 | 11677.00 |
| TUBD1 | 119.00 | 135.00 | 138.00 | 130.67 | 148.00 |
| TUBE1 | 1427.00 | 1227.00 | 1446.00 | 1366.67 | 1705.00 |
| TUBG1 | 739.00 | 775.00 | 627.00 | 713.67 | 923.00 |
| TUBG2 | 783.00 | 804.00 | 797.00 | 794.67 | 725.00 |
| TUBGCP2 | 1729.00 | 1653.00 | 1784.00 | 1722.00 | 1907.00 |
| TUBGCP3 | 373.00 | 362.00 | 407.00 | 380.67 | 356.00 |
| TUBGCP4 | 451.00 | 409.00 | 530.00 | 463.33 | 428.00 |
| TUBGCP5 | 31.00 | 48.00 | 48.00 | 42.33 | 41.00 |
| TUBGCP6 | 1920.00 | 1912.00 | 1957.00 | 1929.67 | 1784.00 |
| TUFM | 6201.00 | 6117.00 | 6167.00 | 6161.67 | 7287.00 |
| TUFT1 | 157.00 | 128.00 | 169.00 | 151.33 | 276.00 |
| TUG1 | 6710.00 | 6381.00 | 6996.00 | 6695.67 | 6108.00 |
| TULP3 | 801.00 | 835.00 | 808.00 | 814.67 | 781.00 |
| TULP4 | 793.00 | 693.00 | 768.00 | 751.33 | 759.00 |
| TUSC2 | 563.00 | 504.00 | 595.00 | 554.00 | 638.00 |
| TUSC3 | 1615.00 | 1633.00 | 1742.00 | 1663.33 | 1954.00 |
| TUT1 | 923.00 | 837.00 | 944.00 | 901.33 | 897.00 |
| TVP23B | 669.00 | 695.00 | 680.00 | 681.33 | 869.00 |
| TVP23C | 251.00 | 287.00 | 260.00 | 266.00 | 236.00 |
| TWF1 | 1296.00 | 1115.00 | 1475.00 | 1295.33 | 1748.00 |
| TWF2 | 1421.00 | 1503.00 | 1351.00 | 1425.00 | 1531.00 |
| TWIST1 | 266.00 | 254.00 | 221.00 | 247.00 | 330.00 |
| TWIST2 | 318.00 | 304.00 | 313.00 | 311.67 | 351.00 |
| TWISTNB | 1009.00 | 845.00 | 1149.00 | 1001.00 | 1119.00 |
| TWSG1 | 1233.00 | 1161.00 | 1311.00 | 1235.00 | 1155.00 |

| | | | | | |
|---|---|---|---|---|---|
| TXK | 76.00 | 54.00 | 72.00 | 67.33 | 55.00 |
| TXLNA | 4013.00 | 3936.00 | 4266.00 | 4071.67 | 3928.00 |
| TXLNB | 19.00 | 10.00 | 15.00 | 14.67 | 20.00 |
| TXLNG | 1937.00 | 1804.00 | 2137.00 | 1959.33 | 1952.00 |
| TXN | 2653.00 | 2668.00 | 2802.00 | 2707.67 | 4234.00 |
| TXN2 | 862.00 | 878.00 | 985.00 | 908.33 | 1161.00 |
| TXNDC11 | 871.00 | 863.00 | 916.00 | 883.33 | 831.00 |
| TXNDC12 | 1991.00 | 1864.00 | 1890.00 | 1915.00 | 2499.00 |
| TXNDC15 | 1321.00 | 1325.00 | 1289.00 | 1311.67 | 1443.00 |
| TXNDC16 | 140.00 | 120.00 | 131.00 | 130.33 | 131.00 |
| TXNDC17 | 770.00 | 814.00 | 809.00 | 797.67 | 1159.00 |
| TXNDC5 | 278.00 | 230.00 | 259.00 | 255.67 | 209.00 |
| TXNDC9 | 624.00 | 597.00 | 707.00 | 642.67 | 798.00 |
| TXNIP | 230.00 | 167.00 | 143.00 | 180.00 | 277.00 |
| TXNL1 | 907.00 | 855.00 | 1042.00 | 934.67 | 1263.00 |
| TXNL4A | 650.00 | 593.00 | 727.00 | 656.67 | 962.00 |
| TXNL4B | 512.00 | 441.00 | 527.00 | 493.33 | 568.00 |
| TXNRD1 | 14967.00 | 13796.00 | 14736.00 | 14499.67 | 18305.00 |
| TXNRD2 | 1546.00 | 1591.00 | 1769.00 | 1635.33 | 1913.00 |
| TXNRD3 | 210.00 | 182.00 | 204.00 | 198.67 | 167.00 |
| TYK2 | 2397.00 | 2431.00 | 2543.00 | 2457.00 | 2642.00 |
| TYMP | 20.00 | 20.00 | 19.00 | 19.67 | 25.00 |
| TYMS | 995.00 | 971.00 | 924.00 | 963.33 | 2332.00 |
| TYMSOS | 15.00 | 13.00 | 7.00 | 11.67 | 8.00 |
| TYRO3 | 1231.00 | 1271.00 | 1216.00 | 1239.33 | 1165.00 |
| TYSND1 | 1134.00 | 1139.00 | 1268.00 | 1180.33 | 1305.00 |
| TYW1 | 768.00 | 738.00 | 822.00 | 776.00 | 808.00 |
| TYW1B | 43.00 | 56.00 | 58.00 | 52.33 | 70.00 |
| TYW3 | 1201.00 | 1050.00 | 1305.00 | 1185.33 | 1423.00 |
| TYW5 | 475.00 | 450.00 | 512.00 | 479.00 | 429.00 |
| U2AF1 | 47.00 | 30.00 | 32.00 | 36.33 | 34.00 |
| U2AF1L4 | 170.00 | 186.00 | 159.00 | 171.67 | 188.00 |
| U2AF2 | 5247.00 | 5274.00 | 5510.00 | 5343.67 | 6143.00 |
| U2SURP | 4088.00 | 3850.00 | 4449.00 | 4129.00 | 5010.00 |
| UACA | 8665.00 | 8465.00 | 9125.00 | 8751.67 | 8831.00 |
| UAP1 | 8030.00 | 7225.00 | 8008.00 | 7754.33 | 16758.00 |
| UAP1L1 | 417.00 | 445.00 | 492.00 | 451.33 | 423.00 |
| UBA1 | 10545.00 | 10162.00 | 10665.00 | 10457.33 | 11019.00 |
| UBA2 | 2451.00 | 2265.00 | 2722.00 | 2479.33 | 2525.00 |
| UBA3 | 1197.00 | 1036.00 | 1197.00 | 1143.33 | 1289.00 |
| UBA5 | 1070.00 | 1035.00 | 1107.00 | 1070.67 | 1306.00 |
| UBA52 | 5357.00 | 5227.00 | 5850.00 | 5478.00 | 7184.00 |
| UBA6 | 1201.00 | 1136.00 | 1123.00 | 1153.33 | 1179.00 |
| UBA6-AS1 | 229.00 | 220.00 | 187.00 | 212.00 | 241.00 |
| UBA7 | 57.00 | 68.00 | 89.00 | 71.33 | 62.00 |
| UBAC1 | 738.00 | 683.00 | 835.00 | 752.00 | 868.00 |
| UBAC2 | 1429.00 | 1542.00 | 1423.00 | 1464.67 | 1643.00 |
| UBAC2-AS1 | 43.00 | 47.00 | 43.00 | 44.33 | 47.00 |

| | | | | | |
|---|---|---|---|---|---|
| UBALD1 | 568.00 | 584.00 | 627.00 | 593.00 | 666.00 |
| UBALD2 | 629.00 | 562.00 | 590.00 | 593.67 | 576.00 |
| UBAP1 | 703.00 | 698.00 | 784.00 | 728.33 | 694.00 |
| UBAP1L | 63.00 | 64.00 | 54.00 | 60.33 | 50.00 |
| UBAP2 | 1420.00 | 1324.00 | 1375.00 | 1373.00 | 1269.00 |
| UBAP2L | 4287.00 | 4206.00 | 4554.00 | 4349.00 | 3856.00 |
| UBASH3B | 1247.00 | 1077.00 | 1274.00 | 1199.33 | 1570.00 |
| UBB | 9839.00 | 9531.00 | 10010.00 | 9793.33 | 12588.00 |
| UBC | 7627.00 | 7490.00 | 8277.00 | 7798.00 | 8359.00 |
| UBE2A | 630.00 | 602.00 | 641.00 | 624.33 | 763.00 |
| UBE2B | 654.00 | 589.00 | 741.00 | 661.33 | 771.00 |
| UBE2C | 1135.00 | 1132.00 | 1193.00 | 1153.33 | 1706.00 |
| UBE2D1 | 449.00 | 347.00 | 420.00 | 405.33 | 525.00 |
| UBE2D2 | 1324.00 | 1306.00 | 1432.00 | 1354.00 | 1727.00 |
| UBE2D3 | 4271.00 | 3976.00 | 4488.00 | 4245.00 | 4894.00 |
| UBE2D4 | 500.00 | 425.00 | 482.00 | 469.00 | 641.00 |
| UBE2E1 | 2019.00 | 1958.00 | 2185.00 | 2054.00 | 2743.00 |
| UBE2E2 | 453.00 | 454.00 | 542.00 | 483.00 | 721.00 |
| UBE2E3 | 828.00 | 820.00 | 895.00 | 847.67 | 1124.00 |
| UBE2E4P | 40.00 | 33.00 | 30.00 | 34.33 | 38.00 |
| UBE2F | 423.00 | 374.00 | 387.00 | 394.67 | 507.00 |
| UBE2G1 | 1186.00 | 1134.00 | 1213.00 | 1177.67 | 1530.00 |
| UBE2G2 | 3690.00 | 3523.00 | 3822.00 | 3678.33 | 3885.00 |
| UBE2H | 2714.00 | 2459.00 | 2805.00 | 2659.33 | 2220.00 |
| UBE2I | 1947.00 | 2032.00 | 2058.00 | 2012.33 | 2656.00 |
| UBE2J1 | 1302.00 | 1168.00 | 1318.00 | 1262.67 | 1582.00 |
| UBE2J2 | 1124.00 | 1144.00 | 1188.00 | 1152.00 | 1266.00 |
| UBE2K | 2823.00 | 2729.00 | 3067.00 | 2873.00 | 3346.00 |
| UBE2L3 | 1275.00 | 1174.00 | 1261.00 | 1236.67 | 1527.00 |
| UBE2L6 | 78.00 | 95.00 | 108.00 | 93.67 | 102.00 |
| UBE2M | 1278.00 | 1276.00 | 1412.00 | 1322.00 | 1804.00 |
| UBE2MP1 | 54.00 | 42.00 | 52.00 | 49.33 | 60.00 |
| UBE2N | 1559.00 | 1541.00 | 1605.00 | 1568.33 | 2034.00 |
| UBE2O | 1618.00 | 1783.00 | 1750.00 | 1717.00 | 1465.00 |
| UBE2Q1 | 1725.00 | 1704.00 | 1721.00 | 1716.67 | 1965.00 |
| UBE2Q2 | 666.00 | 616.00 | 747.00 | 676.33 | 810.00 |
| UBE2Q2P1 | 56.00 | 35.00 | 46.00 | 45.67 | 34.00 |
| UBE2R2 | 1874.00 | 1679.00 | 1910.00 | 1821.00 | 2019.00 |
| UBE2S | 1406.00 | 1344.00 | 1469.00 | 1406.33 | 1955.00 |
| UBE2T | 383.00 | 374.00 | 370.00 | 375.67 | 619.00 |
| UBE2V1 | 89.00 | 82.00 | 93.00 | 88.00 | 78.00 |
| UBE2V2 | 839.00 | 781.00 | 914.00 | 844.67 | 1249.00 |
| UBE2W | 945.00 | 863.00 | 1138.00 | 982.00 | 1185.00 |
| UBE2Z | 2915.00 | 2800.00 | 3051.00 | 2922.00 | 3373.00 |
| UBE3A | 2199.00 | 2046.00 | 2399.00 | 2214.67 | 2339.00 |
| UBE3B | 1561.00 | 1507.00 | 1574.00 | 1547.33 | 1067.00 |
| UBE3C | 3731.00 | 3448.00 | 3780.00 | 3653.00 | 2924.00 |
| UBE3D | 191.00 | 174.00 | 195.00 | 186.67 | 281.00 |

| | | | | | |
|---|---|---|---|---|---|
| UBE4A | 2240.00 | 2013.00 | 2231.00 | 2161.33 | 2112.00 |
| UBE4B | 1421.00 | 1343.00 | 1432.00 | 1398.67 | 1282.00 |
| UBFD1 | 1748.00 | 1509.00 | 1738.00 | 1665.00 | 1649.00 |
| UBIAD1 | 763.00 | 779.00 | 744.00 | 762.00 | 797.00 |
| UBL3 | 709.00 | 586.00 | 758.00 | 684.33 | 639.00 |
| UBL4A | 1038.00 | 1002.00 | 1023.00 | 1021.00 | 1207.00 |
| UBL5 | 856.00 | 938.00 | 955.00 | 916.33 | 1195.00 |
| UBL7 | 934.00 | 933.00 | 1028.00 | 965.00 | 1255.00 |
| UBL7-AS1 | 86.00 | 88.00 | 113.00 | 95.67 | 101.00 |
| UBLCP1 | 956.00 | 818.00 | 1015.00 | 929.67 | 1112.00 |
| UBN1 | 1702.00 | 1661.00 | 1861.00 | 1741.33 | 1362.00 |
| UBN2 | 1162.00 | 1159.00 | 1288.00 | 1203.00 | 958.00 |
| UBOX5 | 297.00 | 246.00 | 261.00 | 268.00 | 295.00 |
| UBP1 | 2677.00 | 2618.00 | 2902.00 | 2732.33 | 3252.00 |
| UBQLN1 | 2483.00 | 2329.00 | 2731.00 | 2514.33 | 3078.00 |
| UBQLN2 | 433.00 | 446.00 | 497.00 | 458.67 | 512.00 |
| UBQLN4 | 1401.00 | 1374.00 | 1469.00 | 1414.67 | 1337.00 |
| UBQLNL | 7.00 | 10.00 | 20.00 | 12.33 | 19.00 |
| UBR1 | 1125.00 | 1013.00 | 1189.00 | 1109.00 | 1167.00 |
| UBR2 | 1544.00 | 1356.00 | 1602.00 | 1500.67 | 1308.00 |
| UBR3 | 917.00 | 773.00 | 901.00 | 863.67 | 777.00 |
| UBR4 | 7322.00 | 6739.00 | 7576.00 | 7212.33 | 5143.00 |
| UBR5 | 5061.00 | 4524.00 | 5380.00 | 4988.33 | 4115.00 |
| UBR5-AS1 | 35.00 | 31.00 | 32.00 | 32.67 | 30.00 |
| UBTD1 | 460.00 | 404.00 | 402.00 | 422.00 | 450.00 |
| UBTD2 | 426.00 | 418.00 | 484.00 | 442.67 | 554.00 |
| UBTF | 2182.00 | 2234.00 | 2426.00 | 2280.67 | 2419.00 |
| UBXN1 | 1840.00 | 1914.00 | 1978.00 | 1910.67 | 2042.00 |
| UBXN11 | 342.00 | 419.00 | 340.00 | 367.00 | 334.00 |
| UBXN2A | 189.00 | 173.00 | 191.00 | 184.33 | 213.00 |
| UBXN2B | 735.00 | 659.00 | 823.00 | 739.00 | 775.00 |
| UBXN4 | 3333.00 | 3254.00 | 3507.00 | 3364.67 | 3459.00 |
| UBXN6 | 1535.00 | 1496.00 | 1634.00 | 1555.00 | 1748.00 |
| UBXN7 | 511.00 | 415.00 | 480.00 | 468.67 | 332.00 |
| UBXN8 | 515.00 | 513.00 | 511.00 | 513.00 | 593.00 |
| UCHL1 | 12287.00 | 12251.00 | 12616.00 | 12384.67 | 15036.00 |
| UCHL3 | 849.00 | 838.00 | 873.00 | 853.33 | 1074.00 |
| UCHL5 | 1129.00 | 1069.00 | 1177.00 | 1125.00 | 1483.00 |
| UCK1 | 480.00 | 487.00 | 450.00 | 472.33 | 492.00 |
| UCK2 | 1980.00 | 1863.00 | 2131.00 | 1991.33 | 2624.00 |
| UCKL1 | 1218.00 | 1255.00 | 1388.00 | 1287.00 | 1286.00 |
| UCKL1-AS1 | 41.00 | 44.00 | 68.00 | 51.00 | 49.00 |
| UCN | 43.00 | 48.00 | 49.00 | 46.67 | 29.00 |
| UCN2 | 17.00 | 16.00 | 17.00 | 16.67 | 6.00 |
| UCP2 | 3.00 | 5.00 | 7.00 | 5.00 | 9.00 |
| UCP3 | 30.00 | 20.00 | 23.00 | 24.33 | 18.00 |
| UEVLD | 293.00 | 228.00 | 281.00 | 267.33 | 258.00 |
| UFC1 | 1209.00 | 1204.00 | 1375.00 | 1262.67 | 1650.00 |

| | | | | | |
|---|---|---|---|---|---|
| UFD1L | 517.00 | 507.00 | 592.00 | 538.67 | 718.00 |
| UFL1 | 1525.00 | 1274.00 | 1454.00 | 1417.67 | 1460.00 |
| UFM1 | 1689.00 | 1587.00 | 1892.00 | 1722.67 | 2147.00 |
| UFSP1 | 34.00 | 42.00 | 44.00 | 40.00 | 49.00 |
| UFSP2 | 363.00 | 312.00 | 376.00 | 350.33 | 448.00 |
| UGCG | 5696.00 | 5371.00 | 5598.00 | 5555.00 | 5717.00 |
| UGDH | 2061.00 | 1969.00 | 2141.00 | 2057.00 | 1660.00 |
| UGDH-AS1 | 127.00 | 87.00 | 131.00 | 115.00 | 96.00 |
| UGGT1 | 4352.00 | 3964.00 | 4200.00 | 4172.00 | 3568.00 |
| UGGT2 | 1155.00 | 1068.00 | 1252.00 | 1158.33 | 1135.00 |
| UGP2 | 1349.00 | 1243.00 | 1328.00 | 1306.67 | 1241.00 |
| UGT8 | 266.00 | 230.00 | 258.00 | 251.33 | 237.00 |
| UHMK1 | 3307.00 | 2775.00 | 3475.00 | 3185.67 | 2569.00 |
| UHRF1 | 1216.00 | 1156.00 | 1120.00 | 1164.00 | 1530.00 |
| UHRF1BP1 | 1921.00 | 1798.00 | 1963.00 | 1894.00 | 1768.00 |
| UHRF1BP1L | 368.00 | 314.00 | 388.00 | 356.67 | 387.00 |
| UHRF2 | 1813.00 | 1796.00 | 1894.00 | 1834.33 | 1815.00 |
| UIMC1 | 1030.00 | 1009.00 | 1074.00 | 1037.67 | 1137.00 |
| ULBP1 | 35.00 | 21.00 | 32.00 | 29.33 | 17.00 |
| ULBP2 | 235.00 | 195.00 | 186.00 | 205.33 | 209.00 |
| ULBP3 | 87.00 | 86.00 | 77.00 | 83.33 | 64.00 |
| ULK1 | 1025.00 | 1066.00 | 1146.00 | 1079.00 | 983.00 |
| ULK2 | 397.00 | 417.00 | 405.00 | 406.33 | 372.00 |
| ULK3 | 1250.00 | 1263.00 | 1256.00 | 1256.33 | 1541.00 |
| ULK4 | 102.00 | 94.00 | 122.00 | 106.00 | 129.00 |
| UMAD1 | 268.00 | 191.00 | 266.00 | 241.67 | 256.00 |
| UMPS | 955.00 | 872.00 | 916.00 | 914.33 | 992.00 |
| UNC119 | 434.00 | 446.00 | 489.00 | 456.33 | 578.00 |
| UNC119B | 1559.00 | 1442.00 | 1531.00 | 1510.67 | 1542.00 |
| UNC13A | 8.00 | 1.00 | 2.00 | 3.67 | 3.00 |
| UNC13B | 983.00 | 930.00 | 1095.00 | 1002.67 | 821.00 |
| UNC13D | 67.00 | 68.00 | 67.00 | 67.33 | 65.00 |
| UNC45A | 1832.00 | 1857.00 | 2049.00 | 1912.67 | 1925.00 |
| UNC50 | 1242.00 | 1203.00 | 1259.00 | 1234.67 | 1334.00 |
| UNC5C | 5.00 | 3.00 | 7.00 | 5.00 | 3.00 |
| UNC5CL | 44.00 | 41.00 | 45.00 | 43.33 | 20.00 |
| UNC93B1 | 499.00 | 505.00 | 518.00 | 507.33 | 552.00 |
| UNG | 285.00 | 302.00 | 328.00 | 305.00 | 525.00 |
| UNK | 967.00 | 960.00 | 1023.00 | 983.33 | 1184.00 |
| UNKL | 681.00 | 756.00 | 712.00 | 716.33 | 896.00 |
| UOX | 3.00 | 3.00 | 2.00 | 2.67 | 5.00 |
| UPF1 | 3285.00 | 3128.00 | 3347.00 | 3253.33 | 3259.00 |
| UPF2 | 1565.00 | 1479.00 | 1584.00 | 1542.67 | 1699.00 |
| UPF3A | 851.00 | 890.00 | 948.00 | 896.33 | 930.00 |
| UPF3B | 461.00 | 425.00 | 420.00 | 435.33 | 538.00 |
| UPK3B | 3.00 | 0.00 | 3.00 | 2.00 | 1.00 |
| UPK3BL | 21.00 | 31.00 | 17.00 | 23.00 | 40.00 |
| UPP1 | 2979.00 | 2956.00 | 2994.00 | 2976.33 | 3242.00 |

| | | | | | |
|---|---|---|---|---|---|
| UPRT | 179.00 | 170.00 | 188.00 | 179.00 | 191.00 |
| UQCC1 | 1966.00 | 1989.00 | 2148.00 | 2034.33 | 2073.00 |
| UQCC2 | 690.00 | 734.00 | 760.00 | 728.00 | 781.00 |
| UQCC3 | 302.00 | 346.00 | 365.00 | 337.67 | 394.00 |
| UQCR10 | 689.00 | 676.00 | 698.00 | 687.67 | 1022.00 |
| UQCR11 | 1203.00 | 1177.00 | 1242.00 | 1207.33 | 1674.00 |
| UQCRB | 2833.00 | 2821.00 | 2857.00 | 2837.00 | 3837.00 |
| UQCRC1 | 2468.00 | 2382.00 | 2504.00 | 2451.33 | 2619.00 |
| UQCRC2 | 2586.00 | 2343.00 | 2782.00 | 2570.33 | 2955.00 |
| UQCRFS1 | 861.00 | 819.00 | 1018.00 | 899.33 | 1267.00 |
| UQCRH | 2563.00 | 2715.00 | 2979.00 | 2752.33 | 4114.00 |
| UQCRHL | 98.00 | 113.00 | 138.00 | 116.33 | 139.00 |
| UQCRQ | 1121.00 | 1238.00 | 1219.00 | 1192.67 | 1588.00 |
| URB1 | 3775.00 | 3479.00 | 3952.00 | 3735.33 | 3193.00 |
| URB1-AS1 | 58.00 | 55.00 | 41.00 | 51.33 | 53.00 |
| URB2 | 848.00 | 807.00 | 872.00 | 842.33 | 809.00 |
| URGCP | 1598.00 | 1625.00 | 1873.00 | 1698.67 | 1915.00 |
| URI1 | 1436.00 | 1355.00 | 1586.00 | 1459.00 | 1811.00 |
| URM1 | 1513.00 | 1398.00 | 1563.00 | 1491.33 | 1865.00 |
| UROD | 2623.00 | 2745.00 | 2746.00 | 2704.67 | 3227.00 |
| UROS | 710.00 | 673.00 | 729.00 | 704.00 | 806.00 |
| USB1 | 975.00 | 917.00 | 1026.00 | 972.67 | 1029.00 |
| USE1 | 408.00 | 381.00 | 414.00 | 401.00 | 497.00 |
| USF1 | 839.00 | 876.00 | 994.00 | 903.00 | 1011.00 |
| USF2 | 1012.00 | 1041.00 | 1130.00 | 1061.00 | 1233.00 |
| USH2A | 7.00 | 4.00 | 5.00 | 5.33 | 7.00 |
| USMG5 | 1275.00 | 1263.00 | 1297.00 | 1278.33 | 1822.00 |
| USO1 | 2164.00 | 2032.00 | 2204.00 | 2133.33 | 2211.00 |
| USP1 | 1237.00 | 1134.00 | 1199.00 | 1190.00 | 1585.00 |
| USP10 | 3311.00 | 3030.00 | 3414.00 | 3251.67 | 3505.00 |
| USP11 | 1471.00 | 1501.00 | 1608.00 | 1526.67 | 1607.00 |
| USP12 | 955.00 | 846.00 | 1008.00 | 936.33 | 824.00 |
| USP12-AS2 | 3.00 | 8.00 | 6.00 | 5.67 | 3.00 |
| USP13 | 1206.00 | 1167.00 | 1341.00 | 1238.00 | 1450.00 |
| USP14 | 4473.00 | 4033.00 | 4557.00 | 4354.33 | 5243.00 |
| USP15 | 1375.00 | 1233.00 | 1424.00 | 1344.00 | 1438.00 |
| USP16 | 1010.00 | 1031.00 | 1120.00 | 1053.67 | 1216.00 |
| USP18 | 17.00 | 11.00 | 23.00 | 17.00 | 17.00 |
| USP19 | 2374.00 | 2287.00 | 2559.00 | 2406.67 | 2232.00 |
| USP20 | 1465.00 | 1521.00 | 1497.00 | 1494.33 | 1540.00 |
| USP21 | 1007.00 | 1046.00 | 1093.00 | 1048.67 | 1146.00 |
| USP22 | 5126.00 | 4829.00 | 5368.00 | 5107.67 | 5242.00 |
| USP24 | 2991.00 | 2684.00 | 3031.00 | 2902.00 | 2699.00 |
| USP25 | 836.00 | 832.00 | 1023.00 | 897.00 | 1109.00 |
| USP27X | 90.00 | 104.00 | 119.00 | 104.33 | 129.00 |
| USP27X-AS1 | 19.00 | 21.00 | 23.00 | 21.00 | 17.00 |
| USP28 | 918.00 | 955.00 | 960.00 | 944.33 | 878.00 |
| USP2-AS1 | 17.00 | 25.00 | 23.00 | 21.67 | 16.00 |

| | | | | | |
|---|---|---|---|---|---|
| USP3 | 1357.00 | 1316.00 | 1389.00 | 1354.00 | 1491.00 |
| USP30 | 319.00 | 348.00 | 344.00 | 337.00 | 336.00 |
| USP31 | 725.00 | 671.00 | 877.00 | 757.67 | 804.00 |
| USP32 | 1791.00 | 1684.00 | 1925.00 | 1800.00 | 1752.00 |
| USP32P1 | 2157.00 | 2170.00 | 2280.00 | 2202.33 | 1930.00 |
| USP32P2 | 14.00 | 19.00 | 30.00 | 21.00 | 20.00 |
| USP33 | 1738.00 | 1625.00 | 1815.00 | 1726.00 | 1959.00 |
| USP34 | 3186.00 | 2992.00 | 3428.00 | 3202.00 | 3199.00 |
| USP35 | 292.00 | 279.00 | 303.00 | 291.33 | 242.00 |
| USP36 | 2800.00 | 2586.00 | 2882.00 | 2756.00 | 2115.00 |
| USP37 | 286.00 | 285.00 | 333.00 | 301.33 | 290.00 |
| USP38 | 395.00 | 325.00 | 407.00 | 375.67 | 330.00 |
| USP39 | 1113.00 | 1091.00 | 1168.00 | 1124.00 | 1343.00 |
| USP3-AS1 | 35.00 | 27.00 | 27.00 | 29.67 | 24.00 |
| USP4 | 1623.00 | 1532.00 | 1680.00 | 1611.67 | 1671.00 |
| USP40 | 2321.00 | 2153.00 | 2392.00 | 2288.67 | 2283.00 |
| USP42 | 610.00 | 572.00 | 640.00 | 607.33 | 619.00 |
| USP45 | 837.00 | 716.00 | 870.00 | 807.67 | 640.00 |
| USP46 | 542.00 | 501.00 | 561.00 | 534.67 | 547.00 |
| USP46-AS1 | 25.00 | 27.00 | 17.00 | 23.00 | 26.00 |
| USP47 | 2221.00 | 1970.00 | 2324.00 | 2171.67 | 2378.00 |
| USP48 | 1934.00 | 1923.00 | 2100.00 | 1985.67 | 2036.00 |
| USP49 | 484.00 | 550.00 | 521.00 | 518.33 | 452.00 |
| USP5 | 2608.00 | 2372.00 | 2561.00 | 2513.67 | 2659.00 |
| USP51 | 33.00 | 34.00 | 40.00 | 35.67 | 47.00 |
| USP53 | 1386.00 | 1286.00 | 1473.00 | 1381.67 | 1314.00 |
| USP54 | 1045.00 | 992.00 | 1065.00 | 1034.00 | 923.00 |
| USP6 | 43.00 | 47.00 | 33.00 | 41.00 | 61.00 |
| USP6NL | 569.00 | 550.00 | 626.00 | 581.67 | 666.00 |
| USP7 | 3194.00 | 3083.00 | 3438.00 | 3238.33 | 3370.00 |
| USP8 | 1295.00 | 1119.00 | 1391.00 | 1268.33 | 1344.00 |
| USP9X | 3799.00 | 3556.00 | 4199.00 | 3851.33 | 3432.00 |
| USPL1 | 800.00 | 803.00 | 873.00 | 825.33 | 774.00 |
| UST | 951.00 | 932.00 | 914.00 | 932.33 | 927.00 |
| UTP11L | 921.00 | 794.00 | 914.00 | 876.33 | 1181.00 |
| UTP14A | 1369.00 | 1345.00 | 1647.00 | 1453.67 | 1705.00 |
| UTP14C | 863.00 | 788.00 | 856.00 | 835.67 | 744.00 |
| UTP15 | 658.00 | 630.00 | 708.00 | 665.33 | 762.00 |
| UTP18 | 1902.00 | 1793.00 | 2086.00 | 1927.00 | 2277.00 |
| UTP20 | 2042.00 | 1977.00 | 2075.00 | 2031.33 | 1871.00 |
| UTP23 | 598.00 | 546.00 | 663.00 | 602.33 | 763.00 |
| UTP3 | 889.00 | 953.00 | 912.00 | 918.00 | 1126.00 |
| UTP6 | 1535.00 | 1502.00 | 1620.00 | 1552.33 | 1885.00 |
| UTRN | 2008.00 | 1616.00 | 1926.00 | 1850.00 | 1026.00 |
| UVRAG | 802.00 | 710.00 | 778.00 | 763.33 | 851.00 |
| UVSSA | 516.00 | 437.00 | 538.00 | 497.00 | 445.00 |
| UXS1 | 864.00 | 831.00 | 902.00 | 865.67 | 1024.00 |
| UXT | 574.00 | 508.00 | 550.00 | 544.00 | 643.00 |

| | | | | | |
|---|---|---|---|---|---|
| UXT-AS1 | 8.00 | 11.00 | 12.00 | 10.33 | 7.00 |
| VAC14 | 1240.00 | 1220.00 | 1343.00 | 1267.67 | 1217.00 |
| VAMP1 | 482.00 | 498.00 | 521.00 | 500.33 | 433.00 |
| VAMP2 | 1593.00 | 1537.00 | 1751.00 | 1627.00 | 1962.00 |
| VAMP3 | 1356.00 | 1249.00 | 1483.00 | 1362.67 | 1637.00 |
| VAMP4 | 752.00 | 705.00 | 813.00 | 756.67 | 770.00 |
| VAMP5 | 23.00 | 16.00 | 21.00 | 20.00 | 26.00 |
| VAMP8 | 5.00 | 5.00 | 2.00 | 4.00 | 1.00 |
| VANGL1 | 1029.00 | 938.00 | 986.00 | 984.33 | 947.00 |
| VAPA | 3846.00 | 3457.00 | 4207.00 | 3836.67 | 4861.00 |
| VAPB | 1598.00 | 1454.00 | 1618.00 | 1556.67 | 1808.00 |
| VARS | 10.00 | 7.00 | 2.00 | 6.33 | 5.00 |
| VARS2 | 181.00 | 162.00 | 162.00 | 168.33 | 138.00 |
| VASH1 | 166.00 | 163.00 | 189.00 | 172.67 | 172.00 |
| VASH2 | 7.00 | 3.00 | 5.00 | 5.00 | 6.00 |
| VASP | 1810.00 | 1748.00 | 1775.00 | 1777.67 | 1756.00 |
| VAT1 | 5429.00 | 5398.00 | 5665.00 | 5497.33 | 4464.00 |
| VAV2 | 1012.00 | 1001.00 | 1161.00 | 1058.00 | 982.00 |
| VAV3 | 4.00 | 5.00 | 6.00 | 5.00 | 3.00 |
| VAX2 | 45.00 | 64.00 | 47.00 | 52.00 | 74.00 |
| VBP1 | 826.00 | 720.00 | 859.00 | 801.67 | 1158.00 |
| VCAM1 | 1807.00 | 1757.00 | 1768.00 | 1777.33 | 2633.00 |
| VCAN | 483.00 | 421.00 | 495.00 | 466.33 | 416.00 |
| VCL | 11048.00 | 10190.00 | 11219.00 | 10819.00 | 9974.00 |
| VCP | 7581.00 | 7431.00 | 7625.00 | 7545.67 | 8400.00 |
| VCPIP1 | 690.00 | 615.00 | 729.00 | 678.00 | 671.00 |
| VCPKMT | 273.00 | 278.00 | 310.00 | 287.00 | 315.00 |
| VDAC1 | 4663.00 | 4345.00 | 4830.00 | 4612.67 | 6886.00 |
| VDAC2 | 2344.00 | 2390.00 | 2514.00 | 2416.00 | 3189.00 |
| VDAC3 | 1390.00 | 1364.00 | 1424.00 | 1392.67 | 1841.00 |
| VDR | 192.00 | 164.00 | 184.00 | 180.00 | 99.00 |
| VEGFA | 8362.00 | 8343.00 | 8572.00 | 8425.67 | 6382.00 |
| VEGFB | 1332.00 | 1421.00 | 1475.00 | 1409.33 | 1425.00 |
| VEGFC | 1760.00 | 1673.00 | 1660.00 | 1697.67 | 1457.00 |
| VEPH1 | 1461.00 | 1243.00 | 1532.00 | 1412.00 | 1815.00 |
| VEZF1 | 1679.00 | 1567.00 | 1877.00 | 1707.67 | 1794.00 |
| VEZT | 1326.00 | 1295.00 | 1348.00 | 1323.00 | 1474.00 |
| VGF | 214.00 | 263.00 | 241.00 | 239.33 | 190.00 |
| VGLL2 | 49.00 | 44.00 | 42.00 | 45.00 | 62.00 |
| VGLL3 | 627.00 | 521.00 | 745.00 | 631.00 | 402.00 |
| VGLL4 | 1039.00 | 1033.00 | 1209.00 | 1093.67 | 1105.00 |
| VHL | 1216.00 | 1230.00 | 1262.00 | 1236.00 | 1425.00 |
| VILL | 55.00 | 53.00 | 54.00 | 54.00 | 59.00 |
| VIM | 101688.00 | 98800.00 | 108880.00 | 103122.67 | 138701.00 |
| VIM-AS1 | 75.00 | 103.00 | 93.00 | 90.33 | 137.00 |
| VIMP | 1199.00 | 1141.00 | 1113.00 | 1151.00 | 1255.00 |
| VIP | 17.00 | 13.00 | 18.00 | 16.00 | 10.00 |
| VIPAS39 | 576.00 | 609.00 | 664.00 | 616.33 | 646.00 |

| | | | | | |
|---|---|---|---|---|---|
| VIPR1 | 157.00 | 170.00 | 162.00 | 163.00 | 62.00 |
| VKORC1 | 2292.00 | 2251.00 | 2055.00 | 2199.33 | 2385.00 |
| VKORC1L1 | 1281.00 | 1244.00 | 1347.00 | 1290.67 | 1391.00 |
| VLDLR | 1518.00 | 1480.00 | 1430.00 | 1476.00 | 1327.00 |
| VLDLR-AS1 | 122.00 | 102.00 | 108.00 | 110.67 | 155.00 |
| VMA21 | 1027.00 | 960.00 | 1118.00 | 1035.00 | 1526.00 |
| VMAC | 101.00 | 89.00 | 110.00 | 100.00 | 67.00 |
| VMO1 | 7.00 | 4.00 | 14.00 | 8.33 | 9.00 |
| VMP1 | 3925.00 | 3917.00 | 4067.00 | 3969.67 | 4298.00 |
| VN1R1 | 6.00 | 6.00 | 7.00 | 6.33 | 2.00 |
| VNN1 | 345.00 | 344.00 | 384.00 | 357.67 | 486.00 |
| VOPP1 | 807.00 | 827.00 | 817.00 | 817.00 | 907.00 |
| VPRBP | 1340.00 | 1289.00 | 1483.00 | 1370.67 | 1328.00 |
| VPS11 | 950.00 | 859.00 | 1060.00 | 956.33 | 995.00 |
| VPS13A | 1457.00 | 1410.00 | 1551.00 | 1472.67 | 1246.00 |
| VPS13B | 1530.00 | 1406.00 | 1699.00 | 1545.00 | 1266.00 |
| VPS13C | 1782.00 | 1684.00 | 1977.00 | 1814.33 | 1661.00 |
| VPS13D | 1245.00 | 1137.00 | 1239.00 | 1207.00 | 929.00 |
| VPS16 | 1100.00 | 1182.00 | 1211.00 | 1164.33 | 1289.00 |
| VPS18 | 739.00 | 764.00 | 820.00 | 774.33 | 689.00 |
| VPS25 | 1354.00 | 1277.00 | 1408.00 | 1346.33 | 1594.00 |
| VPS26A | 1759.00 | 1596.00 | 1883.00 | 1746.00 | 2107.00 |
| VPS26B | 825.00 | 749.00 | 865.00 | 813.00 | 844.00 |
| VPS28 | 1964.00 | 1922.00 | 2038.00 | 1974.67 | 2530.00 |
| VPS29 | 927.00 | 941.00 | 1036.00 | 968.00 | 1282.00 |
| VPS33A | 780.00 | 753.00 | 801.00 | 778.00 | 835.00 |
| VPS33B | 374.00 | 390.00 | 353.00 | 372.33 | 351.00 |
| VPS35 | 2624.00 | 2388.00 | 2722.00 | 2578.00 | 3017.00 |
| VPS36 | 1115.00 | 1132.00 | 1305.00 | 1184.00 | 1365.00 |
| VPS37A | 607.00 | 549.00 | 674.00 | 610.00 | 675.00 |
| VPS37B | 1196.00 | 1227.00 | 1345.00 | 1256.00 | 1340.00 |
| VPS37C | 518.00 | 469.00 | 538.00 | 508.33 | 637.00 |
| VPS37D | 28.00 | 26.00 | 34.00 | 29.33 | 49.00 |
| VPS39 | 2033.00 | 1830.00 | 1970.00 | 1944.33 | 1718.00 |
| VPS41 | 1378.00 | 1227.00 | 1415.00 | 1340.00 | 1382.00 |
| VPS45 | 747.00 | 815.00 | 802.00 | 788.00 | 932.00 |
| VPS4A | 1874.00 | 1834.00 | 2002.00 | 1903.33 | 2088.00 |
| VPS4B | 893.00 | 749.00 | 903.00 | 848.33 | 855.00 |
| VPS50 | 586.00 | 547.00 | 703.00 | 612.00 | 637.00 |
| VPS51 | 2130.00 | 2257.00 | 2447.00 | 2278.00 | 2454.00 |
| VPS52 | 53.00 | 41.00 | 38.00 | 44.00 | 53.00 |
| VPS53 | 704.00 | 671.00 | 647.00 | 674.00 | 722.00 |
| VPS54 | 1176.00 | 1035.00 | 1224.00 | 1145.00 | 1240.00 |
| VPS72 | 1058.00 | 1022.00 | 1078.00 | 1052.67 | 1331.00 |
| VPS8 | 1219.00 | 1242.00 | 1277.00 | 1246.00 | 1130.00 |
| VPS9D1 | 485.00 | 506.00 | 494.00 | 495.00 | 522.00 |
| VPS9D1-AS1 | 719.00 | 677.00 | 756.00 | 717.33 | 640.00 |
| VRK1 | 653.00 | 617.00 | 604.00 | 624.67 | 884.00 |

| | | | | | |
|---|---|---|---|---|---|
| VRK2 | 499.00 | 473.00 | 578.00 | 516.67 | 588.00 |
| VRK3 | 615.00 | 641.00 | 594.00 | 616.67 | 599.00 |
| VSIG10 | 572.00 | 580.00 | 607.00 | 586.33 | 615.00 |
| VSIG10L | 19.00 | 23.00 | 14.00 | 18.67 | 20.00 |
| VSIG2 | 5.00 | 5.00 | 8.00 | 6.00 | 5.00 |
| VSIG8 | 8.00 | 7.00 | 3.00 | 6.00 | 5.00 |
| VTA1 | 799.00 | 664.00 | 769.00 | 744.00 | 1126.00 |
| VTI1A | 628.00 | 629.00 | 740.00 | 665.67 | 599.00 |
| VTI1B | 898.00 | 891.00 | 1030.00 | 939.67 | 1026.00 |
| VTN | 45.00 | 39.00 | 34.00 | 39.33 | 34.00 |
| VWA5A | 5.00 | 17.00 | 11.00 | 11.00 | 18.00 |
| VWA8 | 843.00 | 794.00 | 868.00 | 835.00 | 794.00 |
| VWA9 | 1291.00 | 1220.00 | 1308.00 | 1273.00 | 1494.00 |
| VWCE | 5.00 | 9.00 | 5.00 | 6.33 | 12.00 |
| VWDE | 5.00 | 7.00 | 18.00 | 10.00 | 5.00 |
| VWF | 4.00 | 0.00 | 2.00 | 2.00 | 3.00 |
| WAC | 4314.00 | 4153.00 | 4697.00 | 4388.00 | 4781.00 |
| WAC-AS1 | 600.00 | 583.00 | 599.00 | 594.00 | 673.00 |
| WAPL | 1898.00 | 1740.00 | 2050.00 | 1896.00 | 1952.00 |
| WARS | 5469.00 | 5021.00 | 5429.00 | 5306.33 | 5751.00 |
| WARS2 | 295.00 | 277.00 | 298.00 | 290.00 | 353.00 |
| WAS | 6.00 | 5.00 | 1.00 | 4.00 | 0.00 |
| WASF1 | 824.00 | 674.00 | 798.00 | 765.33 | 725.00 |
| WASF2 | 2429.00 | 2306.00 | 2675.00 | 2470.00 | 1871.00 |
| WASF3 | 556.00 | 561.00 | 649.00 | 588.67 | 619.00 |
| WASH1 | 226.00 | 234.00 | 233.00 | 231.00 | 222.00 |
| WASH2P | 149.00 | 200.00 | 208.00 | 185.67 | 189.00 |
| WASH3P | 220.00 | 236.00 | 171.00 | 209.00 | 198.00 |
| WASH5P | 12.00 | 23.00 | 13.00 | 16.00 | 12.00 |
| WASH7P | 117.00 | 119.00 | 132.00 | 122.67 | 125.00 |
| WASL | 962.00 | 790.00 | 1081.00 | 944.33 | 933.00 |
| WBP1 | 160.00 | 140.00 | 203.00 | 167.67 | 152.00 |
| WBP11 | 2179.00 | 2003.00 | 2197.00 | 2126.33 | 2448.00 |
| WBP1L | 826.00 | 814.00 | 805.00 | 815.00 | 806.00 |
| WBP2 | 1825.00 | 1780.00 | 1784.00 | 1796.33 | 1942.00 |
| WBP4 | 343.00 | 346.00 | 360.00 | 349.67 | 412.00 |
| WBP5 | 1396.00 | 1515.00 | 1541.00 | 1484.00 | 1756.00 |
| WBSCR16 | 917.00 | 877.00 | 897.00 | 897.00 | 1055.00 |
| WBSCR22 | 2062.00 | 1996.00 | 1990.00 | 2016.00 | 2452.00 |
| WBSCR27 | 621.00 | 562.00 | 666.00 | 616.33 | 641.00 |
| WDFY1 | 1736.00 | 1495.00 | 1914.00 | 1715.00 | 1487.00 |
| WDFY2 | 1383.00 | 1243.00 | 1267.00 | 1297.67 | 1585.00 |
| WDFY3 | 626.00 | 622.00 | 622.00 | 623.33 | 489.00 |
| WDFY3-AS2 | 40.00 | 30.00 | 34.00 | 34.67 | 18.00 |
| WDHD1 | 626.00 | 601.00 | 628.00 | 618.33 | 755.00 |
| WDPCP | 95.00 | 105.00 | 89.00 | 96.33 | 91.00 |
| WDR1 | 5271.00 | 5078.00 | 5454.00 | 5267.67 | 4886.00 |
| WDR11 | 2076.00 | 1823.00 | 2115.00 | 2004.67 | 2018.00 |

| | | | | | |
|---|---|---|---|---|---|
| WDR12 | 1732.00 | 1670.00 | 1830.00 | 1744.00 | 2219.00 |
| WDR13 | 912.00 | 882.00 | 914.00 | 902.67 | 1047.00 |
| WDR17 | 463.00 | 414.00 | 516.00 | 464.33 | 559.00 |
| WDR18 | 1042.00 | 920.00 | 1066.00 | 1009.33 | 1310.00 |
| WDR19 | 2252.00 | 2081.00 | 2277.00 | 2203.33 | 1828.00 |
| WDR20 | 475.00 | 426.00 | 456.00 | 452.33 | 467.00 |
| WDR24 | 441.00 | 544.00 | 531.00 | 505.33 | 520.00 |
| WDR25 | 189.00 | 161.00 | 171.00 | 173.67 | 161.00 |
| WDR26 | 3362.00 | 3225.00 | 3598.00 | 3395.00 | 3428.00 |
| WDR27 | 2874.00 | 2885.00 | 2781.00 | 2846.67 | 2302.00 |
| WDR3 | 3166.00 | 3003.00 | 3357.00 | 3175.33 | 3431.00 |
| WDR31 | 51.00 | 59.00 | 73.00 | 61.00 | 55.00 |
| WDR33 | 2414.00 | 2444.00 | 2507.00 | 2455.00 | 2379.00 |
| WDR34 | 700.00 | 732.00 | 745.00 | 725.67 | 1121.00 |
| WDR35 | 2259.00 | 2050.00 | 2434.00 | 2247.67 | 2267.00 |
| WDR36 | 2537.00 | 2238.00 | 2613.00 | 2462.67 | 2313.00 |
| WDR37 | 988.00 | 939.00 | 1022.00 | 983.00 | 955.00 |
| WDR4 | 525.00 | 481.00 | 502.00 | 502.67 | 422.00 |
| WDR41 | 516.00 | 419.00 | 501.00 | 478.67 | 548.00 |
| WDR43 | 2761.00 | 2519.00 | 3114.00 | 2798.00 | 3072.00 |
| WDR44 | 637.00 | 522.00 | 585.00 | 581.33 | 472.00 |
| WDR45 | 1057.00 | 1064.00 | 1174.00 | 1098.33 | 1112.00 |
| WDR45B | 2432.00 | 2397.00 | 2441.00 | 2423.33 | 3039.00 |
| WDR46 | 118.00 | 153.00 | 115.00 | 128.67 | 146.00 |
| WDR47 | 362.00 | 332.00 | 401.00 | 365.00 | 373.00 |
| WDR48 | 852.00 | 819.00 | 985.00 | 885.33 | 963.00 |
| WDR5 | 1900.00 | 1960.00 | 1957.00 | 1939.00 | 2318.00 |
| WDR53 | 344.00 | 326.00 | 318.00 | 329.33 | 378.00 |
| WDR54 | 750.00 | 772.00 | 755.00 | 759.00 | 1057.00 |
| WDR55 | 707.00 | 766.00 | 775.00 | 749.33 | 674.00 |
| WDR59 | 1854.00 | 1970.00 | 2109.00 | 1977.67 | 2190.00 |
| WDR5B | 601.00 | 667.00 | 695.00 | 654.33 | 591.00 |
| WDR6 | 6399.00 | 6451.00 | 6881.00 | 6577.00 | 6087.00 |
| WDR60 | 597.00 | 543.00 | 668.00 | 602.67 | 546.00 |
| WDR61 | 708.00 | 660.00 | 731.00 | 699.67 | 853.00 |
| WDR62 | 643.00 | 658.00 | 658.00 | 653.00 | 782.00 |
| WDR63 | 11.00 | 7.00 | 13.00 | 10.33 | 10.00 |
| WDR64 | 3.00 | 3.00 | 0.00 | 2.00 | 0.00 |
| WDR66 | 374.00 | 322.00 | 434.00 | 376.67 | 1083.00 |
| WDR7 | 334.00 | 327.00 | 413.00 | 358.00 | 319.00 |
| WDR70 | 838.00 | 820.00 | 886.00 | 848.00 | 989.00 |
| WDR72 | 35.00 | 17.00 | 28.00 | 26.67 | 31.00 |
| WDR73 | 1286.00 | 1346.00 | 1256.00 | 1296.00 | 1319.00 |
| WDR74 | 1528.00 | 1578.00 | 1672.00 | 1592.67 | 1692.00 |
| WDR75 | 2791.00 | 2683.00 | 3043.00 | 2839.00 | 3176.00 |
| WDR76 | 140.00 | 140.00 | 124.00 | 134.67 | 224.00 |
| WDR77 | 1893.00 | 1768.00 | 1831.00 | 1830.67 | 2236.00 |
| WDR78 | 63.00 | 79.00 | 89.00 | 77.00 | 98.00 |

| | | | | | |
|---|---|---|---|---|---|
| WDR81 | 311.00 | 255.00 | 271.00 | 279.00 | 318.00 |
| WDR82 | 1706.00 | 1603.00 | 1827.00 | 1712.00 | 1749.00 |
| WDR83 | 195.00 | 198.00 | 205.00 | 199.33 | 269.00 |
| WDR83OS | 1482.00 | 1462.00 | 1456.00 | 1466.67 | 1846.00 |
| WDR89 | 492.00 | 450.00 | 489.00 | 477.00 | 505.00 |
| WDR90 | 1366.00 | 1398.00 | 1360.00 | 1374.67 | 1486.00 |
| WDR91 | 626.00 | 636.00 | 680.00 | 647.33 | 555.00 |
| WDR92 | 434.00 | 365.00 | 434.00 | 411.00 | 466.00 |
| WDSUB1 | 329.00 | 304.00 | 356.00 | 329.67 | 396.00 |
| WDTC1 | 921.00 | 994.00 | 999.00 | 971.33 | 1000.00 |
| WDYHV1 | 327.00 | 278.00 | 336.00 | 313.67 | 451.00 |
| WEE1 | 439.00 | 439.00 | 477.00 | 451.67 | 590.00 |
| WFDC3 | 7.00 | 6.00 | 2.00 | 5.00 | 6.00 |
| WFIKKN1 | 20.00 | 29.00 | 18.00 | 22.33 | 24.00 |
| WFS1 | 1010.00 | 1043.00 | 1012.00 | 1021.67 | 886.00 |
| WHAMM | 241.00 | 211.00 | 243.00 | 231.67 | 216.00 |
| WHAMMP1 | 85.00 | 118.00 | 114.00 | 105.67 | 87.00 |
| WHAMMP2 | 5.00 | 2.00 | 2.00 | 3.00 | 5.00 |
| WHAMMP3 | 83.00 | 108.00 | 112.00 | 101.00 | 124.00 |
| WHSC1 | 8622.00 | 8251.00 | 8732.00 | 8535.00 | 10057.00 |
| WHSC1L1 | 1291.00 | 1106.00 | 1289.00 | 1228.67 | 1116.00 |
| WIBG | 275.00 | 267.00 | 259.00 | 267.00 | 293.00 |
| WIPF1 | 703.00 | 644.00 | 810.00 | 719.00 | 683.00 |
| WIPF2 | 905.00 | 856.00 | 923.00 | 894.67 | 784.00 |
| WIPI1 | 556.00 | 544.00 | 565.00 | 555.00 | 580.00 |
| WIPI2 | 1995.00 | 1828.00 | 1975.00 | 1932.67 | 2061.00 |
| WISP1 | 25.00 | 17.00 | 29.00 | 23.67 | 39.00 |
| WIZ | 1157.00 | 1086.00 | 1202.00 | 1148.33 | 1219.00 |
| WLS | 4596.00 | 4494.00 | 4728.00 | 4606.00 | 4523.00 |
| WNK1 | 6964.00 | 6214.00 | 7463.00 | 6880.33 | 5614.00 |
| WNK3 | 8.00 | 6.00 | 8.00 | 7.33 | 8.00 |
| WNK4 | 2505.00 | 2514.00 | 2776.00 | 2598.33 | 1281.00 |
| WNT10A | 3.00 | 5.00 | 4.00 | 4.00 | 1.00 |
| WNT10B | 7.00 | 5.00 | 10.00 | 7.33 | 20.00 |
| WNT2B | 93.00 | 80.00 | 92.00 | 88.33 | 76.00 |
| WNT5A | 11.00 | 5.00 | 12.00 | 9.33 | 11.00 |
| WNT5B | 438.00 | 479.00 | 487.00 | 468.00 | 1043.00 |
| WNT7B | 4.00 | 2.00 | 9.00 | 5.00 | 0.00 |
| WRAP53 | 484.00 | 462.00 | 449.00 | 465.00 | 523.00 |
| WRAP73 | 642.00 | 656.00 | 657.00 | 651.67 | 703.00 |
| WRB | 427.00 | 390.00 | 462.00 | 426.33 | 586.00 |
| WRN | 870.00 | 783.00 | 941.00 | 864.67 | 934.00 |
| WRNIP1 | 2171.00 | 2106.00 | 2277.00 | 2184.67 | 2545.00 |
| WSB1 | 1880.00 | 1711.00 | 1808.00 | 1799.67 | 2061.00 |
| WSB2 | 812.00 | 702.00 | 823.00 | 779.00 | 1012.00 |
| WT1 | 33.00 | 30.00 | 23.00 | 28.67 | 43.00 |
| WT1-AS | 13.00 | 21.00 | 22.00 | 18.67 | 18.00 |
| WTAP | 1193.00 | 1175.00 | 1348.00 | 1238.67 | 1463.00 |

| | | | | | |
|---|---|---|---|---|---|
| WTAPP1 | 92.00 | 92.00 | 94.00 | 92.67 | 162.00 |
| WTH3DI | 18.00 | 14.00 | 24.00 | 18.67 | 18.00 |
| WTIP | 11.00 | 18.00 | 7.00 | 12.00 | 16.00 |
| WWC1 | 2609.00 | 2475.00 | 2651.00 | 2578.33 | 3350.00 |
| WWC2 | 1039.00 | 942.00 | 1106.00 | 1029.00 | 1011.00 |
| WWC2-AS2 | 5.00 | 6.00 | 9.00 | 6.67 | 6.00 |
| WWC3 | 1053.00 | 1045.00 | 1078.00 | 1058.67 | 819.00 |
| WWP1 | 781.00 | 658.00 | 850.00 | 763.00 | 869.00 |
| WWP2 | 814.00 | 896.00 | 887.00 | 865.67 | 837.00 |
| WWTR1 | 607.00 | 550.00 | 631.00 | 596.00 | 482.00 |
| WWTR1-AS1 | 9.00 | 7.00 | 12.00 | 9.33 | 14.00 |
| XAB2 | 1693.00 | 1654.00 | 1629.00 | 1658.67 | 1606.00 |
| XAF1 | 49.00 | 40.00 | 59.00 | 49.33 | 71.00 |
| XBP1 | 6581.00 | 6282.00 | 6159.00 | 6340.67 | 7182.00 |
| XDH | 189.00 | 136.00 | 163.00 | 162.67 | 618.00 |
| XIAP | 1159.00 | 1005.00 | 1300.00 | 1154.67 | 1060.00 |
| XIRP2 | 329.00 | 271.00 | 318.00 | 306.00 | 302.00 |
| XIST | 22549.00 | 22088.00 | 24496.00 | 23044.33 | 17212.00 |
| XK | 15.00 | 10.00 | 8.00 | 11.00 | 22.00 |
| XKR9 | 6.00 | 1.00 | 1.00 | 2.67 | 1.00 |
| XPA | 162.00 | 154.00 | 159.00 | 158.33 | 200.00 |
| XPC | 1510.00 | 1483.00 | 1665.00 | 1552.67 | 1274.00 |
| XPNPEP1 | 1317.00 | 1240.00 | 1348.00 | 1301.67 | 1451.00 |
| XPNPEP3 | 651.00 | 664.00 | 722.00 | 679.00 | 766.00 |
| XPO1 | 10271.00 | 10192.00 | 11526.00 | 10663.00 | 10837.00 |
| XPO4 | 1234.00 | 1023.00 | 1300.00 | 1185.67 | 1249.00 |
| XPO5 | 2884.00 | 2726.00 | 3161.00 | 2923.67 | 2971.00 |
| XPO6 | 3302.00 | 3372.00 | 3547.00 | 3407.00 | 3560.00 |
| XPO7 | 1460.00 | 1330.00 | 1573.00 | 1454.33 | 1387.00 |
| XPOT | 10375.00 | 9381.00 | 11076.00 | 10277.33 | 11245.00 |
| XPR1 | 1321.00 | 1181.00 | 1318.00 | 1273.33 | 1231.00 |
| XRCC1 | 844.00 | 832.00 | 852.00 | 842.67 | 861.00 |
| XRCC2 | 469.00 | 450.00 | 439.00 | 452.67 | 492.00 |
| XRCC3 | 790.00 | 842.00 | 841.00 | 824.33 | 948.00 |
| XRCC4 | 260.00 | 239.00 | 240.00 | 246.33 | 332.00 |
| XRCC5 | 7584.00 | 6958.00 | 7975.00 | 7505.67 | 8841.00 |
| XRCC6 | 6915.00 | 6443.00 | 7071.00 | 6809.67 | 7905.00 |
| XRN1 | 1243.00 | 1022.00 | 1259.00 | 1174.67 | 968.00 |
| XRN2 | 4443.00 | 3939.00 | 4537.00 | 4306.33 | 4645.00 |
| XRRA1 | 632.00 | 639.00 | 702.00 | 657.67 | 532.00 |
| XXYLT1 | 704.00 | 758.00 | 757.00 | 739.67 | 724.00 |
| XYLB | 231.00 | 231.00 | 229.00 | 230.33 | 219.00 |
| XYLT2 | 2046.00 | 2042.00 | 2156.00 | 2081.33 | 1823.00 |
| YAE1D1 | 287.00 | 349.00 | 371.00 | 335.67 | 428.00 |
| YAF2 | 263.00 | 237.00 | 280.00 | 260.00 | 333.00 |
| YAP1 | 14182.00 | 13148.00 | 15479.00 | 14269.67 | 14930.00 |
| YARS | 6774.00 | 6518.00 | 6926.00 | 6739.33 | 7668.00 |
| YARS2 | 823.00 | 823.00 | 868.00 | 838.00 | 920.00 |

| | | | | | |
|---|---|---|---|---|---|
| YBEY | 267.00 | 207.00 | 238.00 | 237.33 | 249.00 |
| YBX1 | 10758.00 | 10092.00 | 11133.00 | 10661.00 | 15796.00 |
| YBX3 | 4450.00 | 4369.00 | 4716.00 | 4511.67 | 6045.00 |
| YBX3P1 | 28.00 | 21.00 | 25.00 | 24.67 | 34.00 |
| YDJC | 610.00 | 627.00 | 635.00 | 624.00 | 757.00 |
| YEATS2 | 2225.00 | 2031.00 | 2433.00 | 2229.67 | 2245.00 |
| YEATS4 | 457.00 | 438.00 | 461.00 | 452.00 | 626.00 |
| YES1 | 2651.00 | 2389.00 | 2882.00 | 2640.67 | 3906.00 |
| YIF1A | 2399.00 | 2342.00 | 2132.00 | 2291.00 | 2730.00 |
| YIF1B | 1372.00 | 1294.00 | 1210.00 | 1292.00 | 1120.00 |
| YIPF1 | 528.00 | 493.00 | 501.00 | 507.33 | 582.00 |
| YIPF2 | 838.00 | 847.00 | 860.00 | 848.33 | 895.00 |
| YIPF3 | 4056.00 | 4000.00 | 3733.00 | 3929.67 | 3968.00 |
| YIPF4 | 914.00 | 803.00 | 854.00 | 857.00 | 876.00 |
| YIPF5 | 1501.00 | 1482.00 | 1538.00 | 1507.00 | 1583.00 |
| YIPF6 | 2228.00 | 2148.00 | 2256.00 | 2210.67 | 1979.00 |
| YJEFN3 | 358.00 | 421.00 | 431.00 | 403.33 | 308.00 |
| YKT6 | 2592.00 | 2513.00 | 2890.00 | 2665.00 | 2946.00 |
| YLPM1 | 2327.00 | 2134.00 | 2413.00 | 2291.33 | 2031.00 |
| YME1L1 | 4361.00 | 3954.00 | 4629.00 | 4314.67 | 5037.00 |
| YOD1 | 578.00 | 541.00 | 724.00 | 614.33 | 459.00 |
| YPEL1 | 7.00 | 9.00 | 11.00 | 9.00 | 14.00 |
| YPEL2 | 54.00 | 55.00 | 49.00 | 52.67 | 64.00 |
| YPEL3 | 225.00 | 191.00 | 230.00 | 215.33 | 193.00 |
| YPEL4 | 7.00 | 5.00 | 2.00 | 4.67 | 3.00 |
| YPEL5 | 626.00 | 556.00 | 680.00 | 620.67 | 677.00 |
| YRDC | 797.00 | 700.00 | 830.00 | 775.67 | 1049.00 |
| YTHDC1 | 4.00 | 1.00 | 2.00 | 2.33 | 1.00 |
| YTHDC2 | 1496.00 | 1407.00 | 1551.00 | 1484.67 | 1427.00 |
| YTHDF1 | 1447.00 | 1418.00 | 1465.00 | 1443.33 | 1648.00 |
| YTHDF2 | 2018.00 | 2041.00 | 2073.00 | 2044.00 | 2396.00 |
| YTHDF3 | 2102.00 | 1856.00 | 2346.00 | 2101.33 | 2272.00 |
| YTHDF3-AS1 | 6.00 | 2.00 | 9.00 | 5.67 | 19.00 |
| YWHAB | 5555.00 | 5439.00 | 5865.00 | 5619.67 | 7114.00 |
| YWHAE | 31.00 | 36.00 | 38.00 | 35.00 | 51.00 |
| YWHAEP1 | 9.00 | 7.00 | 8.00 | 8.00 | 9.00 |
| YWHAG | 3695.00 | 3342.00 | 3782.00 | 3606.33 | 4014.00 |
| YWHAH | 915.00 | 857.00 | 926.00 | 899.33 | 1187.00 |
| YWHAQ | 4483.00 | 4179.00 | 4678.00 | 4446.67 | 5430.00 |
| YWHAZ | 10152.00 | 9189.00 | 11040.00 | 10127.00 | 12545.00 |
| YY1 | 1390.00 | 1234.00 | 1520.00 | 1381.33 | 1538.00 |
| YY1AP1 | 1104.00 | 1018.00 | 1082.00 | 1068.00 | 1038.00 |
| YY2 | 95.00 | 89.00 | 114.00 | 99.33 | 83.00 |
| ZADH2 | 787.00 | 720.00 | 916.00 | 807.67 | 877.00 |
| ZAK | 1148.00 | 967.00 | 1188.00 | 1101.00 | 1087.00 |
| ZAP70 | 4.00 | 3.00 | 0.00 | 2.33 | 0.00 |
| ZAR1 | 3.00 | 0.00 | 0.00 | 1.00 | 1.00 |
| ZBBX | 8.00 | 13.00 | 5.00 | 8.67 | 9.00 |

| | | | | | |
|---|---|---|---|---|---|
| ZBED3 | 118.00 | 107.00 | 109.00 | 111.33 | 69.00 |
| ZBED3-AS1 | 30.00 | 57.00 | 49.00 | 45.33 | 57.00 |
| ZBED4 | 1411.00 | 1317.00 | 1513.00 | 1413.67 | 1362.00 |
| ZBED5 | 1279.00 | 1281.00 | 1365.00 | 1308.33 | 1298.00 |
| ZBED5-AS1 | 61.00 | 68.00 | 67.00 | 65.33 | 87.00 |
| ZBED6 | 288.00 | 190.00 | 209.00 | 229.00 | 129.00 |
| ZBED8 | 264.00 | 206.00 | 238.00 | 236.00 | 262.00 |
| ZBTB1 | 1126.00 | 1149.00 | 1190.00 | 1155.00 | 1200.00 |
| ZBTB10 | 498.00 | 395.00 | 563.00 | 485.33 | 490.00 |
| ZBTB11 | 1201.00 | 1086.00 | 1237.00 | 1174.67 | 1169.00 |
| ZBTB11-AS1 | 80.00 | 92.00 | 76.00 | 82.67 | 84.00 |
| ZBTB12 | 20.00 | 7.00 | 8.00 | 11.67 | 13.00 |
| ZBTB14 | 582.00 | 600.00 | 604.00 | 595.33 | 658.00 |
| ZBTB17 | 841.00 | 896.00 | 884.00 | 873.67 | 959.00 |
| ZBTB18 | 1075.00 | 1080.00 | 1224.00 | 1126.33 | 1159.00 |
| ZBTB2 | 505.00 | 529.00 | 559.00 | 531.00 | 509.00 |
| ZBTB20 | 499.00 | 477.00 | 445.00 | 473.67 | 323.00 |
| ZBTB21 | 956.00 | 904.00 | 1101.00 | 987.00 | 896.00 |
| ZBTB22 | 13.00 | 5.00 | 3.00 | 7.00 | 10.00 |
| ZBTB24 | 508.00 | 433.00 | 499.00 | 480.00 | 545.00 |
| ZBTB25 | 223.00 | 235.00 | 249.00 | 235.67 | 256.00 |
| ZBTB26 | 286.00 | 293.00 | 349.00 | 309.33 | 236.00 |
| ZBTB3 | 106.00 | 97.00 | 91.00 | 98.00 | 74.00 |
| ZBTB33 | 518.00 | 523.00 | 617.00 | 552.67 | 559.00 |
| ZBTB34 | 348.00 | 333.00 | 420.00 | 367.00 | 339.00 |
| ZBTB37 | 50.00 | 32.00 | 51.00 | 44.33 | 43.00 |
| ZBTB38 | 4342.00 | 3663.00 | 4484.00 | 4163.00 | 4016.00 |
| ZBTB39 | 521.00 | 491.00 | 564.00 | 525.33 | 493.00 |
| ZBTB4 | 2005.00 | 1896.00 | 2264.00 | 2055.00 | 1754.00 |
| ZBTB40 | 3427.00 | 3335.00 | 3621.00 | 3461.00 | 2907.00 |
| ZBTB41 | 374.00 | 298.00 | 376.00 | 349.33 | 308.00 |
| ZBTB42 | 25.00 | 25.00 | 31.00 | 27.00 | 17.00 |
| ZBTB43 | 497.00 | 549.00 | 608.00 | 551.33 | 601.00 |
| ZBTB44 | 549.00 | 616.00 | 718.00 | 627.67 | 636.00 |
| ZBTB45 | 210.00 | 226.00 | 258.00 | 231.33 | 219.00 |
| ZBTB46 | 14.00 | 6.00 | 13.00 | 11.00 | 5.00 |
| ZBTB47 | 584.00 | 574.00 | 629.00 | 595.67 | 500.00 |
| ZBTB48 | 602.00 | 660.00 | 710.00 | 657.33 | 692.00 |
| ZBTB49 | 111.00 | 101.00 | 115.00 | 109.00 | 138.00 |
| ZBTB5 | 478.00 | 514.00 | 593.00 | 528.33 | 458.00 |
| ZBTB6 | 296.00 | 257.00 | 323.00 | 292.00 | 285.00 |
| ZBTB7A | 960.00 | 972.00 | 1101.00 | 1011.00 | 969.00 |
| ZBTB7B | 681.00 | 602.00 | 630.00 | 637.67 | 640.00 |
| ZBTB8A | 109.00 | 108.00 | 121.00 | 112.67 | 109.00 |
| ZBTB8OS | 318.00 | 335.00 | 311.00 | 321.33 | 441.00 |
| ZBTB9 | 25.00 | 33.00 | 23.00 | 27.00 | 17.00 |
| ZC2HC1A | 339.00 | 244.00 | 359.00 | 314.00 | 299.00 |
| ZC2HC1C | 15.00 | 15.00 | 26.00 | 18.67 | 20.00 |

| | | | | | |
|---|---|---|---|---|---|
| ZC3H10 | 143.00 | 147.00 | 161.00 | 150.33 | 143.00 |
| ZC3H11A | 4290.00 | 4194.00 | 4916.00 | 4466.67 | 4904.00 |
| ZC3H12A | 241.00 | 217.00 | 203.00 | 220.33 | 199.00 |
| ZC3H12C | 1062.00 | 976.00 | 1214.00 | 1084.00 | 866.00 |
| ZC3H12D | 5.00 | 1.00 | 4.00 | 3.33 | 5.00 |
| ZC3H13 | 1787.00 | 1726.00 | 1898.00 | 1803.67 | 1509.00 |
| ZC3H14 | 2023.00 | 2021.00 | 2136.00 | 2060.00 | 2268.00 |
| ZC3H15 | 2640.00 | 2464.00 | 2830.00 | 2644.67 | 3481.00 |
| ZC3H18 | 1987.00 | 1920.00 | 1963.00 | 1956.67 | 1826.00 |
| ZC3H3 | 430.00 | 383.00 | 385.00 | 399.33 | 374.00 |
| ZC3H4 | 1220.00 | 1129.00 | 1082.00 | 1143.67 | 1130.00 |
| ZC3H6 | 382.00 | 351.00 | 402.00 | 378.33 | 307.00 |
| ZC3H7A | 1545.00 | 1420.00 | 1581.00 | 1515.33 | 1745.00 |
| ZC3H7B | 3558.00 | 3485.00 | 3665.00 | 3569.33 | 4116.00 |
| ZC3H8 | 499.00 | 498.00 | 549.00 | 515.33 | 625.00 |
| ZC3HAV1 | 1133.00 | 1140.00 | 1172.00 | 1148.33 | 1123.00 |
| ZC3HAV1L | 287.00 | 264.00 | 266.00 | 272.33 | 241.00 |
| ZC3HC1 | 764.00 | 805.00 | 798.00 | 789.00 | 1007.00 |
| ZCCHC10 | 153.00 | 139.00 | 173.00 | 155.00 | 188.00 |
| ZCCHC11 | 1674.00 | 1619.00 | 1748.00 | 1680.33 | 1536.00 |
| ZCCHC14 | 1558.00 | 1489.00 | 1728.00 | 1591.67 | 1493.00 |
| ZCCHC17 | 649.00 | 599.00 | 638.00 | 628.67 | 721.00 |
| ZCCHC18 | 226.00 | 232.00 | 213.00 | 223.67 | 272.00 |
| ZCCHC2 | 323.00 | 314.00 | 385.00 | 340.67 | 350.00 |
| ZCCHC24 | 142.00 | 171.00 | 191.00 | 168.00 | 90.00 |
| ZCCHC3 | 836.00 | 791.00 | 886.00 | 837.67 | 1100.00 |
| ZCCHC4 | 301.00 | 307.00 | 325.00 | 311.00 | 447.00 |
| ZCCHC6 | 587.00 | 566.00 | 668.00 | 607.00 | 539.00 |
| ZCCHC7 | 890.00 | 916.00 | 1078.00 | 961.33 | 1137.00 |
| ZCCHC8 | 1368.00 | 1269.00 | 1323.00 | 1320.00 | 1519.00 |
| ZCCHC9 | 1092.00 | 1074.00 | 1307.00 | 1157.67 | 1602.00 |
| ZCRB1 | 788.00 | 727.00 | 810.00 | 775.00 | 920.00 |
| ZCWPW1 | 224.00 | 209.00 | 244.00 | 225.67 | 221.00 |
| ZCWPW2 | 18.00 | 20.00 | 15.00 | 17.67 | 21.00 |
| ZDBF2 | 992.00 | 916.00 | 998.00 | 968.67 | 745.00 |
| ZDHHC1 | 261.00 | 302.00 | 283.00 | 282.00 | 295.00 |
| ZDHHC11 | 71.00 | 76.00 | 95.00 | 80.67 | 50.00 |
| ZDHHC12 | 332.00 | 348.00 | 273.00 | 317.67 | 447.00 |
| ZDHHC13 | 419.00 | 440.00 | 444.00 | 434.33 | 515.00 |
| ZDHHC14 | 539.00 | 471.00 | 533.00 | 514.33 | 482.00 |
| ZDHHC16 | 1033.00 | 1016.00 | 995.00 | 1014.67 | 1083.00 |
| ZDHHC17 | 1090.00 | 1013.00 | 1041.00 | 1048.00 | 901.00 |
| ZDHHC18 | 748.00 | 784.00 | 771.00 | 767.67 | 818.00 |
| ZDHHC2 | 386.00 | 414.00 | 518.00 | 439.33 | 414.00 |
| ZDHHC20 | 879.00 | 814.00 | 943.00 | 878.67 | 901.00 |
| ZDHHC21 | 1194.00 | 1133.00 | 1299.00 | 1208.67 | 1343.00 |
| ZDHHC23 | 408.00 | 420.00 | 547.00 | 458.33 | 613.00 |
| ZDHHC24 | 724.00 | 710.00 | 666.00 | 700.00 | 846.00 |

| | | | | | |
|---|---|---|---|---|---|
| ZDHHC3 | 1184.00 | 1154.00 | 1180.00 | 1172.67 | 1046.00 |
| ZDHHC4 | 923.00 | 955.00 | 1001.00 | 959.67 | 1127.00 |
| ZDHHC5 | 3917.00 | 3729.00 | 3756.00 | 3800.67 | 3490.00 |
| ZDHHC6 | 2070.00 | 2125.00 | 2148.00 | 2114.33 | 2303.00 |
| ZDHHC7 | 3280.00 | 3141.00 | 3292.00 | 3237.67 | 3188.00 |
| ZDHHC8 | 2118.00 | 2036.00 | 2121.00 | 2091.67 | 2117.00 |
| ZDHHC9 | 1356.00 | 1342.00 | 1400.00 | 1366.00 | 1421.00 |
| ZEB1 | 2302.00 | 2120.00 | 2586.00 | 2336.00 | 2462.00 |
| ZEB1-AS1 | 347.00 | 322.00 | 354.00 | 341.00 | 369.00 |
| ZEB2 | 1768.00 | 1573.00 | 1843.00 | 1728.00 | 1678.00 |
| ZER1 | 1073.00 | 1047.00 | 1051.00 | 1057.00 | 867.00 |
| ZFAND1 | 1447.00 | 1391.00 | 1599.00 | 1479.00 | 1929.00 |
| ZFAND2A | 385.00 | 347.00 | 343.00 | 358.33 | 449.00 |
| ZFAND2B | 428.00 | 440.00 | 402.00 | 423.33 | 467.00 |
| ZFAND3 | 2182.00 | 2129.00 | 2232.00 | 2181.00 | 2366.00 |
| ZFAND4 | 129.00 | 116.00 | 155.00 | 133.33 | 186.00 |
| ZFAND5 | 1962.00 | 1923.00 | 2185.00 | 2023.33 | 2028.00 |
| ZFAND6 | 862.00 | 751.00 | 925.00 | 846.00 | 1055.00 |
| ZFAS1 | 4093.00 | 3995.00 | 4487.00 | 4191.67 | 5537.00 |
| ZFAT | 315.00 | 264.00 | 381.00 | 320.00 | 301.00 |
| ZFC3H1 | 3641.00 | 3492.00 | 3875.00 | 3669.33 | 3195.00 |
| ZFHX2 | 32.00 | 26.00 | 37.00 | 31.67 | 28.00 |
| ZFHX3 | 193.00 | 221.00 | 221.00 | 211.67 | 137.00 |
| ZFHX4 | 1349.00 | 1105.00 | 1410.00 | 1288.00 | 1008.00 |
| ZFHX4-AS1 | 19.00 | 10.00 | 31.00 | 20.00 | 16.00 |
| ZFP1 | 259.00 | 268.00 | 262.00 | 263.00 | 290.00 |
| ZFP14 | 16.00 | 18.00 | 22.00 | 18.67 | 21.00 |
| ZFP2 | 4.00 | 4.00 | 5.00 | 4.33 | 5.00 |
| ZFP28 | 52.00 | 36.00 | 57.00 | 48.33 | 38.00 |
| ZFP30 | 108.00 | 83.00 | 88.00 | 93.00 | 102.00 |
| ZFP36 | 135.00 | 118.00 | 141.00 | 131.33 | 136.00 |
| ZFP36L1 | 3778.00 | 3575.00 | 4071.00 | 3808.00 | 2637.00 |
| ZFP36L2 | 677.00 | 583.00 | 676.00 | 645.33 | 801.00 |
| ZFP37 | 137.00 | 91.00 | 144.00 | 124.00 | 109.00 |
| ZFP41 | 963.00 | 784.00 | 925.00 | 890.67 | 643.00 |
| ZFP62 | 1416.00 | 1407.00 | 1633.00 | 1485.33 | 1567.00 |
| ZFP64 | 343.00 | 339.00 | 360.00 | 347.33 | 412.00 |
| ZFP69 | 136.00 | 136.00 | 142.00 | 138.00 | 172.00 |
| ZFP69B | 211.00 | 199.00 | 232.00 | 214.00 | 230.00 |
| ZFP82 | 25.00 | 19.00 | 19.00 | 21.00 | 24.00 |
| ZFP90 | 549.00 | 530.00 | 554.00 | 544.33 | 539.00 |
| ZFP91 | 1613.00 | 1444.00 | 1867.00 | 1641.33 | 1769.00 |
| ZFPL1 | 1173.00 | 1196.00 | 1261.00 | 1210.00 | 1417.00 |
| ZFPM1 | 164.00 | 156.00 | 169.00 | 163.00 | 203.00 |
| ZFPM2 | 113.00 | 106.00 | 107.00 | 108.67 | 221.00 |
| ZFPM2-AS1 | 294.00 | 297.00 | 273.00 | 288.00 | 197.00 |
| ZFR | 2862.00 | 2558.00 | 2982.00 | 2800.67 | 2873.00 |
| ZFX | 1065.00 | 1002.00 | 1184.00 | 1083.67 | 1146.00 |

| | | | | | |
|---|---|---|---|---|---|
| ZFYVE1 | 442.00 | 420.00 | 450.00 | 437.33 | 348.00 |
| ZFYVE16 | 873.00 | 837.00 | 935.00 | 881.67 | 814.00 |
| ZFYVE19 | 531.00 | 573.00 | 542.00 | 548.67 | 619.00 |
| ZFYVE21 | 527.00 | 560.00 | 567.00 | 551.33 | 522.00 |
| ZFYVE26 | 1187.00 | 1173.00 | 1255.00 | 1205.00 | 1025.00 |
| ZFYVE27 | 1945.00 | 1955.00 | 1969.00 | 1956.33 | 1844.00 |
| ZFYVE28 | 78.00 | 79.00 | 67.00 | 74.67 | 46.00 |
| ZFYVE9 | 1111.00 | 1072.00 | 1149.00 | 1110.67 | 1033.00 |
| ZGLP1 | 203.00 | 227.00 | 211.00 | 213.67 | 187.00 |
| ZGPAT | 1326.00 | 1486.00 | 1538.00 | 1450.00 | 1507.00 |
| ZGRF1 | 390.00 | 352.00 | 358.00 | 366.67 | 334.00 |
| ZHX1 | 706.00 | 631.00 | 766.00 | 701.00 | 616.00 |
| ZHX1-C8orf76 | 30.00 | 30.00 | 37.00 | 32.33 | 26.00 |
| ZHX2 | 203.00 | 220.00 | 227.00 | 216.67 | 211.00 |
| ZHX3 | 2330.00 | 2273.00 | 2391.00 | 2331.33 | 2621.00 |
| ZIC1 | 201.00 | 235.00 | 291.00 | 242.33 | 274.00 |
| ZIC2 | 204.00 | 173.00 | 200.00 | 192.33 | 226.00 |
| ZIC4 | 7.00 | 11.00 | 9.00 | 9.00 | 10.00 |
| ZIC5 | 127.00 | 72.00 | 131.00 | 110.00 | 121.00 |
| ZIK1 | 15.00 | 19.00 | 17.00 | 17.00 | 14.00 |
| ZKSCAN1 | 3735.00 | 3506.00 | 4085.00 | 3775.33 | 3396.00 |
| ZKSCAN2 | 350.00 | 287.00 | 356.00 | 331.00 | 303.00 |
| ZKSCAN3 | 142.00 | 151.00 | 166.00 | 153.00 | 169.00 |
| ZKSCAN4 | 183.00 | 212.00 | 210.00 | 201.67 | 208.00 |
| ZKSCAN5 | 565.00 | 542.00 | 579.00 | 562.00 | 588.00 |
| ZKSCAN8 | 2328.00 | 2221.00 | 2603.00 | 2384.00 | 1597.00 |
| ZMAT1 | 366.00 | 383.00 | 392.00 | 380.33 | 444.00 |
| ZMAT2 | 838.00 | 777.00 | 840.00 | 818.33 | 980.00 |
| ZMAT3 | 411.00 | 386.00 | 476.00 | 424.33 | 443.00 |
| ZMAT5 | 320.00 | 299.00 | 348.00 | 322.33 | 349.00 |
| ZMIZ1 | 1663.00 | 1611.00 | 1716.00 | 1663.33 | 1166.00 |
| ZMIZ1-AS1 | 33.00 | 36.00 | 34.00 | 34.33 | 29.00 |
| ZMIZ2 | 3146.00 | 3032.00 | 3210.00 | 3129.33 | 3011.00 |
| ZMPSTE24 | 2190.00 | 1974.00 | 2173.00 | 2112.33 | 2144.00 |
| ZMYM1 | 930.00 | 896.00 | 1081.00 | 969.00 | 1164.00 |
| ZMYM2 | 1913.00 | 1921.00 | 2209.00 | 2014.33 | 2073.00 |
| ZMYM3 | 1388.00 | 1293.00 | 1411.00 | 1364.00 | 1171.00 |
| ZMYM4 | 2019.00 | 1787.00 | 2067.00 | 1957.67 | 1988.00 |
| ZMYM5 | 397.00 | 371.00 | 440.00 | 402.67 | 350.00 |
| ZMYM6 | 812.00 | 728.00 | 779.00 | 773.00 | 743.00 |
| ZMYM6NB | 786.00 | 804.00 | 809.00 | 799.67 | 961.00 |
| ZMYND10 | 9.00 | 12.00 | 12.00 | 11.00 | 10.00 |
| ZMYND11 | 2164.00 | 1945.00 | 2386.00 | 2165.00 | 2322.00 |
| ZMYND12 | 3.00 | 11.00 | 15.00 | 9.67 | 7.00 |
| ZMYND19 | 628.00 | 685.00 | 645.00 | 652.67 | 773.00 |
| ZMYND8 | 4413.00 | 4286.00 | 4573.00 | 4424.00 | 4800.00 |
| ZNF10 | 352.00 | 377.00 | 389.00 | 372.67 | 372.00 |
| ZNF101 | 186.00 | 156.00 | 194.00 | 178.67 | 226.00 |

| | | | | | |
|---|---|---|---|---|---|
| ZNF106 | 2315.00 | 2064.00 | 2536.00 | 2305.00 | 2149.00 |
| ZNF107 | 630.00 | 554.00 | 657.00 | 613.67 | 707.00 |
| ZNF112 | 298.00 | 275.00 | 285.00 | 286.00 | 270.00 |
| ZNF117 | 43.00 | 41.00 | 58.00 | 47.33 | 63.00 |
| ZNF12 | 1522.00 | 1501.00 | 1722.00 | 1581.67 | 1511.00 |
| ZNF121 | 325.00 | 258.00 | 313.00 | 298.67 | 208.00 |
| ZNF124 | 192.00 | 175.00 | 200.00 | 189.00 | 253.00 |
| ZNF131 | 1295.00 | 1255.00 | 1433.00 | 1327.67 | 1497.00 |
| ZNF132 | 31.00 | 41.00 | 25.00 | 32.33 | 20.00 |
| ZNF133 | 891.00 | 929.00 | 986.00 | 935.33 | 923.00 |
| ZNF134 | 274.00 | 211.00 | 269.00 | 251.33 | 249.00 |
| ZNF136 | 74.00 | 86.00 | 103.00 | 87.67 | 123.00 |
| ZNF137P | 97.00 | 82.00 | 108.00 | 95.67 | 110.00 |
| ZNF138 | 138.00 | 138.00 | 143.00 | 139.67 | 192.00 |
| ZNF14 | 88.00 | 86.00 | 80.00 | 84.67 | 91.00 |
| ZNF140 | 144.00 | 146.00 | 142.00 | 144.00 | 171.00 |
| ZNF141 | 20.00 | 25.00 | 17.00 | 20.67 | 13.00 |
| ZNF142 | 807.00 | 753.00 | 814.00 | 791.33 | 745.00 |
| ZNF143 | 353.00 | 327.00 | 337.00 | 339.00 | 363.00 |
| ZNF146 | 2778.00 | 2799.00 | 3077.00 | 2884.67 | 3327.00 |
| ZNF148 | 994.00 | 849.00 | 1104.00 | 982.33 | 1181.00 |
| ZNF155 | 160.00 | 175.00 | 206.00 | 180.33 | 188.00 |
| ZNF157 | 6.00 | 5.00 | 2.00 | 4.33 | 2.00 |
| ZNF16 | 252.00 | 291.00 | 273.00 | 272.00 | 305.00 |
| ZNF160 | 310.00 | 333.00 | 378.00 | 340.33 | 383.00 |
| ZNF165 | 15.00 | 18.00 | 15.00 | 16.00 | 19.00 |
| ZNF169 | 167.00 | 173.00 | 182.00 | 174.00 | 117.00 |
| ZNF17 | 97.00 | 80.00 | 103.00 | 93.33 | 54.00 |
| ZNF174 | 292.00 | 292.00 | 306.00 | 296.67 | 333.00 |
| ZNF175 | 359.00 | 440.00 | 428.00 | 409.00 | 317.00 |
| ZNF18 | 354.00 | 460.00 | 415.00 | 409.67 | 550.00 |
| ZNF180 | 195.00 | 162.00 | 208.00 | 188.33 | 209.00 |
| ZNF181 | 257.00 | 224.00 | 242.00 | 241.00 | 176.00 |
| ZNF182 | 378.00 | 343.00 | 384.00 | 368.33 | 334.00 |
| ZNF184 | 186.00 | 158.00 | 204.00 | 182.67 | 219.00 |
| ZNF185 | 4049.00 | 3935.00 | 4151.00 | 4045.00 | 3135.00 |
| ZNF189 | 631.00 | 574.00 | 618.00 | 607.67 | 636.00 |
| ZNF19 | 96.00 | 88.00 | 83.00 | 89.00 | 99.00 |
| ZNF192P1 | 122.00 | 112.00 | 97.00 | 110.33 | 90.00 |
| ZNF195 | 902.00 | 860.00 | 863.00 | 875.00 | 1050.00 |
| ZNF197 | 460.00 | 503.00 | 508.00 | 490.33 | 488.00 |
| ZNF2 | 132.00 | 110.00 | 127.00 | 123.00 | 124.00 |
| ZNF200 | 297.00 | 257.00 | 309.00 | 287.67 | 347.00 |
| ZNF202 | 1335.00 | 1296.00 | 1429.00 | 1353.33 | 1368.00 |
| ZNF204P | 11.00 | 15.00 | 13.00 | 13.00 | 15.00 |
| ZNF205 | 378.00 | 382.00 | 443.00 | 401.00 | 450.00 |
| ZNF205-AS1 | 8.00 | 5.00 | 9.00 | 7.33 | 5.00 |
| ZNF207 | 3449.00 | 3263.00 | 3792.00 | 3501.33 | 3521.00 |

| | | | | | |
|---|---|---|---|---|---|
| ZNF211 | 278.00 | 288.00 | 291.00 | 285.67 | 274.00 |
| ZNF212 | 690.00 | 726.00 | 752.00 | 722.67 | 823.00 |
| ZNF213 | 393.00 | 343.00 | 322.00 | 352.67 | 338.00 |
| ZNF213-AS1 | 231.00 | 222.00 | 221.00 | 224.67 | 210.00 |
| ZNF214 | 34.00 | 31.00 | 36.00 | 33.67 | 38.00 |
| ZNF215 | 407.00 | 381.00 | 360.00 | 382.67 | 405.00 |
| ZNF217 | 1448.00 | 1295.00 | 1555.00 | 1432.67 | 1412.00 |
| ZNF219 | 76.00 | 75.00 | 79.00 | 76.67 | 124.00 |
| ZNF22 | 1318.00 | 1172.00 | 1349.00 | 1279.67 | 1227.00 |
| ZNF221 | 32.00 | 23.00 | 24.00 | 26.33 | 14.00 |
| ZNF222 | 69.00 | 78.00 | 82.00 | 76.33 | 116.00 |
| ZNF223 | 37.00 | 48.00 | 43.00 | 42.67 | 48.00 |
| ZNF224 | 474.00 | 502.00 | 513.00 | 496.33 | 462.00 |
| ZNF225 | 125.00 | 131.00 | 147.00 | 134.33 | 136.00 |
| ZNF226 | 422.00 | 444.00 | 439.00 | 435.00 | 467.00 |
| ZNF227 | 252.00 | 240.00 | 298.00 | 263.33 | 259.00 |
| ZNF229 | 6.00 | 3.00 | 5.00 | 4.67 | 0.00 |
| ZNF23 | 895.00 | 880.00 | 974.00 | 916.33 | 1095.00 |
| ZNF230 | 103.00 | 101.00 | 124.00 | 109.33 | 111.00 |
| ZNF232 | 314.00 | 345.00 | 271.00 | 310.00 | 358.00 |
| ZNF234 | 394.00 | 354.00 | 416.00 | 388.00 | 358.00 |
| ZNF235 | 59.00 | 51.00 | 71.00 | 60.33 | 88.00 |
| ZNF236 | 711.00 | 662.00 | 729.00 | 700.67 | 566.00 |
| ZNF239 | 703.00 | 730.00 | 725.00 | 719.33 | 1070.00 |
| ZNF24 | 1685.00 | 1533.00 | 1813.00 | 1677.00 | 1735.00 |
| ZNF248 | 1260.00 | 1212.00 | 1336.00 | 1269.33 | 1209.00 |
| ZNF25 | 355.00 | 355.00 | 456.00 | 388.67 | 560.00 |
| ZNF250 | 285.00 | 232.00 | 234.00 | 250.33 | 217.00 |
| ZNF252P | 696.00 | 639.00 | 708.00 | 681.00 | 773.00 |
| ZNF252P-AS1 | 26.00 | 30.00 | 22.00 | 26.00 | 27.00 |
| ZNF253 | 19.00 | 26.00 | 33.00 | 26.00 | 28.00 |
| ZNF254 | 96.00 | 94.00 | 132.00 | 107.33 | 105.00 |
| ZNF256 | 23.00 | 17.00 | 15.00 | 18.33 | 16.00 |
| ZNF26 | 745.00 | 754.00 | 768.00 | 755.67 | 957.00 |
| ZNF260 | 276.00 | 267.00 | 300.00 | 281.00 | 315.00 |
| ZNF263 | 1089.00 | 1091.00 | 1143.00 | 1107.67 | 1191.00 |
| ZNF264 | 335.00 | 338.00 | 359.00 | 344.00 | 344.00 |
| ZNF266 | 1071.00 | 1217.00 | 1255.00 | 1181.00 | 1226.00 |
| ZNF267 | 401.00 | 420.00 | 518.00 | 446.33 | 458.00 |
| ZNF268 | 331.00 | 297.00 | 342.00 | 323.33 | 355.00 |
| ZNF271P | 491.00 | 516.00 | 571.00 | 526.00 | 705.00 |
| ZNF273 | 63.00 | 50.00 | 64.00 | 59.00 | 62.00 |
| ZNF274 | 559.00 | 585.00 | 566.00 | 570.00 | 728.00 |
| ZNF275 | 571.00 | 549.00 | 600.00 | 573.33 | 508.00 |
| ZNF276 | 2467.00 | 2630.00 | 2911.00 | 2669.33 | 2162.00 |
| ZNF277 | 379.00 | 293.00 | 388.00 | 353.33 | 475.00 |
| ZNF28 | 137.00 | 114.00 | 165.00 | 138.67 | 182.00 |
| ZNF280C | 225.00 | 196.00 | 253.00 | 224.67 | 240.00 |

| | | | | | |
|---|---|---|---|---|---|
| ZNF280D | 792.00 | 740.00 | 892.00 | 808.00 | 971.00 |
| ZNF281 | 1521.00 | 1474.00 | 1613.00 | 1536.00 | 1511.00 |
| ZNF282 | 1161.00 | 1141.00 | 1294.00 | 1198.67 | 1216.00 |
| ZNF283 | 125.00 | 119.00 | 146.00 | 130.00 | 115.00 |
| ZNF284 | 108.00 | 117.00 | 125.00 | 116.67 | 119.00 |
| ZNF285 | 124.00 | 94.00 | 127.00 | 115.00 | 123.00 |
| ZNF286A | 545.00 | 524.00 | 520.00 | 529.67 | 548.00 |
| ZNF286B | 153.00 | 164.00 | 178.00 | 165.00 | 188.00 |
| ZNF287 | 102.00 | 100.00 | 128.00 | 110.00 | 135.00 |
| ZNF292 | 1007.00 | 909.00 | 1067.00 | 994.33 | 1044.00 |
| ZNF296 | 17.00 | 16.00 | 20.00 | 17.67 | 10.00 |
| ZNF3 | 1966.00 | 2050.00 | 2068.00 | 2028.00 | 2223.00 |
| ZNF30 | 162.00 | 146.00 | 175.00 | 161.00 | 134.00 |
| ZNF300 | 1002.00 | 996.00 | 1022.00 | 1006.67 | 1313.00 |
| ZNF302 | 291.00 | 323.00 | 338.00 | 317.33 | 295.00 |
| ZNF304 | 241.00 | 275.00 | 255.00 | 257.00 | 258.00 |
| ZNF311 | 13.00 | 17.00 | 17.00 | 15.67 | 17.00 |
| ZNF316 | 1525.00 | 1527.00 | 1624.00 | 1558.67 | 1490.00 |
| ZNF317 | 1026.00 | 1000.00 | 1091.00 | 1039.00 | 1020.00 |
| ZNF318 | 1132.00 | 1004.00 | 1196.00 | 1110.67 | 988.00 |
| ZNF319 | 472.00 | 496.00 | 474.00 | 480.67 | 497.00 |
| ZNF32 | 326.00 | 398.00 | 377.00 | 367.00 | 489.00 |
| ZNF320 | 19.00 | 20.00 | 30.00 | 23.00 | 15.00 |
| ZNF322 | 242.00 | 223.00 | 285.00 | 250.00 | 264.00 |
| ZNF324 | 216.00 | 246.00 | 244.00 | 235.33 | 179.00 |
| ZNF324B | 104.00 | 93.00 | 95.00 | 97.33 | 93.00 |
| ZNF326 | 816.00 | 830.00 | 801.00 | 815.67 | 1052.00 |
| ZNF329 | 190.00 | 177.00 | 187.00 | 184.67 | 225.00 |
| ZNF32-AS2 | 4.00 | 4.00 | 5.00 | 4.33 | 3.00 |
| ZNF330 | 656.00 | 699.00 | 673.00 | 676.00 | 937.00 |
| ZNF331 | 246.00 | 273.00 | 288.00 | 269.00 | 300.00 |
| ZNF333 | 598.00 | 603.00 | 648.00 | 616.33 | 630.00 |
| ZNF335 | 1220.00 | 1254.00 | 1371.00 | 1281.67 | 1148.00 |
| ZNF337 | 2624.00 | 2698.00 | 2966.00 | 2762.67 | 2267.00 |
| ZNF337-AS1 | 317.00 | 306.00 | 323.00 | 315.33 | 338.00 |
| ZNF33A | 1113.00 | 1043.00 | 1260.00 | 1138.67 | 971.00 |
| ZNF33B | 757.00 | 768.00 | 824.00 | 783.00 | 771.00 |
| ZNF34 | 217.00 | 166.00 | 185.00 | 189.33 | 194.00 |
| ZNF341 | 178.00 | 148.00 | 158.00 | 161.33 | 196.00 |
| ZNF341-AS1 | 4.00 | 3.00 | 4.00 | 3.67 | 5.00 |
| ZNF343 | 692.00 | 681.00 | 741.00 | 704.67 | 688.00 |
| ZNF345 | 3.00 | 0.00 | 0.00 | 1.00 | 0.00 |
| ZNF346 | 472.00 | 414.00 | 387.00 | 424.33 | 423.00 |
| ZNF35 | 69.00 | 71.00 | 66.00 | 68.67 | 80.00 |
| ZNF350 | 71.00 | 66.00 | 93.00 | 76.67 | 80.00 |
| ZNF354A | 589.00 | 594.00 | 657.00 | 613.33 | 678.00 |
| ZNF354B | 438.00 | 424.00 | 513.00 | 458.33 | 385.00 |
| ZNF354C | 4.00 | 1.00 | 6.00 | 3.67 | 2.00 |

| | | | | | |
|---|---|---|---|---|---|
| ZNF358 | 705.00 | 606.00 | 791.00 | 700.67 | 800.00 |
| ZNF362 | 169.00 | 158.00 | 182.00 | 169.67 | 203.00 |
| ZNF365 | 99.00 | 95.00 | 122.00 | 105.33 | 115.00 |
| ZNF367 | 102.00 | 119.00 | 112.00 | 111.00 | 167.00 |
| ZNF37A | 1637.00 | 1445.00 | 1713.00 | 1598.33 | 1360.00 |
| ZNF37BP | 1516.00 | 1647.00 | 1689.00 | 1617.33 | 1629.00 |
| ZNF383 | 51.00 | 34.00 | 32.00 | 39.00 | 41.00 |
| ZNF384 | 1855.00 | 1805.00 | 1956.00 | 1872.00 | 1883.00 |
| ZNF385A | 67.00 | 57.00 | 61.00 | 61.67 | 67.00 |
| ZNF385D | 190.00 | 170.00 | 226.00 | 195.33 | 128.00 |
| ZNF391 | 74.00 | 61.00 | 51.00 | 62.00 | 52.00 |
| ZNF394 | 325.00 | 325.00 | 312.00 | 320.67 | 415.00 |
| ZNF395 | 813.00 | 756.00 | 803.00 | 790.67 | 727.00 |
| ZNF396 | 7.00 | 15.00 | 16.00 | 12.67 | 12.00 |
| ZNF397 | 199.00 | 189.00 | 211.00 | 199.67 | 270.00 |
| ZNF398 | 392.00 | 359.00 | 451.00 | 400.67 | 334.00 |
| ZNF404 | 76.00 | 62.00 | 82.00 | 73.33 | 68.00 |
| ZNF407 | 453.00 | 417.00 | 500.00 | 456.67 | 420.00 |
| ZNF408 | 328.00 | 322.00 | 364.00 | 338.00 | 370.00 |
| ZNF41 | 381.00 | 381.00 | 440.00 | 400.67 | 326.00 |
| ZNF410 | 858.00 | 800.00 | 858.00 | 838.67 | 1050.00 |
| ZNF414 | 164.00 | 125.00 | 174.00 | 154.33 | 190.00 |
| ZNF416 | 138.00 | 108.00 | 119.00 | 121.67 | 120.00 |
| ZNF417 | 191.00 | 157.00 | 180.00 | 176.00 | 121.00 |
| ZNF418 | 34.00 | 32.00 | 44.00 | 36.67 | 33.00 |
| ZNF419 | 384.00 | 410.00 | 405.00 | 399.67 | 432.00 |
| ZNF420 | 45.00 | 38.00 | 56.00 | 46.33 | 46.00 |
| ZNF425 | 82.00 | 67.00 | 71.00 | 73.33 | 89.00 |
| ZNF426 | 182.00 | 171.00 | 197.00 | 183.33 | 164.00 |
| ZNF428 | 470.00 | 519.00 | 462.00 | 483.67 | 640.00 |
| ZNF429 | 5.00 | 0.00 | 5.00 | 3.33 | 1.00 |
| ZNF43 | 4.00 | 1.00 | 3.00 | 2.67 | 1.00 |
| ZNF430 | 130.00 | 136.00 | 173.00 | 146.33 | 132.00 |
| ZNF431 | 285.00 | 241.00 | 278.00 | 268.00 | 259.00 |
| ZNF432 | 181.00 | 162.00 | 152.00 | 165.00 | 154.00 |
| ZNF433 | 34.00 | 32.00 | 36.00 | 34.00 | 29.00 |
| ZNF436 | 260.00 | 264.00 | 308.00 | 277.33 | 217.00 |
| ZNF436-AS1 | 189.00 | 186.00 | 208.00 | 194.33 | 145.00 |
| ZNF438 | 398.00 | 374.00 | 418.00 | 396.67 | 541.00 |
| ZNF439 | 7.00 | 4.00 | 2.00 | 4.33 | 12.00 |
| ZNF44 | 153.00 | 185.00 | 176.00 | 171.33 | 194.00 |
| ZNF440 | 361.00 | 328.00 | 383.00 | 357.33 | 320.00 |
| ZNF441 | 425.00 | 380.00 | 386.00 | 397.00 | 284.00 |
| ZNF442 | 6.00 | 4.00 | 3.00 | 4.33 | 4.00 |
| ZNF443 | 12.00 | 14.00 | 14.00 | 13.33 | 29.00 |
| ZNF444 | 556.00 | 602.00 | 652.00 | 603.33 | 675.00 |
| ZNF445 | 1064.00 | 961.00 | 1095.00 | 1040.00 | 843.00 |
| ZNF446 | 383.00 | 366.00 | 351.00 | 366.67 | 376.00 |

| | | | | | |
|---|---|---|---|---|---|
| ZNF449 | 248.00 | 267.00 | 275.00 | 263.33 | 272.00 |
| ZNF45 | 281.00 | 301.00 | 281.00 | 287.67 | 319.00 |
| ZNF451 | 1184.00 | 1170.00 | 1354.00 | 1236.00 | 1215.00 |
| ZNF460 | 22.00 | 20.00 | 23.00 | 21.67 | 16.00 |
| ZNF461 | 34.00 | 31.00 | 41.00 | 35.33 | 35.00 |
| ZNF462 | 1003.00 | 951.00 | 1149.00 | 1034.33 | 775.00 |
| ZNF468 | 54.00 | 40.00 | 45.00 | 46.33 | 41.00 |
| ZNF469 | 27.00 | 18.00 | 31.00 | 25.33 | 13.00 |
| ZNF470 | 102.00 | 85.00 | 119.00 | 102.00 | 113.00 |
| ZNF473 | 500.00 | 509.00 | 588.00 | 532.33 | 491.00 |
| ZNF474 | 7.00 | 16.00 | 11.00 | 11.33 | 6.00 |
| ZNF48 | 296.00 | 286.00 | 333.00 | 305.00 | 306.00 |
| ZNF480 | 337.00 | 334.00 | 346.00 | 339.00 | 334.00 |
| ZNF484 | 100.00 | 86.00 | 116.00 | 100.67 | 106.00 |
| ZNF485 | 453.00 | 452.00 | 488.00 | 464.33 | 521.00 |
| ZNF486 | 43.00 | 38.00 | 42.00 | 41.00 | 52.00 |
| ZNF487 | 25.00 | 15.00 | 25.00 | 21.67 | 35.00 |
| ZNF488 | 34.00 | 30.00 | 40.00 | 34.67 | 34.00 |
| ZNF490 | 105.00 | 97.00 | 91.00 | 97.67 | 120.00 |
| ZNF491 | 17.00 | 18.00 | 14.00 | 16.33 | 24.00 |
| ZNF493 | 105.00 | 113.00 | 103.00 | 107.00 | 86.00 |
| ZNF496 | 1020.00 | 953.00 | 1036.00 | 1003.00 | 855.00 |
| ZNF497 | 91.00 | 78.00 | 104.00 | 91.00 | 94.00 |
| ZNF500 | 335.00 | 356.00 | 313.00 | 334.67 | 305.00 |
| ZNF501 | 47.00 | 32.00 | 47.00 | 42.00 | 39.00 |
| ZNF502 | 51.00 | 39.00 | 50.00 | 46.67 | 76.00 |
| ZNF503 | 292.00 | 306.00 | 308.00 | 302.00 | 334.00 |
| ZNF503-AS2 | 39.00 | 28.00 | 44.00 | 37.00 | 28.00 |
| ZNF506 | 63.00 | 51.00 | 60.00 | 58.00 | 64.00 |
| ZNF507 | 1166.00 | 1020.00 | 1304.00 | 1163.33 | 984.00 |
| ZNF510 | 317.00 | 273.00 | 339.00 | 309.67 | 254.00 |
| ZNF511 | 414.00 | 471.00 | 469.00 | 451.33 | 631.00 |
| ZNF512 | 1849.00 | 1772.00 | 1956.00 | 1859.00 | 1987.00 |
| ZNF512B | 2466.00 | 2467.00 | 2563.00 | 2498.67 | 2229.00 |
| ZNF513 | 487.00 | 505.00 | 516.00 | 502.67 | 503.00 |
| ZNF514 | 1331.00 | 1334.00 | 1401.00 | 1355.33 | 1338.00 |
| ZNF516 | 1325.00 | 1333.00 | 1458.00 | 1372.00 | 1122.00 |
| ZNF517 | 69.00 | 77.00 | 73.00 | 73.00 | 84.00 |
| ZNF518A | 1314.00 | 1273.00 | 1493.00 | 1360.00 | 1202.00 |
| ZNF518B | 151.00 | 180.00 | 223.00 | 184.67 | 155.00 |
| ZNF519 | 306.00 | 322.00 | 361.00 | 329.67 | 402.00 |
| ZNF521 | 298.00 | 270.00 | 319.00 | 295.67 | 291.00 |
| ZNF524 | 133.00 | 165.00 | 135.00 | 144.33 | 168.00 |
| ZNF525 | 50.00 | 48.00 | 51.00 | 49.67 | 51.00 |
| ZNF526 | 282.00 | 228.00 | 296.00 | 268.67 | 245.00 |
| ZNF527 | 121.00 | 113.00 | 115.00 | 116.33 | 136.00 |
| ZNF528 | 19.00 | 39.00 | 35.00 | 31.00 | 39.00 |
| ZNF528-AS1 | 10.00 | 8.00 | 9.00 | 9.00 | 15.00 |

| | | | | | |
|---|---|---|---|---|---|
| ZNF529 | 279.00 | 268.00 | 260.00 | 269.00 | 276.00 |
| ZNF529-AS1 | 42.00 | 23.00 | 46.00 | 37.00 | 33.00 |
| ZNF530 | 122.00 | 120.00 | 90.00 | 110.67 | 144.00 |
| ZNF532 | 7.00 | 1.00 | 6.00 | 4.67 | 5.00 |
| ZNF540 | 6.00 | 5.00 | 8.00 | 6.33 | 8.00 |
| ZNF542P | 12.00 | 9.00 | 7.00 | 9.33 | 13.00 |
| ZNF543 | 76.00 | 75.00 | 78.00 | 76.33 | 66.00 |
| ZNF544 | 579.00 | 546.00 | 535.00 | 553.33 | 645.00 |
| ZNF546 | 70.00 | 59.00 | 70.00 | 66.33 | 58.00 |
| ZNF547 | 63.00 | 80.00 | 74.00 | 72.33 | 83.00 |
| ZNF548 | 455.00 | 452.00 | 494.00 | 467.00 | 448.00 |
| ZNF549 | 215.00 | 227.00 | 241.00 | 227.67 | 232.00 |
| ZNF550 | 633.00 | 664.00 | 740.00 | 679.00 | 622.00 |
| ZNF551 | 147.00 | 158.00 | 154.00 | 153.00 | 159.00 |
| ZNF552 | 101.00 | 94.00 | 94.00 | 96.33 | 82.00 |
| ZNF554 | 85.00 | 96.00 | 92.00 | 91.00 | 80.00 |
| ZNF555 | 122.00 | 156.00 | 168.00 | 148.67 | 159.00 |
| ZNF557 | 138.00 | 114.00 | 142.00 | 131.33 | 122.00 |
| ZNF559 | 407.00 | 423.00 | 414.00 | 414.67 | 528.00 |
| ZNF561 | 610.00 | 616.00 | 679.00 | 635.00 | 627.00 |
| ZNF561-AS1 | 39.00 | 45.00 | 36.00 | 40.00 | 40.00 |
| ZNF562 | 565.00 | 534.00 | 641.00 | 580.00 | 545.00 |
| ZNF563 | 6.00 | 8.00 | 6.00 | 6.67 | 8.00 |
| ZNF564 | 162.00 | 152.00 | 196.00 | 170.00 | 178.00 |
| ZNF565 | 71.00 | 87.00 | 97.00 | 85.00 | 87.00 |
| ZNF566 | 153.00 | 154.00 | 145.00 | 150.67 | 175.00 |
| ZNF567 | 101.00 | 104.00 | 115.00 | 106.67 | 110.00 |
| ZNF569 | 79.00 | 70.00 | 78.00 | 75.67 | 76.00 |
| ZNF57 | 107.00 | 116.00 | 111.00 | 111.33 | 108.00 |
| ZNF570 | 14.00 | 17.00 | 24.00 | 18.33 | 10.00 |
| ZNF571 | 29.00 | 26.00 | 32.00 | 29.00 | 48.00 |
| ZNF572 | 80.00 | 97.00 | 98.00 | 91.67 | 80.00 |
| ZNF573 | 14.00 | 20.00 | 23.00 | 19.00 | 9.00 |
| ZNF574 | 399.00 | 381.00 | 437.00 | 405.67 | 424.00 |
| ZNF575 | 16.00 | 14.00 | 20.00 | 16.67 | 25.00 |
| ZNF576 | 103.00 | 125.00 | 119.00 | 115.67 | 117.00 |
| ZNF577 | 323.00 | 301.00 | 362.00 | 328.67 | 310.00 |
| ZNF579 | 328.00 | 288.00 | 340.00 | 318.67 | 368.00 |
| ZNF580 | 589.00 | 546.00 | 613.00 | 582.67 | 633.00 |
| ZNF581 | 445.00 | 508.00 | 564.00 | 505.67 | 556.00 |
| ZNF583 | 80.00 | 42.00 | 37.00 | 53.00 | 59.00 |
| ZNF584 | 202.00 | 225.00 | 233.00 | 220.00 | 275.00 |
| ZNF585A | 106.00 | 87.00 | 94.00 | 95.67 | 104.00 |
| ZNF585B | 38.00 | 36.00 | 29.00 | 34.33 | 42.00 |
| ZNF586 | 112.00 | 125.00 | 137.00 | 124.67 | 122.00 |
| ZNF587 | 1045.00 | 1014.00 | 1156.00 | 1071.67 | 726.00 |
| ZNF587B | 282.00 | 215.00 | 274.00 | 257.00 | 352.00 |
| ZNF589 | 927.00 | 942.00 | 977.00 | 948.67 | 1003.00 |

| | | | | | |
|---|---|---|---|---|---|
| ZNF592 | 1022.00 | 941.00 | 1116.00 | 1026.33 | 863.00 |
| ZNF593 | 435.00 | 456.00 | 493.00 | 461.33 | 587.00 |
| ZNF594 | 431.00 | 427.00 | 428.00 | 428.67 | 363.00 |
| ZNF595 | 138.00 | 162.00 | 133.00 | 144.33 | 171.00 |
| ZNF596 | 83.00 | 78.00 | 101.00 | 87.33 | 93.00 |
| ZNF597 | 90.00 | 94.00 | 93.00 | 92.33 | 102.00 |
| ZNF598 | 3457.00 | 3491.00 | 3641.00 | 3529.67 | 2828.00 |
| ZNF599 | 65.00 | 61.00 | 78.00 | 68.00 | 65.00 |
| ZNF600 | 75.00 | 76.00 | 80.00 | 77.00 | 101.00 |
| ZNF605 | 434.00 | 410.00 | 493.00 | 445.67 | 418.00 |
| ZNF606 | 17.00 | 19.00 | 21.00 | 19.00 | 16.00 |
| ZNF607 | 78.00 | 65.00 | 85.00 | 76.00 | 67.00 |
| ZNF608 | 94.00 | 59.00 | 98.00 | 83.67 | 39.00 |
| ZNF609 | 2072.00 | 1845.00 | 1990.00 | 1969.00 | 1421.00 |
| ZNF610 | 8.00 | 7.00 | 9.00 | 8.00 | 9.00 |
| ZNF611 | 152.00 | 135.00 | 141.00 | 142.67 | 91.00 |
| ZNF613 | 60.00 | 60.00 | 54.00 | 58.00 | 67.00 |
| ZNF614 | 241.00 | 208.00 | 234.00 | 227.67 | 256.00 |
| ZNF615 | 137.00 | 138.00 | 134.00 | 136.33 | 143.00 |
| ZNF616 | 154.00 | 199.00 | 205.00 | 186.00 | 188.00 |
| ZNF618 | 432.00 | 440.00 | 466.00 | 446.00 | 353.00 |
| ZNF619 | 197.00 | 184.00 | 213.00 | 198.00 | 192.00 |
| ZNF620 | 154.00 | 127.00 | 124.00 | 135.00 | 118.00 |
| ZNF621 | 2379.00 | 2173.00 | 2502.00 | 2351.33 | 2073.00 |
| ZNF622 | 1002.00 | 1051.00 | 1052.00 | 1035.00 | 1137.00 |
| ZNF623 | 42.00 | 38.00 | 45.00 | 41.67 | 31.00 |
| ZNF624 | 129.00 | 117.00 | 128.00 | 124.67 | 133.00 |
| ZNF625 | 25.00 | 22.00 | 15.00 | 20.67 | 16.00 |
| ZNF625-ZNF20 | 11.00 | 8.00 | 7.00 | 8.67 | 8.00 |
| ZNF626 | 16.00 | 15.00 | 25.00 | 18.67 | 21.00 |
| ZNF627 | 352.00 | 333.00 | 385.00 | 356.67 | 365.00 |
| ZNF628 | 149.00 | 115.00 | 106.00 | 123.33 | 141.00 |
| ZNF629 | 821.00 | 833.00 | 806.00 | 820.00 | 877.00 |
| ZNF638 | 4019.00 | 3917.00 | 4444.00 | 4126.67 | 4068.00 |
| ZNF639 | 847.00 | 867.00 | 956.00 | 890.00 | 1123.00 |
| ZNF641 | 112.00 | 111.00 | 116.00 | 113.00 | 116.00 |
| ZNF644 | 1253.00 | 1073.00 | 1302.00 | 1209.33 | 1328.00 |
| ZNF646 | 1047.00 | 1119.00 | 1088.00 | 1084.67 | 879.00 |
| ZNF649 | 115.00 | 98.00 | 151.00 | 121.33 | 124.00 |
| ZNF652 | 631.00 | 659.00 | 695.00 | 661.67 | 576.00 |
| ZNF653 | 84.00 | 77.00 | 85.00 | 82.00 | 83.00 |
| ZNF654 | 222.00 | 193.00 | 274.00 | 229.67 | 202.00 |
| ZNF655 | 3259.00 | 3223.00 | 3498.00 | 3326.67 | 2978.00 |
| ZNF658 | 60.00 | 57.00 | 81.00 | 66.00 | 67.00 |
| ZNF658B | 29.00 | 20.00 | 21.00 | 23.33 | 23.00 |
| ZNF660 | 14.00 | 16.00 | 20.00 | 16.67 | 17.00 |
| ZNF664 | 3028.00 | 2926.00 | 3408.00 | 3120.67 | 3284.00 |
| ZNF668 | 330.00 | 352.00 | 383.00 | 355.00 | 323.00 |

| | | | | | |
|---|---|---|---|---|---|
| ZNF669 | 102.00 | 93.00 | 98.00 | 97.67 | 88.00 |
| ZNF670 | 134.00 | 140.00 | 156.00 | 143.33 | 156.00 |
| ZNF671 | 94.00 | 102.00 | 93.00 | 96.33 | 119.00 |
| ZNF672 | 815.00 | 832.00 | 839.00 | 828.67 | 729.00 |
| ZNF674 | 148.00 | 150.00 | 169.00 | 155.67 | 230.00 |
| ZNF674-AS1 | 377.00 | 392.00 | 367.00 | 378.67 | 398.00 |
| ZNF675 | 74.00 | 64.00 | 89.00 | 75.67 | 81.00 |
| ZNF678 | 297.00 | 295.00 | 341.00 | 311.00 | 332.00 |
| ZNF680 | 100.00 | 93.00 | 126.00 | 106.33 | 127.00 |
| ZNF681 | 31.00 | 31.00 | 26.00 | 29.33 | 20.00 |
| ZNF682 | 5.00 | 7.00 | 10.00 | 7.33 | 10.00 |
| ZNF684 | 71.00 | 77.00 | 88.00 | 78.67 | 105.00 |
| ZNF687 | 1061.00 | 865.00 | 901.00 | 942.33 | 927.00 |
| ZNF688 | 239.00 | 245.00 | 245.00 | 243.00 | 251.00 |
| ZNF689 | 289.00 | 231.00 | 249.00 | 256.33 | 268.00 |
| ZNF691 | 441.00 | 453.00 | 497.00 | 463.67 | 491.00 |
| ZNF692 | 1882.00 | 1997.00 | 2028.00 | 1969.00 | 2145.00 |
| ZNF695 | 54.00 | 45.00 | 49.00 | 49.33 | 63.00 |
| ZNF696 | 660.00 | 626.00 | 691.00 | 659.00 | 649.00 |
| ZNF697 | 1493.00 | 1284.00 | 1531.00 | 1436.00 | 1370.00 |
| ZNF699 | 54.00 | 49.00 | 59.00 | 54.00 | 35.00 |
| ZNF7 | 1143.00 | 1122.00 | 1190.00 | 1151.67 | 1172.00 |
| ZNF70 | 140.00 | 146.00 | 134.00 | 140.00 | 100.00 |
| ZNF700 | 675.00 | 740.00 | 765.00 | 726.67 | 874.00 |
| ZNF701 | 220.00 | 204.00 | 181.00 | 201.67 | 139.00 |
| ZNF703 | 952.00 | 904.00 | 1044.00 | 966.67 | 1096.00 |
| ZNF704 | 175.00 | 197.00 | 207.00 | 193.00 | 160.00 |
| ZNF705E | 12.00 | 9.00 | 8.00 | 9.67 | 8.00 |
| ZNF706 | 1114.00 | 1093.00 | 1218.00 | 1141.67 | 1456.00 |
| ZNF708 | 116.00 | 98.00 | 125.00 | 113.00 | 134.00 |
| ZNF709 | 11.00 | 8.00 | 10.00 | 9.67 | 12.00 |
| ZNF71 | 112.00 | 110.00 | 115.00 | 112.33 | 132.00 |
| ZNF710 | 914.00 | 938.00 | 1031.00 | 961.00 | 957.00 |
| ZNF711 | 32.00 | 31.00 | 30.00 | 31.00 | 33.00 |
| ZNF713 | 14.00 | 23.00 | 25.00 | 20.67 | 24.00 |
| ZNF714 | 222.00 | 190.00 | 185.00 | 199.00 | 210.00 |
| ZNF717 | 269.00 | 235.00 | 263.00 | 255.67 | 245.00 |
| ZNF718 | 52.00 | 30.00 | 37.00 | 39.67 | 36.00 |
| ZNF720 | 125.00 | 141.00 | 131.00 | 132.33 | 189.00 |
| ZNF721 | 570.00 | 546.00 | 636.00 | 584.00 | 526.00 |
| ZNF724P | 3.00 | 0.00 | 2.00 | 1.67 | 2.00 |
| ZNF726 | 4.00 | 1.00 | 5.00 | 3.33 | 6.00 |
| ZNF736 | 174.00 | 142.00 | 180.00 | 165.33 | 178.00 |
| ZNF737 | 5.00 | 3.00 | 3.00 | 3.67 | 16.00 |
| ZNF738 | 85.00 | 109.00 | 94.00 | 96.00 | 95.00 |
| ZNF74 | 467.00 | 495.00 | 463.00 | 475.00 | 541.00 |
| ZNF740 | 1205.00 | 1175.00 | 1207.00 | 1195.67 | 1252.00 |
| ZNF746 | 754.00 | 794.00 | 847.00 | 798.33 | 895.00 |

| | | | | | |
|---|---|---|---|---|---|
| ZNF747 | 231.00 | 207.00 | 225.00 | 221.00 | 196.00 |
| ZNF749 | 146.00 | 123.00 | 156.00 | 141.67 | 115.00 |
| ZNF75A | 203.00 | 197.00 | 186.00 | 195.33 | 185.00 |
| ZNF75D | 174.00 | 179.00 | 225.00 | 192.67 | 205.00 |
| ZNF76 | 961.00 | 1007.00 | 1069.00 | 1012.33 | 1040.00 |
| ZNF761 | 512.00 | 483.00 | 543.00 | 512.67 | 496.00 |
| ZNF763 | 14.00 | 13.00 | 14.00 | 13.67 | 6.00 |
| ZNF764 | 202.00 | 230.00 | 237.00 | 223.00 | 241.00 |
| ZNF765 | 95.00 | 74.00 | 98.00 | 89.00 | 68.00 |
| ZNF766 | 244.00 | 244.00 | 260.00 | 249.33 | 265.00 |
| ZNF767P | 1091.00 | 1050.00 | 1097.00 | 1079.33 | 998.00 |
| ZNF768 | 1688.00 | 1745.00 | 1847.00 | 1760.00 | 2412.00 |
| ZNF77 | 130.00 | 132.00 | 141.00 | 134.33 | 120.00 |
| ZNF770 | 1785.00 | 1693.00 | 2118.00 | 1865.33 | 1821.00 |
| ZNF771 | 111.00 | 126.00 | 139.00 | 125.33 | 146.00 |
| ZNF772 | 48.00 | 39.00 | 43.00 | 43.33 | 36.00 |
| ZNF773 | 149.00 | 147.00 | 147.00 | 147.67 | 156.00 |
| ZNF774 | 24.00 | 33.00 | 47.00 | 34.67 | 41.00 |
| ZNF775 | 193.00 | 168.00 | 187.00 | 182.67 | 211.00 |
| ZNF776 | 359.00 | 362.00 | 379.00 | 366.67 | 332.00 |
| ZNF777 | 659.00 | 615.00 | 661.00 | 645.00 | 623.00 |
| ZNF778 | 305.00 | 301.00 | 306.00 | 304.00 | 262.00 |
| ZNF780A | 321.00 | 296.00 | 347.00 | 321.33 | 284.00 |
| ZNF780B | 581.00 | 500.00 | 559.00 | 546.67 | 558.00 |
| ZNF781 | 5.00 | 3.00 | 10.00 | 6.00 | 7.00 |
| ZNF782 | 156.00 | 161.00 | 150.00 | 155.67 | 166.00 |
| ZNF783 | 2720.00 | 2742.00 | 3105.00 | 2855.67 | 2575.00 |
| ZNF784 | 48.00 | 53.00 | 41.00 | 47.33 | 46.00 |
| ZNF785 | 414.00 | 438.00 | 432.00 | 428.00 | 317.00 |
| ZNF786 | 405.00 | 444.00 | 439.00 | 429.33 | 389.00 |
| ZNF787 | 698.00 | 725.00 | 706.00 | 709.67 | 871.00 |
| ZNF788 | 5.00 | 13.00 | 8.00 | 8.67 | 15.00 |
| ZNF789 | 406.00 | 395.00 | 396.00 | 399.00 | 351.00 |
| ZNF79 | 141.00 | 143.00 | 120.00 | 134.67 | 171.00 |
| ZNF791 | 407.00 | 464.00 | 422.00 | 431.00 | 331.00 |
| ZNF792 | 157.00 | 160.00 | 160.00 | 159.00 | 87.00 |
| ZNF799 | 50.00 | 32.00 | 27.00 | 36.33 | 61.00 |
| ZNF8 | 156.00 | 161.00 | 168.00 | 161.67 | 165.00 |
| ZNF800 | 535.00 | 458.00 | 597.00 | 530.00 | 546.00 |
| ZNF804A | 487.00 | 427.00 | 577.00 | 497.00 | 506.00 |
| ZNF805 | 200.00 | 203.00 | 227.00 | 210.00 | 190.00 |
| ZNF808 | 107.00 | 100.00 | 90.00 | 99.00 | 68.00 |
| ZNF81 | 227.00 | 232.00 | 193.00 | 217.33 | 187.00 |
| ZNF812 | 6.00 | 5.00 | 1.00 | 4.00 | 2.00 |
| ZNF814 | 122.00 | 124.00 | 130.00 | 125.33 | 103.00 |
| ZNF815P | 31.00 | 28.00 | 20.00 | 26.33 | 36.00 |
| ZNF816 | 14.00 | 12.00 | 12.00 | 12.67 | 17.00 |
| ZNF821 | 76.00 | 73.00 | 61.00 | 70.00 | 73.00 |

| | | | | | |
|---|---|---|---|---|---|
| ZNF823 | 100.00 | 96.00 | 111.00 | 102.33 | 144.00 |
| ZNF827 | 925.00 | 814.00 | 995.00 | 911.33 | 840.00 |
| ZNF83 | 1829.00 | 1855.00 | 2046.00 | 1910.00 | 1662.00 |
| ZNF830 | 393.00 | 405.00 | 443.00 | 413.67 | 497.00 |
| ZNF833P | 16.00 | 11.00 | 21.00 | 16.00 | 27.00 |
| ZNF836 | 34.00 | 37.00 | 34.00 | 35.00 | 38.00 |
| ZNF837 | 39.00 | 32.00 | 35.00 | 35.33 | 52.00 |
| ZNF839 | 979.00 | 1027.00 | 1011.00 | 1005.67 | 1193.00 |
| ZNF84 | 1222.00 | 1250.00 | 1336.00 | 1269.33 | 1380.00 |
| ZNF841 | 447.00 | 447.00 | 534.00 | 476.00 | 418.00 |
| ZNF844 | 4.00 | 7.00 | 1.00 | 4.00 | 1.00 |
| ZNF845 | 179.00 | 134.00 | 172.00 | 161.67 | 189.00 |
| ZNF846 | 65.00 | 70.00 | 58.00 | 64.33 | 48.00 |
| ZNF850 | 6.00 | 6.00 | 13.00 | 8.33 | 11.00 |
| ZNF852 | 119.00 | 84.00 | 96.00 | 99.67 | 59.00 |
| ZNF860 | 7.00 | 10.00 | 7.00 | 8.00 | 7.00 |
| ZNF862 | 1240.00 | 1313.00 | 1401.00 | 1318.00 | 1066.00 |
| ZNF865 | 439.00 | 424.00 | 509.00 | 457.33 | 431.00 |
| ZNF879 | 10.00 | 12.00 | 3.00 | 8.33 | 7.00 |
| ZNF880 | 70.00 | 62.00 | 56.00 | 62.67 | 73.00 |
| ZNF883 | 187.00 | 174.00 | 193.00 | 184.67 | 206.00 |
| ZNF891 | 70.00 | 80.00 | 75.00 | 75.00 | 71.00 |
| ZNF90 | 34.00 | 27.00 | 32.00 | 31.00 | 44.00 |
| ZNF91 | 123.00 | 118.00 | 131.00 | 124.00 | 100.00 |
| ZNF92 | 295.00 | 231.00 | 278.00 | 268.00 | 332.00 |
| ZNF93 | 35.00 | 36.00 | 42.00 | 37.67 | 38.00 |
| ZNFX1 | 2350.00 | 2206.00 | 2456.00 | 2337.33 | 2024.00 |
| ZNHIT1 | 612.00 | 605.00 | 603.00 | 606.67 | 782.00 |
| ZNHIT2 | 385.00 | 382.00 | 430.00 | 399.00 | 540.00 |
| ZNHIT6 | 1943.00 | 1800.00 | 2039.00 | 1927.33 | 2399.00 |
| ZNRD1 | 57.00 | 49.00 | 52.00 | 52.67 | 72.00 |
| ZNRD1-AS1 | 7.00 | 15.00 | 4.00 | 8.67 | 5.00 |
| ZNRF1 | 296.00 | 292.00 | 323.00 | 303.67 | 282.00 |
| ZNRF2 | 76.00 | 81.00 | 105.00 | 87.33 | 81.00 |
| ZNRF2P1 | 17.00 | 15.00 | 19.00 | 17.00 | 16.00 |
| ZNRF2P2 | 9.00 | 4.00 | 7.00 | 6.67 | 6.00 |
| ZNRF3 | 554.00 | 568.00 | 598.00 | 573.33 | 524.00 |
| ZP1 | 49.00 | 39.00 | 40.00 | 42.67 | 12.00 |
| ZP3 | 51.00 | 39.00 | 44.00 | 44.67 | 81.00 |
| ZPLD1 | 16.00 | 14.00 | 19.00 | 16.33 | 6.00 |
| ZPR1 | 1118.00 | 1117.00 | 1241.00 | 1158.67 | 1284.00 |
| ZRANB1 | 1336.00 | 1258.00 | 1385.00 | 1326.33 | 1319.00 |
| ZRANB2 | 3926.00 | 3781.00 | 4343.00 | 4016.67 | 3935.00 |
| ZRANB2-AS2 | 4.00 | 4.00 | 3.00 | 3.67 | 3.00 |
| ZRANB3 | 311.00 | 328.00 | 307.00 | 315.33 | 365.00 |
| ZRSR2 | 437.00 | 508.00 | 542.00 | 495.67 | 539.00 |
| ZSCAN12 | 522.00 | 433.00 | 484.00 | 479.67 | 489.00 |
| ZSCAN12P1 | 75.00 | 51.00 | 79.00 | 68.33 | 76.00 |

| | | | | | |
|---|---|---|---|---|---|
| ZSCAN16 | 84.00 | 104.00 | 111.00 | 99.67 | 115.00 |
| ZSCAN16-AS1 | 103.00 | 99.00 | 106.00 | 102.67 | 103.00 |
| ZSCAN2 | 170.00 | 185.00 | 208.00 | 187.67 | 189.00 |
| ZSCAN20 | 248.00 | 261.00 | 285.00 | 264.67 | 246.00 |
| ZSCAN21 | 216.00 | 246.00 | 269.00 | 243.67 | 224.00 |
| ZSCAN22 | 113.00 | 130.00 | 146.00 | 129.67 | 138.00 |
| ZSCAN23 | 78.00 | 66.00 | 70.00 | 71.33 | 46.00 |
| ZSCAN25 | 404.00 | 419.00 | 422.00 | 415.00 | 375.00 |
| ZSCAN26 | 489.00 | 470.00 | 582.00 | 513.67 | 606.00 |
| ZSCAN29 | 1252.00 | 1206.00 | 1353.00 | 1270.33 | 1062.00 |
| ZSCAN30 | 1427.00 | 1391.00 | 1586.00 | 1468.00 | 1263.00 |
| ZSCAN31 | 28.00 | 27.00 | 30.00 | 28.33 | 31.00 |
| ZSCAN32 | 556.00 | 551.00 | 528.00 | 545.00 | 585.00 |
| ZSCAN5A | 16.00 | 15.00 | 22.00 | 17.67 | 16.00 |
| ZSCAN9 | 559.00 | 547.00 | 539.00 | 548.33 | 512.00 |
| ZSWIM1 | 189.00 | 195.00 | 203.00 | 195.67 | 157.00 |
| ZSWIM3 | 156.00 | 150.00 | 162.00 | 156.00 | 174.00 |
| ZSWIM4 | 419.00 | 426.00 | 486.00 | 443.67 | 574.00 |
| ZSWIM5 | 211.00 | 173.00 | 240.00 | 208.00 | 168.00 |
| ZSWIM6 | 2043.00 | 1877.00 | 2254.00 | 2058.00 | 1480.00 |
| ZSWIM7 | 165.00 | 166.00 | 174.00 | 168.33 | 310.00 |
| ZSWIM8 | 2501.00 | 2538.00 | 2524.00 | 2521.00 | 2024.00 |
| ZSWIM8-AS1 | 249.00 | 255.00 | 269.00 | 257.67 | 220.00 |
| ZUFSP | 387.00 | 345.00 | 399.00 | 377.00 | 430.00 |
| ZW10 | 726.00 | 582.00 | 688.00 | 665.33 | 773.00 |
| ZWILCH | 948.00 | 858.00 | 958.00 | 921.33 | 1148.00 |
| ZWINT | 992.00 | 1085.00 | 1003.00 | 1026.67 | 1579.00 |
| ZXDA | 81.00 | 74.00 | 103.00 | 86.00 | 76.00 |
| ZXDB | 288.00 | 275.00 | 361.00 | 308.00 | 291.00 |
| ZXDC | 1064.00 | 915.00 | 1116.00 | 1031.67 | 887.00 |
| ZYG11A | 3.00 | 0.00 | 0.00 | 1.00 | 0.00 |
| ZYG11B | 642.00 | 629.00 | 796.00 | 689.00 | 690.00 |
| ZYX | 5671.00 | 5478.00 | 5814.00 | 5654.33 | 5629.00 |
| ZZEF1 | 2047.00 | 2001.00 | 2087.00 | 2045.00 | 1660.00 |
| ZZZ3 | 2467.00 | 2251.00 | 2781.00 | 2499.67 | 2799.00 |

| ES2 siARID1A | | | Fold change (siARID1A/scramble, tagwise dispersions ) | P-value (tagwise dispersions) |
|---|---|---|---|---|
| Expression value 2 | Expression value 3 | Average (Expression value ) | | |
| 79.00 | 64.00 | 70.67 | -1.20 | 0.09 |
| 131.00 | 139.00 | 134.67 | -1.19 | 0.03 |
| 1.00 | 1.00 | 1.33 | -3.79 | 0.01 |
| 13.00 | 13.00 | 14.00 | 1.22 | 0.43 |
| 1596.00 | 1794.00 | 1752.00 | 1.07 | 0.10 |
| 430.00 | 455.00 | 455.33 | -1.29 | 0.00 |
| 64.00 | 89.00 | 81.67 | -1.06 | 0.59 |
| 0.00 | 5.00 | 2.33 | -1.57 | 0.50 |
| 365.00 | 433.00 | 421.00 | 1.14 | 0.04 |
| 149.00 | 180.00 | 157.33 | 1.41 | 0.00 |
| 1440.00 | 1522.00 | 1528.00 | 1.30 | 0.00 |
| 1479.00 | 1584.00 | 1527.33 | -1.17 | 0.00 |
| 43.00 | 63.00 | 61.67 | 1.14 | 0.33 |
| 2984.00 | 3192.00 | 3222.33 | 1.03 | 0.38 |
| 4.00 | 2.00 | 3.67 | -1.55 | 0.35 |
| 822.00 | 800.00 | 833.33 | -1.04 | 0.42 |
| 32.00 | 21.00 | 24.33 | 1.57 | 0.03 |
| 9724.00 | 10225.00 | 10069.33 | -1.07 | 0.03 |
| 1243.00 | 1389.00 | 1330.00 | -1.05 | 0.19 |
| 18.00 | 8.00 | 15.33 | 1.29 | 0.31 |
| 474.00 | 488.00 | 455.33 | 1.04 | 0.61 |
| 837.00 | 951.00 | 892.67 | 1.19 | 0.00 |
| 1694.00 | 2003.00 | 1872.00 | -1.10 | 0.02 |
| 4.00 | 5.00 | 3.33 | -1.11 | 1.00 |
| 4.00 | 3.00 | 3.00 | -2.42 | 0.03 |
| 552.00 | 544.00 | 537.67 | -1.67 | 0.00 |
| 13.00 | 8.00 | 14.33 | 1.62 | 0.07 |
| 144.00 | 152.00 | 158.00 | 1.16 | 0.09 |
| 3.00 | 1.00 | 4.00 | -1.51 | 0.37 |
| 1738.00 | 1626.00 | 1755.33 | -1.10 | 0.04 |
| 8.00 | 10.00 | 11.67 | -1.33 | 0.25 |
| 3.00 | 3.00 | 3.33 | -1.31 | 0.68 |
| 265.00 | 299.00 | 276.00 | 1.04 | 0.55 |
| 236.00 | 205.00 | 232.67 | -1.41 | 0.00 |
| 1.00 | 0.00 | 1.33 | -1.94 | 0.40 |
| 140.00 | 201.00 | 175.33 | 1.02 | 0.83 |
| 469.00 | 552.00 | 505.00 | 1.15 | 0.01 |
| 123.00 | 150.00 | 133.67 | -1.37 | 0.00 |
| 607.00 | 677.00 | 643.00 | -1.08 | 0.12 |
| 735.00 | 742.00 | 744.67 | 1.02 | 0.68 |
| 1575.00 | 1595.00 | 1674.00 | -1.06 | 0.15 |
| 238.00 | 214.00 | 232.00 | -1.49 | 0.00 |
| 332.00 | 291.00 | 320.67 | -1.14 | 0.03 |

| | | | | |
|---|---|---|---|---|
| 970.00 | 929.00 | 968.00 | -1.18 | 0.00 |
| 1.00 | 6.00 | 3.67 | -1.90 | 0.12 |
| 3.00 | 2.00 | 2.33 | 1.14 | 1.00 |
| 790.00 | 891.00 | 847.33 | 1.01 | 0.81 |
| 209.00 | 173.00 | 182.67 | -4.42 | 0.00 |
| 1831.00 | 1827.00 | 1840.33 | -1.01 | 0.81 |
| 2527.00 | 2579.00 | 2594.67 | -1.21 | 0.00 |
| 482.00 | 474.00 | 468.33 | 1.12 | 0.06 |
| 194.00 | 170.00 | 185.00 | -1.07 | 0.38 |
| 981.00 | 1166.00 | 1081.33 | 1.09 | 0.06 |
| 974.00 | 1080.00 | 1082.67 | -1.24 | 0.00 |
| 3112.00 | 3512.00 | 3355.00 | 1.11 | 0.00 |
| 4.00 | 2.00 | 2.00 | -1.49 | 0.61 |
| 3578.00 | 3704.00 | 3734.00 | 1.02 | 0.62 |
| 1133.00 | 1116.00 | 1169.67 | -1.02 | 0.75 |
| 231.00 | 266.00 | 244.33 | -1.07 | 0.44 |
| 781.00 | 924.00 | 863.67 | 1.08 | 0.13 |
| 315.00 | 320.00 | 347.67 | 1.28 | 0.00 |
| 1153.00 | 1138.00 | 1147.33 | -1.07 | 0.14 |
| 413.00 | 455.00 | 435.67 | 1.20 | 0.00 |
| 191.00 | 184.00 | 189.33 | 1.01 | 0.89 |
| 3081.00 | 3099.00 | 3242.33 | 1.05 | 0.19 |
| 222.00 | 256.00 | 243.00 | -1.01 | 0.91 |
| 534.00 | 511.00 | 528.67 | 1.13 | 0.07 |
| 187.00 | 201.00 | 196.33 | 1.03 | 0.70 |
| 237.00 | 289.00 | 274.33 | 1.06 | 0.36 |
| 3623.00 | 3195.00 | 3277.33 | -1.32 | 0.00 |
| 275.00 | 261.00 | 269.00 | 1.26 | 0.00 |
| 645.00 | 586.00 | 635.33 | -1.46 | 0.00 |
| 876.00 | 974.00 | 955.33 | -1.19 | 0.00 |
| 287.00 | 273.00 | 297.67 | 1.18 | 0.02 |
| 174.00 | 137.00 | 147.67 | 1.04 | 0.68 |
| 1404.00 | 1605.00 | 1519.00 | 1.10 | 0.03 |
| 2339.00 | 2663.00 | 2514.00 | 1.05 | 0.25 |
| 75.00 | 48.00 | 64.33 | -1.20 | 0.14 |
| 274.00 | 306.00 | 291.00 | -1.46 | 0.00 |
| 5802.00 | 6082.00 | 6036.33 | 1.24 | 0.00 |
| 6599.00 | 6653.00 | 6549.00 | 1.03 | 0.42 |
| 3.00 | 7.00 | 10.00 | -1.53 | 0.16 |
| 3.00 | 6.00 | 6.67 | 1.50 | 0.31 |
| 1366.00 | 1309.00 | 1357.00 | -1.71 | 0.00 |
| 941.00 | 1101.00 | 1067.67 | 1.12 | 0.01 |
| 855.00 | 874.00 | 891.33 | 1.19 | 0.00 |
| 901.00 | 1018.00 | 1018.33 | -1.09 | 0.08 |
| 117.00 | 124.00 | 114.33 | -1.48 | 0.00 |
| 761.00 | 844.00 | 836.00 | 1.04 | 0.37 |
| 220.00 | 270.00 | 268.00 | 1.53 | 0.00 |
| 36.00 | 35.00 | 29.00 | 1.34 | 0.14 |

| | | | | |
|---|---|---|---|---|
| 35.00 | 36.00 | 33.67 | 1.19 | 0.30 |
| 1149.00 | 1169.00 | 1188.00 | -1.11 | 0.01 |
| 224.00 | 203.00 | 207.00 | -1.10 | 0.31 |
| 384.00 | 433.00 | 423.00 | -1.24 | 0.00 |
| 755.00 | 891.00 | 850.33 | -1.01 | 0.77 |
| 15.00 | 12.00 | 12.67 | 1.38 | 0.23 |
| 953.00 | 1111.00 | 1061.00 | 1.26 | 0.00 |
| 92.00 | 100.00 | 105.33 | -1.08 | 0.42 |
| 550.00 | 606.00 | 581.00 | -1.07 | 0.18 |
| 6258.00 | 6329.00 | 6629.33 | -1.06 | 0.12 |
| 11.00 | 2.00 | 8.00 | 1.18 | 0.67 |
| 1648.00 | 1995.00 | 1754.67 | 1.27 | 0.00 |
| 694.00 | 614.00 | 687.67 | -1.29 | 0.00 |
| 1321.00 | 1443.00 | 1469.00 | 1.24 | 0.00 |
| 297.00 | 384.00 | 359.67 | 1.22 | 0.00 |
| 911.00 | 1145.00 | 999.33 | 1.06 | 0.29 |
| 145.00 | 176.00 | 182.00 | -1.20 | 0.02 |
| 796.00 | 850.00 | 851.67 | 1.05 | 0.26 |
| 737.00 | 741.00 | 773.67 | -1.03 | 0.57 |
| 45.00 | 48.00 | 52.00 | 1.20 | 0.17 |
| 341.00 | 364.00 | 379.33 | 1.00 | 0.97 |
| 535.00 | 588.00 | 607.67 | 1.05 | 0.41 |
| 43.00 | 36.00 | 39.00 | 1.99 | 0.00 |
| 764.00 | 828.00 | 805.33 | -1.03 | 0.58 |
| 5474.00 | 5707.00 | 5777.33 | -1.17 | 0.00 |
| 6575.00 | 6436.00 | 6517.67 | 1.00 | 0.97 |
| 2378.00 | 2449.00 | 2409.67 | -1.09 | 0.04 |
| 2450.00 | 2398.00 | 2463.67 | 1.10 | 0.03 |
| 13.00 | 12.00 | 14.33 | 1.76 | 0.03 |
| 596.00 | 606.00 | 660.67 | 1.32 | 0.00 |
| 136.00 | 164.00 | 153.67 | 1.10 | 0.23 |
| 1328.00 | 1357.00 | 1443.33 | 1.32 | 0.00 |
| 409.00 | 381.00 | 412.67 | 1.01 | 0.87 |
| 583.00 | 637.00 | 626.33 | 1.07 | 0.17 |
| 541.00 | 549.00 | 546.67 | -1.03 | 0.61 |
| 75.00 | 83.00 | 80.00 | 2.51 | 0.00 |
| 277.00 | 290.00 | 285.67 | -1.05 | 0.42 |
| 1659.00 | 1859.00 | 1868.33 | 1.15 | 0.01 |
| 623.00 | 639.00 | 663.00 | -1.15 | 0.01 |
| 202.00 | 165.00 | 208.00 | -1.34 | 0.00 |
| 355.00 | 362.00 | 375.00 | -1.15 | 0.01 |
| 15.00 | 32.00 | 21.67 | -1.28 | 0.20 |
| 175.00 | 194.00 | 219.67 | 1.04 | 0.72 |
| 16.00 | 13.00 | 14.33 | -1.18 | 0.55 |
| 307.00 | 296.00 | 302.33 | -1.33 | 0.00 |
| 558.00 | 590.00 | 620.67 | -1.10 | 0.07 |
| 471.00 | 496.00 | 481.67 | -1.07 | 0.23 |
| 3841.00 | 3955.00 | 3829.00 | 1.07 | 0.12 |

| | | | | |
|---|---|---|---|---|
| 4158.00 | 4652.00 | 4283.00 | 1.07 | 0.21 |
| 425.00 | 458.00 | 447.00 | -1.50 | 0.00 |
| 8.00 | 8.00 | 8.00 | 1.08 | 0.89 |
| 623.00 | 699.00 | 676.33 | -1.51 | 0.00 |
| 66.00 | 53.00 | 61.33 | 1.16 | 0.22 |
| 9.00 | 11.00 | 8.67 | -1.20 | 0.61 |
| 50644.00 | 52800.00 | 54528.00 | 1.22 | 0.00 |
| 40345.00 | 39528.00 | 42073.33 | 1.08 | 0.04 |
| 19.00 | 30.00 | 28.33 | 1.20 | 0.37 |
| 43.00 | 27.00 | 38.00 | -1.68 | 0.00 |
| 24.00 | 14.00 | 18.67 | -1.26 | 0.27 |
| 28.00 | 36.00 | 30.67 | 1.11 | 0.62 |
| 1720.00 | 1931.00 | 1947.00 | 1.16 | 0.00 |
| 6497.00 | 6517.00 | 6559.33 | -1.14 | 0.00 |
| 1.00 | 2.00 | 1.33 | -1.48 | 0.76 |
| 16902.00 | 16469.00 | 16539.67 | 1.01 | 0.75 |
| 1038.00 | 1180.00 | 1161.67 | 1.12 | 0.01 |
| 2570.00 | 2667.00 | 2654.33 | 1.01 | 0.84 |
| 1058.00 | 1157.00 | 1199.33 | -1.07 | 0.18 |
| 6536.00 | 7284.00 | 6898.33 | 1.25 | 0.00 |
| 5976.00 | 6527.00 | 6313.00 | 1.12 | 0.00 |
| 407.00 | 440.00 | 452.33 | 1.12 | 0.05 |
| 7.00 | 6.00 | 6.67 | 1.04 | 1.00 |
| 445.00 | 407.00 | 427.00 | -1.06 | 0.34 |
| 501.00 | 567.00 | 556.00 | 1.13 | 0.02 |
| 550.00 | 565.00 | 572.00 | -1.04 | 0.42 |
| 94.00 | 81.00 | 87.67 | 1.02 | 0.85 |
| 1470.00 | 1413.00 | 1469.00 | 1.03 | 0.54 |
| 905.00 | 828.00 | 882.00 | -1.07 | 0.21 |
| 16.00 | 21.00 | 18.00 | -1.37 | 0.13 |
| 301.00 | 352.00 | 332.67 | 1.03 | 0.64 |
| 504.00 | 567.00 | 552.00 | 1.02 | 0.66 |
| 15.00 | 20.00 | 19.00 | -1.23 | 0.27 |
| 2.00 | 5.00 | 4.00 | 1.18 | 0.84 |
| 66.00 | 57.00 | 70.67 | 1.37 | 0.01 |
| 276.00 | 301.00 | 306.00 | 1.31 | 0.00 |
| 36.00 | 42.00 | 45.67 | 1.05 | 0.78 |
| 135.00 | 153.00 | 159.67 | 1.16 | 0.09 |
| 286.00 | 361.00 | 338.67 | -1.23 | 0.00 |
| 2858.00 | 3030.00 | 2955.00 | 1.03 | 0.41 |
| 73.00 | 74.00 | 71.00 | 1.10 | 0.42 |
| 3202.00 | 3143.00 | 3131.67 | -1.56 | 0.00 |
| 3484.00 | 3366.00 | 3468.00 | -1.61 | 0.00 |
| 1426.00 | 1464.00 | 1433.67 | -1.08 | 0.05 |
| 655.00 | 618.00 | 645.00 | -1.41 | 0.00 |
| 268.00 | 326.00 | 317.33 | 1.02 | 0.79 |
| 2.00 | 9.00 | 3.67 | -1.98 | 0.14 |
| 34.00 | 26.00 | 33.00 | -1.24 | 0.19 |

| | | | | |
|---|---|---|---|---|
| 17.00 | 14.00 | 15.67 | -1.01 | 1.00 |
| 422.00 | 466.00 | 437.00 | 1.10 | 0.13 |
| 951.00 | 912.00 | 927.00 | 1.07 | 0.14 |
| 45.00 | 63.00 | 60.33 | -1.72 | 0.00 |
| 2.00 | 5.00 | 4.33 | -2.08 | 0.07 |
| 23.00 | 30.00 | 37.33 | -1.72 | 0.00 |
| 4955.00 | 4965.00 | 4876.33 | -1.17 | 0.00 |
| 13601.00 | 14414.00 | 14083.00 | -1.39 | 0.00 |
| 865.00 | 793.00 | 795.67 | -1.38 | 0.00 |
| 204.00 | 194.00 | 222.33 | -1.19 | 0.02 |
| 10.00 | 8.00 | 7.67 | -2.53 | 0.00 |
| 368.00 | 424.00 | 376.67 | -1.03 | 0.74 |
| 59.00 | 55.00 | 60.33 | -1.57 | 0.00 |
| 17.00 | 29.00 | 26.00 | 1.37 | 0.10 |
| 2399.00 | 2469.00 | 2539.67 | -1.22 | 0.00 |
| 9.00 | 0.00 | 3.33 | 1.01 | 1.00 |
| 189.00 | 206.00 | 193.33 | -1.18 | 0.02 |
| 150.00 | 161.00 | 150.67 | -1.28 | 0.00 |
| 236.00 | 203.00 | 208.00 | 1.28 | 0.01 |
| 2512.00 | 2710.00 | 2716.67 | 1.00 | 0.94 |
| 41.00 | 48.00 | 47.33 | -1.09 | 0.56 |
| 5.00 | 7.00 | 6.33 | -1.69 | 0.10 |
| 8419.00 | 8139.00 | 8416.67 | -1.16 | 0.00 |
| 22.00 | 25.00 | 23.67 | -1.43 | 0.03 |
| 80.00 | 56.00 | 74.67 | -1.85 | 0.00 |
| 7.00 | 5.00 | 7.33 | 1.14 | 0.76 |
| 190.00 | 204.00 | 205.67 | -1.21 | 0.00 |
| 5.00 | 4.00 | 6.00 | -1.84 | 0.06 |
| 6.00 | 4.00 | 6.33 | -1.54 | 0.21 |
| 5486.00 | 5279.00 | 5345.33 | -1.06 | 0.15 |
| 1564.00 | 1593.00 | 1601.33 | -1.22 | 0.00 |
| 538.00 | 530.00 | 494.67 | -1.18 | 0.03 |
| 964.00 | 1132.00 | 1088.67 | 1.06 | 0.16 |
| 70.00 | 61.00 | 64.33 | -1.13 | 0.28 |
| 233.00 | 251.00 | 258.67 | 1.07 | 0.40 |
| 785.00 | 766.00 | 781.33 | -1.12 | 0.01 |
| 856.00 | 928.00 | 922.00 | 1.08 | 0.07 |
| 310.00 | 308.00 | 305.00 | -1.02 | 0.85 |
| 326.00 | 319.00 | 330.67 | -1.03 | 0.67 |
| 156.00 | 145.00 | 148.67 | 1.01 | 0.95 |
| 477.00 | 506.00 | 493.00 | -1.32 | 0.00 |
| 2118.00 | 2094.00 | 2087.67 | 1.02 | 0.70 |
| 1500.00 | 1502.00 | 1500.00 | -1.24 | 0.00 |
| 1557.00 | 1484.00 | 1569.00 | 1.01 | 0.76 |
| 2.00 | 6.00 | 4.67 | 1.25 | 0.69 |
| 561.00 | 498.00 | 530.33 | -1.11 | 0.09 |
| 7006.00 | 7102.00 | 7166.67 | -1.10 | 0.00 |
| 526.00 | 625.00 | 566.67 | -1.05 | 0.38 |

| | | | | |
|---|---|---|---|---|
| 379.00 | 341.00 | 356.33 | -1.44 | 0.00 |
| 885.00 | 871.00 | 885.00 | -1.10 | 0.04 |
| 152.00 | 174.00 | 167.33 | -1.22 | 0.01 |
| 139.00 | 142.00 | 136.33 | 1.03 | 0.76 |
| 7.00 | 1.00 | 3.33 | -1.21 | 0.84 |
| 1025.00 | 1073.00 | 1053.33 | -1.26 | 0.00 |
| 8.00 | 13.00 | 10.00 | -1.65 | 0.05 |
| 69.00 | 52.00 | 55.00 | -2.60 | 0.00 |
| 24.00 | 31.00 | 21.67 | -1.68 | 0.00 |
| 49.00 | 34.00 | 43.67 | -1.91 | 0.00 |
| 8.00 | 12.00 | 9.33 | -2.76 | 0.00 |
| 75.00 | 82.00 | 77.33 | -1.35 | 0.00 |
| 248.00 | 236.00 | 239.33 | 1.22 | 0.00 |
| 1323.00 | 1450.00 | 1379.33 | -1.02 | 0.66 |
| 0.00 | 5.00 | 2.33 | 1.66 | 0.58 |
| 3444.00 | 3631.00 | 3513.00 | -1.20 | 0.00 |
| 89.00 | 97.00 | 95.00 | -1.19 | 0.06 |
| 1261.00 | 1417.00 | 1387.00 | 1.07 | 0.11 |
| 61.00 | 67.00 | 67.33 | -1.46 | 0.00 |
| 305.00 | 362.00 | 350.33 | 1.00 | 0.99 |
| 1733.00 | 1777.00 | 1831.67 | 1.11 | 0.01 |
| 2653.00 | 2619.00 | 2667.33 | -1.03 | 0.42 |
| 1523.00 | 1644.00 | 1650.67 | -1.10 | 0.03 |
| 12.00 | 6.00 | 9.33 | -1.12 | 0.80 |
| 353.00 | 298.00 | 347.33 | -1.13 | 0.07 |
| 164.00 | 160.00 | 164.00 | -1.32 | 0.00 |
| 3930.00 | 4370.00 | 4239.00 | -1.07 | 0.06 |
| 776.00 | 816.00 | 820.67 | 1.04 | 0.35 |
| 28.00 | 40.00 | 34.33 | -1.10 | 0.57 |
| 1234.00 | 1415.00 | 1386.67 | 1.08 | 0.09 |
| 10.00 | 6.00 | 7.67 | -1.79 | 0.04 |
| 1685.00 | 1794.00 | 1841.33 | -1.65 | 0.00 |
| 1796.00 | 1772.00 | 1862.33 | -1.06 | 0.16 |
| 25.00 | 27.00 | 28.00 | -1.07 | 0.72 |
| 29.00 | 22.00 | 27.67 | -1.00 | 1.00 |
| 532.00 | 661.00 | 604.33 | 1.06 | 0.28 |
| 175.00 | 176.00 | 195.67 | -1.02 | 0.79 |
| 40.00 | 37.00 | 38.67 | 1.36 | 0.05 |
| 2573.00 | 2591.00 | 2638.33 | 1.17 | 0.00 |
| 2259.00 | 2113.00 | 2295.67 | -1.01 | 0.78 |
| 71.00 | 86.00 | 80.00 | 1.43 | 0.00 |
| 3279.00 | 3635.00 | 3598.00 | 1.23 | 0.00 |
| 1953.00 | 2047.00 | 2090.33 | 1.18 | 0.00 |
| 1488.00 | 1615.00 | 1573.67 | 1.10 | 0.02 |
| 8.00 | 8.00 | 8.00 | -1.80 | 0.03 |
| 3189.00 | 3573.00 | 3513.67 | -1.10 | 0.01 |
| 4824.00 | 4991.00 | 4946.67 | -1.48 | 0.00 |
| 903.00 | 1146.00 | 1031.33 | 1.20 | 0.00 |

| | | | | |
|---|---|---|---|---|
| 1167.00 | 1163.00 | 1205.33 | -1.04 | 0.39 |
| 3542.00 | 3572.00 | 3758.33 | 1.07 | 0.09 |
| 4158.00 | 3978.00 | 4048.33 | -1.12 | 0.01 |
| 952.00 | 1049.00 | 986.33 | -1.01 | 0.86 |
| 29.00 | 32.00 | 31.33 | -2.00 | 0.00 |
| 546.00 | 518.00 | 515.33 | -1.35 | 0.00 |
| 16.00 | 10.00 | 11.00 | -1.44 | 0.16 |
| 1558.00 | 1369.00 | 1449.00 | 1.03 | 0.55 |
| 2907.00 | 2993.00 | 2891.67 | -1.10 | 0.07 |
| 1402.00 | 1662.00 | 1565.00 | -1.03 | 0.47 |
| 2672.00 | 2999.00 | 2941.33 | 1.16 | 0.00 |
| 569.00 | 590.00 | 612.33 | 1.06 | 0.28 |
| 2.00 | 0.00 | 1.00 | -2.21 | 0.38 |
| 651.00 | 645.00 | 666.00 | 1.01 | 0.86 |
| 862.00 | 818.00 | 835.67 | 1.04 | 0.46 |
| 797.00 | 806.00 | 812.00 | -1.05 | 0.31 |
| 43.00 | 53.00 | 51.00 | 1.19 | 0.21 |
| 0.00 | 5.00 | 2.33 | -1.43 | 0.66 |
| 12.00 | 15.00 | 15.67 | -1.08 | 0.77 |
| 298.00 | 291.00 | 297.33 | -1.18 | 0.01 |
| 2923.00 | 3040.00 | 3032.33 | 1.00 | 0.96 |
| 200.00 | 224.00 | 218.33 | -1.17 | 0.03 |
| 158.00 | 126.00 | 153.33 | -1.24 | 0.02 |
| 787.00 | 881.00 | 862.00 | -1.06 | 0.22 |
| 94.00 | 111.00 | 103.67 | 1.05 | 0.56 |
| 402.00 | 496.00 | 453.33 | -1.11 | 0.10 |
| 60.00 | 81.00 | 66.00 | 1.04 | 0.77 |
| 67.00 | 101.00 | 89.00 | 1.11 | 0.37 |
| 768.00 | 690.00 | 750.33 | -1.06 | 0.32 |
| 1517.00 | 1536.00 | 1541.33 | 1.05 | 0.19 |
| 179.00 | 213.00 | 194.33 | -1.41 | 0.00 |
| 681.00 | 797.00 | 748.00 | 1.03 | 0.50 |
| 286.00 | 282.00 | 302.33 | -1.12 | 0.06 |
| 719.00 | 766.00 | 730.67 | 1.03 | 0.57 |
| 139.00 | 101.00 | 119.00 | 1.11 | 0.33 |
| 601.00 | 587.00 | 585.33 | -1.11 | 0.04 |
| 544.00 | 482.00 | 504.67 | -1.11 | 0.09 |
| 246.00 | 221.00 | 239.67 | 1.06 | 0.47 |
| 711.00 | 767.00 | 735.00 | -1.19 | 0.00 |
| 51.00 | 55.00 | 53.33 | 1.14 | 0.30 |
| 760.00 | 755.00 | 797.67 | 1.08 | 0.13 |
| 2636.00 | 2535.00 | 2597.67 | -1.04 | 0.31 |
| 2284.00 | 2404.00 | 2383.00 | -1.36 | 0.00 |
| 79.00 | 88.00 | 78.67 | -1.70 | 0.00 |
| 2571.00 | 2753.00 | 2686.67 | 1.15 | 0.00 |
| 1214.00 | 1492.00 | 1361.67 | -1.04 | 0.46 |
| 4638.00 | 4606.00 | 4668.00 | -1.63 | 0.00 |
| 596.00 | 731.00 | 672.67 | 1.34 | 0.00 |

| | | | | |
|---|---|---|---|---|
| 64.00 | 74.00 | 65.33 | 1.05 | 0.68 |
| 851.00 | 762.00 | 840.67 | 1.06 | 0.28 |
| 2256.00 | 2613.00 | 2445.67 | 1.02 | 0.66 |
| 226.00 | 193.00 | 202.00 | -1.09 | 0.28 |
| 6432.00 | 6697.00 | 6834.00 | 1.25 | 0.00 |
| 2334.00 | 2361.00 | 2359.67 | -1.07 | 0.06 |
| 741.00 | 710.00 | 738.33 | -1.18 | 0.00 |
| 337.00 | 305.00 | 308.67 | -1.47 | 0.00 |
| 1056.00 | 1159.00 | 1123.33 | 1.09 | 0.05 |
| 31868.00 | 32656.00 | 31548.33 | -1.40 | 0.00 |
| 2173.00 | 1983.00 | 2093.67 | 1.14 | 0.00 |
| 280.00 | 268.00 | 277.00 | -1.26 | 0.01 |
| 1241.00 | 1214.00 | 1288.67 | -1.18 | 0.00 |
| 2887.00 | 3146.00 | 3155.00 | 1.15 | 0.00 |
| 2882.00 | 3210.00 | 3139.00 | -1.12 | 0.00 |
| 361.00 | 417.00 | 389.00 | 1.10 | 0.13 |
| 4.00 | 0.00 | 3.00 | -1.12 | 1.00 |
| 800.00 | 909.00 | 888.33 | 1.07 | 0.11 |
| 467.00 | 514.00 | 527.33 | -1.08 | 0.13 |
| 1103.00 | 1206.00 | 1154.00 | -1.09 | 0.05 |
| 52.00 | 58.00 | 56.33 | -2.06 | 0.00 |
| 5.00 | 4.00 | 6.00 | -1.96 | 0.04 |
| 40.00 | 64.00 | 55.67 | -1.60 | 0.00 |
| 1471.00 | 1721.00 | 1670.67 | 1.10 | 0.02 |
| 1090.00 | 1238.00 | 1250.67 | 1.04 | 0.37 |
| 894.00 | 947.00 | 960.00 | 1.12 | 0.02 |
| 2482.00 | 2736.00 | 2634.67 | -1.52 | 0.00 |
| 457.00 | 526.00 | 516.00 | -1.03 | 0.58 |
| 4915.00 | 5432.00 | 5312.67 | 1.05 | 0.14 |
| 1093.00 | 1325.00 | 1256.67 | 1.07 | 0.14 |
| 427.00 | 346.00 | 400.67 | 1.05 | 0.50 |
| 878.00 | 1010.00 | 962.00 | -1.05 | 0.32 |
| 1.00 | 0.00 | 0.33 | -6.90 | 0.04 |
| 80.00 | 125.00 | 110.00 | -1.19 | 0.08 |
| 1922.00 | 1934.00 | 1984.33 | -1.11 | 0.00 |
| 791.00 | 818.00 | 845.00 | 1.01 | 0.80 |
| 1821.00 | 1930.00 | 1789.00 | -1.03 | 0.56 |
| 15439.00 | 15613.00 | 15196.67 | -1.30 | 0.00 |
| 2323.00 | 2199.00 | 2184.33 | -1.08 | 0.14 |
| 89.00 | 44.00 | 67.67 | -1.46 | 0.01 |
| 3.00 | 14.00 | 8.67 | 1.28 | 0.48 |
| 7.00 | 20.00 | 11.67 | 1.38 | 0.30 |
| 48.00 | 46.00 | 47.67 | -1.32 | 0.02 |
| 111.00 | 121.00 | 119.00 | 1.61 | 0.00 |
| 854.00 | 1030.00 | 980.00 | -1.11 | 0.03 |
| 1269.00 | 1304.00 | 1334.67 | -1.12 | 0.00 |
| 1455.00 | 1505.00 | 1480.33 | -1.17 | 0.00 |
| 435.00 | 533.00 | 514.33 | 1.00 | 0.97 |

| | | | | |
|---|---|---|---|---|
| 1500.00 | 1643.00 | 1635.00 | 1.10 | 0.01 |
| 1077.00 | 1177.00 | 1194.67 | 1.04 | 0.34 |
| 364.00 | 361.00 | 368.67 | -1.08 | 0.18 |
| 12.00 | 19.00 | 16.00 | 1.57 | 0.06 |
| 1504.00 | 1720.00 | 1757.00 | 1.12 | 0.02 |
| 1764.00 | 1826.00 | 1908.67 | -1.09 | 0.03 |
| 18.00 | 23.00 | 23.67 | -1.38 | 0.05 |
| 9.00 | 1.00 | 6.00 | -1.12 | 0.88 |
| 86.00 | 94.00 | 94.67 | 1.14 | 0.16 |
| 2.00 | 2.00 | 2.00 | -1.33 | 0.79 |
| 54.00 | 49.00 | 54.33 | 1.05 | 0.71 |
| 864.00 | 943.00 | 948.67 | 1.14 | 0.00 |
| 6.00 | 8.00 | 7.67 | -1.79 | 0.04 |
| 9.00 | 18.00 | 14.67 | -1.22 | 0.39 |
| 2726.00 | 2818.00 | 2893.67 | -1.02 | 0.55 |
| 1286.00 | 1308.00 | 1343.67 | 1.08 | 0.06 |
| 2381.00 | 2493.00 | 2541.00 | -1.08 | 0.03 |
| 378.00 | 403.00 | 406.33 | 1.09 | 0.10 |
| 445.00 | 494.00 | 485.00 | -1.08 | 0.13 |
| 1031.00 | 1011.00 | 1063.33 | 1.06 | 0.20 |
| 5.00 | 6.00 | 4.67 | 1.06 | 1.00 |
| 6091.00 | 6912.00 | 6374.00 | 1.08 | 0.07 |
| 933.00 | 925.00 | 946.00 | 1.12 | 0.01 |
| 4798.00 | 4800.00 | 4939.33 | 1.05 | 0.17 |
| 20274.00 | 20876.00 | 21119.00 | 1.65 | 0.00 |
| 599.00 | 625.00 | 614.33 | 1.12 | 0.02 |
| 27.00 | 11.00 | 19.67 | 1.12 | 0.61 |
| 1186.00 | 1248.00 | 1184.33 | -1.04 | 0.44 |
| 6.00 | 2.00 | 4.00 | 1.07 | 1.00 |
| 291.00 | 310.00 | 301.33 | -1.19 | 0.01 |
| 97.00 | 87.00 | 90.67 | -2.35 | 0.00 |
| 387.00 | 378.00 | 396.33 | -1.23 | 0.00 |
| 259.00 | 183.00 | 223.33 | 1.24 | 0.03 |
| 315.00 | 312.00 | 317.00 | -1.16 | 0.01 |
| 1240.00 | 1437.00 | 1375.00 | 1.18 | 0.00 |
| 11.00 | 7.00 | 10.33 | -1.40 | 0.22 |
| 1499.00 | 1553.00 | 1536.33 | 1.17 | 0.00 |
| 12346.00 | 12821.00 | 13149.00 | 1.14 | 0.00 |
| 88.00 | 73.00 | 98.00 | 1.19 | 0.19 |
| 1233.00 | 1241.00 | 1266.67 | -1.00 | 0.95 |
| 212.00 | 316.00 | 253.33 | 1.04 | 0.65 |
| 1224.00 | 1505.00 | 1313.67 | 1.06 | 0.37 |
| 168.00 | 151.00 | 143.67 | -1.29 | 0.02 |
| 469.00 | 461.00 | 489.33 | -1.04 | 0.42 |
| 2107.00 | 2336.00 | 2289.00 | -1.18 | 0.00 |
| 190.00 | 197.00 | 221.00 | 1.41 | 0.00 |
| 11.00 | 23.00 | 13.67 | 1.12 | 0.68 |
| 1011.00 | 1067.00 | 1055.67 | -1.01 | 0.85 |

| | | | | |
|---|---|---|---|---|
| 1651.00 | 1826.00 | 1817.67 | 1.05 | 0.25 |
| 695.00 | 760.00 | 748.67 | 1.27 | 0.00 |
| 658.00 | 737.00 | 714.67 | 1.19 | 0.00 |
| 1167.00 | 1155.00 | 1185.67 | -1.02 | 0.72 |
| 681.00 | 752.00 | 717.33 | -1.27 | 0.00 |
| 242.00 | 237.00 | 248.67 | 1.07 | 0.30 |
| 563.00 | 615.00 | 620.00 | 1.23 | 0.00 |
| 509.00 | 576.00 | 582.00 | 1.09 | 0.11 |
| 212.00 | 245.00 | 238.67 | 1.11 | 0.12 |
| 1558.00 | 1588.00 | 1641.33 | 1.03 | 0.44 |
| 368.00 | 380.00 | 394.67 | -1.03 | 0.61 |
| 396.00 | 401.00 | 413.00 | 1.18 | 0.01 |
| 346.00 | 385.00 | 361.67 | 1.12 | 0.05 |
| 560.00 | 512.00 | 528.00 | -1.03 | 0.57 |
| 9.00 | 8.00 | 12.33 | 1.21 | 0.57 |
| 52.00 | 60.00 | 60.67 | -1.17 | 0.19 |
| 3.00 | 2.00 | 4.00 | -1.10 | 1.00 |
| 19.00 | 16.00 | 15.33 | -1.80 | 0.00 |
| 438.00 | 505.00 | 473.33 | -1.08 | 0.17 |
| 5.00 | 1.00 | 4.00 | -3.06 | 0.00 |
| 7.00 | 10.00 | 8.67 | -1.63 | 0.08 |
| 1128.00 | 1302.00 | 1250.00 | -1.01 | 0.81 |
| 16.00 | 28.00 | 25.67 | 1.26 | 0.27 |
| 23.00 | 41.00 | 28.00 | -1.08 | 0.74 |
| 165.00 | 181.00 | 187.00 | 1.06 | 0.44 |
| 1141.00 | 1255.00 | 1225.33 | 1.26 | 0.00 |
| 68.00 | 60.00 | 71.67 | 1.09 | 0.53 |
| 782.00 | 738.00 | 744.33 | -1.28 | 0.00 |
| 1523.00 | 1699.00 | 1596.00 | 1.06 | 0.18 |
| 1170.00 | 1091.00 | 1196.00 | -1.03 | 0.57 |
| 448.00 | 427.00 | 423.67 | -1.14 | 0.07 |
| 2234.00 | 2200.00 | 2272.33 | 1.03 | 0.40 |
| 412.00 | 391.00 | 418.67 | -1.23 | 0.00 |
| 105.00 | 90.00 | 91.67 | -1.02 | 0.92 |
| 7380.00 | 8035.00 | 7692.33 | -1.20 | 0.00 |
| 152.00 | 125.00 | 144.33 | -1.12 | 0.17 |
| 387.00 | 438.00 | 422.67 | 1.02 | 0.76 |
| 1389.00 | 1549.00 | 1462.00 | -1.07 | 0.09 |
| 4.00 | 2.00 | 2.67 | -1.50 | 0.51 |
| 117.00 | 129.00 | 127.33 | 1.32 | 0.00 |
| 1867.00 | 1896.00 | 1841.00 | -1.24 | 0.00 |
| 7804.00 | 7976.00 | 7885.67 | -1.13 | 0.00 |
| 3201.00 | 3133.00 | 3282.67 | -1.04 | 0.37 |
| 612.00 | 560.00 | 621.67 | -1.32 | 0.00 |
| 400.00 | 465.00 | 479.00 | -1.19 | 0.01 |
| 178.00 | 216.00 | 202.33 | -1.01 | 0.94 |
| 1499.00 | 1665.00 | 1668.33 | 1.03 | 0.47 |
| 263.00 | 302.00 | 292.00 | 1.09 | 0.16 |

| | | | | |
|---|---|---|---|---|
| 1457.00 | 1551.00 | 1480.00 | -1.07 | 0.11 |
| 166.00 | 203.00 | 179.00 | 1.39 | 0.00 |
| 1276.00 | 1399.00 | 1429.67 | 1.16 | 0.00 |
| 1913.00 | 2110.00 | 2142.00 | 1.09 | 0.04 |
| 318.00 | 342.00 | 354.67 | 1.18 | 0.01 |
| 1422.00 | 1548.00 | 1546.33 | 1.10 | 0.02 |
| 875.00 | 833.00 | 881.00 | -1.31 | 0.00 |
| 678.00 | 841.00 | 800.33 | 1.12 | 0.04 |
| 2384.00 | 2636.00 | 2604.67 | 1.02 | 0.63 |
| 1749.00 | 2039.00 | 2007.00 | 1.06 | 0.17 |
| 15.00 | 24.00 | 18.67 | 1.02 | 0.93 |
| 1028.00 | 1063.00 | 1032.67 | -1.18 | 0.00 |
| 990.00 | 1069.00 | 1065.67 | 1.11 | 0.01 |
| 96.00 | 97.00 | 95.33 | 1.92 | 0.00 |
| 756.00 | 790.00 | 797.00 | 1.77 | 0.00 |
| 66.00 | 84.00 | 71.00 | 1.39 | 0.01 |
| 209.00 | 214.00 | 217.33 | 3.47 | 0.00 |
| 1.00 | 1.00 | 2.00 | -1.65 | 0.48 |
| 1080.00 | 988.00 | 1016.00 | 1.02 | 0.68 |
| 1345.00 | 1370.00 | 1283.67 | 1.13 | 0.07 |
| 92.00 | 85.00 | 88.00 | -1.07 | 0.54 |
| 16.00 | 21.00 | 19.33 | -1.26 | 0.23 |
| 0.00 | 1.00 | 1.33 | -4.95 | 0.00 |
| 114.00 | 136.00 | 125.67 | 1.12 | 0.18 |
| 76.00 | 88.00 | 86.67 | -1.17 | 0.13 |
| 2444.00 | 2491.00 | 2491.00 | -1.05 | 0.21 |
| 960.00 | 996.00 | 998.33 | -1.11 | 0.01 |
| 276.00 | 295.00 | 303.00 | 1.03 | 0.69 |
| 22.00 | 30.00 | 26.33 | -1.04 | 0.90 |
| 1286.00 | 1427.00 | 1386.33 | -1.00 | 0.98 |
| 11.00 | 15.00 | 12.67 | -1.23 | 0.47 |
| 261.00 | 241.00 | 271.67 | -1.09 | 0.25 |
| 2539.00 | 2735.00 | 2686.00 | -1.14 | 0.00 |
| 480.00 | 467.00 | 451.00 | -1.37 | 0.00 |
| 398.00 | 548.00 | 496.33 | 1.02 | 0.77 |
| 279.00 | 353.00 | 319.67 | -1.01 | 0.88 |
| 10298.00 | 11585.00 | 11445.00 | -1.11 | 0.00 |
| 1266.00 | 1433.00 | 1442.67 | -1.13 | 0.01 |
| 10089.00 | 9378.00 | 9466.33 | -1.28 | 0.00 |
| 1346.00 | 1365.00 | 1303.67 | 1.03 | 0.65 |
| 1532.00 | 1725.00 | 1694.00 | 1.08 | 0.07 |
| 233.00 | 249.00 | 249.67 | 1.15 | 0.03 |
| 524.00 | 662.00 | 599.33 | -1.07 | 0.19 |
| 922.00 | 979.00 | 1005.33 | -1.03 | 0.52 |
| 304.00 | 334.00 | 328.00 | 1.04 | 0.50 |
| 2306.00 | 2209.00 | 2224.00 | -1.11 | 0.03 |
| 210.00 | 237.00 | 236.33 | -1.08 | 0.23 |
| 254.00 | 248.00 | 250.33 | 1.18 | 0.04 |

| | | | | |
|---|---|---|---|---|
| 319.00 | 266.00 | 307.33 | -1.08 | 0.31 |
| 14.00 | 20.00 | 15.67 | -1.89 | 0.00 |
| 75.00 | 64.00 | 72.33 | -1.70 | 0.00 |
| 53.00 | 54.00 | 57.67 | -1.65 | 0.00 |
| 875.00 | 1057.00 | 967.33 | 1.04 | 0.38 |
| 894.00 | 944.00 | 935.00 | 1.10 | 0.04 |
| 3122.00 | 3647.00 | 3477.67 | 1.06 | 0.12 |
| 186.00 | 216.00 | 214.33 | -1.48 | 0.00 |
| 45.00 | 58.00 | 45.67 | 1.63 | 0.00 |
| 722.00 | 775.00 | 724.33 | -2.07 | 0.00 |
| 157.00 | 149.00 | 147.00 | 1.02 | 0.86 |
| 96.00 | 105.00 | 97.67 | -1.04 | 0.70 |
| 65.00 | 78.00 | 71.00 | 1.16 | 0.18 |
| 58.00 | 61.00 | 62.33 | 1.02 | 0.85 |
| 70.00 | 66.00 | 71.67 | -1.45 | 0.00 |
| 43.00 | 40.00 | 42.67 | 1.05 | 0.76 |
| 92.00 | 109.00 | 109.33 | 1.00 | 1.00 |
| 1603.00 | 1673.00 | 1656.67 | -1.01 | 0.75 |
| 263.00 | 293.00 | 274.00 | -1.29 | 0.00 |
| 309.00 | 400.00 | 363.67 | 1.01 | 0.85 |
| 1.00 | 2.00 | 2.67 | 1.54 | 0.59 |
| 345.00 | 406.00 | 367.67 | 1.09 | 0.17 |
| 249.00 | 325.00 | 295.67 | -1.19 | 0.02 |
| 891.00 | 882.00 | 871.33 | -1.25 | 0.00 |
| 2394.00 | 2488.00 | 2412.33 | -1.26 | 0.00 |
| 5.00 | 7.00 | 5.33 | -1.57 | 0.22 |
| 339.00 | 371.00 | 383.33 | 1.07 | 0.29 |
| 10.00 | 15.00 | 12.00 | 1.36 | 0.26 |
| 6.00 | 6.00 | 8.67 | -1.01 | 1.00 |
| 44.00 | 38.00 | 41.67 | 1.13 | 0.40 |
| 236.00 | 206.00 | 222.00 | 1.11 | 0.15 |
| 890.00 | 892.00 | 891.00 | -1.20 | 0.00 |
| 502.00 | 520.00 | 534.67 | -1.06 | 0.29 |
| 1306.00 | 1234.00 | 1287.67 | -1.06 | 0.16 |
| 1295.00 | 1481.00 | 1475.33 | -1.16 | 0.00 |
| 3848.00 | 4142.00 | 3903.00 | 1.33 | 0.00 |
| 3.00 | 8.00 | 4.00 | -1.01 | 1.00 |
| 1496.00 | 1582.00 | 1563.33 | -1.37 | 0.00 |
| 241.00 | 285.00 | 261.00 | 1.02 | 0.76 |
| 2362.00 | 2361.00 | 2362.33 | -1.27 | 0.00 |
| 18.00 | 17.00 | 18.67 | -1.34 | 0.16 |
| 231.00 | 237.00 | 247.00 | -1.31 | 0.00 |
| 297.00 | 294.00 | 287.67 | -1.16 | 0.03 |
| 3009.00 | 3588.00 | 3457.67 | 1.21 | 0.00 |
| 18.00 | 11.00 | 12.67 | -2.17 | 0.00 |
| 4101.00 | 4708.00 | 4567.67 | 1.28 | 0.00 |
| 1401.00 | 1852.00 | 1566.33 | 1.41 | 0.00 |
| 14141.00 | 15223.00 | 14912.67 | -1.03 | 0.47 |

| | | | | |
|---|---|---|---|---|
| 1688.00 | 1533.00 | 1576.33 | -1.47 | 0.00 |
| 1622.00 | 1808.00 | 1714.00 | -1.35 | 0.00 |
| 4753.00 | 5598.00 | 5424.00 | 1.14 | 0.00 |
| 14.00 | 19.00 | 15.67 | -1.97 | 0.00 |
| 1653.00 | 1651.00 | 1666.00 | 1.07 | 0.09 |
| 26582.00 | 30667.00 | 29725.00 | 1.15 | 0.00 |
| 1312.00 | 1685.00 | 1582.33 | 1.18 | 0.00 |
| 58.00 | 47.00 | 58.33 | -1.02 | 0.92 |
| 1.00 | 2.00 | 1.00 | -2.51 | 0.23 |
| 509.00 | 600.00 | 578.33 | 1.11 | 0.05 |
| 7447.00 | 8198.00 | 8166.67 | -1.27 | 0.00 |
| 2228.00 | 2440.00 | 2377.33 | 1.01 | 0.83 |
| 1948.00 | 2209.00 | 2165.67 | -1.25 | 0.00 |
| 1.00 | 7.00 | 3.33 | 1.61 | 0.49 |
| 54.00 | 79.00 | 72.33 | -1.39 | 0.00 |
| 24.00 | 13.00 | 23.00 | -1.29 | 0.17 |
| 6477.00 | 6695.00 | 6645.67 | -1.07 | 0.07 |
| 720.00 | 981.00 | 872.33 | 1.08 | 0.30 |
| 2219.00 | 2291.00 | 2266.67 | 1.04 | 0.30 |
| 1961.00 | 2081.00 | 2055.67 | -1.00 | 0.91 |
| 232.00 | 240.00 | 254.67 | -1.51 | 0.00 |
| 1403.00 | 1441.00 | 1501.67 | 1.15 | 0.00 |
| 1508.00 | 1550.00 | 1618.00 | 1.10 | 0.02 |
| 1039.00 | 1302.00 | 1194.67 | -1.15 | 0.01 |
| 82.00 | 103.00 | 92.00 | 1.60 | 0.00 |
| 1731.00 | 1641.00 | 1743.00 | -1.03 | 0.52 |
| 226.00 | 217.00 | 227.00 | 1.07 | 0.33 |
| 7402.00 | 7380.00 | 7433.33 | -1.12 | 0.00 |
| 6242.00 | 6221.00 | 6260.33 | -1.05 | 0.14 |
| 1300.00 | 1363.00 | 1443.33 | 1.13 | 0.01 |
| 1272.00 | 1365.00 | 1334.67 | 1.06 | 0.15 |
| 4906.00 | 4931.00 | 5017.33 | -1.06 | 0.11 |
| 1858.00 | 1984.00 | 1930.33 | -1.03 | 0.40 |
| 1551.00 | 1686.00 | 1636.33 | 1.06 | 0.17 |
| 700.00 | 838.00 | 795.67 | 1.13 | 0.01 |
| 49.00 | 49.00 | 48.33 | -1.08 | 0.58 |
| 1166.00 | 1386.00 | 1316.33 | 1.07 | 0.11 |
| 522.00 | 564.00 | 553.00 | 1.03 | 0.53 |
| 704.00 | 729.00 | 742.67 | 1.55 | 0.00 |
| 220.00 | 239.00 | 237.67 | 1.05 | 0.46 |
| 1192.00 | 1106.00 | 1136.33 | -1.08 | 0.12 |
| 360.00 | 404.00 | 379.67 | -1.09 | 0.11 |
| 332.00 | 302.00 | 332.33 | 1.05 | 0.48 |
| 2046.00 | 1974.00 | 2073.00 | -1.01 | 0.89 |
| 376.00 | 400.00 | 383.00 | -1.08 | 0.18 |
| 13.00 | 3.00 | 5.67 | -1.28 | 0.65 |
| 90.00 | 78.00 | 84.00 | -1.01 | 1.00 |
| 688.00 | 721.00 | 702.00 | -1.09 | 0.07 |

| | | | | |
|---|---|---|---|---|
| 631.00 | 690.00 | 668.67 | -1.06 | 0.17 |
| 39.00 | 31.00 | 33.67 | 1.33 | 0.08 |
| 3755.00 | 3971.00 | 3928.00 | -1.10 | 0.01 |
| 943.00 | 802.00 | 949.00 | -1.41 | 0.00 |
| 828.00 | 933.00 | 837.33 | -1.07 | 0.32 |
| 9.00 | 19.00 | 13.33 | 1.40 | 0.20 |
| 0.00 | 0.00 | 0.00 | -14.45 | 0.07 |
| 1825.00 | 1861.00 | 1907.33 | 1.15 | 0.00 |
| 4384.00 | 4851.00 | 4934.67 | 1.05 | 0.25 |
| 644.00 | 750.00 | 725.67 | 1.14 | 0.01 |
| 2095.00 | 2152.00 | 2195.33 | -1.02 | 0.60 |
| 1275.00 | 1189.00 | 1254.33 | -1.04 | 0.47 |
| 2609.00 | 2795.00 | 2706.67 | 1.20 | 0.00 |
| 485.00 | 575.00 | 567.33 | 1.37 | 0.00 |
| 36.00 | 39.00 | 44.00 | 1.09 | 0.59 |
| 141.00 | 182.00 | 163.67 | -1.01 | 0.92 |
| 408.00 | 428.00 | 443.33 | 2.16 | 0.00 |
| 27.00 | 18.00 | 20.67 | 1.39 | 0.15 |
| 6224.00 | 6776.00 | 6609.67 | -1.06 | 0.07 |
| 5020.00 | 5017.00 | 5134.00 | -1.17 | 0.00 |
| 1029.00 | 975.00 | 1083.67 | 1.19 | 0.00 |
| 166.00 | 215.00 | 202.00 | 1.69 | 0.00 |
| 245.00 | 203.00 | 210.00 | 1.36 | 0.00 |
| 32.00 | 42.00 | 34.67 | 2.01 | 0.00 |
| 27.00 | 18.00 | 25.67 | 1.13 | 0.53 |
| 1.00 | 2.00 | 1.33 | -2.63 | 0.12 |
| 8.00 | 2.00 | 5.67 | 1.12 | 0.86 |
| 13.00 | 14.00 | 15.33 | -1.34 | 0.20 |
| 23.00 | 29.00 | 23.67 | -1.49 | 0.02 |
| 224.00 | 273.00 | 258.00 | 1.12 | 0.12 |
| 30.00 | 36.00 | 31.33 | -1.49 | 0.01 |
| 512.00 | 526.00 | 533.33 | -1.03 | 0.59 |
| 368.00 | 416.00 | 385.67 | 1.04 | 0.53 |
| 365.00 | 399.00 | 408.00 | 1.19 | 0.00 |
| 589.00 | 699.00 | 632.67 | 1.11 | 0.18 |
| 465.00 | 497.00 | 505.33 | 1.14 | 0.02 |
| 29550.00 | 32233.00 | 31141.00 | -1.10 | 0.00 |
| 1351.00 | 1505.00 | 1456.33 | 1.04 | 0.30 |
| 1051.00 | 1158.00 | 1097.33 | 1.15 | 0.01 |
| 985.00 | 1021.00 | 1032.00 | 1.00 | 0.99 |
| 2683.00 | 2873.00 | 2992.33 | 1.26 | 0.00 |
| 194.00 | 227.00 | 216.67 | 1.12 | 0.09 |
| 1067.00 | 1203.00 | 1203.33 | 1.08 | 0.12 |
| 57.00 | 49.00 | 58.33 | -1.12 | 0.38 |
| 8.00 | 5.00 | 7.00 | 1.04 | 1.00 |
| 103.00 | 83.00 | 100.67 | -1.05 | 0.69 |
| 84.00 | 84.00 | 92.67 | -1.16 | 0.12 |
| 1272.00 | 1427.00 | 1343.00 | 1.01 | 0.88 |

| | | | | |
|---|---|---|---|---|
| 285.00 | 327.00 | 296.67 | -2.05 | 0.00 |
| 980.00 | 968.00 | 1032.00 | -1.02 | 0.63 |
| 2652.00 | 2582.00 | 2635.67 | -1.07 | 0.09 |
| 111.00 | 124.00 | 110.00 | -1.07 | 0.47 |
| 1564.00 | 1553.00 | 1589.33 | -1.12 | 0.00 |
| 7.00 | 16.00 | 11.00 | -1.53 | 0.09 |
| 3789.00 | 3796.00 | 3807.67 | -1.09 | 0.02 |
| 71.00 | 78.00 | 83.00 | 1.16 | 0.19 |
| 1542.00 | 1477.00 | 1515.00 | 1.00 | 0.96 |
| 6231.00 | 6323.00 | 6608.00 | 1.08 | 0.04 |
| 2735.00 | 2862.00 | 2844.67 | -1.04 | 0.29 |
| 3630.00 | 4251.00 | 4166.33 | 1.04 | 0.34 |
| 1050.00 | 1065.00 | 1136.67 | 1.13 | 0.02 |
| 2186.00 | 2331.00 | 2281.67 | 1.13 | 0.00 |
| 3128.00 | 3231.00 | 3322.00 | -1.20 | 0.00 |
| 1366.00 | 1526.00 | 1509.33 | 1.08 | 0.05 |
| 802.00 | 787.00 | 810.33 | -1.11 | 0.02 |
| 2113.00 | 2026.00 | 2052.33 | -1.04 | 0.37 |
| 2552.00 | 2605.00 | 2539.00 | -1.05 | 0.32 |
| 379.00 | 410.00 | 384.33 | -1.11 | 0.09 |
| 377.00 | 437.00 | 396.33 | 1.00 | 1.00 |
| 1160.00 | 1316.00 | 1316.00 | -1.07 | 0.13 |
| 1112.00 | 1127.00 | 1163.33 | 1.04 | 0.37 |
| 308.00 | 333.00 | 367.00 | 1.16 | 0.07 |
| 3645.00 | 4273.00 | 4067.33 | -1.07 | 0.10 |
| 1257.00 | 1311.00 | 1319.00 | 1.11 | 0.01 |
| 213.00 | 227.00 | 233.67 | -1.08 | 0.26 |
| 18.00 | 19.00 | 19.67 | -1.13 | 0.54 |
| 7.00 | 10.00 | 7.67 | 2.46 | 0.02 |
| 1653.00 | 1795.00 | 1723.67 | -1.08 | 0.06 |
| 866.00 | 967.00 | 930.67 | -1.23 | 0.00 |
| 2766.00 | 3201.00 | 3039.67 | -1.03 | 0.37 |
| 493.00 | 537.00 | 528.67 | 1.39 | 0.00 |
| 21.00 | 32.00 | 21.67 | -1.46 | 0.05 |
| 1778.00 | 1785.00 | 1791.33 | -1.17 | 0.00 |
| 1981.00 | 1976.00 | 2045.33 | -1.10 | 0.01 |
| 151.00 | 102.00 | 130.00 | -1.38 | 0.00 |
| 1269.00 | 1421.00 | 1389.33 | 1.24 | 0.00 |
| 1.00 | 6.00 | 6.00 | -1.29 | 0.55 |
| 134.00 | 148.00 | 138.33 | -1.90 | 0.00 |
| 10.00 | 16.00 | 14.67 | -1.29 | 0.22 |
| 11003.00 | 12695.00 | 11809.33 | -1.01 | 0.89 |
| 176.00 | 183.00 | 191.67 | -1.39 | 0.00 |
| 569.00 | 490.00 | 544.00 | -1.01 | 0.85 |
| 12.00 | 6.00 | 7.33 | -1.19 | 0.68 |
| 644.00 | 651.00 | 655.33 | -1.19 | 0.00 |
| 246.00 | 242.00 | 247.33 | 1.11 | 0.24 |
| 2246.00 | 2020.00 | 2058.00 | -1.28 | 0.00 |

| | | | | |
|---|---|---|---|---|
| 657.00 | 631.00 | 649.33 | -1.18 | 0.00 |
| 9.00 | 6.00 | 11.33 | -1.66 | 0.03 |
| 1682.00 | 1672.00 | 1806.33 | -1.01 | 0.86 |
| 906.00 | 910.00 | 862.00 | -1.00 | 0.98 |
| 1182.00 | 1303.00 | 1200.33 | -1.10 | 0.11 |
| 38.00 | 45.00 | 48.00 | 1.12 | 0.47 |
| 144.00 | 164.00 | 157.67 | -1.16 | 0.04 |
| 27.00 | 19.00 | 22.00 | -1.24 | 0.28 |
| 7035.00 | 7084.00 | 7262.33 | -1.26 | 0.00 |
| 2452.00 | 2787.00 | 2783.00 | 1.19 | 0.00 |
| 2625.00 | 2755.00 | 2782.00 | -1.09 | 0.02 |
| 111.00 | 95.00 | 105.67 | -1.21 | 0.03 |
| 1777.00 | 1723.00 | 1802.00 | -1.14 | 0.00 |
| 1912.00 | 1920.00 | 1900.67 | -1.13 | 0.01 |
| 1058.00 | 976.00 | 1047.33 | -1.88 | 0.00 |
| 1019.00 | 1043.00 | 1025.67 | -1.31 | 0.00 |
| 227.00 | 244.00 | 249.67 | -1.26 | 0.00 |
| 7454.00 | 7777.00 | 7605.67 | -1.22 | 0.00 |
| 906.00 | 973.00 | 1026.67 | 1.00 | 0.94 |
| 7.00 | 23.00 | 14.67 | -1.52 | 0.05 |
| 8.00 | 6.00 | 10.33 | -1.95 | 0.00 |
| 2452.00 | 2693.00 | 2608.33 | -1.20 | 0.00 |
| 103.00 | 93.00 | 103.00 | -1.03 | 0.79 |
| 6.00 | 7.00 | 6.67 | 1.23 | 0.62 |
| 259.00 | 270.00 | 262.00 | 1.23 | 0.00 |
| 503.00 | 504.00 | 510.67 | -1.20 | 0.00 |
| 209.00 | 206.00 | 219.67 | -1.50 | 0.00 |
| 922.00 | 983.00 | 980.67 | -1.03 | 0.43 |
| 1101.00 | 1123.00 | 1104.33 | -1.07 | 0.11 |
| 1128.00 | 1137.00 | 1138.33 | -2.62 | 0.00 |
| 1922.00 | 1776.00 | 1836.00 | -1.09 | 0.08 |
| 695.00 | 711.00 | 691.67 | 1.04 | 0.41 |
| 102.00 | 119.00 | 113.33 | 1.21 | 0.04 |
| 319.00 | 334.00 | 344.33 | -1.49 | 0.00 |
| 357.00 | 406.00 | 376.33 | -1.02 | 0.75 |
| 720.00 | 873.00 | 781.33 | -1.02 | 0.71 |
| 77.00 | 88.00 | 79.33 | -1.61 | 0.00 |
| 328.00 | 319.00 | 347.00 | -2.07 | 0.00 |
| 1305.00 | 1435.00 | 1382.67 | -1.04 | 0.32 |
| 2752.00 | 2866.00 | 2967.00 | 1.07 | 0.09 |
| 4.00 | 14.00 | 10.67 | -1.05 | 0.91 |
| 1275.00 | 1501.00 | 1420.00 | 1.14 | 0.00 |
| 75.00 | 65.00 | 60.33 | -1.75 | 0.00 |
| 384.00 | 417.00 | 398.67 | 1.14 | 0.03 |
| 593.00 | 649.00 | 656.33 | 1.13 | 0.03 |
| 1.00 | 5.00 | 2.33 | -1.70 | 0.38 |
| 202.00 | 228.00 | 211.00 | 1.16 | 0.09 |
| 607.00 | 726.00 | 658.33 | 1.06 | 0.39 |

| | | | | |
|---|---|---|---|---|
| 34.00 | 46.00 | 36.33 | -1.12 | 0.47 |
| 1770.00 | 1631.00 | 1863.67 | 1.10 | 0.07 |
| 868.00 | 892.00 | 951.67 | -1.05 | 0.31 |
| 388.00 | 409.00 | 421.33 | 1.07 | 0.24 |
| 433.00 | 467.00 | 462.67 | 1.13 | 0.02 |
| 660.00 | 658.00 | 658.67 | -1.11 | 0.03 |
| 95.00 | 101.00 | 101.00 | -1.23 | 0.02 |
| 623.00 | 733.00 | 676.00 | 1.26 | 0.00 |
| 293.00 | 347.00 | 305.33 | 1.05 | 0.51 |
| 159.00 | 195.00 | 180.00 | -1.00 | 1.00 |
| 3418.00 | 3373.00 | 3482.00 | 1.08 | 0.06 |
| 1837.00 | 1974.00 | 2052.67 | 1.06 | 0.18 |
| 1454.00 | 1629.00 | 1583.67 | 1.16 | 0.00 |
| 577.00 | 686.00 | 611.00 | 1.12 | 0.07 |
| 396.00 | 391.00 | 406.33 | -1.02 | 0.72 |
| 1837.00 | 1847.00 | 1865.00 | 1.19 | 0.00 |
| 5.00 | 6.00 | 6.00 | -1.07 | 1.00 |
| 884.00 | 1000.00 | 959.33 | 1.12 | 0.01 |
| 255.00 | 285.00 | 253.00 | 1.24 | 0.02 |
| 20.00 | 13.00 | 18.33 | -1.07 | 0.79 |
| 61.00 | 66.00 | 59.33 | 1.22 | 0.14 |
| 1134.00 | 1200.00 | 1199.67 | 1.90 | 0.00 |
| 340.00 | 323.00 | 353.00 | -1.16 | 0.01 |
| 1237.00 | 1296.00 | 1342.33 | 1.07 | 0.12 |
| 194.00 | 206.00 | 212.33 | 1.16 | 0.03 |
| 785.00 | 873.00 | 846.33 | -1.02 | 0.67 |
| 653.00 | 724.00 | 680.67 | -1.11 | 0.03 |
| 1039.00 | 1060.00 | 1064.33 | 1.01 | 0.80 |
| 44.00 | 50.00 | 53.33 | 1.28 | 0.07 |
| 735.00 | 825.00 | 782.00 | -1.06 | 0.20 |
| 799.00 | 933.00 | 860.33 | -1.13 | 0.01 |
| 451.00 | 446.00 | 455.33 | -1.04 | 0.48 |
| 38.00 | 31.00 | 33.67 | -1.35 | 0.03 |
| 226.00 | 239.00 | 242.00 | -1.16 | 0.05 |
| 560.00 | 611.00 | 609.67 | 1.20 | 0.00 |
| 592.00 | 610.00 | 605.67 | -1.12 | 0.02 |
| 177.00 | 194.00 | 185.33 | 1.07 | 0.34 |
| 302.00 | 291.00 | 309.00 | -1.23 | 0.00 |
| 4117.00 | 4631.00 | 4470.67 | 1.91 | 0.00 |
| 3562.00 | 3890.00 | 3935.67 | 1.36 | 0.00 |
| 3575.00 | 3738.00 | 3900.00 | 1.18 | 0.00 |
| 4555.00 | 4832.00 | 4934.33 | 1.14 | 0.00 |
| 2843.00 | 3122.00 | 3187.00 | 1.19 | 0.00 |
| 1617.00 | 1673.00 | 1723.67 | 1.17 | 0.00 |
| 2691.00 | 2901.00 | 2960.00 | 1.38 | 0.00 |
| 880.00 | 930.00 | 955.33 | 1.21 | 0.00 |
| 517.00 | 467.00 | 481.00 | 1.08 | 0.24 |
| 2146.00 | 2098.00 | 2195.67 | 1.00 | 0.93 |

| | | | | |
|---|---|---|---|---|
| 12.00 | 20.00 | 17.33 | 1.65 | 0.04 |
| 60.00 | 89.00 | 74.00 | 1.21 | 0.17 |
| 525.00 | 576.00 | 579.33 | 1.35 | 0.00 |
| 483.00 | 526.00 | 541.33 | -1.00 | 0.97 |
| 483.00 | 541.00 | 540.33 | -1.06 | 0.31 |
| 372.00 | 400.00 | 404.33 | 1.03 | 0.62 |
| 502.00 | 625.00 | 590.33 | -1.21 | 0.00 |
| 63.00 | 71.00 | 72.67 | -1.03 | 0.81 |
| 31.00 | 41.00 | 38.00 | -1.09 | 0.54 |
| 616.00 | 607.00 | 615.33 | -1.25 | 0.00 |
| 512.00 | 447.00 | 469.33 | -1.03 | 0.66 |
| 181.00 | 196.00 | 191.00 | -1.32 | 0.00 |
| 53.00 | 37.00 | 43.00 | 1.30 | 0.07 |
| 146.00 | 143.00 | 141.00 | -1.35 | 0.00 |
| 1469.00 | 1455.00 | 1470.33 | -1.39 | 0.00 |
| 466.00 | 569.00 | 518.67 | 1.10 | 0.08 |
| 59.00 | 58.00 | 68.00 | -1.15 | 0.27 |
| 495.00 | 520.00 | 541.00 | 1.10 | 0.08 |
| 55.00 | 69.00 | 66.33 | -1.13 | 0.28 |
| 792.00 | 867.00 | 831.00 | -2.01 | 0.00 |
| 12.00 | 9.00 | 9.67 | 1.19 | 0.59 |
| 32.00 | 39.00 | 33.67 | 1.31 | 0.10 |
| 6771.00 | 6986.00 | 6864.00 | -1.28 | 0.00 |
| 5.00 | 8.00 | 10.00 | -1.05 | 0.90 |
| 2591.00 | 2929.00 | 2893.33 | 1.09 | 0.03 |
| 33.00 | 38.00 | 34.00 | -2.15 | 0.00 |
| 13876.00 | 15260.00 | 14724.33 | -1.22 | 0.00 |
| 598.00 | 559.00 | 625.33 | -1.01 | 0.93 |
| 170.00 | 183.00 | 178.00 | -1.11 | 0.13 |
| 28.00 | 35.00 | 34.00 | -1.06 | 0.70 |
| 894.00 | 877.00 | 933.33 | -1.04 | 0.40 |
| 455.00 | 475.00 | 470.00 | -1.04 | 0.49 |
| 291.00 | 335.00 | 329.33 | -1.00 | 0.97 |
| 109.00 | 149.00 | 135.67 | 2.70 | 0.00 |
| 570.00 | 593.00 | 603.33 | 1.04 | 0.47 |
| 1187.00 | 1244.00 | 1275.33 | -1.04 | 0.30 |
| 1918.00 | 2008.00 | 1919.33 | -1.00 | 1.00 |
| 532.00 | 533.00 | 549.00 | 1.18 | 0.00 |
| 387.00 | 429.00 | 425.67 | 1.05 | 0.48 |
| 50.00 | 48.00 | 54.67 | -1.47 | 0.00 |
| 1703.00 | 1668.00 | 1657.00 | -1.23 | 0.00 |
| 56.00 | 71.00 | 57.67 | -1.08 | 0.58 |
| 798.00 | 919.00 | 918.33 | 1.10 | 0.04 |
| 956.00 | 920.00 | 963.67 | -1.08 | 0.07 |
| 2.00 | 4.00 | 2.67 | -1.26 | 0.82 |
| 67.00 | 58.00 | 64.67 | -1.73 | 0.00 |
| 761.00 | 836.00 | 818.67 | -1.06 | 0.26 |
| 2.00 | 1.00 | 1.33 | -1.71 | 0.56 |

| | | | | |
|---|---|---|---|---|
| 1061.00 | 1231.00 | 1237.67 | 1.06 | 0.27 |
| 9910.00 | 11134.00 | 11056.00 | 1.07 | 0.07 |
| 1142.00 | 1265.00 | 1255.33 | 1.15 | 0.00 |
| 10069.00 | 10843.00 | 10570.67 | -1.03 | 0.35 |
| 193.00 | 202.00 | 212.67 | 1.16 | 0.05 |
| 55.00 | 65.00 | 61.67 | -1.05 | 0.70 |
| 2206.00 | 2573.00 | 2375.67 | -1.05 | 0.29 |
| 5.00 | 10.00 | 7.33 | -1.47 | 0.24 |
| 1054.00 | 1185.00 | 1165.00 | 1.03 | 0.50 |
| 22.00 | 24.00 | 25.00 | 1.54 | 0.03 |
| 10.00 | 9.00 | 13.00 | 1.90 | 0.02 |
| 142.00 | 147.00 | 143.00 | -1.10 | 0.34 |
| 77.00 | 72.00 | 84.00 | 1.09 | 0.45 |
| 1786.00 | 2103.00 | 2055.33 | 1.15 | 0.01 |
| 3678.00 | 3894.00 | 3878.00 | -1.14 | 0.00 |
| 737.00 | 648.00 | 635.00 | -1.18 | 0.07 |
| 9.00 | 4.00 | 5.33 | -1.51 | 0.29 |
| 2215.00 | 2307.00 | 2329.00 | 1.06 | 0.11 |
| 1693.00 | 1990.00 | 1853.00 | 1.45 | 0.00 |
| 318.00 | 364.00 | 346.00 | -1.04 | 0.47 |
| 1052.00 | 1006.00 | 1046.67 | 1.19 | 0.00 |
| 798.00 | 857.00 | 888.67 | 1.07 | 0.18 |
| 464.00 | 480.00 | 458.67 | 1.14 | 0.03 |
| 922.00 | 1032.00 | 984.00 | 1.05 | 0.47 |
| 20.00 | 32.00 | 28.00 | -1.35 | 0.05 |
| 494.00 | 586.00 | 555.00 | 1.12 | 0.04 |
| 876.00 | 1042.00 | 958.00 | -1.00 | 0.93 |
| 169.00 | 157.00 | 167.67 | 1.04 | 0.60 |
| 9799.00 | 10579.00 | 10834.67 | 1.09 | 0.02 |
| 234.00 | 249.00 | 247.00 | 1.11 | 0.13 |
| 2700.00 | 2927.00 | 2832.67 | -1.11 | 0.00 |
| 72.00 | 107.00 | 104.33 | 1.00 | 1.00 |
| 893.00 | 876.00 | 883.33 | -1.31 | 0.00 |
| 1671.00 | 1697.00 | 1679.67 | -1.05 | 0.27 |
| 9.00 | 13.00 | 13.67 | 1.01 | 1.00 |
| 108.00 | 127.00 | 123.00 | 1.20 | 0.04 |
| 937.00 | 925.00 | 988.33 | 1.07 | 0.15 |
| 1128.00 | 1299.00 | 1246.67 | 1.09 | 0.03 |
| 1197.00 | 1125.00 | 1178.33 | -1.22 | 0.00 |
| 605.00 | 597.00 | 613.33 | -1.04 | 0.41 |
| 687.00 | 755.00 | 733.00 | -1.04 | 0.35 |
| 1310.00 | 1312.00 | 1366.67 | 1.12 | 0.01 |
| 1558.00 | 1474.00 | 1542.67 | -1.26 | 0.00 |
| 1159.00 | 1199.00 | 1188.67 | -1.13 | 0.01 |
| 1278.00 | 1550.00 | 1455.00 | 1.18 | 0.00 |
| 219.00 | 230.00 | 228.67 | -1.29 | 0.00 |
| 1111.00 | 1097.00 | 1148.00 | -1.09 | 0.05 |
| 380.00 | 458.00 | 430.67 | 1.09 | 0.15 |

| | | | | |
|---|---|---|---|---|
| 342.00 | 366.00 | 374.00 | 1.12 | 0.04 |
| 491.00 | 562.00 | 538.00 | -1.29 | 0.00 |
| 1169.00 | 1222.00 | 1210.33 | 1.16 | 0.00 |
| 2027.00 | 1939.00 | 2020.67 | -1.11 | 0.01 |
| 4.00 | 7.00 | 6.67 | -1.66 | 0.11 |
| 1620.00 | 1620.00 | 1704.67 | -1.12 | 0.01 |
| 4644.00 | 5059.00 | 4969.67 | 1.12 | 0.00 |
| 4.00 | 1.00 | 2.67 | -1.26 | 0.82 |
| 1817.00 | 1984.00 | 1935.33 | 1.08 | 0.04 |
| 2888.00 | 3289.00 | 3164.00 | 1.03 | 0.42 |
| 2159.00 | 2559.00 | 2380.00 | -1.22 | 0.00 |
| 1319.00 | 1328.00 | 1343.33 | -1.08 | 0.06 |
| 3333.00 | 3043.00 | 3182.67 | -1.18 | 0.00 |
| 44.00 | 33.00 | 42.00 | -2.45 | 0.00 |
| 841.00 | 966.00 | 1021.67 | 1.16 | 0.02 |
| 44.00 | 25.00 | 37.00 | -1.17 | 0.30 |
| 669.00 | 672.00 | 652.67 | -1.29 | 0.00 |
| 1090.00 | 949.00 | 994.00 | -1.25 | 0.00 |
| 1881.00 | 1914.00 | 1893.00 | 1.13 | 0.01 |
| 849.00 | 1019.00 | 922.00 | -1.07 | 0.25 |
| 2747.00 | 2575.00 | 2768.33 | -1.14 | 0.00 |
| 865.00 | 746.00 | 819.67 | -1.28 | 0.00 |
| 5163.00 | 5075.00 | 4953.33 | -1.00 | 0.96 |
| 3.00 | 6.00 | 6.33 | -1.01 | 1.00 |
| 16743.00 | 17581.00 | 17282.00 | -1.15 | 0.00 |
| 37.00 | 33.00 | 38.67 | -1.07 | 0.67 |
| 48.00 | 68.00 | 55.67 | -3.52 | 0.00 |
| 1148.00 | 1185.00 | 1198.67 | -1.18 | 0.00 |
| 5241.00 | 5402.00 | 5520.67 | 1.05 | 0.19 |
| 32.00 | 45.00 | 44.00 | -1.32 | 0.04 |
| 125.00 | 99.00 | 112.67 | 1.37 | 0.00 |
| 10749.00 | 10483.00 | 10513.00 | -1.07 | 0.12 |
| 2138.00 | 2224.00 | 2154.67 | -1.36 | 0.00 |
| 1460.00 | 1352.00 | 1437.00 | -1.25 | 0.00 |
| 984.00 | 1053.00 | 1007.33 | -1.42 | 0.00 |
| 4121.00 | 4147.00 | 4228.67 | 1.02 | 0.54 |
| 5406.00 | 5809.00 | 5873.33 | 1.06 | 0.11 |
| 14771.00 | 15728.00 | 15955.67 | 1.13 | 0.00 |
| 2325.00 | 2600.00 | 2587.67 | 1.05 | 0.21 |
| 1328.00 | 1313.00 | 1379.00 | 1.19 | 0.00 |
| 1078.00 | 1164.00 | 1264.00 | 1.26 | 0.00 |
| 10.00 | 16.00 | 15.00 | 1.63 | 0.07 |
| 3029.00 | 3256.00 | 3356.00 | 1.21 | 0.00 |
| 1257.00 | 1285.00 | 1362.33 | 1.13 | 0.01 |
| 3258.00 | 3381.00 | 3517.33 | 1.11 | 0.01 |
| 2405.00 | 2783.00 | 2804.00 | 1.25 | 0.00 |
| 1487.00 | 1661.00 | 1657.33 | 1.21 | 0.00 |
| 484.00 | 638.00 | 629.00 | 1.13 | 0.10 |

| | | | | |
|---|---|---|---|---|
| 1267.00 | 1279.00 | 1365.67 | 1.20 | 0.00 |
| 834.00 | 938.00 | 931.00 | 1.23 | 0.00 |
| 1736.00 | 1935.00 | 1970.33 | 1.21 | 0.00 |
| 2072.00 | 2252.00 | 2269.33 | 1.21 | 0.00 |
| 170.00 | 189.00 | 183.33 | 1.15 | 0.07 |
| 434.00 | 474.00 | 484.67 | 1.00 | 0.99 |
| 2169.00 | 2288.00 | 2321.00 | -1.06 | 0.21 |
| 204.00 | 248.00 | 234.33 | -1.06 | 0.45 |
| 2236.00 | 2555.00 | 2439.67 | -1.10 | 0.02 |
| 2030.00 | 1979.00 | 2062.33 | -1.24 | 0.00 |
| 1231.00 | 1356.00 | 1264.67 | -1.07 | 0.12 |
| 2477.00 | 2348.00 | 2474.67 | 1.05 | 0.28 |
| 1519.00 | 1498.00 | 1578.00 | 1.06 | 0.25 |
| 994.00 | 1087.00 | 1110.33 | 1.08 | 0.11 |
| 1667.00 | 1769.00 | 1772.00 | 1.05 | 0.29 |
| 702.00 | 860.00 | 803.67 | 1.05 | 0.35 |
| 199.00 | 203.00 | 197.67 | -1.09 | 0.20 |
| 1026.00 | 1111.00 | 1067.33 | 1.10 | 0.04 |
| 67.00 | 68.00 | 72.67 | -1.56 | 0.00 |
| 88.00 | 111.00 | 99.00 | 1.12 | 0.26 |
| 1171.00 | 1182.00 | 1179.67 | 1.02 | 0.67 |
| 2508.00 | 2707.00 | 2659.33 | 1.13 | 0.00 |
| 3.00 | 2.00 | 3.33 | 1.61 | 0.47 |
| 758.00 | 844.00 | 867.67 | 1.19 | 0.00 |
| 1406.00 | 1460.00 | 1514.00 | 1.06 | 0.17 |
| 105.00 | 103.00 | 105.67 | 1.38 | 0.00 |
| 1328.00 | 1437.00 | 1528.33 | 1.15 | 0.01 |
| 974.00 | 1136.00 | 1143.67 | 1.20 | 0.00 |
| 1276.00 | 1339.00 | 1353.33 | 1.06 | 0.18 |
| 326.00 | 382.00 | 352.00 | -1.23 | 0.00 |
| 413.00 | 395.00 | 415.67 | -1.35 | 0.00 |
| 19.00 | 27.00 | 28.67 | -1.25 | 0.16 |
| 47.00 | 48.00 | 42.00 | 1.15 | 0.35 |
| 53.00 | 67.00 | 53.67 | -1.70 | 0.00 |
| 1716.00 | 1682.00 | 1713.67 | -1.21 | 0.00 |
| 82.00 | 85.00 | 86.67 | 1.33 | 0.01 |
| 11.00 | 11.00 | 10.00 | -1.31 | 0.35 |
| 606.00 | 592.00 | 620.67 | -1.14 | 0.00 |
| 248.00 | 260.00 | 260.67 | -1.10 | 0.11 |
| 692.00 | 747.00 | 764.00 | 1.17 | 0.00 |
| 228.00 | 220.00 | 235.33 | 1.16 | 0.04 |
| 953.00 | 1080.00 | 1088.67 | 1.10 | 0.04 |
| 1801.00 | 2029.00 | 1936.00 | -1.06 | 0.13 |
| 2877.00 | 2886.00 | 2969.33 | 1.05 | 0.23 |
| 349.00 | 372.00 | 363.00 | -1.06 | 0.26 |
| 2554.00 | 2569.00 | 2533.00 | -1.16 | 0.00 |
| 51.00 | 37.00 | 41.33 | 1.07 | 0.68 |
| 2142.00 | 2385.00 | 2162.67 | -1.13 | 0.04 |

| | | | | |
|---|---|---|---|---|
| 507.00 | 479.00 | 500.67 | -1.59 | 0.00 |
| 1557.00 | 1713.00 | 1702.33 | 1.08 | 0.06 |
| 869.00 | 858.00 | 852.33 | -1.19 | 0.00 |
| 1396.00 | 1405.00 | 1384.67 | -1.02 | 0.68 |
| 7569.00 | 7219.00 | 7546.33 | -1.14 | 0.00 |
| 351.00 | 404.00 | 378.67 | 1.05 | 0.39 |
| 742.00 | 760.00 | 745.33 | -1.01 | 0.83 |
| 463.00 | 479.00 | 469.67 | -1.13 | 0.03 |
| 379.00 | 334.00 | 375.67 | -1.14 | 0.03 |
| 1326.00 | 1277.00 | 1339.00 | -1.14 | 0.00 |
| 2408.00 | 2519.00 | 2585.67 | 1.02 | 0.55 |
| 136.00 | 126.00 | 127.00 | 1.02 | 0.83 |
| 132.00 | 153.00 | 142.67 | 1.15 | 0.08 |
| 5055.00 | 5190.00 | 5350.33 | 1.03 | 0.35 |
| 1745.00 | 2033.00 | 2003.00 | 1.33 | 0.00 |
| 1325.00 | 1337.00 | 1415.00 | 1.15 | 0.00 |
| 38.00 | 33.00 | 33.00 | -1.12 | 0.53 |
| 1364.00 | 1490.00 | 1530.33 | 1.30 | 0.00 |
| 19.00 | 10.00 | 15.67 | -1.12 | 0.74 |
| 45.00 | 40.00 | 41.33 | 1.17 | 0.33 |
| 348.00 | 352.00 | 366.67 | -1.05 | 0.38 |
| 69.00 | 71.00 | 75.33 | -1.35 | 0.01 |
| 2779.00 | 2797.00 | 2789.33 | 1.01 | 0.79 |
| 113.00 | 107.00 | 110.33 | -1.02 | 0.84 |
| 2.00 | 1.00 | 2.67 | -10.69 | 0.00 |
| 3.00 | 1.00 | 3.00 | -1.34 | 0.67 |
| 1056.00 | 1152.00 | 1122.33 | 1.05 | 0.20 |
| 10.00 | 16.00 | 16.00 | 1.12 | 0.69 |
| 18027.00 | 16822.00 | 17305.00 | -1.58 | 0.00 |
| 1.00 | 3.00 | 2.33 | -1.84 | 0.27 |
| 866.00 | 943.00 | 966.67 | 1.08 | 0.13 |
| 4776.00 | 5160.00 | 4973.33 | 1.17 | 0.00 |
| 17.00 | 20.00 | 22.00 | 1.25 | 0.29 |
| 97.00 | 103.00 | 100.33 | -1.01 | 0.95 |
| 1529.00 | 1573.00 | 1590.00 | 1.27 | 0.00 |
| 264.00 | 284.00 | 271.67 | 1.17 | 0.03 |
| 1310.00 | 1344.00 | 1360.67 | 1.15 | 0.00 |
| 58.00 | 61.00 | 56.67 | 1.11 | 0.42 |
| 60.00 | 56.00 | 53.00 | -1.25 | 0.07 |
| 12.00 | 14.00 | 16.00 | -1.22 | 0.35 |
| 1106.00 | 1173.00 | 1138.33 | 1.06 | 0.15 |
| 5.00 | 7.00 | 5.67 | 1.51 | 0.35 |
| 1527.00 | 1444.00 | 1568.00 | -1.04 | 0.43 |
| 382.00 | 423.00 | 388.33 | 1.15 | 0.05 |
| 502.00 | 533.00 | 499.33 | -1.00 | 0.99 |
| 493.00 | 505.00 | 500.67 | 1.10 | 0.05 |
| 0.00 | 0.00 | 0.33 | -3.96 | 0.23 |
| 43.00 | 30.00 | 39.33 | -1.65 | 0.00 |

| | | | | |
|---|---|---|---|---|
| 208.00 | 208.00 | 210.33 | 1.41 | 0.00 |
| 49.00 | 41.00 | 47.00 | 1.00 | 1.00 |
| 4.00 | 6.00 | 6.00 | -2.45 | 0.00 |
| 1802.00 | 1594.00 | 1718.00 | -1.07 | 0.19 |
| 1.00 | 7.00 | 6.00 | -1.46 | 0.32 |
| 1281.00 | 1310.00 | 1372.67 | 1.04 | 0.39 |
| 1489.00 | 1346.00 | 1456.00 | -1.09 | 0.05 |
| 361.00 | 405.00 | 411.67 | -1.04 | 0.45 |
| 2200.00 | 1974.00 | 2063.33 | 1.05 | 0.38 |
| 746.00 | 904.00 | 785.67 | 1.36 | 0.00 |
| 820.00 | 771.00 | 821.00 | -1.10 | 0.03 |
| 1723.00 | 1884.00 | 1853.67 | -1.22 | 0.00 |
| 290.00 | 328.00 | 337.00 | 1.05 | 0.47 |
| 72.00 | 69.00 | 72.33 | 1.49 | 0.00 |
| 7.00 | 7.00 | 5.67 | 1.05 | 1.00 |
| 1.00 | 2.00 | 1.67 | -1.58 | 0.59 |
| 1648.00 | 1619.00 | 1673.33 | 1.10 | 0.02 |
| 1135.00 | 1179.00 | 1147.00 | -1.31 | 0.00 |
| 78.00 | 71.00 | 82.67 | -1.24 | 0.03 |
| 3438.00 | 3376.00 | 3468.67 | 1.13 | 0.00 |
| 1957.00 | 2284.00 | 2077.00 | -1.08 | 0.16 |
| 2.00 | 3.00 | 2.67 | -2.46 | 0.04 |
| 89.00 | 102.00 | 95.33 | -1.21 | 0.04 |
| 570.00 | 563.00 | 627.00 | 1.04 | 0.54 |
| 458.00 | 455.00 | 461.33 | 1.13 | 0.02 |
| 730.00 | 868.00 | 821.00 | 1.26 | 0.00 |
| 1127.00 | 1159.00 | 1175.00 | -1.14 | 0.00 |
| 371.00 | 429.00 | 395.00 | 1.14 | 0.03 |
| 1021.00 | 1070.00 | 1044.33 | 1.05 | 0.27 |
| 6.00 | 3.00 | 4.33 | -1.16 | 0.85 |
| 164.00 | 125.00 | 151.33 | -1.22 | 0.03 |
| 810.00 | 918.00 | 893.67 | 1.03 | 0.54 |
| 706.00 | 645.00 | 705.67 | -1.04 | 0.44 |
| 240.00 | 214.00 | 240.33 | -1.16 | 0.04 |
| 143.00 | 108.00 | 133.67 | 1.06 | 0.55 |
| 175.00 | 186.00 | 192.67 | 1.45 | 0.00 |
| 6.00 | 8.00 | 5.00 | -1.54 | 0.28 |
| 445.00 | 419.00 | 456.00 | 1.09 | 0.12 |
| 48.00 | 50.00 | 58.33 | -1.47 | 0.00 |
| 2877.00 | 3019.00 | 3160.33 | 1.15 | 0.00 |
| 573.00 | 571.00 | 589.67 | 1.02 | 0.75 |
| 1590.00 | 1530.00 | 1614.67 | -1.11 | 0.01 |
| 241.00 | 278.00 | 260.33 | 1.17 | 0.02 |
| 37.00 | 19.00 | 27.33 | 1.42 | 0.06 |
| 4998.00 | 5107.00 | 5193.33 | 1.01 | 0.75 |
| 21.00 | 22.00 | 20.00 | -1.74 | 0.00 |
| 66.00 | 79.00 | 76.67 | 1.16 | 0.20 |
| 21.00 | 17.00 | 20.00 | -1.08 | 0.80 |

| | | | | |
|---|---|---|---|---|
| 1088.00 | 1123.00 | 1189.00 | 1.03 | 0.49 |
| 1766.00 | 1994.00 | 1927.33 | -1.02 | 0.62 |
| 3402.00 | 3300.00 | 3346.00 | -1.04 | 0.33 |
| 2470.00 | 2218.00 | 2304.00 | -1.26 | 0.00 |
| 612.00 | 541.00 | 555.67 | -1.23 | 0.00 |
| 130.00 | 124.00 | 127.00 | -1.09 | 0.30 |
| 361.00 | 446.00 | 436.67 | 1.13 | 0.07 |
| 849.00 | 840.00 | 863.67 | -1.12 | 0.01 |
| 741.00 | 825.00 | 793.67 | -1.09 | 0.09 |
| 77.00 | 109.00 | 91.67 | -1.05 | 0.71 |
| 1178.00 | 1258.00 | 1240.33 | -1.17 | 0.00 |
| 329.00 | 378.00 | 360.67 | 1.09 | 0.11 |
| 345.00 | 408.00 | 378.67 | 1.09 | 0.12 |
| 465.00 | 555.00 | 494.67 | -1.03 | 0.68 |
| 272.00 | 339.00 | 299.00 | 1.06 | 0.41 |
| 2376.00 | 2308.00 | 2299.00 | -1.04 | 0.37 |
| 1268.00 | 1196.00 | 1281.00 | -1.20 | 0.00 |
| 1697.00 | 1955.00 | 1817.33 | 1.10 | 0.05 |
| 6686.00 | 6644.00 | 6851.67 | -1.03 | 0.41 |
| 4557.00 | 4767.00 | 4824.67 | 1.27 | 0.00 |
| 3969.00 | 4111.00 | 4111.00 | 1.02 | 0.64 |
| 2.00 | 4.00 | 4.33 | -1.70 | 0.18 |
| 673.00 | 730.00 | 753.33 | 1.14 | 0.01 |
| 337.00 | 342.00 | 357.33 | -1.18 | 0.00 |
| 311.00 | 325.00 | 320.67 | 1.02 | 0.77 |
| 6394.00 | 7049.00 | 6741.33 | 1.02 | 0.59 |
| 698.00 | 689.00 | 745.67 | 1.04 | 0.51 |
| 2077.00 | 2328.00 | 2334.67 | 1.03 | 0.43 |
| 179.00 | 136.00 | 173.67 | -1.07 | 0.52 |
| 30.00 | 22.00 | 23.67 | -1.08 | 0.70 |
| 2057.00 | 2291.00 | 2199.67 | 1.13 | 0.00 |
| 298.00 | 280.00 | 307.67 | -1.03 | 0.62 |
| 480.00 | 434.00 | 436.33 | 1.16 | 0.04 |
| 673.00 | 750.00 | 742.00 | -1.06 | 0.22 |
| 712.00 | 788.00 | 754.33 | 1.15 | 0.00 |
| 14.00 | 21.00 | 14.33 | 1.91 | 0.02 |
| 439.00 | 496.00 | 462.67 | 1.48 | 0.00 |
| 1402.00 | 1767.00 | 1677.33 | 1.85 | 0.00 |
| 1229.00 | 1258.00 | 1305.67 | -1.11 | 0.01 |
| 157.00 | 169.00 | 167.00 | 1.25 | 0.00 |
| 572.00 | 552.00 | 618.33 | 1.23 | 0.00 |
| 1359.00 | 1387.00 | 1411.33 | 1.05 | 0.25 |
| 16.00 | 17.00 | 14.67 | 1.40 | 0.18 |
| 1274.00 | 1447.00 | 1412.00 | -1.02 | 0.64 |
| 1.00 | 13.00 | 7.00 | 2.23 | 0.13 |
| 123.00 | 134.00 | 136.33 | -1.05 | 0.58 |
| 1016.00 | 1086.00 | 1089.00 | -1.05 | 0.31 |
| 1267.00 | 1289.00 | 1308.33 | 1.55 | 0.00 |

| | | | | |
|---|---|---|---|---|
| 681.00 | 723.00 | 754.00 | -1.02 | 0.66 |
| 654.00 | 684.00 | 690.67 | 1.28 | 0.00 |
| 538.00 | 511.00 | 523.00 | -1.02 | 0.74 |
| 1943.00 | 1809.00 | 1839.00 | -1.39 | 0.00 |
| 3699.00 | 4376.00 | 4051.00 | 1.08 | 0.06 |
| 22.00 | 20.00 | 18.67 | -1.70 | 0.00 |
| 449.00 | 457.00 | 450.67 | -1.25 | 0.00 |
| 237.00 | 226.00 | 229.00 | -1.28 | 0.00 |
| 690.00 | 734.00 | 732.00 | -1.06 | 0.23 |
| 7.00 | 3.00 | 4.67 | -1.22 | 0.73 |
| 1338.00 | 1327.00 | 1425.33 | 1.01 | 0.89 |
| 26.00 | 19.00 | 39.00 | 2.05 | 0.03 |
| 26.00 | 8.00 | 16.00 | -1.40 | 0.18 |
| 204.00 | 230.00 | 231.33 | -1.01 | 0.88 |
| 11.00 | 8.00 | 10.33 | -1.40 | 0.21 |
| 4.00 | 4.00 | 3.67 | -2.08 | 0.07 |
| 1706.00 | 1664.00 | 1748.00 | 1.43 | 0.00 |
| 27.00 | 25.00 | 25.67 | -1.51 | 0.01 |
| 32.00 | 12.00 | 22.00 | 1.05 | 0.82 |
| 1233.00 | 1480.00 | 1399.33 | 1.12 | 0.02 |
| 280.00 | 329.00 | 328.00 | 1.03 | 0.62 |
| 653.00 | 747.00 | 693.00 | -1.11 | 0.04 |
| 57.00 | 66.00 | 72.67 | -1.21 | 0.13 |
| 227.00 | 219.00 | 223.00 | -1.34 | 0.00 |
| 133.00 | 151.00 | 146.00 | -1.31 | 0.00 |
| 118.00 | 125.00 | 120.67 | 1.59 | 0.00 |
| 11.00 | 11.00 | 10.00 | 1.14 | 0.70 |
| 695.00 | 795.00 | 760.00 | 1.13 | 0.01 |
| 1294.00 | 1249.00 | 1310.33 | -1.10 | 0.02 |
| 41.00 | 44.00 | 43.00 | -1.17 | 0.25 |
| 82.00 | 85.00 | 94.33 | 1.19 | 0.10 |
| 667.00 | 718.00 | 697.33 | 1.06 | 0.24 |
| 6.00 | 11.00 | 6.67 | -1.11 | 0.88 |
| 30.00 | 56.00 | 40.33 | 1.05 | 0.79 |
| 8.00 | 5.00 | 5.33 | -1.88 | 0.06 |
| 1563.00 | 1532.00 | 1599.33 | -1.54 | 0.00 |
| 10.00 | 11.00 | 9.33 | 1.25 | 0.49 |
| 65.00 | 59.00 | 63.33 | -1.28 | 0.02 |
| 401.00 | 409.00 | 416.00 | 1.04 | 0.45 |
| 3.00 | 1.00 | 4.00 | -1.67 | 0.23 |
| 547.00 | 545.00 | 545.00 | -1.13 | 0.02 |
| 224.00 | 208.00 | 235.33 | 1.33 | 0.00 |
| 415.00 | 383.00 | 391.00 | -1.01 | 0.87 |
| 996.00 | 1011.00 | 1013.67 | -1.09 | 0.06 |
| 448.00 | 493.00 | 506.33 | 1.17 | 0.00 |
| 253.00 | 264.00 | 253.67 | -1.01 | 0.97 |
| 336.00 | 332.00 | 350.00 | -1.04 | 0.52 |
| 0.00 | 5.00 | 2.00 | -1.65 | 0.46 |

| | | | | |
|---|---|---|---|---|
| 8805.00 | 10488.00 | 9985.00 | 1.22 | 0.00 |
| 1975.00 | 2276.00 | 2161.67 | 3.38 | 0.00 |
| 954.00 | 1015.00 | 1040.67 | 1.34 | 0.00 |
| 2186.00 | 2127.00 | 2123.00 | -1.10 | 0.03 |
| 20.00 | 10.00 | 15.67 | 1.29 | 0.32 |
| 930.00 | 1204.00 | 1103.00 | 1.21 | 0.00 |
| 1336.00 | 1277.00 | 1355.33 | 1.06 | 0.23 |
| 5.00 | 7.00 | 6.33 | -1.28 | 0.55 |
| 1944.00 | 1981.00 | 2018.00 | 1.06 | 0.13 |
| 379.00 | 491.00 | 445.00 | 1.19 | 0.01 |
| 507.00 | 628.00 | 601.00 | -1.08 | 0.15 |
| 179.00 | 205.00 | 205.00 | -1.07 | 0.29 |
| 593.00 | 665.00 | 662.67 | 1.16 | 0.00 |
| 313.00 | 275.00 | 314.33 | 1.12 | 0.13 |
| 188.00 | 209.00 | 204.33 | -1.07 | 0.34 |
| 1209.00 | 1487.00 | 1379.67 | 1.15 | 0.00 |
| 419.00 | 503.00 | 484.67 | 1.27 | 0.00 |
| 336.00 | 312.00 | 345.33 | -1.05 | 0.48 |
| 518.00 | 522.00 | 540.67 | 1.00 | 0.93 |
| 23.00 | 27.00 | 27.00 | -2.12 | 0.00 |
| 73.00 | 66.00 | 73.00 | -1.04 | 0.77 |
| 534.00 | 521.00 | 524.00 | -1.36 | 0.00 |
| 518.00 | 525.00 | 525.67 | 1.05 | 0.35 |
| 203.00 | 211.00 | 204.67 | -1.18 | 0.01 |
| 10.00 | 7.00 | 9.67 | -1.08 | 0.90 |
| 65.00 | 88.00 | 77.00 | -1.09 | 0.45 |
| 593.00 | 596.00 | 606.67 | 2.44 | 0.00 |
| 303.00 | 304.00 | 303.67 | -1.28 | 0.00 |
| 643.00 | 640.00 | 613.67 | 1.12 | 0.08 |
| 86.00 | 93.00 | 84.67 | -1.34 | 0.00 |
| 1240.00 | 1329.00 | 1284.67 | -1.15 | 0.00 |
| 2615.00 | 2927.00 | 2786.33 | -1.04 | 0.31 |
| 431.00 | 410.00 | 424.33 | -1.05 | 0.39 |
| 19.00 | 3.00 | 10.33 | -1.85 | 0.02 |
| 233.00 | 236.00 | 220.00 | -1.33 | 0.00 |
| 180.00 | 156.00 | 166.00 | -1.35 | 0.00 |
| 85.00 | 95.00 | 82.67 | -1.19 | 0.11 |
| 3253.00 | 3488.00 | 3415.33 | 1.05 | 0.15 |
| 305.00 | 319.00 | 310.67 | -1.73 | 0.00 |
| 394.00 | 425.00 | 421.00 | 1.25 | 0.00 |
| 1336.00 | 1533.00 | 1469.00 | 1.17 | 0.00 |
| 1882.00 | 2169.00 | 2149.67 | 1.21 | 0.00 |
| 859.00 | 1009.00 | 910.33 | -1.05 | 0.38 |
| 24.00 | 41.00 | 33.00 | -1.26 | 0.12 |
| 265.00 | 251.00 | 270.33 | -1.46 | 0.00 |
| 1560.00 | 1776.00 | 1756.00 | 1.14 | 0.00 |
| 1266.00 | 1303.00 | 1258.00 | -1.13 | 0.01 |
| 79.00 | 95.00 | 90.00 | -1.24 | 0.02 |

| | | | | |
|---|---|---|---|---|
| 251.00 | 239.00 | 260.33 | 1.05 | 0.48 |
| 353.00 | 405.00 | 408.67 | 1.17 | 0.01 |
| 90.00 | 72.00 | 79.33 | -1.02 | 0.91 |
| 3708.00 | 3967.00 | 3933.33 | 1.16 | 0.00 |
| 379.00 | 404.00 | 386.33 | 1.28 | 0.00 |
| 187.00 | 244.00 | 225.67 | 1.21 | 0.01 |
| 260.00 | 289.00 | 305.67 | -1.09 | 0.23 |
| 521.00 | 576.00 | 564.00 | 1.17 | 0.00 |
| 2126.00 | 1934.00 | 2001.33 | -1.18 | 0.00 |
| 278.00 | 279.00 | 265.00 | 1.07 | 0.36 |
| 541.00 | 591.00 | 571.67 | 1.08 | 0.12 |
| 1778.00 | 1836.00 | 1883.33 | -1.02 | 0.53 |
| 982.00 | 1046.00 | 975.33 | 1.03 | 0.56 |
| 378.00 | 473.00 | 424.00 | 1.07 | 0.23 |
| 14.00 | 21.00 | 16.33 | 1.10 | 0.69 |
| 475.00 | 553.00 | 534.67 | 1.14 | 0.02 |
| 917.00 | 913.00 | 930.33 | -1.08 | 0.08 |
| 10.00 | 13.00 | 11.67 | -1.39 | 0.20 |
| 862.00 | 898.00 | 864.00 | -1.13 | 0.01 |
| 1880.00 | 1880.00 | 1918.00 | -1.07 | 0.08 |
| 1415.00 | 1502.00 | 1492.00 | 1.17 | 0.00 |
| 1944.00 | 2326.00 | 2196.33 | -1.02 | 0.67 |
| 1227.00 | 1274.00 | 1285.00 | -1.03 | 0.49 |
| 20.00 | 19.00 | 17.33 | -1.07 | 0.85 |
| 1551.00 | 1675.00 | 1692.33 | -1.16 | 0.00 |
| 20.00 | 21.00 | 19.67 | -1.47 | 0.03 |
| 758.00 | 865.00 | 820.00 | 1.08 | 0.12 |
| 392.00 | 368.00 | 390.33 | 1.16 | 0.01 |
| 1075.00 | 1108.00 | 1131.67 | -1.09 | 0.04 |
| 88.00 | 96.00 | 89.67 | 1.59 | 0.00 |
| 140.00 | 178.00 | 176.33 | -1.15 | 0.10 |
| 339.00 | 409.00 | 364.00 | 1.40 | 0.00 |
| 1339.00 | 1542.00 | 1534.33 | 1.28 | 0.00 |
| 1335.00 | 1480.00 | 1496.33 | -1.02 | 0.64 |
| 1246.00 | 1344.00 | 1354.00 | 1.09 | 0.05 |
| 294.00 | 329.00 | 303.67 | 1.21 | 0.01 |
| 1196.00 | 1431.00 | 1330.67 | -1.04 | 0.42 |
| 669.00 | 810.00 | 771.33 | -1.11 | 0.05 |
| 1697.00 | 1574.00 | 1635.00 | -1.17 | 0.00 |
| 145.00 | 158.00 | 154.33 | -1.25 | 0.00 |
| 37.00 | 28.00 | 32.33 | 1.02 | 0.95 |
| 1070.00 | 1258.00 | 1184.00 | -1.19 | 0.00 |
| 5.00 | 11.00 | 7.33 | 1.20 | 0.64 |
| 754.00 | 879.00 | 816.00 | -1.10 | 0.07 |
| 154.00 | 123.00 | 141.33 | -1.25 | 0.01 |
| 1511.00 | 1363.00 | 1438.00 | -1.27 | 0.00 |
| 22016.00 | 21932.00 | 22876.33 | -1.09 | 0.05 |
| 40.00 | 28.00 | 28.67 | -1.08 | 0.76 |

| | | | | |
|---|---|---|---|---|
| 3.00 | 3.00 | 3.67 | -1.19 | 0.84 |
| 283.00 | 279.00 | 298.00 | -1.27 | 0.00 |
| 2344.00 | 2746.00 | 2583.00 | -1.35 | 0.00 |
| 1577.00 | 1819.00 | 1695.00 | 1.15 | 0.00 |
| 773.00 | 920.00 | 864.67 | 1.10 | 0.08 |
| 219.00 | 194.00 | 222.67 | -1.70 | 0.00 |
| 426.00 | 409.00 | 407.33 | -1.36 | 0.00 |
| 2030.00 | 2112.00 | 2150.33 | 1.04 | 0.24 |
| 1077.00 | 1102.00 | 1115.33 | 1.14 | 0.01 |
| 631.00 | 664.00 | 672.00 | 1.23 | 0.00 |
| 24.00 | 22.00 | 24.00 | -1.45 | 0.02 |
| 335.00 | 327.00 | 328.00 | -1.12 | 0.05 |
| 13.00 | 6.00 | 10.00 | -1.15 | 0.72 |
| 477.00 | 550.00 | 506.33 | -1.20 | 0.00 |
| 9746.00 | 10872.00 | 10920.00 | 1.19 | 0.00 |
| 1332.00 | 1531.00 | 1505.33 | 1.22 | 0.00 |
| 1771.00 | 1910.00 | 1950.67 | 1.16 | 0.00 |
| 542.00 | 640.00 | 606.67 | 1.37 | 0.00 |
| 1183.00 | 1242.00 | 1228.33 | -1.05 | 0.26 |
| 454.00 | 473.00 | 459.67 | 1.00 | 0.94 |
| 174.00 | 196.00 | 194.67 | -1.27 | 0.00 |
| 658.00 | 612.00 | 653.33 | 1.02 | 0.70 |
| 590.00 | 573.00 | 581.33 | -1.03 | 0.58 |
| 229.00 | 273.00 | 252.33 | -1.21 | 0.00 |
| 14.00 | 14.00 | 18.33 | 4.06 | 0.00 |
| 845.00 | 870.00 | 896.00 | -1.01 | 0.80 |
| 1554.00 | 1611.00 | 1597.33 | 1.23 | 0.00 |
| 1332.00 | 1458.00 | 1383.33 | 1.20 | 0.00 |
| 5191.00 | 6009.00 | 5756.33 | 1.00 | 0.96 |
| 336.00 | 370.00 | 367.67 | 1.01 | 0.83 |
| 1289.00 | 1537.00 | 1506.33 | 1.14 | 0.00 |
| 237.00 | 258.00 | 245.00 | 1.19 | 0.02 |
| 3.00 | 6.00 | 3.67 | 1.08 | 1.00 |
| 982.00 | 1017.00 | 1052.67 | 1.03 | 0.47 |
| 159.00 | 125.00 | 151.33 | -1.16 | 0.15 |
| 73.00 | 62.00 | 71.33 | 1.22 | 0.10 |
| 4481.00 | 4706.00 | 4689.00 | 1.28 | 0.00 |
| 2607.00 | 2860.00 | 2865.67 | 1.27 | 0.00 |
| 15.00 | 14.00 | 15.67 | 1.84 | 0.02 |
| 77.00 | 67.00 | 78.67 | 1.06 | 0.64 |
| 19.00 | 34.00 | 28.33 | -1.63 | 0.00 |
| 94.00 | 88.00 | 83.00 | -1.27 | 0.03 |
| 1302.00 | 1341.00 | 1363.00 | 1.03 | 0.56 |
| 95.00 | 77.00 | 88.33 | 1.16 | 0.16 |
| 225.00 | 297.00 | 268.67 | 1.06 | 0.42 |
| 1.00 | 0.00 | 1.00 | -2.51 | 0.23 |
| 19.00 | 21.00 | 24.33 | -1.08 | 0.70 |
| 3.00 | 6.00 | 6.00 | -1.01 | 1.00 |

| | | | | |
|---|---|---|---|---|
| 616.00 | 597.00 | 632.67 | -1.18 | 0.00 |
| 530.00 | 643.00 | 609.33 | 1.03 | 0.53 |
| 36.00 | 43.00 | 47.00 | 1.03 | 0.91 |
| 237.00 | 221.00 | 239.67 | 1.10 | 0.17 |
| 764.00 | 811.00 | 821.00 | 1.02 | 0.74 |
| 654.00 | 668.00 | 667.67 | -1.12 | 0.02 |
| 1066.00 | 1174.00 | 1173.67 | 1.09 | 0.04 |
| 57.00 | 46.00 | 60.00 | -1.18 | 0.16 |
| 393.00 | 428.00 | 438.33 | 1.02 | 0.76 |
| 461.00 | 456.00 | 469.00 | -1.07 | 0.35 |
| 760.00 | 762.00 | 770.33 | 1.08 | 0.10 |
| 2608.00 | 3030.00 | 2884.33 | 1.19 | 0.00 |
| 4.00 | 6.00 | 3.67 | 1.34 | 0.65 |
| 6.00 | 9.00 | 7.00 | 1.15 | 0.75 |
| 1883.00 | 1998.00 | 2032.67 | 1.10 | 0.01 |
| 431.00 | 497.00 | 480.67 | 1.33 | 0.00 |
| 83.00 | 118.00 | 102.00 | 1.12 | 0.26 |
| 444.00 | 513.00 | 509.00 | 1.03 | 0.57 |
| 278.00 | 282.00 | 295.67 | 1.19 | 0.01 |
| 435.00 | 538.00 | 536.67 | 1.03 | 0.65 |
| 1356.00 | 1337.00 | 1378.33 | 1.02 | 0.57 |
| 109.00 | 130.00 | 129.33 | 1.03 | 0.75 |
| 3.00 | 5.00 | 4.33 | -1.77 | 0.14 |
| 1357.00 | 1301.00 | 1363.67 | -1.18 | 0.00 |
| 44.00 | 54.00 | 53.33 | -1.05 | 0.66 |
| 1386.00 | 1437.00 | 1478.33 | 1.23 | 0.00 |
| 538.00 | 706.00 | 621.00 | 1.04 | 0.57 |
| 536.00 | 701.00 | 645.33 | 1.12 | 0.05 |
| 255.00 | 247.00 | 269.33 | 1.00 | 1.00 |
| 198.00 | 214.00 | 208.33 | 1.24 | 0.00 |
| 863.00 | 944.00 | 907.67 | 1.04 | 0.43 |
| 67.00 | 76.00 | 72.33 | 1.21 | 0.12 |
| 987.00 | 1062.00 | 1070.00 | -1.03 | 0.47 |
| 47.00 | 44.00 | 50.00 | -1.07 | 0.60 |
| 291.00 | 350.00 | 335.67 | 1.03 | 0.67 |
| 222.00 | 252.00 | 236.67 | -1.10 | 0.20 |
| 230.00 | 262.00 | 248.00 | 1.03 | 0.68 |
| 785.00 | 885.00 | 855.33 | 1.25 | 0.00 |
| 181.00 | 215.00 | 222.67 | 1.06 | 0.52 |
| 401.00 | 387.00 | 411.00 | -1.04 | 0.46 |
| 971.00 | 1104.00 | 1081.00 | 1.08 | 0.11 |
| 55.00 | 60.00 | 60.33 | -1.24 | 0.06 |
| 453.00 | 450.00 | 481.67 | -1.05 | 0.34 |
| 2046.00 | 2432.00 | 2372.67 | 1.16 | 0.00 |
| 428.00 | 405.00 | 437.67 | 1.04 | 0.51 |
| 920.00 | 998.00 | 1039.33 | 1.20 | 0.00 |
| 50.00 | 60.00 | 53.67 | 1.14 | 0.31 |
| 11.00 | 21.00 | 15.33 | 1.01 | 1.00 |

| | | | | |
|---|---|---|---|---|
| 4.00 | 4.00 | 3.00 | -1.12 | 1.00 |
| 387.00 | 414.00 | 408.00 | -1.04 | 0.47 |
| 330.00 | 314.00 | 332.67 | 1.22 | 0.01 |
| 256.00 | 314.00 | 307.33 | -1.06 | 0.42 |
| 10.00 | 13.00 | 12.33 | -1.37 | 0.22 |
| 283.00 | 267.00 | 290.00 | -1.12 | 0.13 |
| 294.00 | 293.00 | 315.33 | 1.04 | 0.53 |
| 457.00 | 521.00 | 496.00 | 1.13 | 0.01 |
| 10.00 | 4.00 | 6.67 | -1.21 | 0.66 |
| 4913.00 | 4995.00 | 5049.67 | -1.16 | 0.00 |
| 3.00 | 6.00 | 5.67 | 2.32 | 0.07 |
| 5.00 | 1.00 | 3.00 | -1.98 | 0.13 |
| 224.00 | 212.00 | 210.00 | -1.01 | 0.88 |
| 158.00 | 182.00 | 180.33 | 1.47 | 0.00 |
| 6.00 | 14.00 | 13.00 | 1.37 | 0.30 |
| 888.00 | 925.00 | 967.67 | 1.16 | 0.00 |
| 87.00 | 92.00 | 89.00 | 1.09 | 0.42 |
| 1741.00 | 2149.00 | 1950.00 | 1.09 | 0.07 |
| 3232.00 | 2914.00 | 3109.67 | -1.10 | 0.03 |
| 365.00 | 390.00 | 397.67 | 1.17 | 0.02 |
| 920.00 | 905.00 | 931.67 | -1.25 | 0.00 |
| 449.00 | 482.00 | 480.33 | -1.02 | 0.64 |
| 1000.00 | 1090.00 | 1062.33 | 1.10 | 0.07 |
| 260.00 | 242.00 | 265.67 | -1.59 | 0.00 |
| 26.00 | 34.00 | 29.00 | 1.04 | 0.90 |
| 102.00 | 104.00 | 107.67 | 1.55 | 0.00 |
| 271.00 | 251.00 | 283.67 | 1.02 | 0.75 |
| 60.00 | 68.00 | 70.33 | -1.07 | 0.57 |
| 107.00 | 75.00 | 98.67 | -1.01 | 0.92 |
| 71.00 | 87.00 | 84.33 | -1.11 | 0.28 |
| 12.00 | 24.00 | 16.67 | 2.13 | 0.00 |
| 22.00 | 17.00 | 20.67 | -1.29 | 0.21 |
| 2117.00 | 2208.00 | 2171.67 | -1.36 | 0.00 |
| 283.00 | 296.00 | 310.00 | -1.05 | 0.43 |
| 335.00 | 408.00 | 399.33 | 1.31 | 0.00 |
| 197.00 | 178.00 | 186.00 | -1.07 | 0.38 |
| 1601.00 | 1555.00 | 1630.67 | 1.05 | 0.19 |
| 83.00 | 71.00 | 71.00 | 1.21 | 0.12 |
| 576.00 | 608.00 | 620.00 | 1.19 | 0.00 |
| 776.00 | 824.00 | 830.67 | -1.03 | 0.48 |
| 2086.00 | 2189.00 | 2140.67 | -1.03 | 0.39 |
| 463.00 | 516.00 | 516.33 | 1.22 | 0.00 |
| 25.00 | 31.00 | 27.67 | 1.12 | 0.59 |
| 230.00 | 244.00 | 262.33 | 1.11 | 0.17 |
| 569.00 | 574.00 | 605.00 | 1.15 | 0.00 |
| 11.00 | 7.00 | 9.33 | -2.48 | 0.00 |
| 406.00 | 517.00 | 477.67 | 1.11 | 0.08 |
| 30.00 | 34.00 | 35.67 | 1.02 | 0.95 |

| | | | | |
|---|---|---|---|---|
| 559.00 | 569.00 | 575.67 | -1.02 | 0.72 |
| 485.00 | 537.00 | 540.33 | 1.16 | 0.00 |
| 553.00 | 508.00 | 566.00 | 1.19 | 0.00 |
| 360.00 | 420.00 | 408.67 | 1.06 | 0.33 |
| 8.00 | 11.00 | 9.67 | 1.50 | 0.20 |
| 1808.00 | 1833.00 | 1878.00 | 1.12 | 0.00 |
| 317.00 | 356.00 | 349.67 | -1.08 | 0.19 |
| 192.00 | 164.00 | 179.33 | 1.19 | 0.03 |
| 2305.00 | 2319.00 | 2435.33 | 1.14 | 0.00 |
| 255.00 | 259.00 | 264.33 | 1.09 | 0.17 |
| 918.00 | 926.00 | 1010.67 | 1.13 | 0.02 |
| 597.00 | 634.00 | 629.00 | -1.12 | 0.01 |
| 52.00 | 30.00 | 38.67 | -1.12 | 0.51 |
| 575.00 | 576.00 | 578.67 | 1.27 | 0.00 |
| 421.00 | 434.00 | 453.67 | -1.19 | 0.00 |
| 382.00 | 407.00 | 391.67 | 1.05 | 0.56 |
| 485.00 | 475.00 | 508.67 | 1.05 | 0.36 |
| 23.00 | 19.00 | 20.00 | -1.18 | 0.41 |
| 10.00 | 9.00 | 11.67 | -1.45 | 0.13 |
| 274.00 | 327.00 | 307.67 | 1.30 | 0.00 |
| 1833.00 | 2070.00 | 1938.67 | 1.09 | 0.05 |
| 1059.00 | 1202.00 | 1153.33 | 1.07 | 0.18 |
| 26.00 | 22.00 | 26.00 | 1.01 | 1.00 |
| 11.00 | 12.00 | 14.67 | -1.72 | 0.00 |
| 735.00 | 768.00 | 774.67 | 1.07 | 0.11 |
| 58.00 | 66.00 | 65.33 | 2.44 | 0.00 |
| 463.00 | 538.00 | 527.33 | 1.25 | 0.00 |
| 428.00 | 413.00 | 437.67 | 1.03 | 0.55 |
| 734.00 | 854.00 | 836.33 | -1.02 | 0.66 |
| 194.00 | 181.00 | 201.00 | 1.14 | 0.08 |
| 442.00 | 426.00 | 463.67 | 1.04 | 0.45 |
| 36.00 | 47.00 | 42.33 | -1.05 | 0.73 |
| 598.00 | 643.00 | 661.00 | 1.12 | 0.03 |
| 4.00 | 1.00 | 1.67 | -1.58 | 0.59 |
| 400.00 | 382.00 | 386.67 | 1.08 | 0.20 |
| 124.00 | 137.00 | 142.67 | -1.25 | 0.00 |
| 498.00 | 468.00 | 503.33 | 1.10 | 0.09 |
| 887.00 | 979.00 | 952.67 | 1.13 | 0.01 |
| 49.00 | 41.00 | 48.67 | -1.26 | 0.07 |
| 68.00 | 65.00 | 67.67 | -1.11 | 0.35 |
| 20.00 | 16.00 | 19.00 | 1.48 | 0.07 |
| 1013.00 | 1124.00 | 1112.33 | -1.02 | 0.65 |
| 441.00 | 406.00 | 434.00 | 1.03 | 0.56 |
| 7202.00 | 7216.00 | 7474.67 | 1.19 | 0.00 |
| 124.00 | 140.00 | 151.33 | 1.01 | 0.97 |
| 511.00 | 586.00 | 576.00 | 1.08 | 0.13 |
| 49.00 | 43.00 | 52.67 | 1.21 | 0.17 |
| 8.00 | 8.00 | 11.67 | -1.10 | 0.74 |

| | | | | |
|---|---|---|---|---|
| 154.00 | 162.00 | 172.67 | 1.00 | 0.98 |
| 201.00 | 247.00 | 231.67 | 1.12 | 0.10 |
| 5728.00 | 6183.00 | 6287.00 | 1.28 | 0.00 |
| 76.00 | 79.00 | 73.67 | 1.45 | 0.00 |
| 34.00 | 34.00 | 32.33 | -2.28 | 0.00 |
| 19.00 | 31.00 | 29.33 | -1.11 | 0.54 |
| 318.00 | 357.00 | 353.00 | -1.10 | 0.07 |
| 17.00 | 18.00 | 17.00 | -1.35 | 0.16 |
| 30.00 | 58.00 | 43.33 | 1.07 | 0.71 |
| 245.00 | 252.00 | 247.00 | -1.15 | 0.03 |
| 481.00 | 415.00 | 424.33 | -1.54 | 0.00 |
| 29.00 | 29.00 | 29.00 | -1.32 | 0.08 |
| 909.00 | 966.00 | 931.67 | -1.10 | 0.04 |
| 35.00 | 45.00 | 42.00 | 1.08 | 0.63 |
| 8.00 | 13.00 | 11.67 | 2.43 | 0.00 |
| 330.00 | 280.00 | 309.33 | 1.37 | 0.00 |
| 1382.00 | 1412.00 | 1466.67 | 1.15 | 0.00 |
| 348.00 | 350.00 | 386.00 | -1.19 | 0.01 |
| 184.00 | 193.00 | 202.67 | -1.15 | 0.04 |
| 85.00 | 108.00 | 101.00 | 1.31 | 0.01 |
| 547.00 | 542.00 | 551.67 | 1.00 | 0.98 |
| 1082.00 | 1257.00 | 1205.33 | 1.13 | 0.01 |
| 208.00 | 223.00 | 210.33 | 1.08 | 0.36 |
| 5.00 | 7.00 | 6.33 | -1.07 | 1.00 |
| 991.00 | 958.00 | 979.00 | -1.12 | 0.01 |
| 4.00 | 16.00 | 11.67 | 1.11 | 0.81 |
| 298.00 | 381.00 | 357.67 | 1.24 | 0.00 |
| 683.00 | 695.00 | 694.33 | -1.34 | 0.00 |
| 1320.00 | 1292.00 | 1299.33 | -1.27 | 0.00 |
| 1030.00 | 939.00 | 984.67 | -1.07 | 0.20 |
| 551.00 | 514.00 | 541.33 | -1.06 | 0.25 |
| 1572.00 | 1725.00 | 1623.33 | 1.14 | 0.00 |
| 8.00 | 12.00 | 10.67 | 1.13 | 0.71 |
| 7.00 | 5.00 | 6.33 | 1.04 | 1.00 |
| 294.00 | 359.00 | 332.00 | 1.14 | 0.07 |
| 420.00 | 474.00 | 459.33 | -1.04 | 0.46 |
| 289.00 | 350.00 | 353.33 | 1.05 | 0.55 |
| 27.00 | 30.00 | 32.33 | 1.54 | 0.02 |
| 474.00 | 594.00 | 543.33 | -1.15 | 0.01 |
| 2.00 | 4.00 | 3.67 | -1.99 | 0.09 |
| 14.00 | 27.00 | 20.00 | 1.02 | 1.00 |
| 753.00 | 831.00 | 829.33 | -1.04 | 0.41 |
| 452.00 | 496.00 | 479.67 | 1.11 | 0.04 |
| 5.00 | 0.00 | 3.33 | -1.11 | 1.00 |
| 261.00 | 283.00 | 304.00 | 1.22 | 0.01 |
| 85.00 | 118.00 | 104.00 | 1.23 | 0.04 |
| 83.00 | 88.00 | 89.33 | 1.08 | 0.44 |
| 23.00 | 8.00 | 14.00 | -1.37 | 0.20 |

| | | | | |
|---|---|---|---|---|
| 7.00 | 6.00 | 8.67 | 1.07 | 0.89 |
| 26.00 | 14.00 | 21.67 | 2.27 | 0.00 |
| 290.00 | 365.00 | 342.00 | 1.41 | 0.00 |
| 1420.00 | 1576.00 | 1522.67 | 1.01 | 0.87 |
| 64.00 | 65.00 | 66.00 | 1.20 | 0.11 |
| 1.00 | 7.00 | 4.33 | -2.53 | 0.01 |
| 155.00 | 163.00 | 169.67 | 1.10 | 0.20 |
| 57.00 | 44.00 | 48.00 | 1.02 | 0.87 |
| 345.00 | 364.00 | 363.33 | 1.01 | 0.90 |
| 204.00 | 203.00 | 202.67 | -1.48 | 0.00 |
| 547.00 | 614.00 | 564.33 | 1.07 | 0.27 |
| 107.00 | 122.00 | 126.67 | -1.10 | 0.28 |
| 164.00 | 172.00 | 177.67 | 1.14 | 0.09 |
| 27.00 | 20.00 | 21.00 | 1.63 | 0.02 |
| 22.00 | 19.00 | 21.33 | 1.37 | 0.12 |
| 56.00 | 62.00 | 63.00 | -2.02 | 0.00 |
| 7.00 | 7.00 | 8.33 | 1.17 | 0.67 |
| 271.00 | 323.00 | 309.00 | 1.22 | 0.01 |
| 120.00 | 113.00 | 124.00 | -1.01 | 0.92 |
| 489.00 | 523.00 | 520.33 | 1.06 | 0.25 |
| 258.00 | 330.00 | 299.33 | 1.26 | 0.00 |
| 88.00 | 124.00 | 114.67 | 1.01 | 0.95 |
| 13.00 | 14.00 | 14.33 | -1.20 | 0.49 |
| 319.00 | 343.00 | 334.67 | 1.19 | 0.00 |
| 30.00 | 29.00 | 26.00 | -1.06 | 0.83 |
| 327.00 | 284.00 | 315.67 | 1.22 | 0.01 |
| 190.00 | 229.00 | 216.67 | 1.53 | 0.00 |
| 2.00 | 5.00 | 5.00 | -2.66 | 0.00 |
| 1337.00 | 1462.00 | 1426.67 | 1.14 | 0.00 |
| 587.00 | 655.00 | 634.67 | 1.02 | 0.72 |
| 1352.00 | 1624.00 | 1469.00 | -1.17 | 0.00 |
| 757.00 | 810.00 | 792.67 | 1.11 | 0.03 |
| 1233.00 | 1420.00 | 1376.67 | 1.38 | 0.00 |
| 517.00 | 574.00 | 560.33 | 1.13 | 0.04 |
| 574.00 | 562.00 | 551.67 | -1.27 | 0.00 |
| 665.00 | 661.00 | 676.00 | -1.17 | 0.00 |
| 1492.00 | 1519.00 | 1519.00 | -1.03 | 0.50 |
| 15.00 | 20.00 | 20.00 | -1.60 | 0.01 |
| 29.00 | 36.00 | 35.00 | -1.30 | 0.06 |
| 189.00 | 232.00 | 211.00 | 1.10 | 0.18 |
| 798.00 | 763.00 | 834.67 | 1.03 | 0.65 |
| 1352.00 | 1381.00 | 1405.67 | 1.00 | 0.91 |
| 2783.00 | 2722.00 | 2733.00 | 1.20 | 0.00 |
| 270.00 | 298.00 | 301.00 | -1.45 | 0.00 |
| 61.00 | 58.00 | 57.67 | 1.01 | 0.96 |
| 4.00 | 4.00 | 4.33 | -1.92 | 0.08 |
| 340.00 | 392.00 | 373.67 | -1.01 | 0.82 |
| 24.00 | 32.00 | 34.33 | -1.02 | 0.95 |

| | | | | |
|---|---|---|---|---|
| 49.00 | 25.00 | 29.00 | 3.74 | 0.00 |
| 22.00 | 33.00 | 26.33 | 1.20 | 0.34 |
| 29.00 | 28.00 | 30.67 | 4.47 | 0.00 |
| 3826.00 | 4528.00 | 4416.00 | 1.10 | 0.03 |
| 2879.00 | 2581.00 | 2772.00 | -1.10 | 0.04 |
| 277.00 | 294.00 | 291.33 | -1.04 | 0.58 |
| 403.00 | 498.00 | 477.33 | 1.19 | 0.00 |
| 512.00 | 488.00 | 520.33 | -1.01 | 0.81 |
| 53.00 | 78.00 | 74.33 | -1.19 | 0.13 |
| 501.00 | 547.00 | 543.67 | -1.05 | 0.34 |
| 946.00 | 990.00 | 1003.67 | 1.03 | 0.44 |
| 2077.00 | 2304.00 | 2284.67 | 1.15 | 0.00 |
| 111.00 | 112.00 | 118.00 | -1.08 | 0.38 |
| 13.00 | 13.00 | 14.33 | -1.46 | 0.09 |
| 167.00 | 195.00 | 179.00 | 1.02 | 0.84 |
| 14.00 | 23.00 | 18.00 | -1.41 | 0.08 |
| 886.00 | 992.00 | 963.33 | 1.32 | 0.00 |
| 7.00 | 15.00 | 14.67 | -1.14 | 0.68 |
| 1501.00 | 1576.00 | 1586.33 | 1.10 | 0.01 |
| 138.00 | 172.00 | 157.33 | -1.10 | 0.22 |
| 9.00 | 6.00 | 7.33 | -1.42 | 0.29 |
| 142.00 | 149.00 | 147.67 | -1.12 | 0.18 |
| 47.00 | 47.00 | 48.00 | 1.09 | 0.53 |
| 58.00 | 70.00 | 67.67 | -1.11 | 0.34 |
| 418.00 | 457.00 | 467.00 | 1.22 | 0.00 |
| 629.00 | 845.00 | 806.67 | 1.10 | 0.16 |
| 121.00 | 134.00 | 127.33 | -1.01 | 0.90 |
| 463.00 | 416.00 | 456.33 | 1.10 | 0.10 |
| 309.00 | 348.00 | 352.33 | 1.15 | 0.05 |
| 1564.00 | 1652.00 | 1702.00 | -1.01 | 0.74 |
| 23.00 | 29.00 | 23.33 | -1.06 | 0.88 |
| 1095.00 | 1033.00 | 1139.67 | 1.12 | 0.03 |
| 142.00 | 186.00 | 178.00 | 1.10 | 0.29 |
| 502.00 | 601.00 | 579.33 | 1.24 | 0.00 |
| 19.00 | 24.00 | 21.00 | -1.43 | 0.04 |
| 38.00 | 34.00 | 32.33 | -1.12 | 0.56 |
| 22.00 | 20.00 | 23.67 | -1.06 | 0.75 |
| 2.00 | 3.00 | 5.00 | -1.68 | 0.17 |
| 135.00 | 162.00 | 140.33 | -1.06 | 0.54 |
| 441.00 | 523.00 | 488.33 | 1.23 | 0.00 |
| 12.00 | 11.00 | 14.67 | -1.40 | 0.11 |
| 11.00 | 8.00 | 12.00 | -1.15 | 0.66 |
| 1997.00 | 2080.00 | 2107.33 | 1.01 | 0.86 |
| 540.00 | 542.00 | 536.33 | -1.08 | 0.18 |
| 1032.00 | 1139.00 | 1123.33 | 1.15 | 0.00 |
| 157.00 | 185.00 | 183.67 | 1.16 | 0.06 |
| 252.00 | 235.00 | 262.67 | -1.10 | 0.18 |
| 116.00 | 105.00 | 113.67 | -1.10 | 0.28 |

| | | | | |
|---|---|---|---|---|
| 940.00 | 912.00 | 968.33 | 1.12 | 0.01 |
| 22.00 | 9.00 | 18.00 | -1.05 | 0.93 |
| 79.00 | 85.00 | 84.33 | -1.16 | 0.12 |
| 293.00 | 354.00 | 319.00 | -1.20 | 0.00 |
| 21.00 | 15.00 | 16.00 | 1.35 | 0.21 |
| 63.00 | 61.00 | 61.67 | -1.22 | 0.12 |
| 139.00 | 160.00 | 164.33 | -1.13 | 0.13 |
| 300.00 | 315.00 | 314.33 | -1.02 | 0.74 |
| 16.00 | 10.00 | 15.67 | 1.08 | 0.76 |
| 399.00 | 567.00 | 487.33 | 1.13 | 0.10 |
| 1420.00 | 1626.00 | 1513.33 | 1.03 | 0.51 |
| 181.00 | 175.00 | 180.67 | -1.00 | 1.00 |
| 1388.00 | 1376.00 | 1372.67 | -1.07 | 0.12 |
| 202.00 | 220.00 | 222.33 | -1.45 | 0.00 |
| 131.00 | 148.00 | 147.67 | -1.15 | 0.09 |
| 7.00 | 6.00 | 5.67 | -1.30 | 0.55 |
| 36.00 | 34.00 | 32.67 | 1.04 | 0.79 |
| 507.00 | 566.00 | 525.33 | 1.07 | 0.26 |
| 831.00 | 922.00 | 824.00 | 1.07 | 0.33 |
| 3514.00 | 3677.00 | 3850.33 | -1.29 | 0.00 |
| 291.00 | 270.00 | 302.33 | -1.20 | 0.00 |
| 406.00 | 452.00 | 456.00 | -1.13 | 0.03 |
| 124.00 | 107.00 | 111.00 | 1.36 | 0.00 |
| 94.00 | 122.00 | 106.67 | -1.15 | 0.14 |
| 115.00 | 116.00 | 119.00 | -1.56 | 0.00 |
| 1020.00 | 1055.00 | 1010.00 | -1.45 | 0.00 |
| 893.00 | 912.00 | 902.67 | -1.07 | 0.15 |
| 16.00 | 15.00 | 18.00 | -1.13 | 0.59 |
| 1285.00 | 1473.00 | 1410.33 | 1.13 | 0.00 |
| 1927.00 | 2311.00 | 2260.00 | 1.35 | 0.00 |
| 5544.00 | 5443.00 | 5507.00 | -1.13 | 0.00 |
| 8.00 | 6.00 | 6.33 | -1.64 | 0.13 |
| 12.00 | 24.00 | 19.33 | -1.14 | 0.55 |
| 401.00 | 374.00 | 396.67 | 1.13 | 0.07 |
| 38.00 | 36.00 | 36.33 | -1.11 | 0.50 |
| 1143.00 | 1155.00 | 1162.33 | -1.42 | 0.00 |
| 1584.00 | 1824.00 | 1779.33 | 1.06 | 0.22 |
| 67.00 | 77.00 | 66.67 | 1.30 | 0.07 |
| 3957.00 | 4340.00 | 4135.00 | 1.22 | 0.00 |
| 21.00 | 19.00 | 19.67 | -1.17 | 0.49 |
| 2417.00 | 2643.00 | 2601.00 | 1.22 | 0.00 |
| 11620.00 | 13996.00 | 13319.67 | -1.11 | 0.01 |
| 2330.00 | 2460.00 | 2477.33 | 1.16 | 0.00 |
| 274.00 | 262.00 | 277.33 | -1.24 | 0.00 |
| 7.00 | 15.00 | 11.67 | -1.42 | 0.17 |
| 16778.00 | 17174.00 | 17450.67 | -1.08 | 0.07 |
| 14789.00 | 14885.00 | 15062.67 | -1.04 | 0.25 |
| 410.00 | 444.00 | 448.67 | 1.07 | 0.19 |

| | | | | |
|---|---|---|---|---|
| 20.00 | 12.00 | 16.67 | -1.15 | 0.58 |
| 747.00 | 787.00 | 792.00 | 1.20 | 0.00 |
| 97.00 | 94.00 | 95.33 | -1.09 | 0.43 |
| 356.00 | 346.00 | 368.67 | -1.08 | 0.20 |
| 1279.00 | 1338.00 | 1355.33 | 1.02 | 0.71 |
| 484.00 | 545.00 | 548.00 | 1.02 | 0.68 |
| 24.00 | 32.00 | 30.33 | 1.34 | 0.11 |
| 104.00 | 97.00 | 102.33 | 1.14 | 0.17 |
| 304.00 | 298.00 | 294.00 | -1.16 | 0.02 |
| 1202.00 | 1257.00 | 1237.00 | 1.04 | 0.36 |
| 99.00 | 124.00 | 129.00 | 1.01 | 0.96 |
| 1274.00 | 1199.00 | 1285.67 | -1.00 | 1.00 |
| 1597.00 | 1529.00 | 1537.00 | -1.07 | 0.19 |
| 1616.00 | 1719.00 | 1653.67 | -1.02 | 0.65 |
| 274.00 | 374.00 | 370.33 | 1.07 | 0.44 |
| 4724.00 | 4652.00 | 4905.67 | -1.21 | 0.00 |
| 2612.00 | 2885.00 | 2692.00 | 1.01 | 0.73 |
| 2925.00 | 2888.00 | 2970.33 | -1.07 | 0.12 |
| 25400.00 | 27588.00 | 26609.33 | -1.15 | 0.00 |
| 6999.00 | 7284.00 | 7414.67 | 1.04 | 0.21 |
| 609.00 | 668.00 | 672.67 | -1.18 | 0.00 |
| 18.00 | 21.00 | 19.00 | 1.86 | 0.01 |
| 2718.00 | 2796.00 | 2837.67 | -1.06 | 0.12 |
| 672.00 | 602.00 | 656.33 | -1.13 | 0.02 |
| 172.00 | 168.00 | 164.67 | -1.28 | 0.00 |
| 8.00 | 10.00 | 10.00 | -1.61 | 0.06 |
| 2772.00 | 2557.00 | 2740.33 | -1.08 | 0.07 |
| 11032.00 | 11414.00 | 11455.33 | -1.11 | 0.00 |
| 108.00 | 101.00 | 106.00 | -1.14 | 0.14 |
| 286.00 | 240.00 | 270.33 | -1.48 | 0.00 |
| 630.00 | 664.00 | 650.33 | -1.15 | 0.00 |
| 3643.00 | 3793.00 | 3881.33 | 1.11 | 0.00 |
| 3.00 | 1.00 | 1.33 | -1.24 | 1.00 |
| 6587.00 | 6804.00 | 6753.33 | 1.10 | 0.01 |
| 4540.00 | 4548.00 | 4732.00 | -1.01 | 0.80 |
| 131.00 | 146.00 | 158.67 | -1.45 | 0.00 |
| 51.00 | 70.00 | 65.33 | -1.07 | 0.58 |
| 4291.00 | 4823.00 | 4870.00 | 1.21 | 0.00 |
| 1249.00 | 1478.00 | 1438.67 | 1.22 | 0.00 |
| 2078.00 | 2046.00 | 2174.00 | -1.05 | 0.27 |
| 1510.00 | 1437.00 | 1520.67 | -1.06 | 0.17 |
| 112.00 | 128.00 | 130.67 | -1.31 | 0.00 |
| 2650.00 | 3256.00 | 2974.33 | 1.31 | 0.00 |
| 117.00 | 132.00 | 126.33 | -1.00 | 1.00 |
| 45.00 | 22.00 | 30.67 | -1.54 | 0.02 |
| 277.00 | 330.00 | 294.33 | -1.06 | 0.39 |
| 289.00 | 258.00 | 304.33 | 1.40 | 0.00 |
| 960.00 | 994.00 | 1023.67 | -1.08 | 0.13 |

| | | | | |
|---|---|---|---|---|
| 1905.00 | 1899.00 | 1922.00 | 1.06 | 0.18 |
| 370.00 | 377.00 | 377.67 | 1.22 | 0.01 |
| 254.00 | 265.00 | 265.33 | -1.04 | 0.50 |
| 1146.00 | 1186.00 | 1202.33 | 1.01 | 0.72 |
| 93.00 | 84.00 | 80.33 | 1.29 | 0.03 |
| 5.00 | 4.00 | 3.67 | -1.28 | 0.69 |
| 4.00 | 3.00 | 2.67 | -1.86 | 0.22 |
| 26.00 | 34.00 | 36.00 | -1.01 | 1.00 |
| 2751.00 | 2833.00 | 2825.33 | -1.02 | 0.63 |
| 2060.00 | 2114.00 | 2064.67 | 1.01 | 0.75 |
| 352.00 | 382.00 | 347.00 | 1.12 | 0.13 |
| 34.00 | 35.00 | 32.33 | 1.35 | 0.07 |
| 478.00 | 566.00 | 530.67 | -1.06 | 0.29 |
| 384.00 | 401.00 | 394.33 | 1.03 | 0.62 |
| 566.00 | 519.00 | 534.33 | 1.04 | 0.53 |
| 53.00 | 69.00 | 63.67 | 1.06 | 0.67 |
| 8.00 | 8.00 | 9.33 | -1.44 | 0.19 |
| 935.00 | 995.00 | 969.00 | 1.06 | 0.22 |
| 570.00 | 650.00 | 643.67 | 1.35 | 0.00 |
| 1939.00 | 2002.00 | 2055.00 | -1.02 | 0.62 |
| 13.00 | 11.00 | 8.00 | 1.40 | 0.52 |
| 104.00 | 87.00 | 95.67 | 1.06 | 0.57 |
| 319.00 | 345.00 | 336.67 | -1.10 | 0.09 |
| 736.00 | 917.00 | 838.33 | 1.01 | 0.81 |
| 700.00 | 775.00 | 729.00 | -1.04 | 0.40 |
| 232.00 | 215.00 | 232.33 | -1.04 | 0.55 |
| 122.00 | 146.00 | 131.00 | 1.90 | 0.00 |
| 3264.00 | 3558.00 | 3449.67 | 1.06 | 0.12 |
| 357.00 | 366.00 | 399.00 | 1.09 | 0.17 |
| 192.00 | 170.00 | 198.00 | 1.09 | 0.26 |
| 109.00 | 143.00 | 130.00 | 1.00 | 1.00 |
| 26.00 | 16.00 | 21.33 | -1.17 | 0.46 |
| 28.00 | 35.00 | 31.33 | -1.02 | 0.89 |
| 7.00 | 5.00 | 4.67 | 1.25 | 0.70 |
| 13972.00 | 14950.00 | 14594.67 | 1.32 | 0.00 |
| 1065.00 | 1226.00 | 1142.00 | 1.25 | 0.00 |
| 269.00 | 240.00 | 259.00 | -1.26 | 0.00 |
| 926.00 | 939.00 | 935.33 | -1.18 | 0.00 |
| 1872.00 | 2190.00 | 2119.33 | 1.36 | 0.00 |
| 1650.00 | 1630.00 | 1574.00 | -1.15 | 0.01 |
| 830.00 | 857.00 | 851.33 | 1.07 | 0.22 |
| 662.00 | 768.00 | 720.67 | 1.09 | 0.07 |
| 67.00 | 68.00 | 60.00 | -1.15 | 0.31 |
| 984.00 | 1074.00 | 1110.67 | 1.12 | 0.02 |
| 76.00 | 92.00 | 83.33 | 1.62 | 0.00 |
| 22.00 | 21.00 | 19.67 | 1.06 | 0.79 |
| 495.00 | 652.00 | 566.00 | 1.21 | 0.00 |
| 207.00 | 200.00 | 198.67 | 1.04 | 0.66 |

| | | | | |
|---|---|---|---|---|
| 251.00 | 347.00 | 334.67 | 1.06 | 0.47 |
| 244.00 | 297.00 | 267.00 | -1.05 | 0.55 |
| 48.00 | 84.00 | 57.33 | -1.11 | 0.56 |
| 312.00 | 333.00 | 316.33 | -1.03 | 0.66 |
| 154.00 | 179.00 | 168.00 | -1.04 | 0.60 |
| 1161.00 | 1162.00 | 1182.67 | -1.14 | 0.00 |
| 209.00 | 201.00 | 208.67 | 1.21 | 0.04 |
| 4869.00 | 5509.00 | 5451.00 | 1.23 | 0.00 |
| 39.00 | 35.00 | 30.67 | -1.02 | 0.95 |
| 565.00 | 616.00 | 601.67 | 1.01 | 0.86 |
| 2563.00 | 2693.00 | 2586.00 | 1.15 | 0.00 |
| 2376.00 | 2332.00 | 2397.67 | 1.08 | 0.04 |
| 304.00 | 298.00 | 308.67 | -1.11 | 0.09 |
| 141.00 | 149.00 | 145.00 | 1.17 | 0.06 |
| 448.00 | 439.00 | 483.00 | 1.25 | 0.00 |
| 1340.00 | 1362.00 | 1382.33 | -1.09 | 0.03 |
| 1248.00 | 1295.00 | 1331.00 | -1.18 | 0.00 |
| 569.00 | 563.00 | 562.67 | 1.08 | 0.15 |
| 3557.00 | 4149.00 | 3899.00 | -1.01 | 0.71 |
| 1914.00 | 2037.00 | 2042.33 | 1.01 | 0.88 |
| 3935.00 | 4072.00 | 3955.67 | -1.23 | 0.00 |
| 561.00 | 594.00 | 628.33 | -1.01 | 0.85 |
| 840.00 | 939.00 | 956.33 | 1.12 | 0.02 |
| 203.00 | 208.00 | 222.67 | 1.05 | 0.46 |
| 290.00 | 308.00 | 312.00 | 1.07 | 0.26 |
| 34.00 | 40.00 | 36.33 | 1.21 | 0.25 |
| 22.00 | 19.00 | 23.00 | 1.54 | 0.04 |
| 224.00 | 263.00 | 259.67 | -1.05 | 0.48 |
| 203.00 | 225.00 | 224.00 | 1.08 | 0.31 |
| 208.00 | 266.00 | 244.00 | 1.57 | 0.00 |
| 15.00 | 17.00 | 14.67 | 1.49 | 0.11 |
| 296.00 | 369.00 | 328.33 | 1.35 | 0.00 |
| 198.00 | 202.00 | 205.00 | -1.07 | 0.33 |
| 301.00 | 377.00 | 372.67 | -1.10 | 0.17 |
| 6.00 | 4.00 | 7.00 | -1.35 | 0.41 |
| 310.00 | 371.00 | 359.67 | 1.14 | 0.03 |
| 381.00 | 462.00 | 438.67 | 1.20 | 0.00 |
| 147.00 | 211.00 | 180.33 | 1.05 | 0.61 |
| 18.00 | 20.00 | 18.33 | 1.50 | 0.07 |
| 119.00 | 132.00 | 126.33 | 1.16 | 0.10 |
| 996.00 | 1080.00 | 1118.67 | 1.13 | 0.01 |
| 17.00 | 17.00 | 21.00 | 1.26 | 0.34 |
| 160.00 | 191.00 | 180.00 | 1.09 | 0.23 |
| 331.00 | 361.00 | 336.33 | -1.04 | 0.54 |
| 851.00 | 994.00 | 957.00 | -1.18 | 0.00 |
| 84.00 | 59.00 | 75.00 | -1.04 | 0.81 |
| 559.00 | 513.00 | 542.67 | -1.41 | 0.00 |
| 1424.00 | 1524.00 | 1508.33 | 1.10 | 0.01 |

| | | | | |
|---|---|---|---|---|
| 182.00 | 161.00 | 183.67 | 1.24 | 0.01 |
| 4089.00 | 5084.00 | 4743.00 | -1.05 | 0.24 |
| 376.00 | 427.00 | 394.67 | -1.27 | 0.00 |
| 19.00 | 15.00 | 18.00 | -1.57 | 0.01 |
| 26.00 | 35.00 | 29.67 | -1.59 | 0.00 |
| 110.00 | 135.00 | 134.00 | -1.34 | 0.00 |
| 15.00 | 20.00 | 17.00 | 1.92 | 0.01 |
| 200.00 | 163.00 | 176.33 | -1.38 | 0.00 |
| 183.00 | 187.00 | 191.67 | 1.36 | 0.00 |
| 360.00 | 407.00 | 403.33 | -1.07 | 0.35 |
| 6.00 | 5.00 | 7.00 | -1.49 | 0.22 |
| 145.00 | 148.00 | 144.67 | 1.30 | 0.00 |
| 0.00 | 0.00 | 0.00 | -27.86 | 0.00 |
| 14.00 | 10.00 | 14.33 | -1.29 | 0.28 |
| 43.00 | 36.00 | 42.00 | 1.22 | 0.20 |
| 151.00 | 127.00 | 141.33 | -1.22 | 0.08 |
| 167.00 | 158.00 | 174.00 | -1.33 | 0.00 |
| 144.00 | 168.00 | 169.00 | 1.09 | 0.34 |
| 4.00 | 4.00 | 4.00 | -1.42 | 0.47 |
| 45.00 | 51.00 | 47.33 | 1.11 | 0.47 |
| 133.00 | 131.00 | 130.67 | -1.35 | 0.00 |
| 5.00 | 8.00 | 5.33 | -1.20 | 0.74 |
| 95.00 | 81.00 | 99.33 | 1.22 | 0.07 |
| 8.00 | 4.00 | 7.00 | 1.29 | 0.52 |
| 468.00 | 468.00 | 485.67 | -1.08 | 0.15 |
| 67.00 | 84.00 | 76.67 | -1.01 | 0.93 |
| 600.00 | 695.00 | 665.00 | -1.02 | 0.72 |
| 94.00 | 119.00 | 108.00 | 1.25 | 0.03 |
| 10.00 | 6.00 | 9.67 | -2.68 | 0.00 |
| 33.00 | 34.00 | 39.00 | -1.50 | 0.00 |
| 310.00 | 310.00 | 299.33 | -1.02 | 0.79 |
| 545.00 | 530.00 | 578.67 | -1.01 | 0.84 |
| 67.00 | 98.00 | 95.67 | 1.08 | 0.54 |
| 565.00 | 568.00 | 580.33 | 1.08 | 0.10 |
| 290.00 | 340.00 | 331.33 | -1.01 | 0.89 |
| 56.00 | 65.00 | 75.00 | 1.18 | 0.22 |
| 18.00 | 15.00 | 14.33 | -1.27 | 0.33 |
| 345.00 | 348.00 | 351.33 | 1.29 | 0.00 |
| 132.00 | 160.00 | 133.67 | 1.09 | 0.51 |
| 0.00 | 0.00 | 0.67 | -1.88 | 0.71 |
| 2.00 | 5.00 | 3.67 | 2.57 | 0.13 |
| 33.00 | 37.00 | 35.33 | -1.53 | 0.00 |
| 489.00 | 555.00 | 538.00 | 1.14 | 0.01 |
| 5163.00 | 5584.00 | 5517.67 | 1.02 | 0.47 |
| 1821.00 | 1935.00 | 1921.00 | -1.21 | 0.00 |
| 338.00 | 397.00 | 383.33 | 1.05 | 0.38 |
| 347.00 | 430.00 | 408.00 | 1.14 | 0.03 |
| 1111.00 | 1047.00 | 1086.67 | -1.20 | 0.00 |

| | | | | |
|---|---|---|---|---|
| 542.00 | 588.00 | 596.33 | 1.29 | 0.00 |
| 612.00 | 788.00 | 750.67 | 1.23 | 0.00 |
| 1735.00 | 1898.00 | 1839.00 | 1.14 | 0.01 |
| 274.00 | 311.00 | 307.33 | 1.21 | 0.00 |
| 20.00 | 26.00 | 22.67 | 1.03 | 0.93 |
| 584.00 | 667.00 | 643.67 | -1.10 | 0.06 |
| 74.00 | 67.00 | 77.00 | -1.33 | 0.01 |
| 158.00 | 201.00 | 178.00 | 1.15 | 0.08 |
| 3.00 | 9.00 | 7.00 | -1.77 | 0.06 |
| 388.00 | 433.00 | 462.33 | 1.01 | 0.86 |
| 680.00 | 766.00 | 732.67 | 1.32 | 0.00 |
| 40.00 | 56.00 | 48.67 | 2.18 | 0.00 |
| 7.00 | 6.00 | 8.67 | 1.81 | 0.09 |
| 617.00 | 716.00 | 685.00 | 1.20 | 0.00 |
| 582.00 | 611.00 | 629.00 | 1.05 | 0.36 |
| 63.00 | 72.00 | 66.00 | -1.04 | 0.79 |
| 8015.00 | 8619.00 | 8334.00 | 1.48 | 0.00 |
| 723.00 | 830.00 | 774.00 | 1.12 | 0.02 |
| 227.00 | 255.00 | 261.67 | -1.10 | 0.17 |
| 9.00 | 11.00 | 8.67 | -1.01 | 1.00 |
| 1659.00 | 1618.00 | 1705.67 | 1.81 | 0.00 |
| 453.00 | 477.00 | 460.00 | 1.03 | 0.54 |
| 1934.00 | 2014.00 | 2043.33 | 1.02 | 0.65 |
| 15.00 | 7.00 | 11.67 | 1.63 | 0.09 |
| 1471.00 | 1571.00 | 1503.00 | -1.07 | 0.11 |
| 75.00 | 77.00 | 77.33 | 1.52 | 0.00 |
| 451.00 | 409.00 | 422.67 | -1.19 | 0.01 |
| 533.00 | 566.00 | 546.00 | 1.17 | 0.00 |
| 645.00 | 717.00 | 723.33 | 1.12 | 0.03 |
| 577.00 | 645.00 | 606.67 | 1.10 | 0.07 |
| 21.00 | 22.00 | 22.67 | -1.03 | 0.93 |
| 1721.00 | 1773.00 | 1760.67 | -1.17 | 0.00 |
| 260.00 | 282.00 | 286.33 | 1.18 | 0.01 |
| 10.00 | 11.00 | 10.00 | -1.61 | 0.06 |
| 449.00 | 484.00 | 484.33 | -1.05 | 0.35 |
| 32.00 | 32.00 | 33.33 | -1.29 | 0.08 |
| 1.00 | 1.00 | 0.67 | -7.41 | 0.00 |
| 10.00 | 5.00 | 7.33 | 1.27 | 0.53 |
| 87.00 | 120.00 | 107.00 | 1.74 | 0.00 |
| 158.00 | 177.00 | 167.67 | -1.11 | 0.17 |
| 803.00 | 880.00 | 852.67 | 1.10 | 0.07 |
| 34.00 | 40.00 | 41.67 | 1.28 | 0.10 |
| 642.00 | 749.00 | 701.33 | 1.28 | 0.00 |
| 2681.00 | 3092.00 | 2988.67 | 1.22 | 0.00 |
| 1523.00 | 1700.00 | 1736.67 | -1.04 | 0.41 |
| 983.00 | 1136.00 | 1072.67 | 1.18 | 0.00 |
| 11.00 | 18.00 | 12.00 | 1.22 | 0.49 |
| 1058.00 | 1311.00 | 1240.00 | 1.22 | 0.00 |

| | | | | |
|---|---|---|---|---|
| 13754.00 | 12964.00 | 13388.33 | 1.04 | 0.35 |
| 412.00 | 457.00 | 480.00 | 1.26 | 0.00 |
| 994.00 | 1138.00 | 1110.67 | -1.03 | 0.42 |
| 122.00 | 126.00 | 135.00 | 1.17 | 0.09 |
| 192.00 | 260.00 | 230.67 | 1.66 | 0.00 |
| 498.00 | 578.00 | 551.00 | 1.20 | 0.00 |
| 1333.00 | 1477.00 | 1484.33 | 1.03 | 0.57 |
| 276.00 | 286.00 | 280.00 | -1.00 | 1.00 |
| 1226.00 | 1348.00 | 1330.33 | 1.24 | 0.00 |
| 2610.00 | 2565.00 | 2689.67 | 1.08 | 0.07 |
| 446.00 | 531.00 | 515.00 | 1.05 | 0.42 |
| 505.00 | 455.00 | 500.67 | -1.07 | 0.25 |
| 997.00 | 934.00 | 992.67 | 1.00 | 0.96 |
| 3273.00 | 4003.00 | 3779.00 | -1.00 | 0.92 |
| 6472.00 | 6970.00 | 7076.67 | -1.19 | 0.00 |
| 395.00 | 394.00 | 394.33 | 1.03 | 0.59 |
| 1529.00 | 1606.00 | 1555.00 | -1.01 | 0.83 |
| 1549.00 | 1968.00 | 1774.67 | -1.03 | 0.58 |
| 14.00 | 9.00 | 10.33 | -1.69 | 0.03 |
| 2181.00 | 2423.00 | 2368.33 | 1.12 | 0.00 |
| 577.00 | 601.00 | 598.67 | 1.23 | 0.00 |
| 207.00 | 274.00 | 225.33 | 1.02 | 0.85 |
| 469.00 | 545.00 | 506.33 | 1.19 | 0.00 |
| 395.00 | 428.00 | 417.00 | -1.24 | 0.00 |
| 33.00 | 35.00 | 33.00 | 1.57 | 0.01 |
| 19.00 | 28.00 | 23.33 | 1.23 | 0.31 |
| 21.00 | 25.00 | 22.67 | -1.68 | 0.00 |
| 791.00 | 939.00 | 892.67 | -1.07 | 0.16 |
| 155.00 | 201.00 | 179.00 | 1.25 | 0.01 |
| 15.00 | 19.00 | 16.67 | -1.15 | 0.58 |
| 685.00 | 746.00 | 717.67 | -1.08 | 0.10 |
| 6370.00 | 7267.00 | 7000.67 | 1.10 | 0.01 |
| 13168.00 | 14383.00 | 14360.33 | 1.14 | 0.00 |
| 8232.00 | 8808.00 | 8983.33 | 1.23 | 0.00 |
| 11731.00 | 13390.00 | 13009.33 | 1.25 | 0.00 |
| 9400.00 | 10817.00 | 10442.33 | 1.13 | 0.00 |
| 24.00 | 24.00 | 30.33 | 1.42 | 0.07 |
| 180.00 | 286.00 | 236.00 | 1.22 | 0.03 |
| 41.00 | 71.00 | 54.67 | -1.16 | 0.35 |
| 9149.00 | 10250.00 | 10046.00 | 1.16 | 0.00 |
| 7336.00 | 8180.00 | 8119.33 | 1.29 | 0.00 |
| 496.00 | 570.00 | 553.67 | 1.19 | 0.00 |
| 188.00 | 230.00 | 215.33 | 1.26 | 0.01 |
| 23.00 | 22.00 | 20.33 | -1.05 | 0.87 |
| 2264.00 | 2375.00 | 2226.00 | -1.06 | 0.27 |
| 7.00 | 5.00 | 5.67 | -1.77 | 0.09 |
| 7581.00 | 7476.00 | 7933.00 | -1.10 | 0.02 |
| 2844.00 | 3217.00 | 3103.33 | 1.08 | 0.03 |

| | | | | |
|---|---|---|---|---|
| 8384.00 | 9279.00 | 8839.33 | 1.15 | 0.00 |
| 78.00 | 76.00 | 87.33 | -1.44 | 0.00 |
| 90.00 | 87.00 | 83.67 | 1.02 | 0.84 |
| 129.00 | 95.00 | 115.67 | -1.23 | 0.03 |
| 10.00 | 16.00 | 15.33 | 1.03 | 1.00 |
| 3.00 | 10.00 | 8.00 | 1.94 | 0.08 |
| 1.00 | 1.00 | 1.00 | -4.01 | 0.02 |
| 1966.00 | 2037.00 | 2066.00 | -1.30 | 0.00 |
| 933.00 | 950.00 | 978.33 | -1.19 | 0.00 |
| 1406.00 | 1439.00 | 1499.67 | 1.01 | 0.76 |
| 828.00 | 983.00 | 893.33 | 1.11 | 0.06 |
| 1784.00 | 1929.00 | 1922.33 | 1.05 | 0.18 |
| 35.00 | 30.00 | 31.00 | 1.31 | 0.12 |
| 2673.00 | 2516.00 | 2707.67 | 1.43 | 0.00 |
| 9.00 | 3.00 | 9.00 | -1.20 | 0.62 |
| 6.00 | 16.00 | 11.67 | -1.13 | 0.78 |
| 760.00 | 773.00 | 778.00 | 1.01 | 0.85 |
| 27293.00 | 28346.00 | 28215.00 | 1.13 | 0.00 |
| 3128.00 | 3544.00 | 3400.33 | -1.12 | 0.00 |
| 3326.00 | 3456.00 | 3381.33 | 1.17 | 0.00 |
| 2174.00 | 2206.00 | 2257.67 | -1.08 | 0.04 |
| 278.00 | 338.00 | 307.00 | -1.15 | 0.09 |
| 24973.00 | 24538.00 | 25305.33 | 1.19 | 0.00 |
| 9036.00 | 8837.00 | 9293.67 | -1.39 | 0.00 |
| 1617.00 | 1506.00 | 1562.33 | 1.11 | 0.03 |
| 37.00 | 54.00 | 49.67 | -1.09 | 0.51 |
| 107.00 | 117.00 | 119.67 | 2.70 | 0.00 |
| 5541.00 | 5620.00 | 5774.33 | 1.07 | 0.12 |
| 0.00 | 1.00 | 1.33 | -3.10 | 0.05 |
| 74.00 | 96.00 | 84.33 | -1.53 | 0.00 |
| 381.00 | 347.00 | 383.67 | 1.95 | 0.00 |
| 986.00 | 1075.00 | 1100.33 | -1.13 | 0.01 |
| 32.00 | 28.00 | 31.33 | -1.14 | 0.38 |
| 21.00 | 26.00 | 22.00 | 1.38 | 0.10 |
| 38.00 | 32.00 | 34.67 | 1.21 | 0.25 |
| 1460.00 | 1262.00 | 1384.00 | -1.04 | 0.46 |
| 188.00 | 183.00 | 200.67 | -1.34 | 0.00 |
| 146.00 | 152.00 | 152.33 | 1.03 | 0.68 |
| 694.00 | 681.00 | 682.67 | -1.20 | 0.00 |
| 3096.00 | 3224.00 | 3316.00 | 1.18 | 0.00 |
| 184.00 | 236.00 | 193.33 | 1.04 | 0.66 |
| 398.00 | 372.00 | 395.33 | -1.01 | 0.88 |
| 1.00 | 2.00 | 1.33 | -1.94 | 0.39 |
| 1475.00 | 1783.00 | 1695.33 | 1.02 | 0.70 |
| 2772.00 | 2998.00 | 2966.67 | 1.28 | 0.00 |
| 14.00 | 20.00 | 17.33 | 1.65 | 0.03 |
| 880.00 | 1073.00 | 992.67 | 1.05 | 0.29 |
| 356.00 | 418.00 | 409.33 | 1.40 | 0.00 |

| | | | | |
|---|---|---|---|---|
| 1646.00 | 1688.00 | 1751.33 | 1.04 | 0.34 |
| 181.00 | 257.00 | 227.33 | 1.32 | 0.00 |
| 145.00 | 161.00 | 165.33 | -1.04 | 0.63 |
| 2871.00 | 3150.00 | 3030.67 | 1.06 | 0.18 |
| 1251.00 | 1258.00 | 1327.67 | 1.06 | 0.17 |
| 4816.00 | 5167.00 | 5251.00 | -1.09 | 0.03 |
| 269.00 | 341.00 | 321.33 | 1.03 | 0.69 |
| 7.00 | 5.00 | 9.67 | -1.26 | 0.49 |
| 628.00 | 685.00 | 660.00 | 1.18 | 0.00 |
| 2285.00 | 2570.00 | 2506.33 | 1.08 | 0.05 |
| 2079.00 | 2222.00 | 2145.67 | -1.04 | 0.28 |
| 3221.00 | 3269.00 | 3272.67 | -1.22 | 0.00 |
| 3969.00 | 3956.00 | 4155.33 | -1.08 | 0.04 |
| 794.00 | 777.00 | 826.00 | -1.31 | 0.00 |
| 5959.00 | 6620.00 | 6418.00 | -1.03 | 0.35 |
| 475.00 | 416.00 | 458.00 | -1.09 | 0.13 |
| 1909.00 | 1957.00 | 2008.00 | -1.06 | 0.12 |
| 656.00 | 706.00 | 686.67 | 1.13 | 0.01 |
| 433.00 | 474.00 | 479.00 | 1.40 | 0.00 |
| 1746.00 | 1979.00 | 1879.67 | 1.07 | 0.11 |
| 527.00 | 590.00 | 578.33 | 1.29 | 0.00 |
| 1016.00 | 1101.00 | 1075.67 | 1.34 | 0.00 |
| 1109.00 | 1165.00 | 1101.33 | 1.17 | 0.01 |
| 325.00 | 335.00 | 335.33 | 1.18 | 0.02 |
| 508.00 | 533.00 | 553.00 | 1.27 | 0.00 |
| 907.00 | 1072.00 | 1053.00 | 1.41 | 0.00 |
| 1587.00 | 1748.00 | 1711.00 | 1.36 | 0.00 |
| 1295.00 | 1588.00 | 1522.00 | 1.24 | 0.00 |
| 1460.00 | 1488.00 | 1531.67 | 1.11 | 0.01 |
| 944.00 | 990.00 | 1020.00 | 1.22 | 0.00 |
| 41.00 | 52.00 | 45.00 | -1.31 | 0.03 |
| 2.00 | 0.00 | 1.00 | -1.61 | 0.75 |
| 264.00 | 233.00 | 242.33 | -1.13 | 0.12 |
| 2557.00 | 2557.00 | 2538.67 | -2.01 | 0.00 |
| 8.00 | 12.00 | 8.67 | -1.24 | 0.53 |
| 86.00 | 85.00 | 79.67 | -1.03 | 0.87 |
| 3146.00 | 3524.00 | 3329.33 | -1.18 | 0.00 |
| 961.00 | 854.00 | 943.00 | -2.32 | 0.00 |
| 6.00 | 1.00 | 3.33 | -2.09 | 0.08 |
| 1346.00 | 1293.00 | 1315.33 | -2.52 | 0.00 |
| 28.00 | 21.00 | 26.33 | -1.78 | 0.00 |
| 132.00 | 148.00 | 130.67 | -1.45 | 0.00 |
| 1087.00 | 1009.00 | 1093.67 | -1.09 | 0.06 |
| 1713.00 | 2109.00 | 1933.33 | 1.27 | 0.00 |
| 2054.00 | 2195.00 | 2278.00 | -1.09 | 0.06 |
| 129.00 | 113.00 | 126.33 | -1.16 | 0.07 |
| 596.00 | 635.00 | 636.00 | -1.02 | 0.73 |
| 2437.00 | 2459.00 | 2356.33 | -1.07 | 0.21 |

| | | | | |
|---|---|---|---|---|
| 1700.00 | 1912.00 | 1796.67 | -1.12 | 0.01 |
| 435.00 | 447.00 | 446.33 | 1.13 | 0.03 |
| 12.00 | 13.00 | 10.67 | -2.01 | 0.00 |
| 2673.00 | 2655.00 | 2800.67 | -1.04 | 0.32 |
| 2009.00 | 2452.00 | 2238.33 | 1.12 | 0.02 |
| 44.00 | 39.00 | 40.00 | -1.03 | 0.86 |
| 657.00 | 685.00 | 674.67 | 1.02 | 0.65 |
| 850.00 | 1030.00 | 987.00 | 1.27 | 0.00 |
| 134.00 | 91.00 | 107.67 | 1.16 | 0.20 |
| 842.00 | 857.00 | 873.67 | 1.13 | 0.00 |
| 439.00 | 415.00 | 452.67 | -1.05 | 0.49 |
| 4101.00 | 4064.00 | 4270.67 | 1.14 | 0.00 |
| 364.00 | 421.00 | 431.00 | 1.08 | 0.27 |
| 196.00 | 183.00 | 193.67 | 1.09 | 0.22 |
| 9.00 | 5.00 | 6.33 | -1.48 | 0.26 |
| 1049.00 | 1225.00 | 1194.33 | -1.06 | 0.19 |
| 1317.00 | 1389.00 | 1377.67 | 1.01 | 0.76 |
| 5301.00 | 5889.00 | 5996.00 | -1.06 | 0.21 |
| 3693.00 | 4069.00 | 3833.00 | 1.10 | 0.06 |
| 7.00 | 4.00 | 4.33 | -2.07 | 0.04 |
| 400.00 | 510.00 | 466.33 | 1.17 | 0.01 |
| 851.00 | 875.00 | 913.33 | 1.05 | 0.26 |
| 415.00 | 415.00 | 402.67 | 1.07 | 0.29 |
| 71.00 | 85.00 | 80.33 | 1.32 | 0.01 |
| 21.00 | 19.00 | 18.00 | 1.44 | 0.11 |
| 77.00 | 78.00 | 82.00 | -1.10 | 0.31 |
| 2.00 | 6.00 | 5.67 | 1.19 | 0.73 |
| 22.00 | 8.00 | 12.67 | -1.69 | 0.03 |
| 156.00 | 187.00 | 187.00 | 1.37 | 0.00 |
| 529.00 | 638.00 | 572.33 | -1.20 | 0.00 |
| 3900.00 | 3989.00 | 4167.67 | 1.04 | 0.33 |
| 258.00 | 290.00 | 288.67 | 1.10 | 0.12 |
| 522.00 | 554.00 | 592.67 | 1.21 | 0.00 |
| 315.00 | 331.00 | 342.67 | -1.58 | 0.00 |
| 101.00 | 100.00 | 92.33 | -1.34 | 0.00 |
| 80.00 | 94.00 | 94.67 | 1.38 | 0.00 |
| 79.00 | 82.00 | 84.33 | 1.14 | 0.21 |
| 349.00 | 430.00 | 440.67 | 1.44 | 0.00 |
| 59.00 | 72.00 | 67.67 | 1.25 | 0.06 |
| 135.00 | 137.00 | 153.00 | 1.07 | 0.47 |
| 101.00 | 130.00 | 123.67 | 1.06 | 0.52 |
| 343.00 | 390.00 | 402.00 | 1.07 | 0.32 |
| 591.00 | 691.00 | 646.67 | -1.10 | 0.04 |
| 388.00 | 379.00 | 396.67 | -1.12 | 0.05 |
| 7.00 | 5.00 | 5.33 | 1.05 | 1.00 |
| 6.00 | 11.00 | 7.67 | -1.49 | 0.20 |
| 1398.00 | 1278.00 | 1337.67 | -1.19 | 0.00 |
| 628.00 | 653.00 | 664.00 | 1.29 | 0.00 |

| | | | | |
|---|---|---|---|---|
| 5556.00 | 5918.00 | 5888.33 | 1.06 | 0.09 |
| 838.00 | 914.00 | 918.67 | 1.02 | 0.65 |
| 8.00 | 12.00 | 12.33 | -1.26 | 0.34 |
| 353.00 | 383.00 | 402.33 | -1.49 | 0.00 |
| 2422.00 | 2301.00 | 2431.00 | 1.34 | 0.00 |
| 38.00 | 55.00 | 50.00 | 1.09 | 0.56 |
| 55.00 | 59.00 | 56.33 | -1.12 | 0.39 |
| 1721.00 | 1833.00 | 1831.00 | 1.07 | 0.10 |
| 1914.00 | 2125.00 | 2098.33 | 1.06 | 0.13 |
| 1011.00 | 1085.00 | 1076.67 | 1.17 | 0.00 |
| 899.00 | 943.00 | 953.00 | 1.13 | 0.01 |
| 7.00 | 12.00 | 9.00 | 1.16 | 0.68 |
| 15.00 | 14.00 | 16.33 | -1.26 | 0.31 |
| 225.00 | 250.00 | 238.33 | -1.08 | 0.26 |
| 19.00 | 23.00 | 20.33 | -1.33 | 0.13 |
| 3937.00 | 4189.00 | 4139.33 | -1.05 | 0.18 |
| 49.00 | 50.00 | 51.33 | -1.30 | 0.03 |
| 113.00 | 91.00 | 94.33 | -1.59 | 0.00 |
| 531.00 | 487.00 | 515.00 | -1.74 | 0.00 |
| 6.00 | 1.00 | 3.33 | -1.89 | 0.15 |
| 60.00 | 60.00 | 58.33 | -1.44 | 0.00 |
| 324.00 | 362.00 | 341.33 | 1.15 | 0.02 |
| 2576.00 | 2625.00 | 2750.33 | -1.03 | 0.51 |
| 510.00 | 496.00 | 518.67 | -1.09 | 0.07 |
| 452.00 | 395.00 | 429.33 | 1.00 | 0.98 |
| 321.00 | 409.00 | 363.00 | 1.13 | 0.05 |
| 937.00 | 1003.00 | 984.67 | 1.07 | 0.11 |
| 3917.00 | 4329.00 | 4223.33 | 1.06 | 0.09 |
| 414.00 | 523.00 | 469.33 | 1.27 | 0.00 |
| 116.00 | 157.00 | 143.67 | 1.31 | 0.00 |
| 662.00 | 786.00 | 729.00 | -1.12 | 0.02 |
| 431.00 | 598.00 | 522.33 | 1.28 | 0.00 |
| 329.00 | 374.00 | 358.67 | 1.19 | 0.00 |
| 126.00 | 132.00 | 139.67 | 1.38 | 0.00 |
| 892.00 | 1034.00 | 988.00 | 1.28 | 0.00 |
| 654.00 | 671.00 | 689.67 | 1.12 | 0.06 |
| 117.00 | 117.00 | 124.00 | 1.28 | 0.01 |
| 289.00 | 325.00 | 323.33 | 1.21 | 0.00 |
| 1077.00 | 1067.00 | 1147.00 | -1.03 | 0.61 |
| 592.00 | 730.00 | 683.33 | 1.66 | 0.00 |
| 319.00 | 354.00 | 340.00 | 1.16 | 0.01 |
| 362.00 | 445.00 | 442.33 | 1.27 | 0.00 |
| 909.00 | 957.00 | 936.33 | -1.07 | 0.10 |
| 109.00 | 105.00 | 108.33 | 1.35 | 0.00 |
| 995.00 | 1124.00 | 1075.67 | 1.08 | 0.06 |
| 768.00 | 881.00 | 862.00 | -1.01 | 0.75 |
| 246.00 | 268.00 | 259.00 | 1.01 | 0.92 |
| 759.00 | 792.00 | 771.33 | 1.01 | 0.83 |

| | | | | |
|---|---|---|---|---|
| 587.00 | 685.00 | 620.00 | 1.15 | 0.02 |
| 1350.00 | 1501.00 | 1410.67 | -1.18 | 0.00 |
| 326.00 | 341.00 | 360.67 | 1.14 | 0.03 |
| 2526.00 | 2717.00 | 2638.00 | 1.03 | 0.45 |
| 15.00 | 11.00 | 14.33 | 1.01 | 1.00 |
| 1812.00 | 1856.00 | 1835.67 | -1.16 | 0.00 |
| 16.00 | 17.00 | 18.67 | -1.19 | 0.38 |
| 64.00 | 54.00 | 59.33 | 1.06 | 0.65 |
| 1083.00 | 1094.00 | 1037.00 | 1.09 | 0.16 |
| 2899.00 | 2911.00 | 2890.33 | -1.34 | 0.00 |
| 1247.00 | 1463.00 | 1383.33 | -1.03 | 0.41 |
| 553.00 | 602.00 | 602.33 | -1.11 | 0.03 |
| 1112.00 | 1168.00 | 1119.33 | -1.07 | 0.14 |
| 1160.00 | 1306.00 | 1224.33 | 1.08 | 0.10 |
| 438.00 | 516.00 | 480.00 | 1.01 | 0.90 |
| 1119.00 | 1430.00 | 1294.33 | 1.43 | 0.00 |
| 3357.00 | 3520.00 | 3604.67 | 1.05 | 0.23 |
| 247.00 | 320.00 | 289.67 | 1.09 | 0.24 |
| 583.00 | 639.00 | 628.67 | 1.09 | 0.08 |
| 415.00 | 493.00 | 446.00 | 1.01 | 0.91 |
| 349.00 | 411.00 | 401.00 | -1.01 | 0.88 |
| 130.00 | 141.00 | 148.67 | -1.02 | 0.83 |
| 229.00 | 217.00 | 220.33 | 1.18 | 0.02 |
| 427.00 | 450.00 | 445.33 | 1.07 | 0.30 |
| 539.00 | 635.00 | 597.33 | 1.09 | 0.07 |
| 29.00 | 37.00 | 30.33 | 1.06 | 0.72 |
| 658.00 | 755.00 | 707.67 | -1.24 | 0.00 |
| 328.00 | 408.00 | 352.33 | -1.12 | 0.11 |
| 198.00 | 245.00 | 218.33 | -1.17 | 0.02 |
| 777.00 | 967.00 | 918.33 | -1.11 | 0.04 |
| 459.00 | 542.00 | 490.00 | 1.04 | 0.52 |
| 1079.00 | 1213.00 | 1181.67 | -1.05 | 0.22 |
| 4030.00 | 3817.00 | 4066.33 | 1.01 | 0.76 |
| 1749.00 | 1880.00 | 1843.67 | -1.12 | 0.00 |
| 13.00 | 28.00 | 24.00 | 1.11 | 0.70 |
| 5408.00 | 5381.00 | 5527.00 | 1.02 | 0.61 |
| 1408.00 | 1432.00 | 1489.67 | -1.01 | 0.85 |
| 2597.00 | 2608.00 | 2595.67 | 1.17 | 0.00 |
| 651.00 | 726.00 | 711.00 | -1.04 | 0.42 |
| 376.00 | 438.00 | 437.00 | -1.14 | 0.03 |
| 710.00 | 816.00 | 787.33 | 1.24 | 0.00 |
| 2.00 | 7.00 | 5.00 | -1.41 | 0.42 |
| 834.00 | 843.00 | 870.67 | 1.28 | 0.00 |
| 637.00 | 814.00 | 753.33 | -1.01 | 0.85 |
| 27.00 | 36.00 | 29.67 | 1.13 | 0.51 |
| 811.00 | 880.00 | 868.33 | -1.16 | 0.00 |
| 202.00 | 249.00 | 233.67 | -1.31 | 0.00 |
| 13.00 | 10.00 | 12.67 | -1.73 | 0.01 |

| | | | | |
|---|---|---|---|---|
| 809.00 | 903.00 | 867.67 | -1.32 | 0.00 |
| 71.00 | 74.00 | 76.33 | 1.22 | 0.07 |
| 49.00 | 54.00 | 48.67 | -1.32 | 0.02 |
| 175.00 | 205.00 | 197.67 | -1.25 | 0.00 |
| 62.00 | 57.00 | 64.67 | -1.46 | 0.00 |
| 4.00 | 3.00 | 4.33 | 1.16 | 0.84 |
| 1397.00 | 1544.00 | 1411.67 | 1.13 | 0.04 |
| 1035.00 | 1174.00 | 1135.67 | 1.20 | 0.00 |
| 484.00 | 531.00 | 532.33 | -1.05 | 0.33 |
| 17943.00 | 19702.00 | 20112.33 | 1.21 | 0.00 |
| 27.00 | 34.00 | 27.67 | -1.09 | 0.72 |
| 1119.00 | 1289.00 | 1233.67 | -1.12 | 0.02 |
| 1260.00 | 1338.00 | 1340.00 | -1.09 | 0.03 |
| 2.00 | 3.00 | 3.33 | -1.40 | 0.55 |
| 8.00 | 11.00 | 11.67 | 1.11 | 0.72 |
| 3.00 | 7.00 | 4.67 | -1.08 | 1.00 |
| 2261.00 | 2647.00 | 2524.67 | 1.00 | 0.92 |
| 94.00 | 95.00 | 92.33 | 1.01 | 0.97 |
| 24.00 | 23.00 | 24.67 | 1.20 | 0.36 |
| 133.00 | 169.00 | 152.33 | 1.04 | 0.64 |
| 203.00 | 193.00 | 207.33 | 1.06 | 0.42 |
| 17.00 | 29.00 | 24.00 | 1.13 | 0.63 |
| 8.00 | 5.00 | 5.67 | 1.39 | 0.49 |
| 1102.00 | 1087.00 | 1196.00 | -1.01 | 0.77 |
| 97.00 | 131.00 | 120.00 | 1.05 | 0.64 |
| 755.00 | 830.00 | 809.67 | 1.11 | 0.06 |
| 250.00 | 326.00 | 304.00 | 1.47 | 0.00 |
| 720.00 | 715.00 | 714.00 | -1.01 | 0.92 |
| 641.00 | 723.00 | 752.67 | 1.12 | 0.10 |
| 4125.00 | 4744.00 | 4749.67 | 1.15 | 0.00 |
| 1921.00 | 2165.00 | 2113.00 | 1.23 | 0.00 |
| 484.00 | 455.00 | 504.33 | 1.21 | 0.00 |
| 187.00 | 235.00 | 243.33 | 1.20 | 0.06 |
| 266.00 | 314.00 | 304.67 | 1.15 | 0.04 |
| 656.00 | 705.00 | 699.33 | 1.07 | 0.18 |
| 1887.00 | 2070.00 | 1943.33 | -1.05 | 0.34 |
| 2043.00 | 2178.00 | 2194.00 | 1.05 | 0.14 |
| 2719.00 | 2774.00 | 2751.67 | -1.28 | 0.00 |
| 5689.00 | 5728.00 | 5844.00 | -1.01 | 0.83 |
| 6544.00 | 6713.00 | 6582.67 | -1.18 | 0.00 |
| 7.00 | 8.00 | 7.33 | -1.15 | 0.78 |
| 1426.00 | 1462.00 | 1428.67 | -1.32 | 0.00 |
| 1880.00 | 1957.00 | 1870.33 | -1.01 | 0.88 |
| 2170.00 | 2349.00 | 2225.67 | -1.17 | 0.00 |
| 1460.00 | 1664.00 | 1550.67 | -1.11 | 0.02 |
| 412.00 | 516.00 | 471.33 | 1.43 | 0.00 |
| 623.00 | 637.00 | 669.00 | 1.33 | 0.00 |
| 1869.00 | 1820.00 | 1850.67 | -1.03 | 0.43 |

| | | | | |
|---|---|---|---|---|
| 455.00 | 492.00 | 527.00 | 1.11 | 0.17 |
| 41.00 | 41.00 | 42.00 | 2.14 | 0.00 |
| 428.00 | 534.00 | 487.00 | -1.03 | 0.64 |
| 260.00 | 303.00 | 303.00 | -1.00 | 1.00 |
| 1747.00 | 1768.00 | 1821.00 | -1.01 | 0.83 |
| 697.00 | 653.00 | 688.67 | -1.04 | 0.44 |
| 14.00 | 11.00 | 15.00 | -1.17 | 0.56 |
| 16.00 | 35.00 | 24.00 | -1.18 | 0.35 |
| 28.00 | 17.00 | 22.33 | -1.07 | 0.81 |
| 7.00 | 13.00 | 8.33 | 1.53 | 0.22 |
| 1099.00 | 1278.00 | 1149.00 | 1.13 | 0.04 |
| 767.00 | 939.00 | 846.00 | -1.10 | 0.07 |
| 1674.00 | 1636.00 | 1748.67 | 1.03 | 0.43 |
| 1135.00 | 1155.00 | 1227.00 | -1.00 | 0.96 |
| 325.00 | 350.00 | 332.00 | -1.01 | 0.96 |
| 982.00 | 1100.00 | 1084.00 | 1.12 | 0.01 |
| 644.00 | 812.00 | 754.67 | 1.08 | 0.15 |
| 97.00 | 99.00 | 102.67 | 1.22 | 0.04 |
| 649.00 | 709.00 | 713.67 | 1.09 | 0.08 |
| 1751.00 | 1823.00 | 1847.33 | -1.04 | 0.31 |
| 1498.00 | 1712.00 | 1693.00 | 1.23 | 0.00 |
| 244.00 | 221.00 | 235.00 | -1.11 | 0.18 |
| 827.00 | 893.00 | 906.00 | 1.14 | 0.00 |
| 256.00 | 254.00 | 256.67 | -1.33 | 0.00 |
| 28.00 | 44.00 | 33.33 | -1.01 | 0.95 |
| 2975.00 | 3817.00 | 3453.00 | 1.12 | 0.03 |
| 939.00 | 906.00 | 937.67 | -1.14 | 0.00 |
| 3707.00 | 3530.00 | 3700.33 | -1.34 | 0.00 |
| 2979.00 | 2982.00 | 2983.00 | -1.36 | 0.00 |
| 902.00 | 956.00 | 978.67 | 1.14 | 0.00 |
| 1253.00 | 1441.00 | 1357.67 | 1.14 | 0.00 |
| 2036.00 | 2282.00 | 2197.33 | 1.00 | 0.92 |
| 3.00 | 4.00 | 3.33 | -1.21 | 0.83 |
| 26.00 | 25.00 | 24.00 | -1.10 | 0.71 |
| 40.00 | 62.00 | 51.67 | -1.65 | 0.00 |
| 967.00 | 1055.00 | 1032.67 | -1.14 | 0.00 |
| 40.00 | 41.00 | 46.00 | -1.33 | 0.02 |
| 207.00 | 228.00 | 212.33 | 1.42 | 0.00 |
| 4.00 | 1.00 | 2.33 | -2.25 | 0.10 |
| 38.00 | 35.00 | 40.33 | -1.11 | 0.48 |
| 16.00 | 14.00 | 19.33 | -1.47 | 0.04 |
| 886.00 | 1013.00 | 974.67 | 1.09 | 0.05 |
| 2115.00 | 2112.00 | 2102.33 | -1.05 | 0.31 |
| 251.00 | 244.00 | 253.67 | -1.41 | 0.00 |
| 931.00 | 938.00 | 945.67 | -1.16 | 0.00 |
| 573.00 | 602.00 | 592.67 | -1.19 | 0.00 |
| 41.00 | 38.00 | 35.00 | 1.16 | 0.40 |
| 3.00 | 10.00 | 7.33 | 1.44 | 0.34 |

| | | | | |
|---|---|---|---|---|
| 227.00 | 217.00 | 214.00 | 1.20 | 0.02 |
| 883.00 | 1085.00 | 988.00 | 1.02 | 0.67 |
| 526.00 | 520.00 | 520.00 | 1.01 | 0.89 |
| 1384.00 | 1376.00 | 1429.33 | 1.01 | 0.82 |
| 1905.00 | 1934.00 | 2025.67 | -1.05 | 0.20 |
| 2404.00 | 2750.00 | 2669.67 | 1.09 | 0.02 |
| 745.00 | 803.00 | 787.00 | -1.14 | 0.00 |
| 683.00 | 730.00 | 725.67 | 1.06 | 0.23 |
| 2499.00 | 2644.00 | 2702.00 | 1.02 | 0.54 |
| 1133.00 | 1253.00 | 1271.67 | 1.07 | 0.13 |
| 291.00 | 344.00 | 350.00 | 1.48 | 0.00 |
| 1174.00 | 1191.00 | 1236.00 | -1.01 | 0.76 |
| 1216.00 | 1109.00 | 1173.00 | -1.27 | 0.00 |
| 2.00 | 5.00 | 6.00 | 1.59 | 0.28 |
| 110.00 | 147.00 | 137.33 | 1.11 | 0.27 |
| 84.00 | 59.00 | 70.67 | 1.84 | 0.00 |
| 0.00 | 0.00 | 0.00 | -11.76 | 0.14 |
| 6.00 | 5.00 | 7.33 | -1.06 | 1.00 |
| 315.00 | 346.00 | 331.67 | 1.21 | 0.00 |
| 786.00 | 817.00 | 841.33 | 1.13 | 0.01 |
| 527.00 | 569.00 | 565.67 | 1.04 | 0.48 |
| 4479.00 | 4749.00 | 4891.00 | 1.02 | 0.68 |
| 31.00 | 16.00 | 28.33 | 1.37 | 0.10 |
| 466.00 | 456.00 | 462.33 | -1.07 | 0.23 |
| 727.00 | 765.00 | 769.00 | 1.14 | 0.01 |
| 492.00 | 526.00 | 497.67 | 1.26 | 0.00 |
| 4.00 | 6.00 | 6.33 | 1.17 | 0.74 |
| 1652.00 | 1792.00 | 1697.00 | 1.06 | 0.15 |
| 3.00 | 7.00 | 6.67 | -1.16 | 0.77 |
| 1702.00 | 1620.00 | 1676.33 | 1.29 | 0.00 |
| 51.00 | 38.00 | 46.33 | 1.78 | 0.00 |
| 3358.00 | 3163.00 | 3305.67 | -1.09 | 0.03 |
| 1292.00 | 1543.00 | 1399.33 | 1.28 | 0.00 |
| 1175.00 | 1227.00 | 1220.33 | 1.10 | 0.02 |
| 8242.00 | 8688.00 | 8576.00 | -1.19 | 0.00 |
| 4696.00 | 4982.00 | 4781.33 | -1.04 | 0.35 |
| 8.00 | 13.00 | 8.67 | -2.01 | 0.01 |
| 260.00 | 288.00 | 286.33 | 1.19 | 0.02 |
| 315.00 | 321.00 | 322.33 | 1.05 | 0.46 |
| 788.00 | 884.00 | 886.33 | 1.39 | 0.00 |
| 847.00 | 950.00 | 1014.67 | 1.45 | 0.00 |
| 1567.00 | 1558.00 | 1555.67 | -1.09 | 0.04 |
| 1138.00 | 1223.00 | 1152.67 | 1.02 | 0.69 |
| 1159.00 | 1153.00 | 1189.67 | -1.08 | 0.10 |
| 14.00 | 17.00 | 17.00 | 2.62 | 0.00 |
| 1582.00 | 1740.00 | 1713.33 | 1.12 | 0.01 |
| 148.00 | 137.00 | 152.67 | -2.18 | 0.00 |
| 259.00 | 246.00 | 264.67 | -1.21 | 0.01 |

| | | | | |
|---|---|---|---|---|
| 1854.00 | 1909.00 | 1884.00 | -1.09 | 0.02 |
| 323.00 | 336.00 | 330.00 | -1.47 | 0.00 |
| 554.00 | 551.00 | 558.67 | -1.05 | 0.35 |
| 1124.00 | 987.00 | 1050.67 | -1.30 | 0.00 |
| 4276.00 | 3872.00 | 4245.00 | -1.15 | 0.00 |
| 813.00 | 924.00 | 919.00 | 2.30 | 0.00 |
| 712.00 | 767.00 | 755.33 | 1.15 | 0.01 |
| 3629.00 | 4331.00 | 3983.33 | 1.11 | 0.01 |
| 105.00 | 73.00 | 100.00 | 1.13 | 0.29 |
| 837.00 | 865.00 | 882.33 | -1.37 | 0.00 |
| 5.00 | 6.00 | 7.00 | 2.52 | 0.03 |
| 10.00 | 7.00 | 6.67 | -1.51 | 0.23 |
| 13.00 | 25.00 | 18.00 | 1.52 | 0.07 |
| 8.00 | 10.00 | 9.33 | -1.01 | 1.00 |
| 1.00 | 3.00 | 2.33 | -2.11 | 0.14 |
| 2414.00 | 2576.00 | 2512.00 | 1.04 | 0.31 |
| 6.00 | 5.00 | 7.00 | -1.11 | 0.88 |
| 3333.00 | 3394.00 | 3496.67 | 1.16 | 0.00 |
| 13.00 | 20.00 | 15.00 | 1.38 | 0.20 |
| 693.00 | 604.00 | 666.33 | -1.17 | 0.00 |
| 3.00 | 5.00 | 2.67 | -2.83 | 0.01 |
| 3.00 | 0.00 | 1.67 | -2.33 | 0.15 |
| 280.00 | 264.00 | 289.33 | 1.04 | 0.58 |
| 640.00 | 805.00 | 750.00 | -1.26 | 0.00 |
| 6.00 | 17.00 | 15.33 | 1.36 | 0.26 |
| 27.00 | 29.00 | 31.67 | 2.45 | 0.00 |
| 1.00 | 0.00 | 1.67 | -1.58 | 0.59 |
| 3.00 | 4.00 | 2.67 | -1.13 | 1.00 |
| 726.00 | 752.00 | 767.67 | 1.18 | 0.00 |
| 330.00 | 445.00 | 407.00 | -1.08 | 0.26 |
| 5.00 | 3.00 | 3.33 | -1.50 | 0.44 |
| 24.00 | 19.00 | 24.00 | 1.03 | 0.94 |
| 6981.00 | 7263.00 | 7100.00 | 1.18 | 0.00 |
| 2052.00 | 2271.00 | 2183.67 | -1.02 | 0.62 |
| 2295.00 | 2392.00 | 2264.67 | -1.08 | 0.15 |
| 1019.00 | 986.00 | 992.00 | -1.41 | 0.00 |
| 52.00 | 36.00 | 42.00 | -1.79 | 0.00 |
| 2506.00 | 2838.00 | 2627.67 | 1.02 | 0.64 |
| 2036.00 | 2369.00 | 2289.00 | -1.02 | 0.68 |
| 0.00 | 2.00 | 0.67 | -3.57 | 0.12 |
| 12.00 | 9.00 | 12.67 | -1.01 | 1.00 |
| 1010.00 | 1229.00 | 1150.00 | -1.06 | 0.20 |
| 368.00 | 457.00 | 443.67 | -1.08 | 0.21 |
| 1104.00 | 1203.00 | 1178.00 | -1.46 | 0.00 |
| 2528.00 | 2372.00 | 2362.33 | -1.02 | 0.75 |
| 1005.00 | 951.00 | 1018.67 | -1.12 | 0.02 |
| 926.00 | 994.00 | 973.33 | 1.06 | 0.23 |
| 1015.00 | 971.00 | 995.67 | 1.09 | 0.05 |

| | | | | |
|---|---|---|---|---|
| 2051.00 | 2359.00 | 2367.33 | 1.11 | 0.02 |
| 610.00 | 640.00 | 617.67 | -1.06 | 0.29 |
| 297.00 | 324.00 | 347.00 | 1.05 | 0.57 |
| 506.00 | 493.00 | 490.33 | 1.00 | 0.97 |
| 780.00 | 731.00 | 802.67 | 1.18 | 0.00 |
| 2296.00 | 2312.00 | 2339.33 | -1.08 | 0.07 |
| 861.00 | 927.00 | 948.00 | 1.05 | 0.31 |
| 732.00 | 850.00 | 813.33 | 1.05 | 0.33 |
| 411.00 | 407.00 | 413.67 | 1.42 | 0.00 |
| 5225.00 | 5615.00 | 5415.00 | -1.20 | 0.00 |
| 7.00 | 4.00 | 6.33 | -1.51 | 0.38 |
| 810.00 | 942.00 | 936.67 | -1.35 | 0.00 |
| 2762.00 | 2986.00 | 3059.33 | 1.12 | 0.01 |
| 618.00 | 622.00 | 672.33 | 1.08 | 0.18 |
| 13028.00 | 13686.00 | 13339.67 | 1.00 | 0.93 |
| 217.00 | 226.00 | 223.33 | -1.76 | 0.00 |
| 66.00 | 83.00 | 78.00 | -1.39 | 0.00 |
| 559.00 | 627.00 | 612.67 | 1.15 | 0.00 |
| 4657.00 | 4570.00 | 4726.33 | -1.07 | 0.06 |
| 11.00 | 8.00 | 8.67 | -1.13 | 0.79 |
| 34.00 | 36.00 | 35.67 | -1.15 | 0.34 |
| 65.00 | 61.00 | 61.33 | 1.05 | 0.70 |
| 22.00 | 19.00 | 21.33 | -1.01 | 1.00 |
| 1110.00 | 1375.00 | 1267.33 | 1.15 | 0.00 |
| 93.00 | 91.00 | 99.00 | 1.36 | 0.00 |
| 552.00 | 609.00 | 621.00 | 1.19 | 0.00 |
| 534.00 | 576.00 | 588.33 | 1.28 | 0.00 |
| 76.00 | 108.00 | 105.67 | 1.08 | 0.49 |
| 2320.00 | 2465.00 | 2435.00 | 1.14 | 0.00 |
| 1844.00 | 2054.00 | 2035.67 | 1.12 | 0.01 |
| 1738.00 | 2038.00 | 1970.33 | 1.23 | 0.00 |
| 54.00 | 25.00 | 43.00 | 1.06 | 0.77 |
| 958.00 | 861.00 | 943.00 | -1.14 | 0.01 |
| 947.00 | 875.00 | 897.33 | -1.16 | 0.01 |
| 2558.00 | 2596.00 | 2573.33 | 1.06 | 0.17 |
| 1339.00 | 1364.00 | 1454.00 | -1.05 | 0.35 |
| 37.00 | 42.00 | 40.67 | 1.00 | 1.00 |
| 1873.00 | 1927.00 | 1938.67 | -1.16 | 0.00 |
| 1147.00 | 1389.00 | 1263.33 | -1.03 | 0.53 |
| 12.00 | 17.00 | 13.33 | -1.49 | 0.08 |
| 8076.00 | 8802.00 | 8978.33 | 1.12 | 0.00 |
| 1896.00 | 1966.00 | 1994.67 | 1.01 | 0.71 |
| 347.00 | 436.00 | 409.00 | 1.09 | 0.14 |
| 18.00 | 38.00 | 26.33 | -1.68 | 0.00 |
| 22.00 | 12.00 | 17.33 | -1.28 | 0.22 |
| 2283.00 | 2660.00 | 2543.00 | 1.21 | 0.00 |
| 10.00 | 12.00 | 10.67 | -1.17 | 0.64 |
| 183.00 | 222.00 | 201.67 | -1.49 | 0.00 |

| | | | | |
|---|---|---|---|---|
| 1085.00 | 1158.00 | 1191.67 | 1.18 | 0.00 |
| 60.00 | 41.00 | 50.67 | -1.37 | 0.02 |
| 75.00 | 71.00 | 72.33 | 1.15 | 0.21 |
| 202.00 | 273.00 | 231.33 | 1.08 | 0.32 |
| 0.00 | 4.00 | 1.67 | -2.52 | 0.10 |
| 799.00 | 775.00 | 808.33 | -1.05 | 0.25 |
| 4908.00 | 5809.00 | 5511.00 | 1.38 | 0.00 |
| 136.00 | 179.00 | 157.67 | -1.14 | 0.12 |
| 354.00 | 283.00 | 319.67 | -1.10 | 0.18 |
| 646.00 | 672.00 | 675.67 | -1.14 | 0.00 |
| 551.00 | 501.00 | 520.67 | -1.09 | 0.13 |
| 5020.00 | 4800.00 | 4848.00 | -1.08 | 0.10 |
| 791.00 | 826.00 | 849.00 | 1.12 | 0.01 |
| 1580.00 | 1774.00 | 1702.67 | 1.07 | 0.10 |
| 1333.00 | 1461.00 | 1441.00 | 1.03 | 0.42 |
| 61.00 | 61.00 | 59.33 | -1.35 | 0.01 |
| 571.00 | 589.00 | 610.00 | -1.10 | 0.06 |
| 1177.00 | 1280.00 | 1239.00 | 1.09 | 0.07 |
| 398.00 | 461.00 | 426.67 | 1.04 | 0.49 |
| 1070.00 | 1306.00 | 1226.33 | 1.10 | 0.05 |
| 685.00 | 750.00 | 737.67 | 1.04 | 0.42 |
| 957.00 | 992.00 | 1028.67 | 1.03 | 0.45 |
| 1207.00 | 1339.00 | 1338.67 | 1.12 | 0.01 |
| 2618.00 | 2784.00 | 2820.33 | 1.05 | 0.21 |
| 1395.00 | 1390.00 | 1434.33 | 1.07 | 0.11 |
| 230.00 | 227.00 | 229.67 | -1.11 | 0.11 |
| 16.00 | 9.00 | 10.00 | -1.98 | 0.01 |
| 1196.00 | 1306.00 | 1302.00 | 1.24 | 0.00 |
| 1681.00 | 1693.00 | 1702.33 | 1.24 | 0.00 |
| 1.00 | 6.00 | 3.00 | 1.26 | 0.81 |
| 372.00 | 461.00 | 424.00 | -1.03 | 0.66 |
| 38.00 | 51.00 | 41.00 | 1.50 | 0.01 |
| 68.00 | 73.00 | 69.33 | -1.61 | 0.00 |
| 1570.00 | 1649.00 | 1642.00 | -1.02 | 0.65 |
| 101.00 | 114.00 | 104.67 | -1.37 | 0.00 |
| 133.00 | 156.00 | 153.33 | -1.29 | 0.00 |
| 48.00 | 62.00 | 53.00 | -1.46 | 0.00 |
| 606.00 | 752.00 | 700.67 | 1.23 | 0.00 |
| 1702.00 | 1770.00 | 1787.67 | -1.02 | 0.62 |
| 2173.00 | 2454.00 | 2415.67 | -1.01 | 0.72 |
| 243.00 | 262.00 | 290.67 | 1.31 | 0.00 |
| 814.00 | 850.00 | 911.00 | 1.10 | 0.07 |
| 527.00 | 621.00 | 631.00 | -1.01 | 0.86 |
| 750.00 | 871.00 | 881.67 | 1.17 | 0.00 |
| 659.00 | 635.00 | 657.00 | 1.04 | 0.44 |
| 2190.00 | 2265.00 | 2366.67 | -1.10 | 0.02 |
| 175.00 | 169.00 | 167.33 | 1.09 | 0.27 |
| 25.00 | 25.00 | 24.00 | -1.03 | 1.00 |

| | | | | |
|---|---|---|---|---|
| 652.00 | 703.00 | 676.00 | -1.29 | 0.00 |
| 624.00 | 683.00 | 676.67 | 1.01 | 0.86 |
| 771.00 | 874.00 | 856.00 | -1.15 | 0.00 |
| 1509.00 | 1650.00 | 1595.33 | -1.03 | 0.52 |
| 1520.00 | 1615.00 | 1599.33 | 1.06 | 0.17 |
| 502.00 | 637.00 | 575.33 | 1.05 | 0.46 |
| 487.00 | 539.00 | 534.00 | 1.01 | 0.84 |
| 390.00 | 366.00 | 368.67 | -1.12 | 0.09 |
| 680.00 | 705.00 | 706.67 | 1.10 | 0.05 |
| 16.00 | 21.00 | 19.33 | 1.78 | 0.01 |
| 6.00 | 11.00 | 6.67 | -1.51 | 0.22 |
| 3157.00 | 3156.00 | 2995.67 | -2.33 | 0.00 |
| 4153.00 | 4586.00 | 4422.33 | -1.74 | 0.00 |
| 978.00 | 992.00 | 1056.67 | -1.02 | 0.64 |
| 174.00 | 168.00 | 167.33 | -1.27 | 0.00 |
| 3085.00 | 3171.00 | 3139.67 | -1.05 | 0.24 |
| 28.00 | 27.00 | 26.67 | -1.19 | 0.31 |
| 11878.00 | 12840.00 | 12523.67 | -1.80 | 0.00 |
| 2.00 | 2.00 | 1.67 | -1.58 | 0.61 |
| 3.00 | 5.00 | 3.67 | -4.66 | 0.00 |
| 5.00 | 4.00 | 4.00 | -1.51 | 0.37 |
| 67.00 | 68.00 | 69.00 | 1.13 | 0.26 |
| 3749.00 | 3811.00 | 3917.00 | -1.07 | 0.04 |
| 3.00 | 0.00 | 1.33 | -3.79 | 0.01 |
| 2032.00 | 2147.00 | 2049.00 | -1.45 | 0.00 |
| 4759.00 | 4762.00 | 4644.33 | -1.26 | 0.00 |
| 0.00 | 1.00 | 0.33 | -3.95 | 0.23 |
| 597.00 | 585.00 | 561.00 | 1.02 | 0.75 |
| 4.00 | 9.00 | 7.00 | -1.39 | 0.34 |
| 1783.00 | 1997.00 | 1915.67 | -1.34 | 0.00 |
| 1120.00 | 1070.00 | 1089.00 | -1.50 | 0.00 |
| 1752.00 | 1709.00 | 1684.33 | -2.32 | 0.00 |
| 2010.00 | 2205.00 | 2107.33 | 1.08 | 0.06 |
| 17556.00 | 17873.00 | 17879.00 | 1.04 | 0.26 |
| 32751.00 | 33591.00 | 33660.67 | -1.03 | 0.42 |
| 8976.00 | 9206.00 | 9081.00 | -1.73 | 0.00 |
| 206.00 | 219.00 | 214.67 | -1.64 | 0.00 |
| 13.00 | 19.00 | 13.33 | -1.89 | 0.00 |
| 40.00 | 44.00 | 44.33 | 1.01 | 1.00 |
| 147.00 | 142.00 | 148.67 | 1.12 | 0.16 |
| 13515.00 | 13854.00 | 14053.33 | 1.03 | 0.37 |
| 2589.00 | 2429.00 | 2555.67 | -1.05 | 0.26 |
| 415.00 | 403.00 | 400.33 | 1.12 | 0.07 |
| 17.00 | 28.00 | 24.00 | -1.69 | 0.00 |
| 185.00 | 218.00 | 205.33 | 1.04 | 0.58 |
| 316.00 | 349.00 | 356.33 | 1.02 | 0.80 |
| 546.00 | 617.00 | 610.67 | 1.03 | 0.59 |
| 355.00 | 348.00 | 380.67 | -1.36 | 0.00 |

| | | | | |
|---|---|---|---|---|
| 1131.00 | 1322.00 | 1345.33 | 1.11 | 0.05 |
| 415.00 | 498.00 | 474.00 | -1.01 | 0.87 |
| 966.00 | 1182.00 | 1157.67 | 1.14 | 0.02 |
| 588.00 | 600.00 | 599.67 | -1.05 | 0.28 |
| 311.00 | 385.00 | 361.67 | 1.24 | 0.00 |
| 340.00 | 381.00 | 405.33 | 1.06 | 0.45 |
| 4224.00 | 4139.00 | 4370.00 | 1.08 | 0.04 |
| 212.00 | 184.00 | 198.33 | 1.13 | 0.10 |
| 7107.00 | 6834.00 | 7026.00 | -1.13 | 0.00 |
| 4907.00 | 5527.00 | 5309.00 | 1.03 | 0.33 |
| 4382.00 | 4668.00 | 4533.33 | 1.00 | 0.95 |
| 1818.00 | 1900.00 | 1979.67 | 1.05 | 0.23 |
| 3664.00 | 4031.00 | 3947.67 | -1.05 | 0.13 |
| 1626.00 | 1766.00 | 1724.67 | 1.08 | 0.04 |
| 856.00 | 866.00 | 922.67 | 1.34 | 0.00 |
| 1472.00 | 1656.00 | 1590.00 | 1.16 | 0.00 |
| 1434.00 | 1596.00 | 1632.33 | 1.20 | 0.00 |
| 498.00 | 580.00 | 556.33 | 1.05 | 0.38 |
| 1501.00 | 1807.00 | 1760.00 | 1.13 | 0.01 |
| 2606.00 | 2745.00 | 2814.33 | 1.20 | 0.00 |
| 1799.00 | 1773.00 | 1899.33 | -1.03 | 0.49 |
| 997.00 | 1103.00 | 1070.00 | 1.01 | 0.88 |
| 1202.00 | 1355.00 | 1357.00 | 1.03 | 0.55 |
| 1970.00 | 2079.00 | 2139.67 | 1.07 | 0.11 |
| 284.00 | 298.00 | 305.33 | -1.08 | 0.18 |
| 104.00 | 111.00 | 112.67 | 1.10 | 0.29 |
| 567.00 | 694.00 | 676.00 | 1.16 | 0.01 |
| 387.00 | 357.00 | 390.33 | 1.23 | 0.00 |
| 159.00 | 184.00 | 193.00 | 1.38 | 0.00 |
| 905.00 | 896.00 | 970.00 | -1.02 | 0.70 |
| 548.00 | 657.00 | 640.67 | 1.15 | 0.02 |
| 374.00 | 420.00 | 423.67 | -1.03 | 0.71 |
| 269.00 | 266.00 | 288.00 | 1.05 | 0.41 |
| 993.00 | 1129.00 | 1097.00 | 1.03 | 0.41 |
| 4.00 | 10.00 | 8.33 | -1.18 | 0.70 |
| 28.00 | 16.00 | 23.00 | -1.12 | 0.59 |
| 2235.00 | 2253.00 | 2334.67 | 1.03 | 0.43 |
| 5288.00 | 5717.00 | 5513.67 | 1.10 | 0.01 |
| 36.00 | 38.00 | 40.00 | 1.20 | 0.22 |
| 240.00 | 274.00 | 252.00 | -2.18 | 0.00 |
| 27.00 | 22.00 | 22.33 | -1.03 | 1.00 |
| 8321.00 | 8801.00 | 9061.67 | 1.29 | 0.00 |
| 325.00 | 280.00 | 306.33 | 1.09 | 0.19 |
| 280.00 | 249.00 | 253.67 | -1.09 | 0.25 |
| 636.00 | 770.00 | 714.67 | 1.02 | 0.76 |
| 197.00 | 180.00 | 210.33 | 1.11 | 0.28 |
| 1079.00 | 1222.00 | 1192.33 | 1.07 | 0.09 |
| 152.00 | 216.00 | 202.33 | 1.22 | 0.03 |

| | | | | |
|---|---|---|---|---|
| 347.00 | 430.00 | 423.67 | 1.25 | 0.00 |
| 257.00 | 266.00 | 268.33 | 1.06 | 0.39 |
| 628.00 | 637.00 | 656.67 | 1.01 | 0.78 |
| 418.00 | 465.00 | 454.33 | 1.07 | 0.19 |
| 5866.00 | 6390.00 | 6435.33 | 1.18 | 0.00 |
| 1548.00 | 1697.00 | 1703.67 | 1.24 | 0.00 |
| 1885.00 | 2123.00 | 2141.00 | 1.18 | 0.00 |
| 1141.00 | 1532.00 | 1394.67 | -1.89 | 0.00 |
| 1034.00 | 1132.00 | 1198.33 | 1.11 | 0.05 |
| 0.00 | 0.00 | 0.33 | -3.22 | 0.38 |
| 2004.00 | 2174.00 | 2225.67 | 1.21 | 0.00 |
| 1421.00 | 1559.00 | 1670.33 | 1.16 | 0.01 |
| 1869.00 | 1961.00 | 2066.67 | 1.10 | 0.04 |
| 1446.00 | 1669.00 | 1702.00 | 1.23 | 0.00 |
| 3560.00 | 4008.00 | 4136.00 | 1.14 | 0.01 |
| 3331.00 | 3701.00 | 3772.33 | 1.16 | 0.00 |
| 10087.00 | 10483.00 | 10495.67 | -2.52 | 0.00 |
| 4.00 | 4.00 | 3.67 | -3.33 | 0.00 |
| 2.00 | 3.00 | 3.00 | -1.55 | 0.41 |
| 15.00 | 9.00 | 10.33 | 1.13 | 0.70 |
| 2601.00 | 2681.00 | 2600.00 | -1.18 | 0.00 |
| 32.00 | 28.00 | 35.67 | 1.01 | 1.00 |
| 14.00 | 13.00 | 12.67 | -1.33 | 0.26 |
| 7.00 | 2.00 | 5.00 | -1.34 | 0.52 |
| 279.00 | 352.00 | 309.00 | -1.05 | 0.61 |
| 1.00 | 1.00 | 0.67 | -4.00 | 0.07 |
| 39.00 | 47.00 | 40.67 | 1.35 | 0.05 |
| 265.00 | 280.00 | 256.33 | -1.20 | 0.02 |
| 1511.00 | 1452.00 | 1419.67 | 1.40 | 0.00 |
| 1.00 | 4.00 | 3.33 | 1.09 | 1.00 |
| 257.00 | 236.00 | 237.00 | -1.29 | 0.00 |
| 4679.00 | 4740.00 | 4982.00 | 1.00 | 0.95 |
| 383.00 | 445.00 | 411.67 | 1.32 | 0.00 |
| 2154.00 | 2360.00 | 2196.67 | -1.02 | 0.66 |
| 66.00 | 59.00 | 64.00 | 1.50 | 0.00 |
| 17.00 | 26.00 | 20.67 | -1.35 | 0.09 |
| 32.00 | 32.00 | 33.00 | -2.24 | 0.00 |
| 797.00 | 982.00 | 918.67 | 1.21 | 0.00 |
| 1070.00 | 1174.00 | 1120.00 | 1.09 | 0.05 |
| 355.00 | 352.00 | 346.33 | -1.16 | 0.01 |
| 682.00 | 706.00 | 694.33 | -1.04 | 0.47 |
| 3274.00 | 3226.00 | 3343.33 | -1.03 | 0.40 |
| 2378.00 | 2500.00 | 2459.67 | 1.11 | 0.01 |
| 1245.00 | 1413.00 | 1389.33 | 1.16 | 0.00 |
| 2324.00 | 2161.00 | 2323.67 | -1.03 | 0.54 |
| 1068.00 | 1090.00 | 1135.33 | 1.12 | 0.01 |
| 0.00 | 1.00 | 0.67 | -3.14 | 0.19 |
| 3049.00 | 3877.00 | 3462.00 | 1.07 | 0.19 |

| | | | | |
|---|---|---|---|---|
| 2356.00 | 2462.00 | 2473.67 | -1.11 | 0.00 |
| 55.00 | 64.00 | 60.00 | 1.54 | 0.00 |
| 55.00 | 68.00 | 58.67 | -1.23 | 0.08 |
| 31.00 | 43.00 | 36.67 | 1.12 | 0.53 |
| 521.00 | 553.00 | 556.67 | 1.21 | 0.00 |
| 492.00 | 462.00 | 523.33 | 1.18 | 0.01 |
| 515.00 | 500.00 | 533.00 | -1.27 | 0.00 |
| 9.00 | 14.00 | 11.67 | -1.21 | 0.51 |
| 1.00 | 6.00 | 9.67 | 1.63 | 0.31 |
| 433.00 | 362.00 | 359.00 | -1.31 | 0.01 |
| 363.00 | 366.00 | 368.00 | 1.29 | 0.00 |
| 1515.00 | 1493.00 | 1500.33 | -1.08 | 0.07 |
| 549.00 | 535.00 | 548.67 | -1.16 | 0.00 |
| 536.00 | 609.00 | 598.67 | -1.02 | 0.70 |
| 985.00 | 943.00 | 952.00 | 1.06 | 0.26 |
| 1423.00 | 1652.00 | 1525.00 | 1.11 | 0.03 |
| 592.00 | 555.00 | 588.67 | -1.08 | 0.13 |
| 379.00 | 349.00 | 367.67 | -2.26 | 0.00 |
| 8769.00 | 7668.00 | 8137.33 | -1.07 | 0.19 |
| 420.00 | 389.00 | 428.33 | 1.02 | 0.79 |
| 878.00 | 894.00 | 872.00 | -1.27 | 0.00 |
| 1900.00 | 1754.00 | 1784.67 | -1.17 | 0.00 |
| 762.00 | 925.00 | 860.33 | 1.10 | 0.06 |
| 596.00 | 561.00 | 549.33 | -1.12 | 0.17 |
| 2134.00 | 2757.00 | 2385.33 | -1.06 | 0.29 |
| 355.00 | 397.00 | 382.00 | 1.22 | 0.00 |
| 14.00 | 11.00 | 15.33 | 2.49 | 0.00 |
| 608.00 | 607.00 | 630.67 | -1.21 | 0.00 |
| 644.00 | 650.00 | 676.00 | -1.04 | 0.45 |
| 667.00 | 834.00 | 789.33 | 1.09 | 0.12 |
| 7.00 | 1.00 | 3.33 | -1.11 | 1.00 |
| 58.00 | 48.00 | 49.00 | -1.31 | 0.14 |
| 5989.00 | 6419.00 | 6063.00 | -1.36 | 0.00 |
| 6.00 | 1.00 | 2.33 | 1.16 | 1.00 |
| 40.00 | 75.00 | 56.33 | 1.21 | 0.23 |
| 278.00 | 318.00 | 311.33 | -1.25 | 0.00 |
| 234.00 | 308.00 | 282.67 | 1.04 | 0.63 |
| 300.00 | 398.00 | 327.33 | 1.06 | 0.48 |
| 37.00 | 37.00 | 44.67 | -4.55 | 0.00 |
| 83.00 | 91.00 | 82.00 | -1.12 | 0.38 |
| 1539.00 | 1550.00 | 1595.33 | 1.02 | 0.64 |
| 464.00 | 470.00 | 491.67 | 1.19 | 0.00 |
| 902.00 | 1022.00 | 1009.00 | 1.15 | 0.00 |
| 2.00 | 0.00 | 1.33 | -1.24 | 1.00 |
| 425.00 | 526.00 | 473.67 | -1.06 | 0.32 |
| 1004.00 | 1083.00 | 1073.33 | 1.21 | 0.00 |
| 1016.00 | 1013.00 | 1034.67 | -1.23 | 0.00 |
| 641.00 | 706.00 | 726.33 | -1.07 | 0.17 |

| | | | | |
|---|---|---|---|---|
| 287.00 | 292.00 | 295.00 | -1.16 | 0.01 |
| 308.00 | 307.00 | 303.33 | -1.12 | 0.07 |
| 35.00 | 54.00 | 45.33 | -1.25 | 0.08 |
| 8439.00 | 8369.00 | 8595.67 | -1.05 | 0.15 |
| 358.00 | 334.00 | 370.33 | -1.35 | 0.00 |
| 1247.00 | 1250.00 | 1310.67 | -1.17 | 0.00 |
| 727.00 | 697.00 | 727.33 | -1.12 | 0.03 |
| 6.00 | 5.00 | 5.33 | 1.12 | 0.86 |
| 789.00 | 937.00 | 868.33 | 1.04 | 0.40 |
| 279.00 | 287.00 | 290.33 | -1.07 | 0.23 |
| 255.00 | 267.00 | 268.00 | -1.01 | 0.94 |
| 3.00 | 12.00 | 6.33 | -1.07 | 1.00 |
| 941.00 | 1037.00 | 981.33 | -1.07 | 0.14 |
| 68.00 | 76.00 | 84.33 | -1.29 | 0.06 |
| 498.00 | 513.00 | 536.00 | 1.25 | 0.00 |
| 3.00 | 9.00 | 6.00 | -1.23 | 0.64 |
| 30.00 | 20.00 | 23.67 | -1.17 | 0.48 |
| 698.00 | 826.00 | 759.00 | 1.16 | 0.01 |
| 369.00 | 401.00 | 387.67 | -1.02 | 0.76 |
| 3898.00 | 4115.00 | 4090.67 | 1.05 | 0.18 |
| 494.00 | 555.00 | 548.67 | 1.24 | 0.00 |
| 32.00 | 35.00 | 36.67 | 1.08 | 0.66 |
| 12.00 | 7.00 | 12.00 | 1.41 | 0.22 |
| 11589.00 | 12169.00 | 11700.00 | 1.04 | 0.32 |
| 5675.00 | 6452.00 | 6196.33 | 1.15 | 0.00 |
| 129.00 | 156.00 | 150.00 | -2.02 | 0.00 |
| 1.00 | 2.00 | 1.67 | -1.77 | 0.43 |
| 0.00 | 3.00 | 1.33 | -3.10 | 0.06 |
| 281.00 | 308.00 | 300.33 | 1.29 | 0.00 |
| 1.00 | 1.00 | 1.00 | -1.91 | 0.51 |
| 1284.00 | 1353.00 | 1302.67 | -1.07 | 0.12 |
| 2054.00 | 2084.00 | 2121.67 | 1.15 | 0.00 |
| 9.00 | 11.00 | 11.00 | -1.23 | 0.49 |
| 64.00 | 71.00 | 69.67 | 2.45 | 0.00 |
| 4842.00 | 5095.00 | 4909.00 | 1.28 | 0.00 |
| 4631.00 | 4731.00 | 4764.00 | -1.13 | 0.00 |
| 157.00 | 110.00 | 127.33 | -1.08 | 0.49 |
| 596.00 | 561.00 | 568.67 | -1.42 | 0.00 |
| 1809.00 | 1744.00 | 1829.67 | -1.02 | 0.67 |
| 1228.00 | 1359.00 | 1314.67 | 1.17 | 0.00 |
| 3715.00 | 3925.00 | 3869.00 | 1.09 | 0.02 |
| 999.00 | 955.00 | 986.00 | 1.02 | 0.69 |
| 479.00 | 434.00 | 460.67 | 1.04 | 0.52 |
| 235.00 | 183.00 | 212.00 | -3.06 | 0.00 |
| 5.00 | 6.00 | 4.67 | -1.71 | 0.16 |
| 789.00 | 850.00 | 831.67 | 1.01 | 0.87 |
| 253.00 | 252.00 | 263.00 | -1.61 | 0.00 |
| 720.00 | 758.00 | 764.00 | 1.12 | 0.04 |

| | | | | |
|---|---|---|---|---|
| 8490.00 | 8880.00 | 9113.33 | 1.30 | 0.00 |
| 81.00 | 110.00 | 102.67 | 1.43 | 0.00 |
| 609.00 | 593.00 | 615.33 | 1.04 | 0.37 |
| 2721.00 | 2271.00 | 2470.00 | 1.17 | 0.01 |
| 9.00 | 13.00 | 12.33 | -1.07 | 0.91 |
| 1688.00 | 1787.00 | 1866.33 | 1.08 | 0.10 |
| 1048.00 | 1043.00 | 1072.33 | 1.10 | 0.03 |
| 498.00 | 607.00 | 578.33 | 1.17 | 0.00 |
| 1306.00 | 1404.00 | 1390.33 | 1.11 | 0.01 |
| 1684.00 | 1790.00 | 1737.67 | -1.08 | 0.05 |
| 19.00 | 17.00 | 17.33 | 1.60 | 0.04 |
| 6.00 | 8.00 | 7.00 | -1.30 | 0.48 |
| 5.00 | 8.00 | 7.67 | 1.72 | 0.14 |
| 1130.00 | 1147.00 | 1176.00 | 1.00 | 0.95 |
| 0.00 | 2.00 | 1.33 | -1.01 | 1.00 |
| 7.00 | 1.00 | 3.67 | 1.20 | 0.83 |
| 9.00 | 5.00 | 7.67 | 1.41 | 0.35 |
| 1970.00 | 1952.00 | 2068.33 | 1.03 | 0.43 |
| 189.00 | 165.00 | 180.67 | -1.00 | 1.00 |
| 1451.00 | 1566.00 | 1552.33 | 1.01 | 0.77 |
| 819.00 | 880.00 | 816.33 | -1.01 | 0.86 |
| 597.00 | 647.00 | 672.00 | -1.13 | 0.02 |
| 469.00 | 472.00 | 486.00 | 1.10 | 0.08 |
| 11.00 | 3.00 | 5.67 | 1.05 | 1.00 |
| 1171.00 | 1144.00 | 1177.00 | 1.02 | 0.60 |
| 2767.00 | 3024.00 | 3024.00 | 1.05 | 0.14 |
| 213.00 | 205.00 | 204.33 | 1.02 | 0.82 |
| 915.00 | 908.00 | 947.00 | 1.07 | 0.10 |
| 3267.00 | 3062.00 | 3164.00 | -1.10 | 0.03 |
| 327.00 | 375.00 | 375.33 | 1.19 | 0.00 |
| 533.00 | 526.00 | 522.00 | 1.19 | 0.00 |
| 6.00 | 6.00 | 6.33 | 1.33 | 0.49 |
| 65131.00 | 70618.00 | 69997.67 | 1.49 | 0.00 |
| 117.00 | 130.00 | 127.33 | -1.27 | 0.00 |
| 917.00 | 962.00 | 951.67 | 1.05 | 0.22 |
| 1339.00 | 1437.00 | 1415.67 | 1.13 | 0.00 |
| 1164.00 | 1087.00 | 1170.33 | 1.05 | 0.26 |
| 4621.00 | 4852.00 | 4758.33 | -1.19 | 0.00 |
| 2160.00 | 2431.00 | 2372.33 | 1.17 | 0.00 |
| 5965.00 | 6420.00 | 6267.67 | 1.02 | 0.60 |
| 322.00 | 343.00 | 350.00 | 1.16 | 0.02 |
| 1165.00 | 1242.00 | 1245.33 | 1.10 | 0.02 |
| 51.00 | 42.00 | 47.33 | -1.45 | 0.00 |
| 13.00 | 13.00 | 11.67 | 1.08 | 0.82 |
| 975.00 | 986.00 | 1000.33 | -1.17 | 0.00 |
| 2221.00 | 2414.00 | 2378.00 | -1.09 | 0.02 |
| 1690.00 | 1905.00 | 1864.67 | -1.02 | 0.67 |
| 1559.00 | 1736.00 | 1625.00 | -1.04 | 0.43 |

| | | | | |
|---|---|---|---|---|
| 26.00 | 35.00 | 33.33 | -1.52 | 0.00 |
| 6646.00 | 6793.00 | 6959.33 | -1.06 | 0.13 |
| 31330.00 | 30445.00 | 31558.33 | -1.03 | 0.39 |
| 3913.00 | 4327.00 | 4283.00 | 1.45 | 0.00 |
| 8152.00 | 8045.00 | 8349.00 | -1.14 | 0.00 |
| 805.00 | 794.00 | 787.67 | -1.20 | 0.00 |
| 835.00 | 843.00 | 861.00 | 1.22 | 0.00 |
| 817.00 | 807.00 | 851.33 | 1.22 | 0.00 |
| 4040.00 | 4118.00 | 4305.67 | -1.07 | 0.13 |
| 92.00 | 96.00 | 99.33 | 1.20 | 0.05 |
| 17.00 | 35.00 | 29.33 | 1.14 | 0.51 |
| 1.00 | 0.00 | 0.67 | -8.69 | 0.00 |
| 1904.00 | 1937.00 | 1948.67 | -1.12 | 0.01 |
| 35989.00 | 36646.00 | 37559.67 | -1.09 | 0.01 |
| 169.00 | 167.00 | 175.00 | -1.14 | 0.06 |
| 982.00 | 938.00 | 955.33 | 1.06 | 0.22 |
| 282.00 | 294.00 | 303.33 | 1.14 | 0.05 |
| 1009.00 | 981.00 | 1023.00 | 1.07 | 0.11 |
| 102.00 | 96.00 | 104.67 | 1.59 | 0.00 |
| 294.00 | 297.00 | 293.00 | 1.18 | 0.02 |
| 1398.00 | 1395.00 | 1452.00 | -1.07 | 0.08 |
| 1430.00 | 1470.00 | 1546.67 | 1.07 | 0.16 |
| 2818.00 | 2983.00 | 2976.00 | 1.04 | 0.35 |
| 2490.00 | 2880.00 | 2788.67 | 1.07 | 0.08 |
| 1836.00 | 2097.00 | 2028.33 | 1.10 | 0.01 |
| 2062.00 | 2094.00 | 2097.00 | 1.05 | 0.26 |
| 2356.00 | 2481.00 | 2418.67 | 1.01 | 0.75 |
| 597.00 | 703.00 | 645.67 | 1.10 | 0.09 |
| 1912.00 | 2005.00 | 2004.00 | -1.14 | 0.00 |
| 1807.00 | 1695.00 | 1799.67 | -1.28 | 0.00 |
| 2.00 | 1.00 | 2.00 | -1.81 | 0.34 |
| 455.00 | 548.00 | 521.67 | 1.08 | 0.16 |
| 1458.00 | 1327.00 | 1426.00 | -1.22 | 0.00 |
| 440.00 | 498.00 | 520.67 | 1.14 | 0.04 |
| 933.00 | 1019.00 | 1013.33 | -1.01 | 0.84 |
| 698.00 | 839.00 | 803.33 | 1.20 | 0.00 |
| 735.00 | 768.00 | 767.33 | 1.13 | 0.02 |
| 191.00 | 255.00 | 227.00 | 1.10 | 0.20 |
| 1648.00 | 2113.00 | 1911.33 | 2.10 | 0.00 |
| 968.00 | 1116.00 | 1075.33 | 1.60 | 0.00 |
| 939.00 | 1175.00 | 1121.67 | 1.40 | 0.00 |
| 45019.00 | 46665.00 | 45599.00 | 1.84 | 0.00 |
| 23052.00 | 23538.00 | 24053.33 | 2.05 | 0.00 |
| 490.00 | 664.00 | 575.33 | -1.12 | 0.07 |
| 332.00 | 386.00 | 368.67 | -1.08 | 0.19 |
| 157.00 | 167.00 | 178.67 | 1.04 | 0.67 |
| 305.00 | 315.00 | 329.33 | -1.03 | 0.66 |
| 241.00 | 321.00 | 289.00 | 1.25 | 0.01 |

| | | | | |
|---|---|---|---|---|
| 246.00 | 270.00 | 253.00 | -1.06 | 0.40 |
| 1037.00 | 1183.00 | 1187.33 | 1.01 | 0.86 |
| 20.00 | 23.00 | 22.00 | -1.29 | 0.16 |
| 5.00 | 9.00 | 6.67 | -1.26 | 0.56 |
| 592.00 | 587.00 | 613.00 | -1.02 | 0.62 |
| 1033.00 | 1023.00 | 1048.00 | -1.26 | 0.00 |
| 495.00 | 510.00 | 494.00 | -1.31 | 0.00 |
| 276.00 | 289.00 | 296.00 | 1.04 | 0.51 |
| 137.00 | 162.00 | 163.33 | 1.33 | 0.00 |
| 1100.00 | 1208.00 | 1185.33 | 1.12 | 0.01 |
| 1135.00 | 1200.00 | 1196.67 | -1.09 | 0.02 |
| 592.00 | 621.00 | 643.33 | -1.07 | 0.15 |
| 773.00 | 714.00 | 763.33 | 1.06 | 0.26 |
| 3.00 | 3.00 | 3.67 | 1.20 | 0.83 |
| 2570.00 | 2396.00 | 2564.67 | -1.15 | 0.00 |
| 368.00 | 366.00 | 372.67 | 1.17 | 0.01 |
| 276.00 | 198.00 | 230.33 | -1.29 | 0.00 |
| 2228.00 | 2210.00 | 2357.67 | -1.10 | 0.09 |
| 2846.00 | 2678.00 | 2794.00 | 1.06 | 0.16 |
| 3120.00 | 3202.00 | 3281.33 | 1.14 | 0.00 |
| 3576.00 | 4405.00 | 4107.67 | 1.36 | 0.00 |
| 1531.00 | 1541.00 | 1570.33 | 1.36 | 0.00 |
| 341.00 | 332.00 | 336.67 | -1.09 | 0.25 |
| 1273.00 | 1304.00 | 1334.67 | -1.16 | 0.00 |
| 884.00 | 911.00 | 890.33 | -1.24 | 0.00 |
| 16.00 | 20.00 | 18.33 | -1.45 | 0.05 |
| 4.00 | 9.00 | 5.67 | -1.65 | 0.14 |
| 8.00 | 5.00 | 6.00 | -1.84 | 0.06 |
| 1063.00 | 1066.00 | 1070.67 | 1.13 | 0.00 |
| 22.00 | 26.00 | 23.67 | 1.88 | 0.00 |
| 287.00 | 274.00 | 288.00 | -1.10 | 0.14 |
| 218.00 | 223.00 | 240.33 | -1.18 | 0.02 |
| 1.00 | 2.00 | 3.33 | -1.31 | 0.69 |
| 238.00 | 272.00 | 264.67 | 1.32 | 0.00 |
| 478.00 | 491.00 | 518.67 | -1.06 | 0.23 |
| 8.00 | 1.00 | 4.00 | -2.89 | 0.00 |
| 525.00 | 434.00 | 477.33 | -1.09 | 0.20 |
| 22.00 | 23.00 | 22.67 | -1.18 | 0.38 |
| 47.00 | 59.00 | 72.67 | 1.24 | 0.22 |
| 513.00 | 523.00 | 539.00 | 1.04 | 0.37 |
| 1546.00 | 1438.00 | 1511.33 | 1.15 | 0.00 |
| 51875.00 | 56185.00 | 55901.33 | 1.00 | 0.91 |
| 22.00 | 20.00 | 21.33 | 1.29 | 0.20 |
| 305.00 | 345.00 | 362.33 | 1.02 | 0.85 |
| 428.00 | 423.00 | 426.00 | -1.01 | 0.89 |
| 2247.00 | 2269.00 | 2344.67 | -1.34 | 0.00 |
| 759.00 | 754.00 | 771.67 | -1.20 | 0.00 |
| 16.00 | 11.00 | 13.00 | 1.66 | 0.06 |

| | | | | |
|---|---|---|---|---|
| 722.00 | 725.00 | 774.00 | -1.03 | 0.52 |
| 134.00 | 151.00 | 144.00 | -1.36 | 0.00 |
| 3.00 | 8.00 | 4.00 | -1.09 | 1.00 |
| 839.00 | 765.00 | 819.67 | -1.89 | 0.00 |
| 493.00 | 494.00 | 507.67 | -1.08 | 0.11 |
| 29.00 | 43.00 | 37.67 | 1.07 | 0.67 |
| 49.00 | 73.00 | 63.67 | 1.10 | 0.45 |
| 21.00 | 12.00 | 14.33 | -1.88 | 0.03 |
| 8.00 | 5.00 | 5.33 | 1.93 | 0.16 |
| 2655.00 | 2520.00 | 2761.33 | 1.08 | 0.14 |
| 1549.00 | 1529.00 | 1544.33 | -1.12 | 0.01 |
| 484.00 | 494.00 | 489.00 | -1.37 | 0.00 |
| 952.00 | 1073.00 | 1048.33 | -1.29 | 0.00 |
| 544.00 | 547.00 | 572.67 | -1.02 | 0.71 |
| 847.00 | 947.00 | 974.33 | -1.04 | 0.40 |
| 3299.00 | 3409.00 | 3530.67 | -1.05 | 0.19 |
| 2975.00 | 3293.00 | 3307.00 | 1.16 | 0.00 |
| 8.00 | 4.00 | 8.33 | -1.13 | 0.79 |
| 1186.00 | 1220.00 | 1267.00 | -1.10 | 0.03 |
| 2371.00 | 2704.00 | 2588.00 | 1.10 | 0.02 |
| 2785.00 | 2815.00 | 2816.33 | 1.28 | 0.00 |
| 35.00 | 28.00 | 33.00 | -1.18 | 0.27 |
| 1339.00 | 1456.00 | 1478.67 | 2.04 | 0.00 |
| 23.00 | 20.00 | 25.00 | 1.12 | 0.64 |
| 5063.00 | 5493.00 | 5450.00 | 1.18 | 0.00 |
| 1822.00 | 1925.00 | 1939.00 | 1.10 | 0.02 |
| 789.00 | 838.00 | 858.00 | 1.29 | 0.00 |
| 831.00 | 893.00 | 871.00 | 1.02 | 0.70 |
| 28.00 | 31.00 | 25.67 | -2.36 | 0.00 |
| 1056.00 | 1254.00 | 1247.67 | 1.27 | 0.00 |
| 8.00 | 5.00 | 8.00 | -1.14 | 0.79 |
| 5302.00 | 5285.00 | 5365.67 | -1.24 | 0.00 |
| 139.00 | 115.00 | 136.33 | 1.13 | 0.17 |
| 75.00 | 57.00 | 65.33 | 1.08 | 0.55 |
| 1133.00 | 1155.00 | 1188.00 | 1.18 | 0.00 |
| 17.00 | 19.00 | 23.33 | 1.11 | 0.63 |
| 362.00 | 390.00 | 387.33 | 1.12 | 0.05 |
| 568.00 | 565.00 | 575.33 | -1.04 | 0.43 |
| 1015.00 | 1041.00 | 1048.33 | -1.08 | 0.08 |
| 907.00 | 995.00 | 1011.00 | -1.06 | 0.21 |
| 684.00 | 743.00 | 738.00 | 1.12 | 0.01 |
| 2533.00 | 2888.00 | 2827.67 | 1.19 | 0.00 |
| 6.00 | 5.00 | 4.67 | 1.25 | 0.70 |
| 970.00 | 1040.00 | 1039.67 | 1.10 | 0.03 |
| 817.00 | 935.00 | 901.67 | -1.08 | 0.08 |
| 920.00 | 1016.00 | 956.33 | -1.10 | 0.09 |
| 550.00 | 574.00 | 593.00 | 1.01 | 0.90 |
| 117.00 | 98.00 | 111.33 | -1.11 | 0.23 |

| | | | | |
|---|---|---|---|---|
| 1267.00 | 1373.00 | 1371.67 | -1.17 | 0.00 |
| 1026.00 | 1070.00 | 1103.00 | 1.14 | 0.00 |
| 3242.00 | 3218.00 | 3207.67 | -1.03 | 0.51 |
| 4086.00 | 4570.00 | 4430.67 | -1.14 | 0.00 |
| 9.00 | 9.00 | 13.00 | 1.13 | 0.74 |
| 608.00 | 557.00 | 623.00 | -1.09 | 0.14 |
| 792.00 | 789.00 | 790.00 | -1.10 | 0.04 |
| 26872.00 | 26397.00 | 26390.67 | 1.09 | 0.03 |
| 216.00 | 156.00 | 175.33 | -1.11 | 0.34 |
| 20.00 | 6.00 | 12.67 | 1.10 | 0.74 |
| 698.00 | 666.00 | 682.67 | -1.27 | 0.00 |
| 18.00 | 17.00 | 19.67 | 1.26 | 0.30 |
| 374.00 | 450.00 | 411.00 | 1.32 | 0.00 |
| 1.00 | 0.00 | 2.00 | -1.65 | 0.46 |
| 233.00 | 226.00 | 230.67 | 1.32 | 0.00 |
| 680.00 | 792.00 | 756.00 | 1.23 | 0.00 |
| 401.00 | 410.00 | 414.00 | 1.13 | 0.02 |
| 633.00 | 750.00 | 699.33 | 1.12 | 0.03 |
| 783.00 | 898.00 | 840.00 | -1.05 | 0.34 |
| 281.00 | 330.00 | 315.67 | -1.12 | 0.04 |
| 727.00 | 746.00 | 739.00 | 1.00 | 0.93 |
| 126.00 | 132.00 | 133.67 | 1.24 | 0.01 |
| 10.00 | 14.00 | 14.33 | -1.08 | 0.76 |
| 1833.00 | 2064.00 | 2032.67 | 1.08 | 0.09 |
| 4437.00 | 4613.00 | 4638.67 | -1.08 | 0.02 |
| 2344.00 | 2614.00 | 2577.67 | -1.00 | 0.97 |
| 526.00 | 518.00 | 573.00 | 1.14 | 0.04 |
| 1557.00 | 1616.00 | 1644.00 | 1.04 | 0.44 |
| 1063.00 | 1058.00 | 1095.00 | -1.16 | 0.00 |
| 299.00 | 322.00 | 320.00 | 1.19 | 0.00 |
| 384.00 | 432.00 | 416.67 | 1.05 | 0.39 |
| 827.00 | 922.00 | 862.00 | 1.21 | 0.00 |
| 251.00 | 309.00 | 304.00 | -1.20 | 0.01 |
| 224.00 | 226.00 | 227.33 | -1.01 | 0.89 |
| 887.00 | 1082.00 | 1001.00 | 1.01 | 0.86 |
| 3685.00 | 4233.00 | 4157.00 | 1.22 | 0.00 |
| 820.00 | 819.00 | 888.00 | 1.02 | 0.69 |
| 725.00 | 758.00 | 764.33 | -1.14 | 0.00 |
| 3880.00 | 4211.00 | 4194.33 | 1.29 | 0.00 |
| 5063.00 | 4907.00 | 4980.00 | -1.07 | 0.09 |
| 283.00 | 313.00 | 308.67 | 1.28 | 0.00 |
| 1167.00 | 1406.00 | 1306.00 | -1.18 | 0.00 |
| 1202.00 | 1316.00 | 1271.67 | 1.03 | 0.45 |
| 44.00 | 60.00 | 51.67 | -1.27 | 0.05 |
| 267.00 | 320.00 | 294.67 | 1.19 | 0.01 |
| 996.00 | 1169.00 | 1197.67 | -1.12 | 0.04 |
| 1577.00 | 1767.00 | 1744.33 | 1.07 | 0.09 |
| 28.00 | 36.00 | 35.67 | 1.88 | 0.00 |

| | | | | |
|---|---|---|---|---|
| 14.00 | 17.00 | 15.67 | 1.05 | 0.84 |
| 10988.00 | 10678.00 | 11196.00 | -1.05 | 0.18 |
| 36.00 | 26.00 | 32.67 | -1.52 | 0.00 |
| 3277.00 | 2774.00 | 2858.00 | -1.09 | 0.22 |
| 1176.00 | 1187.00 | 1212.67 | 1.14 | 0.01 |
| 3269.00 | 3875.00 | 3721.33 | 1.14 | 0.00 |
| 992.00 | 1083.00 | 1068.33 | -1.38 | 0.00 |
| 2012.00 | 2109.00 | 2091.67 | 1.10 | 0.01 |
| 62.00 | 61.00 | 68.00 | 1.04 | 0.79 |
| 62.00 | 74.00 | 77.33 | -1.08 | 0.45 |
| 0.00 | 0.00 | 0.00 | -11.75 | 0.13 |
| 947.00 | 972.00 | 972.67 | -1.03 | 0.51 |
| 12094.00 | 12767.00 | 12729.67 | -1.03 | 0.32 |
| 3175.00 | 3408.00 | 3366.00 | 1.14 | 0.00 |
| 821.00 | 839.00 | 818.00 | -1.01 | 0.85 |
| 524.00 | 515.00 | 537.33 | 1.35 | 0.00 |
| 844.00 | 988.00 | 933.67 | 1.08 | 0.11 |
| 9595.00 | 10076.00 | 9634.67 | 1.11 | 0.03 |
| 1939.00 | 2076.00 | 2046.33 | -1.00 | 0.95 |
| 1.00 | 1.00 | 1.67 | -1.01 | 1.00 |
| 204.00 | 268.00 | 247.67 | -1.15 | 0.04 |
| 2334.00 | 2672.00 | 2497.33 | 1.07 | 0.09 |
| 298.00 | 349.00 | 341.67 | 1.09 | 0.17 |
| 1390.00 | 1455.00 | 1455.67 | -1.05 | 0.18 |
| 1948.00 | 2330.00 | 2218.33 | 1.22 | 0.00 |
| 1.00 | 4.00 | 2.33 | -1.43 | 0.63 |
| 9060.00 | 9627.00 | 9390.67 | 1.01 | 0.79 |
| 2.00 | 5.00 | 3.67 | 1.20 | 0.83 |
| 1522.00 | 1654.00 | 1629.67 | -1.03 | 0.45 |
| 4721.00 | 5158.00 | 5065.00 | 1.01 | 0.87 |
| 2455.00 | 2811.00 | 2628.33 | 1.10 | 0.03 |
| 1440.00 | 1503.00 | 1499.00 | 1.09 | 0.03 |
| 827.00 | 879.00 | 901.33 | 1.24 | 0.00 |
| 27146.00 | 31617.00 | 29894.00 | 1.06 | 0.08 |
| 1579.00 | 1732.00 | 1732.00 | 1.19 | 0.00 |
| 808.00 | 848.00 | 864.67 | -1.12 | 0.01 |
| 112.00 | 99.00 | 110.00 | -1.04 | 0.66 |
| 12.00 | 13.00 | 14.67 | -1.13 | 0.63 |
| 2268.00 | 2156.00 | 2243.33 | -1.04 | 0.35 |
| 1208.00 | 1374.00 | 1361.67 | 1.06 | 0.19 |
| 3034.00 | 3329.00 | 3258.67 | 1.03 | 0.40 |
| 674.00 | 745.00 | 705.00 | 1.21 | 0.00 |
| 442.00 | 482.00 | 479.33 | 1.06 | 0.29 |
| 2246.00 | 2317.00 | 2320.33 | -1.05 | 0.19 |
| 135.00 | 142.00 | 135.00 | 1.24 | 0.01 |
| 109.00 | 155.00 | 138.33 | -1.27 | 0.00 |
| 477.00 | 495.00 | 490.33 | 1.03 | 0.61 |
| 1229.00 | 1335.00 | 1376.67 | 1.13 | 0.01 |

| | | | | |
|---|---|---|---|---|
| 801.00 | 872.00 | 867.00 | 1.03 | 0.47 |
| 278.00 | 228.00 | 272.00 | -1.08 | 0.35 |
| 2235.00 | 2309.00 | 2405.33 | -1.12 | 0.00 |
| 2427.00 | 2403.00 | 2488.33 | -1.07 | 0.10 |
| 3441.00 | 3769.00 | 3583.33 | 1.32 | 0.00 |
| 463.00 | 422.00 | 466.33 | 1.06 | 0.37 |
| 162.00 | 204.00 | 173.67 | 1.14 | 0.13 |
| 0.00 | 0.00 | 0.33 | -3.22 | 0.39 |
| 466.00 | 449.00 | 479.33 | -1.44 | 0.00 |
| 42.00 | 51.00 | 44.00 | 1.03 | 0.91 |
| 1069.00 | 1203.00 | 1215.33 | -1.01 | 0.85 |
| 605.00 | 608.00 | 626.00 | -1.11 | 0.07 |
| 2329.00 | 2199.00 | 2329.67 | -1.25 | 0.00 |
| 722.00 | 714.00 | 713.33 | -1.12 | 0.02 |
| 1661.00 | 1637.00 | 1639.00 | -1.17 | 0.00 |
| 675.00 | 655.00 | 645.67 | -1.15 | 0.02 |
| 512.00 | 576.00 | 540.33 | 1.04 | 0.54 |
| 21.00 | 13.00 | 18.67 | -1.18 | 0.49 |
| 2779.00 | 3254.00 | 3131.67 | 1.17 | 0.00 |
| 1573.00 | 1877.00 | 1718.67 | 1.24 | 0.00 |
| 4.00 | 3.00 | 4.33 | 1.27 | 0.68 |
| 297.00 | 399.00 | 343.67 | 1.33 | 0.00 |
| 20.00 | 14.00 | 21.00 | 1.13 | 0.60 |
| 338.00 | 311.00 | 339.00 | -1.01 | 0.86 |
| 168.00 | 192.00 | 188.67 | 1.20 | 0.02 |
| 36.00 | 38.00 | 39.00 | 1.21 | 0.24 |
| 574.00 | 651.00 | 654.00 | 1.25 | 0.00 |
| 2650.00 | 2449.00 | 2557.67 | -1.05 | 0.25 |
| 1503.00 | 1631.00 | 1623.33 | 1.01 | 0.88 |
| 1590.00 | 1682.00 | 1697.00 | -1.11 | 0.00 |
| 1530.00 | 1687.00 | 1688.33 | 1.17 | 0.00 |
| 211.00 | 255.00 | 236.67 | -1.01 | 0.93 |
| 481.00 | 504.00 | 538.33 | 1.03 | 0.66 |
| 1205.00 | 1136.00 | 1125.33 | 1.21 | 0.00 |
| 368.00 | 368.00 | 377.67 | 1.01 | 0.92 |
| 1809.00 | 1873.00 | 1860.67 | -1.07 | 0.11 |
| 73.00 | 72.00 | 75.67 | -1.20 | 0.07 |
| 125.00 | 174.00 | 139.00 | -1.13 | 0.22 |
| 806.00 | 808.00 | 841.33 | -1.09 | 0.05 |
| 2651.00 | 2499.00 | 2648.00 | 1.05 | 0.26 |
| 24.00 | 34.00 | 27.67 | 1.15 | 0.50 |
| 332.00 | 367.00 | 363.33 | -1.14 | 0.02 |
| 1737.00 | 1843.00 | 1825.33 | -1.09 | 0.03 |
| 426.00 | 386.00 | 427.33 | 1.15 | 0.02 |
| 1285.00 | 1443.00 | 1397.00 | -1.02 | 0.58 |
| 365.00 | 409.00 | 394.33 | -1.15 | 0.02 |
| 1166.00 | 1093.00 | 1124.67 | -1.33 | 0.00 |
| 123.00 | 164.00 | 139.33 | -1.06 | 0.51 |

| | | | | |
|---|---|---|---|---|
| 46.00 | 79.00 | 65.33 | -1.75 | 0.00 |
| 786.00 | 866.00 | 778.67 | -1.19 | 0.01 |
| 43.00 | 45.00 | 48.67 | -1.23 | 0.13 |
| 473.00 | 424.00 | 486.67 | -1.26 | 0.00 |
| 961.00 | 1032.00 | 1051.00 | -1.02 | 0.71 |
| 1127.00 | 1293.00 | 1304.67 | 1.19 | 0.00 |
| 2441.00 | 2349.00 | 2416.67 | -1.26 | 0.00 |
| 14113.00 | 14714.00 | 14683.00 | -1.10 | 0.01 |
| 581.00 | 609.00 | 622.67 | -1.15 | 0.00 |
| 2.00 | 3.00 | 2.67 | -1.50 | 0.52 |
| 238.00 | 331.00 | 293.67 | 1.17 | 0.04 |
| 330.00 | 307.00 | 326.33 | 1.21 | 0.00 |
| 461.00 | 499.00 | 499.00 | 1.19 | 0.00 |
| 1336.00 | 1397.00 | 1408.67 | 1.02 | 0.56 |
| 290.00 | 308.00 | 324.33 | -1.08 | 0.20 |
| 74.00 | 58.00 | 75.67 | 1.30 | 0.03 |
| 115.00 | 136.00 | 132.00 | 1.49 | 0.00 |
| 3.00 | 2.00 | 2.00 | -1.17 | 1.00 |
| 132.00 | 166.00 | 165.67 | -1.23 | 0.01 |
| 162.00 | 168.00 | 172.00 | 1.19 | 0.03 |
| 249.00 | 243.00 | 256.33 | 1.03 | 0.66 |
| 92.00 | 116.00 | 106.67 | 1.40 | 0.00 |
| 1846.00 | 1848.00 | 1914.33 | 1.21 | 0.00 |
| 328.00 | 396.00 | 383.67 | 1.14 | 0.07 |
| 594.00 | 565.00 | 570.67 | -1.09 | 0.12 |
| 3388.00 | 3945.00 | 3703.00 | 1.11 | 0.01 |
| 1048.00 | 1142.00 | 1102.67 | -1.06 | 0.21 |
| 2589.00 | 2661.00 | 2712.00 | -1.00 | 0.94 |
| 1068.00 | 1135.00 | 1166.33 | 1.03 | 0.60 |
| 2642.00 | 2630.00 | 2689.00 | -1.03 | 0.41 |
| 1224.00 | 1335.00 | 1308.00 | -1.19 | 0.00 |
| 630.00 | 643.00 | 656.00 | -1.06 | 0.23 |
| 1236.00 | 1249.00 | 1258.67 | 1.04 | 0.34 |
| 1201.00 | 1136.00 | 1204.33 | -1.07 | 0.16 |
| 2132.00 | 2225.00 | 2188.00 | -1.06 | 0.20 |
| 1194.00 | 1361.00 | 1327.00 | 1.02 | 0.70 |
| 991.00 | 1179.00 | 1093.33 | 1.07 | 0.13 |
| 36.00 | 39.00 | 40.00 | -1.12 | 0.44 |
| 950.00 | 979.00 | 936.00 | -1.07 | 0.17 |
| 5452.00 | 6031.00 | 5827.00 | 1.12 | 0.00 |
| 707.00 | 819.00 | 788.67 | 1.11 | 0.03 |
| 3246.00 | 3388.00 | 3312.00 | -1.00 | 0.99 |
| 381.00 | 419.00 | 411.33 | 1.10 | 0.12 |
| 7.00 | 9.00 | 5.33 | 1.13 | 0.86 |
| 436.00 | 482.00 | 430.67 | 1.25 | 0.00 |
| 1772.00 | 2009.00 | 1868.67 | -1.04 | 0.40 |
| 130.00 | 156.00 | 148.00 | -1.18 | 0.04 |
| 3787.00 | 3772.00 | 3751.67 | -1.15 | 0.00 |

| | | | | |
|---|---|---|---|---|
| 1239.00 | 1296.00 | 1289.67 | -1.00 | 0.97 |
| 516.00 | 596.00 | 584.33 | 1.16 | 0.00 |
| 8.00 | 19.00 | 13.00 | -1.27 | 0.36 |
| 1375.00 | 1447.00 | 1439.33 | -1.32 | 0.00 |
| 6.00 | 2.00 | 3.00 | -1.66 | 0.37 |
| 1036.00 | 1066.00 | 1021.00 | -1.09 | 0.08 |
| 468.00 | 481.00 | 489.00 | -1.15 | 0.01 |
| 14.00 | 12.00 | 11.00 | -1.04 | 1.00 |
| 676.00 | 698.00 | 722.00 | 1.00 | 1.00 |
| 1.00 | 0.00 | 0.67 | -1.44 | 1.00 |
| 1628.00 | 1759.00 | 1719.00 | 1.07 | 0.12 |
| 1141.00 | 1288.00 | 1223.33 | 1.01 | 0.77 |
| 853.00 | 770.00 | 829.33 | -1.15 | 0.00 |
| 7.00 | 2.00 | 3.00 | -1.77 | 0.24 |
| 5.00 | 8.00 | 6.67 | -1.41 | 0.32 |
| 192.00 | 170.00 | 184.33 | -1.48 | 0.00 |
| 243.00 | 263.00 | 251.33 | -1.17 | 0.03 |
| 315.00 | 339.00 | 325.00 | 1.21 | 0.00 |
| 3569.00 | 3924.00 | 3945.00 | 1.19 | 0.00 |
| 36.00 | 27.00 | 30.67 | -1.19 | 0.29 |
| 149.00 | 157.00 | 156.00 | -1.14 | 0.10 |
| 52.00 | 80.00 | 69.33 | -1.00 | 1.00 |
| 7837.00 | 8621.00 | 8526.67 | 1.03 | 0.44 |
| 118.00 | 166.00 | 152.33 | -1.11 | 0.27 |
| 12970.00 | 12717.00 | 13093.67 | -1.58 | 0.00 |
| 1027.00 | 1189.00 | 1125.00 | 1.13 | 0.01 |
| 4914.00 | 5021.00 | 4961.67 | -1.00 | 0.99 |
| 1577.00 | 1777.00 | 1701.67 | 1.13 | 0.00 |
| 247.00 | 272.00 | 283.00 | 1.05 | 0.49 |
| 44.00 | 42.00 | 47.00 | -1.12 | 0.39 |
| 15.00 | 18.00 | 15.67 | 1.25 | 0.34 |
| 2929.00 | 3220.00 | 3105.67 | 1.03 | 0.36 |
| 19.00 | 30.00 | 24.00 | 1.65 | 0.02 |
| 603.00 | 609.00 | 613.00 | -1.08 | 0.12 |
| 1303.00 | 1228.00 | 1306.67 | -1.11 | 0.02 |
| 3.00 | 1.00 | 1.67 | -2.52 | 0.10 |
| 395.00 | 380.00 | 393.00 | 1.03 | 0.61 |
| 290.00 | 343.00 | 327.67 | -1.09 | 0.17 |
| 2.00 | 2.00 | 1.33 | -1.01 | 1.00 |
| 2055.00 | 1991.00 | 2099.00 | -1.32 | 0.00 |
| 3.00 | 8.00 | 6.00 | 3.37 | 0.01 |
| 739.00 | 844.00 | 786.00 | 1.43 | 0.00 |
| 1.00 | 2.00 | 1.67 | -2.14 | 0.22 |
| 126.00 | 109.00 | 124.67 | 1.38 | 0.00 |
| 5.00 | 2.00 | 5.33 | -1.51 | 0.28 |
| 2328.00 | 2464.00 | 2470.33 | 1.01 | 0.70 |
| 3.00 | 2.00 | 2.00 | -1.81 | 0.34 |
| 778.00 | 850.00 | 863.00 | 1.12 | 0.02 |

| | | | | |
|---|---|---|---|---|
| 81.00 | 116.00 | 99.00 | 1.45 | 0.00 |
| 11.00 | 7.00 | 10.00 | 2.09 | 0.03 |
| 2.00 | 5.00 | 6.00 | 1.59 | 0.29 |
| 29.00 | 22.00 | 28.67 | 1.51 | 0.02 |
| 626.00 | 633.00 | 656.00 | -1.01 | 0.86 |
| 50.00 | 54.00 | 51.00 | -1.24 | 0.10 |
| 13.00 | 13.00 | 14.67 | -2.07 | 0.00 |
| 1304.00 | 1234.00 | 1334.00 | -1.20 | 0.00 |
| 50.00 | 63.00 | 60.67 | -1.16 | 0.20 |
| 2807.00 | 3294.00 | 3134.00 | -1.08 | 0.05 |
| 5.00 | 2.00 | 4.33 | 1.58 | 0.39 |
| 999.00 | 812.00 | 894.67 | -1.20 | 0.00 |
| 693.00 | 721.00 | 694.00 | -1.13 | 0.02 |
| 1087.00 | 1262.00 | 1132.67 | -1.03 | 0.63 |
| 498.00 | 586.00 | 553.67 | 1.22 | 0.00 |
| 413.00 | 434.00 | 429.33 | 1.04 | 0.47 |
| 12.00 | 6.00 | 10.00 | -1.08 | 0.91 |
| 1602.00 | 1732.00 | 1709.33 | -1.01 | 0.88 |
| 150.00 | 117.00 | 139.00 | -1.67 | 0.00 |
| 534.00 | 427.00 | 489.67 | -1.45 | 0.00 |
| 20.00 | 24.00 | 19.67 | -1.35 | 0.13 |
| 595.00 | 709.00 | 677.00 | 1.00 | 0.94 |
| 223.00 | 193.00 | 202.33 | -1.15 | 0.07 |
| 348.00 | 338.00 | 329.00 | -1.22 | 0.00 |
| 40.00 | 46.00 | 37.00 | 1.17 | 0.40 |
| 10.00 | 18.00 | 13.33 | 1.04 | 0.91 |
| 3624.00 | 4347.00 | 4037.33 | 1.12 | 0.00 |
| 1855.00 | 2110.00 | 2099.33 | 1.09 | 0.05 |
| 3.00 | 1.00 | 3.00 | -1.45 | 0.53 |
| 1361.00 | 1291.00 | 1425.00 | -1.09 | 0.06 |
| 2506.00 | 2652.00 | 2651.00 | 1.01 | 0.90 |
| 2067.00 | 1974.00 | 2114.33 | 1.16 | 0.00 |
| 6.00 | 8.00 | 7.33 | -1.51 | 0.19 |
| 622.00 | 772.00 | 676.33 | -1.07 | 0.29 |
| 655.00 | 658.00 | 670.33 | -1.02 | 0.73 |
| 253.00 | 293.00 | 283.67 | -1.26 | 0.00 |
| 181.00 | 147.00 | 162.33 | 1.14 | 0.13 |
| 6571.00 | 6786.00 | 6908.67 | 1.21 | 0.00 |
| 598.00 | 644.00 | 598.33 | -1.04 | 0.54 |
| 1268.00 | 1352.00 | 1335.00 | -1.09 | 0.02 |
| 6048.00 | 6327.00 | 6064.67 | -1.00 | 0.99 |
| 2261.00 | 2386.00 | 2363.00 | 1.07 | 0.06 |
| 7.00 | 7.00 | 7.67 | 1.72 | 0.14 |
| 2707.00 | 2813.00 | 2749.67 | -1.01 | 0.77 |
| 1266.00 | 1243.00 | 1283.67 | -1.07 | 0.08 |
| 304.00 | 396.00 | 370.33 | 1.17 | 0.02 |
| 951.00 | 963.00 | 969.67 | -1.17 | 0.00 |
| 237.00 | 266.00 | 279.00 | 1.09 | 0.24 |

| | | | | |
|---|---|---|---|---|
| 834.00 | 805.00 | 823.67 | -1.03 | 0.59 |
| 674.00 | 842.00 | 818.33 | 1.14 | 0.03 |
| 1255.00 | 1473.00 | 1396.67 | 1.09 | 0.05 |
| 934.00 | 1086.00 | 1047.67 | 1.15 | 0.00 |
| 340.00 | 297.00 | 323.33 | 1.06 | 0.38 |
| 441.00 | 540.00 | 517.67 | 1.14 | 0.02 |
| 257.00 | 281.00 | 272.00 | 1.11 | 0.11 |
| 133.00 | 169.00 | 156.00 | -1.16 | 0.05 |
| 1180.00 | 1129.00 | 1145.33 | -1.04 | 0.38 |
| 161.00 | 176.00 | 181.33 | 1.08 | 0.31 |
| 62.00 | 80.00 | 65.67 | -1.12 | 0.36 |
| 838.00 | 833.00 | 876.33 | -1.07 | 0.15 |
| 2346.00 | 2421.00 | 2443.67 | -1.01 | 0.82 |
| 3.00 | 10.00 | 6.67 | 2.40 | 0.04 |
| 216.00 | 166.00 | 195.67 | 1.00 | 1.00 |
| 1744.00 | 1933.00 | 1931.33 | 1.10 | 0.02 |
| 1796.00 | 2054.00 | 2087.00 | 1.15 | 0.00 |
| 1120.00 | 1272.00 | 1301.67 | 1.01 | 0.86 |
| 3.00 | 8.00 | 4.67 | 1.25 | 0.70 |
| 246.00 | 282.00 | 274.00 | -1.13 | 0.04 |
| 237.00 | 228.00 | 261.00 | -1.00 | 1.00 |
| 181.00 | 174.00 | 171.00 | -1.20 | 0.05 |
| 377.00 | 378.00 | 406.33 | 1.06 | 0.37 |
| 73.00 | 74.00 | 76.00 | -1.22 | 0.04 |
| 1006.00 | 973.00 | 1020.00 | -1.14 | 0.01 |
| 88.00 | 116.00 | 105.00 | 1.18 | 0.09 |
| 1358.00 | 1316.00 | 1368.00 | -1.03 | 0.54 |
| 1663.00 | 1622.00 | 1680.67 | -1.38 | 0.00 |
| 227.00 | 231.00 | 226.33 | 1.07 | 0.35 |
| 1250.00 | 1296.00 | 1273.33 | -1.16 | 0.00 |
| 2414.00 | 2700.00 | 2624.33 | 1.11 | 0.01 |
| 12.00 | 18.00 | 16.33 | -1.20 | 0.40 |
| 2829.00 | 2792.00 | 2911.33 | -1.07 | 0.06 |
| 26.00 | 27.00 | 25.67 | 1.26 | 0.19 |
| 6934.00 | 7010.00 | 7091.67 | -1.13 | 0.00 |
| 3.00 | 3.00 | 2.33 | -4.17 | 0.00 |
| 4525.00 | 4643.00 | 4659.00 | 1.13 | 0.00 |
| 394.00 | 436.00 | 417.33 | -1.24 | 0.00 |
| 202.00 | 214.00 | 206.00 | -1.23 | 0.00 |
| 924.00 | 930.00 | 967.00 | -1.04 | 0.38 |
| 844.00 | 941.00 | 974.67 | 1.27 | 0.00 |
| 869.00 | 1020.00 | 1006.67 | 1.14 | 0.01 |
| 1778.00 | 2004.00 | 1940.33 | 1.04 | 0.33 |
| 10.00 | 4.00 | 6.33 | 1.10 | 0.87 |
| 2233.00 | 2209.00 | 2166.33 | -1.08 | 0.11 |
| 2012.00 | 2145.00 | 2028.33 | 1.01 | 0.82 |
| 331.00 | 336.00 | 342.33 | -1.29 | 0.00 |
| 184.00 | 158.00 | 160.33 | -1.57 | 0.00 |

| | | | | |
|---|---|---|---|---|
| 1389.00 | 1411.00 | 1428.00 | -1.52 | 0.00 |
| 2889.00 | 3081.00 | 2878.00 | 1.02 | 0.69 |
| 1396.00 | 1489.00 | 1401.67 | -1.08 | 0.13 |
| 826.00 | 827.00 | 834.00 | -1.23 | 0.00 |
| 1873.00 | 1750.00 | 1743.67 | 1.06 | 0.30 |
| 781.00 | 734.00 | 769.00 | -1.62 | 0.00 |
| 5.00 | 11.00 | 7.67 | 1.19 | 0.65 |
| 388.00 | 436.00 | 421.00 | 1.06 | 0.27 |
| 114.00 | 128.00 | 130.33 | 1.07 | 0.51 |
| 3.00 | 1.00 | 3.00 | -1.23 | 0.83 |
| 158.00 | 167.00 | 176.00 | -1.31 | 0.00 |
| 27.00 | 35.00 | 33.00 | -1.04 | 0.85 |
| 14.00 | 23.00 | 15.00 | -1.05 | 0.93 |
| 4.00 | 3.00 | 3.33 | -2.87 | 0.00 |
| 609.00 | 696.00 | 665.33 | -1.09 | 0.08 |
| 659.00 | 620.00 | 650.33 | 1.01 | 0.93 |
| 378.00 | 446.00 | 431.33 | 1.01 | 0.94 |
| 393.00 | 422.00 | 423.67 | -1.24 | 0.00 |
| 554.00 | 491.00 | 515.00 | -1.07 | 0.30 |
| 1599.00 | 1579.00 | 1629.00 | -1.08 | 0.05 |
| 2268.00 | 2434.00 | 2431.00 | -1.14 | 0.00 |
| 308.00 | 340.00 | 338.00 | 1.21 | 0.00 |
| 66.00 | 73.00 | 67.00 | -1.11 | 0.36 |
| 2170.00 | 2194.00 | 2248.33 | -1.02 | 0.66 |
| 372.00 | 392.00 | 387.67 | -1.13 | 0.03 |
| 733.00 | 650.00 | 707.67 | 1.02 | 0.70 |
| 1263.00 | 1267.00 | 1329.67 | -1.02 | 0.59 |
| 658.00 | 719.00 | 733.67 | 1.11 | 0.04 |
| 1309.00 | 1411.00 | 1451.00 | 1.09 | 0.07 |
| 238.00 | 275.00 | 268.00 | 1.03 | 0.64 |
| 6.00 | 23.00 | 15.67 | 1.21 | 0.49 |
| 1183.00 | 1208.00 | 1264.00 | -1.11 | 0.02 |
| 1174.00 | 1458.00 | 1382.33 | 1.19 | 0.00 |
| 911.00 | 996.00 | 1020.00 | 1.08 | 0.15 |
| 310.00 | 303.00 | 309.33 | 1.15 | 0.05 |
| 1534.00 | 1539.00 | 1575.00 | 1.02 | 0.55 |
| 2726.00 | 2643.00 | 2837.33 | -1.06 | 0.14 |
| 519.00 | 565.00 | 524.33 | -1.15 | 0.03 |
| 6344.00 | 6308.00 | 6449.67 | -1.08 | 0.02 |
| 12.00 | 11.00 | 10.33 | -1.21 | 0.56 |
| 12.00 | 4.00 | 8.00 | -1.01 | 1.00 |
| 3422.00 | 3695.00 | 3488.00 | -1.19 | 0.00 |
| 81.00 | 85.00 | 95.00 | -1.23 | 0.03 |
| 579.00 | 813.00 | 693.67 | -1.03 | 0.62 |
| 159.00 | 203.00 | 183.67 | 1.39 | 0.00 |
| 173.00 | 200.00 | 192.67 | -1.00 | 0.96 |
| 7.00 | 1.00 | 5.33 | -1.01 | 1.00 |
| 31.00 | 64.00 | 45.33 | 1.05 | 0.78 |

| | | | | |
|---|---|---|---|---|
| 1245.00 | 1393.00 | 1316.33 | -1.15 | 0.00 |
| 1182.00 | 1400.00 | 1292.33 | -1.05 | 0.29 |
| 637.00 | 733.00 | 705.67 | 1.24 | 0.00 |
| 397.00 | 400.00 | 408.67 | 1.15 | 0.02 |
| 0.00 | 2.00 | 1.67 | -1.97 | 0.37 |
| 745.00 | 751.00 | 783.00 | -1.05 | 0.36 |
| 1317.00 | 1357.00 | 1333.00 | -1.28 | 0.00 |
| 15.00 | 13.00 | 14.33 | -1.06 | 0.92 |
| 1295.00 | 1440.00 | 1386.67 | 1.12 | 0.00 |
| 3.00 | 4.00 | 2.33 | -1.29 | 0.81 |
| 1182.00 | 1066.00 | 1137.00 | 1.05 | 0.28 |
| 1300.00 | 1304.00 | 1331.67 | 1.11 | 0.01 |
| 8783.00 | 9543.00 | 9658.33 | -1.31 | 0.00 |
| 16.00 | 11.00 | 14.00 | -1.80 | 0.00 |
| 2.00 | 7.00 | 4.33 | -1.62 | 0.25 |
| 0.00 | 2.00 | 1.67 | -2.33 | 0.15 |
| 1061.00 | 1048.00 | 1122.67 | 1.10 | 0.05 |
| 899.00 | 908.00 | 966.00 | 1.00 | 0.98 |
| 1983.00 | 2084.00 | 2034.33 | -1.09 | 0.02 |
| 70.00 | 56.00 | 62.00 | -1.33 | 0.01 |
| 55.00 | 48.00 | 47.67 | -1.03 | 0.87 |
| 0.00 | 0.00 | 1.33 | -1.73 | 0.60 |
| 265.00 | 320.00 | 295.00 | 1.68 | 0.00 |
| 1188.00 | 1080.00 | 1144.00 | 1.01 | 0.81 |
| 3.00 | 6.00 | 5.33 | 1.12 | 0.86 |
| 36.00 | 42.00 | 38.67 | -1.10 | 0.53 |
| 977.00 | 940.00 | 954.33 | -1.05 | 0.30 |
| 352.00 | 329.00 | 328.33 | -1.00 | 0.97 |
| 735.00 | 849.00 | 798.33 | 1.05 | 0.34 |
| 10604.00 | 11499.00 | 10964.67 | -1.20 | 0.00 |
| 2297.00 | 2763.00 | 2646.33 | 1.16 | 0.00 |
| 80.00 | 80.00 | 82.33 | 1.12 | 0.29 |
| 514.00 | 522.00 | 518.33 | -1.27 | 0.00 |
| 1105.00 | 1223.00 | 1242.33 | 1.13 | 0.01 |
| 313.00 | 362.00 | 352.00 | 1.09 | 0.17 |
| 718.00 | 798.00 | 771.67 | 1.31 | 0.00 |
| 639.00 | 699.00 | 676.00 | -1.01 | 0.81 |
| 212.00 | 234.00 | 238.33 | 1.20 | 0.01 |
| 382.00 | 434.00 | 438.00 | 1.21 | 0.00 |
| 463.00 | 558.00 | 555.67 | -1.02 | 0.74 |
| 67.00 | 73.00 | 65.33 | 1.17 | 0.18 |
| 141.00 | 119.00 | 138.00 | -1.18 | 0.05 |
| 35.00 | 37.00 | 34.67 | -1.09 | 0.57 |
| 1345.00 | 1467.00 | 1543.67 | -1.28 | 0.00 |
| 549.00 | 536.00 | 555.67 | -1.29 | 0.00 |
| 1142.00 | 1287.00 | 1281.33 | 1.18 | 0.00 |
| 3077.00 | 3287.00 | 3376.67 | -1.07 | 0.12 |
| 455.00 | 498.00 | 505.67 | 1.05 | 0.45 |

| | | | | |
|---|---|---|---|---|
| 1074.00 | 1073.00 | 1142.67 | 1.08 | 0.09 |
| 489.00 | 594.00 | 567.00 | 1.12 | 0.04 |
| 4328.00 | 4716.00 | 4638.67 | 1.27 | 0.00 |
| 172.00 | 179.00 | 177.67 | -1.49 | 0.00 |
| 396.00 | 434.00 | 409.00 | 1.18 | 0.00 |
| 785.00 | 917.00 | 884.00 | 1.09 | 0.06 |
| 9.00 | 7.00 | 9.00 | -1.12 | 0.80 |
| 409.00 | 429.00 | 419.33 | -1.24 | 0.00 |
| 88.00 | 81.00 | 85.00 | 1.30 | 0.01 |
| 46.00 | 54.00 | 55.00 | 1.20 | 0.17 |
| 21.00 | 24.00 | 24.33 | 1.11 | 0.62 |
| 415.00 | 498.00 | 490.33 | -1.06 | 0.32 |
| 117.00 | 123.00 | 120.67 | 1.07 | 0.45 |
| 57.00 | 57.00 | 62.67 | -1.07 | 0.56 |
| 1928.00 | 1968.00 | 1967.33 | -1.33 | 0.00 |
| 640.00 | 625.00 | 627.33 | -2.53 | 0.00 |
| 4933.00 | 5005.00 | 5200.67 | 1.12 | 0.00 |
| 1.00 | 2.00 | 1.33 | -2.40 | 0.19 |
| 3559.00 | 4319.00 | 4010.33 | -1.22 | 0.00 |
| 988.00 | 1030.00 | 1050.33 | 1.07 | 0.11 |
| 1551.00 | 1809.00 | 1756.33 | 1.15 | 0.00 |
| 8.00 | 7.00 | 5.67 | 1.19 | 0.73 |
| 147.00 | 137.00 | 142.33 | -1.20 | 0.02 |
| 2829.00 | 2653.00 | 2860.67 | -1.02 | 0.59 |
| 1649.00 | 1716.00 | 1768.00 | 1.02 | 0.68 |
| 2224.00 | 2146.00 | 2252.33 | -1.23 | 0.00 |
| 54.00 | 52.00 | 53.67 | 1.19 | 0.16 |
| 605.00 | 588.00 | 612.67 | 1.06 | 0.29 |
| 8.00 | 2.00 | 7.00 | 1.03 | 1.00 |
| 28265.00 | 27908.00 | 27575.00 | -1.07 | 0.14 |
| 39.00 | 41.00 | 44.67 | 1.10 | 0.52 |
| 1860.00 | 2150.00 | 2020.67 | 1.09 | 0.04 |
| 982.00 | 1118.00 | 1138.67 | 1.16 | 0.00 |
| 2612.00 | 2946.00 | 2770.33 | -1.05 | 0.21 |
| 3463.00 | 3750.00 | 3507.00 | -1.13 | 0.01 |
| 442.00 | 554.00 | 501.33 | 1.09 | 0.17 |
| 2341.00 | 2699.00 | 2726.00 | 1.08 | 0.11 |
| 721.00 | 864.00 | 815.67 | -1.15 | 0.00 |
| 1128.00 | 1273.00 | 1336.67 | 1.07 | 0.28 |
| 639.00 | 754.00 | 733.67 | 1.05 | 0.35 |
| 236.00 | 376.00 | 316.33 | 1.07 | 0.47 |
| 994.00 | 1094.00 | 1061.67 | -1.03 | 0.49 |
| 129.00 | 114.00 | 120.67 | -1.11 | 0.22 |
| 655.00 | 719.00 | 675.33 | -1.21 | 0.00 |
| 558.00 | 543.00 | 541.33 | -1.11 | 0.06 |
| 360.00 | 444.00 | 407.00 | 1.05 | 0.44 |
| 2.00 | 0.00 | 1.00 | -1.31 | 1.00 |
| 46.00 | 48.00 | 48.67 | 1.31 | 0.05 |

| | | | | |
|---|---|---|---|---|
| 145.00 | 177.00 | 165.33 | -1.17 | 0.03 |
| 944.00 | 1083.00 | 1025.33 | 1.06 | 0.22 |
| 286.00 | 341.00 | 313.67 | 1.01 | 0.89 |
| 984.00 | 1148.00 | 1090.00 | 1.24 | 0.00 |
| 692.00 | 723.00 | 758.33 | 1.48 | 0.00 |
| 36.00 | 37.00 | 38.67 | 3.16 | 0.00 |
| 1899.00 | 2097.00 | 2054.67 | 1.11 | 0.01 |
| 2291.00 | 2462.00 | 2506.67 | 1.02 | 0.54 |
| 879.00 | 944.00 | 942.00 | 1.15 | 0.00 |
| 454.00 | 491.00 | 504.00 | 1.10 | 0.07 |
| 2980.00 | 3261.00 | 3113.67 | -1.68 | 0.00 |
| 120.00 | 165.00 | 135.33 | 2.96 | 0.00 |
| 868.00 | 914.00 | 939.67 | -1.09 | 0.06 |
| 676.00 | 712.00 | 704.33 | -1.00 | 0.96 |
| 28.00 | 62.00 | 45.67 | 1.84 | 0.00 |
| 475.00 | 471.00 | 476.67 | 1.11 | 0.03 |
| 766.00 | 762.00 | 779.33 | -1.10 | 0.03 |
| 562.00 | 599.00 | 572.67 | 1.16 | 0.00 |
| 17.00 | 16.00 | 15.67 | -1.61 | 0.02 |
| 18.00 | 20.00 | 17.67 | 1.24 | 0.33 |
| 12.00 | 10.00 | 10.67 | -1.64 | 0.04 |
| 4.00 | 2.00 | 2.33 | 1.36 | 0.78 |
| 0.00 | 0.00 | 0.33 | -2.49 | 0.64 |
| 71.00 | 106.00 | 86.67 | -1.13 | 0.25 |
| 1834.00 | 2272.00 | 2087.00 | -1.04 | 0.39 |
| 3915.00 | 4463.00 | 4367.67 | 1.41 | 0.00 |
| 536.00 | 564.00 | 571.33 | 1.03 | 0.65 |
| 946.00 | 1031.00 | 1027.67 | 1.19 | 0.00 |
| 1265.00 | 1268.00 | 1275.00 | 1.05 | 0.23 |
| 2561.00 | 2490.00 | 2538.67 | -1.20 | 0.00 |
| 146.00 | 124.00 | 137.00 | 1.22 | 0.02 |
| 430.00 | 508.00 | 482.67 | 1.37 | 0.00 |
| 1244.00 | 1438.00 | 1399.67 | 1.22 | 0.00 |
| 603.00 | 680.00 | 668.67 | -1.05 | 0.38 |
| 153.00 | 110.00 | 130.00 | 1.26 | 0.02 |
| 1289.00 | 1446.00 | 1460.00 | -1.20 | 0.00 |
| 564.00 | 599.00 | 607.33 | 1.17 | 0.00 |
| 7771.00 | 8822.00 | 8689.67 | 1.09 | 0.01 |
| 364.00 | 357.00 | 392.33 | 1.52 | 0.00 |
| 699.00 | 776.00 | 776.67 | 1.50 | 0.00 |
| 577.00 | 523.00 | 580.67 | 1.11 | 0.06 |
| 1969.00 | 2262.00 | 2096.33 | 1.40 | 0.00 |
| 1.00 | 2.00 | 1.00 | -2.21 | 0.35 |
| 10.00 | 11.00 | 11.33 | 1.02 | 1.00 |
| 306.00 | 315.00 | 323.33 | -1.33 | 0.00 |
| 1059.00 | 1109.00 | 1119.33 | -1.08 | 0.04 |
| 1944.00 | 1924.00 | 1980.00 | -1.09 | 0.02 |
| 3052.00 | 2993.00 | 3067.33 | 1.02 | 0.62 |

| | | | | |
|---|---|---|---|---|
| 1563.00 | 1644.00 | 1607.67 | -1.01 | 0.78 |
| 2333.00 | 2524.00 | 2379.33 | 1.02 | 0.60 |
| 2711.00 | 3005.00 | 3022.33 | 1.12 | 0.01 |
| 17.00 | 24.00 | 18.00 | -5.86 | 0.00 |
| 2951.00 | 3094.00 | 3104.33 | 1.60 | 0.00 |
| 52.00 | 49.00 | 54.67 | 1.42 | 0.01 |
| 135.00 | 143.00 | 139.67 | 1.09 | 0.32 |
| 3.00 | 1.00 | 2.00 | -1.33 | 0.79 |
| 1090.00 | 1179.00 | 1165.67 | -1.12 | 0.01 |
| 1157.00 | 1264.00 | 1141.00 | -1.01 | 0.94 |
| 345.00 | 384.00 | 376.67 | 1.13 | 0.03 |
| 113106.00 | 120887.00 | 122926.67 | 1.18 | 0.00 |
| 6.00 | 5.00 | 7.33 | -1.60 | 0.12 |
| 6796.00 | 7655.00 | 7842.33 | 1.19 | 0.00 |
| 15.00 | 13.00 | 19.00 | -1.07 | 0.82 |
| 10.00 | 7.00 | 6.00 | 1.60 | 0.29 |
| 514.00 | 608.00 | 571.67 | 1.37 | 0.00 |
| 24995.00 | 26908.00 | 27305.33 | 1.15 | 0.00 |
| 66894.00 | 64842.00 | 66668.33 | -1.01 | 0.82 |
| 1064.00 | 1059.00 | 1072.33 | 1.17 | 0.00 |
| 339.00 | 403.00 | 393.33 | 1.11 | 0.15 |
| 322.00 | 375.00 | 368.00 | -1.05 | 0.40 |
| 74.00 | 73.00 | 83.00 | 1.58 | 0.00 |
| 173.00 | 144.00 | 158.67 | -1.00 | 1.00 |
| 2.00 | 1.00 | 1.33 | -1.94 | 0.39 |
| 185.00 | 195.00 | 201.33 | -1.22 | 0.00 |
| 1712.00 | 1720.00 | 1695.67 | -1.02 | 0.71 |
| 119.00 | 131.00 | 131.00 | 1.11 | 0.20 |
| 8.00 | 14.00 | 10.00 | 1.40 | 0.27 |
| 195.00 | 272.00 | 232.67 | 1.16 | 0.06 |
| 1487.00 | 1716.00 | 1671.00 | 1.38 | 0.00 |
| 494.00 | 488.00 | 519.67 | -1.09 | 0.12 |
| 1226.00 | 1374.00 | 1342.33 | -1.08 | 0.05 |
| 11.00 | 15.00 | 13.00 | 1.16 | 0.66 |
| 2757.00 | 2858.00 | 2923.33 | 1.24 | 0.00 |
| 1.00 | 1.00 | 2.00 | -1.49 | 0.62 |
| 28.00 | 30.00 | 31.00 | 1.08 | 0.72 |
| 108.00 | 96.00 | 100.67 | -1.05 | 0.59 |
| 305.00 | 397.00 | 373.00 | -1.02 | 0.82 |
| 762.00 | 795.00 | 770.67 | -1.43 | 0.00 |
| 268.00 | 265.00 | 265.67 | -1.21 | 0.00 |
| 11.00 | 5.00 | 8.00 | -1.26 | 0.51 |
| 1841.00 | 1933.00 | 1854.00 | 1.05 | 0.29 |
| 1058.00 | 1097.00 | 1081.33 | 1.05 | 0.24 |
| 3.00 | 3.00 | 2.67 | -1.74 | 0.29 |
| 3528.00 | 3830.00 | 3752.67 | 1.06 | 0.08 |
| 13.00 | 11.00 | 9.33 | -2.12 | 0.00 |
| 3.00 | 8.00 | 5.33 | 1.21 | 0.72 |

| | | | | |
|---|---|---|---|---|
| 27.00 | 28.00 | 29.67 | 1.03 | 0.88 |
| 8363.00 | 6619.00 | 6993.33 | -1.39 | 0.00 |
| 1236.00 | 1345.00 | 1331.33 | 1.15 | 0.00 |
| 24.00 | 20.00 | 25.00 | 1.54 | 0.03 |
| 1669.00 | 1577.00 | 1711.33 | 1.18 | 0.00 |
| 23.00 | 22.00 | 20.67 | -1.03 | 1.00 |
| 1787.00 | 1832.00 | 1828.67 | 1.05 | 0.19 |
| 2493.00 | 2670.00 | 2612.33 | -1.02 | 0.71 |
| 2561.00 | 2571.00 | 2615.67 | 1.15 | 0.00 |
| 909.00 | 781.00 | 849.33 | -1.38 | 0.00 |
| 12.00 | 18.00 | 17.33 | 1.14 | 0.63 |
| 2980.00 | 2866.00 | 2973.67 | 1.39 | 0.00 |
| 0.00 | 3.00 | 1.33 | -1.48 | 0.76 |
| 72.00 | 106.00 | 88.00 | 1.21 | 0.08 |
| 1375.00 | 1442.00 | 1406.00 | -1.06 | 0.14 |
| 1234.00 | 1205.00 | 1246.33 | 1.17 | 0.00 |
| 2061.00 | 2466.00 | 2376.00 | 1.03 | 0.56 |
| 166.00 | 203.00 | 198.00 | -1.04 | 0.54 |
| 77.00 | 83.00 | 82.33 | 1.14 | 0.19 |
| 27.00 | 47.00 | 34.33 | 1.19 | 0.33 |
| 13704.00 | 15438.00 | 15534.00 | 1.11 | 0.01 |
| 771.00 | 708.00 | 750.00 | -1.23 | 0.00 |
| 2121.00 | 2483.00 | 2335.67 | 1.21 | 0.00 |
| 1137.00 | 1280.00 | 1291.00 | 1.12 | 0.01 |
| 11.00 | 14.00 | 13.00 | 1.28 | 0.35 |
| 3009.00 | 3383.00 | 3406.67 | 1.10 | 0.02 |
| 2327.00 | 2374.00 | 2384.33 | -1.01 | 0.80 |
| 566.00 | 534.00 | 520.00 | 1.01 | 0.92 |
| 487.00 | 451.00 | 471.33 | -1.11 | 0.05 |
| 2495.00 | 2531.00 | 2526.67 | -1.05 | 0.18 |
| 1207.00 | 1239.00 | 1270.33 | 1.11 | 0.01 |
| 1070.00 | 1050.00 | 1059.67 | 1.04 | 0.41 |
| 754.00 | 902.00 | 855.00 | 1.23 | 0.00 |
| 750.00 | 829.00 | 828.00 | -1.11 | 0.02 |
| 1557.00 | 1567.00 | 1601.33 | -1.08 | 0.04 |
| 1879.00 | 1946.00 | 1960.67 | -1.01 | 0.79 |
| 2462.00 | 2821.00 | 2763.00 | 1.30 | 0.00 |
| 3717.00 | 4199.00 | 4096.67 | -1.26 | 0.00 |
| 9848.00 | 10641.00 | 10614.33 | 1.02 | 0.55 |
| 14152.00 | 14662.00 | 14023.67 | -1.01 | 0.90 |
| 7152.00 | 7480.00 | 7524.33 | 1.10 | 0.00 |
| 6985.00 | 7588.00 | 7569.33 | 1.15 | 0.00 |
| 11636.00 | 12808.00 | 12838.33 | 1.15 | 0.00 |
| 2075.00 | 2288.00 | 2310.33 | 1.15 | 0.00 |
| 3265.00 | 3557.00 | 3551.67 | 1.12 | 0.00 |
| 5269.00 | 5910.00 | 5945.33 | 1.19 | 0.00 |
| 3881.00 | 4222.00 | 4344.00 | 1.10 | 0.03 |
| 1951.00 | 2291.00 | 2232.33 | 1.12 | 0.01 |

| | | | | |
|---|---|---|---|---|
| 281.00 | 287.00 | 304.33 | 1.01 | 0.95 |
| 2104.00 | 2288.00 | 2335.67 | 1.21 | 0.00 |
| 5900.00 | 6225.00 | 6295.67 | 1.09 | 0.02 |
| 4822.00 | 5357.00 | 5343.33 | 1.23 | 0.00 |
| 13374.00 | 15657.00 | 15124.33 | 1.02 | 0.56 |
| 2341.00 | 2447.00 | 2488.67 | 1.11 | 0.00 |
| 8764.00 | 8896.00 | 8891.33 | 1.06 | 0.17 |
| 499.00 | 509.00 | 523.67 | 1.19 | 0.00 |
| 922.00 | 904.00 | 947.00 | 1.09 | 0.05 |
| 25.00 | 25.00 | 25.67 | 1.05 | 0.88 |
| 3694.00 | 3929.00 | 4025.67 | 1.19 | 0.00 |
| 1587.00 | 1581.00 | 1579.33 | 1.09 | 0.04 |
| 49.00 | 52.00 | 62.00 | 1.41 | 0.01 |
| 539.00 | 543.00 | 544.67 | -1.04 | 0.39 |
| 18393.00 | 18596.00 | 18615.00 | -1.03 | 0.39 |
| 17907.00 | 19251.00 | 18938.67 | 1.03 | 0.36 |
| 3461.00 | 3560.00 | 3573.33 | -1.11 | 0.00 |
| 5054.00 | 5052.00 | 5284.33 | 1.08 | 0.03 |
| 5156.00 | 5397.00 | 5411.33 | 1.01 | 0.86 |
| 11719.00 | 12243.00 | 12723.00 | 1.20 | 0.00 |
| 296.00 | 280.00 | 291.33 | 1.02 | 0.80 |
| 346.00 | 357.00 | 389.67 | 1.22 | 0.00 |
| 5832.00 | 6757.00 | 6534.67 | 1.19 | 0.00 |
| 4441.00 | 4620.00 | 4857.67 | 1.09 | 0.04 |
| 157.00 | 192.00 | 191.67 | 1.10 | 0.32 |
| 1561.00 | 1696.00 | 1698.33 | 1.01 | 0.81 |
| 2139.00 | 2045.00 | 2161.33 | 1.17 | 0.00 |
| 59.00 | 76.00 | 70.00 | 1.20 | 0.12 |
| 527.00 | 564.00 | 550.33 | -1.15 | 0.01 |
| 264.00 | 367.00 | 324.67 | 1.05 | 0.49 |
| 292.00 | 287.00 | 291.67 | -1.20 | 0.00 |
| 2.00 | 12.00 | 7.33 | -1.06 | 1.00 |
| 63.00 | 74.00 | 68.00 | 1.43 | 0.00 |
| 2041.00 | 1980.00 | 2012.00 | -1.54 | 0.00 |
| 780.00 | 766.00 | 814.33 | 1.06 | 0.19 |
| 6283.00 | 6796.00 | 6591.00 | 1.24 | 0.00 |
| 1681.00 | 1862.00 | 1798.00 | -1.15 | 0.00 |
| 726.00 | 708.00 | 730.67 | 1.03 | 0.52 |
| 3470.00 | 4134.00 | 3810.33 | 1.17 | 0.00 |
| 12.00 | 15.00 | 14.33 | 1.06 | 0.83 |
| 843.00 | 905.00 | 865.67 | -1.01 | 0.78 |
| 100.00 | 136.00 | 124.67 | -1.12 | 0.20 |
| 160.00 | 132.00 | 153.33 | 1.11 | 0.24 |
| 777.00 | 961.00 | 821.67 | 1.10 | 0.17 |
| 528.00 | 617.00 | 610.67 | -1.10 | 0.08 |
| 802.00 | 813.00 | 795.00 | -1.12 | 0.04 |
| 581.00 | 599.00 | 643.67 | -1.05 | 0.36 |
| 3610.00 | 3960.00 | 4003.67 | 1.07 | 0.11 |

| | | | | |
|---|---|---|---|---|
| 290.00 | 319.00 | 312.00 | 1.58 | 0.00 |
| 19.00 | 26.00 | 22.00 | 1.91 | 0.00 |
| 535.00 | 538.00 | 551.33 | 1.05 | 0.29 |
| 4638.00 | 4939.00 | 4848.33 | 1.15 | 0.00 |
| 1471.00 | 1623.00 | 1545.00 | 1.08 | 0.12 |
| 60.00 | 74.00 | 65.33 | 1.07 | 0.57 |
| 1321.00 | 1396.00 | 1379.33 | 1.04 | 0.27 |
| 633.00 | 654.00 | 649.67 | 1.16 | 0.00 |
| 243.00 | 285.00 | 274.00 | 1.09 | 0.21 |
| 1334.00 | 1433.00 | 1475.67 | 1.14 | 0.00 |
| 606.00 | 740.00 | 719.00 | 1.11 | 0.07 |
| 4.00 | 5.00 | 4.33 | -1.32 | 0.59 |
| 3887.00 | 4368.00 | 4086.67 | -1.14 | 0.00 |
| 1184.00 | 1134.00 | 1155.67 | -1.09 | 0.09 |
| 781.00 | 923.00 | 900.00 | 1.18 | 0.00 |
| 931.00 | 943.00 | 1019.67 | -1.10 | 0.09 |
| 348.00 | 421.00 | 417.67 | -1.06 | 0.42 |
| 1939.00 | 1979.00 | 2090.33 | 1.23 | 0.00 |
| 582.00 | 689.00 | 668.67 | 1.12 | 0.03 |
| 1980.00 | 1912.00 | 1992.67 | 1.17 | 0.00 |
| 639.00 | 673.00 | 686.33 | -1.09 | 0.07 |
| 187.00 | 175.00 | 202.67 | -1.10 | 0.25 |
| 1148.00 | 1266.00 | 1319.00 | 1.10 | 0.06 |
| 691.00 | 836.00 | 782.33 | 1.02 | 0.63 |
| 202.00 | 187.00 | 195.67 | -1.27 | 0.00 |
| 1036.00 | 1086.00 | 1087.33 | 1.19 | 0.00 |
| 155.00 | 156.00 | 165.00 | -1.59 | 0.00 |
| 761.00 | 760.00 | 770.67 | 1.10 | 0.04 |
| 82.00 | 84.00 | 81.33 | -1.18 | 0.11 |
| 843.00 | 824.00 | 882.33 | -1.16 | 0.00 |
| 118.00 | 102.00 | 115.67 | -1.76 | 0.00 |
| 1714.00 | 1779.00 | 1794.33 | -1.12 | 0.00 |
| 1427.00 | 1486.00 | 1473.67 | -1.01 | 0.90 |
| 73.00 | 64.00 | 68.67 | 1.02 | 0.86 |
| 155.00 | 136.00 | 148.33 | -1.41 | 0.00 |
| 32631.00 | 35618.00 | 35067.00 | 1.45 | 0.00 |
| 213.00 | 258.00 | 249.00 | 1.31 | 0.00 |
| 1846.00 | 1874.00 | 1949.33 | 1.20 | 0.00 |
| 1687.00 | 1873.00 | 1745.33 | 1.02 | 0.77 |
| 4988.00 | 5524.00 | 5429.00 | 1.05 | 0.20 |
| 718.00 | 710.00 | 687.33 | -1.21 | 0.00 |
| 164.00 | 161.00 | 177.00 | 1.09 | 0.36 |
| 3.00 | 7.00 | 7.00 | 1.46 | 0.34 |
| 311.00 | 295.00 | 317.67 | -1.31 | 0.00 |
| 12014.00 | 11995.00 | 12291.00 | -1.31 | 0.00 |
| 908.00 | 890.00 | 931.33 | -1.17 | 0.00 |
| 6.00 | 10.00 | 8.67 | 2.51 | 0.01 |
| 495.00 | 579.00 | 562.33 | 1.09 | 0.13 |

| | | | | |
|---|---|---|---|---|
| 14.00 | 17.00 | 14.67 | 1.49 | 0.11 |
| 33084.00 | 34386.00 | 35228.00 | 1.04 | 0.25 |
| 3.00 | 7.00 | 4.67 | -1.01 | 1.00 |
| 1212.00 | 1354.00 | 1375.33 | -1.05 | 0.38 |
| 596.00 | 649.00 | 685.67 | 1.03 | 0.61 |
| 805.00 | 938.00 | 903.33 | 1.18 | 0.00 |
| 3528.00 | 3900.00 | 3839.67 | 1.05 | 0.14 |
| 83.00 | 88.00 | 89.00 | 1.73 | 0.00 |
| 209.00 | 209.00 | 227.33 | -1.04 | 0.54 |
| 89.00 | 124.00 | 102.33 | -1.18 | 0.08 |
| 1741.00 | 1911.00 | 1813.00 | 1.00 | 0.92 |
| 89.00 | 103.00 | 96.67 | 2.00 | 0.00 |
| 3.00 | 8.00 | 4.67 | 1.15 | 0.85 |
| 331.00 | 406.00 | 347.00 | -1.03 | 0.71 |
| 1728.00 | 1892.00 | 1884.33 | 1.14 | 0.00 |
| 1024.00 | 997.00 | 1062.67 | -1.12 | 0.01 |
| 81.00 | 87.00 | 84.33 | 1.03 | 0.80 |
| 158.00 | 153.00 | 145.33 | -3.43 | 0.00 |
| 26.00 | 32.00 | 33.00 | 1.37 | 0.07 |
| 3474.00 | 3517.00 | 3519.00 | -1.02 | 0.60 |
| 264.00 | 261.00 | 247.00 | -1.04 | 0.63 |
| 2881.00 | 2901.00 | 2975.33 | -1.10 | 0.01 |
| 1106.00 | 908.00 | 1003.33 | 1.10 | 0.13 |
| 912.00 | 1154.00 | 1111.33 | 1.09 | 0.11 |
| 1271.00 | 1448.00 | 1382.67 | 1.10 | 0.02 |
| 1739.00 | 1634.00 | 1655.67 | -1.34 | 0.00 |
| 3.00 | 5.00 | 3.67 | 1.08 | 1.00 |
| 1822.00 | 1782.00 | 1794.67 | -1.26 | 0.00 |
| 187.00 | 211.00 | 200.33 | -1.35 | 0.00 |
| 298.00 | 260.00 | 281.33 | -1.32 | 0.00 |
| 868.00 | 793.00 | 824.33 | -1.17 | 0.00 |
| 536.00 | 491.00 | 529.67 | -1.05 | 0.40 |
| 3749.00 | 4315.00 | 4035.33 | 1.14 | 0.00 |
| 1911.00 | 2007.00 | 1954.33 | 1.25 | 0.00 |
| 11.00 | 13.00 | 14.67 | -1.17 | 0.51 |
| 726.00 | 741.00 | 770.67 | 1.09 | 0.06 |
| 97.00 | 118.00 | 120.00 | 1.12 | 0.25 |
| 441.00 | 468.00 | 454.67 | -1.06 | 0.23 |
| 713.00 | 754.00 | 739.67 | -1.09 | 0.08 |
| 507.00 | 521.00 | 504.67 | -1.02 | 0.74 |
| 1.00 | 0.00 | 2.33 | 1.65 | 0.60 |
| 20.00 | 21.00 | 16.33 | 1.46 | 0.13 |
| 6539.00 | 6642.00 | 6632.33 | 1.07 | 0.07 |
| 3.00 | 6.00 | 5.00 | -1.34 | 0.51 |
| 1.00 | 2.00 | 1.00 | -2.21 | 0.35 |
| 13229.00 | 13380.00 | 13418.00 | -1.25 | 0.00 |
| 3178.00 | 3342.00 | 3273.00 | 1.49 | 0.00 |
| 168.00 | 184.00 | 161.33 | 1.40 | 0.00 |

| | | | | |
|---|---|---|---|---|
| 14.00 | 21.00 | 20.00 | 3.43 | 0.00 |
| 30.00 | 41.00 | 34.00 | 1.24 | 0.20 |
| 169.00 | 141.00 | 157.33 | 1.28 | 0.00 |
| 1927.00 | 1797.00 | 1883.67 | -1.70 | 0.00 |
| 1931.00 | 1943.00 | 1965.67 | -1.01 | 0.75 |
| 6.00 | 10.00 | 7.67 | -2.23 | 0.00 |
| 407.00 | 412.00 | 433.33 | -1.29 | 0.00 |
| 0.00 | 5.00 | 3.67 | 1.08 | 1.00 |
| 269.00 | 264.00 | 273.00 | 1.21 | 0.01 |
| 116.00 | 119.00 | 115.67 | -1.19 | 0.04 |
| 1063.00 | 1220.00 | 1140.67 | -1.22 | 0.00 |
| 1897.00 | 1989.00 | 1999.67 | 1.03 | 0.35 |
| 1024.00 | 998.00 | 1036.33 | -1.26 | 0.00 |
| 295.00 | 252.00 | 281.67 | -1.15 | 0.04 |
| 1.00 | 1.00 | 0.67 | -4.86 | 0.03 |
| 9113.00 | 9900.00 | 9551.00 | -1.03 | 0.35 |
| 1280.00 | 1414.00 | 1378.33 | 1.09 | 0.04 |
| 798.00 | 888.00 | 827.00 | -1.03 | 0.61 |
| 450.00 | 481.00 | 477.00 | -1.06 | 0.30 |
| 16.00 | 10.00 | 15.33 | 1.03 | 0.92 |
| 2259.00 | 2403.00 | 2284.33 | -1.03 | 0.48 |
| 1119.00 | 1205.00 | 1223.00 | 1.08 | 0.09 |
| 2479.00 | 2653.00 | 2511.67 | 1.01 | 0.86 |
| 254.00 | 290.00 | 275.00 | -1.09 | 0.18 |
| 48.00 | 61.00 | 61.67 | 1.28 | 0.09 |
| 1876.00 | 2023.00 | 1985.00 | -1.05 | 0.25 |
| 1.00 | 2.00 | 2.00 | -2.60 | 0.06 |
| 21.00 | 34.00 | 25.00 | 1.09 | 0.71 |
| 1458.00 | 1409.00 | 1380.00 | -1.17 | 0.00 |
| 2047.00 | 2131.00 | 2206.67 | -1.11 | 0.01 |
| 1672.00 | 1546.00 | 1606.00 | -1.33 | 0.00 |
| 1504.00 | 1512.00 | 1553.33 | 1.03 | 0.46 |
| 451.00 | 500.00 | 458.67 | -1.09 | 0.17 |
| 73.00 | 75.00 | 65.67 | 1.06 | 0.61 |
| 306.00 | 349.00 | 313.33 | 1.04 | 0.62 |
| 428.00 | 462.00 | 441.67 | 1.46 | 0.00 |
| 551.00 | 551.00 | 547.00 | -1.07 | 0.21 |
| 235.00 | 277.00 | 264.00 | 1.05 | 0.42 |
| 2540.00 | 2965.00 | 2671.00 | 1.37 | 0.00 |
| 1491.00 | 1378.00 | 1478.33 | 1.05 | 0.23 |
| 7954.00 | 7393.00 | 7817.67 | 1.05 | 0.25 |
| 1907.00 | 1996.00 | 1982.67 | 1.12 | 0.00 |
| 5717.00 | 5761.00 | 5896.67 | 1.04 | 0.33 |
| 1798.00 | 2147.00 | 2179.33 | 1.35 | 0.00 |
| 594.00 | 710.00 | 652.00 | 1.16 | 0.01 |
| 483.00 | 593.00 | 555.67 | 1.15 | 0.01 |
| 1160.00 | 1198.00 | 1244.00 | 1.03 | 0.57 |
| 26.00 | 22.00 | 30.00 | -1.80 | 0.00 |

| | | | | |
|---|---|---|---|---|
| 229.00 | 274.00 | 254.00 | 1.08 | 0.29 |
| 187.00 | 227.00 | 216.67 | 1.22 | 0.00 |
| 1.00 | 0.00 | 0.67 | -3.14 | 0.19 |
| 2.00 | 5.00 | 2.33 | -1.97 | 0.20 |
| 37.00 | 73.00 | 52.33 | 1.04 | 0.83 |
| 1518.00 | 1524.00 | 1541.00 | 1.03 | 0.45 |
| 3184.00 | 3145.00 | 3237.67 | 1.19 | 0.00 |
| 3671.00 | 3834.00 | 3745.00 | -1.15 | 0.00 |
| 493.00 | 465.00 | 488.00 | 1.10 | 0.10 |
| 1023.00 | 1061.00 | 1052.67 | -1.07 | 0.11 |
| 890.00 | 665.00 | 708.33 | -1.43 | 0.00 |
| 3948.00 | 4034.00 | 4010.00 | 1.16 | 0.00 |
| 288.00 | 253.00 | 269.00 | -1.15 | 0.05 |
| 6126.00 | 6627.00 | 6657.00 | 1.14 | 0.00 |
| 1680.00 | 1922.00 | 1855.33 | 1.05 | 0.24 |
| 15064.00 | 16102.00 | 16224.33 | -1.09 | 0.01 |
| 13.00 | 27.00 | 20.67 | -1.26 | 0.23 |
| 185.00 | 190.00 | 197.00 | -1.21 | 0.00 |
| 28.00 | 9.00 | 17.33 | 1.39 | 0.21 |
| 77.00 | 73.00 | 89.33 | -2.28 | 0.00 |
| 359.00 | 408.00 | 380.33 | 1.05 | 0.43 |
| 92.00 | 110.00 | 102.33 | 1.33 | 0.00 |
| 1464.00 | 1786.00 | 1745.00 | 1.11 | 0.03 |
| 1559.00 | 1767.00 | 1633.00 | 1.12 | 0.01 |
| 24404.00 | 26826.00 | 26262.33 | 1.43 | 0.00 |
| 1792.00 | 1873.00 | 1839.67 | 1.18 | 0.00 |
| 8.00 | 7.00 | 8.33 | 1.63 | 0.16 |
| 20.00 | 26.00 | 23.00 | 1.41 | 0.09 |
| 26.00 | 17.00 | 21.67 | -1.04 | 0.87 |
| 11.00 | 2.00 | 6.67 | -1.96 | 0.02 |
| 16.00 | 24.00 | 21.67 | 1.68 | 0.02 |
| 1902.00 | 2144.00 | 2126.33 | -1.06 | 0.12 |
| 16.00 | 11.00 | 14.67 | 1.49 | 0.11 |
| 3449.00 | 3671.00 | 3582.67 | -1.17 | 0.00 |
| 4692.00 | 4495.00 | 4518.33 | 1.05 | 0.32 |
| 3.00 | 5.00 | 3.00 | -1.55 | 0.41 |
| 420.00 | 498.00 | 493.00 | -1.11 | 0.08 |
| 4015.00 | 4684.00 | 4480.00 | 1.19 | 0.00 |
| 1666.00 | 1830.00 | 1783.00 | 1.08 | 0.06 |
| 629.00 | 628.00 | 665.67 | -1.01 | 0.82 |
| 331.00 | 409.00 | 389.00 | -1.00 | 1.00 |
| 346.00 | 433.00 | 455.67 | -1.13 | 0.16 |
| 2140.00 | 2946.00 | 2499.67 | 1.01 | 0.89 |
| 5116.00 | 5115.00 | 5123.33 | 1.01 | 0.81 |
| 1267.00 | 1313.00 | 1301.00 | 1.04 | 0.39 |
| 1125.00 | 1203.00 | 1127.00 | 1.15 | 0.01 |
| 3.00 | 4.00 | 5.33 | -1.14 | 0.87 |
| 149.00 | 173.00 | 148.33 | -1.19 | 0.09 |

| | | | | |
|---|---|---|---|---|
| 983.00 | 925.00 | 951.67 | 1.22 | 0.00 |
| 2805.00 | 3001.00 | 2973.00 | 1.12 | 0.01 |
| 553.00 | 589.00 | 601.67 | -1.17 | 0.00 |
| 1842.00 | 1926.00 | 1931.00 | 1.34 | 0.00 |
| 317.00 | 277.00 | 316.33 | -1.06 | 0.43 |
| 68.00 | 71.00 | 70.33 | 1.02 | 0.85 |
| 4595.00 | 3442.00 | 3836.67 | -1.54 | 0.00 |
| 606.00 | 656.00 | 692.33 | -1.06 | 0.35 |
| 176.00 | 210.00 | 215.67 | -1.07 | 0.41 |
| 689.00 | 750.00 | 733.00 | -1.01 | 0.82 |
| 542.00 | 600.00 | 591.67 | 1.06 | 0.28 |
| 466.00 | 454.00 | 488.33 | 1.06 | 0.30 |
| 7899.00 | 8822.00 | 8697.33 | 1.08 | 0.04 |
| 2.00 | 4.00 | 4.67 | -1.09 | 1.00 |
| 711.00 | 699.00 | 717.00 | -1.18 | 0.00 |
| 373.00 | 342.00 | 372.00 | -1.31 | 0.00 |
| 316.00 | 385.00 | 350.00 | 1.40 | 0.00 |
| 93.00 | 95.00 | 92.33 | 1.04 | 0.70 |
| 908.00 | 1051.00 | 991.00 | 1.05 | 0.35 |
| 1251.00 | 1364.00 | 1324.00 | -1.08 | 0.05 |
| 1064.00 | 1007.00 | 1055.67 | -1.14 | 0.00 |
| 75.00 | 67.00 | 79.00 | 1.06 | 0.64 |
| 1877.00 | 1773.00 | 1824.00 | -1.05 | 0.27 |
| 1003.00 | 1213.00 | 1068.00 | 1.13 | 0.04 |
| 302.00 | 344.00 | 315.00 | 1.09 | 0.18 |
| 194.00 | 164.00 | 167.00 | -1.29 | 0.01 |
| 1801.00 | 1992.00 | 1968.67 | -1.27 | 0.00 |
| 507.00 | 541.00 | 529.00 | -1.14 | 0.00 |
| 580.00 | 652.00 | 619.67 | -1.03 | 0.57 |
| 688.00 | 719.00 | 790.00 | 1.08 | 0.22 |
| 2038.00 | 2187.00 | 2196.67 | -1.08 | 0.03 |
| 1165.00 | 1259.00 | 1229.67 | 1.08 | 0.05 |
| 799.00 | 900.00 | 913.67 | 1.16 | 0.00 |
| 446.00 | 462.00 | 495.67 | 1.03 | 0.58 |
| 507.00 | 666.00 | 614.33 | 1.07 | 0.27 |
| 380.00 | 400.00 | 387.67 | 1.13 | 0.03 |
| 905.00 | 1017.00 | 1039.00 | 1.16 | 0.01 |
| 1213.00 | 1399.00 | 1382.33 | 1.23 | 0.00 |
| 598.00 | 761.00 | 683.67 | 1.13 | 0.04 |
| 109.00 | 112.00 | 112.33 | -1.21 | 0.02 |
| 4739.00 | 4741.00 | 4866.33 | -1.20 | 0.00 |
| 2941.00 | 3085.00 | 3097.33 | -1.18 | 0.00 |
| 2628.00 | 2872.00 | 2833.67 | 1.07 | 0.10 |
| 1718.00 | 1830.00 | 1742.67 | -1.18 | 0.00 |
| 9.00 | 9.00 | 7.33 | -1.06 | 1.00 |
| 5.00 | 7.00 | 6.33 | -1.12 | 0.88 |
| 664.00 | 711.00 | 696.00 | -1.07 | 0.16 |
| 742.00 | 855.00 | 824.33 | -1.12 | 0.01 |

| | | | | |
|---|---|---|---|---|
| 6.00 | 14.00 | 12.00 | 1.85 | 0.04 |
| 2146.00 | 2282.00 | 2297.67 | -1.12 | 0.00 |
| 1008.00 | 1187.00 | 1158.33 | 1.37 | 0.00 |
| 3025.00 | 3549.00 | 3289.33 | 1.19 | 0.00 |
| 14.00 | 13.00 | 14.67 | -1.11 | 0.79 |
| 145.00 | 154.00 | 149.33 | 1.31 | 0.00 |
| 82.00 | 70.00 | 77.33 | 1.07 | 0.56 |
| 2388.00 | 2547.00 | 2473.33 | -1.71 | 0.00 |
| 483.00 | 510.00 | 489.33 | -2.21 | 0.00 |
| 163.00 | 172.00 | 170.33 | 1.03 | 0.72 |
| 523.00 | 546.00 | 543.33 | -1.44 | 0.00 |
| 88.00 | 93.00 | 96.00 | 1.09 | 0.37 |
| 43.00 | 41.00 | 47.33 | 1.09 | 0.54 |
| 8.00 | 10.00 | 9.33 | -1.55 | 0.11 |
| 0.00 | 0.00 | 0.00 | -11.74 | 0.13 |
| 913.00 | 960.00 | 984.00 | 1.06 | 0.21 |
| 1192.00 | 1179.00 | 1225.67 | -1.03 | 0.42 |
| 73.00 | 91.00 | 77.00 | 1.21 | 0.09 |
| 916.00 | 1026.00 | 1088.33 | 2.40 | 0.00 |
| 39.00 | 42.00 | 46.33 | 3.24 | 0.00 |
| 3728.00 | 3820.00 | 3943.00 | 1.15 | 0.00 |
| 3556.00 | 3349.00 | 3490.33 | -1.03 | 0.41 |
| 977.00 | 1011.00 | 1003.33 | -1.07 | 0.13 |
| 462.00 | 466.00 | 492.67 | -1.63 | 0.00 |
| 1717.00 | 1882.00 | 1885.67 | 1.25 | 0.00 |
| 2565.00 | 2413.00 | 2523.33 | -1.04 | 0.42 |
| 1002.00 | 1088.00 | 1121.00 | -1.01 | 0.89 |
| 418.00 | 468.00 | 450.67 | 1.04 | 0.51 |
| 599.00 | 606.00 | 653.67 | -1.15 | 0.02 |
| 337.00 | 335.00 | 345.67 | 1.14 | 0.02 |
| 72.00 | 89.00 | 86.33 | 1.04 | 0.77 |
| 219.00 | 232.00 | 235.33 | 1.02 | 0.80 |
| 13.00 | 7.00 | 8.67 | -1.40 | 0.32 |
| 692.00 | 731.00 | 735.67 | 1.24 | 0.00 |
| 752.00 | 657.00 | 734.67 | -1.24 | 0.00 |
| 239.00 | 254.00 | 254.67 | 1.15 | 0.02 |
| 173.00 | 183.00 | 180.33 | 1.09 | 0.30 |
| 646.00 | 690.00 | 693.67 | -1.03 | 0.53 |
| 182.00 | 229.00 | 247.00 | 1.30 | 0.02 |
| 29.00 | 33.00 | 31.67 | -1.85 | 0.00 |
| 1.00 | 0.00 | 1.33 | -2.87 | 0.08 |
| 1.00 | 0.00 | 2.00 | -2.44 | 0.08 |
| 28.00 | 31.00 | 28.33 | -1.78 | 0.00 |
| 2.00 | 5.00 | 4.33 | -1.47 | 0.39 |
| 705.00 | 817.00 | 782.00 | -1.36 | 0.00 |
| 217.00 | 203.00 | 211.67 | -1.39 | 0.00 |
| 99.00 | 95.00 | 95.67 | -1.14 | 0.27 |
| 168.00 | 162.00 | 178.00 | 1.28 | 0.01 |

| | | | | |
|---|---|---|---|---|
| 622.00 | 755.00 | 718.33 | 1.28 | 0.00 |
| 115.00 | 140.00 | 135.67 | 1.77 | 0.00 |
| 1260.00 | 1199.00 | 1263.33 | -1.12 | 0.00 |
| 611.00 | 617.00 | 637.67 | 1.16 | 0.00 |
| 7.00 | 6.00 | 7.00 | -1.06 | 1.00 |
| 221.00 | 213.00 | 218.33 | 1.14 | 0.06 |
| 373.00 | 396.00 | 376.00 | -1.04 | 0.54 |
| 243.00 | 243.00 | 256.33 | 1.01 | 0.87 |
| 4530.00 | 4804.00 | 4645.67 | 1.00 | 0.91 |
| 296.00 | 404.00 | 378.00 | 1.12 | 0.12 |
| 350.00 | 389.00 | 383.33 | -1.05 | 0.34 |
| 287.00 | 270.00 | 277.67 | -1.14 | 0.07 |
| 931.00 | 1219.00 | 1095.00 | 1.06 | 0.27 |
| 32.00 | 37.00 | 35.33 | 1.34 | 0.08 |
| 36.00 | 21.00 | 32.33 | 1.18 | 0.34 |
| 1278.00 | 1493.00 | 1374.00 | 1.24 | 0.00 |
| 306.00 | 413.00 | 356.33 | -1.06 | 0.36 |
| 457.00 | 536.00 | 538.33 | -1.13 | 0.04 |
| 448.00 | 484.00 | 488.00 | -1.06 | 0.24 |
| 145.00 | 144.00 | 150.33 | -1.13 | 0.09 |
| 1821.00 | 1679.00 | 1705.33 | 1.24 | 0.00 |
| 10634.00 | 10806.00 | 11010.00 | -1.13 | 0.00 |
| 7.00 | 3.00 | 5.67 | 1.28 | 0.59 |
| 740.00 | 786.00 | 796.33 | -1.02 | 0.71 |
| 29.00 | 23.00 | 27.67 | -1.10 | 0.61 |
| 10.00 | 11.00 | 10.00 | -1.35 | 0.29 |
| 69.00 | 71.00 | 72.33 | -1.14 | 0.20 |
| 70.00 | 93.00 | 78.67 | -1.10 | 0.49 |
| 111.00 | 100.00 | 108.33 | 1.76 | 0.00 |
| 269.00 | 336.00 | 320.00 | 1.08 | 0.30 |
| 13.00 | 18.00 | 18.33 | 1.46 | 0.10 |
| 22.00 | 14.00 | 15.00 | 1.04 | 0.92 |
| 2470.00 | 2373.00 | 2468.67 | -1.01 | 0.88 |
| 43.00 | 46.00 | 44.00 | 1.11 | 0.47 |
| 1101.00 | 1073.00 | 1110.00 | -1.08 | 0.07 |
| 414.00 | 449.00 | 427.00 | -1.02 | 0.73 |
| 1475.00 | 1709.00 | 1684.00 | 1.35 | 0.00 |
| 6.00 | 6.00 | 5.67 | -1.83 | 0.07 |
| 5.00 | 6.00 | 5.67 | 1.83 | 0.18 |
| 397.00 | 451.00 | 419.33 | 1.00 | 0.97 |
| 27.00 | 24.00 | 27.00 | -1.45 | 0.02 |
| 671.00 | 747.00 | 738.33 | -1.14 | 0.00 |
| 22.00 | 15.00 | 22.00 | 1.23 | 0.34 |
| 462.00 | 468.00 | 496.00 | 1.02 | 0.77 |
| 21.00 | 31.00 | 25.00 | 1.04 | 0.94 |
| 259.00 | 287.00 | 274.00 | -1.03 | 0.71 |
| 22.00 | 22.00 | 24.00 | 1.20 | 0.36 |
| 29.00 | 26.00 | 25.00 | -1.27 | 0.17 |

| | | | | |
|---|---|---|---|---|
| 815.00 | 820.00 | 833.00 | -1.18 | 0.00 |
| 608.00 | 699.00 | 638.00 | -1.01 | 0.87 |
| 928.00 | 931.00 | 997.00 | -1.19 | 0.00 |
| 284.00 | 337.00 | 312.33 | -1.10 | 0.10 |
| 90.00 | 108.00 | 102.33 | -1.11 | 0.28 |
| 767.00 | 876.00 | 904.33 | 1.33 | 0.00 |
| 10.00 | 29.00 | 16.67 | -1.07 | 0.87 |
| 626.00 | 577.00 | 596.00 | 1.06 | 0.34 |
| 6.00 | 18.00 | 11.33 | -1.87 | 0.01 |
| 1192.00 | 1346.00 | 1271.33 | -1.02 | 0.62 |
| 3896.00 | 3928.00 | 4020.00 | -1.01 | 0.80 |
| 120.00 | 129.00 | 125.00 | 1.11 | 0.23 |
| 5625.00 | 5450.00 | 5585.33 | -1.13 | 0.00 |
| 151.00 | 117.00 | 130.67 | -1.10 | 0.33 |
| 205.00 | 172.00 | 177.67 | -1.02 | 0.89 |
| 556.00 | 644.00 | 635.00 | -1.02 | 0.71 |
| 7.00 | 5.00 | 6.00 | -1.73 | 0.09 |
| 315.00 | 282.00 | 301.33 | -1.04 | 0.52 |
| 325.00 | 368.00 | 368.67 | 1.26 | 0.00 |
| 297.00 | 324.00 | 298.00 | 1.26 | 0.00 |
| 484.00 | 602.00 | 546.00 | -1.01 | 0.86 |
| 565.00 | 673.00 | 644.00 | 1.05 | 0.34 |
| 430.00 | 488.00 | 459.00 | 1.18 | 0.00 |
| 4.00 | 5.00 | 6.00 | -1.62 | 0.17 |
| 368.00 | 395.00 | 369.33 | -1.23 | 0.00 |
| 45.00 | 22.00 | 31.33 | 1.41 | 0.08 |
| 16.00 | 22.00 | 18.33 | -1.36 | 0.15 |
| 148.00 | 144.00 | 157.00 | 1.02 | 0.80 |
| 114.00 | 137.00 | 121.67 | -1.08 | 0.40 |
| 12.00 | 24.00 | 17.00 | 1.09 | 0.71 |
| 626.00 | 844.00 | 748.67 | 1.23 | 0.00 |
| 9.00 | 6.00 | 7.33 | -1.01 | 1.00 |
| 11.00 | 2.00 | 4.67 | -1.28 | 0.66 |
| 1835.00 | 2119.00 | 2036.33 | -1.03 | 0.56 |
| 534.00 | 566.00 | 562.67 | -1.06 | 0.25 |
| 1265.00 | 1384.00 | 1351.67 | -1.07 | 0.12 |
| 465.00 | 475.00 | 517.33 | 1.09 | 0.22 |
| 1664.00 | 1743.00 | 1803.67 | 1.09 | 0.04 |
| 848.00 | 800.00 | 781.33 | -1.27 | 0.00 |
| 1753.00 | 1882.00 | 1797.33 | -1.05 | 0.29 |
| 4.00 | 2.00 | 3.00 | -1.33 | 0.67 |
| 18.00 | 19.00 | 21.67 | 1.28 | 0.24 |
| 223.00 | 218.00 | 224.67 | 1.04 | 0.54 |
| 388.00 | 493.00 | 462.67 | -1.27 | 0.00 |
| 205.00 | 238.00 | 236.00 | -1.06 | 0.39 |
| 704.00 | 703.00 | 754.00 | 1.15 | 0.01 |
| 500.00 | 499.00 | 532.33 | 1.09 | 0.08 |
| 531.00 | 581.00 | 596.00 | 1.14 | 0.02 |

| | | | | |
|---|---|---|---|---|
| 2750.00 | 3094.00 | 2885.00 | 1.04 | 0.36 |
| 4023.00 | 4237.00 | 4123.33 | -1.12 | 0.00 |
| 1567.00 | 1524.00 | 1554.33 | 1.04 | 0.41 |
| 1630.00 | 1727.00 | 1666.00 | -1.54 | 0.00 |
| 537.00 | 531.00 | 553.00 | 1.13 | 0.02 |
| 3532.00 | 3678.00 | 3671.00 | 1.18 | 0.00 |
| 2.00 | 1.00 | 2.00 | 1.44 | 0.76 |
| 14.00 | 14.00 | 15.00 | -2.35 | 0.00 |
| 0.00 | 1.00 | 0.33 | -8.37 | 0.01 |
| 1642.00 | 1678.00 | 1700.67 | -1.03 | 0.45 |
| 277.00 | 319.00 | 289.67 | -1.26 | 0.00 |
| 538.00 | 559.00 | 570.67 | -1.20 | 0.00 |
| 827.00 | 953.00 | 944.67 | 1.18 | 0.00 |
| 484.00 | 466.00 | 476.00 | 1.13 | 0.03 |
| 24.00 | 43.00 | 32.00 | 1.10 | 0.65 |
| 739.00 | 728.00 | 759.67 | -1.04 | 0.42 |
| 1818.00 | 1940.00 | 1923.00 | -1.17 | 0.00 |
| 1817.00 | 1806.00 | 1843.67 | -1.15 | 0.00 |
| 1379.00 | 1434.00 | 1452.00 | -1.20 | 0.00 |
| 4.00 | 8.00 | 5.67 | -1.66 | 0.15 |
| 397.00 | 469.00 | 464.67 | 1.05 | 0.49 |
| 55.00 | 84.00 | 83.00 | -1.14 | 0.30 |
| 5.00 | 7.00 | 6.00 | -1.07 | 1.00 |
| 350.00 | 328.00 | 357.00 | 1.05 | 0.41 |
| 22.00 | 20.00 | 22.00 | 1.38 | 0.10 |
| 14.00 | 10.00 | 9.33 | 1.61 | 0.25 |
| 263.00 | 303.00 | 284.00 | -1.15 | 0.02 |
| 44.00 | 57.00 | 48.00 | 1.01 | 0.96 |
| 159.00 | 166.00 | 163.67 | 1.02 | 0.86 |
| 80.00 | 95.00 | 84.67 | -1.08 | 0.46 |
| 162.00 | 155.00 | 177.67 | 1.01 | 0.89 |
| 306.00 | 342.00 | 331.00 | 1.15 | 0.03 |
| 113.00 | 118.00 | 119.00 | 1.36 | 0.00 |
| 68.00 | 95.00 | 87.00 | 1.58 | 0.00 |
| 15.00 | 12.00 | 14.67 | 1.01 | 1.00 |
| 44.00 | 45.00 | 46.33 | 1.32 | 0.04 |
| 905.00 | 1013.00 | 1007.67 | 1.08 | 0.09 |
| 673.00 | 772.00 | 726.67 | 1.08 | 0.13 |
| 13.00 | 19.00 | 18.00 | 1.01 | 1.00 |
| 20.00 | 8.00 | 15.33 | 1.37 | 0.25 |
| 777.00 | 713.00 | 775.33 | -1.08 | 0.14 |
| 1541.00 | 1761.00 | 1638.67 | 1.29 | 0.00 |
| 480.00 | 397.00 | 445.00 | -1.14 | 0.04 |
| 330.00 | 384.00 | 376.67 | 1.11 | 0.11 |
| 62.00 | 63.00 | 63.67 | 1.05 | 0.71 |
| 194.00 | 230.00 | 206.33 | -1.15 | 0.08 |
| 24.00 | 17.00 | 23.00 | -1.41 | 0.04 |
| 4.00 | 6.00 | 5.00 | -1.08 | 1.00 |

| | | | | |
|---|---|---|---|---|
| 2.00 | 2.00 | 1.67 | -2.14 | 0.23 |
| 18.00 | 11.00 | 11.67 | -1.33 | 0.32 |
| 1500.00 | 1690.00 | 1618.33 | 1.23 | 0.00 |
| 1985.00 | 2123.00 | 2168.33 | -1.05 | 0.23 |
| 233.00 | 245.00 | 248.33 | 1.43 | 0.00 |
| 192.00 | 162.00 | 182.33 | 1.30 | 0.00 |
| 527.00 | 516.00 | 558.33 | -1.06 | 0.34 |
| 1068.00 | 1127.00 | 1115.00 | -1.14 | 0.00 |
| 1067.00 | 1011.00 | 1022.00 | 1.10 | 0.09 |
| 267.00 | 263.00 | 293.00 | 1.25 | 0.00 |
| 5.00 | 2.00 | 5.67 | 1.28 | 0.60 |
| 241.00 | 222.00 | 229.67 | -1.04 | 0.63 |
| 145.00 | 116.00 | 123.00 | -1.15 | 0.24 |
| 273.00 | 269.00 | 278.00 | 1.29 | 0.00 |
| 2092.00 | 1955.00 | 2096.33 | -1.28 | 0.00 |
| 11.00 | 10.00 | 9.33 | 1.82 | 0.07 |
| 88.00 | 98.00 | 95.00 | -1.01 | 0.97 |
| 31.00 | 38.00 | 34.00 | -1.12 | 0.44 |
| 1.00 | 0.00 | 0.67 | -1.86 | 0.69 |
| 232.00 | 196.00 | 214.00 | -1.19 | 0.02 |
| 0.00 | 0.00 | 0.00 | -22.48 | 0.01 |
| 208.00 | 273.00 | 261.67 | -1.20 | 0.01 |
| 133.00 | 141.00 | 148.33 | 1.20 | 0.04 |
| 226.00 | 248.00 | 257.67 | 1.32 | 0.00 |
| 41.00 | 67.00 | 51.67 | 1.64 | 0.00 |
| 68.00 | 58.00 | 62.67 | 1.28 | 0.06 |
| 767.00 | 744.00 | 729.00 | 1.07 | 0.31 |
| 1123.00 | 1223.00 | 1249.33 | -1.19 | 0.00 |
| 115.00 | 157.00 | 144.00 | -1.10 | 0.28 |
| 565.00 | 837.00 | 721.33 | 1.01 | 0.89 |
| 80.00 | 89.00 | 84.00 | 1.10 | 0.44 |
| 7.00 | 3.00 | 4.67 | 1.25 | 0.69 |
| 202.00 | 207.00 | 206.33 | 1.36 | 0.00 |
| 0.00 | 4.00 | 1.67 | -1.96 | 0.31 |
| 14.00 | 5.00 | 10.67 | 1.02 | 1.00 |
| 680.00 | 745.00 | 732.00 | 1.26 | 0.00 |
| 1.00 | 2.00 | 2.67 | -1.01 | 1.00 |
| 601.00 | 567.00 | 580.33 | -1.20 | 0.00 |
| 108.00 | 124.00 | 113.00 | 2.40 | 0.00 |
| 274.00 | 265.00 | 276.67 | -1.21 | 0.00 |
| 2.00 | 5.00 | 2.67 | -1.74 | 0.29 |
| 1.00 | 0.00 | 1.33 | -2.87 | 0.08 |
| 309.00 | 339.00 | 333.33 | -1.33 | 0.00 |
| 495.00 | 497.00 | 515.33 | 1.00 | 0.95 |
| 88.00 | 99.00 | 99.33 | 1.20 | 0.07 |
| 312.00 | 261.00 | 301.33 | -1.38 | 0.00 |
| 198.00 | 258.00 | 231.33 | 1.02 | 0.77 |
| 41.00 | 46.00 | 44.33 | -1.19 | 0.19 |

| | | | | |
|---|---|---|---|---|
| 147.00 | 147.00 | 147.67 | -1.20 | 0.04 |
| 84.00 | 77.00 | 85.67 | -1.03 | 0.75 |
| 239.00 | 243.00 | 254.33 | 1.04 | 0.52 |
| 664.00 | 624.00 | 680.00 | -1.01 | 0.81 |
| 705.00 | 712.00 | 698.00 | 1.19 | 0.00 |
| 1636.00 | 1955.00 | 1785.00 | 1.05 | 0.36 |
| 671.00 | 799.00 | 764.33 | 1.07 | 0.14 |
| 12.00 | 12.00 | 10.67 | 1.05 | 0.90 |
| 474.00 | 521.00 | 509.00 | -1.05 | 0.39 |
| 778.00 | 993.00 | 940.67 | 1.16 | 0.01 |
| 763.00 | 779.00 | 830.33 | 1.18 | 0.00 |
| 1580.00 | 1791.00 | 1702.67 | 1.09 | 0.03 |
| 1014.00 | 1118.00 | 1084.67 | 1.25 | 0.00 |
| 192.00 | 252.00 | 241.33 | 1.03 | 0.70 |
| 321.00 | 408.00 | 394.67 | -1.09 | 0.16 |
| 1529.00 | 1665.00 | 1632.67 | 1.05 | 0.31 |
| 64.00 | 79.00 | 71.67 | -1.06 | 0.62 |
| 390.00 | 374.00 | 389.33 | 1.13 | 0.04 |
| 945.00 | 1004.00 | 989.67 | 1.13 | 0.01 |
| 326.00 | 355.00 | 349.33 | 1.14 | 0.03 |
| 456.00 | 450.00 | 491.00 | 1.12 | 0.05 |
| 1118.00 | 1186.00 | 1196.67 | 1.21 | 0.00 |
| 1575.00 | 1763.00 | 1686.33 | 1.37 | 0.00 |
| 618.00 | 835.00 | 760.00 | 1.19 | 0.00 |
| 273.00 | 348.00 | 309.00 | 1.03 | 0.67 |
| 7.00 | 12.00 | 8.00 | -1.64 | 0.09 |
| 166.00 | 205.00 | 187.33 | -1.06 | 0.49 |
| 1898.00 | 2357.00 | 2079.00 | 1.17 | 0.00 |
| 1122.00 | 1105.00 | 1129.33 | -1.16 | 0.00 |
| 0.00 | 1.00 | 0.67 | -1.86 | 0.69 |
| 732.00 | 667.00 | 729.33 | 1.03 | 0.59 |
| 148.00 | 157.00 | 154.67 | -1.30 | 0.00 |
| 236.00 | 244.00 | 256.00 | 1.28 | 0.00 |
| 1740.00 | 1832.00 | 1862.67 | 1.05 | 0.24 |
| 2512.00 | 2806.00 | 2740.00 | 1.03 | 0.42 |
| 97.00 | 150.00 | 129.33 | 1.05 | 0.66 |
| 18.00 | 17.00 | 18.33 | -1.36 | 0.11 |
| 10027.00 | 9426.00 | 9607.67 | -1.10 | 0.02 |
| 2292.00 | 2333.00 | 2472.67 | -1.00 | 0.95 |
| 1045.00 | 1133.00 | 1142.67 | -1.09 | 0.05 |
| 1909.00 | 2180.00 | 2035.00 | -1.04 | 0.37 |
| 428.00 | 462.00 | 456.33 | 1.01 | 0.87 |
| 551.00 | 564.00 | 577.00 | 1.02 | 0.77 |
| 1804.00 | 1914.00 | 1820.67 | -1.21 | 0.00 |
| 1.00 | 6.00 | 2.67 | 1.12 | 1.00 |
| 1280.00 | 1425.00 | 1307.33 | 1.94 | 0.00 |
| 8774.00 | 9242.00 | 9596.67 | 1.09 | 0.02 |
| 52.00 | 57.00 | 57.67 | 1.07 | 0.61 |

| | | | | |
|---|---|---|---|---|
| 11.00 | 16.00 | 13.33 | -1.39 | 0.17 |
| 691.00 | 680.00 | 704.67 | 1.01 | 0.86 |
| 2672.00 | 3084.00 | 3134.00 | 1.27 | 0.00 |
| 987.00 | 1022.00 | 1046.33 | -1.52 | 0.00 |
| 29.00 | 22.00 | 28.00 | -1.11 | 0.57 |
| 17.00 | 25.00 | 20.67 | -1.68 | 0.00 |
| 1669.00 | 1736.00 | 1671.33 | -1.81 | 0.00 |
| 5371.00 | 5639.00 | 5363.67 | -1.95 | 0.00 |
| 1928.00 | 1657.00 | 1839.33 | -1.28 | 0.00 |
| 1158.00 | 1147.00 | 1188.00 | 1.05 | 0.23 |
| 429.00 | 452.00 | 463.00 | 1.01 | 0.88 |
| 12.00 | 15.00 | 15.33 | -1.23 | 0.36 |
| 422.00 | 402.00 | 405.67 | 1.08 | 0.21 |
| 336.00 | 324.00 | 362.00 | -1.15 | 0.02 |
| 543.00 | 556.00 | 545.67 | -1.03 | 0.61 |
| 749.00 | 766.00 | 750.00 | -1.10 | 0.08 |
| 1053.00 | 1051.00 | 1102.67 | -1.20 | 0.00 |
| 519.00 | 503.00 | 510.33 | -1.50 | 0.00 |
| 369.00 | 430.00 | 410.00 | 1.06 | 0.34 |
| 424.00 | 423.00 | 437.33 | -1.15 | 0.01 |
| 10.00 | 5.00 | 8.00 | -1.80 | 0.03 |
| 7.00 | 4.00 | 6.67 | 1.16 | 0.75 |
| 647.00 | 782.00 | 718.00 | 1.03 | 0.60 |
| 735.00 | 749.00 | 761.33 | 1.15 | 0.00 |
| 447.00 | 544.00 | 511.00 | -1.20 | 0.00 |
| 1041.00 | 1039.00 | 1089.00 | -1.02 | 0.61 |
| 216.00 | 235.00 | 232.00 | 1.09 | 0.21 |
| 242.00 | 216.00 | 244.33 | -1.39 | 0.00 |
| 515.00 | 454.00 | 486.33 | -1.07 | 0.28 |
| 1534.00 | 1730.00 | 1683.67 | 1.10 | 0.02 |
| 23.00 | 21.00 | 21.67 | 1.14 | 0.48 |
| 36.00 | 32.00 | 37.33 | -1.10 | 0.50 |
| 831.00 | 818.00 | 871.00 | -1.00 | 0.96 |
| 2477.00 | 2647.00 | 2652.67 | -1.08 | 0.02 |
| 549.00 | 534.00 | 518.33 | -1.03 | 0.68 |
| 720.00 | 838.00 | 808.33 | 1.11 | 0.03 |
| 17.00 | 10.00 | 16.00 | 1.28 | 0.30 |
| 115.00 | 98.00 | 114.00 | 1.20 | 0.12 |
| 1021.00 | 1057.00 | 1046.00 | 1.05 | 0.25 |
| 346.00 | 350.00 | 348.33 | 1.02 | 0.70 |
| 912.00 | 980.00 | 951.33 | -1.05 | 0.32 |
| 1397.00 | 1514.00 | 1524.67 | -1.06 | 0.13 |
| 655.00 | 665.00 | 681.33 | 1.21 | 0.00 |
| 279.00 | 298.00 | 289.33 | 1.13 | 0.06 |
| 601.00 | 576.00 | 621.00 | 1.04 | 0.46 |
| 79.00 | 81.00 | 79.67 | -1.13 | 0.26 |
| 964.00 | 1140.00 | 1056.33 | -1.04 | 0.44 |
| 429.00 | 513.00 | 474.67 | 1.41 | 0.00 |

| | | | | |
|---|---|---|---|---|
| 832.00 | 863.00 | 870.67 | -1.11 | 0.02 |
| 629.00 | 629.00 | 627.67 | 1.08 | 0.11 |
| 228.00 | 276.00 | 252.67 | 1.27 | 0.00 |
| 374.00 | 442.00 | 412.00 | 1.05 | 0.47 |
| 664.00 | 702.00 | 682.67 | 1.12 | 0.02 |
| 487.00 | 462.00 | 517.00 | -1.01 | 0.89 |
| 48.00 | 71.00 | 54.67 | -1.23 | 0.10 |
| 426.00 | 551.00 | 495.67 | 1.31 | 0.00 |
| 18.00 | 21.00 | 23.67 | 1.93 | 0.00 |
| 1217.00 | 1200.00 | 1237.33 | 1.01 | 0.73 |
| 150.00 | 196.00 | 183.00 | -1.01 | 0.92 |
| 1195.00 | 1342.00 | 1309.33 | -1.06 | 0.15 |
| 1379.00 | 1495.00 | 1450.00 | -1.19 | 0.00 |
| 1633.00 | 1771.00 | 1716.67 | -1.10 | 0.02 |
| 664.00 | 739.00 | 754.00 | -1.03 | 0.49 |
| 3.00 | 3.00 | 2.67 | -2.22 | 0.08 |
| 1787.00 | 1750.00 | 1908.33 | -1.01 | 0.90 |
| 333.00 | 347.00 | 338.00 | 1.09 | 0.14 |
| 656.00 | 642.00 | 642.67 | -1.20 | 0.00 |
| 221.00 | 217.00 | 236.67 | -1.16 | 0.03 |
| 15.00 | 14.00 | 15.67 | -1.40 | 0.12 |
| 932.00 | 974.00 | 984.00 | 1.12 | 0.01 |
| 5.00 | 12.00 | 9.67 | -1.22 | 0.55 |
| 450.00 | 392.00 | 463.33 | -1.14 | 0.06 |
| 12.00 | 18.00 | 11.67 | -1.35 | 0.28 |
| 691.00 | 836.00 | 755.67 | 1.10 | 0.07 |
| 886.00 | 949.00 | 923.00 | -1.24 | 0.00 |
| 663.00 | 743.00 | 687.33 | 1.01 | 0.91 |
| 7.00 | 5.00 | 6.67 | -2.85 | 0.00 |
| 2.00 | 2.00 | 2.33 | -5.00 | 0.00 |
| 108.00 | 78.00 | 96.33 | 1.01 | 0.97 |
| 983.00 | 1078.00 | 1077.33 | -1.08 | 0.05 |
| 2485.00 | 2730.00 | 2719.67 | 1.14 | 0.00 |
| 541.00 | 714.00 | 636.33 | 1.11 | 0.08 |
| 359.00 | 378.00 | 379.67 | -1.00 | 0.97 |
| 201.00 | 247.00 | 237.33 | 1.23 | 0.00 |
| 156.00 | 170.00 | 163.67 | -1.19 | 0.01 |
| 266.00 | 254.00 | 279.00 | -1.06 | 0.36 |
| 730.00 | 830.00 | 802.67 | 1.08 | 0.09 |
| 1105.00 | 1212.00 | 1154.00 | 1.02 | 0.64 |
| 643.00 | 687.00 | 633.67 | -1.04 | 0.50 |
| 958.00 | 1057.00 | 1032.67 | 1.26 | 0.00 |
| 10.00 | 6.00 | 7.67 | 2.03 | 0.06 |
| 634.00 | 630.00 | 645.33 | -1.07 | 0.18 |
| 44.00 | 26.00 | 37.33 | -2.40 | 0.00 |
| 5.00 | 10.00 | 5.33 | -3.30 | 0.00 |
| 1343.00 | 1289.00 | 1325.00 | 1.12 | 0.01 |
| 1472.00 | 1728.00 | 1601.67 | -1.11 | 0.02 |

| | | | | |
|---|---|---|---|---|
| 319.00 | 368.00 | 353.33 | -1.62 | 0.00 |
| 5.00 | 3.00 | 4.00 | -1.59 | 0.29 |
| 6.00 | 7.00 | 8.00 | 1.07 | 0.89 |
| 3274.00 | 3744.00 | 3651.67 | 1.49 | 0.00 |
| 610.00 | 652.00 | 658.33 | -1.04 | 0.39 |
| 8.00 | 6.00 | 9.00 | -1.31 | 0.39 |
| 2112.00 | 2284.00 | 2267.33 | 1.09 | 0.02 |
| 425.00 | 485.00 | 461.33 | 1.34 | 0.00 |
| 176.00 | 224.00 | 204.33 | 2.24 | 0.00 |
| 10.00 | 7.00 | 9.00 | -2.20 | 0.00 |
| 139.00 | 113.00 | 117.00 | 1.67 | 0.00 |
| 100.00 | 112.00 | 113.67 | 1.15 | 0.15 |
| 4.00 | 9.00 | 5.67 | -1.89 | 0.05 |
| 698.00 | 818.00 | 725.00 | -1.16 | 0.03 |
| 5.00 | 1.00 | 3.33 | 1.22 | 0.82 |
| 36.00 | 29.00 | 33.67 | -1.27 | 0.09 |
| 2827.00 | 3177.00 | 2952.67 | 1.22 | 0.00 |
| 116.00 | 114.00 | 123.33 | 1.03 | 0.80 |
| 3211.00 | 3268.00 | 3267.33 | -1.53 | 0.00 |
| 808.00 | 1080.00 | 982.67 | 1.04 | 0.49 |
| 1042.00 | 1193.00 | 1133.33 | 1.18 | 0.00 |
| 18.00 | 11.00 | 13.33 | 1.46 | 0.16 |
| 446.00 | 427.00 | 446.00 | -1.20 | 0.00 |
| 350.00 | 378.00 | 369.33 | 1.25 | 0.00 |
| 1537.00 | 1672.00 | 1702.67 | 1.15 | 0.00 |
| 35.00 | 22.00 | 29.67 | -1.10 | 0.59 |
| 37.00 | 45.00 | 41.67 | -1.31 | 0.04 |
| 62.00 | 68.00 | 70.33 | -1.31 | 0.01 |
| 910.00 | 938.00 | 967.00 | -1.24 | 0.00 |
| 2976.00 | 3086.00 | 3176.33 | -1.18 | 0.00 |
| 244.00 | 236.00 | 253.33 | -1.02 | 0.72 |
| 1565.00 | 1545.00 | 1600.67 | 1.02 | 0.58 |
| 730.00 | 742.00 | 768.00 | 1.20 | 0.00 |
| 1.00 | 1.00 | 1.67 | -4.59 | 0.00 |
| 1909.00 | 2017.00 | 2072.67 | 1.07 | 0.07 |
| 104.00 | 115.00 | 110.67 | -1.20 | 0.03 |
| 555.00 | 600.00 | 600.00 | -1.18 | 0.00 |
| 364.00 | 412.00 | 392.33 | -1.26 | 0.00 |
| 912.00 | 1084.00 | 1007.33 | 1.00 | 1.00 |
| 7.00 | 10.00 | 7.00 | -1.58 | 0.14 |
| 1109.00 | 1236.00 | 1213.67 | 1.10 | 0.03 |
| 173.00 | 188.00 | 177.00 | -1.39 | 0.00 |
| 647.00 | 717.00 | 700.00 | 1.02 | 0.67 |
| 4.00 | 1.00 | 2.67 | -1.50 | 0.51 |
| 83.00 | 74.00 | 77.00 | -1.17 | 0.14 |
| 295.00 | 266.00 | 286.67 | 1.22 | 0.00 |
| 4366.00 | 4303.00 | 4347.33 | 1.14 | 0.00 |
| 12009.00 | 12549.00 | 12500.00 | -1.20 | 0.00 |

| | | | | |
|---|---|---|---|---|
| 2084.00 | 2192.00 | 2203.33 | 1.06 | 0.18 |
| 1212.00 | 1123.00 | 1156.00 | -1.13 | 0.01 |
| 1587.00 | 1695.00 | 1647.00 | -1.06 | 0.15 |
| 6398.00 | 7089.00 | 7122.67 | -1.04 | 0.38 |
| 41.00 | 45.00 | 49.67 | 1.25 | 0.11 |
| 8.00 | 7.00 | 10.33 | 1.09 | 0.81 |
| 1656.00 | 1679.00 | 1725.00 | 1.09 | 0.08 |
| 1391.00 | 1595.00 | 1606.00 | 1.09 | 0.07 |
| 3799.00 | 3870.00 | 3992.67 | 1.12 | 0.00 |
| 579.00 | 641.00 | 649.67 | 1.51 | 0.00 |
| 441.00 | 543.00 | 516.67 | -1.01 | 0.86 |
| 3875.00 | 4014.00 | 4003.67 | 1.05 | 0.20 |
| 1838.00 | 1836.00 | 1808.33 | -1.09 | 0.07 |
| 51.00 | 51.00 | 45.00 | 1.17 | 0.30 |
| 664.00 | 700.00 | 708.67 | -1.29 | 0.00 |
| 1669.00 | 1688.00 | 1662.00 | -1.06 | 0.20 |
| 1036.00 | 1146.00 | 1153.67 | -1.05 | 0.29 |
| 1072.00 | 1015.00 | 1082.67 | -1.16 | 0.00 |
| 282.00 | 342.00 | 331.33 | -1.10 | 0.16 |
| 876.00 | 939.00 | 915.67 | 1.03 | 0.49 |
| 3584.00 | 3438.00 | 3558.67 | -1.18 | 0.00 |
| 269.00 | 333.00 | 308.00 | 1.02 | 0.79 |
| 22.00 | 27.00 | 27.33 | -1.31 | 0.09 |
| 23.00 | 21.00 | 20.33 | -1.39 | 0.07 |
| 38.00 | 48.00 | 44.00 | -2.29 | 0.00 |
| 679.00 | 714.00 | 700.67 | -1.58 | 0.00 |
| 31.00 | 22.00 | 23.33 | -1.14 | 0.55 |
| 2.00 | 4.00 | 2.33 | -1.15 | 1.00 |
| 28.00 | 28.00 | 29.67 | 1.02 | 0.94 |
| 4.00 | 4.00 | 7.00 | -1.92 | 0.03 |
| 34.00 | 42.00 | 35.67 | 1.08 | 0.65 |
| 15.00 | 31.00 | 26.67 | -1.23 | 0.24 |
| 51.00 | 69.00 | 66.00 | -1.05 | 0.69 |
| 58.00 | 56.00 | 60.67 | -1.44 | 0.00 |
| 54.00 | 77.00 | 70.33 | -1.40 | 0.00 |
| 102.00 | 108.00 | 104.33 | 1.10 | 0.31 |
| 100.00 | 91.00 | 94.67 | -1.32 | 0.00 |
| 20.00 | 27.00 | 27.33 | 1.04 | 0.89 |
| 307.00 | 355.00 | 346.00 | 1.12 | 0.06 |
| 34287.00 | 30735.00 | 31672.33 | -1.31 | 0.00 |
| 15634.00 | 14628.00 | 14998.33 | -1.18 | 0.00 |
| 6674.00 | 6102.00 | 6186.33 | -2.59 | 0.00 |
| 9.00 | 18.00 | 13.67 | 1.22 | 0.44 |
| 906.00 | 817.00 | 876.67 | 1.02 | 0.71 |
| 651.00 | 757.00 | 712.00 | -1.01 | 0.87 |
| 8.00 | 12.00 | 8.67 | -2.05 | 0.00 |
| 52.00 | 47.00 | 56.67 | -1.15 | 0.24 |
| 901.00 | 1044.00 | 965.33 | 1.00 | 0.93 |

| | | | | |
|---|---|---|---|---|
| 21.00 | 20.00 | 20.67 | 1.20 | 0.36 |
| 8.00 | 4.00 | 8.00 | -1.68 | 0.08 |
| 2447.00 | 2442.00 | 2523.67 | -1.06 | 0.17 |
| 371.00 | 370.00 | 395.67 | 1.01 | 0.86 |
| 434.00 | 413.00 | 390.33 | -1.21 | 0.02 |
| 285.00 | 266.00 | 276.33 | 1.39 | 0.00 |
| 898.00 | 930.00 | 945.33 | -1.84 | 0.00 |
| 3128.00 | 3527.00 | 3292.33 | 1.14 | 0.00 |
| 1898.00 | 1780.00 | 1852.33 | -1.31 | 0.00 |
| 49.00 | 61.00 | 64.67 | 1.01 | 0.97 |
| 713.00 | 745.00 | 730.67 | 1.03 | 0.47 |
| 82899.00 | 81936.00 | 80570.33 | 1.24 | 0.00 |
| 63.00 | 65.00 | 65.67 | 1.01 | 0.96 |
| 595.00 | 659.00 | 631.33 | 1.08 | 0.13 |
| 932.00 | 966.00 | 986.33 | -1.09 | 0.06 |
| 783.00 | 905.00 | 839.67 | 1.12 | 0.02 |
| 2967.00 | 3405.00 | 3200.33 | -1.13 | 0.00 |
| 1657.00 | 1800.00 | 1713.33 | 1.08 | 0.08 |
| 1429.00 | 1514.00 | 1417.67 | -1.08 | 0.17 |
| 64.00 | 84.00 | 73.33 | -1.29 | 0.02 |
| 2.00 | 3.00 | 2.67 | 1.12 | 1.00 |
| 1026.00 | 1060.00 | 1043.00 | -1.10 | 0.05 |
| 167.00 | 163.00 | 159.67 | -1.07 | 0.39 |
| 1250.00 | 1337.00 | 1368.67 | 1.02 | 0.57 |
| 22.00 | 22.00 | 25.33 | 1.10 | 0.69 |
| 1624.00 | 1669.00 | 1672.00 | -1.08 | 0.05 |
| 3.00 | 2.00 | 2.67 | -1.38 | 0.65 |
| 63.00 | 66.00 | 65.67 | 1.32 | 0.02 |
| 29.00 | 34.00 | 31.67 | -1.19 | 0.25 |
| 95.00 | 81.00 | 94.33 | -1.25 | 0.02 |
| 12410.00 | 12555.00 | 13058.00 | -1.10 | 0.00 |
| 3102.00 | 2815.00 | 2814.33 | -1.09 | 0.18 |
| 23.00 | 20.00 | 23.67 | -1.10 | 0.58 |
| 644.00 | 631.00 | 670.00 | -1.42 | 0.00 |
| 328.00 | 359.00 | 339.00 | 1.03 | 0.65 |
| 122.00 | 124.00 | 136.67 | 1.17 | 0.08 |
| 22.00 | 19.00 | 26.67 | 1.23 | 0.28 |
| 4304.00 | 4480.00 | 4585.67 | 1.54 | 0.00 |
| 12.00 | 17.00 | 16.00 | 1.43 | 0.13 |
| 67.00 | 102.00 | 83.67 | -1.05 | 0.69 |
| 32.00 | 19.00 | 27.00 | -1.35 | 0.07 |
| 65.00 | 83.00 | 70.67 | 1.14 | 0.30 |
| 18.00 | 16.00 | 16.00 | -1.47 | 0.06 |
| 5.00 | 8.00 | 8.00 | -1.26 | 0.51 |
| 529.00 | 508.00 | 525.67 | 2.13 | 0.00 |
| 695.00 | 691.00 | 701.33 | 1.24 | 0.00 |
| 105.00 | 136.00 | 123.33 | 1.09 | 0.34 |
| 12.00 | 23.00 | 17.67 | -1.20 | 0.37 |

| | | | | |
|---|---|---|---|---|
| 700.00 | 733.00 | 698.67 | -1.22 | 0.00 |
| 2310.00 | 2658.00 | 2488.33 | -1.08 | 0.05 |
| 2279.00 | 2379.00 | 2314.33 | -1.25 | 0.00 |
| 2542.00 | 2506.00 | 2637.67 | -1.00 | 0.97 |
| 1360.00 | 1333.00 | 1383.67 | 1.02 | 0.71 |
| 2394.00 | 2483.00 | 2512.67 | 1.12 | 0.01 |
| 655.00 | 787.00 | 732.67 | 1.21 | 0.00 |
| 548.00 | 535.00 | 558.33 | -1.26 | 0.00 |
| 30.00 | 32.00 | 35.00 | 1.28 | 0.15 |
| 141.00 | 174.00 | 166.33 | -1.12 | 0.14 |
| 644.00 | 669.00 | 652.67 | -1.18 | 0.00 |
| 286.00 | 308.00 | 302.33 | -1.01 | 0.83 |
| 198.00 | 176.00 | 193.00 | -1.35 | 0.00 |
| 6.00 | 6.00 | 4.00 | -1.50 | 0.40 |
| 1589.00 | 1552.00 | 1624.00 | -1.03 | 0.46 |
| 118.00 | 124.00 | 119.67 | -1.27 | 0.00 |
| 1.00 | 2.00 | 1.00 | -1.31 | 1.00 |
| 797.00 | 815.00 | 844.33 | -1.11 | 0.03 |
| 36.00 | 33.00 | 35.00 | 1.57 | 0.01 |
| 4.00 | 3.00 | 3.67 | 1.52 | 0.49 |
| 630.00 | 630.00 | 669.33 | -1.14 | 0.01 |
| 937.00 | 954.00 | 957.00 | -1.08 | 0.08 |
| 1129.00 | 1132.00 | 1145.00 | -1.05 | 0.29 |
| 251.00 | 329.00 | 287.00 | 1.17 | 0.03 |
| 1023.00 | 1297.00 | 1161.33 | 1.14 | 0.01 |
| 505.00 | 512.00 | 487.33 | -1.80 | 0.00 |
| 84.00 | 93.00 | 97.33 | 1.09 | 0.41 |
| 300.00 | 295.00 | 299.33 | 1.24 | 0.00 |
| 2.00 | 2.00 | 2.33 | 1.35 | 0.78 |
| 49.00 | 45.00 | 45.67 | -1.09 | 0.54 |
| 0.00 | 1.00 | 0.67 | -4.85 | 0.02 |
| 229.00 | 228.00 | 235.67 | -1.04 | 0.58 |
| 342.00 | 424.00 | 361.00 | -1.08 | 0.29 |
| 41.00 | 60.00 | 55.67 | -1.18 | 0.17 |
| 381.00 | 393.00 | 379.33 | 1.07 | 0.23 |
| 81.00 | 79.00 | 80.67 | 1.03 | 0.77 |
| 8.00 | 9.00 | 9.33 | 1.45 | 0.25 |
| 17.00 | 22.00 | 22.33 | -1.21 | 0.31 |
| 2477.00 | 2596.00 | 2619.67 | -1.43 | 0.00 |
| 205.00 | 245.00 | 235.33 | 1.18 | 0.04 |
| 827.00 | 832.00 | 841.67 | -1.23 | 0.00 |
| 4501.00 | 4790.00 | 4662.00 | 1.20 | 0.00 |
| 12.00 | 18.00 | 15.33 | -1.04 | 1.00 |
| 1167.00 | 1271.00 | 1238.67 | 1.02 | 0.62 |
| 0.00 | 0.00 | 0.00 | -17.11 | 0.03 |
| 6.00 | 10.00 | 8.33 | -1.01 | 1.00 |
| 137.00 | 152.00 | 146.67 | 1.05 | 0.60 |
| 423.00 | 469.00 | 441.67 | 1.01 | 0.86 |

| | | | | |
|---|---|---|---|---|
| 185.00 | 168.00 | 169.00 | -1.14 | 0.18 |
| 122.00 | 121.00 | 124.67 | 1.04 | 0.69 |
| 1561.00 | 1896.00 | 1675.67 | 1.03 | 0.56 |
| 17.00 | 11.00 | 13.33 | -1.19 | 0.53 |
| 2098.00 | 2012.00 | 2100.67 | 1.32 | 0.00 |
| 796.00 | 791.00 | 830.67 | 1.06 | 0.19 |
| 188.00 | 227.00 | 204.33 | -1.01 | 0.87 |
| 8.00 | 19.00 | 16.00 | -1.08 | 0.78 |
| 35.00 | 30.00 | 32.33 | -2.46 | 0.00 |
| 3461.00 | 3553.00 | 3551.33 | 1.41 | 0.00 |
| 5489.00 | 5629.00 | 5604.00 | -1.05 | 0.22 |
| 296.00 | 266.00 | 298.00 | -1.63 | 0.00 |
| 6.00 | 14.00 | 10.00 | -1.68 | 0.04 |
| 36.00 | 37.00 | 38.67 | 1.24 | 0.18 |
| 5954.00 | 6317.00 | 6511.67 | -1.02 | 0.60 |
| 67.00 | 55.00 | 64.67 | -1.09 | 0.50 |
| 11503.00 | 11676.00 | 12077.00 | -1.30 | 0.00 |
| 699.00 | 666.00 | 702.67 | -1.07 | 0.13 |
| 2.00 | 3.00 | 2.67 | -2.47 | 0.04 |
| 1105.00 | 1216.00 | 1185.67 | 1.00 | 1.00 |
| 756.00 | 792.00 | 814.00 | 1.21 | 0.00 |
| 2137.00 | 2416.00 | 2357.33 | -1.06 | 0.12 |
| 34.00 | 37.00 | 36.67 | -1.12 | 0.42 |
| 3451.00 | 3809.00 | 3629.00 | -1.06 | 0.11 |
| 1797.00 | 1948.00 | 1828.00 | 1.02 | 0.62 |
| 138.00 | 141.00 | 148.00 | 1.13 | 0.15 |
| 4900.00 | 5286.00 | 5224.33 | -1.03 | 0.45 |
| 569.00 | 542.00 | 561.67 | -1.11 | 0.04 |
| 255.00 | 329.00 | 300.00 | 1.25 | 0.00 |
| 347.00 | 403.00 | 399.33 | 1.08 | 0.26 |
| 58.00 | 69.00 | 59.67 | 1.53 | 0.00 |
| 1764.00 | 1676.00 | 1724.33 | -1.31 | 0.00 |
| 10094.00 | 11795.00 | 11342.33 | 1.14 | 0.00 |
| 1.00 | 8.00 | 5.67 | 1.11 | 0.87 |
| 146.00 | 125.00 | 143.67 | 1.02 | 0.87 |
| 236.00 | 169.00 | 188.33 | -1.13 | 0.29 |
| 409.00 | 436.00 | 419.67 | 1.00 | 0.95 |
| 4253.00 | 4289.00 | 4432.67 | 1.10 | 0.01 |
| 180.00 | 186.00 | 193.33 | 1.10 | 0.20 |
| 496.00 | 518.00 | 518.00 | 1.28 | 0.00 |
| 270.00 | 272.00 | 285.67 | 1.17 | 0.01 |
| 3893.00 | 4187.00 | 4201.33 | 1.16 | 0.00 |
| 1236.00 | 1373.00 | 1337.33 | -1.13 | 0.01 |
| 3167.00 | 3279.00 | 3303.00 | 1.18 | 0.00 |
| 2.00 | 2.00 | 1.67 | 1.21 | 1.00 |
| 1292.00 | 1130.00 | 1167.33 | -1.16 | 0.02 |
| 2287.00 | 2503.00 | 2442.33 | 1.08 | 0.07 |
| 996.00 | 1072.00 | 1068.33 | -1.01 | 0.72 |

| | | | | |
|---|---|---|---|---|
| 319.00 | 394.00 | 369.00 | -1.15 | 0.02 |
| 1034.00 | 1115.00 | 1122.00 | -1.06 | 0.17 |
| 370.00 | 397.00 | 364.67 | -1.06 | 0.44 |
| 194.00 | 196.00 | 180.33 | -1.16 | 0.10 |
| 187.00 | 189.00 | 185.33 | -1.02 | 0.86 |
| 2728.00 | 2874.00 | 2811.00 | -1.01 | 0.71 |
| 287.00 | 290.00 | 293.67 | -1.23 | 0.00 |
| 790.00 | 834.00 | 826.00 | -1.10 | 0.02 |
| 7.00 | 6.00 | 5.67 | -1.19 | 0.75 |
| 800.00 | 798.00 | 817.33 | -1.09 | 0.05 |
| 6965.00 | 7338.00 | 7461.00 | 1.08 | 0.07 |
| 805.00 | 978.00 | 863.00 | 1.07 | 0.27 |
| 4546.00 | 5107.00 | 4754.67 | 1.20 | 0.00 |
| 1538.00 | 1653.00 | 1588.33 | -1.22 | 0.00 |
| 2133.00 | 2121.00 | 2222.33 | 1.03 | 0.51 |
| 880.00 | 909.00 | 955.00 | -1.15 | 0.00 |
| 4455.00 | 4158.00 | 4361.67 | -1.09 | 0.05 |
| 174.00 | 195.00 | 182.67 | -1.30 | 0.00 |
| 253.00 | 242.00 | 264.33 | -1.56 | 0.00 |
| 4.00 | 6.00 | 7.67 | 1.13 | 0.77 |
| 2316.00 | 2342.00 | 2557.33 | 1.09 | 0.09 |
| 483.00 | 561.00 | 569.00 | -1.05 | 0.38 |
| 865.00 | 925.00 | 939.33 | 1.12 | 0.01 |
| 20.00 | 25.00 | 23.33 | 1.13 | 0.57 |
| 821.00 | 1113.00 | 925.33 | 1.15 | 0.05 |
| 483.00 | 592.00 | 591.33 | 1.22 | 0.00 |
| 853.00 | 810.00 | 778.00 | -1.57 | 0.00 |
| 126.00 | 153.00 | 137.00 | -1.17 | 0.05 |
| 24.00 | 20.00 | 25.67 | -1.34 | 0.07 |
| 90.00 | 71.00 | 79.67 | 1.11 | 0.34 |
| 10.00 | 17.00 | 14.33 | 2.61 | 0.00 |
| 232.00 | 304.00 | 287.33 | -2.75 | 0.00 |
| 72.00 | 88.00 | 78.67 | 1.82 | 0.00 |
| 31.00 | 35.00 | 33.00 | 1.19 | 0.30 |
| 11.00 | 5.00 | 7.67 | -1.45 | 0.24 |
| 3.00 | 3.00 | 4.00 | -1.42 | 0.47 |
| 93.00 | 90.00 | 96.00 | 1.70 | 0.00 |
| 81.00 | 75.00 | 84.33 | 1.13 | 0.27 |
| 1823.00 | 2059.00 | 2076.00 | 1.11 | 0.01 |
| 977.00 | 971.00 | 1039.00 | 1.00 | 1.00 |
| 989.00 | 1009.00 | 1020.67 | 1.19 | 0.00 |
| 1442.00 | 1378.00 | 1426.00 | -1.22 | 0.00 |
| 39.00 | 19.00 | 33.33 | -1.33 | 0.14 |
| 218.00 | 263.00 | 236.67 | -1.39 | 0.00 |
| 1530.00 | 1614.00 | 1696.33 | -1.10 | 0.04 |
| 395.00 | 383.00 | 412.33 | 1.35 | 0.00 |
| 450.00 | 480.00 | 486.67 | -1.09 | 0.08 |
| 11.00 | 5.00 | 8.33 | -1.33 | 0.37 |

| | | | | |
|---|---|---|---|---|
| 1285.00 | 1247.00 | 1302.00 | 1.01 | 0.80 |
| 1369.00 | 1245.00 | 1299.67 | 1.18 | 0.00 |
| 1962.00 | 1920.00 | 1964.33 | -1.30 | 0.00 |
| 1107.00 | 1254.00 | 1210.67 | 1.12 | 0.01 |
| 141.00 | 147.00 | 156.33 | 1.13 | 0.15 |
| 3.00 | 5.00 | 3.33 | -1.40 | 0.56 |
| 22.00 | 8.00 | 14.00 | -1.75 | 0.01 |
| 184.00 | 235.00 | 238.33 | 1.38 | 0.00 |
| 332.00 | 336.00 | 338.00 | 1.61 | 0.00 |
| 4007.00 | 3842.00 | 3911.67 | -1.28 | 0.00 |
| 11.00 | 5.00 | 7.67 | 1.33 | 0.44 |
| 9.00 | 5.00 | 5.67 | -2.70 | 0.00 |
| 6.00 | 3.00 | 3.33 | -2.09 | 0.08 |
| 3193.00 | 3075.00 | 3140.33 | -1.09 | 0.03 |
| 800.00 | 888.00 | 833.67 | -1.43 | 0.00 |
| 11.00 | 15.00 | 12.33 | 1.22 | 0.48 |
| 4.00 | 5.00 | 5.67 | -1.25 | 0.63 |
| 340.00 | 296.00 | 342.00 | -1.08 | 0.32 |
| 433.00 | 433.00 | 475.00 | 1.14 | 0.05 |
| 52.00 | 74.00 | 65.33 | 1.28 | 0.05 |
| 20471.00 | 21497.00 | 20997.00 | -1.24 | 0.00 |
| 323.00 | 318.00 | 312.33 | -1.03 | 0.77 |
| 5.00 | 6.00 | 6.67 | 1.77 | 0.16 |
| 67710.00 | 70448.00 | 73444.67 | 1.13 | 0.00 |
| 156.00 | 144.00 | 163.00 | 1.92 | 0.00 |
| 1192.00 | 1292.00 | 1246.00 | -1.07 | 0.11 |
| 683.00 | 806.00 | 811.67 | 1.14 | 0.02 |
| 78.00 | 97.00 | 85.67 | -1.08 | 0.46 |
| 27.00 | 26.00 | 25.67 | 1.58 | 0.02 |
| 122.00 | 116.00 | 120.00 | -1.32 | 0.00 |
| 13618.00 | 14592.00 | 14340.00 | -1.03 | 0.29 |
| 3583.00 | 3507.00 | 3532.33 | 1.12 | 0.01 |
| 20.00 | 19.00 | 20.67 | 1.74 | 0.01 |
| 312.00 | 313.00 | 324.00 | -1.01 | 0.92 |
| 603.00 | 696.00 | 644.00 | 1.19 | 0.00 |
| 11277.00 | 12692.00 | 13164.33 | 1.12 | 0.03 |
| 2034.00 | 2132.00 | 2167.33 | -1.12 | 0.00 |
| 13.00 | 4.00 | 7.67 | -2.40 | 0.00 |
| 55.00 | 66.00 | 60.67 | -1.25 | 0.05 |
| 743.00 | 676.00 | 760.00 | -1.01 | 0.86 |
| 179.00 | 217.00 | 213.00 | 1.01 | 0.93 |
| 56.00 | 43.00 | 43.00 | -1.35 | 0.04 |
| 318.00 | 291.00 | 307.00 | -1.51 | 0.00 |
| 32.00 | 28.00 | 32.00 | -1.67 | 0.00 |
| 1836.00 | 2128.00 | 2035.67 | -1.01 | 0.84 |
| 2825.00 | 2804.00 | 2885.67 | 1.05 | 0.15 |
| 23.00 | 2.00 | 10.33 | -1.25 | 0.63 |
| 600.00 | 614.00 | 618.67 | 1.14 | 0.01 |

| | | | | |
|---|---|---|---|---|
| 823.00 | 851.00 | 889.67 | 1.01 | 0.79 |
| 3.00 | 8.00 | 8.00 | -1.06 | 1.00 |
| 76.00 | 65.00 | 72.33 | -1.41 | 0.00 |
| 119.00 | 99.00 | 110.67 | 1.20 | 0.06 |
| 199.00 | 188.00 | 206.33 | -1.13 | 0.08 |
| 14.00 | 9.00 | 13.67 | -1.18 | 0.54 |
| 1910.00 | 1819.00 | 1922.67 | -1.24 | 0.00 |
| 34.00 | 49.00 | 48.33 | -1.33 | 0.05 |
| 1099.00 | 1113.00 | 1121.00 | 1.20 | 0.00 |
| 9.00 | 7.00 | 8.00 | 1.47 | 0.28 |
| 1736.00 | 1865.00 | 1839.33 | 1.10 | 0.02 |
| 3727.00 | 3691.00 | 3695.33 | -1.15 | 0.00 |
| 164.00 | 191.00 | 180.67 | 1.38 | 0.00 |
| 17.00 | 25.00 | 22.33 | 1.73 | 0.01 |
| 117.00 | 103.00 | 119.00 | 1.19 | 0.07 |
| 4.00 | 9.00 | 7.00 | -1.11 | 0.88 |
| 16.00 | 19.00 | 24.33 | 1.28 | 0.24 |
| 221.00 | 239.00 | 241.33 | 1.15 | 0.04 |
| 464.00 | 515.00 | 542.00 | 1.16 | 0.02 |
| 687.00 | 686.00 | 687.67 | -1.05 | 0.33 |
| 1062.00 | 1078.00 | 1055.67 | -1.09 | 0.08 |
| 413.00 | 476.00 | 453.00 | 1.06 | 0.33 |
| 785.00 | 819.00 | 820.67 | 1.23 | 0.00 |
| 59.00 | 77.00 | 68.67 | 1.33 | 0.02 |
| 795.00 | 830.00 | 828.00 | -1.02 | 0.62 |
| 928.00 | 1099.00 | 1012.00 | 1.43 | 0.00 |
| 6017.00 | 5690.00 | 5845.67 | -1.20 | 0.00 |
| 9.00 | 15.00 | 14.67 | -1.33 | 0.18 |
| 677.00 | 692.00 | 709.67 | -1.07 | 0.20 |
| 3.00 | 12.00 | 8.67 | 1.81 | 0.09 |
| 344.00 | 349.00 | 345.00 | 1.37 | 0.00 |
| 95.00 | 143.00 | 122.00 | 1.23 | 0.04 |
| 377.00 | 453.00 | 420.33 | 1.12 | 0.05 |
| 5.00 | 1.00 | 4.33 | 1.16 | 0.84 |
| 436.00 | 429.00 | 441.67 | 1.01 | 0.89 |
| 1177.00 | 1258.00 | 1267.67 | 1.12 | 0.01 |
| 293.00 | 286.00 | 289.00 | -1.22 | 0.00 |
| 524.00 | 438.00 | 490.67 | -2.17 | 0.00 |
| 28.00 | 12.00 | 23.67 | -1.41 | 0.06 |
| 2484.00 | 2629.00 | 2606.33 | 1.01 | 0.84 |
| 6585.00 | 7382.00 | 7316.67 | 1.13 | 0.00 |
| 911.00 | 935.00 | 930.33 | 1.74 | 0.00 |
| 625.00 | 670.00 | 681.67 | -1.12 | 0.01 |
| 371.00 | 365.00 | 348.33 | 1.04 | 0.59 |
| 57.00 | 48.00 | 52.00 | -1.27 | 0.05 |
| 133.00 | 133.00 | 142.33 | -1.48 | 0.00 |
| 36.00 | 27.00 | 31.00 | 1.03 | 0.83 |
| 719.00 | 763.00 | 779.67 | -1.17 | 0.00 |

| | | | | |
|---|---|---|---|---|
| 267.00 | 244.00 | 246.67 | 1.12 | 0.13 |
| 1262.00 | 1217.00 | 1248.00 | 1.01 | 0.82 |
| 1260.00 | 1343.00 | 1314.00 | 1.03 | 0.51 |
| 256.00 | 297.00 | 295.33 | 1.08 | 0.25 |
| 267.00 | 265.00 | 289.67 | 1.18 | 0.01 |
| 479.00 | 606.00 | 584.67 | -1.03 | 0.68 |
| 16.00 | 9.00 | 16.67 | 1.29 | 0.32 |
| 775.00 | 947.00 | 910.67 | 1.06 | 0.29 |
| 1295.00 | 1325.00 | 1390.00 | -1.01 | 0.83 |
| 557.00 | 480.00 | 537.00 | 1.06 | 0.32 |
| 33.00 | 52.00 | 41.00 | 1.17 | 0.34 |
| 1638.00 | 1823.00 | 1744.00 | 1.16 | 0.00 |
| 1078.00 | 1133.00 | 1167.00 | -1.02 | 0.69 |
| 100.00 | 88.00 | 95.67 | -1.12 | 0.23 |
| 501.00 | 507.00 | 525.33 | -1.06 | 0.21 |
| 2493.00 | 2711.00 | 2579.33 | -1.00 | 0.98 |
| 1067.00 | 1167.00 | 1132.67 | -1.20 | 0.00 |
| 2730.00 | 2721.00 | 2818.00 | -1.19 | 0.00 |
| 1205.00 | 1240.00 | 1249.67 | -1.05 | 0.29 |
| 1348.00 | 1242.00 | 1328.00 | -1.19 | 0.00 |
| 42.00 | 43.00 | 47.67 | 1.02 | 0.91 |
| 890.00 | 958.00 | 981.67 | 1.29 | 0.00 |
| 1240.00 | 1281.00 | 1259.33 | -1.14 | 0.01 |
| 2424.00 | 2789.00 | 2683.00 | 1.17 | 0.00 |
| 17.00 | 10.00 | 13.67 | -1.04 | 1.00 |
| 1.00 | 0.00 | 2.00 | -1.17 | 1.00 |
| 6.00 | 11.00 | 8.00 | 1.18 | 0.66 |
| 553.00 | 613.00 | 600.67 | 1.02 | 0.65 |
| 597.00 | 568.00 | 599.00 | -1.10 | 0.07 |
| 1208.00 | 1152.00 | 1201.67 | -1.87 | 0.00 |
| 2213.00 | 2220.00 | 2308.67 | -1.29 | 0.00 |
| 901.00 | 984.00 | 991.67 | -1.22 | 0.00 |
| 4.00 | 1.00 | 4.67 | -1.51 | 0.33 |
| 4077.00 | 4477.00 | 4483.67 | 1.09 | 0.03 |
| 100.00 | 120.00 | 104.67 | 1.25 | 0.03 |
| 3.00 | 9.00 | 5.00 | 1.23 | 0.71 |
| 82.00 | 99.00 | 95.33 | 1.06 | 0.58 |
| 460.00 | 462.00 | 469.67 | -1.13 | 0.03 |
| 966.00 | 1080.00 | 1044.33 | -1.10 | 0.03 |
| 4668.00 | 4787.00 | 4907.00 | -1.03 | 0.44 |
| 1102.00 | 1223.00 | 1177.33 | -1.12 | 0.01 |
| 21.00 | 30.00 | 25.00 | 1.80 | 0.00 |
| 142.00 | 151.00 | 147.33 | -1.16 | 0.06 |
| 714.00 | 706.00 | 738.67 | 1.10 | 0.05 |
| 410.00 | 604.00 | 511.67 | 1.24 | 0.00 |
| 253.00 | 304.00 | 285.33 | 1.49 | 0.00 |
| 270.00 | 311.00 | 311.33 | 1.42 | 0.00 |
| 660.00 | 653.00 | 645.67 | 1.29 | 0.00 |

| | | | | |
|---|---|---|---|---|
| 1397.00 | 1391.00 | 1405.33 | -1.05 | 0.22 |
| 3.00 | 1.00 | 1.67 | -3.46 | 0.01 |
| 2020.00 | 2060.00 | 2098.00 | -1.04 | 0.27 |
| 1208.00 | 1234.00 | 1242.67 | -1.04 | 0.33 |
| 149.00 | 152.00 | 155.33 | 1.16 | 0.06 |
| 23.00 | 30.00 | 30.00 | 1.55 | 0.02 |
| 1043.00 | 1079.00 | 1046.67 | -1.08 | 0.07 |
| 9.00 | 6.00 | 10.33 | -1.27 | 0.42 |
| 0.00 | 0.00 | 0.00 | -22.46 | 0.01 |
| 1.00 | 4.00 | 4.00 | 1.46 | 0.51 |
| 220.00 | 209.00 | 220.00 | -1.09 | 0.21 |
| 799.00 | 953.00 | 874.67 | -1.03 | 0.60 |
| 86.00 | 108.00 | 105.00 | 1.06 | 0.59 |
| 350.00 | 386.00 | 386.00 | -1.02 | 0.77 |
| 2705.00 | 2931.00 | 2895.33 | -1.19 | 0.00 |
| 172.00 | 213.00 | 190.00 | 1.02 | 0.82 |
| 256.00 | 303.00 | 278.33 | -1.10 | 0.16 |
| 890.00 | 925.00 | 899.67 | -1.53 | 0.00 |
| 39.00 | 24.00 | 36.67 | 2.00 | 0.00 |
| 1435.00 | 1630.00 | 1577.67 | 1.06 | 0.16 |
| 5367.00 | 5727.00 | 5583.67 | -1.33 | 0.00 |
| 5.00 | 3.00 | 4.67 | -1.44 | 0.40 |
| 1015.00 | 1024.00 | 1033.67 | -1.32 | 0.00 |
| 1153.00 | 1021.00 | 1075.67 | -1.08 | 0.16 |
| 607.00 | 585.00 | 614.00 | -1.11 | 0.05 |
| 92.00 | 111.00 | 107.33 | -1.15 | 0.12 |
| 1183.00 | 1175.00 | 1189.00 | -1.19 | 0.00 |
| 4.00 | 15.00 | 8.67 | 1.02 | 1.00 |
| 113.00 | 125.00 | 127.33 | -1.66 | 0.00 |
| 39.00 | 42.00 | 43.00 | -1.94 | 0.00 |
| 25.00 | 21.00 | 23.00 | 1.18 | 0.50 |
| 986.00 | 939.00 | 957.67 | -1.24 | 0.00 |
| 472.00 | 560.00 | 547.33 | 1.09 | 0.11 |
| 2646.00 | 2708.00 | 2763.67 | -1.09 | 0.03 |
| 3930.00 | 4310.00 | 4253.33 | 1.51 | 0.00 |
| 951.00 | 1116.00 | 1085.33 | 1.12 | 0.01 |
| 473.00 | 526.00 | 500.67 | 1.10 | 0.06 |
| 1162.00 | 1340.00 | 1317.33 | 1.43 | 0.00 |
| 372.00 | 460.00 | 447.33 | 1.19 | 0.01 |
| 1869.00 | 2086.00 | 2093.33 | 1.17 | 0.00 |
| 556.00 | 582.00 | 602.33 | 1.13 | 0.02 |
| 4181.00 | 4611.00 | 4371.00 | 1.24 | 0.00 |
| 1437.00 | 1617.00 | 1573.33 | 1.03 | 0.55 |
| 238.00 | 250.00 | 259.67 | -1.33 | 0.00 |
| 961.00 | 944.00 | 1035.00 | 1.23 | 0.00 |
| 352.00 | 312.00 | 347.67 | -1.18 | 0.01 |
| 558.00 | 595.00 | 558.33 | -1.10 | 0.10 |
| 3504.00 | 3594.00 | 3703.67 | 1.11 | 0.00 |

| | | | | |
|---|---|---|---|---|
| 2753.00 | 2920.00 | 2893.67 | 1.15 | 0.00 |
| 270.00 | 274.00 | 263.33 | -1.35 | 0.00 |
| 8.00 | 15.00 | 11.33 | 1.15 | 0.63 |
| 35.00 | 45.00 | 42.33 | 1.08 | 0.63 |
| 849.00 | 903.00 | 911.33 | 1.18 | 0.00 |
| 3.00 | 6.00 | 5.33 | 1.12 | 0.86 |
| 1.00 | 1.00 | 2.00 | -2.13 | 0.17 |
| 561.00 | 643.00 | 636.33 | 1.23 | 0.00 |
| 2178.00 | 2418.00 | 2387.67 | -1.05 | 0.18 |
| 785.00 | 761.00 | 821.67 | 1.06 | 0.29 |
| 354.00 | 397.00 | 377.67 | 1.08 | 0.20 |
| 374.00 | 354.00 | 375.33 | 1.08 | 0.19 |
| 20.00 | 27.00 | 26.33 | -1.31 | 0.09 |
| 271.00 | 294.00 | 284.00 | 1.03 | 0.70 |
| 1054.00 | 999.00 | 1072.33 | 1.17 | 0.00 |
| 1454.00 | 1773.00 | 1603.00 | 1.26 | 0.00 |
| 2.00 | 4.00 | 2.00 | -4.19 | 0.00 |
| 428.00 | 374.00 | 404.67 | -1.25 | 0.00 |
| 360.00 | 472.00 | 453.33 | 1.31 | 0.00 |
| 872.00 | 893.00 | 920.33 | -1.14 | 0.00 |
| 253.00 | 294.00 | 293.00 | -1.12 | 0.06 |
| 47.00 | 42.00 | 49.00 | -1.01 | 0.96 |
| 1297.00 | 1515.00 | 1477.67 | -1.14 | 0.00 |
| 2262.00 | 2362.00 | 2346.67 | 1.18 | 0.00 |
| 1586.00 | 1576.00 | 1577.33 | 1.02 | 0.62 |
| 2063.00 | 1969.00 | 2005.33 | -1.02 | 0.63 |
| 1279.00 | 1328.00 | 1300.33 | 1.08 | 0.15 |
| 3.00 | 1.00 | 2.00 | 1.17 | 1.00 |
| 281.00 | 291.00 | 286.33 | 1.16 | 0.03 |
| 3375.00 | 3281.00 | 3410.33 | 1.01 | 0.80 |
| 1662.00 | 1783.00 | 1725.33 | 1.32 | 0.00 |
| 867.00 | 933.00 | 902.67 | 1.12 | 0.04 |
| 8565.00 | 9062.00 | 9359.00 | 1.05 | 0.17 |
| 20.00 | 24.00 | 20.00 | -1.13 | 0.61 |
| 5.00 | 8.00 | 5.00 | -1.14 | 0.86 |
| 17.00 | 15.00 | 14.00 | 1.59 | 0.08 |
| 6015.00 | 6168.00 | 6285.00 | 1.13 | 0.00 |
| 192.00 | 199.00 | 198.33 | 1.00 | 1.00 |
| 2241.00 | 2363.00 | 2407.33 | -1.00 | 0.94 |
| 33001.00 | 35083.00 | 35998.00 | 1.03 | 0.38 |
| 787.00 | 782.00 | 825.00 | -1.26 | 0.00 |
| 885.00 | 918.00 | 883.00 | 1.15 | 0.03 |
| 331.00 | 341.00 | 341.33 | -1.06 | 0.32 |
| 807.00 | 889.00 | 833.33 | -1.19 | 0.00 |
| 40.00 | 45.00 | 53.33 | 1.13 | 0.44 |
| 758.00 | 849.00 | 865.33 | 1.11 | 0.04 |
| 2476.00 | 2546.00 | 2487.67 | -1.09 | 0.08 |
| 5.00 | 7.00 | 6.67 | -1.66 | 0.11 |

| | | | | |
|---|---|---|---|---|
| 48.00 | 46.00 | 50.00 | 1.15 | 0.32 |
| 204.00 | 206.00 | 210.67 | 1.48 | 0.00 |
| 1702.00 | 1969.00 | 1923.00 | 1.19 | 0.00 |
| 10.00 | 13.00 | 13.67 | 1.03 | 1.00 |
| 2318.00 | 2590.00 | 2560.00 | 1.23 | 0.00 |
| 3225.00 | 3745.00 | 3609.00 | 1.03 | 0.39 |
| 1086.00 | 1174.00 | 1165.67 | 1.06 | 0.17 |
| 1.00 | 5.00 | 3.67 | -1.64 | 0.28 |
| 1167.00 | 1252.00 | 1289.00 | 1.07 | 0.11 |
| 2743.00 | 2628.00 | 2819.33 | 1.01 | 0.90 |
| 794.00 | 812.00 | 808.33 | 1.09 | 0.06 |
| 2704.00 | 2980.00 | 2847.67 | 1.07 | 0.16 |
| 965.00 | 1066.00 | 994.33 | -1.06 | 0.22 |
| 1104.00 | 1048.00 | 1102.67 | -1.04 | 0.42 |
| 32.00 | 47.00 | 46.00 | -1.17 | 0.25 |
| 8.00 | 3.00 | 4.33 | -1.62 | 0.24 |
| 8.00 | 16.00 | 14.33 | -1.01 | 1.00 |
| 3756.00 | 3662.00 | 3606.00 | -1.74 | 0.00 |
| 693.00 | 663.00 | 708.67 | -1.14 | 0.01 |
| 2712.00 | 2854.00 | 2811.67 | -1.04 | 0.23 |
| 42.00 | 39.00 | 37.67 | -1.15 | 0.37 |
| 90.00 | 87.00 | 91.67 | -1.11 | 0.30 |
| 2676.00 | 3012.00 | 2871.00 | -1.17 | 0.00 |
| 2442.00 | 2500.00 | 2441.00 | -1.07 | 0.11 |
| 937.00 | 1037.00 | 1035.00 | 1.06 | 0.26 |
| 8.00 | 6.00 | 9.00 | -1.64 | 0.07 |
| 1.00 | 5.00 | 2.33 | -1.43 | 0.64 |
| 6.00 | 4.00 | 6.33 | 1.17 | 0.74 |
| 31.00 | 34.00 | 35.00 | -1.44 | 0.01 |
| 729.00 | 830.00 | 810.67 | 1.17 | 0.00 |
| 3.00 | 0.00 | 2.67 | -1.38 | 0.65 |
| 3423.00 | 4234.00 | 4091.33 | -1.14 | 0.01 |
| 2359.00 | 2869.00 | 2835.33 | -1.14 | 0.01 |
| 5.00 | 0.00 | 2.00 | -2.75 | 0.04 |
| 1.00 | 4.00 | 3.33 | -1.50 | 0.43 |
| 0.00 | 5.00 | 3.00 | -1.88 | 0.18 |
| 2378.00 | 2316.00 | 2403.67 | -1.43 | 0.00 |
| 1665.00 | 1751.00 | 1704.00 | -1.02 | 0.56 |
| 3529.00 | 3615.00 | 3553.67 | -1.01 | 0.87 |
| 1055.00 | 1097.00 | 1114.33 | 1.11 | 0.02 |
| 608.00 | 638.00 | 640.33 | 1.14 | 0.01 |
| 3126.00 | 3318.00 | 3172.33 | 1.17 | 0.00 |
| 1287.00 | 1247.00 | 1230.67 | -1.24 | 0.00 |
| 1519.00 | 1564.00 | 1576.67 | -1.04 | 0.27 |
| 581.00 | 644.00 | 649.33 | 1.08 | 0.14 |
| 1234.00 | 1294.00 | 1282.33 | -1.08 | 0.07 |
| 1877.00 | 1846.00 | 1915.00 | 1.06 | 0.14 |
| 805.00 | 910.00 | 882.67 | 1.02 | 0.60 |

| | | | | |
|---|---|---|---|---|
| 979.00 | 1031.00 | 1043.00 | -1.13 | 0.00 |
| 1927.00 | 2065.00 | 2057.00 | 1.14 | 0.00 |
| 2713.00 | 2794.00 | 2858.67 | 1.11 | 0.00 |
| 3036.00 | 3042.00 | 3178.33 | -1.08 | 0.09 |
| 324.00 | 354.00 | 337.67 | 1.06 | 0.29 |
| 1217.00 | 1326.00 | 1234.00 | -1.34 | 0.00 |
| 189.00 | 238.00 | 218.33 | -1.19 | 0.01 |
| 3255.00 | 3405.00 | 3378.33 | -1.09 | 0.01 |
| 207.00 | 234.00 | 227.00 | 1.01 | 0.93 |
| 365.00 | 472.00 | 411.33 | 1.00 | 0.97 |
| 441.00 | 473.00 | 464.33 | 1.00 | 0.97 |
| 1372.00 | 1387.00 | 1359.33 | -1.18 | 0.00 |
| 261.00 | 254.00 | 279.33 | -1.07 | 0.35 |
| 1167.00 | 1388.00 | 1349.33 | 1.14 | 0.01 |
| 1751.00 | 1633.00 | 1712.00 | -1.27 | 0.00 |
| 0.00 | 1.00 | 2.33 | -1.15 | 1.00 |
| 1555.00 | 1729.00 | 1686.00 | -1.01 | 0.74 |
| 2777.00 | 3157.00 | 3043.33 | -1.02 | 0.52 |
| 116.00 | 116.00 | 113.00 | 1.15 | 0.14 |
| 258.00 | 236.00 | 262.67 | 1.07 | 0.34 |
| 4.00 | 4.00 | 5.67 | -1.36 | 0.44 |
| 128.00 | 119.00 | 110.00 | -1.36 | 0.01 |
| 1238.00 | 1420.00 | 1372.00 | -1.09 | 0.04 |
| 8.00 | 6.00 | 11.33 | -1.08 | 0.83 |
| 533.00 | 582.00 | 564.33 | 1.05 | 0.37 |
| 900.00 | 992.00 | 968.00 | 1.21 | 0.00 |
| 3.00 | 7.00 | 6.67 | 1.23 | 0.63 |
| 583.00 | 594.00 | 574.00 | -1.00 | 1.00 |
| 2443.00 | 2656.00 | 2682.67 | 1.14 | 0.00 |
| 532.00 | 633.00 | 609.67 | 1.06 | 0.23 |
| 31.00 | 40.00 | 37.33 | 1.31 | 0.09 |
| 13.00 | 17.00 | 15.00 | 1.20 | 0.45 |
| 127.00 | 112.00 | 127.00 | 1.34 | 0.00 |
| 689.00 | 902.00 | 808.33 | 1.03 | 0.62 |
| 506.00 | 414.00 | 462.33 | -1.09 | 0.23 |
| 567.00 | 649.00 | 631.33 | 1.20 | 0.00 |
| 907.00 | 849.00 | 882.67 | -1.92 | 0.00 |
| 3.00 | 5.00 | 4.00 | -1.67 | 0.22 |
| 2912.00 | 2956.00 | 2937.33 | -1.07 | 0.08 |
| 1398.00 | 1571.00 | 1574.33 | -1.13 | 0.00 |
| 226.00 | 198.00 | 213.00 | -1.57 | 0.00 |
| 1419.00 | 1433.00 | 1502.33 | -1.10 | 0.02 |
| 469.00 | 473.00 | 479.67 | -1.10 | 0.07 |
| 41.00 | 69.00 | 54.00 | -1.13 | 0.34 |
| 1150.00 | 1290.00 | 1334.00 | 1.03 | 0.58 |
| 12.00 | 10.00 | 11.67 | -1.79 | 0.01 |
| 107.00 | 118.00 | 113.67 | -1.51 | 0.00 |
| 96.00 | 119.00 | 111.00 | -1.47 | 0.00 |

| | | | | |
|---|---|---|---|---|
| 22.00 | 20.00 | 23.00 | -1.42 | 0.03 |
| 42.00 | 23.00 | 36.67 | -1.75 | 0.00 |
| 10.00 | 3.00 | 7.33 | 2.13 | 0.06 |
| 23.00 | 18.00 | 22.67 | 1.08 | 0.74 |
| 810.00 | 746.00 | 776.33 | -1.18 | 0.00 |
| 103.00 | 102.00 | 107.33 | -1.15 | 0.12 |
| 477.00 | 448.00 | 504.33 | -1.30 | 0.00 |
| 883.00 | 834.00 | 874.00 | -1.03 | 0.58 |
| 1005.00 | 1031.00 | 1001.00 | 1.12 | 0.02 |
| 37.00 | 66.00 | 50.33 | -1.07 | 0.67 |
| 297.00 | 288.00 | 292.67 | -1.08 | 0.22 |
| 10.00 | 8.00 | 11.33 | -1.16 | 0.65 |
| 200.00 | 198.00 | 203.00 | 1.09 | 0.23 |
| 1439.00 | 1467.00 | 1513.67 | -1.12 | 0.00 |
| 104.00 | 101.00 | 102.67 | -2.13 | 0.00 |
| 0.00 | 0.00 | 0.00 | -11.74 | 0.13 |
| 461.00 | 539.00 | 483.67 | -1.02 | 0.71 |
| 4.00 | 3.00 | 4.67 | 1.92 | 0.20 |
| 44.00 | 43.00 | 42.00 | -1.45 | 0.00 |
| 125.00 | 133.00 | 127.67 | -1.32 | 0.00 |
| 6.00 | 6.00 | 5.67 | 1.19 | 0.72 |
| 71.00 | 58.00 | 67.00 | -1.08 | 0.52 |
| 4.00 | 3.00 | 3.33 | -1.31 | 0.68 |
| 138.00 | 130.00 | 137.33 | -1.04 | 0.68 |
| 104.00 | 92.00 | 97.33 | -1.13 | 0.20 |
| 393.00 | 418.00 | 408.33 | -1.15 | 0.01 |
| 18.00 | 8.00 | 13.67 | -1.67 | 0.02 |
| 725.00 | 612.00 | 645.00 | -1.31 | 0.00 |
| 166.00 | 153.00 | 160.67 | -1.08 | 0.34 |
| 5.00 | 2.00 | 2.67 | -1.01 | 1.00 |
| 2188.00 | 2279.00 | 2324.33 | 1.01 | 0.80 |
| 711.00 | 799.00 | 797.33 | -1.03 | 0.61 |
| 1851.00 | 1897.00 | 1892.67 | -1.21 | 0.00 |
| 444.00 | 533.00 | 483.00 | 1.27 | 0.00 |
| 20.00 | 19.00 | 19.00 | -1.22 | 0.37 |
| 2174.00 | 2128.00 | 2241.33 | -1.07 | 0.10 |
| 2370.00 | 2410.00 | 2522.67 | 1.13 | 0.00 |
| 2041.00 | 2118.00 | 2190.33 | 1.21 | 0.00 |
| 2667.00 | 2776.00 | 2859.33 | 1.18 | 0.00 |
| 1134.00 | 1115.00 | 1142.67 | -1.09 | 0.04 |
| 89.00 | 104.00 | 103.33 | -1.17 | 0.08 |
| 111.00 | 133.00 | 120.33 | 1.05 | 0.60 |
| 395.00 | 351.00 | 386.00 | 1.05 | 0.41 |
| 2778.00 | 2875.00 | 2867.33 | -1.09 | 0.01 |
| 65.00 | 95.00 | 87.00 | 1.05 | 0.71 |
| 1931.00 | 1989.00 | 1996.00 | 1.05 | 0.19 |
| 41.00 | 35.00 | 44.00 | -1.35 | 0.02 |
| 92.00 | 86.00 | 82.33 | 1.03 | 0.79 |

| | | | | |
|---|---|---|---|---|
| 27.00 | 30.00 | 29.33 | -1.04 | 0.83 |
| 1736.00 | 1790.00 | 1852.00 | 1.14 | 0.00 |
| 7.00 | 3.00 | 5.67 | 1.39 | 0.47 |
| 4.00 | 10.00 | 6.33 | -1.54 | 0.21 |
| 0.00 | 0.00 | 0.00 | -11.74 | 0.13 |
| 2.00 | 0.00 | 1.00 | -2.81 | 0.15 |
| 14.00 | 19.00 | 17.00 | -2.06 | 0.00 |
| 11.00 | 20.00 | 15.67 | -1.81 | 0.00 |
| 9.00 | 15.00 | 12.33 | -1.64 | 0.03 |
| 71.00 | 82.00 | 75.33 | -1.32 | 0.01 |
| 4356.00 | 4424.00 | 4486.67 | -1.10 | 0.01 |
| 19.00 | 18.00 | 20.33 | -1.36 | 0.08 |
| 1365.00 | 1480.00 | 1467.33 | -1.15 | 0.00 |
| 165.00 | 157.00 | 168.33 | -1.14 | 0.10 |
| 213.00 | 249.00 | 235.67 | -1.14 | 0.04 |
| 976.00 | 1054.00 | 1065.33 | 1.35 | 0.00 |
| 0.00 | 1.00 | 1.33 | -2.40 | 0.19 |
| 3.00 | 1.00 | 3.00 | -3.18 | 0.00 |
| 5.00 | 2.00 | 5.33 | 3.00 | 0.03 |
| 11.00 | 7.00 | 9.33 | 1.38 | 0.35 |
| 9534.00 | 9883.00 | 9956.00 | -1.32 | 0.00 |
| 1132.00 | 1163.00 | 1224.67 | 1.19 | 0.00 |
| 1087.00 | 1239.00 | 1226.33 | 1.23 | 0.00 |
| 465.00 | 429.00 | 444.00 | -1.02 | 0.80 |
| 2166.00 | 2442.00 | 2411.67 | 1.11 | 0.01 |
| 881.00 | 868.00 | 872.00 | 1.06 | 0.20 |
| 372.00 | 407.00 | 389.00 | -1.05 | 0.36 |
| 4.00 | 8.00 | 7.00 | -1.68 | 0.09 |
| 721.00 | 834.00 | 812.33 | 1.27 | 0.00 |
| 444.00 | 492.00 | 489.33 | -1.20 | 0.00 |
| 1346.00 | 1302.00 | 1313.33 | -1.42 | 0.00 |
| 15.00 | 14.00 | 16.67 | 1.03 | 0.92 |
| 268.00 | 259.00 | 266.67 | 1.04 | 0.52 |
| 1025.00 | 1113.00 | 1089.33 | 1.07 | 0.12 |
| 1653.00 | 1622.00 | 1646.33 | -1.08 | 0.09 |
| 310.00 | 267.00 | 305.67 | 1.23 | 0.01 |
| 323.00 | 331.00 | 344.00 | 1.19 | 0.01 |
| 5094.00 | 5620.00 | 5469.67 | 1.01 | 0.87 |
| 414.00 | 348.00 | 393.33 | -1.08 | 0.29 |
| 791.00 | 765.00 | 800.67 | 1.22 | 0.00 |
| 2610.00 | 2980.00 | 2906.67 | -1.04 | 0.26 |
| 162.00 | 211.00 | 188.00 | 1.23 | 0.01 |
| 378.00 | 441.00 | 406.67 | 1.13 | 0.05 |
| 2207.00 | 2298.00 | 2398.00 | -1.02 | 0.67 |
| 50.00 | 55.00 | 61.00 | 1.02 | 0.92 |
| 621.00 | 611.00 | 669.67 | -1.06 | 0.25 |
| 197.00 | 187.00 | 197.67 | 1.09 | 0.33 |
| 3811.00 | 3983.00 | 4215.33 | 1.08 | 0.09 |

| | | | | |
|---|---|---|---|---|
| 77.00 | 65.00 | 78.67 | 1.15 | 0.23 |
| 3762.00 | 4148.00 | 4151.00 | 1.19 | 0.00 |
| 215.00 | 200.00 | 230.33 | -1.10 | 0.36 |
| 536.00 | 619.00 | 587.00 | 1.40 | 0.00 |
| 1235.00 | 1442.00 | 1301.67 | 1.26 | 0.00 |
| 576.00 | 648.00 | 627.00 | 1.14 | 0.01 |
| 512.00 | 603.00 | 589.00 | 1.11 | 0.05 |
| 601.00 | 597.00 | 619.00 | 1.11 | 0.04 |
| 566.00 | 631.00 | 612.00 | 1.17 | 0.00 |
| 1667.00 | 1749.00 | 1695.00 | -1.05 | 0.21 |
| 1095.00 | 1357.00 | 1320.67 | 1.17 | 0.00 |
| 1272.00 | 1327.00 | 1345.00 | 1.19 | 0.00 |
| 13.00 | 4.00 | 10.67 | -1.08 | 0.91 |
| 6.00 | 12.00 | 7.00 | 1.04 | 1.00 |
| 1550.00 | 1778.00 | 1673.33 | 1.17 | 0.00 |
| 97.00 | 102.00 | 104.33 | -1.26 | 0.01 |
| 468.00 | 523.00 | 524.00 | 1.10 | 0.13 |
| 1085.00 | 912.00 | 949.67 | -1.10 | 0.18 |
| 92.00 | 86.00 | 98.33 | -1.01 | 0.94 |
| 452.00 | 472.00 | 469.67 | -1.11 | 0.03 |
| 494.00 | 498.00 | 527.00 | -1.07 | 0.17 |
| 31.00 | 12.00 | 19.67 | -1.52 | 0.06 |
| 43.00 | 38.00 | 41.33 | -1.33 | 0.03 |
| 1753.00 | 2001.00 | 1963.00 | 1.16 | 0.00 |
| 3336.00 | 3446.00 | 3366.00 | -1.14 | 0.00 |
| 1735.00 | 1770.00 | 1844.00 | 1.00 | 0.95 |
| 1768.00 | 2095.00 | 1946.00 | -1.03 | 0.55 |
| 840.00 | 690.00 | 743.00 | 1.17 | 0.03 |
| 1262.00 | 1323.00 | 1354.33 | 1.03 | 0.49 |
| 683.00 | 862.00 | 845.00 | 1.18 | 0.01 |
| 1018.00 | 930.00 | 940.00 | -1.24 | 0.00 |
| 670.00 | 811.00 | 749.00 | 1.00 | 0.93 |
| 5411.00 | 5734.00 | 5869.67 | -1.18 | 0.00 |
| 878.00 | 935.00 | 924.67 | -1.04 | 0.35 |
| 4040.00 | 4459.00 | 4262.33 | 1.13 | 0.00 |
| 9.00 | 1.00 | 5.00 | 1.34 | 0.57 |
| 392.00 | 381.00 | 406.67 | 1.21 | 0.00 |
| 512.00 | 527.00 | 528.00 | -1.01 | 0.83 |
| 8.00 | 12.00 | 11.00 | 1.79 | 0.05 |
| 64.00 | 44.00 | 53.33 | -1.09 | 0.54 |
| 877.00 | 841.00 | 881.67 | -1.01 | 0.80 |
| 5.00 | 14.00 | 8.67 | 1.03 | 1.00 |
| 27.00 | 18.00 | 24.33 | 1.20 | 0.33 |
| 144.00 | 146.00 | 151.67 | 1.17 | 0.05 |
| 1969.00 | 2094.00 | 2043.67 | 1.03 | 0.45 |
| 1570.00 | 1673.00 | 1720.67 | 1.02 | 0.69 |
| 236.00 | 309.00 | 258.33 | 1.28 | 0.00 |
| 2562.00 | 2643.00 | 2711.00 | 1.01 | 0.74 |

| | | | | |
|---|---|---|---|---|
| 167.00 | 228.00 | 208.33 | 1.01 | 0.89 |
| 418.00 | 400.00 | 416.00 | -1.35 | 0.00 |
| 17.00 | 20.00 | 20.00 | 1.14 | 0.58 |
| 325.00 | 380.00 | 368.00 | 1.05 | 0.50 |
| 11.00 | 23.00 | 14.00 | 1.12 | 0.70 |
| 1477.00 | 1550.00 | 1480.67 | -1.03 | 0.59 |
| 69.00 | 91.00 | 85.00 | 1.05 | 0.66 |
| 1232.00 | 1187.00 | 1267.00 | 1.21 | 0.00 |
| 445.00 | 395.00 | 433.33 | -1.02 | 0.76 |
| 29.00 | 37.00 | 34.67 | 1.07 | 0.74 |
| 1144.00 | 1189.00 | 1150.67 | 1.05 | 0.28 |
| 798.00 | 825.00 | 832.00 | -1.06 | 0.19 |
| 66.00 | 104.00 | 82.33 | -1.14 | 0.23 |
| 1398.00 | 1414.00 | 1477.33 | 1.05 | 0.23 |
| 1169.00 | 1091.00 | 1170.67 | -1.04 | 0.40 |
| 85.00 | 107.00 | 107.67 | -1.15 | 0.14 |
| 433.00 | 488.00 | 482.33 | 1.14 | 0.02 |
| 2205.00 | 2543.00 | 2469.00 | 1.15 | 0.00 |
| 1105.00 | 1147.00 | 1146.67 | 1.17 | 0.00 |
| 2388.00 | 2704.00 | 2632.67 | -1.02 | 0.57 |
| 4612.00 | 5340.00 | 5324.00 | 1.34 | 0.00 |
| 109.00 | 104.00 | 115.00 | 1.75 | 0.00 |
| 9108.00 | 10056.00 | 10265.67 | 1.22 | 0.00 |
| 67.00 | 60.00 | 60.33 | 1.23 | 0.10 |
| 2900.00 | 2590.00 | 2696.33 | -1.36 | 0.00 |
| 479.00 | 473.00 | 468.33 | 1.09 | 0.15 |
| 447.00 | 593.00 | 535.67 | 1.12 | 0.09 |
| 476.00 | 430.00 | 416.33 | -1.04 | 0.74 |
| 3870.00 | 4045.00 | 4033.33 | 1.10 | 0.01 |
| 229.00 | 207.00 | 216.00 | 1.35 | 0.00 |
| 429.00 | 516.00 | 473.33 | 1.08 | 0.25 |
| 664.00 | 728.00 | 704.33 | 1.22 | 0.00 |
| 571.00 | 568.00 | 577.67 | 1.27 | 0.00 |
| 4020.00 | 4435.00 | 4387.33 | 1.10 | 0.01 |
| 2010.00 | 2196.00 | 2191.67 | 1.14 | 0.00 |
| 556.00 | 568.00 | 584.00 | 1.09 | 0.07 |
| 506.00 | 480.00 | 520.00 | -1.13 | 0.04 |
| 107.00 | 132.00 | 131.67 | -1.65 | 0.00 |
| 4.00 | 10.00 | 7.00 | 1.85 | 0.12 |
| 79.00 | 63.00 | 73.67 | 1.36 | 0.01 |
| 7.00 | 8.00 | 7.00 | 1.58 | 0.24 |
| 44.00 | 43.00 | 47.67 | 1.15 | 0.33 |
| 80.00 | 91.00 | 90.33 | 1.16 | 0.14 |
| 1990.00 | 2364.00 | 2231.33 | 1.03 | 0.51 |
| 1038.00 | 1324.00 | 1233.33 | -1.01 | 0.91 |
| 62.00 | 81.00 | 77.33 | 1.25 | 0.05 |
| 12.00 | 9.00 | 11.67 | 2.01 | 0.02 |
| 102.00 | 83.00 | 95.00 | -1.05 | 0.66 |

| | | | | |
|---|---|---|---|---|
| 1059.00 | 1189.00 | 1184.33 | 1.35 | 0.00 |
| 277.00 | 306.00 | 322.00 | 1.26 | 0.00 |
| 488.00 | 474.00 | 488.67 | 1.22 | 0.00 |
| 395.00 | 491.00 | 449.33 | 1.04 | 0.51 |
| 751.00 | 772.00 | 789.33 | 1.12 | 0.01 |
| 737.00 | 733.00 | 752.33 | -1.04 | 0.42 |
| 1684.00 | 1842.00 | 1785.67 | 1.09 | 0.03 |
| 524.00 | 577.00 | 587.67 | 1.05 | 0.40 |
| 179.00 | 185.00 | 203.67 | 1.26 | 0.01 |
| 3.00 | 1.00 | 3.33 | -1.31 | 0.68 |
| 2920.00 | 3130.00 | 3230.67 | 1.20 | 0.00 |
| 183.00 | 208.00 | 210.67 | 1.85 | 0.00 |
| 3.00 | 0.00 | 2.00 | -1.17 | 1.00 |
| 25.00 | 21.00 | 23.00 | 1.33 | 0.13 |
| 6512.00 | 6209.00 | 6668.00 | -1.06 | 0.14 |
| 652.00 | 649.00 | 666.33 | -1.02 | 0.63 |
| 5.00 | 1.00 | 3.00 | -1.66 | 0.32 |
| 1450.00 | 1581.00 | 1547.33 | 1.28 | 0.00 |
| 99.00 | 92.00 | 96.67 | -1.08 | 0.44 |
| 402.00 | 460.00 | 460.00 | 1.03 | 0.64 |
| 2645.00 | 2518.00 | 2555.00 | -1.09 | 0.05 |
| 576.00 | 619.00 | 634.00 | 1.43 | 0.00 |
| 229.00 | 259.00 | 246.00 | 1.08 | 0.25 |
| 17.00 | 5.00 | 12.33 | -1.15 | 0.68 |
| 1.00 | 4.00 | 2.33 | -1.56 | 0.49 |
| 7.00 | 22.00 | 15.67 | 1.08 | 0.86 |
| 3691.00 | 4068.00 | 4048.00 | 1.05 | 0.16 |
| 65.00 | 63.00 | 64.00 | -2.11 | 0.00 |
| 1.00 | 0.00 | 0.33 | -5.41 | 0.08 |
| 2381.00 | 2545.00 | 2526.00 | 1.01 | 0.74 |
| 979.00 | 995.00 | 1049.33 | -1.27 | 0.00 |
| 38.00 | 53.00 | 54.00 | -1.02 | 0.89 |
| 3618.00 | 3974.00 | 3950.67 | 1.17 | 0.00 |
| 1142.00 | 1244.00 | 1251.00 | -1.01 | 0.88 |
| 435.00 | 438.00 | 442.33 | -1.05 | 0.32 |
| 407.00 | 370.00 | 389.67 | -1.31 | 0.00 |
| 1027.00 | 958.00 | 1049.67 | -1.30 | 0.00 |
| 3345.00 | 3305.00 | 3484.67 | -1.09 | 0.03 |
| 374.00 | 459.00 | 424.33 | 1.05 | 0.37 |
| 833.00 | 899.00 | 897.67 | 1.32 | 0.00 |
| 1111.00 | 1180.00 | 1212.00 | 1.25 | 0.00 |
| 356.00 | 432.00 | 420.33 | 1.05 | 0.41 |
| 7661.00 | 7914.00 | 7986.00 | 1.10 | 0.01 |
| 1387.00 | 1584.00 | 1505.67 | 1.10 | 0.04 |
| 941.00 | 1065.00 | 1037.33 | 1.30 | 0.00 |
| 489.00 | 508.00 | 517.67 | 1.09 | 0.11 |
| 454.00 | 529.00 | 516.00 | -1.05 | 0.46 |
| 266.00 | 281.00 | 287.00 | 1.03 | 0.68 |

| | | | | |
|---|---|---|---|---|
| 12092.00 | 12173.00 | 12388.33 | -1.05 | 0.17 |
| 123.00 | 139.00 | 126.67 | 1.01 | 0.90 |
| 3914.00 | 4080.00 | 4055.33 | -1.07 | 0.06 |
| 603.00 | 666.00 | 624.67 | 1.00 | 0.94 |
| 606.00 | 537.00 | 541.33 | -1.18 | 0.02 |
| 1165.00 | 1256.00 | 1214.33 | -1.11 | 0.01 |
| 1973.00 | 2182.00 | 2108.00 | -1.12 | 0.00 |
| 1091.00 | 1213.00 | 1212.33 | 1.15 | 0.00 |
| 407.00 | 435.00 | 452.00 | -1.06 | 0.31 |
| 460.00 | 510.00 | 499.33 | -1.00 | 0.95 |
| 3725.00 | 3795.00 | 3724.33 | -1.14 | 0.00 |
| 394.00 | 421.00 | 404.33 | -1.01 | 0.83 |
| 864.00 | 861.00 | 898.67 | -1.25 | 0.00 |
| 1448.00 | 1283.00 | 1330.00 | -1.42 | 0.00 |
| 1003.00 | 1016.00 | 1005.33 | 1.24 | 0.00 |
| 1759.00 | 1604.00 | 1609.67 | -1.51 | 0.00 |
| 430.00 | 463.00 | 458.67 | 1.14 | 0.01 |
| 64.00 | 85.00 | 74.00 | 1.05 | 0.66 |
| 359.00 | 438.00 | 415.33 | 1.57 | 0.00 |
| 205.00 | 216.00 | 233.00 | -1.02 | 0.79 |
| 887.00 | 891.00 | 874.67 | -1.08 | 0.10 |
| 1184.00 | 1070.00 | 1126.67 | 1.05 | 0.29 |
| 1426.00 | 1450.00 | 1445.67 | -1.29 | 0.00 |
| 1.00 | 3.00 | 2.00 | -2.12 | 0.17 |
| 1151.00 | 1224.00 | 1158.00 | -1.13 | 0.01 |
| 1501.00 | 1509.00 | 1472.33 | -1.31 | 0.00 |
| 392.00 | 475.00 | 464.33 | -1.28 | 0.00 |
| 183.00 | 143.00 | 163.67 | -1.66 | 0.00 |
| 44.00 | 50.00 | 48.33 | -1.11 | 0.42 |
| 25.00 | 20.00 | 22.67 | 1.07 | 0.74 |
| 176.00 | 168.00 | 158.67 | -1.07 | 0.48 |
| 2883.00 | 3147.00 | 3070.00 | -1.06 | 0.07 |
| 90.00 | 101.00 | 92.67 | 1.61 | 0.00 |
| 108.00 | 119.00 | 119.00 | 1.00 | 1.00 |
| 2057.00 | 1965.00 | 2123.33 | -1.02 | 0.73 |
| 861.00 | 891.00 | 869.67 | -1.06 | 0.23 |
| 115.00 | 122.00 | 120.00 | -1.58 | 0.00 |
| 0.00 | 0.00 | 0.00 | -11.75 | 0.13 |
| 94.00 | 99.00 | 96.33 | 1.26 | 0.02 |
| 109.00 | 143.00 | 133.67 | 1.66 | 0.00 |
| 16.00 | 22.00 | 21.33 | -1.22 | 0.31 |
| 34.00 | 33.00 | 35.33 | 1.77 | 0.00 |
| 4992.00 | 4875.00 | 5075.33 | -1.10 | 0.01 |
| 6.00 | 11.00 | 8.33 | -1.49 | 0.17 |
| 4979.00 | 5048.00 | 5118.00 | 1.06 | 0.15 |
| 446.00 | 472.00 | 477.00 | -1.06 | 0.23 |
| 535.00 | 656.00 | 640.67 | 1.01 | 0.88 |
| 100.00 | 88.00 | 98.67 | 1.07 | 0.55 |

| | | | | |
|---|---|---|---|---|
| 446.00 | 486.00 | 499.67 | -1.03 | 0.52 |
| 54.00 | 66.00 | 64.33 | -1.04 | 0.78 |
| 417.00 | 409.00 | 406.33 | 1.16 | 0.03 |
| 47.00 | 59.00 | 55.33 | 1.30 | 0.04 |
| 3.00 | 1.00 | 1.67 | -1.20 | 1.00 |
| 11.00 | 2.00 | 6.33 | -2.05 | 0.02 |
| 734.00 | 748.00 | 746.33 | 1.07 | 0.16 |
| 7712.00 | 7620.00 | 8035.67 | -1.16 | 0.00 |
| 1427.00 | 1422.00 | 1465.67 | -1.10 | 0.02 |
| 204.00 | 191.00 | 202.33 | -1.41 | 0.00 |
| 275.00 | 335.00 | 322.67 | 1.45 | 0.00 |
| 449.00 | 544.00 | 496.33 | 1.09 | 0.13 |
| 95.00 | 108.00 | 106.00 | -1.09 | 0.37 |
| 9.00 | 10.00 | 11.67 | -1.93 | 0.00 |
| 12.00 | 13.00 | 14.67 | -1.06 | 0.92 |
| 217.00 | 257.00 | 254.67 | 1.20 | 0.01 |
| 1355.00 | 1529.00 | 1485.33 | 1.18 | 0.00 |
| 1367.00 | 1457.00 | 1473.00 | 1.03 | 0.42 |
| 82.00 | 73.00 | 75.33 | -1.18 | 0.14 |
| 1433.00 | 1623.00 | 1634.00 | 1.20 | 0.00 |
| 416.00 | 525.00 | 476.00 | 1.14 | 0.02 |
| 113.00 | 129.00 | 125.00 | -1.39 | 0.00 |
| 152.00 | 154.00 | 160.33 | -1.08 | 0.32 |
| 10.00 | 17.00 | 16.33 | 1.50 | 0.11 |
| 7084.00 | 7425.00 | 6972.33 | -1.24 | 0.00 |
| 14.00 | 20.00 | 15.67 | 1.92 | 0.01 |
| 1830.00 | 1809.00 | 1851.33 | -1.12 | 0.00 |
| 1186.00 | 1423.00 | 1374.33 | 1.24 | 0.00 |
| 598.00 | 597.00 | 672.33 | 1.10 | 0.15 |
| 396.00 | 425.00 | 418.00 | 1.23 | 0.00 |
| 454.00 | 501.00 | 503.33 | -1.00 | 0.95 |
| 4699.00 | 5171.00 | 5269.00 | 1.15 | 0.00 |
| 488.00 | 569.00 | 556.33 | 1.10 | 0.09 |
| 617.00 | 616.00 | 623.33 | 1.14 | 0.00 |
| 1272.00 | 1293.00 | 1248.00 | -1.32 | 0.00 |
| 634.00 | 662.00 | 656.33 | -1.26 | 0.00 |
| 925.00 | 1058.00 | 971.67 | -1.18 | 0.00 |
| 13.00 | 24.00 | 15.33 | 1.03 | 0.92 |
| 6139.00 | 5619.00 | 5658.33 | -1.02 | 0.73 |
| 772.00 | 896.00 | 815.67 | -1.04 | 0.45 |
| 1021.00 | 1100.00 | 1127.67 | -1.01 | 0.75 |
| 458.00 | 549.00 | 524.33 | 1.21 | 0.00 |
| 13.00 | 24.00 | 20.00 | -1.08 | 0.73 |
| 15.00 | 16.00 | 14.33 | 1.15 | 0.59 |
| 4.00 | 0.00 | 2.00 | -1.65 | 0.47 |
| 3.00 | 2.00 | 4.33 | -1.17 | 0.85 |
| 1.00 | 3.00 | 2.33 | 2.16 | 0.35 |
| 3.00 | 4.00 | 2.67 | -1.01 | 1.00 |

| | | | | |
|---|---|---|---|---|
| 16.00 | 11.00 | 16.67 | -1.09 | 0.71 |
| 3.00 | 2.00 | 2.33 | -1.97 | 0.20 |
| 182.00 | 208.00 | 211.33 | -1.08 | 0.24 |
| 6.00 | 13.00 | 11.67 | -1.04 | 0.91 |
| 0.00 | 1.00 | 0.33 | -3.95 | 0.22 |
| 2.00 | 4.00 | 2.33 | -1.15 | 1.00 |
| 9.00 | 7.00 | 9.33 | -1.19 | 0.63 |
| 15.00 | 14.00 | 12.67 | -1.38 | 0.20 |
| 1.00 | 3.00 | 2.33 | -2.53 | 0.05 |
| 11.00 | 19.00 | 17.33 | 1.46 | 0.12 |
| 3.00 | 3.00 | 2.67 | 1.30 | 0.79 |
| 50.00 | 44.00 | 46.00 | 1.00 | 1.00 |
| 4.00 | 8.00 | 5.33 | -1.57 | 0.22 |
| 715.00 | 639.00 | 666.33 | -1.57 | 0.00 |
| 1220.00 | 1093.00 | 1129.00 | -1.21 | 0.00 |
| 462.00 | 424.00 | 458.33 | -1.03 | 0.65 |
| 1556.00 | 1737.00 | 1724.00 | 1.18 | 0.00 |
| 4661.00 | 4647.00 | 4743.67 | -1.07 | 0.05 |
| 5513.00 | 5759.00 | 5634.67 | 1.15 | 0.00 |
| 153.00 | 139.00 | 141.33 | -1.42 | 0.00 |
| 439.00 | 504.00 | 510.00 | -1.06 | 0.36 |
| 2567.00 | 2451.00 | 2565.33 | -1.06 | 0.28 |
| 344.00 | 365.00 | 358.00 | -1.03 | 0.69 |
| 691.00 | 700.00 | 724.33 | -1.18 | 0.00 |
| 12.00 | 23.00 | 15.33 | 1.96 | 0.01 |
| 137.00 | 121.00 | 130.33 | 1.52 | 0.00 |
| 2.00 | 9.00 | 6.00 | 1.04 | 1.00 |
| 1.00 | 2.00 | 1.67 | -1.01 | 1.00 |
| 1195.00 | 1158.00 | 1196.67 | -1.03 | 0.44 |
| 46.00 | 33.00 | 37.00 | 1.71 | 0.00 |
| 2533.00 | 3020.00 | 2802.33 | 1.10 | 0.02 |
| 219.00 | 267.00 | 248.67 | -1.11 | 0.11 |
| 9.00 | 8.00 | 8.67 | -1.55 | 0.12 |
| 4908.00 | 5016.00 | 5137.00 | -1.18 | 0.00 |
| 64.00 | 66.00 | 61.00 | -1.66 | 0.00 |
| 677.00 | 834.00 | 790.67 | 1.12 | 0.02 |
| 630.00 | 664.00 | 703.00 | 1.23 | 0.00 |
| 65.00 | 61.00 | 62.33 | -1.10 | 0.47 |
| 3.00 | 4.00 | 5.00 | 1.22 | 0.71 |
| 929.00 | 968.00 | 938.00 | 1.02 | 0.63 |
| 1683.00 | 1750.00 | 1802.33 | 1.12 | 0.01 |
| 31721.00 | 31469.00 | 32780.00 | -1.29 | 0.00 |
| 4948.00 | 5469.00 | 5223.33 | -1.04 | 0.28 |
| 3999.00 | 4986.00 | 4639.00 | 1.32 | 0.00 |
| 2937.00 | 3560.00 | 3436.00 | 1.39 | 0.00 |
| 548.00 | 598.00 | 614.33 | 1.22 | 0.00 |
| 642.00 | 616.00 | 659.33 | -1.05 | 0.34 |
| 1238.00 | 1349.00 | 1307.67 | 1.02 | 0.57 |

| | | | | |
|---|---|---|---|---|
| 565.00 | 660.00 | 637.67 | 1.01 | 0.91 |
| 3196.00 | 3497.00 | 3446.00 | 1.25 | 0.00 |
| 2416.00 | 2761.00 | 2748.33 | 1.16 | 0.00 |
| 754.00 | 946.00 | 879.67 | -1.04 | 0.45 |
| 18.00 | 32.00 | 22.33 | -1.03 | 0.94 |
| 1852.00 | 2090.00 | 2034.67 | 1.01 | 0.75 |
| 414.00 | 517.00 | 471.33 | 1.49 | 0.00 |
| 1644.00 | 1717.00 | 1735.33 | -1.10 | 0.01 |
| 827.00 | 888.00 | 876.00 | -1.03 | 0.50 |
| 59.00 | 64.00 | 62.00 | -1.55 | 0.00 |
| 923.00 | 1063.00 | 1002.67 | 1.13 | 0.01 |
| 498.00 | 577.00 | 565.67 | 1.63 | 0.00 |
| 876.00 | 948.00 | 960.00 | 1.22 | 0.00 |
| 1213.00 | 1310.00 | 1313.00 | -1.06 | 0.12 |
| 83.00 | 111.00 | 105.00 | 1.44 | 0.00 |
| 4.00 | 0.00 | 2.00 | -1.01 | 1.00 |
| 3619.00 | 3717.00 | 3784.33 | 1.18 | 0.00 |
| 10339.00 | 11704.00 | 11622.00 | 1.17 | 0.00 |
| 209.00 | 227.00 | 219.33 | -1.06 | 0.42 |
| 2293.00 | 2876.00 | 2659.00 | 1.39 | 0.00 |
| 25453.00 | 31085.00 | 29178.33 | 1.19 | 0.00 |
| 9446.00 | 11207.00 | 10770.67 | 1.06 | 0.15 |
| 4697.00 | 5199.00 | 5168.00 | 1.26 | 0.00 |
| 12817.00 | 14545.00 | 14386.33 | 1.17 | 0.00 |
| 5015.00 | 5863.00 | 5716.67 | 1.21 | 0.00 |
| 5888.00 | 6935.00 | 6602.00 | 1.06 | 0.11 |
| 7179.00 | 7629.00 | 7699.67 | 1.12 | 0.00 |
| 13032.00 | 15928.00 | 14743.00 | 1.02 | 0.61 |
| 34.00 | 29.00 | 34.33 | 1.15 | 0.37 |
| 5390.00 | 6450.00 | 6173.33 | 1.14 | 0.00 |
| 13541.00 | 15091.00 | 14770.67 | 1.09 | 0.01 |
| 4602.00 | 4903.00 | 4971.33 | 1.13 | 0.00 |
| 716.00 | 686.00 | 699.33 | 1.08 | 0.11 |
| 6806.00 | 7641.00 | 7297.67 | 1.17 | 0.00 |
| 4279.00 | 4943.00 | 4631.33 | 1.17 | 0.00 |
| 15129.00 | 16462.00 | 16139.00 | 1.23 | 0.00 |
| 2069.00 | 2613.00 | 2329.33 | -1.12 | 0.03 |
| 4732.00 | 4670.00 | 4680.00 | 1.09 | 0.02 |
| 1195.00 | 1384.00 | 1339.67 | 1.03 | 0.42 |
| 36.00 | 24.00 | 34.00 | -1.08 | 0.61 |
| 347.00 | 390.00 | 341.33 | 1.24 | 0.01 |
| 51.00 | 57.00 | 46.67 | -1.01 | 1.00 |
| 1198.00 | 1218.00 | 1254.67 | 1.10 | 0.02 |
| 216.00 | 240.00 | 237.00 | -1.23 | 0.00 |
| 664.00 | 644.00 | 661.33 | -1.19 | 0.00 |
| 723.00 | 843.00 | 788.00 | -1.00 | 0.94 |
| 1127.00 | 1231.00 | 1272.33 | -1.01 | 0.90 |
| 19.00 | 22.00 | 19.67 | -1.35 | 0.12 |

| | | | | |
|---|---|---|---|---|
| 680.00 | 703.00 | 732.67 | -1.15 | 0.00 |
| 129.00 | 127.00 | 146.00 | 1.15 | 0.13 |
| 291.00 | 302.00 | 299.67 | 1.39 | 0.00 |
| 92.00 | 88.00 | 93.00 | 1.49 | 0.00 |
| 153.00 | 139.00 | 152.33 | 1.06 | 0.52 |
| 6.00 | 3.00 | 6.67 | -1.36 | 0.44 |
| 25.00 | 27.00 | 24.00 | -1.18 | 0.38 |
| 143.00 | 164.00 | 159.33 | 1.08 | 0.34 |
| 138.00 | 160.00 | 156.67 | 1.14 | 0.14 |
| 70.00 | 91.00 | 83.00 | -1.04 | 0.72 |
| 17.00 | 11.00 | 15.67 | -1.21 | 0.45 |
| 133.00 | 173.00 | 153.33 | 1.20 | 0.09 |
| 4.00 | 4.00 | 6.67 | 2.14 | 0.08 |
| 48.00 | 33.00 | 45.67 | -1.17 | 0.28 |
| 1273.00 | 1504.00 | 1476.00 | 1.37 | 0.00 |
| 965.00 | 1068.00 | 1067.67 | -1.07 | 0.11 |
| 475.00 | 548.00 | 533.33 | 1.09 | 0.14 |
| 357.00 | 355.00 | 370.00 | -1.02 | 0.69 |
| 776.00 | 739.00 | 766.67 | -1.09 | 0.05 |
| 496.00 | 543.00 | 550.67 | 1.16 | 0.00 |
| 209.00 | 216.00 | 223.67 | 1.00 | 0.98 |
| 2361.00 | 2309.00 | 2395.33 | 1.01 | 0.84 |
| 558.00 | 568.00 | 584.00 | -1.05 | 0.39 |
| 24.00 | 29.00 | 30.00 | 1.25 | 0.23 |
| 38.00 | 51.00 | 48.33 | 1.25 | 0.11 |
| 44.00 | 47.00 | 48.67 | 1.13 | 0.36 |
| 243.00 | 270.00 | 259.67 | 1.06 | 0.35 |
| 63.00 | 88.00 | 72.33 | 1.09 | 0.48 |
| 337.00 | 385.00 | 373.33 | -1.14 | 0.01 |
| 358.00 | 421.00 | 399.00 | -1.19 | 0.00 |
| 154.00 | 180.00 | 179.67 | 1.15 | 0.07 |
| 31.00 | 27.00 | 31.00 | -1.07 | 0.75 |
| 6.00 | 6.00 | 7.67 | -1.40 | 0.29 |
| 96.00 | 94.00 | 94.33 | 1.28 | 0.01 |
| 241.00 | 259.00 | 250.00 | -1.05 | 0.46 |
| 192.00 | 199.00 | 198.67 | 1.04 | 0.54 |
| 272.00 | 286.00 | 296.00 | 1.01 | 0.86 |
| 264.00 | 229.00 | 265.33 | 1.04 | 0.66 |
| 127.00 | 97.00 | 120.33 | 1.25 | 0.02 |
| 4.00 | 12.00 | 7.33 | 1.03 | 1.00 |
| 5779.00 | 6590.00 | 6234.33 | -1.14 | 0.00 |
| 1346.00 | 1257.00 | 1360.33 | -1.11 | 0.01 |
| 447.00 | 458.00 | 471.67 | 1.46 | 0.00 |
| 639.00 | 773.00 | 713.67 | 1.39 | 0.00 |
| 1105.00 | 1078.00 | 1138.67 | -1.02 | 0.72 |
| 1318.00 | 1530.00 | 1423.33 | -1.03 | 0.49 |
| 1178.00 | 1189.00 | 1227.67 | -1.21 | 0.00 |
| 721.00 | 730.00 | 745.00 | 1.12 | 0.01 |

| | | | | |
|---|---|---|---|---|
| 574.00 | 577.00 | 595.33 | -1.02 | 0.63 |
| 209.00 | 231.00 | 249.33 | -1.09 | 0.26 |
| 8.00 | 10.00 | 10.67 | 1.31 | 0.36 |
| 118.00 | 141.00 | 127.33 | 1.31 | 0.00 |
| 1.00 | 0.00 | 0.33 | -2.48 | 0.64 |
| 3.00 | 2.00 | 4.33 | 1.41 | 0.53 |
| 111.00 | 134.00 | 125.33 | -2.02 | 0.00 |
| 530.00 | 529.00 | 541.33 | -1.41 | 0.00 |
| 377.00 | 496.00 | 466.67 | 1.26 | 0.00 |
| 529.00 | 481.00 | 513.33 | -1.01 | 0.88 |
| 0.00 | 1.00 | 1.33 | -2.17 | 0.28 |
| 1646.00 | 1843.00 | 1725.33 | 1.14 | 0.00 |
| 39.00 | 36.00 | 33.67 | 1.06 | 0.85 |
| 15.00 | 3.00 | 8.00 | 1.48 | 0.35 |
| 19.00 | 21.00 | 21.33 | -1.09 | 0.68 |
| 308.00 | 330.00 | 325.00 | 1.12 | 0.06 |
| 12.00 | 3.00 | 6.00 | 1.20 | 0.78 |
| 42.00 | 43.00 | 37.33 | -1.54 | 0.00 |
| 2164.00 | 2370.00 | 2412.33 | 1.25 | 0.00 |
| 34.00 | 20.00 | 29.33 | -1.52 | 0.01 |
| 164.00 | 250.00 | 228.33 | 1.13 | 0.23 |
| 6.00 | 8.00 | 7.00 | -2.20 | 0.00 |
| 197.00 | 177.00 | 204.00 | 1.05 | 0.57 |
| 30.00 | 25.00 | 30.33 | -1.18 | 0.32 |
| 706.00 | 809.00 | 807.00 | 1.18 | 0.00 |
| 1061.00 | 1146.00 | 1158.33 | 1.17 | 0.00 |
| 2151.00 | 2311.00 | 2281.33 | 1.14 | 0.00 |
| 9.00 | 3.00 | 5.00 | 1.14 | 0.85 |
| 60.00 | 43.00 | 49.67 | -1.28 | 0.05 |
| 1.00 | 11.00 | 6.33 | 1.84 | 0.16 |
| 1668.00 | 1846.00 | 1768.67 | 1.07 | 0.13 |
| 22.00 | 20.00 | 20.67 | -1.11 | 0.67 |
| 85.00 | 79.00 | 85.33 | 1.09 | 0.43 |
| 47.00 | 55.00 | 47.33 | -1.32 | 0.03 |
| 105.00 | 127.00 | 127.00 | 1.01 | 0.91 |
| 1.00 | 4.00 | 4.00 | -1.42 | 0.47 |
| 932.00 | 968.00 | 991.33 | -1.09 | 0.08 |
| 518.00 | 600.00 | 567.00 | 1.10 | 0.06 |
| 2429.00 | 2348.00 | 2485.33 | -1.10 | 0.01 |
| 631.00 | 740.00 | 705.00 | 1.02 | 0.65 |
| 47.00 | 36.00 | 39.67 | -1.09 | 0.65 |
| 1050.00 | 1047.00 | 1081.67 | -1.26 | 0.00 |
| 3.00 | 12.00 | 7.00 | -1.30 | 0.49 |
| 24744.00 | 28105.00 | 26702.00 | 1.25 | 0.00 |
| 50340.00 | 54586.00 | 53394.33 | 1.09 | 0.01 |
| 33538.00 | 38419.00 | 36810.67 | -1.08 | 0.03 |
| 140.00 | 142.00 | 146.33 | -1.21 | 0.01 |
| 39.00 | 43.00 | 40.00 | -1.22 | 0.13 |

| | | | | |
|---|---|---|---|---|
| 3353.00 | 3714.00 | 3507.00 | -1.02 | 0.64 |
| 1370.00 | 1478.00 | 1488.33 | 1.09 | 0.03 |
| 29.00 | 20.00 | 22.00 | -1.29 | 0.20 |
| 5.00 | 7.00 | 7.67 | -1.27 | 0.50 |
| 100.00 | 109.00 | 105.67 | 1.07 | 0.53 |
| 4845.00 | 5420.00 | 5267.33 | 1.23 | 0.00 |
| 232.00 | 305.00 | 278.00 | 1.06 | 0.40 |
| 22752.00 | 25401.00 | 24559.67 | -1.25 | 0.00 |
| 27127.00 | 29243.00 | 28769.00 | 1.02 | 0.59 |
| 16733.00 | 18226.00 | 17934.00 | 1.12 | 0.00 |
| 2657.00 | 2780.00 | 2912.33 | 1.37 | 0.00 |
| 235.00 | 301.00 | 288.67 | 1.19 | 0.01 |
| 444.00 | 529.00 | 528.33 | 1.13 | 0.06 |
| 880.00 | 801.00 | 884.33 | 1.02 | 0.64 |
| 15018.00 | 16594.00 | 15952.67 | 1.11 | 0.00 |
| 1078.00 | 1260.00 | 1294.67 | 1.16 | 0.01 |
| 1833.00 | 1471.00 | 1617.33 | -1.42 | 0.00 |
| 3516.00 | 3964.00 | 3806.33 | 1.16 | 0.00 |
| 971.00 | 1024.00 | 1010.67 | 1.07 | 0.12 |
| 1327.00 | 1520.00 | 1455.33 | 1.11 | 0.01 |
| 312.00 | 316.00 | 302.33 | 1.05 | 0.47 |
| 4.00 | 2.00 | 2.67 | -1.26 | 0.82 |
| 1.00 | 1.00 | 0.67 | -4.00 | 0.07 |
| 563.00 | 548.00 | 576.33 | -1.08 | 0.09 |
| 240.00 | 307.00 | 270.33 | 1.06 | 0.45 |
| 310.00 | 313.00 | 320.67 | 1.13 | 0.04 |
| 578.00 | 681.00 | 675.33 | -1.02 | 0.79 |
| 3.00 | 7.00 | 4.67 | 1.92 | 0.20 |
| 3391.00 | 3222.00 | 3254.33 | -1.26 | 0.00 |
| 0.00 | 5.00 | 3.00 | -2.09 | 0.09 |
| 358.00 | 334.00 | 362.33 | -1.09 | 0.15 |
| 8022.00 | 7202.00 | 7450.67 | -1.26 | 0.00 |
| 4.00 | 4.00 | 4.00 | -1.10 | 1.00 |
| 12.00 | 8.00 | 8.67 | -1.44 | 0.22 |
| 1162.00 | 1146.00 | 1173.67 | -1.03 | 0.60 |
| 212.00 | 183.00 | 190.33 | -1.23 | 0.01 |
| 1731.00 | 1673.00 | 1757.33 | -1.24 | 0.00 |
| 61.00 | 81.00 | 71.33 | 1.08 | 0.54 |
| 196.00 | 204.00 | 220.67 | 1.16 | 0.05 |
| 6418.00 | 6642.00 | 6518.33 | -1.20 | 0.00 |
| 457.00 | 509.00 | 522.67 | 1.05 | 0.44 |
| 8893.00 | 9421.00 | 9306.00 | -1.05 | 0.16 |
| 3899.00 | 4018.00 | 3952.67 | 1.15 | 0.00 |
| 185.00 | 184.00 | 189.00 | -1.19 | 0.03 |
| 8.00 | 7.00 | 5.67 | 1.83 | 0.18 |
| 1916.00 | 2003.00 | 1905.67 | -1.08 | 0.13 |
| 533.00 | 555.00 | 558.00 | 1.14 | 0.01 |
| 154.00 | 142.00 | 146.00 | 1.21 | 0.02 |

| | | | | |
|---|---|---|---|---|
| 5240.00 | 5464.00 | 5490.67 | 1.06 | 0.13 |
| 166.00 | 164.00 | 176.00 | 1.24 | 0.01 |
| 2.00 | 3.00 | 5.00 | -1.15 | 0.86 |
| 1406.00 | 1445.00 | 1446.00 | -1.13 | 0.00 |
| 2244.00 | 2410.00 | 2316.67 | -1.09 | 0.03 |
| 1272.00 | 1394.00 | 1358.00 | -1.06 | 0.12 |
| 289.00 | 265.00 | 279.33 | -1.12 | 0.10 |
| 1874.00 | 1971.00 | 1929.33 | 1.02 | 0.57 |
| 409.00 | 442.00 | 460.67 | 1.11 | 0.08 |
| 268.00 | 288.00 | 300.00 | 1.16 | 0.06 |
| 10.00 | 5.00 | 7.00 | 1.09 | 0.88 |
| 378.00 | 431.00 | 442.33 | -1.06 | 0.31 |
| 1225.00 | 1361.00 | 1289.00 | -1.02 | 0.67 |
| 1574.00 | 1771.00 | 1720.33 | 1.08 | 0.04 |
| 8.00 | 7.00 | 7.67 | 1.60 | 0.20 |
| 430.00 | 444.00 | 452.67 | 1.34 | 0.00 |
| 962.00 | 1102.00 | 1037.00 | 1.07 | 0.12 |
| 1392.00 | 1481.00 | 1545.33 | -1.06 | 0.18 |
| 625.00 | 695.00 | 694.33 | 1.00 | 1.00 |
| 477.00 | 535.00 | 519.00 | 1.10 | 0.08 |
| 37.00 | 52.00 | 48.00 | 1.02 | 0.91 |
| 59.00 | 94.00 | 73.00 | 1.13 | 0.37 |
| 1868.00 | 1848.00 | 1800.00 | -1.41 | 0.00 |
| 301.00 | 299.00 | 319.00 | -1.32 | 0.00 |
| 2496.00 | 2554.00 | 2691.33 | 1.13 | 0.00 |
| 881.00 | 997.00 | 983.33 | 1.32 | 0.00 |
| 1073.00 | 1099.00 | 1111.00 | -1.09 | 0.04 |
| 61.00 | 48.00 | 58.00 | -1.17 | 0.20 |
| 1012.00 | 1123.00 | 1109.33 | -1.14 | 0.00 |
| 560.00 | 576.00 | 572.33 | 1.02 | 0.71 |
| 5144.00 | 5726.00 | 5599.00 | -1.17 | 0.00 |
| 1.00 | 18.00 | 12.00 | 1.01 | 1.00 |
| 20.00 | 17.00 | 20.00 | -2.10 | 0.00 |
| 156.00 | 131.00 | 154.67 | 1.24 | 0.02 |
| 490.00 | 595.00 | 546.67 | -1.08 | 0.20 |
| 35.00 | 51.00 | 45.33 | -2.23 | 0.00 |
| 116.00 | 109.00 | 125.00 | 1.29 | 0.01 |
| 98.00 | 105.00 | 107.00 | 1.21 | 0.04 |
| 7.00 | 29.00 | 15.00 | 1.20 | 0.61 |
| 64.00 | 56.00 | 59.33 | -1.02 | 0.88 |
| 171.00 | 179.00 | 182.00 | 1.03 | 0.68 |
| 660.00 | 704.00 | 724.00 | 1.03 | 0.62 |
| 77.00 | 62.00 | 68.67 | 1.11 | 0.37 |
| 731.00 | 826.00 | 849.00 | 1.12 | 0.04 |
| 910.00 | 846.00 | 942.33 | 1.10 | 0.07 |
| 1128.00 | 1133.00 | 1108.33 | 1.15 | 0.02 |
| 346.00 | 376.00 | 386.00 | 1.19 | 0.00 |
| 269.00 | 346.00 | 309.67 | -1.39 | 0.00 |

| | | | | |
|---|---|---|---|---|
| 1021.00 | 1103.00 | 1115.67 | -1.07 | 0.12 |
| 62.00 | 38.00 | 46.67 | 1.04 | 0.82 |
| 2240.00 | 2464.00 | 2417.00 | 1.13 | 0.00 |
| 2438.00 | 2596.00 | 2610.33 | 1.13 | 0.00 |
| 384.00 | 387.00 | 398.33 | 1.04 | 0.50 |
| 843.00 | 902.00 | 885.67 | -1.07 | 0.12 |
| 1170.00 | 1199.00 | 1211.67 | -1.17 | 0.00 |
| 377.00 | 366.00 | 382.33 | -1.15 | 0.01 |
| 166.00 | 167.00 | 182.33 | 1.12 | 0.16 |
| 359.00 | 345.00 | 368.67 | -1.02 | 0.73 |
| 294.00 | 287.00 | 290.33 | -1.00 | 0.96 |
| 510.00 | 525.00 | 531.33 | 1.05 | 0.32 |
| 350.00 | 381.00 | 384.00 | 1.06 | 0.29 |
| 770.00 | 893.00 | 882.67 | 1.15 | 0.01 |
| 411.00 | 520.00 | 492.00 | 1.11 | 0.07 |
| 1063.00 | 1350.00 | 1245.67 | -1.19 | 0.00 |
| 534.00 | 689.00 | 628.67 | 1.03 | 0.65 |
| 348.00 | 446.00 | 407.67 | 1.15 | 0.03 |
| 1237.00 | 1425.00 | 1418.67 | -1.01 | 0.79 |
| 6.00 | 1.00 | 3.33 | -1.40 | 0.55 |
| 2037.00 | 2096.00 | 2075.33 | -1.15 | 0.00 |
| 2333.00 | 2317.00 | 2335.67 | -1.09 | 0.03 |
| 5886.00 | 5991.00 | 6086.67 | -1.02 | 0.49 |
| 65.00 | 88.00 | 76.67 | 1.25 | 0.07 |
| 3028.00 | 3437.00 | 3216.67 | 1.09 | 0.05 |
| 9536.00 | 9209.00 | 9087.00 | -1.34 | 0.00 |
| 14.00 | 11.00 | 13.67 | 1.74 | 0.04 |
| 682.00 | 676.00 | 714.33 | 5.80 | 0.00 |
| 16.00 | 44.00 | 32.67 | -1.26 | 0.17 |
| 469.00 | 536.00 | 527.00 | 1.16 | 0.03 |
| 310.00 | 327.00 | 304.33 | -1.55 | 0.00 |
| 27.00 | 32.00 | 29.00 | 1.13 | 0.55 |
| 112.00 | 66.00 | 100.67 | -2.42 | 0.00 |
| 571.00 | 589.00 | 593.00 | 1.03 | 0.54 |
| 160.00 | 208.00 | 180.67 | -1.13 | 0.18 |
| 4.00 | 4.00 | 4.33 | -1.09 | 1.00 |
| 737.00 | 751.00 | 742.00 | -1.44 | 0.00 |
| 0.00 | 2.00 | 1.00 | -5.51 | 0.00 |
| 96.00 | 86.00 | 89.33 | -1.11 | 0.31 |
| 1.00 | 5.00 | 3.00 | -1.99 | 0.13 |
| 1024.00 | 1138.00 | 1116.67 | -1.24 | 0.00 |
| 2103.00 | 2484.00 | 2384.67 | 1.03 | 0.51 |
| 1230.00 | 1518.00 | 1421.33 | 1.16 | 0.01 |
| 1914.00 | 2109.00 | 1971.00 | 1.02 | 0.70 |
| 2716.00 | 2685.00 | 2774.00 | -1.18 | 0.00 |
| 140.00 | 136.00 | 148.67 | -1.46 | 0.00 |
| 306.00 | 331.00 | 333.00 | -1.01 | 0.93 |
| 59.00 | 62.00 | 55.00 | -1.39 | 0.00 |

| | | | | |
|---|---|---|---|---|
| 207.00 | 214.00 | 223.00 | -1.32 | 0.00 |
| 596.00 | 747.00 | 653.00 | 1.06 | 0.35 |
| 538.00 | 510.00 | 530.00 | 1.22 | 0.00 |
| 106.00 | 107.00 | 103.67 | -6.39 | 0.00 |
| 65.00 | 63.00 | 68.33 | -1.63 | 0.00 |
| 681.00 | 747.00 | 743.67 | 1.87 | 0.00 |
| 5.00 | 11.00 | 8.67 | -1.28 | 0.45 |
| 3.00 | 4.00 | 2.33 | -1.84 | 0.27 |
| 214.00 | 225.00 | 210.00 | -1.10 | 0.24 |
| 1617.00 | 1473.00 | 1542.33 | -1.05 | 0.34 |
| 125.00 | 157.00 | 143.33 | -1.27 | 0.01 |
| 8.00 | 10.00 | 10.33 | 1.02 | 1.00 |
| 17.00 | 10.00 | 12.00 | -1.90 | 0.00 |
| 7.00 | 12.00 | 12.67 | -1.28 | 0.30 |
| 64.00 | 59.00 | 68.67 | -1.10 | 0.41 |
| 632.00 | 667.00 | 658.33 | -1.04 | 0.44 |
| 548.00 | 453.00 | 513.33 | 1.08 | 0.20 |
| 486.00 | 422.00 | 456.67 | -1.04 | 0.49 |
| 8.00 | 13.00 | 10.67 | -1.36 | 0.26 |
| 575.00 | 511.00 | 526.67 | -1.69 | 0.00 |
| 31.00 | 44.00 | 39.00 | -1.03 | 0.90 |
| 101.00 | 123.00 | 124.00 | 1.21 | 0.04 |
| 9994.00 | 11007.00 | 11012.00 | 1.70 | 0.00 |
| 48.00 | 36.00 | 44.00 | -1.09 | 0.54 |
| 985.00 | 1106.00 | 1034.67 | -1.05 | 0.29 |
| 78.00 | 65.00 | 73.33 | -1.56 | 0.00 |
| 12.00 | 6.00 | 14.67 | -2.73 | 0.00 |
| 84.00 | 92.00 | 93.00 | -1.28 | 0.01 |
| 73.00 | 64.00 | 75.33 | 1.70 | 0.00 |
| 7.00 | 10.00 | 10.67 | 1.09 | 0.80 |
| 47.00 | 52.00 | 52.67 | -1.78 | 0.00 |
| 703.00 | 749.00 | 726.33 | 1.19 | 0.00 |
| 5.00 | 0.00 | 3.00 | -1.12 | 1.00 |
| 3.00 | 2.00 | 3.33 | 1.60 | 0.48 |
| 837.00 | 761.00 | 784.00 | -1.40 | 0.00 |
| 30.00 | 48.00 | 48.00 | 2.11 | 0.00 |
| 13.00 | 14.00 | 14.67 | 1.03 | 0.92 |
| 26.00 | 21.00 | 26.67 | -1.16 | 0.41 |
| 763.00 | 785.00 | 800.00 | -1.21 | 0.00 |
| 63.00 | 88.00 | 73.33 | 2.08 | 0.00 |
| 11.00 | 9.00 | 9.67 | -3.39 | 0.00 |
| 5500.00 | 5526.00 | 5386.00 | -1.01 | 0.84 |
| 44.00 | 78.00 | 58.00 | 1.56 | 0.00 |
| 126.00 | 144.00 | 132.00 | -3.73 | 0.00 |
| 4449.00 | 4881.00 | 4814.00 | 1.20 | 0.00 |
| 11927.00 | 12129.00 | 12206.67 | 1.00 | 0.95 |
| 363.00 | 361.00 | 357.00 | 1.01 | 0.89 |
| 3144.00 | 3294.00 | 3389.33 | 1.18 | 0.00 |

| | | | | |
|---|---|---|---|---|
| 715.00 | 771.00 | 777.67 | 1.05 | 0.28 |
| 1251.00 | 1220.00 | 1323.33 | -1.05 | 0.30 |
| 77.00 | 71.00 | 78.67 | 1.18 | 0.13 |
| 291.00 | 387.00 | 349.67 | 1.06 | 0.42 |
| 2344.00 | 2572.00 | 2549.67 | 1.14 | 0.00 |
| 1247.00 | 1326.00 | 1340.00 | 1.19 | 0.00 |
| 1276.00 | 1296.00 | 1334.00 | 1.05 | 0.26 |
| 759.00 | 932.00 | 858.67 | 1.12 | 0.02 |
| 90.00 | 71.00 | 86.33 | 1.28 | 0.02 |
| 1041.00 | 1206.00 | 1137.33 | 1.26 | 0.00 |
| 3460.00 | 3812.00 | 3597.33 | 1.01 | 0.88 |
| 1896.00 | 1880.00 | 1984.33 | 1.08 | 0.06 |
| 8704.00 | 9308.00 | 9325.67 | 1.00 | 0.99 |
| 39.00 | 32.00 | 33.00 | -1.27 | 0.13 |
| 45.00 | 54.00 | 44.67 | -1.54 | 0.00 |
| 62.00 | 83.00 | 71.33 | 1.03 | 0.79 |
| 598.00 | 610.00 | 631.33 | -1.01 | 0.88 |
| 12.00 | 23.00 | 16.00 | -1.01 | 1.00 |
| 958.00 | 971.00 | 969.67 | 1.07 | 0.15 |
| 103.00 | 102.00 | 98.67 | -1.32 | 0.00 |
| 2339.00 | 2288.00 | 2357.33 | -1.16 | 0.00 |
| 217.00 | 236.00 | 232.67 | 1.09 | 0.18 |
| 94.00 | 107.00 | 105.67 | 1.22 | 0.04 |
| 296.00 | 392.00 | 370.67 | 1.13 | 0.08 |
| 490.00 | 542.00 | 539.67 | -1.15 | 0.00 |
| 1003.00 | 953.00 | 1030.33 | -1.04 | 0.45 |
| 40.00 | 30.00 | 33.33 | 1.06 | 0.69 |
| 2892.00 | 2682.00 | 2701.00 | 1.03 | 0.53 |
| 14.00 | 7.00 | 10.67 | -1.08 | 0.91 |
| 5.00 | 2.00 | 3.67 | -1.55 | 0.36 |
| 262.00 | 250.00 | 258.67 | -1.87 | 0.00 |
| 190.00 | 184.00 | 178.00 | 1.03 | 0.77 |
| 1372.00 | 1309.00 | 1351.67 | -1.10 | 0.03 |
| 38.00 | 36.00 | 36.67 | -1.82 | 0.00 |
| 182.00 | 217.00 | 203.33 | 1.57 | 0.00 |
| 561.00 | 677.00 | 638.00 | -1.04 | 0.48 |
| 1396.00 | 1313.00 | 1397.00 | 1.12 | 0.01 |
| 633.00 | 792.00 | 743.00 | -1.07 | 0.21 |
| 1052.00 | 1031.00 | 1072.33 | -1.05 | 0.20 |
| 251.00 | 247.00 | 241.33 | -1.78 | 0.00 |
| 442.00 | 429.00 | 459.00 | -1.04 | 0.42 |
| 747.00 | 762.00 | 753.33 | 1.09 | 0.07 |
| 1014.00 | 1043.00 | 1054.00 | -1.15 | 0.00 |
| 699.00 | 665.00 | 679.67 | -1.15 | 0.00 |
| 20.00 | 11.00 | 18.33 | -1.60 | 0.02 |
| 420.00 | 477.00 | 455.00 | -1.08 | 0.13 |
| 5113.00 | 5073.00 | 5225.33 | -1.17 | 0.00 |
| 77.00 | 91.00 | 78.00 | 3.07 | 0.00 |

| | | | | |
|---|---|---|---|---|
| 1164.00 | 1227.00 | 1208.67 | 1.37 | 0.00 |
| 356.00 | 359.00 | 359.00 | -1.16 | 0.01 |
| 3.00 | 9.00 | 6.67 | 1.77 | 0.16 |
| 3675.00 | 3610.00 | 3651.00 | -1.12 | 0.00 |
| 827.00 | 892.00 | 892.33 | 1.07 | 0.13 |
| 398.00 | 438.00 | 436.67 | 1.01 | 0.90 |
| 1159.00 | 1243.00 | 1186.00 | 1.02 | 0.68 |
| 7.00 | 8.00 | 9.33 | 1.37 | 0.32 |
| 429.00 | 450.00 | 452.67 | -1.09 | 0.11 |
| 7.00 | 7.00 | 7.00 | -1.49 | 0.22 |
| 511.00 | 526.00 | 531.00 | -1.18 | 0.00 |
| 1086.00 | 1156.00 | 1128.67 | 1.08 | 0.05 |
| 31.00 | 67.00 | 50.00 | 1.04 | 0.84 |
| 783.00 | 875.00 | 849.33 | 1.10 | 0.05 |
| 1465.00 | 1679.00 | 1628.00 | -1.04 | 0.31 |
| 407.00 | 413.00 | 434.00 | 1.03 | 0.59 |
| 154.00 | 191.00 | 188.67 | 1.19 | 0.03 |
| 402.00 | 427.00 | 433.67 | -1.06 | 0.27 |
| 861.00 | 884.00 | 913.00 | -1.06 | 0.16 |
| 1786.00 | 1985.00 | 1989.33 | -1.00 | 0.93 |
| 1.00 | 1.00 | 2.00 | -3.39 | 0.01 |
| 252.00 | 320.00 | 273.33 | -1.03 | 0.74 |
| 250.00 | 279.00 | 260.33 | -1.00 | 1.00 |
| 1106.00 | 1228.00 | 1164.67 | -1.10 | 0.03 |
| 2152.00 | 2215.00 | 2275.00 | 1.05 | 0.28 |
| 6070.00 | 6478.00 | 6408.67 | 1.10 | 0.01 |
| 5.00 | 9.00 | 8.67 | -1.32 | 0.38 |
| 8596.00 | 9387.00 | 9020.67 | 1.04 | 0.37 |
| 1891.00 | 2001.00 | 1933.67 | 1.00 | 0.92 |
| 2428.00 | 2448.00 | 2413.00 | -1.13 | 0.00 |
| 8.00 | 10.00 | 9.00 | 2.37 | 0.02 |
| 286.00 | 338.00 | 311.33 | 1.05 | 0.44 |
| 453.00 | 489.00 | 513.33 | 1.08 | 0.21 |
| 707.00 | 890.00 | 816.33 | -1.09 | 0.13 |
| 1208.00 | 1379.00 | 1359.33 | 1.06 | 0.20 |
| 148.00 | 178.00 | 176.33 | 1.12 | 0.16 |
| 9.00 | 27.00 | 17.67 | -1.36 | 0.12 |
| 419.00 | 357.00 | 388.00 | -1.11 | 0.15 |
| 559.00 | 568.00 | 585.33 | 1.30 | 0.00 |
| 127.00 | 129.00 | 124.67 | 1.02 | 0.87 |
| 198.00 | 243.00 | 210.67 | -1.21 | 0.01 |
| 23.00 | 34.00 | 27.00 | 1.42 | 0.06 |
| 75.00 | 75.00 | 77.00 | 1.13 | 0.24 |
| 6683.00 | 6876.00 | 6647.00 | -1.17 | 0.00 |
| 8.00 | 8.00 | 7.33 | -2.19 | 0.00 |
| 2621.00 | 2697.00 | 2719.33 | -1.16 | 0.00 |
| 651.00 | 606.00 | 610.33 | -1.19 | 0.00 |
| 70.00 | 74.00 | 71.00 | -1.01 | 0.93 |

| | | | | |
|---|---|---|---|---|
| 7.00 | 4.00 | 7.67 | -1.19 | 0.68 |
| 16.00 | 10.00 | 14.33 | 1.15 | 0.59 |
| 6034.00 | 5734.00 | 6185.33 | 1.08 | 0.05 |
| 189.00 | 204.00 | 191.33 | 1.07 | 0.38 |
| 227.00 | 265.00 | 240.67 | -1.08 | 0.27 |
| 12.00 | 6.00 | 8.67 | -1.13 | 0.79 |
| 762.00 | 827.00 | 815.33 | 1.28 | 0.00 |
| 970.00 | 1148.00 | 1042.67 | -1.25 | 0.00 |
| 237.00 | 237.00 | 256.00 | -1.12 | 0.09 |
| 303.00 | 274.00 | 308.33 | -1.03 | 0.72 |
| 699.00 | 699.00 | 739.67 | 1.13 | 0.02 |
| 2682.00 | 2799.00 | 2802.67 | 1.03 | 0.51 |
| 1054.00 | 1049.00 | 1051.33 | -1.25 | 0.00 |
| 1539.00 | 1619.00 | 1630.00 | -1.01 | 0.85 |
| 7.00 | 5.00 | 5.33 | 1.12 | 0.86 |
| 256.00 | 288.00 | 294.67 | -1.11 | 0.10 |
| 5.00 | 8.00 | 7.00 | 1.15 | 0.75 |
| 675.00 | 693.00 | 694.67 | -1.35 | 0.00 |
| 4192.00 | 4199.00 | 4181.00 | 1.16 | 0.00 |
| 728.00 | 693.00 | 697.67 | 1.19 | 0.00 |
| 132.00 | 118.00 | 126.33 | -1.20 | 0.03 |
| 30.00 | 47.00 | 36.33 | -1.06 | 0.71 |
| 327.00 | 404.00 | 384.00 | 1.14 | 0.05 |
| 165.00 | 216.00 | 196.00 | 1.06 | 0.48 |
| 752.00 | 847.00 | 846.67 | -1.07 | 0.15 |
| 44.00 | 38.00 | 43.33 | -1.43 | 0.00 |
| 461.00 | 499.00 | 525.00 | 1.10 | 0.14 |
| 733.00 | 738.00 | 756.33 | -1.22 | 0.00 |
| 30.00 | 46.00 | 40.00 | -1.09 | 0.56 |
| 82.00 | 104.00 | 98.00 | 1.06 | 0.61 |
| 124.00 | 134.00 | 131.33 | 1.11 | 0.20 |
| 473.00 | 482.00 | 499.00 | 1.38 | 0.00 |
| 404.00 | 407.00 | 452.67 | 1.03 | 0.71 |
| 59.00 | 78.00 | 64.67 | 1.20 | 0.14 |
| 2333.00 | 2379.00 | 2392.33 | 1.01 | 0.71 |
| 348.00 | 366.00 | 373.33 | 1.03 | 0.65 |
| 133.00 | 124.00 | 144.00 | -1.43 | 0.00 |
| 1715.00 | 1792.00 | 1803.67 | 1.01 | 0.73 |
| 1602.00 | 1640.00 | 1641.33 | -1.01 | 0.84 |
| 471.00 | 458.00 | 472.33 | -1.02 | 0.73 |
| 1881.00 | 1742.00 | 1902.67 | -1.12 | 0.02 |
| 310.00 | 294.00 | 292.00 | -1.69 | 0.00 |
| 1318.00 | 1395.00 | 1429.00 | 1.53 | 0.00 |
| 6.00 | 2.00 | 4.33 | -5.03 | 0.00 |
| 2477.00 | 2691.00 | 2507.67 | -1.76 | 0.00 |
| 5.00 | 3.00 | 3.33 | -1.31 | 0.68 |
| 21453.00 | 21658.00 | 21837.00 | -2.01 | 0.00 |
| 2254.00 | 2520.00 | 2352.67 | 1.05 | 0.30 |

| | | | | |
|---|---|---|---|---|
| 5732.00 | 5880.00 | 5835.33 | -1.32 | 0.00 |
| 20843.00 | 21591.00 | 20836.67 | 1.20 | 0.00 |
| 30.00 | 55.00 | 48.67 | -1.25 | 0.11 |
| 8.00 | 6.00 | 9.00 | 1.55 | 0.18 |
| 335.00 | 423.00 | 404.67 | 1.06 | 0.39 |
| 0.00 | 0.00 | 0.33 | -3.22 | 0.38 |
| 2155.00 | 2465.00 | 2199.67 | -1.08 | 0.19 |
| 40.00 | 58.00 | 57.33 | -2.02 | 0.00 |
| 31760.00 | 33234.00 | 32453.67 | -1.07 | 0.07 |
| 622.00 | 751.00 | 722.33 | 1.18 | 0.00 |
| 28.00 | 26.00 | 27.67 | -1.18 | 0.32 |
| 2806.00 | 2932.00 | 2936.00 | -1.32 | 0.00 |
| 564.00 | 673.00 | 654.67 | -1.06 | 0.25 |
| 188.00 | 193.00 | 188.33 | -4.46 | 0.00 |
| 356.00 | 418.00 | 413.00 | 1.17 | 0.01 |
| 14.00 | 23.00 | 14.67 | -1.06 | 0.92 |
| 2549.00 | 2517.00 | 2609.33 | -1.97 | 0.00 |
| 3.00 | 1.00 | 2.00 | -2.28 | 0.12 |
| 371.00 | 486.00 | 408.67 | 2.40 | 0.00 |
| 4407.00 | 4700.00 | 4687.33 | -1.14 | 0.00 |
| 12.00 | 19.00 | 17.67 | -1.68 | 0.00 |
| 2577.00 | 2614.00 | 2611.00 | 1.06 | 0.18 |
| 3189.00 | 3089.00 | 3240.67 | -1.17 | 0.00 |
| 1490.00 | 1620.00 | 1657.33 | 1.20 | 0.00 |
| 137.00 | 157.00 | 157.00 | -1.13 | 0.11 |
| 463.00 | 432.00 | 447.67 | 1.26 | 0.00 |
| 3.00 | 0.00 | 1.33 | -1.01 | 1.00 |
| 264.00 | 255.00 | 276.33 | -1.12 | 0.08 |
| 510.00 | 483.00 | 486.00 | 1.13 | 0.05 |
| 15.00 | 13.00 | 14.67 | 1.01 | 1.00 |
| 1310.00 | 1293.00 | 1299.00 | -1.15 | 0.00 |
| 2322.00 | 1914.00 | 2148.33 | -1.57 | 0.00 |
| 4595.00 | 4493.00 | 4613.33 | -1.24 | 0.00 |
| 17.00 | 25.00 | 18.33 | 1.23 | 0.34 |
| 3497.00 | 3373.00 | 3404.33 | -1.19 | 0.00 |
| 1164.00 | 1198.00 | 1174.00 | 1.28 | 0.00 |
| 1070.00 | 1004.00 | 1014.67 | -1.11 | 0.05 |
| 1624.00 | 1607.00 | 1653.00 | -1.06 | 0.18 |
| 2634.00 | 3047.00 | 2894.00 | 1.04 | 0.31 |
| 1360.00 | 1536.00 | 1483.00 | 1.11 | 0.01 |
| 26.00 | 36.00 | 36.67 | -1.48 | 0.01 |
| 714.00 | 859.00 | 786.33 | 1.16 | 0.01 |
| 865.00 | 842.00 | 827.00 | -1.03 | 0.67 |
| 513.00 | 486.00 | 486.67 | 1.03 | 0.59 |
| 1041.00 | 1175.00 | 1091.67 | -2.39 | 0.00 |
| 568.00 | 551.00 | 572.67 | -1.08 | 0.10 |
| 3538.00 | 3839.00 | 3745.67 | -1.09 | 0.02 |
| 415.00 | 573.00 | 477.67 | -1.10 | 0.21 |

| | | | | |
|---|---|---|---|---|
| 4.00 | 6.00 | 4.33 | -1.32 | 0.59 |
| 193.00 | 171.00 | 184.67 | -1.05 | 0.49 |
| 1169.00 | 1149.00 | 1154.00 | 1.01 | 0.83 |
| 636.00 | 643.00 | 632.33 | -1.17 | 0.00 |
| 169.00 | 213.00 | 186.33 | 1.11 | 0.17 |
| 2.00 | 3.00 | 1.67 | -1.76 | 0.44 |
| 0.00 | 0.00 | 1.33 | -1.02 | 1.00 |
| 67.00 | 83.00 | 79.00 | -1.01 | 0.91 |
| 1419.00 | 1443.00 | 1463.67 | 1.07 | 0.10 |
| 3528.00 | 3703.00 | 3631.33 | -1.06 | 0.15 |
| 396.00 | 362.00 | 402.67 | 1.12 | 0.13 |
| 606.00 | 609.00 | 640.00 | -1.06 | 0.19 |
| 82.00 | 93.00 | 91.67 | -1.10 | 0.30 |
| 851.00 | 860.00 | 905.33 | 1.05 | 0.32 |
| 562.00 | 652.00 | 596.67 | -1.29 | 0.00 |
| 1737.00 | 1670.00 | 1756.67 | 1.06 | 0.15 |
| 279.00 | 298.00 | 304.33 | 1.01 | 0.91 |
| 231.00 | 236.00 | 245.67 | 1.11 | 0.12 |
| 165.00 | 155.00 | 179.00 | 1.11 | 0.23 |
| 2136.00 | 2089.00 | 2089.67 | -1.17 | 0.00 |
| 880.00 | 1109.00 | 984.67 | 1.07 | 0.19 |
| 1921.00 | 2006.00 | 2015.00 | 1.19 | 0.00 |
| 9136.00 | 9912.00 | 9421.67 | 1.18 | 0.00 |
| 613.00 | 681.00 | 678.33 | 1.29 | 0.00 |
| 276.00 | 234.00 | 259.33 | -1.27 | 0.00 |
| 7.00 | 12.00 | 8.67 | -1.97 | 0.01 |
| 994.00 | 913.00 | 938.67 | -1.31 | 0.00 |
| 1478.00 | 1498.00 | 1484.67 | -2.47 | 0.00 |
| 295.00 | 260.00 | 284.33 | -1.08 | 0.24 |
| 231.00 | 255.00 | 242.67 | -1.06 | 0.48 |
| 575.00 | 585.00 | 600.33 | 1.07 | 0.17 |
| 1836.00 | 2083.00 | 2085.33 | -1.08 | 0.09 |
| 21.00 | 7.00 | 15.67 | 2.09 | 0.01 |
| 3322.00 | 3593.00 | 3541.00 | 1.13 | 0.00 |
| 3995.00 | 4220.00 | 4276.67 | -1.15 | 0.00 |
| 276.00 | 326.00 | 304.67 | 1.34 | 0.00 |
| 1260.00 | 1277.00 | 1319.33 | -1.02 | 0.64 |
| 987.00 | 914.00 | 911.33 | -1.55 | 0.00 |
| 968.00 | 937.00 | 928.67 | -1.17 | 0.00 |
| 885.00 | 835.00 | 868.00 | -1.10 | 0.04 |
| 644.00 | 660.00 | 668.33 | -1.12 | 0.02 |
| 583.00 | 641.00 | 636.33 | 1.20 | 0.00 |
| 218.00 | 227.00 | 221.33 | -1.07 | 0.33 |
| 131.00 | 127.00 | 129.33 | -1.03 | 0.81 |
| 983.00 | 1061.00 | 1079.67 | 1.02 | 0.65 |
| 524.00 | 658.00 | 604.33 | 1.15 | 0.01 |
| 32.00 | 29.00 | 29.33 | 1.04 | 0.85 |
| 92.00 | 96.00 | 95.67 | 1.05 | 0.67 |

| | | | | |
|---|---|---|---|---|
| 1876.00 | 2140.00 | 2107.67 | 1.07 | 0.11 |
| 25.00 | 45.00 | 39.00 | -1.07 | 0.74 |
| 285.00 | 290.00 | 292.33 | 1.17 | 0.01 |
| 375.00 | 459.00 | 403.00 | 1.09 | 0.18 |
| 107.00 | 72.00 | 92.33 | 1.31 | 0.05 |
| 236.00 | 301.00 | 272.00 | 1.03 | 0.69 |
| 238.00 | 271.00 | 259.33 | -1.27 | 0.00 |
| 1264.00 | 1383.00 | 1395.67 | 1.19 | 0.00 |
| 0.00 | 1.00 | 1.00 | -1.31 | 1.00 |
| 13.00 | 15.00 | 17.00 | 1.43 | 0.14 |
| 2947.00 | 2801.00 | 2955.33 | -1.23 | 0.00 |
| 142.00 | 141.00 | 139.67 | -1.13 | 0.13 |
| 101.00 | 85.00 | 93.67 | -1.13 | 0.21 |
| 82.00 | 86.00 | 90.67 | -1.98 | 0.00 |
| 3.00 | 1.00 | 1.67 | -1.77 | 0.43 |
| 126.00 | 125.00 | 130.00 | 1.00 | 1.00 |
| 15.00 | 20.00 | 23.33 | -1.43 | 0.04 |
| 3831.00 | 3683.00 | 3845.33 | 1.07 | 0.06 |
| 19.00 | 21.00 | 19.67 | 1.66 | 0.02 |
| 122.00 | 93.00 | 98.00 | -1.77 | 0.00 |
| 25.00 | 20.00 | 23.67 | 1.25 | 0.23 |
| 5.00 | 1.00 | 2.67 | -1.25 | 0.82 |
| 5.00 | 4.00 | 5.67 | -1.48 | 0.30 |
| 151.00 | 155.00 | 163.00 | -1.61 | 0.00 |
| 12.00 | 18.00 | 16.67 | -1.10 | 0.64 |
| 490.00 | 499.00 | 524.33 | -1.23 | 0.00 |
| 221.00 | 222.00 | 230.33 | -1.19 | 0.01 |
| 10.00 | 6.00 | 8.00 | -1.43 | 0.25 |
| 326.00 | 328.00 | 335.00 | 1.21 | 0.00 |
| 23.00 | 31.00 | 27.67 | 1.60 | 0.01 |
| 5.00 | 7.00 | 6.33 | 1.24 | 0.61 |
| 2.00 | 8.00 | 4.67 | 1.15 | 0.85 |
| 149.00 | 146.00 | 149.67 | 1.97 | 0.00 |
| 109.00 | 105.00 | 111.67 | -1.07 | 0.43 |
| 394.00 | 418.00 | 424.00 | 2.55 | 0.00 |
| 2.00 | 0.00 | 2.33 | -3.07 | 0.01 |
| 101.00 | 111.00 | 105.00 | -1.61 | 0.00 |
| 1.00 | 5.00 | 2.00 | -1.49 | 0.62 |
| 4054.00 | 4063.00 | 4051.33 | -1.11 | 0.01 |
| 16.00 | 11.00 | 15.33 | -1.23 | 0.39 |
| 153.00 | 179.00 | 169.00 | 1.10 | 0.22 |
| 26.00 | 21.00 | 24.33 | 1.06 | 0.75 |
| 74.00 | 108.00 | 89.00 | 1.03 | 0.77 |
| 418.00 | 444.00 | 462.33 | -1.05 | 0.37 |
| 220.00 | 236.00 | 216.33 | -1.44 | 0.00 |
| 3.00 | 3.00 | 4.33 | -2.83 | 0.00 |
| 1438.00 | 1468.00 | 1434.67 | -1.15 | 0.00 |
| 3.00 | 5.00 | 4.67 | 1.15 | 0.85 |

| | | | | |
|---|---|---|---|---|
| 5.00 | 13.00 | 8.00 | -1.55 | 0.14 |
| 50.00 | 46.00 | 48.00 | -1.27 | 0.05 |
| 211.00 | 257.00 | 224.67 | -1.02 | 0.84 |
| 92.00 | 101.00 | 101.33 | -1.33 | 0.00 |
| 9.00 | 7.00 | 8.33 | -2.45 | 0.00 |
| 438.00 | 426.00 | 422.33 | 1.03 | 0.67 |
| 1231.00 | 1314.00 | 1291.67 | -1.13 | 0.00 |
| 378.00 | 422.00 | 418.00 | -1.13 | 0.02 |
| 1113.00 | 1268.00 | 1171.67 | 1.27 | 0.00 |
| 546.00 | 560.00 | 587.67 | -1.01 | 0.88 |
| 209.00 | 199.00 | 207.33 | -1.13 | 0.09 |
| 9.00 | 10.00 | 10.67 | -1.36 | 0.26 |
| 263.00 | 268.00 | 271.67 | -1.09 | 0.14 |
| 669.00 | 732.00 | 718.67 | 1.02 | 0.66 |
| 11.00 | 10.00 | 12.33 | 1.04 | 0.91 |
| 527.00 | 561.00 | 579.00 | -1.03 | 0.49 |
| 1258.00 | 1284.00 | 1299.00 | 1.01 | 0.89 |
| 142.00 | 135.00 | 147.00 | -1.15 | 0.08 |
| 7.00 | 13.00 | 9.67 | -1.15 | 0.71 |
| 798.00 | 911.00 | 882.33 | -1.02 | 0.67 |
| 209.00 | 259.00 | 231.67 | -1.05 | 0.48 |
| 596.00 | 620.00 | 634.00 | -1.04 | 0.46 |
| 194.00 | 216.00 | 220.00 | 1.06 | 0.39 |
| 528.00 | 494.00 | 503.67 | -1.15 | 0.02 |
| 6.00 | 12.00 | 9.00 | 1.39 | 0.32 |
| 409.00 | 508.00 | 465.00 | 1.04 | 0.49 |
| 347.00 | 355.00 | 376.00 | 1.13 | 0.04 |
| 1613.00 | 1595.00 | 1598.67 | 1.08 | 0.06 |
| 3514.00 | 3729.00 | 3771.33 | -1.07 | 0.13 |
| 5450.00 | 5486.00 | 5520.67 | 1.02 | 0.69 |
| 459.00 | 513.00 | 491.00 | -1.02 | 0.78 |
| 2451.00 | 2546.00 | 2546.33 | 1.11 | 0.00 |
| 959.00 | 975.00 | 961.67 | 1.34 | 0.00 |
| 2652.00 | 2552.00 | 2597.67 | -1.02 | 0.66 |
| 771.00 | 732.00 | 748.67 | 1.04 | 0.39 |
| 1398.00 | 1530.00 | 1464.67 | 1.02 | 0.60 |
| 2210.00 | 2028.00 | 2143.33 | -1.09 | 0.06 |
| 2347.00 | 2401.00 | 2432.33 | 1.06 | 0.11 |
| 40.00 | 49.00 | 53.00 | 1.13 | 0.42 |
| 684.00 | 697.00 | 710.33 | -1.17 | 0.00 |
| 933.00 | 996.00 | 974.00 | -1.02 | 0.72 |
| 61.00 | 68.00 | 63.67 | 1.16 | 0.20 |
| 1347.00 | 1481.00 | 1372.00 | 1.00 | 0.98 |
| 1762.00 | 1784.00 | 1803.00 | -1.03 | 0.45 |
| 5563.00 | 5228.00 | 5478.33 | -1.18 | 0.00 |
| 686.00 | 713.00 | 701.67 | -1.03 | 0.49 |
| 1116.00 | 949.00 | 1041.00 | -1.38 | 0.00 |
| 1102.00 | 1225.00 | 1131.33 | -1.05 | 0.36 |

| | | | | |
|---|---|---|---|---|
| 76.00 | 71.00 | 74.67 | -1.14 | 0.23 |
| 79.00 | 62.00 | 67.33 | 1.05 | 0.72 |
| 2082.00 | 1931.00 | 2020.67 | 1.01 | 0.91 |
| 831.00 | 817.00 | 874.67 | 1.02 | 0.74 |
| 4.00 | 1.00 | 3.33 | -1.40 | 0.55 |
| 4.00 | 5.00 | 10.33 | -1.13 | 0.78 |
| 118.00 | 170.00 | 141.00 | -1.03 | 0.78 |
| 3725.00 | 4466.00 | 4249.67 | 1.20 | 0.00 |
| 137.00 | 153.00 | 153.33 | 1.22 | 0.02 |
| 29.00 | 29.00 | 29.33 | 1.52 | 0.02 |
| 2230.00 | 2117.00 | 2182.00 | -1.22 | 0.00 |
| 4634.00 | 4475.00 | 4584.33 | 1.03 | 0.42 |
| 2586.00 | 2954.00 | 2804.67 | 1.15 | 0.00 |
| 5136.00 | 4751.00 | 4843.00 | -1.22 | 0.00 |
| 292.00 | 400.00 | 373.67 | 1.55 | 0.00 |
| 1952.00 | 1960.00 | 2034.33 | -1.11 | 0.00 |
| 1829.00 | 1835.00 | 1875.00 | -1.30 | 0.00 |
| 1660.00 | 1669.00 | 1699.33 | 1.00 | 0.92 |
| 879.00 | 911.00 | 929.33 | 1.03 | 0.54 |
| 4.00 | 6.00 | 5.00 | -4.18 | 0.00 |
| 860.00 | 833.00 | 842.33 | -1.06 | 0.25 |
| 2319.00 | 2238.00 | 2306.67 | -1.07 | 0.14 |
| 178.00 | 156.00 | 156.67 | -1.28 | 0.01 |
| 3558.00 | 3834.00 | 3711.33 | -1.08 | 0.03 |
| 698.00 | 723.00 | 706.67 | 1.13 | 0.02 |
| 232.00 | 201.00 | 227.00 | -1.07 | 0.36 |
| 715.00 | 697.00 | 700.33 | -1.36 | 0.00 |
| 740.00 | 660.00 | 690.00 | 1.01 | 0.93 |
| 1039.00 | 1255.00 | 1153.00 | -1.06 | 0.24 |
| 2012.00 | 1325.00 | 1407.67 | -1.30 | 0.11 |
| 20.00 | 28.00 | 22.00 | 1.08 | 0.74 |
| 187.00 | 194.00 | 194.33 | 1.03 | 0.67 |
| 584.00 | 651.00 | 650.67 | -1.18 | 0.00 |
| 3968.00 | 4298.00 | 4248.67 | -1.12 | 0.00 |
| 682.00 | 716.00 | 711.67 | -1.05 | 0.29 |
| 537.00 | 505.00 | 533.00 | 1.02 | 0.67 |
| 96.00 | 105.00 | 104.67 | -1.37 | 0.00 |
| 1573.00 | 1769.00 | 1636.33 | -1.09 | 0.06 |
| 199.00 | 231.00 | 228.00 | -1.19 | 0.01 |
| 1766.00 | 1963.00 | 1873.00 | -1.27 | 0.00 |
| 510.00 | 544.00 | 494.33 | 1.18 | 0.05 |
| 12.00 | 22.00 | 18.33 | -1.11 | 0.61 |
| 1474.00 | 1643.00 | 1687.33 | -1.03 | 0.48 |
| 260.00 | 236.00 | 230.67 | -1.05 | 0.59 |
| 423.00 | 391.00 | 405.33 | -1.22 | 0.00 |
| 908.00 | 1042.00 | 1003.67 | 1.10 | 0.02 |
| 2.00 | 4.00 | 2.33 | -2.80 | 0.02 |
| 68.00 | 71.00 | 64.33 | -1.25 | 0.05 |

| | | | | |
|---|---|---|---|---|
| 97.00 | 78.00 | 88.33 | -1.34 | 0.00 |
| 351.00 | 408.00 | 406.67 | 1.05 | 0.44 |
| 2545.00 | 2683.00 | 2608.33 | -1.04 | 0.32 |
| 661.00 | 511.00 | 559.00 | -1.87 | 0.00 |
| 901.00 | 997.00 | 942.67 | 1.02 | 0.63 |
| 580.00 | 684.00 | 636.00 | -1.25 | 0.00 |
| 44.00 | 52.00 | 46.00 | -1.01 | 1.00 |
| 934.00 | 1102.00 | 1029.00 | 1.27 | 0.00 |
| 1174.00 | 1155.00 | 1177.33 | -1.19 | 0.00 |
| 4414.00 | 4285.00 | 4270.67 | -1.11 | 0.02 |
| 758.00 | 809.00 | 807.33 | -1.10 | 0.03 |
| 241.00 | 258.00 | 249.00 | 1.19 | 0.01 |
| 201.00 | 242.00 | 235.67 | 1.51 | 0.00 |
| 1223.00 | 1031.00 | 1075.33 | -1.12 | 0.09 |
| 1.00 | 1.00 | 0.67 | -4.85 | 0.02 |
| 1048.00 | 1073.00 | 1032.33 | 1.13 | 0.02 |
| 133.00 | 145.00 | 148.00 | 1.12 | 0.18 |
| 405.00 | 475.00 | 449.67 | 1.05 | 0.38 |
| 155.00 | 171.00 | 161.33 | 1.20 | 0.02 |
| 73.00 | 55.00 | 66.33 | -1.52 | 0.00 |
| 199.00 | 148.00 | 157.67 | -1.26 | 0.05 |
| 129.00 | 167.00 | 147.33 | -1.24 | 0.01 |
| 1495.00 | 1602.00 | 1615.00 | -1.08 | 0.04 |
| 373.00 | 326.00 | 338.00 | -1.01 | 0.96 |
| 565.00 | 615.00 | 571.67 | -1.12 | 0.06 |
| 2070.00 | 2033.00 | 2045.67 | -1.15 | 0.00 |
| 1376.00 | 1590.00 | 1494.67 | 1.28 | 0.00 |
| 0.00 | 0.00 | 0.00 | -9.06 | 0.27 |
| 1223.00 | 1217.00 | 1218.67 | -1.03 | 0.50 |
| 407.00 | 369.00 | 384.67 | -1.18 | 0.00 |
| 1278.00 | 1359.00 | 1285.33 | 1.23 | 0.00 |
| 533.00 | 625.00 | 584.67 | 1.38 | 0.00 |
| 475.00 | 532.00 | 485.67 | -1.08 | 0.21 |
| 23.00 | 16.00 | 22.33 | -1.89 | 0.00 |
| 953.00 | 1168.00 | 1097.00 | 1.25 | 0.00 |
| 1170.00 | 1217.00 | 1241.33 | 1.25 | 0.00 |
| 1336.00 | 1414.00 | 1319.00 | -1.10 | 0.08 |
| 4712.00 | 4857.00 | 4909.00 | -1.02 | 0.64 |
| 622.00 | 686.00 | 667.67 | 1.18 | 0.00 |
| 1370.00 | 1519.00 | 1430.67 | 1.00 | 0.97 |
| 1262.00 | 1406.00 | 1296.33 | 1.03 | 0.57 |
| 659.00 | 601.00 | 624.00 | -1.68 | 0.00 |
| 1472.00 | 1620.00 | 1589.33 | 1.21 | 0.00 |
| 1321.00 | 1636.00 | 1488.33 | 1.17 | 0.00 |
| 267.00 | 305.00 | 305.33 | 1.13 | 0.07 |
| 152.00 | 149.00 | 148.67 | -1.16 | 0.11 |
| 1073.00 | 1094.00 | 1076.33 | 1.18 | 0.00 |
| 1564.00 | 1747.00 | 1700.67 | 1.22 | 0.00 |

| | | | | |
|---|---|---|---|---|
| 641.00 | 814.00 | 696.67 | -1.01 | 0.89 |
| 1281.00 | 1347.00 | 1309.33 | 1.01 | 0.77 |
| 667.00 | 587.00 | 617.33 | 1.13 | 0.06 |
| 1296.00 | 1552.00 | 1424.33 | 1.22 | 0.00 |
| 29.00 | 32.00 | 26.33 | 1.73 | 0.01 |
| 4436.00 | 4698.00 | 4404.00 | -1.03 | 0.54 |
| 178.00 | 196.00 | 183.00 | -1.39 | 0.00 |
| 46.00 | 49.00 | 49.00 | -1.13 | 0.35 |
| 733.00 | 786.00 | 757.67 | -1.23 | 0.00 |
| 149.00 | 162.00 | 167.33 | 1.12 | 0.22 |
| 46.00 | 34.00 | 36.33 | -1.45 | 0.01 |
| 582.00 | 665.00 | 635.00 | 1.02 | 0.69 |
| 8.00 | 11.00 | 7.67 | 1.03 | 1.00 |
| 2357.00 | 2328.00 | 2456.67 | -1.19 | 0.00 |
| 1895.00 | 1994.00 | 2037.33 | -1.03 | 0.48 |
| 509.00 | 628.00 | 592.00 | -1.01 | 0.87 |
| 5221.00 | 4900.00 | 4986.67 | -1.14 | 0.00 |
| 20.00 | 34.00 | 25.00 | -1.74 | 0.00 |
| 9.00 | 18.00 | 12.33 | -5.01 | 0.00 |
| 187.00 | 222.00 | 200.33 | 1.02 | 0.85 |
| 381.00 | 424.00 | 428.33 | -1.11 | 0.06 |
| 1865.00 | 1943.00 | 1958.33 | 1.04 | 0.29 |
| 2456.00 | 2562.00 | 2573.33 | -1.04 | 0.21 |
| 653.00 | 653.00 | 659.33 | -1.21 | 0.00 |
| 686.00 | 772.00 | 752.33 | 1.01 | 0.86 |
| 77.00 | 74.00 | 82.00 | 1.09 | 0.47 |
| 312.00 | 305.00 | 321.67 | -1.10 | 0.09 |
| 25.00 | 25.00 | 31.00 | -1.12 | 0.51 |
| 8.00 | 16.00 | 15.67 | 1.28 | 0.35 |
| 95.00 | 153.00 | 133.00 | -1.08 | 0.43 |
| 1132.00 | 1205.00 | 1227.00 | -1.01 | 0.81 |
| 606.00 | 540.00 | 590.33 | 1.80 | 0.00 |
| 1559.00 | 1691.00 | 1625.67 | 1.03 | 0.50 |
| 42.00 | 69.00 | 59.67 | 1.19 | 0.26 |
| 348.00 | 373.00 | 364.33 | -1.12 | 0.03 |
| 30.00 | 49.00 | 41.33 | 1.08 | 0.64 |
| 2198.00 | 2372.00 | 2324.67 | -1.02 | 0.52 |
| 1012.00 | 1029.00 | 1032.00 | -1.09 | 0.06 |
| 265.00 | 258.00 | 264.00 | 1.07 | 0.32 |
| 33.00 | 45.00 | 39.67 | 1.00 | 1.00 |
| 1247.00 | 1292.00 | 1297.00 | -1.07 | 0.11 |
| 557.00 | 673.00 | 611.67 | 1.01 | 0.93 |
| 1170.00 | 1225.00 | 1266.33 | 1.10 | 0.02 |
| 817.00 | 916.00 | 874.67 | -1.04 | 0.38 |
| 18.00 | 19.00 | 15.67 | -1.63 | 0.02 |
| 107.00 | 126.00 | 125.00 | 1.13 | 0.27 |
| 290.00 | 344.00 | 326.67 | 1.02 | 0.80 |
| 1424.00 | 1533.00 | 1525.67 | 1.08 | 0.04 |

| | | | | |
|---|---|---|---|---|
| 9.00 | 33.00 | 21.33 | 1.06 | 0.89 |
| 28.00 | 17.00 | 26.33 | 1.89 | 0.00 |
| 220.00 | 250.00 | 239.00 | 1.09 | 0.26 |
| 1047.00 | 1027.00 | 1082.00 | -1.09 | 0.06 |
| 789.00 | 695.00 | 759.33 | 1.07 | 0.22 |
| 354.00 | 391.00 | 361.67 | 1.04 | 0.51 |
| 571.00 | 583.00 | 581.00 | -1.03 | 0.62 |
| 1853.00 | 1729.00 | 1777.67 | -1.17 | 0.00 |
| 362.00 | 466.00 | 443.33 | -1.07 | 0.32 |
| 18.00 | 16.00 | 17.33 | 1.46 | 0.10 |
| 416.00 | 440.00 | 407.00 | -1.15 | 0.04 |
| 117.00 | 124.00 | 128.33 | -1.06 | 0.51 |
| 148.00 | 136.00 | 157.67 | -1.09 | 0.34 |
| 189.00 | 212.00 | 193.67 | -1.03 | 0.74 |
| 996.00 | 1001.00 | 1021.33 | -1.31 | 0.00 |
| 685.00 | 755.00 | 731.33 | -1.19 | 0.00 |
| 49.00 | 64.00 | 59.33 | -1.03 | 0.81 |
| 60.00 | 47.00 | 52.33 | 1.27 | 0.09 |
| 1513.00 | 1523.00 | 1526.00 | -1.18 | 0.00 |
| 324.00 | 348.00 | 325.33 | -1.10 | 0.24 |
| 12.00 | 16.00 | 15.00 | -1.08 | 0.84 |
| 2228.00 | 2550.00 | 2432.33 | -1.03 | 0.44 |
| 2899.00 | 3154.00 | 3022.00 | 1.03 | 0.53 |
| 771.00 | 859.00 | 825.67 | 1.04 | 0.35 |
| 46.00 | 41.00 | 46.00 | 1.04 | 0.82 |
| 385.00 | 421.00 | 407.67 | 1.11 | 0.07 |
| 2235.00 | 2408.00 | 2347.33 | 1.07 | 0.10 |
| 47.00 | 32.00 | 38.67 | -1.38 | 0.03 |
| 11.00 | 16.00 | 14.00 | 1.42 | 0.17 |
| 3.00 | 8.00 | 7.67 | 1.03 | 1.00 |
| 73.00 | 92.00 | 85.33 | 1.09 | 0.39 |
| 34.00 | 17.00 | 29.67 | -1.51 | 0.01 |
| 8.00 | 11.00 | 7.67 | -1.74 | 0.07 |
| 2.00 | 1.00 | 1.33 | -4.72 | 0.00 |
| 52.00 | 58.00 | 64.33 | 1.40 | 0.01 |
| 2954.00 | 2816.00 | 2807.00 | -1.26 | 0.00 |
| 1936.00 | 1816.00 | 1870.00 | -1.23 | 0.00 |
| 1307.00 | 1436.00 | 1350.00 | -1.09 | 0.08 |
| 4718.00 | 3596.00 | 3923.00 | -1.43 | 0.00 |
| 1055.00 | 1027.00 | 1016.67 | -1.29 | 0.00 |
| 88.00 | 63.00 | 80.33 | -1.08 | 0.45 |
| 471.00 | 462.00 | 461.00 | 1.23 | 0.00 |
| 609.00 | 723.00 | 714.00 | 1.22 | 0.00 |
| 1394.00 | 1532.00 | 1473.00 | 1.12 | 0.01 |
| 2165.00 | 2404.00 | 2354.67 | 1.00 | 0.98 |
| 3868.00 | 4211.00 | 4150.67 | 1.30 | 0.00 |
| 1232.00 | 1465.00 | 1367.67 | 1.25 | 0.00 |
| 3186.00 | 3579.00 | 3495.33 | 1.16 | 0.00 |

| | | | | |
|---|---|---|---|---|
| 174.00 | 180.00 | 165.00 | 1.03 | 0.78 |
| 2150.00 | 2104.00 | 2150.00 | -1.08 | 0.05 |
| 9437.00 | 9785.00 | 9651.00 | 1.07 | 0.05 |
| 2.00 | 7.00 | 5.00 | 1.06 | 1.00 |
| 266.00 | 334.00 | 296.00 | 1.04 | 0.51 |
| 865.00 | 997.00 | 925.00 | 1.02 | 0.74 |
| 313.00 | 325.00 | 328.67 | -1.08 | 0.19 |
| 52.00 | 48.00 | 47.33 | -1.59 | 0.00 |
| 635.00 | 731.00 | 752.00 | 1.02 | 0.70 |
| 674.00 | 854.00 | 786.67 | 1.17 | 0.00 |
| 250.00 | 230.00 | 244.33 | -1.40 | 0.00 |
| 0.00 | 4.00 | 2.33 | -1.98 | 0.20 |
| 1910.00 | 2012.00 | 2059.67 | 1.03 | 0.44 |
| 1209.00 | 1372.00 | 1303.00 | -1.08 | 0.06 |
| 985.00 | 1146.00 | 1077.33 | 1.09 | 0.09 |
| 435.00 | 466.00 | 470.67 | 1.05 | 0.33 |
| 229.00 | 281.00 | 268.00 | 2.36 | 0.00 |
| 2951.00 | 3281.00 | 3311.00 | -1.22 | 0.00 |
| 3205.00 | 3389.00 | 3474.00 | -1.74 | 0.00 |
| 220.00 | 249.00 | 255.67 | -1.35 | 0.00 |
| 788.00 | 838.00 | 865.67 | -1.21 | 0.00 |
| 571.00 | 565.00 | 594.67 | -1.30 | 0.00 |
| 8529.00 | 8811.00 | 9187.67 | -1.88 | 0.00 |
| 12.00 | 14.00 | 13.67 | -1.80 | 0.00 |
| 0.00 | 3.00 | 1.00 | -5.51 | 0.00 |
| 85.00 | 71.00 | 88.00 | -2.07 | 0.00 |
| 206.00 | 148.00 | 222.00 | 7.20 | 0.00 |
| 191.00 | 93.00 | 127.33 | 3.12 | 0.00 |
| 11.00 | 23.00 | 16.00 | 1.10 | 0.78 |
| 86.00 | 103.00 | 98.33 | 2.99 | 0.00 |
| 27.00 | 23.00 | 24.00 | 3.06 | 0.00 |
| 5.00 | 1.00 | 2.67 | -2.22 | 0.08 |
| 2.00 | 2.00 | 1.33 | -5.18 | 0.00 |
| 1338.00 | 1424.00 | 1489.33 | 1.12 | 0.02 |
| 37.00 | 39.00 | 41.00 | 1.03 | 0.86 |
| 406.00 | 375.00 | 395.33 | -1.03 | 0.69 |
| 5890.00 | 5951.00 | 5823.00 | 1.04 | 0.35 |
| 44.00 | 59.00 | 55.67 | -1.09 | 0.47 |
| 1617.00 | 1667.00 | 1758.00 | 1.03 | 0.54 |
| 5.00 | 0.00 | 4.00 | 1.07 | 1.00 |
| 3.00 | 1.00 | 1.67 | -1.77 | 0.43 |
| 304.00 | 338.00 | 310.67 | 1.14 | 0.05 |
| 696.00 | 779.00 | 738.67 | -1.23 | 0.00 |
| 267.00 | 255.00 | 259.33 | -1.55 | 0.00 |
| 648.00 | 696.00 | 688.33 | 1.11 | 0.02 |
| 647.00 | 738.00 | 729.00 | 1.02 | 0.72 |
| 399.00 | 427.00 | 421.33 | -1.03 | 0.52 |
| 129.00 | 99.00 | 110.33 | 1.15 | 0.23 |

| | | | | |
|---|---|---|---|---|
| 974.00 | 1064.00 | 1023.33 | 1.00 | 1.00 |
| 539.00 | 624.00 | 599.67 | 1.19 | 0.00 |
| 10.00 | 9.00 | 7.33 | -1.01 | 1.00 |
| 208.00 | 239.00 | 244.67 | 1.28 | 0.00 |
| 136.00 | 111.00 | 131.33 | -1.09 | 0.33 |
| 18.00 | 14.00 | 22.67 | 1.48 | 0.08 |
| 445.00 | 418.00 | 416.00 | -2.18 | 0.00 |
| 4922.00 | 5392.00 | 5158.67 | -1.05 | 0.19 |
| 2764.00 | 2616.00 | 2712.33 | -1.36 | 0.00 |
| 18755.00 | 20188.00 | 19862.33 | -1.42 | 0.00 |
| 4847.00 | 5041.00 | 5093.00 | -1.41 | 0.00 |
| 0.00 | 6.00 | 2.67 | -1.62 | 0.39 |
| 24771.00 | 26014.00 | 25954.67 | -1.13 | 0.00 |
| 6.00 | 6.00 | 6.33 | 1.33 | 0.49 |
| 19532.00 | 20479.00 | 20056.33 | -1.05 | 0.20 |
| 1192.00 | 1215.00 | 1236.67 | 1.11 | 0.01 |
| 5596.00 | 6028.00 | 5850.67 | -1.49 | 0.00 |
| 2911.00 | 3062.00 | 2959.00 | -1.13 | 0.00 |
| 1584.00 | 1651.00 | 1758.00 | 1.01 | 0.87 |
| 543.00 | 645.00 | 613.67 | 1.20 | 0.00 |
| 386.00 | 452.00 | 442.67 | 1.21 | 0.00 |
| 698.00 | 776.00 | 791.67 | 1.06 | 0.28 |
| 919.00 | 1083.00 | 1084.33 | 1.05 | 0.37 |
| 14.00 | 12.00 | 12.67 | 1.95 | 0.02 |
| 1715.00 | 1775.00 | 1734.67 | 1.08 | 0.07 |
| 451.00 | 503.00 | 483.00 | 1.17 | 0.00 |
| 747.00 | 797.00 | 800.00 | 1.10 | 0.02 |
| 2986.00 | 3084.00 | 3138.67 | -1.08 | 0.04 |
| 5666.00 | 5809.00 | 5882.67 | 1.10 | 0.02 |
| 10797.00 | 10641.00 | 10806.33 | -1.01 | 0.85 |
| 327.00 | 366.00 | 340.00 | 1.01 | 0.83 |
| 1970.00 | 2281.00 | 2112.67 | 1.07 | 0.17 |
| 2461.00 | 2667.00 | 2653.00 | 1.05 | 0.16 |
| 1484.00 | 1558.00 | 1580.67 | -1.04 | 0.33 |
| 699.00 | 837.00 | 805.00 | 1.10 | 0.07 |
| 10595.00 | 11497.00 | 11063.67 | -1.01 | 0.74 |
| 1316.00 | 1409.00 | 1383.33 | -1.03 | 0.44 |
| 2905.00 | 3142.00 | 3026.00 | 1.06 | 0.13 |
| 3282.00 | 3343.00 | 3388.67 | -1.28 | 0.00 |
| 452.00 | 410.00 | 439.00 | -1.53 | 0.00 |
| 180.00 | 192.00 | 194.33 | 5.12 | 0.00 |
| 725.00 | 756.00 | 750.33 | -1.06 | 0.24 |
| 479.00 | 463.00 | 487.67 | -1.00 | 0.97 |
| 849.00 | 854.00 | 848.33 | 1.33 | 0.00 |
| 713.00 | 713.00 | 745.00 | 1.28 | 0.00 |
| 2269.00 | 2467.00 | 2349.00 | 1.03 | 0.52 |
| 79.00 | 88.00 | 84.67 | 1.05 | 0.65 |
| 119.00 | 140.00 | 134.00 | -1.08 | 0.37 |

| | | | | |
|---|---|---|---|---|
| 121.00 | 132.00 | 133.00 | -1.04 | 0.62 |
| 37.00 | 39.00 | 35.67 | 1.11 | 0.55 |
| 180.00 | 191.00 | 191.67 | 1.03 | 0.64 |
| 617.00 | 658.00 | 656.67 | 1.14 | 0.01 |
| 369.00 | 355.00 | 382.67 | 1.05 | 0.41 |
| 5.00 | 15.00 | 9.67 | 1.06 | 0.90 |
| 284.00 | 287.00 | 293.00 | -1.02 | 0.82 |
| 534.00 | 561.00 | 533.33 | -1.08 | 0.22 |
| 114.00 | 137.00 | 125.33 | -1.09 | 0.29 |
| 59.00 | 46.00 | 49.33 | -1.52 | 0.00 |
| 2.00 | 1.00 | 1.00 | -1.61 | 0.73 |
| 117.00 | 100.00 | 108.00 | 1.65 | 0.00 |
| 578.00 | 617.00 | 612.33 | 1.02 | 0.63 |
| 1419.00 | 1365.00 | 1452.33 | -1.28 | 0.00 |
| 10915.00 | 12490.00 | 12145.00 | 1.48 | 0.00 |
| 19473.00 | 21396.00 | 21481.67 | 1.28 | 0.00 |
| 95.00 | 96.00 | 101.33 | 1.47 | 0.00 |
| 15.00 | 13.00 | 16.00 | 1.21 | 0.41 |
| 1315.00 | 1281.00 | 1284.33 | -1.57 | 0.00 |
| 1443.00 | 1239.00 | 1318.33 | 1.30 | 0.00 |
| 269.00 | 276.00 | 293.33 | -1.01 | 0.88 |
| 577.00 | 564.00 | 545.67 | -1.20 | 0.00 |
| 47.00 | 56.00 | 56.33 | 1.02 | 0.92 |
| 311.00 | 297.00 | 320.00 | 1.31 | 0.00 |
| 7.00 | 10.00 | 9.67 | 1.06 | 0.90 |
| 2086.00 | 2061.00 | 2135.33 | -1.10 | 0.01 |
| 813.00 | 883.00 | 806.33 | -1.05 | 0.40 |
| 112.00 | 122.00 | 104.67 | -1.56 | 0.00 |
| 12.00 | 14.00 | 13.00 | 1.47 | 0.15 |
| 695.00 | 825.00 | 771.67 | 1.17 | 0.00 |
| 353.00 | 409.00 | 386.33 | 1.07 | 0.23 |
| 2188.00 | 2344.00 | 2302.00 | 1.04 | 0.32 |
| 827.00 | 861.00 | 860.33 | 1.12 | 0.02 |
| 2576.00 | 2728.00 | 2766.67 | -1.03 | 0.34 |
| 560.00 | 616.00 | 622.67 | 1.11 | 0.03 |
| 2186.00 | 2463.00 | 2413.67 | -1.07 | 0.06 |
| 25.00 | 29.00 | 29.67 | -1.39 | 0.03 |
| 5774.00 | 6785.00 | 6849.33 | 1.08 | 0.13 |
| 749.00 | 782.00 | 828.00 | -1.07 | 0.16 |
| 15193.00 | 16346.00 | 16194.67 | -1.15 | 0.00 |
| 927.00 | 1014.00 | 997.67 | -1.17 | 0.00 |
| 1.00 | 7.00 | 3.33 | 1.22 | 0.82 |
| 91.00 | 103.00 | 105.00 | 1.22 | 0.05 |
| 1.00 | 0.00 | 0.67 | -3.58 | 0.12 |
| 1417.00 | 1467.00 | 1488.33 | 1.07 | 0.14 |
| 221.00 | 252.00 | 230.33 | 1.38 | 0.00 |
| 458.00 | 471.00 | 462.33 | -1.53 | 0.00 |
| 0.00 | 2.00 | 0.67 | -3.14 | 0.19 |

| | | | | |
|---|---|---|---|---|
| 23.00 | 21.00 | 22.67 | 1.03 | 0.93 |
| 337.00 | 354.00 | 365.00 | -1.32 | 0.00 |
| 1306.00 | 1339.00 | 1359.33 | -1.03 | 0.48 |
| 115.00 | 114.00 | 119.00 | 1.05 | 0.58 |
| 6.00 | 3.00 | 3.33 | 1.23 | 0.82 |
| 679.00 | 659.00 | 712.33 | -1.12 | 0.03 |
| 2.00 | 3.00 | 2.33 | -1.56 | 0.49 |
| 965.00 | 1048.00 | 1028.00 | -1.01 | 0.82 |
| 7802.00 | 8056.00 | 8221.67 | -1.13 | 0.00 |
| 590.00 | 590.00 | 583.33 | -1.43 | 0.00 |
| 9.00 | 12.00 | 15.00 | -1.24 | 0.34 |
| 888.00 | 955.00 | 924.00 | 1.19 | 0.00 |
| 840.00 | 730.00 | 779.33 | -1.09 | 0.14 |
| 465.00 | 592.00 | 534.00 | -1.07 | 0.28 |
| 1095.00 | 1084.00 | 1130.00 | -1.13 | 0.00 |
| 5.00 | 0.00 | 2.00 | -1.01 | 1.00 |
| 2072.00 | 1967.00 | 2038.67 | -1.04 | 0.32 |
| 10.00 | 6.00 | 7.33 | 1.27 | 0.53 |
| 568.00 | 609.00 | 631.00 | -1.08 | 0.15 |
| 992.00 | 1116.00 | 1085.33 | -1.03 | 0.52 |
| 1477.00 | 1407.00 | 1465.67 | -1.35 | 0.00 |
| 316.00 | 315.00 | 327.67 | 1.06 | 0.32 |
| 1009.00 | 1111.00 | 1056.67 | 1.26 | 0.00 |
| 86.00 | 72.00 | 76.33 | 1.15 | 0.20 |
| 195.00 | 213.00 | 224.67 | 1.12 | 0.15 |
| 196.00 | 228.00 | 211.33 | 1.10 | 0.18 |
| 259.00 | 276.00 | 267.00 | 1.16 | 0.02 |
| 48.00 | 76.00 | 63.00 | -1.18 | 0.16 |
| 440.00 | 548.00 | 507.33 | 1.30 | 0.00 |
| 261.00 | 318.00 | 292.33 | 1.26 | 0.00 |
| 627.00 | 629.00 | 656.33 | -1.03 | 0.56 |
| 23.00 | 31.00 | 24.67 | -1.20 | 0.34 |
| 222.00 | 248.00 | 267.33 | 1.15 | 0.10 |
| 240.00 | 248.00 | 263.67 | -1.13 | 0.06 |
| 61.00 | 54.00 | 60.67 | 1.38 | 0.01 |
| 636.00 | 713.00 | 703.67 | -1.16 | 0.00 |
| 78.00 | 73.00 | 77.00 | -1.28 | 0.01 |
| 8.00 | 13.00 | 12.00 | 1.22 | 0.47 |
| 44.00 | 58.00 | 52.67 | 1.00 | 1.00 |
| 20.00 | 18.00 | 17.33 | 1.07 | 0.84 |
| 16.00 | 19.00 | 21.67 | 1.03 | 1.00 |
| 15.00 | 18.00 | 17.67 | 1.45 | 0.10 |
| 3.00 | 2.00 | 3.00 | -1.66 | 0.31 |
| 337.00 | 362.00 | 356.33 | -1.61 | 0.00 |
| 241.00 | 254.00 | 238.67 | 1.02 | 0.76 |
| 1.00 | 3.00 | 1.67 | -1.77 | 0.43 |
| 5.00 | 5.00 | 4.67 | -1.08 | 1.00 |
| 331.00 | 352.00 | 367.00 | -1.07 | 0.21 |

| | | | | |
|---|---|---|---|---|
| 2.00 | 2.00 | 2.00 | -1.17 | 1.00 |
| 16.00 | 22.00 | 22.00 | 1.96 | 0.00 |
| 5.00 | 5.00 | 5.00 | 1.23 | 0.71 |
| 1.00 | 3.00 | 2.67 | -1.13 | 1.00 |
| 7.00 | 5.00 | 6.00 | 1.36 | 0.48 |
| 8.00 | 3.00 | 5.67 | -1.71 | 0.11 |
| 52.00 | 67.00 | 61.00 | 1.73 | 0.00 |
| 8.00 | 3.00 | 6.33 | -1.43 | 0.31 |
| 307.00 | 330.00 | 346.33 | 1.11 | 0.13 |
| 114.00 | 142.00 | 147.00 | -1.27 | 0.01 |
| 360.00 | 381.00 | 380.67 | -1.88 | 0.00 |
| 187.00 | 207.00 | 205.00 | -1.35 | 0.00 |
| 7.00 | 14.00 | 12.67 | 1.20 | 0.57 |
| 4.00 | 6.00 | 5.33 | -1.45 | 0.35 |
| 5.00 | 3.00 | 3.33 | 1.39 | 0.64 |
| 7.00 | 2.00 | 4.33 | -1.01 | 1.00 |
| 6.00 | 3.00 | 3.33 | 1.39 | 0.64 |
| 36.00 | 44.00 | 35.67 | 1.29 | 0.12 |
| 171.00 | 169.00 | 184.00 | 1.22 | 0.01 |
| 109.00 | 133.00 | 124.67 | -1.06 | 0.48 |
| 54.00 | 48.00 | 51.00 | -1.05 | 0.75 |
| 486.00 | 578.00 | 525.33 | 1.11 | 0.07 |
| 1.00 | 2.00 | 1.33 | -1.01 | 1.00 |
| 86.00 | 94.00 | 91.67 | 1.01 | 0.91 |
| 0.00 | 0.00 | 0.00 | -11.75 | 0.13 |
| 633.00 | 730.00 | 756.33 | 1.21 | 0.01 |
| 0.00 | 5.00 | 2.67 | 1.89 | 0.40 |
| 16.00 | 9.00 | 14.33 | 1.28 | 0.32 |
| 2.00 | 4.00 | 2.33 | -2.12 | 0.15 |
| 83.00 | 108.00 | 96.67 | 1.11 | 0.29 |
| 0.00 | 0.00 | 0.00 | -14.44 | 0.07 |
| 7.00 | 2.00 | 5.67 | -1.13 | 0.87 |
| 576.00 | 629.00 | 646.33 | -1.22 | 0.00 |
| 3.00 | 1.00 | 2.00 | -1.17 | 1.00 |
| 7.00 | 10.00 | 13.67 | -1.12 | 0.69 |
| 8.00 | 8.00 | 5.33 | -1.25 | 0.69 |
| 39.00 | 54.00 | 45.33 | 1.01 | 1.00 |
| 2.00 | 1.00 | 1.00 | -2.21 | 0.35 |
| 45.00 | 49.00 | 46.00 | -1.26 | 0.07 |
| 114.00 | 120.00 | 130.67 | -1.16 | 0.08 |
| 73.00 | 83.00 | 74.00 | 1.03 | 0.84 |
| 60.00 | 66.00 | 65.67 | -1.31 | 0.01 |
| 69.00 | 54.00 | 60.00 | -1.52 | 0.00 |
| 2.00 | 1.00 | 2.33 | -1.29 | 0.81 |
| 110.00 | 133.00 | 118.67 | -1.25 | 0.01 |
| 8.00 | 10.00 | 8.67 | -1.32 | 0.38 |
| 443.00 | 486.00 | 432.67 | -1.40 | 0.00 |
| 79.00 | 96.00 | 92.00 | 1.31 | 0.01 |

| | | | | |
|---|---|---|---|---|
| 3.00 | 4.00 | 3.67 | -1.01 | 1.00 |
| 95.00 | 97.00 | 114.67 | -1.20 | 0.08 |
| 19.00 | 18.00 | 16.33 | 1.21 | 0.42 |
| 10003.00 | 10458.00 | 10021.33 | 1.01 | 0.75 |
| 1.00 | 1.00 | 3.00 | 2.72 | 0.18 |
| 87.00 | 99.00 | 99.67 | 1.05 | 0.68 |
| 23.00 | 18.00 | 25.67 | -1.24 | 0.26 |
| 47.00 | 38.00 | 42.00 | 1.12 | 0.46 |
| 1336.00 | 1421.00 | 1434.67 | -1.05 | 0.22 |
| 11.00 | 21.00 | 16.67 | 1.33 | 0.26 |
| 7.00 | 10.00 | 10.33 | -1.50 | 0.12 |
| 31.00 | 32.00 | 35.67 | -1.68 | 0.00 |
| 617.00 | 644.00 | 663.33 | -1.30 | 0.00 |
| 0.00 | 0.00 | 0.00 | -9.06 | 0.27 |
| 12.00 | 15.00 | 12.67 | 1.07 | 0.82 |
| 2.00 | 0.00 | 0.67 | -1.86 | 0.69 |
| 32.00 | 29.00 | 31.33 | -1.48 | 0.01 |
| 1843.00 | 1976.00 | 2005.33 | -1.65 | 0.00 |
| 13.00 | 16.00 | 14.00 | 1.38 | 0.21 |
| 1746.00 | 1918.00 | 1923.33 | 2.13 | 0.00 |
| 927.00 | 979.00 | 988.67 | 1.08 | 0.09 |
| 11.00 | 9.00 | 11.67 | -2.05 | 0.00 |
| 197.00 | 227.00 | 222.33 | -1.14 | 0.06 |
| 36.00 | 33.00 | 35.67 | -1.30 | 0.07 |
| 6.00 | 22.00 | 14.00 | -1.11 | 0.70 |
| 28.00 | 18.00 | 27.33 | -1.41 | 0.03 |
| 10.00 | 10.00 | 10.00 | 1.18 | 0.60 |
| 36.00 | 43.00 | 40.67 | -1.25 | 0.09 |
| 4.00 | 7.00 | 6.67 | -1.06 | 1.00 |
| 9.00 | 14.00 | 14.00 | -1.11 | 0.68 |
| 195.00 | 231.00 | 222.00 | 1.02 | 0.80 |
| 28.00 | 35.00 | 29.33 | 1.01 | 1.00 |
| 118.00 | 104.00 | 123.00 | -1.25 | 0.01 |
| 6.00 | 8.00 | 6.00 | -1.12 | 0.87 |
| 17.00 | 13.00 | 14.33 | -1.34 | 0.21 |
| 75.00 | 72.00 | 80.00 | -1.25 | 0.02 |
| 1.00 | 8.00 | 5.67 | -1.83 | 0.07 |
| 158.00 | 151.00 | 167.33 | -1.06 | 0.45 |
| 221.00 | 303.00 | 280.67 | 1.18 | 0.03 |
| 70.00 | 54.00 | 65.33 | -1.07 | 0.61 |
| 32.00 | 65.00 | 55.33 | 1.39 | 0.04 |
| 51.00 | 53.00 | 53.00 | 1.48 | 0.00 |
| 36.00 | 56.00 | 49.33 | 1.03 | 0.91 |
| 41.00 | 41.00 | 44.67 | -1.07 | 0.61 |
| 41.00 | 20.00 | 35.00 | 1.00 | 1.00 |
| 17.00 | 19.00 | 15.67 | -1.65 | 0.01 |
| 203.00 | 243.00 | 230.00 | 1.00 | 0.98 |
| 372.00 | 414.00 | 412.00 | 1.12 | 0.06 |

| | | | | |
|---|---|---|---|---|
| 125.00 | 112.00 | 116.67 | 1.33 | 0.00 |
| 6.00 | 9.00 | 6.00 | -1.73 | 0.10 |
| 14.00 | 7.00 | 10.33 | 1.22 | 0.53 |
| 55.00 | 47.00 | 53.00 | 1.31 | 0.05 |
| 29.00 | 37.00 | 31.67 | 1.05 | 0.83 |
| 263.00 | 273.00 | 276.00 | -1.16 | 0.01 |
| 0.00 | 1.00 | 0.67 | -2.72 | 0.29 |
| 127.00 | 169.00 | 159.67 | 1.06 | 0.50 |
| 886.00 | 1043.00 | 1002.33 | 1.16 | 0.00 |
| 138.00 | 153.00 | 149.00 | -1.43 | 0.00 |
| 2861.00 | 2704.00 | 2767.33 | -1.27 | 0.00 |
| 57.00 | 82.00 | 70.67 | 1.17 | 0.19 |
| 463.00 | 568.00 | 508.33 | -1.30 | 0.00 |
| 56.00 | 74.00 | 68.33 | 1.35 | 0.01 |
| 11.00 | 2.00 | 6.00 | 1.18 | 0.74 |
| 93.00 | 112.00 | 97.33 | -1.16 | 0.11 |
| 14.00 | 21.00 | 22.00 | 1.32 | 0.23 |
| 76.00 | 69.00 | 75.33 | 1.11 | 0.36 |
| 18.00 | 16.00 | 20.33 | -1.23 | 0.26 |
| 6.00 | 8.00 | 7.00 | 1.15 | 0.75 |
| 0.00 | 2.00 | 1.67 | -2.15 | 0.22 |
| 245.00 | 227.00 | 238.33 | -1.31 | 0.00 |
| 2.00 | 0.00 | 1.67 | -3.09 | 0.03 |
| 9.00 | 6.00 | 6.67 | -1.01 | 1.00 |
| 522.00 | 575.00 | 567.33 | -1.13 | 0.01 |
| 16.00 | 23.00 | 22.67 | -1.72 | 0.00 |
| 3.00 | 6.00 | 4.33 | -1.17 | 0.85 |
| 2.00 | 1.00 | 2.67 | -1.86 | 0.22 |
| 64.00 | 61.00 | 68.00 | 1.20 | 0.14 |
| 402.00 | 471.00 | 450.33 | 1.21 | 0.00 |
| 37.00 | 38.00 | 41.00 | 1.05 | 0.78 |
| 7.00 | 8.00 | 8.00 | -1.01 | 1.00 |
| 32.00 | 41.00 | 39.33 | -1.08 | 0.59 |
| 1.00 | 5.00 | 2.00 | -1.49 | 0.61 |
| 24.00 | 25.00 | 28.67 | -1.20 | 0.27 |
| 115.00 | 77.00 | 88.67 | -1.56 | 0.00 |
| 218.00 | 237.00 | 234.33 | -1.25 | 0.00 |
| 6.00 | 6.00 | 4.67 | -1.93 | 0.07 |
| 6.00 | 10.00 | 8.33 | 1.07 | 0.89 |
| 39.00 | 29.00 | 42.67 | 1.08 | 0.65 |
| 107.00 | 142.00 | 130.00 | -1.42 | 0.00 |
| 212.00 | 252.00 | 254.00 | 1.04 | 0.66 |
| 11.00 | 21.00 | 21.00 | -1.08 | 0.75 |
| 36.00 | 50.00 | 43.67 | -1.11 | 0.47 |
| 0.00 | 3.00 | 1.33 | -1.48 | 0.77 |
| 15.00 | 14.00 | 19.00 | 1.47 | 0.10 |
| 2.00 | 4.00 | 4.00 | -1.42 | 0.47 |
| 15.00 | 21.00 | 15.00 | -1.15 | 0.63 |

| | | | | |
|---|---|---|---|---|
| 349.00 | 442.00 | 409.33 | 1.14 | 0.03 |
| 0.00 | 2.00 | 0.67 | -5.27 | 0.01 |
| 318.00 | 385.00 | 357.67 | -1.03 | 0.55 |
| 2.00 | 0.00 | 1.33 | -1.24 | 1.00 |
| 16.00 | 8.00 | 12.67 | -1.14 | 0.67 |
| 127.00 | 164.00 | 165.00 | 1.24 | 0.03 |
| 47.00 | 51.00 | 60.33 | -1.59 | 0.00 |
| 54.00 | 61.00 | 64.67 | 1.03 | 0.82 |
| 208.00 | 197.00 | 197.00 | 2.01 | 0.00 |
| 3.00 | 4.00 | 3.33 | -2.09 | 0.08 |
| 925.00 | 869.00 | 937.00 | -1.12 | 0.02 |
| 547.00 | 583.00 | 562.00 | 1.55 | 0.00 |
| 34.00 | 24.00 | 31.67 | 1.20 | 0.29 |
| 122.00 | 124.00 | 127.33 | -1.28 | 0.00 |
| 12.00 | 19.00 | 15.33 | 1.11 | 0.68 |
| 97.00 | 110.00 | 103.33 | -1.02 | 0.83 |
| 12.00 | 18.00 | 15.00 | -1.06 | 0.92 |
| 130.00 | 120.00 | 137.00 | -1.04 | 0.62 |
| 0.00 | 1.00 | 0.33 | -2.48 | 0.64 |
| 2.00 | 3.00 | 2.33 | -1.56 | 0.49 |
| 12.00 | 18.00 | 16.00 | 1.13 | 0.61 |
| 2.00 | 4.00 | 3.00 | -1.55 | 0.41 |
| 6.00 | 0.00 | 2.67 | -1.13 | 1.00 |
| 532.00 | 729.00 | 634.67 | -1.07 | 0.28 |
| 8.00 | 6.00 | 9.00 | -1.72 | 0.04 |
| 131.00 | 138.00 | 138.00 | 1.93 | 0.00 |
| 7.00 | 2.00 | 4.33 | -1.92 | 0.08 |
| 17.00 | 12.00 | 15.00 | -1.44 | 0.08 |
| 1.00 | 3.00 | 2.00 | -1.49 | 0.61 |
| 0.00 | 3.00 | 1.67 | -1.77 | 0.43 |
| 50.00 | 52.00 | 53.33 | -1.61 | 0.00 |
| 323.00 | 362.00 | 364.33 | -1.38 | 0.00 |
| 598.00 | 647.00 | 664.00 | 1.07 | 0.17 |
| 17.00 | 24.00 | 22.67 | -1.08 | 0.70 |
| 17.00 | 27.00 | 23.00 | 1.02 | 1.00 |
| 296.00 | 317.00 | 319.67 | 1.01 | 0.94 |
| 9.00 | 9.00 | 8.67 | 1.16 | 0.67 |
| 2.00 | 1.00 | 2.67 | -2.95 | 0.01 |
| 7.00 | 16.00 | 12.67 | 2.06 | 0.01 |
| 355.00 | 414.00 | 413.67 | 1.00 | 0.98 |
| 4.00 | 9.00 | 7.33 | -1.74 | 0.06 |
| 8.00 | 11.00 | 10.00 | 1.02 | 1.00 |
| 41.00 | 53.00 | 44.00 | 1.09 | 0.56 |
| 5.00 | 7.00 | 8.00 | -1.47 | 0.21 |
| 58.00 | 64.00 | 62.67 | -1.27 | 0.02 |
| 5.00 | 0.00 | 1.67 | 1.59 | 0.75 |
| 0.00 | 0.00 | 0.00 | -14.45 | 0.07 |
| 17.00 | 14.00 | 13.00 | -1.35 | 0.23 |

| | | | | |
|---|---|---|---|---|
| 32.00 | 22.00 | 30.67 | -1.93 | 0.00 |
| 115.00 | 136.00 | 120.00 | 1.37 | 0.00 |
| 3.00 | 5.00 | 3.67 | -1.55 | 0.35 |
| 10.00 | 7.00 | 7.33 | -1.69 | 0.08 |
| 37.00 | 28.00 | 31.67 | -1.07 | 0.74 |
| 2.00 | 1.00 | 1.67 | -1.39 | 0.78 |
| 6.00 | 4.00 | 5.33 | 1.31 | 0.58 |
| 8.00 | 8.00 | 7.33 | -1.60 | 0.12 |
| 70.00 | 58.00 | 68.00 | 1.05 | 0.68 |
| 4.00 | 1.00 | 3.33 | -1.40 | 0.55 |
| 57.00 | 55.00 | 60.00 | 1.11 | 0.39 |
| 1.00 | 3.00 | 2.33 | -1.70 | 0.37 |
| 112.00 | 127.00 | 125.00 | 1.01 | 0.91 |
| 58.00 | 46.00 | 54.00 | -1.12 | 0.37 |
| 61.00 | 85.00 | 77.33 | 1.31 | 0.02 |
| 505.00 | 588.00 | 586.33 | 1.25 | 0.00 |
| 1122.00 | 1202.00 | 1182.33 | -1.34 | 0.00 |
| 38.00 | 24.00 | 34.67 | -1.22 | 0.22 |
| 47.00 | 83.00 | 62.33 | -1.00 | 1.00 |
| 3.00 | 0.00 | 2.00 | -4.82 | 0.00 |
| 10.00 | 5.00 | 7.33 | -1.06 | 1.00 |
| 1.00 | 2.00 | 2.33 | -1.01 | 1.00 |
| 205.00 | 179.00 | 203.67 | -1.19 | 0.02 |
| 655.00 | 734.00 | 740.67 | -1.15 | 0.01 |
| 10.00 | 10.00 | 10.33 | -1.11 | 0.81 |
| 957.00 | 1054.00 | 1019.33 | -1.20 | 0.00 |
| 5.00 | 2.00 | 3.67 | -2.35 | 0.02 |
| 30.00 | 42.00 | 37.00 | -1.49 | 0.01 |
| 218.00 | 263.00 | 244.67 | -1.05 | 0.53 |
| 3339.00 | 3524.00 | 3500.67 | -1.08 | 0.03 |
| 18.00 | 19.00 | 20.33 | 1.09 | 0.72 |
| 14.00 | 15.00 | 17.00 | 1.12 | 0.69 |
| 245.00 | 275.00 | 270.00 | -1.05 | 0.42 |
| 19.00 | 24.00 | 25.33 | 2.32 | 0.00 |
| 50.00 | 54.00 | 61.00 | 1.29 | 0.05 |
| 1115.00 | 1095.00 | 1134.33 | 1.08 | 0.06 |
| 1206.00 | 1186.00 | 1246.33 | -1.19 | 0.00 |
| 23.00 | 27.00 | 26.67 | -1.12 | 0.54 |
| 534.00 | 625.00 | 593.67 | 1.16 | 0.00 |
| 5.00 | 12.00 | 10.33 | -1.18 | 0.64 |
| 6175.00 | 6519.00 | 6223.00 | -1.06 | 0.21 |
| 4370.00 | 4244.00 | 4425.00 | -1.37 | 0.00 |
| 767.00 | 776.00 | 784.00 | -1.12 | 0.02 |
| 1257.00 | 1303.00 | 1249.67 | 1.05 | 0.26 |
| 995.00 | 970.00 | 1026.00 | -1.25 | 0.00 |
| 586.00 | 685.00 | 658.67 | -1.04 | 0.36 |
| 311.00 | 338.00 | 283.00 | -1.11 | 0.38 |
| 50.00 | 66.00 | 58.33 | -1.14 | 0.31 |

| | | | | |
|---|---|---|---|---|
| 1761.00 | 1794.00 | 1842.67 | -1.10 | 0.02 |
| 731.00 | 795.00 | 780.67 | -1.11 | 0.02 |
| 23.00 | 17.00 | 20.00 | 1.18 | 0.41 |
| 3624.00 | 3908.00 | 3831.33 | 1.10 | 0.01 |
| 1249.00 | 1403.00 | 1313.00 | 1.54 | 0.00 |
| 6471.00 | 6017.00 | 6278.33 | -1.14 | 0.00 |
| 1.00 | 1.00 | 1.00 | -2.51 | 0.23 |
| 1875.00 | 1989.00 | 2047.33 | 1.21 | 0.00 |
| 901.00 | 1008.00 | 952.00 | -1.09 | 0.04 |
| 3.00 | 2.00 | 2.00 | -1.17 | 1.00 |
| 1013.00 | 1047.00 | 973.33 | -1.23 | 0.00 |
| 142.00 | 165.00 | 153.33 | 1.36 | 0.00 |
| 286.00 | 259.00 | 250.33 | -1.40 | 0.00 |
| 26.00 | 53.00 | 40.67 | 1.14 | 0.46 |
| 132.00 | 213.00 | 168.00 | -1.09 | 0.42 |
| 3.00 | 2.00 | 2.33 | 2.16 | 0.35 |
| 195.00 | 179.00 | 192.00 | -1.01 | 0.91 |
| 7.00 | 7.00 | 9.00 | 2.02 | 0.04 |
| 6.00 | 8.00 | 7.33 | 1.14 | 0.76 |
| 5.00 | 10.00 | 6.00 | -2.06 | 0.02 |
| 44.00 | 58.00 | 56.67 | -1.23 | 0.09 |
| 36.00 | 35.00 | 38.33 | -1.02 | 0.90 |
| 2.00 | 8.00 | 5.00 | -1.54 | 0.28 |
| 37.00 | 47.00 | 47.00 | -1.61 | 0.00 |
| 8.00 | 12.00 | 10.00 | -1.48 | 0.14 |
| 0.00 | 3.00 | 1.67 | -1.01 | 1.00 |
| 6.00 | 3.00 | 3.67 | -2.79 | 0.00 |
| 16.00 | 22.00 | 16.67 | -1.37 | 0.14 |
| 1389.00 | 1281.00 | 1333.00 | -1.38 | 0.00 |
| 14.00 | 17.00 | 15.33 | -1.73 | 0.01 |
| 1.00 | 2.00 | 1.33 | -2.40 | 0.19 |
| 72.00 | 56.00 | 62.33 | -1.36 | 0.01 |
| 1.00 | 2.00 | 3.33 | 1.38 | 0.64 |
| 73.00 | 75.00 | 75.00 | -1.27 | 0.02 |
| 326.00 | 302.00 | 311.67 | -1.21 | 0.00 |
| 34.00 | 32.00 | 32.33 | 1.18 | 0.35 |
| 78.00 | 48.00 | 53.00 | -1.71 | 0.00 |
| 3.00 | 6.00 | 3.67 | -1.37 | 0.56 |
| 87.00 | 89.00 | 89.33 | -1.41 | 0.00 |
| 3.00 | 2.00 | 2.33 | -2.27 | 0.15 |
| 167.00 | 199.00 | 189.00 | -1.07 | 0.32 |
| 25.00 | 27.00 | 25.33 | -1.41 | 0.03 |
| 244.00 | 235.00 | 230.67 | -1.05 | 0.53 |
| 68.00 | 74.00 | 67.67 | -1.00 | 1.00 |
| 22.00 | 31.00 | 30.33 | 1.02 | 1.00 |
| 26.00 | 28.00 | 25.33 | -1.04 | 0.94 |
| 1475.00 | 1578.00 | 1613.67 | -1.26 | 0.00 |
| 14.00 | 8.00 | 11.67 | 1.02 | 1.00 |

| | | | | |
|---|---|---|---|---|
| 17.00 | 10.00 | 12.00 | -1.12 | 0.74 |
| 21.00 | 13.00 | 20.00 | -1.06 | 0.73 |
| 1.00 | 1.00 | 1.67 | -2.52 | 0.11 |
| 0.00 | 0.00 | 0.00 | -9.06 | 0.27 |
| 76.00 | 82.00 | 80.00 | -1.07 | 0.54 |
| 62.00 | 83.00 | 74.67 | -1.14 | 0.21 |
| 2.00 | 5.00 | 3.67 | -2.70 | 0.01 |
| 7.00 | 5.00 | 7.00 | -2.01 | 0.01 |
| 37.00 | 32.00 | 36.33 | -1.13 | 0.39 |
| 18.00 | 13.00 | 15.00 | 1.03 | 0.92 |
| 27.00 | 30.00 | 23.33 | -1.66 | 0.00 |
| 150.00 | 195.00 | 190.00 | -1.31 | 0.00 |
| 193.00 | 212.00 | 224.33 | 1.02 | 0.80 |
| 22.00 | 24.00 | 24.67 | 1.06 | 0.81 |
| 100.00 | 125.00 | 114.00 | -1.09 | 0.30 |
| 59.00 | 76.00 | 61.00 | -1.15 | 0.27 |
| 61.00 | 51.00 | 65.00 | 1.04 | 0.83 |
| 32.00 | 53.00 | 40.33 | -2.32 | 0.00 |
| 73.00 | 56.00 | 76.33 | -1.24 | 0.10 |
| 4.00 | 6.00 | 4.67 | -1.58 | 0.25 |
| 5.00 | 15.00 | 8.33 | 1.44 | 0.31 |
| 39.00 | 37.00 | 35.33 | -1.59 | 0.00 |
| 302.00 | 310.00 | 330.00 | -1.32 | 0.00 |
| 32.00 | 31.00 | 31.00 | 1.76 | 0.00 |
| 86.00 | 92.00 | 90.00 | 1.24 | 0.03 |
| 15.00 | 23.00 | 18.33 | 1.01 | 1.00 |
| 21.00 | 34.00 | 28.33 | -1.19 | 0.30 |
| 30.00 | 42.00 | 35.33 | 1.34 | 0.08 |
| 5.00 | 19.00 | 10.00 | 1.28 | 0.48 |
| 75.00 | 95.00 | 95.33 | -1.01 | 0.98 |
| 6.00 | 14.00 | 12.33 | -1.07 | 0.83 |
| 3.00 | 1.00 | 1.67 | -1.95 | 0.31 |
| 17.00 | 14.00 | 13.67 | -1.18 | 0.60 |
| 33.00 | 33.00 | 37.33 | 1.41 | 0.03 |
| 26.00 | 32.00 | 29.00 | -1.32 | 0.07 |
| 0.00 | 2.00 | 1.00 | -3.71 | 0.04 |
| 21.00 | 37.00 | 32.33 | -1.39 | 0.03 |
| 8.00 | 11.00 | 15.00 | 1.38 | 0.26 |
| 14.00 | 17.00 | 16.00 | 1.52 | 0.08 |
| 52.00 | 53.00 | 49.67 | -1.12 | 0.41 |
| 70.00 | 84.00 | 87.67 | 1.12 | 0.37 |
| 80.00 | 79.00 | 79.00 | 1.29 | 0.02 |
| 56.00 | 32.00 | 47.33 | -1.12 | 0.43 |
| 2.00 | 4.00 | 3.00 | -1.55 | 0.41 |
| 64.00 | 73.00 | 68.00 | 1.31 | 0.02 |
| 37.00 | 35.00 | 35.67 | -1.12 | 0.45 |
| 5.00 | 11.00 | 7.33 | -2.37 | 0.00 |
| 36.00 | 26.00 | 28.00 | -1.17 | 0.38 |

| | | | | |
|---|---|---|---|---|
| 3.00 | 6.00 | 3.67 | -1.46 | 0.45 |
| 6.00 | 4.00 | 6.33 | -1.33 | 0.47 |
| 13.00 | 22.00 | 17.00 | 1.43 | 0.13 |
| 35.00 | 25.00 | 26.00 | 1.15 | 0.45 |
| 3.00 | 1.00 | 1.67 | -2.14 | 0.22 |
| 3.00 | 4.00 | 4.00 | -1.01 | 1.00 |
| 20.00 | 27.00 | 25.33 | -1.08 | 0.68 |
| 30.00 | 28.00 | 32.00 | 1.39 | 0.06 |
| 35.00 | 39.00 | 40.67 | -1.14 | 0.35 |
| 8.00 | 1.00 | 3.00 | 1.74 | 0.50 |
| 309.00 | 342.00 | 309.33 | -1.17 | 0.04 |
| 30.00 | 27.00 | 29.33 | 1.21 | 0.29 |
| 90.00 | 121.00 | 104.67 | 1.93 | 0.00 |
| 0.00 | 2.00 | 0.67 | -7.82 | 0.00 |
| 29.00 | 32.00 | 30.67 | 1.33 | 0.11 |
| 1.00 | 0.00 | 1.00 | -7.01 | 0.00 |
| 169.00 | 195.00 | 195.33 | -1.12 | 0.12 |
| 3.00 | 3.00 | 5.33 | 1.30 | 0.58 |
| 31.00 | 40.00 | 41.00 | 1.01 | 1.00 |
| 47.00 | 47.00 | 48.00 | -1.10 | 0.45 |
| 2515.00 | 2887.00 | 2776.67 | -1.06 | 0.12 |
| 234.00 | 212.00 | 233.67 | -1.12 | 0.15 |
| 29.00 | 64.00 | 42.33 | -1.07 | 0.72 |
| 190.00 | 149.00 | 165.00 | -1.18 | 0.06 |
| 79.00 | 74.00 | 72.33 | 1.03 | 0.80 |
| 37.00 | 29.00 | 36.00 | -1.32 | 0.05 |
| 13.00 | 20.00 | 15.00 | 1.70 | 0.04 |
| 20.00 | 25.00 | 20.33 | 1.00 | 1.00 |
| 135.00 | 160.00 | 155.00 | 1.08 | 0.39 |
| 24.00 | 31.00 | 29.33 | -2.12 | 0.00 |
| 30.00 | 25.00 | 29.33 | -1.23 | 0.17 |
| 7.00 | 5.00 | 5.00 | -2.00 | 0.04 |
| 81.00 | 87.00 | 95.00 | 1.01 | 0.95 |
| 1244.00 | 1237.00 | 1277.00 | -1.44 | 0.00 |
| 5.00 | 6.00 | 6.00 | 1.46 | 0.37 |
| 13.00 | 21.00 | 18.00 | -1.48 | 0.04 |
| 9.00 | 7.00 | 10.67 | 1.16 | 0.64 |
| 28.00 | 21.00 | 27.00 | -1.33 | 0.07 |
| 27.00 | 45.00 | 31.67 | -1.27 | 0.14 |
| 193.00 | 172.00 | 204.00 | -1.08 | 0.36 |
| 14.00 | 18.00 | 14.00 | -2.21 | 0.02 |
| 79.00 | 87.00 | 92.33 | 1.01 | 0.94 |
| 44.00 | 20.00 | 29.33 | -1.13 | 0.53 |
| 51.00 | 65.00 | 69.00 | 1.03 | 0.84 |
| 76.00 | 97.00 | 94.67 | -1.08 | 0.48 |
| 50.00 | 46.00 | 47.00 | 1.12 | 0.42 |
| 10.00 | 15.00 | 10.67 | -1.39 | 0.23 |
| 9.00 | 14.00 | 15.67 | -1.59 | 0.02 |

| | | | | |
|---|---|---|---|---|
| 278.00 | 275.00 | 287.00 | -1.02 | 0.69 |
| 10.00 | 5.00 | 11.33 | 1.66 | 0.10 |
| 25.00 | 28.00 | 23.67 | 1.07 | 0.77 |
| 5.00 | 7.00 | 9.33 | -1.19 | 0.63 |
| 34.00 | 47.00 | 42.67 | 1.23 | 0.15 |
| 25.00 | 22.00 | 20.67 | -1.25 | 0.28 |
| 2.00 | 5.00 | 3.00 | 1.26 | 0.81 |
| 131.00 | 111.00 | 112.33 | 1.37 | 0.00 |
| 3.00 | 0.00 | 1.00 | -1.61 | 0.74 |
| 14.00 | 10.00 | 15.00 | -1.08 | 0.85 |
| 12.00 | 5.00 | 9.67 | 1.19 | 0.59 |
| 3.00 | 5.00 | 4.33 | -2.45 | 0.01 |
| 27.00 | 21.00 | 23.33 | -1.24 | 0.27 |
| 2464.00 | 2164.00 | 2367.33 | -1.16 | 0.00 |
| 8.00 | 14.00 | 13.67 | 1.34 | 0.30 |
| 3.00 | 5.00 | 5.33 | 1.56 | 0.34 |
| 2.00 | 8.00 | 6.67 | 1.31 | 0.51 |
| 26.00 | 21.00 | 23.67 | -1.14 | 0.49 |
| 4.00 | 7.00 | 7.00 | -1.54 | 0.18 |
| 43.00 | 45.00 | 41.67 | -1.11 | 0.45 |
| 86.00 | 75.00 | 87.33 | -1.13 | 0.23 |
| 4.00 | 6.00 | 5.00 | -1.54 | 0.27 |
| 197.00 | 183.00 | 212.33 | 1.19 | 0.04 |
| 463.00 | 579.00 | 510.00 | -1.24 | 0.00 |
| 208.00 | 258.00 | 228.33 | -1.41 | 0.00 |
| 1.00 | 2.00 | 1.00 | -2.21 | 0.36 |
| 95.00 | 113.00 | 108.33 | -1.22 | 0.02 |
| 6.00 | 8.00 | 6.67 | 1.23 | 0.62 |
| 11.00 | 26.00 | 18.33 | -1.51 | 0.03 |
| 1.00 | 5.00 | 2.00 | -5.46 | 0.00 |
| 1.00 | 1.00 | 2.33 | 1.66 | 0.56 |
| 4.00 | 5.00 | 4.00 | -1.34 | 0.58 |
| 116.00 | 135.00 | 119.67 | -1.01 | 0.98 |
| 9.00 | 5.00 | 6.67 | -1.26 | 0.56 |
| 59.00 | 43.00 | 50.33 | -1.27 | 0.06 |
| 2.00 | 1.00 | 1.67 | -2.52 | 0.10 |
| 23.00 | 33.00 | 23.33 | -1.04 | 0.95 |
| 16.00 | 41.00 | 32.67 | -2.13 | 0.00 |
| 7.00 | 6.00 | 6.00 | 1.18 | 0.73 |
| 0.00 | 1.00 | 0.33 | -5.42 | 0.07 |
| 10.00 | 7.00 | 10.67 | -1.42 | 0.19 |
| 7.00 | 13.00 | 8.33 | -1.09 | 0.89 |
| 0.00 | 4.00 | 2.67 | -1.98 | 0.16 |
| 61.00 | 66.00 | 64.67 | 1.22 | 0.09 |
| 14.00 | 17.00 | 16.33 | -1.40 | 0.11 |
| 17.00 | 42.00 | 26.00 | -1.39 | 0.08 |
| 1.00 | 5.00 | 5.33 | -1.45 | 0.37 |
| 24.00 | 27.00 | 22.67 | 1.37 | 0.13 |

| | | | | |
|---|---|---|---|---|
| 287.00 | 298.00 | 310.67 | 2.18 | 0.00 |
| 3.00 | 4.00 | 5.00 | -1.21 | 0.73 |
| 48.00 | 47.00 | 48.33 | -1.14 | 0.31 |
| 2.00 | 1.00 | 2.00 | -1.49 | 0.61 |
| 42.00 | 27.00 | 37.00 | 1.37 | 0.06 |
| 2.00 | 1.00 | 3.33 | 1.22 | 0.82 |
| 0.00 | 1.00 | 1.00 | -3.41 | 0.06 |
| 1.00 | 6.00 | 2.67 | -1.98 | 0.18 |
| 10.00 | 14.00 | 15.67 | 1.71 | 0.04 |
| 99.00 | 123.00 | 119.00 | 1.29 | 0.01 |
| 2.00 | 13.00 | 8.00 | 1.03 | 1.00 |
| 5.00 | 1.00 | 2.67 | -1.25 | 0.82 |
| 32.00 | 41.00 | 39.00 | -1.14 | 0.38 |
| 12.00 | 15.00 | 10.67 | 1.26 | 0.45 |
| 16.00 | 8.00 | 12.33 | -1.96 | 0.00 |
| 21.00 | 13.00 | 15.00 | 1.23 | 0.40 |
| 29.00 | 24.00 | 27.33 | 1.87 | 0.00 |
| 29.00 | 18.00 | 27.67 | -1.37 | 0.06 |
| 68.00 | 90.00 | 79.00 | 1.34 | 0.01 |
| 1.00 | 2.00 | 3.67 | -1.02 | 1.00 |
| 24.00 | 18.00 | 21.67 | 1.07 | 0.73 |
| 3.00 | 3.00 | 6.00 | -1.29 | 0.62 |
| 45.00 | 61.00 | 48.00 | -1.01 | 0.96 |
| 20.00 | 11.00 | 16.33 | 1.01 | 1.00 |
| 99.00 | 114.00 | 109.33 | -1.12 | 0.20 |
| 155.00 | 137.00 | 148.00 | -1.01 | 0.94 |
| 10.00 | 15.00 | 12.33 | 1.40 | 0.22 |
| 4.00 | 8.00 | 6.67 | 1.31 | 0.51 |
| 13.00 | 27.00 | 18.33 | -1.38 | 0.12 |
| 13.00 | 12.00 | 13.67 | -1.16 | 0.60 |
| 35.00 | 43.00 | 38.67 | 1.33 | 0.07 |
| 9.00 | 10.00 | 11.00 | 1.47 | 0.19 |
| 3.00 | 8.00 | 4.33 | -2.07 | 0.05 |
| 4.00 | 8.00 | 6.33 | -1.17 | 0.76 |
| 4.00 | 3.00 | 3.33 | -1.99 | 0.10 |
| 1.00 | 6.00 | 5.00 | -1.21 | 0.73 |
| 1.00 | 0.00 | 0.33 | -3.22 | 0.38 |
| 4.00 | 1.00 | 3.67 | -1.37 | 0.57 |
| 9.00 | 6.00 | 9.33 | 1.02 | 1.00 |
| 4.00 | 2.00 | 2.67 | -1.13 | 1.00 |
| 126.00 | 139.00 | 146.33 | -1.07 | 0.43 |
| 1.00 | 3.00 | 3.00 | 1.45 | 0.61 |
| 24.00 | 26.00 | 20.33 | 1.26 | 0.30 |
| 15.00 | 7.00 | 10.33 | -1.36 | 0.34 |
| 3.00 | 2.00 | 2.33 | -1.56 | 0.49 |
| 44.00 | 60.00 | 54.67 | 1.31 | 0.04 |
| 5.00 | 16.00 | 9.33 | 1.45 | 0.27 |
| 10.00 | 8.00 | 11.33 | -1.81 | 0.01 |

| | | | | |
|---|---|---|---|---|
| 0.00 | 1.00 | 0.33 | -3.22 | 0.39 |
| 1.00 | 4.00 | 4.00 | 1.18 | 0.84 |
| 9.00 | 8.00 | 7.67 | 1.13 | 0.77 |
| 33.00 | 31.00 | 28.00 | -1.20 | 0.30 |
| 7.00 | 1.00 | 4.00 | -1.59 | 0.29 |
| 3.00 | 1.00 | 1.67 | -1.76 | 0.43 |
| 28.00 | 22.00 | 22.33 | -1.37 | 0.07 |
| 4.00 | 8.00 | 4.33 | -1.46 | 0.40 |
| 12.00 | 8.00 | 10.00 | -1.81 | 0.01 |
| 4.00 | 6.00 | 4.00 | -1.66 | 0.23 |
| 24.00 | 32.00 | 31.33 | 1.34 | 0.11 |
| 18.00 | 16.00 | 17.67 | -1.30 | 0.22 |
| 29.00 | 17.00 | 24.33 | 1.56 | 0.02 |
| 682.00 | 746.00 | 737.00 | 1.07 | 0.16 |
| 0.00 | 2.00 | 0.67 | -1.44 | 1.00 |
| 4.00 | 9.00 | 7.00 | -1.11 | 0.88 |
| 31.00 | 20.00 | 27.67 | -1.35 | 0.06 |
| 34.00 | 27.00 | 30.67 | -1.22 | 0.18 |
| 7.00 | 14.00 | 8.67 | -1.09 | 0.90 |
| 30.00 | 32.00 | 31.33 | -1.14 | 0.39 |
| 82.00 | 62.00 | 76.00 | 2.22 | 0.00 |
| 0.00 | 0.00 | 0.33 | -3.96 | 0.23 |
| 0.00 | 5.00 | 2.67 | -1.14 | 1.00 |
| 1.00 | 1.00 | 3.33 | 1.38 | 0.65 |
| 10.00 | 5.00 | 8.33 | 1.63 | 0.17 |
| 2.00 | 5.00 | 4.00 | -1.26 | 0.71 |
| 11.00 | 6.00 | 7.67 | -1.32 | 0.42 |
| 21.00 | 19.00 | 20.00 | -3.21 | 0.00 |
| 195.00 | 208.00 | 209.67 | -1.14 | 0.04 |
| 11.00 | 2.00 | 6.00 | -1.34 | 0.48 |
| 41.00 | 59.00 | 53.67 | 1.01 | 1.00 |
| 20.00 | 23.00 | 22.67 | 1.03 | 1.00 |
| 4.00 | 8.00 | 4.33 | -1.92 | 0.08 |
| 10.00 | 6.00 | 8.33 | 1.07 | 0.89 |
| 68.00 | 75.00 | 71.67 | -1.28 | 0.02 |
| 6.00 | 3.00 | 6.00 | -1.73 | 0.09 |
| 22.00 | 26.00 | 23.00 | 1.10 | 0.68 |
| 29.00 | 19.00 | 25.00 | 1.16 | 0.47 |
| 20.00 | 23.00 | 22.33 | 1.61 | 0.02 |
| 371.00 | 455.00 | 430.00 | 1.10 | 0.12 |
| 25.00 | 16.00 | 21.33 | -1.61 | 0.01 |
| 0.00 | 7.00 | 4.00 | 1.30 | 0.68 |
| 3.00 | 12.00 | 7.33 | -1.06 | 1.00 |
| 9.00 | 9.00 | 7.33 | 1.54 | 0.25 |
| 19.00 | 10.00 | 15.33 | -1.58 | 0.03 |
| 6.00 | 8.00 | 6.33 | -1.12 | 0.88 |
| 2.00 | 5.00 | 2.33 | -2.11 | 0.14 |
| 1.00 | 0.00 | 1.33 | -1.01 | 1.00 |

| | | | | |
|---|---|---|---|---|
| 9.00 | 12.00 | 9.67 | 2.17 | 0.02 |
| 182.00 | 227.00 | 206.33 | 1.42 | 0.00 |
| 44.00 | 48.00 | 54.33 | 1.06 | 0.73 |
| 19.00 | 29.00 | 25.00 | -1.08 | 0.65 |
| 2.00 | 4.00 | 3.33 | -1.50 | 0.43 |
| 24.00 | 19.00 | 19.67 | 1.00 | 1.00 |
| 7.00 | 8.00 | 8.67 | 2.78 | 0.01 |
| 216.00 | 258.00 | 237.00 | 1.04 | 0.62 |
| 428.00 | 433.00 | 448.00 | -1.65 | 0.00 |
| 1.00 | 2.00 | 1.67 | -1.96 | 0.31 |
| 26.00 | 25.00 | 24.67 | -1.43 | 0.04 |
| 334.00 | 375.00 | 360.33 | -1.22 | 0.00 |
| 186.00 | 189.00 | 202.33 | -1.41 | 0.00 |
| 74.00 | 93.00 | 88.67 | 1.13 | 0.26 |
| 1.00 | 2.00 | 2.00 | -1.33 | 0.79 |
| 9.00 | 17.00 | 14.67 | 1.24 | 0.46 |
| 1.00 | 9.00 | 6.33 | -1.85 | 0.05 |
| 3.00 | 1.00 | 3.00 | -1.34 | 0.67 |
| 16.00 | 16.00 | 17.33 | -1.09 | 0.78 |
| 14.00 | 17.00 | 14.33 | 1.28 | 0.32 |
| 7.00 | 7.00 | 7.33 | -1.38 | 0.34 |
| 304.00 | 300.00 | 315.67 | -1.32 | 0.00 |
| 3.00 | 3.00 | 4.67 | -1.30 | 0.61 |
| 7.00 | 4.00 | 4.67 | -1.30 | 0.61 |
| 7.00 | 10.00 | 8.33 | -1.17 | 0.69 |
| 61.00 | 44.00 | 54.00 | 1.13 | 0.38 |
| 1691.00 | 1900.00 | 1907.00 | -1.12 | 0.01 |
| 565.00 | 643.00 | 635.33 | -1.04 | 0.38 |
| 850.00 | 875.00 | 887.00 | -1.13 | 0.00 |
| 151.00 | 190.00 | 171.67 | -1.01 | 0.86 |
| 84.00 | 69.00 | 76.00 | -1.33 | 0.00 |
| 6.00 | 6.00 | 4.67 | 1.25 | 0.70 |
| 811.00 | 759.00 | 784.33 | -1.44 | 0.00 |
| 171.00 | 183.00 | 192.33 | -1.13 | 0.10 |
| 71.00 | 56.00 | 62.67 | -1.33 | 0.01 |
| 434.00 | 490.00 | 463.67 | -1.17 | 0.00 |
| 44.00 | 57.00 | 61.67 | -1.13 | 0.35 |
| 26.00 | 47.00 | 37.67 | 2.70 | 0.00 |
| 209.00 | 227.00 | 229.67 | 1.06 | 0.44 |
| 3.00 | 12.00 | 6.33 | 1.68 | 0.22 |
| 10.00 | 7.00 | 7.67 | -1.97 | 0.01 |
| 12.00 | 16.00 | 13.67 | -1.06 | 0.92 |
| 3.00 | 2.00 | 1.67 | -1.58 | 0.59 |
| 2.00 | 4.00 | 3.00 | -2.53 | 0.02 |
| 8.00 | 9.00 | 8.00 | 1.24 | 0.55 |
| 391.00 | 391.00 | 408.00 | -1.12 | 0.05 |
| 146.00 | 133.00 | 135.67 | -1.14 | 0.12 |
| 30.00 | 43.00 | 35.33 | 1.26 | 0.17 |

| | | | | |
|---|---|---|---|---|
| 181.00 | 169.00 | 186.00 | 1.37 | 0.00 |
| 88.00 | 100.00 | 104.67 | -1.80 | 0.00 |
| 656.00 | 725.00 | 720.33 | -1.07 | 0.17 |
| 378.00 | 461.00 | 426.33 | -1.23 | 0.00 |
| 10.00 | 7.00 | 10.67 | -1.36 | 0.28 |
| 0.00 | 0.00 | 0.00 | -17.09 | 0.03 |
| 162.00 | 196.00 | 171.67 | -1.15 | 0.10 |
| 15.00 | 18.00 | 16.33 | 1.05 | 0.84 |
| 25.00 | 19.00 | 22.33 | -1.10 | 0.71 |
| 217.00 | 198.00 | 206.00 | -1.09 | 0.30 |
| 307.00 | 286.00 | 304.33 | 1.19 | 0.01 |
| 6.00 | 15.00 | 11.33 | 1.19 | 0.54 |
| 97.00 | 110.00 | 101.67 | -1.03 | 0.79 |
| 280.00 | 279.00 | 300.00 | -1.06 | 0.35 |
| 229.00 | 230.00 | 239.00 | -1.19 | 0.02 |
| 1.00 | 4.00 | 2.33 | -1.01 | 1.00 |
| 6.00 | 9.00 | 6.67 | -1.46 | 0.26 |
| 77.00 | 87.00 | 78.33 | 1.20 | 0.10 |
| 41.00 | 34.00 | 38.33 | -1.23 | 0.19 |
| 48.00 | 45.00 | 48.33 | -1.12 | 0.36 |
| 95.00 | 121.00 | 128.00 | -1.01 | 0.90 |
| 374.00 | 395.00 | 379.00 | -1.14 | 0.02 |
| 136.00 | 150.00 | 148.00 | 1.13 | 0.11 |
| 11.00 | 8.00 | 9.00 | -1.05 | 1.00 |
| 31.00 | 18.00 | 27.00 | -1.11 | 0.58 |
| 383.00 | 478.00 | 426.33 | 1.35 | 0.00 |
| 514.00 | 503.00 | 511.67 | -1.23 | 0.00 |
| 53.00 | 52.00 | 58.67 | 1.17 | 0.27 |
| 0.00 | 0.00 | 0.00 | -9.06 | 0.27 |
| 6.00 | 4.00 | 5.00 | -2.53 | 0.00 |
| 11.00 | 14.00 | 11.33 | 1.29 | 0.38 |
| 6.00 | 5.00 | 6.00 | -1.95 | 0.03 |
| 305.00 | 326.00 | 315.67 | -1.10 | 0.11 |
| 8.00 | 3.00 | 5.00 | -1.14 | 0.86 |
| 11.00 | 5.00 | 9.67 | 1.19 | 0.59 |
| 43.00 | 62.00 | 49.00 | -1.07 | 0.71 |
| 6.00 | 9.00 | 8.33 | -1.57 | 0.14 |
| 34.00 | 35.00 | 30.00 | 1.31 | 0.12 |
| 24.00 | 21.00 | 20.67 | -1.16 | 0.46 |
| 9.00 | 9.00 | 9.00 | 1.11 | 0.78 |
| 11.00 | 17.00 | 15.00 | 1.34 | 0.23 |
| 3.00 | 4.00 | 5.00 | 1.81 | 0.22 |
| 34.00 | 29.00 | 36.67 | 1.20 | 0.27 |
| 137.00 | 150.00 | 143.67 | 1.01 | 0.91 |
| 47.00 | 58.00 | 47.00 | -1.04 | 0.79 |
| 6.00 | 6.00 | 8.00 | 1.38 | 0.36 |
| 0.00 | 0.00 | 0.67 | -2.29 | 0.47 |
| 4.00 | 6.00 | 5.33 | -1.57 | 0.23 |

| | | | | |
|---|---|---|---|---|
| 18.00 | 22.00 | 24.33 | -1.28 | 0.14 |
| 5.00 | 32.00 | 40.00 | 2.31 | 0.02 |
| 38.00 | 50.00 | 46.33 | 1.22 | 0.16 |
| 31.00 | 38.00 | 30.33 | -1.15 | 0.43 |
| 15.00 | 30.00 | 21.33 | 1.15 | 0.55 |
| 637.00 | 729.00 | 705.33 | -1.05 | 0.29 |
| 1968.00 | 1965.00 | 1966.67 | -1.26 | 0.00 |
| 17.00 | 14.00 | 18.67 | 1.15 | 0.57 |
| 7.00 | 2.00 | 4.00 | -1.75 | 0.19 |
| 1.00 | 3.00 | 2.67 | 1.12 | 1.00 |
| 10.00 | 18.00 | 15.33 | -2.39 | 0.00 |
| 63.00 | 85.00 | 84.67 | 1.11 | 0.41 |
| 57.00 | 50.00 | 62.00 | -1.33 | 0.01 |
| 104.00 | 80.00 | 93.67 | -1.14 | 0.19 |
| 1.00 | 1.00 | 0.67 | -4.00 | 0.07 |
| 34.00 | 24.00 | 36.67 | 1.27 | 0.18 |
| 340.00 | 350.00 | 369.00 | -1.10 | 0.11 |
| 55.00 | 74.00 | 69.67 | 1.29 | 0.04 |
| 224.00 | 256.00 | 243.33 | -1.04 | 0.50 |
| 86.00 | 90.00 | 84.33 | -1.25 | 0.03 |
| 86.00 | 73.00 | 93.33 | -1.08 | 0.50 |
| 20.00 | 25.00 | 23.33 | 2.15 | 0.00 |
| 72.00 | 73.00 | 78.33 | 1.21 | 0.09 |
| 62.00 | 42.00 | 56.67 | 1.04 | 0.78 |
| 16.00 | 17.00 | 17.33 | -1.30 | 0.21 |
| 118.00 | 106.00 | 125.00 | -1.16 | 0.09 |
| 33.00 | 25.00 | 28.33 | -1.18 | 0.33 |
| 0.00 | 3.00 | 1.67 | -1.58 | 0.59 |
| 103.00 | 140.00 | 129.33 | 1.25 | 0.02 |
| 16.00 | 14.00 | 16.00 | 1.21 | 0.40 |
| 4.00 | 3.00 | 2.67 | -1.86 | 0.22 |
| 17.00 | 31.00 | 25.33 | -1.25 | 0.17 |
| 362.00 | 393.00 | 400.00 | -1.17 | 0.00 |
| 12.00 | 19.00 | 17.33 | -1.79 | 0.00 |
| 108.00 | 98.00 | 108.00 | -1.16 | 0.09 |
| 104.00 | 94.00 | 105.67 | 1.03 | 0.78 |
| 26.00 | 33.00 | 29.00 | 1.21 | 0.29 |
| 619.00 | 600.00 | 585.33 | -1.54 | 0.00 |
| 8.00 | 1.00 | 6.00 | -1.18 | 0.76 |
| 103.00 | 94.00 | 109.67 | -1.43 | 0.00 |
| 22.00 | 23.00 | 21.00 | -1.29 | 0.18 |
| 0.00 | 0.00 | 0.67 | -2.29 | 0.46 |
| 101.00 | 103.00 | 102.33 | -1.04 | 0.72 |
| 132.00 | 177.00 | 160.67 | 1.22 | 0.02 |
| 271.00 | 296.00 | 286.33 | 1.03 | 0.63 |
| 484.00 | 581.00 | 546.00 | 1.19 | 0.00 |
| 10.00 | 16.00 | 16.00 | 1.00 | 1.00 |
| 23.00 | 21.00 | 28.00 | -1.05 | 0.79 |

| | | | | |
|---|---|---|---|---|
| 1.00 | 3.00 | 3.00 | -2.53 | 0.02 |
| 4.00 | 3.00 | 3.00 | 1.10 | 1.00 |
| 16.00 | 25.00 | 20.33 | -1.03 | 0.86 |
| 260.00 | 289.00 | 283.67 | 1.32 | 0.00 |
| 22.00 | 11.00 | 22.33 | -1.24 | 0.31 |
| 201.00 | 250.00 | 240.00 | 1.10 | 0.21 |
| 257.00 | 311.00 | 298.67 | -1.07 | 0.30 |
| 315.00 | 331.00 | 344.67 | 1.10 | 0.14 |
| 101.00 | 114.00 | 112.33 | -1.05 | 0.64 |
| 7.00 | 13.00 | 11.00 | 2.01 | 0.02 |
| 5052.00 | 5213.00 | 5239.33 | 1.00 | 0.94 |
| 1540.00 | 1562.00 | 1550.00 | -1.08 | 0.05 |
| 540.00 | 574.00 | 579.67 | 1.04 | 0.43 |
| 659.00 | 649.00 | 649.00 | -1.27 | 0.00 |
| 24623.00 | 27771.00 | 27022.67 | 2.42 | 0.00 |
| 2926.00 | 2863.00 | 2904.00 | -1.10 | 0.02 |
| 600.00 | 686.00 | 667.00 | 1.10 | 0.04 |
| 11210.00 | 11607.00 | 11539.00 | -1.23 | 0.00 |
| 347.00 | 376.00 | 354.67 | -1.53 | 0.00 |
| 174.00 | 182.00 | 184.00 | -1.33 | 0.00 |
| 1036.00 | 1052.00 | 1042.00 | -1.15 | 0.00 |
| 136.00 | 125.00 | 141.67 | -1.08 | 0.39 |
| 196.00 | 200.00 | 196.67 | 1.06 | 0.48 |
| 73.00 | 73.00 | 68.00 | -1.16 | 0.20 |
| 50.00 | 44.00 | 44.67 | -1.09 | 0.55 |
| 364.00 | 351.00 | 358.67 | -1.14 | 0.03 |
| 365.00 | 355.00 | 362.33 | -1.29 | 0.00 |
| 1599.00 | 1553.00 | 1612.67 | 1.10 | 0.03 |
| 2371.00 | 2423.00 | 2415.00 | 1.17 | 0.00 |
| 1555.00 | 1589.00 | 1570.00 | 1.10 | 0.02 |
| 3435.00 | 3489.00 | 3466.33 | -1.26 | 0.00 |
| 5.00 | 12.00 | 10.00 | -1.08 | 0.90 |
| 2644.00 | 2749.00 | 2648.33 | -1.02 | 0.69 |
| 39.00 | 52.00 | 48.67 | 1.27 | 0.08 |
| 637.00 | 556.00 | 606.00 | -1.06 | 0.29 |
| 5.00 | 9.00 | 5.00 | -1.08 | 1.00 |
| 14.00 | 8.00 | 11.33 | 2.98 | 0.00 |
| 1259.00 | 1238.00 | 1298.67 | -2.17 | 0.00 |
| 1054.00 | 1069.00 | 1032.00 | -1.09 | 0.08 |
| 719.00 | 687.00 | 729.33 | 1.04 | 0.42 |
| 68.00 | 97.00 | 80.33 | 1.14 | 0.25 |
| 445.00 | 365.00 | 390.00 | -1.17 | 0.04 |
| 984.00 | 910.00 | 983.33 | -1.43 | 0.00 |
| 103.00 | 101.00 | 94.67 | -1.05 | 0.69 |
| 232.00 | 256.00 | 253.67 | 1.01 | 0.92 |
| 2317.00 | 2192.00 | 2351.33 | 1.10 | 0.02 |
| 319.00 | 320.00 | 334.33 | 1.44 | 0.00 |
| 14.00 | 17.00 | 15.00 | -1.24 | 0.39 |

| | | | | |
|---|---|---|---|---|
| 728.00 | 823.00 | 810.00 | 1.12 | 0.01 |
| 626.00 | 636.00 | 636.00 | -1.27 | 0.00 |
| 524.00 | 585.00 | 575.00 | -1.23 | 0.00 |
| 257.00 | 255.00 | 264.67 | 1.09 | 0.19 |
| 9.00 | 14.00 | 13.33 | 1.04 | 0.91 |
| 2718.00 | 2430.00 | 2486.00 | -1.50 | 0.00 |
| 4208.00 | 4016.00 | 4163.00 | -1.29 | 0.00 |
| 1123.00 | 1232.00 | 1176.67 | 1.20 | 0.00 |
| 1314.00 | 1403.00 | 1379.00 | -1.05 | 0.21 |
| 0.00 | 0.00 | 0.00 | -22.49 | 0.01 |
| 14.00 | 20.00 | 15.33 | -1.23 | 0.39 |
| 19.00 | 24.00 | 22.67 | 1.18 | 0.44 |
| 396.00 | 395.00 | 393.67 | -1.27 | 0.00 |
| 4.00 | 7.00 | 6.33 | 1.04 | 1.00 |
| 430.00 | 401.00 | 399.67 | 1.08 | 0.23 |
| 129.00 | 127.00 | 130.00 | -1.15 | 0.10 |
| 1296.00 | 1405.00 | 1326.00 | -1.08 | 0.09 |
| 2178.00 | 2284.00 | 2200.00 | -1.02 | 0.61 |
| 6269.00 | 6239.00 | 6396.33 | -1.06 | 0.16 |
| 4949.00 | 5293.00 | 5130.67 | 1.04 | 0.31 |
| 369.00 | 416.00 | 405.00 | 1.15 | 0.01 |
| 158.00 | 176.00 | 165.67 | 1.04 | 0.64 |
| 1586.00 | 1563.00 | 1670.33 | -1.13 | 0.01 |
| 189.00 | 190.00 | 194.33 | 1.14 | 0.07 |
| 5.00 | 11.00 | 10.33 | 1.13 | 0.70 |
| 1357.00 | 1550.00 | 1464.00 | -1.10 | 0.01 |
| 183.00 | 200.00 | 203.67 | 1.31 | 0.00 |
| 1110.00 | 1116.00 | 1152.00 | 1.07 | 0.10 |
| 82.00 | 88.00 | 85.67 | -1.11 | 0.30 |
| 119.00 | 98.00 | 119.67 | -1.05 | 0.60 |
| 265.00 | 308.00 | 293.33 | -1.02 | 0.70 |
| 98.00 | 120.00 | 113.67 | -1.05 | 0.59 |
| 37.00 | 48.00 | 42.67 | -1.08 | 0.60 |
| 26.00 | 33.00 | 25.33 | 1.34 | 0.15 |
| 184.00 | 202.00 | 196.67 | 1.11 | 0.18 |
| 1.00 | 5.00 | 2.33 | -1.70 | 0.37 |
| 110.00 | 119.00 | 128.00 | -1.05 | 0.63 |
| 26.00 | 19.00 | 23.33 | -1.04 | 0.87 |
| 89.00 | 101.00 | 105.00 | 1.06 | 0.52 |
| 270.00 | 254.00 | 267.67 | -1.12 | 0.06 |
| 24.00 | 22.00 | 23.67 | -1.11 | 0.58 |
| 609.00 | 644.00 | 640.00 | 1.16 | 0.00 |
| 2529.00 | 2607.00 | 2620.00 | -1.13 | 0.00 |
| 717.00 | 767.00 | 757.00 | 1.12 | 0.01 |
| 451.00 | 465.00 | 463.33 | 1.01 | 0.93 |
| 14.00 | 36.00 | 25.67 | 1.54 | 0.08 |
| 1215.00 | 1386.00 | 1328.67 | 1.07 | 0.12 |
| 24.00 | 18.00 | 20.00 | -1.03 | 1.00 |

| | | | | |
|---|---|---|---|---|
| 190.00 | 236.00 | 211.00 | 1.06 | 0.44 |
| 39.00 | 46.00 | 43.67 | -1.39 | 0.01 |
| 348.00 | 380.00 | 380.00 | -1.03 | 0.64 |
| 1696.00 | 1949.00 | 1804.33 | 1.13 | 0.02 |
| 4783.00 | 5250.00 | 5147.00 | 1.08 | 0.02 |
| 48.00 | 61.00 | 55.33 | -1.24 | 0.07 |
| 9.00 | 18.00 | 16.33 | -1.49 | 0.04 |
| 14.00 | 8.00 | 12.33 | 1.01 | 1.00 |
| 49.00 | 74.00 | 55.67 | -1.36 | 0.02 |
| 185.00 | 203.00 | 208.67 | -1.32 | 0.00 |
| 5.00 | 2.00 | 5.67 | 1.65 | 0.26 |
| 14660.00 | 15719.00 | 16475.67 | 1.03 | 0.45 |
| 70.00 | 69.00 | 77.00 | 1.01 | 0.94 |
| 2646.00 | 2504.00 | 2557.33 | -1.35 | 0.00 |
| 446.00 | 428.00 | 425.33 | 1.16 | 0.03 |
| 925.00 | 939.00 | 920.00 | -1.09 | 0.09 |
| 2195.00 | 2232.00 | 2255.67 | -1.02 | 0.55 |
| 487.00 | 514.00 | 526.67 | -1.21 | 0.00 |
| 431.00 | 498.00 | 474.00 | 1.41 | 0.00 |
| 1.00 | 3.00 | 2.67 | 1.30 | 0.79 |
| 2947.00 | 3178.00 | 3019.67 | 1.03 | 0.47 |
| 1791.00 | 2092.00 | 1987.67 | 1.38 | 0.00 |
| 6.00 | 10.00 | 7.00 | 1.15 | 0.76 |
| 24.00 | 13.00 | 23.33 | 1.27 | 0.33 |
| 5.00 | 5.00 | 7.00 | 1.47 | 0.32 |
| 1104.00 | 1070.00 | 1079.33 | -1.25 | 0.00 |
| 98.00 | 128.00 | 117.00 | 1.08 | 0.44 |
| 9.00 | 6.00 | 8.33 | -1.01 | 1.00 |
| 0.00 | 2.00 | 0.67 | -2.29 | 0.49 |
| 811.00 | 851.00 | 867.67 | -1.14 | 0.00 |
| 2.00 | 1.00 | 2.00 | -2.44 | 0.08 |
| 128.00 | 153.00 | 141.33 | 1.42 | 0.00 |
| 762.00 | 757.00 | 781.67 | 1.08 | 0.09 |
| 0.00 | 1.00 | 1.67 | -2.90 | 0.05 |
| 2063.00 | 2215.00 | 2170.33 | -1.06 | 0.10 |
| 530.00 | 581.00 | 608.67 | 1.20 | 0.00 |
| 459.00 | 470.00 | 516.33 | 1.09 | 0.21 |
| 287.00 | 331.00 | 303.33 | -1.32 | 0.00 |
| 242.00 | 289.00 | 276.00 | 1.23 | 0.01 |
| 30.00 | 35.00 | 30.67 | 1.07 | 0.78 |
| 1535.00 | 1571.00 | 1619.67 | -1.02 | 0.57 |
| 417.00 | 470.00 | 467.00 | 1.23 | 0.00 |
| 2041.00 | 2240.00 | 2270.33 | 1.13 | 0.00 |
| 1122.00 | 1387.00 | 1348.00 | 1.29 | 0.00 |
| 287.00 | 295.00 | 297.00 | 1.25 | 0.00 |
| 948.00 | 970.00 | 1048.67 | 1.06 | 0.24 |
| 1391.00 | 1576.00 | 1507.33 | 1.06 | 0.23 |
| 99.00 | 133.00 | 120.00 | -1.24 | 0.01 |

| | | | | |
|---|---|---|---|---|
| 20.00 | 21.00 | 24.00 | 1.11 | 0.68 |
| 93.00 | 115.00 | 103.00 | -1.34 | 0.00 |
| 2115.00 | 2251.00 | 2217.33 | 1.05 | 0.19 |
| 2316.00 | 2361.00 | 2459.67 | -1.21 | 0.00 |
| 578.00 | 459.00 | 492.67 | -1.47 | 0.00 |
| 242.00 | 228.00 | 226.67 | -1.38 | 0.00 |
| 6632.00 | 6615.00 | 6667.67 | -1.14 | 0.00 |
| 3928.00 | 3859.00 | 3934.00 | -1.77 | 0.00 |
| 3338.00 | 3424.00 | 3481.33 | -1.33 | 0.00 |
| 3236.00 | 3073.00 | 3196.00 | -1.07 | 0.11 |
| 10.00 | 7.00 | 8.33 | -1.53 | 0.14 |
| 1548.00 | 1707.00 | 1593.67 | -1.02 | 0.61 |
| 1475.00 | 1814.00 | 1705.00 | 1.11 | 0.02 |
| 845.00 | 921.00 | 919.00 | 1.00 | 1.00 |
| 212.00 | 224.00 | 204.67 | -1.00 | 1.00 |
| 9239.00 | 11594.00 | 10774.00 | -1.07 | 0.11 |
| 12.00 | 15.00 | 14.67 | -2.50 | 0.00 |
| 18.00 | 20.00 | 23.67 | -1.38 | 0.06 |
| 377.00 | 446.00 | 429.33 | 1.24 | 0.00 |
| 282.00 | 302.00 | 312.00 | -1.50 | 0.00 |
| 1623.00 | 1533.00 | 1515.67 | -1.38 | 0.00 |
| 0.00 | 2.00 | 1.67 | -2.33 | 0.15 |
| 44.00 | 51.00 | 49.33 | 1.25 | 0.11 |
| 10.00 | 9.00 | 8.33 | 1.12 | 0.78 |
| 11.00 | 14.00 | 10.67 | -1.03 | 1.00 |
| 447.00 | 537.00 | 495.33 | 1.55 | 0.00 |
| 3486.00 | 3688.00 | 3735.00 | -1.14 | 0.00 |
| 676.00 | 670.00 | 734.33 | 1.13 | 0.07 |
| 716.00 | 835.00 | 823.33 | 1.17 | 0.00 |
| 34.00 | 38.00 | 37.67 | -1.04 | 0.81 |
| 14.00 | 15.00 | 21.67 | -1.02 | 0.94 |
| 90.00 | 131.00 | 112.67 | 1.06 | 0.58 |
| 30.00 | 27.00 | 31.33 | -2.40 | 0.00 |
| 103.00 | 122.00 | 110.00 | 1.57 | 0.00 |
| 1150.00 | 1426.00 | 1242.00 | 1.40 | 0.00 |
| 885.00 | 888.00 | 962.33 | 1.01 | 0.82 |
| 502.00 | 666.00 | 600.33 | 1.03 | 0.58 |
| 166.00 | 190.00 | 189.33 | -1.96 | 0.00 |
| 471.00 | 527.00 | 520.00 | 1.22 | 0.00 |
| 855.00 | 955.00 | 936.00 | 1.15 | 0.00 |
| 488.00 | 564.00 | 566.33 | 1.07 | 0.23 |
| 217.00 | 273.00 | 251.00 | 1.07 | 0.34 |
| 893.00 | 1076.00 | 977.00 | 1.05 | 0.35 |
| 24.00 | 30.00 | 26.67 | 1.48 | 0.05 |
| 283.00 | 281.00 | 292.67 | -1.09 | 0.22 |
| 232.00 | 281.00 | 266.00 | 1.19 | 0.02 |
| 593.00 | 716.00 | 657.00 | -1.11 | 0.06 |
| 229.00 | 277.00 | 255.67 | -1.12 | 0.08 |

| | | | | |
|---|---|---|---|---|
| 462.00 | 514.00 | 488.67 | -1.39 | 0.00 |
| 6.00 | 7.00 | 6.67 | 1.31 | 0.51 |
| 379.00 | 388.00 | 407.33 | 1.19 | 0.00 |
| 428.00 | 483.00 | 485.33 | 1.06 | 0.26 |
| 2132.00 | 2104.00 | 2186.00 | -1.12 | 0.00 |
| 1536.00 | 1585.00 | 1571.00 | -1.24 | 0.00 |
| 262.00 | 245.00 | 258.67 | -1.04 | 0.51 |
| 4102.00 | 3935.00 | 4060.67 | -1.08 | 0.07 |
| 3.00 | 4.00 | 4.33 | 1.79 | 0.27 |
| 13.00 | 10.00 | 12.33 | 1.35 | 0.28 |
| 3.00 | 0.00 | 2.00 | -2.77 | 0.04 |
| 18899.00 | 19725.00 | 19239.67 | -1.04 | 0.26 |
| 1180.00 | 1163.00 | 1239.67 | -1.02 | 0.63 |
| 22.00 | 19.00 | 19.00 | -1.03 | 1.00 |
| 1081.00 | 1136.00 | 1168.67 | 1.10 | 0.02 |
| 704.00 | 760.00 | 757.67 | 1.54 | 0.00 |
| 481.00 | 527.00 | 552.00 | 1.00 | 0.96 |
| 520.00 | 598.00 | 589.00 | 1.08 | 0.11 |
| 1188.00 | 1138.00 | 1185.67 | -1.33 | 0.00 |
| 733.00 | 866.00 | 816.00 | 1.12 | 0.02 |
| 2758.00 | 2984.00 | 2992.67 | 1.06 | 0.15 |
| 21.00 | 15.00 | 18.33 | -2.07 | 0.00 |
| 1922.00 | 1944.00 | 1979.33 | 1.22 | 0.00 |
| 3375.00 | 3407.00 | 3416.67 | -1.13 | 0.00 |
| 119.00 | 141.00 | 132.00 | 1.27 | 0.01 |
| 298.00 | 342.00 | 342.00 | -1.18 | 0.02 |
| 1125.00 | 1235.00 | 1276.00 | -1.13 | 0.01 |
| 9.00 | 9.00 | 10.33 | 1.38 | 0.28 |
| 38.00 | 39.00 | 42.67 | 1.03 | 0.86 |
| 52.00 | 49.00 | 52.33 | 1.06 | 0.68 |
| 118.00 | 121.00 | 122.33 | 1.14 | 0.13 |
| 353.00 | 436.00 | 408.00 | 1.06 | 0.37 |
| 4.00 | 4.00 | 3.00 | -1.88 | 0.18 |
| 112.00 | 109.00 | 111.00 | 1.42 | 0.00 |
| 29.00 | 26.00 | 27.00 | 1.31 | 0.15 |
| 21.00 | 19.00 | 20.33 | -1.47 | 0.04 |
| 1.00 | 2.00 | 2.67 | -2.47 | 0.04 |
| 209.00 | 187.00 | 190.67 | 1.12 | 0.15 |
| 13.00 | 13.00 | 17.33 | -1.06 | 0.79 |
| 9.00 | 19.00 | 11.33 | -1.07 | 0.91 |
| 2610.00 | 2430.00 | 2544.33 | -1.00 | 0.97 |
| 3388.00 | 3446.00 | 3557.67 | -1.12 | 0.00 |
| 52.00 | 28.00 | 46.67 | 1.00 | 1.00 |
| 864.00 | 996.00 | 985.00 | 1.13 | 0.01 |
| 104.00 | 77.00 | 88.00 | 1.21 | 0.15 |
| 1616.00 | 1709.00 | 1658.67 | -1.11 | 0.01 |
| 156.00 | 155.00 | 164.33 | -1.10 | 0.23 |
| 1102.00 | 1444.00 | 1279.33 | -1.22 | 0.00 |

| | | | | |
|---|---|---|---|---|
| 391.00 | 440.00 | 418.33 | 1.09 | 0.11 |
| 5.00 | 2.00 | 3.67 | -3.33 | 0.00 |
| 781.00 | 921.00 | 947.67 | 1.21 | 0.00 |
| 606.00 | 839.00 | 788.33 | 1.46 | 0.00 |
| 2495.00 | 2791.00 | 2586.67 | -1.09 | 0.07 |
| 2.00 | 7.00 | 6.33 | 4.36 | 0.00 |
| 805.00 | 910.00 | 866.67 | 1.16 | 0.00 |
| 3852.00 | 4679.00 | 4294.00 | -1.45 | 0.00 |
| 5.00 | 5.00 | 4.33 | -1.32 | 0.59 |
| 374.00 | 436.00 | 437.00 | -1.06 | 0.33 |
| 592.00 | 623.00 | 593.00 | -1.26 | 0.00 |
| 435.00 | 421.00 | 459.67 | -1.02 | 0.78 |
| 1179.00 | 1072.00 | 1143.33 | -1.10 | 0.05 |
| 1256.00 | 1380.00 | 1308.33 | -1.14 | 0.00 |
| 402.00 | 403.00 | 381.33 | -1.23 | 0.01 |
| 127.00 | 91.00 | 122.00 | -1.71 | 0.00 |
| 12.00 | 12.00 | 17.00 | -1.35 | 0.17 |
| 746.00 | 812.00 | 759.00 | -1.01 | 0.79 |
| 871.00 | 890.00 | 826.00 | -1.12 | 0.10 |
| 3376.00 | 3340.00 | 3446.33 | -1.16 | 0.00 |
| 31.00 | 52.00 | 42.67 | 1.25 | 0.13 |
| 0.00 | 1.00 | 0.33 | -3.95 | 0.23 |
| 3831.00 | 4332.00 | 3971.33 | -1.24 | 0.00 |
| 1139.00 | 1114.00 | 1145.00 | -1.32 | 0.00 |
| 1728.00 | 1756.00 | 1729.67 | -1.09 | 0.05 |
| 627.00 | 556.00 | 595.33 | -1.16 | 0.01 |
| 2589.00 | 2376.00 | 2575.33 | -1.25 | 0.00 |
| 587.00 | 608.00 | 626.33 | -1.12 | 0.03 |
| 1031.00 | 1075.00 | 1131.00 | -1.01 | 0.87 |
| 876.00 | 1039.00 | 946.33 | 1.11 | 0.04 |
| 41.00 | 67.00 | 51.00 | 1.06 | 0.71 |
| 86.00 | 86.00 | 81.33 | 1.05 | 0.66 |
| 1588.00 | 1668.00 | 1701.33 | 1.01 | 0.83 |
| 43.00 | 40.00 | 42.67 | -1.08 | 0.60 |
| 1.00 | 0.00 | 3.33 | 1.21 | 0.85 |
| 86.00 | 70.00 | 74.00 | 1.25 | 0.06 |
| 2357.00 | 2327.00 | 2286.67 | -1.45 | 0.00 |
| 34.00 | 32.00 | 40.33 | 1.49 | 0.02 |
| 2109.00 | 2369.00 | 2404.33 | 1.14 | 0.00 |
| 73.00 | 60.00 | 74.67 | 1.24 | 0.08 |
| 874.00 | 895.00 | 884.00 | -1.10 | 0.05 |
| 135.00 | 134.00 | 133.33 | -1.35 | 0.00 |
| 1525.00 | 1516.00 | 1581.67 | 1.02 | 0.58 |
| 2601.00 | 2723.00 | 2757.67 | 1.08 | 0.03 |
| 1188.00 | 1164.00 | 1259.67 | -1.04 | 0.43 |
| 704.00 | 701.00 | 713.00 | -1.15 | 0.00 |
| 486.00 | 517.00 | 497.67 | 1.03 | 0.63 |
| 1162.00 | 1182.00 | 1218.67 | 1.00 | 1.00 |

| | | | | |
|---|---|---|---|---|
| 186.00 | 186.00 | 177.00 | -1.52 | 0.00 |
| 481.00 | 514.00 | 530.00 | -1.08 | 0.12 |
| 1718.00 | 1732.00 | 1728.67 | -1.07 | 0.07 |
| 842.00 | 898.00 | 897.00 | -1.11 | 0.01 |
| 443.00 | 466.00 | 490.67 | 1.04 | 0.50 |
| 1541.00 | 1615.00 | 1583.33 | 1.05 | 0.23 |
| 922.00 | 1091.00 | 981.33 | -1.22 | 0.00 |
| 961.00 | 1085.00 | 1066.67 | -1.21 | 0.00 |
| 696.00 | 770.00 | 741.67 | -1.20 | 0.00 |
| 233.00 | 219.00 | 223.00 | 1.31 | 0.00 |
| 445.00 | 480.00 | 505.00 | 1.16 | 0.02 |
| 1589.00 | 1836.00 | 1733.00 | 1.01 | 0.76 |
| 416.00 | 487.00 | 475.67 | 1.62 | 0.00 |
| 26.00 | 23.00 | 23.33 | 1.06 | 0.75 |
| 127.00 | 117.00 | 119.33 | -1.42 | 0.00 |
| 6995.00 | 6804.00 | 6906.00 | -1.28 | 0.00 |
| 8.00 | 14.00 | 14.67 | -1.01 | 1.00 |
| 1137.00 | 1087.00 | 1145.00 | 1.06 | 0.17 |
| 1261.00 | 1526.00 | 1365.33 | -1.02 | 0.70 |
| 9559.00 | 9572.00 | 9630.33 | 1.05 | 0.19 |
| 1828.00 | 2060.00 | 1959.67 | 1.03 | 0.48 |
| 2.00 | 2.00 | 1.33 | -2.63 | 0.12 |
| 107.00 | 90.00 | 101.00 | 1.19 | 0.08 |
| 15.00 | 11.00 | 12.67 | 1.49 | 0.14 |
| 1704.00 | 1746.00 | 1753.00 | -1.16 | 0.00 |
| 5.00 | 5.00 | 6.33 | 1.68 | 0.21 |
| 1546.00 | 1757.00 | 1677.67 | -1.25 | 0.00 |
| 542.00 | 601.00 | 561.33 | 1.03 | 0.62 |
| 2398.00 | 2602.00 | 2532.33 | 1.06 | 0.12 |
| 1.00 | 3.00 | 2.67 | -2.71 | 0.02 |
| 291.00 | 306.00 | 319.67 | -1.06 | 0.33 |
| 861.00 | 958.00 | 935.33 | -1.14 | 0.00 |
| 1048.00 | 1038.00 | 1117.67 | -1.20 | 0.00 |
| 1533.00 | 1758.00 | 1661.00 | 1.04 | 0.39 |
| 1889.00 | 2049.00 | 2074.67 | 1.05 | 0.24 |
| 736.00 | 681.00 | 743.00 | 1.05 | 0.33 |
| 1421.00 | 1444.00 | 1444.00 | 1.13 | 0.01 |
| 304.00 | 314.00 | 316.00 | -1.02 | 0.78 |
| 1019.00 | 1175.00 | 1153.33 | -1.02 | 0.64 |
| 209.00 | 222.00 | 218.33 | -1.00 | 0.98 |
| 11.00 | 5.00 | 12.00 | -1.21 | 0.53 |
| 2902.00 | 3189.00 | 3247.33 | -1.29 | 0.00 |
| 884.00 | 890.00 | 898.67 | 1.11 | 0.01 |
| 3063.00 | 3145.00 | 3168.67 | 1.28 | 0.00 |
| 2120.00 | 2079.00 | 2191.67 | -1.00 | 0.96 |
| 932.00 | 903.00 | 913.67 | -1.38 | 0.00 |
| 759.00 | 733.00 | 765.00 | 1.07 | 0.11 |
| 361.00 | 432.00 | 431.33 | -1.10 | 0.13 |

| | | | | |
|---|---|---|---|---|
| 1274.00 | 1269.00 | 1285.00 | -1.13 | 0.01 |
| 2876.00 | 3139.00 | 3138.67 | 1.22 | 0.00 |
| 411.00 | 379.00 | 401.00 | -1.04 | 0.55 |
| 153.00 | 176.00 | 173.67 | 1.03 | 0.71 |
| 25.00 | 19.00 | 19.33 | 1.10 | 0.65 |
| 10.00 | 8.00 | 9.67 | 1.67 | 0.12 |
| 93.00 | 72.00 | 79.67 | 1.64 | 0.00 |
| 4065.00 | 4167.00 | 4171.67 | 1.47 | 0.00 |
| 954.00 | 1030.00 | 1037.67 | 1.04 | 0.40 |
| 4878.00 | 5445.00 | 5264.67 | -1.08 | 0.02 |
| 2025.00 | 2540.00 | 2248.00 | -1.00 | 0.96 |
| 66.00 | 46.00 | 52.33 | -1.24 | 0.11 |
| 529.00 | 499.00 | 541.00 | -1.12 | 0.03 |
| 2409.00 | 2635.00 | 2554.67 | 1.00 | 0.97 |
| 1570.00 | 1524.00 | 1663.67 | 1.38 | 0.00 |
| 291.00 | 415.00 | 356.00 | 1.11 | 0.19 |
| 1433.00 | 1469.00 | 1475.67 | 1.03 | 0.40 |
| 1656.00 | 1837.00 | 1815.67 | -1.13 | 0.00 |
| 530.00 | 535.00 | 537.67 | -1.16 | 0.00 |
| 11613.00 | 12331.00 | 12281.67 | 1.04 | 0.24 |
| 769.00 | 823.00 | 817.33 | -1.06 | 0.14 |
| 2189.00 | 2194.00 | 2179.00 | -1.11 | 0.01 |
| 15.00 | 11.00 | 11.67 | -2.19 | 0.00 |
| 5.00 | 3.00 | 6.00 | 1.36 | 0.48 |
| 2815.00 | 2972.00 | 2953.00 | -1.32 | 0.00 |
| 219.00 | 247.00 | 228.33 | -1.13 | 0.06 |
| 658.00 | 676.00 | 654.00 | 1.13 | 0.03 |
| 492.00 | 644.00 | 594.67 | 1.09 | 0.13 |
| 12570.00 | 13712.00 | 13017.67 | -1.07 | 0.07 |
| 1337.00 | 1524.00 | 1468.67 | 1.16 | 0.00 |
| 12.00 | 3.00 | 6.33 | 1.17 | 0.75 |
| 16.00 | 8.00 | 11.67 | -1.18 | 0.58 |
| 101.00 | 114.00 | 98.33 | -1.78 | 0.00 |
| 91.00 | 60.00 | 74.00 | 1.42 | 0.01 |
| 4.00 | 3.00 | 3.33 | -1.50 | 0.43 |
| 7685.00 | 8896.00 | 8229.00 | 1.16 | 0.00 |
| 1254.00 | 1240.00 | 1317.33 | -1.16 | 0.00 |
| 2994.00 | 2931.00 | 2933.00 | -1.38 | 0.00 |
| 1061.00 | 1119.00 | 1161.33 | -1.08 | 0.08 |
| 2221.00 | 2387.00 | 2424.00 | 1.12 | 0.01 |
| 256.00 | 305.00 | 291.67 | -1.19 | 0.02 |
| 2002.00 | 2042.00 | 2058.33 | -1.12 | 0.00 |
| 710.00 | 805.00 | 800.00 | 1.05 | 0.38 |
| 2805.00 | 3021.00 | 3095.00 | 1.02 | 0.71 |
| 723.00 | 781.00 | 785.33 | -1.04 | 0.42 |
| 235.00 | 186.00 | 220.67 | -1.16 | 0.08 |
| 1131.00 | 1083.00 | 1151.67 | -1.12 | 0.01 |
| 301.00 | 343.00 | 317.67 | 1.23 | 0.00 |

| | | | | |
|---|---|---|---|---|
| 157.00 | 164.00 | 165.00 | -1.12 | 0.12 |
| 29.00 | 29.00 | 28.67 | 1.39 | 0.06 |
| 432.00 | 498.00 | 467.33 | 1.06 | 0.29 |
| 3631.00 | 4015.00 | 3784.33 | -1.04 | 0.44 |
| 43.00 | 33.00 | 41.33 | -1.07 | 0.64 |
| 329.00 | 329.00 | 340.67 | -1.38 | 0.00 |
| 256.00 | 284.00 | 264.67 | 1.30 | 0.00 |
| 226.00 | 222.00 | 234.67 | -1.04 | 0.60 |
| 427.00 | 402.00 | 413.67 | 1.03 | 0.62 |
| 21.00 | 16.00 | 20.33 | -1.91 | 0.00 |
| 608.00 | 625.00 | 627.67 | -1.44 | 0.00 |
| 976.00 | 1075.00 | 1039.33 | -1.02 | 0.60 |
| 6054.00 | 5840.00 | 6100.67 | -1.16 | 0.00 |
| 1008.00 | 952.00 | 959.00 | 1.12 | 0.04 |
| 1624.00 | 1666.00 | 1697.33 | 1.02 | 0.55 |
| 0.00 | 6.00 | 3.33 | 3.03 | 0.11 |
| 49.00 | 32.00 | 38.00 | -2.67 | 0.00 |
| 261.00 | 275.00 | 277.33 | 1.23 | 0.00 |
| 650.00 | 601.00 | 649.67 | 1.03 | 0.61 |
| 768.00 | 820.00 | 795.00 | 1.01 | 0.79 |
| 1363.00 | 1444.00 | 1424.67 | 1.08 | 0.05 |
| 94.00 | 102.00 | 107.00 | -1.11 | 0.23 |
| 6.00 | 6.00 | 5.33 | -2.13 | 0.02 |
| 25.00 | 24.00 | 31.33 | -1.12 | 0.49 |
| 16618.00 | 15236.00 | 15718.67 | 1.01 | 0.76 |
| 180.00 | 195.00 | 198.67 | 1.04 | 0.62 |
| 15684.00 | 17220.00 | 16937.67 | 1.14 | 0.00 |
| 422.00 | 522.00 | 459.00 | 1.47 | 0.00 |
| 1988.00 | 2093.00 | 2046.67 | 1.14 | 0.00 |
| 3049.00 | 3259.00 | 3221.00 | 1.17 | 0.00 |
| 3272.00 | 3281.00 | 3294.67 | -1.10 | 0.01 |
| 178.00 | 179.00 | 176.67 | -1.17 | 0.03 |
| 2360.00 | 2550.00 | 2379.00 | 1.03 | 0.51 |
| 1132.00 | 1166.00 | 1163.33 | -1.09 | 0.06 |
| 1732.00 | 1842.00 | 1748.33 | 1.26 | 0.00 |
| 5310.00 | 5853.00 | 5727.67 | 1.36 | 0.00 |
| 1522.00 | 1908.00 | 1737.67 | 1.16 | 0.00 |
| 318.00 | 351.00 | 340.00 | 1.08 | 0.24 |
| 1608.00 | 1782.00 | 1770.67 | 1.14 | 0.00 |
| 19.00 | 26.00 | 23.33 | -1.07 | 0.69 |
| 598.00 | 611.00 | 619.33 | -1.17 | 0.00 |
| 855.00 | 886.00 | 889.00 | 1.11 | 0.01 |
| 214.00 | 236.00 | 221.33 | -1.16 | 0.02 |
| 336.00 | 377.00 | 356.67 | 1.02 | 0.78 |
| 23.00 | 18.00 | 24.33 | 1.16 | 0.48 |
| 1330.00 | 1384.00 | 1428.33 | 1.09 | 0.03 |
| 867.00 | 837.00 | 844.00 | 1.23 | 0.00 |
| 1085.00 | 1239.00 | 1201.67 | 1.04 | 0.31 |

| | | | | |
|---|---|---|---|---|
| 180.00 | 162.00 | 177.00 | -1.30 | 0.00 |
| 637.00 | 657.00 | 665.33 | -1.03 | 0.53 |
| 767.00 | 871.00 | 828.33 | 1.10 | 0.03 |
| 2.00 | 0.00 | 1.00 | -2.51 | 0.23 |
| 793.00 | 941.00 | 847.67 | 1.17 | 0.01 |
| 35.00 | 25.00 | 30.00 | -2.41 | 0.00 |
| 80.00 | 106.00 | 96.00 | -1.22 | 0.03 |
| 2884.00 | 3179.00 | 3209.33 | 1.23 | 0.00 |
| 36.00 | 28.00 | 34.33 | 1.39 | 0.04 |
| 4194.00 | 4283.00 | 4375.67 | 1.26 | 0.00 |
| 1790.00 | 1714.00 | 1803.00 | 1.13 | 0.02 |
| 325.00 | 404.00 | 373.00 | 1.09 | 0.15 |
| 1067.00 | 1208.00 | 1171.33 | 1.07 | 0.14 |
| 1935.00 | 2153.00 | 2135.67 | 1.00 | 0.91 |
| 3555.00 | 3662.00 | 3573.33 | -1.14 | 0.00 |
| 37.00 | 38.00 | 38.00 | -1.21 | 0.17 |
| 805.00 | 850.00 | 863.33 | 1.18 | 0.00 |
| 1301.00 | 1537.00 | 1440.00 | 1.20 | 0.00 |
| 29.00 | 36.00 | 37.33 | -1.99 | 0.00 |
| 1020.00 | 1192.00 | 1175.33 | 1.18 | 0.00 |
| 1318.00 | 1557.00 | 1527.00 | 1.22 | 0.00 |
| 1062.00 | 1264.00 | 1115.33 | 1.10 | 0.16 |
| 1163.00 | 1114.00 | 1143.33 | -1.26 | 0.00 |
| 468.00 | 458.00 | 474.67 | 1.02 | 0.76 |
| 2434.00 | 2276.00 | 2298.67 | -1.14 | 0.01 |
| 663.00 | 766.00 | 732.67 | 1.24 | 0.00 |
| 233.00 | 223.00 | 246.00 | 1.13 | 0.09 |
| 1660.00 | 1523.00 | 1594.00 | -1.25 | 0.00 |
| 98.00 | 100.00 | 90.67 | -1.14 | 0.22 |
| 2051.00 | 2228.00 | 2077.33 | -1.08 | 0.17 |
| 2778.00 | 2854.00 | 2728.33 | -1.10 | 0.06 |
| 1433.00 | 1484.00 | 1474.00 | -1.03 | 0.53 |
| 1.00 | 2.00 | 1.00 | -2.51 | 0.25 |
| 1833.00 | 1941.00 | 1929.67 | -1.02 | 0.67 |
| 661.00 | 647.00 | 646.33 | -1.02 | 0.66 |
| 868.00 | 954.00 | 930.00 | 1.04 | 0.35 |
| 296.00 | 284.00 | 320.33 | 1.04 | 0.57 |
| 257.00 | 361.00 | 342.33 | -1.01 | 0.93 |
| 677.00 | 704.00 | 702.00 | 1.11 | 0.02 |
| 493.00 | 616.00 | 584.67 | 1.01 | 0.83 |
| 1585.00 | 1531.00 | 1569.33 | 1.10 | 0.03 |
| 1259.00 | 1405.00 | 1339.00 | -1.11 | 0.01 |
| 2114.00 | 2027.00 | 2128.67 | -1.09 | 0.04 |
| 532.00 | 545.00 | 555.00 | -1.08 | 0.10 |
| 203.00 | 232.00 | 224.67 | -1.05 | 0.51 |
| 502.00 | 549.00 | 552.67 | 1.14 | 0.01 |
| 441.00 | 493.00 | 491.33 | -1.01 | 0.86 |
| 584.00 | 548.00 | 576.00 | -1.01 | 0.93 |

| | | | | |
|---|---|---|---|---|
| 318.00 | 324.00 | 347.33 | 1.29 | 0.00 |
| 141.00 | 170.00 | 164.00 | 1.06 | 0.46 |
| 474.00 | 469.00 | 494.67 | 1.18 | 0.00 |
| 489.00 | 526.00 | 514.33 | 1.15 | 0.01 |
| 248.00 | 266.00 | 266.67 | -1.03 | 0.73 |
| 1518.00 | 1700.00 | 1712.33 | 1.17 | 0.00 |
| 194.00 | 209.00 | 208.67 | -1.11 | 0.17 |
| 1147.00 | 1257.00 | 1203.67 | -1.00 | 0.99 |
| 0.00 | 3.00 | 1.33 | -3.33 | 0.03 |
| 103.00 | 96.00 | 104.00 | 1.28 | 0.02 |
| 4.00 | 8.00 | 7.00 | -1.06 | 1.00 |
| 198.00 | 220.00 | 221.33 | -1.10 | 0.18 |
| 9.00 | 12.00 | 11.33 | 1.20 | 0.54 |
| 1377.00 | 1379.00 | 1399.33 | -1.21 | 0.00 |
| 6.00 | 3.00 | 3.67 | -2.71 | 0.01 |
| 1519.00 | 1550.00 | 1487.00 | -1.44 | 0.00 |
| 202.00 | 189.00 | 200.33 | -1.47 | 0.00 |
| 8.00 | 18.00 | 13.67 | 1.06 | 0.83 |
| 13.00 | 12.00 | 10.33 | 1.45 | 0.23 |
| 131.00 | 166.00 | 152.00 | 1.12 | 0.15 |
| 21.00 | 26.00 | 27.33 | -1.40 | 0.03 |
| 392.00 | 432.00 | 423.00 | -1.00 | 0.98 |
| 1.00 | 0.00 | 2.33 | -2.39 | 0.07 |
| 121.00 | 144.00 | 138.67 | -1.07 | 0.36 |
| 1277.00 | 1432.00 | 1426.67 | 1.34 | 0.00 |
| 549.00 | 572.00 | 591.00 | 1.32 | 0.00 |
| 3210.00 | 3228.00 | 3355.00 | -1.12 | 0.00 |
| 1813.00 | 1811.00 | 1859.67 | -1.08 | 0.05 |
| 313.00 | 357.00 | 338.67 | -1.30 | 0.00 |
| 945.00 | 1022.00 | 1008.00 | 1.30 | 0.00 |
| 1948.00 | 1883.00 | 1951.67 | 1.00 | 0.99 |
| 26.00 | 34.00 | 30.67 | 2.01 | 0.00 |
| 20.00 | 17.00 | 16.00 | 1.96 | 0.01 |
| 17.00 | 19.00 | 18.00 | -1.11 | 0.72 |
| 5109.00 | 5322.00 | 5236.00 | -1.41 | 0.00 |
| 1313.00 | 1517.00 | 1458.67 | 1.19 | 0.00 |
| 473.00 | 559.00 | 525.00 | -1.20 | 0.00 |
| 2451.00 | 2752.00 | 2694.67 | 1.20 | 0.00 |
| 618.00 | 589.00 | 635.67 | -1.03 | 0.63 |
| 159.00 | 161.00 | 167.67 | -1.06 | 0.46 |
| 382.00 | 399.00 | 428.67 | 1.04 | 0.52 |
| 524.00 | 624.00 | 596.67 | 1.22 | 0.00 |
| 24.00 | 25.00 | 32.67 | -1.28 | 0.15 |
| 1431.00 | 1462.00 | 1527.00 | -1.06 | 0.19 |
| 396.00 | 457.00 | 435.33 | 1.07 | 0.22 |
| 472.00 | 548.00 | 525.33 | 1.08 | 0.12 |
| 1140.00 | 1261.00 | 1190.00 | 1.06 | 0.21 |
| 1362.00 | 1439.00 | 1455.33 | 1.01 | 0.90 |

| | | | | |
|---|---|---|---|---|
| 202.00 | 248.00 | 244.00 | 1.20 | 0.02 |
| 47.00 | 43.00 | 44.00 | -1.01 | 1.00 |
| 259.00 | 299.00 | 290.00 | 1.44 | 0.00 |
| 260.00 | 299.00 | 293.00 | -1.07 | 0.27 |
| 2.00 | 2.00 | 2.67 | -1.62 | 0.39 |
| 355.00 | 358.00 | 376.33 | -1.16 | 0.01 |
| 596.00 | 683.00 | 670.00 | 1.07 | 0.15 |
| 48.00 | 53.00 | 56.00 | -1.13 | 0.28 |
| 603.00 | 606.00 | 601.33 | 1.23 | 0.00 |
| 733.00 | 779.00 | 742.00 | 1.13 | 0.02 |
| 2635.00 | 2864.00 | 2846.67 | 1.00 | 0.96 |
| 1022.00 | 1145.00 | 1092.67 | 1.05 | 0.27 |
| 809.00 | 981.00 | 962.67 | 1.17 | 0.00 |
| 452.00 | 481.00 | 451.33 | 1.14 | 0.03 |
| 0.00 | 1.00 | 0.33 | -5.43 | 0.07 |
| 6.00 | 6.00 | 7.00 | -1.87 | 0.03 |
| 703.00 | 802.00 | 771.00 | 1.12 | 0.01 |
| 4874.00 | 4930.00 | 5093.33 | 1.36 | 0.00 |
| 859.00 | 861.00 | 869.67 | 1.05 | 0.38 |
| 117.00 | 154.00 | 135.67 | -1.08 | 0.39 |
| 1094.00 | 1149.00 | 1167.33 | 1.17 | 0.00 |
| 324.00 | 336.00 | 327.33 | 1.16 | 0.02 |
| 699.00 | 820.00 | 778.67 | 1.12 | 0.02 |
| 71.00 | 87.00 | 82.67 | -1.14 | 0.23 |
| 893.00 | 866.00 | 894.33 | -1.25 | 0.00 |
| 219.00 | 217.00 | 210.33 | 1.07 | 0.35 |
| 1560.00 | 1753.00 | 1740.67 | 1.17 | 0.00 |
| 1376.00 | 1364.00 | 1404.00 | -1.48 | 0.00 |
| 569.00 | 490.00 | 509.00 | -1.19 | 0.01 |
| 1984.00 | 1907.00 | 1976.67 | -1.26 | 0.00 |
| 119.00 | 110.00 | 122.33 | 1.35 | 0.00 |
| 1002.00 | 1148.00 | 1068.33 | 1.06 | 0.22 |
| 2623.00 | 2763.00 | 2804.00 | -1.01 | 0.89 |
| 35.00 | 50.00 | 46.33 | -1.27 | 0.06 |
| 2489.00 | 2519.00 | 2527.33 | -1.00 | 0.92 |
| 6113.00 | 6104.00 | 6530.33 | -1.05 | 0.20 |
| 512.00 | 636.00 | 602.33 | 1.13 | 0.03 |
| 1472.00 | 1474.00 | 1466.33 | -1.33 | 0.00 |
| 258.00 | 276.00 | 285.67 | -1.20 | 0.00 |
| 870.00 | 838.00 | 874.33 | -1.17 | 0.00 |
| 656.00 | 622.00 | 675.00 | -1.15 | 0.00 |
| 90.00 | 82.00 | 85.00 | -1.55 | 0.00 |
| 465.00 | 519.00 | 524.33 | -1.10 | 0.08 |
| 163.00 | 153.00 | 155.67 | -1.20 | 0.02 |
| 1166.00 | 1252.00 | 1225.67 | -1.09 | 0.05 |
| 2287.00 | 2367.00 | 2371.33 | -1.14 | 0.00 |
| 1280.00 | 1373.00 | 1362.33 | 1.03 | 0.50 |
| 7.00 | 13.00 | 11.33 | 1.51 | 0.16 |

| | | | | |
|---|---|---|---|---|
| 3853.00 | 3751.00 | 3972.67 | -1.03 | 0.50 |
| 1975.00 | 1948.00 | 1949.00 | -1.07 | 0.14 |
| 432.00 | 443.00 | 440.00 | -1.83 | 0.00 |
| 15.00 | 16.00 | 15.33 | -1.36 | 0.17 |
| 0.00 | 0.00 | 2.33 | -1.17 | 1.00 |
| 622.00 | 665.00 | 685.33 | -1.15 | 0.00 |
| 13.00 | 21.00 | 17.00 | -1.21 | 0.41 |
| 114.00 | 89.00 | 103.00 | -1.07 | 0.53 |
| 135.00 | 176.00 | 166.00 | 1.22 | 0.02 |
| 7041.00 | 7693.00 | 7434.33 | -1.05 | 0.15 |
| 5799.00 | 5712.00 | 5980.67 | 1.05 | 0.19 |
| 719.00 | 900.00 | 880.33 | 1.08 | 0.22 |
| 926.00 | 862.00 | 897.33 | -1.16 | 0.00 |
| 1615.00 | 1878.00 | 1837.67 | 1.05 | 0.32 |
| 180.00 | 202.00 | 185.33 | 1.26 | 0.01 |
| 950.00 | 994.00 | 1032.67 | -1.10 | 0.04 |
| 2681.00 | 2900.00 | 2842.33 | -1.23 | 0.00 |
| 5.00 | 15.00 | 12.00 | -1.02 | 1.00 |
| 43.00 | 33.00 | 40.00 | 1.01 | 1.00 |
| 1475.00 | 1584.00 | 1453.00 | 1.01 | 0.93 |
| 955.00 | 898.00 | 954.33 | -1.29 | 0.00 |
| 214.00 | 245.00 | 250.00 | 1.02 | 0.78 |
| 1557.00 | 1633.00 | 1652.33 | -1.18 | 0.00 |
| 8457.00 | 8089.00 | 8282.00 | -1.23 | 0.00 |
| 2940.00 | 2761.00 | 2868.33 | -1.33 | 0.00 |
| 116.00 | 97.00 | 108.00 | 1.25 | 0.02 |
| 1044.00 | 1088.00 | 1050.33 | -1.25 | 0.00 |
| 381.00 | 339.00 | 356.33 | -1.33 | 0.00 |
| 883.00 | 899.00 | 937.67 | 1.04 | 0.48 |
| 699.00 | 769.00 | 754.67 | 1.05 | 0.31 |
| 123.00 | 204.00 | 160.33 | -1.10 | 0.40 |
| 422.00 | 480.00 | 472.33 | -1.34 | 0.00 |
| 529.00 | 586.00 | 599.67 | -1.00 | 0.94 |
| 11.00 | 14.00 | 13.00 | -1.76 | 0.01 |
| 800.00 | 801.00 | 839.00 | -1.74 | 0.00 |
| 1403.00 | 1363.00 | 1399.33 | -1.05 | 0.32 |
| 127.00 | 148.00 | 144.33 | -1.50 | 0.00 |
| 386.00 | 457.00 | 460.00 | 1.07 | 0.31 |
| 643.00 | 720.00 | 687.33 | 1.12 | 0.02 |
| 727.00 | 751.00 | 773.33 | -1.02 | 0.64 |
| 516.00 | 642.00 | 585.00 | 1.00 | 0.95 |
| 5.00 | 13.00 | 8.67 | 1.81 | 0.09 |
| 358.00 | 366.00 | 382.00 | 1.06 | 0.31 |
| 767.00 | 738.00 | 827.67 | 1.03 | 0.65 |
| 345.00 | 364.00 | 337.67 | -1.03 | 0.68 |
| 1903.00 | 2134.00 | 2101.00 | 1.05 | 0.24 |
| 3922.00 | 3635.00 | 3873.67 | -1.09 | 0.04 |
| 703.00 | 724.00 | 788.67 | 1.10 | 0.11 |

| | | | | |
|---|---|---|---|---|
| 1258.00 | 1239.00 | 1256.33 | 1.02 | 0.70 |
| 1172.00 | 1357.00 | 1303.33 | 1.12 | 0.01 |
| 11.00 | 8.00 | 7.00 | -2.03 | 0.03 |
| 434.00 | 426.00 | 440.33 | 1.21 | 0.00 |
| 48.00 | 49.00 | 52.33 | -1.14 | 0.32 |
| 429.00 | 544.00 | 502.33 | 1.12 | 0.07 |
| 3462.00 | 4337.00 | 4045.67 | -1.36 | 0.00 |
| 1.00 | 11.00 | 6.67 | 1.40 | 0.41 |
| 13.00 | 8.00 | 8.33 | -1.09 | 0.89 |
| 3.00 | 1.00 | 3.00 | -1.77 | 0.24 |
| 1.00 | 5.00 | 3.67 | -1.01 | 1.00 |
| 14.00 | 16.00 | 11.67 | 1.19 | 0.61 |
| 45.00 | 72.00 | 56.67 | -1.25 | 0.08 |
| 0.00 | 5.00 | 1.67 | -7.42 | 0.00 |
| 7.00 | 5.00 | 6.67 | -1.07 | 1.00 |
| 12.00 | 7.00 | 8.00 | 1.18 | 0.66 |
| 0.00 | 0.00 | 0.00 | -9.06 | 0.27 |
| 86.00 | 111.00 | 102.33 | -1.09 | 0.34 |
| 14.00 | 24.00 | 15.33 | 1.51 | 0.12 |
| 36.00 | 34.00 | 29.67 | -1.78 | 0.00 |
| 54.00 | 68.00 | 72.67 | -1.02 | 0.88 |
| 9.00 | 4.00 | 6.67 | -1.01 | 1.00 |
| 172.00 | 262.00 | 217.33 | -1.05 | 0.57 |
| 572.00 | 701.00 | 613.00 | -1.06 | 0.43 |
| 37.00 | 56.00 | 37.33 | -1.32 | 0.18 |
| 7.00 | 1.00 | 3.67 | -1.01 | 1.00 |
| 0.00 | 1.00 | 0.67 | -3.57 | 0.11 |
| 53.00 | 61.00 | 67.00 | -1.05 | 0.70 |
| 24.00 | 17.00 | 19.00 | 1.20 | 0.40 |
| 13.00 | 15.00 | 17.67 | -1.92 | 0.00 |
| 606.00 | 667.00 | 670.67 | -1.27 | 0.00 |
| 41.00 | 55.00 | 51.33 | -1.49 | 0.00 |
| 4.00 | 3.00 | 5.33 | -4.37 | 0.00 |
| 0.00 | 1.00 | 1.00 | -4.31 | 0.01 |
| 2.00 | 1.00 | 1.67 | -1.39 | 0.78 |
| 41.00 | 55.00 | 55.33 | 1.26 | 0.10 |
| 0.00 | 0.00 | 0.00 | -9.06 | 0.27 |
| 0.00 | 0.00 | 1.00 | -1.92 | 0.52 |
| 1.00 | 0.00 | 1.00 | -3.11 | 0.10 |
| 2.00 | 2.00 | 2.00 | -1.97 | 0.24 |
| 3.00 | 4.00 | 3.67 | -1.90 | 0.13 |
| 35.00 | 35.00 | 38.33 | -1.28 | 0.08 |
| 4.00 | 2.00 | 4.00 | -1.58 | 0.32 |
| 5.00 | 4.00 | 7.00 | -1.96 | 0.05 |
| 19.00 | 19.00 | 22.33 | -1.27 | 0.19 |
| 228.00 | 267.00 | 266.00 | 1.05 | 0.45 |
| 42.00 | 34.00 | 34.00 | 1.40 | 0.05 |
| 3.00 | 4.00 | 2.67 | -1.86 | 0.22 |

| | | | | |
|---|---|---|---|---|
| 1.00 | 2.00 | 1.33 | -2.17 | 0.27 |
| 3.00 | 4.00 | 2.67 | -1.98 | 0.16 |
| 6.00 | 3.00 | 4.00 | -2.16 | 0.04 |
| 25.00 | 27.00 | 29.33 | -1.29 | 0.12 |
| 0.00 | 3.00 | 1.00 | -1.61 | 0.73 |
| 12.00 | 35.00 | 23.00 | -1.92 | 0.00 |
| 3.00 | 4.00 | 3.00 | -2.85 | 0.01 |
| 5.00 | 5.00 | 5.00 | 1.06 | 1.00 |
| 287.00 | 361.00 | 347.67 | -1.39 | 0.00 |
| 0.00 | 0.00 | 0.33 | -4.69 | 0.13 |
| 4.00 | 3.00 | 2.67 | -1.74 | 0.29 |
| 12.00 | 11.00 | 11.33 | 1.34 | 0.31 |
| 4.00 | 5.00 | 4.33 | -1.47 | 0.38 |
| 0.00 | 0.00 | 0.00 | -9.06 | 0.27 |
| 1888.00 | 1809.00 | 1969.00 | 1.10 | 0.03 |
| 2.00 | 0.00 | 1.67 | -1.77 | 0.43 |
| 4.00 | 1.00 | 1.67 | -2.52 | 0.10 |
| 4.00 | 3.00 | 3.00 | -2.42 | 0.03 |
| 0.00 | 6.00 | 4.33 | -1.01 | 1.00 |
| 30.00 | 42.00 | 41.00 | 1.17 | 0.32 |
| 6.00 | 3.00 | 6.67 | 1.04 | 1.00 |
| 5.00 | 0.00 | 1.67 | -2.52 | 0.11 |
| 1.00 | 1.00 | 0.67 | -6.98 | 0.00 |
| 1.00 | 0.00 | 0.67 | -4.85 | 0.02 |
| 3.00 | 6.00 | 4.00 | -1.42 | 0.47 |
| 4.00 | 1.00 | 3.67 | 1.08 | 1.00 |
| 7.00 | 10.00 | 8.67 | 1.59 | 0.17 |
| 121.00 | 121.00 | 119.00 | -1.14 | 0.15 |
| 4.00 | 1.00 | 2.67 | -1.62 | 0.39 |
| 4.00 | 7.00 | 4.33 | -2.08 | 0.05 |
| 5.00 | 8.00 | 5.67 | 2.05 | 0.11 |
| 23.00 | 25.00 | 22.67 | -1.07 | 0.75 |
| 5.00 | 12.00 | 9.67 | 2.16 | 0.02 |
| 10.00 | 7.00 | 9.00 | -1.75 | 0.03 |
| 26.00 | 19.00 | 21.00 | -1.24 | 0.30 |
| 3.00 | 3.00 | 3.33 | -1.50 | 0.43 |
| 3005.00 | 3055.00 | 3202.33 | 1.13 | 0.00 |
| 147.00 | 139.00 | 145.67 | -1.21 | 0.01 |
| 27.00 | 32.00 | 29.33 | -1.06 | 0.79 |
| 227.00 | 243.00 | 259.33 | -1.06 | 0.40 |
| 19.00 | 13.00 | 16.00 | 1.47 | 0.11 |
| 1.00 | 5.00 | 2.67 | 1.30 | 0.79 |
| 2.00 | 0.00 | 1.00 | -1.91 | 0.52 |
| 3.00 | 1.00 | 2.00 | 1.17 | 1.00 |
| 11.00 | 14.00 | 15.67 | 1.92 | 0.01 |
| 2.00 | 5.00 | 2.67 | -1.13 | 1.00 |
| 25.00 | 20.00 | 23.67 | -1.51 | 0.03 |
| 8.00 | 9.00 | 8.00 | -1.30 | 0.43 |

| | | | | |
|---|---|---|---|---|
| 3.00 | 2.00 | 2.00 | -1.33 | 0.79 |
| 25.00 | 17.00 | 19.33 | -1.34 | 0.13 |
| 15.00 | 8.00 | 9.33 | 1.53 | 0.20 |
| 4.00 | 1.00 | 2.00 | -3.08 | 0.02 |
| 136.00 | 158.00 | 131.00 | -1.53 | 0.00 |
| 10.00 | 7.00 | 9.33 | 1.10 | 0.79 |
| 8.00 | 1.00 | 4.00 | -1.59 | 0.30 |
| 9.00 | 1.00 | 4.33 | -1.62 | 0.25 |
| 1.00 | 0.00 | 0.67 | -2.72 | 0.29 |
| 0.00 | 0.00 | 0.67 | -3.58 | 0.12 |
| 8.00 | 9.00 | 6.33 | -1.43 | 0.32 |
| 13.00 | 6.00 | 9.67 | 1.30 | 0.42 |
| 1.00 | 1.00 | 1.33 | -1.01 | 1.00 |
| 3.00 | 4.00 | 3.67 | -1.90 | 0.12 |
| 1.00 | 7.00 | 4.00 | -1.18 | 0.85 |
| 1.00 | 1.00 | 2.00 | -1.49 | 0.61 |
| 9.00 | 21.00 | 13.00 | -1.17 | 0.67 |
| 55.00 | 60.00 | 62.67 | -1.76 | 0.00 |
| 829.00 | 986.00 | 965.67 | -1.17 | 0.00 |
| 1808.00 | 2174.00 | 2077.67 | 1.09 | 0.05 |
| 1.00 | 0.00 | 0.67 | -2.29 | 0.46 |
| 0.00 | 0.00 | 0.00 | -25.19 | 0.00 |
| 0.00 | 1.00 | 0.33 | -6.16 | 0.04 |
| 5.00 | 0.00 | 2.00 | -1.49 | 0.62 |
| 5.00 | 4.00 | 4.67 | -1.72 | 0.20 |
| 3.00 | 6.00 | 4.67 | 1.51 | 0.41 |
| 4.00 | 2.00 | 3.00 | -1.23 | 0.83 |
| 19.00 | 15.00 | 15.67 | 1.54 | 0.08 |
| 17.00 | 10.00 | 10.33 | -1.43 | 0.19 |
| 6.00 | 6.00 | 5.33 | 1.93 | 0.16 |
| 10.00 | 5.00 | 8.00 | -1.68 | 0.07 |
| 3.00 | 2.00 | 1.67 | -1.20 | 1.00 |
| 2.00 | 0.00 | 1.67 | -3.10 | 0.03 |
| 27.00 | 30.00 | 29.00 | -1.49 | 0.01 |
| 188.00 | 185.00 | 188.00 | -1.16 | 0.04 |
| 38.00 | 28.00 | 34.33 | -1.37 | 0.03 |
| 5.00 | 18.00 | 11.67 | -1.59 | 0.04 |
| 25.00 | 13.00 | 15.67 | -1.57 | 0.04 |
| 19.00 | 15.00 | 18.67 | 1.02 | 0.93 |
| 10.00 | 5.00 | 6.33 | -2.32 | 0.00 |
| 19.00 | 24.00 | 22.33 | -1.04 | 0.87 |
| 10.00 | 9.00 | 8.67 | -2.97 | 0.00 |
| 9.00 | 10.00 | 9.67 | -2.19 | 0.00 |
| 56.00 | 46.00 | 57.00 | -1.16 | 0.22 |
| 2.00 | 3.00 | 3.00 | -1.12 | 1.00 |
| 0.00 | 0.00 | 0.00 | -11.75 | 0.13 |
| 1.00 | 1.00 | 1.67 | -1.96 | 0.31 |
| 2.00 | 0.00 | 1.00 | -5.82 | 0.00 |

| | | | | |
|---|---|---|---|---|
| 10.00 | 3.00 | 5.00 | 1.83 | 0.25 |
| 2.00 | 1.00 | 1.33 | -1.24 | 1.00 |
| 3.00 | 3.00 | 3.00 | -1.55 | 0.41 |
| 7.00 | 4.00 | 5.00 | -1.34 | 0.51 |
| 2.00 | 0.00 | 0.67 | -2.71 | 0.30 |
| 20.00 | 20.00 | 19.67 | -1.30 | 0.18 |
| 2.00 | 1.00 | 1.00 | -3.71 | 0.04 |
| 6.00 | 2.00 | 4.00 | -1.34 | 0.58 |
| 4.00 | 2.00 | 4.00 | -1.42 | 0.47 |
| 3.00 | 7.00 | 7.00 | 1.70 | 0.17 |
| 57.00 | 54.00 | 54.33 | -2.02 | 0.00 |
| 7.00 | 14.00 | 9.33 | -1.26 | 0.47 |
| 6.00 | 4.00 | 4.00 | -2.16 | 0.04 |
| 2.00 | 5.00 | 2.67 | -2.58 | 0.03 |
| 0.00 | 0.00 | 0.00 | -11.76 | 0.14 |
| 5.00 | 5.00 | 4.33 | -1.32 | 0.59 |
| 3.00 | 0.00 | 2.33 | -1.56 | 0.49 |
| 25.00 | 45.00 | 42.67 | -1.03 | 0.84 |
| 2.00 | 4.00 | 3.67 | 1.52 | 0.49 |
| 10.00 | 10.00 | 12.67 | 1.17 | 0.57 |
| 5.00 | 5.00 | 5.67 | -1.25 | 0.63 |
| 56.00 | 50.00 | 48.33 | 1.06 | 0.78 |
| 6.00 | 3.00 | 3.67 | -1.55 | 0.36 |
| 1.00 | 0.00 | 0.33 | -3.95 | 0.23 |
| 1.00 | 0.00 | 1.00 | -2.51 | 0.24 |
| 1.00 | 2.00 | 3.33 | -1.60 | 0.34 |
| 1.00 | 2.00 | 2.33 | -2.53 | 0.05 |
| 8.00 | 11.00 | 10.00 | -1.18 | 0.63 |
| 9.00 | 4.00 | 9.00 | -1.12 | 0.81 |
| 116.00 | 127.00 | 115.33 | -1.22 | 0.03 |
| 210.00 | 233.00 | 238.33 | -1.61 | 0.00 |
| 37.00 | 55.00 | 49.33 | -1.28 | 0.06 |
| 3.00 | 6.00 | 4.00 | -3.64 | 0.00 |
| 1.00 | 1.00 | 1.00 | -3.11 | 0.10 |
| 18.00 | 2.00 | 10.33 | -1.20 | 0.66 |
| 1315.00 | 1550.00 | 1491.00 | -1.04 | 0.40 |
| 371.00 | 454.00 | 443.33 | 1.19 | 0.00 |
| 896.00 | 1101.00 | 983.67 | 1.02 | 0.75 |
| 535.00 | 639.00 | 620.67 | 1.10 | 0.08 |
| 140.00 | 150.00 | 147.67 | -1.32 | 0.00 |
| 2.00 | 2.00 | 2.33 | 1.35 | 0.78 |
| 5094.00 | 5082.00 | 5059.00 | -1.00 | 1.00 |
| 1147.00 | 1332.00 | 1303.00 | 1.11 | 0.02 |
| 502.00 | 501.00 | 501.33 | -1.07 | 0.18 |
| 594.00 | 539.00 | 560.33 | -1.11 | 0.08 |
| 2090.00 | 2282.00 | 2199.67 | -1.00 | 0.92 |
| 102.00 | 88.00 | 92.00 | -1.00 | 1.00 |
| 532.00 | 587.00 | 593.33 | -1.13 | 0.01 |

| | | | | |
|---|---|---|---|---|
| 1349.00 | 1313.00 | 1383.00 | -1.39 | 0.00 |
| 1776.00 | 1899.00 | 1920.67 | -1.03 | 0.48 |
| 941.00 | 967.00 | 1003.00 | -1.03 | 0.52 |
| 794.00 | 809.00 | 826.00 | -1.08 | 0.08 |
| 141.00 | 144.00 | 137.00 | 1.15 | 0.11 |
| 4973.00 | 4631.00 | 4722.00 | -1.24 | 0.00 |
| 411.00 | 434.00 | 436.00 | 1.22 | 0.00 |
| 2792.00 | 2942.00 | 3013.33 | 1.09 | 0.02 |
| 877.00 | 962.00 | 946.33 | 1.12 | 0.01 |
| 654.00 | 759.00 | 735.00 | -1.07 | 0.17 |
| 512.00 | 521.00 | 535.00 | -1.14 | 0.01 |
| 2485.00 | 2544.00 | 2544.33 | 1.05 | 0.17 |
| 1839.00 | 2056.00 | 1997.33 | 1.03 | 0.39 |
| 132.00 | 139.00 | 148.33 | 1.07 | 0.41 |
| 148.00 | 181.00 | 159.33 | 1.17 | 0.05 |
| 3235.00 | 3627.00 | 3407.67 | -1.09 | 0.03 |
| 26.00 | 21.00 | 26.00 | 1.16 | 0.42 |
| 4.00 | 7.00 | 4.33 | -1.47 | 0.39 |
| 8719.00 | 8893.00 | 9139.67 | -1.44 | 0.00 |
| 987.00 | 986.00 | 1054.67 | 1.23 | 0.00 |
| 1212.00 | 1246.00 | 1306.00 | 1.17 | 0.00 |
| 1895.00 | 1868.00 | 1937.33 | -1.31 | 0.00 |
| 235.00 | 212.00 | 230.33 | -1.02 | 0.75 |
| 67.00 | 68.00 | 68.33 | 1.48 | 0.00 |
| 265.00 | 280.00 | 287.33 | 1.10 | 0.12 |
| 262.00 | 281.00 | 265.67 | 1.02 | 0.74 |
| 2141.00 | 2405.00 | 2397.33 | 1.19 | 0.00 |
| 275.00 | 304.00 | 289.67 | -1.01 | 0.85 |
| 10640.00 | 10682.00 | 10668.33 | 1.42 | 0.00 |
| 993.00 | 965.00 | 965.67 | 1.24 | 0.00 |
| 842639.00 | 936150.00 | 906546.67 | -1.04 | 0.23 |
| 15.00 | 11.00 | 13.67 | 1.19 | 0.59 |
| 5.00 | 10.00 | 6.67 | 1.16 | 0.75 |
| 3.00 | 1.00 | 3.00 | -1.55 | 0.41 |
| 14950.00 | 15961.00 | 15567.67 | -1.18 | 0.00 |
| 10.00 | 8.00 | 10.33 | -1.14 | 0.72 |
| 395.00 | 411.00 | 378.33 | -1.13 | 0.10 |
| 276.00 | 309.00 | 289.67 | -1.23 | 0.00 |
| 178.00 | 187.00 | 179.00 | -1.17 | 0.04 |
| 1998.00 | 1897.00 | 2001.00 | -1.94 | 0.00 |
| 653.00 | 723.00 | 728.67 | -1.15 | 0.00 |
| 17.00 | 18.00 | 19.00 | -1.72 | 0.00 |
| 82.00 | 101.00 | 84.33 | -1.29 | 0.01 |
| 2406.00 | 2686.00 | 2556.67 | -1.80 | 0.00 |
| 2997.00 | 3256.00 | 3231.67 | -1.02 | 0.50 |
| 650.00 | 770.00 | 715.00 | 1.22 | 0.00 |
| 2.00 | 1.00 | 1.33 | -1.48 | 0.76 |
| 791.00 | 928.00 | 901.33 | 1.27 | 0.00 |

| | | | | |
|---|---|---|---|---|
| 138.00 | 215.00 | 177.67 | 1.51 | 0.00 |
| 132.00 | 124.00 | 126.00 | 1.44 | 0.00 |
| 1072.00 | 1119.00 | 1102.67 | -1.00 | 0.98 |
| 760.00 | 747.00 | 751.33 | 1.09 | 0.09 |
| 691.00 | 784.00 | 720.33 | 1.23 | 0.00 |
| 331.00 | 358.00 | 376.67 | -1.01 | 0.93 |
| 810.00 | 813.00 | 827.00 | -1.26 | 0.00 |
| 16.00 | 6.00 | 13.00 | 1.32 | 0.30 |
| 233.00 | 217.00 | 233.33 | -1.11 | 0.19 |
| 30.00 | 42.00 | 39.00 | 1.07 | 0.67 |
| 1205.00 | 1217.00 | 1200.00 | -1.42 | 0.00 |
| 492.00 | 504.00 | 524.33 | -1.09 | 0.10 |
| 1034.00 | 1175.00 | 1173.33 | 1.18 | 0.00 |
| 385.00 | 392.00 | 414.00 | 1.01 | 0.87 |
| 0.00 | 0.00 | 0.00 | -11.74 | 0.13 |
| 3570.00 | 3569.00 | 3582.00 | -1.19 | 0.00 |
| 23872.00 | 26037.00 | 26647.67 | 1.24 | 0.00 |
| 418.00 | 411.00 | 431.00 | -1.10 | 0.08 |
| 877.00 | 880.00 | 887.33 | -1.20 | 0.00 |
| 1028.00 | 1135.00 | 1082.00 | -1.03 | 0.53 |
| 1987.00 | 2041.00 | 2070.33 | -1.08 | 0.02 |
| 641.00 | 706.00 | 658.00 | 1.07 | 0.19 |
| 567.00 | 683.00 | 658.33 | 1.17 | 0.00 |
| 2300.00 | 2515.00 | 2507.00 | 1.12 | 0.00 |
| 7589.00 | 8902.00 | 8664.00 | -1.01 | 0.84 |
| 117.00 | 137.00 | 136.67 | -1.07 | 0.38 |
| 125.00 | 133.00 | 132.67 | -1.19 | 0.04 |
| 189.00 | 216.00 | 209.00 | 1.28 | 0.00 |
| 1.00 | 2.00 | 2.00 | -2.13 | 0.17 |
| 469.00 | 436.00 | 458.00 | -1.16 | 0.00 |
| 291.00 | 330.00 | 334.00 | 1.01 | 0.86 |
| 338.00 | 344.00 | 339.67 | -1.12 | 0.04 |
| 281.00 | 275.00 | 294.33 | 1.02 | 0.76 |
| 1219.00 | 1231.00 | 1250.00 | -1.17 | 0.00 |
| 1641.00 | 1789.00 | 1716.67 | 1.33 | 0.00 |
| 5.00 | 5.00 | 3.33 | -2.46 | 0.03 |
| 471.00 | 575.00 | 558.33 | -1.02 | 0.69 |
| 371.00 | 430.00 | 412.00 | 1.00 | 0.98 |
| 1296.00 | 1301.00 | 1398.67 | 1.03 | 0.65 |
| 3379.00 | 3530.00 | 3417.33 | -1.22 | 0.00 |
| 1084.00 | 1070.00 | 1091.33 | -1.14 | 0.00 |
| 859.00 | 1098.00 | 1003.33 | 1.03 | 0.53 |
| 861.00 | 1100.00 | 1026.33 | 1.11 | 0.06 |
| 933.00 | 1065.00 | 1000.33 | -1.13 | 0.01 |
| 583.00 | 593.00 | 558.33 | 1.01 | 0.90 |
| 516.00 | 552.00 | 557.67 | -1.03 | 0.54 |
| 5.00 | 5.00 | 6.33 | 1.84 | 0.14 |
| 913.00 | 949.00 | 964.33 | 1.18 | 0.00 |

| | | | | |
|---|---|---|---|---|
| 121.00 | 131.00 | 127.00 | 1.10 | 0.26 |
| 167.00 | 122.00 | 157.00 | 1.56 | 0.00 |
| 60.00 | 39.00 | 48.33 | 1.17 | 0.32 |
| 855.00 | 847.00 | 857.67 | -1.19 | 0.00 |
| 1252.00 | 1448.00 | 1342.33 | 1.02 | 0.65 |
| 1028.00 | 1122.00 | 1042.67 | 1.22 | 0.00 |
| 1167.00 | 1168.00 | 1181.67 | 1.19 | 0.00 |
| 180.00 | 214.00 | 183.33 | -1.06 | 0.55 |
| 430.00 | 469.00 | 466.00 | 1.05 | 0.31 |
| 6859.00 | 6784.00 | 6762.67 | -1.27 | 0.00 |
| 593.00 | 593.00 | 610.33 | 1.01 | 0.80 |
| 1079.00 | 1107.00 | 1164.67 | -1.30 | 0.00 |
| 351.00 | 349.00 | 365.00 | 1.04 | 0.46 |
| 20.00 | 13.00 | 15.67 | -2.53 | 0.00 |
| 2739.00 | 2535.00 | 2657.67 | -1.09 | 0.05 |
| 235.00 | 237.00 | 252.00 | -1.33 | 0.00 |
| 6.00 | 15.00 | 11.33 | 1.95 | 0.03 |
| 84.00 | 63.00 | 75.33 | -1.01 | 0.97 |
| 8.00 | 6.00 | 6.00 | 1.36 | 0.48 |
| 465.00 | 450.00 | 453.00 | -1.46 | 0.00 |
| 1189.00 | 1500.00 | 1353.67 | 1.09 | 0.09 |
| 89.00 | 129.00 | 115.00 | -1.09 | 0.35 |
| 2752.00 | 2898.00 | 2987.33 | 1.19 | 0.00 |
| 718.00 | 778.00 | 798.67 | 1.07 | 0.18 |
| 1123.00 | 1173.00 | 1188.00 | 1.17 | 0.00 |
| 1218.00 | 1437.00 | 1349.00 | -1.05 | 0.29 |
| 18.00 | 7.00 | 12.67 | -1.30 | 0.31 |
| 1310.00 | 1009.00 | 1191.33 | -1.56 | 0.00 |
| 5.00 | 7.00 | 7.67 | 1.19 | 0.65 |
| 406.00 | 504.00 | 475.33 | 1.16 | 0.01 |
| 1016.00 | 1076.00 | 1118.67 | 1.07 | 0.17 |
| 1747.00 | 1899.00 | 1876.33 | 1.21 | 0.00 |
| 1847.00 | 2061.00 | 2077.33 | 1.28 | 0.00 |
| 972.00 | 1189.00 | 1153.33 | 1.18 | 0.00 |
| 602.00 | 676.00 | 679.33 | 1.08 | 0.17 |
| 1435.00 | 1572.00 | 1600.00 | 1.15 | 0.00 |
| 691.00 | 775.00 | 782.67 | 1.15 | 0.01 |
| 1254.00 | 1372.00 | 1362.33 | 1.28 | 0.00 |
| 917.00 | 920.00 | 953.67 | 1.18 | 0.00 |
| 1171.00 | 1282.00 | 1270.33 | 1.19 | 0.00 |
| 801.00 | 840.00 | 865.33 | 1.18 | 0.00 |
| 813.00 | 860.00 | 915.00 | 1.16 | 0.01 |
| 812.00 | 888.00 | 893.00 | 1.27 | 0.00 |
| 482.00 | 497.00 | 510.33 | 1.18 | 0.00 |
| 63.00 | 73.00 | 81.33 | 1.05 | 0.71 |
| 2059.00 | 2318.00 | 2279.33 | 1.11 | 0.01 |
| 599.00 | 682.00 | 683.00 | 1.06 | 0.29 |
| 808.00 | 961.00 | 923.67 | 1.06 | 0.22 |

| | | | | |
|---|---|---|---|---|
| 3062.00 | 3549.00 | 3506.00 | 1.17 | 0.00 |
| 1410.00 | 1421.00 | 1472.67 | 1.11 | 0.02 |
| 812.00 | 1012.00 | 941.00 | 1.09 | 0.09 |
| 1171.00 | 1303.00 | 1357.33 | 1.11 | 0.06 |
| 633.00 | 696.00 | 743.00 | 1.18 | 0.02 |
| 868.00 | 943.00 | 927.67 | 1.12 | 0.01 |
| 555.00 | 624.00 | 622.67 | 1.14 | 0.01 |
| 1786.00 | 1892.00 | 1908.67 | 1.11 | 0.01 |
| 1070.00 | 1189.00 | 1177.33 | 1.10 | 0.02 |
| 515.00 | 566.00 | 579.67 | 1.14 | 0.03 |
| 1595.00 | 1731.00 | 1755.67 | 1.09 | 0.03 |
| 608.00 | 659.00 | 658.00 | 1.14 | 0.01 |
| 508.00 | 562.00 | 568.67 | 1.13 | 0.02 |
| 985.00 | 1203.00 | 1112.33 | 1.03 | 0.56 |
| 879.00 | 947.00 | 965.67 | 1.01 | 0.89 |
| 649.00 | 736.00 | 700.67 | 1.20 | 0.00 |
| 112.00 | 136.00 | 134.00 | 1.20 | 0.05 |
| 57.00 | 68.00 | 63.00 | 1.05 | 0.70 |
| 397.00 | 509.00 | 474.00 | 1.09 | 0.18 |
| 1035.00 | 1130.00 | 1138.67 | 1.27 | 0.00 |
| 369.00 | 422.00 | 428.00 | 1.43 | 0.00 |
| 1237.00 | 1223.00 | 1260.00 | -1.01 | 0.82 |
| 465.00 | 487.00 | 503.00 | 1.27 | 0.00 |
| 1875.00 | 2038.00 | 2095.33 | 1.15 | 0.00 |
| 1294.00 | 1390.00 | 1406.00 | 1.23 | 0.00 |
| 358.00 | 455.00 | 454.33 | 1.21 | 0.02 |
| 363.00 | 354.00 | 399.67 | 1.26 | 0.00 |
| 333.00 | 342.00 | 356.00 | 1.04 | 0.54 |
| 627.00 | 631.00 | 656.00 | 1.12 | 0.03 |
| 1465.00 | 1558.00 | 1593.67 | 1.17 | 0.00 |
| 1512.00 | 1533.00 | 1568.67 | 1.09 | 0.03 |
| 431.00 | 466.00 | 462.33 | -1.11 | 0.03 |
| 619.00 | 635.00 | 663.33 | 1.04 | 0.53 |
| 421.00 | 440.00 | 453.67 | -1.08 | 0.12 |
| 821.00 | 933.00 | 924.00 | 1.14 | 0.00 |
| 2954.00 | 2854.00 | 3006.33 | 1.09 | 0.03 |
| 329.00 | 380.00 | 389.33 | 1.10 | 0.15 |
| 557.00 | 570.00 | 578.67 | 1.28 | 0.00 |
| 207.00 | 242.00 | 248.33 | 1.13 | 0.15 |
| 1360.00 | 1554.00 | 1542.33 | 1.04 | 0.38 |
| 1407.00 | 1603.00 | 1641.33 | 1.08 | 0.12 |
| 1050.00 | 1224.00 | 1169.33 | 1.10 | 0.02 |
| 1149.00 | 1205.00 | 1213.00 | 1.20 | 0.00 |
| 86.00 | 76.00 | 95.00 | 1.22 | 0.10 |
| 2400.00 | 2574.00 | 2570.67 | -1.11 | 0.00 |
| 962.00 | 1031.00 | 1018.67 | 1.22 | 0.00 |
| 1771.00 | 1843.00 | 1877.33 | -1.02 | 0.59 |
| 470.00 | 569.00 | 564.00 | 1.14 | 0.03 |

| | | | | |
|---|---|---|---|---|
| 892.00 | 962.00 | 953.67 | 1.21 | 0.00 |
| 390.00 | 451.00 | 429.33 | -1.09 | 0.09 |
| 218.00 | 234.00 | 227.67 | -1.10 | 0.13 |
| 829.00 | 1012.00 | 981.33 | 1.11 | 0.06 |
| 1661.00 | 1793.00 | 1811.33 | 1.23 | 0.00 |
| 1328.00 | 1460.00 | 1484.33 | 1.15 | 0.00 |
| 91.00 | 89.00 | 106.00 | 1.00 | 1.00 |
| 1751.00 | 1946.00 | 1980.67 | 1.05 | 0.27 |
| 870.00 | 1039.00 | 985.33 | 1.37 | 0.00 |
| 805.00 | 967.00 | 929.33 | 1.22 | 0.00 |
| 1036.00 | 1192.00 | 1150.33 | 1.32 | 0.00 |
| 706.00 | 868.00 | 833.67 | 1.02 | 0.73 |
| 942.00 | 918.00 | 974.33 | 1.06 | 0.16 |
| 1374.00 | 1578.00 | 1574.67 | 1.16 | 0.00 |
| 3.00 | 8.00 | 8.00 | -1.18 | 0.69 |
| 562.00 | 652.00 | 671.67 | -1.30 | 0.00 |
| 1150.00 | 1301.00 | 1274.33 | 1.10 | 0.03 |
| 448.00 | 506.00 | 477.33 | -1.03 | 0.67 |
| 26.00 | 20.00 | 18.67 | 1.53 | 0.08 |
| 65.00 | 81.00 | 75.00 | -1.24 | 0.04 |
| 1206.00 | 1574.00 | 1423.67 | 1.09 | 0.10 |
| 449.00 | 415.00 | 426.67 | -1.05 | 0.45 |
| 12.00 | 13.00 | 15.67 | -1.19 | 0.44 |
| 1734.00 | 1858.00 | 1808.33 | 1.00 | 0.93 |
| 359.00 | 372.00 | 351.33 | -1.06 | 0.39 |
| 1839.00 | 2171.00 | 2042.00 | -1.02 | 0.64 |
| 909.00 | 997.00 | 959.33 | 1.14 | 0.03 |
| 1136.00 | 1321.00 | 1251.00 | -1.13 | 0.00 |
| 322.00 | 361.00 | 349.00 | -1.05 | 0.38 |
| 638.00 | 714.00 | 686.00 | 1.01 | 0.79 |
| 4.00 | 3.00 | 3.00 | 1.11 | 1.00 |
| 21783.00 | 22160.00 | 22031.00 | -1.11 | 0.00 |
| 29.00 | 31.00 | 32.00 | 1.11 | 0.57 |
| 152.00 | 156.00 | 175.00 | 1.20 | 0.05 |
| 194.00 | 179.00 | 191.33 | 1.08 | 0.31 |
| 362.00 | 373.00 | 392.33 | -1.01 | 0.93 |
| 948.00 | 872.00 | 908.67 | 1.06 | 0.25 |
| 370.00 | 371.00 | 374.67 | 1.07 | 0.24 |
| 138.00 | 117.00 | 141.00 | -1.21 | 0.03 |
| 16.00 | 8.00 | 11.33 | -1.45 | 0.14 |
| 82.00 | 75.00 | 86.00 | -1.16 | 0.13 |
| 4.00 | 7.00 | 4.67 | 2.23 | 0.12 |
| 0.00 | 0.00 | 0.33 | -5.43 | 0.08 |
| 90.00 | 89.00 | 90.67 | -1.04 | 0.73 |
| 96.00 | 105.00 | 111.00 | -1.04 | 0.72 |
| 88.00 | 101.00 | 101.67 | -1.12 | 0.21 |
| 2397.00 | 2562.00 | 2687.67 | 1.19 | 0.00 |
| 1.00 | 0.00 | 0.33 | -3.21 | 0.38 |

| | | | | |
|---|---|---|---|---|
| 216.00 | 238.00 | 249.33 | 1.25 | 0.00 |
| 4061.00 | 4276.00 | 4567.33 | -1.46 | 0.00 |
| 2751.00 | 2677.00 | 2788.33 | 1.02 | 0.53 |
| 2670.00 | 2940.00 | 2898.00 | 1.05 | 0.24 |
| 631.00 | 768.00 | 732.67 | 1.08 | 0.14 |
| 3196.00 | 3527.00 | 3409.00 | 1.15 | 0.00 |
| 594.00 | 795.00 | 685.33 | 1.28 | 0.00 |
| 4753.00 | 4929.00 | 5041.67 | -1.04 | 0.31 |
| 1843.00 | 1880.00 | 1972.00 | 1.21 | 0.00 |
| 4163.00 | 4109.00 | 4152.33 | -1.05 | 0.19 |
| 98.00 | 154.00 | 139.67 | -1.03 | 0.78 |
| 8202.00 | 7961.00 | 7933.00 | 1.09 | 0.04 |
| 311.00 | 416.00 | 377.67 | 1.04 | 0.61 |
| 198.00 | 294.00 | 261.67 | 1.05 | 0.60 |
| 898.00 | 1060.00 | 1025.00 | 1.09 | 0.05 |
| 993.00 | 1061.00 | 1057.00 | -1.08 | 0.05 |
| 477.00 | 440.00 | 488.00 | -1.21 | 0.00 |
| 641.00 | 714.00 | 725.33 | 1.17 | 0.00 |
| 246.00 | 327.00 | 303.33 | 1.05 | 0.59 |
| 445.00 | 431.00 | 467.00 | 1.22 | 0.00 |
| 710.00 | 815.00 | 810.67 | 1.19 | 0.00 |
| 719.00 | 713.00 | 763.33 | -1.04 | 0.49 |
| 155.00 | 202.00 | 187.67 | 1.27 | 0.00 |
| 1405.00 | 1602.00 | 1560.00 | -1.01 | 0.82 |
| 1146.00 | 1100.00 | 1156.33 | -1.14 | 0.01 |
| 3044.00 | 3454.00 | 3354.00 | 1.15 | 0.00 |
| 3221.00 | 3405.00 | 3411.33 | 1.08 | 0.03 |
| 5859.00 | 6461.00 | 6411.67 | 1.20 | 0.00 |
| 142.00 | 178.00 | 163.00 | 1.36 | 0.00 |
| 214.00 | 239.00 | 223.67 | -1.02 | 0.77 |
| 52.00 | 51.00 | 53.00 | 1.62 | 0.00 |
| 651.00 | 734.00 | 723.33 | -1.08 | 0.12 |
| 1826.00 | 2109.00 | 2077.67 | 1.13 | 0.00 |
| 368.00 | 457.00 | 446.00 | 1.08 | 0.25 |
| 102.00 | 107.00 | 105.67 | 1.07 | 0.50 |
| 147.00 | 181.00 | 166.33 | -1.15 | 0.06 |
| 1565.00 | 1759.00 | 1736.67 | 1.08 | 0.05 |
| 32.00 | 33.00 | 29.33 | -1.01 | 1.00 |
| 292.00 | 308.00 | 318.67 | 1.00 | 0.99 |
| 960.00 | 1025.00 | 987.67 | 1.18 | 0.00 |
| 761.00 | 694.00 | 759.33 | 1.05 | 0.35 |
| 1930.00 | 1986.00 | 2029.00 | 1.07 | 0.10 |
| 1000.00 | 965.00 | 984.00 | -1.13 | 0.00 |
| 1685.00 | 1753.00 | 1773.00 | 1.00 | 0.93 |
| 785.00 | 854.00 | 791.67 | 1.22 | 0.01 |
| 0.00 | 2.00 | 1.67 | -4.03 | 0.00 |
| 520.00 | 485.00 | 496.33 | -1.02 | 0.79 |
| 75.00 | 61.00 | 70.67 | -2.48 | 0.00 |

| | | | | |
|---|---|---|---|---|
| 918.00 | 1069.00 | 1037.33 | 1.04 | 0.36 |
| 2264.00 | 2276.00 | 2267.67 | -1.09 | 0.03 |
| 888.00 | 1071.00 | 998.00 | 1.12 | 0.01 |
| 1106.00 | 1267.00 | 1200.33 | -1.00 | 0.95 |
| 530.00 | 550.00 | 567.33 | -1.04 | 0.48 |
| 324.00 | 347.00 | 340.00 | -1.06 | 0.43 |
| 12.00 | 13.00 | 17.00 | 1.29 | 0.32 |
| 1223.00 | 1326.00 | 1267.67 | -1.10 | 0.02 |
| 121.00 | 84.00 | 110.33 | -1.19 | 0.07 |
| 877.00 | 843.00 | 887.33 | -1.49 | 0.00 |
| 566.00 | 631.00 | 602.00 | -1.10 | 0.05 |
| 22.00 | 31.00 | 21.33 | -1.25 | 0.32 |
| 282.00 | 330.00 | 321.00 | 1.08 | 0.25 |
| 962.00 | 1183.00 | 1137.33 | 1.16 | 0.00 |
| 1115.00 | 1184.00 | 1164.00 | -1.02 | 0.57 |
| 61.00 | 57.00 | 61.00 | -1.27 | 0.05 |
| 674.00 | 701.00 | 695.67 | -1.16 | 0.01 |
| 1302.00 | 1388.00 | 1398.67 | -1.04 | 0.31 |
| 98.00 | 105.00 | 102.33 | 1.18 | 0.10 |
| 54.00 | 72.00 | 69.00 | 1.10 | 0.44 |
| 2071.00 | 2225.00 | 2250.00 | 1.07 | 0.12 |
| 32.00 | 35.00 | 35.33 | -1.22 | 0.17 |
| 21.00 | 42.00 | 31.33 | -1.19 | 0.32 |
| 1088.00 | 1142.00 | 1149.67 | 1.02 | 0.66 |
| 916.00 | 1024.00 | 1027.33 | 1.09 | 0.13 |
| 647.00 | 726.00 | 706.00 | 1.09 | 0.05 |
| 314.00 | 332.00 | 320.00 | 1.01 | 0.87 |
| 706.00 | 703.00 | 746.67 | -1.04 | 0.41 |
| 125.00 | 146.00 | 143.67 | -1.09 | 0.25 |
| 1698.00 | 1841.00 | 1829.67 | -1.09 | 0.02 |
| 27.00 | 27.00 | 28.33 | 1.58 | 0.01 |
| 594.00 | 614.00 | 618.67 | 1.04 | 0.37 |
| 156.00 | 208.00 | 194.67 | -1.10 | 0.24 |
| 378.00 | 435.00 | 403.67 | -1.14 | 0.02 |
| 826.00 | 835.00 | 873.00 | 1.09 | 0.07 |
| 1957.00 | 1791.00 | 1929.33 | 1.02 | 0.62 |
| 2.00 | 1.00 | 1.67 | -1.39 | 0.78 |
| 597.00 | 602.00 | 598.67 | -1.06 | 0.38 |
| 25.00 | 31.00 | 29.33 | 2.69 | 0.00 |
| 4583.00 | 4402.00 | 4515.00 | -1.09 | 0.04 |
| 1050.00 | 1367.00 | 1182.33 | 1.32 | 0.00 |
| 470.00 | 579.00 | 558.00 | 1.54 | 0.00 |
| 6668.00 | 7061.00 | 7093.00 | 1.12 | 0.00 |
| 6.00 | 12.00 | 9.33 | 1.44 | 0.25 |
| 2501.00 | 2635.00 | 2523.00 | -1.18 | 0.00 |
| 4.00 | 6.00 | 5.67 | -1.30 | 0.53 |
| 213.00 | 236.00 | 230.00 | 1.03 | 0.76 |
| 325.00 | 332.00 | 346.67 | -1.07 | 0.21 |

| | | | | |
|---|---|---|---|---|
| 2728.00 | 2564.00 | 2728.33 | 1.05 | 0.34 |
| 1743.00 | 1968.00 | 1838.33 | 1.16 | 0.00 |
| 2022.00 | 2184.00 | 2153.67 | -1.63 | 0.00 |
| 429.00 | 425.00 | 450.00 | 1.20 | 0.00 |
| 1275.00 | 1262.00 | 1233.33 | -1.01 | 0.90 |
| 100.00 | 99.00 | 105.67 | 1.90 | 0.00 |
| 1.00 | 3.00 | 3.33 | -1.31 | 0.68 |
| 135.00 | 138.00 | 135.00 | -1.12 | 0.18 |
| 7.00 | 6.00 | 7.33 | 1.03 | 1.00 |
| 14519.00 | 14052.00 | 14043.00 | -1.57 | 0.00 |
| 6002.00 | 6626.00 | 6765.67 | 1.18 | 0.00 |
| 5563.00 | 6251.00 | 6251.00 | 1.21 | 0.00 |
| 54.00 | 60.00 | 61.67 | -1.10 | 0.40 |
| 3427.00 | 3954.00 | 3948.00 | 1.08 | 0.08 |
| 1142.00 | 1300.00 | 1318.33 | -1.07 | 0.13 |
| 108.00 | 118.00 | 113.67 | -1.23 | 0.01 |
| 3409.00 | 3290.00 | 3296.33 | -1.24 | 0.00 |
| 8.00 | 8.00 | 8.33 | 1.23 | 0.56 |
| 42.00 | 31.00 | 38.00 | -1.51 | 0.00 |
| 47.00 | 38.00 | 42.33 | 1.12 | 0.47 |
| 845.00 | 923.00 | 936.33 | 1.02 | 0.65 |
| 7959.00 | 7651.00 | 7764.33 | -1.09 | 0.05 |
| 4.00 | 0.00 | 3.33 | -1.40 | 0.55 |
| 29.00 | 20.00 | 24.67 | -1.28 | 0.18 |
| 631.00 | 565.00 | 592.00 | -1.22 | 0.00 |
| 1.00 | 15.00 | 8.67 | -1.59 | 0.26 |
| 2.00 | 0.00 | 0.67 | -4.42 | 0.05 |
| 2961.00 | 3150.00 | 3067.33 | -1.01 | 0.79 |
| 2401.00 | 2352.00 | 2394.33 | -1.08 | 0.04 |
| 172.00 | 152.00 | 168.00 | -1.31 | 0.00 |
| 897.00 | 846.00 | 873.67 | -1.21 | 0.00 |
| 2.00 | 1.00 | 2.33 | -1.70 | 0.37 |
| 1572.00 | 1790.00 | 1627.67 | -1.34 | 0.00 |
| 26.00 | 14.00 | 19.67 | -1.40 | 0.09 |
| 6.00 | 5.00 | 5.33 | 1.56 | 0.34 |
| 1334.00 | 1534.00 | 1422.00 | -1.09 | 0.08 |
| 15.00 | 22.00 | 19.33 | -1.47 | 0.03 |
| 394.00 | 365.00 | 352.67 | -1.29 | 0.00 |
| 3625.00 | 3430.00 | 3568.00 | -1.10 | 0.02 |
| 42709.00 | 41137.00 | 41155.00 | 1.06 | 0.21 |
| 27.00 | 23.00 | 24.33 | -1.63 | 0.00 |
| 0.00 | 1.00 | 1.33 | -4.25 | 0.01 |
| 366.00 | 390.00 | 390.00 | -1.80 | 0.00 |
| 183.00 | 173.00 | 183.33 | 1.11 | 0.15 |
| 67.00 | 30.00 | 49.00 | -4.35 | 0.00 |
| 10.00 | 10.00 | 13.67 | 1.22 | 0.51 |
| 88.00 | 94.00 | 91.33 | 1.02 | 0.81 |
| 2790.00 | 3440.00 | 3098.67 | 1.00 | 0.97 |

| | | | | |
|---|---|---|---|---|
| 2.00 | 2.00 | 1.33 | -2.17 | 0.27 |
| 735.00 | 732.00 | 758.33 | -1.19 | 0.00 |
| 116.00 | 108.00 | 114.33 | -1.02 | 0.84 |
| 573.00 | 774.00 | 673.00 | 1.38 | 0.00 |
| 480.00 | 590.00 | 555.33 | 1.19 | 0.00 |
| 1212.00 | 1326.00 | 1328.00 | 1.15 | 0.00 |
| 905.00 | 884.00 | 923.67 | 1.06 | 0.19 |
| 744.00 | 810.00 | 727.00 | 1.01 | 0.87 |
| 33.00 | 36.00 | 31.00 | 1.16 | 0.36 |
| 1672.00 | 2006.00 | 1879.00 | 1.01 | 0.81 |
| 218.00 | 244.00 | 227.33 | -1.16 | 0.07 |
| 235.00 | 267.00 | 260.00 | -1.17 | 0.01 |
| 125.00 | 119.00 | 128.67 | 1.04 | 0.68 |
| 1685.00 | 1880.00 | 1908.33 | 1.19 | 0.00 |
| 1592.00 | 1783.00 | 1663.67 | 1.13 | 0.01 |
| 406.00 | 512.00 | 460.33 | -1.10 | 0.12 |
| 1753.00 | 2017.00 | 1980.33 | 1.14 | 0.00 |
| 1849.00 | 1972.00 | 1884.67 | 1.01 | 0.78 |
| 511.00 | 608.00 | 555.33 | 1.11 | 0.11 |
| 625.00 | 663.00 | 636.00 | 1.07 | 0.21 |
| 933.00 | 959.00 | 1003.67 | 1.24 | 0.00 |
| 1232.00 | 1291.00 | 1294.67 | -1.02 | 0.62 |
| 3427.00 | 3844.00 | 3709.33 | 1.03 | 0.46 |
| 531.00 | 577.00 | 576.33 | -1.02 | 0.72 |
| 181.00 | 157.00 | 166.00 | -1.08 | 0.41 |
| 4.00 | 0.00 | 1.33 | -3.09 | 0.06 |
| 29.00 | 28.00 | 31.33 | -1.00 | 1.00 |
| 1193.00 | 1478.00 | 1396.00 | 1.11 | 0.03 |
| 967.00 | 1024.00 | 1077.00 | 1.54 | 0.00 |
| 2002.00 | 2265.00 | 2135.33 | -1.27 | 0.00 |
| 693.00 | 665.00 | 718.67 | 1.04 | 0.44 |
| 11729.00 | 13091.00 | 13271.00 | 1.16 | 0.00 |
| 2890.00 | 2664.00 | 2814.33 | -1.08 | 0.05 |
| 884.00 | 856.00 | 891.33 | -1.20 | 0.00 |
| 1100.00 | 1065.00 | 1109.00 | -1.06 | 0.18 |
| 788.00 | 978.00 | 924.33 | 1.11 | 0.06 |
| 6670.00 | 7201.00 | 7154.00 | -1.21 | 0.00 |
| 1848.00 | 2151.00 | 2064.67 | 1.16 | 0.00 |
| 275.00 | 271.00 | 280.67 | 1.03 | 0.73 |
| 792.00 | 756.00 | 778.00 | -1.15 | 0.00 |
| 409.00 | 454.00 | 458.67 | -1.18 | 0.00 |
| 1274.00 | 1416.00 | 1390.33 | -1.01 | 0.79 |
| 905.00 | 1006.00 | 970.67 | -1.04 | 0.33 |
| 4.00 | 2.00 | 2.33 | -2.79 | 0.02 |
| 81.00 | 72.00 | 74.00 | -1.09 | 0.44 |
| 274.00 | 250.00 | 270.00 | -1.14 | 0.03 |
| 12.00 | 4.00 | 9.67 | 1.24 | 0.50 |
| 513.00 | 443.00 | 474.00 | -1.31 | 0.00 |

| | | | | |
|---|---|---|---|---|
| 4035.00 | 4730.00 | 4481.33 | -1.02 | 0.68 |
| 0.00 | 1.00 | 0.33 | -5.43 | 0.08 |
| 375.00 | 411.00 | 391.00 | -1.01 | 0.89 |
| 657.00 | 647.00 | 689.00 | 1.05 | 0.35 |
| 15387.00 | 17412.00 | 16973.33 | 1.10 | 0.01 |
| 24.00 | 32.00 | 32.33 | 1.16 | 0.41 |
| 59.00 | 82.00 | 72.00 | 1.11 | 0.37 |
| 203.00 | 235.00 | 231.67 | 1.14 | 0.06 |
| 7.00 | 8.00 | 7.67 | -1.01 | 1.00 |
| 889.00 | 915.00 | 929.33 | 1.01 | 0.90 |
| 12.00 | 11.00 | 14.67 | 1.40 | 0.18 |
| 432.00 | 532.00 | 493.33 | -1.12 | 0.03 |
| 4.00 | 6.00 | 5.33 | -1.76 | 0.11 |
| 1156.00 | 1401.00 | 1322.33 | 1.00 | 0.99 |
| 15.00 | 11.00 | 12.33 | 1.01 | 1.00 |
| 641.00 | 759.00 | 736.00 | -1.04 | 0.46 |
| 529.00 | 528.00 | 575.67 | -1.02 | 0.72 |
| 6972.00 | 8256.00 | 7742.67 | 1.06 | 0.12 |
| 523.00 | 540.00 | 528.33 | 1.04 | 0.52 |
| 3099.00 | 3663.00 | 3485.33 | 1.20 | 0.00 |
| 50.00 | 50.00 | 52.00 | 1.19 | 0.18 |
| 2315.00 | 2478.00 | 2410.00 | -1.05 | 0.16 |
| 599.00 | 604.00 | 612.33 | 1.00 | 0.98 |
| 0.00 | 0.00 | 0.00 | -14.45 | 0.07 |
| 164.00 | 178.00 | 187.67 | -1.11 | 0.18 |
| 1666.00 | 1690.00 | 1673.67 | -1.01 | 0.88 |
| 1744.00 | 1801.00 | 1890.33 | -1.07 | 0.12 |
| 23.00 | 16.00 | 19.67 | 1.04 | 0.86 |
| 2746.00 | 2733.00 | 2718.33 | -1.65 | 0.00 |
| 3107.00 | 2977.00 | 3019.00 | -1.54 | 0.00 |
| 3786.00 | 3999.00 | 3867.67 | 1.10 | 0.03 |
| 1505.00 | 1618.00 | 1548.67 | -1.01 | 0.85 |
| 350.00 | 331.00 | 345.00 | -1.29 | 0.00 |
| 84.00 | 89.00 | 85.67 | -1.03 | 0.81 |
| 2599.00 | 2551.00 | 2670.00 | -1.19 | 0.00 |
| 11.00 | 11.00 | 11.00 | -1.26 | 0.43 |
| 1852.00 | 2099.00 | 2002.33 | 1.18 | 0.00 |
| 366.00 | 356.00 | 342.00 | -1.22 | 0.02 |
| 53.00 | 58.00 | 57.00 | -1.36 | 0.02 |
| 312.00 | 332.00 | 327.33 | -1.14 | 0.01 |
| 412.00 | 421.00 | 422.00 | -1.20 | 0.00 |
| 411.00 | 453.00 | 404.67 | -1.03 | 0.71 |
| 2006.00 | 2238.00 | 2149.33 | 1.01 | 0.77 |
| 189.00 | 216.00 | 213.67 | 1.04 | 0.59 |
| 29.00 | 33.00 | 26.67 | -1.24 | 0.24 |
| 325.00 | 333.00 | 315.00 | -1.00 | 0.97 |
| 1.00 | 0.00 | 0.33 | -4.69 | 0.13 |
| 3.00 | 12.00 | 5.00 | 1.35 | 0.65 |

| | | | | |
|---|---|---|---|---|
| 1412.00 | 1588.00 | 1520.33 | 1.07 | 0.07 |
| 1175.00 | 1269.00 | 1213.67 | 1.15 | 0.00 |
| 2965.00 | 3151.00 | 3075.67 | -1.13 | 0.00 |
| 217.00 | 231.00 | 228.67 | -1.30 | 0.00 |
| 44.00 | 45.00 | 53.33 | -1.07 | 0.62 |
| 484.00 | 504.00 | 477.67 | 1.59 | 0.00 |
| 4979.00 | 5097.00 | 5063.67 | 1.02 | 0.66 |
| 986.00 | 1035.00 | 987.67 | -1.06 | 0.27 |
| 955.00 | 1157.00 | 1069.67 | 1.42 | 0.00 |
| 1953.00 | 2233.00 | 2115.33 | 1.19 | 0.00 |
| 654.00 | 810.00 | 748.33 | 1.26 | 0.00 |
| 731.00 | 818.00 | 821.00 | 1.11 | 0.04 |
| 1675.00 | 1923.00 | 1799.33 | 1.03 | 0.55 |
| 3114.00 | 3290.00 | 3391.33 | 1.14 | 0.00 |
| 379.00 | 360.00 | 387.67 | 1.13 | 0.03 |
| 1570.00 | 1490.00 | 1593.33 | -1.15 | 0.00 |
| 3472.00 | 3334.00 | 3427.67 | 1.58 | 0.00 |
| 11.00 | 9.00 | 10.33 | -1.47 | 0.18 |
| 644.00 | 724.00 | 693.00 | 1.01 | 0.79 |
| 22.00 | 28.00 | 27.33 | 1.49 | 0.03 |
| 909.00 | 890.00 | 910.00 | -1.16 | 0.00 |
| 5130.00 | 5416.00 | 5209.00 | 1.07 | 0.17 |
| 8.00 | 6.00 | 6.67 | 1.50 | 0.30 |
| 102.00 | 69.00 | 82.33 | 1.05 | 0.67 |
| 1299.00 | 1207.00 | 1270.33 | -1.27 | 0.00 |
| 597.00 | 550.00 | 604.33 | -1.26 | 0.00 |
| 23000.00 | 26269.00 | 25027.00 | 1.14 | 0.00 |
| 4229.00 | 4137.00 | 4263.33 | 1.05 | 0.25 |
| 386.00 | 381.00 | 378.67 | -1.17 | 0.01 |
| 566.00 | 485.00 | 483.67 | -1.12 | 0.29 |
| 1141.00 | 1108.00 | 1122.00 | -1.25 | 0.00 |
| 3192.00 | 3366.00 | 3348.33 | -1.03 | 0.43 |
| 1423.00 | 1496.00 | 1500.00 | -1.04 | 0.33 |
| 1503.00 | 1408.00 | 1427.67 | -1.21 | 0.00 |
| 684.00 | 659.00 | 670.00 | -1.21 | 0.00 |
| 2227.00 | 2189.00 | 2160.67 | -1.11 | 0.04 |
| 5.00 | 7.00 | 5.33 | 1.12 | 0.86 |
| 7478.00 | 6737.00 | 7143.33 | -1.26 | 0.00 |
| 639.00 | 628.00 | 584.00 | -1.23 | 0.01 |
| 2.00 | 1.00 | 2.33 | -1.15 | 1.00 |
| 2110.00 | 2071.00 | 2119.67 | -1.08 | 0.05 |
| 5177.00 | 5294.00 | 5325.00 | -1.21 | 0.00 |
| 3126.00 | 3240.00 | 3217.33 | 1.19 | 0.00 |
| 924.00 | 1238.00 | 1129.33 | 1.32 | 0.00 |
| 1131.00 | 1250.00 | 1233.67 | -1.10 | 0.02 |
| 1994.00 | 2201.00 | 2133.00 | 1.18 | 0.00 |
| 1053.00 | 1112.00 | 1065.33 | 1.01 | 0.85 |
| 81.00 | 90.00 | 78.00 | -1.13 | 0.29 |

| | | | | |
|---|---|---|---|---|
| 2528.00 | 2515.00 | 2584.33 | -1.17 | 0.00 |
| 15.00 | 6.00 | 7.67 | 1.14 | 0.79 |
| 424.00 | 431.00 | 419.33 | -1.11 | 0.09 |
| 986.00 | 991.00 | 1011.33 | -1.01 | 0.87 |
| 12.00 | 15.00 | 14.00 | 1.01 | 1.00 |
| 3288.00 | 3143.00 | 3189.00 | -1.50 | 0.00 |
| 894.00 | 841.00 | 866.33 | -1.26 | 0.00 |
| 6.00 | 3.00 | 3.00 | -2.20 | 0.08 |
| 608.00 | 706.00 | 682.67 | 1.19 | 0.00 |
| 1395.00 | 1559.00 | 1529.67 | 1.05 | 0.20 |
| 982.00 | 1025.00 | 1080.33 | 1.02 | 0.70 |
| 542.00 | 650.00 | 642.33 | 1.21 | 0.00 |
| 1577.00 | 1651.00 | 1715.33 | 1.17 | 0.00 |
| 404.00 | 417.00 | 442.33 | 1.09 | 0.19 |
| 1890.00 | 2039.00 | 2134.00 | 1.18 | 0.00 |
| 54.00 | 61.00 | 57.67 | -1.05 | 0.73 |
| 691.00 | 954.00 | 854.33 | 1.15 | 0.02 |
| 728.00 | 794.00 | 833.33 | 1.36 | 0.00 |
| 622.00 | 742.00 | 725.33 | 1.22 | 0.00 |
| 1675.00 | 1732.00 | 1767.00 | 1.17 | 0.00 |
| 1150.00 | 1323.00 | 1328.33 | 1.25 | 0.00 |
| 256.00 | 276.00 | 292.00 | -1.03 | 0.66 |
| 596.00 | 648.00 | 663.00 | 1.23 | 0.00 |
| 354.00 | 371.00 | 397.00 | 1.10 | 0.13 |
| 656.00 | 742.00 | 739.67 | 1.10 | 0.04 |
| 275.00 | 291.00 | 295.33 | 1.27 | 0.00 |
| 343.00 | 414.00 | 387.67 | 1.03 | 0.61 |
| 557.00 | 761.00 | 695.00 | 1.06 | 0.41 |
| 341.00 | 407.00 | 412.00 | 1.25 | 0.00 |
| 1047.00 | 1267.00 | 1218.00 | 1.09 | 0.07 |
| 897.00 | 898.00 | 960.33 | 1.16 | 0.00 |
| 1218.00 | 1205.00 | 1342.67 | 1.20 | 0.00 |
| 37.00 | 44.00 | 40.33 | 1.06 | 0.72 |
| 638.00 | 646.00 | 691.00 | 1.13 | 0.02 |
| 1195.00 | 1299.00 | 1380.33 | 1.09 | 0.11 |
| 1162.00 | 1278.00 | 1249.33 | 1.36 | 0.00 |
| 821.00 | 916.00 | 938.00 | 1.25 | 0.00 |
| 974.00 | 1075.00 | 1096.33 | 1.15 | 0.01 |
| 1923.00 | 2083.00 | 2124.33 | 1.20 | 0.00 |
| 3513.00 | 3974.00 | 3968.00 | 1.05 | 0.26 |
| 368.00 | 410.00 | 417.67 | 1.37 | 0.00 |
| 281.00 | 333.00 | 315.00 | 1.13 | 0.05 |
| 1699.00 | 1894.00 | 1828.00 | 1.16 | 0.00 |
| 3247.00 | 3541.00 | 3593.33 | 1.10 | 0.01 |
| 851.00 | 1001.00 | 990.00 | 1.17 | 0.00 |
| 738.00 | 856.00 | 847.00 | 1.18 | 0.00 |
| 1593.00 | 1709.00 | 1803.00 | 1.26 | 0.00 |
| 999.00 | 1098.00 | 1098.67 | 1.31 | 0.00 |

| | | | | |
|---|---|---|---|---|
| 867.00 | 890.00 | 954.67 | 1.15 | 0.02 |
| 870.00 | 915.00 | 942.00 | 1.07 | 0.15 |
| 2791.00 | 2914.00 | 3000.33 | 1.05 | 0.23 |
| 2041.00 | 2361.00 | 2375.33 | 1.28 | 0.00 |
| 12.00 | 23.00 | 14.67 | -1.10 | 0.77 |
| 537.00 | 614.00 | 603.33 | -1.04 | 0.39 |
| 15052.00 | 17571.00 | 16894.33 | -1.18 | 0.00 |
| 1.00 | 8.00 | 7.67 | -1.49 | 0.16 |
| 2.00 | 1.00 | 2.33 | -1.84 | 0.27 |
| 27.00 | 34.00 | 35.00 | 1.31 | 0.14 |
| 548.00 | 520.00 | 570.00 | -1.03 | 0.55 |
| 870.00 | 893.00 | 925.33 | 1.07 | 0.12 |
| 967.00 | 1040.00 | 1051.33 | 1.03 | 0.47 |
| 791.00 | 931.00 | 842.00 | 1.25 | 0.00 |
| 2740.00 | 3038.00 | 2854.33 | 1.11 | 0.02 |
| 1778.00 | 1948.00 | 1839.67 | -1.12 | 0.01 |
| 911.00 | 1043.00 | 1079.33 | 1.11 | 0.06 |
| 4.00 | 0.00 | 2.33 | -1.01 | 1.00 |
| 268.00 | 253.00 | 282.67 | 1.07 | 0.36 |
| 197.00 | 197.00 | 194.67 | -1.18 | 0.03 |
| 38.00 | 49.00 | 52.00 | -1.08 | 0.57 |
| 422.00 | 476.00 | 483.33 | 1.15 | 0.02 |
| 111.00 | 144.00 | 134.67 | 1.35 | 0.00 |
| 344.00 | 460.00 | 386.33 | -1.06 | 0.45 |
| 128.00 | 141.00 | 138.33 | -1.31 | 0.00 |
| 87.00 | 124.00 | 104.67 | 1.04 | 0.77 |
| 544.00 | 608.00 | 581.67 | 1.21 | 0.00 |
| 417.00 | 552.00 | 507.67 | 1.31 | 0.00 |
| 536.00 | 517.00 | 530.33 | -1.01 | 0.88 |
| 13.00 | 19.00 | 17.33 | 1.09 | 0.77 |
| 1309.00 | 1237.00 | 1307.00 | -1.33 | 0.00 |
| 1077.00 | 1416.00 | 1225.00 | 1.21 | 0.00 |
| 92.00 | 68.00 | 81.67 | -1.56 | 0.00 |
| 3704.00 | 3847.00 | 3810.33 | -1.19 | 0.00 |
| 1618.00 | 1688.00 | 1663.33 | 1.10 | 0.02 |
| 1603.00 | 1657.00 | 1692.00 | 1.04 | 0.35 |
| 2395.00 | 2608.00 | 2599.67 | 1.01 | 0.83 |
| 116.00 | 141.00 | 123.33 | -1.00 | 1.00 |
| 1616.00 | 1882.00 | 1777.67 | 1.01 | 0.80 |
| 2304.00 | 2449.00 | 2392.67 | 1.10 | 0.01 |
| 534.00 | 576.00 | 560.67 | 1.58 | 0.00 |
| 732.00 | 770.00 | 790.33 | 1.27 | 0.00 |
| 514.00 | 522.00 | 510.33 | -1.31 | 0.00 |
| 91.00 | 109.00 | 103.33 | 1.04 | 0.68 |
| 2250.00 | 2424.00 | 2394.33 | 1.29 | 0.00 |
| 1994.00 | 1972.00 | 1980.00 | -1.12 | 0.01 |
| 352.00 | 278.00 | 308.67 | -1.24 | 0.00 |
| 194.00 | 209.00 | 200.67 | -1.17 | 0.02 |

| | | | | |
|---|---|---|---|---|
| 1990.00 | 2141.00 | 2154.00 | -1.12 | 0.00 |
| 5.00 | 5.00 | 4.33 | -2.30 | 0.02 |
| 1012.00 | 1096.00 | 1045.33 | 1.02 | 0.68 |
| 114.00 | 84.00 | 98.33 | 1.35 | 0.01 |
| 2087.00 | 2217.00 | 2126.00 | -1.10 | 0.06 |
| 3465.00 | 3405.00 | 3596.00 | 1.18 | 0.00 |
| 2412.00 | 2714.00 | 2500.67 | -1.18 | 0.00 |
| 30.00 | 28.00 | 26.00 | 1.03 | 0.88 |
| 2223.00 | 2344.00 | 2323.67 | -1.03 | 0.43 |
| 455.00 | 471.00 | 472.67 | -1.34 | 0.00 |
| 845.00 | 757.00 | 840.00 | -1.08 | 0.13 |
| 10241.00 | 9935.00 | 10127.33 | -1.16 | 0.00 |
| 2471.00 | 2728.00 | 2654.67 | 1.03 | 0.42 |
| 2884.00 | 3373.00 | 3202.33 | 1.71 | 0.00 |
| 262.00 | 256.00 | 282.00 | 1.18 | 0.02 |
| 1717.00 | 1799.00 | 1803.00 | -1.03 | 0.50 |
| 1449.00 | 1351.00 | 1370.33 | -1.13 | 0.02 |
| 377.00 | 409.00 | 421.00 | -1.16 | 0.01 |
| 502.00 | 455.00 | 481.00 | -1.03 | 0.57 |
| 1225.00 | 1394.00 | 1287.00 | 1.17 | 0.00 |
| 2521.00 | 2611.00 | 2663.00 | 1.33 | 0.00 |
| 736.00 | 784.00 | 765.33 | 1.38 | 0.00 |
| 509.00 | 629.00 | 608.67 | 1.23 | 0.00 |
| 84.00 | 87.00 | 94.00 | -1.31 | 0.00 |
| 224.00 | 242.00 | 249.00 | -1.02 | 0.79 |
| 33.00 | 31.00 | 29.67 | -1.06 | 0.78 |
| 1125.00 | 1331.00 | 1253.67 | -1.16 | 0.00 |
| 653.00 | 641.00 | 651.33 | -1.18 | 0.00 |
| 820.00 | 891.00 | 884.67 | 1.01 | 0.82 |
| 406.00 | 455.00 | 463.67 | 1.21 | 0.00 |
| 1806.00 | 1801.00 | 1806.67 | -1.01 | 0.82 |
| 614.00 | 746.00 | 677.67 | -1.02 | 0.70 |
| 1940.00 | 2019.00 | 1997.67 | -1.15 | 0.00 |
| 441.00 | 539.00 | 498.33 | 1.22 | 0.00 |
| 879.00 | 1007.00 | 1021.67 | 1.02 | 0.77 |
| 248.00 | 305.00 | 284.33 | -1.13 | 0.05 |
| 681.00 | 678.00 | 730.00 | 1.10 | 0.06 |
| 77.00 | 89.00 | 84.67 | -2.29 | 0.00 |
| 841.00 | 839.00 | 832.00 | 1.40 | 0.00 |
| 3907.00 | 4379.00 | 4372.33 | 1.21 | 0.00 |
| 824.00 | 1013.00 | 936.00 | -1.00 | 0.96 |
| 1723.00 | 1867.00 | 1898.33 | 1.13 | 0.00 |
| 248.00 | 275.00 | 284.33 | -1.09 | 0.18 |
| 5.00 | 9.00 | 6.33 | 1.17 | 0.74 |
| 3.00 | 1.00 | 3.00 | -1.87 | 0.21 |
| 14.00 | 17.00 | 17.67 | 1.53 | 0.07 |
| 386.00 | 439.00 | 397.33 | -1.01 | 0.90 |
| 661.00 | 802.00 | 743.00 | -1.03 | 0.55 |

| | | | | |
|---|---|---|---|---|
| 15.00 | 9.00 | 12.67 | 1.07 | 0.82 |
| 1687.00 | 1856.00 | 1876.33 | 1.13 | 0.00 |
| 81.00 | 113.00 | 90.67 | -2.20 | 0.00 |
| 470.00 | 469.00 | 477.33 | 1.08 | 0.13 |
| 15.00 | 10.00 | 9.67 | -1.01 | 1.00 |
| 612.00 | 633.00 | 635.67 | -1.30 | 0.00 |
| 207.00 | 226.00 | 222.00 | 1.10 | 0.15 |
| 4.00 | 6.00 | 5.67 | -1.36 | 0.44 |
| 981.00 | 1048.00 | 1042.00 | 1.06 | 0.17 |
| 1606.00 | 1771.00 | 1790.33 | 1.13 | 0.01 |
| 85.00 | 73.00 | 86.00 | 1.20 | 0.10 |
| 11.00 | 18.00 | 15.67 | -1.65 | 0.01 |
| 1573.00 | 1795.00 | 1654.33 | 1.04 | 0.38 |
| 377.00 | 327.00 | 373.00 | -1.02 | 0.79 |
| 869.00 | 950.00 | 915.00 | -1.11 | 0.04 |
| 1372.00 | 1501.00 | 1503.33 | 1.20 | 0.00 |
| 1180.00 | 1221.00 | 1218.67 | 1.11 | 0.01 |
| 2032.00 | 2289.00 | 2204.67 | 1.15 | 0.00 |
| 222.00 | 205.00 | 197.00 | -1.26 | 0.01 |
| 138.00 | 131.00 | 129.67 | -1.03 | 0.74 |
| 574.00 | 550.00 | 558.00 | -1.20 | 0.00 |
| 113.00 | 98.00 | 96.67 | -1.18 | 0.15 |
| 2036.00 | 2226.00 | 2095.33 | -1.08 | 0.09 |
| 363.00 | 438.00 | 399.67 | -1.15 | 0.01 |
| 1457.00 | 1491.00 | 1565.67 | 1.27 | 0.00 |
| 213.00 | 250.00 | 229.67 | 1.01 | 0.90 |
| 55.00 | 54.00 | 54.00 | -1.08 | 0.59 |
| 2674.00 | 2661.00 | 2743.67 | -1.29 | 0.00 |
| 678.00 | 795.00 | 773.00 | 1.01 | 0.83 |
| 1212.00 | 1233.00 | 1293.67 | 1.16 | 0.00 |
| 369.00 | 384.00 | 391.00 | 1.07 | 0.21 |
| 3.00 | 5.00 | 5.00 | -2.60 | 0.00 |
| 423.00 | 462.00 | 461.67 | 1.11 | 0.04 |
| 633.00 | 576.00 | 625.67 | -1.15 | 0.01 |
| 475.00 | 513.00 | 498.33 | 1.09 | 0.09 |
| 3580.00 | 4236.00 | 3991.33 | -1.18 | 0.00 |
| 18.00 | 22.00 | 19.33 | -1.29 | 0.22 |
| 107.00 | 81.00 | 98.67 | 1.24 | 0.06 |
| 187.00 | 131.00 | 169.67 | 1.13 | 0.22 |
| 20.00 | 18.00 | 19.33 | 1.06 | 0.79 |
| 96.00 | 105.00 | 102.00 | -1.24 | 0.02 |
| 30.00 | 29.00 | 29.33 | 1.29 | 0.14 |
| 104.00 | 128.00 | 111.67 | -1.12 | 0.25 |
| 977.00 | 995.00 | 988.67 | 1.16 | 0.00 |
| 79.00 | 76.00 | 78.33 | 1.14 | 0.23 |
| 2738.00 | 2647.00 | 2792.00 | -1.17 | 0.00 |
| 12.00 | 8.00 | 10.00 | -1.01 | 1.00 |
| 557.00 | 631.00 | 591.00 | -1.34 | 0.00 |

| | | | | |
|---|---|---|---|---|
| 1425.00 | 1368.00 | 1407.00 | 1.13 | 0.00 |
| 311.00 | 355.00 | 336.67 | -1.31 | 0.00 |
| 2801.00 | 2736.00 | 2840.33 | -1.02 | 0.52 |
| 3.00 | 3.00 | 2.67 | -1.25 | 0.82 |
| 4.00 | 4.00 | 3.33 | -1.31 | 0.68 |
| 904.00 | 919.00 | 953.33 | 1.38 | 0.00 |
| 486.00 | 432.00 | 456.67 | 1.11 | 0.10 |
| 225.00 | 251.00 | 244.33 | 1.26 | 0.00 |
| 3.00 | 2.00 | 2.33 | -1.01 | 1.00 |
| 2266.00 | 2514.00 | 2427.67 | 1.16 | 0.00 |
| 1843.00 | 2036.00 | 2049.67 | 1.31 | 0.00 |
| 571.00 | 548.00 | 598.00 | 1.10 | 0.06 |
| 576.00 | 566.00 | 617.00 | 1.02 | 0.72 |
| 1705.00 | 1787.00 | 1851.33 | 1.16 | 0.00 |
| 414.00 | 343.00 | 406.00 | -1.06 | 0.39 |
| 491.00 | 526.00 | 547.33 | -1.03 | 0.56 |
| 263.00 | 301.00 | 279.00 | 1.19 | 0.01 |
| 334.00 | 389.00 | 377.33 | 1.07 | 0.24 |
| 2352.00 | 2479.00 | 2471.67 | 1.44 | 0.00 |
| 457.00 | 525.00 | 510.33 | 1.06 | 0.25 |
| 2874.00 | 2770.00 | 2826.67 | 1.04 | 0.38 |
| 2821.00 | 2909.00 | 2851.67 | -1.36 | 0.00 |
| 24.00 | 18.00 | 22.00 | 3.06 | 0.00 |
| 107.00 | 147.00 | 134.67 | 1.10 | 0.31 |
| 1425.00 | 1536.00 | 1455.00 | -1.10 | 0.05 |
| 428.00 | 422.00 | 414.67 | -1.14 | 0.07 |
| 578.00 | 524.00 | 564.00 | 1.04 | 0.52 |
| 1997.00 | 1997.00 | 2053.33 | 1.10 | 0.01 |
| 3918.00 | 4248.00 | 4149.00 | 1.06 | 0.10 |
| 2380.00 | 2727.00 | 2630.67 | 1.07 | 0.07 |
| 663.00 | 602.00 | 657.33 | 1.08 | 0.14 |
| 120.00 | 142.00 | 124.00 | 1.10 | 0.41 |
| 226.00 | 319.00 | 286.00 | 1.10 | 0.23 |
| 19.00 | 26.00 | 21.00 | 1.02 | 0.93 |
| 1105.00 | 1219.00 | 1209.33 | 1.18 | 0.00 |
| 1665.00 | 1811.00 | 1772.33 | 1.11 | 0.00 |
| 703.00 | 725.00 | 747.33 | -1.02 | 0.68 |
| 597.00 | 676.00 | 692.33 | -1.27 | 0.00 |
| 14.00 | 8.00 | 8.67 | -1.50 | 0.19 |
| 1735.00 | 1685.00 | 1782.33 | -1.12 | 0.00 |
| 3378.00 | 3508.00 | 3559.00 | -1.00 | 0.92 |
| 1423.00 | 1730.00 | 1660.00 | 1.23 | 0.00 |
| 1478.00 | 1645.00 | 1594.67 | -1.02 | 0.67 |
| 763.00 | 927.00 | 881.67 | 1.14 | 0.05 |
| 6035.00 | 6541.00 | 6439.67 | 1.08 | 0.02 |
| 1315.00 | 1352.00 | 1310.00 | -1.05 | 0.26 |
| 164.00 | 211.00 | 187.00 | -1.09 | 0.30 |
| 599.00 | 536.00 | 554.67 | -1.24 | 0.00 |

| | | | | |
|---|---|---|---|---|
| 11096.00 | 11478.00 | 11497.00 | 1.04 | 0.26 |
| 1461.00 | 1601.00 | 1706.67 | 1.32 | 0.00 |
| 5712.00 | 6089.00 | 5919.33 | 1.03 | 0.42 |
| 292.00 | 321.00 | 317.00 | 1.09 | 0.13 |
| 1271.00 | 1354.00 | 1417.67 | 1.09 | 0.05 |
| 3080.00 | 3400.00 | 3343.67 | -1.02 | 0.52 |
| 7490.00 | 8309.00 | 8271.67 | 1.07 | 0.05 |
| 3919.00 | 4661.00 | 4482.67 | 1.15 | 0.00 |
| 1923.00 | 1620.00 | 1669.33 | -1.56 | 0.00 |
| 21.00 | 17.00 | 16.33 | -1.01 | 1.00 |
| 179.00 | 163.00 | 169.33 | -1.08 | 0.35 |
| 1243.00 | 1359.00 | 1389.00 | 1.16 | 0.00 |
| 1568.00 | 1457.00 | 1490.00 | -1.31 | 0.00 |
| 11614.00 | 10882.00 | 10865.00 | -1.27 | 0.00 |
| 145.00 | 109.00 | 127.67 | -1.09 | 0.43 |
| 3.00 | 6.00 | 5.33 | 1.42 | 0.45 |
| 195.00 | 182.00 | 190.67 | -1.62 | 0.00 |
| 4.00 | 5.00 | 4.00 | 1.07 | 1.00 |
| 0.00 | 1.00 | 0.33 | -2.48 | 0.64 |
| 155.00 | 209.00 | 178.67 | 1.26 | 0.01 |
| 17.00 | 8.00 | 14.33 | -1.53 | 0.06 |
| 83.00 | 61.00 | 79.67 | -1.06 | 0.65 |
| 2.00 | 1.00 | 2.33 | -1.43 | 0.63 |
| 15.00 | 20.00 | 20.00 | -1.01 | 1.00 |
| 11.00 | 12.00 | 11.00 | -1.44 | 0.16 |
| 7288.00 | 7730.00 | 7777.67 | 1.13 | 0.00 |
| 78.00 | 78.00 | 83.67 | -1.09 | 0.42 |
| 361.00 | 430.00 | 379.67 | -1.01 | 0.86 |
| 4098.00 | 4100.00 | 4128.67 | -1.21 | 0.00 |
| 3.00 | 2.00 | 3.00 | -1.23 | 0.83 |
| 506.00 | 564.00 | 557.33 | -1.29 | 0.00 |
| 3697.00 | 3812.00 | 3855.67 | -1.03 | 0.35 |
| 33.00 | 46.00 | 40.33 | 1.13 | 0.46 |
| 2010.00 | 2280.00 | 2171.00 | 1.04 | 0.29 |
| 56.00 | 72.00 | 71.33 | 1.03 | 0.88 |
| 28.00 | 32.00 | 36.00 | 2.16 | 0.00 |
| 171.00 | 182.00 | 170.00 | -1.01 | 0.92 |
| 289.00 | 368.00 | 342.67 | -1.17 | 0.01 |
| 200.00 | 306.00 | 251.33 | 1.16 | 0.15 |
| 556.00 | 514.00 | 539.67 | -1.26 | 0.00 |
| 16.00 | 34.00 | 30.33 | 1.19 | 0.42 |
| 66.00 | 68.00 | 69.67 | -1.38 | 0.00 |
| 51.00 | 57.00 | 58.00 | -1.18 | 0.16 |
| 65.00 | 87.00 | 80.00 | -1.31 | 0.01 |
| 115.00 | 136.00 | 133.67 | -1.31 | 0.00 |
| 379.00 | 365.00 | 406.33 | -1.06 | 0.39 |
| 3.00 | 4.00 | 3.00 | -1.88 | 0.18 |
| 7.00 | 4.00 | 10.00 | -1.75 | 0.03 |

| | | | | |
|---|---|---|---|---|
| 19.00 | 13.00 | 12.67 | -1.35 | 0.23 |
| 184.00 | 209.00 | 196.67 | 1.18 | 0.02 |
| 3573.00 | 3452.00 | 3562.67 | -1.13 | 0.00 |
| 25535.00 | 30050.00 | 29188.67 | 1.25 | 0.00 |
| 1508.00 | 1523.00 | 1536.67 | 1.34 | 0.00 |
| 84.00 | 80.00 | 91.00 | 1.13 | 0.25 |
| 482.00 | 504.00 | 519.33 | -1.24 | 0.00 |
| 274.00 | 291.00 | 283.33 | -1.06 | 0.33 |
| 565.00 | 603.00 | 618.67 | -1.02 | 0.62 |
| 1417.00 | 1539.00 | 1543.00 | 1.16 | 0.00 |
| 1711.00 | 1828.00 | 1800.00 | -1.01 | 0.89 |
| 135.00 | 107.00 | 111.00 | -1.38 | 0.00 |
| 10.00 | 15.00 | 10.00 | -1.88 | 0.01 |
| 78.00 | 77.00 | 84.67 | -1.01 | 0.90 |
| 72.00 | 111.00 | 98.67 | -1.04 | 0.74 |
| 14.00 | 10.00 | 11.33 | 2.21 | 0.01 |
| 2160.00 | 2256.00 | 2259.67 | -1.15 | 0.00 |
| 414.00 | 356.00 | 412.33 | 1.05 | 0.52 |
| 274.00 | 311.00 | 300.00 | 1.17 | 0.02 |
| 1438.00 | 1573.00 | 1520.33 | 1.14 | 0.01 |
| 995.00 | 1044.00 | 1054.33 | 1.08 | 0.06 |
| 186.00 | 158.00 | 187.33 | 1.05 | 0.61 |
| 0.00 | 0.00 | 0.00 | -9.06 | 0.27 |
| 549.00 | 736.00 | 669.67 | -1.32 | 0.00 |
| 1677.00 | 1682.00 | 1655.00 | -1.07 | 0.12 |
| 563.00 | 611.00 | 615.33 | 1.15 | 0.01 |
| 13.00 | 8.00 | 9.00 | -1.86 | 0.02 |
| 963.00 | 1033.00 | 996.33 | 1.32 | 0.00 |
| 507.00 | 531.00 | 533.00 | 1.10 | 0.11 |
| 3338.00 | 3461.00 | 3407.67 | 1.03 | 0.50 |
| 124.00 | 144.00 | 130.67 | -1.66 | 0.00 |
| 862.00 | 893.00 | 885.33 | -1.01 | 0.87 |
| 1115.00 | 1226.00 | 1213.00 | -1.10 | 0.03 |
| 228.00 | 239.00 | 237.00 | -1.16 | 0.03 |
| 202.00 | 191.00 | 195.00 | -1.17 | 0.03 |
| 21.00 | 21.00 | 20.00 | -1.01 | 1.00 |
| 4.00 | 17.00 | 10.00 | 1.02 | 1.00 |
| 4.00 | 12.00 | 9.67 | 1.35 | 0.34 |
| 162.00 | 161.00 | 164.67 | -1.23 | 0.00 |
| 45.00 | 42.00 | 46.67 | 1.14 | 0.35 |
| 2091.00 | 2384.00 | 2270.33 | 1.27 | 0.00 |
| 151.00 | 188.00 | 169.33 | -1.12 | 0.14 |
| 484.00 | 460.00 | 493.67 | -1.06 | 0.24 |
| 2177.00 | 2393.00 | 2295.33 | 1.04 | 0.27 |
| 1049.00 | 1053.00 | 1089.67 | -1.15 | 0.00 |
| 5716.00 | 6243.00 | 6123.67 | 1.03 | 0.32 |
| 363.00 | 374.00 | 363.67 | -1.01 | 0.85 |
| 439.00 | 472.00 | 473.67 | 1.20 | 0.00 |

| | | | | |
|---|---|---|---|---|
| 449.00 | 492.00 | 472.33 | -1.04 | 0.48 |
| 1890.00 | 1743.00 | 1765.00 | -1.05 | 0.37 |
| 64.00 | 69.00 | 61.33 | -1.41 | 0.00 |
| 138.00 | 176.00 | 166.33 | 1.06 | 0.50 |
| 1622.00 | 1750.00 | 1659.67 | -1.27 | 0.00 |
| 1516.00 | 1611.00 | 1571.33 | 1.62 | 0.00 |
| 795.00 | 854.00 | 815.33 | -1.24 | 0.00 |
| 4.00 | 6.00 | 7.33 | -1.01 | 1.00 |
| 3.00 | 4.00 | 2.67 | -1.01 | 1.00 |
| 4.00 | 2.00 | 3.67 | -2.26 | 0.03 |
| 7677.00 | 8069.00 | 8029.00 | -1.70 | 0.00 |
| 2903.00 | 2813.00 | 2833.67 | 1.11 | 0.02 |
| 522.00 | 530.00 | 522.33 | -1.01 | 0.85 |
| 1084.00 | 1018.00 | 1106.00 | -1.11 | 0.03 |
| 124.00 | 121.00 | 116.67 | -1.18 | 0.07 |
| 1.00 | 4.00 | 3.00 | -1.01 | 1.00 |
| 102.00 | 94.00 | 94.00 | -1.48 | 0.00 |
| 1345.00 | 1569.00 | 1563.00 | 1.21 | 0.00 |
| 1938.00 | 1815.00 | 1826.33 | -1.20 | 0.00 |
| 550.00 | 570.00 | 592.33 | 1.13 | 0.01 |
| 34.00 | 35.00 | 36.33 | 1.19 | 0.28 |
| 1326.00 | 1544.00 | 1483.33 | 1.03 | 0.43 |
| 3.00 | 2.00 | 3.00 | -1.99 | 0.13 |
| 478.00 | 540.00 | 535.00 | 1.13 | 0.02 |
| 2318.00 | 2447.00 | 2496.67 | 1.00 | 0.99 |
| 792.00 | 873.00 | 876.33 | 1.09 | 0.06 |
| 466.00 | 505.00 | 504.33 | 1.03 | 0.55 |
| 996.00 | 1090.00 | 1062.00 | 1.20 | 0.00 |
| 1999.00 | 1958.00 | 2053.33 | -1.22 | 0.00 |
| 1092.00 | 1283.00 | 1256.67 | 1.13 | 0.01 |
| 3413.00 | 3653.00 | 3572.00 | 1.03 | 0.38 |
| 139.00 | 174.00 | 158.00 | 1.35 | 0.00 |
| 862.00 | 893.00 | 904.00 | -1.02 | 0.70 |
| 460.00 | 487.00 | 508.67 | 1.12 | 0.06 |
| 679.00 | 779.00 | 788.67 | -1.12 | 0.03 |
| 592.00 | 642.00 | 667.00 | -1.10 | 0.08 |
| 536.00 | 534.00 | 570.33 | -1.14 | 0.01 |
| 6.00 | 6.00 | 5.00 | 1.06 | 1.00 |
| 766.00 | 916.00 | 891.00 | 1.16 | 0.00 |
| 2131.00 | 2292.00 | 2244.33 | -1.11 | 0.00 |
| 917.00 | 1035.00 | 1006.33 | 1.20 | 0.00 |
| 853.00 | 988.00 | 980.00 | 1.12 | 0.02 |
| 591.00 | 644.00 | 624.67 | 1.09 | 0.07 |
| 2253.00 | 2389.00 | 2409.33 | -1.13 | 0.00 |
| 470.00 | 474.00 | 488.67 | 1.18 | 0.00 |
| 25374.00 | 25568.00 | 25336.33 | -1.10 | 0.01 |
| 105.00 | 115.00 | 118.33 | 1.14 | 0.16 |
| 24.00 | 23.00 | 19.33 | 1.07 | 0.83 |

| | | | | |
|---|---|---|---|---|
| 480.00 | 481.00 | 514.33 | 1.07 | 0.29 |
| 3344.00 | 3499.00 | 3540.00 | -1.08 | 0.01 |
| 1049.00 | 1079.00 | 1119.00 | 1.04 | 0.44 |
| 840.00 | 922.00 | 901.33 | -2.31 | 0.00 |
| 2347.00 | 2377.00 | 2371.00 | -1.30 | 0.00 |
| 21.00 | 20.00 | 16.33 | -2.10 | 0.00 |
| 262.00 | 263.00 | 284.67 | 1.12 | 0.08 |
| 4.00 | 5.00 | 6.00 | -1.18 | 0.75 |
| 190.00 | 194.00 | 194.67 | -1.02 | 0.80 |
| 226.00 | 221.00 | 223.33 | -1.17 | 0.02 |
| 1181.00 | 1314.00 | 1237.00 | -1.12 | 0.01 |
| 349.00 | 335.00 | 345.00 | 1.06 | 0.35 |
| 800.00 | 770.00 | 828.33 | 1.05 | 0.34 |
| 425.00 | 469.00 | 465.33 | 1.08 | 0.14 |
| 3135.00 | 3102.00 | 3202.67 | -1.17 | 0.00 |
| 1580.00 | 1783.00 | 1743.33 | 1.27 | 0.00 |
| 4340.00 | 4909.00 | 4750.00 | 1.19 | 0.00 |
| 5625.00 | 5954.00 | 5991.00 | 1.21 | 0.00 |
| 1431.00 | 1555.00 | 1552.00 | 1.20 | 0.00 |
| 559.00 | 634.00 | 633.33 | 1.09 | 0.08 |
| 2792.00 | 2770.00 | 2853.33 | 1.05 | 0.20 |
| 344.00 | 393.00 | 392.33 | 1.31 | 0.00 |
| 4.00 | 6.00 | 4.67 | -1.15 | 0.86 |
| 323.00 | 362.00 | 353.33 | 1.20 | 0.00 |
| 431.00 | 526.00 | 473.67 | 1.18 | 0.01 |
| 295.00 | 355.00 | 338.67 | 1.22 | 0.00 |
| 35.00 | 28.00 | 29.67 | 1.19 | 0.34 |
| 503.00 | 532.00 | 535.00 | 1.31 | 0.00 |
| 1763.00 | 1765.00 | 1783.67 | -1.13 | 0.00 |
| 322.00 | 359.00 | 346.33 | 1.05 | 0.43 |
| 122.00 | 116.00 | 126.67 | -1.07 | 0.43 |
| 46.00 | 48.00 | 54.00 | 1.30 | 0.06 |
| 488.00 | 505.00 | 535.33 | 1.14 | 0.03 |
| 594.00 | 631.00 | 677.33 | 1.13 | 0.05 |
| 2965.00 | 3333.00 | 3195.67 | 1.17 | 0.00 |
| 1435.00 | 1372.00 | 1481.00 | -1.00 | 0.92 |
| 118.00 | 118.00 | 119.00 | 1.36 | 0.00 |
| 571.00 | 658.00 | 613.67 | 1.09 | 0.10 |
| 3039.00 | 3316.00 | 3327.00 | 1.09 | 0.01 |
| 79.00 | 91.00 | 91.00 | -1.16 | 0.11 |
| 29.00 | 28.00 | 26.00 | 1.24 | 0.23 |
| 501.00 | 512.00 | 514.33 | 1.02 | 0.68 |
| 14.00 | 26.00 | 20.33 | -1.13 | 0.56 |
| 900.00 | 1172.00 | 1086.33 | 1.46 | 0.00 |
| 285.00 | 320.00 | 327.00 | 1.15 | 0.04 |
| 3540.00 | 3792.00 | 3667.67 | -1.16 | 0.00 |
| 9901.00 | 9911.00 | 9959.00 | -1.28 | 0.00 |
| 1771.00 | 1892.00 | 1891.67 | -1.13 | 0.00 |

| | | | | |
|---|---|---|---|---|
| 602.00 | 600.00 | 603.00 | 1.13 | 0.01 |
| 2490.00 | 2781.00 | 2683.00 | 1.07 | 0.07 |
| 1414.00 | 1471.00 | 1504.33 | 1.07 | 0.09 |
| 3001.00 | 3099.00 | 3030.67 | 1.04 | 0.38 |
| 1718.00 | 1631.00 | 1606.00 | 1.25 | 0.00 |
| 1312.00 | 1439.00 | 1390.00 | 1.05 | 0.29 |
| 3104.00 | 3190.00 | 3129.33 | -1.01 | 0.73 |
| 4500.00 | 4636.00 | 4557.00 | 1.11 | 0.00 |
| 1887.00 | 1718.00 | 1775.33 | -1.22 | 0.00 |
| 539.00 | 610.00 | 612.67 | 1.13 | 0.03 |
| 1011.00 | 1113.00 | 1088.00 | 1.14 | 0.00 |
| 1074.00 | 1238.00 | 1182.33 | 1.22 | 0.00 |
| 1576.00 | 1633.00 | 1622.00 | 1.13 | 0.00 |
| 215.00 | 217.00 | 251.67 | 1.06 | 0.56 |
| 758.00 | 930.00 | 840.33 | 1.19 | 0.00 |
| 1640.00 | 1706.00 | 1730.33 | 1.05 | 0.25 |
| 8.00 | 12.00 | 13.67 | 1.19 | 0.58 |
| 1066.00 | 1178.00 | 1164.33 | 1.05 | 0.33 |
| 1737.00 | 2010.00 | 1933.00 | 1.13 | 0.00 |
| 2083.00 | 2234.00 | 2237.67 | 1.13 | 0.00 |
| 3454.00 | 3637.00 | 3572.00 | -1.08 | 0.03 |
| 1366.00 | 1508.00 | 1445.67 | 1.12 | 0.01 |
| 836.00 | 960.00 | 925.33 | -1.02 | 0.63 |
| 233.00 | 253.00 | 267.67 | -1.75 | 0.00 |
| 1193.00 | 1335.00 | 1272.00 | 1.18 | 0.00 |
| 1184.00 | 1371.00 | 1357.67 | 1.31 | 0.00 |
| 1531.00 | 1663.00 | 1719.00 | 1.17 | 0.00 |
| 23.00 | 7.00 | 16.33 | 1.10 | 0.72 |
| 313.00 | 292.00 | 329.67 | -1.01 | 0.87 |
| 5.00 | 10.00 | 9.33 | 1.31 | 0.41 |
| 378.00 | 390.00 | 407.33 | -1.02 | 0.82 |
| 18.00 | 21.00 | 21.33 | -1.09 | 0.62 |
| 854.00 | 890.00 | 887.67 | 1.15 | 0.00 |
| 1699.00 | 1865.00 | 1865.33 | -1.13 | 0.00 |
| 15.00 | 10.00 | 13.67 | -1.48 | 0.08 |
| 875.00 | 877.00 | 894.67 | 1.09 | 0.06 |
| 13.00 | 14.00 | 13.33 | -1.06 | 0.91 |
| 764.00 | 751.00 | 731.00 | -1.22 | 0.00 |
| 38.00 | 58.00 | 46.00 | 1.64 | 0.00 |
| 55.00 | 58.00 | 62.67 | 1.01 | 1.00 |
| 573.00 | 550.00 | 583.67 | 1.12 | 0.02 |
| 128.00 | 187.00 | 148.33 | 1.26 | 0.04 |
| 27.00 | 22.00 | 27.00 | -1.53 | 0.02 |
| 1789.00 | 1684.00 | 1771.67 | -1.22 | 0.00 |
| 585.00 | 707.00 | 675.33 | 1.13 | 0.02 |
| 3.00 | 3.00 | 4.33 | 1.58 | 0.39 |
| 4.00 | 3.00 | 5.00 | 1.62 | 0.32 |
| 82.00 | 80.00 | 85.33 | 1.25 | 0.04 |

| | | | | |
|---|---|---|---|---|
| 1737.00 | 1925.00 | 1911.67 | -1.08 | 0.06 |
| 8855.00 | 9182.00 | 9500.67 | 1.11 | 0.00 |
| 996.00 | 1054.00 | 1089.67 | 1.23 | 0.00 |
| 23.00 | 18.00 | 24.33 | 1.47 | 0.06 |
| 623.00 | 603.00 | 604.67 | 1.09 | 0.14 |
| 160.00 | 125.00 | 139.00 | -1.62 | 0.00 |
| 1780.00 | 1816.00 | 1873.33 | -1.02 | 0.62 |
| 3.00 | 6.00 | 3.67 | 1.52 | 0.49 |
| 71.00 | 95.00 | 85.67 | 1.13 | 0.29 |
| 4100.00 | 4253.00 | 4311.33 | 1.10 | 0.01 |
| 1996.00 | 2214.00 | 2282.33 | 1.21 | 0.00 |
| 21.00 | 18.00 | 20.33 | 1.06 | 0.79 |
| 12.00 | 10.00 | 15.67 | 1.08 | 0.87 |
| 704.00 | 686.00 | 704.67 | -1.03 | 0.53 |
| 8171.00 | 8748.00 | 8853.00 | 1.02 | 0.60 |
| 3147.00 | 3494.00 | 3390.00 | -1.01 | 0.81 |
| 1504.00 | 1808.00 | 1715.00 | -1.01 | 0.85 |
| 6.00 | 6.00 | 5.00 | -1.21 | 0.73 |
| 1452.00 | 1683.00 | 1630.00 | -1.17 | 0.00 |
| 2501.00 | 2447.00 | 2492.00 | 1.02 | 0.62 |
| 58.00 | 87.00 | 75.67 | 1.48 | 0.00 |
| 1601.00 | 1790.00 | 1724.33 | 1.05 | 0.20 |
| 664.00 | 683.00 | 708.67 | -1.16 | 0.00 |
| 1089.00 | 1056.00 | 1095.67 | -1.07 | 0.14 |
| 1055.00 | 1117.00 | 1121.33 | -1.02 | 0.67 |
| 435.00 | 491.00 | 464.00 | -1.10 | 0.10 |
| 3.00 | 8.00 | 7.00 | 1.15 | 0.76 |
| 295.00 | 265.00 | 305.00 | -1.16 | 0.03 |
| 26573.00 | 30272.00 | 29265.33 | -1.06 | 0.07 |
| 136.00 | 161.00 | 154.33 | 1.15 | 0.09 |
| 252.00 | 253.00 | 259.67 | -1.08 | 0.27 |
| 2730.00 | 3091.00 | 3044.33 | 1.16 | 0.00 |
| 15.00 | 28.00 | 22.67 | -1.24 | 0.25 |
| 35.00 | 36.00 | 35.33 | 2.88 | 0.00 |
| 799.00 | 927.00 | 881.33 | -1.05 | 0.31 |
| 18.00 | 26.00 | 19.33 | 1.96 | 0.00 |
| 227.00 | 225.00 | 240.00 | -1.01 | 0.88 |
| 731.00 | 865.00 | 839.67 | -1.02 | 0.75 |
| 127.00 | 138.00 | 136.00 | 1.11 | 0.19 |
| 2067.00 | 2245.00 | 2135.67 | 1.03 | 0.43 |
| 485.00 | 443.00 | 466.00 | -1.00 | 0.99 |
| 27.00 | 20.00 | 24.33 | 4.71 | 0.00 |
| 300.00 | 281.00 | 281.67 | -1.11 | 0.14 |
| 13.00 | 9.00 | 10.33 | 2.72 | 0.00 |
| 511.00 | 564.00 | 557.33 | -1.37 | 0.00 |
| 1.00 | 5.00 | 3.00 | -2.31 | 0.05 |
| 749.00 | 761.00 | 808.67 | 1.05 | 0.30 |
| 7.00 | 7.00 | 5.33 | 1.21 | 0.73 |

| | | | | |
|---|---|---|---|---|
| 0.00 | 0.00 | 0.67 | -1.87 | 0.70 |
| 10.00 | 7.00 | 7.67 | -1.36 | 0.38 |
| 38.00 | 28.00 | 33.67 | -1.09 | 0.56 |
| 39.00 | 54.00 | 43.33 | 1.44 | 0.02 |
| 24.00 | 25.00 | 21.00 | -1.53 | 0.02 |
| 18.00 | 19.00 | 19.00 | -1.07 | 0.86 |
| 708.00 | 737.00 | 730.00 | 1.08 | 0.12 |
| 1222.00 | 1077.00 | 1134.67 | -1.32 | 0.00 |
| 297.00 | 314.00 | 315.67 | -1.30 | 0.00 |
| 3732.00 | 4080.00 | 4020.00 | -1.05 | 0.13 |
| 190.00 | 199.00 | 202.00 | 1.48 | 0.00 |
| 1096.00 | 1161.00 | 1162.33 | 1.04 | 0.33 |
| 959.00 | 1103.00 | 1081.67 | 1.15 | 0.00 |
| 705.00 | 765.00 | 754.00 | 1.08 | 0.13 |
| 938.00 | 1038.00 | 1034.33 | 1.22 | 0.00 |
| 982.00 | 1120.00 | 1129.00 | 1.17 | 0.01 |
| 1852.00 | 1926.00 | 2018.33 | -1.12 | 0.01 |
| 233.00 | 209.00 | 237.33 | 1.01 | 0.92 |
| 854.00 | 793.00 | 844.67 | -1.06 | 0.25 |
| 5988.00 | 6422.00 | 6303.33 | -1.16 | 0.00 |
| 1635.00 | 1555.00 | 1617.67 | -1.07 | 0.11 |
| 128.00 | 124.00 | 132.67 | -1.02 | 0.80 |
| 402.00 | 340.00 | 386.67 | 1.00 | 0.98 |
| 513.00 | 529.00 | 519.33 | 1.17 | 0.00 |
| 978.00 | 915.00 | 957.00 | 1.12 | 0.02 |
| 943.00 | 965.00 | 936.67 | 1.01 | 0.88 |
| 1206.00 | 1328.00 | 1301.33 | -1.06 | 0.15 |
| 393.00 | 374.00 | 397.00 | -1.04 | 0.48 |
| 1015.00 | 1102.00 | 1081.00 | 1.28 | 0.00 |
| 497.00 | 463.00 | 487.67 | -1.42 | 0.00 |
| 1105.00 | 1357.00 | 1179.67 | -1.01 | 0.91 |
| 1565.00 | 1664.00 | 1669.67 | -1.07 | 0.09 |
| 28.00 | 43.00 | 39.33 | -2.14 | 0.00 |
| 78.00 | 76.00 | 78.00 | 1.12 | 0.33 |
| 956.00 | 1047.00 | 1059.33 | -1.05 | 0.31 |
| 51.00 | 37.00 | 48.33 | 1.03 | 0.87 |
| 806.00 | 988.00 | 949.33 | -1.14 | 0.01 |
| 453.00 | 540.00 | 499.00 | 1.08 | 0.17 |
| 10.00 | 10.00 | 12.67 | 1.01 | 1.00 |
| 167.00 | 161.00 | 180.33 | -1.21 | 0.01 |
| 693.00 | 810.00 | 752.33 | 1.12 | 0.02 |
| 5970.00 | 6046.00 | 6048.67 | -1.32 | 0.00 |
| 5.00 | 12.00 | 10.00 | -1.32 | 0.36 |
| 40.00 | 46.00 | 44.67 | 1.11 | 0.46 |
| 1072.00 | 1223.00 | 1284.00 | 1.13 | 0.04 |
| 1326.00 | 1295.00 | 1365.67 | 1.09 | 0.03 |
| 479.00 | 518.00 | 527.00 | 1.45 | 0.00 |
| 1378.00 | 1425.00 | 1375.33 | 1.17 | 0.00 |

| | | | | |
|---|---|---|---|---|
| 10.00 | 1.00 | 6.00 | -2.77 | 0.00 |
| 0.00 | 0.00 | 0.00 | -11.75 | 0.13 |
| 2644.00 | 2609.00 | 2729.67 | 1.05 | 0.26 |
| 111.00 | 151.00 | 146.33 | -1.13 | 0.23 |
| 725.00 | 863.00 | 792.00 | -1.12 | 0.02 |
| 1508.00 | 1653.00 | 1573.33 | -1.06 | 0.22 |
| 1203.00 | 1272.00 | 1266.33 | -1.01 | 0.90 |
| 763.00 | 994.00 | 878.33 | 1.11 | 0.08 |
| 663.00 | 775.00 | 724.33 | 1.15 | 0.01 |
| 672.00 | 692.00 | 672.33 | -1.05 | 0.38 |
| 619.00 | 663.00 | 622.67 | -1.07 | 0.25 |
| 66.00 | 73.00 | 67.67 | 1.19 | 0.13 |
| 310.00 | 338.00 | 342.00 | -1.06 | 0.31 |
| 83.00 | 57.00 | 69.67 | 1.15 | 0.25 |
| 36.00 | 41.00 | 39.00 | -1.65 | 0.00 |
| 2128.00 | 2441.00 | 2444.00 | 1.02 | 0.69 |
| 2028.00 | 2202.00 | 2140.00 | 1.24 | 0.00 |
| 9.00 | 12.00 | 10.33 | 1.60 | 0.13 |
| 1.00 | 3.00 | 2.33 | -1.57 | 0.50 |
| 1.00 | 0.00 | 1.33 | -3.33 | 0.03 |
| 1363.00 | 1395.00 | 1428.67 | 1.03 | 0.51 |
| 387.00 | 400.00 | 403.00 | 1.02 | 0.68 |
| 736.00 | 810.00 | 782.33 | 1.15 | 0.01 |
| 476.00 | 535.00 | 530.33 | -1.07 | 0.15 |
| 1538.00 | 1584.00 | 1572.33 | -1.08 | 0.03 |
| 402.00 | 475.00 | 433.67 | -1.10 | 0.12 |
| 194.00 | 209.00 | 211.00 | -1.39 | 0.00 |
| 417.00 | 498.00 | 486.00 | -1.02 | 0.73 |
| 50.00 | 34.00 | 39.67 | 1.11 | 0.52 |
| 6.00 | 6.00 | 8.67 | -1.28 | 0.46 |
| 9.00 | 7.00 | 7.67 | 1.26 | 0.54 |
| 143.00 | 120.00 | 120.00 | -1.18 | 0.15 |
| 124.00 | 114.00 | 119.00 | -1.15 | 0.12 |
| 8.00 | 9.00 | 7.33 | 1.03 | 1.00 |
| 1.00 | 5.00 | 2.67 | -3.43 | 0.00 |
| 2351.00 | 2618.00 | 2510.67 | -1.13 | 0.00 |
| 1345.00 | 1441.00 | 1429.00 | -1.08 | 0.06 |
| 3499.00 | 3723.00 | 3702.00 | -1.11 | 0.01 |
| 1175.00 | 1193.00 | 1214.67 | -1.01 | 0.80 |
| 2637.00 | 2733.00 | 2732.00 | 1.03 | 0.44 |
| 788.00 | 825.00 | 834.33 | -1.09 | 0.04 |
| 43.00 | 64.00 | 54.00 | 1.48 | 0.00 |
| 21091.00 | 20918.00 | 21336.00 | -1.10 | 0.02 |
| 302.00 | 272.00 | 301.33 | -1.17 | 0.01 |
| 6006.00 | 6704.00 | 6566.00 | 1.20 | 0.00 |
| 134.00 | 163.00 | 153.00 | 1.11 | 0.24 |
| 515.00 | 453.00 | 503.67 | 1.02 | 0.69 |
| 42209.00 | 45257.00 | 43499.00 | 1.14 | 0.00 |

| | | | | |
|---|---|---|---|---|
| 4297.00 | 4331.00 | 4466.33 | -1.06 | 0.10 |
| 157.00 | 186.00 | 171.00 | 1.16 | 0.08 |
| 7935.00 | 8779.00 | 8722.33 | -1.00 | 0.94 |
| 874.00 | 940.00 | 949.67 | -1.01 | 0.91 |
| 34.00 | 33.00 | 30.33 | -1.22 | 0.21 |
| 1.00 | 1.00 | 2.33 | -1.29 | 0.81 |
| 261.00 | 340.00 | 305.67 | 1.02 | 0.78 |
| 2017.00 | 1849.00 | 1890.00 | -1.22 | 0.00 |
| 1293.00 | 1179.00 | 1264.33 | -1.18 | 0.00 |
| 12.00 | 20.00 | 20.33 | -1.46 | 0.05 |
| 1128.00 | 1082.00 | 1116.67 | -1.03 | 0.48 |
| 529.00 | 558.00 | 565.67 | -1.03 | 0.56 |
| 0.00 | 2.00 | 1.33 | -2.63 | 0.12 |
| 797.00 | 850.00 | 847.00 | -1.02 | 0.69 |
| 4047.00 | 4637.00 | 4381.33 | 1.03 | 0.52 |
| 1259.00 | 1366.00 | 1340.00 | 1.18 | 0.00 |
| 351.00 | 375.00 | 387.67 | 1.29 | 0.00 |
| 347.00 | 384.00 | 377.00 | -1.07 | 0.19 |
| 606.00 | 580.00 | 560.67 | -1.44 | 0.00 |
| 8.00 | 5.00 | 8.33 | 3.93 | 0.00 |
| 550.00 | 489.00 | 536.33 | 1.01 | 0.80 |
| 4284.00 | 4705.00 | 4585.33 | 1.21 | 0.00 |
| 1620.00 | 1713.00 | 1764.00 | 1.10 | 0.03 |
| 1132.00 | 1144.00 | 1181.67 | 1.19 | 0.00 |
| 83.00 | 90.00 | 85.00 | -1.05 | 0.65 |
| 1006.00 | 1229.00 | 1137.67 | 1.11 | 0.02 |
| 1319.00 | 1430.00 | 1418.00 | 1.16 | 0.00 |
| 2240.00 | 2361.00 | 2340.33 | -1.01 | 0.87 |
| 158.00 | 164.00 | 149.67 | -1.03 | 0.73 |
| 692.00 | 696.00 | 717.00 | 1.16 | 0.00 |
| 4.00 | 6.00 | 4.00 | 1.66 | 0.37 |
| 479.00 | 592.00 | 563.00 | -1.03 | 0.62 |
| 370.00 | 403.00 | 397.67 | 1.31 | 0.00 |
| 69.00 | 69.00 | 69.67 | -1.16 | 0.18 |
| 2.00 | 8.00 | 4.67 | -1.72 | 0.17 |
| 3629.00 | 3765.00 | 3687.33 | -1.27 | 0.00 |
| 0.00 | 0.00 | 0.00 | -17.12 | 0.03 |
| 1707.00 | 1751.00 | 1752.67 | -1.65 | 0.00 |
| 2197.00 | 2191.00 | 2316.33 | -1.38 | 0.00 |
| 904.00 | 1171.00 | 1062.00 | -1.04 | 0.40 |
| 54.00 | 49.00 | 55.67 | -1.29 | 0.03 |
| 265.00 | 285.00 | 293.00 | 1.15 | 0.02 |
| 653.00 | 661.00 | 677.00 | 1.10 | 0.03 |
| 308.00 | 270.00 | 267.00 | -1.25 | 0.02 |
| 30.00 | 21.00 | 22.67 | 1.03 | 0.94 |
| 500.00 | 480.00 | 493.33 | 1.00 | 1.00 |
| 150.00 | 186.00 | 162.33 | -1.24 | 0.01 |
| 45.00 | 36.00 | 40.67 | 1.24 | 0.14 |

| | | | | |
|---|---|---|---|---|
| 791.00 | 904.00 | 849.67 | 1.03 | 0.57 |
| 559.00 | 541.00 | 528.67 | 1.04 | 0.56 |
| 1196.00 | 1368.00 | 1298.00 | -1.10 | 0.02 |
| 389.00 | 367.00 | 374.00 | 1.34 | 0.00 |
| 3804.00 | 4061.00 | 3965.00 | 1.12 | 0.01 |
| 478.00 | 502.00 | 490.00 | 1.04 | 0.41 |
| 136.00 | 143.00 | 134.33 | -2.04 | 0.00 |
| 1506.00 | 1326.00 | 1293.00 | -1.55 | 0.00 |
| 749.00 | 780.00 | 777.00 | 1.04 | 0.41 |
| 683.00 | 748.00 | 730.00 | 1.04 | 0.35 |
| 697.00 | 772.00 | 718.00 | -1.02 | 0.79 |
| 489.00 | 568.00 | 543.00 | 1.39 | 0.00 |
| 58.00 | 47.00 | 48.67 | 1.03 | 0.83 |
| 201.00 | 205.00 | 214.33 | -1.35 | 0.00 |
| 965.00 | 977.00 | 1030.33 | 1.06 | 0.23 |
| 733.00 | 626.00 | 670.00 | -1.38 | 0.00 |
| 1953.00 | 2074.00 | 2019.00 | -1.07 | 0.06 |
| 14.00 | 18.00 | 16.00 | -1.12 | 0.70 |
| 133.00 | 85.00 | 106.67 | -1.07 | 0.56 |
| 34.00 | 26.00 | 28.33 | 1.10 | 0.61 |
| 60.00 | 89.00 | 80.00 | 1.21 | 0.12 |
| 3.00 | 1.00 | 3.00 | -1.23 | 0.83 |
| 15.00 | 5.00 | 8.67 | 1.35 | 0.40 |
| 664.00 | 778.00 | 724.00 | 1.06 | 0.21 |
| 131.00 | 132.00 | 119.33 | -1.42 | 0.00 |
| 29.00 | 30.00 | 34.33 | -1.08 | 0.61 |
| 2934.00 | 3319.00 | 3355.33 | 1.14 | 0.00 |
| 700.00 | 751.00 | 773.33 | 1.07 | 0.23 |
| 3.00 | 0.00 | 1.33 | -2.86 | 0.08 |
| 5.00 | 3.00 | 3.00 | -2.09 | 0.09 |
| 3412.00 | 3688.00 | 3525.33 | 1.24 | 0.00 |
| 700.00 | 694.00 | 740.33 | -1.14 | 0.02 |
| 210.00 | 246.00 | 226.67 | 1.03 | 0.67 |
| 379.00 | 351.00 | 370.33 | 1.09 | 0.16 |
| 772.00 | 881.00 | 832.67 | -1.09 | 0.08 |
| 180.00 | 242.00 | 226.67 | 1.11 | 0.19 |
| 734.00 | 791.00 | 795.33 | 1.02 | 0.65 |
| 998.00 | 967.00 | 1028.33 | -1.15 | 0.00 |
| 2170.00 | 2113.00 | 2143.00 | -1.27 | 0.00 |
| 1338.00 | 1478.00 | 1482.33 | -1.18 | 0.00 |
| 480.00 | 600.00 | 537.00 | -1.09 | 0.13 |
| 245.00 | 227.00 | 237.33 | -1.21 | 0.00 |
| 450.00 | 526.00 | 486.67 | 1.24 | 0.00 |
| 183.00 | 211.00 | 197.33 | -1.02 | 0.83 |
| 0.00 | 2.00 | 2.33 | -1.56 | 0.49 |
| 57.00 | 77.00 | 67.00 | 1.22 | 0.11 |
| 1754.00 | 1844.00 | 1869.67 | -1.23 | 0.00 |
| 921.00 | 1022.00 | 1027.00 | 1.15 | 0.00 |

| | | | | |
|---|---|---|---|---|
| 414.00 | 415.00 | 432.00 | 1.02 | 0.71 |
| 4.00 | 12.00 | 7.67 | -1.06 | 1.00 |
| 1111.00 | 1127.00 | 1144.33 | -1.06 | 0.17 |
| 22.00 | 18.00 | 26.00 | -1.11 | 0.63 |
| 1041.00 | 1132.00 | 1116.67 | -1.03 | 0.49 |
| 416.00 | 410.00 | 391.67 | -1.08 | 0.27 |
| 531.00 | 563.00 | 555.67 | -1.07 | 0.16 |
| 17.00 | 20.00 | 19.33 | -1.33 | 0.11 |
| 8.00 | 8.00 | 8.67 | -1.24 | 0.53 |
| 143.00 | 162.00 | 151.67 | 1.12 | 0.16 |
| 1552.00 | 1957.00 | 1776.00 | -1.03 | 0.51 |
| 16.00 | 44.00 | 28.67 | -1.16 | 0.43 |
| 706.00 | 789.00 | 771.00 | -1.00 | 0.99 |
| 243.00 | 281.00 | 282.00 | -1.10 | 0.17 |
| 100.00 | 107.00 | 108.00 | -1.22 | 0.02 |
| 471.00 | 614.00 | 566.67 | 1.08 | 0.21 |
| 449.00 | 597.00 | 530.33 | 1.59 | 0.00 |
| 127.00 | 147.00 | 142.00 | 1.13 | 0.14 |
| 1397.00 | 1508.00 | 1430.00 | -1.12 | 0.01 |
| 32.00 | 16.00 | 21.67 | -1.03 | 1.00 |
| 472.00 | 539.00 | 532.67 | 1.71 | 0.00 |
| 1014.00 | 948.00 | 991.00 | -1.25 | 0.00 |
| 850.00 | 871.00 | 882.33 | 1.01 | 0.75 |
| 8.00 | 12.00 | 9.67 | -1.29 | 0.41 |
| 1192.00 | 1235.00 | 1284.67 | 1.35 | 0.00 |
| 252.00 | 303.00 | 292.00 | 1.01 | 0.87 |
| 4846.00 | 5565.00 | 5480.00 | 1.14 | 0.00 |
| 195.00 | 200.00 | 218.67 | -1.20 | 0.01 |
| 10122.00 | 11236.00 | 11333.33 | 1.07 | 0.08 |
| 5.00 | 13.00 | 8.67 | -1.28 | 0.49 |
| 6.00 | 10.00 | 7.33 | -1.78 | 0.04 |
| 602.00 | 594.00 | 631.00 | -1.08 | 0.14 |
| 294.00 | 369.00 | 337.67 | -1.23 | 0.00 |
| 1062.00 | 1067.00 | 1088.00 | 1.06 | 0.12 |
| 0.00 | 0.00 | 0.67 | -2.30 | 0.46 |
| 39.00 | 40.00 | 38.67 | -1.27 | 0.08 |
| 240.00 | 281.00 | 263.00 | 1.08 | 0.29 |
| 14.00 | 21.00 | 17.67 | 1.27 | 0.27 |
| 9.00 | 7.00 | 8.00 | 1.38 | 0.36 |
| 9.00 | 16.00 | 13.00 | -2.53 | 0.00 |
| 47.00 | 50.00 | 46.67 | -1.33 | 0.02 |
| 2.00 | 3.00 | 2.67 | -1.38 | 0.65 |
| 6.00 | 4.00 | 3.67 | -1.99 | 0.09 |
| 11.00 | 10.00 | 12.00 | -1.21 | 0.51 |
| 6.00 | 4.00 | 5.67 | -1.01 | 1.00 |
| 24.00 | 17.00 | 19.67 | 1.38 | 0.13 |
| 32.00 | 34.00 | 33.67 | 1.12 | 0.58 |
| 50.00 | 68.00 | 60.33 | 1.02 | 0.93 |

| | | | | |
|---|---|---|---|---|
| 49.00 | 63.00 | 56.67 | -1.25 | 0.06 |
| 44.00 | 43.00 | 41.67 | -1.01 | 0.95 |
| 2.00 | 3.00 | 1.67 | -1.58 | 0.59 |
| 0.00 | 2.00 | 1.67 | -2.34 | 0.15 |
| 30.00 | 39.00 | 39.33 | -1.36 | 0.02 |
| 8.00 | 4.00 | 5.67 | 2.33 | 0.07 |
| 2.00 | 4.00 | 4.67 | -1.37 | 0.50 |
| 57.00 | 52.00 | 56.00 | 1.31 | 0.03 |
| 29.00 | 52.00 | 43.33 | 1.29 | 0.09 |
| 12.00 | 12.00 | 13.33 | 1.19 | 0.49 |
| 9.00 | 12.00 | 11.33 | -1.07 | 0.91 |
| 19.00 | 17.00 | 15.33 | 1.37 | 0.20 |
| 16.00 | 23.00 | 18.67 | -1.79 | 0.00 |
| 51.00 | 49.00 | 47.33 | -2.33 | 0.00 |
| 154.00 | 155.00 | 160.67 | -2.09 | 0.00 |
| 4.00 | 3.00 | 3.67 | -1.99 | 0.09 |
| 20.00 | 20.00 | 21.67 | -1.11 | 0.56 |
| 9.00 | 13.00 | 9.67 | -1.25 | 0.47 |
| 8.00 | 11.00 | 9.33 | -2.08 | 0.00 |
| 12.00 | 12.00 | 11.67 | -1.18 | 0.58 |
| 48.00 | 34.00 | 38.00 | 1.28 | 0.12 |
| 9.00 | 16.00 | 11.33 | -1.07 | 0.91 |
| 5.00 | 2.00 | 3.67 | -3.68 | 0.00 |
| 29.00 | 38.00 | 34.33 | -1.72 | 0.00 |
| 25.00 | 18.00 | 17.33 | -1.48 | 0.05 |
| 20.00 | 9.00 | 11.00 | -1.73 | 0.04 |
| 26.00 | 28.00 | 26.67 | -1.44 | 0.02 |
| 152.00 | 94.00 | 121.67 | -1.77 | 0.00 |
| 10.00 | 16.00 | 15.67 | -1.10 | 0.71 |
| 15.00 | 10.00 | 11.67 | -2.10 | 0.00 |
| 499.00 | 437.00 | 446.67 | -1.99 | 0.00 |
| 5.00 | 5.00 | 5.00 | -2.73 | 0.00 |
| 5.00 | 6.00 | 5.00 | -3.19 | 0.00 |
| 3451.00 | 3291.00 | 3545.33 | -1.04 | 0.34 |
| 269.00 | 311.00 | 309.33 | -1.39 | 0.00 |
| 1664.00 | 1908.00 | 1795.67 | -1.14 | 0.00 |
| 455.00 | 456.00 | 490.67 | 1.10 | 0.12 |
| 642.00 | 646.00 | 681.00 | -1.06 | 0.25 |
| 801.00 | 960.00 | 857.67 | -1.01 | 0.84 |
| 391.00 | 441.00 | 422.33 | 1.31 | 0.00 |
| 1391.00 | 1518.00 | 1526.33 | 1.02 | 0.67 |
| 470.00 | 479.00 | 514.67 | -1.09 | 0.20 |
| 4299.00 | 4597.00 | 4588.33 | 1.02 | 0.68 |
| 359.00 | 342.00 | 348.00 | -1.37 | 0.00 |
| 1590.00 | 1902.00 | 1783.00 | 1.01 | 0.85 |
| 1562.00 | 1654.00 | 1683.67 | 1.18 | 0.00 |
| 768.00 | 843.00 | 806.67 | -1.04 | 0.41 |
| 633.00 | 881.00 | 758.33 | 1.12 | 0.12 |

| | | | | |
|---|---|---|---|---|
| 1169.00 | 1479.00 | 1394.33 | 1.41 | 0.00 |
| 2031.00 | 2474.00 | 2263.67 | 1.19 | 0.00 |
| 2122.00 | 2242.00 | 2176.00 | -1.12 | 0.00 |
| 1965.00 | 2194.00 | 2060.00 | -1.19 | 0.00 |
| 943.00 | 966.00 | 958.33 | -1.30 | 0.00 |
| 3386.00 | 3309.00 | 3394.67 | -1.44 | 0.00 |
| 2233.00 | 2436.00 | 2374.33 | -1.05 | 0.19 |
| 4822.00 | 5321.00 | 5270.67 | -1.20 | 0.00 |
| 503.00 | 576.00 | 573.00 | 1.14 | 0.01 |
| 63.00 | 47.00 | 53.67 | -1.07 | 0.62 |
| 2.00 | 5.00 | 3.67 | -2.08 | 0.06 |
| 92.00 | 107.00 | 107.67 | -1.45 | 0.00 |
| 1163.00 | 1113.00 | 1165.00 | -1.08 | 0.09 |
| 332.00 | 350.00 | 352.33 | 1.19 | 0.00 |
| 1454.00 | 1453.00 | 1436.67 | -1.26 | 0.00 |
| 452.00 | 466.00 | 467.33 | -1.04 | 0.44 |
| 535.00 | 510.00 | 513.33 | -1.08 | 0.19 |
| 663.00 | 652.00 | 680.33 | 1.00 | 0.95 |
| 2719.00 | 2752.00 | 2952.00 | 1.12 | 0.02 |
| 1.00 | 2.00 | 1.00 | -2.81 | 0.15 |
| 789.00 | 914.00 | 871.00 | 1.24 | 0.00 |
| 5559.00 | 5733.00 | 5578.67 | -1.14 | 0.00 |
| 1065.00 | 1108.00 | 1127.67 | 1.07 | 0.09 |
| 1657.00 | 1780.00 | 1806.33 | 1.20 | 0.00 |
| 248.00 | 268.00 | 290.67 | 1.28 | 0.00 |
| 2105.00 | 2609.00 | 2391.00 | -1.15 | 0.00 |
| 148.00 | 191.00 | 172.67 | 1.06 | 0.51 |
| 1084.00 | 1181.00 | 1238.00 | 1.23 | 0.00 |
| 3117.00 | 3554.00 | 3557.00 | 1.18 | 0.00 |
| 4297.00 | 5150.00 | 4827.33 | 1.06 | 0.16 |
| 38.00 | 41.00 | 36.00 | 1.12 | 0.49 |
| 870.00 | 962.00 | 957.67 | 1.08 | 0.09 |
| 243.00 | 267.00 | 251.33 | -1.05 | 0.45 |
| 377.00 | 412.00 | 406.33 | 1.15 | 0.01 |
| 38.00 | 50.00 | 47.33 | 1.17 | 0.27 |
| 2610.00 | 2680.00 | 2647.67 | -1.12 | 0.01 |
| 17.00 | 14.00 | 15.33 | -1.21 | 0.45 |
| 22.00 | 17.00 | 16.00 | 1.10 | 0.70 |
| 680.00 | 769.00 | 736.00 | 1.03 | 0.46 |
| 1246.00 | 1407.00 | 1305.33 | 1.12 | 0.01 |
| 128.00 | 118.00 | 129.33 | -1.23 | 0.01 |
| 16.00 | 17.00 | 14.00 | -1.47 | 0.09 |
| 628.00 | 622.00 | 619.33 | 1.28 | 0.00 |
| 453.00 | 451.00 | 466.67 | -1.16 | 0.01 |
| 192.00 | 230.00 | 218.00 | 1.06 | 0.45 |
| 8.00 | 8.00 | 8.67 | -1.59 | 0.10 |
| 705.00 | 820.00 | 783.00 | -1.08 | 0.10 |
| 2632.00 | 2570.00 | 2626.67 | -1.13 | 0.00 |

| | | | | |
|---|---|---|---|---|
| 99.00 | 114.00 | 103.67 | 1.10 | 0.31 |
| 462.00 | 478.00 | 490.00 | 1.08 | 0.12 |
| 10.00 | 6.00 | 6.67 | 1.16 | 0.75 |
| 712.00 | 808.00 | 783.00 | 1.02 | 0.63 |
| 178.00 | 219.00 | 201.00 | 2.13 | 0.00 |
| 761.00 | 848.00 | 828.67 | -1.12 | 0.01 |
| 69.00 | 77.00 | 74.33 | 1.17 | 0.18 |
| 752.00 | 835.00 | 813.67 | -1.74 | 0.00 |
| 10.00 | 13.00 | 13.00 | -2.23 | 0.00 |
| 2101.00 | 2440.00 | 2325.33 | -1.11 | 0.01 |
| 119.00 | 121.00 | 121.00 | 1.07 | 0.43 |
| 992.00 | 1015.00 | 998.33 | -1.35 | 0.00 |
| 32.00 | 37.00 | 32.67 | -1.95 | 0.00 |
| 70.00 | 81.00 | 76.00 | -1.43 | 0.00 |
| 1163.00 | 1280.00 | 1268.00 | 1.05 | 0.26 |
| 966.00 | 1045.00 | 1044.67 | 1.23 | 0.00 |
| 731.00 | 928.00 | 854.00 | 1.21 | 0.00 |
| 13518.00 | 15089.00 | 14383.67 | -1.09 | 0.02 |
| 1267.00 | 1475.00 | 1378.67 | -1.05 | 0.25 |
| 12696.00 | 13762.00 | 13471.67 | -1.08 | 0.05 |
| 1286.00 | 1318.00 | 1334.33 | 1.04 | 0.37 |
| 13287.00 | 14411.00 | 14197.67 | 1.03 | 0.41 |
| 119.00 | 137.00 | 140.67 | -1.12 | 0.19 |
| 688.00 | 749.00 | 730.67 | 1.32 | 0.00 |
| 269.00 | 259.00 | 273.67 | -1.05 | 0.44 |
| 174.00 | 188.00 | 187.33 | -1.05 | 0.47 |
| 13.00 | 10.00 | 12.33 | 1.58 | 0.10 |
| 646.00 | 714.00 | 752.00 | -1.27 | 0.00 |
| 495.00 | 441.00 | 489.33 | 1.15 | 0.03 |
| 35.00 | 40.00 | 34.33 | -1.75 | 0.00 |
| 563.00 | 610.00 | 632.00 | -1.05 | 0.34 |
| 517.00 | 497.00 | 520.00 | 1.03 | 0.58 |
| 1878.00 | 1952.00 | 2010.67 | 1.14 | 0.00 |
| 3430.00 | 3997.00 | 3795.00 | -1.14 | 0.00 |
| 660.00 | 673.00 | 653.00 | -1.19 | 0.00 |
| 634.00 | 599.00 | 602.67 | -1.30 | 0.00 |
| 0.00 | 0.00 | 0.67 | -6.55 | 0.00 |
| 2089.00 | 1972.00 | 1983.67 | -1.21 | 0.00 |
| 544.00 | 581.00 | 591.33 | 1.18 | 0.00 |
| 5390.00 | 5836.00 | 5660.67 | 1.06 | 0.13 |
| 694.00 | 725.00 | 688.67 | 1.07 | 0.21 |
| 415.00 | 502.00 | 465.00 | 1.38 | 0.00 |
| 429.00 | 504.00 | 468.33 | 1.13 | 0.03 |
| 95.00 | 86.00 | 97.33 | -1.09 | 0.38 |
| 398.00 | 354.00 | 386.00 | -1.51 | 0.00 |
| 1673.00 | 1618.00 | 1731.00 | -1.23 | 0.00 |
| 489.00 | 499.00 | 525.33 | 1.14 | 0.02 |
| 889.00 | 1029.00 | 1036.00 | 1.06 | 0.27 |

| | | | | |
|---|---|---|---|---|
| 16.00 | 12.00 | 11.67 | -2.04 | 0.00 |
| 4.00 | 0.00 | 2.67 | -1.74 | 0.30 |
| 1938.00 | 1989.00 | 1914.00 | 1.04 | 0.45 |
| 2.00 | 4.00 | 3.33 | -2.48 | 0.02 |
| 21.00 | 10.00 | 18.00 | -1.33 | 0.18 |
| 1.00 | 0.00 | 0.67 | -2.29 | 0.46 |
| 4925.00 | 5100.00 | 5258.67 | -1.19 | 0.00 |
| 1029.00 | 1063.00 | 1047.33 | -1.22 | 0.00 |
| 5727.00 | 5817.00 | 5905.33 | -1.28 | 0.00 |
| 3379.00 | 3723.00 | 3739.33 | 1.11 | 0.01 |
| 4.00 | 3.00 | 4.00 | -1.09 | 1.00 |
| 168.00 | 173.00 | 173.33 | 1.30 | 0.00 |
| 863.00 | 901.00 | 940.33 | -1.02 | 0.74 |
| 1300.00 | 1360.00 | 1349.67 | 1.15 | 0.00 |
| 217.00 | 293.00 | 261.67 | 1.11 | 0.17 |
| 31.00 | 31.00 | 34.00 | 1.08 | 0.63 |
| 112.00 | 93.00 | 105.33 | -1.63 | 0.00 |
| 307.00 | 352.00 | 340.33 | 1.01 | 0.93 |
| 2839.00 | 2816.00 | 2875.00 | 1.03 | 0.48 |
| 757.00 | 710.00 | 768.67 | 1.04 | 0.37 |
| 822.00 | 852.00 | 868.67 | 1.04 | 0.33 |
| 635.00 | 595.00 | 634.67 | -1.12 | 0.02 |
| 339.00 | 286.00 | 310.33 | -1.28 | 0.00 |
| 366.00 | 349.00 | 376.33 | 1.08 | 0.24 |
| 2381.00 | 2482.00 | 2475.33 | 1.09 | 0.03 |
| 2838.00 | 3110.00 | 3063.00 | 1.13 | 0.00 |
| 292.00 | 320.00 | 327.33 | 1.19 | 0.05 |
| 114.00 | 103.00 | 117.33 | -1.02 | 0.82 |
| 803.00 | 823.00 | 824.33 | -1.16 | 0.00 |
| 678.00 | 763.00 | 754.67 | 1.06 | 0.21 |
| 197.00 | 239.00 | 207.67 | 1.10 | 0.25 |
| 428.00 | 459.00 | 474.33 | 1.09 | 0.12 |
| 29.00 | 33.00 | 35.67 | 1.13 | 0.47 |
| 269.00 | 284.00 | 288.33 | 1.02 | 0.80 |
| 398.00 | 416.00 | 414.00 | 1.13 | 0.04 |
| 485.00 | 480.00 | 512.33 | 1.01 | 0.88 |
| 341.00 | 385.00 | 388.00 | -1.10 | 0.09 |
| 980.00 | 1023.00 | 1028.67 | 1.00 | 0.92 |
| 707.00 | 750.00 | 743.00 | 1.15 | 0.01 |
| 573.00 | 565.00 | 562.67 | -1.02 | 0.75 |
| 500.00 | 496.00 | 514.00 | 1.07 | 0.17 |
| 1239.00 | 1320.00 | 1322.00 | -1.14 | 0.00 |
| 329.00 | 333.00 | 344.00 | 2.56 | 0.00 |
| 2767.00 | 2804.00 | 2818.00 | -1.14 | 0.00 |
| 223.00 | 266.00 | 255.33 | 1.29 | 0.00 |
| 6.00 | 6.00 | 6.67 | -1.46 | 0.26 |
| 4650.00 | 4523.00 | 4622.00 | -1.05 | 0.20 |
| 2428.00 | 2566.00 | 2671.67 | 1.18 | 0.00 |

| | | | | |
|---|---|---|---|---|
| 1918.00 | 2247.00 | 2285.00 | 1.18 | 0.00 |
| 577.00 | 674.00 | 666.67 | 1.16 | 0.00 |
| 3545.00 | 4236.00 | 4214.33 | 1.13 | 0.01 |
| 0.00 | 0.00 | 0.00 | -17.09 | 0.03 |
| 388.00 | 427.00 | 410.67 | -1.06 | 0.26 |
| 1641.00 | 1616.00 | 1688.67 | 1.10 | 0.02 |
| 89.00 | 121.00 | 107.33 | -1.49 | 0.00 |
| 2078.00 | 2010.00 | 2096.00 | -1.04 | 0.34 |
| 3383.00 | 3546.00 | 3586.00 | -1.01 | 0.89 |
| 9671.00 | 9793.00 | 9868.00 | -1.15 | 0.00 |
| 21053.00 | 22231.00 | 22898.67 | 1.23 | 0.00 |
| 5689.00 | 6053.00 | 6048.67 | 1.30 | 0.00 |
| 19.00 | 26.00 | 22.00 | -1.07 | 0.74 |
| 1572.00 | 1611.00 | 1715.67 | 1.20 | 0.00 |
| 13.00 | 11.00 | 12.00 | 1.68 | 0.07 |
| 7.00 | 5.00 | 6.67 | -1.11 | 0.88 |
| 986.00 | 958.00 | 992.00 | 1.24 | 0.00 |
| 440.00 | 541.00 | 477.00 | -1.17 | 0.02 |
| 718.00 | 679.00 | 746.67 | 1.15 | 0.01 |
| 190.00 | 158.00 | 167.00 | -1.38 | 0.00 |
| 262.00 | 281.00 | 293.00 | -1.03 | 0.63 |
| 280.00 | 304.00 | 292.00 | 1.12 | 0.11 |
| 256.00 | 223.00 | 239.00 | -1.09 | 0.20 |
| 3380.00 | 3557.00 | 3517.33 | -1.01 | 0.86 |
| 523.00 | 524.00 | 526.67 | -1.12 | 0.04 |
| 181.00 | 201.00 | 189.00 | 1.01 | 0.90 |
| 10459.00 | 11421.00 | 11364.00 | 1.20 | 0.00 |
| 1842.00 | 1797.00 | 1888.33 | 1.08 | 0.05 |
| 642.00 | 681.00 | 686.67 | -1.01 | 0.75 |
| 1701.00 | 1886.00 | 1852.00 | 1.19 | 0.00 |
| 132.00 | 123.00 | 132.00 | -1.25 | 0.00 |
| 1595.00 | 1887.00 | 1788.00 | -1.06 | 0.13 |
| 37.00 | 35.00 | 31.00 | -1.24 | 0.42 |
| 709.00 | 565.00 | 609.00 | -1.90 | 0.00 |
| 32.00 | 42.00 | 39.67 | -1.07 | 0.69 |
| 762.00 | 710.00 | 770.67 | -1.10 | 0.06 |
| 419.00 | 458.00 | 460.00 | -1.48 | 0.00 |
| 110.00 | 113.00 | 103.67 | -1.06 | 0.59 |
| 1608.00 | 1499.00 | 1543.33 | 1.35 | 0.00 |
| 1067.00 | 1066.00 | 1093.33 | -1.01 | 0.83 |
| 1432.00 | 1420.00 | 1448.67 | -1.14 | 0.00 |
| 300.00 | 293.00 | 290.33 | 1.13 | 0.08 |
| 518.00 | 567.00 | 559.67 | 1.05 | 0.37 |
| 930.00 | 973.00 | 926.33 | 1.06 | 0.26 |
| 667.00 | 800.00 | 749.00 | 1.11 | 0.03 |
| 3186.00 | 3420.00 | 3515.33 | 1.17 | 0.00 |
| 5202.00 | 5469.00 | 5605.67 | 1.15 | 0.00 |
| 546.00 | 510.00 | 535.67 | -1.36 | 0.00 |

| | | | | |
|---|---|---|---|---|
| 3178.00 | 3259.00 | 3335.33 | -1.04 | 0.25 |
| 1259.00 | 1322.00 | 1244.33 | -1.11 | 0.09 |
| 65.00 | 59.00 | 66.33 | 1.14 | 0.28 |
| 56.00 | 66.00 | 68.67 | -1.26 | 0.04 |
| 722.00 | 763.00 | 758.67 | -1.09 | 0.06 |
| 121.00 | 127.00 | 131.67 | -1.04 | 0.64 |
| 905.00 | 959.00 | 936.00 | -1.26 | 0.00 |
| 561.00 | 589.00 | 589.67 | 1.03 | 0.53 |
| 1457.00 | 1734.00 | 1639.00 | 1.18 | 0.00 |
| 1004.00 | 1037.00 | 1055.33 | 1.18 | 0.00 |
| 783.00 | 805.00 | 781.67 | -1.11 | 0.03 |
| 1426.00 | 1335.00 | 1378.33 | -1.05 | 0.32 |
| 2022.00 | 2245.00 | 2145.33 | -1.01 | 0.75 |
| 788.00 | 778.00 | 816.33 | -1.12 | 0.02 |
| 1410.00 | 1648.00 | 1605.67 | 1.07 | 0.13 |
| 1686.00 | 1770.00 | 1702.33 | -1.10 | 0.06 |
| 635.00 | 637.00 | 688.33 | 1.27 | 0.00 |
| 942.00 | 1070.00 | 1007.33 | 1.12 | 0.02 |
| 115.00 | 116.00 | 119.33 | -1.15 | 0.11 |
| 1031.00 | 1123.00 | 1076.00 | -1.32 | 0.00 |
| 12678.00 | 13545.00 | 13638.33 | 1.10 | 0.00 |
| 1712.00 | 2106.00 | 1860.67 | 1.01 | 0.91 |
| 634.00 | 672.00 | 655.00 | -1.13 | 0.01 |
| 659.00 | 683.00 | 671.00 | -1.16 | 0.00 |
| 948.00 | 1075.00 | 1037.00 | -1.07 | 0.11 |
| 20.00 | 26.00 | 19.67 | -1.34 | 0.15 |
| 679.00 | 710.00 | 726.00 | -1.09 | 0.05 |
| 6659.00 | 7220.00 | 7103.00 | -1.16 | 0.00 |
| 179.00 | 173.00 | 190.00 | -1.25 | 0.00 |
| 1742.00 | 1639.00 | 1746.00 | -1.13 | 0.01 |
| 258.00 | 340.00 | 286.33 | -3.44 | 0.00 |
| 92.00 | 115.00 | 112.00 | -1.05 | 0.60 |
| 324.00 | 313.00 | 295.33 | -1.32 | 0.00 |
| 558.00 | 617.00 | 593.33 | -1.11 | 0.04 |
| 53.00 | 61.00 | 53.33 | -1.21 | 0.12 |
| 943.00 | 1013.00 | 1076.00 | -1.02 | 0.69 |
| 28.00 | 32.00 | 36.67 | 1.18 | 0.33 |
| 531.00 | 531.00 | 533.00 | 1.05 | 0.33 |
| 866.00 | 960.00 | 909.00 | 1.24 | 0.00 |
| 506.00 | 513.00 | 532.00 | 1.02 | 0.66 |
| 144.00 | 119.00 | 134.67 | 1.07 | 0.50 |
| 10.00 | 7.00 | 10.67 | -1.99 | 0.01 |
| 627.00 | 704.00 | 675.67 | -1.11 | 0.04 |
| 1.00 | 2.00 | 1.67 | -2.33 | 0.15 |
| 987.00 | 919.00 | 975.67 | -1.09 | 0.05 |
| 308.00 | 308.00 | 309.67 | 1.39 | 0.00 |
| 1426.00 | 1343.00 | 1396.00 | -1.23 | 0.00 |
| 59.00 | 68.00 | 67.00 | -1.26 | 0.06 |

| | | | | |
|---|---|---|---|---|
| 106.00 | 118.00 | 128.00 | -1.22 | 0.02 |
| 1756.00 | 1814.00 | 1861.00 | -1.04 | 0.35 |
| 9.00 | 4.00 | 5.33 | -1.76 | 0.11 |
| 288.00 | 319.00 | 311.00 | -1.10 | 0.08 |
| 560.00 | 625.00 | 601.33 | 1.01 | 0.89 |
| 832.00 | 913.00 | 960.00 | 1.07 | 0.19 |
| 479.00 | 473.00 | 476.67 | 1.01 | 0.91 |
| 479.00 | 540.00 | 536.00 | 1.14 | 0.01 |
| 2706.00 | 2875.00 | 2772.00 | 1.02 | 0.59 |
| 209.00 | 226.00 | 226.00 | -1.06 | 0.53 |
| 26.00 | 27.00 | 29.67 | 1.19 | 0.33 |
| 472.00 | 487.00 | 508.67 | -1.21 | 0.00 |
| 7058.00 | 6333.00 | 6734.00 | -1.21 | 0.00 |
| 773.00 | 821.00 | 852.33 | 1.07 | 0.20 |
| 170.00 | 262.00 | 221.67 | 1.07 | 0.48 |
| 502.00 | 522.00 | 528.33 | -1.04 | 0.52 |
| 187.00 | 202.00 | 213.00 | 1.14 | 0.09 |
| 670.00 | 677.00 | 727.67 | 1.06 | 0.31 |
| 570.00 | 614.00 | 630.67 | 1.06 | 0.31 |
| 935.00 | 950.00 | 964.67 | -1.23 | 0.00 |
| 424.00 | 418.00 | 444.33 | 1.16 | 0.01 |
| 2507.00 | 2850.00 | 2669.00 | 1.01 | 0.82 |
| 215.00 | 188.00 | 221.67 | -1.12 | 0.15 |
| 549.00 | 623.00 | 606.00 | -1.12 | 0.02 |
| 807.00 | 833.00 | 779.67 | -1.07 | 0.28 |
| 1786.00 | 1677.00 | 1740.00 | -1.16 | 0.00 |
| 227.00 | 229.00 | 252.00 | 1.14 | 0.07 |
| 977.00 | 914.00 | 993.67 | -1.02 | 0.64 |
| 999.00 | 981.00 | 1019.00 | -1.04 | 0.41 |
| 4458.00 | 4143.00 | 4329.67 | -1.07 | 0.09 |
| 1159.00 | 1239.00 | 1249.00 | -1.10 | 0.03 |
| 719.00 | 716.00 | 758.67 | -1.02 | 0.65 |
| 255.00 | 277.00 | 273.67 | 1.01 | 0.90 |
| 160.00 | 179.00 | 181.33 | -1.01 | 0.92 |
| 755.00 | 760.00 | 811.00 | 1.03 | 0.62 |
| 49.00 | 44.00 | 44.67 | 1.00 | 1.00 |
| 1097.00 | 1269.00 | 1157.33 | -1.19 | 0.00 |
| 15.00 | 27.00 | 19.33 | 1.27 | 0.34 |
| 590.00 | 614.00 | 583.00 | 1.02 | 0.68 |
| 551.00 | 526.00 | 538.33 | -1.06 | 0.39 |
| 677.00 | 722.00 | 692.67 | 1.03 | 0.60 |
| 1251.00 | 1172.00 | 1231.33 | -1.25 | 0.00 |
| 30.00 | 34.00 | 33.67 | 1.87 | 0.00 |
| 169.00 | 161.00 | 154.67 | 1.17 | 0.10 |
| 25.00 | 15.00 | 18.33 | 1.23 | 0.34 |
| 593.00 | 566.00 | 553.00 | 1.03 | 0.70 |
| 1401.00 | 1297.00 | 1371.33 | -1.16 | 0.00 |
| 62.00 | 57.00 | 57.67 | -1.01 | 0.96 |

| | | | | |
|---|---|---|---|---|
| 1344.00 | 1409.00 | 1421.00 | -1.13 | 0.00 |
| 1701.00 | 1987.00 | 1794.67 | -1.07 | 0.18 |
| 7.00 | 6.00 | 7.00 | -1.35 | 0.40 |
| 22.00 | 31.00 | 29.33 | 1.26 | 0.23 |
| 966.00 | 1056.00 | 1037.33 | 1.18 | 0.00 |
| 147.00 | 195.00 | 169.67 | 1.25 | 0.01 |
| 1375.00 | 1367.00 | 1411.33 | 1.23 | 0.00 |
| 792.00 | 843.00 | 856.00 | 1.04 | 0.34 |
| 389.00 | 458.00 | 450.00 | 1.20 | 0.00 |
| 18.00 | 9.00 | 11.67 | -1.41 | 0.18 |
| 956.00 | 963.00 | 986.00 | -1.31 | 0.00 |
| 285.00 | 311.00 | 299.00 | -1.12 | 0.06 |
| 34.00 | 29.00 | 32.67 | -2.24 | 0.00 |
| 242.00 | 252.00 | 257.67 | 1.05 | 0.51 |
| 1365.00 | 1285.00 | 1378.00 | -1.26 | 0.00 |
| 723.00 | 717.00 | 758.00 | -1.09 | 0.08 |
| 2391.00 | 2586.00 | 2556.67 | -1.04 | 0.32 |
| 1045.00 | 1042.00 | 1048.67 | -1.15 | 0.00 |
| 35.00 | 30.00 | 31.67 | -1.18 | 0.30 |
| 31.00 | 33.00 | 29.00 | -1.14 | 0.51 |
| 53.00 | 67.00 | 56.67 | 1.48 | 0.00 |
| 1085.00 | 1120.00 | 1150.33 | -1.16 | 0.00 |
| 696.00 | 849.00 | 826.33 | 1.34 | 0.00 |
| 1526.00 | 1601.00 | 1583.67 | -1.03 | 0.46 |
| 184.00 | 162.00 | 185.33 | 1.04 | 0.67 |
| 1974.00 | 2268.00 | 2159.00 | 1.15 | 0.00 |
| 290.00 | 351.00 | 330.00 | 1.04 | 0.53 |
| 852.00 | 882.00 | 880.33 | -1.28 | 0.00 |
| 857.00 | 847.00 | 874.67 | -1.32 | 0.00 |
| 32.00 | 30.00 | 31.67 | -1.07 | 0.73 |
| 3588.00 | 3675.00 | 3706.67 | -1.20 | 0.00 |
| 10.00 | 12.00 | 11.67 | -1.07 | 0.91 |
| 211.00 | 180.00 | 214.33 | 1.18 | 0.05 |
| 44.00 | 46.00 | 46.67 | 1.04 | 0.82 |
| 4.00 | 2.00 | 2.33 | 1.67 | 0.56 |
| 38.00 | 51.00 | 44.67 | -1.12 | 0.42 |
| 766.00 | 863.00 | 854.00 | -1.02 | 0.65 |
| 1673.00 | 1776.00 | 1679.67 | -1.07 | 0.21 |
| 3773.00 | 3429.00 | 3629.00 | -1.43 | 0.00 |
| 10.00 | 3.00 | 7.00 | 1.29 | 0.52 |
| 56.00 | 46.00 | 50.00 | -1.01 | 0.96 |
| 126.00 | 143.00 | 128.33 | -1.20 | 0.03 |
| 245.00 | 249.00 | 253.33 | -1.22 | 0.01 |
| 1271.00 | 1418.00 | 1395.33 | -1.05 | 0.24 |
| 4.00 | 4.00 | 4.33 | 1.07 | 1.00 |
| 51.00 | 50.00 | 55.67 | 1.20 | 0.17 |
| 261.00 | 295.00 | 291.00 | -1.06 | 0.30 |
| 1143.00 | 1218.00 | 1201.33 | 1.43 | 0.00 |

| | | | | |
|---|---|---|---|---|
| 25.00 | 26.00 | 26.33 | 1.03 | 0.94 |
| 197.00 | 207.00 | 221.00 | 1.24 | 0.01 |
| 24481.00 | 25363.00 | 25594.67 | -1.02 | 0.50 |
| 545.00 | 594.00 | 579.67 | 1.28 | 0.00 |
| 3884.00 | 3708.00 | 3932.00 | -1.05 | 0.19 |
| 1868.00 | 2046.00 | 1920.67 | 1.03 | 0.61 |
| 10.00 | 9.00 | 11.00 | 1.79 | 0.05 |
| 681.00 | 803.00 | 790.00 | 1.06 | 0.27 |
| 544.00 | 498.00 | 543.67 | -1.20 | 0.00 |
| 70.00 | 82.00 | 72.00 | 1.56 | 0.00 |
| 1924.00 | 2069.00 | 2028.33 | 1.02 | 0.54 |
| 241.00 | 313.00 | 287.00 | -1.05 | 0.44 |
| 0.00 | 0.00 | 0.00 | -11.75 | 0.13 |
| 545.00 | 521.00 | 555.67 | -1.21 | 0.00 |
| 132.00 | 147.00 | 137.33 | -1.13 | 0.12 |
| 305.00 | 342.00 | 317.33 | -1.19 | 0.01 |
| 23.00 | 6.00 | 17.33 | -1.14 | 0.67 |
| 43.00 | 43.00 | 46.67 | 1.04 | 0.82 |
| 246.00 | 263.00 | 264.67 | -1.28 | 0.00 |
| 4.00 | 7.00 | 7.00 | -1.20 | 0.67 |
| 5.00 | 6.00 | 4.67 | -1.08 | 1.00 |
| 1172.00 | 1196.00 | 1242.67 | 1.18 | 0.00 |
| 64.00 | 54.00 | 66.67 | 1.03 | 0.82 |
| 9.00 | 5.00 | 8.00 | 1.80 | 0.11 |
| 43.00 | 49.00 | 54.67 | 1.04 | 0.80 |
| 552.00 | 702.00 | 628.33 | -1.17 | 0.01 |
| 891.00 | 1036.00 | 979.00 | -1.25 | 0.00 |
| 829.00 | 855.00 | 843.67 | -1.03 | 0.60 |
| 41990.00 | 46924.00 | 45649.67 | 1.09 | 0.03 |
| 14768.00 | 15160.00 | 15573.33 | -1.46 | 0.00 |
| 11599.00 | 11641.00 | 12009.67 | -1.04 | 0.38 |
| 7.00 | 12.00 | 8.33 | 1.23 | 0.56 |
| 1690.00 | 1068.00 | 1348.00 | -1.55 | 0.00 |
| 639.00 | 690.00 | 643.33 | -1.13 | 0.06 |
| 125.00 | 105.00 | 117.67 | 1.26 | 0.05 |
| 2284.00 | 2197.00 | 2299.67 | -1.31 | 0.00 |
| 159.00 | 188.00 | 175.67 | -1.14 | 0.08 |
| 34.00 | 52.00 | 41.33 | -1.08 | 0.67 |
| 214.00 | 228.00 | 232.33 | -1.18 | 0.01 |
| 28.00 | 36.00 | 36.67 | 1.93 | 0.00 |
| 521.00 | 554.00 | 540.00 | -1.40 | 0.00 |
| 3721.00 | 3658.00 | 3811.00 | -1.19 | 0.00 |
| 13.00 | 8.00 | 10.00 | -1.38 | 0.26 |
| 13.00 | 4.00 | 7.00 | 1.71 | 0.20 |
| 70.00 | 65.00 | 68.67 | -1.03 | 0.79 |
| 197.00 | 229.00 | 203.67 | 1.02 | 0.81 |
| 37.00 | 28.00 | 32.00 | -2.24 | 0.00 |
| 40.00 | 49.00 | 45.67 | -1.12 | 0.38 |

| | | | | |
|---|---|---|---|---|
| 103.00 | 98.00 | 111.00 | 1.05 | 0.63 |
| 1002.00 | 1042.00 | 1065.00 | -1.19 | 0.00 |
| 3435.00 | 3481.00 | 3565.33 | -1.22 | 0.00 |
| 541.00 | 509.00 | 528.67 | 1.23 | 0.00 |
| 18738.00 | 17575.00 | 17990.67 | -1.76 | 0.00 |
| 37.00 | 39.00 | 36.67 | -1.11 | 0.46 |
| 403.00 | 531.00 | 468.67 | -1.04 | 0.50 |
| 1375.00 | 1447.00 | 1446.67 | 1.13 | 0.01 |
| 316.00 | 419.00 | 362.33 | 1.14 | 0.08 |
| 194.00 | 179.00 | 202.67 | 1.02 | 0.81 |
| 1044.00 | 1085.00 | 1041.67 | -1.22 | 0.00 |
| 21.00 | 16.00 | 20.00 | -2.96 | 0.00 |
| 12.00 | 7.00 | 9.00 | 1.47 | 0.24 |
| 607.00 | 729.00 | 649.67 | -1.10 | 0.10 |
| 85.00 | 85.00 | 93.00 | -1.16 | 0.10 |
| 2857.00 | 3011.00 | 3010.00 | 1.07 | 0.05 |
| 121.00 | 121.00 | 133.00 | -1.06 | 0.49 |
| 293.00 | 300.00 | 303.00 | -1.11 | 0.06 |
| 942.00 | 843.00 | 912.33 | -1.13 | 0.01 |
| 609.00 | 580.00 | 596.00 | 1.02 | 0.73 |
| 1786.00 | 1691.00 | 1810.33 | -1.65 | 0.00 |
| 51.00 | 35.00 | 40.00 | -1.13 | 0.45 |
| 1546.00 | 1451.00 | 1521.67 | -1.05 | 0.23 |
| 413.00 | 420.00 | 435.33 | -1.09 | 0.12 |
| 113.00 | 116.00 | 121.00 | -1.25 | 0.01 |
| 430.00 | 527.00 | 497.33 | 1.04 | 0.54 |
| 965.00 | 898.00 | 985.33 | 1.02 | 0.72 |
| 144.00 | 121.00 | 137.33 | -1.09 | 0.29 |
| 544.00 | 515.00 | 500.33 | -1.49 | 0.00 |
| 2030.00 | 2019.00 | 2066.67 | -1.18 | 0.00 |
| 357.00 | 328.00 | 338.00 | -1.11 | 0.10 |
| 223.00 | 197.00 | 206.33 | 1.09 | 0.25 |
| 468.00 | 475.00 | 504.33 | -1.08 | 0.16 |
| 687.00 | 718.00 | 767.67 | -1.02 | 0.77 |
| 680.00 | 716.00 | 716.00 | 1.13 | 0.01 |
| 10.00 | 6.00 | 7.33 | -1.06 | 1.00 |
| 560.00 | 591.00 | 582.33 | 1.02 | 0.68 |
| 1665.00 | 1666.00 | 1729.00 | 1.14 | 0.00 |
| 1347.00 | 1360.00 | 1418.00 | 1.18 | 0.00 |
| 2280.00 | 2508.00 | 2402.00 | 1.31 | 0.00 |
| 440.00 | 488.00 | 477.00 | 1.04 | 0.41 |
| 395.00 | 500.00 | 452.33 | 1.06 | 0.30 |
| 0.00 | 1.00 | 0.67 | -3.57 | 0.11 |
| 5513.00 | 5767.00 | 5694.33 | -1.28 | 0.00 |
| 17968.00 | 20137.00 | 18896.00 | 1.00 | 0.97 |
| 5463.00 | 5743.00 | 5803.33 | -1.10 | 0.01 |
| 1809.00 | 1797.00 | 1865.33 | 1.05 | 0.31 |
| 1724.00 | 1950.00 | 1902.67 | 1.08 | 0.05 |

| | | | | |
|---|---|---|---|---|
| 189.00 | 198.00 | 197.67 | 1.17 | 0.05 |
| 7991.00 | 8771.00 | 8543.67 | 1.07 | 0.05 |
| 75.00 | 89.00 | 86.00 | 1.10 | 0.42 |
| 663.00 | 724.00 | 724.33 | 1.08 | 0.12 |
| 5.00 | 14.00 | 13.00 | -1.07 | 0.84 |
| 80.00 | 105.00 | 99.33 | -1.43 | 0.00 |
| 3426.00 | 3453.00 | 3570.67 | 1.01 | 0.86 |
| 2.00 | 1.00 | 2.00 | -2.12 | 0.17 |
| 4177.00 | 3983.00 | 4045.00 | -1.42 | 0.00 |
| 5501.00 | 5013.00 | 5268.67 | 1.29 | 0.00 |
| 651.00 | 583.00 | 642.67 | -1.25 | 0.00 |
| 2371.00 | 2093.00 | 2156.33 | -1.59 | 0.00 |
| 110.00 | 90.00 | 103.00 | -1.47 | 0.00 |
| 1303.00 | 1311.00 | 1302.67 | -1.62 | 0.00 |
| 731.00 | 641.00 | 714.67 | -1.46 | 0.00 |
| 142.00 | 125.00 | 132.67 | -1.06 | 0.56 |
| 2193.00 | 2088.00 | 2158.67 | -1.61 | 0.00 |
| 428.00 | 539.00 | 470.00 | 1.25 | 0.00 |
| 1564.00 | 1932.00 | 1808.00 | 1.34 | 0.00 |
| 5.00 | 3.00 | 5.00 | -1.34 | 0.51 |
| 156.00 | 173.00 | 172.67 | -1.46 | 0.00 |
| 195.00 | 220.00 | 226.67 | 1.11 | 0.19 |
| 8.00 | 8.00 | 7.67 | -1.19 | 0.68 |
| 17.00 | 15.00 | 16.67 | -1.44 | 0.08 |
| 711.00 | 734.00 | 739.00 | -1.07 | 0.11 |
| 413.00 | 460.00 | 471.33 | 1.11 | 0.11 |
| 891.00 | 906.00 | 908.67 | -1.03 | 0.52 |
| 440.00 | 444.00 | 449.33 | -1.06 | 0.25 |
| 1878.00 | 2048.00 | 2034.67 | -1.01 | 0.74 |
| 1169.00 | 1358.00 | 1328.00 | -1.02 | 0.70 |
| 1183.00 | 1531.00 | 1393.33 | 1.09 | 0.08 |
| 563.00 | 571.00 | 555.67 | 1.03 | 0.63 |
| 152.00 | 140.00 | 152.33 | 1.02 | 0.87 |
| 304.00 | 355.00 | 344.00 | 1.34 | 0.00 |
| 2.00 | 6.00 | 4.33 | 1.79 | 0.27 |
| 165.00 | 172.00 | 163.33 | -1.13 | 0.13 |
| 17.00 | 40.00 | 32.00 | 1.49 | 0.04 |
| 63.00 | 56.00 | 61.00 | 1.07 | 0.55 |
| 13.00 | 19.00 | 14.33 | 1.21 | 0.45 |
| 13.00 | 10.00 | 11.00 | 1.05 | 0.90 |
| 616.00 | 625.00 | 656.67 | 1.26 | 0.00 |
| 2.00 | 3.00 | 2.67 | 1.54 | 0.59 |
| 5053.00 | 6003.00 | 5651.33 | -1.16 | 0.00 |
| 434.00 | 419.00 | 453.33 | -1.10 | 0.08 |
| 560.00 | 506.00 | 534.67 | -1.03 | 0.68 |
| 1.00 | 1.00 | 1.33 | -2.87 | 0.08 |
| 1144.00 | 1163.00 | 1206.33 | -1.03 | 0.53 |
| 181.00 | 173.00 | 170.67 | -1.48 | 0.00 |

| | | | | |
|---|---|---|---|---|
| 10.00 | 10.00 | 9.67 | -1.01 | 1.00 |
| 688.00 | 637.00 | 678.67 | 1.23 | 0.00 |
| 8860.00 | 10764.00 | 9988.00 | 1.03 | 0.48 |
| 831.00 | 898.00 | 925.67 | 1.01 | 0.81 |
| 2603.00 | 3035.00 | 2971.33 | 1.40 | 0.00 |
| 1088.00 | 1117.00 | 1144.67 | -1.29 | 0.00 |
| 199.00 | 201.00 | 194.33 | -1.55 | 0.00 |
| 340.00 | 346.00 | 375.67 | 1.00 | 0.99 |
| 2811.00 | 2589.00 | 2725.00 | -1.17 | 0.00 |
| 27.00 | 29.00 | 31.33 | -1.03 | 0.89 |
| 408.00 | 512.00 | 462.00 | -1.12 | 0.05 |
| 715.00 | 666.00 | 719.00 | 1.02 | 0.71 |
| 1472.00 | 1757.00 | 1661.00 | 1.09 | 0.03 |
| 191.00 | 244.00 | 220.33 | -1.27 | 0.00 |
| 1431.00 | 1818.00 | 1694.00 | 1.14 | 0.01 |
| 482.00 | 486.00 | 508.67 | 1.18 | 0.00 |
| 510.00 | 643.00 | 591.67 | -1.07 | 0.24 |
| 361.00 | 443.00 | 419.33 | 1.12 | 0.05 |
| 8.00 | 0.00 | 4.33 | -1.47 | 0.40 |
| 3916.00 | 3743.00 | 3955.00 | -1.13 | 0.00 |
| 2253.00 | 2310.00 | 2343.00 | 1.12 | 0.01 |
| 2984.00 | 3061.00 | 2952.00 | -1.29 | 0.00 |
| 1559.00 | 1661.00 | 1650.33 | -1.25 | 0.00 |
| 2772.00 | 2903.00 | 2920.33 | 1.06 | 0.08 |
| 962.00 | 1016.00 | 997.00 | -1.29 | 0.00 |
| 3079.00 | 3415.00 | 3359.33 | -1.08 | 0.02 |
| 519.00 | 632.00 | 579.00 | 1.13 | 0.02 |
| 1041.00 | 1172.00 | 1144.67 | 1.21 | 0.00 |
| 454.00 | 469.00 | 471.33 | 1.11 | 0.04 |
| 1438.00 | 1465.00 | 1459.67 | 1.13 | 0.00 |
| 3025.00 | 3098.00 | 3236.00 | 1.17 | 0.00 |
| 479.00 | 565.00 | 543.00 | 1.23 | 0.00 |
| 664.00 | 715.00 | 735.33 | 1.04 | 0.47 |
| 2198.00 | 2231.00 | 2353.00 | 1.08 | 0.05 |
| 1269.00 | 1224.00 | 1239.33 | -1.10 | 0.03 |
| 1849.00 | 1914.00 | 1877.33 | 1.04 | 0.40 |
| 196.00 | 150.00 | 181.67 | 1.43 | 0.00 |
| 1219.00 | 1263.00 | 1310.67 | 1.26 | 0.00 |
| 612.00 | 590.00 | 651.00 | 1.27 | 0.00 |
| 1677.00 | 1730.00 | 1783.67 | -1.16 | 0.00 |
| 905.00 | 1018.00 | 983.67 | -1.11 | 0.01 |
| 265.00 | 323.00 | 295.00 | -1.03 | 0.68 |
| 808.00 | 896.00 | 887.67 | -1.07 | 0.13 |
| 829.00 | 951.00 | 929.67 | -1.07 | 0.13 |
| 915.00 | 972.00 | 998.00 | -1.06 | 0.24 |
| 1609.00 | 1769.00 | 1786.67 | 1.02 | 0.67 |
| 91.00 | 113.00 | 102.33 | -1.32 | 0.00 |
| 542.00 | 603.00 | 578.33 | 1.22 | 0.00 |

| | | | | |
|---|---|---|---|---|
| 2114.00 | 1952.00 | 1963.67 | -1.07 | 0.21 |
| 1968.00 | 2141.00 | 2101.00 | 1.05 | 0.21 |
| 1096.00 | 1268.00 | 1254.00 | 1.03 | 0.53 |
| 1811.00 | 1921.00 | 1992.00 | 1.13 | 0.00 |
| 808.00 | 813.00 | 857.33 | 1.21 | 0.00 |
| 5785.00 | 5069.00 | 5290.00 | -1.23 | 0.00 |
| 3175.00 | 3731.00 | 3556.67 | 1.05 | 0.24 |
| 1490.00 | 1523.00 | 1639.33 | 1.20 | 0.00 |
| 781.00 | 758.00 | 784.00 | 1.23 | 0.00 |
| 1967.00 | 1982.00 | 2077.00 | 1.09 | 0.03 |
| 488.00 | 559.00 | 539.33 | 1.13 | 0.02 |
| 1091.00 | 1091.00 | 1154.00 | 1.19 | 0.00 |
| 1381.00 | 1491.00 | 1549.00 | 1.05 | 0.31 |
| 296.00 | 333.00 | 340.33 | 1.23 | 0.00 |
| 222.00 | 219.00 | 234.67 | 1.20 | 0.01 |
| 30.00 | 28.00 | 27.00 | -1.26 | 0.18 |
| 217.00 | 270.00 | 251.33 | -1.10 | 0.14 |
| 848.00 | 923.00 | 955.00 | 1.21 | 0.00 |
| 862.00 | 980.00 | 1015.67 | 1.11 | 0.07 |
| 65.00 | 44.00 | 52.33 | 1.18 | 0.25 |
| 2609.00 | 2641.00 | 2597.67 | -1.03 | 0.45 |
| 650.00 | 654.00 | 649.33 | 1.04 | 0.46 |
| 1051.00 | 1061.00 | 1088.67 | 1.13 | 0.01 |
| 880.00 | 863.00 | 869.00 | -1.01 | 0.92 |
| 1455.00 | 1656.00 | 1583.00 | 1.03 | 0.50 |
| 515.00 | 605.00 | 562.00 | 1.14 | 0.02 |
| 810.00 | 1085.00 | 957.67 | 1.09 | 0.20 |
| 294.00 | 259.00 | 305.67 | 1.05 | 0.52 |
| 1081.00 | 1107.00 | 1128.00 | 1.02 | 0.68 |
| 312.00 | 353.00 | 363.00 | 1.29 | 0.00 |
| 2246.00 | 2171.00 | 2258.00 | -1.14 | 0.00 |
| 978.00 | 779.00 | 863.67 | -1.29 | 0.00 |
| 1273.00 | 1092.00 | 1209.00 | -1.21 | 0.00 |
| 1.00 | 1.00 | 1.33 | -3.79 | 0.01 |
| 57.00 | 50.00 | 58.33 | -1.86 | 0.00 |
| 2506.00 | 2483.00 | 2535.67 | -1.11 | 0.00 |
| 495.00 | 520.00 | 508.33 | -1.01 | 0.88 |
| 41.00 | 37.00 | 32.33 | 1.09 | 0.68 |
| 1212.00 | 1333.00 | 1345.67 | 1.22 | 0.00 |
| 1181.00 | 1273.00 | 1253.67 | -1.14 | 0.00 |
| 1124.00 | 1125.00 | 1156.00 | -1.44 | 0.00 |
| 2260.00 | 2148.00 | 2294.00 | 1.04 | 0.40 |
| 668.00 | 747.00 | 730.00 | 1.04 | 0.44 |
| 442.00 | 499.00 | 501.67 | 1.12 | 0.04 |
| 400.00 | 475.00 | 480.00 | 1.10 | 0.21 |
| 652.00 | 621.00 | 646.67 | 1.23 | 0.00 |
| 15.00 | 12.00 | 16.33 | -1.43 | 0.12 |
| 673.00 | 705.00 | 741.67 | 1.28 | 0.00 |

| | | | | |
|---|---|---|---|---|
| 2130.00 | 2165.00 | 2219.67 | -1.09 | 0.06 |
| 363.00 | 332.00 | 367.00 | -1.35 | 0.00 |
| 839.00 | 862.00 | 909.00 | 1.03 | 0.56 |
| 8.00 | 9.00 | 12.33 | 1.51 | 0.19 |
| 700.00 | 856.00 | 781.33 | -1.18 | 0.00 |
| 1995.00 | 1977.00 | 2071.67 | 1.02 | 0.58 |
| 39.00 | 39.00 | 39.33 | 1.35 | 0.05 |
| 1082.00 | 1208.00 | 1197.00 | 1.08 | 0.05 |
| 99.00 | 110.00 | 111.33 | -1.39 | 0.00 |
| 133.00 | 153.00 | 155.33 | -1.07 | 0.37 |
| 2849.00 | 3475.00 | 3320.00 | 1.17 | 0.00 |
| 754.00 | 861.00 | 815.00 | 1.20 | 0.00 |
| 186.00 | 193.00 | 193.33 | -1.25 | 0.00 |
| 297.00 | 340.00 | 332.67 | 1.01 | 0.91 |
| 1646.00 | 1631.00 | 1702.67 | -1.32 | 0.00 |
| 1653.00 | 1550.00 | 1694.00 | 1.06 | 0.29 |
| 29.00 | 32.00 | 32.33 | -1.35 | 0.05 |
| 1044.00 | 1060.00 | 1078.00 | -1.01 | 0.83 |
| 565.00 | 590.00 | 568.00 | -1.05 | 0.34 |
| 68.00 | 50.00 | 59.67 | -1.07 | 0.61 |
| 321.00 | 283.00 | 299.33 | -1.52 | 0.00 |
| 2773.00 | 2817.00 | 3007.67 | -1.05 | 0.22 |
| 376.00 | 407.00 | 405.67 | -1.31 | 0.00 |
| 10.00 | 15.00 | 14.00 | 1.04 | 0.92 |
| 415.00 | 389.00 | 383.67 | 1.02 | 0.73 |
| 697.00 | 771.00 | 740.00 | 1.17 | 0.00 |
| 269.00 | 281.00 | 291.00 | 1.34 | 0.00 |
| 693.00 | 824.00 | 813.00 | 1.18 | 0.00 |
| 1343.00 | 1192.00 | 1367.00 | 1.14 | 0.02 |
| 7218.00 | 7539.00 | 7443.67 | -1.13 | 0.00 |
| 141.00 | 189.00 | 165.67 | -1.08 | 0.44 |
| 202.00 | 211.00 | 214.67 | -1.23 | 0.00 |
| 12.00 | 14.00 | 11.33 | 1.24 | 0.45 |
| 3109.00 | 3360.00 | 3164.33 | -1.34 | 0.00 |
| 30.00 | 28.00 | 30.67 | 1.21 | 0.30 |
| 1142.00 | 1270.00 | 1220.33 | -1.03 | 0.42 |
| 9805.00 | 11042.00 | 11112.67 | 1.21 | 0.00 |
| 42.00 | 37.00 | 38.00 | 1.46 | 0.02 |
| 0.00 | 0.00 | 0.00 | -11.74 | 0.13 |
| 10831.00 | 11240.00 | 11468.33 | -1.13 | 0.00 |
| 782.00 | 745.00 | 770.33 | 1.13 | 0.03 |
| 928.00 | 1056.00 | 1045.00 | 1.24 | 0.00 |
| 1048.00 | 1147.00 | 1170.33 | 1.13 | 0.01 |
| 109.00 | 119.00 | 122.67 | -1.42 | 0.00 |
| 2620.00 | 2626.00 | 2713.33 | 1.04 | 0.24 |
| 1084.00 | 1254.00 | 1181.67 | 1.11 | 0.01 |
| 754.00 | 971.00 | 896.33 | 1.29 | 0.00 |
| 1013.00 | 1225.00 | 1192.67 | 1.24 | 0.00 |

| | | | | |
|---|---|---|---|---|
| 774.00 | 791.00 | 829.67 | -1.03 | 0.60 |
| 501.00 | 555.00 | 566.67 | 1.14 | 0.02 |
| 755.00 | 899.00 | 870.00 | 1.11 | 0.03 |
| 55.00 | 50.00 | 54.33 | 1.12 | 0.39 |
| 562.00 | 607.00 | 595.00 | -1.07 | 0.16 |
| 909.00 | 1019.00 | 940.00 | 1.18 | 0.01 |
| 1117.00 | 1222.00 | 1196.33 | 1.08 | 0.05 |
| 634.00 | 699.00 | 682.33 | 1.04 | 0.35 |
| 265.00 | 305.00 | 298.33 | 1.20 | 0.00 |
| 462.00 | 506.00 | 472.67 | 1.14 | 0.03 |
| 2037.00 | 2016.00 | 2046.00 | -1.10 | 0.02 |
| 3600.00 | 3825.00 | 3826.33 | 1.17 | 0.00 |
| 134.00 | 153.00 | 144.33 | -1.05 | 0.58 |
| 11.00 | 31.00 | 24.33 | -1.03 | 0.89 |
| 455.00 | 503.00 | 483.67 | 1.10 | 0.14 |
| 65.00 | 45.00 | 48.67 | -1.71 | 0.00 |
| 578.00 | 624.00 | 612.67 | -1.03 | 0.50 |
| 25.00 | 17.00 | 21.67 | -1.15 | 0.50 |
| 1031.00 | 943.00 | 1001.00 | -1.06 | 0.23 |
| 786.00 | 898.00 | 890.00 | -1.07 | 0.17 |
| 2124.00 | 2100.00 | 2251.67 | 1.20 | 0.00 |
| 1985.00 | 2315.00 | 2167.33 | 1.19 | 0.00 |
| 4528.00 | 4985.00 | 4888.67 | 1.20 | 0.00 |
| 101.00 | 76.00 | 86.67 | 1.00 | 1.00 |
| 1449.00 | 1574.00 | 1482.00 | 1.02 | 0.67 |
| 399.00 | 398.00 | 386.67 | -1.30 | 0.00 |
| 1919.00 | 1972.00 | 2042.33 | -1.10 | 0.02 |
| 283.00 | 311.00 | 304.33 | -1.15 | 0.01 |
| 302.00 | 280.00 | 292.33 | -1.11 | 0.12 |
| 4701.00 | 5237.00 | 5175.67 | 1.15 | 0.00 |
| 9402.00 | 9538.00 | 9818.67 | -1.18 | 0.00 |
| 2624.00 | 2593.00 | 2637.67 | 1.02 | 0.70 |
| 1415.00 | 1488.00 | 1525.67 | 1.01 | 0.87 |
| 0.00 | 0.00 | 1.33 | -2.17 | 0.28 |
| 4.00 | 1.00 | 3.00 | -1.34 | 0.67 |
| 685.00 | 825.00 | 762.67 | 1.10 | 0.06 |
| 358.00 | 417.00 | 424.33 | -1.07 | 0.31 |
| 1143.00 | 1117.00 | 1165.33 | -1.23 | 0.00 |
| 74.00 | 44.00 | 57.33 | -1.07 | 0.67 |
| 18.00 | 22.00 | 21.33 | 1.11 | 0.66 |
| 19.00 | 11.00 | 18.33 | -1.18 | 0.40 |
| 950.00 | 974.00 | 1026.00 | 1.19 | 0.00 |
| 14.00 | 13.00 | 11.33 | 2.74 | 0.00 |
| 700.00 | 685.00 | 707.67 | -1.03 | 0.55 |
| 830.00 | 749.00 | 814.67 | 1.44 | 0.00 |
| 128.00 | 142.00 | 139.00 | -1.04 | 0.65 |
| 597.00 | 600.00 | 619.00 | -1.21 | 0.00 |
| 255.00 | 247.00 | 269.33 | 1.04 | 0.59 |

| | | | | |
|---|---|---|---|---|
| 740.00 | 783.00 | 800.00 | 1.15 | 0.00 |
| 1012.00 | 1068.00 | 1073.33 | 1.11 | 0.02 |
| 319.00 | 401.00 | 365.33 | 1.04 | 0.56 |
| 3299.00 | 3150.00 | 3257.67 | 1.06 | 0.18 |
| 3175.00 | 3715.00 | 3574.00 | 1.08 | 0.06 |
| 1008.00 | 1066.00 | 1054.33 | 1.15 | 0.00 |
| 3259.00 | 3427.00 | 3495.67 | 1.08 | 0.03 |
| 492.00 | 489.00 | 501.33 | -1.12 | 0.03 |
| 1636.00 | 1833.00 | 1800.67 | 1.18 | 0.00 |
| 8.00 | 15.00 | 11.00 | -1.26 | 0.43 |
| 1909.00 | 2007.00 | 2035.67 | -1.14 | 0.00 |
| 321.00 | 326.00 | 313.67 | -1.10 | 0.22 |
| 125.00 | 138.00 | 134.33 | -1.05 | 0.53 |
| 621.00 | 685.00 | 679.67 | 1.20 | 0.00 |
| 2034.00 | 2141.00 | 2169.33 | -1.01 | 0.72 |
| 542.00 | 648.00 | 621.00 | 1.03 | 0.60 |
| 700.00 | 789.00 | 766.00 | 1.00 | 1.00 |
| 1761.00 | 1938.00 | 1907.67 | 1.26 | 0.00 |
| 1073.00 | 1120.00 | 1142.33 | 1.05 | 0.28 |
| 877.00 | 985.00 | 965.00 | 1.12 | 0.01 |
| 566.00 | 656.00 | 610.67 | -1.19 | 0.00 |
| 1597.00 | 1805.00 | 1731.33 | 1.20 | 0.00 |
| 234.00 | 283.00 | 275.33 | -1.21 | 0.00 |
| 90.00 | 125.00 | 113.67 | -1.03 | 0.77 |
| 1368.00 | 1530.00 | 1458.67 | 1.28 | 0.00 |
| 1683.00 | 1765.00 | 1805.00 | 1.13 | 0.00 |
| 2193.00 | 2398.00 | 2343.33 | 1.02 | 0.51 |
| 254.00 | 201.00 | 236.67 | -1.45 | 0.00 |
| 728.00 | 814.00 | 755.67 | 1.21 | 0.00 |
| 1727.00 | 1764.00 | 1788.67 | 1.03 | 0.47 |
| 1658.00 | 1822.00 | 1757.67 | 1.05 | 0.22 |
| 1148.00 | 1242.00 | 1252.67 | 1.13 | 0.00 |
| 836.00 | 930.00 | 935.00 | 1.15 | 0.00 |
| 2791.00 | 2789.00 | 2868.33 | -1.02 | 0.50 |
| 1372.00 | 1374.00 | 1438.00 | -1.02 | 0.59 |
| 2309.00 | 2483.00 | 2362.33 | 1.08 | 0.07 |
| 3472.00 | 3512.00 | 3488.33 | -1.16 | 0.00 |
| 4655.00 | 4726.00 | 4817.00 | 1.19 | 0.00 |
| 597.00 | 581.00 | 583.00 | 1.09 | 0.13 |
| 1012.00 | 1203.00 | 1176.00 | 1.06 | 0.21 |
| 978.00 | 1039.00 | 1063.33 | 1.06 | 0.21 |
| 547.00 | 563.00 | 583.67 | 1.04 | 0.45 |
| 650.00 | 637.00 | 674.33 | -1.03 | 0.49 |
| 173.00 | 176.00 | 173.33 | 1.18 | 0.04 |
| 146.00 | 185.00 | 174.67 | 1.33 | 0.00 |
| 756.00 | 631.00 | 759.33 | 1.02 | 0.80 |
| 1.00 | 4.00 | 2.00 | -2.28 | 0.12 |
| 2112.00 | 2217.00 | 2203.33 | 1.23 | 0.00 |

| | | | | |
|---|---|---|---|---|
| 21.00 | 26.00 | 21.33 | -1.04 | 0.93 |
| 350.00 | 421.00 | 421.00 | 1.01 | 0.93 |
| 1326.00 | 1356.00 | 1436.67 | 1.10 | 0.04 |
| 172.00 | 135.00 | 149.00 | -1.61 | 0.00 |
| 1413.00 | 1412.00 | 1466.33 | -1.28 | 0.00 |
| 84.00 | 80.00 | 90.33 | 2.32 | 0.00 |
| 272.00 | 324.00 | 303.33 | 1.06 | 0.35 |
| 14.00 | 20.00 | 16.67 | -1.34 | 0.16 |
| 8.00 | 10.00 | 8.67 | -1.36 | 0.32 |
| 63.00 | 61.00 | 66.00 | 1.29 | 0.03 |
| 507.00 | 613.00 | 578.00 | 1.02 | 0.78 |
| 715.00 | 686.00 | 685.67 | -1.19 | 0.00 |
| 1604.00 | 1625.00 | 1634.67 | -1.09 | 0.02 |
| 154.00 | 157.00 | 159.33 | -1.19 | 0.02 |
| 10.00 | 13.00 | 13.33 | -1.29 | 0.33 |
| 519.00 | 591.00 | 570.67 | -1.09 | 0.08 |
| 6052.00 | 6776.00 | 6823.67 | 1.24 | 0.00 |
| 1639.00 | 1801.00 | 1835.67 | 1.22 | 0.00 |
| 2839.00 | 3392.00 | 3235.67 | 1.17 | 0.00 |
| 4464.00 | 4506.00 | 4661.67 | 1.09 | 0.03 |
| 5179.00 | 5540.00 | 5635.67 | 1.20 | 0.00 |
| 4314.00 | 4959.00 | 4977.33 | 1.16 | 0.00 |
| 793.00 | 835.00 | 867.00 | -1.04 | 0.36 |
| 2556.00 | 2696.00 | 2745.33 | 1.30 | 0.00 |
| 133.00 | 110.00 | 127.67 | -1.01 | 0.90 |
| 851.00 | 846.00 | 868.33 | 1.03 | 0.50 |
| 2664.00 | 2812.00 | 2707.00 | 1.00 | 0.95 |
| 3159.00 | 3020.00 | 3103.33 | -1.02 | 0.69 |
| 6.00 | 4.00 | 3.67 | -1.54 | 0.36 |
| 100.00 | 92.00 | 93.00 | 1.08 | 0.56 |
| 126.00 | 123.00 | 131.00 | -1.32 | 0.00 |
| 253.00 | 243.00 | 250.33 | -1.40 | 0.00 |
| 140.00 | 147.00 | 141.33 | -1.47 | 0.00 |
| 285.00 | 363.00 | 330.00 | 1.34 | 0.00 |
| 641.00 | 620.00 | 652.00 | 1.34 | 0.00 |
| 391.00 | 479.00 | 455.33 | 1.30 | 0.00 |
| 2253.00 | 2822.00 | 2487.67 | -1.02 | 0.72 |
| 548.00 | 597.00 | 558.33 | -1.06 | 0.29 |
| 499.00 | 514.00 | 513.67 | -1.04 | 0.43 |
| 1233.00 | 1242.00 | 1247.67 | -1.11 | 0.01 |
| 691.00 | 694.00 | 711.00 | -1.05 | 0.27 |
| 1129.00 | 1270.00 | 1188.33 | 1.04 | 0.38 |
| 1479.00 | 1611.00 | 1609.33 | -1.00 | 0.93 |
| 2437.00 | 2833.00 | 2691.67 | 1.06 | 0.12 |
| 2728.00 | 3093.00 | 2887.00 | -1.24 | 0.00 |
| 394.00 | 443.00 | 416.00 | -1.02 | 0.72 |
| 458.00 | 408.00 | 381.00 | 1.02 | 0.88 |
| 32.00 | 22.00 | 25.00 | -1.32 | 0.11 |

| | | | | |
|---|---|---|---|---|
| 150.00 | 216.00 | 187.67 | 1.51 | 0.00 |
| 6537.00 | 6333.00 | 6323.00 | 1.10 | 0.03 |
| 810.00 | 827.00 | 839.67 | 1.08 | 0.07 |
| 263.00 | 282.00 | 296.33 | 1.75 | 0.00 |
| 768.00 | 759.00 | 762.33 | 1.16 | 0.00 |
| 1.00 | 2.00 | 1.00 | -4.61 | 0.01 |
| 217.00 | 199.00 | 208.67 | -1.32 | 0.00 |
| 1544.00 | 1822.00 | 1682.00 | 1.14 | 0.00 |
| 91.00 | 114.00 | 100.67 | 1.27 | 0.01 |
| 11214.00 | 12078.00 | 11909.67 | -1.03 | 0.43 |
| 201.00 | 200.00 | 205.33 | -1.12 | 0.09 |
| 319.00 | 345.00 | 347.00 | -1.38 | 0.00 |
| 1915.00 | 1900.00 | 1936.33 | 1.10 | 0.02 |
| 14369.00 | 15469.00 | 14655.33 | -1.02 | 0.59 |
| 135.00 | 127.00 | 124.67 | -1.29 | 0.01 |
| 45.00 | 40.00 | 39.33 | 1.03 | 0.86 |
| 33.00 | 28.00 | 32.00 | 1.02 | 0.94 |
| 372.00 | 405.00 | 432.33 | 1.01 | 0.91 |
| 27.00 | 44.00 | 38.67 | -1.28 | 0.08 |
| 1333.00 | 1538.00 | 1443.00 | 1.04 | 0.37 |
| 153.00 | 199.00 | 176.00 | -1.16 | 0.05 |
| 331.00 | 354.00 | 355.00 | -1.30 | 0.00 |
| 103.00 | 113.00 | 101.00 | 1.34 | 0.01 |
| 3405.00 | 3624.00 | 3747.67 | 1.17 | 0.00 |
| 1451.00 | 1550.00 | 1525.33 | -1.12 | 0.00 |
| 947.00 | 1097.00 | 1036.67 | 1.17 | 0.00 |
| 2651.00 | 2707.00 | 2710.33 | 1.05 | 0.24 |
| 623.00 | 693.00 | 697.67 | 1.01 | 0.81 |
| 849.00 | 849.00 | 873.00 | 1.04 | 0.33 |
| 243.00 | 276.00 | 265.00 | -1.04 | 0.56 |
| 15207.00 | 15565.00 | 15796.00 | 1.32 | 0.00 |
| 51.00 | 56.00 | 50.67 | -1.96 | 0.00 |
| 84.00 | 90.00 | 91.00 | 1.10 | 0.37 |
| 2106.00 | 2138.00 | 2229.00 | 1.13 | 0.00 |
| 1.00 | 3.00 | 2.33 | -1.15 | 1.00 |
| 24.00 | 36.00 | 33.67 | -1.26 | 0.11 |
| 48.00 | 51.00 | 54.33 | 1.06 | 0.74 |
| 559.00 | 595.00 | 596.00 | -1.06 | 0.17 |
| 851.00 | 900.00 | 951.67 | 1.10 | 0.07 |
| 1380.00 | 1396.00 | 1415.00 | -1.02 | 0.71 |
| 84.00 | 72.00 | 72.33 | 1.03 | 0.80 |
| 353.00 | 314.00 | 337.00 | -1.11 | 0.08 |
| 6.00 | 3.00 | 4.67 | -1.93 | 0.06 |
| 339.00 | 320.00 | 340.33 | 1.10 | 0.12 |
| 3722.00 | 3795.00 | 3856.67 | 1.11 | 0.00 |
| 1377.00 | 1642.00 | 1563.67 | 1.01 | 0.86 |
| 19.00 | 11.00 | 15.00 | -1.84 | 0.00 |
| 1166.00 | 1266.00 | 1254.67 | 1.03 | 0.42 |

| | | | | |
|---|---|---|---|---|
| 3019.00 | 3562.00 | 3403.00 | 1.04 | 0.30 |
| 1493.00 | 1894.00 | 1763.67 | -1.03 | 0.52 |
| 748.00 | 841.00 | 819.00 | 1.02 | 0.71 |
| 3596.00 | 4040.00 | 3795.00 | 1.11 | 0.01 |
| 412.00 | 416.00 | 444.00 | -1.14 | 0.02 |
| 3452.00 | 4007.00 | 3664.00 | -1.07 | 0.15 |
| 3115.00 | 3451.00 | 3301.67 | 1.05 | 0.15 |
| 25.00 | 11.00 | 16.00 | -1.07 | 0.86 |
| 915.00 | 976.00 | 983.33 | 1.16 | 0.00 |
| 396.00 | 451.00 | 436.67 | 1.13 | 0.02 |
| 647.00 | 721.00 | 706.67 | 1.13 | 0.01 |
| 589.00 | 671.00 | 663.67 | 1.02 | 0.70 |
| 2724.00 | 2738.00 | 2673.00 | 1.18 | 0.00 |
| 943.00 | 850.00 | 889.33 | -1.07 | 0.21 |
| 1094.00 | 1207.00 | 1189.67 | 1.29 | 0.00 |
| 815.00 | 806.00 | 840.00 | 1.04 | 0.39 |
| 1358.00 | 1267.00 | 1367.33 | -1.17 | 0.00 |
| 168.00 | 248.00 | 193.00 | 1.03 | 0.80 |
| 28.00 | 24.00 | 23.33 | 1.41 | 0.09 |
| 226.00 | 221.00 | 229.67 | 1.19 | 0.01 |
| 8.00 | 8.00 | 9.00 | 1.39 | 0.31 |
| 8.00 | 9.00 | 8.67 | 2.10 | 0.04 |
| 74.00 | 56.00 | 71.00 | 1.15 | 0.25 |
| 37.00 | 53.00 | 49.00 | 1.35 | 0.03 |
| 284.00 | 287.00 | 296.00 | 1.17 | 0.01 |
| 118.00 | 112.00 | 117.33 | -1.03 | 0.80 |
| 1317.00 | 1494.00 | 1419.33 | 1.05 | 0.27 |
| 15.00 | 20.00 | 18.00 | -1.07 | 0.85 |
| 9616.00 | 8645.00 | 9096.00 | -1.11 | 0.02 |
| 8286.00 | 7508.00 | 7758.00 | -1.27 | 0.00 |
| 238.00 | 224.00 | 231.33 | -1.00 | 1.00 |
| 5.00 | 13.00 | 9.00 | 1.06 | 0.89 |
| 6.00 | 1.00 | 6.00 | 1.11 | 0.88 |
| 16.00 | 5.00 | 11.33 | -1.28 | 0.39 |
| 56.00 | 49.00 | 52.00 | -1.38 | 0.01 |
| 58.00 | 55.00 | 59.00 | -1.04 | 0.81 |
| 1203.00 | 1333.00 | 1275.00 | -1.10 | 0.04 |
| 1123.00 | 1196.00 | 1193.33 | -1.05 | 0.24 |
| 4333.00 | 4178.00 | 4306.00 | -1.41 | 0.00 |
| 10.00 | 22.00 | 17.33 | -1.19 | 0.42 |
| 621.00 | 681.00 | 720.33 | -1.05 | 0.43 |
| 0.00 | 2.00 | 2.67 | -1.62 | 0.39 |
| 6.00 | 9.00 | 6.00 | 1.04 | 1.00 |
| 373.00 | 322.00 | 389.00 | -1.30 | 0.00 |
| 2.00 | 3.00 | 2.33 | -1.43 | 0.63 |
| 706.00 | 737.00 | 709.00 | -1.05 | 0.34 |
| 202.00 | 229.00 | 216.33 | -1.08 | 0.23 |
| 11.00 | 5.00 | 8.67 | -1.17 | 0.70 |

| | | | | |
|---|---|---|---|---|
| 131.00 | 136.00 | 136.33 | 1.07 | 0.39 |
| 26432.00 | 27764.00 | 28004.67 | -1.27 | 0.00 |
| 7932.00 | 8689.00 | 8593.00 | 1.19 | 0.00 |
| 36.00 | 32.00 | 31.67 | 1.29 | 0.15 |
| 8.00 | 10.00 | 12.00 | -1.60 | 0.04 |
| 521.00 | 510.00 | 495.33 | -1.61 | 0.00 |
| 704.00 | 755.00 | 737.67 | -1.18 | 0.00 |
| 1400.00 | 1568.00 | 1520.33 | -1.03 | 0.40 |
| 1234.00 | 1197.00 | 1234.67 | 1.11 | 0.01 |
| 827.00 | 823.00 | 894.33 | 1.10 | 0.12 |
| 33.00 | 42.00 | 37.00 | -1.08 | 0.64 |
| 13.00 | 18.00 | 20.00 | 3.42 | 0.00 |
| 11.00 | 9.00 | 9.00 | 1.76 | 0.09 |
| 1958.00 | 2121.00 | 2139.67 | -1.16 | 0.00 |
| 89.00 | 105.00 | 103.33 | -1.00 | 1.00 |
| 24.00 | 22.00 | 24.33 | -1.04 | 0.88 |
| 16.00 | 18.00 | 22.00 | 2.30 | 0.00 |
| 44.00 | 38.00 | 42.33 | 1.46 | 0.01 |
| 2500.00 | 3053.00 | 2791.00 | 1.33 | 0.00 |
| 542.00 | 582.00 | 594.67 | 1.11 | 0.04 |
| 3283.00 | 4094.00 | 3918.33 | 1.21 | 0.00 |
| 2139.00 | 2377.00 | 2433.00 | 1.23 | 0.00 |
| 1190.00 | 1491.00 | 1446.67 | 1.12 | 0.04 |
| 1252.00 | 1495.00 | 1465.00 | -1.01 | 0.88 |
| 2107.00 | 2525.00 | 2438.00 | 1.19 | 0.00 |
| 2036.00 | 2228.00 | 2279.67 | 1.14 | 0.00 |
| 1840.00 | 2091.00 | 2035.00 | 1.12 | 0.00 |
| 4941.00 | 5115.00 | 5287.00 | 1.15 | 0.00 |
| 2224.00 | 2372.00 | 2466.33 | 1.20 | 0.00 |
| 324.00 | 278.00 | 307.33 | 1.13 | 0.07 |
| 2057.00 | 2266.00 | 2193.67 | 1.26 | 0.00 |
| 3946.00 | 4156.00 | 4393.00 | 1.19 | 0.00 |
| 2401.00 | 2597.00 | 2672.33 | 1.15 | 0.00 |
| 2446.00 | 2684.00 | 2689.33 | 1.27 | 0.00 |
| 2005.00 | 2191.00 | 2238.67 | 1.15 | 0.00 |
| 1206.00 | 1354.00 | 1347.33 | 1.17 | 0.00 |
| 1642.00 | 1885.00 | 1880.67 | 1.19 | 0.00 |
| 2063.00 | 2477.00 | 2340.00 | 1.19 | 0.00 |
| 373.00 | 488.00 | 466.33 | 1.45 | 0.00 |
| 1793.00 | 1899.00 | 1946.33 | 1.16 | 0.00 |
| 3513.00 | 3778.00 | 3916.33 | 1.19 | 0.00 |
| 1433.00 | 1716.00 | 1610.00 | 1.11 | 0.01 |
| 2727.00 | 2904.00 | 2759.33 | 1.12 | 0.01 |
| 1125.00 | 1255.00 | 1264.00 | 1.09 | 0.10 |
| 2090.00 | 2124.00 | 2160.33 | 1.14 | 0.00 |
| 1445.00 | 1754.00 | 1634.33 | 1.11 | 0.02 |
| 2056.00 | 2457.00 | 2378.67 | 1.21 | 0.00 |
| 2823.00 | 3179.00 | 3153.00 | 1.24 | 0.00 |

| | | | | |
|---|---|---|---|---|
| 11454.00 | 12407.00 | 12158.33 | -1.12 | 0.00 |
| 2745.00 | 2735.00 | 2903.67 | 1.14 | 0.00 |
| 2802.00 | 3095.00 | 3103.67 | 1.23 | 0.00 |
| 777.00 | 863.00 | 859.33 | 1.12 | 0.01 |
| 612.00 | 675.00 | 674.33 | -1.01 | 0.89 |
| 1510.00 | 1697.00 | 1669.67 | 1.20 | 0.00 |
| 92.00 | 81.00 | 87.67 | -1.42 | 0.00 |
| 2147.00 | 2395.00 | 2326.67 | 1.26 | 0.00 |
| 1864.00 | 2020.00 | 2084.33 | 1.13 | 0.00 |
| 1002.00 | 1030.00 | 1080.33 | 1.23 | 0.00 |
| 963.00 | 1118.00 | 1105.33 | 1.03 | 0.50 |
| 1027.00 | 1132.00 | 1161.33 | 1.12 | 0.02 |
| 2368.00 | 2538.00 | 2492.67 | 1.04 | 0.32 |
| 2360.00 | 2515.00 | 2430.67 | -1.05 | 0.26 |
| 1913.00 | 1882.00 | 1994.00 | 1.01 | 0.76 |
| 833.00 | 936.00 | 916.33 | 1.20 | 0.00 |
| 1099.00 | 1176.00 | 1214.00 | 1.09 | 0.06 |
| 1503.00 | 1579.00 | 1671.00 | 1.08 | 0.11 |
| 124.00 | 115.00 | 123.00 | -1.30 | 0.00 |
| 354.00 | 379.00 | 383.67 | 1.18 | 0.00 |
| 694.00 | 793.00 | 741.33 | 1.13 | 0.02 |
| 1146.00 | 1251.00 | 1285.33 | 1.20 | 0.00 |
| 11.00 | 10.00 | 11.33 | -1.52 | 0.09 |
| 661.00 | 744.00 | 743.67 | 1.12 | 0.02 |
| 149.00 | 131.00 | 139.33 | 1.05 | 0.60 |
| 172.00 | 163.00 | 173.00 | 1.26 | 0.00 |
| 2.00 | 8.00 | 4.00 | -1.58 | 0.33 |
| 1307.00 | 1538.00 | 1359.67 | -1.12 | 0.06 |
| 7927.00 | 7943.00 | 8115.00 | 1.09 | 0.01 |
| 728.00 | 985.00 | 866.67 | 1.13 | 0.05 |
| 1175.00 | 1423.00 | 1279.33 | 1.04 | 0.50 |
| 25.00 | 19.00 | 17.67 | 1.01 | 1.00 |
| 310.00 | 278.00 | 299.67 | -1.16 | 0.02 |
| 2249.00 | 2556.00 | 2483.33 | -1.01 | 0.82 |
| 137.00 | 130.00 | 131.67 | -1.17 | 0.06 |
| 17.00 | 14.00 | 13.00 | -1.27 | 0.41 |
| 3786.00 | 3721.00 | 3786.00 | -1.01 | 0.81 |
| 195.00 | 173.00 | 181.00 | -1.05 | 0.57 |
| 1290.00 | 1426.00 | 1388.67 | 1.15 | 0.00 |
| 13.00 | 10.00 | 11.33 | 1.59 | 0.11 |
| 5.00 | 7.00 | 6.00 | 1.36 | 0.48 |
| 665.00 | 806.00 | 722.33 | 1.23 | 0.00 |
| 27.00 | 20.00 | 23.33 | 1.06 | 0.74 |
| 54.00 | 70.00 | 63.33 | -1.79 | 0.00 |
| 4.00 | 4.00 | 4.33 | -1.85 | 0.11 |
| 211.00 | 212.00 | 219.67 | 1.34 | 0.00 |
| 1397.00 | 1499.00 | 1480.00 | 1.03 | 0.39 |
| 8.00 | 8.00 | 8.33 | -1.50 | 0.19 |

| | | | | |
|---|---|---|---|---|
| 6914.00 | 7971.00 | 7726.33 | 1.27 | 0.00 |
| 1100.00 | 1223.00 | 1118.33 | -1.10 | 0.07 |
| 2172.00 | 2072.00 | 2146.67 | -1.14 | 0.00 |
| 14.00 | 15.00 | 14.67 | -1.10 | 0.76 |
| 908.00 | 1073.00 | 1010.33 | 1.10 | 0.05 |
| 411.00 | 513.00 | 475.00 | -1.04 | 0.47 |
| 244.00 | 252.00 | 235.00 | -3.25 | 0.00 |
| 419.00 | 522.00 | 472.67 | 1.03 | 0.66 |
| 0.00 | 0.00 | 0.00 | -11.74 | 0.13 |
| 42.00 | 45.00 | 44.00 | -1.40 | 0.01 |
| 3041.00 | 3084.00 | 3175.33 | -1.15 | 0.00 |
| 86.00 | 74.00 | 76.33 | 1.06 | 0.58 |
| 7.00 | 6.00 | 6.67 | -1.26 | 0.56 |
| 387.00 | 427.00 | 419.33 | -1.25 | 0.00 |
| 11784.00 | 13393.00 | 13096.33 | 1.40 | 0.00 |
| 1586.00 | 1736.00 | 1737.67 | 1.15 | 0.00 |
| 1163.00 | 1302.00 | 1285.33 | -1.01 | 0.78 |
| 6.00 | 10.00 | 7.67 | -1.19 | 0.68 |
| 263.00 | 311.00 | 292.33 | -1.02 | 0.78 |
| 2013.00 | 2494.00 | 2357.33 | 1.20 | 0.00 |
| 5030.00 | 5681.00 | 5550.67 | 1.22 | 0.00 |
| 11.00 | 10.00 | 10.00 | -1.31 | 0.35 |
| 1012.00 | 994.00 | 1022.67 | -1.06 | 0.18 |
| 697.00 | 754.00 | 742.67 | 1.13 | 0.01 |
| 2145.00 | 2195.00 | 2216.67 | -1.04 | 0.25 |
| 4505.00 | 4538.00 | 4507.00 | 1.11 | 0.01 |
| 2884.00 | 3165.00 | 3088.33 | 1.18 | 0.00 |
| 857.00 | 854.00 | 848.00 | -1.04 | 0.46 |
| 4522.00 | 4146.00 | 4203.33 | -1.15 | 0.01 |
| 1204.00 | 1256.00 | 1326.33 | -1.14 | 0.01 |
| 1168.00 | 1423.00 | 1375.00 | 1.17 | 0.00 |
| 2.00 | 1.00 | 3.67 | -1.46 | 0.45 |
| 503.00 | 549.00 | 533.33 | -1.02 | 0.68 |
| 592.00 | 550.00 | 561.33 | 1.21 | 0.00 |
| 1787.00 | 1648.00 | 1709.67 | -1.28 | 0.00 |
| 369.00 | 331.00 | 337.00 | 1.00 | 0.97 |
| 331.00 | 416.00 | 376.67 | 1.20 | 0.00 |
| 22.00 | 18.00 | 24.00 | 1.77 | 0.01 |
| 1074.00 | 1089.00 | 1097.33 | -1.03 | 0.48 |
| 2069.00 | 2038.00 | 2118.00 | 1.08 | 0.04 |
| 494.00 | 470.00 | 496.33 | -1.13 | 0.02 |
| 55.00 | 50.00 | 55.33 | -1.24 | 0.07 |
| 163.00 | 126.00 | 137.00 | -1.17 | 0.12 |
| 545.00 | 579.00 | 566.33 | -1.19 | 0.00 |
| 15695.00 | 15009.00 | 15282.00 | -1.11 | 0.01 |
| 1624.00 | 1405.00 | 1463.33 | -1.06 | 0.38 |
| 68.00 | 65.00 | 66.67 | 1.32 | 0.02 |
| 144.00 | 131.00 | 140.00 | 1.24 | 0.01 |

| | | | | |
|---|---|---|---|---|
| 754.00 | 750.00 | 740.33 | -1.15 | 0.02 |
| 6.00 | 2.00 | 3.67 | -1.28 | 0.70 |
| 784.00 | 755.00 | 758.33 | -1.29 | 0.00 |
| 0.00 | 0.00 | 0.00 | -11.75 | 0.13 |
| 305.00 | 337.00 | 320.00 | 1.08 | 0.22 |
| 295.00 | 315.00 | 304.00 | -1.32 | 0.00 |
| 5.00 | 5.00 | 4.33 | 1.41 | 0.53 |
| 18.00 | 10.00 | 14.67 | -1.29 | 0.29 |
| 1865.00 | 1678.00 | 1802.33 | -1.21 | 0.00 |
| 0.00 | 0.00 | 0.33 | -4.69 | 0.13 |
| 190.00 | 215.00 | 217.00 | 2.30 | 0.00 |
| 4451.00 | 4464.00 | 4498.33 | 1.22 | 0.00 |
| 172.00 | 190.00 | 178.00 | 1.10 | 0.27 |
| 1150.00 | 1204.00 | 1245.67 | 1.10 | 0.05 |
| 158.00 | 220.00 | 202.33 | 1.28 | 0.00 |
| 374.00 | 409.00 | 426.67 | 1.14 | 0.03 |
| 1164.00 | 1379.00 | 1372.67 | 1.26 | 0.00 |
| 16506.00 | 15473.00 | 16205.67 | 1.18 | 0.00 |
| 7.00 | 7.00 | 7.67 | 1.50 | 0.26 |
| 39914.00 | 46389.00 | 43895.67 | 1.22 | 0.00 |
| 66.00 | 80.00 | 84.67 | 1.27 | 0.05 |
| 2237.00 | 2291.00 | 2227.33 | 1.00 | 0.99 |
| 2802.00 | 3116.00 | 2977.33 | 1.09 | 0.03 |
| 423.00 | 509.00 | 468.67 | -1.06 | 0.32 |
| 2351.00 | 2575.00 | 2517.33 | -1.10 | 0.01 |
| 25.00 | 27.00 | 29.00 | 1.24 | 0.22 |
| 833.00 | 925.00 | 958.33 | 1.02 | 0.75 |
| 164.00 | 162.00 | 154.33 | -1.02 | 0.82 |
| 170.00 | 181.00 | 187.67 | 1.27 | 0.00 |
| 1039.00 | 1156.00 | 1123.67 | 1.04 | 0.28 |
| 599.00 | 616.00 | 585.67 | 1.36 | 0.00 |
| 162.00 | 148.00 | 161.33 | 1.11 | 0.22 |
| 5748.00 | 5640.00 | 5743.33 | -1.63 | 0.00 |
| 2.00 | 6.00 | 3.33 | -1.11 | 1.00 |
| 792.00 | 762.00 | 814.33 | 1.09 | 0.08 |
| 2156.00 | 2087.00 | 2182.67 | -1.14 | 0.00 |
| 1012.00 | 1019.00 | 1018.67 | 1.08 | 0.11 |
| 3.00 | 3.00 | 7.00 | 2.23 | 0.08 |
| 1.00 | 6.00 | 2.67 | -1.50 | 0.51 |
| 1159.00 | 1299.00 | 1286.67 | -1.04 | 0.34 |
| 11.00 | 7.00 | 7.33 | -1.10 | 0.89 |
| 48.00 | 51.00 | 50.33 | -1.27 | 0.05 |
| 21.00 | 20.00 | 19.33 | -1.03 | 1.00 |
| 1541.00 | 1714.00 | 1666.67 | 1.07 | 0.07 |
| 324.00 | 338.00 | 346.33 | 1.12 | 0.06 |
| 1154.00 | 1457.00 | 1365.33 | 1.45 | 0.00 |
| 349.00 | 348.00 | 345.67 | -1.02 | 0.83 |
| 1065.00 | 1095.00 | 1127.00 | -1.02 | 0.68 |

| | | | | |
|---|---|---|---|---|
| 6049.00 | 5806.00 | 5982.33 | -1.25 | 0.00 |
| 4.00 | 7.00 | 5.33 | -1.20 | 0.74 |
| 607.00 | 635.00 | 641.00 | -1.40 | 0.00 |
| 236.00 | 205.00 | 232.00 | 1.25 | 0.00 |
| 224.00 | 212.00 | 209.67 | 1.02 | 0.80 |
| 7965.00 | 8075.00 | 8258.33 | -1.12 | 0.00 |
| 34.00 | 25.00 | 33.33 | 1.28 | 0.14 |
| 181.00 | 175.00 | 195.67 | -1.04 | 0.58 |
| 1.00 | 0.00 | 0.33 | -3.95 | 0.23 |
| 4445.00 | 4613.00 | 4741.33 | -1.06 | 0.07 |
| 2627.00 | 2876.00 | 2961.00 | 1.13 | 0.00 |
| 4473.00 | 4486.00 | 4501.33 | -1.12 | 0.00 |
| 191.00 | 210.00 | 200.33 | 1.96 | 0.00 |
| 18.00 | 12.00 | 14.00 | -1.23 | 0.44 |
| 399.00 | 444.00 | 394.33 | 1.10 | 0.23 |
| 1225.00 | 1267.00 | 1269.67 | 1.08 | 0.06 |
| 554.00 | 586.00 | 595.33 | 1.20 | 0.00 |
| 336.00 | 357.00 | 378.00 | -1.26 | 0.00 |
| 26.00 | 19.00 | 25.00 | 1.34 | 0.13 |
| 6.00 | 4.00 | 3.67 | -1.90 | 0.12 |
| 3961.00 | 4179.00 | 4166.00 | -1.00 | 0.93 |
| 441.00 | 441.00 | 452.00 | 1.23 | 0.00 |
| 2923.00 | 3343.00 | 3016.33 | -1.00 | 0.99 |
| 832.00 | 830.00 | 841.67 | -1.10 | 0.03 |
| 518.00 | 521.00 | 554.67 | -1.18 | 0.00 |
| 218.00 | 217.00 | 219.67 | 1.34 | 0.00 |
| 1799.00 | 1996.00 | 1961.67 | -1.02 | 0.55 |
| 136.00 | 143.00 | 154.67 | -1.53 | 0.00 |
| 587.00 | 666.00 | 626.00 | 1.11 | 0.03 |
| 1970.00 | 2006.00 | 1964.33 | 1.14 | 0.00 |
| 27284.00 | 29244.00 | 28648.33 | -1.39 | 0.00 |
| 3315.00 | 3371.00 | 3384.67 | -1.13 | 0.00 |
| 1334.00 | 1434.00 | 1432.00 | -1.13 | 0.00 |
| 1344.00 | 1542.00 | 1475.33 | 1.11 | 0.02 |
| 296.00 | 347.00 | 331.33 | 1.35 | 0.00 |
| 558.00 | 596.00 | 587.00 | 1.05 | 0.36 |
| 1182.00 | 1183.00 | 1183.67 | -1.01 | 0.82 |
| 2267.00 | 2334.00 | 2364.67 | -1.09 | 0.02 |
| 1145.00 | 1081.00 | 1183.00 | -1.04 | 0.41 |
| 2491.00 | 2896.00 | 2730.33 | 1.03 | 0.44 |
| 1136.00 | 1276.00 | 1285.67 | 1.14 | 0.01 |
| 1071.00 | 1106.00 | 1148.67 | 1.18 | 0.00 |
| 22.00 | 28.00 | 30.33 | -1.03 | 0.95 |
| 290.00 | 293.00 | 292.00 | 1.08 | 0.23 |
| 875.00 | 961.00 | 910.33 | 1.01 | 0.78 |
| 21.00 | 16.00 | 22.33 | 1.32 | 0.23 |
| 2.00 | 7.00 | 5.33 | -1.32 | 0.52 |
| 1431.00 | 1239.00 | 1370.33 | -1.23 | 0.00 |

| | | | | |
|---|---|---|---|---|
| 1125.00 | 1305.00 | 1263.67 | 1.08 | 0.07 |
| 2853.00 | 2760.00 | 2868.00 | -1.09 | 0.03 |
| 1583.00 | 1791.00 | 1725.33 | 1.12 | 0.02 |
| 198.00 | 221.00 | 221.33 | 1.09 | 0.21 |
| 1266.00 | 1489.00 | 1430.33 | 1.12 | 0.02 |
| 1883.00 | 2261.00 | 2179.00 | 1.13 | 0.01 |
| 3378.00 | 3355.00 | 3587.00 | -1.01 | 0.88 |
| 21.00 | 16.00 | 20.67 | -1.05 | 0.86 |
| 628.00 | 741.00 | 696.67 | 1.06 | 0.25 |
| 1035.00 | 1173.00 | 1174.00 | -1.16 | 0.00 |
| 642.00 | 645.00 | 644.67 | -1.05 | 0.42 |
| 1148.00 | 1319.00 | 1325.33 | -1.03 | 0.56 |
| 27.00 | 18.00 | 26.33 | -1.47 | 0.02 |
| 134.00 | 162.00 | 156.00 | -1.01 | 0.98 |
| 205.00 | 199.00 | 203.67 | 1.37 | 0.00 |
| 324.00 | 363.00 | 338.33 | 1.09 | 0.14 |
| 1067.00 | 1125.00 | 1144.00 | -1.08 | 0.07 |
| 1923.00 | 2384.00 | 2180.00 | 1.17 | 0.00 |
| 393.00 | 451.00 | 451.00 | -1.02 | 0.78 |
| 23.00 | 49.00 | 44.00 | 1.49 | 0.03 |
| 63.00 | 69.00 | 69.67 | 1.05 | 0.72 |
| 1279.00 | 1265.00 | 1297.67 | 1.01 | 0.87 |
| 896.00 | 873.00 | 946.67 | 1.10 | 0.07 |
| 152.00 | 120.00 | 144.33 | 1.04 | 0.71 |
| 1598.00 | 1719.00 | 1750.67 | 1.01 | 0.72 |
| 1392.00 | 1326.00 | 1414.67 | 1.07 | 0.11 |
| 94.00 | 55.00 | 76.33 | -1.35 | 0.04 |
| 220.00 | 221.00 | 226.00 | -1.01 | 0.87 |
| 35.00 | 24.00 | 27.00 | 1.38 | 0.08 |
| 21.00 | 19.00 | 25.00 | 1.25 | 0.26 |
| 7578.00 | 7779.00 | 7712.67 | 1.14 | 0.00 |
| 6.00 | 2.00 | 5.00 | -1.61 | 0.21 |
| 39.00 | 41.00 | 46.67 | 1.20 | 0.21 |
| 1733.00 | 1782.00 | 1825.00 | -1.03 | 0.46 |
| 1594.00 | 1736.00 | 1667.33 | -1.04 | 0.31 |
| 8.00 | 13.00 | 11.00 | 1.35 | 0.30 |
| 722.00 | 765.00 | 754.00 | 1.23 | 0.00 |
| 4.00 | 8.00 | 6.33 | -1.43 | 0.32 |
| 3.00 | 1.00 | 3.00 | -1.12 | 1.00 |
| 179.00 | 159.00 | 178.33 | 1.07 | 0.40 |
| 455.00 | 420.00 | 446.67 | -1.14 | 0.01 |
| 46.00 | 29.00 | 31.67 | -1.47 | 0.05 |
| 386.00 | 383.00 | 427.67 | 1.25 | 0.00 |
| 1069.00 | 1245.00 | 1194.00 | 1.08 | 0.11 |
| 1227.00 | 1171.00 | 1241.33 | -1.12 | 0.01 |
| 1903.00 | 2021.00 | 2005.00 | 1.05 | 0.22 |
| 1716.00 | 1957.00 | 1859.00 | 1.01 | 0.88 |
| 43.00 | 59.00 | 60.00 | 1.30 | 0.05 |

| | | | | |
|---|---|---|---|---|
| 5987.00 | 6024.00 | 6199.67 | 1.12 | 0.00 |
| 837.00 | 863.00 | 892.67 | 1.10 | 0.04 |
| 600.00 | 677.00 | 639.33 | -1.34 | 0.00 |
| 518.00 | 541.00 | 561.33 | -1.07 | 0.20 |
| 31.00 | 18.00 | 28.33 | 2.03 | 0.00 |
| 634.00 | 583.00 | 654.00 | -1.18 | 0.01 |
| 2417.00 | 2742.00 | 2667.67 | 1.09 | 0.02 |
| 645.00 | 633.00 | 676.33 | 1.09 | 0.08 |
| 697.00 | 789.00 | 809.67 | 1.09 | 0.14 |
| 1175.00 | 1247.00 | 1204.67 | -1.06 | 0.15 |
| 463.00 | 487.00 | 492.33 | 1.14 | 0.02 |
| 1148.00 | 1303.00 | 1268.00 | 1.06 | 0.16 |
| 523.00 | 462.00 | 505.33 | -1.00 | 1.00 |
| 6414.00 | 7801.00 | 7415.00 | 1.10 | 0.02 |
| 190.00 | 226.00 | 213.33 | 1.11 | 0.14 |
| 80.00 | 76.00 | 79.33 | -1.19 | 0.08 |
| 435.00 | 424.00 | 448.33 | -1.01 | 0.84 |
| 474.00 | 535.00 | 524.67 | -1.03 | 0.60 |
| 2525.00 | 2517.00 | 2620.00 | 1.08 | 0.04 |
| 5498.00 | 6216.00 | 6105.33 | 1.17 | 0.00 |
| 1566.00 | 1544.00 | 1636.00 | -1.04 | 0.29 |
| 896.00 | 873.00 | 939.00 | 1.16 | 0.00 |
| 1403.00 | 1583.00 | 1496.00 | 1.27 | 0.00 |
| 736.00 | 861.00 | 800.67 | 1.15 | 0.01 |
| 50.00 | 65.00 | 61.33 | -1.07 | 0.54 |
| 752.00 | 872.00 | 829.00 | -1.05 | 0.28 |
| 6296.00 | 7178.00 | 6735.00 | 1.17 | 0.00 |
| 5.00 | 4.00 | 5.67 | -1.42 | 0.36 |
| 24.00 | 16.00 | 23.00 | 1.54 | 0.04 |
| 2811.00 | 2753.00 | 2890.33 | 1.16 | 0.00 |
| 2698.00 | 2895.00 | 2913.00 | 1.13 | 0.00 |
| 2006.00 | 2315.00 | 2177.00 | -1.08 | 0.06 |
| 161.00 | 186.00 | 178.67 | 1.71 | 0.00 |
| 387.00 | 475.00 | 418.00 | 1.08 | 0.27 |
| 109.00 | 80.00 | 81.67 | -1.90 | 0.00 |
| 235.00 | 207.00 | 227.67 | 1.06 | 0.47 |
| 613.00 | 713.00 | 713.00 | 1.16 | 0.01 |
| 275.00 | 323.00 | 294.33 | 1.14 | 0.05 |
| 892.00 | 960.00 | 960.00 | -1.11 | 0.01 |
| 335.00 | 400.00 | 371.33 | 1.29 | 0.00 |
| 393.00 | 401.00 | 406.67 | 1.11 | 0.06 |
| 464.00 | 411.00 | 401.00 | -1.45 | 0.00 |
| 276.00 | 319.00 | 300.00 | -1.01 | 0.85 |
| 11.00 | 18.00 | 16.67 | 1.17 | 0.54 |
| 1332.00 | 1538.00 | 1533.67 | 1.10 | 0.04 |
| 12.00 | 13.00 | 14.33 | -1.65 | 0.02 |
| 9.00 | 12.00 | 10.00 | 1.28 | 0.42 |
| 26.00 | 35.00 | 32.67 | 1.19 | 0.32 |

| | | | | |
|---|---|---|---|---|
| 1764.00 | 1790.00 | 1835.67 | -1.04 | 0.34 |
| 7.00 | 19.00 | 13.00 | -1.50 | 0.09 |
| 1521.00 | 1369.00 | 1427.00 | -1.26 | 0.00 |
| 8325.00 | 8860.00 | 8513.33 | 1.06 | 0.17 |
| 1281.00 | 1523.00 | 1425.33 | 1.12 | 0.01 |
| 982.00 | 1011.00 | 1048.33 | 1.10 | 0.04 |
| 2200.00 | 2275.00 | 2282.67 | 1.01 | 0.74 |
| 374.00 | 377.00 | 364.67 | -1.05 | 0.43 |
| 22.00 | 17.00 | 22.00 | -1.21 | 0.30 |
| 540.00 | 566.00 | 552.33 | -1.02 | 0.70 |
| 2577.00 | 2664.00 | 2623.33 | -1.08 | 0.05 |
| 925.00 | 911.00 | 938.00 | 1.16 | 0.00 |
| 61.00 | 62.00 | 61.67 | -1.05 | 0.74 |
| 659.00 | 715.00 | 676.00 | -1.36 | 0.00 |
| 2886.00 | 3057.00 | 3118.67 | 1.12 | 0.00 |
| 12.00 | 13.00 | 10.67 | 1.02 | 1.00 |
| 8.00 | 7.00 | 5.67 | -1.01 | 1.00 |
| 9.00 | 6.00 | 7.33 | -1.15 | 0.78 |
| 6490.00 | 7364.00 | 7271.67 | 1.25 | 0.00 |
| 1494.00 | 1730.00 | 1675.33 | 1.23 | 0.00 |
| 672.00 | 757.00 | 718.00 | -1.03 | 0.62 |
| 357.00 | 363.00 | 355.33 | -1.03 | 0.66 |
| 3796.00 | 4306.00 | 4056.00 | -1.03 | 0.39 |
| 1043.00 | 1088.00 | 1122.33 | 1.01 | 0.80 |
| 1444.00 | 1613.00 | 1571.67 | -1.07 | 0.10 |
| 979.00 | 1024.00 | 1040.00 | 1.05 | 0.23 |
| 5141.00 | 5229.00 | 5350.00 | 1.01 | 0.74 |
| 866.00 | 995.00 | 1015.33 | -1.06 | 0.25 |
| 580.00 | 790.00 | 717.67 | 1.19 | 0.01 |
| 1361.00 | 1530.00 | 1487.00 | 1.14 | 0.00 |
| 6045.00 | 5892.00 | 5948.00 | 1.19 | 0.00 |
| 981.00 | 1058.00 | 1039.00 | 1.07 | 0.12 |
| 449.00 | 501.00 | 488.67 | 1.10 | 0.08 |
| 2964.00 | 3255.00 | 3273.00 | 1.65 | 0.00 |
| 476.00 | 591.00 | 544.33 | 1.16 | 0.01 |
| 22.00 | 32.00 | 21.67 | -1.41 | 0.07 |
| 1440.00 | 1462.00 | 1464.00 | 1.07 | 0.09 |
| 569.00 | 615.00 | 576.33 | -1.47 | 0.00 |
| 350.00 | 369.00 | 368.67 | -1.09 | 0.10 |
| 6.00 | 5.00 | 6.00 | -1.40 | 0.38 |
| 182.00 | 196.00 | 200.33 | 1.23 | 0.00 |
| 572.00 | 603.00 | 583.67 | -1.17 | 0.00 |
| 183.00 | 220.00 | 207.00 | 1.07 | 0.37 |
| 2734.00 | 2929.00 | 2851.00 | 1.17 | 0.00 |
| 7.00 | 8.00 | 6.00 | -2.22 | 0.01 |
| 529.00 | 617.00 | 607.33 | -1.45 | 0.00 |
| 20.00 | 22.00 | 29.33 | 1.17 | 0.49 |
| 91.00 | 89.00 | 88.67 | -1.26 | 0.02 |

| | | | | |
|---|---|---|---|---|
| 198.00 | 207.00 | 218.00 | -1.38 | 0.00 |
| 58.00 | 84.00 | 78.00 | 1.67 | 0.00 |
| 5.00 | 1.00 | 5.33 | -1.14 | 0.87 |
| 2530.00 | 2881.00 | 2765.00 | 1.11 | 0.00 |
| 689.00 | 765.00 | 749.67 | 1.09 | 0.06 |
| 3038.00 | 3268.00 | 3121.33 | 1.18 | 0.00 |
| 1494.00 | 1787.00 | 1652.33 | 1.24 | 0.00 |
| 1427.00 | 1339.00 | 1384.33 | -1.19 | 0.00 |
| 6.00 | 10.00 | 5.67 | 1.28 | 0.60 |
| 13.00 | 11.00 | 11.00 | 1.41 | 0.24 |
| 1703.00 | 1718.00 | 1717.67 | -1.02 | 0.68 |
| 45.00 | 45.00 | 49.67 | 1.03 | 0.83 |
| 4.00 | 8.00 | 5.67 | -1.48 | 0.29 |
| 1.00 | 8.00 | 4.67 | 1.06 | 1.00 |
| 5.00 | 7.00 | 5.67 | 1.28 | 0.59 |
| 602.00 | 588.00 | 617.67 | -1.22 | 0.00 |
| 216.00 | 235.00 | 220.67 | -1.58 | 0.00 |
| 8.00 | 4.00 | 6.67 | -1.06 | 1.00 |
| 60.00 | 75.00 | 65.00 | 1.20 | 0.12 |
| 7.00 | 15.00 | 11.00 | 1.62 | 0.11 |
| 3.00 | 1.00 | 3.33 | -1.40 | 0.55 |
| 686.00 | 651.00 | 707.67 | 1.08 | 0.11 |
| 1.00 | 0.00 | 1.00 | -3.41 | 0.06 |
| 763.00 | 751.00 | 772.00 | -1.09 | 0.06 |
| 349.00 | 415.00 | 421.67 | 1.09 | 0.20 |
| 1.00 | 2.00 | 2.00 | -1.97 | 0.25 |
| 6.00 | 4.00 | 7.67 | 1.33 | 0.44 |
| 1682.00 | 1788.00 | 1716.00 | -1.18 | 0.00 |
| 88.00 | 111.00 | 110.00 | -1.13 | 0.22 |
| 1680.00 | 1699.00 | 1717.67 | -1.02 | 0.65 |
| 517.00 | 611.00 | 589.33 | -1.10 | 0.17 |
| 826.00 | 898.00 | 841.00 | 1.17 | 0.01 |
| 2338.00 | 2410.00 | 2352.00 | -1.01 | 0.87 |
| 794.00 | 890.00 | 829.33 | -1.05 | 0.29 |
| 2653.00 | 2916.00 | 2894.00 | 1.12 | 0.00 |
| 588.00 | 630.00 | 632.33 | 1.08 | 0.12 |
| 2372.00 | 2584.00 | 2538.33 | -1.10 | 0.01 |
| 3571.00 | 3825.00 | 3859.00 | 1.07 | 0.05 |
| 1023.00 | 1155.00 | 1161.33 | 1.17 | 0.00 |
| 469.00 | 471.00 | 471.67 | 1.23 | 0.00 |
| 2318.00 | 2423.00 | 2445.00 | -1.40 | 0.00 |
| 524.00 | 575.00 | 554.33 | -1.06 | 0.26 |
| 19.00 | 23.00 | 19.00 | -1.17 | 0.50 |
| 5.00 | 3.00 | 3.33 | -1.11 | 1.00 |
| 87.00 | 89.00 | 87.33 | -1.11 | 0.29 |
| 57.00 | 66.00 | 73.33 | 1.12 | 0.37 |
| 548.00 | 631.00 | 592.33 | 1.25 | 0.00 |
| 2755.00 | 2883.00 | 2804.00 | -1.07 | 0.09 |

| | | | | |
|---|---|---|---|---|
| 1791.00 | 1882.00 | 1903.33 | 1.02 | 0.64 |
| 2396.00 | 2487.00 | 2495.67 | 1.00 | 0.91 |
| 856.00 | 987.00 | 904.00 | -1.07 | 0.21 |
| 1150.00 | 1222.00 | 1231.00 | 1.15 | 0.00 |
| 518.00 | 549.00 | 553.00 | 1.02 | 0.68 |
| 1740.00 | 1784.00 | 1783.00 | -1.00 | 0.94 |
| 2322.00 | 2567.00 | 2511.33 | 1.08 | 0.05 |
| 481.00 | 571.00 | 540.33 | 1.10 | 0.07 |
| 1900.00 | 1787.00 | 1829.33 | -1.01 | 0.78 |
| 12.00 | 14.00 | 14.33 | 1.09 | 0.75 |
| 1217.00 | 1370.00 | 1334.33 | -1.04 | 0.39 |
| 736.00 | 758.00 | 792.67 | -1.18 | 0.00 |
| 18.00 | 25.00 | 20.33 | -1.21 | 0.34 |
| 4214.00 | 5091.00 | 4729.67 | 1.01 | 0.80 |
| 1806.00 | 2012.00 | 1885.67 | 1.06 | 0.14 |
| 97.00 | 119.00 | 110.00 | 1.04 | 0.70 |
| 944.00 | 970.00 | 961.00 | 1.14 | 0.01 |
| 1639.00 | 1777.00 | 1736.00 | 1.08 | 0.03 |
| 2418.00 | 2660.00 | 2642.67 | 1.28 | 0.00 |
| 2558.00 | 2532.00 | 2521.00 | -1.27 | 0.00 |
| 738.00 | 876.00 | 842.33 | 1.10 | 0.07 |
| 400.00 | 420.00 | 430.00 | 1.08 | 0.17 |
| 9795.00 | 11833.00 | 11356.00 | -1.09 | 0.05 |
| 1346.00 | 1492.00 | 1497.33 | -1.06 | 0.14 |
| 646.00 | 787.00 | 743.33 | -1.05 | 0.42 |
| 1146.00 | 1209.00 | 1236.33 | 1.09 | 0.04 |
| 164.00 | 214.00 | 186.67 | 1.38 | 0.00 |
| 18.00 | 22.00 | 21.67 | -1.67 | 0.00 |
| 269.00 | 294.00 | 288.00 | 1.21 | 0.00 |
| 9.00 | 6.00 | 6.33 | -1.32 | 0.48 |
| 435.00 | 507.00 | 494.00 | -1.08 | 0.16 |
| 564.00 | 602.00 | 614.33 | -1.04 | 0.40 |
| 2515.00 | 2741.00 | 2728.33 | -1.10 | 0.01 |
| 2037.00 | 2331.00 | 2242.33 | -1.04 | 0.29 |
| 422.00 | 469.00 | 458.33 | 1.01 | 0.94 |
| 1876.00 | 1952.00 | 1993.67 | 1.18 | 0.00 |
| 1660.00 | 1891.00 | 1784.67 | 1.12 | 0.02 |
| 633.00 | 638.00 | 636.00 | -1.22 | 0.00 |
| 437.00 | 484.00 | 450.67 | 1.03 | 0.68 |
| 25.00 | 23.00 | 24.33 | 1.03 | 0.88 |
| 6110.00 | 6805.00 | 6527.33 | 1.16 | 0.00 |
| 285.00 | 362.00 | 345.33 | 1.02 | 0.77 |
| 575.00 | 637.00 | 614.33 | 1.17 | 0.00 |
| 6.00 | 3.00 | 4.67 | 1.51 | 0.41 |
| 963.00 | 914.00 | 937.33 | -1.17 | 0.00 |
| 95.00 | 104.00 | 107.00 | -1.06 | 0.57 |
| 929.00 | 955.00 | 943.33 | -1.08 | 0.08 |
| 412.00 | 464.00 | 480.67 | 1.17 | 0.02 |

| | | | | |
|---|---|---|---|---|
| 622.00 | 743.00 | 685.00 | -1.01 | 0.87 |
| 370.00 | 332.00 | 329.33 | 1.01 | 0.91 |
| 2031.00 | 2062.00 | 2115.00 | 1.49 | 0.00 |
| 10.00 | 11.00 | 11.33 | 1.15 | 0.62 |
| 23.00 | 38.00 | 25.00 | -1.12 | 0.67 |
| 871.00 | 922.00 | 891.33 | 1.07 | 0.16 |
| 136.00 | 155.00 | 139.00 | -1.08 | 0.37 |
| 2345.00 | 2531.00 | 2466.33 | 1.14 | 0.00 |
| 2345.00 | 2706.00 | 2631.00 | 1.17 | 0.00 |
| 690.00 | 681.00 | 706.33 | 1.03 | 0.48 |
| 897.00 | 936.00 | 956.67 | 1.01 | 0.80 |
| 287.00 | 315.00 | 300.00 | 1.12 | 0.13 |
| 667.00 | 638.00 | 700.33 | 1.14 | 0.01 |
| 9977.00 | 10148.00 | 10120.33 | 1.06 | 0.12 |
| 2689.00 | 3021.00 | 2873.33 | 1.12 | 0.00 |
| 4948.00 | 5158.00 | 5217.33 | -1.18 | 0.00 |
| 636.00 | 704.00 | 656.00 | 1.04 | 0.47 |
| 17.00 | 27.00 | 24.33 | -1.08 | 0.65 |
| 814.00 | 964.00 | 892.00 | -1.06 | 0.22 |
| 379.00 | 400.00 | 407.33 | 1.04 | 0.51 |
| 4204.00 | 4449.00 | 4517.00 | 1.10 | 0.03 |
| 8.00 | 5.00 | 7.00 | 1.29 | 0.52 |
| 94.00 | 55.00 | 75.00 | 1.25 | 0.10 |
| 7.00 | 11.00 | 12.00 | 1.01 | 1.00 |
| 24.00 | 28.00 | 22.67 | -1.78 | 0.00 |
| 3090.00 | 3380.00 | 3225.67 | 1.13 | 0.00 |
| 75.00 | 67.00 | 76.33 | 1.10 | 0.47 |
| 243.00 | 218.00 | 234.33 | -1.13 | 0.06 |
| 2827.00 | 3068.00 | 3022.67 | 1.15 | 0.00 |
| 1200.00 | 1240.00 | 1284.00 | 1.15 | 0.01 |
| 1366.00 | 1315.00 | 1369.67 | -1.06 | 0.22 |
| 1.00 | 4.00 | 2.67 | 1.12 | 1.00 |
| 62.00 | 66.00 | 71.67 | -1.01 | 0.93 |
| 1007.00 | 1113.00 | 1058.67 | 1.17 | 0.00 |
| 1087.00 | 1093.00 | 1104.67 | 1.15 | 0.00 |
| 2969.00 | 3084.00 | 3111.33 | 1.02 | 0.73 |
| 72.00 | 67.00 | 76.33 | -1.10 | 0.39 |
| 88.00 | 61.00 | 72.00 | -1.34 | 0.01 |
| 2611.00 | 2431.00 | 2649.33 | 1.03 | 0.53 |
| 632.00 | 654.00 | 708.00 | 1.54 | 0.00 |
| 259.00 | 266.00 | 281.67 | 1.10 | 0.25 |
| 138.00 | 170.00 | 158.00 | -1.37 | 0.00 |
| 45.00 | 49.00 | 44.67 | -2.47 | 0.00 |
| 551.00 | 459.00 | 500.00 | -1.21 | 0.00 |
| 14.00 | 11.00 | 12.67 | -1.36 | 0.22 |
| 27.00 | 33.00 | 34.33 | -1.72 | 0.00 |
| 30.00 | 34.00 | 31.67 | -1.43 | 0.01 |
| 3489.00 | 3420.00 | 3614.00 | 1.03 | 0.40 |

| | | | | |
|---|---|---|---|---|
| 1328.00 | 1508.00 | 1443.00 | -1.07 | 0.11 |
| 22.00 | 12.00 | 17.33 | -1.88 | 0.00 |
| 3075.00 | 3201.00 | 3231.00 | -1.07 | 0.06 |
| 2255.00 | 1986.00 | 2164.00 | -1.17 | 0.00 |
| 39.00 | 50.00 | 47.67 | -1.47 | 0.00 |
| 979.00 | 1147.00 | 1071.00 | 1.08 | 0.08 |
| 1605.00 | 1578.00 | 1604.33 | -1.14 | 0.00 |
| 929.00 | 1039.00 | 1024.00 | -1.03 | 0.52 |
| 522.00 | 559.00 | 529.33 | -1.23 | 0.00 |
| 1439.00 | 1360.00 | 1441.33 | -1.00 | 0.92 |
| 1702.00 | 1916.00 | 1882.67 | 1.02 | 0.65 |
| 894.00 | 847.00 | 907.67 | 1.12 | 0.01 |
| 2532.00 | 2803.00 | 2661.67 | 1.13 | 0.00 |
| 522.00 | 642.00 | 616.00 | 1.44 | 0.00 |
| 582.00 | 772.00 | 674.00 | 1.31 | 0.00 |
| 976.00 | 1154.00 | 1141.67 | 1.19 | 0.00 |
| 758.00 | 852.00 | 845.00 | 1.23 | 0.00 |
| 69.00 | 66.00 | 72.00 | 1.04 | 0.72 |
| 101.00 | 87.00 | 96.33 | -1.22 | 0.03 |
| 447.00 | 559.00 | 504.33 | 1.24 | 0.00 |
| 1541.00 | 1506.00 | 1613.67 | -1.13 | 0.01 |
| 5.00 | 4.00 | 6.67 | -1.66 | 0.11 |
| 1702.00 | 1719.00 | 1704.00 | -1.01 | 0.83 |
| 1282.00 | 1240.00 | 1297.00 | 1.05 | 0.24 |
| 4.00 | 6.00 | 6.00 | 2.46 | 0.05 |
| 853.00 | 891.00 | 944.33 | 1.14 | 0.01 |
| 1792.00 | 1968.00 | 1899.67 | 1.18 | 0.00 |
| 384.00 | 371.00 | 392.67 | -1.19 | 0.00 |
| 17.00 | 15.00 | 16.00 | 1.81 | 0.02 |
| 513.00 | 600.00 | 571.00 | 1.08 | 0.29 |
| 45.00 | 54.00 | 47.67 | 1.19 | 0.22 |
| 1398.00 | 1619.00 | 1523.67 | 1.27 | 0.00 |
| 571.00 | 606.00 | 571.33 | -1.04 | 0.52 |
| 3130.00 | 3174.00 | 3193.33 | 1.11 | 0.00 |
| 761.00 | 849.00 | 852.67 | 1.06 | 0.19 |
| 241.00 | 303.00 | 261.00 | 1.07 | 0.44 |
| 2.00 | 4.00 | 2.67 | -1.62 | 0.39 |
| 31.00 | 34.00 | 30.67 | -1.05 | 0.84 |
| 54.00 | 35.00 | 46.67 | 1.70 | 0.00 |
| 401.00 | 405.00 | 412.67 | -1.01 | 0.90 |
| 13.00 | 16.00 | 15.00 | -1.01 | 1.00 |
| 352.00 | 353.00 | 375.67 | -1.39 | 0.00 |
| 6256.00 | 6776.00 | 6831.33 | 1.07 | 0.06 |
| 2.00 | 2.00 | 2.33 | -1.98 | 0.20 |
| 311.00 | 250.00 | 255.67 | -1.55 | 0.00 |
| 13.00 | 11.00 | 10.00 | -1.51 | 0.12 |
| 1291.00 | 1414.00 | 1412.00 | -1.18 | 0.00 |
| 1261.00 | 1246.00 | 1241.33 | -1.22 | 0.00 |

| | | | | |
|---|---|---|---|---|
| 173.00 | 198.00 | 196.33 | -1.19 | 0.02 |
| 31.00 | 16.00 | 24.00 | 1.15 | 0.53 |
| 69.00 | 66.00 | 73.33 | -1.23 | 0.05 |
| 311.00 | 305.00 | 318.67 | 1.25 | 0.00 |
| 110.00 | 125.00 | 121.33 | 1.07 | 0.50 |
| 566.00 | 628.00 | 631.00 | 1.13 | 0.01 |
| 2.00 | 5.00 | 3.67 | -1.46 | 0.46 |
| 2561.00 | 2808.00 | 2832.67 | 1.46 | 0.00 |
| 2785.00 | 3224.00 | 3085.00 | 2.78 | 0.00 |
| 174.00 | 177.00 | 178.00 | 1.08 | 0.31 |
| 1.00 | 0.00 | 1.00 | -4.92 | 0.01 |
| 4.00 | 10.00 | 7.00 | -1.06 | 1.00 |
| 1725.00 | 1842.00 | 1797.00 | -1.04 | 0.32 |
| 518.00 | 525.00 | 534.33 | 1.10 | 0.08 |
| 660.00 | 806.00 | 734.33 | 1.13 | 0.03 |
| 873.00 | 849.00 | 899.00 | -1.69 | 0.00 |
| 974.00 | 1019.00 | 1028.00 | -1.25 | 0.00 |
| 85.00 | 67.00 | 82.00 | -1.61 | 0.00 |
| 8.00 | 3.00 | 5.00 | -1.47 | 0.33 |
| 1759.00 | 1979.00 | 2021.67 | 1.14 | 0.00 |
| 85.00 | 107.00 | 108.00 | -1.22 | 0.03 |
| 689.00 | 728.00 | 751.67 | 1.26 | 0.00 |
| 5822.00 | 6211.00 | 6247.33 | 1.16 | 0.00 |
| 100.00 | 106.00 | 99.67 | -1.79 | 0.00 |
| 466.00 | 474.00 | 476.67 | -1.08 | 0.12 |
| 291.00 | 302.00 | 303.33 | 1.11 | 0.07 |
| 3064.00 | 3503.00 | 3268.33 | -1.05 | 0.23 |
| 3151.00 | 3393.00 | 3520.00 | 1.12 | 0.01 |
| 99.00 | 104.00 | 112.67 | 1.19 | 0.07 |
| 598.00 | 619.00 | 652.00 | 1.03 | 0.56 |
| 1.00 | 7.00 | 3.33 | -1.79 | 0.20 |
| 239.00 | 269.00 | 275.67 | 1.16 | 0.05 |
| 931.00 | 1036.00 | 1033.33 | 1.01 | 0.86 |
| 822.00 | 852.00 | 867.67 | 1.06 | 0.24 |
| 2481.00 | 2418.00 | 2613.67 | -1.08 | 0.08 |
| 97.00 | 93.00 | 108.00 | 1.02 | 0.83 |
| 8.00 | 13.00 | 11.00 | 1.25 | 0.45 |
| 110.00 | 124.00 | 109.67 | 1.22 | 0.06 |
| 12.00 | 9.00 | 8.00 | -1.30 | 0.44 |
| 10.00 | 12.00 | 12.67 | 1.95 | 0.02 |
| 959.00 | 931.00 | 948.33 | -1.18 | 0.00 |
| 1750.00 | 1756.00 | 1863.67 | -1.03 | 0.53 |
| 428.00 | 527.00 | 473.67 | 1.07 | 0.25 |
| 1149.00 | 1208.00 | 1170.67 | -1.27 | 0.00 |
| 1634.00 | 1827.00 | 1676.33 | 1.05 | 0.36 |
| 475.00 | 519.00 | 512.33 | -1.25 | 0.00 |
| 180.00 | 199.00 | 203.67 | -1.10 | 0.18 |
| 1.00 | 1.00 | 1.67 | -1.39 | 0.78 |

| | | | | |
|---|---|---|---|---|
| 1078.00 | 1324.00 | 1170.33 | 1.05 | 0.39 |
| 20.00 | 34.00 | 22.67 | -1.25 | 0.28 |
| 60.00 | 101.00 | 81.33 | 1.06 | 0.70 |
| 3.00 | 3.00 | 3.33 | 1.61 | 0.46 |
| 1048.00 | 993.00 | 1081.33 | -1.29 | 0.00 |
| 2186.00 | 2324.00 | 2343.00 | 1.29 | 0.00 |
| 702.00 | 647.00 | 700.00 | -1.43 | 0.00 |
| 65.00 | 68.00 | 63.67 | -1.09 | 0.45 |
| 614.00 | 786.00 | 729.33 | 1.08 | 0.18 |
| 1075.00 | 1261.00 | 1205.33 | 1.21 | 0.00 |
| 1070.00 | 1260.00 | 1195.00 | 1.01 | 0.84 |
| 1529.00 | 1727.00 | 1628.33 | -1.00 | 1.00 |
| 872.00 | 921.00 | 968.33 | -1.06 | 0.23 |
| 1104.00 | 1245.00 | 1199.67 | 1.01 | 0.77 |
| 66.00 | 75.00 | 75.33 | 1.16 | 0.20 |
| 506.00 | 568.00 | 572.00 | 1.04 | 0.54 |
| 563.00 | 582.00 | 607.00 | 1.09 | 0.12 |
| 445.00 | 498.00 | 481.00 | -1.08 | 0.13 |
| 1904.00 | 2075.00 | 1954.67 | 1.04 | 0.43 |
| 15.00 | 14.00 | 14.67 | 1.17 | 0.53 |
| 1129.00 | 1365.00 | 1299.67 | 1.13 | 0.01 |
| 121.00 | 111.00 | 113.00 | 1.20 | 0.06 |
| 1131.00 | 1236.00 | 1211.67 | 1.14 | 0.00 |
| 235.00 | 278.00 | 268.67 | 1.45 | 0.00 |
| 190.00 | 231.00 | 212.33 | 1.26 | 0.00 |
| 697.00 | 706.00 | 712.00 | 1.00 | 0.97 |
| 657.00 | 723.00 | 729.00 | 1.14 | 0.01 |
| 343.00 | 394.00 | 384.33 | 1.22 | 0.00 |
| 5.00 | 36.00 | 22.67 | 1.68 | 0.18 |
| 21.00 | 81.00 | 55.33 | 1.54 | 0.10 |
| 160.00 | 374.00 | 278.33 | -1.22 | 0.41 |
| 145.00 | 400.00 | 282.00 | -1.56 | 0.10 |
| 2.00 | 10.00 | 6.00 | 1.93 | 0.13 |
| 11.00 | 12.00 | 11.67 | 1.11 | 0.72 |
| 30.00 | 23.00 | 28.33 | 1.52 | 0.02 |
| 735.00 | 694.00 | 702.33 | 1.05 | 0.41 |
| 282.00 | 294.00 | 306.67 | 1.21 | 0.01 |
| 780.00 | 990.00 | 904.67 | 1.28 | 0.00 |
| 445.00 | 467.00 | 493.33 | 1.16 | 0.01 |
| 6.00 | 13.00 | 7.33 | -1.10 | 0.89 |
| 1528.00 | 1544.00 | 1629.00 | -1.13 | 0.00 |
| 89.00 | 126.00 | 115.00 | 1.05 | 0.66 |
| 59.00 | 76.00 | 66.67 | 1.31 | 0.03 |
| 576.00 | 578.00 | 580.33 | 1.10 | 0.05 |
| 1.00 | 0.00 | 0.33 | -3.21 | 0.41 |
| 5933.00 | 6416.00 | 6463.67 | 1.22 | 0.00 |
| 1766.00 | 1751.00 | 1797.67 | -1.13 | 0.00 |
| 765.00 | 728.00 | 734.33 | -1.03 | 0.61 |

| | | | | |
|---|---|---|---|---|
| 838.00 | 1035.00 | 928.00 | -1.17 | 0.00 |
| 859.00 | 849.00 | 832.00 | -1.13 | 0.03 |
| 3.00 | 3.00 | 2.67 | -1.50 | 0.51 |
| 242.00 | 256.00 | 258.00 | 1.06 | 0.34 |
| 1890.00 | 2038.00 | 2089.00 | 1.14 | 0.00 |
| 360.00 | 367.00 | 372.00 | -1.20 | 0.00 |
| 1437.00 | 1406.00 | 1435.67 | -1.16 | 0.00 |
| 15.00 | 13.00 | 14.00 | 1.12 | 0.67 |
| 1392.00 | 1363.00 | 1393.00 | -1.06 | 0.14 |
| 204.00 | 205.00 | 214.67 | 1.41 | 0.00 |
| 1053.00 | 1077.00 | 1099.67 | 1.16 | 0.00 |
| 92.00 | 96.00 | 91.00 | -1.12 | 0.25 |
| 1197.00 | 1187.00 | 1188.33 | 1.12 | 0.02 |
| 750.00 | 678.00 | 706.67 | 1.06 | 0.27 |
| 0.00 | 1.00 | 1.33 | -1.48 | 0.76 |
| 628.00 | 704.00 | 706.67 | 1.12 | 0.02 |
| 444.00 | 595.00 | 503.00 | 1.38 | 0.00 |
| 1030.00 | 1206.00 | 1154.00 | -1.07 | 0.12 |
| 83.00 | 104.00 | 100.00 | 1.15 | 0.18 |
| 853.00 | 944.00 | 913.67 | 1.15 | 0.00 |
| 516.00 | 484.00 | 515.33 | 1.14 | 0.01 |
| 134.00 | 135.00 | 130.67 | 1.17 | 0.09 |
| 1615.00 | 1726.00 | 1698.33 | -1.03 | 0.48 |
| 1081.00 | 1345.00 | 1282.33 | -1.09 | 0.09 |
| 2907.00 | 3152.00 | 3065.33 | -1.01 | 0.73 |
| 58.00 | 43.00 | 45.00 | -1.11 | 0.53 |
| 18.00 | 28.00 | 18.67 | -1.06 | 0.88 |
| 113.00 | 102.00 | 101.33 | -1.24 | 0.02 |
| 962.00 | 913.00 | 975.00 | 1.02 | 0.62 |
| 1887.00 | 1905.00 | 1988.00 | -1.09 | 0.03 |
| 391.00 | 388.00 | 397.67 | -1.04 | 0.49 |
| 686.00 | 739.00 | 719.00 | -1.06 | 0.16 |
| 757.00 | 826.00 | 791.00 | 1.12 | 0.01 |
| 6.00 | 12.00 | 6.67 | -1.46 | 0.28 |
| 844.00 | 1028.00 | 990.00 | 1.04 | 0.44 |
| 162.00 | 183.00 | 174.33 | -1.06 | 0.39 |
| 652.00 | 620.00 | 675.00 | -1.08 | 0.13 |
| 1438.00 | 1489.00 | 1531.33 | -1.07 | 0.07 |
| 803.00 | 1017.00 | 906.00 | 1.04 | 0.46 |
| 552.00 | 645.00 | 600.67 | 1.09 | 0.08 |
| 591.00 | 670.00 | 640.67 | 1.37 | 0.00 |
| 1308.00 | 1316.00 | 1302.00 | -1.09 | 0.06 |
| 642.00 | 683.00 | 697.00 | -1.07 | 0.15 |
| 37.00 | 51.00 | 46.67 | 1.11 | 0.46 |
| 4955.00 | 4519.00 | 4652.00 | -1.30 | 0.00 |
| 487.00 | 453.00 | 466.33 | -1.07 | 0.24 |
| 962.00 | 975.00 | 1002.33 | 1.17 | 0.00 |
| 2556.00 | 2718.00 | 2710.67 | -1.12 | 0.00 |

| | | | | |
|---|---|---|---|---|
| 675.00 | 765.00 | 746.33 | 1.04 | 0.39 |
| 232.00 | 290.00 | 270.33 | -1.09 | 0.18 |
| 11.00 | 12.00 | 13.00 | 1.13 | 0.65 |
| 762.00 | 927.00 | 872.67 | 1.07 | 0.19 |
| 0.00 | 0.00 | 0.00 | -11.75 | 0.13 |
| 1284.00 | 1436.00 | 1415.33 | -1.29 | 0.00 |
| 469.00 | 383.00 | 406.33 | -1.26 | 0.00 |
| 454.00 | 567.00 | 550.00 | 1.06 | 0.35 |
| 1687.00 | 1661.00 | 1737.33 | -1.04 | 0.39 |
| 1075.00 | 1020.00 | 1078.00 | -1.07 | 0.10 |
| 121.00 | 128.00 | 131.00 | 1.15 | 0.09 |
| 576.00 | 610.00 | 636.33 | 1.06 | 0.29 |
| 450.00 | 463.00 | 457.33 | -1.00 | 0.98 |
| 747.00 | 714.00 | 747.33 | 1.08 | 0.09 |
| 2274.00 | 2311.00 | 2318.67 | -1.11 | 0.01 |
| 832.00 | 934.00 | 911.00 | -1.04 | 0.36 |
| 1630.00 | 1516.00 | 1613.67 | -1.05 | 0.28 |
| 16.00 | 23.00 | 21.00 | 1.11 | 0.66 |
| 976.00 | 1137.00 | 1069.00 | 1.07 | 0.14 |
| 1293.00 | 1352.00 | 1433.33 | 1.07 | 0.17 |
| 481.00 | 505.00 | 505.00 | 1.03 | 0.49 |
| 629.00 | 704.00 | 677.67 | 1.10 | 0.06 |
| 105.00 | 139.00 | 137.00 | 1.06 | 0.61 |
| 643.00 | 736.00 | 682.33 | 1.07 | 0.27 |
| 2.00 | 2.00 | 5.67 | -1.25 | 0.67 |
| 36.00 | 44.00 | 40.67 | -1.17 | 0.25 |
| 1853.00 | 1991.00 | 1933.67 | -1.14 | 0.00 |
| 358.00 | 390.00 | 393.00 | 1.08 | 0.19 |
| 591.00 | 680.00 | 635.67 | -1.09 | 0.08 |
| 1450.00 | 1584.00 | 1575.67 | 1.09 | 0.04 |
| 1021.00 | 992.00 | 1043.00 | -1.07 | 0.15 |
| 3050.00 | 3184.00 | 3264.00 | 1.18 | 0.00 |
| 2.00 | 0.00 | 1.33 | -1.94 | 0.39 |
| 1.00 | 3.00 | 2.67 | -1.01 | 1.00 |
| 41.00 | 37.00 | 32.00 | -1.60 | 0.01 |
| 56.00 | 76.00 | 69.67 | -1.05 | 0.63 |
| 1459.00 | 1420.00 | 1397.67 | -1.07 | 0.19 |
| 87.00 | 96.00 | 84.67 | 1.04 | 0.74 |
| 448.00 | 433.00 | 490.00 | 1.16 | 0.04 |
| 1692.00 | 1744.00 | 1758.67 | -1.04 | 0.29 |
| 1627.00 | 1767.00 | 1628.67 | 1.04 | 0.52 |
| 36.00 | 40.00 | 37.33 | -1.05 | 0.81 |
| 1137.00 | 1090.00 | 1064.33 | -1.13 | 0.04 |
| 88.00 | 81.00 | 93.67 | -1.09 | 0.40 |
| 65.00 | 71.00 | 68.00 | -1.29 | 0.02 |
| 660.00 | 709.00 | 736.00 | 1.11 | 0.09 |
| 21.00 | 21.00 | 21.00 | -1.36 | 0.10 |
| 167.00 | 232.00 | 195.67 | 1.30 | 0.00 |

| | | | | |
|---|---|---|---|---|
| 22.00 | 31.00 | 26.00 | 1.04 | 0.88 |
| 25.00 | 8.00 | 13.00 | 1.15 | 0.72 |
| 88.00 | 69.00 | 81.67 | -2.41 | 0.00 |
| 16.00 | 6.00 | 10.67 | -1.20 | 0.57 |
| 25.00 | 49.00 | 38.00 | -1.37 | 0.03 |
| 233.00 | 296.00 | 266.33 | 1.08 | 0.25 |
| 281.00 | 315.00 | 333.67 | 1.11 | 0.15 |
| 100.00 | 105.00 | 99.67 | 1.33 | 0.01 |
| 1540.00 | 1535.00 | 1569.33 | 1.15 | 0.00 |
| 666.00 | 829.00 | 788.33 | 1.16 | 0.01 |
| 581.00 | 662.00 | 659.67 | 1.21 | 0.00 |
| 12.00 | 19.00 | 14.00 | 2.16 | 0.01 |
| 1228.00 | 1347.00 | 1357.00 | -1.02 | 0.65 |
| 1379.00 | 1452.00 | 1438.33 | -1.17 | 0.00 |
| 487.00 | 462.00 | 486.67 | 1.04 | 0.45 |
| 825.00 | 929.00 | 935.67 | 1.10 | 0.05 |
| 236.00 | 230.00 | 259.00 | 1.06 | 0.39 |
| 1334.00 | 1437.00 | 1426.00 | 1.04 | 0.36 |
| 17.00 | 33.00 | 25.67 | 1.24 | 0.31 |
| 1122.00 | 1272.00 | 1239.67 | 1.20 | 0.00 |
| 1462.00 | 1650.00 | 1620.00 | 1.18 | 0.00 |
| 296.00 | 297.00 | 299.33 | 1.46 | 0.00 |
| 0.00 | 2.00 | 1.00 | -4.01 | 0.02 |
| 442.00 | 498.00 | 476.00 | -1.09 | 0.09 |
| 956.00 | 1025.00 | 1014.00 | 1.28 | 0.00 |
| 17200.00 | 17532.00 | 19064.67 | 1.09 | 0.06 |
| 12445.00 | 13641.00 | 14047.00 | 1.17 | 0.00 |
| 17272.00 | 19342.00 | 19428.67 | 1.15 | 0.00 |
| 19810.00 | 21770.00 | 22511.33 | 1.18 | 0.00 |
| 26781.00 | 25896.00 | 27512.00 | 1.15 | 0.00 |
| 29638.00 | 31874.00 | 32589.00 | 1.11 | 0.00 |
| 72.00 | 57.00 | 70.00 | -1.06 | 0.62 |
| 2270.00 | 2216.00 | 2537.67 | 1.15 | 0.03 |
| 0.00 | 0.00 | 1.00 | -1.32 | 1.00 |
| 48.00 | 56.00 | 48.00 | 1.19 | 0.21 |
| 9475.00 | 11028.00 | 10990.33 | 1.13 | 0.00 |
| 16476.00 | 18147.00 | 18666.33 | 1.10 | 0.03 |
| 1078.00 | 1216.00 | 1213.67 | 1.11 | 0.02 |
| 11357.00 | 12153.00 | 12668.33 | 1.11 | 0.02 |
| 7455.00 | 7568.00 | 7994.33 | 1.15 | 0.00 |
| 22660.00 | 25559.00 | 25622.67 | 1.13 | 0.00 |
| 36.00 | 45.00 | 38.33 | 1.29 | 0.11 |
| 694.00 | 792.00 | 827.33 | 1.14 | 0.04 |
| 7.00 | 6.00 | 6.33 | -1.43 | 0.31 |
| 5606.00 | 6549.00 | 6420.67 | 1.19 | 0.00 |
| 3790.00 | 4288.00 | 4419.67 | 1.12 | 0.03 |
| 14220.00 | 15931.00 | 16667.33 | 1.11 | 0.04 |
| 7305.00 | 8028.00 | 8433.33 | 1.08 | 0.14 |

| | | | | |
|---|---|---|---|---|
| 86.00 | 116.00 | 102.67 | -1.11 | 0.27 |
| 147.00 | 198.00 | 174.67 | 1.02 | 0.87 |
| 132.00 | 105.00 | 121.67 | 1.11 | 0.28 |
| 74.00 | 74.00 | 80.67 | 1.19 | 0.10 |
| 7055.00 | 7706.00 | 7949.33 | 1.17 | 0.00 |
| 21274.00 | 23529.00 | 24264.67 | 1.18 | 0.00 |
| 797.00 | 1023.00 | 985.33 | 1.30 | 0.00 |
| 12879.00 | 14804.00 | 15048.67 | 1.17 | 0.00 |
| 13547.00 | 14904.00 | 15298.33 | 1.09 | 0.03 |
| 7596.00 | 8075.00 | 8554.00 | 1.01 | 0.80 |
| 11248.00 | 12220.00 | 12668.33 | 1.11 | 0.01 |
| 39756.00 | 43586.00 | 44583.33 | 1.09 | 0.04 |
| 14989.00 | 16341.00 | 16903.67 | 1.12 | 0.01 |
| 9731.00 | 10993.00 | 11668.00 | 1.15 | 0.02 |
| 11262.00 | 12557.00 | 13073.00 | 1.17 | 0.00 |
| 412.00 | 339.00 | 367.33 | -1.00 | 0.97 |
| 4904.00 | 5343.00 | 5494.00 | 1.13 | 0.00 |
| 8.00 | 14.00 | 12.67 | 1.69 | 0.06 |
| 7172.00 | 7838.00 | 8024.67 | 1.20 | 0.00 |
| 10239.00 | 11118.00 | 11484.33 | 1.21 | 0.00 |
| 7854.00 | 8286.00 | 8708.67 | 1.17 | 0.00 |
| 391.00 | 401.00 | 407.67 | -1.06 | 0.28 |
| 1415.00 | 1676.00 | 1669.33 | 1.03 | 0.55 |
| 19051.00 | 21177.00 | 22264.67 | 1.17 | 0.00 |
| 18781.00 | 20513.00 | 21621.33 | 1.14 | 0.01 |
| 5315.00 | 6351.00 | 6440.00 | 1.26 | 0.00 |
| 3961.00 | 4498.00 | 4637.67 | 1.11 | 0.04 |
| 771.00 | 707.00 | 770.67 | 1.41 | 0.00 |
| 31148.00 | 34068.00 | 34880.00 | 1.11 | 0.01 |
| 1038.00 | 1225.00 | 1285.00 | 1.16 | 0.03 |
| 28320.00 | 32064.00 | 32465.67 | 1.16 | 0.00 |
| 17027.00 | 19288.00 | 19217.00 | 1.19 | 0.00 |
| 11321.00 | 13360.00 | 13292.00 | 1.21 | 0.00 |
| 24896.00 | 27765.00 | 28138.00 | 1.13 | 0.00 |
| 1560.00 | 1444.00 | 1471.33 | 1.15 | 0.01 |
| 35761.00 | 38701.00 | 39907.00 | 1.21 | 0.00 |
| 11697.00 | 12795.00 | 13170.33 | 1.17 | 0.00 |
| 34907.00 | 37027.00 | 37910.00 | 1.13 | 0.00 |
| 58.00 | 45.00 | 55.33 | -1.23 | 0.10 |
| 22256.00 | 25555.00 | 25739.67 | 1.14 | 0.00 |
| 11119.00 | 12131.00 | 12500.33 | 1.17 | 0.00 |
| 8462.00 | 8888.00 | 8829.67 | -1.03 | 0.43 |
| 13928.00 | 14025.00 | 13803.00 | -1.03 | 0.45 |
| 634.00 | 661.00 | 654.67 | 1.01 | 0.79 |
| 34.00 | 33.00 | 37.33 | 1.57 | 0.01 |
| 480.00 | 484.00 | 532.67 | 1.08 | 0.23 |
| 1020.00 | 1209.00 | 1164.33 | 1.11 | 0.02 |
| 301.00 | 360.00 | 363.33 | 1.23 | 0.00 |

| | | | | |
|---|---|---|---|---|
| 332.00 | 399.00 | 368.00 | 1.19 | 0.00 |
| 4.00 | 25.00 | 16.33 | 1.15 | 0.77 |
| 1307.00 | 1526.00 | 1451.33 | 1.11 | 0.01 |
| 1065.00 | 1214.00 | 1215.00 | -1.10 | 0.04 |
| 1364.00 | 1240.00 | 1316.33 | -1.12 | 0.02 |
| 7.00 | 4.00 | 4.67 | -1.15 | 0.86 |
| 818.00 | 881.00 | 902.00 | 1.25 | 0.00 |
| 65.00 | 77.00 | 78.67 | 1.04 | 0.77 |
| 11784.00 | 13109.00 | 13782.00 | 1.11 | 0.04 |
| 18181.00 | 19862.00 | 20599.00 | 1.14 | 0.00 |
| 8658.00 | 9729.00 | 10009.67 | 1.13 | 0.01 |
| 14776.00 | 16567.00 | 17011.33 | 1.16 | 0.00 |
| 46.00 | 57.00 | 53.00 | -1.08 | 0.55 |
| 4394.00 | 4612.00 | 4845.67 | 1.11 | 0.01 |
| 10363.00 | 11861.00 | 12195.67 | 1.12 | 0.02 |
| 8.00 | 11.00 | 8.33 | -1.57 | 0.11 |
| 10790.00 | 11800.00 | 12176.00 | 1.09 | 0.04 |
| 14.00 | 11.00 | 13.33 | -1.24 | 0.41 |
| 165.00 | 199.00 | 197.67 | 1.20 | 0.02 |
| 76.00 | 87.00 | 97.00 | 1.04 | 0.78 |
| 13714.00 | 14436.00 | 14851.33 | 1.09 | 0.02 |
| 1123.00 | 1260.00 | 1232.33 | 1.10 | 0.03 |
| 30829.00 | 34299.00 | 34743.67 | 1.31 | 0.00 |
| 14385.00 | 16347.00 | 16704.00 | 1.15 | 0.00 |
| 8360.00 | 9036.00 | 9718.00 | 1.07 | 0.20 |
| 13522.00 | 15299.00 | 15731.67 | 1.20 | 0.00 |
| 14231.00 | 16244.00 | 16202.00 | 1.25 | 0.00 |
| 9010.00 | 10631.00 | 10848.67 | 1.15 | 0.01 |
| 193.00 | 229.00 | 241.67 | 1.12 | 0.19 |
| 5673.00 | 5985.00 | 6395.67 | 1.08 | 0.12 |
| 11059.00 | 12692.00 | 12811.33 | 1.15 | 0.00 |
| 851.00 | 1008.00 | 1016.67 | 1.11 | 0.07 |
| 2108.00 | 2293.00 | 2328.67 | 1.12 | 0.00 |
| 6031.00 | 5931.00 | 6791.00 | 1.14 | 0.04 |
| 15.00 | 33.00 | 21.33 | -1.22 | 0.37 |
| 38390.00 | 41151.00 | 42877.67 | 1.14 | 0.00 |
| 8709.00 | 9982.00 | 9920.67 | 1.17 | 0.00 |
| 37782.00 | 41498.00 | 42273.00 | 1.18 | 0.00 |
| 10099.00 | 10624.00 | 11336.67 | 1.02 | 0.66 |
| 33401.00 | 36020.00 | 35524.67 | 1.19 | 0.00 |
| 356.00 | 375.00 | 372.00 | 1.01 | 0.90 |
| 4719.00 | 4281.00 | 4543.33 | 1.08 | 0.07 |
| 18.00 | 11.00 | 15.67 | -1.01 | 1.00 |
| 1058.00 | 1217.00 | 1142.33 | -1.11 | 0.01 |
| 2089.00 | 2065.00 | 2118.67 | 1.10 | 0.01 |
| 66.00 | 81.00 | 68.67 | -1.09 | 0.47 |
| 101.00 | 137.00 | 121.67 | -1.20 | 0.04 |
| 784.00 | 943.00 | 871.67 | -1.06 | 0.19 |

| | | | | |
|---|---|---|---|---|
| 555.00 | 566.00 | 612.00 | 1.02 | 0.79 |
| 560.00 | 585.00 | 581.00 | -1.14 | 0.00 |
| 123.00 | 129.00 | 122.67 | -1.09 | 0.36 |
| 15681.00 | 16619.00 | 17501.67 | 1.18 | 0.00 |
| 19495.00 | 21442.00 | 21932.00 | 1.14 | 0.00 |
| 378.00 | 429.00 | 413.33 | 1.14 | 0.02 |
| 13398.00 | 14148.00 | 14812.67 | 1.11 | 0.01 |
| 27.00 | 37.00 | 35.33 | -1.67 | 0.00 |
| 1970.00 | 2099.00 | 2230.67 | 1.13 | 0.02 |
| 33.00 | 56.00 | 49.33 | -1.06 | 0.66 |
| 1756.00 | 1645.00 | 1676.67 | -1.06 | 0.23 |
| 948.00 | 1052.00 | 1047.33 | -1.01 | 0.89 |
| 322.00 | 377.00 | 356.33 | 1.02 | 0.81 |
| 699.00 | 744.00 | 753.67 | 1.07 | 0.15 |
| 642.00 | 706.00 | 721.00 | 1.04 | 0.41 |
| 1751.00 | 1947.00 | 1968.00 | 1.21 | 0.00 |
| 10.00 | 7.00 | 10.00 | -1.28 | 0.42 |
| 880.00 | 884.00 | 906.00 | 1.13 | 0.00 |
| 276.00 | 323.00 | 301.33 | -1.17 | 0.01 |
| 670.00 | 596.00 | 640.67 | -1.11 | 0.05 |
| 11.00 | 8.00 | 11.00 | -1.01 | 1.00 |
| 310.00 | 328.00 | 349.00 | 1.02 | 0.77 |
| 1528.00 | 1841.00 | 1717.67 | 1.25 | 0.00 |
| 6165.00 | 6615.00 | 6441.00 | -1.34 | 0.00 |
| 925.00 | 954.00 | 936.67 | -1.19 | 0.00 |
| 4.00 | 4.00 | 4.00 | 1.18 | 0.83 |
| 1926.00 | 2073.00 | 2083.00 | 1.43 | 0.00 |
| 1922.00 | 2511.00 | 2279.67 | 1.70 | 0.00 |
| 596.00 | 615.00 | 612.00 | 1.18 | 0.00 |
| 19.00 | 12.00 | 14.67 | 1.60 | 0.07 |
| 240.00 | 312.00 | 290.67 | -1.11 | 0.14 |
| 22.00 | 12.00 | 15.33 | -1.51 | 0.05 |
| 111.00 | 88.00 | 105.33 | -1.02 | 0.84 |
| 946.00 | 1038.00 | 1030.33 | -1.10 | 0.02 |
| 1168.00 | 1328.00 | 1279.00 | 1.09 | 0.09 |
| 3058.00 | 2971.00 | 3013.33 | -1.06 | 0.12 |
| 876.00 | 1104.00 | 1039.67 | 1.03 | 0.52 |
| 2275.00 | 2322.00 | 2277.33 | -1.00 | 0.94 |
| 784.00 | 879.00 | 866.33 | 1.09 | 0.06 |
| 737.00 | 748.00 | 766.33 | 1.09 | 0.07 |
| 176.00 | 182.00 | 185.00 | -1.27 | 0.00 |
| 395.00 | 405.00 | 414.00 | 1.02 | 0.77 |
| 1204.00 | 1340.00 | 1307.00 | 1.04 | 0.34 |
| 1888.00 | 2000.00 | 2031.33 | 1.11 | 0.01 |
| 2.00 | 4.00 | 3.33 | -1.31 | 0.68 |
| 2800.00 | 3042.00 | 3031.67 | -1.00 | 0.97 |
| 434.00 | 464.00 | 426.00 | 1.02 | 0.79 |
| 400.00 | 417.00 | 409.00 | 1.03 | 0.55 |

| | | | | |
|---|---|---|---|---|
| 58.00 | 45.00 | 47.67 | -1.56 | 0.00 |
| 655.00 | 739.00 | 677.67 | -1.05 | 0.48 |
| 97.00 | 74.00 | 92.67 | -1.08 | 0.50 |
| 4452.00 | 4895.00 | 4822.67 | 1.12 | 0.00 |
| 3492.00 | 4026.00 | 3969.33 | 1.24 | 0.00 |
| 86.00 | 79.00 | 80.33 | -1.33 | 0.00 |
| 5.00 | 6.00 | 5.67 | -1.13 | 0.87 |
| 16.00 | 15.00 | 14.33 | 1.36 | 0.22 |
| 1489.00 | 1587.00 | 1600.67 | 1.25 | 0.00 |
| 414.00 | 438.00 | 441.67 | 1.12 | 0.03 |
| 1490.00 | 1812.00 | 1754.33 | 1.08 | 0.12 |
| 1388.00 | 1538.00 | 1538.67 | -1.16 | 0.00 |
| 1943.00 | 2074.00 | 2091.33 | 1.13 | 0.00 |
| 1495.00 | 1671.00 | 1634.67 | 1.19 | 0.00 |
| 18.00 | 16.00 | 18.33 | 1.18 | 0.44 |
| 1791.00 | 1962.00 | 1942.00 | 1.11 | 0.00 |
| 156.00 | 144.00 | 148.00 | -1.03 | 0.76 |
| 6.00 | 19.00 | 14.00 | 2.05 | 0.01 |
| 975.00 | 995.00 | 1009.67 | 1.01 | 0.80 |
| 2131.00 | 2427.00 | 2428.67 | 1.12 | 0.01 |
| 336.00 | 398.00 | 390.00 | 1.06 | 0.37 |
| 156.00 | 197.00 | 183.00 | 1.23 | 0.01 |
| 680.00 | 644.00 | 658.67 | 1.63 | 0.00 |
| 88.00 | 80.00 | 90.33 | -1.51 | 0.00 |
| 6502.00 | 6392.00 | 6638.33 | 1.03 | 0.45 |
| 26566.00 | 26351.00 | 27220.33 | 1.11 | 0.01 |
| 247.00 | 340.00 | 296.67 | 1.23 | 0.01 |
| 48.00 | 29.00 | 38.00 | 1.21 | 0.24 |
| 11.00 | 3.00 | 5.67 | 1.29 | 0.60 |
| 360.00 | 423.00 | 400.67 | -1.07 | 0.26 |
| 692.00 | 722.00 | 746.00 | -1.09 | 0.07 |
| 490.00 | 550.00 | 529.33 | -1.20 | 0.00 |
| 842.00 | 1001.00 | 959.33 | -1.30 | 0.00 |
| 380.00 | 364.00 | 376.00 | -1.16 | 0.01 |
| 9.00 | 8.00 | 8.67 | 1.22 | 0.57 |
| 26.00 | 46.00 | 35.33 | 1.36 | 0.08 |
| 2026.00 | 2170.00 | 2106.33 | -1.02 | 0.66 |
| 2.00 | 5.00 | 4.00 | 1.46 | 0.51 |
| 327.00 | 391.00 | 347.00 | 1.57 | 0.00 |
| 864.00 | 920.00 | 914.33 | -1.22 | 0.00 |
| 583.00 | 604.00 | 622.67 | -1.13 | 0.02 |
| 12.00 | 16.00 | 15.67 | -1.68 | 0.01 |
| 2665.00 | 2578.00 | 2685.33 | -1.44 | 0.00 |
| 2748.00 | 3202.00 | 3071.00 | 1.13 | 0.00 |
| 2581.00 | 2835.00 | 2786.67 | 1.19 | 0.00 |
| 807.00 | 925.00 | 906.33 | 1.13 | 0.01 |
| 216.00 | 256.00 | 250.67 | -1.09 | 0.20 |
| 601.00 | 592.00 | 634.67 | 1.25 | 0.00 |

| | | | | |
|---|---|---|---|---|
| 291.00 | 342.00 | 324.00 | 1.11 | 0.18 |
| 239.00 | 311.00 | 271.33 | 1.24 | 0.00 |
| 158.00 | 178.00 | 169.33 | 1.11 | 0.18 |
| 340.00 | 286.00 | 325.67 | -1.07 | 0.31 |
| 4.00 | 7.00 | 5.67 | -1.36 | 0.46 |
| 627.00 | 664.00 | 680.33 | -1.01 | 0.87 |
| 2640.00 | 2798.00 | 2753.33 | 1.16 | 0.00 |
| 71.00 | 74.00 | 74.00 | -1.21 | 0.06 |
| 277.00 | 310.00 | 327.00 | -1.17 | 0.03 |
| 1783.00 | 1951.00 | 2001.00 | 1.09 | 0.05 |
| 1101.00 | 1253.00 | 1269.33 | 1.13 | 0.01 |
| 7.00 | 7.00 | 8.33 | -1.05 | 1.00 |
| 3760.00 | 3844.00 | 4134.67 | 1.15 | 0.00 |
| 52.00 | 52.00 | 56.33 | -1.25 | 0.05 |
| 3.00 | 5.00 | 4.67 | -1.58 | 0.25 |
| 3.00 | 2.00 | 3.67 | -1.01 | 1.00 |
| 0.00 | 2.00 | 2.33 | -1.15 | 1.00 |
| 3830.00 | 4120.00 | 4278.00 | -1.24 | 0.00 |
| 8.00 | 13.00 | 13.33 | 1.35 | 0.30 |
| 711.00 | 797.00 | 790.00 | -1.13 | 0.01 |
| 218.00 | 242.00 | 231.00 | 1.03 | 0.77 |
| 80.00 | 82.00 | 80.00 | -1.25 | 0.02 |
| 567.00 | 598.00 | 583.33 | -1.17 | 0.00 |
| 584.00 | 667.00 | 637.00 | 1.21 | 0.00 |
| 329.00 | 332.00 | 346.00 | 1.28 | 0.00 |
| 1154.00 | 1239.00 | 1221.67 | 1.08 | 0.06 |
| 3775.00 | 4422.00 | 4028.33 | -1.19 | 0.00 |
| 2319.00 | 2550.00 | 2520.67 | 1.14 | 0.00 |
| 2923.00 | 3175.00 | 3096.00 | 1.05 | 0.18 |
| 1470.00 | 1589.00 | 1535.33 | 1.00 | 0.99 |
| 69.00 | 83.00 | 76.00 | -1.19 | 0.12 |
| 408.00 | 438.00 | 440.00 | 1.04 | 0.44 |
| 257.00 | 225.00 | 255.00 | 1.11 | 0.24 |
| 239.00 | 234.00 | 237.00 | -1.29 | 0.00 |
| 38.00 | 52.00 | 44.00 | 1.10 | 0.52 |
| 17.00 | 14.00 | 18.67 | 1.02 | 1.00 |
| 434.00 | 544.00 | 511.33 | -1.06 | 0.28 |
| 3011.00 | 3216.00 | 3166.00 | 1.08 | 0.02 |
| 1992.00 | 1901.00 | 1924.33 | -1.17 | 0.00 |
| 166.00 | 142.00 | 153.67 | 1.06 | 0.45 |
| 1374.00 | 1429.00 | 1407.67 | -1.07 | 0.10 |
| 561.00 | 604.00 | 555.67 | -1.03 | 0.62 |
| 262.00 | 285.00 | 284.00 | -1.31 | 0.00 |
| 527.00 | 572.00 | 540.33 | -1.01 | 0.91 |
| 771.00 | 817.00 | 821.00 | 1.11 | 0.02 |
| 7.00 | 12.00 | 10.00 | 1.18 | 0.60 |
| 644.00 | 637.00 | 649.00 | -1.03 | 0.53 |
| 1231.00 | 1513.00 | 1489.33 | 1.15 | 0.02 |

| | | | | |
|---|---|---|---|---|
| 72.00 | 83.00 | 87.33 | -1.46 | 0.00 |
| 194.00 | 206.00 | 205.67 | 1.33 | 0.00 |
| 1046.00 | 1070.00 | 1100.67 | 1.07 | 0.14 |
| 563.00 | 510.00 | 525.67 | -1.08 | 0.25 |
| 9.00 | 13.00 | 11.00 | -1.53 | 0.09 |
| 672.00 | 660.00 | 671.33 | 1.07 | 0.27 |
| 1923.00 | 2223.00 | 2186.67 | 1.09 | 0.03 |
| 953.00 | 991.00 | 1019.67 | 1.08 | 0.08 |
| 1732.00 | 1783.00 | 1761.33 | -1.08 | 0.04 |
| 63.00 | 73.00 | 65.33 | -1.20 | 0.12 |
| 789.00 | 777.00 | 802.00 | -1.02 | 0.66 |
| 574.00 | 727.00 | 691.33 | 1.11 | 0.07 |
| 9456.00 | 10003.00 | 10144.33 | 1.04 | 0.23 |
| 391.00 | 450.00 | 445.33 | 1.07 | 0.24 |
| 3383.00 | 3597.00 | 3581.67 | 1.11 | 0.01 |
| 1414.00 | 1605.00 | 1504.67 | 1.04 | 0.40 |
| 2800.00 | 2981.00 | 2832.00 | 1.29 | 0.00 |
| 512.00 | 579.00 | 536.00 | 1.15 | 0.01 |
| 317.00 | 359.00 | 349.00 | -1.27 | 0.00 |
| 790.00 | 736.00 | 829.00 | -1.10 | 0.08 |
| 588.00 | 540.00 | 562.67 | -1.06 | 0.30 |
| 3.00 | 7.00 | 6.67 | 1.50 | 0.31 |
| 876.00 | 867.00 | 925.33 | -1.14 | 0.00 |
| 287.00 | 320.00 | 304.67 | 1.08 | 0.20 |
| 13.00 | 14.00 | 11.67 | 1.15 | 0.63 |
| 313.00 | 331.00 | 339.00 | 1.03 | 0.68 |
| 1625.00 | 1975.00 | 1847.33 | 1.00 | 0.96 |
| 623.00 | 768.00 | 742.67 | 1.13 | 0.03 |
| 2143.00 | 2021.00 | 2117.00 | -1.17 | 0.00 |
| 1190.00 | 1094.00 | 1135.67 | -1.14 | 0.01 |
| 64.00 | 66.00 | 69.00 | -1.16 | 0.16 |
| 6.00 | 7.00 | 6.67 | 1.16 | 0.75 |
| 1592.00 | 1691.00 | 1643.33 | 1.06 | 0.15 |
| 53.00 | 37.00 | 42.33 | -1.39 | 0.02 |
| 242.00 | 228.00 | 231.67 | -1.18 | 0.03 |
| 726.00 | 796.00 | 757.00 | 1.18 | 0.00 |
| 1407.00 | 1402.00 | 1425.67 | -1.12 | 0.00 |
| 2235.00 | 2560.00 | 2379.67 | -1.08 | 0.05 |
| 1158.00 | 1257.00 | 1237.00 | -1.06 | 0.15 |
| 1754.00 | 1948.00 | 1899.67 | 1.02 | 0.52 |
| 255.00 | 296.00 | 273.33 | -1.14 | 0.05 |
| 1049.00 | 1104.00 | 1004.67 | 1.23 | 0.00 |
| 183.00 | 197.00 | 188.33 | 1.12 | 0.16 |
| 1249.00 | 1146.00 | 1213.67 | -1.12 | 0.02 |
| 1717.00 | 1482.00 | 1663.33 | -1.12 | 0.04 |
| 62.00 | 70.00 | 72.00 | 1.46 | 0.00 |
| 664.00 | 670.00 | 684.00 | 1.24 | 0.00 |
| 197.00 | 187.00 | 194.67 | -1.04 | 0.62 |

| | | | | |
|---|---|---|---|---|
| 2492.00 | 2528.00 | 2569.67 | -1.19 | 0.00 |
| 227.00 | 234.00 | 245.67 | 1.06 | 0.41 |
| 7.00 | 12.00 | 11.33 | -1.96 | 0.00 |
| 2333.00 | 2387.00 | 2398.33 | -1.02 | 0.63 |
| 4612.00 | 4469.00 | 4607.00 | 1.06 | 0.14 |
| 782.00 | 758.00 | 779.00 | -1.21 | 0.00 |
| 2.00 | 0.00 | 0.67 | -2.29 | 0.46 |
| 124.00 | 124.00 | 133.00 | -1.05 | 0.55 |
| 26.00 | 38.00 | 32.67 | -1.00 | 1.00 |
| 11.00 | 19.00 | 15.00 | -1.24 | 0.38 |
| 2.00 | 1.00 | 3.00 | -1.45 | 0.55 |
| 0.00 | 3.00 | 2.67 | 1.30 | 0.79 |
| 10.00 | 12.00 | 13.33 | -1.39 | 0.13 |
| 28.00 | 33.00 | 27.67 | -1.37 | 0.05 |
| 4.00 | 0.00 | 3.00 | -1.12 | 1.00 |
| 4.00 | 0.00 | 2.67 | -1.38 | 0.65 |
| 1.00 | 3.00 | 3.00 | 1.10 | 1.00 |
| 1.00 | 1.00 | 2.67 | 1.29 | 0.80 |
| 9.00 | 5.00 | 7.00 | 1.21 | 0.63 |
| 231.00 | 190.00 | 219.00 | 1.02 | 0.80 |
| 1439.00 | 1504.00 | 1495.33 | 1.22 | 0.00 |
| 10920.00 | 10336.00 | 10661.00 | 1.15 | 0.00 |
| 229.00 | 204.00 | 207.33 | 1.10 | 0.28 |
| 21.00 | 19.00 | 20.33 | 1.00 | 1.00 |
| 928.00 | 1154.00 | 1068.67 | 1.12 | 0.04 |
| 405.00 | 447.00 | 449.33 | -1.03 | 0.67 |
| 31.00 | 43.00 | 42.33 | -1.12 | 0.40 |
| 7.00 | 10.00 | 6.33 | 1.85 | 0.15 |
| 377.00 | 499.00 | 424.67 | 1.10 | 0.19 |
| 18.00 | 20.00 | 19.67 | 1.02 | 0.93 |
| 1333.00 | 1317.00 | 1369.00 | -1.03 | 0.50 |
| 317.00 | 368.00 | 358.00 | 1.25 | 0.00 |
| 174.00 | 171.00 | 169.00 | 1.13 | 0.11 |
| 1.00 | 2.00 | 1.00 | -1.01 | 1.00 |
| 2.00 | 6.00 | 4.00 | -1.75 | 0.17 |
| 38.00 | 38.00 | 40.67 | 1.32 | 0.07 |
| 5.00 | 4.00 | 4.33 | -1.32 | 0.59 |
| 172.00 | 170.00 | 162.67 | -2.15 | 0.00 |
| 3.00 | 3.00 | 2.67 | -2.95 | 0.01 |
| 699.00 | 719.00 | 702.67 | -1.44 | 0.00 |
| 22.00 | 16.00 | 18.00 | -1.48 | 0.04 |
| 86.00 | 96.00 | 101.33 | 1.13 | 0.24 |
| 14.00 | 18.00 | 16.00 | 1.01 | 1.00 |
| 390.00 | 410.00 | 428.00 | -1.29 | 0.00 |
| 814.00 | 864.00 | 872.67 | 1.05 | 0.32 |
| 269.00 | 255.00 | 269.00 | -1.10 | 0.12 |
| 865.00 | 1091.00 | 1009.67 | 1.20 | 0.00 |
| 12.00 | 25.00 | 20.00 | 2.17 | 0.00 |

| | | | | |
|---|---|---|---|---|
| 1009.00 | 1018.00 | 1036.00 | 1.14 | 0.00 |
| 1252.00 | 1271.00 | 1331.67 | -1.03 | 0.51 |
| 6.00 | 15.00 | 12.33 | 1.50 | 0.23 |
| 8158.00 | 8157.00 | 8260.00 | -1.06 | 0.10 |
| 633.00 | 679.00 | 700.33 | -1.01 | 0.88 |
| 546.00 | 601.00 | 578.33 | 1.08 | 0.10 |
| 9.00 | 4.00 | 7.67 | 1.19 | 0.65 |
| 57.00 | 54.00 | 55.00 | 1.26 | 0.07 |
| 76.00 | 91.00 | 94.33 | -1.04 | 0.73 |
| 2906.00 | 3100.00 | 3125.67 | -1.02 | 0.54 |
| 845.00 | 892.00 | 860.00 | 1.10 | 0.05 |
| 307.00 | 385.00 | 355.33 | -1.05 | 0.49 |
| 1107.00 | 1252.00 | 1231.33 | 1.09 | 0.04 |
| 8781.00 | 8620.00 | 8769.00 | 1.49 | 0.00 |
| 49.00 | 55.00 | 55.67 | -1.68 | 0.00 |
| 1025.00 | 976.00 | 1019.67 | -1.21 | 0.00 |
| 1237.00 | 1211.00 | 1276.00 | 1.31 | 0.00 |
| 2906.00 | 3232.00 | 3134.00 | 1.01 | 0.75 |
| 107.00 | 108.00 | 112.33 | 1.32 | 0.02 |
| 2674.00 | 2946.00 | 2963.67 | 1.07 | 0.08 |
| 29.00 | 19.00 | 23.33 | 1.46 | 0.17 |
| 636.00 | 744.00 | 655.00 | 1.11 | 0.11 |
| 709.00 | 719.00 | 728.67 | -1.01 | 0.90 |
| 378.00 | 436.00 | 428.33 | 1.18 | 0.01 |
| 4659.00 | 4456.00 | 4667.33 | -1.01 | 0.87 |
| 2466.00 | 2844.00 | 2664.33 | 1.06 | 0.12 |
| 172.00 | 172.00 | 172.33 | 1.06 | 0.41 |
| 502.00 | 581.00 | 580.33 | 1.09 | 0.13 |
| 143.00 | 175.00 | 172.00 | 1.19 | 0.03 |
| 180.00 | 181.00 | 197.67 | 1.28 | 0.00 |
| 331.00 | 323.00 | 326.00 | -1.28 | 0.00 |
| 4.00 | 2.00 | 2.00 | -1.64 | 0.51 |
| 19.00 | 12.00 | 14.67 | -1.38 | 0.16 |
| 1264.00 | 1415.00 | 1401.67 | 1.19 | 0.00 |
| 1203.00 | 1132.00 | 1213.33 | -1.06 | 0.22 |
| 841.00 | 918.00 | 944.00 | -1.35 | 0.00 |
| 21.00 | 17.00 | 19.33 | -1.22 | 0.35 |
| 619.00 | 639.00 | 688.00 | 1.01 | 0.83 |
| 170.00 | 145.00 | 162.67 | -1.02 | 0.88 |
| 3334.00 | 3361.00 | 3567.33 | 1.14 | 0.00 |
| 487.00 | 569.00 | 548.00 | 1.21 | 0.00 |
| 2117.00 | 2101.00 | 2201.00 | 1.01 | 0.72 |
| 2210.00 | 2152.00 | 2229.00 | -1.19 | 0.00 |
| 40.00 | 55.00 | 52.00 | -1.05 | 0.68 |
| 248.00 | 217.00 | 254.33 | -1.09 | 0.25 |
| 4246.00 | 4090.00 | 4110.67 | -1.21 | 0.00 |
| 119.00 | 124.00 | 135.67 | -1.10 | 0.28 |
| 554.00 | 506.00 | 541.67 | 1.08 | 0.17 |

| | | | | |
|---|---|---|---|---|
| 1149.00 | 1299.00 | 1279.33 | -1.29 | 0.00 |
| 1155.00 | 1322.00 | 1241.33 | -1.06 | 0.16 |
| 1566.00 | 1783.00 | 1656.00 | 1.01 | 0.81 |
| 1742.00 | 1829.00 | 1806.33 | 1.03 | 0.49 |
| 1849.00 | 1858.00 | 1822.33 | 1.05 | 0.31 |
| 701.00 | 779.00 | 710.00 | -1.30 | 0.00 |
| 726.00 | 812.00 | 810.67 | -1.09 | 0.10 |
| 16.00 | 23.00 | 17.33 | -1.29 | 0.25 |
| 3870.00 | 3778.00 | 3886.00 | -1.17 | 0.00 |
| 921.00 | 1042.00 | 986.33 | -1.28 | 0.00 |
| 2186.00 | 2145.00 | 2208.67 | -1.18 | 0.00 |
| 707.00 | 644.00 | 704.33 | -1.42 | 0.00 |
| 15395.00 | 15258.00 | 15425.67 | -1.20 | 0.00 |
| 431.00 | 385.00 | 415.00 | -1.08 | 0.18 |
| 906.00 | 1087.00 | 1062.33 | 1.08 | 0.17 |
| 886.00 | 968.00 | 994.33 | 1.16 | 0.00 |
| 3355.00 | 3739.00 | 3509.33 | 1.08 | 0.06 |
| 5172.00 | 5261.00 | 5296.67 | 1.07 | 0.07 |
| 1508.00 | 1608.00 | 1593.67 | 1.09 | 0.02 |
| 801.00 | 800.00 | 783.33 | -1.09 | 0.15 |
| 5.00 | 4.00 | 6.00 | -1.34 | 0.45 |
| 2058.00 | 2327.00 | 2246.67 | 1.27 | 0.00 |
| 3063.00 | 3025.00 | 2969.67 | -1.31 | 0.00 |
| 401.00 | 379.00 | 395.00 | -1.79 | 0.00 |
| 34.00 | 25.00 | 33.67 | 1.51 | 0.02 |
| 538.00 | 553.00 | 563.67 | 1.16 | 0.00 |
| 15.00 | 25.00 | 19.67 | 1.00 | 1.00 |
| 525.00 | 547.00 | 577.33 | -1.05 | 0.34 |
| 15.00 | 13.00 | 15.67 | -1.57 | 0.03 |
| 2030.00 | 2066.00 | 2107.33 | 1.22 | 0.00 |
| 1570.00 | 1837.00 | 1681.67 | -1.21 | 0.00 |
| 77.00 | 73.00 | 76.00 | -1.96 | 0.00 |
| 1470.00 | 1802.00 | 1588.33 | 1.84 | 0.00 |
| 29.00 | 25.00 | 24.67 | -1.26 | 0.21 |
| 31.00 | 19.00 | 24.67 | -2.13 | 0.00 |
| 125.00 | 134.00 | 129.00 | 1.40 | 0.00 |
| 883.00 | 852.00 | 871.67 | -1.65 | 0.00 |
| 2067.00 | 1974.00 | 2074.00 | -1.54 | 0.00 |
| 128.00 | 148.00 | 139.67 | -1.15 | 0.08 |
| 722.00 | 783.00 | 768.67 | -1.05 | 0.27 |
| 46.00 | 42.00 | 47.33 | -1.17 | 0.23 |
| 1196.00 | 1209.00 | 1159.67 | 1.38 | 0.00 |
| 16.00 | 21.00 | 19.00 | -1.23 | 0.35 |
| 23.00 | 28.00 | 25.00 | -1.12 | 0.56 |
| 59.00 | 66.00 | 57.33 | -1.85 | 0.00 |
| 241.00 | 269.00 | 256.67 | 1.07 | 0.31 |
| 18.00 | 9.00 | 12.00 | -1.65 | 0.03 |
| 4188.00 | 4196.00 | 4254.00 | -2.09 | 0.00 |

| | | | | |
|---|---|---|---|---|
| 62.00 | 83.00 | 78.00 | 1.14 | 0.23 |
| 607.00 | 656.00 | 618.67 | 1.07 | 0.24 |
| 817.00 | 904.00 | 894.67 | 1.10 | 0.03 |
| 340.00 | 342.00 | 377.33 | 1.03 | 0.68 |
| 12.00 | 19.00 | 16.33 | -1.71 | 0.00 |
| 1327.00 | 1329.00 | 1335.00 | -1.03 | 0.46 |
| 2355.00 | 2704.00 | 2614.67 | 1.06 | 0.14 |
| 275.00 | 420.00 | 336.33 | -1.01 | 0.92 |
| 48.00 | 49.00 | 51.67 | -1.11 | 0.41 |
| 14.00 | 13.00 | 15.00 | -1.10 | 0.76 |
| 1441.00 | 1604.00 | 1538.33 | 1.19 | 0.00 |
| 2681.00 | 2834.00 | 2758.00 | 1.19 | 0.00 |
| 3870.00 | 4028.00 | 3997.33 | 1.28 | 0.00 |
| 7272.00 | 8148.00 | 7879.67 | 1.05 | 0.13 |
| 30.00 | 55.00 | 45.33 | -1.26 | 0.08 |
| 2926.00 | 3597.00 | 3397.00 | 1.10 | 0.03 |
| 1576.00 | 1687.00 | 1700.33 | -1.09 | 0.03 |
| 2376.00 | 2335.00 | 2385.67 | -1.13 | 0.00 |
| 1876.00 | 2026.00 | 2047.67 | 1.13 | 0.00 |
| 1472.00 | 1546.00 | 1585.67 | 1.13 | 0.00 |
| 3090.00 | 2748.00 | 2910.67 | -1.05 | 0.36 |
| 270.00 | 301.00 | 279.33 | 1.09 | 0.23 |
| 6.00 | 7.00 | 7.00 | 1.09 | 0.88 |
| 21.00 | 19.00 | 21.00 | -1.16 | 0.46 |
| 330.00 | 385.00 | 362.33 | -1.26 | 0.00 |
| 7.00 | 7.00 | 6.33 | -1.27 | 0.55 |
| 395.00 | 391.00 | 412.67 | 1.20 | 0.00 |
| 133.00 | 127.00 | 128.67 | -1.11 | 0.21 |
| 8606.00 | 9597.00 | 9308.00 | 1.23 | 0.00 |
| 3376.00 | 3574.00 | 3745.33 | 1.06 | 0.16 |
| 211.00 | 291.00 | 257.67 | 1.11 | 0.18 |
| 5604.00 | 6344.00 | 6059.67 | -1.04 | 0.30 |
| 935.00 | 874.00 | 925.33 | -5.36 | 0.00 |
| 4481.00 | 4505.00 | 4415.33 | -1.12 | 0.01 |
| 2.00 | 6.00 | 3.67 | -1.46 | 0.45 |
| 333.00 | 352.00 | 337.67 | -1.24 | 0.00 |
| 2981.00 | 3278.00 | 3200.33 | 1.10 | 0.01 |
| 2.00 | 6.00 | 6.00 | -1.01 | 1.00 |
| 16.00 | 20.00 | 23.67 | 1.88 | 0.00 |
| 87.00 | 115.00 | 104.00 | 2.79 | 0.00 |
| 123.00 | 117.00 | 128.67 | 1.63 | 0.00 |
| 697.00 | 748.00 | 769.33 | 1.02 | 0.78 |
| 22695.00 | 21655.00 | 21503.33 | 1.63 | 0.00 |
| 1846.00 | 1968.00 | 2030.00 | 1.06 | 0.17 |
| 1223.00 | 1320.00 | 1303.33 | 1.14 | 0.00 |
| 1817.00 | 2118.00 | 2076.33 | 1.29 | 0.00 |
| 5.00 | 5.00 | 5.33 | -1.26 | 0.62 |
| 163.00 | 159.00 | 164.33 | 1.07 | 0.41 |

| | | | | |
|---|---|---|---|---|
| 63267.00 | 61942.00 | 62961.67 | -1.32 | 0.00 |
| 650.00 | 586.00 | 601.67 | -1.19 | 0.01 |
| 756.00 | 716.00 | 770.00 | 1.24 | 0.00 |
| 7655.00 | 8427.00 | 8358.00 | -1.22 | 0.00 |
| 124.00 | 115.00 | 128.67 | 1.01 | 0.90 |
| 230.00 | 258.00 | 266.67 | 1.05 | 0.53 |
| 1379.00 | 1502.00 | 1457.00 | 1.12 | 0.01 |
| 200.00 | 198.00 | 206.00 | 1.09 | 0.21 |
| 0.00 | 8.00 | 4.33 | -2.15 | 0.03 |
| 4.00 | 8.00 | 6.00 | -2.45 | 0.00 |
| 119.00 | 148.00 | 141.67 | -1.04 | 0.68 |
| 3931.00 | 4212.00 | 4168.00 | 1.01 | 0.71 |
| 206.00 | 263.00 | 237.67 | -1.24 | 0.00 |
| 237.00 | 260.00 | 238.67 | -1.04 | 0.55 |
| 9591.00 | 10492.00 | 10483.33 | 1.18 | 0.00 |
| 88.00 | 76.00 | 77.67 | -1.54 | 0.00 |
| 1122.00 | 1017.00 | 1067.33 | -1.18 | 0.00 |
| 897.00 | 765.00 | 823.33 | -1.23 | 0.00 |
| 1659.00 | 1628.00 | 1646.67 | -1.25 | 0.00 |
| 962.00 | 975.00 | 988.67 | 1.12 | 0.01 |
| 717.00 | 792.00 | 784.67 | -1.09 | 0.05 |
| 3826.00 | 3951.00 | 4000.67 | -1.18 | 0.00 |
| 706.00 | 731.00 | 746.67 | -1.04 | 0.37 |
| 1340.00 | 1347.00 | 1346.00 | -1.03 | 0.48 |
| 1309.00 | 1453.00 | 1424.33 | -1.04 | 0.28 |
| 178.00 | 164.00 | 188.33 | 1.24 | 0.01 |
| 1385.00 | 1380.00 | 1424.33 | -1.11 | 0.01 |
| 294.00 | 331.00 | 326.67 | -1.09 | 0.14 |
| 144.00 | 161.00 | 161.00 | 1.02 | 0.85 |
| 27.00 | 29.00 | 33.33 | 1.11 | 0.59 |
| 1572.00 | 1752.00 | 1672.67 | -1.05 | 0.21 |
| 646.00 | 613.00 | 639.67 | -1.27 | 0.00 |
| 8101.00 | 8233.00 | 8342.33 | -1.02 | 0.62 |
| 2863.00 | 2756.00 | 2786.67 | -1.04 | 0.42 |
| 1574.00 | 1576.00 | 1603.00 | -1.02 | 0.60 |
| 2800.00 | 3186.00 | 3029.00 | 1.14 | 0.00 |
| 9386.00 | 11106.00 | 10489.67 | -1.04 | 0.26 |
| 7850.00 | 8481.00 | 8338.00 | 1.06 | 0.09 |
| 7532.00 | 8040.00 | 7883.67 | 1.06 | 0.09 |
| 1569.00 | 1682.00 | 1698.00 | 1.06 | 0.18 |
| 1156.00 | 1187.00 | 1246.33 | 1.13 | 0.01 |
| 2660.00 | 3023.00 | 3064.33 | 1.60 | 0.00 |
| 764.00 | 816.00 | 833.00 | -1.00 | 0.93 |
| 386.00 | 384.00 | 383.33 | -1.00 | 0.96 |
| 543.00 | 552.00 | 558.33 | -1.09 | 0.08 |
| 7808.00 | 8874.00 | 8362.00 | 1.22 | 0.00 |
| 168.00 | 212.00 | 195.00 | 1.23 | 0.01 |
| 14024.00 | 13367.00 | 13743.33 | 1.08 | 0.04 |

| | | | | |
|---|---|---|---|---|
| 862.00 | 969.00 | 941.67 | -1.13 | 0.00 |
| 1537.00 | 1673.00 | 1644.33 | -1.15 | 0.00 |
| 88.00 | 105.00 | 98.00 | 1.09 | 0.34 |
| 250.00 | 230.00 | 252.00 | 1.35 | 0.00 |
| 5.00 | 3.00 | 3.33 | -2.48 | 0.02 |
| 1085.00 | 1085.00 | 1097.33 | 1.15 | 0.00 |
| 233.00 | 211.00 | 226.33 | -1.05 | 0.47 |
| 961.00 | 851.00 | 929.67 | 1.48 | 0.00 |
| 1334.00 | 1395.00 | 1431.33 | 1.20 | 0.00 |
| 299.00 | 353.00 | 309.33 | -1.16 | 0.05 |
| 720.00 | 656.00 | 698.00 | 1.04 | 0.48 |
| 886.00 | 944.00 | 935.33 | -1.08 | 0.06 |
| 0.00 | 1.00 | 0.67 | -3.15 | 0.19 |
| 19.00 | 26.00 | 19.67 | 1.42 | 0.11 |
| 5219.00 | 6136.00 | 5842.67 | 1.50 | 0.00 |
| 3.00 | 6.00 | 6.00 | -1.23 | 0.64 |
| 369.00 | 406.00 | 388.33 | -1.82 | 0.00 |
| 3.00 | 5.00 | 4.67 | 2.23 | 0.12 |
| 646.00 | 728.00 | 672.33 | -1.23 | 0.00 |
| 544.00 | 563.00 | 543.67 | -1.04 | 0.52 |
| 61.00 | 50.00 | 54.67 | 1.03 | 0.84 |
| 252.00 | 280.00 | 254.67 | -1.26 | 0.00 |
| 273.00 | 289.00 | 293.67 | 1.14 | 0.04 |
| 423.00 | 509.00 | 470.33 | 1.26 | 0.00 |
| 2318.00 | 2141.00 | 2279.67 | 1.21 | 0.00 |
| 756.00 | 928.00 | 836.00 | -1.28 | 0.00 |
| 939.00 | 902.00 | 955.67 | -1.19 | 0.00 |
| 3226.00 | 3228.00 | 3332.33 | -1.19 | 0.00 |
| 1341.00 | 1341.00 | 1417.00 | -1.11 | 0.01 |
| 1650.00 | 1767.00 | 1782.33 | 1.05 | 0.25 |
| 397.00 | 433.00 | 418.33 | -1.07 | 0.28 |
| 1616.00 | 1599.00 | 1665.67 | -1.18 | 0.00 |
| 53.00 | 72.00 | 60.67 | -1.19 | 0.13 |
| 2910.00 | 2810.00 | 2952.67 | -1.47 | 0.00 |
| 0.00 | 0.00 | 0.33 | -9.11 | 0.00 |
| 16.00 | 11.00 | 15.00 | 1.43 | 0.15 |
| 1169.00 | 1185.00 | 1221.33 | -1.08 | 0.05 |
| 1.00 | 6.00 | 2.67 | -1.62 | 0.39 |
| 37.00 | 39.00 | 35.67 | 1.12 | 0.51 |
| 1130.00 | 1305.00 | 1237.33 | 1.11 | 0.05 |
| 255.00 | 263.00 | 253.00 | -1.02 | 0.75 |
| 2477.00 | 2651.00 | 2774.33 | -1.01 | 0.76 |
| 237.00 | 209.00 | 235.67 | -1.07 | 0.34 |
| 8415.00 | 8312.00 | 8180.33 | -1.26 | 0.00 |
| 2218.00 | 2272.00 | 2250.33 | -1.89 | 0.00 |
| 1313.00 | 1482.00 | 1433.00 | 1.12 | 0.01 |
| 253.00 | 321.00 | 300.00 | -1.29 | 0.00 |
| 2897.00 | 2937.00 | 2995.33 | -1.20 | 0.00 |

| | | | | |
|---|---|---|---|---|
| 507.00 | 518.00 | 527.67 | -1.03 | 0.57 |
| 294.00 | 234.00 | 280.00 | -1.82 | 0.00 |
| 1285.00 | 1304.00 | 1343.00 | 1.16 | 0.00 |
| 16.00 | 12.00 | 13.67 | -1.90 | 0.00 |
| 11.00 | 11.00 | 14.67 | 1.20 | 0.52 |
| 37.00 | 27.00 | 39.00 | 1.28 | 0.12 |
| 1479.00 | 1560.00 | 1535.67 | 1.04 | 0.30 |
| 1342.00 | 1377.00 | 1431.33 | 1.04 | 0.36 |
| 4178.00 | 4323.00 | 4316.67 | -1.16 | 0.00 |
| 4347.00 | 4350.00 | 4406.67 | -1.20 | 0.00 |
| 825.00 | 751.00 | 786.00 | -1.53 | 0.00 |
| 658.00 | 689.00 | 658.67 | -1.09 | 0.12 |
| 92.00 | 108.00 | 107.33 | -1.17 | 0.09 |
| 714.00 | 719.00 | 712.33 | -1.24 | 0.00 |
| 7.00 | 5.00 | 7.67 | -1.19 | 0.68 |
| 238.00 | 232.00 | 228.00 | 1.08 | 0.33 |
| 1580.00 | 1767.00 | 1678.33 | 1.64 | 0.00 |
| 834.00 | 1066.00 | 977.67 | -1.15 | 0.01 |
| 908.00 | 698.00 | 801.67 | -1.25 | 0.00 |
| 1180.00 | 1047.00 | 1113.00 | -1.23 | 0.00 |
| 924.00 | 951.00 | 981.67 | -1.11 | 0.01 |
| 910.00 | 765.00 | 831.00 | 1.10 | 0.13 |
| 6.00 | 3.00 | 4.00 | -1.50 | 0.37 |
| 5794.00 | 5830.00 | 6062.33 | -1.11 | 0.00 |
| 46.00 | 32.00 | 39.33 | -1.71 | 0.00 |
| 55.00 | 53.00 | 50.00 | -1.06 | 0.68 |
| 21.00 | 34.00 | 23.67 | -1.51 | 0.02 |
| 692.00 | 811.00 | 745.33 | 1.18 | 0.00 |
| 20.00 | 20.00 | 20.33 | -1.29 | 0.19 |
| 1145.00 | 1242.00 | 1286.00 | 1.14 | 0.01 |
| 236.00 | 256.00 | 257.00 | 1.75 | 0.00 |
| 181.00 | 241.00 | 207.67 | 1.19 | 0.03 |
| 110.00 | 116.00 | 122.67 | 1.14 | 0.13 |
| 3355.00 | 3198.00 | 3377.67 | 1.13 | 0.01 |
| 1219.00 | 1289.00 | 1314.67 | 1.03 | 0.45 |
| 443.00 | 434.00 | 453.67 | 1.32 | 0.00 |
| 10157.00 | 10619.00 | 10880.67 | 1.03 | 0.45 |
| 658.00 | 803.00 | 734.33 | 1.07 | 0.23 |
| 540.00 | 679.00 | 618.33 | 1.06 | 0.35 |
| 418.00 | 409.00 | 405.33 | -1.29 | 0.00 |
| 595.00 | 631.00 | 605.33 | -1.09 | 0.09 |
| 492.00 | 549.00 | 544.00 | 1.15 | 0.01 |
| 362.00 | 387.00 | 382.00 | -1.05 | 0.36 |
| 688.00 | 765.00 | 722.33 | 1.09 | 0.13 |
| 36.00 | 42.00 | 41.00 | 1.70 | 0.00 |
| 33.00 | 23.00 | 26.00 | -1.05 | 0.83 |
| 1307.00 | 1660.00 | 1527.00 | -1.00 | 0.99 |
| 667.00 | 710.00 | 690.67 | -1.33 | 0.00 |

| | | | | |
|---|---|---|---|---|
| 428.00 | 437.00 | 476.00 | 1.06 | 0.38 |
| 648.00 | 712.00 | 717.00 | 1.14 | 0.01 |
| 342.00 | 361.00 | 343.33 | -1.38 | 0.00 |
| 420.00 | 452.00 | 459.00 | 1.50 | 0.00 |
| 179.00 | 145.00 | 180.00 | -2.42 | 0.00 |
| 5.00 | 5.00 | 4.33 | -2.00 | 0.06 |
| 4669.00 | 4575.00 | 4779.67 | 1.04 | 0.33 |
| 63.00 | 85.00 | 72.67 | 1.03 | 0.83 |
| 455.00 | 450.00 | 439.00 | -1.14 | 0.02 |
| 635.00 | 647.00 | 636.00 | -1.08 | 0.11 |
| 1382.00 | 1593.00 | 1482.00 | -1.04 | 0.42 |
| 1608.00 | 1702.00 | 1713.67 | 1.00 | 0.95 |
| 16.00 | 15.00 | 14.00 | 1.18 | 0.51 |
| 69.00 | 104.00 | 84.33 | 1.10 | 0.43 |
| 306.00 | 355.00 | 334.67 | 1.07 | 0.23 |
| 1461.00 | 1454.00 | 1425.33 | -1.03 | 0.60 |
| 1948.00 | 1876.00 | 1942.00 | -1.24 | 0.00 |
| 732.00 | 743.00 | 770.00 | -1.17 | 0.00 |
| 1539.00 | 1581.00 | 1545.33 | -1.41 | 0.00 |
| 45.00 | 52.00 | 48.00 | -1.30 | 0.03 |
| 2825.00 | 2521.00 | 2652.67 | -1.15 | 0.00 |
| 657.00 | 591.00 | 621.00 | -1.17 | 0.00 |
| 138.00 | 133.00 | 131.33 | -1.89 | 0.00 |
| 3.00 | 1.00 | 2.00 | 1.17 | 1.00 |
| 588.00 | 716.00 | 689.67 | 1.05 | 0.44 |
| 556.00 | 576.00 | 609.00 | -1.07 | 0.18 |
| 340.00 | 414.00 | 390.00 | -1.10 | 0.08 |
| 15.00 | 13.00 | 12.33 | 1.65 | 0.08 |
| 639.00 | 640.00 | 649.67 | -1.01 | 0.89 |
| 647.00 | 673.00 | 693.00 | -1.06 | 0.34 |
| 743.00 | 934.00 | 878.00 | -1.02 | 0.74 |
| 650.00 | 591.00 | 642.67 | 1.08 | 0.19 |
| 453.00 | 469.00 | 452.67 | 1.07 | 0.26 |
| 609.00 | 747.00 | 662.33 | -1.07 | 0.37 |
| 303.00 | 313.00 | 316.33 | 1.08 | 0.25 |
| 645.00 | 819.00 | 763.00 | 1.43 | 0.00 |
| 481.00 | 537.00 | 519.00 | 1.22 | 0.00 |
| 10.00 | 10.00 | 10.67 | -1.23 | 0.50 |
| 1051.00 | 1132.00 | 1063.67 | -1.20 | 0.00 |
| 1946.00 | 1859.00 | 1929.33 | 1.04 | 0.35 |
| 68.00 | 57.00 | 61.33 | -1.06 | 0.61 |
| 724.00 | 908.00 | 821.33 | -1.04 | 0.55 |
| 2.00 | 3.00 | 3.67 | -1.01 | 1.00 |
| 2537.00 | 2840.00 | 2750.00 | 1.07 | 0.11 |
| 10.00 | 12.00 | 11.67 | -1.36 | 0.24 |
| 2455.00 | 2729.00 | 2721.00 | 1.14 | 0.00 |
| 24.00 | 30.00 | 24.67 | 1.07 | 0.75 |
| 839.00 | 829.00 | 840.67 | 1.25 | 0.00 |

| | | | | |
|---|---|---|---|---|
| 3.00 | 7.00 | 4.00 | -1.01 | 1.00 |
| 480.00 | 457.00 | 486.33 | 1.07 | 0.22 |
| 903.00 | 997.00 | 948.67 | 1.02 | 0.71 |
| 4283.00 | 4913.00 | 4755.67 | 1.17 | 0.00 |
| 479.00 | 464.00 | 494.00 | -1.14 | 0.02 |
| 2.00 | 2.00 | 4.33 | -1.09 | 1.00 |
| 201.00 | 259.00 | 237.33 | -1.09 | 0.25 |
| 1538.00 | 1506.00 | 1555.00 | -1.14 | 0.00 |
| 669.00 | 759.00 | 696.33 | 1.05 | 0.37 |
| 1131.00 | 1008.00 | 1124.00 | -1.07 | 0.14 |
| 2113.00 | 2157.00 | 2079.33 | 1.00 | 0.94 |
| 1416.00 | 1443.00 | 1461.67 | 1.06 | 0.11 |
| 1736.00 | 1745.00 | 1830.33 | -1.05 | 0.25 |
| 634.00 | 672.00 | 671.33 | -1.31 | 0.00 |
| 5.00 | 3.00 | 3.00 | -1.77 | 0.24 |
| 26.00 | 43.00 | 34.67 | 1.13 | 0.50 |
| 501.00 | 509.00 | 533.67 | 1.01 | 0.94 |
| 4376.00 | 4959.00 | 4668.33 | -1.05 | 0.19 |
| 80.00 | 72.00 | 79.33 | 1.39 | 0.01 |
| 47.00 | 50.00 | 44.33 | -1.61 | 0.00 |
| 80.00 | 55.00 | 62.67 | -1.12 | 0.42 |
| 51.00 | 48.00 | 49.67 | -1.28 | 0.04 |
| 193.00 | 192.00 | 189.67 | -1.07 | 0.36 |
| 555.00 | 543.00 | 565.00 | 1.11 | 0.05 |
| 466.00 | 401.00 | 464.67 | -1.20 | 0.00 |
| 25.00 | 29.00 | 27.33 | -1.01 | 0.94 |
| 75.00 | 93.00 | 85.33 | -1.29 | 0.01 |
| 200.00 | 196.00 | 209.00 | 1.02 | 0.78 |
| 734.00 | 896.00 | 804.67 | 1.44 | 0.00 |
| 1602.00 | 1630.00 | 1612.00 | -1.20 | 0.00 |
| 306.00 | 325.00 | 306.67 | -1.85 | 0.00 |
| 0.00 | 1.00 | 3.00 | -1.35 | 0.69 |
| 618.00 | 745.00 | 693.33 | -1.09 | 0.11 |
| 1157.00 | 1187.00 | 1196.00 | -1.03 | 0.50 |
| 828.00 | 864.00 | 864.67 | -1.09 | 0.06 |
| 289.00 | 347.00 | 329.33 | 1.28 | 0.00 |
| 232.00 | 256.00 | 267.33 | 1.06 | 0.40 |
| 46.00 | 37.00 | 37.00 | -1.61 | 0.00 |
| 1388.00 | 1489.00 | 1482.00 | 1.11 | 0.01 |
| 2488.00 | 2590.00 | 2568.33 | -1.04 | 0.30 |
| 14500.00 | 15681.00 | 15504.67 | -1.13 | 0.00 |
| 10295.00 | 11143.00 | 11142.00 | 1.25 | 0.00 |
| 1008.00 | 941.00 | 960.00 | -1.12 | 0.02 |
| 21.00 | 25.00 | 23.00 | -1.41 | 0.04 |
| 7.00 | 7.00 | 5.33 | -2.13 | 0.02 |
| 4.00 | 0.00 | 1.67 | -3.28 | 0.02 |
| 346.00 | 345.00 | 361.00 | -1.48 | 0.00 |
| 3.00 | 2.00 | 2.67 | 1.12 | 1.00 |

| | | | | |
|---|---|---|---|---|
| 60.00 | 33.00 | 51.33 | 1.14 | 0.45 |
| 5.00 | 3.00 | 2.67 | -1.86 | 0.22 |
| 7.00 | 9.00 | 6.00 | -2.17 | 0.01 |
| 464.00 | 456.00 | 477.33 | -1.47 | 0.00 |
| 88.00 | 81.00 | 90.00 | -1.10 | 0.34 |
| 148.00 | 176.00 | 161.33 | -1.48 | 0.00 |
| 2343.00 | 2297.00 | 2301.67 | -1.15 | 0.00 |
| 111.00 | 126.00 | 117.33 | -1.60 | 0.00 |
| 622.00 | 631.00 | 607.00 | -1.15 | 0.02 |
| 2.00 | 2.00 | 1.33 | -2.63 | 0.13 |
| 455.00 | 475.00 | 512.00 | 1.20 | 0.00 |
| 385.00 | 467.00 | 442.33 | 1.18 | 0.01 |
| 1200.00 | 1303.00 | 1292.00 | 1.19 | 0.00 |
| 945.00 | 1102.00 | 1027.00 | 1.08 | 0.10 |
| 959.00 | 1088.00 | 1072.33 | 1.27 | 0.00 |
| 155.00 | 162.00 | 172.00 | -1.16 | 0.08 |
| 570.00 | 578.00 | 585.67 | 1.27 | 0.00 |
| 416.00 | 538.00 | 502.67 | 1.06 | 0.31 |
| 405.00 | 431.00 | 478.00 | 1.09 | 0.27 |
| 398.00 | 403.00 | 398.67 | 1.19 | 0.00 |
| 124.00 | 177.00 | 153.67 | 1.26 | 0.02 |
| 73.00 | 75.00 | 86.67 | 1.44 | 0.00 |
| 99.00 | 93.00 | 95.33 | 1.00 | 1.00 |
| 928.00 | 889.00 | 949.67 | 1.05 | 0.37 |
| 164.00 | 176.00 | 173.67 | -1.05 | 0.49 |
| 1535.00 | 1774.00 | 1659.00 | 1.19 | 0.00 |
| 506.00 | 519.00 | 511.67 | 1.08 | 0.22 |
| 388.00 | 354.00 | 388.33 | -1.45 | 0.00 |
| 263.00 | 254.00 | 287.00 | 1.03 | 0.70 |
| 625.00 | 705.00 | 700.33 | -1.42 | 0.00 |
| 652.00 | 762.00 | 731.33 | -1.02 | 0.76 |
| 917.00 | 1049.00 | 1039.00 | -1.14 | 0.00 |
| 8917.00 | 9883.00 | 9999.00 | 1.12 | 0.00 |
| 335.00 | 344.00 | 329.00 | -1.00 | 1.00 |
| 1477.00 | 1712.00 | 1602.00 | 1.21 | 0.00 |
| 339.00 | 420.00 | 400.33 | 1.18 | 0.01 |
| 50.00 | 69.00 | 58.00 | -1.03 | 0.85 |
| 298.00 | 259.00 | 289.33 | 1.10 | 0.15 |
| 2292.00 | 2637.00 | 2485.00 | 1.07 | 0.10 |
| 966.00 | 1005.00 | 1019.67 | -1.38 | 0.00 |
| 1034.00 | 1103.00 | 1092.67 | 1.04 | 0.32 |
| 2187.00 | 2210.00 | 2372.67 | -1.09 | 0.05 |
| 296.00 | 346.00 | 339.00 | 1.20 | 0.01 |
| 302.00 | 360.00 | 345.33 | 1.31 | 0.00 |
| 89.00 | 108.00 | 103.67 | -1.16 | 0.16 |
| 357.00 | 370.00 | 362.67 | 1.13 | 0.03 |
| 733.00 | 808.00 | 802.33 | 1.04 | 0.40 |
| 217.00 | 199.00 | 215.00 | 2.66 | 0.00 |

| | | | | |
|---|---|---|---|---|
| 1129.00 | 1266.00 | 1220.00 | 1.03 | 0.52 |
| 4179.00 | 4499.00 | 4511.33 | 1.18 | 0.00 |
| 218.00 | 238.00 | 243.00 | 1.02 | 0.79 |
| 116.00 | 97.00 | 107.00 | -1.06 | 0.54 |
| 14.00 | 21.00 | 22.33 | -1.17 | 0.40 |
| 18.00 | 28.00 | 25.33 | 1.44 | 0.06 |
| 5.00 | 3.00 | 4.00 | -1.18 | 0.85 |
| 136.00 | 123.00 | 140.33 | -1.24 | 0.01 |
| 395.00 | 417.00 | 433.00 | -1.06 | 0.26 |
| 1090.00 | 1130.00 | 1094.33 | -1.02 | 0.70 |
| 7.00 | 15.00 | 10.33 | -1.05 | 1.00 |
| 774.00 | 708.00 | 764.00 | -1.49 | 0.00 |
| 14.00 | 13.00 | 14.33 | -1.34 | 0.21 |
| 271.00 | 272.00 | 278.00 | -1.27 | 0.00 |
| 1703.00 | 1538.00 | 1649.67 | -1.01 | 0.88 |
| 66.00 | 81.00 | 72.00 | 1.20 | 0.11 |
| 49.00 | 62.00 | 57.00 | -1.04 | 0.77 |
| 2766.00 | 2979.00 | 2976.33 | 1.15 | 0.00 |
| 98.00 | 71.00 | 87.33 | 1.15 | 0.26 |
| 2.00 | 5.00 | 6.33 | -1.59 | 0.20 |
| 28.00 | 23.00 | 25.67 | 1.09 | 0.63 |
| 2026.00 | 2203.00 | 2199.33 | -1.20 | 0.00 |
| 93.00 | 59.00 | 73.67 | -1.39 | 0.01 |
| 2.00 | 1.00 | 1.67 | -1.77 | 0.43 |
| 50.00 | 75.00 | 58.33 | 1.42 | 0.01 |
| 623.00 | 709.00 | 651.67 | -1.27 | 0.00 |
| 4.00 | 10.00 | 8.00 | -1.80 | 0.03 |
| 758.00 | 886.00 | 839.00 | -1.03 | 0.49 |
| 1.00 | 1.00 | 1.00 | -3.41 | 0.06 |
| 849.00 | 819.00 | 871.33 | -1.55 | 0.00 |
| 211.00 | 181.00 | 199.33 | -1.15 | 0.10 |
| 365.00 | 366.00 | 365.67 | -1.18 | 0.01 |
| 473.00 | 458.00 | 451.00 | -1.06 | 0.40 |
| 72.00 | 48.00 | 53.33 | -1.16 | 0.38 |
| 2014.00 | 2006.00 | 2014.33 | 1.09 | 0.04 |
| 1049.00 | 1092.00 | 1100.33 | -1.04 | 0.33 |
| 2225.00 | 2409.00 | 2278.00 | -1.03 | 0.51 |
| 2463.00 | 2752.00 | 2637.33 | 1.13 | 0.00 |
| 1286.00 | 1253.00 | 1286.00 | -1.17 | 0.00 |
| 141.00 | 167.00 | 162.00 | -1.02 | 0.85 |
| 1595.00 | 1622.00 | 1610.33 | -1.12 | 0.01 |
| 733.00 | 841.00 | 823.00 | -1.00 | 0.97 |
| 730.00 | 781.00 | 800.67 | -1.00 | 0.96 |
| 911.00 | 977.00 | 901.67 | 1.13 | 0.04 |
| 1072.00 | 1050.00 | 1097.67 | -1.05 | 0.31 |
| 1450.00 | 1589.00 | 1541.67 | 1.07 | 0.12 |
| 831.00 | 953.00 | 937.67 | 1.03 | 0.58 |
| 2502.00 | 2451.00 | 2540.67 | -1.01 | 0.81 |

| | | | | |
|---|---|---|---|---|
| 789.00 | 895.00 | 853.67 | 1.07 | 0.11 |
| 1181.00 | 1289.00 | 1223.33 | -1.07 | 0.16 |
| 630.00 | 606.00 | 654.00 | -1.17 | 0.01 |
| 2514.00 | 2345.00 | 2478.33 | -1.08 | 0.07 |
| 835.00 | 702.00 | 755.00 | 1.08 | 0.20 |
| 797.00 | 809.00 | 825.67 | 1.14 | 0.00 |
| 3.00 | 0.00 | 1.33 | -2.17 | 0.27 |
| 1347.00 | 1282.00 | 1337.33 | 1.04 | 0.36 |
| 569.00 | 539.00 | 567.00 | 1.00 | 1.00 |
| 1720.00 | 1685.00 | 1674.00 | -1.05 | 0.31 |
| 278.00 | 276.00 | 273.00 | 1.02 | 0.73 |
| 414.00 | 425.00 | 428.33 | -1.34 | 0.00 |
| 10.00 | 10.00 | 10.33 | 2.01 | 0.03 |
| 1604.00 | 1635.00 | 1666.00 | -1.01 | 0.81 |
| 4.00 | 6.00 | 6.00 | -1.57 | 0.19 |
| 1338.00 | 1338.00 | 1330.67 | 1.14 | 0.01 |
| 985.00 | 884.00 | 986.67 | -1.14 | 0.01 |
| 307.00 | 347.00 | 314.00 | 1.27 | 0.00 |
| 1086.00 | 1204.00 | 1236.33 | 1.07 | 0.15 |
| 688.00 | 714.00 | 718.00 | -1.16 | 0.00 |
| 361.00 | 391.00 | 380.33 | 1.02 | 0.70 |
| 455.00 | 396.00 | 440.33 | -1.98 | 0.00 |
| 1716.00 | 1811.00 | 1790.33 | 1.08 | 0.07 |
| 688.00 | 699.00 | 730.00 | 1.03 | 0.54 |
| 17198.00 | 17744.00 | 17298.00 | -1.18 | 0.00 |
| 4861.00 | 4912.00 | 4856.67 | -1.09 | 0.02 |
| 22576.00 | 25475.00 | 24137.33 | -1.08 | 0.05 |
| 77.00 | 87.00 | 81.33 | 1.40 | 0.00 |
| 7.00 | 11.00 | 9.33 | -2.09 | 0.00 |
| 1110.00 | 1340.00 | 1274.33 | 1.01 | 0.84 |
| 1097.00 | 1077.00 | 1117.33 | -1.23 | 0.00 |
| 577.00 | 673.00 | 636.67 | -1.15 | 0.01 |
| 3270.00 | 3245.00 | 3318.67 | -1.03 | 0.47 |
| 1485.00 | 1646.00 | 1587.33 | 1.06 | 0.22 |
| 538.00 | 545.00 | 557.33 | 1.02 | 0.68 |
| 2927.00 | 2810.00 | 2965.33 | -1.20 | 0.00 |
| 5407.00 | 5302.00 | 5245.00 | -1.13 | 0.01 |
| 785.00 | 810.00 | 808.33 | 1.07 | 0.14 |
| 212.00 | 257.00 | 257.33 | 1.30 | 0.00 |
| 4.00 | 5.00 | 5.00 | -1.08 | 1.00 |
| 2185.00 | 2248.00 | 2188.67 | -1.02 | 0.60 |
| 450.00 | 465.00 | 447.33 | 1.09 | 0.11 |
| 1644.00 | 1413.00 | 1567.00 | -1.03 | 0.55 |
| 12949.00 | 14027.00 | 14128.00 | -1.13 | 0.00 |
| 1733.00 | 1632.00 | 1664.00 | -1.32 | 0.00 |
| 118.00 | 151.00 | 135.67 | -1.32 | 0.00 |
| 1188.00 | 1104.00 | 1195.00 | 1.08 | 0.09 |
| 1202.00 | 1204.00 | 1279.67 | 1.02 | 0.62 |

| | | | | |
|---|---|---|---|---|
| 1.00 | 6.00 | 2.33 | 1.36 | 0.78 |
| 6946.00 | 7298.00 | 7217.33 | -1.08 | 0.03 |
| 1031.00 | 1092.00 | 1057.00 | 1.05 | 0.24 |
| 844.00 | 829.00 | 822.67 | -1.38 | 0.00 |
| 16.00 | 16.00 | 14.33 | 1.51 | 0.11 |
| 601.00 | 706.00 | 646.00 | -1.07 | 0.26 |
| 107.00 | 107.00 | 107.67 | 1.15 | 0.14 |
| 610.00 | 524.00 | 571.00 | -1.26 | 0.00 |
| 813.00 | 900.00 | 881.33 | 1.08 | 0.08 |
| 185.00 | 187.00 | 183.33 | -1.10 | 0.18 |
| 104.00 | 126.00 | 116.33 | -1.59 | 0.00 |
| 323.00 | 286.00 | 335.67 | -1.14 | 0.08 |
| 63.00 | 44.00 | 55.67 | 2.19 | 0.00 |
| 713.00 | 877.00 | 829.67 | -1.03 | 0.50 |
| 6727.00 | 6427.00 | 6842.67 | -1.07 | 0.10 |
| 141.00 | 109.00 | 127.67 | -1.26 | 0.01 |
| 49.00 | 52.00 | 52.33 | 1.09 | 0.52 |
| 11.00 | 16.00 | 10.00 | 1.02 | 1.00 |
| 3058.00 | 3406.00 | 3080.33 | -1.12 | 0.08 |
| 568.00 | 543.00 | 558.33 | -1.03 | 0.59 |
| 21.00 | 32.00 | 24.33 | 1.12 | 0.53 |
| 1086.00 | 1137.00 | 1146.33 | -1.03 | 0.54 |
| 3250.00 | 3318.00 | 3412.33 | -1.04 | 0.28 |
| 1213.00 | 1454.00 | 1298.67 | 1.02 | 0.70 |
| 2414.00 | 2474.00 | 2504.00 | -1.12 | 0.00 |
| 2.00 | 4.00 | 5.00 | -1.08 | 1.00 |
| 4278.00 | 4398.00 | 4408.67 | -1.08 | 0.03 |
| 200.00 | 214.00 | 208.67 | -1.37 | 0.00 |
| 949.00 | 915.00 | 933.00 | -1.33 | 0.00 |
| 1570.00 | 1477.00 | 1581.67 | 1.00 | 0.96 |
| 519.00 | 476.00 | 502.00 | -1.14 | 0.02 |
| 8680.00 | 8323.00 | 8568.33 | -1.15 | 0.00 |
| 3162.00 | 3534.00 | 3229.00 | -1.53 | 0.00 |
| 281.00 | 289.00 | 287.67 | 1.14 | 0.02 |
| 935.00 | 1052.00 | 956.67 | -1.38 | 0.00 |
| 2.00 | 7.00 | 3.33 | -1.01 | 1.00 |
| 24136.00 | 22831.00 | 23780.67 | -1.07 | 0.12 |
| 57.00 | 53.00 | 47.67 | 1.20 | 0.25 |
| 37.00 | 50.00 | 44.67 | -1.44 | 0.00 |
| 1995.00 | 2001.00 | 2042.00 | -1.24 | 0.00 |
| 305.00 | 341.00 | 328.00 | -1.06 | 0.34 |
| 21.00 | 29.00 | 25.00 | -1.34 | 0.08 |
| 33.00 | 41.00 | 35.33 | -1.93 | 0.00 |
| 1.00 | 1.00 | 2.00 | -1.97 | 0.25 |
| 1366.00 | 1377.00 | 1384.00 | -1.08 | 0.05 |
| 7.00 | 2.00 | 4.00 | -2.32 | 0.02 |
| 0.00 | 2.00 | 0.67 | -4.00 | 0.07 |
| 245.00 | 263.00 | 257.33 | -1.36 | 0.00 |

| | | | | |
|---|---|---|---|---|
| 1524.00 | 1515.00 | 1548.67 | -1.33 | 0.00 |
| 24.00 | 25.00 | 24.00 | 1.27 | 0.20 |
| 424.00 | 465.00 | 434.00 | 1.10 | 0.10 |
| 1002.00 | 994.00 | 1037.00 | -2.04 | 0.00 |
| 648.00 | 631.00 | 646.00 | -1.12 | 0.02 |
| 605.00 | 594.00 | 595.67 | -1.17 | 0.00 |
| 3701.00 | 3768.00 | 3774.33 | -1.04 | 0.31 |
| 1065.00 | 1094.00 | 1090.67 | -1.23 | 0.00 |
| 56.00 | 52.00 | 53.67 | 1.16 | 0.26 |
| 468.00 | 495.00 | 495.33 | -1.04 | 0.49 |
| 0.00 | 2.00 | 1.33 | -2.17 | 0.27 |
| 49.00 | 47.00 | 50.33 | -1.39 | 0.01 |
| 1566.00 | 1699.00 | 1648.00 | -1.03 | 0.52 |
| 329.00 | 378.00 | 360.67 | 1.40 | 0.00 |
| 1567.00 | 1636.00 | 1608.33 | 1.00 | 0.91 |
| 1022.00 | 1120.00 | 1084.33 | 1.28 | 0.00 |
| 326.00 | 399.00 | 363.00 | -1.09 | 0.16 |
| 5.00 | 1.00 | 3.67 | -1.90 | 0.12 |
| 10.00 | 24.00 | 13.67 | 1.61 | 0.15 |
| 2791.00 | 2746.00 | 2679.00 | -1.25 | 0.00 |
| 14.00 | 5.00 | 10.33 | 1.22 | 0.52 |
| 115.00 | 139.00 | 135.00 | -1.29 | 0.00 |
| 897.00 | 985.00 | 974.33 | 1.25 | 0.00 |
| 1112.00 | 1152.00 | 1156.00 | 1.29 | 0.00 |
| 1497.00 | 1441.00 | 1481.33 | -1.45 | 0.00 |
| 7.00 | 6.00 | 6.00 | 1.18 | 0.73 |
| 858.00 | 835.00 | 839.00 | 1.18 | 0.00 |
| 1641.00 | 1992.00 | 1768.67 | 1.01 | 0.93 |
| 1101.00 | 1159.00 | 1120.33 | 1.03 | 0.47 |
| 364.00 | 372.00 | 370.67 | 1.06 | 0.28 |
| 823.00 | 1127.00 | 1017.33 | 1.04 | 0.52 |
| 3372.00 | 3880.00 | 3696.67 | -1.00 | 0.95 |
| 1123.00 | 1279.00 | 1213.00 | 1.01 | 0.91 |
| 585.00 | 644.00 | 618.67 | -1.05 | 0.36 |
| 91.00 | 69.00 | 77.00 | -1.21 | 0.18 |
| 19.00 | 23.00 | 18.67 | -1.07 | 0.86 |
| 247.00 | 266.00 | 256.00 | 1.17 | 0.02 |
| 1001.00 | 1121.00 | 1116.67 | 1.12 | 0.02 |
| 3887.00 | 4160.00 | 4100.00 | -1.82 | 0.00 |
| 1097.00 | 1229.00 | 1165.00 | -1.04 | 0.43 |
| 1804.00 | 2062.00 | 1906.67 | -1.04 | 0.38 |
| 16.00 | 27.00 | 22.33 | 1.27 | 0.26 |
| 97.00 | 102.00 | 100.00 | -1.61 | 0.00 |
| 151.00 | 136.00 | 141.33 | 1.17 | 0.06 |
| 1263.00 | 1275.00 | 1299.67 | 1.15 | 0.00 |
| 746.00 | 756.00 | 784.33 | 1.11 | 0.02 |
| 1572.00 | 1731.00 | 1735.67 | 1.26 | 0.00 |
| 1021.00 | 1135.00 | 1078.33 | 1.25 | 0.00 |

| | | | | |
|---|---|---|---|---|
| 849.00 | 943.00 | 916.33 | -1.02 | 0.66 |
| 4626.00 | 4473.00 | 4502.00 | 1.04 | 0.33 |
| 2096.00 | 2339.00 | 2218.00 | 1.10 | 0.01 |
| 1502.00 | 1740.00 | 1615.67 | 1.04 | 0.42 |
| 805.00 | 887.00 | 858.00 | 1.07 | 0.16 |
| 3125.00 | 2929.00 | 2929.33 | -1.05 | 0.34 |
| 2562.00 | 2495.00 | 2549.67 | -1.14 | 0.00 |
| 3319.00 | 3474.00 | 3531.67 | 1.11 | 0.00 |
| 2739.00 | 2811.00 | 2844.33 | 1.07 | 0.06 |
| 134.00 | 117.00 | 134.00 | -1.45 | 0.00 |
| 1923.00 | 2117.00 | 2145.33 | 1.12 | 0.01 |
| 4323.00 | 4395.00 | 4367.67 | -1.01 | 0.78 |
| 1209.00 | 1360.00 | 1279.00 | 1.18 | 0.00 |
| 2696.00 | 3190.00 | 2972.67 | 1.11 | 0.01 |
| 5441.00 | 6517.00 | 5999.33 | 1.27 | 0.00 |
| 982.00 | 1183.00 | 1051.33 | 1.02 | 0.73 |
| 6.00 | 7.00 | 8.00 | -1.10 | 0.89 |
| 1612.00 | 1835.00 | 1759.67 | 1.01 | 0.85 |
| 2887.00 | 3266.00 | 3010.33 | 1.01 | 0.84 |
| 62.00 | 73.00 | 68.67 | 1.41 | 0.01 |
| 97.00 | 100.00 | 110.67 | 1.49 | 0.00 |
| 1613.00 | 1500.00 | 1521.67 | -1.24 | 0.00 |
| 3736.00 | 4117.00 | 3844.00 | -1.05 | 0.27 |
| 56.00 | 61.00 | 53.67 | -1.47 | 0.00 |
| 38.00 | 36.00 | 33.00 | 1.06 | 0.74 |
| 42.00 | 46.00 | 38.67 | -1.71 | 0.00 |
| 6.00 | 9.00 | 7.67 | -1.75 | 0.05 |
| 139.00 | 161.00 | 174.00 | 1.01 | 0.92 |
| 3960.00 | 3890.00 | 3958.33 | -1.08 | 0.05 |
| 2205.00 | 2082.00 | 2203.00 | 1.19 | 0.00 |
| 2286.00 | 2267.00 | 2327.00 | -1.11 | 0.00 |
| 8.00 | 5.00 | 5.33 | 1.13 | 0.86 |
| 1037.00 | 1049.00 | 1066.33 | -1.00 | 0.97 |
| 1688.00 | 1807.00 | 1817.67 | 1.16 | 0.00 |
| 108.00 | 109.00 | 113.67 | 1.16 | 0.11 |
| 94.00 | 155.00 | 126.33 | 1.01 | 0.94 |
| 559.00 | 653.00 | 628.67 | -1.18 | 0.00 |
| 1025.00 | 1122.00 | 1091.33 | 1.39 | 0.00 |
| 125.00 | 134.00 | 139.33 | -1.05 | 0.51 |
| 1309.00 | 1554.00 | 1486.00 | 1.18 | 0.00 |
| 9.00 | 3.00 | 6.00 | -1.18 | 0.76 |
| 401.00 | 462.00 | 441.00 | 1.18 | 0.00 |
| 296.00 | 335.00 | 329.00 | 1.10 | 0.11 |
| 26.00 | 37.00 | 29.00 | 1.45 | 0.04 |
| 96.00 | 97.00 | 94.00 | 1.00 | 1.00 |
| 141.00 | 195.00 | 193.33 | -1.11 | 0.34 |
| 971.00 | 958.00 | 1016.00 | 1.21 | 0.00 |
| 133.00 | 150.00 | 148.00 | 1.12 | 0.15 |

| | | | | |
|---|---|---|---|---|
| 47.00 | 54.00 | 51.00 | 1.16 | 0.26 |
| 1154.00 | 1378.00 | 1281.33 | 1.09 | 0.06 |
| 209.00 | 221.00 | 209.00 | 1.05 | 0.50 |
| 506.00 | 510.00 | 537.33 | 1.05 | 0.31 |
| 40.00 | 49.00 | 42.67 | -1.08 | 0.62 |
| 502.00 | 544.00 | 559.67 | -1.41 | 0.00 |
| 1211.00 | 1195.00 | 1244.33 | -1.09 | 0.07 |
| 229.00 | 215.00 | 226.33 | 1.25 | 0.00 |
| 0.00 | 0.00 | 0.67 | -8.26 | 0.00 |
| 1431.00 | 1418.00 | 1457.00 | -1.15 | 0.00 |
| 161.00 | 198.00 | 185.67 | 1.04 | 0.60 |
| 3515.00 | 4020.00 | 3944.00 | 1.38 | 0.00 |
| 7424.00 | 7405.00 | 7547.67 | -1.06 | 0.13 |
| 1644.00 | 1773.00 | 1733.67 | 1.12 | 0.00 |
| 510.00 | 597.00 | 575.67 | -1.00 | 0.98 |
| 1093.00 | 1015.00 | 1046.33 | -1.08 | 0.13 |
| 9496.00 | 10794.00 | 10430.33 | 1.25 | 0.00 |
| 647.00 | 642.00 | 659.33 | 1.10 | 0.04 |
| 476.00 | 466.00 | 510.67 | 1.09 | 0.21 |
| 449.00 | 467.00 | 454.67 | -1.07 | 0.29 |
| 975.00 | 1035.00 | 1045.67 | -1.06 | 0.15 |
| 21.00 | 34.00 | 35.00 | 1.04 | 0.90 |
| 1035.00 | 1139.00 | 1091.67 | -1.10 | 0.03 |
| 0.00 | 1.00 | 0.33 | -4.68 | 0.13 |
| 52.00 | 57.00 | 62.00 | -1.30 | 0.02 |
| 1501.00 | 1629.00 | 1575.67 | 1.04 | 0.33 |
| 173.00 | 207.00 | 188.33 | 1.47 | 0.00 |
| 557.00 | 544.00 | 551.00 | 1.09 | 0.11 |
| 551.00 | 553.00 | 594.00 | 1.02 | 0.73 |
| 81.00 | 70.00 | 72.67 | 1.15 | 0.24 |
| 1729.00 | 1917.00 | 1867.33 | -1.06 | 0.13 |
| 657.00 | 744.00 | 727.67 | 1.05 | 0.29 |
| 688.00 | 706.00 | 712.67 | 1.00 | 1.00 |
| 1205.00 | 1229.00 | 1302.00 | -1.15 | 0.00 |
| 149.00 | 182.00 | 181.00 | 1.06 | 0.52 |
| 350.00 | 434.00 | 416.67 | 1.04 | 0.53 |
| 199.00 | 224.00 | 219.00 | -1.42 | 0.00 |
| 66.00 | 58.00 | 72.33 | 1.39 | 0.01 |
| 21.00 | 23.00 | 24.67 | -1.61 | 0.00 |
| 7385.00 | 7108.00 | 7179.33 | -1.05 | 0.29 |
| 773.00 | 946.00 | 898.00 | 1.12 | 0.02 |
| 5300.00 | 5929.00 | 6006.00 | 1.06 | 0.17 |
| 286.00 | 300.00 | 327.00 | -1.08 | 0.37 |
| 416.00 | 489.00 | 485.00 | -1.03 | 0.56 |
| 735.00 | 881.00 | 864.00 | -1.04 | 0.46 |
| 1302.00 | 1347.00 | 1406.67 | 1.21 | 0.00 |
| 2103.00 | 2123.00 | 2306.33 | -1.00 | 0.96 |
| 1733.00 | 1878.00 | 1952.67 | 1.18 | 0.00 |

| | | | | |
|---|---|---|---|---|
| 220.00 | 244.00 | 243.67 | -1.02 | 0.81 |
| 423.00 | 481.00 | 485.33 | -1.09 | 0.14 |
| 479.00 | 526.00 | 507.33 | -1.03 | 0.56 |
| 186.00 | 227.00 | 216.67 | -1.06 | 0.38 |
| 6.00 | 6.00 | 8.33 | -1.09 | 0.89 |
| 4345.00 | 5013.00 | 5006.67 | -1.10 | 0.03 |
| 1384.00 | 1594.00 | 1463.00 | -1.10 | 0.04 |
| 1116.00 | 1197.00 | 1267.67 | 1.05 | 0.31 |
| 1780.00 | 2011.00 | 2074.00 | 1.21 | 0.00 |
| 2892.00 | 2876.00 | 3041.33 | -1.00 | 1.00 |
| 2068.00 | 2061.00 | 2216.33 | 1.05 | 0.25 |
| 54.00 | 37.00 | 51.00 | -1.00 | 1.00 |
| 355.00 | 456.00 | 422.00 | -1.00 | 0.96 |
| 159.00 | 144.00 | 155.00 | 1.09 | 0.36 |
| 34.00 | 52.00 | 44.67 | 1.03 | 0.86 |
| 28.00 | 23.00 | 32.33 | 1.21 | 0.31 |
| 1.00 | 4.00 | 2.33 | -2.25 | 0.10 |
| 4.00 | 3.00 | 4.33 | 1.07 | 1.00 |
| 4.00 | 10.00 | 10.67 | 1.01 | 1.00 |
| 5.00 | 9.00 | 11.33 | -1.19 | 0.51 |
| 4.00 | 4.00 | 4.00 | 1.30 | 0.67 |
| 46.00 | 56.00 | 50.33 | -1.09 | 0.49 |
| 2.00 | 0.00 | 0.67 | -1.86 | 0.69 |
| 196.00 | 217.00 | 207.33 | -1.04 | 0.55 |
| 13.00 | 9.00 | 9.67 | 1.89 | 0.05 |
| 16.00 | 25.00 | 23.00 | -1.28 | 0.17 |
| 5.00 | 10.00 | 6.00 | 1.11 | 0.87 |
| 3.00 | 3.00 | 3.00 | -1.55 | 0.41 |
| 25.00 | 51.00 | 39.00 | -1.07 | 0.69 |
| 96.00 | 117.00 | 112.67 | -1.16 | 0.08 |
| 16.00 | 11.00 | 17.00 | -1.43 | 0.07 |
| 38.00 | 40.00 | 40.33 | 1.02 | 0.95 |
| 7.00 | 6.00 | 6.00 | 1.26 | 0.61 |
| 1.00 | 2.00 | 3.00 | -2.09 | 0.10 |
| 10.00 | 8.00 | 11.67 | -1.39 | 0.21 |
| 118.00 | 139.00 | 138.67 | -1.05 | 0.57 |
| 105.00 | 130.00 | 115.33 | 1.02 | 0.82 |
| 52.00 | 69.00 | 62.00 | 1.52 | 0.00 |
| 221.00 | 319.00 | 274.33 | 1.09 | 0.29 |
| 26.00 | 26.00 | 24.67 | 1.01 | 0.94 |
| 22.00 | 37.00 | 37.00 | -1.45 | 0.01 |
| 43.00 | 37.00 | 39.33 | -1.27 | 0.07 |
| 46.00 | 40.00 | 51.00 | -1.18 | 0.21 |
| 3.00 | 4.00 | 6.33 | -1.96 | 0.03 |
| 38.00 | 35.00 | 35.67 | -1.82 | 0.00 |
| 32.00 | 33.00 | 31.33 | -1.07 | 0.68 |
| 42.00 | 51.00 | 50.33 | -1.14 | 0.31 |
| 4.00 | 7.00 | 7.33 | -1.24 | 0.58 |

| | | | | |
|---|---|---|---|---|
| 30.00 | 36.00 | 33.00 | -1.01 | 0.95 |
| 17.00 | 18.00 | 20.00 | -1.13 | 0.55 |
| 2.00 | 4.00 | 2.00 | -1.17 | 1.00 |
| 5.00 | 2.00 | 7.33 | -1.84 | 0.06 |
| 16.00 | 28.00 | 20.67 | -1.24 | 0.27 |
| 46.00 | 37.00 | 42.33 | -1.19 | 0.20 |
| 56.00 | 48.00 | 47.67 | -1.28 | 0.06 |
| 28.00 | 15.00 | 22.33 | -1.12 | 0.64 |
| 151.00 | 184.00 | 172.00 | -1.11 | 0.22 |
| 59.00 | 74.00 | 72.00 | -1.14 | 0.21 |
| 49.00 | 57.00 | 57.00 | -1.12 | 0.34 |
| 51.00 | 39.00 | 52.00 | -1.18 | 0.20 |
| 24.00 | 37.00 | 32.33 | -1.06 | 0.74 |
| 17.00 | 43.00 | 36.00 | 1.09 | 0.71 |
| 103.00 | 87.00 | 103.00 | -1.13 | 0.23 |
| 37.00 | 39.00 | 37.33 | -1.18 | 0.25 |
| 22.00 | 19.00 | 19.67 | -1.05 | 0.87 |
| 256.00 | 282.00 | 290.00 | -1.05 | 0.44 |
| 33.00 | 45.00 | 38.33 | -1.06 | 0.72 |
| 50.00 | 46.00 | 48.33 | -1.04 | 0.79 |
| 52.00 | 68.00 | 60.33 | 1.02 | 0.89 |
| 10.00 | 6.00 | 7.00 | -1.77 | 0.06 |
| 41.00 | 41.00 | 41.67 | 1.20 | 0.24 |
| 212.00 | 217.00 | 230.33 | -1.11 | 0.10 |
| 142.00 | 151.00 | 149.33 | 1.09 | 0.28 |
| 9.00 | 14.00 | 10.00 | 1.34 | 0.35 |
| 32.00 | 33.00 | 33.67 | 1.07 | 0.70 |
| 17.00 | 8.00 | 14.00 | -1.08 | 0.84 |
| 3.00 | 3.00 | 2.33 | -1.01 | 1.00 |
| 14.00 | 7.00 | 8.67 | -1.13 | 0.80 |
| 190.00 | 205.00 | 212.33 | 1.05 | 0.52 |
| 5.00 | 3.00 | 5.33 | -1.82 | 0.08 |
| 12.00 | 11.00 | 8.33 | 1.37 | 0.42 |
| 59.00 | 66.00 | 61.00 | -1.36 | 0.01 |
| 70.00 | 68.00 | 65.00 | -1.06 | 0.66 |
| 64.00 | 75.00 | 77.00 | -1.12 | 0.27 |
| 36.00 | 28.00 | 40.00 | 1.01 | 0.96 |
| 45.00 | 64.00 | 60.00 | 1.05 | 0.74 |
| 37.00 | 44.00 | 42.33 | 1.00 | 1.00 |
| 61.00 | 64.00 | 78.67 | -1.25 | 0.12 |
| 7.00 | 7.00 | 7.67 | -1.27 | 0.50 |
| 79.00 | 111.00 | 97.00 | -1.21 | 0.04 |
| 3.00 | 3.00 | 2.00 | -1.65 | 0.46 |
| 0.00 | 2.00 | 1.33 | -1.48 | 0.76 |
| 2.00 | 7.00 | 4.00 | -1.58 | 0.30 |
| 8.00 | 5.00 | 6.67 | -1.26 | 0.57 |
| 9.00 | 9.00 | 8.00 | -1.22 | 0.60 |
| 2.00 | 3.00 | 2.33 | -1.56 | 0.49 |

| | | | | |
|---|---|---|---|---|
| 7.00 | 3.00 | 5.00 | -1.21 | 0.73 |
| 17.00 | 5.00 | 10.00 | -1.11 | 0.81 |
| 34.00 | 18.00 | 22.67 | -1.73 | 0.01 |
| 13.00 | 13.00 | 13.00 | 2.11 | 0.01 |
| 0.00 | 3.00 | 1.67 | -1.39 | 0.78 |
| 59.00 | 44.00 | 49.00 | -1.42 | 0.01 |
| 0.00 | 4.00 | 2.00 | -1.33 | 0.79 |
| 10.00 | 5.00 | 7.67 | 1.73 | 0.14 |
| 6.00 | 5.00 | 7.00 | -1.30 | 0.48 |
| 63.00 | 41.00 | 49.00 | -1.11 | 0.51 |
| 9.00 | 16.00 | 10.00 | -1.08 | 0.90 |
| 49.00 | 85.00 | 69.33 | -1.01 | 0.94 |
| 87.00 | 96.00 | 98.33 | -1.04 | 0.64 |
| 5.00 | 5.00 | 5.67 | -1.25 | 0.64 |
| 11.00 | 9.00 | 16.67 | -1.18 | 0.53 |
| 6.00 | 11.00 | 9.33 | -1.73 | 0.03 |
| 18.00 | 22.00 | 19.00 | -1.14 | 0.60 |
| 23.00 | 34.00 | 27.67 | -1.05 | 0.77 |
| 17.00 | 13.00 | 17.67 | -1.16 | 0.49 |
| 22.00 | 16.00 | 22.33 | 1.16 | 0.49 |
| 8.00 | 5.00 | 7.33 | -1.74 | 0.06 |
| 32.00 | 39.00 | 40.00 | -1.17 | 0.28 |
| 16.00 | 11.00 | 14.00 | -1.04 | 1.00 |
| 24.00 | 27.00 | 29.67 | -1.13 | 0.46 |
| 14.00 | 12.00 | 11.67 | -1.38 | 0.20 |
| 6.00 | 5.00 | 5.33 | -1.45 | 0.35 |
| 22.00 | 32.00 | 24.00 | -1.07 | 0.77 |
| 51.00 | 74.00 | 63.00 | -1.01 | 0.96 |
| 13.00 | 9.00 | 14.33 | -1.32 | 0.21 |
| 23.00 | 21.00 | 23.00 | -1.22 | 0.24 |
| 164.00 | 170.00 | 173.33 | 1.04 | 0.66 |
| 16.00 | 19.00 | 19.33 | 1.27 | 0.29 |
| 15.00 | 8.00 | 12.67 | -1.28 | 0.35 |
| 59.00 | 53.00 | 60.33 | -1.01 | 0.93 |
| 32.00 | 24.00 | 29.33 | -1.11 | 0.57 |
| 5.00 | 7.00 | 9.67 | 1.35 | 0.42 |
| 52.00 | 71.00 | 69.33 | 1.04 | 0.77 |
| 21.00 | 24.00 | 22.00 | -1.03 | 1.00 |
| 35.00 | 33.00 | 37.00 | 1.20 | 0.25 |
| 233.00 | 200.00 | 216.33 | -1.04 | 0.64 |
| 25.00 | 15.00 | 20.00 | -1.16 | 0.50 |
| 29.00 | 34.00 | 34.00 | -1.10 | 0.52 |
| 26.00 | 29.00 | 29.67 | -1.25 | 0.14 |
| 190.00 | 215.00 | 228.33 | 1.18 | 0.05 |
| 26.00 | 32.00 | 29.00 | -1.09 | 0.57 |
| 6.00 | 8.00 | 6.67 | 1.62 | 0.22 |
| 5.00 | 5.00 | 9.00 | -1.17 | 0.62 |
| 17.00 | 34.00 | 30.00 | 1.19 | 0.39 |

| | | | | |
|---|---|---|---|---|
| 0.00 | 6.00 | 3.00 | -1.01 | 1.00 |
| 16.00 | 7.00 | 13.67 | 1.01 | 1.00 |
| 3.00 | 3.00 | 2.67 | -4.28 | 0.00 |
| 28.00 | 32.00 | 30.67 | 1.01 | 1.00 |
| 147.00 | 146.00 | 149.67 | 1.14 | 0.10 |
| 148.00 | 162.00 | 141.00 | -1.02 | 0.89 |
| 158.00 | 222.00 | 190.33 | -1.14 | 0.09 |
| 89.00 | 93.00 | 89.33 | -1.34 | 0.00 |
| 24.00 | 33.00 | 28.33 | 1.37 | 0.08 |
| 170.00 | 207.00 | 177.67 | -1.05 | 0.61 |
| 78.00 | 88.00 | 87.00 | -1.15 | 0.13 |
| 45.00 | 39.00 | 39.67 | 1.19 | 0.25 |
| 3.00 | 8.00 | 5.33 | 1.21 | 0.72 |
| 85.00 | 86.00 | 86.33 | -1.07 | 0.50 |
| 3.00 | 0.00 | 1.33 | -2.40 | 0.19 |
| 27.00 | 30.00 | 31.33 | -1.24 | 0.16 |
| 8.00 | 9.00 | 11.00 | 1.08 | 0.81 |
| 1.00 | 5.00 | 2.67 | -1.01 | 1.00 |
| 27.00 | 25.00 | 30.67 | -1.16 | 0.40 |
| 26.00 | 38.00 | 30.00 | 1.15 | 0.42 |
| 3.00 | 2.00 | 4.33 | -1.01 | 1.00 |
| 32.00 | 38.00 | 40.67 | -1.27 | 0.07 |
| 46.00 | 80.00 | 66.00 | 1.01 | 0.95 |
| 1.00 | 2.00 | 2.00 | -1.33 | 0.79 |
| 3.00 | 3.00 | 2.67 | -2.10 | 0.11 |
| 37.00 | 44.00 | 39.33 | 1.05 | 0.76 |
| 11.00 | 15.00 | 14.33 | 1.28 | 0.32 |
| 26.00 | 32.00 | 26.67 | 1.05 | 0.82 |
| 11.00 | 6.00 | 9.33 | -1.19 | 0.62 |
| 0.00 | 6.00 | 2.67 | -1.14 | 1.00 |
| 29.00 | 24.00 | 29.00 | 1.10 | 0.66 |
| 10.00 | 11.00 | 10.00 | 1.47 | 0.21 |
| 1.00 | 5.00 | 5.67 | -1.49 | 0.31 |
| 3.00 | 9.00 | 5.00 | -1.28 | 0.62 |
| 62.00 | 39.00 | 48.67 | 1.05 | 0.80 |
| 1.00 | 1.00 | 3.67 | -1.02 | 1.00 |
| 36.00 | 41.00 | 37.67 | -1.22 | 0.15 |
| 53.00 | 63.00 | 63.67 | -1.22 | 0.06 |
| 77.00 | 96.00 | 91.33 | 1.05 | 0.69 |
| 30.00 | 40.00 | 32.00 | 1.28 | 0.13 |
| 386.00 | 427.00 | 423.00 | -1.05 | 0.37 |
| 18.00 | 26.00 | 24.00 | -1.11 | 0.54 |
| 79.00 | 67.00 | 75.00 | -1.24 | 0.04 |
| 115.00 | 124.00 | 112.67 | -1.09 | 0.37 |
| 2.00 | 0.00 | 0.67 | -2.71 | 0.30 |
| 35.00 | 55.00 | 38.33 | 1.26 | 0.20 |
| 15.00 | 13.00 | 18.67 | 1.04 | 1.00 |
| 6.00 | 7.00 | 9.67 | 1.88 | 0.07 |

| | | | | |
|---|---|---|---|---|
| 34.00 | 43.00 | 41.00 | 1.12 | 0.47 |
| 6.00 | 6.00 | 8.33 | -1.37 | 0.31 |
| 14.00 | 26.00 | 17.67 | -1.01 | 1.00 |
| 22.00 | 19.00 | 17.00 | -1.31 | 0.20 |
| 223.00 | 279.00 | 267.67 | -1.11 | 0.14 |
| 22.00 | 21.00 | 21.33 | 1.07 | 0.73 |
| 1.00 | 2.00 | 1.33 | -3.10 | 0.06 |
| 19.00 | 28.00 | 22.33 | 1.12 | 0.55 |
| 556.00 | 537.00 | 569.33 | 1.06 | 0.31 |
| 505.00 | 563.00 | 552.00 | -1.11 | 0.04 |
| 238.00 | 270.00 | 271.33 | -1.47 | 0.00 |
| 7580.00 | 7445.00 | 7477.67 | -1.11 | 0.00 |
| 246.00 | 284.00 | 280.00 | 1.30 | 0.00 |
| 531.00 | 640.00 | 625.00 | 1.26 | 0.00 |
| 319.00 | 329.00 | 328.00 | 1.27 | 0.00 |
| 1182.00 | 1247.00 | 1271.33 | 1.11 | 0.01 |
| 624.00 | 804.00 | 718.00 | 1.00 | 0.96 |
| 3719.00 | 4133.00 | 4033.00 | 1.02 | 0.61 |
| 1984.00 | 2122.00 | 2201.00 | 1.14 | 0.00 |
| 987.00 | 1086.00 | 1102.67 | 1.11 | 0.02 |
| 6072.00 | 6378.00 | 6591.33 | 1.21 | 0.00 |
| 1658.00 | 1957.00 | 1936.33 | 1.17 | 0.00 |
| 2136.00 | 2351.00 | 2436.67 | 1.26 | 0.00 |
| 1502.00 | 1680.00 | 1688.00 | 1.40 | 0.00 |
| 1959.00 | 2259.00 | 2320.67 | 1.17 | 0.00 |
| 1793.00 | 2045.00 | 2012.00 | 1.19 | 0.00 |
| 1423.00 | 1615.00 | 1662.33 | 1.34 | 0.00 |
| 1142.00 | 1419.00 | 1379.33 | 1.27 | 0.00 |
| 1028.00 | 1164.00 | 1217.00 | 1.23 | 0.00 |
| 130.00 | 137.00 | 138.33 | 1.14 | 0.11 |
| 129.00 | 123.00 | 120.00 | 1.09 | 0.37 |
| 983.00 | 1048.00 | 1018.33 | 1.11 | 0.01 |
| 1935.00 | 2225.00 | 2224.67 | 1.27 | 0.00 |
| 605.00 | 658.00 | 643.00 | 1.18 | 0.00 |
| 1797.00 | 2123.00 | 2014.00 | 1.09 | 0.04 |
| 2217.00 | 2528.00 | 2412.67 | -1.12 | 0.00 |
| 47.00 | 80.00 | 61.00 | 1.09 | 0.51 |
| 364.00 | 357.00 | 368.00 | -1.06 | 0.31 |
| 1116.00 | 1138.00 | 1185.67 | 1.16 | 0.00 |
| 1132.00 | 1194.00 | 1164.00 | 1.04 | 0.31 |
| 2544.00 | 2746.00 | 2693.67 | 1.02 | 0.66 |
| 250.00 | 339.00 | 303.00 | 1.18 | 0.03 |
| 1782.00 | 2025.00 | 2027.67 | 1.10 | 0.03 |
| 361.00 | 407.00 | 391.67 | 1.17 | 0.01 |
| 1653.00 | 1603.00 | 1596.67 | -1.16 | 0.00 |
| 1534.00 | 1770.00 | 1740.67 | 1.18 | 0.00 |
| 427.00 | 420.00 | 418.33 | -1.09 | 0.14 |
| 5.00 | 0.00 | 2.00 | 1.44 | 0.76 |

| | | | | |
|---|---|---|---|---|
| 342.00 | 343.00 | 347.67 | 1.29 | 0.00 |
| 526.00 | 578.00 | 560.00 | 1.26 | 0.00 |
| 1116.00 | 1156.00 | 1179.00 | -1.13 | 0.00 |
| 373.00 | 408.00 | 382.00 | -1.36 | 0.00 |
| 12.00 | 14.00 | 10.67 | -1.20 | 0.57 |
| 7.00 | 5.00 | 4.67 | 1.15 | 0.85 |
| 1723.00 | 1902.00 | 1930.33 | 1.05 | 0.26 |
| 405.00 | 389.00 | 409.00 | -1.19 | 0.01 |
| 39.00 | 45.00 | 48.33 | -1.22 | 0.11 |
| 1146.00 | 1155.00 | 1157.33 | -1.39 | 0.00 |
| 374.00 | 417.00 | 406.00 | 1.31 | 0.00 |
| 2708.00 | 3224.00 | 3101.00 | 1.12 | 0.01 |
| 1610.00 | 1808.00 | 1752.33 | 1.05 | 0.25 |
| 528.00 | 586.00 | 582.00 | 1.20 | 0.00 |
| 1047.00 | 988.00 | 1058.33 | -1.04 | 0.37 |
| 1438.00 | 1599.00 | 1539.33 | -1.15 | 0.00 |
| 2633.00 | 2895.00 | 2790.00 | 1.03 | 0.42 |
| 148.00 | 186.00 | 164.33 | -1.06 | 0.45 |
| 9.00 | 14.00 | 11.33 | 1.39 | 0.25 |
| 583.00 | 646.00 | 629.00 | 1.20 | 0.00 |
| 67.00 | 64.00 | 63.33 | -1.04 | 0.80 |
| 1005.00 | 1032.00 | 1045.33 | -1.07 | 0.11 |
| 1104.00 | 1276.00 | 1221.33 | -1.08 | 0.05 |
| 604.00 | 586.00 | 582.00 | -1.04 | 0.50 |
| 318.00 | 361.00 | 338.67 | -1.18 | 0.00 |
| 56.00 | 75.00 | 64.00 | -1.12 | 0.34 |
| 2058.00 | 2399.00 | 2332.33 | 1.21 | 0.00 |
| 654.00 | 663.00 | 668.67 | 1.23 | 0.00 |
| 889.00 | 644.00 | 719.67 | -1.35 | 0.00 |
| 376.00 | 396.00 | 365.67 | -1.39 | 0.00 |
| 6.00 | 5.00 | 4.67 | -1.15 | 0.86 |
| 6932.00 | 7387.00 | 7022.33 | -1.10 | 0.02 |
| 6.00 | 1.00 | 3.67 | -1.73 | 0.21 |
| 67.00 | 76.00 | 81.67 | -1.44 | 0.00 |
| 830.00 | 782.00 | 835.67 | 1.04 | 0.41 |
| 5.00 | 11.00 | 7.33 | -1.51 | 0.19 |
| 239.00 | 256.00 | 257.00 | -1.70 | 0.00 |
| 519.00 | 596.00 | 562.33 | -1.38 | 0.00 |
| 1199.00 | 1176.00 | 1195.33 | 1.27 | 0.00 |
| 798.00 | 793.00 | 790.00 | -1.04 | 0.46 |
| 1152.00 | 1342.00 | 1239.67 | -1.07 | 0.15 |
| 760.00 | 772.00 | 763.33 | -1.06 | 0.31 |
| 0.00 | 4.00 | 1.67 | 1.57 | 0.74 |
| 337.00 | 397.00 | 368.00 | 1.25 | 0.00 |
| 1798.00 | 1723.00 | 1789.33 | 1.17 | 0.00 |
| 95.00 | 107.00 | 105.00 | -1.37 | 0.00 |
| 8.00 | 13.00 | 11.67 | -1.21 | 0.51 |
| 8.00 | 10.00 | 10.67 | -1.42 | 0.18 |

| | | | | |
|---|---|---|---|---|
| 10.00 | 15.00 | 12.33 | 1.72 | 0.06 |
| 6.00 | 6.00 | 5.00 | 1.34 | 0.57 |
| 512.00 | 587.00 | 576.00 | -1.04 | 0.55 |
| 8.00 | 11.00 | 10.33 | -1.62 | 0.09 |
| 4.00 | 10.00 | 7.00 | -2.15 | 0.01 |
| 35.00 | 32.00 | 36.33 | -1.50 | 0.00 |
| 2091.00 | 2287.00 | 2272.33 | 1.42 | 0.00 |
| 95.00 | 98.00 | 102.67 | 1.46 | 0.00 |
| 1986.00 | 1973.00 | 1974.67 | -1.14 | 0.00 |
| 1957.00 | 1940.00 | 1988.67 | -1.07 | 0.09 |
| 282.00 | 349.00 | 326.00 | -1.02 | 0.74 |
| 103.00 | 143.00 | 126.00 | -1.32 | 0.00 |
| 700.00 | 781.00 | 766.33 | -1.06 | 0.20 |
| 447.00 | 404.00 | 448.67 | -1.04 | 0.46 |
| 25.00 | 25.00 | 29.67 | -1.10 | 0.58 |
| 1621.00 | 1796.00 | 1688.00 | 1.12 | 0.02 |
| 344.00 | 400.00 | 378.67 | 1.01 | 0.86 |
| 12.00 | 4.00 | 9.00 | 1.11 | 0.79 |
| 9.00 | 10.00 | 8.67 | 1.42 | 0.30 |
| 19.00 | 24.00 | 20.00 | 1.44 | 0.08 |
| 110.00 | 148.00 | 149.33 | 1.17 | 0.13 |
| 64.00 | 88.00 | 80.00 | -1.13 | 0.22 |
| 216.00 | 272.00 | 236.67 | 1.43 | 0.00 |
| 292.00 | 357.00 | 342.33 | 1.13 | 0.05 |
| 125.00 | 104.00 | 123.00 | -1.80 | 0.00 |
| 1258.00 | 1380.00 | 1362.67 | 1.03 | 0.46 |
| 0.00 | 3.00 | 3.33 | -1.21 | 0.84 |
| 2302.00 | 2611.00 | 2560.00 | -1.07 | 0.12 |
| 28.00 | 34.00 | 35.00 | -1.23 | 0.16 |
| 3514.00 | 3732.00 | 3597.00 | -1.11 | 0.01 |
| 1.00 | 3.00 | 2.33 | -1.29 | 0.81 |
| 233.00 | 291.00 | 286.33 | 1.11 | 0.18 |
| 26.00 | 34.00 | 31.33 | 1.20 | 0.31 |
| 5446.00 | 5657.00 | 5678.33 | -1.07 | 0.14 |
| 544.00 | 584.00 | 552.00 | 1.42 | 0.00 |
| 7.00 | 9.00 | 8.00 | 1.80 | 0.10 |
| 963.00 | 1017.00 | 955.33 | -1.21 | 0.00 |
| 50.00 | 53.00 | 55.00 | -1.09 | 0.49 |
| 528.00 | 560.00 | 576.33 | 1.09 | 0.10 |
| 1749.00 | 1662.00 | 1798.67 | -1.23 | 0.00 |
| 67.00 | 58.00 | 70.00 | -1.08 | 0.52 |
| 139.00 | 172.00 | 167.33 | -1.15 | 0.10 |
| 233.00 | 289.00 | 276.67 | 1.43 | 0.00 |
| 165.00 | 180.00 | 183.67 | -1.20 | 0.02 |
| 12.00 | 6.00 | 10.67 | 1.84 | 0.05 |
| 22.00 | 16.00 | 19.67 | 1.35 | 0.16 |
| 211.00 | 223.00 | 213.00 | 1.01 | 0.92 |
| 488.00 | 549.00 | 535.67 | -1.04 | 0.39 |

| | | | | |
|---|---|---|---|---|
| 69.00 | 65.00 | 75.33 | 1.12 | 0.34 |
| 25.00 | 37.00 | 32.00 | -1.80 | 0.00 |
| 178.00 | 203.00 | 190.00 | 1.03 | 0.68 |
| 8.00 | 15.00 | 11.33 | 1.29 | 0.38 |
| 5056.00 | 5614.00 | 5565.67 | 1.26 | 0.00 |
| 2011.00 | 2180.00 | 2145.00 | -1.01 | 0.77 |
| 123.00 | 126.00 | 125.33 | 1.23 | 0.01 |
| 5.00 | 8.00 | 7.33 | 1.65 | 0.18 |
| 1963.00 | 2069.00 | 2165.67 | 1.15 | 0.00 |
| 562.00 | 633.00 | 631.00 | 1.15 | 0.03 |
| 3796.00 | 4057.00 | 3907.67 | 1.29 | 0.00 |
| 1150.00 | 1459.00 | 1325.33 | 1.05 | 0.30 |
| 4.00 | 4.00 | 4.67 | -1.58 | 0.25 |
| 1.00 | 2.00 | 2.00 | -1.97 | 0.24 |
| 88.00 | 75.00 | 90.67 | 1.11 | 0.33 |
| 917.00 | 959.00 | 987.67 | -1.08 | 0.07 |
| 7.00 | 15.00 | 9.67 | -1.33 | 0.40 |
| 130.00 | 161.00 | 152.00 | -1.09 | 0.34 |
| 2292.00 | 2221.00 | 2307.67 | -1.41 | 0.00 |
| 2873.00 | 2586.00 | 2611.00 | -1.50 | 0.00 |
| 128.00 | 166.00 | 150.33 | 1.05 | 0.64 |
| 1792.00 | 1797.00 | 1792.67 | -1.14 | 0.00 |
| 1550.00 | 1933.00 | 1797.67 | 1.04 | 0.46 |
| 25.00 | 25.00 | 26.67 | -1.03 | 0.88 |
| 1715.00 | 1904.00 | 1896.00 | 1.07 | 0.07 |
| 2969.00 | 2949.00 | 3104.33 | -1.10 | 0.02 |
| 2632.00 | 2757.00 | 2868.33 | -1.29 | 0.00 |
| 608.00 | 578.00 | 626.67 | -1.01 | 0.87 |
| 404.00 | 376.00 | 396.33 | -1.13 | 0.03 |
| 921.00 | 965.00 | 949.00 | -1.06 | 0.17 |
| 1706.00 | 1864.00 | 1819.67 | -1.06 | 0.16 |
| 26.00 | 26.00 | 31.33 | -1.00 | 1.00 |
| 735.00 | 841.00 | 814.33 | -1.09 | 0.05 |
| 608.00 | 683.00 | 619.00 | 1.48 | 0.00 |
| 5.00 | 1.00 | 2.67 | -2.10 | 0.11 |
| 0.00 | 1.00 | 0.67 | -2.72 | 0.29 |
| 30.00 | 19.00 | 23.67 | -1.34 | 0.10 |
| 2562.00 | 2860.00 | 2757.67 | 1.04 | 0.37 |
| 323.00 | 369.00 | 352.33 | -1.10 | 0.09 |
| 1617.00 | 1450.00 | 1623.33 | -1.04 | 0.50 |
| 4.00 | 2.00 | 4.33 | 1.07 | 1.00 |
| 14.00 | 17.00 | 17.33 | -2.12 | 0.00 |
| 1916.00 | 1908.00 | 1997.00 | -1.32 | 0.00 |
| 168.00 | 159.00 | 155.33 | -2.44 | 0.00 |
| 3.00 | 4.00 | 3.67 | -1.99 | 0.09 |
| 6.00 | 4.00 | 5.33 | -1.51 | 0.28 |
| 1132.00 | 1270.00 | 1247.00 | 1.12 | 0.01 |
| 662.00 | 763.00 | 709.00 | 1.22 | 0.00 |

| | | | | |
|---|---|---|---|---|
| 125.00 | 114.00 | 126.67 | 1.55 | 0.00 |
| 1015.00 | 1073.00 | 1038.00 | 1.25 | 0.00 |
| 1378.00 | 1421.00 | 1484.67 | -1.17 | 0.00 |
| 961.00 | 988.00 | 1016.33 | 1.11 | 0.01 |
| 567.00 | 569.00 | 597.67 | 1.44 | 0.00 |
| 1912.00 | 2308.00 | 2052.67 | -1.26 | 0.00 |
| 2022.00 | 2058.00 | 2076.67 | 1.18 | 0.00 |
| 287.00 | 324.00 | 306.00 | 1.03 | 0.65 |
| 298.00 | 263.00 | 288.67 | 1.21 | 0.00 |
| 102.00 | 124.00 | 113.33 | 1.49 | 0.00 |
| 5155.00 | 5045.00 | 5245.33 | 1.13 | 0.00 |
| 3006.00 | 2938.00 | 3136.67 | -1.17 | 0.00 |
| 79.00 | 68.00 | 65.67 | -1.60 | 0.00 |
| 594.00 | 569.00 | 594.33 | 1.05 | 0.32 |
| 162.00 | 251.00 | 206.33 | -1.22 | 0.02 |
| 302.00 | 349.00 | 329.67 | 1.32 | 0.00 |
| 823.00 | 787.00 | 826.00 | -1.10 | 0.05 |
| 240.00 | 321.00 | 288.33 | -1.01 | 0.88 |
| 580.00 | 565.00 | 607.67 | -1.14 | 0.01 |
| 5010.00 | 5073.00 | 4970.00 | -1.15 | 0.00 |
| 0.00 | 1.00 | 1.00 | -3.11 | 0.10 |
| 3173.00 | 3131.00 | 3080.67 | -1.42 | 0.00 |
| 3837.00 | 3601.00 | 3709.00 | -1.03 | 0.51 |
| 2.00 | 2.00 | 1.33 | -4.02 | 0.01 |
| 155.00 | 196.00 | 180.33 | -1.14 | 0.07 |
| 2039.00 | 2273.00 | 2166.00 | 1.02 | 0.66 |
| 1215.00 | 1107.00 | 1134.67 | -1.11 | 0.05 |
| 476.00 | 399.00 | 431.00 | -1.12 | 0.08 |
| 422.00 | 555.00 | 481.33 | 1.31 | 0.00 |
| 596.00 | 561.00 | 569.00 | -1.01 | 0.82 |
| 40.00 | 40.00 | 41.33 | 1.11 | 0.50 |
| 581.00 | 620.00 | 615.00 | -1.12 | 0.04 |
| 1689.00 | 1770.00 | 1767.67 | 1.06 | 0.14 |
| 239.00 | 292.00 | 277.00 | -1.10 | 0.14 |
| 6501.00 | 5973.00 | 6343.33 | -1.31 | 0.00 |
| 828.00 | 917.00 | 952.33 | 1.03 | 0.61 |
| 405.00 | 444.00 | 427.33 | 1.07 | 0.20 |
| 1948.00 | 1819.00 | 1981.00 | -1.14 | 0.00 |
| 3740.00 | 3114.00 | 3323.33 | -1.34 | 0.00 |
| 446.00 | 457.00 | 448.67 | 1.03 | 0.56 |
| 274.00 | 243.00 | 266.00 | 1.58 | 0.00 |
| 9.00 | 9.00 | 8.33 | -1.41 | 0.26 |
| 3649.00 | 3841.00 | 3763.33 | 1.01 | 0.88 |
| 2436.00 | 2322.00 | 2346.33 | 1.02 | 0.62 |
| 2866.00 | 3267.00 | 3135.00 | 1.02 | 0.69 |
| 587.00 | 715.00 | 646.00 | 1.08 | 0.22 |
| 1768.00 | 1703.00 | 1789.33 | 1.08 | 0.05 |
| 533.00 | 483.00 | 514.33 | 1.11 | 0.06 |

| | | | | |
|---|---|---|---|---|
| 2325.00 | 2141.00 | 2225.67 | 1.01 | 0.79 |
| 1489.00 | 1393.00 | 1498.00 | -1.23 | 0.00 |
| 83.00 | 92.00 | 84.67 | -1.26 | 0.01 |
| 57.00 | 49.00 | 54.67 | -1.15 | 0.32 |
| 30.00 | 28.00 | 26.33 | -1.34 | 0.12 |
| 6.00 | 9.00 | 7.00 | -1.72 | 0.07 |
| 8920.00 | 8939.00 | 9190.33 | 1.36 | 0.00 |
| 1394.00 | 1671.00 | 1654.67 | 1.23 | 0.00 |
| 1729.00 | 1935.00 | 1922.67 | 1.18 | 0.00 |
| 1835.00 | 2017.00 | 2060.67 | 1.06 | 0.16 |
| 173.00 | 171.00 | 165.33 | -1.45 | 0.00 |
| 657.00 | 736.00 | 711.00 | 1.03 | 0.57 |
| 1496.00 | 1714.00 | 1655.00 | -1.01 | 0.88 |
| 1750.00 | 1822.00 | 1854.00 | 1.05 | 0.21 |
| 2197.00 | 2227.00 | 2250.67 | 1.08 | 0.07 |
| 3004.00 | 3621.00 | 3440.67 | 1.25 | 0.00 |
| 3940.00 | 4261.00 | 4158.67 | 1.14 | 0.00 |
| 825.00 | 845.00 | 840.67 | 1.14 | 0.01 |
| 3124.00 | 3107.00 | 3222.33 | -1.04 | 0.30 |
| 2563.00 | 2405.00 | 2559.33 | 1.06 | 0.20 |
| 7067.00 | 7419.00 | 7272.33 | 1.21 | 0.00 |
| 976.00 | 1038.00 | 1043.33 | -1.27 | 0.00 |
| 158.00 | 201.00 | 178.33 | 1.03 | 0.73 |
| 331.00 | 353.00 | 364.33 | 1.07 | 0.22 |
| 2677.00 | 2775.00 | 2803.00 | -1.16 | 0.00 |
| 25399.00 | 21471.00 | 23187.33 | -1.40 | 0.00 |
| 6.00 | 5.00 | 5.67 | -1.48 | 0.29 |
| 17.00 | 22.00 | 21.33 | 1.03 | 0.93 |
| 3019.00 | 3222.00 | 3222.67 | 1.03 | 0.42 |
| 9229.00 | 10300.00 | 9944.00 | 1.15 | 0.00 |
| 3030.00 | 3738.00 | 3476.67 | 1.10 | 0.02 |
| 6652.00 | 8364.00 | 7692.67 | 1.07 | 0.10 |
| 7201.00 | 8450.00 | 8079.00 | 1.07 | 0.06 |
| 5926.00 | 6615.00 | 6569.67 | 1.16 | 0.00 |
| 2523.00 | 2663.00 | 2654.33 | -1.01 | 0.87 |
| 4740.00 | 5389.00 | 5246.33 | -1.07 | 0.05 |
| 7211.00 | 8537.00 | 8200.00 | -1.04 | 0.31 |
| 3482.00 | 3872.00 | 3880.67 | 1.10 | 0.02 |
| 700.00 | 789.00 | 763.33 | 1.03 | 0.52 |
| 2766.00 | 3015.00 | 2996.67 | 1.13 | 0.00 |
| 1886.00 | 1784.00 | 1893.33 | 1.19 | 0.00 |
| 1807.00 | 2272.00 | 2103.00 | -1.16 | 0.00 |
| 1101.00 | 1227.00 | 1214.33 | -1.05 | 0.20 |
| 262.00 | 320.00 | 316.33 | 1.09 | 0.24 |
| 3278.00 | 3844.00 | 3702.67 | 1.18 | 0.00 |
| 622.00 | 711.00 | 704.67 | 1.22 | 0.00 |
| 113.00 | 101.00 | 120.67 | 1.12 | 0.24 |
| 1135.00 | 1157.00 | 1186.67 | 1.04 | 0.32 |

| | | | | |
|---|---|---|---|---|
| 1058.00 | 1017.00 | 1056.67 | 1.06 | 0.17 |
| 10.00 | 11.00 | 11.67 | -1.19 | 0.58 |
| 29.00 | 29.00 | 28.67 | -1.45 | 0.01 |
| 3622.00 | 3958.00 | 3633.33 | 1.11 | 0.08 |
| 1608.00 | 1497.00 | 1539.33 | -1.23 | 0.00 |
| 565.00 | 588.00 | 587.67 | -1.15 | 0.00 |
| 173.00 | 158.00 | 182.33 | -1.16 | 0.05 |
| 742.00 | 733.00 | 804.00 | 1.15 | 0.01 |
| 8.00 | 5.00 | 8.67 | -1.28 | 0.45 |
| 0.00 | 3.00 | 1.33 | -3.56 | 0.02 |
| 6911.00 | 7249.00 | 7258.67 | -1.03 | 0.39 |
| 5969.00 | 6139.00 | 6348.00 | 1.16 | 0.00 |
| 27839.00 | 27802.00 | 28188.67 | 1.39 | 0.00 |
| 2284.00 | 2200.00 | 2342.00 | 1.07 | 0.17 |
| 79.00 | 93.00 | 86.00 | 1.15 | 0.16 |
| 6002.00 | 6336.00 | 6274.67 | 1.17 | 0.00 |
| 685.00 | 742.00 | 761.67 | 1.19 | 0.00 |
| 46.00 | 42.00 | 43.67 | -1.13 | 0.38 |
| 1673.00 | 1725.00 | 1823.33 | 1.16 | 0.00 |
| 9.00 | 10.00 | 9.33 | -1.51 | 0.13 |
| 570.00 | 622.00 | 602.33 | 1.23 | 0.00 |
| 3183.00 | 3359.00 | 3369.33 | 1.17 | 0.00 |
| 46.00 | 46.00 | 56.33 | 1.45 | 0.01 |
| 312.00 | 348.00 | 357.67 | -1.06 | 0.30 |
| 47.00 | 41.00 | 53.33 | 1.01 | 1.00 |
| 16.00 | 14.00 | 13.33 | 2.06 | 0.01 |
| 2133.00 | 1915.00 | 1997.33 | -1.48 | 0.00 |
| 536.00 | 598.00 | 568.67 | -1.02 | 0.65 |
| 509.00 | 549.00 | 574.67 | -1.08 | 0.14 |
| 279.00 | 270.00 | 272.67 | 1.02 | 0.78 |
| 29.00 | 38.00 | 32.33 | -1.12 | 0.48 |
| 126.00 | 102.00 | 113.33 | -1.13 | 0.19 |
| 510.00 | 589.00 | 554.33 | -1.66 | 0.00 |
| 53.00 | 44.00 | 52.00 | -1.68 | 0.00 |
| 548.00 | 570.00 | 556.67 | -1.20 | 0.00 |
| 446.00 | 420.00 | 433.33 | 1.14 | 0.02 |
| 14.00 | 13.00 | 14.67 | 1.14 | 0.59 |
| 2.00 | 1.00 | 1.33 | -3.33 | 0.03 |
| 191.00 | 175.00 | 198.33 | -1.40 | 0.00 |
| 533.00 | 480.00 | 536.67 | -1.01 | 0.84 |
| 907.00 | 944.00 | 999.33 | 1.10 | 0.08 |
| 874.00 | 812.00 | 855.33 | -1.15 | 0.00 |
| 43.00 | 38.00 | 38.67 | -1.27 | 0.08 |
| 1188.00 | 1217.00 | 1221.67 | 1.06 | 0.17 |
| 8.00 | 5.00 | 6.67 | -1.96 | 0.02 |
| 222.00 | 271.00 | 253.67 | 1.08 | 0.29 |
| 71.00 | 108.00 | 89.67 | 1.04 | 0.77 |
| 24.00 | 28.00 | 28.33 | -1.58 | 0.00 |

| | | | | |
|---|---|---|---|---|
| 5.00 | 1.00 | 2.67 | -1.74 | 0.29 |
| 128.00 | 148.00 | 131.33 | -1.17 | 0.08 |
| 73.00 | 69.00 | 70.67 | -1.07 | 0.54 |
| 1027.00 | 1023.00 | 1020.33 | -1.05 | 0.25 |
| 2213.00 | 2468.00 | 2311.67 | 1.02 | 0.70 |
| 254.00 | 278.00 | 278.33 | 1.10 | 0.16 |
| 37.00 | 19.00 | 25.33 | -1.13 | 0.59 |
| 211.00 | 194.00 | 190.67 | -1.22 | 0.04 |
| 102.00 | 77.00 | 84.67 | -1.12 | 0.34 |
| 180.00 | 176.00 | 186.67 | -1.01 | 0.96 |
| 5.00 | 6.00 | 6.33 | 1.10 | 0.87 |
| 677.00 | 710.00 | 723.00 | -1.19 | 0.00 |
| 980.00 | 1084.00 | 1039.67 | 1.10 | 0.03 |
| 646.00 | 757.00 | 702.33 | 1.07 | 0.19 |
| 75.00 | 87.00 | 80.00 | -1.15 | 0.19 |
| 1067.00 | 1077.00 | 1097.00 | 1.10 | 0.04 |
| 846.00 | 942.00 | 947.33 | -1.01 | 0.87 |
| 95.00 | 93.00 | 102.33 | 1.28 | 0.02 |
| 182.00 | 164.00 | 202.33 | -1.02 | 0.83 |
| 2301.00 | 2465.00 | 2449.33 | -1.13 | 0.00 |
| 1570.00 | 1431.00 | 1540.00 | -1.05 | 0.28 |
| 1245.00 | 1339.00 | 1290.00 | -1.02 | 0.69 |
| 398.00 | 423.00 | 424.00 | 1.27 | 0.00 |
| 42.00 | 51.00 | 46.00 | -1.20 | 0.16 |
| 3326.00 | 3625.00 | 3572.67 | 1.10 | 0.00 |
| 210.00 | 223.00 | 212.67 | -1.11 | 0.12 |
| 504.00 | 476.00 | 480.00 | -1.01 | 0.92 |
| 538.00 | 504.00 | 496.67 | -1.50 | 0.00 |
| 1495.00 | 1624.00 | 1596.00 | 1.05 | 0.26 |
| 1049.00 | 1104.00 | 1084.67 | -1.22 | 0.00 |
| 1114.00 | 1086.00 | 1118.67 | -1.01 | 0.87 |
| 23.00 | 17.00 | 20.67 | -1.71 | 0.00 |
| 168.00 | 162.00 | 162.67 | 1.04 | 0.64 |
| 1508.00 | 1564.00 | 1550.33 | -1.10 | 0.01 |
| 3510.00 | 3552.00 | 3628.00 | -1.18 | 0.00 |
| 2687.00 | 2850.00 | 2811.00 | 1.11 | 0.00 |
| 666.00 | 696.00 | 688.00 | 1.04 | 0.41 |
| 15.00 | 13.00 | 13.00 | 1.07 | 0.83 |
| 3499.00 | 3291.00 | 3437.00 | -1.28 | 0.00 |
| 12099.00 | 11723.00 | 12419.67 | -1.33 | 0.00 |
| 1739.00 | 2027.00 | 1965.33 | 1.55 | 0.00 |
| 1383.00 | 1425.00 | 1453.00 | 1.62 | 0.00 |
| 1022.00 | 1134.00 | 1082.67 | 1.25 | 0.00 |
| 4158.00 | 4142.00 | 4144.00 | -1.25 | 0.00 |
| 666.00 | 778.00 | 742.00 | 1.24 | 0.00 |
| 1072.00 | 1014.00 | 1038.67 | -1.20 | 0.00 |
| 365.00 | 370.00 | 382.33 | 1.02 | 0.78 |
| 9563.00 | 10370.00 | 10224.00 | 1.07 | 0.05 |

| | | | | |
|---|---|---|---|---|
| 790.00 | 744.00 | 788.67 | -1.46 | 0.00 |
| 1359.00 | 1484.00 | 1442.00 | 1.09 | 0.03 |
| 712.00 | 729.00 | 770.00 | 1.08 | 0.13 |
| 588.00 | 681.00 | 677.33 | 1.01 | 0.87 |
| 1630.00 | 1847.00 | 1775.67 | 1.14 | 0.00 |
| 398.00 | 496.00 | 453.67 | -1.23 | 0.00 |
| 1867.00 | 1927.00 | 1971.00 | 1.03 | 0.48 |
| 2499.00 | 2593.00 | 2707.00 | 1.04 | 0.30 |
| 420.00 | 494.00 | 444.67 | 1.20 | 0.01 |
| 565.00 | 659.00 | 610.67 | 1.20 | 0.00 |
| 50.00 | 46.00 | 48.00 | -1.13 | 0.39 |
| 40.00 | 42.00 | 40.67 | 1.11 | 0.49 |
| 179.00 | 190.00 | 189.33 | -1.13 | 0.10 |
| 1402.00 | 1392.00 | 1414.00 | -1.09 | 0.03 |
| 1147.00 | 1228.00 | 1226.33 | -1.21 | 0.00 |
| 1003.00 | 1105.00 | 1116.33 | -1.11 | 0.02 |
| 597.00 | 681.00 | 638.33 | 1.24 | 0.00 |
| 1198.00 | 1217.00 | 1230.67 | 1.02 | 0.62 |
| 1613.00 | 1698.00 | 1613.00 | -1.16 | 0.00 |
| 969.00 | 990.00 | 1032.67 | -1.35 | 0.00 |
| 10.00 | 12.00 | 14.00 | 1.53 | 0.13 |
| 6.00 | 5.00 | 8.67 | -1.29 | 0.46 |
| 3023.00 | 3469.00 | 3434.33 | 1.55 | 0.00 |
| 1329.00 | 1375.00 | 1398.00 | 1.38 | 0.00 |
| 1614.00 | 1461.00 | 1562.00 | 1.22 | 0.00 |
| 328.00 | 276.00 | 311.67 | -1.01 | 0.90 |
| 4489.00 | 4687.00 | 4933.67 | 1.26 | 0.00 |
| 77.00 | 105.00 | 89.67 | -1.16 | 0.13 |
| 13.00 | 5.00 | 7.67 | 2.03 | 0.07 |
| 295.00 | 247.00 | 280.33 | -1.22 | 0.03 |
| 3.00 | 2.00 | 3.00 | 1.10 | 1.00 |
| 709.00 | 698.00 | 776.33 | 1.23 | 0.00 |
| 832.00 | 1006.00 | 987.00 | -1.01 | 0.89 |
| 584.00 | 633.00 | 631.33 | -1.08 | 0.11 |
| 3443.00 | 3867.00 | 3783.33 | -1.15 | 0.00 |
| 675.00 | 683.00 | 723.67 | 1.28 | 0.00 |
| 1002.00 | 979.00 | 989.67 | -1.16 | 0.00 |
| 1207.00 | 1376.00 | 1327.33 | -1.81 | 0.00 |
| 880.00 | 950.00 | 865.67 | 1.00 | 0.98 |
| 582.00 | 652.00 | 625.33 | 1.06 | 0.21 |
| 1902.00 | 1835.00 | 1947.67 | 1.02 | 0.67 |
| 218.00 | 193.00 | 203.67 | -1.18 | 0.02 |
| 4887.00 | 5312.00 | 5221.00 | -1.12 | 0.00 |
| 7517.00 | 8194.00 | 7900.00 | -1.01 | 0.73 |
| 1845.00 | 1953.00 | 1979.00 | -1.03 | 0.37 |
| 858.00 | 966.00 | 999.00 | 1.12 | 0.05 |
| 31.00 | 17.00 | 24.67 | -1.33 | 0.12 |
| 786.00 | 821.00 | 810.67 | 1.15 | 0.00 |

| | | | | |
|---|---|---|---|---|
| 3846.00 | 4503.00 | 4270.33 | -1.11 | 0.01 |
| 31.00 | 43.00 | 37.67 | -1.09 | 0.55 |
| 425.00 | 403.00 | 408.00 | -1.24 | 0.00 |
| 464.00 | 465.00 | 475.67 | 1.09 | 0.12 |
| 31.00 | 23.00 | 24.67 | -1.28 | 0.15 |
| 1.00 | 6.00 | 3.00 | 1.45 | 0.62 |
| 859.00 | 890.00 | 922.00 | -1.64 | 0.00 |
| 480.00 | 520.00 | 517.00 | -1.61 | 0.00 |
| 790.00 | 947.00 | 882.00 | 1.06 | 0.19 |
| 1160.00 | 1225.00 | 1197.00 | -1.09 | 0.08 |
| 815.00 | 778.00 | 851.00 | -1.09 | 0.09 |
| 467.00 | 478.00 | 494.00 | -1.12 | 0.02 |
| 797.00 | 867.00 | 848.00 | -1.10 | 0.05 |
| 853.00 | 1025.00 | 979.33 | 1.14 | 0.01 |
| 345.00 | 398.00 | 392.67 | 1.21 | 0.00 |
| 1355.00 | 1361.00 | 1409.00 | -1.09 | 0.02 |
| 4.00 | 4.00 | 3.33 | -1.60 | 0.33 |
| 720.00 | 886.00 | 841.00 | 1.07 | 0.20 |
| 194.00 | 209.00 | 212.00 | 1.09 | 0.23 |
| 720.00 | 730.00 | 707.00 | -1.12 | 0.06 |
| 270.00 | 303.00 | 294.33 | -1.02 | 0.70 |
| 279.00 | 307.00 | 290.33 | 4.05 | 0.00 |
| 379.00 | 508.00 | 442.33 | 1.14 | 0.05 |
| 915.00 | 943.00 | 976.33 | 1.10 | 0.03 |
| 2181.00 | 2578.00 | 2452.00 | 1.18 | 0.00 |
| 425.00 | 512.00 | 473.00 | 1.05 | 0.39 |
| 1039.00 | 1191.00 | 1176.67 | 1.20 | 0.00 |
| 1865.00 | 2032.00 | 1938.67 | -1.03 | 0.53 |
| 104.00 | 137.00 | 119.33 | -1.37 | 0.00 |
| 1631.00 | 1739.00 | 1677.67 | -1.13 | 0.00 |
| 1085.00 | 1134.00 | 1185.67 | 1.06 | 0.18 |
| 574.00 | 501.00 | 530.33 | -1.22 | 0.00 |
| 112.00 | 118.00 | 122.00 | 1.22 | 0.03 |
| 738.00 | 712.00 | 755.33 | -1.00 | 0.96 |
| 3417.00 | 3459.00 | 3461.33 | -1.21 | 0.00 |
| 814.00 | 910.00 | 906.33 | 1.29 | 0.00 |
| 207.00 | 200.00 | 197.67 | 1.22 | 0.01 |
| 3.00 | 3.00 | 5.33 | 1.05 | 1.00 |
| 61.00 | 58.00 | 60.33 | -3.08 | 0.00 |
| 685.00 | 649.00 | 683.00 | -1.09 | 0.11 |
| 7478.00 | 6964.00 | 7476.67 | 1.09 | 0.05 |
| 1511.00 | 1724.00 | 1735.33 | 1.27 | 0.00 |
| 26.00 | 27.00 | 28.67 | 1.23 | 0.24 |
| 1616.00 | 1844.00 | 1835.00 | 1.24 | 0.00 |
| 1932.00 | 2217.00 | 2163.00 | 1.25 | 0.00 |
| 3766.00 | 4049.00 | 3956.33 | -1.14 | 0.00 |
| 1579.00 | 1768.00 | 1703.00 | -1.14 | 0.01 |
| 302.00 | 354.00 | 344.67 | 1.05 | 0.52 |

| | | | | |
|---|---|---|---|---|
| 5177.00 | 5487.00 | 5350.67 | 1.12 | 0.00 |
| 1168.00 | 1231.00 | 1226.00 | 1.00 | 0.95 |
| 539.00 | 651.00 | 639.67 | 1.01 | 0.82 |
| 926.00 | 1035.00 | 1035.00 | 1.03 | 0.47 |
| 3012.00 | 2848.00 | 2933.33 | -1.12 | 0.01 |
| 3755.00 | 3873.00 | 3835.33 | -1.12 | 0.00 |
| 2338.00 | 2579.00 | 2576.33 | -1.14 | 0.00 |
| 1137.00 | 1344.00 | 1289.33 | 1.04 | 0.33 |
| 574.00 | 514.00 | 577.33 | 1.04 | 0.62 |
| 402.00 | 378.00 | 397.33 | 1.25 | 0.00 |
| 8213.00 | 8329.00 | 8341.00 | -1.12 | 0.01 |
| 1327.00 | 1399.00 | 1425.33 | 1.14 | 0.00 |
| 112.00 | 103.00 | 115.00 | -1.09 | 0.37 |
| 6.00 | 6.00 | 6.33 | -1.17 | 0.76 |
| 95.00 | 103.00 | 93.33 | -1.08 | 0.48 |
| 488.00 | 475.00 | 489.67 | -1.10 | 0.06 |
| 437.00 | 473.00 | 481.67 | 1.14 | 0.02 |
| 479.00 | 520.00 | 505.00 | 1.22 | 0.00 |
| 764.00 | 811.00 | 805.00 | -1.06 | 0.24 |
| 237.00 | 218.00 | 238.33 | -1.10 | 0.15 |
| 1091.00 | 1302.00 | 1146.33 | 1.08 | 0.20 |
| 176.00 | 225.00 | 213.67 | -1.01 | 0.88 |
| 894.00 | 860.00 | 871.67 | -1.34 | 0.00 |
| 86.00 | 106.00 | 105.33 | 1.34 | 0.00 |
| 121.00 | 101.00 | 108.33 | 1.18 | 0.09 |
| 936.00 | 966.00 | 952.00 | 1.02 | 0.62 |
| 554.00 | 565.00 | 575.67 | -1.05 | 0.34 |
| 45.00 | 62.00 | 53.33 | -1.73 | 0.00 |
| 424.00 | 487.00 | 475.33 | 1.54 | 0.00 |
| 1126.00 | 1138.00 | 1149.00 | 1.19 | 0.00 |
| 721.00 | 863.00 | 833.00 | 1.16 | 0.00 |
| 37.00 | 37.00 | 41.00 | 1.30 | 0.10 |
| 151.00 | 147.00 | 155.67 | -1.11 | 0.15 |
| 1175.00 | 1381.00 | 1318.00 | 1.15 | 0.00 |
| 35.00 | 44.00 | 43.33 | 1.24 | 0.15 |
| 4.00 | 5.00 | 4.33 | 2.07 | 0.17 |
| 28.00 | 40.00 | 38.00 | 1.20 | 0.26 |
| 37.00 | 40.00 | 42.67 | -1.02 | 0.91 |
| 4.00 | 7.00 | 4.33 | -1.01 | 1.00 |
| 854.00 | 810.00 | 865.33 | -1.04 | 0.43 |
| 124.00 | 104.00 | 119.00 | 1.13 | 0.19 |
| 917.00 | 1021.00 | 1026.33 | 1.14 | 0.01 |
| 2644.00 | 2803.00 | 2826.00 | -1.02 | 0.53 |
| 333.00 | 409.00 | 391.67 | 1.20 | 0.00 |
| 7686.00 | 8932.00 | 8278.67 | 1.13 | 0.00 |
| 1315.00 | 1424.00 | 1362.00 | -1.09 | 0.05 |
| 424.00 | 414.00 | 417.67 | 1.14 | 0.02 |
| 149.00 | 127.00 | 129.00 | -1.62 | 0.00 |

| | | | | |
|---|---|---|---|---|
| 43.00 | 42.00 | 47.33 | 1.11 | 0.46 |
| 90.00 | 76.00 | 77.33 | -1.82 | 0.00 |
| 185.00 | 199.00 | 201.33 | -1.52 | 0.00 |
| 1291.00 | 1286.00 | 1294.00 | 1.11 | 0.06 |
| 10.00 | 7.00 | 9.00 | 1.02 | 1.00 |
| 332.00 | 331.00 | 323.00 | -1.12 | 0.09 |
| 1420.00 | 1397.00 | 1431.00 | -1.46 | 0.00 |
| 793.00 | 806.00 | 772.33 | 1.08 | 0.17 |
| 349.00 | 385.00 | 377.00 | -1.14 | 0.02 |
| 1046.00 | 862.00 | 946.00 | -1.70 | 0.00 |
| 17.00 | 10.00 | 11.33 | -1.48 | 0.13 |
| 0.00 | 1.00 | 0.67 | -5.28 | 0.01 |
| 10.00 | 2.00 | 6.33 | 1.43 | 0.39 |
| 35.00 | 35.00 | 41.33 | -1.06 | 0.69 |
| 2771.00 | 3283.00 | 3114.33 | 1.11 | 0.02 |
| 11.00 | 6.00 | 9.67 | 1.14 | 0.69 |
| 644.00 | 766.00 | 723.00 | 1.04 | 0.51 |
| 515.00 | 472.00 | 506.00 | 1.01 | 0.87 |
| 894.00 | 890.00 | 889.00 | 1.01 | 0.82 |
| 113.00 | 109.00 | 108.67 | 1.03 | 0.79 |
| 46.00 | 77.00 | 61.33 | 1.26 | 0.09 |
| 1.00 | 2.00 | 1.00 | -2.21 | 0.35 |
| 5.00 | 9.00 | 8.00 | 1.38 | 0.36 |
| 163.00 | 161.00 | 155.00 | 1.52 | 0.00 |
| 20.00 | 21.00 | 27.33 | 2.67 | 0.00 |
| 667.00 | 821.00 | 776.33 | 1.14 | 0.02 |
| 159.00 | 188.00 | 180.67 | -1.36 | 0.00 |
| 117.00 | 119.00 | 119.33 | 1.02 | 0.84 |
| 2474.00 | 2434.00 | 2501.00 | -1.27 | 0.00 |
| 3409.00 | 3453.00 | 3538.00 | 1.05 | 0.13 |
| 1553.00 | 1338.00 | 1423.67 | -1.34 | 0.00 |
| 542.00 | 602.00 | 578.67 | -1.06 | 0.23 |
| 1608.00 | 1710.00 | 1668.00 | 1.02 | 0.57 |
| 397.00 | 471.00 | 439.00 | -1.06 | 0.30 |
| 2096.00 | 2091.00 | 2040.67 | -1.62 | 0.00 |
| 1131.00 | 1136.00 | 1162.00 | -1.08 | 0.05 |
| 8335.00 | 9195.00 | 8989.00 | 1.09 | 0.02 |
| 562.00 | 643.00 | 634.00 | 1.10 | 0.07 |
| 6.00 | 7.00 | 5.67 | -4.57 | 0.00 |
| 283.00 | 304.00 | 301.33 | 1.09 | 0.17 |
| 10.00 | 17.00 | 13.67 | 1.04 | 0.92 |
| 525.00 | 457.00 | 504.33 | -1.07 | 0.28 |
| 2423.00 | 2565.00 | 2605.00 | 1.13 | 0.00 |
| 1905.00 | 1908.00 | 1845.67 | -1.20 | 0.00 |
| 724.00 | 793.00 | 801.67 | 1.18 | 0.00 |
| 560.00 | 699.00 | 659.67 | 1.04 | 0.52 |
| 392.00 | 512.00 | 478.67 | -1.04 | 0.53 |
| 243.00 | 258.00 | 245.67 | 1.32 | 0.00 |

| | | | | |
|---|---|---|---|---|
| 16.00 | 12.00 | 13.67 | 1.04 | 0.92 |
| 358.00 | 397.00 | 388.00 | 1.04 | 0.48 |
| 32.00 | 24.00 | 31.33 | -1.15 | 0.38 |
| 460.00 | 500.00 | 518.00 | 1.24 | 0.00 |
| 2013.00 | 2150.00 | 2180.00 | -1.11 | 0.01 |
| 1789.00 | 2169.00 | 2024.33 | -1.01 | 0.86 |
| 3.00 | 2.00 | 2.33 | -1.70 | 0.37 |
| 1560.00 | 1611.00 | 1588.33 | -1.06 | 0.18 |
| 278.00 | 276.00 | 275.33 | 1.01 | 0.94 |
| 146.00 | 117.00 | 124.33 | 1.01 | 0.96 |
| 599.00 | 634.00 | 641.00 | 1.08 | 0.12 |
| 240.00 | 267.00 | 253.33 | 1.22 | 0.00 |
| 629.00 | 658.00 | 682.33 | -1.03 | 0.50 |
| 1497.00 | 1570.00 | 1580.67 | -1.04 | 0.25 |
| 412.00 | 461.00 | 449.67 | 1.06 | 0.25 |
| 1780.00 | 2056.00 | 2002.33 | -1.01 | 0.85 |
| 509.00 | 603.00 | 558.33 | -1.00 | 0.99 |
| 1.00 | 1.00 | 0.67 | -7.02 | 0.01 |
| 670.00 | 826.00 | 780.67 | 1.18 | 0.00 |
| 43.00 | 47.00 | 50.33 | -1.50 | 0.00 |
| 4116.00 | 4347.00 | 4435.67 | -1.00 | 0.90 |
| 688.00 | 769.00 | 792.00 | 1.24 | 0.00 |
| 2.00 | 6.00 | 5.00 | -1.01 | 1.00 |
| 2852.00 | 3210.00 | 3189.33 | 1.13 | 0.00 |
| 201.00 | 245.00 | 236.00 | 1.04 | 0.57 |
| 655.00 | 756.00 | 695.00 | -1.28 | 0.00 |
| 1037.00 | 924.00 | 978.33 | -1.64 | 0.00 |
| 347.00 | 381.00 | 376.00 | -1.02 | 0.79 |
| 464.00 | 406.00 | 444.33 | 1.01 | 0.84 |
| 701.00 | 804.00 | 781.00 | 1.11 | 0.02 |
| 1726.00 | 1841.00 | 1726.00 | -1.04 | 0.53 |
| 1570.00 | 1467.00 | 1524.33 | -1.32 | 0.00 |
| 1852.00 | 2169.00 | 2055.67 | 1.33 | 0.00 |
| 1.00 | 0.00 | 0.67 | -2.29 | 0.46 |
| 4.00 | 5.00 | 6.00 | -1.73 | 0.10 |
| 20.00 | 26.00 | 23.00 | -1.19 | 0.35 |
| 223.00 | 212.00 | 215.33 | -1.11 | 0.13 |
| 342.00 | 415.00 | 368.33 | -1.02 | 0.73 |
| 54.00 | 65.00 | 65.33 | -1.12 | 0.31 |
| 1562.00 | 1678.00 | 1622.00 | -1.11 | 0.03 |
| 814.00 | 772.00 | 805.00 | 1.03 | 0.54 |
| 4032.00 | 4492.00 | 4302.00 | -1.02 | 0.53 |
| 12.00 | 10.00 | 11.00 | 1.35 | 0.30 |
| 9601.00 | 11045.00 | 10799.33 | 1.02 | 0.59 |
| 1140.00 | 1148.00 | 1174.67 | -1.07 | 0.08 |
| 210.00 | 216.00 | 214.33 | -1.14 | 0.04 |
| 1.00 | 2.00 | 4.33 | 1.56 | 0.43 |
| 5.00 | 17.00 | 11.33 | -1.49 | 0.14 |

| | | | | |
|---|---|---|---|---|
| 100.00 | 113.00 | 108.00 | -1.10 | 0.27 |
| 203.00 | 259.00 | 238.33 | 1.17 | 0.03 |
| 924.00 | 1165.00 | 1140.33 | 1.18 | 0.01 |
| 946.00 | 866.00 | 918.67 | -1.15 | 0.00 |
| 335.00 | 366.00 | 358.33 | 1.36 | 0.00 |
| 2600.00 | 2900.00 | 2812.33 | 1.05 | 0.23 |
| 181.00 | 170.00 | 183.33 | 1.14 | 0.09 |
| 1056.00 | 1037.00 | 1124.67 | -1.14 | 0.01 |
| 1328.00 | 1368.00 | 1382.33 | -1.10 | 0.02 |
| 271.00 | 260.00 | 277.67 | 1.10 | 0.26 |
| 1133.00 | 1154.00 | 1169.67 | -1.10 | 0.02 |
| 268.00 | 274.00 | 278.33 | -1.06 | 0.37 |
| 983.00 | 896.00 | 963.67 | -1.02 | 0.65 |
| 1790.00 | 1682.00 | 1744.67 | 1.03 | 0.54 |
| 784.00 | 871.00 | 810.33 | 1.04 | 0.44 |
| 1959.00 | 1746.00 | 1833.33 | -1.30 | 0.00 |
| 445.00 | 434.00 | 490.67 | -1.07 | 0.32 |
| 60.00 | 77.00 | 67.00 | 1.35 | 0.01 |
| 233.00 | 210.00 | 225.67 | 1.03 | 0.66 |
| 1245.00 | 1321.00 | 1295.00 | -1.29 | 0.00 |
| 487.00 | 470.00 | 474.00 | 1.06 | 0.28 |
| 433.00 | 382.00 | 424.67 | 1.22 | 0.00 |
| 2683.00 | 3019.00 | 2868.00 | 1.56 | 0.00 |
| 713.00 | 670.00 | 719.33 | -1.16 | 0.00 |
| 411.00 | 455.00 | 439.33 | 1.11 | 0.06 |
| 1005.00 | 953.00 | 974.67 | -1.51 | 0.00 |
| 177.00 | 209.00 | 197.67 | 1.09 | 0.23 |
| 446.00 | 539.00 | 501.00 | 1.11 | 0.06 |
| 284.00 | 372.00 | 319.67 | 1.02 | 0.75 |
| 4.00 | 4.00 | 3.00 | 1.11 | 1.00 |
| 967.00 | 1113.00 | 1031.00 | 1.16 | 0.00 |
| 2412.00 | 2252.00 | 2294.33 | -1.08 | 0.13 |
| 197.00 | 219.00 | 210.00 | -1.09 | 0.20 |
| 458.00 | 501.00 | 491.33 | -1.38 | 0.00 |
| 341.00 | 377.00 | 354.00 | -1.06 | 0.39 |
| 724.00 | 767.00 | 751.33 | -1.07 | 0.14 |
| 4157.00 | 3969.00 | 4091.33 | -1.12 | 0.00 |
| 1627.00 | 1852.00 | 1862.67 | 1.31 | 0.00 |
| 1011.00 | 1201.00 | 1160.00 | 1.17 | 0.00 |
| 510.00 | 551.00 | 575.33 | 1.07 | 0.19 |
| 253.00 | 267.00 | 264.33 | 1.08 | 0.25 |
| 950.00 | 938.00 | 912.67 | -1.08 | 0.17 |
| 711.00 | 858.00 | 819.00 | 1.18 | 0.00 |
| 577.00 | 662.00 | 620.00 | -1.04 | 0.46 |
| 468.00 | 522.00 | 496.00 | -1.20 | 0.00 |
| 1014.00 | 1059.00 | 1065.33 | 1.14 | 0.00 |
| 135.00 | 110.00 | 125.33 | -1.15 | 0.12 |
| 674.00 | 640.00 | 698.00 | -1.18 | 0.00 |

| | | | | |
|---|---|---|---|---|
| 2323.00 | 2716.00 | 2575.00 | 1.12 | 0.01 |
| 1213.00 | 1338.00 | 1330.00 | -1.02 | 0.69 |
| 1107.00 | 1100.00 | 1129.67 | 1.03 | 0.49 |
| 408.00 | 466.00 | 456.67 | 1.17 | 0.00 |
| 607.00 | 728.00 | 695.67 | 1.24 | 0.00 |
| 1021.00 | 1003.00 | 1046.67 | -1.11 | 0.01 |
| 3043.00 | 3518.00 | 3441.00 | -1.02 | 0.54 |
| 797.00 | 971.00 | 883.67 | -1.00 | 0.96 |
| 71.00 | 69.00 | 73.33 | -1.21 | 0.06 |
| 342.00 | 403.00 | 372.00 | 1.00 | 0.96 |
| 72.00 | 85.00 | 84.67 | -1.17 | 0.17 |
| 2099.00 | 2160.00 | 2174.33 | -1.38 | 0.00 |
| 23.00 | 40.00 | 39.00 | -1.19 | 0.29 |
| 22.00 | 31.00 | 25.00 | -1.01 | 1.00 |
| 197.00 | 181.00 | 198.67 | -1.14 | 0.06 |
| 1206.00 | 1075.00 | 1121.67 | -1.24 | 0.00 |
| 62.00 | 70.00 | 69.67 | 1.40 | 0.00 |
| 521.00 | 618.00 | 567.00 | -1.01 | 0.81 |
| 38.00 | 57.00 | 48.67 | 1.24 | 0.12 |
| 5.00 | 2.00 | 3.33 | -4.43 | 0.00 |
| 3707.00 | 3983.00 | 3957.67 | 1.17 | 0.00 |
| 353.00 | 328.00 | 364.33 | -1.30 | 0.00 |
| 6.00 | 7.00 | 8.33 | 1.23 | 0.56 |
| 363.00 | 408.00 | 387.33 | -1.19 | 0.00 |
| 334.00 | 399.00 | 377.00 | -1.08 | 0.17 |
| 2063.00 | 2412.00 | 2361.33 | 1.08 | 0.08 |
| 1228.00 | 1510.00 | 1460.67 | 1.13 | 0.01 |
| 72.00 | 81.00 | 80.00 | -1.00 | 1.00 |
| 407.00 | 451.00 | 419.33 | -1.12 | 0.08 |
| 1288.00 | 1628.00 | 1533.00 | 1.21 | 0.00 |
| 1289.00 | 1377.00 | 1435.67 | 1.09 | 0.06 |
| 1756.00 | 1907.00 | 1886.67 | -1.06 | 0.11 |
| 59.00 | 77.00 | 77.33 | -1.18 | 0.14 |
| 3040.00 | 3523.00 | 3339.33 | 1.05 | 0.21 |
| 1983.00 | 1993.00 | 1954.00 | 1.06 | 0.30 |
| 10.00 | 16.00 | 12.33 | 1.04 | 0.91 |
| 2386.00 | 2384.00 | 2442.67 | -1.09 | 0.02 |
| 2594.00 | 2474.00 | 2600.67 | 1.10 | 0.01 |
| 331.00 | 411.00 | 353.67 | -1.00 | 1.00 |
| 39.00 | 26.00 | 34.67 | 1.13 | 0.50 |
| 13.00 | 11.00 | 12.00 | 1.36 | 0.27 |
| 482.00 | 551.00 | 517.33 | -1.02 | 0.71 |
| 914.00 | 1022.00 | 1031.00 | 1.07 | 0.19 |
| 913.00 | 1001.00 | 983.00 | 1.06 | 0.19 |
| 836.00 | 825.00 | 891.67 | -1.26 | 0.00 |
| 10.00 | 4.00 | 9.67 | 3.93 | 0.00 |
| 88.00 | 100.00 | 108.67 | 1.88 | 0.00 |
| 4178.00 | 4222.00 | 4209.67 | -1.02 | 0.54 |

| | | | | |
|---|---|---|---|---|
| 8.00 | 2.00 | 4.33 | -1.24 | 0.72 |
| 6402.00 | 7045.00 | 7059.00 | 1.24 | 0.00 |
| 8.00 | 9.00 | 9.33 | 1.44 | 0.25 |
| 303.00 | 269.00 | 281.67 | -1.04 | 0.58 |
| 76.00 | 92.00 | 80.67 | -1.31 | 0.02 |
| 361.00 | 370.00 | 371.00 | -1.15 | 0.01 |
| 214.00 | 217.00 | 222.67 | 1.14 | 0.05 |
| 894.00 | 928.00 | 935.00 | -1.08 | 0.08 |
| 218.00 | 241.00 | 217.33 | 1.02 | 0.81 |
| 1714.00 | 1658.00 | 1706.00 | -1.03 | 0.49 |
| 480.00 | 542.00 | 522.00 | -1.04 | 0.42 |
| 737.00 | 845.00 | 808.00 | -1.16 | 0.00 |
| 1016.00 | 1156.00 | 1150.00 | 1.10 | 0.04 |
| 306.00 | 346.00 | 335.33 | 1.05 | 0.50 |
| 17.00 | 21.00 | 22.33 | -1.81 | 0.00 |
| 100.00 | 111.00 | 112.67 | -1.07 | 0.44 |
| 283.00 | 305.00 | 302.00 | 1.31 | 0.00 |
| 123.00 | 135.00 | 128.33 | -1.05 | 0.57 |
| 2120.00 | 2261.00 | 2150.67 | -1.32 | 0.00 |
| 83.00 | 86.00 | 88.33 | 1.05 | 0.66 |
| 244.00 | 252.00 | 272.33 | -1.30 | 0.00 |
| 874.00 | 868.00 | 886.00 | 1.07 | 0.21 |
| 83.00 | 93.00 | 89.67 | -1.33 | 0.00 |
| 1042.00 | 917.00 | 1002.00 | -1.05 | 0.30 |
| 725.00 | 645.00 | 683.33 | -1.30 | 0.00 |
| 2252.00 | 2196.00 | 2359.67 | 1.07 | 0.27 |
| 7.00 | 7.00 | 12.33 | -1.59 | 0.12 |
| 10.00 | 2.00 | 4.67 | 1.06 | 1.00 |
| 115.00 | 137.00 | 125.67 | 1.13 | 0.15 |
| 180.00 | 231.00 | 211.00 | -1.03 | 0.63 |
| 47.00 | 37.00 | 43.33 | -1.22 | 0.14 |
| 1042.00 | 1096.00 | 1100.33 | 1.02 | 0.65 |
| 43.00 | 50.00 | 53.67 | -1.15 | 0.27 |
| 218.00 | 259.00 | 245.67 | -1.60 | 0.00 |
| 680.00 | 731.00 | 692.00 | -1.49 | 0.00 |
| 1169.00 | 1190.00 | 1170.67 | -1.06 | 0.18 |
| 205.00 | 214.00 | 196.33 | -1.27 | 0.01 |
| 1246.00 | 1220.00 | 1222.00 | -1.20 | 0.00 |
| 656.00 | 742.00 | 691.33 | 1.18 | 0.00 |
| 709.00 | 779.00 | 757.33 | 1.09 | 0.04 |
| 1206.00 | 1304.00 | 1286.67 | 1.20 | 0.00 |
| 265.00 | 266.00 | 270.67 | 1.16 | 0.02 |
| 251.00 | 269.00 | 279.00 | -1.27 | 0.00 |
| 393.00 | 419.00 | 431.00 | -1.18 | 0.00 |
| 76.00 | 59.00 | 65.00 | -1.44 | 0.00 |
| 52.00 | 51.00 | 53.67 | -1.15 | 0.25 |
| 250.00 | 289.00 | 277.33 | -1.10 | 0.12 |
| 1068.00 | 1209.00 | 1113.00 | -1.16 | 0.00 |

| | | | | |
|---|---|---|---|---|
| 1570.00 | 1609.00 | 1606.67 | 1.16 | 0.00 |
| 2.00 | 4.00 | 3.67 | -1.37 | 0.56 |
| 4.00 | 3.00 | 5.00 | -1.21 | 0.73 |
| 114.00 | 152.00 | 131.33 | 1.71 | 0.00 |
| 23.00 | 30.00 | 23.67 | -1.48 | 0.02 |
| 1430.00 | 1375.00 | 1352.00 | 1.01 | 0.82 |
| 20.00 | 15.00 | 16.33 | -1.12 | 0.71 |
| 35.00 | 34.00 | 37.67 | 1.16 | 0.36 |
| 2651.00 | 2594.00 | 2641.67 | -1.09 | 0.04 |
| 1223.00 | 1174.00 | 1250.00 | -1.03 | 0.47 |
| 390.00 | 522.00 | 484.33 | 1.35 | 0.00 |
| 12.00 | 38.00 | 19.00 | 1.25 | 0.52 |
| 80.00 | 85.00 | 88.67 | 1.32 | 0.01 |
| 1.00 | 4.00 | 2.00 | -1.65 | 0.46 |
| 1955.00 | 2226.00 | 2128.00 | 1.23 | 0.00 |
| 350.00 | 394.00 | 376.67 | 1.07 | 0.24 |
| 12.00 | 12.00 | 13.00 | -1.61 | 0.03 |
| 283.00 | 299.00 | 299.00 | -1.03 | 0.67 |
| 10.00 | 5.00 | 6.00 | -1.45 | 0.31 |
| 802.00 | 796.00 | 804.33 | 1.13 | 0.01 |
| 507.00 | 595.00 | 580.00 | 1.08 | 0.16 |
| 670.00 | 787.00 | 760.67 | 1.12 | 0.03 |
| 977.00 | 1111.00 | 1058.00 | 1.13 | 0.00 |
| 1929.00 | 2140.00 | 2121.33 | 1.19 | 0.00 |
| 925.00 | 1003.00 | 935.33 | 1.20 | 0.00 |
| 2431.00 | 2502.00 | 2583.33 | 1.04 | 0.33 |
| 17.00 | 8.00 | 10.67 | -1.14 | 0.73 |
| 2690.00 | 2911.00 | 2888.33 | 1.01 | 0.79 |
| 826.00 | 864.00 | 866.00 | -1.10 | 0.03 |
| 13298.00 | 15379.00 | 14492.67 | -1.24 | 0.00 |
| 6616.00 | 7133.00 | 6959.67 | 1.30 | 0.00 |
| 3.00 | 3.00 | 3.67 | -2.53 | 0.01 |
| 57.00 | 69.00 | 59.00 | 1.07 | 0.61 |
| 5603.00 | 5861.00 | 5755.33 | -1.16 | 0.00 |
| 5.00 | 5.00 | 5.33 | -1.26 | 0.63 |
| 331.00 | 350.00 | 360.67 | 1.15 | 0.01 |
| 1433.00 | 1483.00 | 1472.33 | -1.18 | 0.00 |
| 971.00 | 942.00 | 1009.67 | -1.09 | 0.07 |
| 396.00 | 352.00 | 369.67 | -1.37 | 0.00 |
| 15.00 | 12.00 | 15.00 | 1.11 | 0.68 |
| 22.00 | 11.00 | 16.33 | 1.10 | 0.80 |
| 1044.00 | 1029.00 | 1083.00 | -2.00 | 0.00 |
| 654.00 | 640.00 | 638.00 | 1.02 | 0.71 |
| 6969.00 | 6572.00 | 6638.33 | -1.08 | 0.10 |
| 311.00 | 312.00 | 297.00 | -1.10 | 0.20 |
| 30.00 | 19.00 | 25.00 | 1.08 | 0.72 |
| 1089.00 | 977.00 | 1065.00 | 1.01 | 0.82 |
| 880.00 | 928.00 | 955.33 | -1.21 | 0.00 |

| | | | | |
|---|---|---|---|---|
| 660.00 | 638.00 | 688.33 | 1.45 | 0.00 |
| 33.00 | 36.00 | 30.67 | -1.20 | 0.27 |
| 18292.00 | 18211.00 | 18294.67 | -1.01 | 0.70 |
| 3.00 | 10.00 | 4.67 | -1.29 | 0.61 |
| 7267.00 | 6795.00 | 6914.33 | -1.35 | 0.00 |
| 774.00 | 845.00 | 817.67 | 1.00 | 0.98 |
| 1516.00 | 1613.00 | 1565.33 | -1.10 | 0.02 |
| 257.00 | 329.00 | 283.33 | 1.23 | 0.01 |
| 154.00 | 187.00 | 161.00 | 1.17 | 0.09 |
| 673.00 | 641.00 | 673.67 | 1.09 | 0.08 |
| 136.00 | 164.00 | 150.00 | 1.06 | 0.45 |
| 377.00 | 397.00 | 392.67 | 1.06 | 0.25 |
| 1326.00 | 1371.00 | 1390.67 | 1.10 | 0.02 |
| 435.00 | 552.00 | 501.00 | 1.20 | 0.00 |
| 237.00 | 279.00 | 277.00 | 1.14 | 0.05 |
| 275.00 | 341.00 | 310.67 | 1.15 | 0.03 |
| 28.00 | 36.00 | 33.33 | -1.12 | 0.43 |
| 44.00 | 55.00 | 56.00 | -1.03 | 0.85 |
| 76.00 | 65.00 | 72.00 | -1.30 | 0.01 |
| 1341.00 | 1721.00 | 1562.67 | 1.21 | 0.00 |
| 1632.00 | 1726.00 | 1666.33 | -1.02 | 0.64 |
| 21.00 | 10.00 | 13.00 | 1.38 | 0.32 |
| 819.00 | 860.00 | 890.00 | -1.00 | 0.97 |
| 17.00 | 26.00 | 20.33 | -1.18 | 0.45 |
| 7.00 | 4.00 | 6.33 | 1.04 | 1.00 |
| 8.00 | 10.00 | 7.33 | -2.60 | 0.00 |
| 516.00 | 571.00 | 554.33 | 1.02 | 0.75 |
| 1082.00 | 1113.00 | 1121.67 | 1.01 | 0.78 |
| 12.00 | 9.00 | 9.67 | -2.56 | 0.00 |
| 417.00 | 430.00 | 434.67 | 1.19 | 0.01 |
| 602.00 | 736.00 | 689.33 | -1.04 | 0.41 |
| 94.00 | 76.00 | 81.33 | -1.03 | 0.79 |
| 2599.00 | 2973.00 | 2870.67 | 1.07 | 0.07 |
| 1572.00 | 1604.00 | 1574.67 | -1.04 | 0.38 |
| 266.00 | 276.00 | 286.00 | 1.07 | 0.31 |
| 1332.00 | 1470.00 | 1405.00 | 1.12 | 0.01 |
| 989.00 | 1006.00 | 1009.33 | 1.25 | 0.00 |
| 955.00 | 1055.00 | 1054.67 | 1.16 | 0.00 |
| 2120.00 | 2197.00 | 2210.00 | 1.06 | 0.11 |
| 315.00 | 295.00 | 301.33 | 1.03 | 0.71 |
| 4068.00 | 4371.00 | 4200.00 | 1.03 | 0.44 |
| 0.00 | 0.00 | 0.33 | -4.69 | 0.13 |
| 121.00 | 126.00 | 123.67 | 1.31 | 0.00 |
| 1274.00 | 1166.00 | 1183.33 | -1.68 | 0.00 |
| 1.00 | 5.00 | 3.00 | -1.34 | 0.67 |
| 284.00 | 245.00 | 277.67 | -1.51 | 0.00 |
| 1025.00 | 1194.00 | 1122.00 | 1.12 | 0.01 |
| 460.00 | 500.00 | 500.33 | 1.01 | 0.91 |

| | | | | |
|---|---|---|---|---|
| 907.00 | 857.00 | 941.00 | -1.05 | 0.35 |
| 471.00 | 571.00 | 522.33 | -1.12 | 0.04 |
| 699.00 | 635.00 | 690.00 | -1.12 | 0.08 |
| 364.00 | 410.00 | 415.33 | 1.02 | 0.74 |
| 2104.00 | 2721.00 | 2430.67 | -1.09 | 0.10 |
| 7.00 | 12.00 | 8.00 | -2.09 | 0.00 |
| 3783.00 | 4349.00 | 4286.33 | 1.05 | 0.24 |
| 377.00 | 322.00 | 343.33 | -1.13 | 0.11 |
| 14.00 | 9.00 | 12.67 | -2.07 | 0.00 |
| 388.00 | 409.00 | 418.33 | -1.00 | 1.00 |
| 1.00 | 0.00 | 0.67 | -3.14 | 0.18 |
| 533.00 | 567.00 | 570.67 | 1.04 | 0.43 |
| 12390.00 | 12624.00 | 12692.00 | -1.20 | 0.00 |
| 199.00 | 250.00 | 230.67 | 1.27 | 0.00 |
| 359.00 | 370.00 | 392.33 | 1.08 | 0.18 |
| 240.00 | 270.00 | 266.00 | -1.09 | 0.18 |
| 183.00 | 196.00 | 189.67 | 1.05 | 0.53 |
| 6.00 | 1.00 | 5.67 | -1.30 | 0.54 |
| 24.00 | 22.00 | 23.33 | -1.01 | 1.00 |
| 140.00 | 158.00 | 154.67 | -1.04 | 0.59 |
| 31.00 | 25.00 | 28.00 | 1.24 | 0.25 |
| 1450.00 | 1600.00 | 1522.67 | 1.20 | 0.00 |
| 417.00 | 491.00 | 485.33 | 1.17 | 0.01 |
| 644.00 | 664.00 | 660.33 | -1.01 | 0.82 |
| 799.00 | 875.00 | 897.67 | 1.22 | 0.00 |
| 1060.00 | 1181.00 | 1188.00 | 1.29 | 0.00 |
| 763.00 | 751.00 | 782.33 | 1.03 | 0.58 |
| 566.00 | 605.00 | 624.00 | 1.16 | 0.00 |
| 1479.00 | 1484.00 | 1590.67 | 1.20 | 0.00 |
| 312.00 | 378.00 | 349.33 | -1.11 | 0.10 |
| 1522.00 | 1581.00 | 1623.67 | 1.02 | 0.65 |
| 1027.00 | 1102.00 | 1073.33 | 1.14 | 0.00 |
| 181.00 | 232.00 | 219.33 | 1.03 | 0.68 |
| 323.00 | 383.00 | 378.00 | 1.20 | 0.00 |
| 610.00 | 682.00 | 711.67 | 1.14 | 0.04 |
| 1945.00 | 2100.00 | 2105.67 | 1.03 | 0.50 |
| 7532.00 | 7786.00 | 8133.00 | -1.01 | 0.91 |
| 5391.00 | 4829.00 | 5185.33 | -1.19 | 0.00 |
| 3657.00 | 3698.00 | 3764.33 | -1.18 | 0.00 |
| 7.00 | 8.00 | 8.00 | -3.09 | 0.00 |
| 8.00 | 13.00 | 14.33 | -1.56 | 0.07 |
| 565.00 | 631.00 | 620.67 | 1.00 | 0.94 |
| 763.00 | 878.00 | 814.00 | 1.08 | 0.10 |
| 37.00 | 25.00 | 36.00 | -1.08 | 0.63 |
| 224.00 | 256.00 | 266.67 | 1.09 | 0.27 |
| 1339.00 | 1487.00 | 1484.00 | 1.21 | 0.00 |
| 71.00 | 83.00 | 74.00 | -1.21 | 0.13 |
| 1428.00 | 1590.00 | 1584.67 | -1.08 | 0.04 |

| | | | | |
|---|---|---|---|---|
| 67.00 | 69.00 | 68.67 | -1.22 | 0.06 |
| 2119.00 | 2111.00 | 2087.00 | 1.28 | 0.00 |
| 412.00 | 430.00 | 452.67 | 1.65 | 0.00 |
| 900.00 | 998.00 | 981.67 | -1.07 | 0.09 |
| 1247.00 | 1214.00 | 1266.00 | 1.03 | 0.44 |
| 6393.00 | 6354.00 | 6539.00 | -1.17 | 0.00 |
| 130.00 | 145.00 | 140.00 | 1.22 | 0.02 |
| 2247.00 | 2292.00 | 2315.67 | -1.11 | 0.00 |
| 2238.00 | 2265.00 | 2272.67 | 1.04 | 0.28 |
| 666.00 | 633.00 | 664.00 | -1.19 | 0.00 |
| 227.00 | 290.00 | 255.00 | 1.26 | 0.00 |
| 945.00 | 1012.00 | 994.00 | 1.10 | 0.08 |
| 1139.00 | 1175.00 | 1148.00 | 1.04 | 0.39 |
| 2.00 | 2.00 | 2.67 | -1.74 | 0.30 |
| 2.00 | 3.00 | 2.67 | -1.13 | 1.00 |
| 13543.00 | 13021.00 | 13146.33 | -1.50 | 0.00 |
| 409.00 | 394.00 | 397.00 | 1.38 | 0.00 |
| 20.00 | 25.00 | 22.33 | 1.16 | 0.44 |
| 14.00 | 23.00 | 20.33 | 1.02 | 1.00 |
| 9.00 | 10.00 | 9.33 | -5.73 | 0.00 |
| 582.00 | 556.00 | 563.33 | 1.23 | 0.00 |
| 0.00 | 0.00 | 0.33 | -6.16 | 0.04 |
| 488.00 | 446.00 | 463.67 | 1.20 | 0.00 |
| 1002.00 | 1033.00 | 1017.67 | 1.08 | 0.08 |
| 570.00 | 607.00 | 611.00 | 1.15 | 0.00 |
| 21439.00 | 22085.00 | 23166.33 | 2.13 | 0.00 |
| 473.00 | 461.00 | 471.33 | 1.40 | 0.00 |
| 45.00 | 39.00 | 38.67 | -1.69 | 0.00 |
| 35.00 | 37.00 | 37.33 | 1.55 | 0.01 |
| 7.00 | 6.00 | 7.67 | 2.03 | 0.06 |
| 6.00 | 8.00 | 5.67 | -1.95 | 0.04 |
| 228.00 | 200.00 | 217.33 | -1.06 | 0.45 |
| 3362.00 | 3469.00 | 3467.33 | 1.19 | 0.00 |
| 2447.00 | 2342.00 | 2473.00 | -1.08 | 0.06 |
| 3971.00 | 4287.00 | 4173.67 | -1.07 | 0.06 |
| 10219.00 | 10982.00 | 10577.33 | 1.11 | 0.00 |
| 3960.00 | 3951.00 | 4030.33 | -1.19 | 0.00 |
| 424.00 | 523.00 | 514.67 | 1.24 | 0.00 |
| 1027.00 | 1236.00 | 1270.33 | 1.21 | 0.00 |
| 2795.00 | 2760.00 | 2825.33 | 1.03 | 0.47 |
| 1120.00 | 1374.00 | 1315.67 | 1.10 | 0.06 |
| 24079.00 | 23971.00 | 24494.67 | -1.15 | 0.00 |
| 0.00 | 0.00 | 0.33 | -10.58 | 0.00 |
| 16.00 | 7.00 | 15.33 | -1.72 | 0.01 |
| 25.00 | 15.00 | 24.00 | -1.59 | 0.00 |
| 379.00 | 352.00 | 365.00 | 1.39 | 0.00 |
| 0.00 | 0.00 | 0.00 | -33.26 | 0.00 |
| 412.00 | 472.00 | 450.00 | 1.06 | 0.27 |

| | | | | |
|---|---|---|---|---|
| 619.00 | 776.00 | 721.33 | 1.04 | 0.45 |
| 66.00 | 81.00 | 77.67 | 1.17 | 0.16 |
| 256.00 | 295.00 | 280.33 | -1.15 | 0.02 |
| 2588.00 | 2645.00 | 2676.00 | 1.13 | 0.00 |
| 762.00 | 875.00 | 817.00 | -1.24 | 0.00 |
| 321.00 | 273.00 | 307.67 | 1.07 | 0.35 |
| 564.00 | 598.00 | 593.33 | -1.13 | 0.01 |
| 789.00 | 838.00 | 819.67 | -1.00 | 0.96 |
| 6520.00 | 6449.00 | 6533.67 | 1.03 | 0.52 |
| 4649.00 | 4575.00 | 4633.67 | 1.09 | 0.02 |
| 2041.00 | 1890.00 | 2058.00 | 1.20 | 0.00 |
| 3477.00 | 3162.00 | 3402.67 | 1.01 | 0.90 |
| 2483.00 | 2899.00 | 2661.67 | 1.04 | 0.36 |
| 8.00 | 18.00 | 11.33 | -1.19 | 0.55 |
| 1312.00 | 1480.00 | 1392.00 | 1.09 | 0.03 |
| 14.00 | 19.00 | 14.00 | -1.06 | 0.92 |
| 77.00 | 45.00 | 57.00 | 1.01 | 0.97 |
| 4664.00 | 4754.00 | 4990.67 | -1.09 | 0.07 |
| 18.00 | 19.00 | 20.67 | -1.36 | 0.10 |
| 1254.00 | 1312.00 | 1370.00 | 1.05 | 0.28 |
| 49.00 | 38.00 | 43.33 | -1.27 | 0.08 |
| 744.00 | 685.00 | 713.67 | -1.23 | 0.00 |
| 206.00 | 188.00 | 190.67 | 1.11 | 0.21 |
| 1889.00 | 2067.00 | 1933.33 | -1.07 | 0.16 |
| 1202.00 | 1242.00 | 1291.33 | 1.10 | 0.03 |
| 525.00 | 504.00 | 529.00 | 1.10 | 0.13 |
| 137.00 | 181.00 | 167.00 | -1.16 | 0.05 |
| 217.00 | 337.00 | 290.00 | -1.32 | 0.00 |
| 3015.00 | 2988.00 | 3180.67 | 1.07 | 0.16 |
| 301.00 | 412.00 | 373.00 | 1.06 | 0.41 |
| 582.00 | 550.00 | 554.00 | -1.16 | 0.01 |
| 1768.00 | 1765.00 | 1797.67 | -1.09 | 0.04 |
| 253.00 | 256.00 | 267.67 | 1.26 | 0.00 |
| 75.00 | 83.00 | 73.67 | -1.10 | 0.46 |
| 432.00 | 468.00 | 474.00 | 1.10 | 0.06 |
| 1748.00 | 1707.00 | 1750.00 | -1.23 | 0.00 |
| 32.00 | 21.00 | 27.67 | 1.67 | 0.01 |
| 13985.00 | 14696.00 | 14034.00 | -1.03 | 0.48 |
| 254.00 | 280.00 | 278.67 | 1.10 | 0.11 |
| 636.00 | 622.00 | 665.00 | 1.15 | 0.01 |
| 1467.00 | 1491.00 | 1483.33 | -1.14 | 0.00 |
| 303.00 | 384.00 | 351.00 | 1.17 | 0.02 |
| 6589.00 | 6580.00 | 6778.67 | 1.03 | 0.49 |
| 2572.00 | 2329.00 | 2418.33 | -1.24 | 0.00 |
| 2162.00 | 2252.00 | 2315.33 | -1.06 | 0.13 |
| 2.00 | 2.00 | 2.67 | -1.62 | 0.40 |
| 397.00 | 525.00 | 481.67 | -1.04 | 0.47 |
| 203.00 | 221.00 | 228.00 | 1.05 | 0.47 |

| | | | | |
|---|---|---|---|---|
| 268.00 | 339.00 | 298.67 | 1.09 | 0.29 |
| 436.00 | 483.00 | 482.33 | 1.02 | 0.76 |
| 829.00 | 837.00 | 888.67 | 1.00 | 1.00 |
| 33.00 | 24.00 | 29.67 | -1.14 | 0.45 |
| 335.00 | 354.00 | 377.33 | 1.12 | 0.07 |
| 124.00 | 110.00 | 122.67 | -1.04 | 0.63 |
| 202.00 | 183.00 | 205.00 | 1.48 | 0.00 |
| 26.00 | 33.00 | 29.33 | 1.13 | 0.50 |
| 1528.00 | 1518.00 | 1580.00 | 1.08 | 0.05 |
| 954.00 | 1085.00 | 1082.67 | 1.03 | 0.62 |
| 311.00 | 369.00 | 362.33 | 1.53 | 0.00 |
| 1676.00 | 1883.00 | 1907.00 | 1.19 | 0.00 |
| 2242.00 | 2318.00 | 2431.33 | 1.11 | 0.02 |
| 327.00 | 351.00 | 370.67 | -1.18 | 0.00 |
| 31.00 | 19.00 | 27.67 | 1.51 | 0.04 |
| 438.00 | 411.00 | 438.33 | -1.22 | 0.00 |
| 2984.00 | 3167.00 | 3204.67 | -1.13 | 0.00 |
| 2762.00 | 2774.00 | 2887.00 | 1.39 | 0.00 |
| 336.00 | 315.00 | 327.00 | -1.10 | 0.13 |
| 213.00 | 228.00 | 227.00 | 1.40 | 0.00 |
| 1140.00 | 1180.00 | 1184.67 | -1.14 | 0.00 |
| 1192.00 | 1292.00 | 1228.00 | -1.06 | 0.21 |
| 249.00 | 265.00 | 283.00 | 1.01 | 0.92 |
| 1343.00 | 1441.00 | 1378.67 | -1.19 | 0.00 |
| 1499.00 | 1710.00 | 1666.00 | 1.00 | 0.97 |
| 1868.00 | 2184.00 | 2074.67 | 1.11 | 0.01 |
| 918.00 | 912.00 | 983.33 | 1.09 | 0.08 |
| 587.00 | 742.00 | 661.00 | 1.07 | 0.26 |
| 3.00 | 5.00 | 3.33 | -1.31 | 0.68 |
| 283.00 | 290.00 | 295.00 | 1.21 | 0.00 |
| 826.00 | 939.00 | 882.67 | 1.11 | 0.03 |
| 259.00 | 271.00 | 268.00 | 1.39 | 0.00 |
| 86.00 | 102.00 | 105.33 | 1.36 | 0.00 |
| 429.00 | 458.00 | 471.00 | 1.15 | 0.04 |
| 463.00 | 462.00 | 486.67 | 1.08 | 0.19 |
| 346.00 | 437.00 | 407.00 | 1.12 | 0.07 |
| 28.00 | 21.00 | 28.00 | -1.48 | 0.01 |
| 663.00 | 621.00 | 662.33 | 1.09 | 0.14 |
| 828.00 | 873.00 | 873.33 | 1.20 | 0.00 |
| 237.00 | 302.00 | 272.33 | -1.29 | 0.00 |
| 3753.00 | 3798.00 | 3756.33 | 1.03 | 0.43 |
| 135.00 | 143.00 | 149.00 | 1.32 | 0.00 |
| 462.00 | 494.00 | 504.67 | -1.05 | 0.38 |
| 36.00 | 43.00 | 40.67 | 1.03 | 0.88 |
| 24.00 | 11.00 | 19.67 | -1.17 | 0.51 |
| 1735.00 | 1530.00 | 1669.33 | -1.10 | 0.06 |
| 736.00 | 713.00 | 729.33 | -1.09 | 0.07 |
| 168.00 | 167.00 | 178.00 | -1.13 | 0.09 |

| | | | | |
|---|---|---|---|---|
| 3521.00 | 3439.00 | 3579.00 | -1.02 | 0.68 |
| 337.00 | 392.00 | 379.33 | -1.01 | 0.91 |
| 243.00 | 267.00 | 277.33 | -1.04 | 0.59 |
| 615.00 | 728.00 | 720.00 | 1.11 | 0.12 |
| 602.00 | 655.00 | 634.67 | 1.10 | 0.05 |
| 10.00 | 5.00 | 6.67 | 1.77 | 0.16 |
| 4.00 | 3.00 | 2.33 | -1.29 | 0.81 |
| 851.00 | 773.00 | 800.33 | 1.00 | 0.97 |
| 113.00 | 125.00 | 116.33 | -1.41 | 0.00 |
| 1067.00 | 1047.00 | 1080.67 | -1.38 | 0.00 |
| 489.00 | 558.00 | 553.00 | 1.02 | 0.68 |
| 417.00 | 441.00 | 427.33 | 1.00 | 0.98 |
| 4100.00 | 4213.00 | 4050.00 | -1.07 | 0.12 |
| 80.00 | 92.00 | 82.67 | 1.56 | 0.00 |
| 1295.00 | 1320.00 | 1345.67 | 1.40 | 0.00 |
| 880.00 | 860.00 | 896.33 | 1.09 | 0.05 |
| 423.00 | 455.00 | 453.67 | -1.10 | 0.09 |
| 691.00 | 657.00 | 727.67 | -1.05 | 0.37 |
| 877.00 | 988.00 | 973.00 | -1.01 | 0.76 |
| 594.00 | 641.00 | 651.00 | 1.05 | 0.51 |
| 2760.00 | 2952.00 | 2825.33 | 1.11 | 0.01 |
| 1597.00 | 1655.00 | 1641.00 | -1.56 | 0.00 |
| 150.00 | 163.00 | 157.00 | -1.34 | 0.00 |
| 22.00 | 25.00 | 24.33 | 1.01 | 1.00 |
| 294.00 | 279.00 | 286.67 | 1.01 | 0.84 |
| 1105.00 | 1148.00 | 1204.00 | 1.05 | 0.30 |
| 27.00 | 21.00 | 21.00 | 1.27 | 0.25 |
| 718.00 | 742.00 | 758.33 | -1.01 | 0.79 |
| 295.00 | 294.00 | 297.33 | 1.21 | 0.01 |
| 2414.00 | 2720.00 | 2685.00 | 1.14 | 0.00 |
| 409.00 | 355.00 | 404.67 | -1.12 | 0.06 |
| 38.00 | 37.00 | 35.33 | -1.22 | 0.16 |
| 142.00 | 144.00 | 146.67 | 1.05 | 0.54 |
| 64.00 | 52.00 | 59.33 | -1.46 | 0.00 |
| 1001.00 | 1193.00 | 1132.67 | 1.03 | 0.54 |
| 1.00 | 0.00 | 1.67 | -1.39 | 0.78 |
| 10.00 | 16.00 | 12.33 | -1.42 | 0.15 |
| 210.00 | 232.00 | 230.00 | 1.03 | 0.69 |
| 535.00 | 524.00 | 551.00 | 1.32 | 0.00 |
| 522.00 | 572.00 | 569.00 | -1.01 | 0.78 |
| 21.00 | 9.00 | 21.33 | 2.61 | 0.00 |
| 2416.00 | 2283.00 | 2322.67 | 1.02 | 0.62 |
| 1066.00 | 1013.00 | 1055.67 | -1.05 | 0.26 |
| 2254.00 | 2239.00 | 2282.67 | -1.13 | 0.00 |
| 87.00 | 107.00 | 94.33 | -1.29 | 0.01 |
| 0.00 | 2.00 | 1.00 | -3.41 | 0.07 |
| 640.00 | 643.00 | 650.00 | 1.02 | 0.66 |
| 60.00 | 76.00 | 62.00 | -1.05 | 0.71 |

| | | | | |
|---|---|---|---|---|
| 352.00 | 353.00 | 349.33 | -1.97 | 0.00 |
| 394.00 | 396.00 | 421.33 | 1.07 | 0.26 |
| 842.00 | 892.00 | 939.67 | 1.15 | 0.01 |
| 6378.00 | 6182.00 | 6485.67 | -1.06 | 0.10 |
| 16.00 | 15.00 | 16.00 | -1.06 | 0.85 |
| 556.00 | 559.00 | 580.33 | 1.04 | 0.47 |
| 1285.00 | 1286.00 | 1360.00 | 1.14 | 0.00 |
| 1205.00 | 1359.00 | 1329.33 | 1.07 | 0.11 |
| 255.00 | 297.00 | 290.67 | -1.15 | 0.02 |
| 19.00 | 14.00 | 14.67 | -1.13 | 0.69 |
| 5416.00 | 5624.00 | 5392.67 | -1.23 | 0.00 |
| 6594.00 | 7175.00 | 7004.00 | 1.06 | 0.06 |
| 23.00 | 23.00 | 22.00 | 1.14 | 0.49 |
| 885.00 | 849.00 | 887.33 | 1.10 | 0.04 |
| 1764.00 | 1734.00 | 1793.33 | -1.17 | 0.00 |
| 1022.00 | 1117.00 | 1130.33 | -1.04 | 0.44 |
| 910.00 | 889.00 | 938.00 | -1.06 | 0.27 |
| 1043.00 | 1222.00 | 1152.33 | 1.04 | 0.37 |
| 478.00 | 531.00 | 530.33 | -1.06 | 0.26 |
| 4167.00 | 4199.00 | 4312.33 | -1.33 | 0.00 |
| 907.00 | 987.00 | 971.00 | -1.31 | 0.00 |
| 90.00 | 94.00 | 102.67 | -1.06 | 0.58 |
| 166.00 | 191.00 | 179.33 | -1.08 | 0.30 |
| 6.00 | 16.00 | 9.33 | 2.09 | 0.04 |
| 1021.00 | 1028.00 | 1058.33 | 1.18 | 0.00 |
| 1973.00 | 2185.00 | 2175.00 | 1.16 | 0.00 |
| 4.00 | 1.00 | 2.33 | -2.77 | 0.04 |
| 197.00 | 182.00 | 190.00 | -1.12 | 0.13 |
| 2.00 | 3.00 | 2.67 | 1.30 | 0.79 |
| 76.00 | 88.00 | 82.33 | 1.04 | 0.75 |
| 425.00 | 503.00 | 489.67 | 1.05 | 0.40 |
| 407.00 | 544.00 | 487.00 | 1.29 | 0.00 |
| 527.00 | 498.00 | 540.33 | -1.10 | 0.05 |
| 46.00 | 44.00 | 46.00 | 1.53 | 0.00 |
| 27.00 | 51.00 | 33.33 | -1.24 | 0.31 |
| 664.00 | 749.00 | 742.33 | -1.09 | 0.11 |
| 2441.00 | 2460.00 | 2538.33 | -1.03 | 0.43 |
| 56.00 | 43.00 | 45.00 | -1.66 | 0.00 |
| 289.00 | 300.00 | 313.00 | 1.04 | 0.52 |
| 355.00 | 399.00 | 398.00 | 1.27 | 0.00 |
| 932.00 | 851.00 | 911.67 | -1.28 | 0.00 |
| 1326.00 | 1194.00 | 1292.67 | -1.12 | 0.04 |
| 674.00 | 674.00 | 678.00 | 1.24 | 0.00 |
| 268.00 | 309.00 | 295.33 | -1.16 | 0.01 |
| 681.00 | 737.00 | 717.67 | -1.01 | 0.87 |
| 1252.00 | 1383.00 | 1340.00 | 1.27 | 0.00 |
| 890.00 | 1033.00 | 1022.67 | 1.18 | 0.00 |
| 645.00 | 712.00 | 736.00 | 1.04 | 0.46 |

| | | | | |
|---|---|---|---|---|
| 214.00 | 170.00 | 197.67 | 1.06 | 0.48 |
| 40.00 | 29.00 | 32.00 | 1.20 | 0.30 |
| 289.00 | 299.00 | 303.33 | 1.03 | 0.58 |
| 555.00 | 625.00 | 630.00 | -1.04 | 0.42 |
| 51.00 | 72.00 | 71.00 | -1.16 | 0.23 |
| 6.00 | 11.00 | 8.00 | -1.43 | 0.26 |
| 154.00 | 134.00 | 142.67 | -1.09 | 0.34 |
| 1577.00 | 1692.00 | 1641.67 | -1.02 | 0.62 |
| 1368.00 | 1539.00 | 1412.67 | -1.13 | 0.01 |
| 12.00 | 10.00 | 12.33 | -1.80 | 0.01 |
| 1540.00 | 1554.00 | 1607.00 | -1.06 | 0.17 |
| 334.00 | 330.00 | 324.67 | -1.20 | 0.00 |
| 1619.00 | 1665.00 | 1716.67 | -1.05 | 0.32 |
| 169.00 | 181.00 | 177.33 | -1.18 | 0.02 |
| 32.00 | 31.00 | 32.33 | 1.14 | 0.48 |
| 790.00 | 772.00 | 802.00 | 1.37 | 0.00 |
| 568.00 | 546.00 | 577.33 | 1.19 | 0.00 |
| 318.00 | 326.00 | 347.67 | 1.04 | 0.51 |
| 788.00 | 891.00 | 837.33 | 1.13 | 0.02 |
| 178.00 | 125.00 | 147.67 | -1.78 | 0.00 |
| 1104.00 | 1255.00 | 1158.67 | 1.02 | 0.72 |
| 133.00 | 146.00 | 138.00 | -1.04 | 0.63 |
| 321.00 | 321.00 | 320.33 | -1.13 | 0.07 |
| 849.00 | 893.00 | 867.00 | 1.10 | 0.06 |
| 14.00 | 11.00 | 14.67 | -1.54 | 0.04 |
| 2338.00 | 2650.00 | 2486.00 | 1.39 | 0.00 |
| 47.00 | 65.00 | 59.67 | 1.55 | 0.00 |
| 28.00 | 27.00 | 25.67 | -1.92 | 0.00 |
| 798.00 | 840.00 | 811.33 | -1.14 | 0.02 |
| 13.00 | 22.00 | 15.00 | 1.20 | 0.46 |
| 1.00 | 4.00 | 2.67 | -1.62 | 0.39 |
| 13742.00 | 14846.00 | 15765.33 | 1.34 | 0.00 |
| 6.00 | 9.00 | 9.00 | 1.39 | 0.31 |
| 3777.00 | 4035.00 | 4356.33 | 1.41 | 0.00 |
| 6.00 | 8.00 | 5.67 | -1.01 | 1.00 |
| 299.00 | 291.00 | 291.00 | -1.09 | 0.17 |
| 575.00 | 676.00 | 623.67 | -1.07 | 0.21 |
| 676.00 | 709.00 | 686.33 | 1.26 | 0.00 |
| 1207.00 | 1192.00 | 1235.00 | 1.15 | 0.00 |
| 1134.00 | 1127.00 | 1152.33 | -1.10 | 0.02 |
| 2057.00 | 2126.00 | 2082.33 | 1.22 | 0.00 |
| 2581.00 | 2745.00 | 2743.67 | 1.10 | 0.01 |
| 3105.00 | 3783.00 | 3493.67 | -1.01 | 0.84 |
| 1119.00 | 998.00 | 1045.67 | 1.07 | 0.29 |
| 7650.00 | 7100.00 | 7177.67 | -1.46 | 0.00 |
| 575.00 | 618.00 | 616.67 | -1.07 | 0.18 |
| 973.00 | 940.00 | 991.00 | -1.39 | 0.00 |
| 666.00 | 684.00 | 684.33 | -1.19 | 0.00 |

| | | | | |
|---|---|---|---|---|
| 3.00 | 0.00 | 1.00 | -3.40 | 0.07 |
| 89.00 | 117.00 | 107.33 | 1.18 | 0.15 |
| 281.00 | 313.00 | 323.33 | 1.30 | 0.00 |
| 474.00 | 440.00 | 438.33 | -1.10 | 0.14 |
| 4584.00 | 4809.00 | 4778.67 | -1.25 | 0.00 |
| 2.00 | 3.00 | 2.00 | -2.60 | 0.06 |
| 4327.00 | 4167.00 | 4272.67 | -1.32 | 0.00 |
| 155.00 | 150.00 | 150.00 | -1.03 | 0.72 |
| 14.00 | 7.00 | 10.67 | -1.14 | 0.72 |
| 8804.00 | 9214.00 | 9455.00 | 1.10 | 0.02 |
| 795.00 | 751.00 | 801.33 | 1.15 | 0.00 |
| 2361.00 | 2343.00 | 2407.00 | -1.17 | 0.00 |
| 768.00 | 739.00 | 755.67 | -1.23 | 0.00 |
| 694.00 | 696.00 | 730.33 | 1.06 | 0.33 |
| 494.00 | 590.00 | 579.00 | 1.98 | 0.00 |
| 18.00 | 35.00 | 26.00 | -1.30 | 0.12 |
| 203.00 | 161.00 | 193.00 | 1.13 | 0.14 |
| 5.00 | 5.00 | 6.00 | -2.12 | 0.02 |
| 32.00 | 45.00 | 40.00 | -1.22 | 0.15 |
| 19.00 | 24.00 | 21.33 | -1.15 | 0.51 |
| 5.00 | 5.00 | 7.00 | 2.25 | 0.05 |
| 55.00 | 73.00 | 74.67 | 1.01 | 1.00 |
| 4.00 | 4.00 | 4.33 | -1.54 | 0.30 |
| 427.00 | 390.00 | 386.33 | -1.11 | 0.14 |
| 1798.00 | 1751.00 | 1826.00 | -1.30 | 0.00 |
| 592.00 | 619.00 | 631.33 | 1.09 | 0.07 |
| 5.00 | 6.00 | 5.00 | 1.62 | 0.31 |
| 1673.00 | 1655.00 | 1735.33 | -1.14 | 0.00 |
| 1142.00 | 1367.00 | 1249.33 | -1.01 | 0.81 |
| 3511.00 | 3469.00 | 3555.67 | -1.14 | 0.00 |
| 2435.00 | 2520.00 | 2452.33 | 1.02 | 0.65 |
| 12.00 | 16.00 | 14.67 | 1.11 | 0.67 |
| 3.00 | 4.00 | 3.00 | -1.55 | 0.41 |
| 56.00 | 65.00 | 58.33 | -1.11 | 0.41 |
| 5393.00 | 6024.00 | 5533.00 | 1.02 | 0.79 |
| 4007.00 | 4127.00 | 4181.67 | -1.03 | 0.44 |
| 2316.00 | 2519.00 | 2439.00 | -1.12 | 0.00 |
| 5532.00 | 5265.00 | 5377.33 | -1.10 | 0.02 |
| 2669.00 | 2710.00 | 2704.67 | -1.12 | 0.00 |
| 634.00 | 659.00 | 637.00 | -1.18 | 0.00 |
| 337.00 | 327.00 | 338.33 | -1.24 | 0.00 |
| 2.00 | 4.00 | 3.33 | -1.31 | 0.68 |
| 399.00 | 393.00 | 398.00 | -1.91 | 0.00 |
| 2149.00 | 1954.00 | 2031.00 | -1.17 | 0.00 |
| 147.00 | 166.00 | 148.67 | -1.31 | 0.00 |
| 4.00 | 5.00 | 6.33 | -1.75 | 0.08 |
| 1241.00 | 1168.00 | 1200.00 | -1.03 | 0.49 |
| 49.00 | 54.00 | 49.33 | 1.52 | 0.00 |

| | | | | |
|---|---|---|---|---|
| 676.00 | 803.00 | 764.33 | 1.16 | 0.00 |
| 798.00 | 858.00 | 831.67 | -1.03 | 0.51 |
| 19.00 | 15.00 | 15.33 | 1.33 | 0.23 |
| 541.00 | 534.00 | 561.33 | 1.01 | 0.80 |
| 948.00 | 977.00 | 1007.33 | 1.23 | 0.00 |
| 908.00 | 869.00 | 887.33 | -1.18 | 0.00 |
| 4348.00 | 4451.00 | 4615.33 | 1.15 | 0.00 |
| 1270.00 | 1329.00 | 1331.00 | 1.36 | 0.00 |
| 6317.00 | 6881.00 | 6920.33 | 1.02 | 0.65 |
| 1638.00 | 1801.00 | 1804.00 | 1.20 | 0.00 |
| 471.00 | 470.00 | 489.00 | 1.05 | 0.33 |
| 1288.00 | 1496.00 | 1526.00 | 1.27 | 0.00 |
| 885.00 | 996.00 | 1013.33 | -1.30 | 0.00 |
| 2035.00 | 2127.00 | 2251.67 | 1.17 | 0.00 |
| 3152.00 | 3430.00 | 3335.00 | 1.10 | 0.02 |
| 1576.00 | 1496.00 | 1584.00 | 1.06 | 0.27 |
| 16.00 | 21.00 | 19.33 | -1.38 | 0.08 |
| 4375.00 | 4997.00 | 4675.00 | -1.07 | 0.07 |
| 1512.00 | 1667.00 | 1616.33 | 1.08 | 0.08 |
| 57.00 | 64.00 | 55.33 | 1.02 | 0.88 |
| 1.00 | 1.00 | 1.33 | -1.24 | 1.00 |
| 4295.00 | 4805.00 | 4466.00 | 1.18 | 0.00 |
| 3483.00 | 3922.00 | 3679.00 | 1.09 | 0.05 |
| 791.00 | 854.00 | 817.33 | -1.02 | 0.74 |
| 710.00 | 656.00 | 709.33 | -1.18 | 0.00 |
| 1534.00 | 1725.00 | 1632.00 | 1.03 | 0.49 |
| 778.00 | 901.00 | 851.00 | 1.07 | 0.15 |
| 42.00 | 45.00 | 40.00 | -1.00 | 1.00 |
| 1744.00 | 1785.00 | 1802.00 | 1.07 | 0.15 |
| 1445.00 | 1402.00 | 1419.67 | 1.13 | 0.01 |
| 1811.00 | 1871.00 | 1888.33 | 1.07 | 0.07 |
| 1038.00 | 1050.00 | 1073.33 | -1.10 | 0.05 |
| 543.00 | 463.00 | 511.00 | 1.01 | 0.91 |
| 963.00 | 869.00 | 915.67 | 1.18 | 0.00 |
| 1984.00 | 1786.00 | 1878.67 | 1.13 | 0.02 |
| 97.00 | 100.00 | 93.67 | 1.50 | 0.00 |
| 1494.00 | 1367.00 | 1469.33 | -1.25 | 0.00 |
| 1088.00 | 1148.00 | 1126.00 | -1.07 | 0.09 |
| 1894.00 | 1873.00 | 1901.33 | 1.46 | 0.00 |
| 1.00 | 0.00 | 0.33 | -5.42 | 0.07 |
| 2170.00 | 2137.00 | 2133.33 | -1.10 | 0.03 |
| 1281.00 | 1370.00 | 1358.33 | -1.10 | 0.01 |
| 651.00 | 694.00 | 710.33 | -1.06 | 0.27 |
| 236.00 | 255.00 | 259.33 | 1.01 | 0.89 |
| 111.00 | 170.00 | 135.00 | -1.32 | 0.00 |
| 149.00 | 150.00 | 155.00 | -1.89 | 0.00 |
| 567.00 | 552.00 | 591.00 | 1.11 | 0.04 |
| 50.00 | 40.00 | 51.00 | 1.19 | 0.23 |

| | | | | |
|---|---|---|---|---|
| 7.00 | 8.00 | 11.67 | -2.22 | 0.00 |
| 898.00 | 993.00 | 920.67 | 1.38 | 0.00 |
| 2160.00 | 2357.00 | 2232.00 | -2.14 | 0.00 |
| 3366.00 | 3309.00 | 3346.33 | -1.52 | 0.00 |
| 3174.00 | 3333.00 | 3269.67 | -1.09 | 0.02 |
| 843.00 | 991.00 | 925.33 | 1.30 | 0.00 |
| 28.00 | 27.00 | 29.33 | 1.29 | 0.14 |
| 4520.00 | 4635.00 | 4769.33 | 1.07 | 0.04 |
| 140.00 | 160.00 | 156.00 | 1.07 | 0.39 |
| 803.00 | 849.00 | 867.00 | -1.05 | 0.26 |
| 21.00 | 34.00 | 28.67 | 1.43 | 0.05 |
| 10407.00 | 11218.00 | 11390.67 | 1.18 | 0.00 |
| 55.00 | 36.00 | 49.67 | 1.08 | 0.61 |
| 100.00 | 123.00 | 118.33 | 1.21 | 0.06 |
| 6507.00 | 7080.00 | 7062.67 | 1.17 | 0.00 |
| 1328.00 | 1509.00 | 1505.00 | -1.01 | 0.78 |
| 7086.00 | 7830.00 | 7754.00 | 1.24 | 0.00 |
| 149.00 | 203.00 | 184.67 | -1.12 | 0.14 |
| 11416.00 | 12151.00 | 12281.67 | 1.15 | 0.00 |
| 444.00 | 502.00 | 482.00 | 1.15 | 0.01 |
| 14185.00 | 13612.00 | 14048.33 | -1.40 | 0.00 |
| 1996.00 | 2144.00 | 2100.67 | 1.01 | 0.84 |
| 5.00 | 7.00 | 4.33 | -1.01 | 1.00 |
| 11.00 | 2.00 | 7.33 | 1.03 | 1.00 |
| 4639.00 | 4968.00 | 4852.00 | -1.21 | 0.00 |
| 1486.00 | 1502.00 | 1508.00 | -1.12 | 0.00 |
| 4.00 | 6.00 | 4.33 | 1.27 | 0.69 |
| 5.00 | 8.00 | 5.00 | 1.13 | 0.85 |
| 626.00 | 655.00 | 668.33 | -1.15 | 0.00 |
| 526.00 | 539.00 | 545.00 | 1.21 | 0.00 |
| 687.00 | 678.00 | 698.67 | -1.08 | 0.13 |
| 830.00 | 867.00 | 877.67 | 1.01 | 0.84 |
| 404.00 | 400.00 | 408.00 | -1.17 | 0.01 |
| 25140.00 | 28446.00 | 28304.67 | 1.08 | 0.03 |
| 112.00 | 142.00 | 144.00 | -1.10 | 0.32 |
| 4.00 | 1.00 | 2.67 | -1.50 | 0.51 |
| 4.00 | 5.00 | 6.00 | -1.67 | 0.12 |
| 11.00 | 17.00 | 17.00 | -1.09 | 0.71 |
| 16.00 | 20.00 | 14.33 | -1.34 | 0.22 |
| 4159.00 | 4682.00 | 4399.67 | 1.04 | 0.31 |
| 1857.00 | 2489.00 | 2285.33 | 1.16 | 0.01 |
| 3759.00 | 4369.00 | 4246.67 | 1.17 | 0.00 |
| 1412.00 | 1528.00 | 1568.33 | 1.02 | 0.68 |
| 684.00 | 691.00 | 686.33 | -1.02 | 0.65 |
| 238.00 | 234.00 | 250.00 | -1.11 | 0.12 |
| 99.00 | 89.00 | 103.00 | 1.42 | 0.01 |
| 817.00 | 900.00 | 897.00 | 1.08 | 0.13 |
| 1395.00 | 1445.00 | 1415.67 | -1.04 | 0.39 |

| | | | | |
|---|---|---|---|---|
| 389.00 | 424.00 | 404.33 | -1.08 | 0.22 |
| 312.00 | 286.00 | 297.00 | -1.07 | 0.36 |
| 162.00 | 202.00 | 187.67 | -1.11 | 0.17 |
| 623.00 | 661.00 | 678.33 | -1.08 | 0.12 |
| 705.00 | 852.00 | 808.00 | 1.13 | 0.04 |
| 334.00 | 380.00 | 352.00 | -1.16 | 0.01 |
| 2071.00 | 1924.00 | 2034.33 | -1.39 | 0.00 |
| 429.00 | 547.00 | 487.67 | 1.01 | 0.93 |
| 337.00 | 414.00 | 391.67 | 1.17 | 0.06 |
| 1072.00 | 1080.00 | 1108.00 | -1.01 | 0.77 |
| 931.00 | 944.00 | 928.00 | -1.23 | 0.00 |
| 3253.00 | 3810.00 | 3461.00 | -1.10 | 0.05 |
| 82.00 | 90.00 | 88.00 | 1.13 | 0.23 |
| 4640.00 | 4361.00 | 4418.00 | -1.45 | 0.00 |
| 175.00 | 219.00 | 214.00 | 1.11 | 0.22 |
| 48.00 | 39.00 | 42.33 | -1.11 | 0.49 |
| 3580.00 | 3853.00 | 3832.67 | 1.09 | 0.01 |
| 1266.00 | 1431.00 | 1474.00 | 1.17 | 0.00 |
| 799.00 | 812.00 | 789.00 | -1.12 | 0.03 |
| 713.00 | 780.00 | 722.00 | -1.05 | 0.42 |
| 294.00 | 281.00 | 285.00 | -1.27 | 0.00 |
| 615.00 | 705.00 | 703.00 | 1.14 | 0.02 |
| 436.00 | 429.00 | 470.67 | 1.12 | 0.05 |
| 68.00 | 78.00 | 71.67 | 1.07 | 0.54 |
| 633.00 | 668.00 | 682.67 | 1.21 | 0.00 |
| 1125.00 | 1198.00 | 1194.33 | 1.01 | 0.73 |
| 0.00 | 0.00 | 0.00 | -9.06 | 0.27 |
| 635.00 | 788.00 | 728.00 | 1.11 | 0.04 |
| 382.00 | 411.00 | 429.33 | 1.09 | 0.16 |
| 58.00 | 62.00 | 65.33 | -1.23 | 0.07 |
| 591.00 | 598.00 | 623.67 | -1.02 | 0.70 |
| 900.00 | 1017.00 | 972.00 | -1.03 | 0.49 |
| 352.00 | 329.00 | 353.33 | -1.13 | 0.03 |
| 236.00 | 285.00 | 270.33 | -1.06 | 0.38 |
| 1091.00 | 1157.00 | 1134.00 | -1.20 | 0.00 |
| 225.00 | 246.00 | 246.33 | -1.04 | 0.52 |
| 23.00 | 36.00 | 30.67 | -1.06 | 0.73 |
| 9.00 | 9.00 | 8.33 | 1.12 | 0.78 |
| 1271.00 | 1264.00 | 1230.67 | -1.22 | 0.00 |
| 529.00 | 650.00 | 611.33 | -1.35 | 0.00 |
| 316.00 | 335.00 | 367.00 | 1.10 | 0.22 |
| 573.00 | 530.00 | 587.00 | 1.09 | 0.10 |
| 490.00 | 525.00 | 532.33 | -1.38 | 0.00 |
| 4364.00 | 4460.00 | 4651.33 | -1.34 | 0.00 |
| 16.00 | 19.00 | 18.00 | 1.77 | 0.03 |
| 956.00 | 985.00 | 1019.00 | 1.07 | 0.14 |
| 586.00 | 554.00 | 600.00 | -1.04 | 0.46 |
| 769.00 | 803.00 | 793.33 | 1.04 | 0.37 |

| | | | | |
|---|---|---|---|---|
| 532.00 | 588.00 | 604.00 | -1.03 | 0.63 |
| 174.00 | 203.00 | 211.33 | -1.06 | 0.49 |
| 18.00 | 17.00 | 20.33 | 1.07 | 0.79 |
| 275.00 | 267.00 | 281.33 | 1.13 | 0.06 |
| 324.00 | 318.00 | 314.67 | 1.13 | 0.07 |
| 445.00 | 468.00 | 458.00 | 1.16 | 0.00 |
| 3133.00 | 3125.00 | 3239.00 | -1.28 | 0.00 |
| 6993.00 | 7425.00 | 7378.67 | 1.05 | 0.13 |
| 407.00 | 444.00 | 452.00 | -1.16 | 0.01 |
| 549.00 | 616.00 | 591.67 | -1.15 | 0.00 |
| 1206.00 | 1448.00 | 1276.00 | -1.04 | 0.54 |
| 8.00 | 6.00 | 7.33 | -1.65 | 0.09 |
| 269.00 | 347.00 | 321.67 | 1.04 | 0.53 |
| 182.00 | 182.00 | 184.67 | 1.20 | 0.01 |
| 1134.00 | 1264.00 | 1189.67 | 1.10 | 0.04 |
| 1204.00 | 1236.00 | 1242.67 | -1.04 | 0.36 |
| 232.00 | 257.00 | 257.33 | -1.14 | 0.04 |
| 2069.00 | 2135.00 | 2175.00 | 1.01 | 0.79 |
| 4356.00 | 4593.00 | 4507.67 | -1.13 | 0.00 |
| 444.00 | 502.00 | 504.33 | -1.08 | 0.24 |
| 281.00 | 289.00 | 295.33 | -1.10 | 0.09 |
| 47.00 | 45.00 | 48.67 | 1.36 | 0.03 |
| 853.00 | 957.00 | 946.67 | 1.05 | 0.28 |
| 323.00 | 320.00 | 331.67 | -1.39 | 0.00 |
| 217.00 | 219.00 | 224.33 | -1.01 | 0.86 |
| 2237.00 | 2645.00 | 2599.67 | 1.20 | 0.00 |
| 710.00 | 540.00 | 562.33 | -1.42 | 0.00 |
| 753.00 | 787.00 | 779.67 | -1.16 | 0.00 |
| 408.00 | 468.00 | 438.00 | 1.16 | 0.01 |
| 150.00 | 181.00 | 166.00 | 1.12 | 0.14 |
| 160.00 | 161.00 | 169.00 | -1.16 | 0.03 |
| 1794.00 | 1929.00 | 1940.33 | 1.21 | 0.00 |
| 994.00 | 1030.00 | 977.67 | -1.24 | 0.00 |
| 411.00 | 377.00 | 391.00 | -1.01 | 0.83 |
| 85.00 | 80.00 | 86.67 | 1.04 | 0.71 |
| 3550.00 | 3745.00 | 3761.33 | -1.03 | 0.31 |
| 161.00 | 183.00 | 180.33 | 1.46 | 0.00 |
| 9925.00 | 9339.00 | 9508.33 | -1.44 | 0.00 |
| 2106.00 | 2108.00 | 2167.67 | -1.18 | 0.00 |
| 1359.00 | 1559.00 | 1556.67 | -1.03 | 0.52 |
| 1512.00 | 1524.00 | 1543.00 | -1.14 | 0.00 |
| 5271.00 | 5637.00 | 5423.33 | -1.04 | 0.27 |
| 1077.00 | 1244.00 | 1195.33 | 1.32 | 0.00 |
| 515.00 | 541.00 | 541.00 | 1.09 | 0.11 |
| 2775.00 | 2807.00 | 2981.33 | 1.03 | 0.55 |
| 724.00 | 892.00 | 823.33 | 1.14 | 0.04 |
| 797.00 | 927.00 | 911.00 | -1.03 | 0.60 |
| 3446.00 | 3557.00 | 3687.67 | -1.01 | 0.70 |

| | | | | |
|---|---|---|---|---|
| 146.00 | 179.00 | 169.00 | 1.02 | 0.79 |
| 297.00 | 303.00 | 311.67 | 1.03 | 0.66 |
| 769.00 | 878.00 | 842.00 | 1.14 | 0.00 |
| 600.00 | 768.00 | 685.67 | 1.06 | 0.34 |
| 3427.00 | 3922.00 | 3899.33 | 1.14 | 0.00 |
| 364.00 | 371.00 | 370.67 | -1.04 | 0.43 |
| 925.00 | 1269.00 | 1082.67 | -1.02 | 0.76 |
| 769.00 | 866.00 | 816.00 | 1.02 | 0.66 |
| 1069.00 | 1083.00 | 1104.33 | 1.04 | 0.36 |
| 423.00 | 493.00 | 466.33 | 1.25 | 0.00 |
| 300.00 | 275.00 | 299.00 | -1.15 | 0.03 |
| 942.00 | 1054.00 | 1019.33 | 1.06 | 0.21 |
| 1026.00 | 1246.00 | 1189.33 | 1.16 | 0.00 |
| 955.00 | 1032.00 | 1074.67 | 1.05 | 0.30 |
| 463.00 | 483.00 | 483.00 | -1.03 | 0.55 |
| 656.00 | 712.00 | 710.33 | -1.06 | 0.22 |
| 239.00 | 242.00 | 253.00 | 1.04 | 0.52 |
| 953.00 | 1011.00 | 1020.33 | -1.57 | 0.00 |
| 769.00 | 938.00 | 843.33 | 1.04 | 0.43 |
| 2.00 | 2.00 | 1.33 | -1.71 | 0.55 |
| 1138.00 | 1219.00 | 1223.67 | 1.13 | 0.02 |
| 391.00 | 428.00 | 408.67 | 1.01 | 0.87 |
| 140.00 | 179.00 | 155.33 | 1.07 | 0.47 |
| 706.00 | 928.00 | 786.67 | -1.12 | 0.07 |
| 26.00 | 17.00 | 22.33 | -1.11 | 0.67 |
| 31.00 | 40.00 | 43.00 | 1.02 | 1.00 |
| 3117.00 | 3040.00 | 3149.33 | -1.07 | 0.08 |
| 112.00 | 127.00 | 119.33 | -1.02 | 0.85 |
| 784.00 | 717.00 | 784.00 | 1.09 | 0.11 |
| 4.00 | 1.00 | 2.00 | -1.17 | 1.00 |
| 822.00 | 834.00 | 853.67 | -1.12 | 0.01 |
| 4.00 | 7.00 | 4.00 | 1.07 | 1.00 |
| 2878.00 | 3206.00 | 3040.67 | 1.02 | 0.67 |
| 449.00 | 415.00 | 430.67 | -1.20 | 0.00 |
| 810.00 | 870.00 | 916.67 | 1.01 | 0.88 |
| 431.00 | 504.00 | 487.67 | -1.25 | 0.00 |
| 1446.00 | 1480.00 | 1517.67 | -1.28 | 0.00 |
| 2596.00 | 2279.00 | 2382.33 | -1.17 | 0.00 |
| 824.00 | 1061.00 | 922.00 | 1.13 | 0.06 |
| 683.00 | 776.00 | 730.00 | -1.01 | 0.84 |
| 7.00 | 8.00 | 11.00 | -1.14 | 0.65 |
| 1580.00 | 1629.00 | 1641.33 | -1.14 | 0.00 |
| 2029.00 | 1872.00 | 1980.67 | -1.27 | 0.00 |
| 1044.00 | 1105.00 | 1123.67 | -1.57 | 0.00 |
| 31.00 | 26.00 | 30.67 | -1.11 | 0.54 |
| 1006.00 | 1081.00 | 1050.33 | -1.00 | 0.94 |
| 507.00 | 546.00 | 554.67 | 1.05 | 0.29 |
| 933.00 | 1017.00 | 999.33 | -1.25 | 0.00 |

| | | | | |
|---|---|---|---|---|
| 1439.00 | 1756.00 | 1670.67 | 1.18 | 0.00 |
| 591.00 | 640.00 | 604.67 | -1.14 | 0.01 |
| 531.00 | 551.00 | 570.67 | -1.04 | 0.49 |
| 568.00 | 581.00 | 604.33 | 1.15 | 0.00 |
| 1369.00 | 1408.00 | 1414.00 | 1.08 | 0.07 |
| 58.00 | 62.00 | 64.33 | -1.16 | 0.19 |
| 24.00 | 21.00 | 19.33 | 1.54 | 0.05 |
| 186.00 | 204.00 | 206.00 | -1.24 | 0.00 |
| 442.00 | 518.00 | 497.00 | 1.02 | 0.69 |
| 2.00 | 9.00 | 3.67 | -2.35 | 0.02 |
| 899.00 | 866.00 | 907.67 | -1.11 | 0.01 |
| 262.00 | 351.00 | 320.67 | -1.77 | 0.00 |
| 1918.00 | 1999.00 | 2057.33 | 1.15 | 0.00 |
| 207.00 | 236.00 | 232.00 | -1.04 | 0.56 |
| 583.00 | 701.00 | 661.67 | 1.18 | 0.00 |
| 24.00 | 32.00 | 28.00 | -1.28 | 0.13 |
| 3.00 | 2.00 | 3.33 | -1.11 | 1.00 |
| 82.00 | 90.00 | 97.00 | 1.27 | 0.03 |
| 142.00 | 141.00 | 142.00 | 1.23 | 0.02 |
| 119.00 | 116.00 | 122.33 | 1.21 | 0.03 |
| 2429.00 | 2276.00 | 2388.00 | -1.71 | 0.00 |
| 100.00 | 103.00 | 104.33 | -1.31 | 0.00 |
| 1167.00 | 1090.00 | 1145.00 | -1.01 | 0.76 |
| 3.00 | 13.00 | 9.33 | 2.09 | 0.04 |
| 3833.00 | 3783.00 | 3923.67 | 1.05 | 0.31 |
| 475.00 | 505.00 | 506.00 | -1.04 | 0.38 |
| 41.00 | 34.00 | 41.67 | 1.92 | 0.00 |
| 4009.00 | 3941.00 | 4223.00 | 1.01 | 0.76 |
| 1761.00 | 1710.00 | 1752.67 | -1.23 | 0.00 |
| 833.00 | 898.00 | 920.33 | 1.02 | 0.76 |
| 317.00 | 319.00 | 328.00 | -1.24 | 0.00 |
| 5.00 | 4.00 | 4.33 | 1.16 | 0.84 |
| 614.00 | 639.00 | 622.33 | -1.27 | 0.00 |
| 2304.00 | 2243.00 | 2446.33 | 1.14 | 0.00 |
| 1231.00 | 1361.00 | 1287.00 | 1.14 | 0.00 |
| 1014.00 | 1002.00 | 1037.67 | -1.32 | 0.00 |
| 918.00 | 1001.00 | 1007.67 | -1.11 | 0.01 |
| 4879.00 | 5206.00 | 5126.67 | 1.13 | 0.00 |
| 566.00 | 597.00 | 612.67 | 1.13 | 0.01 |
| 936.00 | 878.00 | 931.00 | 1.17 | 0.00 |
| 622.00 | 660.00 | 654.67 | 1.21 | 0.00 |
| 146.00 | 143.00 | 134.67 | -1.28 | 0.02 |
| 735.00 | 751.00 | 797.33 | -1.03 | 0.60 |
| 1.00 | 1.00 | 2.33 | -1.15 | 1.00 |
| 72.00 | 68.00 | 71.67 | -1.02 | 0.90 |
| 175.00 | 211.00 | 208.67 | 1.13 | 0.13 |
| 28.00 | 27.00 | 28.33 | -1.12 | 0.54 |
| 618.00 | 665.00 | 651.67 | -1.12 | 0.01 |

| | | | | |
|---|---|---|---|---|
| 87.00 | 60.00 | 77.33 | -1.19 | 0.14 |
| 253.00 | 246.00 | 232.00 | -1.22 | 0.03 |
| 1342.00 | 1526.00 | 1491.00 | 1.16 | 0.00 |
| 386.00 | 431.00 | 414.33 | -1.09 | 0.15 |
| 824.00 | 969.00 | 903.67 | 1.09 | 0.07 |
| 2116.00 | 2597.00 | 2309.00 | 1.03 | 0.53 |
| 2798.00 | 3032.00 | 2980.00 | -1.22 | 0.00 |
| 1461.00 | 1461.00 | 1510.00 | -1.10 | 0.02 |
| 89.00 | 78.00 | 83.67 | -1.60 | 0.00 |
| 779.00 | 752.00 | 767.67 | 1.03 | 0.51 |
| 387.00 | 368.00 | 386.67 | 1.01 | 0.80 |
| 2.00 | 1.00 | 1.00 | -1.61 | 0.73 |
| 27.00 | 23.00 | 25.33 | 1.04 | 0.88 |
| 488.00 | 639.00 | 568.00 | 1.04 | 0.51 |
| 513.00 | 587.00 | 554.67 | 1.06 | 0.26 |
| 712.00 | 635.00 | 682.67 | -1.06 | 0.26 |
| 149.00 | 191.00 | 174.33 | -1.25 | 0.01 |
| 6.00 | 4.00 | 4.33 | 1.41 | 0.53 |
| 3471.00 | 3530.00 | 3539.00 | -1.07 | 0.05 |
| 143.00 | 168.00 | 146.67 | 1.11 | 0.24 |
| 71.00 | 59.00 | 70.67 | -1.06 | 0.61 |
| 961.00 | 977.00 | 1002.33 | -1.11 | 0.01 |
| 97.00 | 127.00 | 122.33 | -1.13 | 0.18 |
| 465.00 | 554.00 | 502.67 | 1.28 | 0.00 |
| 4364.00 | 4634.00 | 4481.67 | -1.07 | 0.10 |
| 428.00 | 404.00 | 426.67 | -1.18 | 0.00 |
| 0.00 | 0.00 | 0.00 | -9.06 | 0.27 |
| 129.00 | 134.00 | 136.67 | 1.27 | 0.00 |
| 212.00 | 250.00 | 244.67 | 1.14 | 0.06 |
| 32.00 | 32.00 | 32.33 | -1.14 | 0.39 |
| 73.00 | 72.00 | 74.67 | -1.07 | 0.55 |
| 18.00 | 5.00 | 14.67 | -1.45 | 0.22 |
| 331.00 | 345.00 | 332.00 | 1.04 | 0.47 |
| 4.00 | 1.00 | 2.67 | -1.98 | 0.16 |
| 381.00 | 422.00 | 419.33 | -1.11 | 0.04 |
| 969.00 | 959.00 | 1030.00 | 1.15 | 0.01 |
| 502.00 | 589.00 | 571.33 | -1.03 | 0.61 |
| 1.00 | 0.00 | 1.00 | -2.51 | 0.23 |
| 415.00 | 407.00 | 438.00 | -1.08 | 0.18 |
| 448.00 | 491.00 | 472.33 | 1.12 | 0.04 |
| 1208.00 | 1196.00 | 1223.67 | -1.12 | 0.01 |
| 983.00 | 1015.00 | 1010.00 | -1.09 | 0.04 |
| 249.00 | 255.00 | 253.00 | 1.11 | 0.14 |
| 807.00 | 1040.00 | 950.00 | 1.16 | 0.01 |
| 2653.00 | 2791.00 | 2816.00 | 1.11 | 0.00 |
| 92.00 | 114.00 | 99.00 | 1.20 | 0.07 |
| 346.00 | 320.00 | 311.33 | -1.31 | 0.00 |
| 1443.00 | 1403.00 | 1474.33 | 1.05 | 0.22 |

| | | | | |
|---|---|---|---|---|
| 3.00 | 6.00 | 5.67 | -1.59 | 0.25 |
| 115.00 | 81.00 | 93.00 | -1.79 | 0.00 |
| 1176.00 | 1238.00 | 1218.67 | -1.22 | 0.00 |
| 829.00 | 818.00 | 830.67 | -1.13 | 0.01 |
| 147.00 | 156.00 | 150.33 | -1.50 | 0.00 |
| 4.00 | 6.00 | 4.00 | -1.09 | 1.00 |
| 164.00 | 168.00 | 170.00 | -1.38 | 0.00 |
| 30.00 | 56.00 | 38.00 | 1.40 | 0.10 |
| 1044.00 | 1087.00 | 1092.67 | -1.10 | 0.03 |
| 211.00 | 182.00 | 202.00 | -1.60 | 0.00 |
| 2470.00 | 2412.00 | 2432.33 | -1.41 | 0.00 |
| 432.00 | 494.00 | 462.33 | -1.18 | 0.02 |
| 5865.00 | 6280.00 | 6338.00 | 1.21 | 0.00 |
| 6762.00 | 7355.00 | 7276.33 | 1.11 | 0.00 |
| 6.00 | 8.00 | 5.33 | 1.31 | 0.58 |
| 3.00 | 6.00 | 3.33 | 1.91 | 0.31 |
| 13.00 | 22.00 | 18.33 | -1.01 | 1.00 |
| 7.00 | 11.00 | 8.33 | -1.41 | 0.26 |
| 1.00 | 1.00 | 0.67 | -4.42 | 0.04 |
| 54.00 | 55.00 | 60.00 | -1.07 | 0.56 |
| 751.00 | 740.00 | 774.67 | 1.04 | 0.44 |
| 4429.00 | 4947.00 | 4862.33 | 1.10 | 0.02 |
| 9501.00 | 10505.00 | 10561.00 | 1.54 | 0.00 |
| 119.00 | 130.00 | 132.33 | -1.00 | 1.00 |
| 1370.00 | 1618.00 | 1564.33 | 1.13 | 0.01 |
| 760.00 | 843.00 | 842.00 | 1.16 | 0.02 |
| 682.00 | 738.00 | 715.00 | -1.13 | 0.01 |
| 1655.00 | 1733.00 | 1765.00 | 1.01 | 0.78 |
| 433.00 | 409.00 | 399.33 | 1.04 | 0.59 |
| 393.00 | 403.00 | 408.00 | -1.15 | 0.02 |
| 41.00 | 51.00 | 44.33 | 1.03 | 0.87 |
| 1574.00 | 1507.00 | 1621.67 | -1.21 | 0.00 |
| 6291.00 | 6556.00 | 6711.33 | 1.07 | 0.05 |
| 244.00 | 234.00 | 251.33 | 1.64 | 0.00 |
| 5694.00 | 5974.00 | 5925.33 | -1.14 | 0.00 |
| 721.00 | 704.00 | 735.33 | -1.12 | 0.01 |
| 662.00 | 784.00 | 735.00 | -1.04 | 0.47 |
| 530.00 | 637.00 | 601.67 | 1.07 | 0.19 |
| 1808.00 | 2035.00 | 1932.33 | 1.15 | 0.00 |
| 804.00 | 919.00 | 873.33 | -1.05 | 0.32 |
| 739.00 | 737.00 | 781.67 | 1.13 | 0.01 |
| 223.00 | 239.00 | 232.67 | -1.16 | 0.03 |
| 1562.00 | 1743.00 | 1684.33 | 1.29 | 0.00 |
| 1418.00 | 1561.00 | 1503.33 | 1.04 | 0.43 |
| 240.00 | 251.00 | 273.67 | 1.09 | 0.32 |
| 336.00 | 306.00 | 331.00 | 1.05 | 0.44 |
| 1181.00 | 1256.00 | 1185.33 | 1.17 | 0.01 |
| 1155.00 | 1306.00 | 1205.33 | -1.04 | 0.45 |

| | | | | |
|---|---|---|---|---|
| 41.00 | 55.00 | 50.33 | -1.36 | 0.01 |
| 3601.00 | 3776.00 | 3768.33 | -1.09 | 0.01 |
| 13.00 | 15.00 | 16.00 | 1.08 | 0.76 |
| 1844.00 | 2049.00 | 1948.33 | -1.02 | 0.68 |
| 3131.00 | 3463.00 | 3609.33 | 1.31 | 0.00 |
| 1007.00 | 1089.00 | 1085.67 | 1.18 | 0.00 |
| 861.00 | 864.00 | 852.00 | -1.05 | 0.31 |
| 2375.00 | 2225.00 | 2366.33 | 1.22 | 0.00 |
| 1275.00 | 1325.00 | 1347.67 | 1.01 | 0.78 |
| 128.00 | 118.00 | 125.67 | -1.05 | 0.59 |
| 865.00 | 914.00 | 979.33 | 1.21 | 0.00 |
| 253.00 | 236.00 | 232.67 | -1.11 | 0.19 |
| 708.00 | 805.00 | 770.33 | 1.18 | 0.00 |
| 244.00 | 280.00 | 267.00 | 1.46 | 0.00 |
| 989.00 | 1124.00 | 1125.33 | 1.19 | 0.00 |
| 782.00 | 878.00 | 874.00 | 1.31 | 0.00 |
| 519.00 | 523.00 | 536.67 | 1.07 | 0.17 |
| 16776.00 | 18193.00 | 17758.00 | 1.21 | 0.00 |
| 1634.00 | 1606.00 | 1717.67 | 1.04 | 0.42 |
| 156.00 | 183.00 | 168.67 | -1.19 | 0.02 |
| 2357.00 | 2465.00 | 2488.00 | -1.00 | 0.97 |
| 23.00 | 20.00 | 22.67 | 1.14 | 0.49 |
| 1835.00 | 2212.00 | 2126.33 | 2.17 | 0.00 |
| 7.00 | 31.00 | 15.33 | 1.29 | 0.49 |
| 1078.00 | 1023.00 | 1088.67 | -1.15 | 0.00 |
| 1202.00 | 1291.00 | 1266.00 | 1.06 | 0.16 |
| 736.00 | 713.00 | 752.33 | -1.05 | 0.33 |
| 39.00 | 59.00 | 56.00 | 1.05 | 0.74 |
| 1272.00 | 1301.00 | 1332.00 | 1.11 | 0.02 |
| 389.00 | 475.00 | 431.00 | -1.13 | 0.03 |
| 31.00 | 36.00 | 33.67 | -1.09 | 0.57 |
| 157.00 | 137.00 | 160.67 | -1.09 | 0.36 |
| 5495.00 | 5542.00 | 5726.67 | 1.06 | 0.11 |
| 4573.00 | 5367.00 | 4983.33 | 1.19 | 0.00 |
| 7487.00 | 8851.00 | 8389.67 | -1.06 | 0.12 |
| 14267.00 | 15689.00 | 15571.33 | 1.98 | 0.00 |
| 351.00 | 358.00 | 377.33 | -1.21 | 0.00 |
| 10398.00 | 10377.00 | 10598.00 | 1.00 | 1.00 |
| 2026.00 | 2596.00 | 2382.33 | -1.06 | 0.28 |
| 993.00 | 1198.00 | 1160.00 | -1.00 | 0.99 |
| 1096.00 | 1247.00 | 1216.33 | 1.12 | 0.01 |
| 5536.00 | 5938.00 | 6219.33 | 1.12 | 0.01 |
| 1198.00 | 1352.00 | 1243.00 | 1.06 | 0.24 |
| 190.00 | 222.00 | 217.67 | 1.01 | 0.94 |
| 58.00 | 58.00 | 59.33 | -1.22 | 0.10 |
| 725.00 | 801.00 | 798.00 | 1.05 | 0.34 |
| 1592.00 | 1491.00 | 1575.33 | 1.06 | 0.23 |
| 41.00 | 47.00 | 45.00 | 1.00 | 1.00 |

| | | | | |
|---|---|---|---|---|
| 577.00 | 551.00 | 598.00 | -1.01 | 0.92 |
| 517.00 | 519.00 | 537.33 | -1.12 | 0.02 |
| 618.00 | 687.00 | 666.33 | -1.11 | 0.02 |
| 46.00 | 48.00 | 48.00 | -1.27 | 0.05 |
| 1160.00 | 1218.00 | 1215.67 | -1.15 | 0.00 |
| 3956.00 | 3879.00 | 3897.00 | -1.13 | 0.00 |
| 1438.00 | 1673.00 | 1560.33 | 1.28 | 0.00 |
| 9893.00 | 11108.00 | 11196.33 | 1.12 | 0.00 |
| 7721.00 | 8100.00 | 8060.00 | 1.02 | 0.52 |
| 631.00 | 722.00 | 705.33 | 1.11 | 0.02 |
| 609.00 | 774.00 | 718.00 | 1.07 | 0.24 |
| 1288.00 | 1485.00 | 1493.00 | 1.27 | 0.00 |
| 475.00 | 608.00 | 536.00 | 1.31 | 0.00 |
| 1340.00 | 1610.00 | 1559.00 | 1.13 | 0.01 |
| 4267.00 | 4568.00 | 4576.33 | 1.06 | 0.07 |
| 479.00 | 484.00 | 534.67 | 1.12 | 0.08 |
| 2045.00 | 2642.00 | 2476.67 | 1.19 | 0.00 |
| 581.00 | 598.00 | 633.33 | 1.29 | 0.00 |
| 911.00 | 1001.00 | 1012.00 | 1.18 | 0.00 |
| 47.00 | 32.00 | 39.00 | 1.12 | 0.46 |
| 376.00 | 461.00 | 448.00 | 1.12 | 0.09 |
| 1374.00 | 1559.00 | 1487.67 | 1.25 | 0.00 |
| 3383.00 | 3689.00 | 3652.33 | -1.02 | 0.53 |
| 2020.00 | 2052.00 | 2097.33 | -1.28 | 0.00 |
| 2269.00 | 2199.00 | 2374.67 | 1.16 | 0.00 |
| 1383.00 | 1577.00 | 1514.00 | 1.18 | 0.00 |
| 1073.00 | 1106.00 | 1148.33 | -1.02 | 0.67 |
| 2886.00 | 3502.00 | 3244.67 | 1.11 | 0.01 |
| 1200.00 | 1300.00 | 1342.33 | 1.07 | 0.15 |
| 62.00 | 76.00 | 80.00 | -1.19 | 0.13 |
| 1420.00 | 1564.00 | 1596.00 | 1.19 | 0.00 |
| 62.00 | 76.00 | 66.00 | 1.32 | 0.02 |
| 1625.00 | 1838.00 | 1832.33 | 1.15 | 0.00 |
| 1380.00 | 1456.00 | 1433.67 | -1.21 | 0.00 |
| 1628.00 | 1729.00 | 1774.00 | 1.02 | 0.68 |
| 725.00 | 816.00 | 783.67 | 1.14 | 0.00 |
| 35.00 | 33.00 | 34.00 | -1.36 | 0.03 |
| 1840.00 | 1890.00 | 1916.33 | 1.04 | 0.32 |
| 1743.00 | 1886.00 | 1861.33 | 1.31 | 0.00 |
| 459.00 | 545.00 | 541.00 | 1.42 | 0.00 |
| 75.00 | 69.00 | 74.00 | -1.20 | 0.07 |
| 1040.00 | 1182.00 | 1157.00 | 1.35 | 0.00 |
| 1048.00 | 1195.00 | 1142.67 | 1.15 | 0.01 |
| 3064.00 | 3183.00 | 3206.67 | 1.08 | 0.02 |
| 2264.00 | 2489.00 | 2364.00 | 1.06 | 0.19 |
| 1073.00 | 1111.00 | 1083.67 | -1.45 | 0.00 |
| 2717.00 | 2746.00 | 2795.67 | -1.32 | 0.00 |
| 192.00 | 201.00 | 224.67 | 1.18 | 0.04 |

| | | | | |
|---|---|---|---|---|
| 2162.00 | 2154.00 | 2142.67 | -1.02 | 0.64 |
| 1243.00 | 1249.00 | 1258.00 | -1.13 | 0.00 |
| 1679.00 | 1681.00 | 1669.67 | -1.01 | 0.86 |
| 784.00 | 776.00 | 785.67 | 1.02 | 0.74 |
| 643.00 | 648.00 | 643.33 | -1.08 | 0.20 |
| 1108.00 | 1090.00 | 1135.00 | 1.10 | 0.03 |
| 841.00 | 898.00 | 978.00 | 1.05 | 0.46 |
| 990.00 | 1148.00 | 1131.00 | 1.15 | 0.00 |
| 107.00 | 103.00 | 103.67 | 1.07 | 0.47 |
| 991.00 | 1132.00 | 1078.33 | 1.15 | 0.00 |
| 1369.00 | 1315.00 | 1348.67 | -1.31 | 0.00 |
| 859.00 | 987.00 | 934.67 | -1.30 | 0.00 |
| 281.00 | 266.00 | 280.67 | 1.03 | 0.64 |
| 2976.00 | 3110.00 | 3112.67 | 1.12 | 0.00 |
| 2871.00 | 3142.00 | 3030.33 | 1.19 | 0.00 |
| 439.00 | 536.00 | 495.67 | 1.07 | 0.25 |
| 1247.00 | 1228.00 | 1270.67 | -1.13 | 0.00 |
| 22.00 | 13.00 | 18.00 | 1.44 | 0.12 |
| 1126.00 | 1195.00 | 1162.67 | 1.04 | 0.41 |
| 1295.00 | 1399.00 | 1334.00 | -1.14 | 0.00 |
| 961.00 | 926.00 | 888.00 | 1.02 | 0.78 |
| 5963.00 | 5345.00 | 5483.67 | -1.33 | 0.00 |
| 4147.00 | 4646.00 | 4302.67 | -1.17 | 0.00 |
| 45.00 | 41.00 | 38.67 | 1.17 | 0.32 |
| 457.00 | 410.00 | 439.00 | 1.03 | 0.69 |
| 465.00 | 491.00 | 503.33 | 1.12 | 0.03 |
| 2285.00 | 2543.00 | 2415.67 | 1.05 | 0.26 |
| 1857.00 | 1883.00 | 1927.33 | -1.01 | 0.90 |
| 295.00 | 362.00 | 330.33 | -1.13 | 0.12 |
| 186.00 | 266.00 | 221.67 | 1.19 | 0.04 |
| 715.00 | 839.00 | 776.33 | 1.04 | 0.48 |
| 3032.00 | 3478.00 | 3323.00 | -1.03 | 0.46 |
| 1530.00 | 1432.00 | 1570.00 | -1.00 | 0.93 |
| 425.00 | 406.00 | 387.67 | -1.22 | 0.01 |
| 564.00 | 516.00 | 557.67 | 1.07 | 0.20 |
| 12169.00 | 12970.00 | 13391.67 | 1.06 | 0.11 |
| 776.00 | 935.00 | 928.33 | 1.07 | 0.22 |
| 1285.00 | 1548.00 | 1438.67 | 1.26 | 0.00 |
| 493.00 | 459.00 | 481.33 | 1.01 | 0.94 |
| 2342.00 | 2370.00 | 2445.33 | 1.21 | 0.00 |
| 1172.00 | 1133.00 | 1197.00 | -1.09 | 0.05 |
| 44.00 | 46.00 | 46.33 | -1.11 | 0.47 |
| 24.00 | 47.00 | 33.33 | -1.42 | 0.02 |
| 4.00 | 3.00 | 4.33 | -3.82 | 0.00 |
| 17.00 | 7.00 | 11.00 | 2.15 | 0.02 |
| 18.00 | 14.00 | 16.67 | -1.48 | 0.06 |
| 279.00 | 295.00 | 277.33 | 1.03 | 0.73 |
| 1300.00 | 1414.00 | 1454.67 | 1.14 | 0.01 |

| | | | | |
|---|---|---|---|---|
| 609.00 | 658.00 | 661.67 | 1.21 | 0.00 |
| 1506.00 | 1641.00 | 1535.67 | 1.07 | 0.19 |
| 1742.00 | 1907.00 | 1932.00 | 1.11 | 0.02 |
| 62.00 | 40.00 | 50.33 | 1.24 | 0.14 |
| 366.00 | 393.00 | 402.33 | 1.13 | 0.03 |
| 5549.00 | 5780.00 | 5682.00 | 1.01 | 0.79 |
| 1651.00 | 1715.00 | 1675.33 | -1.24 | 0.00 |
| 116.00 | 131.00 | 114.33 | -1.02 | 0.94 |
| 3871.00 | 3844.00 | 3761.00 | -1.12 | 0.02 |
| 979.00 | 1121.00 | 1078.33 | -1.09 | 0.05 |
| 1158.00 | 1322.00 | 1240.33 | -1.07 | 0.13 |
| 258.00 | 266.00 | 253.67 | -1.00 | 1.00 |
| 2643.00 | 2984.00 | 2732.00 | -1.18 | 0.00 |
| 1429.00 | 1479.00 | 1479.33 | 1.25 | 0.00 |
| 1940.00 | 1878.00 | 1862.00 | -1.03 | 0.57 |
| 387.00 | 402.00 | 392.00 | 1.09 | 0.16 |
| 1662.00 | 1834.00 | 1770.33 | -1.05 | 0.19 |
| 972.00 | 1124.00 | 1077.67 | 1.02 | 0.58 |
| 20.00 | 31.00 | 22.67 | -1.31 | 0.17 |
| 209.00 | 221.00 | 213.00 | 1.02 | 0.79 |
| 86.00 | 58.00 | 69.33 | -1.22 | 0.10 |
| 914.00 | 812.00 | 903.00 | -1.21 | 0.00 |
| 398.00 | 402.00 | 390.67 | -1.05 | 0.40 |
| 1266.00 | 1193.00 | 1333.33 | 1.04 | 0.40 |
| 98.00 | 109.00 | 112.00 | 1.04 | 0.66 |
| 255.00 | 271.00 | 260.67 | 1.07 | 0.42 |
| 899.00 | 934.00 | 941.67 | 1.02 | 0.73 |
| 461.00 | 457.00 | 498.67 | 1.08 | 0.21 |
| 1472.00 | 1592.00 | 1535.33 | 1.00 | 0.94 |
| 0.00 | 1.00 | 1.33 | -2.64 | 0.14 |
| 812.00 | 811.00 | 814.67 | -1.25 | 0.00 |
| 51.00 | 49.00 | 55.00 | -1.24 | 0.06 |
| 1770.00 | 1879.00 | 1858.00 | -1.04 | 0.26 |
| 1183.00 | 1297.00 | 1271.33 | 1.02 | 0.71 |
| 2.00 | 4.00 | 3.00 | -1.66 | 0.31 |
| 23.00 | 20.00 | 21.00 | -2.08 | 0.00 |
| 512.00 | 502.00 | 522.00 | 1.02 | 0.77 |
| 460.00 | 435.00 | 473.33 | 1.53 | 0.00 |
| 947.00 | 1108.00 | 1079.67 | 1.08 | 0.08 |
| 807.00 | 952.00 | 885.00 | 1.22 | 0.00 |
| 3.00 | 10.00 | 6.00 | 2.16 | 0.08 |
| 3223.00 | 3315.00 | 3265.67 | -1.01 | 0.82 |
| 1512.00 | 1719.00 | 1643.33 | 1.05 | 0.21 |
| 753.00 | 922.00 | 868.33 | -1.05 | 0.34 |
| 460.00 | 533.00 | 510.33 | 1.16 | 0.01 |
| 1.00 | 1.00 | 1.00 | -1.91 | 0.51 |
| 31.00 | 9.00 | 26.67 | 1.13 | 0.70 |
| 2739.00 | 2969.00 | 2983.33 | -1.01 | 0.73 |

| | | | | |
|---|---|---|---|---|
| 188.00 | 198.00 | 192.33 | 1.06 | 0.41 |
| 1730.00 | 1760.00 | 1854.33 | -1.11 | 0.01 |
| 588.00 | 709.00 | 692.67 | -1.07 | 0.22 |
| 330.00 | 344.00 | 356.00 | 1.04 | 0.54 |
| 726.00 | 900.00 | 882.67 | 1.26 | 0.00 |
| 1167.00 | 1414.00 | 1418.33 | 1.15 | 0.01 |
| 2696.00 | 3335.00 | 3289.33 | 1.14 | 0.02 |
| 2088.00 | 2400.00 | 2369.00 | -1.05 | 0.22 |
| 2494.00 | 2696.00 | 2715.00 | 1.04 | 0.29 |
| 1048.00 | 1127.00 | 1147.33 | 1.26 | 0.00 |
| 2934.00 | 3285.00 | 3444.33 | 1.23 | 0.00 |
| 129.00 | 132.00 | 133.33 | 1.13 | 0.16 |
| 1284.00 | 1281.00 | 1384.33 | 1.14 | 0.01 |
| 3273.00 | 3263.00 | 3243.00 | -1.16 | 0.00 |
| 47.00 | 34.00 | 44.67 | -1.17 | 0.29 |
| 801.00 | 823.00 | 811.00 | -1.05 | 0.26 |
| 1602.00 | 1724.00 | 1747.00 | 1.01 | 0.74 |
| 1611.00 | 1802.00 | 1741.33 | 1.18 | 0.00 |
| 1446.00 | 1636.00 | 1649.00 | 1.09 | 0.06 |
| 2713.00 | 2924.00 | 2954.67 | 1.08 | 0.06 |
| 603.00 | 618.00 | 675.67 | -1.06 | 0.32 |
| 891.00 | 894.00 | 938.00 | -1.05 | 0.25 |
| 379.00 | 436.00 | 437.33 | 1.07 | 0.22 |
| 823.00 | 891.00 | 908.33 | -1.01 | 0.87 |
| 1030.00 | 1007.00 | 1090.00 | 1.01 | 0.80 |
| 5.00 | 0.00 | 4.00 | -1.34 | 0.58 |
| 1320.00 | 1594.00 | 1578.67 | 1.21 | 0.00 |
| 1961.00 | 2310.00 | 2160.67 | -1.00 | 0.98 |
| 1339.00 | 1636.00 | 1520.00 | 1.26 | 0.00 |
| 2952.00 | 3306.00 | 3254.33 | -1.01 | 0.72 |
| 1498.00 | 1478.00 | 1527.67 | -1.01 | 0.76 |
| 799.00 | 978.00 | 867.00 | -1.09 | 0.12 |
| 4.00 | 2.00 | 3.00 | -1.88 | 0.18 |
| 1375.00 | 1443.00 | 1422.67 | 1.14 | 0.00 |
| 4791.00 | 5173.00 | 5069.00 | 1.15 | 0.00 |
| 1349.00 | 1488.00 | 1425.00 | 1.05 | 0.27 |
| 1040.00 | 1160.00 | 1138.67 | 1.07 | 0.13 |
| 20.00 | 17.00 | 18.00 | 1.05 | 0.85 |
| 2140.00 | 2160.00 | 2177.33 | -1.12 | 0.00 |
| 1395.00 | 1387.00 | 1440.67 | -1.05 | 0.22 |
| 926.00 | 1001.00 | 1024.33 | -1.04 | 0.39 |
| 5012.00 | 4628.00 | 4960.67 | -1.04 | 0.30 |
| 2647.00 | 2708.00 | 2684.67 | -1.09 | 0.02 |
| 974.00 | 1005.00 | 1029.33 | 1.13 | 0.01 |
| 89.00 | 116.00 | 111.33 | 1.05 | 0.63 |
| 17.00 | 32.00 | 22.00 | 1.03 | 0.87 |
| 788.00 | 954.00 | 873.33 | -1.10 | 0.06 |
| 18.00 | 23.00 | 19.00 | -1.15 | 0.54 |

| | | | | |
|---|---|---|---|---|
| 1204.00 | 1371.00 | 1355.33 | -1.01 | 0.73 |
| 291.00 | 313.00 | 313.33 | -1.09 | 0.14 |
| 715.00 | 773.00 | 764.00 | -1.00 | 0.96 |
| 1770.00 | 1816.00 | 1779.33 | -1.02 | 0.59 |
| 1527.00 | 1866.00 | 1774.33 | -1.26 | 0.00 |
| 11.00 | 21.00 | 17.33 | -1.23 | 0.34 |
| 1854.00 | 1972.00 | 1928.33 | 1.10 | 0.01 |
| 2948.00 | 3223.00 | 3123.33 | -1.04 | 0.30 |
| 233.00 | 271.00 | 248.67 | -1.19 | 0.01 |
| 2268.00 | 2191.00 | 2191.33 | -1.27 | 0.00 |
| 290.00 | 298.00 | 292.67 | -1.04 | 0.50 |
| 326.00 | 389.00 | 348.33 | -1.09 | 0.21 |
| 1147.00 | 1265.00 | 1251.67 | 1.10 | 0.02 |
| 32.00 | 23.00 | 26.33 | -1.14 | 0.47 |
| 1529.00 | 1649.00 | 1616.33 | -1.01 | 0.79 |
| 2098.00 | 2186.00 | 2189.00 | -1.06 | 0.11 |
| 591.00 | 572.00 | 594.00 | -1.04 | 0.47 |
| 607.00 | 653.00 | 633.33 | -1.29 | 0.00 |
| 494.00 | 506.00 | 515.67 | -1.05 | 0.31 |
| 20.00 | 39.00 | 28.33 | 1.21 | 0.35 |
| 2137.00 | 2429.00 | 2314.67 | 1.05 | 0.21 |
| 1818.00 | 2113.00 | 1989.00 | -1.01 | 0.77 |
| 446.00 | 493.00 | 463.67 | -1.13 | 0.03 |
| 2531.00 | 2504.00 | 2564.67 | 1.01 | 0.85 |
| 43.00 | 36.00 | 42.00 | 1.16 | 0.32 |
| 1331.00 | 1513.00 | 1386.00 | -1.01 | 0.87 |
| 902.00 | 1000.00 | 941.67 | -1.11 | 0.02 |
| 31.00 | 43.00 | 45.00 | 1.08 | 0.68 |
| 571.00 | 667.00 | 634.67 | 1.08 | 0.13 |
| 3035.00 | 3376.00 | 3260.33 | -1.01 | 0.86 |
| 1271.00 | 1355.00 | 1323.33 | 1.03 | 0.49 |
| 3359.00 | 3400.00 | 3397.00 | -1.15 | 0.00 |
| 695.00 | 783.00 | 750.67 | -1.11 | 0.01 |
| 920.00 | 908.00 | 918.33 | -1.03 | 0.55 |
| 961.00 | 1145.00 | 1095.67 | 1.23 | 0.00 |
| 1381.00 | 1632.00 | 1572.67 | 1.07 | 0.18 |
| 794.00 | 882.00 | 806.67 | -1.05 | 0.41 |
| 705.00 | 758.00 | 741.67 | 1.10 | 0.03 |
| 1885.00 | 2175.00 | 2112.33 | 1.08 | 0.06 |
| 2120.00 | 2129.00 | 2040.00 | -1.01 | 0.91 |
| 620.00 | 740.00 | 707.67 | 1.16 | 0.00 |
| 917.00 | 1054.00 | 1032.33 | 1.11 | 0.04 |
| 1530.00 | 1825.00 | 1746.67 | 1.11 | 0.01 |
| 1090.00 | 1223.00 | 1113.00 | -1.68 | 0.00 |
| 791.00 | 797.00 | 813.00 | 1.05 | 0.28 |
| 392.00 | 348.00 | 395.00 | -1.27 | 0.00 |
| 956.00 | 1062.00 | 1014.00 | 1.16 | 0.00 |
| 529.00 | 546.00 | 572.67 | 1.04 | 0.50 |

| | | | | |
|---|---|---|---|---|
| 3.00 | 6.00 | 5.33 | -1.94 | 0.04 |
| 1262.00 | 1174.00 | 1217.67 | -1.05 | 0.26 |
| 378.00 | 399.00 | 403.33 | -1.26 | 0.00 |
| 1568.00 | 1699.00 | 1743.00 | 1.06 | 0.21 |
| 1394.00 | 1461.00 | 1497.33 | 1.08 | 0.05 |
| 701.00 | 789.00 | 753.33 | -1.02 | 0.71 |
| 15.00 | 28.00 | 23.00 | 1.13 | 0.56 |
| 6.00 | 4.00 | 3.67 | -1.10 | 1.00 |
| 962.00 | 998.00 | 969.00 | -1.03 | 0.56 |
| 4035.00 | 4631.00 | 4509.00 | 1.16 | 0.00 |
| 1615.00 | 1633.00 | 1685.33 | 1.07 | 0.09 |
| 6.00 | 4.00 | 5.00 | -1.28 | 0.62 |
| 135.00 | 128.00 | 133.67 | -1.28 | 0.00 |
| 182.00 | 164.00 | 172.67 | -1.01 | 0.90 |
| 5.00 | 6.00 | 5.67 | 1.12 | 0.86 |
| 1477.00 | 1620.00 | 1617.67 | -1.12 | 0.01 |
| 4157.00 | 4029.00 | 4216.67 | -1.32 | 0.00 |
| 946.00 | 940.00 | 956.00 | -1.12 | 0.01 |
| 4.00 | 5.00 | 4.00 | -1.26 | 0.71 |
| 74.00 | 50.00 | 66.00 | 1.25 | 0.11 |
| 886.00 | 1026.00 | 1023.33 | 1.26 | 0.00 |
| 2274.00 | 2668.00 | 2525.00 | 1.40 | 0.00 |
| 382.00 | 459.00 | 419.00 | -1.13 | 0.04 |
| 9944.00 | 9617.00 | 9845.00 | -1.11 | 0.00 |
| 7648.00 | 8417.00 | 8155.00 | 1.07 | 0.06 |
| 623.00 | 685.00 | 659.67 | -1.04 | 0.41 |
| 262.00 | 303.00 | 293.33 | 1.01 | 0.91 |
| 5620.00 | 6154.00 | 6220.00 | 1.33 | 0.00 |
| 2574.00 | 2791.00 | 2851.33 | 1.16 | 0.00 |
| 1481.00 | 1847.00 | 1723.00 | 1.22 | 0.00 |
| 119.00 | 87.00 | 101.67 | -1.79 | 0.00 |
| 5801.00 | 6233.00 | 6138.67 | -1.39 | 0.00 |
| 1218.00 | 1312.00 | 1318.33 | -1.08 | 0.05 |
| 1248.00 | 1296.00 | 1333.67 | -1.29 | 0.00 |
| 1474.00 | 1722.00 | 1670.33 | 1.17 | 0.00 |
| 1636.00 | 1775.00 | 1735.00 | 1.00 | 0.93 |
| 1341.00 | 1432.00 | 1415.67 | 1.06 | 0.16 |
| 157.00 | 158.00 | 168.33 | -1.44 | 0.00 |
| 42.00 | 65.00 | 56.33 | 1.23 | 0.12 |
| 487.00 | 522.00 | 470.33 | -1.35 | 0.00 |
| 1142.00 | 1108.00 | 1118.33 | 1.01 | 0.81 |
| 1200.00 | 1390.00 | 1338.33 | 1.07 | 0.13 |
| 52.00 | 57.00 | 56.00 | 1.02 | 0.88 |
| 117764.00 | 127730.00 | 128065.00 | 1.22 | 0.00 |
| 92.00 | 100.00 | 109.67 | 1.19 | 0.11 |
| 1173.00 | 1124.00 | 1184.00 | 1.01 | 0.77 |
| 9.00 | 16.00 | 11.67 | -1.39 | 0.20 |
| 571.00 | 612.00 | 609.67 | -1.02 | 0.62 |

| | | | | |
|---|---|---|---|---|
| 57.00 | 44.00 | 54.33 | -3.04 | 0.00 |
| 2080.00 | 2068.00 | 2177.67 | -1.03 | 0.59 |
| 1415.00 | 1382.00 | 1396.00 | 1.07 | 0.12 |
| 1257.00 | 1273.00 | 1285.67 | -1.16 | 0.00 |
| 118.00 | 153.00 | 142.00 | 1.26 | 0.01 |
| 1198.00 | 1397.00 | 1373.67 | 1.31 | 0.00 |
| 76.00 | 55.00 | 66.00 | -1.53 | 0.00 |
| 7.00 | 16.00 | 10.67 | 1.26 | 0.46 |
| 3609.00 | 4014.00 | 3973.67 | -1.01 | 0.69 |
| 7.00 | 4.00 | 4.33 | -1.47 | 0.39 |
| 347.00 | 442.00 | 425.00 | 1.17 | 0.02 |
| 948.00 | 892.00 | 915.67 | 1.11 | 0.04 |
| 1339.00 | 1378.00 | 1348.33 | -1.03 | 0.53 |
| 873.00 | 954.00 | 940.67 | -1.03 | 0.53 |
| 1176.00 | 1197.00 | 1206.33 | -1.24 | 0.00 |
| 1169.00 | 1267.00 | 1234.00 | -1.27 | 0.00 |
| 1605.00 | 1793.00 | 1686.33 | -1.09 | 0.05 |
| 989.00 | 893.00 | 937.00 | -1.30 | 0.00 |
| 1099.00 | 1147.00 | 1178.33 | -1.00 | 0.95 |
| 677.00 | 682.00 | 682.67 | -1.15 | 0.00 |
| 1258.00 | 1334.00 | 1395.33 | 1.02 | 0.65 |
| 1740.00 | 1934.00 | 1927.00 | 1.09 | 0.04 |
| 756.00 | 823.00 | 807.67 | -1.02 | 0.66 |
| 1980.00 | 1977.00 | 2162.33 | 1.08 | 0.12 |
| 929.00 | 1153.00 | 1121.33 | 1.14 | 0.02 |
| 779.00 | 837.00 | 817.00 | 1.04 | 0.41 |
| 362.00 | 345.00 | 352.67 | -1.07 | 0.29 |
| 2772.00 | 2789.00 | 2859.33 | 1.10 | 0.01 |
| 1158.00 | 1323.00 | 1282.00 | 1.07 | 0.13 |
| 609.00 | 687.00 | 657.00 | 1.06 | 0.22 |
| 1282.00 | 1280.00 | 1300.67 | 1.02 | 0.59 |
| 594.00 | 561.00 | 597.33 | 1.16 | 0.00 |
| 35.00 | 45.00 | 43.00 | 1.44 | 0.01 |
| 1743.00 | 1811.00 | 1757.33 | -1.12 | 0.01 |
| 1320.00 | 1370.00 | 1357.33 | 1.00 | 0.99 |
| 706.00 | 907.00 | 848.33 | 1.06 | 0.32 |
| 1900.00 | 2047.00 | 2011.67 | 1.04 | 0.24 |
| 885.00 | 906.00 | 882.00 | 1.03 | 0.59 |
| 574.00 | 666.00 | 625.67 | 1.01 | 0.86 |
| 2229.00 | 2214.00 | 2299.00 | -1.00 | 0.92 |
| 48.00 | 48.00 | 49.67 | 1.11 | 0.44 |
| 713.00 | 721.00 | 718.67 | 1.05 | 0.33 |
| 1151.00 | 1252.00 | 1214.33 | 1.05 | 0.28 |
| 1024.00 | 1119.00 | 1158.00 | 1.08 | 0.10 |
| 1037.00 | 1091.00 | 1086.00 | -1.16 | 0.00 |
| 412.00 | 436.00 | 456.67 | -1.10 | 0.09 |
| 568.00 | 660.00 | 622.67 | -1.17 | 0.00 |
| 708.00 | 889.00 | 827.00 | 1.30 | 0.00 |

| | | | | |
|---|---|---|---|---|
| 508.00 | 524.00 | 540.00 | 1.03 | 0.55 |
| 530.00 | 560.00 | 563.00 | -1.11 | 0.04 |
| 549.00 | 566.00 | 576.67 | -1.03 | 0.52 |
| 10.00 | 11.00 | 13.67 | -1.38 | 0.12 |
| 0.00 | 4.00 | 3.00 | -1.99 | 0.13 |
| 4.00 | 7.00 | 5.33 | -1.14 | 0.87 |
| 881.00 | 1038.00 | 1015.00 | 1.34 | 0.00 |
| 579.00 | 584.00 | 587.33 | -1.15 | 0.01 |
| 905.00 | 978.00 | 969.67 | 1.02 | 0.67 |
| 27.00 | 25.00 | 28.67 | -1.39 | 0.03 |
| 22.00 | 16.00 | 18.67 | 1.67 | 0.03 |
| 694.00 | 726.00 | 738.00 | -1.15 | 0.00 |
| 1299.00 | 1394.00 | 1395.67 | 1.08 | 0.05 |
| 9.00 | 11.00 | 10.67 | 1.65 | 0.10 |
| 5.00 | 7.00 | 5.67 | -1.77 | 0.10 |
| 2.00 | 6.00 | 3.67 | 1.76 | 0.34 |
| 4385.00 | 4729.00 | 4631.67 | 1.04 | 0.22 |
| 612.00 | 647.00 | 644.00 | 1.07 | 0.15 |
| 1962.00 | 2045.00 | 1986.33 | 1.04 | 0.40 |
| 5271.00 | 5482.00 | 5501.33 | 1.02 | 0.50 |
| 256.00 | 301.00 | 303.33 | 1.03 | 0.67 |
| 5.00 | 4.00 | 3.00 | -1.34 | 0.67 |
| 730.00 | 820.00 | 758.33 | -1.02 | 0.73 |
| 1968.00 | 1793.00 | 1877.33 | -1.33 | 0.00 |
| 557.00 | 603.00 | 593.00 | -1.01 | 0.92 |
| 209.00 | 204.00 | 211.67 | -1.11 | 0.13 |
| 152.00 | 189.00 | 176.67 | -1.07 | 0.47 |
| 211.00 | 175.00 | 194.67 | -1.09 | 0.39 |
| 11.00 | 9.00 | 10.67 | -1.52 | 0.11 |
| 115.00 | 98.00 | 112.67 | -1.10 | 0.28 |
| 880.00 | 1008.00 | 940.33 | -1.01 | 0.81 |
| 102.00 | 130.00 | 128.00 | -1.33 | 0.00 |
| 2308.00 | 2369.00 | 2375.00 | 1.10 | 0.01 |
| 829.00 | 774.00 | 803.00 | -1.03 | 0.59 |
| 1697.00 | 1689.00 | 1776.00 | -1.03 | 0.53 |
| 377.00 | 451.00 | 413.33 | 1.17 | 0.01 |
| 1464.00 | 1758.00 | 1659.33 | 1.10 | 0.03 |
| 885.00 | 960.00 | 966.67 | 1.06 | 0.18 |
| 2045.00 | 2160.00 | 2219.00 | 1.08 | 0.05 |
| 633.00 | 628.00 | 634.00 | 1.02 | 0.74 |
| 1430.00 | 1508.00 | 1475.00 | -1.18 | 0.00 |
| 1456.00 | 1503.00 | 1514.67 | 1.15 | 0.00 |
| 439.00 | 537.00 | 488.33 | -1.29 | 0.00 |
| 30.00 | 28.00 | 25.33 | -1.38 | 0.05 |
| 683.00 | 800.00 | 746.00 | 1.19 | 0.00 |
| 98.00 | 90.00 | 93.00 | -1.05 | 0.64 |
| 4595.00 | 4707.00 | 4729.33 | -1.13 | 0.00 |
| 1792.00 | 2029.00 | 1946.33 | -1.04 | 0.30 |

| | | | | |
|---|---|---|---|---|
| 1854.00 | 2045.00 | 2039.33 | 1.15 | 0.00 |
| 789.00 | 932.00 | 922.67 | 1.01 | 0.92 |
| 499.00 | 608.00 | 555.33 | 1.18 | 0.00 |
| 1086.00 | 1180.00 | 1192.00 | 1.16 | 0.00 |
| 1697.00 | 1919.00 | 1814.67 | -1.23 | 0.00 |
| 439.00 | 477.00 | 461.00 | 1.01 | 0.92 |
| 445.00 | 485.00 | 483.33 | -1.06 | 0.34 |
| 132.00 | 160.00 | 151.00 | -1.17 | 0.05 |
| 3011.00 | 3416.00 | 3285.00 | -1.05 | 0.19 |
| 2127.00 | 2362.00 | 2263.67 | -1.28 | 0.00 |
| 3281.00 | 3386.00 | 3366.00 | 1.05 | 0.19 |
| 71.00 | 89.00 | 71.67 | 1.16 | 0.27 |
| 2266.00 | 2383.00 | 2342.67 | -1.06 | 0.11 |
| 878.00 | 940.00 | 979.67 | 1.33 | 0.00 |
| 2024.00 | 2405.00 | 2232.00 | -1.02 | 0.66 |
| 2293.00 | 2632.00 | 2412.67 | -1.03 | 0.51 |
| 857.00 | 915.00 | 909.00 | -1.10 | 0.02 |
| 484.00 | 394.00 | 433.33 | -1.17 | 0.02 |
| 523.00 | 526.00 | 532.33 | 1.10 | 0.09 |
| 2901.00 | 3217.00 | 3063.33 | 1.08 | 0.06 |
| 415.00 | 498.00 | 461.67 | -1.28 | 0.00 |
| 932.00 | 968.00 | 1004.00 | -1.11 | 0.02 |
| 2409.00 | 2588.00 | 2678.67 | 1.09 | 0.05 |
| 133.00 | 153.00 | 144.00 | 1.10 | 0.32 |
| 346.00 | 388.00 | 369.00 | -1.00 | 1.00 |
| 896.00 | 928.00 | 929.00 | 1.04 | 0.42 |
| 2059.00 | 2197.00 | 2191.33 | 1.11 | 0.00 |
| 315.00 | 342.00 | 345.00 | 1.03 | 0.61 |
| 834.00 | 819.00 | 903.33 | 1.17 | 0.01 |
| 729.00 | 691.00 | 698.00 | -1.09 | 0.13 |
| 1806.00 | 1941.00 | 1979.00 | -1.01 | 0.74 |
| 508.00 | 534.00 | 544.33 | -1.22 | 0.00 |
| 5743.00 | 5676.00 | 5835.33 | -1.14 | 0.00 |
| 511.00 | 519.00 | 525.33 | -1.16 | 0.00 |
| 745.00 | 792.00 | 796.67 | 1.12 | 0.01 |
| 649.00 | 690.00 | 707.00 | 1.07 | 0.18 |
| 15.00 | 6.00 | 10.33 | -1.01 | 1.00 |
| 0.00 | 0.00 | 0.00 | -17.14 | 0.03 |
| 1026.00 | 963.00 | 1024.00 | 2.69 | 0.00 |
| 300.00 | 324.00 | 314.33 | -1.15 | 0.02 |
| 836.00 | 976.00 | 933.67 | 1.09 | 0.06 |
| 29.00 | 25.00 | 28.33 | 1.05 | 0.83 |
| 1169.00 | 1303.00 | 1263.67 | -1.04 | 0.37 |
| 1545.00 | 1645.00 | 1627.33 | 1.01 | 0.84 |
| 2710.00 | 3029.00 | 2971.67 | 1.03 | 0.38 |
| 211.00 | 287.00 | 240.67 | 1.76 | 0.00 |
| 1742.00 | 1978.00 | 1985.33 | 1.07 | 0.15 |
| 72.00 | 100.00 | 90.00 | 1.15 | 0.20 |

| | | | | |
|---|---|---|---|---|
| 255.00 | 259.00 | 277.33 | -1.02 | 0.77 |
| 1649.00 | 1641.00 | 1679.67 | -1.03 | 0.42 |
| 223.00 | 236.00 | 242.67 | 1.20 | 0.01 |
| 1370.00 | 1545.00 | 1587.00 | 1.06 | 0.23 |
| 463.00 | 516.00 | 494.67 | 1.02 | 0.66 |
| 1273.00 | 1212.00 | 1323.67 | -1.05 | 0.27 |
| 507.00 | 527.00 | 529.67 | -1.24 | 0.00 |
| 436.00 | 439.00 | 447.00 | 1.07 | 0.20 |
| 326.00 | 363.00 | 361.67 | 1.08 | 0.17 |
| 1000.00 | 968.00 | 989.33 | 1.01 | 0.91 |
| 366.00 | 481.00 | 432.67 | 1.36 | 0.00 |
| 507.00 | 596.00 | 564.33 | 1.23 | 0.00 |
| 7.00 | 0.00 | 4.33 | -1.17 | 0.86 |
| 18.00 | 16.00 | 19.33 | -1.17 | 0.40 |
| 814.00 | 915.00 | 871.67 | -1.19 | 0.00 |
| 193.00 | 243.00 | 217.33 | -1.08 | 0.28 |
| 73.00 | 123.00 | 94.33 | -1.14 | 0.33 |
| 13.00 | 5.00 | 7.67 | 2.47 | 0.03 |
| 106.00 | 142.00 | 124.00 | 1.21 | 0.05 |
| 9499.00 | 10011.00 | 9855.67 | 1.14 | 0.00 |
| 1129.00 | 1103.00 | 1116.00 | -1.11 | 0.02 |
| 240.00 | 289.00 | 274.00 | 1.01 | 0.89 |
| 652.00 | 717.00 | 684.00 | -1.06 | 0.24 |
| 901.00 | 806.00 | 830.33 | -1.09 | 0.14 |
| 456.00 | 477.00 | 504.33 | -1.12 | 0.04 |
| 1795.00 | 1906.00 | 1920.67 | -1.02 | 0.59 |
| 54.00 | 49.00 | 47.33 | 1.97 | 0.00 |
| 1090.00 | 1134.00 | 1147.67 | -1.01 | 0.73 |
| 3995.00 | 4164.00 | 4227.33 | -1.11 | 0.00 |
| 6089.00 | 5828.00 | 5843.67 | -1.19 | 0.00 |
| 5.00 | 4.00 | 5.67 | -1.31 | 0.53 |
| 1260.00 | 1237.00 | 1259.33 | -2.09 | 0.00 |
| 2.00 | 0.00 | 1.00 | -3.71 | 0.04 |
| 16.00 | 21.00 | 19.00 | 2.53 | 0.00 |
| 67.00 | 76.00 | 73.00 | -1.23 | 0.04 |
| 9.00 | 8.00 | 9.33 | -1.01 | 1.00 |
| 899.00 | 931.00 | 957.67 | 2.02 | 0.00 |
| 4.00 | 1.00 | 1.67 | -2.89 | 0.05 |
| 463.00 | 513.00 | 499.67 | 1.06 | 0.29 |
| 622.00 | 605.00 | 643.33 | -1.03 | 0.58 |
| 560.00 | 544.00 | 563.33 | 1.30 | 0.00 |
| 792.00 | 942.00 | 889.33 | 1.01 | 0.76 |
| 2311.00 | 2588.00 | 2481.33 | 1.12 | 0.00 |
| 1702.00 | 1985.00 | 1916.00 | 1.05 | 0.27 |
| 874.00 | 896.00 | 927.33 | 1.17 | 0.00 |
| 39.00 | 45.00 | 42.33 | 1.45 | 0.01 |
| 9.00 | 18.00 | 15.00 | -1.26 | 0.29 |
| 1156.00 | 1362.00 | 1327.00 | 1.06 | 0.24 |

| | | | | |
|---|---|---|---|---|
| 47.00 | 57.00 | 88.67 | -1.08 | 0.75 |
| 18.00 | 26.00 | 20.67 | 1.09 | 0.66 |
| 7.00 | 8.00 | 10.33 | -1.18 | 0.65 |
| 3135.00 | 3332.00 | 3272.33 | 1.25 | 0.00 |
| 975.00 | 1045.00 | 1010.33 | -1.03 | 0.50 |
| 12.00 | 12.00 | 10.00 | 1.47 | 0.22 |
| 780.00 | 779.00 | 792.67 | -1.35 | 0.00 |
| 799.00 | 942.00 | 870.00 | 1.13 | 0.03 |
| 745.00 | 813.00 | 798.33 | -1.10 | 0.04 |
| 490.00 | 521.00 | 497.67 | -1.21 | 0.00 |
| 12.00 | 12.00 | 12.67 | 1.33 | 0.28 |
| 1505.00 | 1662.00 | 1591.00 | -1.06 | 0.19 |
| 32.00 | 49.00 | 50.67 | 1.01 | 0.96 |
| 6732.00 | 6772.00 | 6895.33 | 1.07 | 0.08 |
| 592.00 | 622.00 | 610.67 | 3.70 | 0.00 |
| 1254.00 | 1229.00 | 1181.00 | 1.02 | 0.81 |
| 253.00 | 271.00 | 275.33 | -1.13 | 0.06 |
| 16055.00 | 18720.00 | 17329.00 | -1.35 | 0.00 |
| 19.00 | 7.00 | 16.00 | 1.43 | 0.16 |
| 5.00 | 1.00 | 2.33 | -1.15 | 1.00 |
| 165.00 | 185.00 | 183.33 | 1.14 | 0.07 |
| 1168.00 | 1295.00 | 1245.67 | -1.26 | 0.00 |
| 1307.00 | 1367.00 | 1375.00 | 1.04 | 0.28 |
| 596.00 | 685.00 | 682.33 | -1.01 | 0.84 |
| 9683.00 | 11335.00 | 10618.33 | -1.02 | 0.65 |
| 1249.00 | 1364.00 | 1287.33 | 1.07 | 0.17 |
| 2829.00 | 2839.00 | 2879.67 | -1.03 | 0.47 |
| 3414.00 | 3326.00 | 3433.33 | -1.01 | 0.89 |
| 1501.00 | 1457.00 | 1448.33 | -1.01 | 0.78 |
| 10012.00 | 10864.00 | 10707.00 | 1.03 | 0.42 |
| 1228.00 | 1363.00 | 1274.00 | -1.01 | 0.81 |
| 736.00 | 831.00 | 809.33 | -1.06 | 0.21 |
| 443.00 | 529.00 | 488.00 | 1.06 | 0.29 |
| 792.00 | 864.00 | 868.00 | 1.04 | 0.44 |
| 319.00 | 397.00 | 349.33 | 1.40 | 0.00 |
| 7985.00 | 8844.00 | 8556.67 | 1.13 | 0.00 |
| 6932.00 | 7775.00 | 7537.33 | 1.09 | 0.01 |
| 1019.00 | 1133.00 | 1040.00 | -1.14 | 0.02 |
| 4445.00 | 4790.00 | 4626.67 | 1.06 | 0.12 |
| 559.00 | 558.00 | 549.67 | -1.21 | 0.00 |
| 626.00 | 644.00 | 664.67 | -1.13 | 0.01 |
| 219.00 | 211.00 | 216.33 | -1.08 | 0.26 |
| 1674.00 | 1865.00 | 1787.33 | -1.18 | 0.00 |
| 319.00 | 386.00 | 377.67 | 1.11 | 0.16 |
| 253.00 | 373.00 | 319.67 | 1.21 | 0.02 |
| 13698.00 | 15023.00 | 14550.33 | 1.01 | 0.84 |
| 6587.00 | 7013.00 | 7089.33 | 1.04 | 0.28 |
| 778.00 | 848.00 | 848.67 | -1.00 | 0.97 |

| | | | | |
|---|---|---|---|---|
| 246.00 | 258.00 | 251.00 | 1.04 | 0.55 |
| 12947.00 | 14929.00 | 14557.33 | 1.35 | 0.00 |
| 4760.00 | 5352.00 | 5385.67 | 1.18 | 0.00 |
| 27.00 | 37.00 | 32.67 | 1.30 | 0.11 |
| 672.00 | 682.00 | 703.67 | 1.11 | 0.02 |
| 2218.00 | 2092.00 | 2185.00 | -1.03 | 0.47 |
| 511.00 | 582.00 | 573.00 | 1.25 | 0.00 |
| 3668.00 | 4141.00 | 3905.00 | 1.46 | 0.00 |
| 2385.00 | 2389.00 | 2501.33 | 1.07 | 0.14 |
| 1009.00 | 974.00 | 1034.33 | -1.27 | 0.00 |
| 475.00 | 550.00 | 535.67 | 1.04 | 0.47 |
| 783.00 | 760.00 | 812.67 | -1.06 | 0.22 |
| 3389.00 | 3523.00 | 3626.67 | -1.10 | 0.03 |
| 891.00 | 921.00 | 896.00 | 1.03 | 0.52 |
| 1474.00 | 1451.00 | 1502.67 | -1.02 | 0.68 |
| 1903.00 | 1914.00 | 1932.00 | -1.16 | 0.00 |
| 297.00 | 272.00 | 292.33 | -1.40 | 0.00 |
| 2542.00 | 2705.00 | 2731.00 | 1.01 | 0.77 |
| 2120.00 | 2213.00 | 2121.33 | -1.09 | 0.05 |
| 4780.00 | 4988.00 | 4935.00 | 1.13 | 0.00 |
| 419.00 | 536.00 | 471.33 | -1.32 | 0.00 |
| 8.00 | 9.00 | 10.33 | 1.13 | 0.70 |
| 85.00 | 85.00 | 78.00 | 1.46 | 0.00 |
| 193.00 | 165.00 | 183.67 | -1.19 | 0.02 |
| 6.00 | 2.00 | 3.67 | -1.28 | 0.70 |
| 585.00 | 636.00 | 632.67 | 1.01 | 0.91 |
| 902.00 | 844.00 | 931.67 | 1.19 | 0.00 |
| 0.00 | 2.00 | 1.00 | -2.21 | 0.35 |
| 1303.00 | 1532.00 | 1420.67 | -1.06 | 0.18 |
| 1337.00 | 1564.00 | 1516.33 | 1.03 | 0.43 |
| 1966.00 | 2187.00 | 2183.00 | 1.05 | 0.21 |
| 2271.00 | 2343.00 | 2295.33 | 1.08 | 0.09 |
| 4.00 | 6.00 | 9.67 | 1.66 | 0.23 |
| 6476.00 | 6786.00 | 6792.00 | 1.19 | 0.00 |
| 42.00 | 56.00 | 49.67 | 1.40 | 0.02 |
| 6.00 | 18.00 | 11.00 | 1.35 | 0.31 |
| 3696.00 | 3975.00 | 3895.00 | 1.07 | 0.07 |
| 1046.00 | 1175.00 | 1136.00 | 1.25 | 0.00 |
| 4668.00 | 5229.00 | 5109.00 | 1.13 | 0.00 |
| 11067.00 | 12292.00 | 11968.00 | 1.17 | 0.00 |
| 1462.00 | 1591.00 | 1530.33 | 1.10 | 0.04 |
| 941.00 | 990.00 | 989.67 | -1.09 | 0.03 |
| 83.00 | 86.00 | 84.00 | -1.20 | 0.06 |
| 729.00 | 847.00 | 817.67 | -1.00 | 1.00 |
| 1240.00 | 1195.00 | 1174.00 | 1.06 | 0.34 |
| 0.00 | 1.00 | 0.33 | -5.43 | 0.08 |
| 1.00 | 1.00 | 1.00 | -1.01 | 1.00 |
| 11.00 | 15.00 | 11.67 | 1.32 | 0.32 |

| | | | | |
|---|---|---|---|---|
| 84.00 | 95.00 | 82.67 | -1.36 | 0.00 |
| 35.00 | 49.00 | 47.00 | 1.02 | 0.96 |
| 1316.00 | 1256.00 | 1311.33 | -1.09 | 0.04 |
| 1043.00 | 1313.00 | 1218.00 | -1.09 | 0.07 |
| 93.00 | 99.00 | 93.00 | 1.40 | 0.00 |
| 232.00 | 172.00 | 177.67 | -1.29 | 0.09 |
| 222.00 | 263.00 | 249.00 | 1.04 | 0.59 |
| 1090.00 | 1152.00 | 1147.33 | -1.02 | 0.61 |
| 464.00 | 545.00 | 499.67 | 1.02 | 0.79 |
| 1066.00 | 1313.00 | 1182.67 | -1.01 | 0.91 |
| 76.00 | 100.00 | 86.67 | 1.03 | 0.79 |
| 17.00 | 9.00 | 13.00 | 1.10 | 0.76 |
| 554.00 | 599.00 | 603.67 | -1.00 | 0.99 |
| 724.00 | 846.00 | 843.00 | -1.05 | 0.32 |
| 1076.00 | 1210.00 | 1148.33 | 1.01 | 0.88 |
| 516.00 | 534.00 | 519.67 | -1.03 | 0.52 |
| 311.00 | 314.00 | 316.00 | -1.52 | 0.00 |
| 819.00 | 844.00 | 853.00 | -1.17 | 0.00 |
| 13.00 | 7.00 | 10.00 | 1.40 | 0.29 |
| 564.00 | 533.00 | 547.33 | 1.13 | 0.03 |
| 219.00 | 215.00 | 230.00 | -1.04 | 0.57 |
| 287.00 | 286.00 | 269.67 | -1.16 | 0.06 |
| 82.00 | 76.00 | 77.33 | -1.28 | 0.01 |
| 546.00 | 592.00 | 565.67 | 1.01 | 0.83 |
| 311.00 | 316.00 | 322.00 | -1.15 | 0.02 |
| 52.00 | 54.00 | 49.67 | 1.11 | 0.46 |
| 4386.00 | 4747.00 | 4383.00 | 1.04 | 0.45 |
| 461.00 | 506.00 | 486.67 | -1.09 | 0.07 |
| 1838.00 | 1736.00 | 1776.00 | -1.17 | 0.00 |
| 2718.00 | 2934.00 | 2853.00 | -1.23 | 0.00 |
| 298.00 | 401.00 | 335.67 | -1.05 | 0.54 |
| 14.00 | 15.00 | 15.33 | -1.78 | 0.00 |
| 527.00 | 570.00 | 566.00 | 1.01 | 0.81 |
| 588.00 | 668.00 | 630.67 | -1.01 | 0.91 |
| 188.00 | 184.00 | 197.00 | -1.19 | 0.01 |
| 7.00 | 5.00 | 5.67 | -1.94 | 0.04 |
| 444.00 | 478.00 | 474.00 | -1.27 | 0.00 |
| 546.00 | 615.00 | 617.67 | -1.08 | 0.15 |
| 119.00 | 129.00 | 128.67 | 1.16 | 0.07 |
| 421.00 | 446.00 | 441.67 | -1.21 | 0.00 |
| 284.00 | 294.00 | 287.67 | -1.03 | 0.70 |
| 974.00 | 946.00 | 963.00 | -1.06 | 0.19 |
| 550.00 | 538.00 | 576.00 | -1.12 | 0.03 |
| 97.00 | 132.00 | 112.67 | -1.01 | 0.90 |
| 332.00 | 402.00 | 391.67 | 1.20 | 0.01 |
| 19.00 | 12.00 | 16.00 | -1.71 | 0.01 |
| 289.00 | 349.00 | 312.33 | -1.02 | 0.87 |
| 15.00 | 21.00 | 18.67 | -1.01 | 1.00 |

| | | | | |
|---|---|---|---|---|
| 163.00 | 174.00 | 160.00 | 1.05 | 0.54 |
| 4333.00 | 4764.00 | 4667.00 | 1.03 | 0.40 |
| 210.00 | 199.00 | 202.67 | -1.10 | 0.21 |
| 921.00 | 1040.00 | 942.33 | -1.16 | 0.01 |
| 2.00 | 2.00 | 3.00 | -1.12 | 1.00 |
| 1688.00 | 1642.00 | 1613.00 | -1.13 | 0.01 |
| 2012.00 | 2227.00 | 2169.00 | 1.04 | 0.30 |
| 2718.00 | 3282.00 | 3160.33 | 1.18 | 0.00 |
| 1816.00 | 1863.00 | 1835.00 | -1.08 | 0.06 |
| 353.00 | 395.00 | 374.00 | -1.08 | 0.18 |
| 1275.00 | 1222.00 | 1209.00 | 1.04 | 0.46 |
| 271.00 | 324.00 | 300.67 | -1.28 | 0.00 |
| 1440.00 | 1707.00 | 1630.67 | 1.06 | 0.17 |
| 3708.00 | 3835.00 | 3886.33 | 1.07 | 0.03 |
| 522.00 | 576.00 | 574.33 | 1.10 | 0.05 |
| 1144.00 | 1264.00 | 1177.00 | 1.01 | 0.80 |
| 263.00 | 245.00 | 249.67 | -1.10 | 0.15 |
| 814.00 | 876.00 | 899.00 | 1.12 | 0.02 |
| 162.00 | 184.00 | 178.00 | 1.13 | 0.09 |
| 1503.00 | 1588.00 | 1542.33 | -1.10 | 0.01 |
| 1414.00 | 1397.00 | 1434.67 | -1.12 | 0.00 |
| 578.00 | 639.00 | 646.00 | 1.01 | 0.82 |
| 243.00 | 213.00 | 242.67 | 1.07 | 0.37 |
| 356.00 | 340.00 | 348.67 | 1.01 | 0.85 |
| 111.00 | 95.00 | 98.67 | -1.72 | 0.00 |
| 888.00 | 875.00 | 954.33 | 1.12 | 0.02 |
| 354.00 | 384.00 | 395.00 | 1.25 | 0.00 |
| 490.00 | 554.00 | 527.67 | -1.17 | 0.00 |
| 879.00 | 999.00 | 1005.00 | 1.03 | 0.57 |
| 1234.00 | 1420.00 | 1391.00 | 1.04 | 0.40 |
| 1326.00 | 1516.00 | 1481.33 | 1.26 | 0.00 |
| 789.00 | 924.00 | 877.67 | 1.12 | 0.02 |
| 187.00 | 210.00 | 206.00 | -1.11 | 0.11 |
| 23.00 | 20.00 | 21.33 | 1.19 | 0.37 |
| 800.00 | 902.00 | 815.67 | -1.20 | 0.00 |
| 299.00 | 312.00 | 302.00 | 1.06 | 0.41 |
| 49.00 | 49.00 | 49.33 | -1.65 | 0.00 |
| 355.00 | 326.00 | 376.00 | 1.16 | 0.09 |
| 472.00 | 510.00 | 499.00 | 1.13 | 0.01 |
| 422.00 | 412.00 | 438.67 | -1.19 | 0.00 |
| 950.00 | 998.00 | 1010.33 | -1.02 | 0.66 |
| 843.00 | 988.00 | 910.67 | -1.17 | 0.00 |
| 776.00 | 723.00 | 772.33 | -1.01 | 0.88 |
| 429.00 | 453.00 | 432.00 | -1.03 | 0.71 |
| 934.00 | 978.00 | 937.67 | 1.06 | 0.27 |
| 1233.00 | 1500.00 | 1358.67 | 1.11 | 0.03 |
| 487.00 | 588.00 | 562.67 | 1.21 | 0.00 |
| 756.00 | 760.00 | 787.33 | 1.11 | 0.04 |

| | | | | |
|---|---|---|---|---|
| 966.00 | 959.00 | 990.33 | -1.20 | 0.00 |
| 969.00 | 1018.00 | 1038.00 | 1.07 | 0.13 |
| 3566.00 | 3252.00 | 3436.00 | -1.12 | 0.01 |
| 1926.00 | 2202.00 | 2143.67 | -1.00 | 0.97 |
| 2889.00 | 2855.00 | 2977.33 | -1.10 | 0.01 |
| 1855.00 | 1871.00 | 1947.67 | -1.09 | 0.03 |
| 1274.00 | 1226.00 | 1307.00 | -1.06 | 0.17 |
| 2429.00 | 2793.00 | 2561.33 | 1.08 | 0.09 |
| 290.00 | 341.00 | 333.33 | -1.04 | 0.53 |
| 1716.00 | 1676.00 | 1690.00 | -1.03 | 0.46 |
| 864.00 | 852.00 | 861.00 | -1.24 | 0.00 |
| 1548.00 | 1820.00 | 1765.67 | 1.18 | 0.00 |
| 333.00 | 343.00 | 375.00 | 1.03 | 0.70 |
| 361.00 | 406.00 | 411.33 | -1.05 | 0.46 |
| 2178.00 | 2259.00 | 2267.67 | 1.03 | 0.48 |
| 160.00 | 155.00 | 167.00 | 1.24 | 0.01 |
| 1750.00 | 2010.00 | 1929.33 | -1.06 | 0.12 |
| 925.00 | 1038.00 | 1006.00 | 1.17 | 0.00 |
| 4180.00 | 4482.00 | 4733.00 | 1.11 | 0.02 |
| 278.00 | 303.00 | 294.00 | -1.10 | 0.19 |
| 3010.00 | 3315.00 | 3173.33 | -1.17 | 0.00 |
| 25.00 | 25.00 | 26.00 | -1.23 | 0.22 |
| 178.00 | 167.00 | 160.67 | -1.33 | 0.00 |
| 1109.00 | 1089.00 | 1068.67 | -1.22 | 0.00 |
| 20.00 | 15.00 | 17.00 | -1.19 | 0.50 |
| 272.00 | 306.00 | 289.33 | 1.09 | 0.20 |
| 14.00 | 37.00 | 24.00 | 1.27 | 0.29 |
| 3.00 | 0.00 | 2.67 | -1.62 | 0.39 |
| 48.00 | 57.00 | 47.67 | -1.03 | 0.92 |
| 80.00 | 100.00 | 94.00 | -1.00 | 0.97 |
| 97.00 | 109.00 | 114.00 | -1.17 | 0.08 |
| 2533.00 | 2540.00 | 2570.00 | -1.50 | 0.00 |
| 721.00 | 734.00 | 752.00 | 1.15 | 0.00 |
| 115.00 | 111.00 | 111.67 | -1.12 | 0.28 |
| 687.00 | 674.00 | 668.00 | -1.35 | 0.00 |
| 1377.00 | 1574.00 | 1506.00 | 1.00 | 0.99 |
| 434.00 | 400.00 | 415.33 | 1.18 | 0.00 |
| 155.00 | 168.00 | 165.00 | 1.18 | 0.03 |
| 254.00 | 193.00 | 225.67 | 1.04 | 0.60 |
| 20.00 | 19.00 | 21.00 | -1.01 | 1.00 |
| 464.00 | 461.00 | 488.00 | -1.13 | 0.01 |
| 1687.00 | 1864.00 | 1773.33 | 1.07 | 0.17 |
| 1178.00 | 1254.00 | 1283.00 | 1.04 | 0.30 |
| 184.00 | 168.00 | 185.00 | 1.12 | 0.13 |
| 186.00 | 201.00 | 202.67 | 1.84 | 0.00 |
| 158.00 | 189.00 | 181.33 | -1.61 | 0.00 |
| 2927.00 | 3143.00 | 2981.00 | 1.05 | 0.25 |
| 1007.00 | 1079.00 | 1077.33 | -1.02 | 0.66 |

| | | | | |
|---|---|---|---|---|
| 375.00 | 379.00 | 367.33 | -1.21 | 0.00 |
| 831.00 | 891.00 | 845.33 | -1.06 | 0.26 |
| 442.00 | 544.00 | 535.00 | -1.04 | 0.50 |
| 415.00 | 462.00 | 466.33 | -1.20 | 0.00 |
| 925.00 | 995.00 | 981.67 | -1.24 | 0.00 |
| 1709.00 | 1904.00 | 1819.00 | -1.09 | 0.03 |
| 23.00 | 28.00 | 32.33 | -2.34 | 0.00 |
| 984.00 | 1030.00 | 1015.67 | -1.11 | 0.01 |
| 140.00 | 176.00 | 167.67 | -1.29 | 0.00 |
| 1304.00 | 1384.00 | 1398.33 | -1.05 | 0.26 |
| 294.00 | 445.00 | 357.67 | -1.04 | 0.66 |
| 567.00 | 682.00 | 621.67 | -1.14 | 0.01 |
| 23.00 | 25.00 | 24.67 | -1.33 | 0.09 |
| 229.00 | 200.00 | 213.33 | -1.03 | 0.73 |
| 2485.00 | 2601.00 | 2569.00 | 1.09 | 0.02 |
| 245.00 | 252.00 | 257.00 | 1.05 | 0.55 |
| 259.00 | 220.00 | 235.00 | 1.21 | 0.02 |
| 13.00 | 11.00 | 11.33 | 1.24 | 0.45 |
| 130.00 | 144.00 | 131.67 | 1.19 | 0.16 |
| 20.00 | 11.00 | 15.00 | -1.15 | 0.62 |
| 3395.00 | 3585.00 | 3458.67 | -1.10 | 0.01 |
| 244.00 | 292.00 | 279.67 | -1.20 | 0.01 |
| 106.00 | 155.00 | 143.33 | -1.08 | 0.37 |
| 161.00 | 177.00 | 182.00 | -1.12 | 0.11 |
| 576.00 | 613.00 | 592.33 | 1.04 | 0.42 |
| 1755.00 | 1826.00 | 1726.00 | -1.39 | 0.00 |
| 331.00 | 377.00 | 384.00 | -1.01 | 0.92 |
| 830.00 | 950.00 | 920.00 | 1.11 | 0.02 |
| 442.00 | 494.00 | 459.67 | 1.07 | 0.25 |
| 306.00 | 297.00 | 317.33 | -1.03 | 0.64 |
| 1237.00 | 1207.00 | 1203.33 | -1.40 | 0.00 |
| 21.00 | 20.00 | 23.33 | -1.49 | 0.01 |
| 2753.00 | 2584.00 | 2782.67 | -1.14 | 0.00 |
| 2071.00 | 2322.00 | 2179.00 | 1.02 | 0.67 |
| 959.00 | 1265.00 | 1129.33 | 1.15 | 0.01 |
| 1784.00 | 2138.00 | 1998.33 | -1.02 | 0.63 |
| 1151.00 | 1125.00 | 1149.00 | -1.20 | 0.00 |
| 1841.00 | 1973.00 | 1934.00 | -1.03 | 0.53 |
| 327.00 | 409.00 | 362.00 | -1.13 | 0.06 |
| 600.00 | 739.00 | 694.00 | -1.13 | 0.02 |
| 840.00 | 866.00 | 889.00 | 1.10 | 0.04 |
| 3.00 | 5.00 | 6.00 | -1.84 | 0.05 |
| 2197.00 | 2452.00 | 2323.67 | 1.06 | 0.17 |
| 14.00 | 21.00 | 14.00 | 1.43 | 0.26 |
| 605.00 | 708.00 | 695.33 | 1.05 | 0.40 |
| 4343.00 | 4752.00 | 4631.67 | 1.03 | 0.33 |
| 316.00 | 363.00 | 350.33 | -1.08 | 0.18 |
| 204.00 | 211.00 | 213.67 | 1.18 | 0.02 |

| | | | | |
|---|---|---|---|---|
| 2254.00 | 2283.00 | 2228.67 | -1.04 | 0.36 |
| 630.00 | 753.00 | 696.67 | 1.12 | 0.03 |
| 241.00 | 250.00 | 253.67 | -1.14 | 0.03 |
| 49.00 | 61.00 | 57.67 | 1.20 | 0.15 |
| 1383.00 | 1529.00 | 1474.33 | -1.09 | 0.04 |
| 262.00 | 231.00 | 233.67 | -1.29 | 0.00 |
| 185.00 | 232.00 | 223.33 | 1.16 | 0.04 |
| 1227.00 | 1470.00 | 1398.00 | 1.04 | 0.40 |
| 24.00 | 24.00 | 22.67 | -1.44 | 0.04 |
| 734.00 | 914.00 | 857.00 | -1.11 | 0.04 |
| 222.00 | 222.00 | 231.00 | -1.10 | 0.18 |
| 115.00 | 111.00 | 116.33 | 1.31 | 0.00 |
| 83.00 | 96.00 | 96.33 | -1.01 | 0.97 |
| 181.00 | 176.00 | 183.00 | 1.29 | 0.00 |
| 81.00 | 93.00 | 88.33 | 1.03 | 0.81 |
| 123.00 | 185.00 | 159.67 | 1.09 | 0.35 |
| 18.00 | 19.00 | 16.67 | -1.25 | 0.28 |
| 704.00 | 675.00 | 708.00 | -1.13 | 0.01 |
| 307.00 | 367.00 | 345.67 | 1.01 | 0.95 |
| 2779.00 | 3146.00 | 3084.00 | 1.05 | 0.16 |
| 1040.00 | 1267.00 | 1162.67 | 1.17 | 0.00 |
| 176.00 | 174.00 | 179.33 | -1.02 | 0.82 |
| 7.00 | 6.00 | 5.00 | 1.14 | 0.85 |
| 271.00 | 270.00 | 282.00 | 1.02 | 0.75 |
| 345.00 | 381.00 | 369.67 | 1.07 | 0.23 |
| 15.00 | 21.00 | 18.33 | 1.13 | 0.57 |
| 119.00 | 118.00 | 118.00 | -1.49 | 0.00 |
| 90.00 | 88.00 | 77.33 | -1.22 | 0.15 |
| 302.00 | 300.00 | 311.67 | 1.04 | 0.55 |
| 322.00 | 309.00 | 316.00 | -1.31 | 0.00 |
| 441.00 | 473.00 | 488.00 | 1.17 | 0.03 |
| 164.00 | 207.00 | 193.33 | 1.01 | 0.89 |
| 171.00 | 215.00 | 187.33 | -1.30 | 0.00 |
| 310.00 | 364.00 | 336.00 | -1.11 | 0.07 |
| 218.00 | 203.00 | 213.33 | 1.15 | 0.05 |
| 3101.00 | 2928.00 | 3054.67 | -1.34 | 0.00 |
| 574.00 | 623.00 | 611.00 | -1.01 | 0.87 |
| 83.00 | 79.00 | 87.00 | -1.04 | 0.73 |
| 114.00 | 103.00 | 102.33 | -1.09 | 0.41 |
| 843.00 | 1076.00 | 989.67 | 1.11 | 0.04 |
| 442.00 | 499.00 | 476.33 | -1.04 | 0.42 |
| 108.00 | 103.00 | 111.67 | -1.12 | 0.21 |
| 287.00 | 377.00 | 337.00 | 1.16 | 0.04 |
| 1124.00 | 1301.00 | 1264.33 | -1.09 | 0.05 |
| 11.00 | 17.00 | 14.33 | 1.09 | 0.75 |
| 395.00 | 327.00 | 390.67 | -1.04 | 0.59 |
| 4.00 | 2.00 | 3.67 | -1.99 | 0.09 |
| 3346.00 | 3693.00 | 3520.00 | -1.01 | 0.87 |

| | | | | |
|---|---|---|---|---|
| 213.00 | 281.00 | 256.00 | -1.13 | 0.06 |
| 596.00 | 694.00 | 704.33 | -1.04 | 0.45 |
| 287.00 | 340.00 | 321.67 | -1.11 | 0.12 |
| 206.00 | 191.00 | 202.33 | -1.13 | 0.08 |
| 34.00 | 30.00 | 34.00 | -1.00 | 1.00 |
| 343.00 | 450.00 | 399.33 | 1.03 | 0.71 |
| 1268.00 | 1375.00 | 1351.67 | -1.07 | 0.09 |
| 122.00 | 97.00 | 114.33 | 1.47 | 0.00 |
| 1103.00 | 1207.00 | 1179.00 | -1.10 | 0.02 |
| 19.00 | 14.00 | 15.67 | -1.70 | 0.01 |
| 99.00 | 121.00 | 112.00 | 1.45 | 0.00 |
| 50.00 | 52.00 | 50.00 | 1.16 | 0.29 |
| 394.00 | 444.00 | 433.33 | -1.16 | 0.00 |
| 119.00 | 140.00 | 131.67 | -1.03 | 0.71 |
| 369.00 | 388.00 | 408.00 | -1.08 | 0.17 |
| 255.00 | 267.00 | 260.33 | -1.02 | 0.74 |
| 4.00 | 2.00 | 2.00 | -2.28 | 0.12 |
| 829.00 | 909.00 | 944.33 | 1.01 | 0.78 |
| 105.00 | 110.00 | 108.67 | -1.02 | 0.86 |
| 255.00 | 313.00 | 308.67 | -1.02 | 0.78 |
| 296.00 | 410.00 | 354.67 | -1.11 | 0.14 |
| 83.00 | 88.00 | 86.33 | 1.41 | 0.00 |
| 611.00 | 674.00 | 617.00 | -1.15 | 0.02 |
| 853.00 | 1015.00 | 979.33 | 1.34 | 0.00 |
| 1557.00 | 1769.00 | 1687.00 | -1.01 | 0.88 |
| 1032.00 | 1158.00 | 1133.00 | -1.14 | 0.00 |
| 436.00 | 484.00 | 493.33 | 1.25 | 0.00 |
| 211.00 | 245.00 | 224.33 | -1.13 | 0.12 |
| 716.00 | 700.00 | 729.67 | 1.06 | 0.23 |
| 13.00 | 39.00 | 26.33 | -1.01 | 1.00 |
| 25.00 | 22.00 | 25.00 | -1.05 | 0.82 |
| 102.00 | 124.00 | 110.33 | 1.02 | 0.87 |
| 15.00 | 10.00 | 13.67 | -1.36 | 0.20 |
| 743.00 | 813.00 | 837.67 | 1.09 | 0.08 |
| 255.00 | 308.00 | 292.67 | 1.03 | 0.69 |
| 1067.00 | 1096.00 | 1118.00 | -1.00 | 0.91 |
| 272.00 | 312.00 | 309.33 | -1.13 | 0.04 |
| 1024.00 | 1111.00 | 1120.33 | -1.07 | 0.16 |
| 418.00 | 542.00 | 472.67 | 1.05 | 0.53 |
| 336.00 | 347.00 | 346.00 | 1.06 | 0.33 |
| 514.00 | 619.00 | 612.67 | 1.15 | 0.02 |
| 61.00 | 86.00 | 69.67 | 1.17 | 0.20 |
| 606.00 | 659.00 | 664.33 | 1.15 | 0.01 |
| 447.00 | 489.00 | 481.33 | -1.21 | 0.00 |
| 2039.00 | 1968.00 | 2056.33 | -1.31 | 0.00 |
| 378.00 | 400.00 | 417.67 | 1.17 | 0.02 |
| 143.00 | 169.00 | 164.67 | 1.17 | 0.06 |
| 231.00 | 266.00 | 245.67 | 1.08 | 0.27 |

| | | | | |
|---|---|---|---|---|
| 754.00 | 890.00 | 871.67 | 1.06 | 0.23 |
| 1499.00 | 1590.00 | 1533.33 | -1.01 | 0.74 |
| 1097.00 | 1124.00 | 1145.67 | -1.06 | 0.15 |
| 94.00 | 156.00 | 121.67 | -1.08 | 0.47 |
| 112.00 | 133.00 | 121.33 | 1.03 | 0.79 |
| 119.00 | 133.00 | 125.00 | 1.07 | 0.43 |
| 447.00 | 519.00 | 504.67 | -1.07 | 0.23 |
| 129.00 | 142.00 | 153.00 | -1.09 | 0.28 |
| 119.00 | 124.00 | 126.00 | 1.13 | 0.16 |
| 918.00 | 1152.00 | 1038.00 | 1.03 | 0.56 |
| 16.00 | 17.00 | 14.33 | -1.25 | 0.37 |
| 2000.00 | 2207.00 | 2143.33 | 1.04 | 0.29 |
| 157.00 | 138.00 | 143.00 | -1.14 | 0.13 |
| 1071.00 | 1414.00 | 1266.00 | 1.24 | 0.00 |
| 252.00 | 296.00 | 281.00 | -1.15 | 0.03 |
| 236.00 | 274.00 | 256.00 | -1.02 | 0.79 |
| 14.00 | 14.00 | 15.00 | -1.06 | 0.92 |
| 1429.00 | 1426.00 | 1448.33 | -1.09 | 0.03 |
| 968.00 | 1020.00 | 1002.67 | -1.05 | 0.24 |
| 963.00 | 1032.00 | 994.33 | -1.13 | 0.01 |
| 398.00 | 424.00 | 439.67 | -1.11 | 0.06 |
| 372.00 | 402.00 | 421.00 | 1.13 | 0.09 |
| 29.00 | 26.00 | 23.33 | 1.00 | 1.00 |
| 227.00 | 259.00 | 250.00 | -1.01 | 0.87 |
| 161.00 | 165.00 | 168.33 | -1.42 | 0.00 |
| 120.00 | 92.00 | 101.67 | 1.03 | 0.77 |
| 888.00 | 1039.00 | 993.00 | 1.20 | 0.00 |
| 161.00 | 200.00 | 195.33 | 1.04 | 0.62 |
| 2.00 | 8.00 | 4.33 | -1.01 | 1.00 |
| 752.00 | 893.00 | 860.67 | 1.25 | 0.00 |
| 225.00 | 268.00 | 264.33 | -1.03 | 0.63 |
| 523.00 | 624.00 | 592.33 | -1.06 | 0.28 |
| 977.00 | 1087.00 | 1070.67 | -1.21 | 0.00 |
| 1749.00 | 2077.00 | 2031.00 | -1.38 | 0.00 |
| 326.00 | 296.00 | 320.00 | 1.00 | 0.99 |
| 900.00 | 1003.00 | 958.00 | -1.20 | 0.00 |
| 703.00 | 748.00 | 740.67 | -1.07 | 0.11 |
| 160.00 | 166.00 | 173.33 | -1.11 | 0.21 |
| 178.00 | 175.00 | 183.00 | 1.12 | 0.14 |
| 4.00 | 0.00 | 3.00 | -1.23 | 0.83 |
| 613.00 | 696.00 | 665.67 | -1.07 | 0.12 |
| 3.00 | 2.00 | 1.67 | 1.58 | 0.73 |
| 378.00 | 376.00 | 392.33 | -1.10 | 0.16 |
| 76.00 | 78.00 | 78.00 | 1.12 | 0.31 |
| 103.00 | 90.00 | 91.00 | 1.17 | 0.15 |
| 607.00 | 693.00 | 659.33 | 1.06 | 0.22 |
| 396.00 | 476.00 | 419.00 | -1.11 | 0.14 |
| 5.00 | 2.00 | 3.00 | -1.23 | 0.83 |

| | | | | |
|---|---|---|---|---|
| 717.00 | 683.00 | 733.33 | 1.03 | 0.54 |
| 200.00 | 196.00 | 199.67 | 1.16 | 0.03 |
| 83.00 | 123.00 | 107.00 | 1.00 | 1.00 |
| 153.00 | 202.00 | 174.00 | 1.55 | 0.00 |
| 1309.00 | 1463.00 | 1377.33 | -1.17 | 0.00 |
| 1343.00 | 1522.00 | 1498.00 | -1.10 | 0.03 |
| 50.00 | 60.00 | 50.33 | 1.27 | 0.11 |
| 1738.00 | 1757.00 | 1792.67 | -1.06 | 0.12 |
| 83.00 | 66.00 | 72.00 | 1.15 | 0.22 |
| 124.00 | 101.00 | 117.67 | -1.68 | 0.00 |
| 54.00 | 47.00 | 51.00 | -1.23 | 0.10 |
| 311.00 | 335.00 | 353.67 | 1.09 | 0.23 |
| 717.00 | 655.00 | 699.67 | -1.14 | 0.01 |
| 9.00 | 14.00 | 11.67 | -1.10 | 0.82 |
| 254.00 | 316.00 | 280.00 | 1.38 | 0.00 |
| 335.00 | 319.00 | 329.33 | -1.23 | 0.00 |
| 52.00 | 51.00 | 57.00 | -1.30 | 0.02 |
| 444.00 | 365.00 | 409.67 | -1.13 | 0.07 |
| 310.00 | 326.00 | 335.33 | -1.02 | 0.71 |
| 329.00 | 355.00 | 336.67 | -1.20 | 0.00 |
| 858.00 | 944.00 | 950.67 | 1.12 | 0.01 |
| 154.00 | 158.00 | 167.33 | 1.07 | 0.42 |
| 88.00 | 105.00 | 104.33 | -1.18 | 0.07 |
| 124.00 | 133.00 | 126.00 | -1.41 | 0.00 |
| 29.00 | 41.00 | 34.33 | -1.08 | 0.65 |
| 326.00 | 387.00 | 381.67 | -1.06 | 0.30 |
| 30.00 | 46.00 | 40.67 | -1.15 | 0.34 |
| 57.00 | 63.00 | 69.67 | -1.07 | 0.59 |
| 175.00 | 202.00 | 180.33 | -1.03 | 0.74 |
| 547.00 | 563.00 | 583.33 | 1.19 | 0.00 |
| 5.00 | 1.00 | 2.33 | -1.42 | 0.64 |
| 3.00 | 5.00 | 3.00 | 1.11 | 1.00 |
| 154.00 | 198.00 | 161.33 | 1.09 | 0.42 |
| 254.00 | 277.00 | 263.33 | -1.03 | 0.65 |
| 160.00 | 166.00 | 160.00 | -1.05 | 0.59 |
| 20.00 | 28.00 | 25.67 | -1.34 | 0.06 |
| 246.00 | 256.00 | 239.67 | -1.17 | 0.03 |
| 118.00 | 162.00 | 141.67 | -1.39 | 0.00 |
| 439.00 | 467.00 | 482.33 | 1.20 | 0.00 |
| 6.00 | 2.00 | 6.67 | 1.50 | 0.32 |
| 149.00 | 157.00 | 166.67 | -1.04 | 0.59 |
| 349.00 | 367.00 | 345.33 | -1.05 | 0.49 |
| 297.00 | 340.00 | 307.00 | -1.31 | 0.00 |
| 3.00 | 6.00 | 4.33 | -1.01 | 1.00 |
| 17.00 | 19.00 | 21.67 | 1.60 | 0.03 |
| 627.00 | 564.00 | 622.00 | 1.02 | 0.76 |
| 902.00 | 962.00 | 902.33 | -1.17 | 0.00 |
| 351.00 | 329.00 | 352.00 | -1.06 | 0.37 |

| | | | | |
|---|---|---|---|---|
| 211.00 | 239.00 | 240.67 | -1.11 | 0.11 |
| 291.00 | 328.00 | 312.67 | 1.07 | 0.27 |
| 1196.00 | 1305.00 | 1238.67 | -1.01 | 0.84 |
| 24.00 | 20.00 | 20.00 | -1.10 | 0.73 |
| 38.00 | 40.00 | 37.67 | 1.05 | 0.75 |
| 947.00 | 818.00 | 846.67 | -1.23 | 0.00 |
| 58.00 | 36.00 | 45.00 | -1.04 | 0.84 |
| 19.00 | 12.00 | 14.67 | -1.74 | 0.01 |
| 109.00 | 104.00 | 108.67 | 1.05 | 0.58 |
| 486.00 | 463.00 | 480.00 | -1.12 | 0.03 |
| 10.00 | 4.00 | 6.67 | -1.71 | 0.09 |
| 306.00 | 281.00 | 297.67 | -1.04 | 0.56 |
| 322.00 | 356.00 | 337.33 | -1.02 | 0.76 |
| 84.00 | 84.00 | 91.33 | -1.12 | 0.27 |
| 448.00 | 465.00 | 478.00 | 1.02 | 0.77 |
| 49.00 | 56.00 | 52.33 | 1.26 | 0.08 |
| 28.00 | 14.00 | 25.67 | 1.17 | 0.47 |
| 25.00 | 34.00 | 31.00 | -1.13 | 0.46 |
| 89.00 | 91.00 | 100.00 | 1.01 | 0.97 |
| 17.00 | 17.00 | 19.33 | 1.17 | 0.51 |
| 90.00 | 96.00 | 90.67 | -1.20 | 0.06 |
| 931.00 | 832.00 | 872.67 | -1.16 | 0.00 |
| 110.00 | 97.00 | 100.33 | 1.09 | 0.37 |
| 277.00 | 308.00 | 296.67 | -1.14 | 0.04 |
| 62.00 | 38.00 | 46.33 | 1.09 | 0.56 |
| 50.00 | 56.00 | 60.67 | 1.28 | 0.06 |
| 313.00 | 344.00 | 330.33 | 1.08 | 0.20 |
| 36.00 | 34.00 | 32.67 | -1.15 | 0.44 |
| 50.00 | 41.00 | 51.67 | -1.14 | 0.32 |
| 1008.00 | 1139.00 | 1043.67 | -1.13 | 0.03 |
| 240.00 | 274.00 | 256.00 | -1.22 | 0.00 |
| 515.00 | 575.00 | 573.67 | 1.25 | 0.00 |
| 1765.00 | 2006.00 | 1919.33 | 1.02 | 0.63 |
| 2126.00 | 2076.00 | 2143.67 | -1.18 | 0.00 |
| 424.00 | 471.00 | 466.00 | -1.09 | 0.07 |
| 1171.00 | 1313.00 | 1274.00 | -1.08 | 0.05 |
| 1132.00 | 1158.00 | 1137.33 | -1.22 | 0.00 |
| 80.00 | 92.00 | 85.33 | 1.15 | 0.18 |
| 1085.00 | 1282.00 | 1189.67 | -1.16 | 0.00 |
| 175.00 | 160.00 | 163.33 | -1.14 | 0.16 |
| 272.00 | 433.00 | 369.00 | 1.10 | 0.28 |
| 298.00 | 276.00 | 288.33 | -1.04 | 0.57 |
| 147.00 | 155.00 | 156.67 | 1.07 | 0.46 |
| 53.00 | 54.00 | 52.67 | 1.05 | 0.75 |
| 237.00 | 216.00 | 232.67 | -1.17 | 0.03 |
| 123.00 | 115.00 | 124.67 | 1.06 | 0.53 |
| 41.00 | 38.00 | 39.33 | 1.25 | 0.17 |
| 14.00 | 10.00 | 13.00 | 1.42 | 0.19 |

| | | | | |
|---|---|---|---|---|
| 247.00 | 293.00 | 272.00 | -1.00 | 0.97 |
| 42.00 | 36.00 | 37.00 | -1.01 | 1.00 |
| 101.00 | 134.00 | 126.33 | 1.12 | 0.29 |
| 7.00 | 3.00 | 5.00 | 1.06 | 1.00 |
| 5.00 | 9.00 | 7.33 | 1.14 | 0.76 |
| 7.00 | 12.00 | 10.67 | 1.13 | 0.71 |
| 50.00 | 65.00 | 60.33 | -1.28 | 0.02 |
| 617.00 | 595.00 | 619.00 | 1.10 | 0.06 |
| 48.00 | 67.00 | 57.67 | -1.17 | 0.20 |
| 63.00 | 65.00 | 70.33 | -1.04 | 0.70 |
| 428.00 | 425.00 | 433.67 | -1.09 | 0.09 |
| 218.00 | 256.00 | 235.33 | 1.02 | 0.78 |
| 526.00 | 622.00 | 590.00 | -1.17 | 0.00 |
| 167.00 | 156.00 | 160.67 | 1.04 | 0.66 |
| 80.00 | 90.00 | 84.00 | -1.16 | 0.12 |
| 83.00 | 74.00 | 79.00 | -1.17 | 0.13 |
| 119.00 | 116.00 | 131.33 | -1.15 | 0.15 |
| 109.00 | 166.00 | 132.33 | -1.01 | 0.98 |
| 363.00 | 435.00 | 442.00 | 1.05 | 0.51 |
| 602.00 | 628.00 | 619.00 | -1.04 | 0.42 |
| 36.00 | 44.00 | 40.00 | -1.01 | 0.95 |
| 571.00 | 564.00 | 560.00 | -1.05 | 0.41 |
| 5.00 | 6.00 | 6.33 | -1.07 | 1.00 |
| 163.00 | 160.00 | 167.00 | -1.03 | 0.71 |
| 60.00 | 82.00 | 76.33 | -1.13 | 0.28 |
| 136.00 | 155.00 | 155.33 | 1.02 | 0.87 |
| 70.00 | 105.00 | 95.00 | -1.14 | 0.19 |
| 78.00 | 85.00 | 79.67 | 1.04 | 0.70 |
| 72.00 | 107.00 | 95.67 | -1.18 | 0.09 |
| 16.00 | 17.00 | 14.33 | -1.29 | 0.29 |
| 34.00 | 34.00 | 38.67 | 1.31 | 0.07 |
| 76.00 | 85.00 | 80.33 | -1.16 | 0.14 |
| 20.00 | 19.00 | 16.00 | -1.20 | 0.47 |
| 407.00 | 373.00 | 401.33 | -1.02 | 0.69 |
| 16.00 | 15.00 | 18.67 | 1.10 | 0.71 |
| 91.00 | 79.00 | 95.67 | -1.23 | 0.04 |
| 305.00 | 344.00 | 319.67 | -1.04 | 0.53 |
| 371.00 | 346.00 | 361.67 | 1.12 | 0.06 |
| 553.00 | 518.00 | 568.00 | -1.04 | 0.46 |
| 466.00 | 502.00 | 508.00 | -1.01 | 0.88 |
| 50.00 | 50.00 | 53.00 | -1.02 | 0.97 |
| 185.00 | 187.00 | 215.67 | -1.04 | 0.68 |
| 92.00 | 85.00 | 93.67 | -1.04 | 0.74 |
| 30.00 | 35.00 | 35.67 | 1.02 | 0.95 |
| 118.00 | 134.00 | 124.67 | -1.01 | 0.90 |
| 865.00 | 921.00 | 837.33 | -1.29 | 0.00 |
| 253.00 | 307.00 | 304.00 | 1.17 | 0.05 |
| 926.00 | 969.00 | 966.00 | 1.00 | 0.92 |

| | | | | |
|---|---|---|---|---|
| 979.00 | 878.00 | 906.67 | -1.14 | 0.02 |
| 533.00 | 565.00 | 561.67 | 1.20 | 0.00 |
| 397.00 | 330.00 | 363.33 | -1.19 | 0.01 |
| 167.00 | 139.00 | 159.00 | 1.09 | 0.38 |
| 78.00 | 88.00 | 86.33 | -1.03 | 0.84 |
| 75.00 | 82.00 | 86.33 | -1.08 | 0.39 |
| 2573.00 | 2486.00 | 2629.00 | -1.36 | 0.00 |
| 65.00 | 52.00 | 60.67 | -1.13 | 0.29 |
| 67.00 | 88.00 | 85.33 | 1.09 | 0.42 |
| 401.00 | 408.00 | 409.00 | -1.10 | 0.07 |
| 16.00 | 24.00 | 18.67 | -1.03 | 1.00 |
| 90.00 | 87.00 | 81.33 | 1.06 | 0.62 |
| 39.00 | 46.00 | 41.33 | -2.04 | 0.00 |
| 1648.00 | 1502.00 | 1523.67 | -1.31 | 0.00 |
| 12.00 | 5.00 | 8.67 | 1.07 | 0.89 |
| 113.00 | 115.00 | 106.33 | -1.36 | 0.00 |
| 69.00 | 61.00 | 65.67 | 1.12 | 0.34 |
| 236.00 | 249.00 | 247.00 | 1.07 | 0.29 |
| 105.00 | 128.00 | 125.33 | -1.10 | 0.23 |
| 172.00 | 171.00 | 177.00 | -1.06 | 0.45 |
| 360.00 | 315.00 | 342.67 | -1.32 | 0.00 |
| 175.00 | 201.00 | 189.33 | -1.06 | 0.42 |
| 121.00 | 146.00 | 128.33 | -1.07 | 0.51 |
| 2036.00 | 2117.00 | 2075.33 | -1.15 | 0.00 |
| 986.00 | 1107.00 | 1076.67 | 1.03 | 0.57 |
| 27.00 | 35.00 | 31.00 | -1.36 | 0.03 |
| 128.00 | 125.00 | 128.67 | 1.02 | 0.84 |
| 22.00 | 18.00 | 18.67 | -1.12 | 0.60 |
| 9.00 | 9.00 | 8.67 | -1.01 | 1.00 |
| 13.00 | 17.00 | 17.00 | -1.11 | 0.64 |
| 367.00 | 353.00 | 361.67 | 1.00 | 0.98 |
| 121.00 | 108.00 | 123.33 | -1.02 | 0.89 |
| 845.00 | 853.00 | 858.33 | 1.03 | 0.50 |
| 3755.00 | 4372.00 | 4065.00 | -1.03 | 0.49 |
| 875.00 | 991.00 | 996.33 | 1.10 | 0.04 |
| 113.00 | 82.00 | 103.67 | -1.10 | 0.32 |
| 1347.00 | 1508.00 | 1394.33 | 1.14 | 0.01 |
| 928.00 | 812.00 | 873.00 | -1.26 | 0.00 |
| 136.00 | 159.00 | 139.67 | 1.14 | 0.20 |
| 580.00 | 569.00 | 575.00 | -1.17 | 0.00 |
| 55.00 | 68.00 | 68.67 | -1.21 | 0.07 |
| 214.00 | 281.00 | 232.33 | 1.00 | 0.99 |
| 2633.00 | 3066.00 | 2892.33 | -1.17 | 0.00 |
| 37.00 | 50.00 | 51.33 | -1.30 | 0.04 |
| 23.00 | 27.00 | 24.33 | 1.03 | 0.87 |
| 20.00 | 26.00 | 21.00 | 1.24 | 0.28 |
| 2941.00 | 3090.00 | 3105.00 | -1.02 | 0.63 |
| 300.00 | 346.00 | 323.00 | -1.11 | 0.07 |

| | | | | |
|---|---|---|---|---|
| 114.00 | 102.00 | 101.33 | 1.03 | 0.81 |
| 126.00 | 146.00 | 142.67 | -1.02 | 0.83 |
| 91.00 | 124.00 | 111.33 | 1.14 | 0.18 |
| 577.00 | 649.00 | 651.67 | -1.29 | 0.00 |
| 244.00 | 241.00 | 238.33 | 1.51 | 0.00 |
| 344.00 | 386.00 | 376.00 | -1.02 | 0.70 |
| 87.00 | 98.00 | 88.67 | 1.16 | 0.16 |
| 354.00 | 392.00 | 359.33 | 1.14 | 0.04 |
| 105.00 | 144.00 | 125.33 | 1.16 | 0.11 |
| 20.00 | 31.00 | 23.67 | -1.26 | 0.20 |
| 5.00 | 16.00 | 10.33 | 1.38 | 0.29 |
| 84.00 | 99.00 | 96.00 | 1.20 | 0.06 |
| 848.00 | 856.00 | 877.00 | -1.09 | 0.12 |
| 218.00 | 248.00 | 239.00 | -1.03 | 0.63 |
| 261.00 | 257.00 | 262.00 | 1.01 | 0.91 |
| 395.00 | 467.00 | 451.00 | -1.04 | 0.44 |
| 1591.00 | 1697.00 | 1811.00 | -1.11 | 0.05 |
| 76.00 | 78.00 | 72.33 | 1.45 | 0.00 |
| 639.00 | 600.00 | 629.33 | -1.06 | 0.23 |
| 1249.00 | 1408.00 | 1342.33 | -1.08 | 0.07 |
| 50.00 | 41.00 | 42.00 | -1.30 | 0.05 |
| 972.00 | 1049.00 | 1064.33 | -1.10 | 0.02 |
| 113.00 | 96.00 | 103.00 | -1.38 | 0.00 |
| 601.00 | 761.00 | 745.33 | 1.01 | 0.91 |
| 154.00 | 153.00 | 148.67 | -1.37 | 0.00 |
| 951.00 | 898.00 | 981.67 | 1.00 | 0.97 |
| 193.00 | 173.00 | 175.33 | -1.11 | 0.21 |
| 9.00 | 7.00 | 8.00 | -1.22 | 0.59 |
| 1123.00 | 1218.00 | 1265.67 | 1.09 | 0.06 |
| 103.00 | 142.00 | 126.33 | 1.10 | 0.29 |
| 4.00 | 8.00 | 8.00 | -1.22 | 0.59 |
| 102.00 | 121.00 | 118.33 | 1.04 | 0.68 |
| 896.00 | 901.00 | 918.00 | -1.06 | 0.17 |
| 38.00 | 55.00 | 42.00 | 1.34 | 0.07 |
| 29.00 | 25.00 | 26.00 | 1.24 | 0.23 |
| 201.00 | 266.00 | 225.67 | 1.12 | 0.21 |
| 199.00 | 245.00 | 229.67 | -1.13 | 0.07 |
| 32.00 | 48.00 | 38.67 | -1.04 | 0.82 |
| 142.00 | 139.00 | 156.67 | 1.17 | 0.08 |
| 566.00 | 653.00 | 581.67 | -1.01 | 0.83 |
| 1.00 | 0.00 | 1.00 | -1.61 | 0.73 |
| 6.00 | 3.00 | 5.00 | 1.46 | 0.43 |
| 149.00 | 217.00 | 181.33 | 1.08 | 0.39 |
| 3.00 | 2.00 | 7.00 | 1.83 | 0.25 |
| 111.00 | 103.00 | 103.00 | 1.06 | 0.57 |
| 423.00 | 514.00 | 492.67 | 1.02 | 0.75 |
| 1117.00 | 1186.00 | 1185.00 | -1.02 | 0.56 |
| 727.00 | 759.00 | 793.67 | -1.02 | 0.67 |

| | | | | |
|---|---|---|---|---|
| 214.00 | 192.00 | 200.67 | -1.11 | 0.14 |
| 130.00 | 134.00 | 126.33 | -1.13 | 0.15 |
| 201.00 | 228.00 | 204.67 | 1.03 | 0.67 |
| 147.00 | 153.00 | 168.33 | -1.16 | 0.08 |
| 874.00 | 949.00 | 954.33 | -1.08 | 0.09 |
| 466.00 | 497.00 | 486.33 | -1.07 | 0.18 |
| 10.00 | 13.00 | 9.67 | -1.43 | 0.20 |
| 206.00 | 234.00 | 227.00 | 1.00 | 0.96 |
| 86.00 | 121.00 | 91.67 | 1.02 | 0.89 |
| 238.00 | 291.00 | 264.67 | 1.05 | 0.49 |
| 819.00 | 1027.00 | 948.00 | -1.16 | 0.00 |
| 2109.00 | 2019.00 | 2180.00 | 1.22 | 0.00 |
| 134.00 | 130.00 | 128.00 | -1.06 | 0.50 |
| 1724.00 | 2037.00 | 1860.67 | -1.01 | 0.80 |
| 172.00 | 164.00 | 160.67 | 1.27 | 0.01 |
| 51.00 | 49.00 | 45.33 | 1.03 | 0.82 |
| 176.00 | 168.00 | 166.67 | 1.11 | 0.17 |
| 37.00 | 27.00 | 35.00 | -1.00 | 1.00 |
| 175.00 | 158.00 | 181.33 | -1.02 | 0.79 |
| 328.00 | 328.00 | 329.33 | -1.13 | 0.03 |
| 566.00 | 591.00 | 593.33 | -1.10 | 0.03 |
| 275.00 | 243.00 | 260.00 | -1.18 | 0.01 |
| 257.00 | 300.00 | 280.33 | -1.16 | 0.01 |
| 506.00 | 488.00 | 517.33 | -1.07 | 0.20 |
| 3.00 | 12.00 | 7.33 | 1.20 | 0.65 |
| 131.00 | 130.00 | 142.33 | -1.11 | 0.20 |
| 2422.00 | 2398.00 | 2465.00 | -1.17 | 0.00 |
| 54.00 | 39.00 | 46.33 | -1.04 | 0.83 |
| 342.00 | 358.00 | 339.00 | -1.28 | 0.00 |
| 371.00 | 377.00 | 379.00 | -1.15 | 0.01 |
| 688.00 | 754.00 | 771.00 | 1.07 | 0.20 |
| 11.00 | 17.00 | 14.33 | 1.62 | 0.06 |
| 344.00 | 379.00 | 358.00 | -1.13 | 0.03 |
| 109.00 | 101.00 | 127.00 | -1.08 | 0.50 |
| 347.00 | 378.00 | 352.00 | -1.24 | 0.00 |
| 103.00 | 102.00 | 97.33 | -1.65 | 0.00 |
| 38.00 | 46.00 | 48.33 | 1.31 | 0.10 |
| 142.00 | 132.00 | 146.33 | -1.12 | 0.14 |
| 525.00 | 618.00 | 563.00 | 1.05 | 0.42 |
| 457.00 | 504.00 | 489.00 | -1.03 | 0.65 |
| 180.00 | 147.00 | 172.33 | -1.23 | 0.01 |
| 98.00 | 94.00 | 86.67 | -1.16 | 0.22 |
| 173.00 | 174.00 | 178.00 | -1.24 | 0.01 |
| 9.00 | 5.00 | 5.33 | 1.31 | 0.59 |
| 99.00 | 111.00 | 104.33 | -1.22 | 0.02 |
| 26.00 | 25.00 | 29.00 | 1.08 | 0.72 |
| 11.00 | 13.00 | 13.67 | 1.06 | 0.83 |
| 55.00 | 83.00 | 70.33 | -1.01 | 0.93 |

| | | | | |
|---|---|---|---|---|
| 82.00 | 112.00 | 112.67 | 1.08 | 0.48 |
| 806.00 | 837.00 | 827.67 | -1.11 | 0.02 |
| 1362.00 | 1552.00 | 1525.33 | -1.27 | 0.00 |
| 389.00 | 462.00 | 449.33 | 1.07 | 0.23 |
| 13.00 | 21.00 | 20.33 | 1.25 | 0.32 |
| 42.00 | 25.00 | 35.00 | -1.01 | 0.95 |
| 47.00 | 36.00 | 45.00 | 1.26 | 0.12 |
| 944.00 | 1152.00 | 1096.33 | 1.07 | 0.16 |
| 1144.00 | 1324.00 | 1282.67 | -1.00 | 0.92 |
| 399.00 | 486.00 | 434.33 | -1.11 | 0.10 |
| 3.00 | 3.00 | 2.33 | -1.70 | 0.37 |
| 138.00 | 185.00 | 170.67 | 1.04 | 0.66 |
| 54.00 | 57.00 | 53.00 | -1.23 | 0.08 |
| 4.00 | 14.00 | 9.67 | 1.14 | 0.70 |
| 89.00 | 84.00 | 77.33 | -1.30 | 0.04 |
| 14.00 | 10.00 | 10.33 | 1.27 | 0.43 |
| 970.00 | 940.00 | 992.00 | -1.35 | 0.00 |
| 420.00 | 424.00 | 425.00 | -1.09 | 0.12 |
| 6.00 | 4.00 | 5.67 | -1.48 | 0.30 |
| 67.00 | 66.00 | 68.67 | 1.08 | 0.49 |
| 175.00 | 191.00 | 190.67 | 1.02 | 0.81 |
| 67.00 | 83.00 | 73.67 | -1.03 | 0.76 |
| 33.00 | 28.00 | 35.00 | 1.11 | 0.55 |
| 109.00 | 98.00 | 102.33 | -1.23 | 0.02 |
| 252.00 | 334.00 | 306.00 | 1.13 | 0.12 |
| 31.00 | 44.00 | 37.67 | -1.01 | 0.95 |
| 2066.00 | 2075.00 | 2055.00 | -1.15 | 0.00 |
| 657.00 | 645.00 | 694.67 | 1.13 | 0.02 |
| 428.00 | 439.00 | 469.00 | 1.16 | 0.01 |
| 2107.00 | 2378.00 | 2294.67 | 1.17 | 0.00 |
| 58.00 | 57.00 | 62.33 | 1.17 | 0.21 |
| 1.00 | 8.00 | 4.67 | -1.87 | 0.13 |
| 319.00 | 311.00 | 304.00 | -1.01 | 0.90 |
| 96.00 | 115.00 | 97.33 | 1.10 | 0.38 |
| 25.00 | 22.00 | 21.00 | 1.22 | 0.32 |
| 8.00 | 11.00 | 8.33 | 1.23 | 0.56 |
| 595.00 | 559.00 | 559.33 | -1.04 | 0.56 |
| 21.00 | 19.00 | 17.33 | -2.48 | 0.00 |
| 63.00 | 57.00 | 67.00 | 1.48 | 0.00 |
| 6.00 | 6.00 | 6.00 | -2.72 | 0.00 |
| 1136.00 | 1164.00 | 1194.67 | 1.02 | 0.67 |
| 1250.00 | 1362.00 | 1310.33 | -1.03 | 0.54 |
| 3920.00 | 4729.00 | 4194.67 | 1.03 | 0.53 |
| 3.00 | 5.00 | 3.67 | -1.01 | 1.00 |
| 314.00 | 340.00 | 339.67 | 1.06 | 0.33 |
| 436.00 | 472.00 | 482.33 | -1.04 | 0.49 |
| 489.00 | 498.00 | 492.00 | 1.01 | 0.82 |
| 80.00 | 57.00 | 71.00 | 1.03 | 0.84 |

| | | | | |
|---|---|---|---|---|
| 106.00 | 108.00 | 109.67 | 1.09 | 0.38 |
| 102.00 | 117.00 | 107.33 | 1.03 | 0.75 |
| 153.00 | 195.00 | 179.00 | -1.06 | 0.42 |
| 235.00 | 227.00 | 236.00 | -1.14 | 0.04 |
| 180.00 | 201.00 | 201.67 | -1.23 | 0.00 |
| 122.00 | 102.00 | 120.67 | -1.09 | 0.36 |
| 47.00 | 71.00 | 54.67 | -1.32 | 0.03 |
| 333.00 | 386.00 | 364.67 | -1.15 | 0.01 |
| 496.00 | 529.00 | 543.67 | 1.04 | 0.43 |
| 1137.00 | 1118.00 | 1105.67 | -1.16 | 0.00 |
| 1047.00 | 1252.00 | 1187.33 | -1.25 | 0.00 |
| 6.00 | 27.00 | 21.33 | -1.35 | 0.19 |
| 452.00 | 562.00 | 533.00 | -1.04 | 0.50 |
| 19.00 | 24.00 | 19.67 | 1.10 | 0.65 |
| 402.00 | 490.00 | 468.00 | -1.19 | 0.00 |
| 206.00 | 197.00 | 186.67 | -1.06 | 0.52 |
| 173.00 | 170.00 | 172.33 | 1.09 | 0.24 |
| 490.00 | 479.00 | 514.33 | 1.14 | 0.01 |
| 174.00 | 169.00 | 170.33 | -1.23 | 0.01 |
| 1285.00 | 1409.00 | 1391.33 | -1.50 | 0.00 |
| 231.00 | 236.00 | 259.00 | 1.51 | 0.00 |
| 1938.00 | 1894.00 | 1952.00 | -1.31 | 0.00 |
| 190.00 | 212.00 | 207.33 | -1.26 | 0.00 |
| 376.00 | 436.00 | 414.00 | 1.08 | 0.15 |
| 716.00 | 769.00 | 752.67 | 1.12 | 0.04 |
| 938.00 | 1090.00 | 1058.67 | 1.13 | 0.01 |
| 1221.00 | 1523.00 | 1441.00 | 1.38 | 0.00 |
| 82.00 | 76.00 | 78.00 | -1.12 | 0.32 |
| 283.00 | 256.00 | 276.67 | -1.13 | 0.10 |
| 786.00 | 931.00 | 868.00 | -1.20 | 0.00 |
| 0.00 | 1.00 | 0.33 | -2.48 | 0.64 |
| 600.00 | 647.00 | 645.67 | -1.08 | 0.15 |
| 5345.00 | 5367.00 | 5447.00 | -1.05 | 0.14 |
| 1661.00 | 1657.00 | 1659.33 | -1.25 | 0.00 |
| 2563.00 | 2829.00 | 2730.33 | 1.08 | 0.07 |

| FDR p-value correction |
|---|
| 0.31 |
| 0.13 |
| 0.07 |
| 0.97 |
| 0.34 |
| 0.00 |
| 1.00 |
| 1.00 |
| 0.16 |
| 0.00 |
| 0.00 |
| 0.00 |
| 0.81 |
| 0.89 |
| 0.84 |
| 0.95 |
| 0.13 |
| 0.15 |
| 0.55 |
| 0.77 |
| 1.00 |
| 0.00 |
| 0.11 |
| 1.00 |
| 0.14 |
| 0.00 |
| 0.27 |
| 0.30 |
| 0.88 |
| 0.15 |
| 0.68 |
| 1.00 |
| 1.00 |
| 0.00 |
| 0.91 |
| 1.00 |
| 0.07 |
| 0.00 |
| 0.40 |
| 1.00 |
| 0.46 |
| 0.00 |
| 0.13 |

| |
|---|
| 0.00 |
| 0.39 |
| 1.00 |
| 1.00 |
| 0.00 |
| 1.00 |
| 0.00 |
| 0.24 |
| 0.88 |
| 0.24 |
| 0.00 |
| 0.02 |
| 1.00 |
| 1.00 |
| 1.00 |
| 0.97 |
| 0.42 |
| 0.00 |
| 0.45 |
| 0.01 |
| 1.00 |
| 0.54 |
| 1.00 |
| 0.25 |
| 1.00 |
| 0.85 |
| 0.00 |
| 0.00 |
| 0.00 |
| 0.00 |
| 0.09 |
| 1.00 |
| 0.15 |
| 0.66 |
| 0.43 |
| 0.00 |
| 0.00 |
| 0.94 |
| 0.48 |
| 0.77 |
| 0.00 |
| 0.08 |
| 0.00 |
| 0.27 |
| 0.00 |
| 0.87 |
| 0.00 |
| 0.45 |

| |
|---|
| 0.74 |
| 0.07 |
| 0.77 |
| 0.00 |
| 1.00 |
| 0.64 |
| 0.00 |
| 0.95 |
| 0.53 |
| 0.39 |
| 1.00 |
| 0.00 |
| 0.00 |
| 0.00 |
| 0.02 |
| 0.73 |
| 0.11 |
| 0.68 |
| 1.00 |
| 0.52 |
| 1.00 |
| 0.93 |
| 0.00 |
| 1.00 |
| 0.00 |
| 1.00 |
| 0.16 |
| 0.15 |
| 0.14 |
| 0.00 |
| 0.64 |
| 0.00 |
| 1.00 |
| 0.52 |
| 1.00 |
| 0.00 |
| 0.94 |
| 0.03 |
| 0.05 |
| 0.03 |
| 0.04 |
| 0.56 |
| 1.00 |
| 1.00 |
| 0.00 |
| 0.26 |
| 0.62 |
| 0.40 |

| |
|---|
| 0.60 |
| 0.00 |
| 1.00 |
| 0.00 |
| 0.61 |
| 1.00 |
| 0.00 |
| 0.18 |
| 0.88 |
| 0.00 |
| 0.70 |
| 1.00 |
| 0.00 |
| 0.00 |
| 1.00 |
| 1.00 |
| 0.04 |
| 1.00 |
| 0.52 |
| 0.00 |
| 0.02 |
| 0.19 |
| 1.00 |
| 0.82 |
| 0.08 |
| 0.95 |
| 1.00 |
| 1.00 |
| 0.59 |
| 0.41 |
| 1.00 |
| 1.00 |
| 0.70 |
| 1.00 |
| 0.06 |
| 0.01 |
| 1.00 |
| 0.30 |
| 0.01 |
| 0.94 |
| 0.95 |
| 0.00 |
| 0.00 |
| 0.21 |
| 0.00 |
| 1.00 |
| 0.44 |
| 0.56 |

| |
|---|
| 1.00 |
| 0.41 |
| 0.45 |
| 0.00 |
| 0.26 |
| 0.01 |
| 0.00 |
| 0.00 |
| 0.00 |
| 0.08 |
| 0.00 |
| 1.00 |
| 0.00 |
| 0.35 |
| 0.00 |
| 1.00 |
| 0.09 |
| 0.02 |
| 0.06 |
| 1.00 |
| 1.00 |
| 0.34 |
| 0.00 |
| 0.13 |
| 0.00 |
| 1.00 |
| 0.03 |
| 0.22 |
| 0.59 |
| 0.47 |
| 0.00 |
| 0.14 |
| 0.49 |
| 0.71 |
| 0.92 |
| 0.05 |
| 0.26 |
| 1.00 |
| 1.00 |
| 1.00 |
| 0.00 |
| 1.00 |
| 0.00 |
| 1.00 |
| 1.00 |
| 0.30 |
| 0.02 |
| 0.89 |

| |
|---|
| 0.00 |
| 0.16 |
| 0.07 |
| 1.00 |
| 1.00 |
| 0.00 |
| 0.19 |
| 0.00 |
| 0.03 |
| 0.00 |
| 0.00 |
| 0.03 |
| 0.03 |
| 1.00 |
| 1.00 |
| 0.00 |
| 0.25 |
| 0.38 |
| 0.00 |
| 1.00 |
| 0.06 |
| 0.95 |
| 0.14 |
| 1.00 |
| 0.25 |
| 0.00 |
| 0.23 |
| 0.84 |
| 1.00 |
| 0.32 |
| 0.16 |
| 0.00 |
| 0.49 |
| 1.00 |
| 1.00 |
| 0.72 |
| 1.00 |
| 0.21 |
| 0.00 |
| 1.00 |
| 0.01 |
| 0.00 |
| 0.00 |
| 0.08 |
| 0.14 |
| 0.06 |
| 0.00 |
| 0.00 |

| |
|---|
| 0.89 |
| 0.32 |
| 0.04 |
| 1.00 |
| 0.00 |
| 0.00 |
| 0.49 |
| 1.00 |
| 0.25 |
| 1.00 |
| 0.00 |
| 0.72 |
| 0.88 |
| 1.00 |
| 1.00 |
| 0.78 |
| 0.59 |
| 1.00 |
| 1.00 |
| 0.04 |
| 1.00 |
| 0.13 |
| 0.08 |
| 0.61 |
| 1.00 |
| 0.34 |
| 1.00 |
| 0.86 |
| 0.79 |
| 0.55 |
| 0.00 |
| 1.00 |
| 0.23 |
| 1.00 |
| 0.80 |
| 0.16 |
| 0.31 |
| 1.00 |
| 0.00 |
| 0.75 |
| 0.41 |
| 0.77 |
| 0.00 |
| 0.00 |
| 0.00 |
| 1.00 |
| 0.00 |
| 0.00 |

| |
|---|
| 1.00 |
| 0.72 |
| 1.00 |
| 0.73 |
| 0.00 |
| 0.24 |
| 0.00 |
| 0.00 |
| 0.20 |
| 0.00 |
| 0.02 |
| 0.03 |
| 0.00 |
| 0.00 |
| 0.02 |
| 0.42 |
| 1.00 |
| 0.36 |
| 0.42 |
| 0.21 |
| 0.00 |
| 0.18 |
| 0.00 |
| 0.10 |
| 0.87 |
| 0.09 |
| 0.00 |
| 1.00 |
| 0.43 |
| 0.44 |
| 1.00 |
| 0.78 |
| 0.17 |
| 0.28 |
| 0.03 |
| 1.00 |
| 1.00 |
| 0.00 |
| 0.44 |
| 0.05 |
| 1.00 |
| 0.76 |
| 0.11 |
| 0.00 |
| 0.12 |
| 0.02 |
| 0.00 |
| 1.00 |

| |
|---|
| 0.07 |
| 0.82 |
| 0.54 |
| 0.24 |
| 0.11 |
| 0.13 |
| 0.20 |
| 1.00 |
| 0.49 |
| 1.00 |
| 1.00 |
| 0.03 |
| 0.16 |
| 0.89 |
| 1.00 |
| 0.24 |
| 0.14 |
| 0.35 |
| 0.42 |
| 0.57 |
| 1.00 |
| 0.27 |
| 0.07 |
| 0.51 |
| 0.00 |
| 0.08 |
| 1.00 |
| 0.98 |
| 1.00 |
| 0.03 |
| 0.00 |
| 0.00 |
| 0.13 |
| 0.05 |
| 0.00 |
| 0.62 |
| 0.00 |
| 0.00 |
| 0.54 |
| 1.00 |
| 1.00 |
| 0.87 |
| 0.10 |
| 0.95 |
| 0.00 |
| 0.00 |
| 1.00 |
| 1.00 |

| |
|---|
| 0.67 |
| 0.00 |
| 0.01 |
| 1.00 |
| 0.00 |
| 0.76 |
| 0.00 |
| 0.36 |
| 0.39 |
| 0.97 |
| 1.00 |
| 0.04 |
| 0.19 |
| 1.00 |
| 1.00 |
| 0.54 |
| 1.00 |
| 0.02 |
| 0.51 |
| 0.01 |
| 0.28 |
| 1.00 |
| 0.70 |
| 1.00 |
| 0.97 |
| 0.00 |
| 1.00 |
| 0.00 |
| 0.52 |
| 1.00 |
| 0.25 |
| 0.92 |
| 0.01 |
| 1.00 |
| 0.00 |
| 0.50 |
| 1.00 |
| 0.31 |
| 1.00 |
| 0.01 |
| 0.00 |
| 0.00 |
| 0.86 |
| 0.00 |
| 0.06 |
| 1.00 |
| 1.00 |
| 0.49 |

| |
|---|
| 0.38 |
| 0.00 |
| 0.01 |
| 0.15 |
| 0.04 |
| 0.09 |
| 0.00 |
| 0.16 |
| 1.00 |
| 0.50 |
| 1.00 |
| 0.00 |
| 0.07 |
| 0.00 |
| 0.00 |
| 0.05 |
| 0.00 |
| 1.00 |
| 1.00 |
| 0.27 |
| 1.00 |
| 0.64 |
| 0.01 |
| 0.52 |
| 0.42 |
| 0.60 |
| 0.04 |
| 1.00 |
| 1.00 |
| 1.00 |
| 1.00 |
| 0.66 |
| 0.00 |
| 0.00 |
| 1.00 |
| 1.00 |
| 0.02 |
| 0.05 |
| 0.00 |
| 1.00 |
| 0.25 |
| 0.12 |
| 0.54 |
| 1.00 |
| 1.00 |
| 0.13 |
| 0.64 |
| 0.16 |

| |
|---|
| 0.77 |
| 0.01 |
| 0.00 |
| 0.00 |
| 0.89 |
| 0.15 |
| 0.38 |
| 0.00 |
| 0.01 |
| 0.00 |
| 1.00 |
| 1.00 |
| 0.54 |
| 1.00 |
| 0.00 |
| 1.00 |
| 1.00 |
| 1.00 |
| 0.00 |
| 1.00 |
| 1.00 |
| 0.50 |
| 0.09 |
| 0.00 |
| 0.00 |
| 0.61 |
| 0.74 |
| 0.69 |
| 1.00 |
| 0.91 |
| 0.46 |
| 0.00 |
| 0.74 |
| 0.49 |
| 0.01 |
| 0.00 |
| 1.00 |
| 0.00 |
| 1.00 |
| 0.00 |
| 0.48 |
| 0.00 |
| 0.14 |
| 0.00 |
| 0.00 |
| 0.00 |
| 0.00 |
| 1.00 |

| |
|---|
| 0.00 |
| 0.00 |
| 0.01 |
| 0.01 |
| 0.32 |
| 0.00 |
| 0.01 |
| 1.00 |
| 0.64 |
| 0.19 |
| 0.00 |
| 1.00 |
| 0.00 |
| 1.00 |
| 0.02 |
| 0.51 |
| 0.27 |
| 0.75 |
| 0.75 |
| 1.00 |
| 0.00 |
| 0.01 |
| 0.09 |
| 0.04 |
| 0.00 |
| 1.00 |
| 0.81 |
| 0.03 |
| 0.45 |
| 0.06 |
| 0.45 |
| 0.37 |
| 0.91 |
| 0.51 |
| 0.07 |
| 1.00 |
| 0.38 |
| 1.00 |
| 0.00 |
| 1.00 |
| 0.40 |
| 0.38 |
| 1.00 |
| 1.00 |
| 0.53 |
| 1.00 |
| 1.00 |
| 0.25 |

| |
|---|
| 0.51 |
| 0.28 |
| 0.05 |
| 0.00 |
| 0.78 |
| 0.57 |
| 0.26 |
| 0.00 |
| 0.67 |
| 0.07 |
| 1.00 |
| 1.00 |
| 0.00 |
| 0.00 |
| 1.00 |
| 1.00 |
| 0.00 |
| 0.45 |
| 0.25 |
| 0.00 |
| 0.01 |
| 0.00 |
| 0.01 |
| 0.00 |
| 1.00 |
| 0.40 |
| 1.00 |
| 0.56 |
| 0.10 |
| 0.39 |
| 0.05 |
| 1.00 |
| 1.00 |
| 0.01 |
| 0.52 |
| 0.09 |
| 0.02 |
| 0.76 |
| 0.04 |
| 1.00 |
| 0.00 |
| 0.33 |
| 0.39 |
| 0.89 |
| 1.00 |
| 1.00 |
| 0.39 |
| 1.00 |

| |
|---|
| 0.00 |
| 1.00 |
| 0.33 |
| 1.00 |
| 0.02 |
| 0.33 |
| 0.11 |
| 0.54 |
| 1.00 |
| 0.18 |
| 0.73 |
| 0.82 |
| 0.09 |
| 0.01 |
| 0.00 |
| 0.19 |
| 0.11 |
| 0.87 |
| 0.78 |
| 0.30 |
| 1.00 |
| 0.41 |
| 0.87 |
| 0.26 |
| 0.34 |
| 0.05 |
| 0.68 |
| 1.00 |
| 0.11 |
| 0.25 |
| 0.00 |
| 0.88 |
| 0.00 |
| 0.19 |
| 0.00 |
| 0.05 |
| 0.01 |
| 0.00 |
| 1.00 |
| 0.00 |
| 0.61 |
| 1.00 |
| 0.00 |
| 1.00 |
| 1.00 |
| 0.00 |
| 0.65 |
| 0.00 |

| |
|---|
| 0.00 |
| 0.12 |
| 1.00 |
| 1.00 |
| 0.36 |
| 1.00 |
| 0.16 |
| 0.71 |
| 0.00 |
| 0.00 |
| 0.11 |
| 0.14 |
| 0.00 |
| 0.03 |
| 0.00 |
| 0.00 |
| 0.00 |
| 0.00 |
| 1.00 |
| 0.20 |
| 0.03 |
| 0.00 |
| 1.00 |
| 1.00 |
| 0.01 |
| 0.00 |
| 0.00 |
| 0.97 |
| 0.36 |
| 0.00 |
| 0.30 |
| 0.93 |
| 0.18 |
| 0.00 |
| 1.00 |
| 1.00 |
| 0.00 |
| 0.00 |
| 0.78 |
| 0.31 |
| 1.00 |
| 0.02 |
| 0.00 |
| 0.13 |
| 0.14 |
| 0.88 |
| 0.32 |
| 0.89 |

| |
|---|
| 1.00 |
| 0.27 |
| 0.77 |
| 0.66 |
| 0.09 |
| 0.15 |
| 0.10 |
| 0.00 |
| 1.00 |
| 1.00 |
| 0.24 |
| 0.52 |
| 0.00 |
| 0.26 |
| 1.00 |
| 0.00 |
| 1.00 |
| 0.05 |
| 0.11 |
| 1.00 |
| 0.43 |
| 0.00 |
| 0.06 |
| 0.40 |
| 0.14 |
| 1.00 |
| 0.12 |
| 1.00 |
| 0.27 |
| 0.57 |
| 0.04 |
| 1.00 |
| 0.14 |
| 0.21 |
| 0.00 |
| 0.10 |
| 0.82 |
| 0.01 |
| 0.00 |
| 0.00 |
| 0.00 |
| 0.00 |
| 0.00 |
| 0.00 |
| 0.00 |
| 0.00 |
| 0.65 |
| 1.00 |

| |
|---|
| 0.18 |
| 0.51 |
| 0.00 |
| 1.00 |
| 0.77 |
| 1.00 |
| 0.01 |
| 1.00 |
| 1.00 |
| 0.00 |
| 1.00 |
| 0.00 |
| 0.27 |
| 0.00 |
| 0.00 |
| 0.30 |
| 0.70 |
| 0.29 |
| 0.71 |
| 0.00 |
| 1.00 |
| 0.34 |
| 0.00 |
| 1.00 |
| 0.14 |
| 0.00 |
| 0.00 |
| 1.00 |
| 0.42 |
| 1.00 |
| 0.91 |
| 1.00 |
| 1.00 |
| 0.00 |
| 1.00 |
| 0.74 |
| 1.00 |
| 0.01 |
| 1.00 |
| 0.00 |
| 0.00 |
| 1.00 |
| 0.17 |
| 0.26 |
| 1.00 |
| 0.00 |
| 0.68 |
| 1.00 |

| |
|---|
| 0.70 |
| 0.25 |
| 0.01 |
| 0.84 |
| 0.20 |
| 1.00 |
| 0.73 |
| 0.65 |
| 1.00 |
| 0.13 |
| 0.11 |
| 0.83 |
| 0.99 |
| 0.04 |
| 0.00 |
| 0.25 |
| 0.73 |
| 0.37 |
| 0.00 |
| 1.00 |
| 0.00 |
| 0.52 |
| 0.13 |
| 1.00 |
| 0.21 |
| 0.17 |
| 1.00 |
| 1.00 |
| 0.10 |
| 0.42 |
| 0.02 |
| 1.00 |
| 0.00 |
| 0.70 |
| 1.00 |
| 0.18 |
| 0.47 |
| 0.15 |
| 0.00 |
| 0.93 |
| 0.84 |
| 0.07 |
| 0.00 |
| 0.04 |
| 0.00 |
| 0.00 |
| 0.19 |
| 0.46 |

| |
|---|
| 0.17 |
| 0.00 |
| 0.00 |
| 0.07 |
| 0.36 |
| 0.06 |
| 0.01 |
| 1.00 |
| 0.15 |
| 0.94 |
| 0.00 |
| 0.23 |
| 0.00 |
| 0.00 |
| 0.11 |
| 0.75 |
| 0.00 |
| 0.00 |
| 0.04 |
| 0.66 |
| 0.01 |
| 0.00 |
| 1.00 |
| 1.00 |
| 0.00 |
| 1.00 |
| 0.00 |
| 0.00 |
| 0.54 |
| 0.15 |
| 0.01 |
| 0.40 |
| 0.00 |
| 0.00 |
| 0.00 |
| 1.00 |
| 0.36 |
| 0.00 |
| 0.59 |
| 0.00 |
| 0.01 |
| 0.25 |
| 0.00 |
| 0.05 |
| 0.04 |
| 0.00 |
| 0.00 |
| 0.35 |

| |
|---|
| 0.00 |
| 0.00 |
| 0.00 |
| 0.00 |
| 0.25 |
| 1.00 |
| 0.59 |
| 0.99 |
| 0.11 |
| 0.00 |
| 0.39 |
| 0.72 |
| 0.67 |
| 0.36 |
| 0.74 |
| 0.84 |
| 0.57 |
| 0.15 |
| 0.00 |
| 0.68 |
| 1.00 |
| 0.00 |
| 1.00 |
| 0.01 |
| 0.52 |
| 0.01 |
| 0.04 |
| 0.00 |
| 0.53 |
| 0.00 |
| 0.00 |
| 0.49 |
| 0.83 |
| 0.00 |
| 0.00 |
| 0.04 |
| 0.84 |
| 0.03 |
| 0.36 |
| 0.02 |
| 0.15 |
| 0.17 |
| 0.41 |
| 0.63 |
| 0.69 |
| 0.01 |
| 1.00 |
| 0.17 |

| |
|---|
| 0.00 |
| 0.23 |
| 0.00 |
| 1.00 |
| 0.00 |
| 0.90 |
| 1.00 |
| 0.13 |
| 0.15 |
| 0.02 |
| 1.00 |
| 1.00 |
| 0.28 |
| 0.84 |
| 0.00 |
| 0.01 |
| 1.00 |
| 0.00 |
| 1.00 |
| 0.81 |
| 0.89 |
| 0.04 |
| 1.00 |
| 1.00 |
| 0.00 |
| 1.00 |
| 0.57 |
| 1.00 |
| 0.00 |
| 0.70 |
| 0.42 |
| 0.00 |
| 0.73 |
| 1.00 |
| 0.00 |
| 0.13 |
| 0.00 |
| 0.94 |
| 0.26 |
| 0.84 |
| 0.46 |
| 0.84 |
| 0.97 |
| 0.21 |
| 1.00 |
| 0.21 |
| 0.64 |
| 0.00 |

| |
|---|
| 0.00 |
| 1.00 |
| 0.01 |
| 0.54 |
| 0.78 |
| 0.90 |
| 0.19 |
| 1.00 |
| 0.89 |
| 0.00 |
| 0.15 |
| 0.00 |
| 1.00 |
| 0.01 |
| 1.00 |
| 1.00 |
| 0.08 |
| 0.00 |
| 0.14 |
| 0.02 |
| 0.48 |
| 0.16 |
| 0.17 |
| 1.00 |
| 0.11 |
| 0.00 |
| 0.01 |
| 0.14 |
| 0.70 |
| 1.00 |
| 0.14 |
| 1.00 |
| 0.98 |
| 0.18 |
| 1.00 |
| 0.00 |
| 0.72 |
| 0.39 |
| 0.01 |
| 0.01 |
| 1.00 |
| 0.05 |
| 0.08 |
| 0.24 |
| 1.00 |
| 0.01 |
| 0.57 |
| 1.00 |

| |
|---|
| 1.00 |
| 1.00 |
| 0.80 |
| 0.00 |
| 0.01 |
| 0.75 |
| 0.25 |
| 0.04 |
| 0.31 |
| 1.00 |
| 0.00 |
| 0.36 |
| 0.39 |
| 1.00 |
| 0.93 |
| 0.88 |
| 0.00 |
| 0.20 |
| 0.93 |
| 0.00 |
| 1.00 |
| 0.54 |
| 0.04 |
| 0.03 |
| 1.00 |
| 1.00 |
| 1.00 |
| 0.97 |
| 1.00 |
| 1.00 |
| 0.01 |
| 1.00 |
| 0.16 |
| 0.61 |
| 0.02 |
| 0.10 |
| 0.00 |
| 0.00 |
| 0.05 |
| 0.02 |
| 0.01 |
| 0.66 |
| 0.52 |
| 1.00 |
| 0.41 |
| 1.00 |
| 0.77 |
| 0.00 |

| |
|---|
| 1.00 |
| 0.00 |
| 1.00 |
| 0.00 |
| 0.23 |
| 0.02 |
| 0.00 |
| 0.01 |
| 0.64 |
| 1.00 |
| 1.00 |
| 0.12 |
| 0.53 |
| 1.00 |
| 0.60 |
| 0.25 |
| 0.00 |
| 0.05 |
| 1.00 |
| 0.10 |
| 1.00 |
| 0.18 |
| 0.41 |
| 0.00 |
| 0.01 |
| 0.00 |
| 1.00 |
| 0.05 |
| 0.09 |
| 0.66 |
| 0.35 |
| 0.64 |
| 1.00 |
| 1.00 |
| 0.23 |
| 0.00 |
| 1.00 |
| 0.10 |
| 1.00 |
| 0.63 |
| 0.08 |
| 0.00 |
| 1.00 |
| 0.23 |
| 0.03 |
| 1.00 |
| 1.00 |
| 1.00 |

| |
|---|
| 0.00 |
| 0.00 |
| 0.00 |
| 0.13 |
| 0.78 |
| 0.00 |
| 0.63 |
| 1.00 |
| 0.41 |
| 0.03 |
| 0.45 |
| 0.73 |
| 0.01 |
| 0.41 |
| 0.82 |
| 0.02 |
| 0.00 |
| 1.00 |
| 1.00 |
| 0.00 |
| 1.00 |
| 0.00 |
| 0.84 |
| 0.07 |
| 1.00 |
| 1.00 |
| 0.00 |
| 0.00 |
| 0.28 |
| 0.02 |
| 0.01 |
| 0.77 |
| 0.90 |
| 0.08 |
| 0.00 |
| 0.00 |
| 0.37 |
| 0.46 |
| 0.00 |
| 0.00 |
| 0.00 |
| 0.00 |
| 0.88 |
| 0.39 |
| 0.00 |
| 0.01 |
| 0.05 |
| 0.10 |

| |
|---|
| 1.00 |
| 0.07 |
| 1.00 |
| 0.00 |
| 0.00 |
| 0.07 |
| 0.63 |
| 0.01 |
| 0.01 |
| 0.85 |
| 0.39 |
| 1.00 |
| 1.00 |
| 0.64 |
| 1.00 |
| 0.09 |
| 0.30 |
| 0.57 |
| 0.05 |
| 0.28 |
| 0.00 |
| 1.00 |
| 1.00 |
| 1.00 |
| 0.00 |
| 0.14 |
| 0.39 |
| 0.07 |
| 0.18 |
| 0.00 |
| 0.34 |
| 0.00 |
| 0.00 |
| 1.00 |
| 0.18 |
| 0.03 |
| 0.94 |
| 0.20 |
| 0.00 |
| 0.01 |
| 1.00 |
| 0.00 |
| 1.00 |
| 0.25 |
| 0.07 |
| 0.00 |
| 0.19 |
| 1.00 |

| |
|---|
| 1.00 |
| 0.00 |
| 0.00 |
| 0.02 |
| 0.29 |
| 0.00 |
| 0.00 |
| 0.65 |
| 0.06 |
| 0.00 |
| 0.11 |
| 0.19 |
| 1.00 |
| 0.01 |
| 0.00 |
| 0.00 |
| 0.00 |
| 0.00 |
| 0.68 |
| 1.00 |
| 0.00 |
| 1.00 |
| 1.00 |
| 0.01 |
| 0.00 |
| 1.00 |
| 0.00 |
| 0.00 |
| 1.00 |
| 1.00 |
| 0.03 |
| 0.08 |
| 1.00 |
| 1.00 |
| 0.46 |
| 0.35 |
| 0.00 |
| 0.00 |
| 0.08 |
| 1.00 |
| 0.01 |
| 0.15 |
| 1.00 |
| 0.49 |
| 0.95 |
| 0.64 |
| 1.00 |
| 1.00 |

| |
|---|
| 0.00 |
| 1.00 |
| 1.00 |
| 0.51 |
| 1.00 |
| 0.09 |
| 0.15 |
| 0.48 |
| 1.00 |
| 0.84 |
| 0.34 |
| 0.00 |
| 1.00 |
| 1.00 |
| 0.07 |
| 0.00 |
| 0.69 |
| 1.00 |
| 0.03 |
| 1.00 |
| 1.00 |
| 1.00 |
| 0.43 |
| 0.00 |
| 1.00 |
| 0.00 |
| 1.00 |
| 0.20 |
| 1.00 |
| 0.02 |
| 0.97 |
| 0.39 |
| 1.00 |
| 1.00 |
| 1.00 |
| 0.57 |
| 1.00 |
| 0.00 |
| 1.00 |
| 1.00 |
| 0.36 |
| 0.23 |
| 0.82 |
| 0.01 |
| 1.00 |
| 0.00 |
| 0.77 |
| 1.00 |

| |
|---|
| 1.00 |
| 1.00 |
| 0.05 |
| 0.95 |
| 0.61 |
| 0.41 |
| 1.00 |
| 0.07 |
| 1.00 |
| 0.00 |
| 0.26 |
| 0.42 |
| 1.00 |
| 0.00 |
| 0.74 |
| 0.02 |
| 0.95 |
| 0.26 |
| 0.14 |
| 0.09 |
| 0.00 |
| 1.00 |
| 0.26 |
| 0.00 |
| 1.00 |
| 0.00 |
| 1.00 |
| 1.00 |
| 1.00 |
| 0.71 |
| 0.02 |
| 0.59 |
| 0.00 |
| 0.96 |
| 0.00 |
| 0.89 |
| 0.56 |
| 0.40 |
| 0.00 |
| 1.00 |
| 0.90 |
| 0.00 |
| 1.00 |
| 0.51 |
| 0.03 |
| 0.00 |
| 0.29 |
| 1.00 |

| |
|---|
| 1.00 |
| 0.02 |
| 0.02 |
| 0.80 |
| 0.57 |
| 0.01 |
| 0.54 |
| 0.12 |
| 0.01 |
| 0.51 |
| 0.09 |
| 0.05 |
| 1.00 |
| 0.00 |
| 0.01 |
| 1.00 |
| 0.85 |
| 0.93 |
| 0.41 |
| 0.00 |
| 0.21 |
| 0.54 |
| 1.00 |
| 0.03 |
| 0.37 |
| 0.00 |
| 0.00 |
| 1.00 |
| 1.00 |
| 0.30 |
| 0.99 |
| 1.00 |
| 0.13 |
| 1.00 |
| 0.57 |
| 0.03 |
| 0.33 |
| 0.04 |
| 0.25 |
| 0.84 |
| 0.27 |
| 1.00 |
| 1.00 |
| 0.00 |
| 1.00 |
| 0.41 |
| 0.51 |
| 1.00 |

| |
|---|
| 1.00 |
| 0.34 |
| 0.00 |
| 0.02 |
| 0.00 |
| 1.00 |
| 0.27 |
| 0.48 |
| 1.00 |
| 0.14 |
| 0.00 |
| 0.29 |
| 0.15 |
| 1.00 |
| 0.03 |
| 0.00 |
| 0.01 |
| 0.04 |
| 0.16 |
| 0.06 |
| 1.00 |
| 0.04 |
| 0.86 |
| 1.00 |
| 0.05 |
| 1.00 |
| 0.01 |
| 0.00 |
| 0.00 |
| 0.57 |
| 0.68 |
| 0.02 |
| 1.00 |
| 1.00 |
| 0.25 |
| 1.00 |
| 1.00 |
| 0.08 |
| 0.05 |
| 0.31 |
| 1.00 |
| 0.93 |
| 0.16 |
| 1.00 |
| 0.04 |
| 0.16 |
| 0.99 |
| 0.57 |

| |
|---|
| 1.00 |
| 0.00 |
| 0.00 |
| 1.00 |
| 0.38 |
| 0.04 |
| 0.57 |
| 1.00 |
| 1.00 |
| 0.00 |
| 0.70 |
| 0.72 |
| 0.31 |
| 0.11 |
| 0.39 |
| 0.00 |
| 1.00 |
| 0.03 |
| 1.00 |
| 0.67 |
| 0.00 |
| 1.00 |
| 1.00 |
| 0.01 |
| 1.00 |
| 0.04 |
| 0.00 |
| 0.01 |
| 0.01 |
| 1.00 |
| 0.01 |
| 0.12 |
| 0.00 |
| 0.17 |
| 0.00 |
| 0.01 |
| 1.00 |
| 0.05 |
| 0.23 |
| 0.52 |
| 1.00 |
| 1.00 |
| 0.00 |
| 0.00 |
| 1.00 |
| 0.28 |
| 1.00 |
| 1.00 |

| |
|---|
| 0.00 |
| 0.82 |
| 0.00 |
| 0.14 |
| 0.17 |
| 1.00 |
| 0.02 |
| 1.00 |
| 0.42 |
| 0.82 |
| 0.97 |
| 0.00 |
| 0.89 |
| 0.31 |
| 1.00 |
| 0.29 |
| 0.00 |
| 1.00 |
| 0.07 |
| 0.61 |
| 0.73 |
| 0.53 |
| 1.00 |
| 0.81 |
| 0.01 |
| 0.48 |
| 1.00 |
| 0.35 |
| 0.20 |
| 1.00 |
| 1.00 |
| 0.13 |
| 0.73 |
| 0.00 |
| 0.18 |
| 1.00 |
| 1.00 |
| 0.51 |
| 1.00 |
| 0.00 |
| 0.37 |
| 1.00 |
| 1.00 |
| 0.53 |
| 0.01 |
| 0.24 |
| 0.52 |
| 0.71 |

| |
|---|
| 0.06 |
| 1.00 |
| 0.40 |
| 0.02 |
| 0.58 |
| 0.40 |
| 0.42 |
| 1.00 |
| 1.00 |
| 0.33 |
| 1.00 |
| 1.00 |
| 0.40 |
| 0.00 |
| 0.32 |
| 1.00 |
| 1.00 |
| 0.68 |
| 0.80 |
| 0.00 |
| 0.02 |
| 0.12 |
| 0.02 |
| 0.44 |
| 0.00 |
| 0.00 |
| 0.46 |
| 1.00 |
| 0.02 |
| 0.00 |
| 0.01 |
| 0.41 |
| 1.00 |
| 0.25 |
| 1.00 |
| 0.00 |
| 0.61 |
| 0.25 |
| 0.00 |
| 1.00 |
| 0.00 |
| 0.04 |
| 0.00 |
| 0.00 |
| 0.51 |
| 0.25 |
| 0.67 |
| 0.55 |

| |
|---|
| 1.00 |
| 0.00 |
| 0.96 |
| 0.57 |
| 1.00 |
| 1.00 |
| 0.37 |
| 0.50 |
| 0.11 |
| 0.85 |
| 1.00 |
| 1.00 |
| 0.56 |
| 1.00 |
| 0.98 |
| 0.00 |
| 1.00 |
| 0.40 |
| 0.00 |
| 0.59 |
| 0.02 |
| 0.04 |
| 0.40 |
| 0.10 |
| 0.01 |
| 0.23 |
| 0.26 |
| 0.01 |
| 0.42 |
| 0.00 |
| 0.01 |
| 0.02 |
| 1.00 |
| 0.06 |
| 1.00 |
| 0.00 |
| 1.00 |
| 0.00 |
| 0.00 |
| 0.70 |
| 0.50 |
| 0.01 |
| 0.00 |
| 1.00 |
| 0.10 |
| 0.90 |
| 0.00 |
| 0.42 |

| |
|---|
| 0.52 |
| 0.04 |
| 1.00 |
| 1.00 |
| 0.15 |
| 1.00 |
| 0.61 |
| 1.00 |
| 1.00 |
| 1.00 |
| 0.41 |
| 0.25 |
| 0.74 |
| 1.00 |
| 1.00 |
| 1.00 |
| 0.56 |
| 0.61 |
| 0.00 |
| 1.00 |
| 1.00 |
| 1.00 |
| 0.32 |
| 1.00 |
| 0.92 |
| 1.00 |
| 0.00 |
| 0.40 |
| 0.51 |
| 0.68 |
| 1.00 |
| 1.00 |
| 1.00 |
| 1.00 |
| 0.00 |
| 0.00 |
| 0.00 |
| 0.00 |
| 0.00 |
| 0.06 |
| 0.61 |
| 0.26 |
| 0.77 |
| 0.09 |
| 0.00 |
| 1.00 |
| 0.03 |
| 1.00 |

| |
|---|
| 1.00 |
| 1.00 |
| 1.00 |
| 1.00 |
| 1.00 |
| 0.01 |
| 0.16 |
| 0.00 |
| 1.00 |
| 1.00 |
| 0.01 |
| 0.16 |
| 0.32 |
| 0.23 |
| 0.00 |
| 0.12 |
| 0.00 |
| 0.47 |
| 1.00 |
| 1.00 |
| 0.00 |
| 1.00 |
| 0.11 |
| 1.00 |
| 0.69 |
| 0.67 |
| 0.15 |
| 1.00 |
| 0.78 |
| 0.00 |
| 0.38 |
| 0.00 |
| 0.81 |
| 0.51 |
| 0.93 |
| 0.15 |
| 0.02 |
| 1.00 |
| 0.25 |
| 0.33 |
| 0.07 |
| 0.83 |
| 0.63 |
| 1.00 |
| 0.00 |
| 1.00 |
| 0.00 |
| 0.06 |

| |
|---|
| 0.04 |
| 0.65 |
| 0.00 |
| 0.07 |
| 0.02 |
| 0.00 |
| 0.04 |
| 0.00 |
| 0.00 |
| 0.84 |
| 0.61 |
| 0.01 |
| 0.01 |
| 0.72 |
| 0.57 |
| 0.28 |
| 0.00 |
| 0.83 |
| 1.00 |
| 1.00 |
| 0.02 |
| 1.00 |
| 0.25 |
| 1.00 |
| 0.46 |
| 1.00 |
| 1.00 |
| 0.12 |
| 0.00 |
| 0.02 |
| 1.00 |
| 1.00 |
| 1.00 |
| 0.34 |
| 1.00 |
| 0.61 |
| 0.81 |
| 0.00 |
| 1.00 |
| 1.00 |
| 0.41 |
| 0.01 |
| 0.06 |
| 1.00 |
| 0.00 |
| 0.89 |
| 0.12 |
| 0.00 |

| |
|---|
| 0.00 |
| 0.01 |
| 0.03 |
| 0.02 |
| 1.00 |
| 0.24 |
| 0.06 |
| 0.30 |
| 0.23 |
| 1.00 |
| 0.00 |
| 0.00 |
| 0.32 |
| 0.00 |
| 0.85 |
| 1.00 |
| 0.00 |
| 0.10 |
| 0.51 |
| 1.00 |
| 0.00 |
| 1.00 |
| 1.00 |
| 0.32 |
| 0.38 |
| 0.00 |
| 0.06 |
| 0.02 |
| 0.14 |
| 0.25 |
| 1.00 |
| 0.00 |
| 0.06 |
| 0.23 |
| 0.84 |
| 0.28 |
| 0.01 |
| 1.00 |
| 0.00 |
| 0.50 |
| 0.26 |
| 0.35 |
| 0.00 |
| 0.00 |
| 0.94 |
| 0.00 |
| 1.00 |
| 0.00 |

| |
|---|
| 0.83 |
| 0.01 |
| 0.95 |
| 0.31 |
| 0.00 |
| 0.01 |
| 1.00 |
| 1.00 |
| 0.00 |
| 0.25 |
| 0.95 |
| 0.66 |
| 1.00 |
| 1.00 |
| 0.00 |
| 1.00 |
| 1.00 |
| 1.00 |
| 0.14 |
| 0.03 |
| 0.00 |
| 1.00 |
| 0.01 |
| 0.00 |
| 0.05 |
| 0.77 |
| 0.01 |
| 0.48 |
| 0.06 |
| 1.00 |
| 0.36 |
| 0.04 |
| 0.00 |
| 0.00 |
| 0.00 |
| 0.01 |
| 0.26 |
| 0.13 |
| 0.83 |
| 0.00 |
| 0.00 |
| 0.01 |
| 0.03 |
| 1.00 |
| 0.70 |
| 0.30 |
| 0.10 |
| 0.15 |

| |
|---|
| 0.00 |
| 0.00 |
| 1.00 |
| 0.12 |
| 1.00 |
| 0.28 |
| 0.11 |
| 0.00 |
| 0.00 |
| 1.00 |
| 0.22 |
| 0.54 |
| 0.40 |
| 0.00 |
| 1.00 |
| 1.00 |
| 1.00 |
| 0.00 |
| 0.01 |
| 0.00 |
| 0.16 |
| 0.31 |
| 0.00 |
| 0.00 |
| 0.14 |
| 1.00 |
| 0.00 |
| 0.40 |
| 0.21 |
| 0.00 |
| 0.00 |
| 0.04 |
| 0.89 |
| 0.35 |
| 0.66 |
| 1.00 |
| 0.00 |
| 1.00 |
| 0.00 |
| 0.00 |
| 1.00 |
| 1.00 |
| 0.91 |
| 1.00 |
| 0.00 |
| 0.14 |
| 0.74 |
| 0.00 |

| |
|---|
| 0.82 |
| 0.01 |
| 1.00 |
| 0.53 |
| 0.52 |
| 0.13 |
| 1.00 |
| 1.00 |
| 0.01 |
| 0.21 |
| 0.71 |
| 0.00 |
| 0.18 |
| 0.00 |
| 0.84 |
| 0.42 |
| 0.40 |
| 0.05 |
| 0.00 |
| 0.36 |
| 0.00 |
| 0.00 |
| 0.05 |
| 0.09 |
| 0.00 |
| 0.00 |
| 0.00 |
| 0.00 |
| 0.06 |
| 0.00 |
| 0.15 |
| 1.00 |
| 0.39 |
| 0.00 |
| 1.00 |
| 1.00 |
| 0.00 |
| 0.00 |
| 0.28 |
| 0.00 |
| 0.00 |
| 0.00 |
| 0.24 |
| 0.00 |
| 0.24 |
| 0.26 |
| 1.00 |
| 0.59 |

| |
|---|
| 0.05 |
| 0.15 |
| 0.01 |
| 0.79 |
| 0.08 |
| 1.00 |
| 1.00 |
| 0.00 |
| 0.57 |
| 0.02 |
| 1.00 |
| 0.00 |
| 0.70 |
| 0.62 |
| 0.68 |
| 0.55 |
| 1.00 |
| 0.60 |
| 0.22 |
| 0.18 |
| 0.04 |
| 0.69 |
| 0.73 |
| 0.07 |
| 0.37 |
| 0.77 |
| 1.00 |
| 0.13 |
| 0.00 |
| 0.02 |
| 0.81 |
| 0.40 |
| 0.01 |
| 0.00 |
| 0.02 |
| 0.01 |
| 0.59 |
| 0.00 |
| 0.22 |
| 1.00 |
| 1.00 |
| 0.79 |
| 0.17 |
| 0.19 |
| 1.00 |
| 0.57 |
| 0.00 |
| 0.00 |

| |
|---|
| 0.33 |
| 1.00 |
| 0.82 |
| 0.00 |
| 0.00 |
| 1.00 |
| 0.91 |
| 0.34 |
| 0.41 |
| 0.00 |
| 0.05 |
| 1.00 |
| 0.77 |
| 0.69 |
| 0.42 |
| 0.54 |
| 0.14 |
| 0.00 |
| 0.00 |
| 0.45 |
| 0.01 |
| 0.11 |
| 1.00 |
| 0.27 |
| 1.00 |
| 0.21 |
| 0.37 |
| 0.32 |
| 0.00 |
| 0.02 |
| 0.10 |
| 0.01 |
| 0.02 |
| 0.00 |
| 0.00 |
| 0.22 |
| 0.07 |
| 0.01 |
| 1.00 |
| 0.00 |
| 0.06 |
| 0.03 |
| 0.33 |
| 0.01 |
| 0.24 |
| 1.00 |
| 1.00 |
| 1.00 |

| |
|---|
| 0.09 |
| 0.00 |
| 0.15 |
| 1.00 |
| 1.00 |
| 0.00 |
| 0.89 |
| 1.00 |
| 0.48 |
| 0.00 |
| 0.93 |
| 0.12 |
| 0.45 |
| 0.33 |
| 1.00 |
| 0.00 |
| 0.64 |
| 0.65 |
| 0.29 |
| 1.00 |
| 1.00 |
| 1.00 |
| 0.11 |
| 0.75 |
| 0.26 |
| 1.00 |
| 0.00 |
| 0.38 |
| 0.09 |
| 0.16 |
| 1.00 |
| 0.62 |
| 1.00 |
| 0.02 |
| 1.00 |
| 1.00 |
| 1.00 |
| 0.00 |
| 0.94 |
| 0.15 |
| 0.00 |
| 0.95 |
| 0.00 |
| 1.00 |
| 1.00 |
| 0.00 |
| 0.00 |
| 0.07 |

| |
|---|
| 0.00 |
| 0.26 |
| 0.10 |
| 0.01 |
| 0.00 |
| 1.00 |
| 0.18 |
| 0.00 |
| 0.80 |
| 0.00 |
| 1.00 |
| 0.08 |
| 0.12 |
| 1.00 |
| 1.00 |
| 1.00 |
| 1.00 |
| 1.00 |
| 0.85 |
| 1.00 |
| 0.95 |
| 1.00 |
| 1.00 |
| 1.00 |
| 1.00 |
| 0.24 |
| 0.00 |
| 1.00 |
| 0.33 |
| 0.01 |
| 0.00 |
| 0.01 |
| 0.23 |
| 0.16 |
| 0.54 |
| 0.82 |
| 0.45 |
| 0.00 |
| 1.00 |
| 0.00 |
| 1.00 |
| 0.00 |
| 1.00 |
| 0.00 |
| 0.09 |
| 0.00 |
| 0.00 |
| 0.96 |

| |
|---|
| 0.52 |
| 0.00 |
| 1.00 |
| 1.00 |
| 1.00 |
| 0.98 |
| 1.00 |
| 0.83 |
| 1.00 |
| 0.62 |
| 0.16 |
| 0.25 |
| 0.97 |
| 1.00 |
| 1.00 |
| 0.04 |
| 0.47 |
| 0.15 |
| 0.30 |
| 0.76 |
| 0.00 |
| 0.54 |
| 0.02 |
| 0.00 |
| 1.00 |
| 0.12 |
| 0.02 |
| 0.00 |
| 0.00 |
| 0.02 |
| 0.02 |
| 1.00 |
| 1.00 |
| 1.00 |
| 0.00 |
| 0.03 |
| 0.11 |
| 0.00 |
| 0.34 |
| 1.00 |
| 0.16 |
| 0.20 |
| 0.77 |
| 0.00 |
| 0.01 |
| 0.01 |
| 0.92 |
| 0.82 |

| |
|---|
| 0.08 |
| 1.00 |
| 1.00 |
| 1.00 |
| 0.58 |
| 0.09 |
| 0.02 |
| 0.64 |
| 1.00 |
| 0.42 |
| 0.00 |
| 1.00 |
| 0.00 |
| 0.72 |
| 0.71 |
| 0.00 |
| 0.45 |
| 1.00 |
| 0.01 |
| 0.07 |
| 1.00 |
| 1.00 |
| 0.35 |
| 0.63 |
| 0.05 |
| 0.00 |
| 1.00 |
| 0.46 |
| 1.00 |
| 0.00 |
| 0.00 |
| 0.15 |
| 0.00 |
| 0.09 |
| 0.00 |
| 0.83 |
| 0.04 |
| 0.11 |
| 1.00 |
| 0.00 |
| 0.00 |
| 0.17 |
| 1.00 |
| 0.34 |
| 0.00 |
| 0.03 |
| 0.00 |
| 0.04 |

| |
|---|
| 0.11 |
| 0.00 |
| 0.84 |
| 0.00 |
| 0.01 |
| 0.00 |
| 0.03 |
| 0.06 |
| 0.73 |
| 0.00 |
| 0.13 |
| 0.63 |
| 0.26 |
| 1.00 |
| 0.44 |
| 0.77 |
| 1.00 |
| 0.01 |
| 0.57 |
| 0.02 |
| 0.07 |
| 0.46 |
| 1.00 |
| 0.00 |
| 0.68 |
| 0.00 |
| 1.00 |
| 1.00 |
| 0.00 |
| 0.68 |
| 0.97 |
| 1.00 |
| 0.00 |
| 1.00 |
| 0.46 |
| 0.00 |
| 0.00 |
| 1.00 |
| 1.00 |
| 0.38 |
| 1.00 |
| 0.57 |
| 0.60 |
| 0.00 |
| 1.00 |
| 0.10 |
| 0.64 |
| 0.21 |

| |
|---|
| 0.09 |
| 0.74 |
| 1.00 |
| 1.00 |
| 0.01 |
| 0.27 |
| 0.77 |
| 0.80 |
| 0.00 |
| 0.00 |
| 0.89 |
| 0.00 |
| 0.03 |
| 0.53 |
| 1.00 |
| 0.00 |
| 0.01 |
| 0.03 |
| 0.24 |
| 1.00 |
| 0.83 |
| 1.00 |
| 1.00 |
| 0.02 |
| 0.02 |
| 0.02 |
| 0.00 |
| 1.00 |
| 0.00 |
| 0.05 |
| 0.00 |
| 1.00 |
| 0.04 |
| 0.04 |
| 0.51 |
| 0.84 |
| 1.00 |
| 0.00 |
| 1.00 |
| 0.29 |
| 0.02 |
| 1.00 |
| 0.43 |
| 0.01 |
| 0.62 |
| 0.00 |
| 1.00 |
| 0.00 |

| |
|---|
| 0.00 |
| 0.08 |
| 0.59 |
| 0.79 |
| 0.35 |
| 0.67 |
| 0.00 |
| 0.39 |
| 0.53 |
| 0.02 |
| 0.41 |
| 0.35 |
| 0.07 |
| 0.35 |
| 0.94 |
| 0.05 |
| 0.22 |
| 0.25 |
| 1.00 |
| 0.21 |
| 0.94 |
| 0.99 |
| 0.06 |
| 0.59 |
| 0.37 |
| 0.38 |
| 0.03 |
| 0.00 |
| 0.00 |
| 1.00 |
| 1.00 |
| 0.05 |
| 0.00 |
| 1.00 |
| 0.00 |
| 0.00 |
| 0.01 |
| 0.00 |
| 1.00 |
| 1.00 |
| 0.01 |
| 0.26 |
| 1.00 |
| 0.03 |
| 0.98 |
| 0.11 |
| 0.70 |
| 1.00 |

| |
|---|
| 0.00 |
| 1.00 |
| 0.01 |
| 1.00 |
| 0.51 |
| 1.00 |
| 1.00 |
| 0.31 |
| 0.19 |
| 0.05 |
| 0.61 |
| 0.00 |
| 0.00 |
| 1.00 |
| 0.01 |
| 0.64 |
| 0.78 |
| 0.00 |
| 1.00 |
| 0.00 |
| 0.88 |
| 0.68 |
| 0.17 |
| 0.07 |
| 0.00 |
| 0.00 |
| 0.63 |
| 1.00 |
| 0.82 |
| 0.00 |
| 0.00 |
| 0.00 |
| 0.23 |
| 0.69 |
| 0.94 |
| 0.00 |
| 0.00 |
| 0.01 |
| 1.00 |
| 0.49 |
| 0.88 |
| 0.68 |
| 0.27 |
| 0.01 |
| 1.00 |
| 1.00 |
| 1.00 |
| 0.00 |

| |
|---|
| 0.18 |
| 1.00 |
| 0.08 |
| 0.72 |
| 0.01 |
| 1.00 |
| 0.16 |
| 0.34 |
| 0.01 |
| 0.81 |
| 1.00 |
| 0.64 |
| 0.41 |
| 0.16 |
| 0.00 |
| 0.00 |
| 0.00 |
| 0.89 |
| 0.05 |
| 0.00 |
| 1.00 |
| 1.00 |
| 1.00 |
| 0.37 |
| 0.53 |
| 0.74 |
| 0.07 |
| 0.01 |
| 0.00 |
| 1.00 |
| 0.08 |
| 1.00 |
| 0.94 |
| 0.94 |
| 1.00 |
| 1.00 |
| 0.97 |
| 0.05 |
| 0.62 |
| 0.00 |
| 1.00 |
| 0.00 |
| 0.55 |
| 0.68 |
| 1.00 |
| 0.72 |
| 0.30 |
| 0.12 |

| |
|---|
| 0.01 |
| 0.91 |
| 1.00 |
| 0.54 |
| 0.00 |
| 0.00 |
| 0.00 |
| 0.00 |
| 0.21 |
| 0.89 |
| 0.00 |
| 0.05 |
| 0.16 |
| 0.00 |
| 0.05 |
| 0.01 |
| 0.00 |
| 0.00 |
| 0.94 |
| 1.00 |
| 0.00 |
| 1.00 |
| 0.69 |
| 1.00 |
| 1.00 |
| 0.26 |
| 0.21 |
| 0.11 |
| 0.00 |
| 1.00 |
| 0.00 |
| 1.00 |
| 0.00 |
| 1.00 |
| 0.01 |
| 0.33 |
| 0.00 |
| 0.00 |
| 0.21 |
| 0.06 |
| 1.00 |
| 0.92 |
| 0.06 |
| 0.01 |
| 1.00 |
| 0.06 |
| 0.54 |
| 0.55 |

| |
|---|
| 0.03 |
| 0.01 |
| 0.29 |
| 1.00 |
| 0.00 |
| 0.04 |
| 0.00 |
| 1.00 |
| 0.77 |
| 0.05 |
| 0.00 |
| 0.27 |
| 0.02 |
| 1.00 |
| 0.68 |
| 0.12 |
| 0.42 |
| 0.00 |
| 0.55 |
| 1.00 |
| 0.00 |
| 0.02 |
| 0.23 |
| 0.52 |
| 0.74 |
| 0.01 |
| 0.01 |
| 0.00 |
| 1.00 |
| 0.39 |
| 1.00 |
| 0.44 |
| 0.00 |
| 1.00 |
| 0.63 |
| 0.00 |
| 1.00 |
| 1.00 |
| 0.00 |
| 0.88 |
| 1.00 |
| 0.02 |
| 0.01 |
| 1.00 |
| 0.79 |
| 0.00 |
| 0.00 |
| 0.50 |

| |
|---|
| 0.06 |
| 0.27 |
| 0.29 |
| 0.45 |
| 0.00 |
| 0.00 |
| 0.13 |
| 1.00 |
| 0.91 |
| 0.63 |
| 1.00 |
| 1.00 |
| 0.44 |
| 0.23 |
| 0.00 |
| 1.00 |
| 1.00 |
| 0.03 |
| 1.00 |
| 0.53 |
| 0.00 |
| 1.00 |
| 0.61 |
| 0.79 |
| 0.00 |
| 0.00 |
| 0.96 |
| 0.22 |
| 0.00 |
| 1.00 |
| 0.40 |
| 0.00 |
| 1.00 |
| 0.00 |
| 0.00 |
| 0.00 |
| 1.00 |
| 0.00 |
| 1.00 |
| 0.00 |
| 0.08 |
| 1.00 |
| 1.00 |
| 0.00 |
| 0.49 |
| 1.00 |
| 0.00 |
| 0.15 |

| |
|---|
| 0.00 |
| 0.00 |
| 0.88 |
| 0.03 |
| 1.00 |
| 0.35 |
| 0.12 |
| 0.03 |
| 0.05 |
| 0.19 |
| 0.18 |
| 1.00 |
| 0.44 |
| 1.00 |
| 1.00 |
| 1.00 |
| 0.83 |
| 0.97 |
| 1.00 |
| 1.00 |
| 1.00 |
| 0.10 |
| 0.28 |
| 1.00 |
| 1.00 |
| 0.43 |
| 1.00 |
| 0.35 |
| 0.13 |
| 0.02 |
| 0.02 |
| 1.00 |
| 0.00 |
| 0.02 |
| 0.61 |
| 0.02 |
| 0.68 |
| 0.00 |
| 0.00 |
| 1.00 |
| 0.09 |
| 0.08 |
| 0.02 |
| 1.00 |
| 0.00 |
| 0.11 |
| 1.00 |
| 0.96 |

| |
|---|
| 0.03 |
| 0.41 |
| 0.90 |
| 0.00 |
| 0.02 |
| 0.01 |
| 0.00 |
| 0.00 |
| 0.42 |
| 0.21 |
| 1.00 |
| 0.00 |
| 0.03 |
| 0.03 |
| 0.23 |
| 0.61 |
| 0.20 |
| 0.37 |
| 0.00 |
| 0.08 |
| 0.30 |
| 0.49 |
| 0.84 |
| 0.30 |
| 0.07 |
| 0.69 |
| 1.00 |
| 0.33 |
| 0.01 |
| 0.00 |
| 0.82 |
| 0.48 |
| 0.00 |
| 0.17 |
| 1.00 |
| 0.00 |
| 0.09 |
| 0.57 |
| 0.00 |
| 0.00 |
| 0.00 |
| 0.00 |
| 0.00 |
| 0.27 |
| 0.54 |
| 1.00 |
| 1.00 |
| 0.04 |

| |
|---|
| 0.91 |
| 1.00 |
| 0.47 |
| 1.00 |
| 1.00 |
| 0.00 |
| 0.00 |
| 1.00 |
| 0.01 |
| 0.05 |
| 0.10 |
| 0.45 |
| 0.68 |
| 1.00 |
| 0.02 |
| 0.04 |
| 0.02 |
| 0.32 |
| 0.49 |
| 0.00 |
| 0.00 |
| 0.00 |
| 0.67 |
| 0.00 |
| 0.00 |
| 0.19 |
| 0.45 |
| 0.22 |
| 0.03 |
| 0.01 |
| 0.44 |
| 0.08 |
| 1.00 |
| 0.00 |
| 0.64 |
| 0.01 |
| 0.58 |
| 0.89 |
| 0.61 |
| 0.88 |
| 0.02 |
| 1.00 |
| 0.57 |
| 1.00 |
| 1.00 |
| 0.00 |
| 0.00 |
| 0.24 |

| |
|---|
| 1.00 |
| 0.00 |
| 1.00 |
| 0.00 |
| 0.36 |
| 1.00 |
| 1.00 |
| 0.12 |
| 0.48 |
| 0.43 |
| 0.03 |
| 0.00 |
| 0.00 |
| 1.00 |
| 0.91 |
| 0.56 |
| 0.00 |
| 1.00 |
| 0.12 |
| 0.08 |
| 0.00 |
| 0.70 |
| 0.00 |
| 1.00 |
| 0.00 |
| 0.10 |
| 0.00 |
| 1.00 |
| 0.00 |
| 0.00 |
| 1.00 |
| 0.00 |
| 0.51 |
| 1.00 |
| 0.00 |
| 1.00 |
| 0.20 |
| 0.96 |
| 0.29 |
| 0.60 |
| 0.07 |
| 0.00 |
| 1.00 |
| 0.13 |
| 0.30 |
| 0.32 |
| 1.00 |
| 0.64 |

| |
|---|
| 0.00 |
| 0.01 |
| 1.00 |
| 0.00 |
| 1.00 |
| 0.44 |
| 0.18 |
| 0.14 |
| 0.83 |
| 1.00 |
| 0.00 |
| 0.75 |
| 0.00 |
| 1.00 |
| 0.00 |
| 0.00 |
| 0.09 |
| 0.14 |
| 0.82 |
| 0.18 |
| 1.00 |
| 0.07 |
| 1.00 |
| 0.30 |
| 0.08 |
| 1.00 |
| 0.18 |
| 0.98 |
| 0.00 |
| 0.02 |
| 0.90 |
| 0.01 |
| 0.03 |
| 1.00 |
| 1.00 |
| 0.00 |
| 1.00 |
| 0.02 |
| 0.00 |
| 0.31 |
| 0.00 |
| 0.00 |
| 0.99 |
| 0.21 |
| 0.06 |
| 0.18 |
| 0.31 |
| 0.00 |

| |
|---|
| 1.00 |
| 0.53 |
| 0.02 |
| 0.62 |
| 0.04 |
| 0.01 |
| 0.00 |
| 0.07 |
| 1.00 |
| 1.00 |
| 0.42 |
| 1.00 |
| 0.78 |
| 0.00 |
| 1.00 |
| 0.00 |
| 0.36 |
| 0.12 |
| 1.00 |
| 1.00 |
| 0.16 |
| 0.33 |
| 0.51 |
| 0.52 |
| 0.00 |
| 1.00 |
| 1.00 |
| 1.00 |
| 0.99 |
| 1.00 |
| 0.12 |
| 0.13 |
| 0.00 |
| 0.28 |
| 0.00 |
| 0.06 |
| 1.00 |
| 1.00 |
| 0.83 |
| 0.54 |
| 0.92 |
| 0.00 |
| 0.74 |
| 0.54 |
| 0.07 |
| 0.03 |
| 1.00 |
| 0.05 |

| |
|---|
| 1.00 |
| 0.83 |
| 0.03 |
| 0.33 |
| 0.00 |
| 0.87 |
| 0.41 |
| 0.89 |
| 0.00 |
| 1.00 |
| 1.00 |
| 0.26 |
| 0.00 |
| 0.11 |
| 0.00 |
| 0.09 |
| 1.00 |
| 1.00 |
| 0.00 |
| 0.00 |
| 1.00 |
| 0.00 |
| 1.00 |
| 1.00 |
| 0.11 |
| 0.65 |
| 0.00 |
| 0.68 |
| 1.00 |
| 0.03 |
| 0.00 |
| 1.00 |
| 1.00 |
| 0.01 |
| 1.00 |
| 0.37 |
| 0.27 |
| 0.62 |
| 0.21 |
| 0.69 |
| 1.00 |
| 0.08 |
| 0.14 |
| 0.09 |
| 1.00 |
| 0.08 |
| 0.00 |
| 1.00 |

| |
|---|
| 0.00 |
| 0.05 |
| 0.42 |
| 0.00 |
| 1.00 |
| 0.00 |
| 0.00 |
| 0.04 |
| 0.01 |
| 1.00 |
| 0.16 |
| 0.02 |
| 0.00 |
| 1.00 |
| 0.56 |
| 0.13 |
| 0.00 |
| 1.00 |
| 0.06 |
| 0.13 |
| 1.00 |
| 0.00 |
| 0.00 |
| 0.25 |
| 0.40 |
| 0.04 |
| 0.60 |
| 1.00 |
| 1.00 |
| 0.94 |
| 0.00 |
| 0.64 |
| 0.83 |
| 0.47 |
| 0.58 |
| 1.00 |
| 0.42 |
| 0.98 |
| 0.50 |
| 0.01 |
| 0.13 |
| 1.00 |
| 0.40 |
| 1.00 |
| 0.03 |
| 0.91 |
| 0.18 |
| 0.00 |

| |
|---|
| 1.00 |
| 0.03 |
| 0.85 |
| 0.00 |
| 0.86 |
| 0.30 |
| 0.07 |
| 1.00 |
| 1.00 |
| 1.00 |
| 0.39 |
| 1.00 |
| 0.03 |
| 0.66 |
| 0.79 |
| 0.00 |
| 0.12 |
| 0.03 |
| 0.00 |
| 0.74 |
| 0.36 |
| 1.00 |
| 0.98 |
| 0.70 |
| 0.00 |
| 0.07 |
| 1.00 |
| 0.02 |
| 1.00 |
| 0.90 |
| 0.82 |
| 0.85 |
| 0.08 |
| 0.40 |
| 0.09 |
| 0.35 |
| 1.00 |
| 0.50 |
| 1.00 |
| 0.00 |
| 0.06 |
| 0.00 |
| 0.60 |
| 0.00 |
| 0.72 |
| 1.00 |
| 0.82 |
| 0.09 |

| |
|---|
| 0.00 |
| 0.12 |
| 0.74 |
| 0.11 |
| 1.00 |
| 0.35 |
| 0.00 |
| 0.00 |
| 0.56 |
| 0.21 |
| 0.90 |
| 0.02 |
| 0.09 |
| 1.00 |
| 0.00 |
| 1.00 |
| 1.00 |
| 1.00 |
| 0.00 |
| 0.00 |
| 0.41 |
| 1.00 |
| 0.26 |
| 0.03 |
| 0.92 |
| 1.00 |
| 0.02 |
| 0.20 |
| 1.00 |
| 0.22 |
| 1.00 |
| 0.00 |
| 0.54 |
| 0.74 |
| 1.00 |
| 0.00 |
| 0.41 |
| 0.00 |
| 1.00 |
| 0.11 |
| 1.00 |
| 0.22 |
| 0.43 |
| 1.00 |
| 0.29 |
| 0.09 |
| 0.00 |
| 0.64 |

| |
|---|
| 1.00 |
| 0.13 |
| 0.19 |
| 0.01 |
| 0.89 |
| 0.11 |
| 0.36 |
| 0.21 |
| 0.89 |
| 0.77 |
| 0.86 |
| 0.45 |
| 1.00 |
| 0.17 |
| 1.00 |
| 0.11 |
| 0.02 |
| 1.00 |
| 1.00 |
| 0.16 |
| 1.00 |
| 0.20 |
| 0.87 |
| 0.18 |
| 0.03 |
| 0.33 |
| 1.00 |
| 0.00 |
| 0.83 |
| 0.00 |
| 0.04 |
| 0.91 |
| 0.24 |
| 0.54 |
| 0.00 |
| 0.00 |
| 0.00 |
| 0.00 |
| 0.01 |
| 0.89 |
| 0.00 |
| 0.04 |
| 0.80 |
| 1.00 |
| 0.36 |
| 1.00 |
| 0.00 |
| 0.00 |

| |
|---|
| 0.00 |
| 1.00 |
| 0.41 |
| 0.00 |
| 0.75 |
| 0.00 |
| 1.00 |
| 0.70 |
| 1.00 |
| 1.00 |
| 0.00 |
| 1.00 |
| 1.00 |
| 0.03 |
| 0.28 |
| 1.00 |
| 1.00 |
| 0.00 |
| 0.76 |
| 0.19 |
| 0.00 |
| 0.01 |
| 0.85 |
| 1.00 |
| 0.15 |
| 1.00 |
| 1.00 |
| 0.15 |
| 0.25 |
| 1.00 |
| 1.00 |
| 0.09 |
| 0.00 |
| 0.46 |
| 0.19 |
| 1.00 |
| 0.45 |
| 0.13 |
| 0.09 |
| 1.00 |
| 1.00 |
| 0.00 |
| 0.12 |
| 1.00 |
| 0.00 |
| 1.00 |
| 1.00 |
| 1.00 |

| |
|---|
| 0.01 |
| 0.74 |
| 0.00 |
| 0.08 |
| 0.86 |
| 0.86 |
| 0.00 |
| 1.00 |
| 0.03 |
| 1.00 |
| 0.72 |
| 0.05 |
| 0.00 |
| 0.03 |
| 0.66 |
| 0.46 |
| 0.19 |
| 1.00 |
| 0.10 |
| 0.05 |
| 1.00 |
| 1.00 |
| 0.00 |
| 1.00 |
| 1.00 |
| 1.00 |
| 0.76 |
| 1.00 |
| 0.82 |
| 0.00 |
| 0.01 |
| 0.74 |
| 0.00 |
| 0.07 |
| 0.52 |
| 0.00 |
| 1.00 |
| 0.04 |
| 0.01 |
| 1.00 |
| 0.54 |
| 0.20 |
| 1.00 |
| 0.00 |
| 0.00 |
| 0.00 |
| 0.40 |
| 1.00 |

| |
|---|
| 0.32 |
| 0.17 |
| 0.00 |
| 0.00 |
| 0.03 |
| 0.24 |
| 1.00 |
| 0.00 |
| 0.07 |
| 0.52 |
| 1.00 |
| 0.78 |
| 1.00 |
| 1.00 |
| 0.00 |
| 0.00 |
| 0.01 |
| 0.54 |
| 0.00 |
| 0.36 |
| 0.01 |
| 1.00 |
| 0.10 |
| 1.00 |
| 1.00 |
| 0.00 |
| 0.48 |
| 0.73 |
| 1.00 |
| 0.43 |
| 1.00 |
| 0.16 |
| 0.02 |
| 0.59 |
| 0.04 |
| 0.50 |
| 0.37 |
| 0.02 |
| 0.72 |
| 0.84 |
| 1.00 |
| 1.00 |
| 0.61 |
| 0.01 |
| 0.23 |
| 0.97 |
| 1.00 |
| 0.21 |

| |
|---|
| 0.14 |
| 0.61 |
| 1.00 |
| 0.00 |
| 0.00 |
| 0.00 |
| 0.04 |
| 1.00 |
| 0.01 |
| 0.27 |
| 0.00 |
| 0.00 |
| 0.23 |
| 1.00 |
| 0.01 |
| 0.15 |
| 0.12 |
| 0.03 |
| 0.10 |
| 0.80 |
| 0.18 |
| 1.00 |
| 1.00 |
| 0.66 |
| 0.89 |
| 0.00 |
| 1.00 |
| 0.00 |
| 0.62 |
| 0.00 |
| 0.11 |
| 0.00 |
| 0.00 |
| 0.89 |
| 0.11 |
| 0.00 |
| 0.01 |
| 0.07 |
| 0.00 |
| 0.00 |
| 0.23 |
| 0.00 |
| 0.84 |
| 1.00 |
| 0.00 |
| 0.18 |
| 0.10 |
| 1.00 |

| |
|---|
| 1.00 |
| 1.00 |
| 0.03 |
| 0.00 |
| 0.00 |
| 0.05 |
| 0.79 |
| 1.00 |
| 0.04 |
| 1.00 |
| 0.15 |
| 0.00 |
| 0.39 |
| 0.00 |
| 1.00 |
| 0.74 |
| 0.00 |
| 0.00 |
| 1.00 |
| 0.01 |
| 0.47 |
| 0.92 |
| 0.00 |
| 1.00 |
| 0.91 |
| 0.02 |
| 1.00 |
| 0.56 |
| 0.71 |
| 0.24 |
| 0.00 |
| 0.40 |
| 0.21 |
| 1.00 |
| 0.00 |
| 1.00 |
| 1.00 |
| 1.00 |
| 1.00 |
| 0.00 |
| 0.03 |
| 1.00 |
| 0.73 |
| 0.65 |
| 0.74 |
| 0.28 |
| 0.01 |
| 1.00 |

| |
|---|
| 1.00 |
| 0.00 |
| 0.01 |
| 0.13 |
| 0.03 |
| 1.00 |
| 0.55 |
| 1.00 |
| 0.00 |
| 0.00 |
| 1.00 |
| 0.00 |
| 1.00 |
| 0.30 |
| 0.43 |
| 0.00 |
| 1.00 |
| 1.00 |
| 0.56 |
| 0.80 |
| 0.07 |
| 0.00 |
| 0.00 |
| 0.07 |
| 0.83 |
| 0.10 |
| 1.00 |
| 1.00 |
| 0.20 |
| 0.52 |
| 0.07 |
| 0.93 |
| 0.00 |
| 0.09 |
| 0.17 |
| 1.00 |
| 0.00 |
| 0.00 |
| 1.00 |
| 1.00 |
| 0.01 |
| 0.00 |
| 0.00 |
| 0.00 |
| 0.01 |
| 0.00 |
| 0.13 |
| 0.04 |

| |
|---|
| 1.00 |
| 0.00 |
| 0.09 |
| 0.00 |
| 1.00 |
| 0.02 |
| 0.51 |
| 0.02 |
| 0.19 |
| 1.00 |
| 0.00 |
| 0.16 |
| 0.07 |
| 0.90 |
| 0.89 |
| 0.85 |
| 0.01 |
| 0.12 |
| 1.00 |
| 0.00 |
| 1.00 |
| 0.03 |
| 0.00 |
| 0.17 |
| 0.79 |
| 1.00 |
| 0.00 |
| 0.40 |
| 0.04 |
| 1.00 |
| 0.02 |
| 1.00 |
| 0.03 |
| 0.00 |
| 0.56 |
| 0.00 |
| 0.00 |
| 1.00 |
| 0.01 |
| 1.00 |
| 1.00 |
| 0.58 |
| 0.65 |
| 0.51 |
| 0.30 |
| 0.15 |
| 0.86 |
| 0.36 |

| |
|---|
| 0.00 |
| 0.02 |
| 0.74 |
| 0.00 |
| 0.40 |
| 1.00 |
| 0.70 |
| 0.01 |
| 0.58 |
| 0.02 |
| 0.25 |
| 1.00 |
| 0.01 |
| 0.33 |
| 0.01 |
| 0.32 |
| 0.94 |
| 0.00 |
| 0.13 |
| 0.01 |
| 0.25 |
| 0.67 |
| 0.23 |
| 1.00 |
| 0.00 |
| 0.00 |
| 0.00 |
| 0.16 |
| 0.36 |
| 0.02 |
| 0.00 |
| 0.03 |
| 1.00 |
| 1.00 |
| 0.00 |
| 0.00 |
| 0.00 |
| 0.00 |
| 1.00 |
| 0.57 |
| 0.01 |
| 0.86 |
| 0.82 |
| 0.00 |
| 0.00 |
| 0.01 |
| 0.07 |
| 0.41 |

| |
|---|
| 0.37 |
| 0.68 |
| 1.00 |
| 0.89 |
| 1.00 |
| 0.01 |
| 0.43 |
| 0.00 |
| 1.00 |
| 0.30 |
| 1.00 |
| 0.00 |
| 1.00 |
| 1.00 |
| 0.00 |
| 0.06 |
| 1.00 |
| 0.00 |
| 0.25 |
| 1.00 |
| 1.00 |
| 0.06 |
| 0.41 |
| 0.37 |
| 0.11 |
| 0.00 |
| 1.00 |
| 0.00 |
| 0.00 |
| 0.00 |
| 0.03 |
| 0.91 |
| 0.01 |
| 0.00 |
| 1.00 |
| 0.23 |
| 0.66 |
| 0.63 |
| 0.28 |
| 1.00 |
| 1.00 |
| 0.41 |
| 0.26 |
| 1.00 |
| 0.84 |
| 0.00 |
| 0.00 |
| 0.02 |

| |
|---|
| 0.00 |
| 0.57 |
| 0.02 |
| 0.00 |
| 1.00 |
| 0.02 |
| 0.00 |
| 1.00 |
| 0.04 |
| 0.16 |
| 0.00 |
| 0.83 |
| 0.00 |
| 0.17 |
| 0.13 |
| 0.84 |
| 0.15 |
| 1.00 |
| 0.75 |
| 1.00 |
| 1.00 |
| 0.33 |
| 1.00 |
| 0.52 |
| 0.33 |
| 0.66 |
| 0.22 |
| 1.00 |
| 0.03 |
| 0.04 |
| 0.00 |
| 1.00 |
| 0.50 |
| 1.00 |
| 1.00 |
| 0.00 |
| 0.59 |
| 0.95 |
| 0.00 |
| 0.64 |
| 0.67 |
| 0.03 |
| 0.81 |
| 0.00 |
| 0.05 |
| 0.05 |
| 1.00 |
| 0.00 |

| |
|---|
| 0.74 |
| 0.03 |
| 0.55 |
| 0.57 |
| 1.00 |
| 1.00 |
| 0.00 |
| 0.01 |
| 0.33 |
| 0.36 |
| 0.00 |
| 0.00 |
| 0.21 |
| 0.03 |
| 0.65 |
| 0.06 |
| 0.63 |
| 0.03 |
| 0.60 |
| 0.00 |
| 0.96 |
| 0.02 |
| 0.13 |
| 0.07 |
| 0.00 |
| 0.00 |
| 0.49 |
| 0.31 |
| 1.00 |
| 0.11 |
| 0.08 |
| 0.40 |
| 0.37 |
| 0.00 |
| 0.79 |
| 0.94 |
| 0.30 |
| 0.00 |
| 0.23 |
| 1.00 |
| 1.00 |
| 0.47 |
| 1.00 |
| 1.00 |
| 0.91 |
| 0.04 |
| 1.00 |
| 0.33 |

| |
|---|
| 0.00 |
| 0.04 |
| 0.01 |
| 0.00 |
| 0.96 |
| 1.00 |
| 0.00 |
| 0.84 |
| 0.92 |
| 1.00 |
| 0.73 |
| 0.75 |
| 0.16 |
| 1.00 |
| 0.00 |
| 0.00 |
| 0.00 |
| 1.00 |
| 0.84 |
| 0.21 |
| 0.01 |
| 1.00 |
| 0.70 |
| 0.16 |
| 0.53 |
| 0.03 |
| 0.00 |
| 0.03 |
| 1.00 |
| 0.62 |
| 0.14 |
| 0.21 |
| 0.03 |
| 1.00 |
| 0.70 |
| 0.12 |
| 0.04 |
| 0.00 |
| 0.17 |
| 0.11 |
| 0.00 |
| 0.00 |
| 0.35 |
| 0.00 |
| 1.00 |
| 1.00 |
| 0.49 |
| 0.05 |

| |
|---|
| 0.16 |
| 0.03 |
| 0.00 |
| 0.00 |
| 1.00 |
| 0.01 |
| 1.00 |
| 0.00 |
| 0.00 |
| 1.00 |
| 0.00 |
| 0.88 |
| 1.00 |
| 0.36 |
| 0.42 |
| 0.59 |
| 0.95 |
| 0.31 |
| 0.00 |
| 0.00 |
| 0.00 |
| 0.93 |
| 0.42 |
| 0.00 |
| 0.00 |
| 0.95 |
| 1.00 |
| 1.00 |
| 0.08 |
| 0.11 |
| 1.00 |
| 1.00 |
| 0.78 |
| 0.00 |
| 0.00 |
| 0.11 |
| 0.75 |
| 1.00 |
| 0.11 |
| 0.00 |
| 0.29 |
| 0.30 |
| 0.00 |
| 0.90 |
| 0.00 |
| 0.00 |
| 0.70 |
| 0.04 |

| |
|---|
| 0.00 |
| 0.00 |
| 0.03 |
| 0.01 |
| 1.00 |
| 0.24 |
| 1.00 |
| 1.00 |
| 1.00 |
| 0.39 |
| 0.82 |
| 0.27 |
| 0.70 |
| 0.28 |
| 0.82 |
| 0.00 |
| 0.86 |
| 0.18 |
| 0.65 |
| 0.32 |
| 0.00 |
| 0.00 |
| 1.00 |
| 1.00 |
| 1.00 |
| 0.74 |
| 0.56 |
| 1.00 |
| 0.00 |
| 0.75 |
| 0.35 |
| 1.00 |
| 1.00 |
| 1.00 |
| 0.26 |
| 1.00 |
| 0.00 |
| 0.26 |
| 0.52 |
| 1.00 |
| 0.11 |
| 0.02 |
| 0.82 |
| 1.00 |
| 1.00 |
| 1.00 |
| 0.85 |
| 0.51 |

| |
|---|
| 0.00 |
| 1.00 |
| 0.00 |
| 0.34 |
| 0.72 |
| 0.00 |
| 1.00 |
| 0.82 |
| 0.04 |
| 1.00 |
| 1.00 |
| 0.63 |
| 0.01 |
| 0.81 |
| 1.00 |
| 1.00 |
| 0.32 |
| 1.00 |
| 0.00 |
| 0.01 |
| 1.00 |
| 0.83 |
| 0.02 |
| 0.50 |
| 0.01 |
| 0.28 |
| 0.45 |
| 1.00 |
| 0.92 |
| 1.00 |
| 0.00 |
| 1.00 |
| 1.00 |
| 1.00 |
| 0.68 |
| 0.40 |
| 0.62 |
| 0.18 |
| 0.00 |
| 0.73 |
| 1.00 |
| 0.66 |
| 1.00 |
| 0.00 |
| 0.89 |
| 0.04 |
| 0.30 |
| 0.08 |

| |
|---|
| 0.86 |
| 0.03 |
| 0.93 |
| 0.00 |
| 0.09 |
| 0.00 |
| 1.00 |
| 0.00 |
| 0.04 |
| 0.99 |
| 0.00 |
| 0.00 |
| 0.01 |
| 0.12 |
| 1.00 |
| 0.94 |
| 0.00 |
| 0.00 |
| 0.00 |
| 0.45 |
| 1.00 |
| 0.75 |
| 1.00 |
| 0.93 |
| 0.35 |
| 0.67 |
| 0.10 |
| 1.00 |
| 1.00 |
| 1.00 |
| 1.00 |
| 0.13 |
| 0.01 |
| 0.00 |
| 1.00 |
| 0.18 |
| 0.32 |
| 0.42 |
| 1.00 |
| 0.66 |
| 0.44 |
| 0.00 |
| 0.18 |
| 0.37 |
| 1.00 |
| 0.28 |
| 0.17 |
| 1.00 |

| |
|---|
| 0.63 |
| 0.78 |
| 0.00 |
| 0.62 |
| 0.00 |
| 0.02 |
| 0.82 |
| 0.02 |
| 0.32 |
| 0.03 |
| 1.00 |
| 1.00 |
| 0.65 |
| 0.01 |
| 0.00 |
| 0.26 |
| 1.00 |
| 0.98 |
| 1.00 |
| 0.09 |
| 0.05 |
| 0.07 |
| 0.15 |
| 0.00 |
| 0.01 |
| 0.24 |
| 0.77 |
| 0.00 |
| 0.72 |
| 1.00 |
| 0.97 |
| 1.00 |
| 0.00 |
| 0.77 |
| 1.00 |
| 0.00 |
| 1.00 |
| 0.00 |
| 0.00 |
| 0.02 |
| 0.74 |
| 0.29 |
| 0.00 |
| 1.00 |
| 0.25 |
| 0.00 |
| 1.00 |
| 0.54 |

| |
|---|
| 0.16 |
| 1.00 |
| 1.00 |
| 1.00 |
| 0.00 |
| 0.85 |
| 0.45 |
| 1.00 |
| 0.89 |
| 0.04 |
| 0.01 |
| 0.14 |
| 0.00 |
| 1.00 |
| 0.49 |
| 0.77 |
| 1.00 |
| 0.16 |
| 0.08 |
| 0.12 |
| 0.21 |
| 0.00 |
| 0.00 |
| 0.03 |
| 1.00 |
| 0.32 |
| 1.00 |
| 0.02 |
| 0.00 |
| 1.00 |
| 1.00 |
| 0.00 |
| 0.00 |
| 0.65 |
| 0.95 |
| 1.00 |
| 0.38 |
| 0.10 |
| 1.00 |
| 0.18 |
| 0.87 |
| 1.00 |
| 1.00 |
| 0.00 |
| 1.00 |
| 0.00 |
| 0.11 |
| 1.00 |

| |
|---|
| 0.50 |
| 1.00 |
| 0.00 |
| 0.00 |
| 1.00 |
| 0.01 |
| 0.00 |
| 0.00 |
| 0.00 |
| 0.63 |
| 1.00 |
| 0.85 |
| 0.60 |
| 0.11 |
| 1.00 |
| 0.27 |
| 0.00 |
| 0.00 |
| 0.82 |
| 0.03 |
| 0.14 |
| 1.00 |
| 1.00 |
| 0.01 |
| 0.00 |
| 1.00 |
| 0.59 |
| 0.00 |
| 0.71 |
| 0.09 |
| 1.00 |
| 1.00 |
| 1.00 |
| 0.11 |
| 1.00 |
| 0.14 |
| 0.75 |
| 0.40 |
| 0.66 |
| 1.00 |
| 0.78 |
| 0.42 |
| 0.00 |
| 0.23 |
| 1.00 |
| 0.69 |
| 0.97 |
| 0.00 |

| |
|---|
| 0.10 |
| 0.36 |
| 0.01 |
| 1.00 |
| 0.09 |
| 1.00 |
| 0.35 |
| 0.00 |
| 0.02 |
| 1.00 |
| 1.00 |
| 0.47 |
| 0.00 |
| 0.08 |
| 1.00 |
| 0.29 |
| 1.00 |
| 0.43 |
| 0.00 |
| 0.14 |
| 0.39 |
| 0.04 |
| 1.00 |
| 0.24 |
| 0.72 |
| 0.26 |
| 0.00 |
| 1.00 |
| 0.00 |
| 0.00 |
| 1.00 |
| 0.21 |
| 0.00 |
| 0.29 |
| 1.00 |
| 0.02 |
| 0.07 |
| 0.85 |
| 0.33 |
| 1.00 |
| 1.00 |
| 0.00 |
| 0.24 |
| 0.52 |
| 0.00 |
| 0.00 |
| 0.05 |
| 0.11 |

| |
|---|
| 0.00 |
| 0.73 |
| 1.00 |
| 0.00 |
| 0.90 |
| 0.89 |
| 0.11 |
| 0.00 |
| 0.00 |
| 0.01 |
| 0.00 |
| 0.46 |
| 0.21 |
| 0.14 |
| 1.00 |
| 0.33 |
| 0.00 |
| 1.00 |
| 0.00 |
| 1.00 |
| 0.00 |
| 0.48 |
| 0.01 |
| 0.00 |
| 0.01 |
| 1.00 |
| 0.16 |
| 0.05 |
| 0.00 |
| 0.00 |
| 1.00 |
| 1.00 |
| 0.00 |
| 0.01 |
| 0.25 |
| 0.13 |
| 0.00 |
| 0.00 |
| 1.00 |
| 0.45 |
| 0.12 |
| 0.00 |
| 1.00 |
| 1.00 |
| 0.43 |
| 0.02 |
| 0.00 |
| 0.00 |

| |
|---|
| 0.52 |
| 0.06 |
| 0.45 |
| 0.88 |
| 0.38 |
| 1.00 |
| 0.29 |
| 0.26 |
| 0.01 |
| 0.00 |
| 1.00 |
| 0.56 |
| 0.26 |
| 0.75 |
| 0.00 |
| 0.58 |
| 0.73 |
| 0.00 |
| 0.48 |
| 1.00 |
| 0.00 |
| 1.00 |
| 0.30 |
| 0.25 |
| 0.00 |
| 0.00 |
| 1.00 |
| 1.00 |
| 1.00 |
| 0.12 |
| 1.00 |
| 0.65 |
| 1.00 |
| 0.01 |
| 0.01 |
| 0.77 |
| 0.01 |
| 1.00 |
| 0.24 |
| 0.00 |
| 0.00 |
| 0.00 |
| 0.98 |
| 1.00 |
| 1.00 |
| 0.03 |
| 0.65 |
| 1.00 |

| |
|---|
| 0.86 |
| 0.29 |
| 0.50 |
| 1.00 |
| 0.10 |
| 0.00 |
| 0.00 |
| 0.01 |
| 0.00 |
| 1.00 |
| 1.00 |
| 0.00 |
| 1.00 |
| 0.41 |
| 0.24 |
| 0.08 |
| 0.01 |
| 0.29 |
| 0.51 |
| 0.09 |
| 1.00 |
| 0.21 |
| 0.90 |
| 1.00 |
| 1.00 |
| 0.20 |
| 1.00 |
| 0.09 |
| 0.66 |
| 0.08 |
| 0.02 |
| 0.52 |
| 1.00 |
| 0.00 |
| 1.00 |
| 0.29 |
| 0.72 |
| 0.00 |
| 0.42 |
| 1.00 |
| 0.27 |
| 0.75 |
| 0.24 |
| 1.00 |
| 0.00 |
| 0.00 |
| 0.82 |
| 0.88 |

| |
|---|
| 0.00 |
| 0.20 |
| 0.00 |
| 1.00 |
| 1.00 |
| 0.05 |
| 0.00 |
| 0.00 |
| 0.47 |
| 0.45 |
| 0.00 |
| 1.00 |
| 0.00 |
| 0.91 |
| 1.00 |
| 0.03 |
| 1.00 |
| 0.13 |
| 0.05 |
| 1.00 |
| 0.04 |
| 0.30 |
| 0.74 |
| 0.12 |
| 0.06 |
| 0.00 |
| 0.92 |
| 0.01 |
| 1.00 |
| 1.00 |
| 0.11 |
| 1.00 |
| 0.74 |
| 0.51 |
| 0.64 |
| 1.00 |
| 0.68 |
| 0.76 |
| 0.00 |
| 0.18 |
| 0.00 |
| 0.00 |
| 1.00 |
| 1.00 |
| 0.14 |
| 1.00 |
| 1.00 |
| 1.00 |

| |
|---|
| 0.54 |
| 1.00 |
| 1.00 |
| 1.00 |
| 0.00 |
| 0.54 |
| 1.00 |
| 1.00 |
| 0.00 |
| 0.00 |
| 0.60 |
| 0.00 |
| 0.17 |
| 0.52 |
| 1.00 |
| 1.00 |
| 0.00 |
| 0.42 |
| 0.16 |
| 1.00 |
| 0.00 |
| 0.39 |
| 0.94 |
| 0.37 |
| 1.00 |
| 0.46 |
| 1.00 |
| 0.18 |
| 0.01 |
| 0.68 |
| 0.01 |
| 0.00 |
| 0.01 |
| 1.00 |
| 1.00 |
| 0.73 |
| 1.00 |
| 0.03 |
| 0.58 |
| 0.00 |
| 0.07 |
| 0.00 |
| 0.03 |
| 0.00 |
| 1.00 |
| 0.10 |
| 0.25 |
| 1.00 |

| |
|---|
| 0.11 |
| 0.50 |
| 0.98 |
| 0.34 |
| 1.00 |
| 1.00 |
| 0.00 |
| 0.09 |
| 1.00 |
| 0.21 |
| 0.26 |
| 0.70 |
| 0.00 |
| 0.00 |
| 1.00 |
| 0.01 |
| 0.20 |
| 0.00 |
| 0.00 |
| 1.00 |
| 0.31 |
| 0.89 |
| 0.05 |
| 1.00 |
| 0.19 |
| 0.02 |
| 0.00 |
| 0.18 |
| 0.26 |
| 0.82 |
| 0.01 |
| 0.00 |
| 0.00 |
| 0.75 |
| 0.66 |
| 1.00 |
| 0.00 |
| 0.70 |
| 0.07 |
| 1.00 |
| 0.00 |
| 0.00 |
| 0.42 |
| 0.00 |
| 0.16 |
| 0.00 |
| 0.28 |
| 0.87 |

| |
|---|
| 1.00 |
| 0.01 |
| 0.00 |
| 0.07 |
| 0.46 |
| 1.00 |
| 0.05 |
| 0.01 |
| 0.00 |
| 0.00 |
| 0.98 |
| 0.00 |
| 0.28 |
| 0.12 |
| 0.00 |
| 1.00 |
| 1.00 |
| 0.78 |
| 0.19 |
| 0.19 |
| 0.00 |
| 1.00 |
| 0.48 |
| 0.01 |
| 0.00 |
| 0.38 |
| 0.09 |
| 1.00 |
| 0.08 |
| 0.01 |
| 0.74 |
| 0.04 |
| 0.05 |
| 1.00 |
| 0.01 |
| 0.12 |
| 0.03 |
| 0.01 |
| 0.21 |
| 1.00 |
| 1.00 |
| 0.17 |
| 0.00 |
| 0.00 |
| 1.00 |
| 0.45 |
| 1.00 |
| 0.04 |

| |
|---|
| 1.00 |
| 1.00 |
| 0.01 |
| 0.22 |
| 0.28 |
| 1.00 |
| 0.00 |
| 0.20 |
| 0.00 |
| 0.71 |
| 0.08 |
| 0.00 |
| 0.00 |
| 0.06 |
| 0.25 |
| 1.00 |
| 0.65 |
| 0.17 |
| 0.09 |
| 0.81 |
| 0.30 |
| 0.81 |
| 0.00 |
| 0.10 |
| 1.00 |
| 0.00 |
| 0.00 |
| 0.54 |
| 0.57 |
| 0.32 |
| 0.00 |
| 0.17 |
| 0.20 |
| 1.00 |
| 1.00 |
| 0.04 |
| 0.01 |
| 0.00 |
| 0.23 |
| 1.00 |
| 0.01 |
| 0.00 |
| 0.07 |
| 1.00 |
| 0.19 |
| 0.00 |
| 1.00 |
| 0.00 |

| |
|---|
| 0.42 |
| 1.00 |
| 1.00 |
| 0.66 |
| 0.07 |
| 1.00 |
| 0.79 |
| 0.73 |
| 1.00 |
| 0.78 |
| 0.82 |
| 0.00 |
| 1.00 |
| 0.64 |
| 0.59 |
| 1.00 |
| 0.00 |
| 0.00 |
| 0.73 |
| 0.00 |
| 1.00 |
| 0.00 |
| 0.03 |
| 0.01 |
| 1.00 |
| 1.00 |
| 1.00 |
| 1.00 |
| 0.25 |
| 0.00 |
| 0.00 |
| 0.00 |
| 0.80 |
| 0.12 |
| 0.15 |
| 1.00 |
| 1.00 |
| 0.14 |
| 0.13 |
| 0.98 |
| 0.04 |
| 0.02 |
| 0.23 |
| 0.21 |
| 0.02 |
| 0.00 |
| 0.00 |
| 0.00 |

| |
|---|
| 0.62 |
| 0.06 |
| 0.70 |
| 0.80 |
| 0.22 |
| 0.11 |
| 0.27 |
| 0.95 |
| 0.05 |
| 1.00 |
| 0.60 |
| 1.00 |
| 1.00 |
| 1.00 |
| 0.00 |
| 1.00 |
| 0.48 |
| 0.00 |
| 0.00 |
| 0.48 |
| 0.00 |
| 0.92 |
| 0.00 |
| 0.49 |
| 0.22 |
| 0.39 |
| 0.00 |
| 1.00 |
| 0.00 |
| 0.00 |
| 1.00 |
| 0.00 |
| 0.37 |
| 0.12 |
| 0.00 |
| 0.08 |
| 0.23 |
| 0.00 |
| 0.03 |
| 0.00 |
| 0.11 |
| 0.00 |
| 1.00 |
| 0.00 |
| 0.00 |
| 0.04 |
| 0.34 |
| 0.03 |

| |
|---|
| 0.00 |
| 0.00 |
| 1.00 |
| 1.00 |
| 0.00 |
| 1.00 |
| 0.51 |
| 0.00 |
| 0.54 |
| 0.74 |
| 0.57 |
| 0.55 |
| 0.33 |
| 1.00 |
| 0.01 |
| 0.00 |
| 0.01 |
| 0.01 |
| 0.00 |
| 0.01 |
| 0.24 |
| 1.00 |
| 0.03 |
| 0.00 |
| 1.00 |
| 1.00 |
| 0.45 |
| 1.00 |
| 0.12 |
| 1.00 |
| 0.00 |
| 0.17 |
| 0.51 |
| 1.00 |
| 1.00 |
| 0.28 |
| 0.00 |
| 1.00 |
| 1.00 |
| 0.89 |
| 0.00 |
| 0.14 |
| 0.79 |
| 0.00 |
| 0.98 |
| 0.15 |
| 0.30 |
| 0.38 |

| |
|---|
| 0.78 |
| 0.00 |
| 0.00 |
| 1.00 |
| 0.00 |
| 0.90 |
| 0.51 |
| 0.72 |
| 0.36 |
| 1.00 |
| 0.22 |
| 0.48 |
| 0.61 |
| 0.95 |
| 0.66 |
| 0.65 |
| 1.00 |
| 0.00 |
| 0.04 |
| 0.63 |
| 0.87 |
| 0.74 |
| 0.00 |
| 0.36 |
| 0.68 |
| 0.25 |
| 1.00 |
| 1.00 |
| 0.04 |
| 0.01 |
| 1.00 |
| 0.03 |
| 0.06 |
| 0.17 |
| 0.97 |
| 0.52 |
| 0.00 |
| 1.00 |
| 1.00 |
| 0.10 |
| 0.04 |
| 0.00 |
| 0.00 |
| 0.70 |
| 0.45 |
| 0.27 |
| 0.45 |
| 1.00 |

| |
|---|
| 0.01 |
| 0.01 |
| 0.02 |
| 0.31 |
| 0.74 |
| 0.00 |
| 0.07 |
| 0.07 |
| 1.00 |
| 1.00 |
| 1.00 |
| 0.00 |
| 0.84 |
| 0.03 |
| 0.00 |
| 1.00 |
| 1.00 |
| 1.00 |
| 0.43 |
| 0.82 |
| 0.98 |
| 0.04 |
| 0.15 |
| 1.00 |
| 0.87 |
| 0.00 |
| 1.00 |
| 1.00 |
| 0.00 |
| 0.63 |
| 0.32 |
| 1.00 |
| 0.01 |
| 1.00 |
| 0.64 |
| 0.00 |
| 0.00 |
| 0.62 |
| 0.30 |
| 0.03 |
| 0.00 |
| 0.10 |
| 0.27 |
| 0.83 |
| 1.00 |
| 0.06 |
| 0.00 |
| 0.00 |

| |
|---|
| 0.14 |
| 0.00 |
| 0.23 |
| 1.00 |
| 0.01 |
| 0.40 |
| 0.00 |
| 1.00 |
| 0.09 |
| 1.00 |
| 0.61 |
| 1.00 |
| 0.63 |
| 0.03 |
| 0.00 |
| 0.42 |
| 1.00 |
| 0.56 |
| 0.02 |
| 0.00 |
| 1.00 |
| 1.00 |
| 1.00 |
| 1.00 |
| 0.57 |
| 0.04 |
| 0.09 |
| 0.00 |
| 0.83 |
| 1.00 |
| 1.00 |
| 1.00 |
| 0.00 |
| 0.00 |
| 0.88 |
| 0.35 |
| 0.02 |
| 0.00 |
| 0.00 |
| 0.18 |
| 0.28 |
| 1.00 |
| 0.93 |
| 0.07 |
| 1.00 |
| 0.55 |
| 0.11 |
| 1.00 |

| |
|---|
| 1.00 |
| 0.01 |
| 1.00 |
| 0.58 |
| 0.42 |
| 0.46 |
| 0.00 |
| 0.01 |
| 0.14 |
| 0.04 |
| 0.04 |
| 0.28 |
| 0.00 |
| 0.35 |
| 0.16 |
| 0.00 |
| 0.54 |
| 0.01 |
| 0.14 |
| 0.83 |
| 0.00 |
| 0.00 |
| 0.00 |
| 1.00 |
| 0.06 |
| 0.57 |
| 0.86 |
| 0.31 |
| 0.00 |
| 0.02 |
| 0.00 |
| 1.00 |
| 1.00 |
| 0.39 |
| 0.30 |
| 0.03 |
| 0.03 |
| 1.00 |
| 0.73 |
| 0.00 |
| 0.68 |
| 0.05 |
| 0.21 |
| 1.00 |
| 1.00 |
| 0.68 |
| 0.81 |
| 0.32 |

| |
|---|
| 0.63 |
| 0.00 |
| 0.85 |
| 0.00 |
| 0.00 |
| 0.04 |
| 0.19 |
| 0.15 |
| 0.01 |
| 0.59 |
| 0.02 |
| 0.00 |
| 1.00 |
| 1.00 |
| 0.00 |
| 0.05 |
| 0.41 |
| 0.54 |
| 1.00 |
| 0.14 |
| 0.50 |
| 0.24 |
| 0.12 |
| 0.00 |
| 0.00 |
| 1.00 |
| 1.00 |
| 0.14 |
| 1.00 |
| 0.06 |
| 0.00 |
| 1.00 |
| 0.00 |
| 0.84 |
| 0.01 |
| 1.00 |
| 0.01 |
| 1.00 |
| 0.21 |
| 1.00 |
| 1.00 |
| 1.00 |
| 0.80 |
| 0.20 |
| 1.00 |
| 1.00 |
| 0.02 |
| 1.00 |

| |
|---|
| 1.00 |
| 0.00 |
| 1.00 |
| 1.00 |
| 1.00 |
| 1.00 |
| 1.00 |
| 0.00 |
| 1.00 |
| 1.00 |
| 0.72 |
| 0.55 |
| 0.64 |
| 0.64 |
| 0.91 |
| 0.44 |
| 0.11 |
| 0.01 |
| 0.01 |
| 1.00 |
| 0.00 |
| 0.00 |
| 0.00 |
| 0.33 |
| 0.00 |
| 0.45 |
| 0.30 |
| 1.00 |
| 0.04 |
| 0.00 |
| 0.66 |
| 0.00 |
| 0.00 |
| 0.03 |
| 0.00 |
| 0.27 |
| 0.17 |
| 0.00 |
| 0.39 |
| 0.05 |
| 0.64 |
| 0.81 |
| 0.44 |
| 1.00 |
| 1.00 |
| 0.20 |
| 0.10 |
| 1.00 |

| |
|---|
| 0.00 |
| 0.01 |
| 0.00 |
| 1.00 |
| 0.06 |
| 0.95 |
| 0.15 |
| 0.91 |
| 0.04 |
| 1.00 |
| 0.00 |
| 0.00 |
| 1.00 |
| 0.42 |
| 0.43 |
| 1.00 |
| 0.78 |
| 0.00 |
| 0.98 |
| 1.00 |
| 0.21 |
| 0.00 |
| 0.67 |
| 1.00 |
| 1.00 |
| 1.00 |
| 0.49 |
| 0.00 |
| 0.28 |
| 1.00 |
| 0.00 |
| 1.00 |
| 0.00 |
| 1.00 |
| 0.78 |
| 0.00 |
| 0.00 |
| 0.12 |
| 0.87 |
| 0.00 |
| 0.00 |
| 0.93 |
| 0.03 |
| 0.16 |
| 0.00 |
| 0.37 |
| 1.00 |
| 1.00 |

| |
|---|
| 0.50 |
| 1.00 |
| 0.23 |
| 1.00 |
| 0.10 |
| 0.06 |
| 0.02 |
| 0.01 |
| 0.77 |
| 1.00 |
| 0.01 |
| 1.00 |
| 0.00 |
| 0.00 |
| 0.00 |
| 0.00 |
| 0.06 |
| 1.00 |
| 0.00 |
| 1.00 |
| 0.36 |
| 0.73 |
| 0.00 |
| 0.52 |
| 0.05 |
| 0.00 |
| 0.00 |
| 0.00 |
| 0.95 |
| 1.00 |
| 1.00 |
| 0.26 |
| 0.00 |
| 1.00 |
| 1.00 |
| 0.63 |
| 0.00 |
| 0.42 |
| 0.11 |
| 0.00 |
| 0.76 |
| 0.01 |
| 0.06 |
| 0.51 |
| 0.47 |
| 0.63 |
| 1.00 |
| 1.00 |

| |
|---|
| 1.00 |
| 1.00 |
| 0.12 |
| 0.18 |
| 1.00 |
| 0.09 |
| 0.49 |
| 0.00 |
| 0.08 |
| 0.00 |
| 0.00 |
| 0.42 |
| 0.87 |
| 0.02 |
| 1.00 |
| 0.05 |
| 0.00 |
| 0.95 |
| 0.44 |
| 0.00 |
| 0.11 |
| 0.00 |
| 0.78 |
| 0.37 |
| 0.00 |
| 0.06 |
| 0.02 |
| 0.00 |
| 0.45 |
| 0.00 |
| 1.00 |
| 0.00 |
| 0.31 |
| 0.03 |
| 0.00 |
| 0.00 |
| 0.02 |
| 1.00 |
| 1.00 |
| 0.99 |
| 1.00 |
| 0.01 |
| 1.00 |
| 1.00 |
| 1.00 |
| 1.00 |
| 0.84 |
| 1.00 |

| |
|---|
| 1.00 |
| 0.56 |
| 0.65 |
| 1.00 |
| 0.62 |
| 1.00 |
| 1.00 |
| 0.56 |
| 0.20 |
| 0.40 |
| 1.00 |
| 1.00 |
| 0.61 |
| 0.00 |
| 0.01 |
| 1.00 |
| 0.00 |
| 0.21 |
| 0.00 |
| 0.00 |
| 0.85 |
| 0.72 |
| 1.00 |
| 0.00 |
| 0.06 |
| 0.00 |
| 1.00 |
| 1.00 |
| 0.97 |
| 0.02 |
| 0.10 |
| 0.37 |
| 0.39 |
| 0.00 |
| 0.00 |
| 0.10 |
| 0.00 |
| 1.00 |
| 1.00 |
| 1.00 |
| 0.06 |
| 0.00 |
| 0.71 |
| 0.00 |
| 0.00 |
| 0.00 |
| 0.82 |
| 1.00 |

| |
|---|
| 1.00 |
| 0.00 |
| 0.00 |
| 1.00 |
| 1.00 |
| 1.00 |
| 0.00 |
| 0.07 |
| 1.00 |
| 0.00 |
| 0.05 |
| 0.00 |
| 0.00 |
| 0.39 |
| 0.00 |
| 1.00 |
| 0.00 |
| 0.00 |
| 0.95 |
| 0.00 |
| 0.00 |
| 0.45 |
| 0.00 |
| 0.00 |
| 0.00 |
| 0.37 |
| 0.00 |
| 1.00 |
| 0.88 |
| 0.01 |
| 0.05 |
| 0.00 |
| 0.36 |
| 0.00 |
| 0.00 |
| 0.00 |
| 0.12 |
| 0.11 |
| 0.95 |
| 1.00 |
| 0.07 |
| 1.00 |
| 0.11 |
| 0.01 |
| 0.00 |
| 1.00 |
| 1.00 |
| 0.40 |

| |
|---|
| 0.02 |
| 0.42 |
| 0.00 |
| 0.00 |
| 1.00 |
| 0.98 |
| 0.88 |
| 0.82 |
| 0.44 |
| 1.00 |
| 0.99 |
| 0.32 |
| 0.28 |
| 0.71 |
| 0.00 |
| 0.38 |
| 0.44 |
| 1.00 |
| 0.21 |
| 0.02 |
| 1.00 |
| 1.00 |
| 0.90 |
| 0.64 |
| 0.36 |
| 0.86 |
| 0.84 |
| 1.00 |
| 0.07 |
| 0.01 |
| 0.26 |
| 1.00 |
| 0.74 |
| 0.07 |
| 1.00 |
| 1.00 |
| 1.00 |
| 1.00 |
| 0.11 |
| 1.00 |
| 0.00 |
| 0.07 |
| 0.00 |
| 0.00 |
| 1.00 |
| 1.00 |
| 0.00 |
| 0.07 |

| |
|---|
| 1.00 |
| 0.68 |
| 0.86 |
| 0.02 |
| 1.00 |
| 1.00 |
| 0.00 |
| 0.00 |
| 0.01 |
| 1.00 |
| 0.71 |
| 0.02 |
| 1.00 |
| 0.84 |
| 1.00 |
| 0.24 |
| 1.00 |
| 0.01 |
| 0.00 |
| 0.04 |
| 0.63 |
| 0.02 |
| 1.00 |
| 0.78 |
| 0.01 |
| 0.00 |
| 0.00 |
| 1.00 |
| 0.21 |
| 0.48 |
| 0.41 |
| 1.00 |
| 0.97 |
| 0.13 |
| 1.00 |
| 1.00 |
| 0.28 |
| 0.24 |
| 0.07 |
| 1.00 |
| 1.00 |
| 0.00 |
| 1.00 |
| 0.00 |
| 0.03 |
| 0.13 |
| 0.07 |
| 0.42 |

| |
|---|
| 1.00 |
| 0.14 |
| 0.56 |
| 1.00 |
| 1.00 |
| 0.00 |
| 0.91 |
| 0.00 |
| 1.00 |
| 0.00 |
| 0.00 |
| 0.07 |
| 0.24 |
| 1.00 |
| 0.01 |
| 0.06 |
| 0.00 |
| 0.00 |
| 0.38 |
| 0.07 |
| 1.00 |
| 1.00 |
| 0.26 |
| 0.33 |
| 0.99 |
| 0.17 |
| 1.00 |
| 0.57 |
| 0.00 |
| 0.33 |
| 0.46 |
| 0.00 |
| 1.00 |
| 0.61 |
| 1.00 |
| 0.06 |
| 0.00 |
| 1.00 |
| 0.21 |
| 0.00 |
| 0.97 |
| 0.49 |
| 0.00 |
| 0.12 |
| 0.53 |
| 0.41 |
| 0.05 |
| 0.10 |

| |
|---|
| 0.42 |
| 0.04 |
| 1.00 |
| 0.03 |
| 0.12 |
| 0.39 |
| 0.33 |
| 1.00 |
| 0.28 |
| 0.24 |
| 1.00 |
| 0.77 |
| 1.00 |
| 0.18 |
| 0.56 |
| 0.00 |
| 0.40 |
| 0.54 |
| 1.00 |
| 0.29 |
| 1.00 |
| 0.87 |
| 0.00 |
| 0.00 |
| 0.02 |
| 0.00 |
| 0.17 |
| 0.56 |
| 0.01 |
| 1.00 |
| 0.00 |
| 1.00 |
| 0.00 |
| 0.10 |
| 0.56 |
| 0.00 |
| 0.06 |
| 0.17 |
| 1.00 |
| 1.00 |
| 1.00 |
| 1.00 |
| 0.87 |
| 0.17 |
| 0.26 |
| 0.09 |
| 0.02 |
| 0.00 |

| |
|---|
| 0.39 |
| 1.00 |
| 0.01 |
| 0.01 |
| 1.00 |
| 0.39 |
| 0.00 |
| 0.06 |
| 0.48 |
| 1.00 |
| 1.00 |
| 0.78 |
| 0.74 |
| 0.03 |
| 0.27 |
| 0.00 |
| 1.00 |
| 0.12 |
| 1.00 |
| 1.00 |
| 0.00 |
| 0.13 |
| 1.00 |
| 0.27 |
| 0.20 |
| 0.00 |
| 0.18 |
| 0.00 |
| 0.50 |
| 0.13 |
| 0.00 |
| 1.00 |
| 0.00 |
| 1.00 |
| 0.53 |
| 1.00 |
| 0.00 |
| 0.01 |
| 0.77 |
| 0.42 |
| 0.00 |
| 1.00 |
| 0.04 |
| 1.00 |
| 0.00 |
| 0.00 |
| 1.00 |
| 0.03 |

| |
|---|
| 0.00 |
| 0.83 |
| 0.00 |
| 0.00 |
| 0.00 |
| 0.00 |
| 0.99 |
| 0.70 |
| 0.65 |
| 0.83 |
| 0.04 |
| 1.00 |
| 0.02 |
| 0.75 |
| 0.93 |
| 0.98 |
| 0.57 |
| 1.00 |
| 0.69 |
| 0.00 |
| 1.00 |
| 0.17 |
| 0.00 |
| 1.00 |
| 0.74 |
| 0.00 |
| 0.00 |
| 0.06 |
| 0.00 |
| 1.00 |
| 0.00 |
| 0.00 |
| 1.00 |
| 1.00 |
| 0.00 |
| 0.00 |
| 1.00 |
| 0.93 |
| 0.00 |
| 0.00 |
| 0.00 |
| 1.00 |
| 0.03 |
| 0.00 |
| 0.00 |
| 1.00 |
| 1.00 |
| 0.00 |

| |
|---|
| 0.72 |
| 0.74 |
| 0.41 |
| 0.94 |
| 0.00 |
| 0.00 |
| 0.68 |
| 0.11 |
| 0.11 |
| 0.00 |
| 1.00 |
| 0.24 |
| 1.00 |
| 0.42 |
| 0.00 |
| 1.00 |
| 1.00 |
| 1.00 |
| 0.45 |
| 0.01 |
| 0.00 |
| 0.52 |
| 0.15 |
| 0.29 |
| 0.02 |
| 1.00 |
| 1.00 |
| 1.00 |
| 1.00 |
| 0.85 |
| 0.00 |
| 1.00 |
| 0.12 |
| 0.00 |
| 0.00 |
| 1.00 |
| 0.05 |
| 0.60 |
| 0.57 |
| 0.00 |
| 0.95 |
| 0.26 |
| 0.01 |
| 0.03 |
| 0.09 |
| 0.41 |
| 0.00 |
| 0.00 |

| |
|---|
| 0.00 |
| 0.06 |
| 0.48 |
| 0.03 |
| 0.42 |
| 1.00 |
| 1.00 |
| 0.79 |
| 0.36 |
| 0.61 |
| 0.01 |
| 0.20 |
| 1.00 |
| 0.20 |
| 0.77 |
| 1.00 |
| 0.15 |
| 0.70 |
| 0.49 |
| 1.00 |
| 0.04 |
| 1.00 |
| 1.00 |
| 0.12 |
| 0.71 |
| 0.03 |
| 0.89 |
| 0.88 |
| 1.00 |
| 0.02 |
| 0.08 |
| 0.99 |
| 0.59 |
| 0.42 |
| 0.57 |
| 0.47 |
| 0.39 |
| 0.47 |
| 0.00 |
| 1.00 |
| 0.07 |
| 0.23 |
| 0.66 |
| 0.00 |
| 0.02 |
| 0.00 |
| 0.02 |
| 1.00 |

| |
|---|
| 1.00 |
| 1.00 |
| 0.21 |
| 0.89 |
| 0.69 |
| 1.00 |
| 0.00 |
| 0.00 |
| 0.32 |
| 1.00 |
| 0.08 |
| 1.00 |
| 0.00 |
| 1.00 |
| 1.00 |
| 0.35 |
| 1.00 |
| 0.00 |
| 0.00 |
| 0.02 |
| 0.14 |
| 1.00 |
| 0.18 |
| 1.00 |
| 0.47 |
| 0.03 |
| 0.44 |
| 0.00 |
| 1.00 |
| 1.00 |
| 0.56 |
| 0.00 |
| 1.00 |
| 0.44 |
| 1.00 |
| 1.00 |
| 0.00 |
| 1.00 |
| 1.00 |
| 1.00 |
| 0.11 |
| 0.00 |
| 0.00 |
| 0.00 |
| 0.00 |
| 1.00 |
| 0.00 |
| 0.76 |

| |
|---|
| 0.00 |
| 0.00 |
| 0.38 |
| 0.54 |
| 0.90 |
| 0.89 |
| 0.54 |
| 0.00 |
| 0.25 |
| 0.01 |
| 0.79 |
| 0.00 |
| 0.68 |
| 0.00 |
| 0.04 |
| 1.00 |
| 0.00 |
| 0.40 |
| 0.00 |
| 0.00 |
| 0.03 |
| 0.52 |
| 0.00 |
| 0.00 |
| 0.37 |
| 0.00 |
| 1.00 |
| 0.30 |
| 0.18 |
| 1.00 |
| 0.01 |
| 0.00 |
| 0.00 |
| 0.83 |
| 0.00 |
| 0.00 |
| 0.19 |
| 0.53 |
| 0.77 |
| 0.06 |
| 0.06 |
| 0.03 |
| 1.00 |
| 1.00 |
| 0.00 |
| 0.35 |
| 0.08 |
| 0.60 |

| |
|---|
| 1.00 |
| 1.00 |
| 1.00 |
| 0.02 |
| 0.52 |
| 0.98 |
| 1.00 |
| 1.00 |
| 0.33 |
| 0.47 |
| 0.41 |
| 0.56 |
| 0.76 |
| 0.79 |
| 0.00 |
| 0.45 |
| 1.00 |
| 0.40 |
| 0.63 |
| 0.00 |
| 0.54 |
| 0.00 |
| 0.00 |
| 0.00 |
| 0.01 |
| 0.04 |
| 0.00 |
| 0.00 |
| 0.65 |
| 1.00 |
| 0.51 |
| 0.31 |
| 0.04 |
| 0.00 |
| 0.00 |
| 0.00 |
| 1.00 |
| 0.00 |
| 0.02 |
| 0.16 |
| 0.09 |
| 0.01 |
| 0.81 |
| 1.00 |
| 1.00 |
| 0.06 |
| 1.00 |
| 1.00 |

| |
|---|
| 0.37 |
| 1.00 |
| 0.05 |
| 0.52 |
| 0.20 |
| 1.00 |
| 0.00 |
| 0.00 |
| 1.00 |
| 0.44 |
| 0.00 |
| 0.41 |
| 0.59 |
| 0.00 |
| 0.96 |
| 1.00 |
| 0.15 |
| 0.23 |
| 0.10 |
| 0.00 |
| 0.64 |
| 1.00 |
| 0.74 |
| 0.00 |
| 1.00 |
| 0.00 |
| 0.06 |
| 0.67 |
| 0.01 |
| 0.07 |
| 1.00 |
| 1.00 |
| 0.00 |
| 0.97 |
| 0.00 |
| 0.06 |
| 0.00 |
| 1.00 |
| 0.04 |
| 0.90 |
| 0.62 |
| 1.00 |
| 1.00 |
| 0.87 |
| 0.00 |
| 0.01 |
| 0.02 |
| 1.00 |

| |
|---|
| 0.45 |
| 0.19 |
| 1.00 |
| 0.01 |
| 0.00 |
| 1.00 |
| 0.01 |
| 0.10 |
| 0.00 |
| 1.00 |
| 0.30 |
| 0.69 |
| 0.44 |
| 1.00 |
| 1.00 |
| 1.00 |
| 1.00 |
| 0.29 |
| 1.00 |
| 1.00 |
| 1.00 |
| 1.00 |
| 0.90 |
| 0.11 |
| 0.78 |
| 1.00 |
| 0.16 |
| 0.24 |
| 0.41 |
| 1.00 |
| 1.00 |
| 0.02 |
| 0.00 |
| 1.00 |
| 0.91 |
| 1.00 |
| 0.24 |
| 0.36 |
| 0.95 |
| 0.01 |
| 1.00 |
| 0.57 |
| 1.00 |
| 1.00 |
| 0.00 |
| 1.00 |
| 0.00 |
| 0.85 |

| |
|---|
| 0.62 |
| 1.00 |
| 1.00 |
| 1.00 |
| 1.00 |
| 1.00 |
| 1.00 |
| 0.00 |
| 0.08 |
| 0.10 |
| 0.00 |
| 0.95 |
| 0.00 |
| 0.00 |
| 0.00 |
| 0.03 |
| 0.00 |
| 1.00 |
| 1.00 |
| 0.00 |
| 0.68 |
| 0.42 |
| 0.03 |
| 0.14 |
| 0.10 |
| 0.85 |
| 0.00 |
| 1.00 |
| 0.66 |
| 0.37 |
| 1.00 |
| 1.00 |
| 0.01 |
| 0.01 |
| 0.73 |
| 1.00 |
| 0.00 |
| 0.23 |
| 0.04 |
| 0.00 |
| 0.19 |
| 1.00 |
| 1.00 |
| 1.00 |
| 0.00 |
| 0.11 |
| 0.10 |
| 0.20 |

| |
|---|
| 0.01 |
| 0.98 |
| 0.78 |
| 0.00 |
| 1.00 |
| 0.00 |
| 1.00 |
| 0.00 |
| 0.00 |
| 0.10 |
| 0.13 |
| 0.05 |
| 0.00 |
| 0.31 |
| 0.11 |
| 0.09 |
| 0.52 |
| 0.89 |
| 0.10 |
| 0.00 |
| 0.20 |
| 0.07 |
| 0.18 |
| 1.00 |
| 0.22 |
| 0.01 |
| 0.00 |
| 0.70 |
| 1.00 |
| 0.03 |
| 0.00 |
| 0.00 |
| 0.59 |
| 0.00 |
| 0.00 |
| 0.00 |
| 0.29 |
| 1.00 |
| 0.00 |
| 1.00 |
| 1.00 |
| 0.00 |
| 0.00 |
| 0.01 |
| 0.25 |
| 0.36 |
| 0.00 |
| 0.00 |

| |
|---|
| 1.00 |
| 1.00 |
| 0.23 |
| 0.00 |
| 0.04 |
| 1.00 |
| 0.00 |
| 0.84 |
| 0.00 |
| 0.61 |
| 0.07 |
| 1.00 |
| 1.00 |
| 0.00 |
| 1.00 |
| 1.00 |
| 0.03 |
| 0.00 |
| 0.00 |
| 1.00 |
| 0.23 |
| 0.73 |
| 0.60 |
| 0.00 |
| 1.00 |
| 1.00 |
| 0.32 |
| 1.00 |
| 0.84 |
| 0.97 |
| 1.00 |
| 0.00 |
| 1.00 |
| 0.68 |
| 0.14 |
| 1.00 |
| 1.00 |
| 0.23 |
| 0.78 |
| 1.00 |
| 0.38 |
| 1.00 |
| 0.09 |
| 0.89 |
| 0.11 |
| 0.70 |
| 1.00 |
| 0.16 |

| |
|---|
| 1.00 |
| 0.01 |
| 0.68 |
| 0.22 |
| 0.62 |
| 1.00 |
| 1.00 |
| 0.00 |
| 0.78 |
| 0.33 |
| 0.18 |
| 1.00 |
| 0.82 |
| 1.00 |
| 0.00 |
| 0.00 |
| 1.00 |
| 0.30 |
| 0.00 |
| 0.64 |
| 1.00 |
| 0.98 |
| 1.00 |
| 0.84 |
| 1.00 |
| 0.26 |
| 0.35 |
| 0.13 |
| 0.50 |
| 1.00 |
| 0.90 |
| 0.04 |
| 0.25 |
| 0.01 |
| 0.05 |
| 0.00 |
| 0.00 |
| 0.28 |
| 0.00 |
| 0.00 |
| 0.99 |
| 0.00 |
| 0.00 |
| 0.03 |
| 1.00 |
| 0.00 |
| 0.00 |
| 0.00 |

| |
|---|
| 1.00 |
| 0.21 |
| 0.19 |
| 1.00 |
| 1.00 |
| 1.00 |
| 0.54 |
| 0.00 |
| 1.00 |
| 0.02 |
| 0.00 |
| 0.57 |
| 0.97 |
| 0.23 |
| 0.32 |
| 0.81 |
| 0.00 |
| 0.00 |
| 0.00 |
| 0.00 |
| 0.00 |
| 0.00 |
| 0.00 |
| 0.03 |
| 0.01 |
| 0.00 |
| 0.00 |
| 0.00 |
| 1.00 |
| 0.00 |
| 0.00 |
| 0.29 |
| 0.01 |
| 0.09 |
| 1.00 |
| 1.00 |
| 0.84 |
| 1.00 |
| 1.00 |
| 1.00 |
| 0.97 |
| 0.19 |
| 0.00 |
| 0.00 |
| 0.10 |
| 1.00 |
| 1.00 |
| 0.64 |

| |
|---|
| 1.00 |
| 0.00 |
| 1.00 |
| 0.01 |
| 0.80 |
| 0.29 |
| 0.00 |
| 0.55 |
| 0.00 |
| 0.00 |
| 0.00 |
| 0.91 |
| 0.01 |
| 1.00 |
| 0.56 |
| 0.06 |
| 0.00 |
| 0.01 |
| 1.00 |
| 0.00 |
| 0.02 |
| 0.72 |
| 0.87 |
| 0.10 |
| 0.25 |
| 0.02 |
| 0.11 |
| 0.16 |
| 0.08 |
| 1.00 |
| 1.00 |
| 0.51 |
| 0.48 |
| 0.81 |
| 0.25 |
| 1.00 |
| 0.98 |
| 0.41 |
| 0.00 |
| 0.00 |
| 0.00 |
| 0.65 |
| 1.00 |
| 0.00 |
| 0.00 |
| 1.00 |
| 1.00 |
| 0.87 |

| |
|---|
| 1.00 |
| 1.00 |
| 1.00 |
| 0.04 |
| 0.93 |
| 1.00 |
| 1.00 |
| 0.60 |
| 0.73 |
| 0.01 |
| 1.00 |
| 0.00 |
| 1.00 |
| 0.00 |
| 0.00 |
| 0.00 |
| 0.00 |
| 0.92 |
| 0.00 |
| 0.00 |
| 1.00 |
| 0.02 |
| 1.00 |
| 0.00 |
| 1.00 |
| 0.05 |
| 0.91 |
| 0.01 |
| 0.45 |
| 0.01 |
| 0.64 |
| 0.79 |
| 0.08 |
| 0.83 |
| 0.14 |
| 0.24 |
| 0.12 |
| 0.42 |
| 0.48 |
| 0.00 |
| 0.00 |
| 1.00 |
| 0.20 |
| 0.39 |
| 0.42 |
| 0.00 |
| 0.00 |
| 0.54 |

| |
|---|
| 1.00 |
| 0.00 |
| 1.00 |
| 1.00 |
| 1.00 |
| 0.12 |
| 1.00 |
| 1.00 |
| 0.00 |
| 0.00 |
| 0.83 |
| 0.01 |
| 0.43 |
| 0.72 |
| 0.03 |
| 1.00 |
| 0.79 |
| 1.00 |
| 0.46 |
| 1.00 |
| 0.00 |
| 0.78 |
| 0.00 |
| 0.58 |
| 0.47 |
| 0.52 |
| 0.10 |
| 0.48 |
| 0.00 |
| 0.00 |
| 1.00 |
| 0.82 |
| 0.33 |
| 0.23 |
| 0.06 |
| 0.01 |
| 0.06 |
| 1.00 |
| 1.00 |
| 1.00 |
| 1.00 |
| 0.34 |
| 0.78 |
| 0.00 |
| 1.00 |
| 0.96 |
| 1.00 |
| 0.60 |

| |
|---|
| 1.00 |
| 0.02 |
| 1.00 |
| 1.00 |
| 1.00 |
| 0.37 |
| 0.00 |
| 0.78 |
| 0.41 |
| 0.05 |
| 0.00 |
| 0.00 |
| 1.00 |
| 0.83 |
| 1.00 |
| 1.00 |
| 1.00 |
| 0.40 |
| 0.06 |
| 1.00 |
| 1.00 |
| 0.25 |
| 1.00 |
| 1.00 |
| 0.42 |
| 0.03 |
| 0.92 |
| 0.78 |
| 0.46 |
| 0.74 |
| 0.25 |
| 1.00 |
| 0.00 |
| 1.00 |
| 1.00 |
| 1.00 |
| 1.00 |
| 0.84 |
| 0.27 |
| 0.30 |
| 1.00 |
| 0.05 |
| 0.00 |
| 1.00 |
| 0.06 |
| 0.89 |
| 0.00 |
| 0.05 |

| |
|---|
| 1.00 |
| 0.30 |
| 0.95 |
| 1.00 |
| 0.53 |
| 1.00 |
| 0.68 |
| 1.00 |
| 0.62 |
| 0.68 |
| 0.40 |
| 0.00 |
| 0.00 |
| 0.70 |
| 1.00 |
| 1.00 |
| 0.03 |
| 0.00 |
| 0.59 |
| 0.00 |
| 0.31 |
| 0.01 |
| 0.22 |
| 0.25 |
| 1.00 |
| 0.13 |
| 1.00 |
| 0.31 |
| 1.00 |
| 1.00 |
| 1.00 |
| 1.00 |
| 0.05 |
| 1.00 |
| 0.59 |
| 0.10 |
| 0.25 |
| 1.00 |
| 0.13 |
| 1.00 |
| 0.15 |
| 0.02 |
| 1.00 |
| 1.00 |
| 1.00 |
| 0.06 |
| 1.00 |
| 0.24 |

| |
|---|
| 0.02 |
| 0.33 |
| 1.00 |
| 0.18 |
| 1.00 |
| 0.05 |
| 0.74 |
| 1.00 |
| 0.01 |
| 0.00 |
| 0.00 |
| 0.56 |
| 0.00 |
| 0.07 |
| 1.00 |
| 0.38 |
| 0.62 |
| 0.85 |
| 0.68 |
| 1.00 |
| 0.61 |
| 0.00 |
| 0.13 |
| 1.00 |
| 0.05 |
| 0.01 |
| 1.00 |
| 0.61 |
| 0.43 |
| 0.01 |
| 1.00 |
| 1.00 |
| 1.00 |
| 1.00 |
| 0.70 |
| 0.01 |
| 0.00 |
| 0.25 |
| 1.00 |
| 1.00 |
| 0.00 |
| 1.00 |
| 1.00 |
| 1.00 |
| 1.00 |
| 0.35 |
| 1.00 |
| 1.00 |

| |
|---|
| 0.15 |
| 0.08 |
| 1.00 |
| 1.00 |
| 1.00 |
| 0.14 |
| 0.00 |
| 1.00 |
| 0.00 |
| 0.28 |
| 0.10 |
| 0.00 |
| 0.73 |
| 0.02 |
| 1.00 |
| 1.00 |
| 1.00 |
| 1.00 |
| 1.00 |
| 1.00 |
| 1.00 |
| 0.93 |
| 1.00 |
| 0.72 |
| 0.18 |
| 0.00 |
| 0.29 |
| 0.30 |
| 1.00 |
| 0.97 |
| 0.00 |
| 0.00 |
| 0.50 |
| 1.00 |
| 1.00 |
| 1.00 |
| 1.00 |
| 0.05 |
| 0.07 |
| 1.00 |
| 0.23 |
| 1.00 |
| 1.00 |
| 0.59 |
| 0.12 |
| 1.00 |
| 0.26 |
| 0.63 |

| |
|---|
| 0.00 |
| 0.01 |
| 0.84 |
| 0.28 |
| 1.00 |
| 1.00 |
| 1.00 |
| 0.39 |
| 1.00 |
| 1.00 |
| 0.91 |
| 0.87 |
| 1.00 |
| 0.88 |
| 0.09 |
| 0.00 |
| 0.00 |
| 0.62 |
| 1.00 |
| 0.00 |
| 1.00 |
| 1.00 |
| 0.11 |
| 0.04 |
| 1.00 |
| 0.00 |
| 0.11 |
| 0.07 |
| 1.00 |
| 0.15 |
| 1.00 |
| 1.00 |
| 0.95 |
| 0.00 |
| 0.21 |
| 0.24 |
| 0.00 |
| 1.00 |
| 0.02 |
| 1.00 |
| 0.59 |
| 0.00 |
| 0.10 |
| 0.69 |
| 0.00 |
| 0.85 |
| 0.89 |
| 0.76 |

| |
|---|
| 0.08 |
| 0.09 |
| 0.93 |
| 0.06 |
| 0.00 |
| 0.01 |
| 0.63 |
| 0.00 |
| 0.16 |
| 1.00 |
| 0.01 |
| 0.00 |
| 0.00 |
| 1.00 |
| 0.94 |
| 0.84 |
| 1.00 |
| 0.18 |
| 1.00 |
| 0.09 |
| 0.33 |
| 1.00 |
| 0.71 |
| 0.00 |
| 0.44 |
| 1.00 |
| 0.02 |
| 0.44 |
| 0.00 |
| 0.03 |
| 0.54 |
| 0.04 |
| 1.00 |
| 0.08 |
| 0.01 |
| 0.84 |
| 0.01 |
| 1.00 |
| 0.00 |
| 0.46 |
| 0.78 |
| 0.15 |
| 1.00 |
| 1.00 |
| 1.00 |
| 1.00 |
| 0.00 |
| 1.00 |

| |
|---|
| 1.00 |
| 1.00 |
| 0.36 |
| 0.70 |
| 1.00 |
| 0.58 |
| 0.04 |
| 0.07 |
| 0.89 |
| 1.00 |
| 0.02 |
| 0.01 |
| 1.00 |
| 1.00 |
| 0.75 |
| 0.70 |
| 1.00 |
| 0.00 |
| 0.35 |
| 0.67 |
| 0.76 |
| 0.00 |
| 0.00 |
| 0.01 |
| 0.14 |
| 1.00 |
| 0.74 |
| 0.29 |
| 1.00 |
| 1.00 |
| 1.00 |
| 0.77 |
| 1.00 |
| 0.15 |
| 0.26 |
| 0.16 |
| 0.12 |
| 0.69 |
| 0.29 |
| 0.93 |
| 0.86 |
| 0.09 |
| 0.96 |
| 0.94 |
| 0.10 |
| 0.99 |
| 0.01 |
| 0.89 |

| |
|---|
| 1.00 |
| 1.00 |
| 0.41 |
| 0.99 |
| 0.60 |
| 1.00 |
| 1.00 |
| 0.24 |
| 0.84 |
| 1.00 |
| 0.15 |
| 0.73 |
| 0.00 |
| 0.00 |
| 0.38 |
| 0.00 |
| 0.38 |
| 1.00 |
| 1.00 |
| 0.99 |
| 0.40 |
| 0.46 |
| 1.00 |
| 0.24 |
| 1.00 |
| 0.19 |
| 0.17 |
| 1.00 |
| 0.91 |
| 0.00 |
| 0.51 |
| 0.17 |
| 1.00 |
| 0.00 |
| 0.87 |
| 0.16 |
| 1.00 |
| 0.27 |
| 0.44 |
| 0.86 |
| 0.11 |
| 1.00 |
| 1.00 |
| 1.00 |
| 1.00 |
| 0.95 |
| 0.63 |
| 0.10 |

| |
|---|
| 1.00 |
| 0.35 |
| 1.00 |
| 1.00 |
| 0.47 |
| 0.71 |
| 1.00 |
| 0.02 |
| 1.00 |
| 1.00 |
| 1.00 |
| 0.05 |
| 0.70 |
| 0.01 |
| 0.75 |
| 0.81 |
| 1.00 |
| 1.00 |
| 0.53 |
| 1.00 |
| 0.63 |
| 0.70 |
| 0.16 |
| 0.00 |
| 0.00 |
| 0.85 |
| 0.10 |
| 1.00 |
| 0.13 |
| 0.00 |
| 1.00 |
| 1.00 |
| 1.00 |
| 1.00 |
| 0.24 |
| 0.35 |
| 1.00 |
| 0.00 |
| 1.00 |
| 0.27 |
| 0.55 |
| 1.00 |
| 0.49 |
| 0.31 |
| 0.36 |
| 0.27 |
| 0.88 |
| 0.42 |

| |
|---|
| 0.00 |
| 1.00 |
| 0.77 |
| 1.00 |
| 0.24 |
| 1.00 |
| 0.23 |
| 0.54 |
| 0.18 |
| 0.03 |
| 1.00 |
| 1.00 |
| 0.89 |
| 0.99 |
| 0.01 |
| 0.92 |
| 0.01 |
| 0.24 |
| 0.04 |
| 1.00 |
| 1.00 |
| 1.00 |
| 1.00 |
| 1.00 |
| 0.56 |
| 1.00 |
| 0.61 |
| 1.00 |
| 0.40 |
| 1.00 |
| 0.27 |
| 0.55 |
| 0.19 |
| 1.00 |
| 0.36 |
| 1.00 |
| 0.89 |
| 1.00 |
| 1.00 |
| 1.00 |
| 0.96 |
| 1.00 |
| 0.75 |
| 0.83 |
| 1.00 |
| 0.17 |
| 0.71 |
| 0.05 |

| |
|---|
| 0.89 |
| 1.00 |
| 1.00 |
| 0.76 |
| 0.74 |
| 0.97 |
| 0.27 |
| 0.92 |
| 0.07 |
| 0.64 |
| 0.36 |
| 0.61 |
| 0.11 |
| 0.48 |
| 1.00 |
| 1.00 |
| 0.25 |
| 0.53 |
| 1.00 |
| 0.90 |
| 0.00 |
| 0.64 |
| 1.00 |
| 1.00 |
| 0.50 |
| 1.00 |
| 0.95 |
| 0.00 |
| 0.18 |
| 1.00 |
| 1.00 |
| 1.00 |
| 0.29 |
| 1.00 |
| 0.10 |
| 0.32 |
| 1.00 |
| 1.00 |
| 0.11 |
| 0.39 |
| 0.04 |
| 1.00 |
| 1.00 |
| 0.68 |
| 0.13 |
| 1.00 |
| 0.44 |
| 1.00 |

| |
|---|
| 0.11 |
| 0.00 |
| 1.00 |
| 1.00 |
| 0.97 |
| 1.00 |
| 0.04 |
| 1.00 |
| 0.00 |
| 0.76 |
| 0.18 |
| 0.00 |
| 0.00 |
| 0.68 |
| 1.00 |
| 1.00 |
| 0.19 |
| 1.00 |
| 1.00 |
| 0.78 |
| 0.83 |
| 0.00 |
| 1.00 |
| 1.00 |
| 1.00 |
| 0.89 |
| 0.05 |
| 0.89 |
| 0.03 |
| 1.00 |
| 0.03 |
| 1.00 |
| 0.00 |
| 0.35 |
| 0.06 |
| 0.01 |
| 0.84 |
| 0.00 |
| 0.97 |
| 0.61 |
| 0.07 |
| 1.00 |
| 1.00 |
| 0.10 |
| 1.00 |
| 0.21 |
| 0.39 |
| 0.51 |

| |
|---|
| 0.00 |
| 0.00 |
| 0.50 |
| 0.00 |
| 0.72 |
| 0.15 |
| 0.34 |
| 1.00 |
| 1.00 |
| 0.75 |
| 0.05 |
| 1.00 |
| 1.00 |
| 0.84 |
| 0.09 |
| 1.00 |
| 0.69 |
| 0.33 |
| 0.56 |
| 0.86 |
| 1.00 |
| 0.10 |
| 0.38 |
| 1.00 |
| 1.00 |
| 0.00 |
| 0.00 |
| 0.70 |
| 0.70 |
| 0.02 |
| 0.89 |
| 0.14 |
| 0.36 |
| 1.00 |
| 1.00 |
| 1.00 |
| 0.43 |
| 0.39 |
| 1.00 |
| 1.00 |
| 0.63 |
| 0.61 |
| 0.69 |
| 1.00 |
| 1.00 |
| 0.86 |
| 1.00 |
| 0.63 |

| |
|---|
| 0.44 |
| 0.11 |
| 0.49 |
| 0.97 |
| 1.00 |
| 0.73 |
| 0.00 |
| 1.00 |
| 0.56 |
| 1.00 |
| 0.00 |
| 0.94 |
| 0.06 |
| 0.55 |
| 0.27 |
| 0.52 |
| 0.37 |
| 0.16 |
| 1.00 |
| 0.13 |
| 1.00 |
| 0.01 |
| 0.33 |
| 1.00 |
| 0.60 |
| 0.33 |
| 0.80 |
| 1.00 |
| 0.09 |
| 0.92 |
| 0.61 |
| 0.50 |
| 0.02 |
| 0.01 |
| 0.32 |
| 1.00 |
| 0.73 |
| 0.00 |
| 1.00 |
| 0.00 |
| 0.52 |
| 1.00 |
| 1.00 |
| 0.09 |
| 1.00 |
| 0.01 |
| 1.00 |
| 1.00 |

| |
|---|
| 0.11 |
| 1.00 |
| 1.00 |
| 0.00 |
| 0.77 |
| 0.58 |
| 0.76 |
| 0.44 |
| 1.00 |
| 0.11 |
| 1.00 |
| 0.20 |
| 0.97 |
| 0.00 |
| 0.00 |
| 0.10 |
| 0.17 |
| 0.00 |
| 0.00 |
| 0.00 |
| 0.01 |
| 0.91 |
| 1.00 |
| 0.57 |
| 1.00 |
| 0.13 |
| 0.00 |
| 0.12 |
| 0.00 |
| 0.08 |
| 0.00 |
| 1.00 |
| 1.00 |
| 0.29 |
| 0.73 |
| 1.00 |
| 0.01 |
| 0.00 |
| 0.28 |
| 0.95 |
| 0.67 |
| 0.15 |
| 0.00 |
| 1.00 |
| 1.00 |
| 0.11 |
| 0.00 |
| 0.89 |

| |
|---|
| 0.07 |
| 0.00 |
| 0.00 |
| 0.55 |
| 1.00 |
| 0.00 |
| 0.00 |
| 0.00 |
| 0.59 |
| 0.05 |
| 0.90 |
| 0.98 |
| 0.00 |
| 1.00 |
| 0.62 |
| 0.35 |
| 0.30 |
| 1.00 |
| 0.49 |
| 0.77 |
| 0.06 |
| 1.00 |
| 0.03 |
| 0.25 |
| 1.00 |
| 0.07 |
| 0.01 |
| 0.35 |
| 0.75 |
| 1.00 |
| 1.00 |
| 1.00 |
| 1.00 |
| 0.46 |
| 0.52 |
| 0.87 |
| 1.00 |
| 1.00 |
| 1.00 |
| 0.24 |
| 1.00 |
| 0.02 |
| 0.00 |
| 0.05 |
| 1.00 |
| 0.29 |
| 0.38 |
| 1.00 |

| |
|---|
| 0.98 |
| 0.04 |
| 1.00 |
| 0.11 |
| 0.08 |
| 0.25 |
| 0.18 |
| 1.00 |
| 0.11 |
| 0.00 |
| 0.68 |
| 0.99 |
| 1.00 |
| 0.00 |
| 0.14 |
| 0.33 |
| 1.00 |
| 0.00 |
| 0.00 |
| 1.00 |
| 1.00 |
| 0.00 |
| 1.00 |
| 0.80 |
| 0.79 |
| 0.00 |
| 0.97 |
| 1.00 |
| 1.00 |
| 0.01 |
| 0.30 |
| 0.00 |
| 0.31 |
| 0.18 |
| 0.33 |
| 0.02 |
| 0.60 |
| 0.00 |
| 0.03 |
| 1.00 |
| 1.00 |
| 0.00 |
| 0.03 |
| 0.00 |
| 0.00 |
| 0.65 |
| 0.63 |
| 0.07 |

| |
|---|
| 1.00 |
| 0.01 |
| 0.55 |
| 0.00 |
| 0.00 |
| 0.00 |
| 0.00 |
| 0.00 |
| 0.00 |
| 0.36 |
| 0.45 |
| 1.00 |
| 0.10 |
| 1.00 |
| 1.00 |
| 0.36 |
| 0.00 |
| 0.23 |
| 0.00 |
| 0.00 |
| 0.00 |
| 0.46 |
| 0.37 |
| 1.00 |
| 1.00 |
| 0.00 |
| 0.02 |
| 0.26 |
| 0.01 |
| 1.00 |
| 1.00 |
| 1.00 |
| 0.00 |
| 0.00 |
| 0.00 |
| 1.00 |
| 1.00 |
| 0.00 |
| 0.00 |
| 0.01 |
| 0.63 |
| 0.82 |
| 0.84 |
| 0.20 |
| 0.61 |
| 0.11 |
| 0.22 |
| 0.29 |

| |
|---|
| 0.00 |
| 1.00 |
| 0.02 |
| 0.69 |
| 0.01 |
| 0.00 |
| 1.00 |
| 0.25 |
| 0.70 |
| 0.72 |
| 0.18 |
| 0.69 |
| 1.00 |
| 1.00 |
| 0.11 |
| 0.00 |
| 1.00 |
| 0.38 |
| 0.00 |
| 0.09 |
| 0.45 |
| 0.00 |
| 0.00 |
| 0.01 |
| 0.07 |
| 0.08 |
| 0.05 |
| 0.72 |
| 1.00 |
| 1.00 |
| 0.41 |
| 0.87 |
| 0.52 |
| 0.00 |
| 0.46 |
| 0.15 |
| 0.17 |
| 0.47 |
| 1.00 |
| 1.00 |
| 1.00 |
| 0.01 |
| 1.00 |
| 0.06 |
| 0.45 |
| 0.07 |
| 0.62 |
| 0.00 |

| |
|---|
| 0.36 |
| 0.00 |
| 0.02 |
| 0.00 |
| 0.25 |
| 0.02 |
| 0.01 |
| 0.00 |
| 1.00 |
| 0.81 |
| 0.00 |
| 1.00 |
| 0.21 |
| 0.02 |
| 0.04 |
| 0.00 |
| 0.50 |
| 1.00 |
| 0.33 |
| 0.00 |
| 0.42 |
| 0.64 |
| 0.00 |
| 0.00 |
| 0.19 |
| 0.06 |
| 0.00 |
| 0.14 |
| 1.00 |
| 0.16 |
| 1.00 |
| 1.00 |
| 1.00 |
| 1.00 |
| 1.00 |
| 0.22 |
| 0.00 |
| 0.08 |
| 0.03 |
| 0.28 |
| 0.19 |
| 0.00 |
| 1.00 |
| 0.15 |
| 0.97 |
| 0.02 |
| 1.00 |
| 1.00 |

| |
|---|
| 0.00 |
| 0.40 |
| 0.27 |
| 0.07 |
| 1.00 |
| 0.63 |
| 0.00 |
| 0.00 |
| 0.00 |
| 0.00 |
| 0.11 |
| 1.00 |
| 0.00 |
| 1.00 |
| 0.00 |
| 0.00 |
| 1.00 |
| 0.51 |
| 1.00 |
| 0.55 |
| 1.00 |
| 0.40 |
| 0.27 |
| 0.45 |
| 0.00 |
| 0.59 |
| 0.00 |
| 1.00 |
| 0.40 |
| 0.09 |
| 0.81 |
| 0.01 |
| 0.00 |
| 0.90 |
| 0.65 |
| 0.80 |
| 0.07 |
| 1.00 |
| 1.00 |
| 1.00 |
| 1.00 |
| 0.00 |
| 0.07 |
| 0.00 |
| 1.00 |
| 0.00 |
| 0.38 |
| 0.42 |

| |
|---|
| 0.03 |
| 0.00 |
| 1.00 |
| 1.00 |
| 1.00 |
| 0.39 |
| 0.00 |
| 0.00 |
| 0.92 |
| 0.09 |
| 1.00 |
| 0.37 |
| 0.14 |
| 1.00 |
| 0.00 |
| 0.56 |
| 0.91 |
| 0.01 |
| 0.02 |
| 0.65 |
| 0.43 |
| 0.04 |
| 0.00 |
| 1.00 |
| 0.00 |
| 0.23 |
| 0.15 |
| 0.42 |
| 0.27 |
| 0.00 |
| 1.00 |
| 1.00 |
| 0.00 |
| 0.04 |
| 0.97 |
| 0.01 |
| 0.00 |
| 0.00 |
| 0.30 |
| 0.04 |
| 0.08 |
| 0.01 |
| 0.89 |
| 1.00 |
| 0.95 |
| 0.30 |
| 0.05 |
| 0.01 |

| |
|---|
| 0.38 |
| 0.24 |
| 0.73 |
| 0.98 |
| 1.00 |
| 0.00 |
| 0.00 |
| 1.00 |
| 1.00 |
| 0.00 |
| 0.00 |
| 1.00 |
| 0.00 |
| 0.17 |
| 1.00 |
| 0.37 |
| 0.00 |
| 0.01 |
| 1.00 |
| 1.00 |
| 0.21 |
| 0.62 |
| 0.09 |
| 1.00 |
| 1.00 |
| 1.00 |
| 0.00 |
| 0.00 |
| 0.01 |
| 0.00 |
| 0.07 |
| 0.12 |
| 1.00 |
| 0.22 |
| 0.00 |
| 0.00 |
| 0.01 |
| 0.65 |
| 0.02 |
| 1.00 |
| 0.03 |
| 0.07 |
| 0.11 |
| 1.00 |
| 1.00 |
| 0.15 |
| 0.00 |
| 0.77 |

| |
|---|
| 0.00 |
| 1.00 |
| 0.15 |
| 0.64 |
| 0.07 |
| 0.00 |
| 0.14 |
| 0.00 |
| 0.18 |
| 0.00 |
| 0.10 |
| 0.46 |
| 0.44 |
| 1.00 |
| 0.02 |
| 0.51 |
| 0.00 |
| 0.00 |
| 0.00 |
| 0.01 |
| 0.00 |
| 0.50 |
| 0.00 |
| 1.00 |
| 0.06 |
| 0.00 |
| 0.31 |
| 0.00 |
| 0.61 |
| 0.52 |
| 0.25 |
| 1.00 |
| 0.66 |
| 1.00 |
| 1.00 |
| 0.84 |
| 1.00 |
| 1.00 |
| 0.08 |
| 1.00 |
| 0.12 |
| 0.05 |
| 0.16 |
| 0.34 |
| 1.00 |
| 0.04 |
| 1.00 |
| 1.00 |

| |
|---|
| 0.00 |
| 1.00 |
| 0.02 |
| 0.04 |
| 1.00 |
| 0.00 |
| 0.50 |
| 1.00 |
| 0.15 |
| 0.08 |
| 1.00 |
| 0.54 |
| 1.00 |
| 0.00 |
| 0.03 |
| 0.00 |
| 0.00 |
| 1.00 |
| 0.62 |
| 0.46 |
| 0.13 |
| 1.00 |
| 0.26 |
| 0.86 |
| 0.00 |
| 0.00 |
| 0.01 |
| 0.21 |
| 0.00 |
| 0.00 |
| 1.00 |
| 0.00 |
| 0.05 |
| 1.00 |
| 0.00 |
| 0.00 |
| 0.00 |
| 0.00 |
| 1.00 |
| 1.00 |
| 1.00 |
| 0.00 |
| 0.46 |
| 0.54 |
| 0.61 |
| 0.39 |
| 0.58 |
| 1.00 |

| |
|---|
| 0.08 |
| 1.00 |
| 0.00 |
| 0.70 |
| 0.90 |
| 0.04 |
| 0.45 |
| 0.72 |
| 0.00 |
| 0.08 |
| 1.00 |
| 0.69 |
| 0.02 |
| 0.15 |
| 0.27 |
| 0.14 |
| 0.07 |
| 0.00 |
| 0.89 |
| 0.90 |
| 0.00 |
| 0.09 |
| 0.11 |
| 0.63 |
| 0.00 |
| 0.84 |
| 0.00 |
| 0.00 |
| 0.07 |
| 0.00 |
| 0.01 |
| 0.61 |
| 1.00 |
| 0.22 |
| 1.00 |
| 0.58 |
| 0.12 |
| 0.00 |
| 0.03 |
| 0.00 |
| 0.02 |
| 0.00 |
| 0.29 |
| 0.08 |
| 0.21 |
| 0.00 |
| 1.00 |
| 0.48 |

| |
|---|
| 1.00 |
| 0.43 |
| 0.00 |
| 0.50 |
| 1.00 |
| 0.02 |
| 0.93 |
| 1.00 |
| 0.10 |
| 0.45 |
| 0.56 |
| 0.61 |
| 0.01 |
| 0.78 |
| 0.07 |
| 0.18 |
| 0.00 |
| 1.00 |
| 1.00 |
| 1.00 |
| 0.00 |
| 1.00 |
| 0.00 |
| 0.00 |
| 0.00 |
| 0.10 |
| 0.00 |
| 0.00 |
| 1.00 |
| 0.78 |
| 0.91 |
| 0.00 |
| 1.00 |
| 0.05 |
| 0.00 |
| 0.79 |
| 0.00 |
| 0.77 |
| 0.10 |
| 1.00 |
| 1.00 |
| 0.32 |
| 0.76 |
| 1.00 |
| 1.00 |
| 0.64 |
| 0.16 |
| 0.38 |

| |
|---|
| 1.00 |
| 0.04 |
| 0.13 |
| 0.00 |
| 0.78 |
| 0.27 |
| 0.00 |
| 0.94 |
| 1.00 |
| 0.65 |
| 1.00 |
| 1.00 |
| 0.28 |
| 0.00 |
| 1.00 |
| 1.00 |
| 0.70 |
| 0.83 |
| 0.38 |
| 0.00 |
| 1.00 |
| 1.00 |
| 1.00 |
| 0.97 |
| 0.54 |
| 1.00 |
| 0.38 |
| 1.00 |
| 0.92 |
| 0.00 |
| 0.00 |
| 0.00 |
| 0.00 |
| 0.08 |
| 1.00 |
| 0.34 |
| 0.70 |
| 1.00 |
| 0.33 |
| 0.66 |
| 0.42 |
| 0.29 |
| 0.79 |
| 0.19 |
| 0.54 |
| 0.99 |
| 0.19 |
| 0.61 |

| |
|---|
| 0.70 |
| 0.49 |
| 0.16 |
| 0.39 |
| 1.00 |
| 0.00 |
| 0.04 |
| 1.00 |
| 0.00 |
| 0.42 |
| 0.74 |
| 0.77 |
| 0.89 |
| 0.70 |
| 0.12 |
| 0.97 |
| 0.36 |
| 0.14 |
| 1.00 |
| 0.79 |
| 1.00 |
| 0.36 |
| 0.01 |
| 0.11 |
| 1.00 |
| 1.00 |
| 0.51 |
| 0.46 |
| 0.90 |
| 0.19 |
| 0.38 |
| 1.00 |
| 0.11 |
| 0.14 |
| 0.75 |
| 0.97 |
| 0.02 |
| 0.06 |
| 1.00 |
| 0.91 |
| 0.36 |
| 1.00 |
| 1.00 |
| 1.00 |
| 0.08 |
| 1.00 |
| 0.12 |
| 0.96 |

| |
|---|
| 1.00 |
| 0.42 |
| 0.58 |
| 0.08 |
| 0.00 |
| 1.00 |
| 0.75 |
| 0.67 |
| 0.74 |
| 0.39 |
| 0.78 |
| 0.94 |
| 1.00 |
| 0.39 |
| 1.00 |
| 1.00 |
| 1.00 |
| 0.00 |
| 0.01 |
| 0.19 |
| 1.00 |
| 0.03 |
| 0.17 |
| 1.00 |
| 0.57 |
| 0.93 |
| 1.00 |
| 0.30 |
| 0.55 |
| 0.48 |
| 0.26 |
| 1.00 |
| 0.15 |
| 0.04 |
| 0.18 |
| 0.12 |
| 0.18 |
| 0.16 |
| 1.00 |
| 0.02 |
| 1.00 |
| 0.00 |
| 0.01 |
| 0.61 |
| 1.00 |
| 0.42 |
| 0.76 |
| 0.01 |

| |
|---|
| 0.67 |
| 1.00 |
| 0.93 |
| 1.00 |
| 0.74 |
| 0.54 |
| 0.16 |
| 1.00 |
| 1.00 |
| 0.51 |
| 0.00 |
| 1.00 |
| 0.16 |
| 0.13 |
| 0.45 |
| 1.00 |
| 1.00 |
| 1.00 |
| 1.00 |
| 1.00 |
| 1.00 |
| 1.00 |
| 0.86 |
| 0.63 |
| 0.64 |
| 0.82 |
| 0.19 |
| 1.00 |
| 1.00 |
| 0.12 |
| 0.00 |
| 0.24 |
| 0.00 |
| 0.33 |
| 1.00 |
| 0.91 |
| 0.03 |
| 1.00 |
| 0.28 |
| 0.00 |
| 1.00 |
| 1.00 |
| 0.08 |
| 0.53 |
| 0.29 |
| 1.00 |
| 1.00 |
| 0.07 |

| |
|---|
| 0.00 |
| 1.00 |
| 1.00 |
| 0.28 |
| 0.37 |
| 0.00 |
| 0.00 |
| 0.10 |
| 0.05 |
| 0.51 |
| 0.06 |
| 0.50 |
| 0.90 |
| 0.94 |
| 0.21 |
| 0.13 |
| 0.95 |
| 0.90 |
| 0.00 |
| 0.00 |
| 0.01 |
| 0.00 |
| 1.00 |
| 0.01 |
| 0.39 |
| 1.00 |
| 0.00 |
| 1.00 |
| 0.00 |
| 0.00 |
| 0.64 |
| 1.00 |
| 1.00 |
| 0.94 |
| 0.00 |
| 1.00 |
| 0.34 |
| 0.00 |
| 0.15 |
| 0.00 |
| 0.03 |
| 0.01 |
| 0.07 |
| 0.00 |
| 1.00 |
| 0.00 |
| 1.00 |
| 0.00 |

| |
|---|
| 0.00 |
| 0.00 |
| 1.00 |
| 0.31 |
| 0.00 |
| 1.00 |
| 0.00 |
| 0.75 |
| 0.54 |
| 1.00 |
| 0.00 |
| 0.33 |
| 0.01 |
| 1.00 |
| 0.42 |
| 0.00 |
| 0.00 |
| 0.29 |
| 0.00 |
| 1.00 |
| 0.11 |
| 0.55 |
| 0.02 |
| 0.02 |
| 1.00 |
| 0.89 |
| 0.17 |
| 0.00 |
| 0.51 |
| 0.03 |
| 1.00 |
| 0.18 |
| 1.00 |
| 0.00 |
| 0.00 |
| 0.14 |
| 1.00 |
| 1.00 |
| 1.00 |
| 0.00 |
| 0.01 |
| 1.00 |
| 0.23 |
| 0.04 |
| 1.00 |
| 1.00 |
| 0.44 |
| 0.00 |

| |
|---|
| 0.69 |
| 0.00 |
| 0.78 |
| 0.00 |
| 1.00 |
| 0.00 |
| 0.00 |
| 1.00 |
| 0.77 |
| 0.00 |
| 1.00 |
| 0.00 |
| 1.00 |
| 0.00 |
| 0.21 |
| 0.00 |
| 0.13 |
| 1.00 |
| 1.00 |
| 0.00 |
| 0.33 |
| 0.84 |
| 0.00 |
| 0.52 |
| 0.00 |
| 0.74 |
| 0.76 |
| 0.00 |
| 1.00 |
| 0.05 |
| 0.50 |
| 0.00 |
| 0.00 |
| 0.01 |
| 0.51 |
| 0.01 |
| 0.03 |
| 0.00 |
| 0.00 |
| 0.00 |
| 0.00 |
| 0.04 |
| 0.00 |
| 0.01 |
| 1.00 |
| 0.05 |
| 0.73 |
| 0.61 |

| |
|---|
| 0.00 |
| 0.09 |
| 0.33 |
| 0.25 |
| 0.08 |
| 0.05 |
| 0.04 |
| 0.04 |
| 0.11 |
| 0.14 |
| 0.14 |
| 0.04 |
| 0.08 |
| 1.00 |
| 1.00 |
| 0.00 |
| 0.19 |
| 1.00 |
| 0.52 |
| 0.00 |
| 0.00 |
| 1.00 |
| 0.00 |
| 0.01 |
| 0.00 |
| 0.08 |
| 0.01 |
| 1.00 |
| 0.12 |
| 0.00 |
| 0.12 |
| 0.15 |
| 1.00 |
| 0.39 |
| 0.03 |
| 0.15 |
| 0.45 |
| 0.00 |
| 0.47 |
| 0.89 |
| 0.39 |
| 0.11 |
| 0.00 |
| 0.34 |
| 0.02 |
| 0.00 |
| 1.00 |
| 0.13 |

| |
|---|
| 0.00 |
| 0.32 |
| 0.41 |
| 0.23 |
| 0.00 |
| 0.03 |
| 1.00 |
| 0.70 |
| 0.00 |
| 0.00 |
| 0.00 |
| 1.00 |
| 0.49 |
| 0.01 |
| 1.00 |
| 0.00 |
| 0.12 |
| 1.00 |
| 0.28 |
| 0.15 |
| 0.35 |
| 1.00 |
| 0.98 |
| 1.00 |
| 0.89 |
| 1.00 |
| 0.13 |
| 0.02 |
| 0.89 |
| 1.00 |
| 1.00 |
| 0.02 |
| 1.00 |
| 0.19 |
| 0.77 |
| 1.00 |
| 0.67 |
| 0.66 |
| 0.12 |
| 0.43 |
| 0.41 |
| 0.40 |
| 0.28 |
| 1.00 |
| 1.00 |
| 0.60 |
| 0.00 |
| 0.89 |

| |
|---|
| 0.02 |
| 0.00 |
| 1.00 |
| 0.65 |
| 0.43 |
| 0.00 |
| 0.00 |
| 0.78 |
| 0.00 |
| 0.54 |
| 1.00 |
| 0.18 |
| 1.00 |
| 1.00 |
| 0.21 |
| 0.19 |
| 0.00 |
| 0.02 |
| 1.00 |
| 0.01 |
| 0.00 |
| 1.00 |
| 0.02 |
| 1.00 |
| 0.05 |
| 0.00 |
| 0.13 |
| 0.00 |
| 0.00 |
| 1.00 |
| 0.00 |
| 0.39 |
| 0.02 |
| 0.66 |
| 1.00 |
| 0.23 |
| 0.21 |
| 1.00 |
| 1.00 |
| 0.01 |
| 0.84 |
| 0.34 |
| 0.03 |
| 1.00 |
| 0.03 |
| 0.02 |
| 1.00 |
| 0.00 |

| |
|---|
| 0.85 |
| 0.13 |
| 0.06 |
| 1.00 |
| 1.00 |
| 0.97 |
| 0.78 |
| 0.11 |
| 0.27 |
| 0.00 |
| 0.21 |
| 0.78 |
| 0.68 |
| 0.02 |
| 1.00 |
| 0.20 |
| 0.03 |
| 0.77 |
| 0.34 |
| 0.97 |
| 0.40 |
| 0.52 |
| 0.79 |
| 1.00 |
| 0.41 |
| 0.21 |
| 1.00 |
| 0.92 |
| 0.67 |
| 0.11 |
| 0.07 |
| 0.88 |
| 0.65 |
| 0.10 |
| 0.26 |
| 1.00 |
| 1.00 |
| 0.89 |
| 0.00 |
| 0.17 |
| 0.00 |
| 0.00 |
| 0.00 |
| 0.68 |
| 0.00 |
| 1.00 |
| 1.00 |
| 0.60 |

| |
|---|
| 0.82 |
| 0.00 |
| 0.00 |
| 0.01 |
| 1.00 |
| 0.00 |
| 1.00 |
| 0.54 |
| 1.00 |
| 0.00 |
| 0.00 |
| 0.00 |
| 0.92 |
| 0.29 |
| 0.41 |
| 0.07 |
| 0.00 |
| 1.00 |
| 0.01 |
| 1.00 |
| 1.00 |
| 0.20 |
| 1.00 |
| 0.52 |
| 0.00 |
| 0.69 |
| 0.19 |
| 1.00 |
| 0.17 |
| 0.00 |
| 0.00 |
| 0.87 |
| 0.00 |
| 0.32 |
| 0.81 |
| 0.28 |
| 0.13 |
| 0.02 |
| 0.09 |
| 0.59 |
| 0.02 |
| 0.04 |
| 0.00 |
| 0.46 |
| 0.00 |
| 1.00 |
| 1.00 |
| 1.00 |

| |
|---|
| 0.71 |
| 0.00 |
| 1.00 |
| 0.00 |
| 0.01 |
| 0.01 |
| 0.56 |
| 1.00 |
| 0.86 |
| 1.00 |
| 0.27 |
| 0.05 |
| 1.00 |
| 0.00 |
| 0.06 |
| 0.39 |
| 0.01 |
| 1.00 |
| 0.38 |
| 0.60 |
| 0.00 |
| 1.00 |
| 1.00 |
| 1.00 |
| 0.93 |
| 0.22 |
| 1.00 |
| 0.13 |
| 0.00 |
| 0.00 |
| 0.98 |
| 0.00 |
| 0.21 |
| 0.00 |
| 0.53 |
| 0.22 |
| 0.00 |
| 0.00 |
| 1.00 |
| 0.01 |
| 0.01 |
| 1.00 |
| 0.80 |
| 0.11 |
| 0.98 |
| 0.15 |
| 1.00 |
| 0.00 |

| |
|---|
| 1.00 |
| 0.28 |
| 1.00 |
| 0.84 |
| 0.05 |
| 0.93 |
| 0.88 |
| 0.23 |
| 1.00 |
| 1.00 |
| 0.53 |
| 0.13 |
| 0.36 |
| 1.00 |
| 1.00 |
| 1.00 |
| 1.00 |
| 0.39 |
| 1.00 |
| 0.00 |
| 0.53 |
| 0.48 |
| 1.00 |
| 0.26 |
| 0.54 |
| 1.00 |
| 0.40 |
| 1.00 |
| 0.00 |
| 0.00 |
| 0.12 |
| 1.00 |
| 0.00 |
| 1.00 |
| 0.00 |
| 0.96 |
| 0.00 |
| 0.09 |
| 0.09 |
| 0.07 |
| 0.00 |
| 1.00 |
| 1.00 |
| 1.00 |
| 0.65 |
| 1.00 |
| 0.41 |
| 1.00 |

| |
|---|
| 0.27 |
| 0.01 |
| 0.01 |
| 0.00 |
| 1.00 |
| 0.00 |
| 1.00 |
| 0.69 |
| 0.00 |
| 0.00 |
| 0.00 |
| 0.16 |
| 1.00 |
| 0.01 |
| 0.14 |
| 0.00 |
| 0.00 |
| 0.53 |
| 1.00 |
| 0.14 |
| 0.01 |
| 0.51 |
| 0.76 |
| 1.00 |
| 0.00 |
| 0.00 |
| 0.00 |
| 0.68 |
| 0.05 |
| 0.73 |
| 0.00 |
| 0.97 |
| 0.81 |
| 0.00 |
| 0.00 |
| 0.15 |
| 1.00 |
| 0.00 |
| 0.04 |
| 1.00 |
| 0.19 |
| 0.00 |
| 0.00 |
| 0.00 |
| 0.09 |
| 0.00 |
| 1.00 |
| 0.73 |

| |
|---|
| 0.00 |
| 1.00 |
| 0.30 |
| 1.00 |
| 1.00 |
| 0.00 |
| 0.00 |
| 0.28 |
| 0.01 |
| 0.57 |
| 1.00 |
| 0.01 |
| 0.00 |
| 0.55 |
| 0.01 |
| 1.00 |
| 0.11 |
| 0.00 |
| 0.00 |
| 0.00 |
| 0.00 |
| 1.00 |
| 0.00 |
| 0.42 |
| 0.16 |
| 0.00 |
| 1.00 |
| 0.92 |
| 0.02 |
| 0.27 |
| 0.02 |
| 0.01 |
| 1.00 |
| 0.11 |
| 0.36 |
| 0.00 |
| 0.00 |
| 0.03 |
| 0.00 |
| 0.68 |
| 0.00 |
| 0.21 |
| 0.00 |
| 0.07 |
| 0.01 |
| 0.00 |
| 0.00 |
| 0.00 |

| |
|---|
| 0.08 |
| 0.46 |
| 0.64 |
| 0.00 |
| 1.00 |
| 0.89 |
| 0.00 |
| 0.49 |
| 0.70 |
| 0.44 |
| 1.00 |
| 0.38 |
| 1.00 |
| 0.00 |
| 0.10 |
| 0.05 |
| 0.23 |
| 1.00 |
| 0.86 |
| 0.12 |
| 1.00 |
| 0.10 |
| 0.01 |
| 1.00 |
| 0.00 |
| 1.00 |
| 0.00 |
| 0.00 |
| 1.00 |
| 1.00 |
| 0.00 |
| 0.02 |
| 0.00 |
| 0.00 |
| 0.10 |
| 0.83 |
| 1.00 |
| 1.00 |
| 1.00 |
| 0.07 |
| 0.00 |
| 0.00 |
| 0.00 |
| 1.00 |
| 0.00 |
| 0.04 |
| 0.03 |
| 0.09 |

| |
|---|
| 0.01 |
| 0.08 |
| 1.00 |
| 0.03 |
| 0.22 |
| 0.00 |
| 0.00 |
| 1.00 |
| 0.96 |
| 0.00 |
| 0.42 |
| 0.00 |
| 0.95 |
| 0.00 |
| 0.11 |
| 1.00 |
| 0.09 |
| 0.04 |
| 1.00 |
| 0.01 |
| 0.00 |
| 0.00 |
| 0.00 |
| 0.02 |
| 1.00 |
| 1.00 |
| 0.00 |
| 0.00 |
| 1.00 |
| 0.02 |
| 1.00 |
| 1.00 |
| 0.00 |
| 0.00 |
| 1.00 |
| 0.21 |
| 0.24 |
| 0.00 |
| 0.00 |
| 0.00 |
| 1.00 |
| 0.01 |
| 0.54 |
| 1.00 |
| 0.60 |
| 0.27 |
| 1.00 |
| 1.00 |

| |
|---|
| 1.00 |
| 0.03 |
| 0.00 |
| 0.42 |
| 1.00 |
| 0.00 |
| 0.46 |
| 0.98 |
| 0.51 |
| 0.03 |
| 0.36 |
| 0.05 |
| 0.88 |
| 1.00 |
| 0.16 |
| 0.00 |
| 0.07 |
| 0.00 |
| 0.06 |
| 1.00 |
| 0.01 |
| 0.46 |
| 0.31 |
| 0.07 |
| 0.00 |
| 1.00 |
| 1.00 |
| 0.00 |
| 1.00 |
| 0.00 |
| 0.58 |
| 0.01 |
| 0.16 |
| 0.03 |
| 0.33 |
| 0.00 |
| 0.61 |
| 0.25 |
| 0.62 |
| 1.00 |
| 0.08 |
| 0.44 |
| 0.66 |
| 0.01 |
| 0.64 |
| 0.00 |
| 1.00 |
| 0.00 |

| |
|---|
| 0.02 |
| 0.00 |
| 1.00 |
| 1.00 |
| 1.00 |
| 0.00 |
| 0.33 |
| 0.00 |
| 1.00 |
| 0.00 |
| 0.00 |
| 0.25 |
| 1.00 |
| 0.01 |
| 0.91 |
| 1.00 |
| 0.05 |
| 0.65 |
| 0.00 |
| 0.66 |
| 0.89 |
| 0.00 |
| 0.00 |
| 0.77 |
| 0.19 |
| 0.27 |
| 1.00 |
| 0.06 |
| 0.35 |
| 0.26 |
| 0.44 |
| 0.93 |
| 0.63 |
| 1.00 |
| 0.00 |
| 0.03 |
| 1.00 |
| 0.00 |
| 0.55 |
| 0.02 |
| 1.00 |
| 0.00 |
| 1.00 |
| 0.19 |
| 0.11 |
| 0.69 |
| 0.74 |
| 0.01 |

| |
|---|
| 0.68 |
| 0.00 |
| 0.95 |
| 0.42 |
| 0.21 |
| 1.00 |
| 0.20 |
| 0.00 |
| 0.00 |
| 1.00 |
| 0.84 |
| 0.01 |
| 0.00 |
| 0.00 |
| 0.96 |
| 1.00 |
| 0.00 |
| 1.00 |
| 1.00 |
| 0.04 |
| 0.23 |
| 1.00 |
| 1.00 |
| 1.00 |
| 0.49 |
| 0.00 |
| 0.95 |
| 1.00 |
| 0.00 |
| 1.00 |
| 0.00 |
| 0.84 |
| 1.00 |
| 0.73 |
| 1.00 |
| 0.00 |
| 1.00 |
| 0.06 |
| 0.47 |
| 0.00 |
| 0.95 |
| 0.01 |
| 0.48 |
| 0.03 |
| 0.01 |
| 0.90 |
| 0.52 |
| 0.12 |

| |
|---|
| 0.63 |
| 0.09 |
| 0.01 |
| 0.00 |
| 0.00 |
| 0.67 |
| 0.00 |
| 0.81 |
| 1.00 |
| 0.00 |
| 1.00 |
| 0.02 |
| 0.05 |
| 1.00 |
| 1.00 |
| 0.06 |
| 0.00 |
| 1.00 |
| 0.09 |
| 0.04 |
| 0.23 |
| 1.00 |
| 0.70 |
| 0.00 |
| 0.40 |
| 0.04 |
| 0.08 |
| 0.00 |
| 0.36 |
| 1.00 |
| 0.00 |
| 1.00 |
| 0.15 |
| 0.12 |
| 0.13 |
| 1.00 |
| 1.00 |
| 0.83 |
| 0.02 |
| 0.83 |
| 0.00 |
| 0.44 |
| 0.65 |
| 0.71 |
| 0.01 |
| 0.78 |
| 1.00 |
| 0.00 |

| |
|---|
| 1.00 |
| 0.87 |
| 0.01 |
| 1.00 |
| 0.00 |
| 0.00 |
| 0.00 |
| 1.00 |
| 1.00 |
| 0.15 |
| 0.00 |
| 0.08 |
| 1.00 |
| 0.12 |
| 0.26 |
| 1.00 |
| 0.00 |
| 0.00 |
| 0.00 |
| 0.07 |
| 0.72 |
| 0.97 |
| 0.41 |
| 0.08 |
| 1.00 |
| 0.22 |
| 1.00 |
| 0.00 |
| 0.00 |
| 0.04 |
| 0.89 |
| 0.01 |
| 1.00 |
| 0.23 |
| 0.15 |
| 0.30 |
| 0.04 |
| 1.00 |
| 0.02 |
| 0.02 |
| 0.00 |
| 0.09 |
| 0.25 |
| 0.01 |
| 0.02 |
| 0.07 |
| 0.47 |
| 1.00 |

| |
|---|
| 0.73 |
| 0.07 |
| 0.99 |
| 0.00 |
| 0.00 |
| 0.00 |
| 0.29 |
| 1.00 |
| 1.00 |
| 0.10 |
| 0.07 |
| 0.84 |
| 0.82 |
| 0.43 |
| 0.01 |
| 0.00 |
| 0.00 |
| 0.00 |
| 0.00 |
| 0.30 |
| 0.58 |
| 0.00 |
| 1.00 |
| 0.01 |
| 0.03 |
| 0.01 |
| 0.82 |
| 0.00 |
| 0.01 |
| 0.96 |
| 0.97 |
| 0.23 |
| 0.14 |
| 0.20 |
| 0.00 |
| 1.00 |
| 0.02 |
| 0.34 |
| 0.07 |
| 0.37 |
| 0.63 |
| 1.00 |
| 1.00 |
| 0.00 |
| 0.17 |
| 0.01 |
| 0.00 |
| 0.01 |

| |
|---|
| 0.06 |
| 0.26 |
| 0.31 |
| 0.89 |
| 0.00 |
| 0.74 |
| 1.00 |
| 0.03 |
| 0.00 |
| 0.15 |
| 0.02 |
| 0.00 |
| 0.02 |
| 1.00 |
| 0.01 |
| 0.66 |
| 1.00 |
| 0.80 |
| 0.02 |
| 0.00 |
| 0.14 |
| 0.04 |
| 1.00 |
| 0.00 |
| 0.00 |
| 0.00 |
| 0.00 |
| 1.00 |
| 1.00 |
| 0.93 |
| 1.00 |
| 1.00 |
| 0.01 |
| 0.01 |
| 0.28 |
| 0.22 |
| 1.00 |
| 0.01 |
| 0.02 |
| 1.00 |
| 0.11 |
| 0.16 |
| 0.11 |
| 0.00 |
| 0.11 |
| 0.90 |
| 0.78 |
| 0.16 |

| |
|---|
| 0.24 |
| 0.02 |
| 0.00 |
| 0.23 |
| 0.44 |
| 0.00 |
| 1.00 |
| 1.00 |
| 0.74 |
| 0.05 |
| 0.00 |
| 1.00 |
| 1.00 |
| 1.00 |
| 1.00 |
| 1.00 |
| 1.00 |
| 1.00 |
| 0.00 |
| 1.00 |
| 0.01 |
| 0.56 |
| 0.02 |
| 0.44 |
| 1.00 |
| 0.33 |
| 1.00 |
| 0.13 |
| 0.26 |
| 0.30 |
| 0.70 |
| 0.00 |
| 0.66 |
| 0.00 |
| 0.77 |
| 0.02 |
| 1.00 |
| 1.00 |
| 0.56 |
| 0.97 |
| 1.00 |
| 0.00 |
| 0.44 |
| 0.02 |
| 0.00 |
| 0.20 |
| 0.75 |
| 1.00 |

| |
|---|
| 1.00 |
| 0.89 |
| 1.00 |
| 0.09 |
| 0.11 |
| 1.00 |
| 0.39 |
| 0.00 |
| 0.00 |
| 0.42 |
| 0.00 |
| 0.81 |
| 0.01 |
| 0.41 |
| 0.00 |
| 0.04 |
| 0.04 |
| 1.00 |
| 0.67 |
| 0.00 |
| 0.37 |
| 1.00 |
| 1.00 |
| 0.02 |
| 0.09 |
| 1.00 |
| 0.45 |
| 1.00 |
| 0.00 |
| 0.00 |
| 1.00 |
| 0.30 |
| 0.00 |
| 0.80 |
| 0.76 |
| 1.00 |
| 0.04 |
| 0.52 |
| 1.00 |
| 0.07 |
| 0.11 |
| 0.00 |
| 0.86 |
| 1.00 |
| 0.16 |
| 0.14 |
| 0.00 |
| 0.01 |

| |
|---|
| 0.00 |
| 0.42 |
| 0.68 |
| 0.63 |
| 0.10 |
| 0.60 |
| 1.00 |
| 0.28 |
| 0.07 |
| 0.88 |
| 0.66 |
| 0.41 |
| 0.78 |
| 0.66 |
| 0.01 |
| 1.00 |
| 0.00 |
| 0.42 |
| 1.00 |
| 0.15 |
| 1.00 |
| 1.00 |
| 0.04 |
| 0.45 |
| 0.15 |
| 0.38 |
| 0.00 |
| 1.00 |
| 1.00 |
| 1.00 |
| 1.00 |
| 0.45 |
| 0.38 |
| 1.00 |
| 0.01 |
| 0.02 |
| 0.24 |
| 0.04 |
| 1.00 |
| 0.98 |
| 0.18 |
| 0.03 |
| 0.09 |
| 0.06 |
| 0.00 |
| 0.66 |
| 1.00 |
| 0.00 |

| |
|---|
| 0.35 |
| 0.29 |
| 1.00 |
| 1.00 |
| 0.59 |
| 1.00 |
| 1.00 |
| 0.00 |
| 0.00 |
| 0.21 |
| 1.00 |
| 1.00 |
| 0.40 |
| 1.00 |
| 1.00 |
| 0.00 |
| 0.00 |
| 0.56 |
| 0.00 |
| 0.01 |
| 1.00 |
| 0.00 |
| 0.13 |
| 0.00 |
| 1.00 |
| 0.09 |
| 0.00 |
| 1.00 |
| 1.00 |
| 0.01 |
| 0.87 |
| 1.00 |
| 0.00 |
| 0.52 |
| 0.51 |
| 0.00 |
| 0.14 |
| 0.00 |
| 0.00 |
| 0.91 |
| 0.12 |
| 0.10 |
| 0.15 |
| 0.09 |
| 1.00 |
| 1.00 |
| 0.05 |
| 0.45 |

| |
|---|
| 1.00 |
| 1.00 |
| 0.08 |
| 0.00 |
| 0.03 |
| 0.93 |
| 0.00 |
| 0.00 |
| 0.93 |
| 0.83 |
| 1.00 |
| 0.00 |
| 1.00 |
| 0.00 |
| 0.63 |
| 0.00 |
| 0.24 |
| 1.00 |
| 1.00 |
| 1.00 |
| 0.38 |
| 1.00 |
| 0.92 |
| 0.59 |
| 0.00 |
| 1.00 |
| 0.01 |
| 0.64 |
| 0.29 |
| 0.32 |
| 0.00 |
| 0.10 |
| 1.00 |
| 0.49 |
| 0.28 |
| 0.55 |
| 1.00 |
| 0.02 |
| 0.00 |
| 0.00 |
| 0.42 |
| 0.02 |
| 0.00 |
| 1.00 |
| 1.00 |
| 0.37 |
| 0.00 |
| 0.01 |

| |
|---|
| 1.00 |
| 1.00 |
| 0.50 |
| 1.00 |
| 1.00 |
| 0.70 |
| 0.49 |
| 0.38 |
| 1.00 |
| 0.48 |
| 1.00 |
| 0.97 |
| 1.00 |
| 0.50 |
| 0.09 |
| 0.58 |
| 0.00 |
| 0.43 |
| 0.07 |
| 1.00 |
| 0.00 |
| 0.00 |
| 1.00 |
| 0.93 |
| 0.00 |
| 1.00 |
| 0.01 |
| 0.06 |
| 0.28 |
| 1.00 |
| 0.18 |
| 0.43 |
| 0.01 |
| 0.40 |
| 1.00 |
| 0.29 |
| 0.73 |
| 0.70 |
| 0.85 |
| 0.00 |
| 0.10 |
| 1.00 |
| 0.31 |
| 1.00 |
| 1.00 |
| 0.41 |
| 1.00 |
| 1.00 |

| |
|---|
| 0.24 |
| 1.00 |
| 1.00 |
| 0.46 |
| 0.11 |
| 0.26 |
| 1.00 |
| 0.15 |
| 0.33 |
| 1.00 |
| 1.00 |
| 0.57 |
| 0.01 |
| 0.00 |
| 0.00 |
| 0.31 |
| 1.00 |
| 1.00 |
| 0.01 |
| 1.00 |
| 0.40 |
| 1.00 |
| 0.00 |
| 0.00 |
| 0.21 |
| 0.17 |
| 0.08 |
| 0.00 |
| 1.00 |
| 0.00 |
| 0.00 |
| 0.01 |
| 0.00 |
| 0.83 |
| 0.00 |
| 0.01 |
| 0.39 |
| 0.68 |
| 1.00 |
| 0.00 |
| 1.00 |
| 0.58 |
| 1.00 |
| 0.00 |
| 1.00 |
| 0.00 |
| 0.93 |
| 0.39 |

| |
|---|
| 0.00 |
| 0.00 |
| 0.02 |
| 0.00 |
| 0.00 |
| 0.00 |
| 0.55 |
| 0.00 |
| 0.07 |
| 1.00 |
| 0.24 |
| 0.00 |
| 0.30 |
| 0.01 |
| 0.00 |
| 0.97 |
| 0.54 |
| 1.00 |
| 0.11 |
| 0.46 |
| 0.00 |
| 0.01 |
| 0.32 |
| 0.00 |
| 0.01 |
| 0.01 |
| 1.00 |
| 0.00 |
| 0.00 |
| 0.48 |
| 1.00 |
| 0.31 |
| 0.99 |
| 0.04 |
| 0.69 |
| 0.03 |
| 0.99 |
| 1.00 |
| 1.00 |
| 0.06 |
| 0.04 |
| 0.31 |
| 0.00 |
| 0.03 |
| 1.00 |
| 0.33 |
| 0.35 |
| 0.01 |

| |
|---|
| 0.77 |
| 0.39 |
| 1.00 |
| 1.00 |
| 0.00 |
| 0.04 |
| 0.53 |
| 0.00 |
| 0.00 |
| 0.07 |
| 0.96 |
| 0.00 |
| 0.00 |
| 0.00 |
| 0.68 |
| 0.00 |
| 0.00 |
| 0.09 |
| 0.68 |
| 0.21 |
| 0.88 |
| 0.94 |
| 0.56 |
| 0.00 |
| 0.98 |
| 1.00 |
| 0.34 |
| 0.00 |
| 0.12 |
| 0.00 |
| 0.82 |
| 1.00 |
| 0.01 |
| 0.01 |
| 0.02 |
| 0.00 |
| 0.02 |
| 0.00 |
| 0.01 |
| 0.41 |
| 0.59 |
| 0.00 |
| 0.12 |
| 0.88 |
| 0.00 |
| 0.00 |
| 0.11 |
| 0.70 |

| |
|---|
| 0.01 |
| 0.74 |
| 1.00 |
| 0.10 |
| 0.53 |
| 1.00 |
| 0.00 |
| 0.00 |
| 0.00 |
| 0.04 |
| 1.00 |
| 0.01 |
| 1.00 |
| 0.01 |
| 0.51 |
| 1.00 |
| 0.00 |
| 1.00 |
| 1.00 |
| 0.88 |
| 0.79 |
| 0.10 |
| 0.00 |
| 0.64 |
| 0.14 |
| 0.00 |
| 0.21 |
| 1.00 |
| 0.01 |
| 0.60 |
| 0.68 |
| 0.39 |
| 1.00 |
| 1.00 |
| 0.17 |
| 1.00 |
| 0.32 |
| 1.00 |
| 0.05 |
| 1.00 |
| 0.51 |
| 0.01 |
| 0.00 |
| 0.00 |
| 0.00 |
| 0.69 |
| 0.58 |
| 0.00 |

| |
|---|
| 0.01 |
| 0.03 |
| 0.08 |
| 0.15 |
| 0.69 |
| 0.11 |
| 0.00 |
| 0.83 |
| 1.00 |
| 0.00 |
| 0.00 |
| 0.00 |
| 1.00 |
| 0.00 |
| 0.26 |
| 1.00 |
| 0.00 |
| 0.08 |
| 0.05 |
| 0.01 |
| 1.00 |
| 0.37 |
| 0.56 |
| 1.00 |
| 0.17 |
| 1.00 |
| 0.00 |
| 0.19 |
| 1.00 |
| 0.00 |
| 0.03 |
| 0.41 |
| 0.94 |
| 0.00 |
| 1.00 |
| 0.23 |
| 0.00 |
| 1.00 |
| 0.00 |
| 1.00 |
| 0.01 |
| 0.28 |
| 0.87 |
| 0.69 |
| 0.14 |
| 0.00 |
| 0.00 |
| 0.00 |

| |
|---|
| 0.68 |
| 0.32 |
| 0.71 |
| 0.15 |
| 0.22 |
| 1.00 |
| 0.00 |
| 1.00 |
| 0.00 |
| 0.00 |
| 0.13 |
| 0.79 |
| 1.00 |
| 0.08 |
| 0.41 |
| 0.22 |
| 0.00 |
| 0.09 |
| 0.36 |
| 0.00 |
| 0.02 |
| 1.00 |
| 0.03 |
| 0.02 |
| 0.36 |
| 0.46 |
| 0.21 |
| 0.00 |
| 0.01 |
| 0.04 |
| 0.00 |
| 1.00 |
| 0.00 |
| 0.16 |
| 0.39 |
| 1.00 |
| 0.81 |
| 0.81 |
| 0.00 |
| 1.00 |
| 1.00 |
| 0.03 |
| 0.17 |
| 0.46 |
| 0.19 |
| 0.00 |
| 0.00 |
| 0.24 |

| |
|---|
| 0.11 |
| 0.84 |
| 0.36 |
| 0.30 |
| 1.00 |
| 0.55 |
| 1.00 |
| 0.06 |
| 1.00 |
| 1.00 |
| 0.81 |
| 0.00 |
| 0.00 |
| 0.58 |
| 1.00 |
| 1.00 |
| 0.30 |
| 0.78 |
| 0.76 |
| 0.00 |
| 0.06 |
| 1.00 |
| 0.45 |
| 0.08 |
| 0.72 |
| 0.01 |
| 0.25 |
| 1.00 |
| 0.93 |
| 0.32 |
| 0.12 |
| 1.00 |
| 1.00 |
| 1.00 |
| 1.00 |
| 1.00 |
| 0.00 |
| 0.82 |
| 1.00 |
| 0.90 |
| 1.00 |
| 0.00 |
| 0.00 |
| 0.34 |
| 0.83 |
| 1.00 |
| 0.01 |
| 1.00 |

| |
|---|
| 0.01 |
| 0.53 |
| 0.92 |
| 0.63 |
| 0.00 |
| 0.05 |
| 0.00 |
| 0.82 |
| 0.01 |
| 0.53 |
| 0.00 |
| 0.22 |
| 0.00 |
| 1.00 |
| 0.00 |
| 0.28 |
| 0.78 |
| 0.01 |
| 0.74 |
| 1.00 |
| 0.02 |
| 0.00 |
| 0.00 |
| 1.00 |
| 1.00 |
| 0.00 |
| 1.00 |
| 0.00 |
| 0.00 |
| 1.00 |
| 0.00 |
| 1.00 |
| 0.20 |
| 1.00 |
| 1.00 |
| 0.94 |
| 1.00 |
| 0.59 |
| 0.00 |
| 1.00 |
| 1.00 |
| 0.13 |
| 0.04 |
| 0.64 |
| 1.00 |
| 0.51 |
| 0.75 |
| 0.00 |

| |
|---|
| 1.00 |
| 0.04 |
| 1.00 |
| 0.00 |
| 0.55 |
| 1.00 |
| 0.21 |
| 0.70 |
| 0.00 |
| 0.00 |
| 1.00 |
| 0.97 |
| 0.42 |
| 0.00 |
| 0.38 |
| 0.04 |
| 1.00 |
| 1.00 |
| 0.00 |
| 1.00 |
| 1.00 |
| 0.00 |
| 1.00 |
| 0.36 |
| 1.00 |
| 0.04 |
| 0.00 |
| 1.00 |
| 0.13 |
| 0.00 |
| 0.89 |
| 1.00 |
| 0.00 |
| 0.24 |
| 0.20 |
| 0.00 |
| 0.30 |
| 1.00 |
| 0.05 |
| 0.00 |
| 0.00 |
| 0.00 |
| 0.69 |
| 0.58 |
| 1.00 |
| 1.00 |
| 0.00 |
| 0.88 |

| |
|---|
| 1.00 |
| 0.00 |
| 0.00 |
| 0.01 |
| 0.00 |
| 1.00 |
| 1.00 |
| 0.05 |
| 0.28 |
| 1.00 |
| 0.00 |
| 0.00 |
| 0.66 |
| 0.35 |
| 0.34 |
| 0.19 |
| 1.00 |
| 0.24 |
| 0.05 |
| 1.00 |
| 0.00 |
| 0.99 |
| 0.64 |
| 0.39 |
| 0.04 |
| 1.00 |
| 1.00 |
| 0.74 |
| 0.00 |
| 0.00 |
| 0.34 |
| 0.66 |
| 0.49 |
| 1.00 |
| 0.04 |
| 1.00 |
| 1.00 |
| 0.01 |
| 0.00 |
| 0.00 |
| 0.93 |
| 0.76 |
| 0.38 |
| 0.00 |
| 1.00 |
| 0.03 |
| 0.77 |
| 0.20 |

| |
|---|
| 0.20 |
| 0.19 |
| 0.95 |
| 0.40 |
| 1.00 |
| 0.00 |
| 1.00 |
| 0.52 |
| 0.00 |
| 0.00 |
| 0.00 |
| 0.00 |
| 0.00 |
| 0.00 |
| 0.00 |
| 1.00 |
| 0.00 |
| 0.03 |
| 0.00 |
| 1.00 |
| 0.00 |
| 0.54 |
| 1.00 |
| 0.28 |
| 0.37 |
| 0.37 |
| 1.00 |
| 0.67 |
| 1.00 |
| 1.00 |
| 0.27 |
| 1.00 |
| 1.00 |
| 0.00 |
| 0.70 |
| 0.41 |
| 0.16 |
| 1.00 |
| 0.99 |
| 1.00 |
| 0.00 |
| 1.00 |
| 0.00 |
| 0.29 |
| 1.00 |
| 0.29 |
| 1.00 |
| 0.00 |

| |
|---|
| 1.00 |
| 0.00 |
| 1.00 |
| 1.00 |
| 0.00 |
| 0.00 |
| 0.00 |
| 1.00 |
| 0.00 |
| 1.00 |
| 0.21 |
| 1.00 |
| 0.14 |
| 0.00 |
| 0.04 |
| 0.01 |
| 0.65 |
| 0.18 |
| 0.92 |
| 0.01 |
| 0.05 |
| 0.00 |
| 0.00 |
| 0.30 |
| 0.00 |
| 0.11 |
| 0.11 |
| 0.00 |
| 0.17 |
| 0.02 |
| 0.00 |
| 0.00 |
| 1.00 |
| 0.20 |
| 0.14 |
| 0.92 |
| 0.00 |
| 0.00 |
| 0.00 |
| 0.00 |
| 0.07 |
| 1.00 |
| 0.42 |
| 0.41 |
| 0.66 |
| 1.00 |
| 0.01 |
| 0.00 |

| |
|---|
| 0.58 |
| 0.58 |
| 1.00 |
| 0.03 |
| 0.00 |
| 0.00 |
| 0.66 |
| 0.00 |
| 0.00 |
| 0.13 |
| 0.09 |
| 0.00 |
| 0.77 |
| 0.01 |
| 0.05 |
| 0.53 |
| 0.44 |
| 0.00 |
| 0.26 |
| 0.67 |
| 1.00 |
| 1.00 |
| 0.05 |
| 1.00 |
| 1.00 |
| 0.09 |
| 0.57 |
| 1.00 |
| 1.00 |
| 0.00 |
| 0.00 |
| 0.00 |
| 0.00 |
| 0.07 |
| 0.00 |
| 0.03 |
| 1.00 |
| 1.00 |
| 0.00 |
| 0.01 |
| 0.00 |
| 0.91 |
| 0.97 |
| 0.17 |
| 0.60 |
| 0.00 |
| 0.40 |
| 0.00 |

| |
|---|
| 0.22 |
| 0.00 |
| 1.00 |
| 0.56 |
| 0.01 |
| 1.00 |
| 0.21 |
| 0.21 |
| 0.00 |
| 0.88 |
| 0.00 |
| 0.00 |
| 0.01 |
| 1.00 |
| 0.00 |
| 0.73 |
| 0.19 |
| 1.00 |
| 0.83 |
| 1.00 |
| 0.00 |
| 0.61 |
| 0.00 |
| 1.00 |
| 1.00 |
| 0.01 |
| 0.00 |
| 0.02 |
| 0.08 |
| 0.00 |
| 0.97 |
| 0.02 |
| 1.00 |
| 0.00 |
| 0.75 |
| 0.95 |
| 0.00 |
| 0.10 |
| 0.42 |
| 0.02 |
| 0.12 |
| 0.00 |
| 0.03 |
| 0.00 |
| 0.65 |
| 0.08 |
| 0.00 |
| 0.00 |

| |
|---|
| 1.00 |
| 0.09 |
| 0.13 |
| 0.90 |
| 0.49 |
| 0.04 |
| 0.21 |
| 0.83 |
| 0.03 |
| 0.13 |
| 0.08 |
| 0.00 |
| 1.00 |
| 1.00 |
| 0.43 |
| 0.00 |
| 1.00 |
| 1.00 |
| 0.64 |
| 0.51 |
| 0.00 |
| 0.00 |
| 0.00 |
| 1.00 |
| 1.00 |
| 0.00 |
| 0.09 |
| 0.06 |
| 0.40 |
| 0.00 |
| 0.00 |
| 1.00 |
| 1.00 |
| 0.71 |
| 1.00 |
| 0.23 |
| 0.76 |
| 0.00 |
| 1.00 |
| 1.00 |
| 0.91 |
| 0.00 |
| 0.02 |
| 1.00 |
| 0.00 |
| 1.00 |
| 0.00 |
| 1.00 |

| |
|---|
| 0.02 |
| 0.09 |
| 1.00 |
| 0.52 |
| 0.22 |
| 0.00 |
| 0.12 |
| 0.13 |
| 0.00 |
| 0.97 |
| 0.00 |
| 0.60 |
| 1.00 |
| 0.00 |
| 1.00 |
| 1.00 |
| 1.00 |
| 0.00 |
| 0.72 |
| 0.06 |
| 0.00 |
| 0.00 |
| 0.01 |
| 1.00 |
| 0.00 |
| 0.02 |
| 1.00 |
| 0.00 |
| 0.01 |
| 1.00 |
| 0.61 |
| 0.03 |
| 0.01 |
| 1.00 |
| 1.00 |
| 0.27 |
| 0.00 |
| 0.00 |
| 0.41 |
| 0.58 |
| 0.59 |
| 1.00 |
| 1.00 |
| 0.17 |
| 0.00 |
| 1.00 |
| 0.40 |
| 0.00 |

| |
|---|
| 1.00 |
| 1.00 |
| 0.17 |
| 0.00 |
| 0.00 |
| 0.00 |
| 0.84 |
| 0.47 |
| 0.78 |
| 0.15 |
| 1.00 |
| 0.01 |
| 0.09 |
| 0.08 |
| 0.80 |
| 0.29 |
| 0.00 |
| 0.00 |
| 0.00 |
| 0.14 |
| 0.00 |
| 0.01 |
| 0.86 |
| 0.00 |
| 1.00 |
| 1.00 |
| 1.00 |
| 1.00 |
| 0.86 |
| 1.00 |
| 0.00 |
| 0.00 |
| 0.00 |
| 0.00 |
| 0.00 |
| 0.00 |
| 1.00 |
| 0.73 |
| 0.97 |
| 0.08 |
| 0.70 |
| 0.88 |
| 1.00 |
| 0.39 |
| 0.00 |
| 1.00 |
| 1.00 |
| 0.37 |

| |
|---|
| 0.00 |
| 0.13 |
| 0.25 |
| 0.00 |
| 0.02 |
| 0.05 |
| 0.00 |
| 0.03 |
| 0.08 |
| 0.96 |
| 0.33 |
| 0.00 |
| 0.08 |
| 1.00 |
| 0.06 |
| 1.00 |
| 1.00 |
| 1.00 |
| 0.29 |
| 0.88 |
| 0.20 |
| 0.00 |
| 0.04 |
| 0.00 |
| 0.02 |
| 0.00 |
| 0.65 |
| 1.00 |
| 0.80 |
| 1.00 |
| 0.00 |
| 0.00 |
| 0.87 |
| 0.01 |
| 1.00 |
| 0.36 |
| 1.00 |
| 0.51 |
| 0.27 |
| 1.00 |
| 1.00 |
| 0.30 |
| 0.24 |
| 0.39 |
| 0.02 |
| 1.00 |
| 0.02 |
| 0.95 |

| |
|---|
| 0.75 |
| 1.00 |
| 1.00 |
| 0.05 |
| 0.10 |
| 0.46 |
| 0.47 |
| 1.00 |
| 0.01 |
| 0.09 |
| 0.04 |
| 1.00 |
| 0.00 |
| 0.58 |
| 0.00 |
| 0.91 |
| 0.00 |
| 1.00 |
| 0.31 |
| 0.06 |
| 0.77 |
| 0.16 |
| 0.68 |
| 0.14 |
| 0.06 |
| 1.00 |
| 0.71 |
| 1.00 |
| 0.10 |
| 0.00 |
| 1.00 |
| 1.00 |
| 1.00 |
| 0.89 |
| 0.08 |
| 1.00 |
| 0.16 |
| 0.66 |
| 0.00 |
| 0.95 |
| 0.97 |
| 0.91 |
| 1.00 |
| 0.00 |
| 1.00 |
| 0.82 |
| 0.63 |
| 1.00 |

| |
|---|
| 0.90 |
| 0.00 |
| 0.00 |
| 0.47 |
| 0.16 |
| 0.00 |
| 0.00 |
| 0.92 |
| 0.07 |
| 0.39 |
| 1.00 |
| 0.00 |
| 0.33 |
| 0.00 |
| 1.00 |
| 1.00 |
| 0.00 |
| 0.07 |
| 0.00 |
| 0.16 |
| 0.00 |
| 0.00 |
| 0.16 |
| 1.00 |
| 0.00 |
| 0.02 |
| 0.02 |
| 0.00 |
| 0.00 |
| 0.26 |
| 0.00 |
| 0.00 |
| 0.01 |
| 0.00 |
| 0.00 |
| 0.00 |
| 0.00 |
| 0.00 |
| 0.00 |
| 0.00 |
| 0.00 |
| 0.07 |
| 0.05 |
| 0.34 |
| 0.01 |
| 0.11 |
| 0.00 |
| 0.00 |

| |
|---|
| 0.00 |
| 0.01 |
| 0.00 |
| 0.07 |
| 1.00 |
| 0.00 |
| 0.00 |
| 0.00 |
| 0.03 |
| 0.00 |
| 1.00 |
| 0.09 |
| 0.78 |
| 0.68 |
| 1.00 |
| 0.00 |
| 0.23 |
| 0.36 |
| 0.01 |
| 0.03 |
| 0.10 |
| 0.00 |
| 0.32 |
| 0.11 |
| 1.00 |
| 0.02 |
| 0.79 |
| 0.23 |
| 0.06 |
| 0.18 |
| 1.00 |
| 1.00 |
| 0.09 |
| 1.00 |
| 0.23 |
| 0.93 |
| 1.00 |
| 1.00 |
| 0.02 |
| 0.38 |
| 1.00 |
| 0.00 |
| 1.00 |
| 0.00 |
| 0.36 |
| 0.00 |
| 0.90 |
| 0.55 |

| |
|---|
| 0.00 |
| 0.27 |
| 0.01 |
| 1.00 |
| 0.18 |
| 1.00 |
| 0.00 |
| 1.00 |
| 0.42 |
| 0.04 |
| 0.00 |
| 1.00 |
| 1.00 |
| 0.00 |
| 0.00 |
| 0.01 |
| 1.00 |
| 1.00 |
| 1.00 |
| 0.00 |
| 0.00 |
| 0.84 |
| 0.53 |
| 0.07 |
| 0.67 |
| 0.05 |
| 0.00 |
| 1.00 |
| 0.06 |
| 0.04 |
| 0.01 |
| 1.00 |
| 1.00 |
| 0.01 |
| 0.00 |
| 1.00 |
| 0.02 |
| 0.04 |
| 1.00 |
| 0.18 |
| 0.11 |
| 0.25 |
| 0.39 |
| 0.00 |
| 0.04 |
| 0.89 |
| 0.10 |
| 0.06 |

| |
|---|
| 0.09 |
| 1.00 |
| 0.00 |
| 0.42 |
| 0.61 |
| 0.00 |
| 1.00 |
| 0.73 |
| 0.00 |
| 0.42 |
| 0.00 |
| 0.00 |
| 0.69 |
| 0.20 |
| 0.03 |
| 0.14 |
| 0.00 |
| 0.00 |
| 0.69 |
| 0.00 |
| 0.21 |
| 1.00 |
| 0.12 |
| 0.79 |
| 0.05 |
| 0.62 |
| 1.00 |
| 1.00 |
| 0.01 |
| 0.72 |
| 0.00 |
| 0.61 |
| 0.00 |
| 1.00 |
| 0.28 |
| 0.03 |
| 0.36 |
| 0.30 |
| 1.00 |
| 0.82 |
| 1.00 |
| 0.20 |
| 1.00 |
| 0.27 |
| 0.22 |
| 0.00 |
| 1.00 |
| 1.00 |

| |
|---|
| 0.00 |
| 1.00 |
| 0.00 |
| 0.02 |
| 1.00 |
| 0.00 |
| 0.43 |
| 1.00 |
| 0.63 |
| 0.25 |
| 0.03 |
| 0.01 |
| 0.00 |
| 0.97 |
| 0.63 |
| 0.24 |
| 0.01 |
| 0.00 |
| 0.41 |
| 0.40 |
| 1.00 |
| 0.00 |
| 1.00 |
| 0.13 |
| 0.01 |
| 0.00 |
| 1.00 |
| 0.00 |
| 0.13 |
| 0.02 |
| 0.00 |
| 0.01 |
| 0.02 |
| 0.08 |
| 0.00 |
| 0.85 |
| 1.00 |
| 0.11 |
| 0.92 |
| 0.98 |
| 0.03 |
| 0.00 |
| 1.00 |
| 0.64 |
| 1.00 |
| 0.63 |
| 1.00 |
| 0.00 |

| |
|---|
| 0.27 |
| 0.13 |
| 0.11 |
| 0.60 |
| 0.09 |
| 0.04 |
| 1.00 |
| 1.00 |
| 0.66 |
| 0.01 |
| 0.94 |
| 1.00 |
| 0.08 |
| 1.00 |
| 0.00 |
| 0.44 |
| 0.25 |
| 0.01 |
| 1.00 |
| 0.14 |
| 1.00 |
| 1.00 |
| 0.25 |
| 1.00 |
| 1.00 |
| 0.36 |
| 0.18 |
| 1.00 |
| 0.29 |
| 0.68 |
| 0.00 |
| 0.59 |
| 0.59 |
| 1.00 |
| 0.77 |
| 0.75 |
| 0.00 |
| 0.78 |
| 1.00 |
| 0.92 |
| 0.07 |
| 0.21 |
| 0.03 |
| 0.37 |
| 0.05 |
| 0.61 |
| 1.00 |
| 0.21 |

| |
|---|
| 0.01 |
| 0.15 |
| 0.00 |
| 0.56 |
| 0.01 |
| 0.03 |
| 0.11 |
| 0.29 |
| 0.44 |
| 0.47 |
| 0.09 |
| 0.49 |
| 1.00 |
| 0.11 |
| 0.43 |
| 0.28 |
| 1.00 |
| 1.00 |
| 0.17 |
| 0.00 |
| 0.73 |
| 0.02 |
| 0.00 |
| 0.04 |
| 1.00 |
| 0.72 |
| 0.00 |
| 0.85 |
| 0.16 |
| 0.00 |
| 0.01 |
| 0.24 |
| 0.00 |
| 0.69 |
| 0.00 |
| 1.00 |
| 0.05 |
| 0.21 |
| 0.05 |
| 0.00 |
| 0.24 |
| 0.00 |
| 1.00 |
| 1.00 |
| 0.17 |
| 0.08 |
| 0.95 |
| 0.78 |

| |
|---|
| 0.82 |
| 0.31 |
| 0.00 |
| 0.51 |
| 0.07 |
| 0.17 |
| 1.00 |
| 0.97 |
| 0.76 |
| 1.00 |
| 0.20 |
| 0.02 |
| 1.00 |
| 0.00 |
| 0.02 |
| 1.00 |
| 1.00 |
| 1.00 |
| 0.00 |
| 0.00 |
| 1.00 |
| 1.00 |
| 0.90 |
| 1.00 |
| 0.35 |
| 0.63 |
| 1.00 |
| 0.67 |
| 0.04 |
| 0.01 |
| 0.00 |
| 0.38 |
| 0.29 |
| 0.00 |
| 0.04 |
| 0.25 |
| 0.33 |
| 0.00 |
| 0.34 |
| 0.88 |
| 0.03 |
| 0.01 |
| 0.88 |
| 0.00 |
| 0.04 |
| 0.00 |
| 1.00 |
| 0.09 |

| |
|---|
| 0.00 |
| 0.00 |
| 1.00 |
| 0.02 |
| 0.23 |
| 0.00 |
| 0.00 |
| 0.00 |
| 1.00 |
| 0.66 |
| 1.00 |
| 1.00 |
| 0.74 |
| 1.00 |
| 1.00 |
| 0.00 |
| 0.00 |
| 1.00 |
| 0.40 |
| 0.37 |
| 1.00 |
| 0.38 |
| 0.23 |
| 0.22 |
| 0.56 |
| 0.66 |
| 0.99 |
| 0.01 |
| 0.61 |
| 1.00 |
| 0.50 |
| 0.04 |
| 1.00 |
| 0.74 |
| 0.01 |
| 0.40 |
| 0.07 |
| 0.18 |
| 0.01 |
| 0.00 |
| 0.00 |
| 0.68 |
| 1.00 |
| 1.00 |
| 0.73 |
| 0.87 |
| 0.00 |
| 0.32 |

| |
|---|
| 1.00 |
| 1.00 |
| 0.60 |
| 0.01 |
| 1.00 |
| 1.00 |
| 0.21 |
| 0.27 |
| 1.00 |
| 1.00 |
| 0.91 |
| 0.00 |
| 0.82 |
| 1.00 |
| 0.45 |
| 1.00 |
| 0.05 |
| 0.13 |
| 0.00 |
| 0.00 |
| 0.25 |
| 0.50 |
| 0.18 |
| 0.44 |
| 0.95 |
| 0.16 |
| 0.01 |
| 0.01 |
| 0.01 |
| 1.00 |
| 0.48 |
| 0.91 |
| 0.07 |
| 0.73 |
| 1.00 |
| 0.00 |
| 0.08 |
| 0.00 |
| 1.00 |
| 1.00 |
| 0.00 |
| 1.00 |
| 0.01 |
| 0.94 |
| 0.00 |
| 1.00 |
| 0.27 |
| 0.10 |

| |
|---|
| 1.00 |
| 1.00 |
| 0.00 |
| 1.00 |
| 1.00 |
| 0.48 |
| 0.88 |
| 0.00 |
| 0.00 |
| 1.00 |
| 1.00 |
| 0.41 |
| 0.07 |
| 0.40 |
| 0.03 |
| 0.00 |
| 1.00 |
| 1.00 |
| 0.62 |
| 1.00 |
| 0.12 |
| 1.00 |
| 0.34 |
| 1.00 |
| 0.00 |
| 0.02 |
| 1.00 |
| 0.23 |
| 0.01 |
| 0.04 |
| 0.60 |
| 1.00 |
| 1.00 |
| 0.01 |
| 0.02 |
| 1.00 |
| 0.91 |
| 0.06 |
| 1.00 |
| 0.00 |
| 0.67 |
| 0.00 |
| 0.00 |
| 0.02 |
| 0.62 |
| 0.00 |
| 0.06 |
| 0.91 |

| |
|---|
| 0.36 |
| 0.00 |
| 0.22 |
| 0.01 |
| 0.01 |
| 0.29 |
| 0.01 |
| 1.00 |
| 0.00 |
| 1.00 |
| 1.00 |
| 0.06 |
| 0.01 |
| 0.00 |
| 0.00 |
| 0.00 |
| 0.00 |
| 1.00 |
| 0.12 |
| 0.01 |
| 0.04 |
| 0.36 |
| 1.00 |
| 0.64 |
| 0.19 |
| 0.05 |
| 0.00 |
| 0.02 |
| 0.08 |
| 0.74 |
| 0.62 |
| 0.00 |
| 1.00 |
| 0.02 |
| 0.54 |
| 0.98 |
| 0.90 |
| 1.00 |
| 0.00 |
| 1.00 |
| 1.00 |
| 0.00 |
| 0.24 |
| 0.56 |
| 0.00 |
| 0.39 |
| 0.00 |
| 0.00 |

| |
|---|
| 0.08 |
| 1.00 |
| 0.19 |
| 0.01 |
| 1.00 |
| 0.07 |
| 1.00 |
| 0.00 |
| 0.00 |
| 0.77 |
| 0.03 |
| 1.00 |
| 0.78 |
| 0.29 |
| 0.15 |
| 0.00 |
| 0.00 |
| 0.00 |
| 0.81 |
| 0.03 |
| 0.13 |
| 0.00 |
| 0.00 |
| 0.00 |
| 0.40 |
| 0.27 |
| 0.63 |
| 0.04 |
| 0.26 |
| 1.00 |
| 0.56 |
| 0.19 |
| 1.00 |
| 0.66 |
| 0.29 |
| 1.00 |
| 0.99 |
| 0.23 |
| 0.97 |
| 0.10 |
| 0.00 |
| 1.00 |
| 0.66 |
| 0.00 |
| 0.86 |
| 0.00 |
| 0.52 |
| 1.00 |

| |
|---|
| 0.91 |
| 0.72 |
| 1.00 |
| 1.00 |
| 0.00 |
| 0.00 |
| 0.00 |
| 1.00 |
| 0.54 |
| 0.00 |
| 1.00 |
| 1.00 |
| 0.64 |
| 1.00 |
| 0.58 |
| 1.00 |
| 0.39 |
| 0.42 |
| 0.97 |
| 1.00 |
| 0.04 |
| 0.23 |
| 0.01 |
| 0.00 |
| 0.01 |
| 1.00 |
| 0.04 |
| 0.00 |
| 0.52 |
| 0.34 |
| 0.94 |
| 0.35 |
| 0.42 |
| 1.00 |
| 0.11 |
| 0.93 |
| 0.03 |
| 0.00 |
| 0.05 |
| 1.00 |
| 0.02 |
| 1.00 |
| 0.12 |
| 0.19 |
| 0.92 |
| 0.00 |
| 0.01 |
| 1.00 |

| |
|---|
| 0.02 |
| 0.14 |
| 1.00 |
| 0.83 |
| 0.01 |
| 0.01 |
| 0.01 |
| 1.00 |
| 0.44 |
| 0.00 |
| 0.00 |
| 0.66 |
| 0.08 |
| 0.70 |
| 1.00 |
| 0.11 |
| 0.00 |
| 0.39 |
| 0.53 |
| 0.03 |
| 0.06 |
| 0.33 |
| 1.00 |
| 0.30 |
| 1.00 |
| 1.00 |
| 1.00 |
| 0.11 |
| 1.00 |
| 0.13 |
| 1.00 |
| 0.49 |
| 0.06 |
| 0.72 |
| 0.97 |
| 0.91 |
| 0.42 |
| 0.27 |
| 1.00 |
| 0.30 |
| 0.00 |
| 0.22 |
| 0.46 |
| 1.00 |
| 0.00 |
| 0.64 |
| 0.01 |
| 0.01 |

| |
|---|
| 0.90 |
| 0.53 |
| 1.00 |
| 0.54 |
| 0.42 |
| 0.00 |
| 0.02 |
| 0.84 |
| 0.90 |
| 0.33 |
| 0.33 |
| 0.73 |
| 1.00 |
| 0.31 |
| 0.03 |
| 0.85 |
| 0.71 |
| 1.00 |
| 0.45 |
| 0.52 |
| 1.00 |
| 0.24 |
| 1.00 |
| 0.70 |
| 1.00 |
| 0.68 |
| 0.01 |
| 0.54 |
| 0.30 |
| 0.17 |
| 0.46 |
| 0.00 |
| 0.91 |
| 1.00 |
| 0.03 |
| 1.00 |
| 0.55 |
| 1.00 |
| 0.18 |
| 0.74 |
| 1.00 |
| 1.00 |
| 0.18 |
| 0.92 |
| 0.08 |
| 0.30 |
| 0.34 |
| 0.02 |

| |
|---|
| 1.00 |
| 1.00 |
| 0.00 |
| 1.00 |
| 0.12 |
| 0.67 |
| 0.45 |
| 0.03 |
| 0.00 |
| 0.03 |
| 0.00 |
| 0.04 |
| 1.00 |
| 0.00 |
| 1.00 |
| 0.22 |
| 0.91 |
| 0.85 |
| 0.76 |
| 0.00 |
| 0.00 |
| 0.00 |
| 0.11 |
| 0.31 |
| 0.00 |
| 0.23 |
| 0.00 |
| 0.00 |
| 0.00 |
| 0.00 |
| 0.03 |
| 1.00 |
| 0.13 |
| 1.00 |
| 0.60 |
| 0.03 |
| 0.13 |
| 0.11 |
| 0.08 |
| 0.00 |
| 0.01 |
| 0.37 |
| 0.15 |
| 0.77 |
| 0.00 |
| 0.12 |
| 0.16 |
| 0.43 |

| |
|---|
| 0.70 |
| 1.00 |
| 0.73 |
| 0.34 |
| 0.00 |
| 0.00 |
| 0.00 |
| 0.01 |
| 0.13 |
| 1.00 |
| 0.07 |
| 0.18 |
| 0.03 |
| 0.09 |
| 0.01 |
| 1.00 |
| 0.02 |
| 0.23 |
| 0.00 |
| 0.00 |
| 0.00 |
| 0.71 |
| 1.00 |
| 0.02 |
| 0.05 |
| 0.00 |
| 0.18 |
| 0.00 |
| 0.07 |
| 0.12 |
| 0.00 |
| 0.00 |
| 0.00 |
| 0.02 |
| 0.05 |
| 0.00 |
| 0.00 |
| 0.01 |
| 0.34 |
| 0.03 |
| 0.00 |
| 0.96 |
| 0.99 |
| 1.00 |
| 0.03 |
| 0.64 |
| 0.09 |
| 0.02 |

| |
|---|
| 0.03 |
| 1.00 |
| 0.07 |
| 0.18 |
| 0.08 |
| 1.00 |
| 0.00 |
| 1.00 |
| 0.18 |
| 0.02 |
| 0.06 |
| 0.01 |
| 1.00 |
| 0.06 |
| 0.09 |
| 0.37 |
| 0.17 |
| 0.93 |
| 0.11 |
| 1.00 |
| 0.08 |
| 0.12 |
| 0.00 |
| 0.02 |
| 0.57 |
| 0.00 |
| 0.00 |
| 0.05 |
| 0.55 |
| 0.39 |
| 0.01 |
| 0.27 |
| 0.02 |
| 0.15 |
| 0.88 |
| 0.01 |
| 0.00 |
| 0.00 |
| 1.00 |
| 0.00 |
| 1.00 |
| 0.27 |
| 1.00 |
| 0.07 |
| 0.07 |
| 1.00 |
| 0.16 |
| 0.54 |

| |
|---|
| 1.00 |
| 0.02 |
| 0.85 |
| 0.00 |
| 0.01 |
| 0.09 |
| 0.06 |
| 0.00 |
| 0.08 |
| 1.00 |
| 0.63 |
| 1.00 |
| 1.00 |
| 0.46 |
| 0.93 |
| 0.00 |
| 0.94 |
| 0.02 |
| 0.06 |
| 0.19 |
| 1.00 |
| 1.00 |
| 0.00 |
| 0.00 |
| 0.00 |
| 1.00 |
| 0.00 |
| 0.00 |
| 0.01 |
| 0.25 |
| 0.43 |
| 0.21 |
| 1.00 |
| 0.09 |
| 0.32 |
| 0.39 |
| 1.00 |
| 1.00 |
| 0.23 |
| 0.27 |
| 0.02 |
| 1.00 |
| 0.82 |
| 0.06 |
| 1.00 |
| 1.00 |
| 1.00 |
| 1.00 |

| |
|---|
| 0.00 |
| 1.00 |
| 1.00 |
| 0.01 |
| 0.00 |
| 0.02 |
| 1.00 |
| 0.61 |
| 0.00 |
| 0.12 |
| 0.40 |
| 0.01 |
| 0.01 |
| 0.00 |
| 0.98 |
| 0.02 |
| 1.00 |
| 0.07 |
| 1.00 |
| 0.04 |
| 0.87 |
| 0.05 |
| 0.00 |
| 0.00 |
| 1.00 |
| 0.04 |
| 0.03 |
| 0.65 |
| 1.00 |
| 0.68 |
| 0.25 |
| 0.00 |
| 0.00 |
| 0.04 |
| 1.00 |
| 0.28 |
| 1.00 |
| 1.00 |
| 0.00 |
| 0.00 |
| 0.09 |
| 0.03 |
| 0.00 |
| 0.01 |
| 0.00 |
| 0.04 |
| 0.57 |
| 0.00 |

| |
|---|
| 0.54 |
| 0.02 |
| 0.54 |
| 0.76 |
| 1.00 |
| 1.00 |
| 0.00 |
| 0.22 |
| 0.12 |
| 0.21 |
| 0.06 |
| 1.00 |
| 0.02 |
| 0.19 |
| 0.68 |
| 1.00 |
| 1.00 |
| 0.00 |
| 0.75 |
| 0.05 |
| 1.00 |
| 0.09 |
| 0.02 |
| 0.00 |
| 0.00 |
| 0.22 |
| 0.00 |
| 0.00 |
| 0.54 |
| 1.00 |
| 0.39 |
| 0.98 |
| 0.64 |
| 0.01 |
| 1.00 |
| 1.00 |
| 0.72 |
| 0.10 |
| 0.00 |
| 0.99 |
| 0.35 |
| 1.00 |
| 0.00 |
| 1.00 |
| 0.10 |
| 1.00 |
| 1.00 |
| 0.09 |

| |
|---|
| 0.00 |
| 0.00 |
| 0.44 |
| 0.67 |
| 0.31 |
| 0.70 |
| 0.14 |
| 0.30 |
| 0.18 |
| 0.39 |
| 1.00 |
| 0.27 |
| 0.63 |
| 0.65 |
| 0.05 |
| 0.92 |
| 0.00 |
| 0.06 |
| 0.00 |
| 0.30 |
| 0.74 |
| 0.77 |
| 0.02 |
| 0.57 |
| 1.00 |
| 1.00 |
| 1.00 |
| 0.13 |
| 0.00 |
| 0.04 |
| 0.49 |
| 1.00 |
| 0.46 |
| 0.08 |
| 0.12 |
| 0.01 |
| 0.02 |
| 0.21 |
| 0.46 |
| 1.00 |
| 0.21 |
| 0.03 |
| 0.49 |
| 0.11 |
| 0.16 |
| 0.01 |
| 0.00 |
| 1.00 |

| |
|---|
| 0.00 |
| 0.93 |
| 0.01 |
| 1.00 |
| 0.42 |
| 0.00 |
| 1.00 |
| 1.00 |
| 1.00 |
| 0.89 |
| 1.00 |
| 1.00 |
| 0.42 |
| 0.21 |
| 1.00 |
| 1.00 |
| 1.00 |
| 1.00 |
| 1.00 |
| 1.00 |
| 0.00 |
| 0.01 |
| 0.71 |
| 1.00 |
| 0.16 |
| 1.00 |
| 0.92 |
| 0.46 |
| 0.54 |
| 1.00 |
| 1.00 |
| 0.00 |
| 0.36 |
| 1.00 |
| 0.51 |
| 0.25 |
| 1.00 |
| 0.00 |
| 0.05 |
| 0.00 |
| 0.17 |
| 0.64 |
| 1.00 |
| 0.00 |
| 0.78 |
| 0.39 |
| 0.01 |
| 0.01 |

| |
|---|
| 0.03 |
| 1.00 |
| 0.63 |
| 0.34 |
| 1.00 |
| 0.35 |
| 1.00 |
| 0.27 |
| 1.00 |
| 1.00 |
| 0.19 |
| 1.00 |
| 0.18 |
| 0.00 |
| 0.00 |
| 0.00 |
| 0.00 |
| 1.00 |
| 0.10 |
| 0.29 |
| 0.50 |
| 0.37 |
| 1.00 |
| 0.04 |
| 1.00 |
| 0.39 |
| 0.93 |
| 0.41 |
| 0.14 |
| 0.01 |
| 0.00 |
| 1.00 |
| 0.48 |
| 0.00 |
| 0.61 |
| 0.00 |
| 0.84 |
| 1.00 |
| 1.00 |
| 0.01 |
| 0.00 |
| 1.00 |
| 0.00 |
| 1.00 |
| 0.67 |
| 0.00 |
| 0.72 |
| 0.51 |

| |
|---|
| 0.00 |
| 0.48 |
| 1.00 |
| 1.00 |
| 0.76 |
| 0.00 |
| 0.35 |
| 0.68 |
| 0.00 |
| 0.00 |
| 0.00 |
| 0.00 |
| 0.00 |
| 0.54 |
| 0.50 |
| 0.01 |
| 0.24 |
| 0.26 |
| 0.09 |
| 0.45 |
| 1.00 |
| 0.00 |
| 0.00 |
| 0.00 |
| 0.10 |
| 0.03 |
| 1.00 |
| 0.82 |
| 0.14 |
| 0.00 |
| 0.00 |
| 0.00 |
| 0.00 |
| 0.59 |
| 0.00 |
| 0.00 |
| 0.00 |
| 0.00 |
| 0.28 |
| 0.70 |
| 0.63 |
| 0.00 |
| 0.84 |
| 1.00 |
| 0.00 |
| 0.77 |
| 0.14 |
| 0.00 |

| |
|---|
| 0.64 |
| 0.66 |
| 0.12 |
| 1.00 |
| 0.02 |
| 1.00 |
| 0.44 |
| 1.00 |
| 0.93 |
| 1.00 |
| 0.00 |
| 0.00 |
| 0.00 |
| 0.41 |
| 0.27 |
| 0.13 |
| 0.13 |
| 0.01 |
| 0.02 |
| 0.01 |
| 0.85 |
| 0.63 |
| 1.00 |
| 1.00 |
| 0.00 |
| 1.00 |
| 0.01 |
| 0.60 |
| 0.00 |
| 0.48 |
| 0.52 |
| 0.74 |
| 0.00 |
| 0.04 |
| 1.00 |
| 0.01 |
| 0.07 |
| 1.00 |
| 0.02 |
| 0.00 |
| 0.00 |
| 1.00 |
| 0.00 |
| 0.51 |
| 0.02 |
| 0.00 |
| 1.00 |
| 0.93 |

| |
|---|
| 0.00 |
| 0.07 |
| 0.00 |
| 0.00 |
| 1.00 |
| 1.00 |
| 0.06 |
| 0.60 |
| 0.14 |
| 0.01 |
| 1.00 |
| 1.00 |
| 0.01 |
| 1.00 |
| 0.00 |
| 0.00 |
| 0.00 |
| 0.00 |
| 0.00 |
| 0.05 |
| 0.19 |
| 0.00 |
| 0.88 |
| 1.00 |
| 0.71 |
| 0.06 |
| 0.06 |
| 0.45 |
| 1.00 |
| 1.00 |
| 0.58 |
| 0.00 |
| 1.00 |
| 0.95 |
| 1.00 |
| 0.00 |
| 0.68 |
| 0.31 |
| 0.30 |
| 0.52 |
| 0.05 |
| 0.00 |
| 1.00 |
| 1.00 |
| 0.29 |
| 0.00 |
| 0.04 |
| 0.17 |

| |
|---|
| 0.02 |
| 0.00 |
| 0.82 |
| 0.00 |
| 0.09 |
| 0.01 |
| 1.00 |
| 0.00 |
| 0.00 |
| 0.19 |
| 1.00 |
| 0.24 |
| 0.54 |
| 0.36 |
| 0.00 |
| 1.00 |
| 0.00 |
| 0.40 |
| 0.00 |
| 1.00 |
| 1.00 |
| 0.01 |
| 0.16 |
| 0.00 |
| 0.00 |
| 0.00 |
| 0.00 |
| 0.00 |
| 0.08 |
| 0.67 |
| 0.72 |
| 0.00 |
| 0.42 |
| 0.00 |
| 0.02 |
| 0.45 |
| 0.19 |
| 0.91 |
| 1.00 |
| 0.21 |
| 1.00 |
| 1.00 |
| 0.82 |
| 0.00 |
| 0.00 |
| 0.04 |
| 0.00 |
| 0.00 |

| |
|---|
| 1.00 |
| 0.00 |
| 0.00 |
| 0.01 |
| 1.00 |
| 0.40 |
| 0.75 |
| 0.86 |
| 0.00 |
| 0.00 |
| 0.00 |
| 0.39 |
| 0.30 |
| 0.00 |
| 1.00 |
| 0.80 |
| 0.00 |
| 0.03 |
| 0.01 |
| 0.00 |
| 0.07 |
| 0.41 |
| 0.87 |
| 0.02 |
| 0.00 |
| 1.00 |
| 0.10 |
| 0.01 |
| 0.55 |
| 0.05 |
| 0.00 |
| 0.13 |
| 0.41 |
| 0.03 |
| 1.00 |
| 0.00 |
| 1.00 |
| 0.63 |
| 0.83 |
| 0.00 |
| 0.30 |
| 0.03 |
| 0.85 |
| 0.42 |
| 0.01 |
| 1.00 |
| 1.00 |
| 0.00 |

| |
|---|
| 0.89 |
| 0.03 |
| 0.00 |
| 0.00 |
| 0.00 |
| 0.22 |
| 0.80 |
| 1.00 |
| 0.11 |
| 0.38 |
| 0.94 |
| 1.00 |
| 1.00 |
| 0.96 |
| 0.63 |
| 1.00 |
| 0.00 |
| 0.01 |
| 0.00 |
| 0.14 |
| 0.03 |
| 0.03 |
| 0.00 |
| 1.00 |
| 0.98 |
| 0.52 |
| 0.29 |
| 0.28 |
| 1.00 |
| 0.81 |
| 1.00 |
| 0.54 |
| 0.68 |
| 0.88 |
| 0.68 |
| 0.00 |
| 0.00 |
| 1.00 |
| 0.00 |
| 0.84 |
| 1.00 |
| 1.00 |
| 1.00 |
| 0.36 |
| 0.65 |
| 0.01 |
| 1.00 |
| 0.00 |

| |
|---|
| 1.00 |
| 0.61 |
| 1.00 |
| 0.00 |
| 0.08 |
| 1.00 |
| 0.67 |
| 0.00 |
| 0.87 |
| 0.44 |
| 1.00 |
| 0.37 |
| 0.66 |
| 0.00 |
| 0.65 |
| 1.00 |
| 1.00 |
| 0.55 |
| 0.03 |
| 0.00 |
| 0.95 |
| 0.17 |
| 0.86 |
| 0.20 |
| 0.02 |
| 1.00 |
| 0.04 |
| 1.00 |
| 0.00 |
| 0.00 |
| 0.00 |
| 1.00 |
| 0.37 |
| 1.00 |
| 0.23 |
| 0.00 |
| 0.91 |
| 0.00 |
| 0.06 |
| 0.76 |
| 0.00 |
| 0.00 |
| 0.11 |
| 0.18 |
| 0.09 |
| 0.10 |
| 0.00 |
| 1.00 |

| |
|---|
| 0.99 |
| 0.61 |
| 0.05 |
| 0.00 |
| 0.82 |
| 0.00 |
| 0.01 |
| 0.00 |
| 0.08 |
| 0.41 |
| 0.02 |
| 0.07 |
| 0.00 |
| 0.34 |
| 0.00 |
| 0.29 |
| 0.00 |
| 0.77 |
| 0.70 |
| 0.02 |
| 0.08 |
| 0.01 |
| 1.00 |
| 0.87 |
| 1.00 |
| 0.00 |
| 0.61 |
| 0.00 |
| 1.00 |
| 0.00 |
| 1.00 |
| 0.02 |
| 0.02 |
| 1.00 |
| 0.00 |
| 0.04 |
| 1.00 |
| 0.46 |
| 0.33 |
| 0.00 |
| 0.78 |
| 0.19 |
| 0.03 |
| 0.00 |
| 0.49 |
| 0.14 |
| 0.91 |
| 0.00 |

| |
|---|
| 1.00 |
| 0.00 |
| 1.00 |
| 1.00 |
| 0.91 |
| 0.24 |
| 1.00 |
| 0.04 |
| 0.68 |
| 1.00 |
| 1.00 |
| 0.00 |
| 0.59 |
| 0.00 |
| 1.00 |
| 0.36 |
| 1.00 |
| 0.01 |
| 0.68 |
| 0.56 |
| 1.00 |
| 0.00 |
| 0.04 |
| 0.97 |
| 0.06 |
| 0.00 |
| 0.14 |
| 1.00 |
| 0.23 |
| 0.00 |
| 0.33 |
| 0.04 |
| 0.91 |
| 0.88 |
| 0.18 |
| 0.80 |
| 1.00 |
| 0.02 |
| 0.01 |
| 1.00 |
| 0.03 |
| 1.00 |
| 1.00 |
| 0.16 |
| 0.77 |
| 0.40 |
| 1.00 |
| 1.00 |

| |
|---|
| 0.38 |
| 0.48 |
| 0.07 |
| 0.25 |
| 0.58 |
| 0.01 |
| 0.70 |
| 0.86 |
| 1.00 |
| 0.77 |
| 1.00 |
| 0.00 |
| 0.13 |
| 1.00 |
| 0.55 |
| 0.05 |
| 0.07 |
| 0.01 |
| 0.46 |
| 0.02 |
| 1.00 |
| 0.00 |
| 0.25 |
| 1.00 |
| 0.00 |
| 0.10 |
| 0.19 |
| 0.01 |
| 0.02 |
| 1.00 |
| 0.00 |
| 0.03 |
| 1.00 |
| 0.62 |
| 1.00 |
| 0.00 |
| 0.05 |
| 0.42 |
| 0.02 |
| 1.00 |
| 1.00 |
| 0.38 |
| 1.00 |
| 0.01 |
| 0.00 |
| 0.01 |
| 0.32 |
| 1.00 |

| |
|---|
| 1.00 |
| 0.14 |
| 0.64 |
| 0.00 |
| 0.37 |
| 0.68 |
| 0.44 |
| 0.00 |
| 0.28 |
| 0.53 |
| 0.00 |
| 0.30 |
| 0.00 |
| 1.00 |
| 0.34 |
| 0.04 |
| 1.00 |
| 1.00 |
| 0.30 |
| 1.00 |
| 1.00 |
| 1.00 |
| 0.71 |
| 1.00 |
| 0.01 |
| 1.00 |
| 0.12 |
| 0.00 |
| 0.00 |
| 1.00 |
| 0.09 |
| 0.00 |
| 0.00 |
| 0.11 |
| 0.00 |
| 1.00 |
| 0.40 |
| 0.68 |
| 0.02 |
| 0.00 |
| 0.82 |
| 0.30 |
| 0.00 |
| 0.66 |
| 0.20 |
| 0.10 |
| 0.26 |
| 0.00 |

| |
|---|
| 0.00 |
| 0.57 |
| 0.34 |
| 0.00 |
| 0.08 |
| 0.02 |
| 0.77 |
| 0.00 |
| 0.68 |
| 1.00 |
| 0.70 |
| 0.04 |
| 1.00 |
| 0.00 |
| 1.00 |
| 0.00 |
| 0.49 |
| 0.40 |
| 0.45 |
| 0.00 |
| 1.00 |
| 0.01 |
| 0.00 |
| 0.00 |
| 0.00 |
| 1.00 |
| 0.01 |
| 1.00 |
| 1.00 |
| 0.71 |
| 1.00 |
| 1.00 |
| 1.00 |
| 0.86 |
| 0.52 |
| 1.00 |
| 0.08 |
| 0.11 |
| 0.00 |
| 0.96 |
| 0.89 |
| 0.69 |
| 0.00 |
| 0.24 |
| 0.00 |
| 0.10 |
| 0.00 |
| 0.00 |

| |
|---|
| 1.00 |
| 0.81 |
| 0.07 |
| 0.95 |
| 0.49 |
| 0.83 |
| 0.00 |
| 0.01 |
| 0.23 |
| 0.00 |
| 0.05 |
| 1.00 |
| 0.00 |
| 0.07 |
| 0.00 |
| 1.00 |
| 1.00 |
| 1.00 |
| 1.00 |
| 0.04 |
| 0.00 |
| 0.00 |
| 0.70 |
| 0.02 |
| 1.00 |
| 0.00 |
| 0.19 |
| 1.00 |
| 0.19 |
| 0.00 |
| 0.03 |
| 1.00 |
| 1.00 |
| 0.00 |
| 0.38 |
| 1.00 |
| 0.00 |
| 0.00 |
| 1.00 |
| 0.00 |
| 1.00 |
| 0.01 |
| 0.38 |
| 0.15 |
| 1.00 |
| 0.82 |
| 0.00 |
| 0.46 |

| |
|---|
| 0.69 |
| 0.24 |
| 1.00 |
| 0.77 |
| 1.00 |
| 0.00 |
| 0.27 |
| 0.01 |
| 0.00 |
| 0.00 |
| 1.00 |
| 0.00 |
| 0.42 |
| 0.02 |
| 1.00 |
| 0.42 |
| 0.00 |
| 0.17 |
| 0.59 |
| 0.73 |
| 0.46 |
| 1.00 |
| 0.12 |
| 0.42 |
| 0.10 |
| 0.81 |
| 0.00 |
| 0.38 |
| 1.00 |
| 0.64 |
| 0.41 |
| 0.74 |
| 1.00 |
| 0.02 |
| 1.00 |
| 1.00 |
| 0.00 |
| 0.07 |
| 0.01 |
| 0.73 |
| 0.10 |
| 0.51 |
| 0.88 |
| 1.00 |
| 1.00 |
| 0.00 |
| 1.00 |
| 0.00 |

| |
|---|
| 1.00 |
| 0.43 |
| 1.00 |
| 0.89 |
| 1.00 |
| 0.13 |
| 0.16 |
| 0.77 |
| 0.00 |
| 1.00 |
| 0.66 |
| 1.00 |
| 1.00 |
| 0.85 |
| 1.00 |
| 0.77 |
| 0.34 |
| 1.00 |
| 1.00 |
| 1.00 |
| 1.00 |
| 1.00 |
| 1.00 |
| 1.00 |
| 0.22 |
| 0.52 |
| 1.00 |
| 0.94 |
| 1.00 |
| 0.29 |
| 0.26 |
| 1.00 |
| 1.00 |
| 0.33 |
| 0.59 |
| 1.00 |
| 1.00 |
| 0.01 |
| 0.73 |
| 1.00 |
| 0.06 |
| 0.27 |
| 0.59 |
| 0.12 |
| 0.00 |
| 1.00 |
| 0.77 |
| 1.00 |

| |
|---|
| 1.00 |
| 1.00 |
| 1.00 |
| 0.23 |
| 0.70 |
| 0.57 |
| 0.24 |
| 1.00 |
| 0.61 |
| 0.59 |
| 0.83 |
| 0.57 |
| 1.00 |
| 1.00 |
| 0.64 |
| 0.67 |
| 1.00 |
| 0.98 |
| 1.00 |
| 1.00 |
| 1.00 |
| 0.22 |
| 0.64 |
| 0.35 |
| 0.71 |
| 0.83 |
| 1.00 |
| 1.00 |
| 1.00 |
| 1.00 |
| 1.00 |
| 0.29 |
| 0.95 |
| 0.03 |
| 1.00 |
| 0.71 |
| 1.00 |
| 1.00 |
| 1.00 |
| 0.39 |
| 1.00 |
| 0.15 |
| 1.00 |
| 1.00 |
| 0.75 |
| 1.00 |
| 1.00 |
| 1.00 |

| |
|---|
| 1.00 |
| 1.00 |
| 0.03 |
| 0.05 |
| 1.00 |
| 0.04 |
| 1.00 |
| 0.44 |
| 1.00 |
| 1.00 |
| 1.00 |
| 1.00 |
| 1.00 |
| 1.00 |
| 1.00 |
| 0.14 |
| 1.00 |
| 1.00 |
| 1.00 |
| 1.00 |
| 0.22 |
| 0.72 |
| 1.00 |
| 1.00 |
| 0.57 |
| 0.83 |
| 1.00 |
| 1.00 |
| 0.59 |
| 0.66 |
| 1.00 |
| 0.74 |
| 0.83 |
| 1.00 |
| 1.00 |
| 0.95 |
| 1.00 |
| 1.00 |
| 0.68 |
| 1.00 |
| 1.00 |
| 1.00 |
| 0.44 |
| 0.19 |
| 1.00 |
| 0.62 |
| 1.00 |
| 0.90 |

| |
|---|
| 1.00 |
| 1.00 |
| 0.00 |
| 1.00 |
| 0.34 |
| 1.00 |
| 0.32 |
| 0.01 |
| 0.28 |
| 1.00 |
| 0.42 |
| 0.68 |
| 1.00 |
| 1.00 |
| 0.54 |
| 0.48 |
| 1.00 |
| 1.00 |
| 0.91 |
| 0.95 |
| 1.00 |
| 0.27 |
| 1.00 |
| 1.00 |
| 0.38 |
| 1.00 |
| 0.79 |
| 1.00 |
| 1.00 |
| 1.00 |
| 1.00 |
| 0.60 |
| 0.76 |
| 1.00 |
| 1.00 |
| 1.00 |
| 0.46 |
| 0.24 |
| 1.00 |
| 0.42 |
| 0.87 |
| 1.00 |
| 0.16 |
| 0.87 |
| 0.74 |
| 0.57 |
| 1.00 |
| 0.25 |

| |
|---|
| 1.00 |
| 0.77 |
| 1.00 |
| 0.58 |
| 0.44 |
| 1.00 |
| 0.22 |
| 1.00 |
| 0.77 |
| 0.15 |
| 0.00 |
| 0.03 |
| 0.02 |
| 0.00 |
| 0.00 |
| 0.06 |
| 1.00 |
| 1.00 |
| 0.02 |
| 0.10 |
| 0.00 |
| 0.01 |
| 0.00 |
| 0.00 |
| 0.02 |
| 0.00 |
| 0.00 |
| 0.00 |
| 0.00 |
| 0.37 |
| 0.87 |
| 0.07 |
| 0.00 |
| 0.00 |
| 0.16 |
| 0.02 |
| 1.00 |
| 0.77 |
| 0.01 |
| 0.77 |
| 1.00 |
| 0.13 |
| 0.14 |
| 0.05 |
| 0.01 |
| 0.00 |
| 0.44 |
| 1.00 |

| |
|---|
| 0.00 |
| 0.00 |
| 0.01 |
| 0.00 |
| 1.00 |
| 1.00 |
| 0.68 |
| 0.05 |
| 0.38 |
| 0.00 |
| 0.00 |
| 0.05 |
| 0.66 |
| 0.00 |
| 0.87 |
| 0.00 |
| 0.95 |
| 0.99 |
| 0.68 |
| 0.00 |
| 1.00 |
| 0.37 |
| 0.22 |
| 1.00 |
| 0.02 |
| 0.82 |
| 0.00 |
| 0.00 |
| 0.00 |
| 0.00 |
| 1.00 |
| 0.11 |
| 0.59 |
| 0.00 |
| 0.94 |
| 0.55 |
| 0.00 |
| 0.00 |
| 0.00 |
| 1.00 |
| 0.47 |
| 0.77 |
| 1.00 |
| 0.00 |
| 0.00 |
| 0.00 |
| 1.00 |
| 0.53 |

| |
|---|
| 0.22 |
| 1.00 |
| 1.00 |
| 0.32 |
| 0.04 |
| 0.02 |
| 0.00 |
| 0.00 |
| 0.01 |
| 0.32 |
| 1.00 |
| 0.01 |
| 0.57 |
| 1.00 |
| 1.00 |
| 0.10 |
| 1.00 |
| 1.00 |
| 0.76 |
| 0.29 |
| 0.41 |
| 0.62 |
| 0.00 |
| 0.19 |
| 0.00 |
| 1.00 |
| 1.00 |
| 0.39 |
| 0.48 |
| 0.04 |
| 1.00 |
| 0.52 |
| 0.77 |
| 0.43 |
| 0.00 |
| 0.36 |
| 0.00 |
| 1.00 |
| 0.35 |
| 0.00 |
| 1.00 |
| 0.35 |
| 0.00 |
| 0.08 |
| 0.21 |
| 0.48 |
| 1.00 |
| 0.90 |

| |
|---|
| 0.82 |
| 0.00 |
| 1.00 |
| 0.89 |
| 0.00 |
| 1.00 |
| 0.07 |
| 0.54 |
| 0.02 |
| 0.12 |
| 0.00 |
| 0.76 |
| 0.68 |
| 0.66 |
| 0.80 |
| 0.26 |
| 0.92 |
| 0.83 |
| 0.00 |
| 0.00 |
| 1.00 |
| 0.02 |
| 1.00 |
| 1.00 |
| 0.26 |
| 0.08 |
| 0.00 |
| 1.00 |
| 0.14 |
| 0.51 |
| 0.49 |
| 1.00 |
| 0.21 |
| 0.00 |
| 0.38 |
| 0.74 |
| 0.34 |
| 0.87 |
| 0.32 |
| 1.00 |
| 1.00 |
| 0.00 |
| 0.00 |
| 0.00 |
| 0.31 |
| 0.71 |
| 0.04 |
| 0.00 |

| |
|---|
| 0.00 |
| 0.00 |
| 0.00 |
| 0.07 |
| 0.00 |
| 0.00 |
| 0.00 |
| 1.00 |
| 0.02 |
| 0.00 |
| 0.01 |
| 0.00 |
| 0.00 |
| 0.78 |
| 0.11 |
| 0.00 |
| 0.19 |
| 1.00 |
| 0.06 |
| 0.01 |
| 0.33 |
| 0.00 |
| 1.00 |
| 0.05 |
| 0.26 |
| 1.00 |
| 0.19 |
| 0.29 |
| 0.00 |
| 1.00 |
| 1.00 |
| 0.18 |
| 0.44 |
| 0.44 |
| 0.00 |
| 1.00 |
| 0.57 |
| 0.01 |
| 0.00 |
| 1.00 |
| 0.00 |
| 0.68 |
| 1.00 |
| 1.00 |
| 1.00 |
| 0.60 |
| 0.21 |
| 0.23 |

| |
|---|
| 1.00 |
| 0.00 |
| 0.07 |
| 0.78 |
| 0.40 |
| 0.26 |
| 0.00 |
| 0.00 |
| 0.00 |
| 0.48 |
| 0.00 |
| 1.00 |
| 1.00 |
| 0.58 |
| 0.26 |
| 0.00 |
| 0.00 |
| 0.03 |
| 0.74 |
| 0.58 |
| 0.00 |
| 0.00 |
| 1.00 |
| 0.61 |
| 0.00 |
| 0.00 |
| 0.73 |
| 1.00 |
| 0.94 |
| 0.00 |
| 0.10 |
| 0.34 |
| 0.24 |
| 0.00 |
| 1.00 |
| 0.19 |
| 0.77 |
| 0.09 |
| 1.00 |
| 0.00 |
| 0.00 |
| 0.01 |
| 0.57 |
| 0.65 |
| 0.00 |
| 0.00 |
| 0.65 |
| 0.78 |

| |
|---|
| 0.50 |
| 1.00 |
| 0.06 |
| 0.30 |
| 0.00 |
| 0.03 |
| 0.19 |
| 0.05 |
| 1.00 |
| 0.10 |
| 0.90 |
| 0.00 |
| 0.00 |
| 0.51 |
| 0.49 |
| 0.00 |
| 0.00 |
| 0.89 |
| 0.00 |
| 0.41 |
| 0.00 |
| 0.00 |
| 0.07 |
| 0.74 |
| 1.00 |
| 0.06 |
| 0.00 |
| 1.00 |
| 0.44 |
| 1.00 |
| 1.00 |
| 0.56 |
| 0.00 |
| 0.00 |
| 0.00 |
| 0.10 |
| 1.00 |
| 0.15 |
| 0.00 |
| 1.00 |
| 0.28 |
| 0.01 |
| 0.29 |
| 0.50 |
| 0.11 |
| 0.73 |
| 1.00 |
| 0.01 |

| |
|---|
| 0.74 |
| 0.27 |
| 1.00 |
| 0.67 |
| 1.00 |
| 0.49 |
| 1.00 |
| 0.18 |
| 0.82 |
| 1.00 |
| 1.00 |
| 0.00 |
| 0.12 |
| 0.55 |
| 0.56 |
| 0.18 |
| 1.00 |
| 0.08 |
| 1.00 |
| 0.01 |
| 0.72 |
| 1.00 |
| 0.00 |
| 0.48 |
| 0.03 |
| 0.40 |
| 1.00 |
| 0.00 |
| 0.68 |
| 0.00 |
| 1.00 |
| 0.01 |
| 1.00 |
| 0.07 |
| 0.00 |
| 0.02 |
| 0.93 |
| 1.00 |
| 0.00 |
| 0.00 |
| 0.00 |
| 0.00 |
| 0.00 |
| 0.00 |
| 0.00 |
| 0.00 |
| 1.00 |
| 0.22 |

| |
|---|
| 0.00 |
| 0.13 |
| 0.42 |
| 1.00 |
| 0.01 |
| 0.00 |
| 1.00 |
| 0.74 |
| 0.07 |
| 0.01 |
| 0.89 |
| 1.00 |
| 0.35 |
| 0.13 |
| 0.00 |
| 0.08 |
| 0.00 |
| 1.00 |
| 0.02 |
| 0.00 |
| 0.41 |
| 1.00 |
| 0.00 |
| 0.00 |
| 0.00 |
| 1.00 |
| 0.00 |
| 0.41 |
| 0.25 |
| 0.14 |
| 1.00 |
| 0.00 |
| 1.00 |
| 0.36 |
| 0.00 |
| 0.00 |
| 0.01 |
| 0.00 |
| 1.00 |
| 0.60 |
| 1.00 |
| 0.11 |
| 0.00 |
| 1.00 |
| 0.86 |
| 0.18 |
| 0.39 |
| 0.01 |

| |
|---|
| 0.03 |
| 1.00 |
| 0.00 |
| 0.39 |
| 0.47 |
| 1.00 |
| 0.00 |
| 0.00 |
| 0.56 |
| 0.30 |
| 0.31 |
| 0.11 |
| 0.21 |
| 0.05 |
| 0.01 |
| 0.10 |
| 0.81 |
| 0.57 |
| 0.63 |
| 0.23 |
| 1.00 |
| 0.00 |
| 0.20 |
| 0.14 |
| 0.00 |
| 0.89 |
| 0.00 |
| 1.00 |
| 0.00 |
| 0.02 |
| 0.52 |
| 0.01 |
| 0.12 |
| 1.00 |
| 0.00 |
| 0.00 |
| 0.07 |
| 1.00 |
| 0.00 |
| 0.37 |
| 0.21 |
| 0.00 |
| 0.66 |
| 0.00 |
| 0.00 |
| 0.00 |
| 0.03 |
| 1.00 |

| |
|---|
| 0.01 |
| 1.00 |
| 1.00 |
| 1.00 |
| 0.04 |
| 0.01 |
| 0.00 |
| 0.80 |
| 1.00 |
| 0.00 |
| 0.05 |
| 0.01 |
| 0.87 |
| 1.00 |
| 1.00 |
| 0.22 |
| 0.09 |
| 0.00 |
| 0.64 |
| 0.45 |
| 0.57 |
| 1.00 |
| 0.00 |
| 0.02 |
| 0.32 |
| 1.00 |
| 0.82 |
| 0.00 |
| 0.00 |
| 0.00 |
| 0.02 |
| 0.33 |
| 0.45 |
| 0.01 |
| 0.46 |
| 0.51 |
| 0.68 |
| 1.00 |
| 1.00 |
| 0.96 |
| 0.55 |
| 0.04 |
| 1.00 |
| 0.02 |
| 0.03 |
| 0.20 |
| 0.10 |
| 0.00 |

| |
|---|
| 1.00 |
| 0.00 |
| 0.00 |
| 0.24 |
| 1.00 |
| 0.33 |
| 0.00 |
| 0.52 |
| 0.08 |
| 0.00 |
| 0.41 |
| 0.07 |
| 0.90 |
| 1.00 |
| 0.08 |
| 1.00 |
| 1.00 |
| 1.00 |
| 1.00 |
| 1.00 |
| 0.31 |
| 0.84 |
| 0.85 |
| 0.00 |
| 0.00 |
| 0.08 |
| 0.00 |
| 1.00 |
| 0.00 |
| 0.41 |
| 0.00 |
| 0.64 |
| 1.00 |
| 0.74 |
| 0.00 |
| 0.20 |
| 0.11 |
| 0.25 |
| 0.00 |
| 0.50 |
| 1.00 |
| 0.72 |
| 0.01 |
| 0.00 |
| 0.01 |
| 1.00 |
| 1.00 |
| 0.00 |

| |
|---|
| 1.00 |
| 1.00 |
| 0.89 |
| 0.01 |
| 0.05 |
| 1.00 |
| 0.87 |
| 0.52 |
| 1.00 |
| 1.00 |
| 0.39 |
| 0.02 |
| 1.00 |
| 0.68 |
| 0.67 |
| 1.00 |
| 1.00 |
| 0.04 |
| 0.02 |
| 0.01 |
| 1.00 |
| 0.00 |
| 1.00 |
| 0.01 |
| 1.00 |
| 0.00 |
| 0.00 |
| 1.00 |
| 1.00 |
| 0.11 |
| 1.00 |
| 0.00 |
| 0.00 |
| 1.00 |
| 0.34 |
| 0.83 |
| 0.42 |
| 1.00 |
| 0.76 |
| 0.12 |
| 1.00 |
| 1.00 |
| 0.75 |
| 1.00 |
| 0.30 |
| 0.18 |
| 0.96 |
| 0.43 |

| |
|---|
| 0.70 |
| 0.13 |
| 0.03 |
| 0.02 |
| 0.00 |
| 0.64 |
| 0.30 |
| 0.03 |
| 0.10 |
| 0.68 |
| 0.10 |
| 0.87 |
| 1.00 |
| 1.00 |
| 0.97 |
| 0.00 |
| 0.79 |
| 0.06 |
| 1.00 |
| 0.00 |
| 0.72 |
| 0.01 |
| 0.00 |
| 0.01 |
| 0.23 |
| 0.00 |
| 0.63 |
| 0.23 |
| 1.00 |
| 1.00 |
| 0.02 |
| 0.42 |
| 0.57 |
| 0.00 |
| 0.90 |
| 0.43 |
| 0.03 |
| 0.00 |
| 0.01 |
| 0.55 |
| 0.67 |
| 0.52 |
| 0.01 |
| 1.00 |
| 0.00 |
| 0.01 |
| 0.39 |
| 0.01 |

| |
|---|
| 0.06 |
| 1.00 |
| 1.00 |
| 0.02 |
| 0.00 |
| 0.06 |
| 1.00 |
| 1.00 |
| 0.24 |
| 1.00 |
| 0.50 |
| 0.00 |
| 0.73 |
| 1.00 |
| 0.23 |
| 0.00 |
| 0.03 |
| 1.00 |
| 0.40 |
| 0.00 |
| 0.00 |
| 0.00 |
| 1.00 |
| 0.01 |
| 0.52 |
| 0.27 |
| 0.07 |
| 1.00 |
| 0.30 |
| 0.00 |
| 0.24 |
| 0.37 |
| 0.43 |
| 0.60 |
| 0.76 |
| 1.00 |
| 0.09 |
| 0.05 |
| 1.00 |
| 1.00 |
| 0.70 |
| 1.00 |
| 0.55 |
| 0.55 |
| 0.00 |
| 0.00 |
| 0.00 |
| 1.00 |

| |
|---|
| 1.00 |
| 0.00 |
| 0.67 |
| 1.00 |
| 0.08 |
| 0.05 |
| 0.19 |
| 0.28 |
| 1.00 |
| 1.00 |
| 0.95 |
| 0.01 |
| 0.18 |
| 1.00 |
| 0.00 |
| 0.97 |
| 0.00 |
| 1.00 |
| 0.00 |
| 1.00 |
| 0.00 |
| 0.60 |
| 0.01 |
| 0.75 |
| 0.00 |
| 0.71 |
| 0.39 |
| 1.00 |
| 0.47 |
| 1.00 |
| 0.44 |
| 1.00 |
| 0.70 |
| 0.00 |
| 0.00 |
| 0.52 |
| 0.05 |
| 0.00 |
| 0.01 |
| 0.18 |
| 0.00 |
| 0.11 |
| 0.00 |
| 0.02 |
| 0.01 |
| 0.67 |
| 0.39 |
| 0.03 |

| |
|---|
| 0.00 |
| 1.00 |
| 1.00 |
| 0.00 |
| 0.11 |
| 1.00 |
| 1.00 |
| 0.85 |
| 0.16 |
| 1.00 |
| 0.00 |
| 1.00 |
| 0.06 |
| 1.00 |
| 0.00 |
| 0.64 |
| 0.14 |
| 1.00 |
| 0.76 |
| 0.06 |
| 0.48 |
| 0.12 |
| 0.03 |
| 0.00 |
| 0.01 |
| 0.80 |
| 1.00 |
| 1.00 |
| 0.13 |
| 0.00 |
| 0.00 |
| 0.06 |
| 1.00 |
| 0.00 |
| 1.00 |
| 0.07 |
| 0.00 |
| 0.25 |
| 0.00 |
| 1.00 |
| 1.00 |
| 0.00 |
| 1.00 |
| 0.33 |
| 0.57 |
| 1.00 |
| 1.00 |
| 0.00 |

| |
|---|
| 0.00 |
| 0.70 |
| 1.00 |
| 1.00 |
| 0.00 |
| 1.00 |
| 0.08 |
| 0.05 |
| 0.31 |
| 0.30 |
| 0.99 |
| 0.67 |
| 0.10 |
| 0.03 |
| 0.21 |
| 0.14 |
| 0.97 |
| 1.00 |
| 0.05 |
| 0.00 |
| 1.00 |
| 0.78 |
| 1.00 |
| 1.00 |
| 1.00 |
| 0.00 |
| 1.00 |
| 1.00 |
| 0.00 |
| 0.03 |
| 0.94 |
| 1.00 |
| 0.25 |
| 0.89 |
| 0.77 |
| 0.04 |
| 0.00 |
| 0.00 |
| 0.36 |
| 1.00 |
| 0.98 |
| 0.41 |
| 0.02 |
| 0.00 |
| 1.00 |
| 0.00 |
| 0.05 |
| 1.00 |

| |
|---|
| 0.84 |
| 0.16 |
| 0.30 |
| 1.00 |
| 0.35 |
| 0.03 |
| 0.65 |
| 0.37 |
| 0.00 |
| 1.00 |
| 0.54 |
| 0.96 |
| 0.00 |
| 0.01 |
| 0.53 |
| 0.53 |
| 1.00 |
| 1.00 |
| 1.00 |
| 1.00 |
| 0.66 |
| 0.00 |
| 0.04 |
| 1.00 |
| 0.00 |
| 0.00 |
| 1.00 |
| 0.02 |
| 0.00 |
| 0.35 |
| 1.00 |
| 0.02 |
| 1.00 |
| 0.03 |
| 0.16 |
| 1.00 |
| 1.00 |
| 0.00 |
| 0.00 |
| 0.00 |
| 0.27 |
| 1.00 |
| 0.34 |
| 1.00 |
| 0.70 |
| 0.00 |
| 0.41 |
| 0.18 |

| |
|---|
| 0.24 |
| 0.00 |
| 0.00 |
| 0.31 |
| 0.97 |
| 0.00 |
| 0.11 |
| 0.02 |
| 0.73 |
| 0.00 |
| 0.03 |
| 0.27 |
| 0.90 |
| 0.74 |
| 1.00 |
| 0.00 |
| 0.00 |
| 0.97 |
| 1.00 |
| 0.00 |
| 0.00 |
| 0.18 |
| 0.01 |
| 0.30 |
| 0.02 |
| 0.00 |
| 0.00 |
| 0.00 |
| 0.04 |
| 0.24 |
| 0.16 |
| 1.00 |
| 0.00 |
| 0.23 |
| 0.24 |
| 0.03 |
| 0.00 |
| 0.01 |
| 0.02 |
| 1.00 |
| 0.22 |
| 0.00 |
| 0.01 |
| 0.04 |
| 0.03 |
| 0.00 |
| 0.01 |
| 0.70 |

| |
|---|
| 1.00 |
| 0.49 |
| 0.11 |
| 0.01 |
| 0.00 |
| 0.84 |
| 0.05 |
| 1.00 |
| 1.00 |
| 0.10 |
| 0.00 |
| 1.00 |
| 0.85 |
| 1.00 |
| 0.15 |
| 1.00 |
| 1.00 |
| 0.27 |
| 0.35 |
| 0.73 |
| 0.28 |
| 0.00 |
| 0.60 |
| 0.49 |
| 0.13 |
| 0.41 |
| 0.21 |
| 0.02 |
| 0.49 |
| 0.93 |
| 0.05 |
| 0.16 |
| 0.00 |
| 1.00 |
| 0.24 |
| 0.00 |
| 0.05 |
| 1.00 |
| 0.37 |
| 0.05 |
| 0.01 |
| 0.08 |
| 1.00 |
| 0.00 |
| 0.42 |
| 0.91 |
| 1.00 |
| 1.00 |

| |
|---|
| 0.73 |
| 1.00 |
| 1.00 |
| 1.00 |
| 0.26 |
| 1.00 |
| 0.00 |
| 1.00 |
| 0.22 |
| 1.00 |
| 0.00 |
| 0.00 |
| 0.09 |
| 0.03 |
| 0.16 |
| 0.02 |
| 0.00 |
| 0.00 |
| 0.42 |
| 0.00 |
| 0.02 |
| 0.60 |
| 1.00 |
| 0.00 |
| 1.00 |
| 0.06 |
| 0.28 |
| 0.69 |
| 1.00 |
| 0.02 |
| 0.14 |
| 0.00 |
| 0.02 |
| 0.16 |
| 0.55 |
| 0.26 |
| 0.06 |
| 0.43 |
| 0.00 |
| 0.00 |
| 0.97 |
| 0.01 |
| 0.88 |
| 1.00 |
| 1.00 |
| 0.22 |
| 0.26 |
| 0.32 |

| |
|---|
| 1.00 |
| 1.00 |
| 1.00 |
| 0.40 |
| 0.19 |
| 0.48 |
| 1.00 |
| 1.00 |
| 0.00 |
| 0.00 |
| 1.00 |
| 1.00 |
| 0.40 |
| 0.00 |
| 0.00 |
| 0.20 |
| 0.31 |
| 0.87 |
| 1.00 |
| 1.00 |
| 0.07 |
| 0.00 |
| 0.00 |
| 1.00 |
| 1.00 |
| 0.76 |
| 0.66 |
| 1.00 |
| 0.03 |
| 0.01 |
| 0.23 |
| 0.49 |
| 1.00 |
| 0.01 |
| 1.00 |
| 1.00 |
| 0.46 |
| 1.00 |
| 0.00 |
| 1.00 |
| 0.02 |
| 1.00 |
| 0.68 |
| 0.01 |
| 0.05 |
| 0.25 |
| 1.00 |
| 1.00 |

| |
|---|
| 0.00 |
| 0.68 |
| 0.03 |
| 0.35 |
| 1.00 |
| 1.00 |
| 0.02 |
| 0.36 |
| 0.10 |
| 1.00 |
| 0.00 |
| 0.24 |
| 1.00 |
| 0.18 |
| 0.00 |
| 0.99 |
| 0.69 |
| 0.87 |
| 0.68 |
| 0.00 |
| 0.00 |
| 1.00 |
| 0.76 |
| 0.15 |
| 0.00 |
| 0.00 |
| 0.17 |
| 0.41 |
| 1.00 |
| 1.00 |
| 0.91 |
| 0.00 |
| 0.21 |
| 0.02 |
| 0.77 |
| 0.37 |
| 0.96 |
| 0.00 |
| 1.00 |
| 0.01 |
| 0.00 |
| 0.15 |
| 0.00 |
| 0.07 |
| 1.00 |
| 0.00 |
| 0.01 |
| 1.00 |

| |
|---|
| 1.00 |
| 0.74 |
| 1.00 |
| 0.94 |
| 0.63 |
| 0.68 |
| 0.82 |
| 1.00 |
| 0.07 |
| 0.04 |
| 0.51 |
| 0.02 |
| 0.78 |
| 0.11 |
| 1.00 |
| 0.00 |
| 0.00 |
| 1.00 |
| 0.10 |
| 0.00 |
| 1.00 |
| 1.00 |
| 0.26 |
| 0.24 |
| 0.18 |
| 0.00 |
| 0.01 |
| 0.00 |
| 0.10 |
| 1.00 |
| 0.90 |
| 0.00 |
| 0.77 |
| 0.00 |
| 1.00 |
| 0.52 |
| 0.59 |
| 0.00 |
| 0.01 |
| 0.10 |
| 0.00 |
| 0.07 |
| 1.00 |
| 0.74 |
| 0.00 |
| 0.53 |
| 0.00 |
| 0.00 |

| |
|---|
| 0.25 |
| 0.45 |
| 0.00 |
| 0.45 |
| 0.00 |
| 0.22 |
| 0.00 |
| 1.00 |
| 1.00 |
| 0.11 |
| 0.03 |
| 0.00 |
| 0.00 |
| 0.80 |
| 0.00 |
| 0.39 |
| 0.44 |
| 0.08 |
| 0.46 |
| 1.00 |
| 0.21 |
| 1.00 |
| 0.76 |
| 0.45 |
| 0.00 |
| 0.27 |
| 0.78 |
| 0.01 |
| 1.00 |
| 0.00 |
| 1.00 |
| 1.00 |
| 0.93 |
| 0.92 |
| 1.00 |
| 0.98 |
| 0.01 |
| 0.09 |
| 0.01 |
| 0.01 |
| 0.00 |
| 1.00 |
| 0.00 |
| 0.01 |
| 0.01 |
| 0.29 |
| 1.00 |
| 0.02 |

| |
|---|
| 0.01 |
| 1.00 |
| 0.64 |
| 1.00 |
| 0.00 |
| 0.00 |
| 0.00 |
| 0.00 |
| 1.00 |
| 0.00 |
| 0.80 |
| 0.00 |
| 0.00 |
| 0.01 |
| 0.08 |
| 0.69 |
| 0.30 |
| 0.25 |
| 0.28 |
| 1.00 |
| 1.00 |
| 0.00 |
| 0.18 |
| 1.00 |
| 0.01 |
| 1.00 |
| 0.45 |
| 1.00 |
| 0.47 |
| 0.05 |
| 0.25 |
| 0.20 |
| 1.00 |
| 0.02 |
| 0.09 |
| 0.00 |
| 0.00 |
| 0.31 |
| 0.00 |
| 0.27 |
| 0.12 |
| 0.05 |
| 0.70 |
| 1.00 |
| 0.02 |
| 0.00 |
| 0.15 |
| 0.64 |

| |
|---|
| 0.00 |
| 0.00 |
| 0.00 |
| 0.00 |
| 0.10 |
| 0.00 |
| 0.44 |
| 0.17 |
| 0.90 |
| 0.68 |
| 0.20 |
| 0.00 |
| 1.00 |
| 0.23 |
| 0.00 |
| 1.00 |
| 0.00 |
| 0.43 |
| 0.00 |
| 0.05 |
| 0.00 |
| 1.00 |
| 1.00 |
| 1.00 |
| 0.00 |
| 0.02 |
| 1.00 |
| 1.00 |
| 0.02 |
| 0.00 |
| 0.41 |
| 1.00 |
| 0.03 |
| 0.12 |
| 0.79 |
| 1.00 |
| 0.40 |
| 1.00 |
| 0.61 |
| 0.77 |
| 0.05 |
| 0.00 |
| 1.00 |
| 1.00 |
| 0.40 |
| 0.05 |
| 0.41 |
| 0.90 |

| |
|---|
| 0.60 |
| 0.86 |
| 0.51 |
| 0.38 |
| 0.16 |
| 0.05 |
| 0.00 |
| 1.00 |
| 0.24 |
| 1.00 |
| 0.00 |
| 0.21 |
| 0.63 |
| 0.00 |
| 0.61 |
| 1.00 |
| 0.06 |
| 0.02 |
| 0.12 |
| 0.95 |
| 0.00 |
| 0.10 |
| 0.20 |
| 1.00 |
| 0.00 |
| 1.00 |
| 0.70 |
| 0.17 |
| 0.48 |
| 0.25 |
| 1.00 |
| 1.00 |
| 0.12 |
| 0.89 |
| 0.00 |
| 1.00 |
| 1.00 |
| 1.00 |
| 0.00 |
| 0.00 |
| 0.60 |
| 0.36 |
| 0.00 |
| 0.00 |
| 0.12 |
| 0.43 |
| 1.00 |
| 0.87 |

| |
|---|
| 1.00 |
| 1.00 |
| 1.00 |
| 0.22 |
| 0.26 |
| 0.03 |
| 0.00 |
| 0.42 |
| 0.03 |
| 0.02 |
| 1.00 |
| 0.33 |
| 1.00 |
| 0.07 |
| 0.15 |
| 0.85 |
| 0.17 |
| 1.00 |
| 0.01 |
| 0.65 |
| 0.33 |
| 0.13 |
| 0.72 |
| 0.00 |
| 1.00 |
| 0.00 |
| 0.01 |
| 0.00 |
| 0.05 |
| 0.43 |
| 0.14 |
| 0.00 |
| 0.00 |
| 1.00 |
| 1.00 |
| 0.77 |
| 0.00 |
| 0.00 |
| 0.00 |
| 1.00 |
| 0.01 |
| 0.70 |
| 0.00 |
| 0.38 |
| 1.00 |
| 0.18 |
| 1.00 |
| 1.00 |

| |
|---|
| 1.00 |
| 1.00 |
| 0.03 |
| 0.82 |
| 0.01 |
| 0.96 |
| 1.00 |
| 1.00 |
| 0.86 |
| 0.00 |
| 0.12 |
| 0.59 |
| 0.03 |
| 0.75 |
| 1.00 |
| 0.62 |
| 1.00 |
| 0.00 |
| 0.97 |
| 1.00 |
| 0.11 |
| 1.00 |
| 1.00 |
| 0.26 |
| 1.00 |
| 1.00 |
| 0.28 |
| 1.00 |
| 0.37 |
| 1.00 |
| 0.04 |
| 1.00 |
| 1.00 |
| 0.01 |
| 1.00 |
| 0.00 |
| 0.00 |
| 0.02 |
| 0.23 |
| 1.00 |
| 1.00 |
| 0.00 |
| 0.00 |
| 0.00 |
| 1.00 |
| 1.00 |
| 0.74 |
| 0.00 |

| |
|---|
| 0.00 |
| 0.06 |
| 1.00 |
| 0.03 |
| 0.25 |
| 0.56 |
| 0.20 |
| 0.01 |
| 1.00 |
| 0.11 |
| 0.07 |
| 0.00 |
| 0.00 |
| 1.00 |
| 0.01 |
| 0.42 |
| 1.00 |
| 0.12 |
| 0.10 |
| 0.13 |
| 0.00 |
| 0.01 |
| 1.00 |
| 0.17 |
| 0.77 |
| 0.89 |
| 0.00 |
| 1.00 |
| 0.00 |
| 1.00 |
| 0.00 |
| 1.00 |
| 0.00 |
| 0.03 |
| 0.01 |
| 0.00 |
| 0.06 |
| 0.00 |
| 0.06 |
| 0.01 |
| 0.00 |
| 0.09 |
| 1.00 |
| 1.00 |
| 1.00 |
| 0.41 |
| 1.00 |
| 0.06 |

| |
|---|
| 0.43 |
| 0.14 |
| 0.00 |
| 0.45 |
| 0.27 |
| 1.00 |
| 0.00 |
| 0.08 |
| 0.00 |
| 1.00 |
| 1.00 |
| 1.00 |
| 1.00 |
| 1.00 |
| 0.69 |
| 0.69 |
| 0.05 |
| 1.00 |
| 0.21 |
| 0.65 |
| 1.00 |
| 0.07 |
| 0.53 |
| 0.00 |
| 0.34 |
| 0.02 |
| 0.70 |
| 0.03 |
| 0.22 |
| 0.90 |
| 1.00 |
| 0.61 |
| 1.00 |
| 0.49 |
| 0.18 |
| 0.03 |
| 1.00 |
| 0.64 |
| 0.52 |
| 0.16 |
| 0.05 |
| 0.16 |
| 0.43 |
| 0.03 |
| 0.02 |
| 0.27 |
| 0.00 |
| 0.62 |

| |
|---|
| 0.67 |
| 0.00 |
| 0.00 |
| 0.03 |
| 0.00 |
| 1.00 |
| 0.00 |
| 0.33 |
| 0.15 |
| 0.00 |
| 0.00 |
| 0.08 |
| 0.00 |
| 0.01 |
| 1.00 |
| 0.77 |
| 1.00 |
| 0.68 |
| 0.17 |
| 1.00 |
| 0.98 |
| 0.10 |
| 0.00 |
| 1.00 |
| 0.05 |
| 0.11 |
| 0.05 |
| 1.00 |
| 1.00 |
| 0.08 |
| 1.00 |
| 0.00 |
| 0.20 |
| 0.00 |
| 0.00 |
| 0.08 |
| 1.00 |
| 0.55 |
| 0.00 |
| 0.78 |
| 0.07 |
| 0.13 |
| 0.00 |
| 0.97 |
| 0.78 |
| 0.99 |
| 0.04 |
| 1.00 |

| |
|---|
| 0.07 |
| 0.03 |
| 1.00 |
| 1.00 |
| 0.00 |
| 0.00 |
| 0.77 |
| 0.00 |
| 1.00 |
| 1.00 |
| 0.01 |
| 0.55 |
| 0.00 |
| 0.00 |
| 0.00 |
| 0.00 |
| 0.50 |
| 0.00 |
| 0.95 |
| 0.08 |
| 1.00 |
| 1.00 |
| 0.00 |
| 1.00 |
| 0.01 |
| 0.47 |
| 0.81 |
| 1.00 |
| 0.09 |
| 0.15 |
| 1.00 |
| 0.85 |
| 0.36 |
| 0.00 |
| 0.40 |
| 0.00 |
| 0.01 |
| 1.00 |
| 0.71 |
| 1.00 |
| 0.04 |
| 0.06 |
| 0.64 |
| 1.00 |
| 0.35 |
| 0.82 |
| 0.64 |
| 1.00 |

| |
|---|
| 1.00 |
| 0.11 |
| 0.11 |
| 0.20 |
| 0.01 |
| 0.01 |
| 0.00 |
| 0.02 |
| 1.00 |
| 0.11 |
| 0.65 |
| 0.00 |
| 0.00 |
| 0.03 |
| 0.26 |
| 0.30 |
| 0.01 |
| 0.00 |
| 0.00 |
| 1.00 |
| 0.30 |
| 0.00 |
| 1.00 |
| 0.00 |
| 0.01 |
| 0.00 |
| 1.00 |
| 0.05 |
| 0.46 |
| 0.41 |
| 0.00 |
| 0.10 |
| 0.01 |
| 0.00 |
| 1.00 |
| 0.03 |
| 0.14 |
| 0.79 |
| 0.00 |
| 0.00 |
| 0.25 |
| 0.00 |
| 0.04 |
| 0.09 |
| 0.56 |
| 0.00 |
| 0.00 |
| 0.18 |

| |
|---|
| 1.00 |
| 0.02 |
| 1.00 |
| 1.00 |
| 0.58 |
| 0.14 |
| 1.00 |
| 0.01 |
| 1.00 |
| 0.03 |
| 0.00 |
| 0.00 |
| 1.00 |
| 0.00 |
| 0.00 |
| 0.67 |
| 0.02 |
| 0.39 |
| 0.94 |
| 0.03 |
| 1.00 |
| 0.00 |
| 0.00 |
| 0.78 |
| 1.00 |
| 0.12 |
| 0.68 |
| 1.00 |
| 0.39 |
| 0.18 |
| 1.00 |
| 1.00 |
| 1.00 |
| 0.07 |
| 0.56 |
| 0.36 |
| 0.61 |
| 0.00 |
| 1.00 |
| 0.00 |
| 0.19 |
| 1.00 |
| 0.09 |
| 0.00 |
| 0.09 |
| 0.22 |
| 1.00 |
| 0.03 |

| |
|---|
| 0.00 |
| 0.54 |
| 0.09 |
| 0.43 |
| 0.14 |
| 1.00 |
| 0.00 |
| 1.00 |
| 0.08 |
| 0.19 |
| 0.42 |
| 1.00 |
| 0.02 |
| 0.00 |
| 1.00 |
| 0.48 |
| 0.56 |
| 1.00 |
| 0.50 |
| 1.00 |
| 0.35 |
| 0.00 |
| 0.91 |
| 0.92 |
| 1.00 |
| 0.94 |
| 1.00 |
| 0.59 |
| 1.00 |
| 0.44 |
| 0.00 |
| 0.23 |
| 0.69 |
| 1.00 |
| 0.78 |
| 0.00 |
| 1.00 |
| 0.00 |
| 0.29 |
| 0.00 |
| 0.30 |
| 1.00 |
| 0.60 |
| 0.82 |
| 0.05 |
| 1.00 |
| 1.00 |
| 1.00 |

| |
|---|
| 0.92 |
| 0.05 |
| 0.61 |
| 1.00 |
| 0.00 |
| 0.06 |
| 0.08 |
| 0.61 |
| 0.74 |
| 0.00 |
| 0.00 |
| 0.49 |
| 0.05 |
| 0.00 |
| 0.73 |
| 0.68 |
| 1.00 |
| 0.00 |
| 0.23 |
| 0.24 |
| 0.79 |
| 0.67 |
| 0.61 |
| 1.00 |
| 1.00 |
| 1.00 |
| 0.00 |
| 1.00 |
| 0.00 |
| 1.00 |
| 1.00 |
| 0.39 |
| 0.52 |
| 0.01 |
| 0.00 |
| 0.70 |
| 0.41 |
| 1.00 |
| 0.02 |
| 0.62 |
| 0.90 |
| 0.75 |
| 0.10 |
| 0.05 |
| 1.00 |
| 1.00 |
| 0.23 |
| 1.00 |

| |
|---|
| 1.00 |
| 0.43 |
| 1.00 |
| 1.00 |
| 0.00 |
| 0.82 |
| 0.05 |
| 0.75 |
| 0.04 |
| 0.00 |
| 1.00 |
| 0.59 |
| 0.09 |
| 1.00 |
| 1.00 |
| 0.37 |
| 1.00 |
| 0.00 |
| 0.76 |
| 0.84 |
| 0.60 |
| 1.00 |
| 0.15 |
| 1.00 |
| 0.79 |
| 1.00 |
| 0.09 |
| 1.00 |
| 0.42 |
| 1.00 |
| 1.00 |
| 0.00 |
| 0.07 |
| 1.00 |
| 0.00 |
| 0.52 |
| 0.93 |
| 0.14 |
| 0.22 |
| 1.00 |
| 0.03 |
| 0.16 |
| 0.08 |
| 0.00 |
| 0.71 |
| 0.00 |
| 0.01 |
| 1.00 |

| |
|---|
| 0.18 |
| 0.68 |
| 0.00 |
| 0.59 |
| 0.21 |
| 1.00 |
| 1.00 |
| 1.00 |
| 1.00 |
| 0.00 |
| 0.32 |
| 1.00 |
| 0.01 |
| 1.00 |
| 1.00 |
| 0.04 |
| 0.00 |
| 0.05 |
| 1.00 |
| 0.38 |
| 0.00 |
| 0.00 |
| 0.16 |
| 0.03 |
| 0.24 |
| 0.94 |
| 1.00 |
| 0.00 |
| 0.00 |
| 0.00 |
| 0.00 |
| 0.00 |
| 0.20 |
| 0.00 |
| 0.01 |
| 1.00 |
| 0.49 |
| 0.00 |
| 0.39 |
| 0.00 |
| 1.00 |
| 0.41 |
| 1.00 |
| 0.00 |
| 0.38 |
| 1.00 |
| 0.57 |
| 1.00 |

| |
|---|
| 0.00 |
| 1.00 |
| 0.39 |
| 0.00 |
| 0.04 |
| 0.00 |
| 0.00 |
| 1.00 |
| 1.00 |
| 0.89 |
| 0.08 |
| 0.18 |
| 1.00 |
| 1.00 |
| 0.00 |
| 0.00 |
| 0.20 |
| 0.00 |
| 1.00 |
| 0.02 |
| 1.00 |
| 0.17 |
| 1.00 |
| 0.40 |
| 0.08 |
| 0.94 |
| 0.73 |
| 0.07 |
| 0.42 |
| 0.61 |
| 1.00 |
| 0.02 |
| 0.06 |
| 0.05 |
| 1.00 |
| 0.79 |
| 0.66 |
| 1.00 |
| 1.00 |
| 1.00 |
| 0.98 |
| 0.80 |
| 0.72 |
| 0.33 |
| 0.00 |
| 0.31 |
| 0.01 |
| 0.00 |

| |
|---|
| 1.00 |
| 0.17 |
| 1.00 |
| 0.40 |
| 0.41 |
| 1.00 |
| 0.00 |
| 0.04 |
| 1.00 |
| 0.13 |
| 0.12 |
| 0.01 |
| 0.19 |
| 0.34 |
| 0.35 |
| 0.83 |
| 0.61 |
| 0.46 |
| 0.92 |
| 1.00 |
| 1.00 |
| 1.00 |
| 1.00 |
| 0.00 |
| 1.00 |
| 0.41 |
| 1.00 |
| 0.90 |
| 0.37 |
| 0.71 |
| 1.00 |
| 0.02 |
| 0.04 |
| 1.00 |
| 1.00 |
| 0.04 |
| 0.15 |
| 0.52 |
| 0.21 |
| 1.00 |
| 0.01 |
| 0.01 |
| 0.00 |
| 0.21 |
| 0.00 |
| 1.00 |
| 0.00 |
| 0.75 |

| |
|---|
| 0.00 |
| 1.00 |
| 0.03 |
| 0.00 |
| 0.00 |
| 1.00 |
| 0.82 |
| 0.19 |
| 0.55 |
| 0.00 |
| 0.55 |
| 0.70 |
| 0.36 |
| 0.00 |
| 1.00 |
| 1.00 |
| 0.11 |
| 0.10 |
| 0.32 |
| 0.24 |
| 0.00 |
| 0.08 |
| 0.20 |
| 0.79 |
| 1.00 |
| 0.95 |
| 0.03 |
| 1.00 |
| 0.03 |
| 0.42 |
| 1.00 |
| 0.00 |
| 0.00 |
| 0.02 |
| 0.04 |
| 0.54 |
| 1.00 |
| 0.15 |
| 0.00 |
| 0.11 |
| 0.24 |
| 1.00 |
| 0.88 |
| 1.00 |
| 0.89 |
| 0.00 |
| 0.46 |
| 0.57 |

| |
|---|
| 1.00 |
| 0.95 |
| 0.03 |
| 0.63 |
| 1.00 |
| 0.70 |
| 0.00 |
| 0.56 |
| 0.50 |
| 1.00 |
| 0.00 |
| 0.00 |
| 1.00 |
| 0.91 |
| 0.00 |
| 0.71 |
| 0.80 |
| 0.12 |
| 0.21 |
| 0.00 |
| 0.11 |
| 1.00 |
| 0.65 |
| 0.43 |
| 0.16 |
| 1.00 |
| 0.00 |
| 1.00 |
| 0.02 |
| 0.00 |
| 1.00 |
| 0.00 |
| 0.16 |
| 0.00 |
| 0.18 |
| 1.00 |
| 0.00 |
| 0.20 |
| 0.73 |
| 1.00 |
| 0.00 |
| 1.00 |
| 0.01 |
| 0.70 |
| 0.00 |
| 0.07 |
| 0.74 |
| 0.64 |

| |
|---|
| 1.00 |
| 1.00 |
| 1.00 |
| 0.00 |
| 1.00 |
| 0.60 |
| 0.00 |
| 0.13 |
| 0.17 |
| 0.00 |
| 0.71 |
| 0.55 |
| 1.00 |
| 0.28 |
| 0.00 |
| 1.00 |
| 0.23 |
| 0.00 |
| 0.49 |
| 1.00 |
| 0.27 |
| 0.00 |
| 0.72 |
| 1.00 |
| 1.00 |
| 0.50 |
| 1.00 |
| 1.00 |
| 1.00 |
| 0.95 |
| 1.00 |
| 0.60 |
| 0.74 |
| 0.97 |
| 0.00 |
| 0.01 |
| 0.05 |
| 0.11 |
| 0.40 |
| 0.00 |
| 0.06 |
| 0.68 |
| 0.00 |
| 0.49 |
| 0.08 |
| 1.00 |
| 0.71 |
| 1.00 |

| |
|---|
| 1.00 |
| 0.00 |
| 0.00 |
| 0.37 |
| 0.11 |
| 1.00 |
| 0.00 |
| 0.00 |
| 0.45 |
| 0.00 |
| 1.00 |
| 0.61 |
| 0.13 |
| 1.00 |
| 1.00 |
| 0.00 |
| 0.00 |
| 1.00 |
| 0.19 |
| 0.01 |
| 0.00 |
| 1.00 |
| 0.01 |
| 0.11 |
| 1.00 |
| 1.00 |
| 0.01 |
| 0.84 |
| 0.54 |
| 0.96 |
| 0.59 |
| 0.32 |
| 0.62 |
| 0.00 |
| 0.08 |
| 0.77 |
| 0.26 |
| 0.00 |
| 0.00 |
| 0.00 |
| 0.17 |
| 0.13 |
| 0.25 |
| 1.00 |
| 0.83 |
| 0.28 |
| 1.00 |
| 0.79 |

| |
|---|
| 0.01 |
| 1.00 |
| 0.16 |
| 0.25 |
| 0.01 |
| 0.31 |
| 1.00 |
| 1.00 |
| 1.00 |
| 1.00 |
| 1.00 |
| 1.00 |
| 1.00 |
| 0.78 |
| 1.00 |
| 1.00 |
| 0.00 |
| 0.00 |
| 0.74 |
| 0.14 |
| 1.00 |
| 0.24 |
| 0.07 |
| 1.00 |
| 0.09 |
| 1.00 |
| 0.99 |
| 0.27 |
| 0.01 |
| 0.00 |
| 1.00 |
| 0.02 |
| 1.00 |
| 1.00 |
| 0.07 |
| 0.16 |
| 0.00 |
| 0.46 |
| 0.26 |
| 0.00 |
| 1.00 |
| 0.56 |
| 0.13 |
| 1.00 |
| 0.03 |
| 0.04 |
| 1.00 |
| 1.00 |

| |
|---|
| 1.00 |
| 0.91 |
| 0.59 |
| 0.06 |
| 1.00 |
| 0.07 |
| 0.76 |
| 0.00 |
| 0.23 |
| 0.54 |
| 1.00 |
| 0.00 |
| 0.50 |
| 0.14 |
| 0.21 |
| 1.00 |
| 0.45 |
| 0.09 |
| 0.32 |
| 0.07 |
| 0.03 |
| 1.00 |
| 0.87 |
| 1.00 |
| 0.00 |
| 0.11 |
| 0.00 |
| 0.02 |
| 1.00 |
| 0.92 |
| 0.00 |
| 0.08 |
| 0.38 |
| 0.88 |
| 0.01 |
| 0.92 |
| 0.00 |
| 0.31 |
| 0.07 |
| 0.01 |
| 1.00 |
| 0.01 |
| 1.00 |
| 1.00 |
| 0.70 |
| 0.13 |
| 0.02 |
| 0.18 |

| |
|---|
| 0.00 |
| 0.41 |
| 0.06 |
| 1.00 |
| 0.04 |
| 0.12 |
| 0.50 |
| 0.33 |
| 1.00 |
| 1.00 |
| 0.00 |
| 0.00 |
| 1.00 |
| 1.00 |
| 1.00 |
| 0.05 |
| 0.40 |
| 0.00 |
| 0.11 |
| 0.56 |
| 0.00 |
| 0.60 |
| 0.01 |
| 0.01 |
| 1.00 |
| 0.56 |
| 0.73 |
| 0.90 |
| 1.00 |
| 1.00 |
| 0.28 |
| 0.00 |
| 0.02 |
| 0.71 |
| 0.00 |
| 1.00 |
| 0.03 |
| 0.14 |
| 1.00 |
| 1.00 |
| 0.07 |
| 0.50 |
| 0.75 |
| 0.41 |
| 0.00 |
| 0.00 |
| 0.66 |
| 1.00 |

| |
|---|
| 0.01 |
| 0.69 |
| 1.00 |
| 0.00 |
| 0.00 |
| 0.13 |
| 0.00 |
| 0.06 |
| 0.01 |
| 0.68 |
| 1.00 |
| 0.07 |
| 0.32 |
| 1.00 |
| 0.10 |
| 1.00 |
| 0.08 |
| 1.00 |
| 0.49 |
| 1.00 |
| 0.07 |
| 0.03 |
| 0.88 |
| 0.38 |
| 0.94 |
| 0.00 |
| 1.00 |
| 0.10 |
| 0.66 |
| 1.00 |
| 0.00 |
| 0.07 |
| 0.01 |
| 1.00 |
| 0.07 |
| 1.00 |
| 0.00 |
| 1.00 |
| 0.24 |
| 0.09 |
| 0.16 |
| 0.22 |
| 0.51 |
| 0.68 |
| 0.92 |
| 0.81 |
| 0.53 |
| 0.09 |

| |
|---|
| 0.86 |
| 0.13 |
| 0.12 |
| 0.45 |
| 0.17 |
| 0.02 |
| 0.16 |
| 0.91 |
| 0.16 |
| 0.17 |
| 0.52 |
| 0.02 |
| 1.00 |
| 0.00 |
| 1.00 |
| 0.84 |
| 0.72 |
| 0.04 |
| 1.00 |
| 0.49 |
| 0.02 |
| 1.00 |
| 1.00 |
| 1.00 |
| 0.64 |
| 1.00 |
| 0.00 |
| 0.45 |
| 1.00 |
| 0.00 |
| 0.12 |
| 1.00 |
| 0.00 |
| 0.27 |
| 0.20 |
| 0.00 |
| 1.00 |
| 1.00 |
| 0.93 |
| 0.18 |
| 0.95 |
| 0.58 |
| 0.15 |
| 0.19 |
| 1.00 |
| 1.00 |
| 0.30 |
| 1.00 |

| |
|---|
| 0.24 |
| 1.00 |
| 0.40 |
| 0.30 |
| 1.00 |
| 1.00 |
| 0.31 |
| 0.00 |
| 0.10 |
| 0.05 |
| 0.00 |
| 0.73 |
| 0.02 |
| 1.00 |
| 0.50 |
| 1.00 |
| 0.40 |
| 1.00 |
| 1.00 |
| 1.00 |
| 0.43 |
| 0.01 |
| 0.08 |
| 0.00 |
| 1.00 |
| 0.01 |
| 0.00 |
| 0.39 |
| 0.62 |
| 1.00 |
| 1.00 |
| 1.00 |
| 0.57 |
| 0.30 |
| 1.00 |
| 1.00 |
| 0.16 |
| 0.48 |
| 1.00 |
| 0.81 |
| 0.10 |
| 0.58 |
| 0.04 |
| 0.00 |
| 0.00 |
| 0.11 |
| 0.23 |
| 0.70 |

| |
|---|
| 0.63 |
| 1.00 |
| 0.46 |
| 1.00 |
| 1.00 |
| 0.96 |
| 0.63 |
| 0.72 |
| 0.49 |
| 1.00 |
| 0.88 |
| 0.73 |
| 0.41 |
| 0.00 |
| 0.13 |
| 1.00 |
| 1.00 |
| 0.12 |
| 0.65 |
| 0.04 |
| 0.24 |
| 0.30 |
| 1.00 |
| 1.00 |
| 0.00 |
| 1.00 |
| 0.00 |
| 1.00 |
| 1.00 |
| 0.00 |
| 1.00 |
| 0.71 |
| 0.00 |
| 0.00 |
| 1.00 |
| 0.00 |
| 0.38 |
| 0.59 |
| 0.43 |
| 1.00 |
| 0.40 |
| 1.00 |
| 0.47 |
| 0.77 |
| 0.45 |
| 0.61 |
| 0.44 |
| 1.00 |

| |
|---|
| 1.00 |
| 0.14 |
| 1.00 |
| 0.00 |
| 0.00 |
| 0.15 |
| 0.38 |
| 0.40 |
| 0.61 |
| 0.00 |
| 0.33 |
| 0.62 |
| 0.04 |
| 1.00 |
| 0.00 |
| 0.01 |
| 0.09 |
| 0.26 |
| 1.00 |
| 0.01 |
| 0.07 |
| 0.95 |
| 0.26 |
| 0.00 |
| 1.00 |
| 0.76 |
| 0.82 |
| 1.00 |
| 1.00 |
| 0.02 |
| 1.00 |
| 1.00 |
| 0.95 |
| 1.00 |
| 1.00 |
| 0.24 |
| 0.14 |
| 0.00 |
| 0.01 |
| 0.79 |
| 1.00 |
| 1.00 |
| 0.00 |
| 1.00 |
| 0.13 |
| 1.00 |
| 0.02 |
| 0.87 |

| |
|---|
| 0.36 |
| 0.70 |
| 1.00 |
| 1.00 |
| 1.00 |
| 0.01 |
| 1.00 |
| 0.04 |
| 1.00 |
| 0.14 |
| 0.31 |
| 1.00 |
| 1.00 |
| 0.70 |
| 1.00 |
| 0.29 |
| 1.00 |
| 1.00 |
| 1.00 |
| 1.00 |
| 0.23 |
| 0.03 |
| 0.88 |
| 0.16 |
| 1.00 |
| 0.22 |
| 0.56 |
| 0.98 |
| 0.79 |
| 0.15 |
| 0.01 |
| 0.00 |
| 1.00 |
| 0.00 |
| 0.27 |
| 0.21 |
| 0.00 |
| 0.53 |
| 0.01 |
| 0.49 |
| 0.71 |
| 1.00 |
| 1.00 |
| 1.00 |
| 0.13 |
| 1.00 |
| 0.52 |
| 0.54 |

| |
|---|
| 1.00 |
| 1.00 |
| 0.74 |
| 1.00 |
| 1.00 |
| 1.00 |
| 0.11 |
| 0.23 |
| 0.57 |
| 1.00 |
| 0.31 |
| 1.00 |
| 0.02 |
| 1.00 |
| 0.39 |
| 0.41 |
| 0.47 |
| 1.00 |
| 1.00 |
| 0.95 |
| 1.00 |
| 0.92 |
| 1.00 |
| 1.00 |
| 0.71 |
| 1.00 |
| 0.54 |
| 1.00 |
| 0.31 |
| 0.73 |
| 0.27 |
| 0.45 |
| 1.00 |
| 1.00 |
| 1.00 |
| 0.17 |
| 1.00 |
| 0.22 |
| 1.00 |
| 1.00 |
| 1.00 |
| 1.00 |
| 1.00 |
| 1.00 |
| 1.00 |
| 0.00 |
| 0.20 |
| 1.00 |

| |
|---|
| 0.09 |
| 0.00 |
| 0.04 |
| 0.88 |
| 1.00 |
| 0.91 |
| 0.00 |
| 0.73 |
| 0.95 |
| 0.26 |
| 1.00 |
| 1.00 |
| 0.00 |
| 0.00 |
| 1.00 |
| 0.01 |
| 0.82 |
| 0.73 |
| 0.63 |
| 0.99 |
| 0.00 |
| 0.95 |
| 1.00 |
| 0.00 |
| 1.00 |
| 0.14 |
| 1.00 |
| 1.00 |
| 1.00 |
| 1.00 |
| 1.00 |
| 1.00 |
| 1.00 |
| 1.00 |
| 0.18 |
| 0.78 |
| 0.06 |
| 0.00 |
| 0.57 |
| 0.01 |
| 0.27 |
| 1.00 |
| 0.00 |
| 0.17 |
| 1.00 |
| 0.71 |
| 1.00 |
| 0.26 |

| |
|---|
| 1.00 |
| 1.00 |
| 0.53 |
| 0.00 |
| 0.00 |
| 1.00 |
| 0.48 |
| 0.18 |
| 0.37 |
| 0.58 |
| 0.74 |
| 0.22 |
| 0.39 |
| 1.00 |
| 1.00 |
| 0.98 |
| 0.21 |
| 0.01 |
| 0.64 |
| 0.27 |
| 0.21 |
| 0.11 |
| 0.00 |
| 1.00 |
| 0.00 |
| 1.00 |
| 0.59 |
| 1.00 |
| 0.24 |
| 0.74 |
| 1.00 |
| 1.00 |
| 0.52 |
| 0.26 |
| 0.63 |
| 0.60 |
| 0.26 |
| 1.00 |
| 0.30 |
| 1.00 |
| 1.00 |
| 0.97 |
| 0.90 |
| 0.67 |
| 1.00 |
| 1.00 |
| 1.00 |
| 1.00 |

| |
|---|
| 0.43 |
| 0.47 |
| 1.00 |
| 0.30 |
| 0.32 |
| 0.54 |
| 0.57 |
| 1.00 |
| 1.00 |
| 1.00 |
| 0.02 |
| 0.00 |
| 1.00 |
| 1.00 |
| 0.04 |
| 1.00 |
| 0.51 |
| 1.00 |
| 1.00 |
| 0.14 |
| 0.15 |
| 0.05 |
| 0.07 |
| 0.57 |
| 1.00 |
| 0.57 |
| 0.00 |
| 1.00 |
| 0.00 |
| 0.07 |
| 0.56 |
| 0.24 |
| 0.14 |
| 1.00 |
| 0.01 |
| 0.00 |
| 0.35 |
| 0.44 |
| 0.95 |
| 1.00 |
| 0.03 |
| 0.61 |
| 0.03 |
| 1.00 |
| 0.11 |
| 1.00 |
| 1.00 |
| 1.00 |

| |
|---|
| 1.00 |
| 0.10 |
| 0.00 |
| 0.64 |
| 0.78 |
| 1.00 |
| 0.39 |
| 0.48 |
| 1.00 |
| 0.34 |
| 0.88 |
| 1.00 |
| 0.29 |
| 1.00 |
| 0.17 |
| 0.97 |
| 0.00 |
| 0.39 |
| 0.75 |
| 1.00 |
| 1.00 |
| 1.00 |
| 1.00 |
| 0.10 |
| 0.39 |
| 1.00 |
| 0.00 |
| 0.10 |
| 0.06 |
| 0.00 |
| 0.60 |
| 0.41 |
| 1.00 |
| 0.89 |
| 0.79 |
| 1.00 |
| 1.00 |
| 0.00 |
| 0.01 |
| 0.00 |
| 1.00 |
| 1.00 |
| 1.00 |
| 1.00 |
| 0.80 |
| 1.00 |
| 1.00 |
| 1.00 |

| |
|---|
| 0.89 |
| 1.00 |
| 0.95 |
| 0.18 |
| 0.03 |
| 0.86 |
| 0.12 |
| 0.05 |
| 0.97 |
| 0.01 |
| 0.00 |
| 0.56 |
| 1.00 |
| 1.00 |
| 0.01 |
| 1.00 |
| 0.65 |
| 0.07 |
| 0.05 |
| 0.00 |
| 0.00 |
| 0.00 |
| 0.00 |
| 0.45 |
| 0.15 |
| 0.03 |
| 0.00 |
| 0.79 |
| 0.34 |
| 0.00 |
| 1.00 |
| 0.46 |
| 0.43 |
| 0.00 |
| 0.25 |
