## Supplementary Legend for "Targeting Mitochondrial Metabolism in Clear Cell Carcinoma of the Ovaries"

**SUPPLEMENTARY LEGENDS**

**Supplementary Table 1:** Comparison of gene expression profile between scramble and siARID1A in ES2 cells. Each condition contains triplicate samples.

**Supplementary Figure 1. A.** Original of WB shown in Figure 1A (boxed area). **B.** Originals of WBs shown in Figure 2A and 2E. **C.** Amido black performed on scr. and KD ES-2 cell lysates after Seahorse experiments shown in Figures 2B and 2D showing that equal amounts of cells were subjected to measurement of mitochondrial respiration via SeaHorse. **D.** Amido black performed on scr. and KD RMG1 cell lysates after Seahorse experiments shown in Figures 2F and 2H (boxed area). **E.** Original WB shown in Figure 2I (boxed area). **F.** Amido black performed on empty vector (EV) and overexpressing (OE) TOV-21G cell lysates after Seahorse experiments shown in Figures 2J and 2L.

**Supplementary Figure 2. A.** Original of WB shown in Figure 1E (boxed area for ES-2, *left,* and entire WB for RMG1, *right*). **B.** Original of WB shown in Figure 4C (boxed area)

**Supplementary Figure 3.** Colony formation assay in ARID1A-mutated OCCCs derived TOV-21G cell line expressing either Empty Vector (EV) of pcDNA-ARID1A (overexpressing OE)
